# MENDELSEEK: An algorithm that predicts Mendelian Genes and elucidates what makes them special

**DOI:** 10.1101/2025.04.06.647432

**Authors:** Hongyi Zhou, Brice Edelman, Jeffrey Skolnick

## Abstract

Although individual Mendelian diseases—those caused by a single gene—are rare, their collective disease burden is substantial. Identifying the causal gene for each condition is essential for accurate diagnosis and effective treatment. Yet, despite decades of research, the genetic basis of more than half of all known Mendelian diseases remains unresolved. To address this gap, we introduce **MENDELSEEK**, a machine learning framework that predicts Mendelian genes by integrating residue variation scores with pathway participation, Gene Ontology processes, and protein language model features. In benchmarking across 16,946 human genes with 10-fold cross-validation, MENDELSEEK achieved an AUC of 0.869 and an AUPR of 0.737—substantially outperforming the next best methods, ENTPRISE+ENTPRISE-X (AUC 0.781; AUPR 0.626), and REVEL (AUC 0.585; AUPR 0.401). When applied to the full set of 17,858 human genes, MENDELSEEK predicted 1,277 novel Mendelian gene candidates with precision greater than 0.7. Analysis further revealed that Mendelian genes engage in significantly more protein-protein interactions than non-Mendelian genes and are evolutionarily ancient. Together, these results highlight MENDELSEEK as a major advance over existing methods, offering new insights into the biochemical features that distinguish Mendelian from non-Mendelian genes.

## Introduction

Roughly 80% of rare diseases are thought to arise from mutations in a single gene, i.e., they are Mendelian in nature; however, many of their causative genes remain unidentified^1^. Even when the genes are known, as documented in the OMIM database^2^, the mechanisms by which they give rise to the observed disease phenotypes are still poorly understood^3^. Such understanding is fundamental to the further development of precision medicine. Indeed, without knowledge of how a single gene drives a Mendelian disorder, it is unlikely that we will fully comprehend how multiple genes interact to cause non-Mendelian diseases. Consequently, significant efforts have been devoted to identifying the genetic drivers of rare diseases^4,5^. With the advent of low-cost, high-throughput next-generation sequencing (NGS) technologies, the pace of discovery has markedly accelerated, with approximately 170 to 240 Mendelian disease genes identified each year^4^. Nevertheless, exome sequencing alone cannot pinpoint which genes are the true drivers of Mendelian diseases, let alone the specific diseases they cause. For disorders with unknown genetic origins, disease–gene relationships could be uncovered through genome-wide association studies (GWAS)^6^ or bioinformatics and computational approaches^7–15^. GWAS can be applied to both germline and somatic variations, but it requires sufficiently large cohorts to achieve statistical power. Thus, for rare diseases—which by definition affect only a few individuals—GWAS is generally inapplicable. Moreover, GWAS reveals only disease-associated genes, not those that are truly causal^16^. In contrast, computational genome variation annotation tools can prioritize candidate causal genes at the level of an individual patient^7,8,12–15^. Despite their promise, however, many bioinformatics approaches have not been rigorously benchmarked on Mendelian genes, or at best, have been tested only on relatively small protein sets^7–9^.

Accurately predicting which genes are likely to be Mendelian enables the identification of the key features that distinguish them from those genes causing polygenic diseases. Recognizing such Mendelian genes also helps researchers and physicians prioritize those most likely to cause diseases or phenotypes. However, existing state-of-the-art methods often overpredict disease-causing variations, which in turn inflates the number of predicted disease-causing genes. This suggests that either the methods fail to capture the essential characteristics of disease-causing genes, or that machine learning approaches overfit the data, making the features non-transferable to new predictions. When applied to patient data, these methods frequently misrank the true disease-causing gene due to the abundance of false positives. For example, in our earlier work, we evaluated ENTPRISE^12^, SIFT^11^, and PolyPhen2-HDIV^9^ on ten patient samples. On average, ENTPRISE predicted ∼100 disease-causing genes per patient, while SIFT and PolyPhen2-HDIV predicted between 400 and 500 such genes.

To address this limitation, we developed MENDELSEEK, a machine learning framework that predicts Mendelian genes by integrating aggregate residue variation scores with intrinsic gene properties. The aim is not to identify the specific variations causing disease, but rather to determine which genes among the many mutated candidates likely underlie Mendelian diseases or phenotypes. By doing so, MENDELSEEK can filter out false positives produced by variation-based methods. MENDELSEEK accomplishes that by integrating multiple sources of gene-level information, including Reactome pathway data^17^, Gene Ontology (GO) biological processes^18^, and protein language model features^19^, alongside aggregate variation scores. The aggregate variation scores are derived using ENTPRISE^12^ for missense variations and ENTPRISE-X^13^ for frameshift and stop codon variations. This combined approach substantially outperforms methods that rely solely on aggregate variation scores, such as ENTPRISE^12^, ENTPRISE-X^13^, and REVEL^14^ —a meta-predictor that integrates 13 methods including MutPred^20,21^, FATHMM^22^, VEST^7^, PolyPhen^9^, SIFT^11^, PROVEAN^23^, MutationAssessor^24^, MutationTaster^25^, LRT^26^, GERP^27^, SiPhy^28^, phyloP^29^, phastCons^30^, —as well as the more recently developed AlphaMissense^31^.

To evaluate its performance, we benchmarked MENDELSEEK using 10-fold cross-validation on a dataset of 16,946 genes, including 4,823 known Mendelian genes from the OMIM database; all are treated as true positives^2^. MENDELSEEK demonstrated a significant improvement over state-of-the-art approaches in distinguishing Mendelian from non-Mendelian genes. Finally, by analyzing MENDELSEEK’s input features, we elucidate the biochemical characteristics that differentiate Mendelian from non-Mendelian genes.

## Methods

A flowchart of MENDELSEEK is given in Figure 1. The detailed steps are explained below.

**Figure 1:**
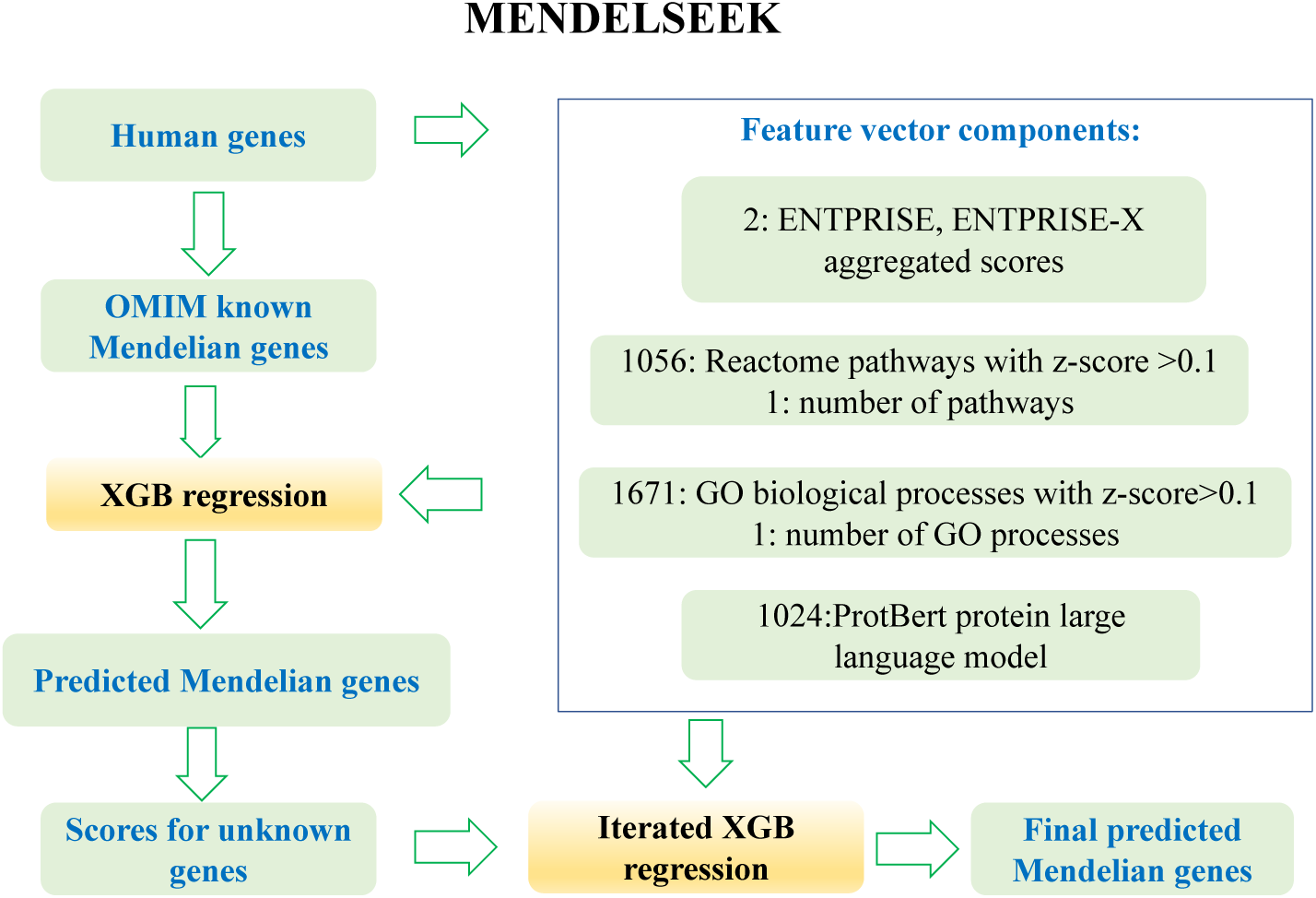
Flowchart of the MENDELSEEK algorithm.

### Calculation of the aggregate gene variation score

As discussed in ^7^, the simplest way of determining the aggregate variation score for a gene is to use the average score of all possible variations of a gene. Reference ^7^ also tested two other ways: Fisher’s method^32^ and Stouffer’s Z-score^33^. Fisher’s method requires a p-value and Stouffer’s Z-score requires a Z-score; however, neither are readily available for many methods. Here, we adopt the average variation score for use in the comparison of different variation annotation methods to MENDELSEEK. In ENTPRISE’s evaluation of the impact of missense variations, we assume that all wildtype residue positions are mutated to all other 19 amino acid types (in a real-life situation, this may not be realizable by a single nucleotide variation, but by combinations of such variations). Then, their variation scores are averaged to provide the aggregate score for a gene. Similarly, in ENTPRISE-X’s evaluation of the impact of nonsense variations, all residue positions are assumed to be mutated to a stop codon, and the resulting scores averaged. For the other variation based methods, their score or rank score as provided by the dbNSFP database (v4.2a)^34^ for a given gene is averaged. For MAVERICK^8^ for which dbNSFP does not provide results, we obtained their published pre-computed whole human genome scores and averaged that score for a given gene. For AlphaMissense, the gene-level average predictions were computed by the authors by taking the mean pathogenicity over all possible missense variants in a transcript (i.e. to all other 19 residue types, calculated the same way as ENTPRISE, see AlphaMissense_gene_hg19.tsv.gz which was downloaded from https://zenodo.org/records/8208688).

### Assignment of a gene to its pathways and GO processes

Pathways and their corresponding associated genes are obtained from the 2,363 distinct pathways in the Reactome database^17^ . Thus, its contribution to the assessment of whether a gene is Mendelian is described by a feature vector with 2,363 components. If a gene is present in a pathway, the component corresponding to this pathway is set to 1; otherwise, it is set to 0. We also include an additional component which is the total number of pathways in which a gene is involved.

The 12,535 unique human biological processes provided by the GO processes of genes are downloaded from http://geneontology.org/docs/download-ontology/^18^. If a gene is involved in a GO process, the component corresponding to that process is set to 1; otherwise, it is set to 0. Again, the total number of processes a gene is involved with is added as an additional feature component. We tested GO molecular functions and cell components and found that the best performing choice is biological process.

The protein large language model (pLLM) embedding of a gene is obtained from^19^ (https://github.com/agemagician/ProtTrans). As was also found by the ligand virtual screening method ConPlex^35^, the ProtBert embeddings model performs better than alternatives. Here, we employ the 1,024-dimensional ProtBert model which is the vector output of the deep learning-based protein language model that embeds/represents a protein sequence. The exact meaning of each component depends on the embedding token dictionary that describes the amino acid sequence used in training the pLLM.

Concatenating all the above features leads to a 15,926-dimensional feature vector for each gene: 2 dimensions are from the ENTPRISE and ENTPRISE-X aggregate scores, 2,364 dimensions are from pathways, 12,536 dimensions are from GO biological processes, and 1,024 dimensions are from the pLLM.

### Mann–Whitney U-test and feature reduction

To trace back the importance of each Reactome pathway and GO process, we employ the Mann–Whitney U-test on each component between Mendelian genes and unknown genes to calculate its z-score^36^. The component values of all genes are ranked according to their values with all tied values set to the average rank. For example, if three genes have a value of 1, they will be ranked 1, 2, 3 (which is ranked 1, or, 2, or 3 is random), their final assigned ranks will be (1+2+3)/3=2. Then, the z-score is calculated using the following equations:

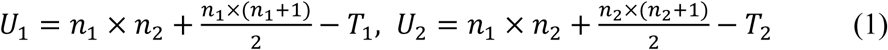

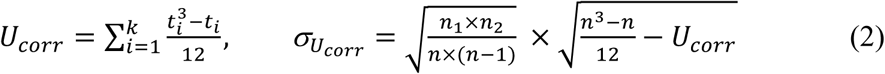

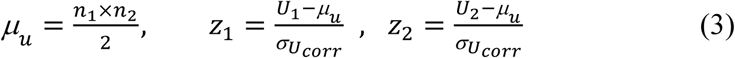

where 𝑛_1,2_are the sample numbers (here gene numbers) of group 1 (here Mendelian) and 2 (unknown); 𝑇_1,2_ are the sums of group 1, 2 ranks; 𝑈_1,2_ are the U-values of group 1 and 2; 𝑈_𝑐𝑜𝑟𝑟_ is a correction value to the U-value for calculating its standard deviation, σ_𝑈𝑐𝑜rr_; 𝑘 is the number of tied ranks and 𝑡_𝑖_ is the number of genes sharing rank 𝑖; *μ*_𝑢_ is the expected U-value of both groups 1 and 2. 𝑧_1,2_ are the z-scores of U-values from Mendelian and unknown genes.

In this work, we use 𝑧_1_ to measure the importance of pathways and GO processes to Mendelian genes. To reduce the number of pathway features (a total of 2,364) and GO processes (a total of 12,536) and to avoid possible overfitting, we only keep those features having a z-score >0.1. The z-score cutoff of 0.1 is empirically determined by scanning a small number of values from 0 to 0.5, where we found that a value of 0.1 results in a very small reduction in performance compared to the full set of features, while providing a significant reduction in the number of features. Thus, the number of pathways is reduced from 2,364 to 1,057, and the number of GO processes is reduced from 12,536 to 1,672. The final total number of features is 3,755 (compared to full set’s15,926 features).

### Machine learning and iterated training

We employed the Extreme Gradient Boosting (XGB) regression machine learning method^37^. XGB is optimized for memory usage and computational efficiency (the sparse matrix caused by many of the pathway and GO process features having 0 value components) and is less likely to cause overfitting. Thus, it is well suited for the large dimensions of the feature vectors used in this work. In practice, the GradientBoostingRegressor was implemented in the Scikit-learn kit^38^ with the following empirical parameters: n_estimators=1000, max_depth=6, and learning_rate=0.05. A known Mendelian gene from the OMIM dataset^2^ was set to an objective regression value of 1.0; otherwise, it is set to 0.0. To reduce the uncertainty of unknown genes that are treated as true negatives (thus, set to 0.0) in training, we utilize the predicted precision score (see Equation 4 below) for the unknown genes as the training values of unknown genes in a second or iterated round of training and prediction. The final predictions are provided by this iterated training model.

### Evaluation of the relative importance of protein-protein interactions

The dataset of genes to be evaluated was combined with the unique genes from the STRING (with a cutoff score of 500)^39^ and HIPPIE (with a cutoff score of 0.5)^40^ databases to assess the relative importance of protein-protein interactions, PPIs. Interestingly, we find that including protein interactomes does not improve performance and thus is not included in the final version. We explain the reason for this lack of sensitivity to the further inclusion of PPIs in the Results section.

### Benchmarking protocol

The above analysis yielded a final set of 17,858 unique genes for evaluation. Of these, 4,823 genes overlap with the OMIM dataset^2^ and are considered to be Mendelian genes. We randomly partition the genes into 10 sets and use 9 sets for training and 1 set for testing. Then, the predictions for the 10 testing sets are combined into a composite set for evaluation and novel Mendelian gene prediction (see Results). In practice, we choose cutoff independent metrics because ranking rather than scoring is often used in practical applications. In addition, for many of the other methods that we considered, there is no appropriate cutoff information available. Commonly used cutoff independent metrics are the area under receiver operating characteristic curve (AUC) and the area under precision-recall curve (AUPR). AUPR is better than AUC for measuring the ability of a method to rank true positives at the top when the dataset is unbalanced and true positives are in the minority class^41^. Here, ∼27% of the total number of genes are known true positives; thus, the total set is unbalanced. For all predictions, we convert the raw score to the predicted precision by:

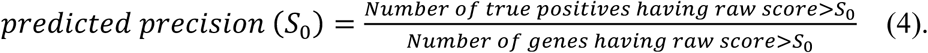

## Results

### Comparison to other methods

We compared MENDELSEEK to other methods, most of which are based on variation scores, e.g. VEST^7^. Table 1 shows the results on the 16,946-consensus gene set for the following evaluated methods besides the ENTPRISE+ENTPRISE-X score: SIFT, PolyPhen2-HDIV, PolyPhen2-HVAR, VEST4, REVEL, PrimateAI, CADD, MAVERICK, and AlphaMissense. MENDELSEEK has an AUPR, AUC and enrichment factor for the top ranked 180 gene predictions (∼top 1%) of 0.737, 0.869 and 3.28, respectively. The second-best method ENTPRISE+ENTPRISE-X has respective values of 0.626, 0.781 and 3.39. MENDELSEEK, which includes ENTPRISE+ENTPRISE-X, performs better than ENTPRISE+ENTPRISE-X alone, with a relative increase in its AUPR of 17.7%. All three measures of these two approaches perform much better than the third best method, REVEL, which is a meta-approach and whose AUPR, AUC and enrichment factor for the top 180 genes are 0.401, 0.585 and 2.53, respectively. The performance of the AlphaMissense method is surprising since it utilizes the most state-of-the-art artificial intelligence (AI) approach^31^; yet, it seems to have significantly overpredicted Mendelian or disease causing genes. Indeed, its AUPR 0.324 is behind the 0.354 result of MAVERICK and less than half of MENDELSEEK’s. Some of the alternative methods even perform worse than random selection (enrichment factors < 1 or AUC < 0.5). Figure 2 shows the AUC and AUPR curves of the compared methods. MENDELSEEK and ENTPRISE+ENTPRISE-X are well separated from the other approaches.

**Figure 2:**
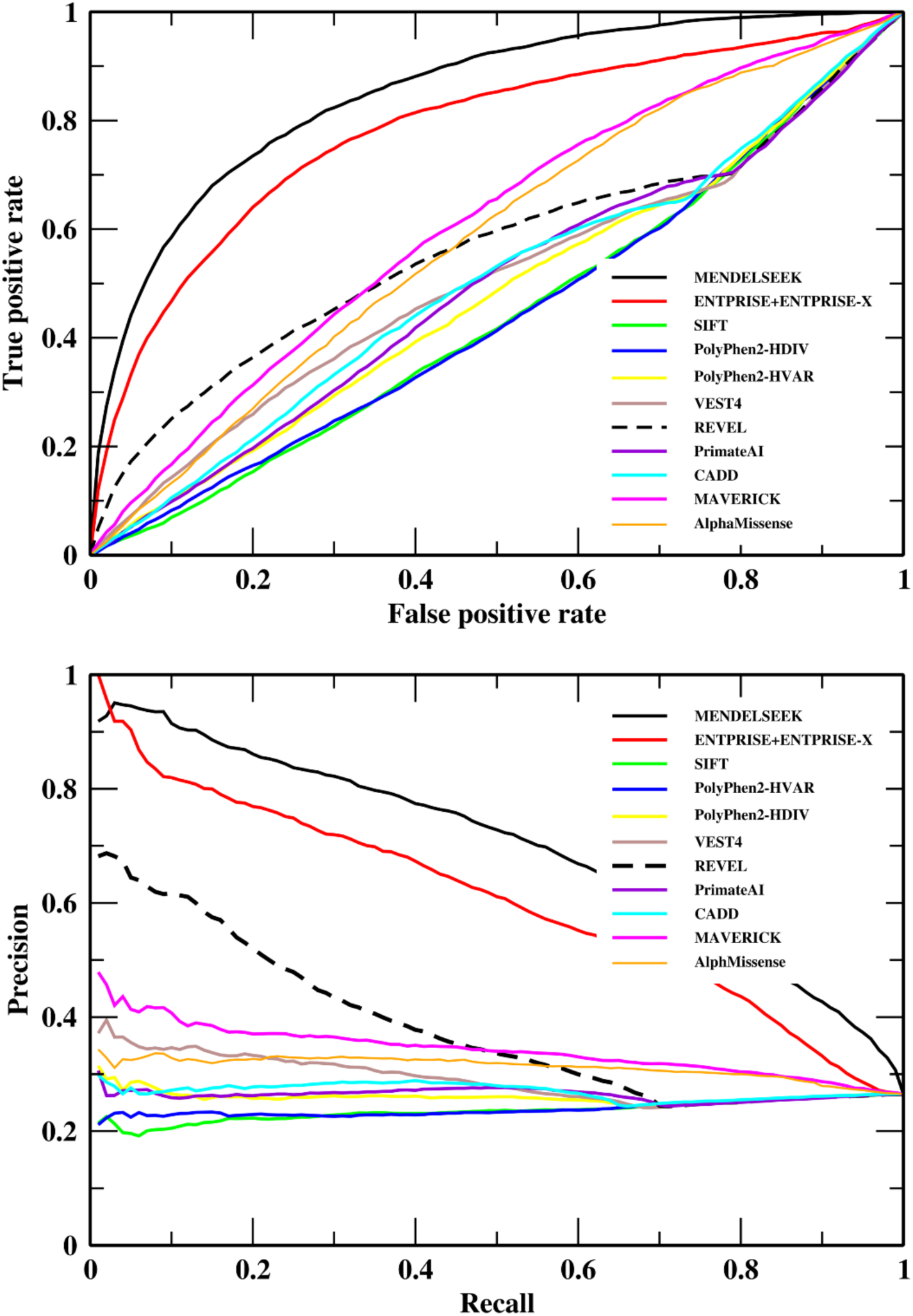
Classification performance of MENDELSEEK on the consensus 16,946 set in comparison to other methods. Receiver Operating Characteristic (upper figure) and precision-recall curve (lower figure).

**Table 1.** Comparison of the performance of different methods on the consensus 16,946 gene set.

| **Method** | **Enrichment**<br>**factor of**<br>**top ranked**<br>**180 genes<sup>a</sup>** | **AUC** | **AUPR** |
| --- | --- | --- | --- |
| MENDELSEEK | 3.28 | 0.869 | 0.737 |
| ENTPRISE+ENTPRISE-X | 3.39 | 0.781 | 0.626 |
| REVEL | 2.53 | 0.585 | 0.401 |
| MAVERICK | 1.56 | 0.597 | 0.354 |
| AlphaMissense | 1.11 | 0.576 | 0.324 |
| VEST4 | 1.50 | 0.527 | 0.310 |
| CADD | 1.05 | 0.529 | 0.292 |
| PolyPhen2-HVAR | 0.73 | 0.473 | 0.260 |
| PrimateAI | 0.87 | 0.469 | 0.256 |
| PolyPhen2-HDIV | 0.72 | 0.474 | 0.250 |
| SIFT | 0.84 | 0.467 | 0.245 |

<sup>a</sup> The maximal possible enrichment factor of the top ranked 180 genes is 3.55.

Table 2 and Figure 3 show the results on the 14,598 hard gene set that excluded the disease-causing training genes of ENTPRISE+ENTPRISE-X from the above consensus set, i.e. in this set the disease causing genes used in ENTPRISE+ENTPRISE-X training are excluded from evaluation to avoid possible bias to ENTPRISE+ENTPRISE-X. Note that the 14,598 genes may still contain training genes used within the other compared methods. Unfortunately, this information is unavailable. The best and second-best methods are again MENDELSEEK with an AUPR=0.489, AUC=0.811, and enrichment factor of the top 180 ranked genes of 4.31 and ENTPRISE+ENTPRISE-X with an AUPR=0.334, AUC=0.683, and an enrichment factor of the top 180 genes of 2.88, respectively. The relative difference of the AUPR is ∼46% (0.489 vs. 0.334). Although those AUPRs are considerably lower than those for the whole consensus set, they remain substantially higher than the next best method REVEL, whose AUPR is 0.225. The enrichment factor of MENDELSEEK is even slightly better than that for the above “easier” set (4.31 vs. 3.28). Nevertheless, the maximal possible enrichment factor for the top 180 genes is 5.70 for the hard set and 3.55 for the easy set. The AlphaMissense’s ranked fourth AUPR 0.219 is behind the 0.225 value provided by REVEL (ranked third).

**Figure 3:**
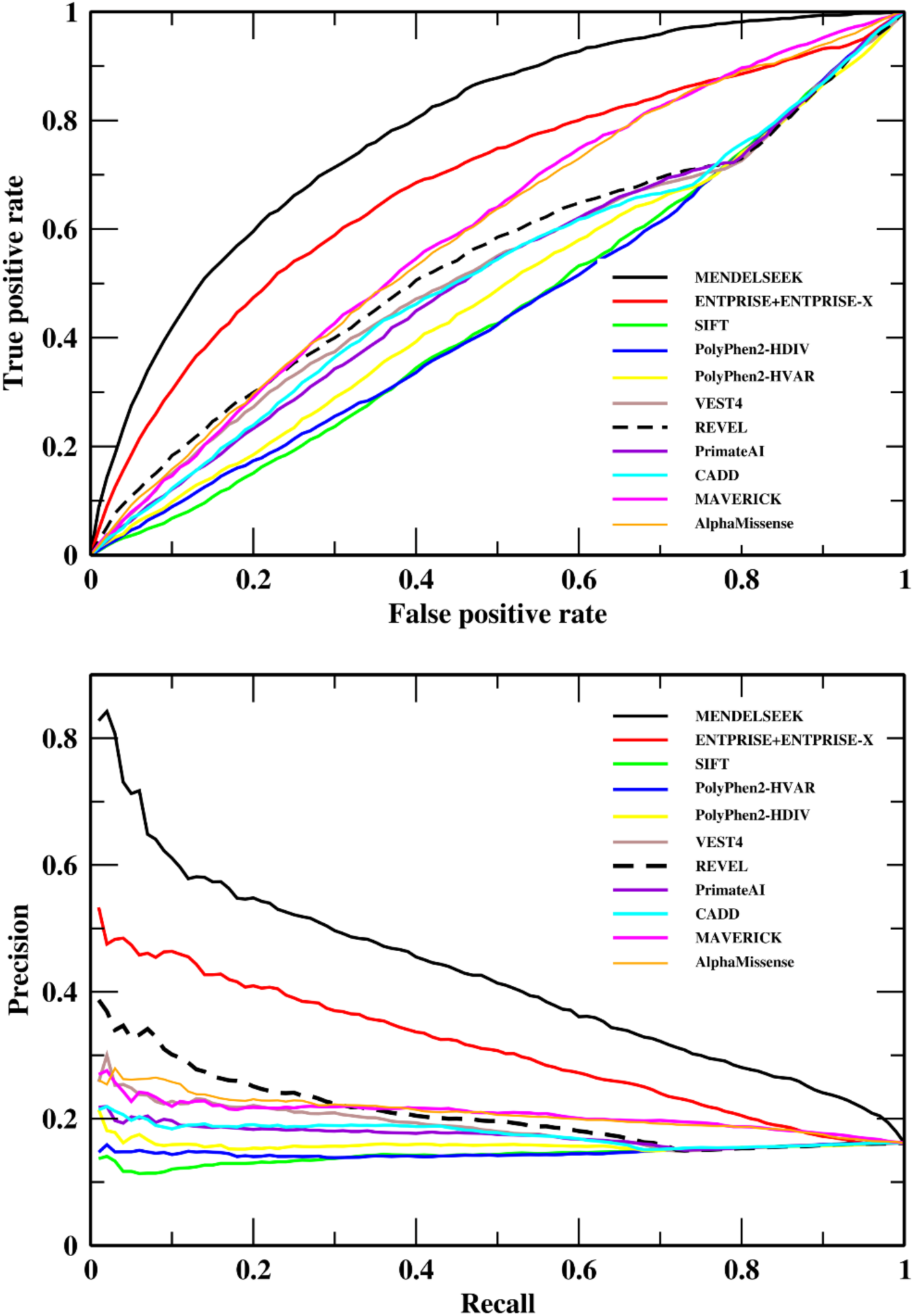
Classification performance of MENDELSEEK on the consensus 14,598 hard set in comparison to other methods. Receiver Operating Characteristic (upper figure) and precision-recall curve (lower figure).

**Table 2.**
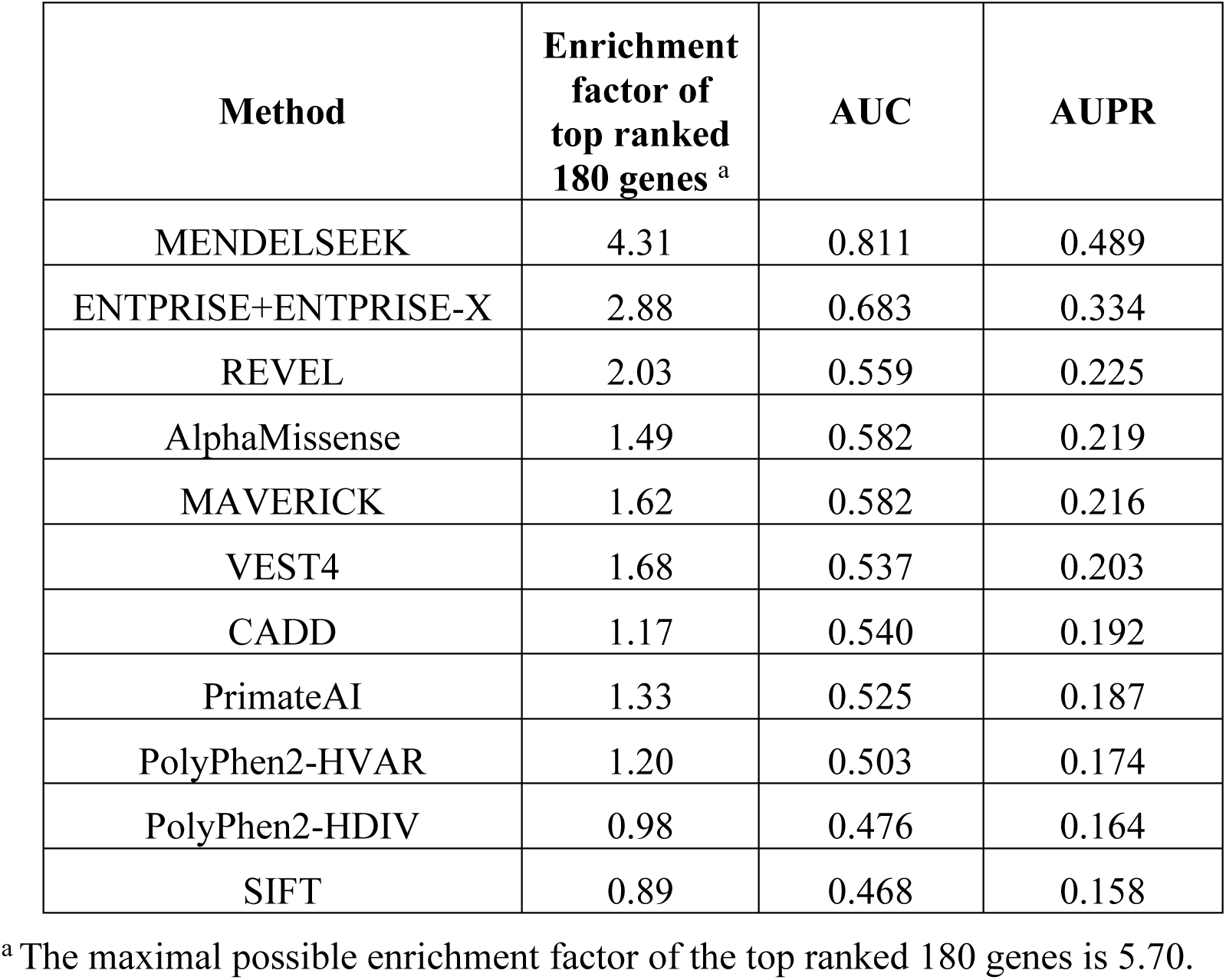
Comparison of the performance of different methods on the 14,598-member hard set.

From Figure 3, we see that MENDELSEEK and ENTPRISE+ENTPRISE-X are again well separated from the other methods, and the gap between MENDELSEEK and ENTPRISE+ENTPRISE-X becomes relatively larger compared to that of the full set (see Figure 2). MENDELSEEK’s whole gene properties contribute more significantly for genes not seen in training for ENTPRISE+ENTPRISE-X. Thus, MENDELSEEK performed significantly better than all other methods for both the whole and hard sets in terms of AUPR, AUC.

### Ablation investigation

To tease out the relative contribution of each component of MENDELSEEK to its performance, an ablation study was performed by removing one component at a time in training and doing a 10-fold cross-validation for the 17,858 gene set. Table 3 shows the results. Without the ENTPRISE+ENTPRISE-X feature component, the AUPR decreases most from 0.739 to 0.646. Exclusion of the Reactome Pathway component results in the smallest decrease of AUPR to 0.731. The next smallest decrease is to 0.729, when the GO process component is ignored. Removing protein language model features results in AUPR reduction to 0.712. Without iterated training, AUPR decreases to 0.713. Thus, the ENTPRISE+ENTPRISE-X score contributes the most to MENDELSEEK by increasing the AUPR from 0.646 to 0.739 (+14%). Iterated training increases the AUPR by ∼3.6%, and the protein language model component increases the AUPR by ∼3.8%. The pathway and GO process components contribute the least, which when included increases the AUPR by ∼1% from 0.729 and 0.731 to 0.739. This small increase is due to their correlations with the ENTPRISE+ENTPRISE-X score and to each other (see below).

**Table 3.** Ablation results on the 17,858 gene set.

| **Method** | **Enrichment factor**<br>**of top ranked 180**<br>**genes <sup>a</sup>** | **AUC** | **AUPR** |
| --- | --- | --- | --- |
| MENDELSEEK | 3.41 | 0.876 | 0.739 |
| ENTPRISE and ENTPRISE-X<br>removed | 3.27 | 0.825 | 0.646 |
| Protein language model removed | 3.44 | 0.854 | 0.712 |
| Iterated training removed | 3.62 | 0.853 | 0.713 |
| GO processes removed | 3.39 | 0.870 | 0.729 |
| Reactome pathways removed | 3.46 | 0.872 | 0.731 |

<sup>a</sup> The maximal possible enrichment factor of the top ranked 180 genes is 3.70.

### Correlations of GO processes and Reactome pathways with the number of protein-protein interactions

The above results indicate that the ENTPRISE+ENTPRISE-X aggregate variation score has the largest contribution to MENDELSEEK’s improved performance. ENTPRISE+ENTPRISE-X scores are mainly determined by the protein’s three-dimensional (3D) structure and structure-pathogenicity relationships learned from existing knowledge^12,13^. The protein language model component embeds tokens describing protein amino acid sequences and encodes intrinsic properties of proteins learned from existing knowledge^19^. As such, we are unable to dissect them further. In contrast, the GO processes and Reactome pathways have biological meaning for each component that they contribute to the feature vector. What, then, are the GO processes and Reactome pathways that most distinguish Mendelian genes from non-Mendelian genes? We analyzed those components using the Mann–Whitney U-test between true positives and the unknown ones (the majority will be true negatives) in the 17,858 dataset (see Equations 1-3)^36^.

We first analyze the possible correlation of the z-scores of the GO processes and Reactome pathways with the maximal number of protein-protein interactions (PPI) of a given process or pathway. A given gene’s number of PPIs is computed from the combined set of the STRING (with a cutoff score of 500)^39^ and HIPPIE (with a cutoff score of 0.5)^40^ databases. The maximal numbers of PPIs for the top 20 z-scores of GO processes and Reactome pathways are given in Tables 4 and 5 respectively (with the full list found in Supplemental Material Table S1 and S2). For the 12,287 GO processes having a maximal number of PPIs (>0), the Pearson’s correlation between z-scores and the corresponding maximal number of PPIs is 0.421 with a p-value of 0. For the 2,362 pathways having a maximal number of PPIs, the correlation coefficient is 0.382 with a p-value 0. These results mean that genes whose GO processes or pathways have a higher number of protein-protein interactions are more likely to be Mendelian (higher z-scores). Since they are reasonably well correlated, inclusion of the number of protein-protein interactions does not improve performance.

**Table 4.** Top 20 GO processes that distinguish Mendelian genes.

| **z-score** | **Minimal LCA <sup>a</sup>** | **Maximal PPIs** | **GO process ID** | **Name** |
| --- | --- | --- | --- | --- |
| 3.92 | 1 | 2458 | GO:0045944 | positive regulation of transcription by RNA polymerase II |
| 3.09 | 1 | 1018 | GO:0045893 | positive regulation of DNA-templated transcription |
| 2.96 | 1 | 2133 | GO:0010628 | positive regulation of gene expression |
| 2.85 | 1 | 370 | GO:0007601 | visual perception |
| 2.27 | 1 | 1196 | GO:0000122 | negative regulation of transcription by RNA polymerase II |
| 2.23 | 1 | 960 | GO:0009410 | response to xenobiotic stimulus |
| 2.17 | 1 | 1018 | GO:0043066 | negative regulation of apoptotic process |
| 1.82 | 1 | 2133 | GO:0008285 | negative regulation of cell population proliferation |
| 1.76 | 1 | 332 | GO:0007605 | sensory perception of sound |
| 1.74 | 1 | 1196 | GO:0008284 | positive regulation of cell population |
| 1.68 | 1 | 2133 | GO:0006468 | protein phosphorylation |
| 1.58 | 1 | 473 | GO:0007420 | brain development |
| 1.58 | 1 | 1018 | GO:0001701 | in utero embryonic development |
| 1.55 | 1 | 2133 | GO:0010629 | negative regulation of gene |
| 1.54 | 1 | 895 | GO:0007507 | heart development |
| 1.53 | 1 | 2458 | GO:0006357 | regulation of transcription by RNA |
| 1.45 | 1 | 287 | GO:0060271 | cilium assembly |
| 1.42 | 1 | 1108 | GO:0007165 | signal transduction |
| 1.35 | 1 | 332 | GO:0001501 | skeletal system development |
| 1.15 | 1 | 680 | GO:0001666 | response to hypoxia |

<sup>a</sup> Minimal LCA (Lowest Common Ancestor) is the minimal value of a gene's LCA as determined in <sup>42</sup> having the same GO process. Gene LCA values are from 1 to 31 with 1 being the oldest.

**Table 5.** Top 20 Reactome pathways that distinguish Mendelian genes.

| **z-score** | **Minimal LCA<sup>a</sup>** | **Maxim** | **Pathway ID** | **Name** |
| --- | --- | --- | --- | --- |
| 10.4 | 1 | 2458 | REACT:R-HSA- | Metabolism |
| 8.14 | 1 | 387 | KEGG:hsa01100 | Metabolic pathways |
| 5.72 | 1 | 2133 | REACT:R-HSA- | Signal Transduction |
| 5.43 | 1 | 2133 | REACT:R-HSA- | Immune System |
| 4.20 | 1 | 2133 | REACT:R-HSA- | Innate Immune System |
| 4.16 | 1 | 2133 | REACT:R-HSA- | Disease |
| 3.88 | 1 | 2133 | REACT:R-HSA- | Metabolism of proteins |
| 3.78 | 1 | 1196 | REACT:R-HSA- | Developmental |
| 3.69 | 1 | 1108 | KEGG:hsa05200 | Pathways in cancer |
| 3.30 | 1 | 666 | REACT:R-HSA- | Transmembrane |
| 3.20 | 1 | 1196 | REACT:R-HSA- | Cytokine Signaling in |
| 2.91 | 1 | 2458 | REACT:R-HSA- | Metabolism of lipids |
| 2.89 | 1 | 2133 | REACT:R-HSA- | Hemostasis |
| 2.68 | 1 | 1196 | REACT:R-HSA- | Axon guidance |
| 2.37 | 1 | 1108 | KEGG:hsa04151 | PI3K-Akt signaling |
| 2.34 | 1 | 1196 | REACT:R-HSA- | Signaling by |
| 2.29 | 1 | 895 | REACT:R-HSA- | Extracellular matrix |
| 2.18 | 1 | 1640 | REACT:R-HSA- | Post-translational |
| 2.14 | 1 | 666 | REACT:R-HSA- | Metabolism of amino |
| 2.12 | 1 | 1196 | REACT:R-HSA- | Signaling by NGF |

<sup>a</sup> Minimal LCA (Lowest Common Ancestor) is the minimal value of a gene's LCA as determined in <sup>42</sup> having the same pathway. Gene LCA values are from 1 to 31 with 1 being the oldest.

### Correlations of GO processes and Reactome pathways with evolutional time

In Tables 4 and 5, we also present the top 20 GO processes and Reactome pathways ranked by their z-scores along with their minimal LCA (Lowest Common Ancestor) evolutional time scale. LCA values range from 1 to 31, with 1 being the oldest, i.e., at the origin of life to 31 for the first cellular organisms (i.e. Prokaryota) as determined in ^42^. The minimal LCA is the minimal value of a gene’s LCA having/involving the same GO process/pathway. For all 12,220 GO processes having minimal LCA values, the Pearson’s correlation between the z-score and the corresponding minimal LCA is -0.215 with a p-value of 0. For the 2,355 pathways having minimal LCA values, the correlation coefficient is -0.153 with a p-value of 8.3 × 10^−1^^4^. Thus, genes likely to be Mendelian are the most ancient. This is intuitively reasonable as ancient genes and their associated functions are likely to be essential for life. As such, their disruption should have a major phenotypical effect on the organism.

Since maximal protein–protein interactions (PPIs) and minimal lowest common ancestors (LCA) of Gene Ontology (GO) processes and pathways are highly correlated with their z-scores— which characterize their ability to distinguish Mendelian genes from non-Mendelian genes—they are also strongly correlated with each other. As a result, adding any of these features to the vector does not improve MENDELSEEK’s performance; these properties are already (but implicitly) encoded in the GO processes, Reactome pathways, and the ENTPRISE+ENTPRISE-X aggregate scores. Indeed, the correlations between the ENTPRISE+ENTPRISE-X score and the LCA or number of PPIs across 17,858 genes are -0.338 and 0.228, respectively, with p-values effectively equal to 0. Direct correlations of Mendelian gene values (set to 1.0 for regression training, and 0.0 for non-Mendelian/unknown genes) with the ENTPRISE+ENTPRISE-X score, LCA, and number of PPIs are 0.477, -0.124, and 0.107, respectively, all with p-values of 0.0. Defining a gene’s maximal z_path_ or z_go proc_ as the maximal z-scores of all pathways or GO processes in which it is involved, we find correlations between the ENTPRISE+ENTPRISE-X score and maximal z_path_ and z_go proc_ of 0.405 and 0.291. Direct correlations of Mendelian gene values with maximal z_path_ and z_go proc_ are 0.221 and 0.201, respectively, compared to 0.477 for the ENTPRISE+ENTPRISE-X score. There is also a significant correlation of 0.096 (p-value = 7.8×10^-38^) between z_path_ and z_go proc_. These findings explain why ENTPRISE+ENTPRISE-X contributes most strongly to MENDELSEEK’s accuracy. When either pathway or GO process features are removed, MENDELSEEK’s AUPR decreases by only ∼1%. If both features are removed, AUPR drops from 0.739 to 0.718 (a 3% reduction). For comparison, AlphaMissense mean score’s correlations with Mendelian gene value, maximal z_path_ and z_go proc_ are 0.130, 0.239, and 0.239, respectively, compared to 0.477, 0.405, and 0.291 for the ENTPRISE+ENTPRISE-X score. This difference demonstrates that the current approach more effectively captures the essential features underlying MENDELSEEK’s superior performance.

### Analysis of the top GO processes and pathways

The top 20 GO process and pathways ranked by z-scores are the oldest with a minimal LCA of 1. The top three of these GO processes are related to gene expression: *positive regulation of transcription by RNA polymerase II, positive regulation of DNA-templated transcription,* and *positive regulation of gene expression.* These processes are essential for life. When these processes malfunction, variations/mutations in genes will happen and cause diseases or even death.

The fourth ranked GO process *visual perception* (GO:0007601) with a z-score of 2.85, has 29 phenotypes caused by 6 genes (BBS4, COL1A1, COL2A1, OAT, RDH11, RPE65) having a LCA of 1. While it may, at first glance, seem odd that visual perception has an LCA=1 (ancient organisms did not have eyes but they could perceive light^43^), these genes also engage in other essential, nonvisual processes. For example, the BBS4 gene (Bardet-Biedl syndrome 4) is a protein-coding gene that plays a role in the development and function of cilia^44^ and involves 49 human GO processes including *gene expression* (GO: 0010467). The COL1A1 gene produces a component of type I collagen that strengthens and supports many tissues in the body^45^; it is involved in 5 human GO processes including *skeletal system development* (GO:0001501). The COL2A1 gene encodes the alpha-1 chain of type II collagen which is essential for the structure and function of cartilage^46^. COL2A1 is involved in 39 GO processes including *visual perception*^47,48^, *sensory perception of sound*^49^*, skeletal system development*^50,51^*, central nervous system development*^52^, as well as other important biological functions. Furthermore, the current OMIM database^2^ lists 15 phenotypes caused by mutations in COL2A1. OAT encodes the ornithine aminotransferase enzyme, that is found in mitochondria where it helps break down ornithine. Ornithine is involved in the urea cycle and in maintaining the balance of amino acids in the body^53^. For RDH11, retinol dehydrogenase, in addition to being an essential gene in the eye, another of its 6 human GO processes involves the cellular detoxification of aldehyde^18,54^. Finally, while RPE65 helps convert light into electrical signals that are sent to the brain, this protein is also involved in 14 human GO processes including the *insulin receptor signaling pathway* (GO:0008286).

In humans, there are 29 phenotypes associated with these genes; many are eye diseases (see Table S3 for the full list) including Retinitis pigmentosa, Leber congenital amaurosis caused by RPE65, Gyrate atrophy of choroid and retina with or without ornithinemia by OAT; Retinal dystrophy caused by RDH11, and Vitreoretinopathy with phalangeal epiphyseal dysplasia (the latter is not eye related) caused by COL2A1. There are also completely non-eye related diseases: Czech dysplasia, Chondrogenesis, type II or hypochondrogenesis, Spondyloperipheral dysplasia by COL2A1; Osteogenesis imperfecta, type I, Caffey disease, Ehlers-Danlos syndrome that are caused by COL1A1. Bardet-Biedl syndrome which is caused by BBS4 affects vision, body weight, genital abnormalities and kidney functions.

The top four pathways of Mendelian genes are *Metabolism* (z-score=10.4), *Metabolic pathways* (z-score=8.1), *Signal Transduction* (z-score=5.7), *Immune System* (z-score=5.4). These generic pathways are crucial because they allow cells to efficiently capture and utilize energy from nutrients, enabling essential functions such as growth, reproduction, maintaining structure, and when uncontrolled, they could result in cancers in some organisms. The top pathway, *Metabolism* (REACT:R-HSA-1430728) is associated with 371 phenotypes caused by genes having a LCA of 1 (see Table S4 for the full list).

### Literature evidence that substantiates the predictions of novel Mendelian genes

The dataset of 17,858 unique genes obtained by combining the genes from the STRING (with a cutoff score of 500)^39^ and HIPPIE (with a cutoff score of 0.5)^40^ databases are also used for novel Mendelian gene prediction (these genes are absent in the OMIM database). We restricted our attention to predictions of genes within this interaction dataset that have known protein-protein interactions, as we have shown that genes having a higher number of protein-protein interactions are more likely to be Mendelian. Equation 4 converts the raw regression score to the predicted precision score. With a predicted precision score cutoff of 0.7, we have 1,277 novel gene predictions (those that are not in the OMIM database). These predictions of novel Mendelian genes are listed in Supplemental Material Table S5.

How can we validate these predictions? To do so, we employ our latest literature mining tool, Valsci, for validation^55^. Valsci is an open-source, self-hostable automated literature-review system that combines retrieval-augmented generation with bibliometric scoring to verify claims against the Semantic Scholar corpus and related scholarly sources. For each query, it retrieves and ranks relevant papers (augmented using citation counts, author h-index, and journal venue) and synthesizes a structured report evaluating each claim with a reasoning analysis and an ordinal 1–5 rating (from “Contradicted” to “Highly Supported”) and traceable citations. In this study, we ran Valsci with an OpenAI-compatible LLM (gpt-5-mini) as the artificial intelligence backend. We asked Valsci if a given gene is likely associated with at least one Mendelian disease based on known literature evidence. We also asked the same question for ∼1,000 randomly chosen known and unknown Mendelian genes, respectively. For the 991 known Mendelian genes, Valsci finds 509 genes with rating score ≥ 3, whereas for the 997 unknown genes Valsci has only 44 genes with rating score ≥ 3. This leads to Valsci’s precision/recall of 0.92/0.51, respectively, assuming the unknown ones are true negatives. The high precision of Valsci means MENDELSEEK has a low false positive rate (0.04) and its validated predictions are highly accurate.

Valsci returns supported literature evidence with a rating score ≥ 3 for 108 genes of the 1,277 novel Mendelian gene predictions. This leads to an effective enrichment factor of 1.9 compared to the 44/997 random unknown set. If we check around the top 1% of the 13,035 unknown genes, or the top 100 predicted novel genes, we get 10 genes with supportive evidence whereas random expected 100*44/997=4.41 genes, this results in an enrichment of 2.3. If we check the top 50(20) novel predictions, we get an enrichment factor of 2.7(4.5). This means higher ranked Mendelian genes are more likely to be recalled. Since the predicted unknown genes have not been curated by the OMIM database, their literature evidence is rare, we cannot expect the recall rate (108/1277=0.08) to be comparable to those of the known Mendelian Genes, 0.51. The 108 genes with Valsci score ≥ 3 are highly accurate based on Valsci’s precision of 0.92 for this purpose.

Examples of the top predicted genes with literature evidence that are not known to the MENDELSEEK algorithm are: ITGB1 (precision=0.90) encoding integrin beta 1, is involved in 61 pathways, and 10 of them are within the top 20 z-scores in Table 5 including Signal Transduction (z-score=5.72) and the Immune System (z-score=5.43). Its dysfunction causes kidney/renal diseases^56^. ND6 has a predicted precision of 0.90 and is involved in 9 pathways including the top two Metabolism (z-score=10.4) and Metabolic pathways (z-score=8.14). Mutations in this gene cause mitochondrial disease^57^ and Leber’s hereditary optic neuropathy (LHON)^58^. RIMS1 (precision=0.90) was documented as causing Cone-rod dystrophies (CORDs)^59^. It is involved in 21 GO processes including visual perception (z-score=2.85). Variants in SORL1 (precision=0.86) have been implicated in familial dementia^60^ and is involved in protein Metabolism (z-score=3.88).

Whether the other gene predictions with no Valsci supportive evidence are false positives or novel true positives is uncertain at this juncture. A full list of predicted genes can be found in Supplemental Material Table S5. Those with Valsci evidence (with scores of 3 or above indicating that at least some evidence in support has been found in the existing literature) for being disease causing are indicated in Table S5 and those that are not (rating score < 3) are marked “NONE”. These can serve as guidance for further bioinformatics/experimental validations/tests. A detailed report of Valsci’s results is included in the Supplemental Material.

### Difference between Mendelian genes and complex disease driving genes

Are Mendelian genes also drivers of complex diseases? Combining the known 4,823 OMIM genes in our test set and the 1,277 predicted Mendelian genes leads to 6,101 putative Mendelian genes. For putative complex disease driving genes, we have previously derived a set of 7,311 genes from 3,608 complex diseases having the gene as a driver^61^. The two sets of genes have 2,532 overlaps. Thus, a subset of the Mendelian genes are also involved in complex diseases (see Table S6). The remaining 3,569 putative Mendelian genes are not complex disease drivers, with 2,834 of these found in OMIM and another 735 predicted. Thus, roughly 60% of this set of Mendelian genes appear to be drivers of a single disease, with the remaining ∼40% being possible drivers of complex diseases as well. This is not surprising in that the malfunction of Mendelian genes is associated with the disruption of key biochemical processes.

## Discussion

In this contribution, we demonstrated that MENDELSEEK significantly outperforms other approaches including the state-of-the-art AlphaMissense^31^, in distinguishing Mendelian genes— those whose variations alone are sufficient to cause disease—from non-Mendelian genes. MENDELSEEK’s predictions are consistently supported by benchmarking tests and corroborating literature. Ablation analysis shows that the ENTPRISE+ENTPRISE-X score, reflecting residue variation, contributes the most to performance, improving AUPR by 14%. These robust results reflect the low false-positive rates of ENTPRISE^12^ and ENTPRISE-X^13^ in predicting disease-causing variants. For example, ENTPRISE exhibits a 10.7% false-positive rate compared to 36.4% for PolyPhen-2-HVAR^9^ on the 1,000 Genomes dataset^12^, while ENTPRISE-X shows an 8.4% false-positive rate compared to 18.6% for VEST-indel^13,62^ on the 1,000 Genomes dataset.

Mann–Whitney U-test analysis of individual GO processes that discriminate Mendelian from non-Mendelian genes reveals that the most discriminating processes (those with large positive z-scores) are typically associated with the oldest genes (lowest LCA values) and genes with a higher number of PPIs. A similar trend is observed for discriminative Reactome pathways. Literature mining indicates that approximately 8% (108/1277) of MENDELSEEK’s novel Mendelian gene predictions have existing literature support, while the remaining candidates represent high-value targets for experimental validation. Future directions include extending MENDELSEEK to predict not only whether a gene is Mendelian, but also its associated phenotypes or symptoms and the mode of inheritance (autosomal dominant or autosomal recessive). This approach assumes that specific phenotypes are linked to particular GO processes or pathways and that artificial intelligence can learn these patterns. A further challenge arises when a single gene gives rise to multiple phenotypes. For instance, COL2A1 is associated with 15 phenotypes in the OMIM database; thus, an important question is whether one can predict which phenotypes manifest themselves in a given patient and whether “protective” genes can prevent certain outcomes. More broadly, genetic modifiers^63^ play critical roles in Mendelian disease phenotypes, but identifying them and understanding their mechanisms remain unresolved challenges.

For non-Mendelian diseases, where dozens to hundreds of gene variants may contribute, the specific causal combinations are often unclear^12^. By contrast, the unimodal nature of Mendelian genes provides valuable insights into the link between genotype and phenotype. Developing tools that map Mendelian genes to their phenotypes will not only advance understanding of these rare disorders but also yield algorithms and principles applicable to complex, non-Mendelian diseases.

## Acknowledgments

This research was supported in part by grant GM-118039 from the Division of General Medical Sciences of the National Institutes of Health. A gift from the Ovarian Cancer Institute is also gratefully acknowledged. We thank Jessica Forness for proofreading and polishing this manuscript and Bartosz Ilkowski for his computational support.

## Author contributions

Conceptualization: JS, HZ

Methodology: HZ, BE, JS

Investigation: HZ, BE, JS

Funding acquisition: JS

Project administration: JS

Supervision: JS

Writing – original draft: HZ

Writing – review and editing: JS, HZ, BE

## Declaration of generative AI and AI-assisted technologies in the writing process

During the preparation of this work the authors have not used generative AI and AI-assisted technologies in the writing process.

## Availability of data and materials

Scripts and necessary inputs for generating the results in this work are available at https://github.com/hzhou3ga/MENDELSEEK.

## Abbreviations

GWAS: genome-wide association studies
GO: Gene Ontology
XGB: Extreme Gradient Boosting
AUC: area under receiver operating characteristic curve
AUPR: area under precision-recall curve
LCA: Lowest Common Ancestor

## Ethics approval and consent to participate

Not applicable

## Consent for publication

Not applicable

## Clinical trial number

Not applicable.

