## Supplementary material for "MENDELSEEK: An algorithm that predicts Mendelian Genes and elucidates what makes them special": ABCF1.txt

# Claim: The gene ABCF1 is likely associated with at least one Mendelian disease.

**Status**: processed

**Overall Rating**: -1

**Explanation**:

No relevant papers were found that support or refute this claim.


## Relevant Papers


## Other Reviewed Papers


### Whole exome sequencing diagnosis of inborn errors of metabolism and other disorders in United Arab Emirates

**Why Not Relevant**: The provided paper content contains a single, general sentence: "Although WES provided molecular diagnoses, the results required careful interpretations and many patients required additional investigations, suggesting this tool is useful when conventional diagnostic methods fail." This sentence does not mention the gene ABCF1, any specific gene, any specific phenotype, or any Mendelian disease. It therefore provides no direct evidence (e.g., case reports, variant data, segregation, statistical association) nor any mechanistic evidence (e.g., functional studies, pathway information) linking ABCF1 to a Mendelian disorder. The statement merely comments on the general utility and interpretive challenges of whole-exome sequencing (WES). Without gene- or disease-specific data, one cannot infer support for or against the claim that "the gene ABCF1 is likely associated with at least one Mendelian disease." Additional reasons for judging relevance as 0.0: (1) no verbatim mention of ABCF1 or any variant therein; (2) no phenotype descriptions, inheritance patterns, or clinical cases; (3) no experimental or mechanistic details that could plausibly connect ABCF1 to disease biology; (4) no statistical or population evidence. In short, the excerpt is unrelated to the claim and insufficient to evaluate it.


[Read Paper](https://www.semanticscholar.org/paper/5b148886669bca04d0a2c5fa1055e7ec2e023b89)


### Simultaneous identification and prioritization of variants in familial, de novo, and somatic genetic disorders with VariantMaster

**Why Not Relevant**: The paper describes a variant-prioritization and detection tool (VariantMaster) and reports successful identification of causative variants in a rare Mendelian disease, but the provided content does not mention ABCF1 at all nor report any results implicating specific genes by name. There is no gene-level data, no variant-level evidence, no phenotypic association, and no functional/mechanistic experiments concerning ABCF1. Therefore the paper does not provide direct evidence that ABCF1 is associated with any Mendelian disease, and only offers generic methodological plausibility (i.e., that such a tool could detect an ABCF1-disease association if present). Because the claim concerns a specific gene, and the paper contains no specific data or statements about ABCF1, its relevance to the claim is very low.


[Read Paper](https://www.semanticscholar.org/paper/a56e2dc7369c76b03ad5e9309093beb98a159719)


### Genome-wide linkage analysis and whole-genome sequencing identify a recurrent SMARCAD1 variant in a unique Chinese family with Basan syndrome

**Why Not Relevant**: The provided paper content does not mention ABCF1 or any specific gene by name. Although it reports that genome-wide linkage analysis and whole-genome sequencing were used to identify a pathogenic gene in a Basan syndrome pedigree, the excerpt omits the identity of that gene and any variant, statistical measures, segregation data, or mechanistic functional evidence. Without the gene name and supporting data, this content cannot be used to support or refute the claim that ABCF1 is associated with a Mendelian disease. In addition, no mechanistic statements (e.g., gene function, protein effect, pathogenic mechanism) relating to ABCF1 are present. For those reasons the relevance to the specific claim is negligible.


[Read Paper](https://www.semanticscholar.org/paper/04dce4679e553ed8eb3c650487ab222c72c2ac5d)


### Whole-Exome Sequencing in Adults With Chronic Kidney Disease

**Why Not Relevant**: The provided paper excerpt does not mention the gene ABCF1, report variants in ABCF1, or present any results linking ABCF1 to Mendelian disease. The content describes study aims, methods, gene lists (including a 287-gene nephropathy list and review of OMIM and ACMG genes), and general limitations of WES, but contains no direct or mechanistic evidence about ABCF1. Without the Supplementary Table 3 or results sections that might list specific gene findings, nothing in the excerpt supports or refutes the claim that ABCF1 is likely associated with a Mendelian disease. The study methodology indicates it could potentially detect Mendelian gene associations, but the absence of ABCF1 in the text means this paper, as provided, is not relevant to establishing an association between ABCF1 and any Mendelian disease.


[Read Paper](https://www.semanticscholar.org/paper/a29bd57964f68ac5f8e1abcafe1112ba488f53f5)


### Combinational analysis of linkage and exome sequencing identifies the causative mutation in a Chinese family with congenital cataract

**Why Not Relevant**: The provided paper content contains only a single general summary sentence about methodology (combining exome sequencing with linkage analysis) and does not mention ABCF1, any specific genes, variants, phenotypes, diseases, or experimental results. There is no direct evidence (no observations, variant associations, segregation data, or statistical measures) and no mechanistic biological evidence linking ABCF1 to a Mendelian disease. Because the excerpt is methodological and agnostic to gene identity, it is insufficient to support or refute the claim that ABCF1 is associated with at least one Mendelian disease.


[Read Paper](https://www.semanticscholar.org/paper/e089af83ca2af616c36c6cf122805858fecce0aa)


### Positional cloning and comprehensive mutation analysis of a Japanese family with lithium-responsive bipolar disorder identifies a novel DOCK5 mutation

**Why Not Relevant**: The provided paper content discusses DOCK5 as a biomarker for mood state, suicidality, and a possible pathological relationship with bipolar disorder (BD). It contains no mention of ABCF1, no genetic or clinical data (e.g., segregating variants, case reports, linkage/association studies, functional assays of ABCF1 variants), and no mechanistic discussion involving ABCF1. Association of a gene with a Mendelian disease typically requires evidence such as pathogenic variants segregating with disease in families, statistically significant case-control or cohort evidence, functional assays showing variant impact consistent with disease mechanism, or curated clinical assertions (ClinVar/OMIM). None of those types of evidence are present in the provided content. Therefore the paper content is irrelevant to the claim that ABCF1 is likely associated with at least one Mendelian disease.


[Read Paper](https://www.semanticscholar.org/paper/8ba88b327eac334913ca440ffacce1e093a36f4d)


### Novel variant of KIF11 associated with MCLMR syndrome

**Why Not Relevant**: This paper does not provide any evidence for or against the claim that ABCF1 is associated with a Mendelian disease. The manuscript text supplied discusses a novel heterozygous frameshift variant in KIF11 causing an autosomal-dominant microcephaly-lymphedema-chorioretinal dysplasia (MCLMR/MLCRD) phenotype. The gene ABCF1 is not mentioned anywhere in the provided text, and all genetic, clinical, and mechanistic statements concern KIF11. Therefore there are no direct data, no mechanistic connections, and no citations in this paper that bear on ABCF1. Additional points relevant to interpretation: - The paper reports a single de novo KIF11 frameshift variant (c.1858_1859delGT (p.Val620Thrfs*6)), notes absence from population and disease databases, and hypothesizes loss-of-function via nonsense-mediated decay or truncation; those observations are specific to KIF11 and cannot be generalized to ABCF1 without independent evidence. - Because the supplied content neither mentions ABCF1 nor provides pathway, interaction, expression, or functional data linking ABCF1 to KIF11 or to the reported phenotype, no mechanistic inference can be drawn about ABCF1 from this paper. - The lack of relevance is not a statement about ABCF1’s true disease association; it only means this particular paper does not inform that claim. Recommended next steps to evaluate the claim: query gene-disease resources (OMIM, ClinVar, HGMD), population constraint metrics (gnomAD pLI/LOEUF), and the primary literature for ABCF1-specific reports or functional studies.


[Read Paper](https://www.semanticscholar.org/paper/598b1c7508865997c295d53a98d38407fbecde29)


### Letter regarding “ SCN9A variant in a family of mixed breed dogs with congenital insensitivity to pain”—Navigating the pathogenicity of candidate gene mutations: Spotlight on paralog Nav genes

**Why Not Relevant**: This paper does not provide any evidence for or against the claim that ABCF1 is associated with a Mendelian disease. The manuscript and letter exclusively discuss variants in SCN9A (Nav1.7) in dogs and related evaluation methods; ABCF1 is not mentioned anywhere in the provided text. Relevant verbatim passages from the paper that demonstrate its focus on SCN9A/Nav1.7 include: "We read the recent publication by Dr. Gutierrez-Quintana et al. titled “SCN9A variant in a family of mixed breed dogs with congenital insensitivity to pain” with great interest."; "In their study, the authors conducted whole-genome sequencing analysis of a dog with congenital insensitivity to pain and identified the pathogenic variant c.C2761T (p.R921C) in SCN9A, which encodes the alpha subunit of voltage-gated neuronal sodium channel Nav1.7 that is mainly expressed in nociceptive neurons."; "Second, the authors compared whole genome sequence data of the affected dog with those of 926 healthy control dogs, and found that the SCN9A-C2761T allele was absent in controls."; "Fifth, there is a case record of the homologous genetic variation Nav1.7-R922C in a human with hereditary sensory and autonomic neuropathy (accession number; VCV000836792) in the ClinVar database."; "However, even if a similar record is found in ClinVar, which has collected millions of human genetic variants from clinical settings, the identification of single record does not necessarily mean that the genetic variant is more likely to cause disease, if it is a single record."; and "It is important to note that the current rating for human Nav1.7-R922C in ClinVar is “uncertain significance.”

Because the paper (and the commentary provided) only addresses SCN9A and related diagnostic and functional-evidence considerations (orthologous alignment, absence in 926 controls, in-silico predictions, ClinVar record, and ACMG-AMP guideline considerations), there is no direct evidence (genetic association, segregation, functional assay, clinical reports) relating ABCF1 to any Mendelian phenotype. There is also no mechanistic discussion involving ABCF1 protein function, pathogenic variant consequences, or disease pathways that could plausibly connect ABCF1 to Mendelian disease. Therefore the document is not relevant to the claim about ABCF1.


[Read Paper](https://www.semanticscholar.org/paper/39dfc4195578b71ad782115a3a552b9b9e2dae03)


### Autosomal Dominant Intellectual Development Disorder-6 (MRD6) Without Seizures Linked to a De Novo Mutation in the grin2b Gene Revealed by Exome Sequencing: A Case Report of a Moroccan Child

**Why Not Relevant**: The provided paper content exclusively discusses a mutation in the grin2b (GRIN2B) gene causing Autosomal dominant intellectual development disorder-6 (MRD6). There is no mention of the gene ABCF1, no data, no variant, no phenotypic association, and no mechanistic discussion involving ABCF1. Therefore the paper contains no direct evidence (e.g., reported ABCF1 variants segregating with disease) nor mechanistic evidence (e.g., functional studies implicating ABCF1) relevant to the claim that ABCF1 is likely associated with a Mendelian disease. Absence of ABCF1 in this content neither supports nor refutes the claim; it simply provides no information about ABCF1. Notes and caveats: (1) This conclusion is based solely on the supplied text; if the full publication contains sections not included here that discuss ABCF1, those were not available for review. (2) Lack of mention in a paper about a different gene (GRIN2B) should not be interpreted as evidence that ABCF1 is not disease-associated — it is a non-evidence circumstance.


[Read Paper](https://www.semanticscholar.org/paper/67d756e0cc302e5d46fdda9df34403f1f2de9a5a)


### Identification Of The Disease-Causing Mutation In Autosomal Dominant Familial Immune Thrombocytopenia By Genome-Wide Linkage Analysis and Whole Genome Sequencing

**Why Not Relevant**: This paper provides no evidence supporting the claim that ABCF1 is likely associated with at least one Mendelian disease. The study focuses on a four-generation family with childhood immune thrombocytopenia and reports a validated candidate disease-causing variant in NOS3 (c.1267G>A) with supporting functional data. ABCF1 is not mentioned anywhere in the provided content; the authors filtered variants and prioritized NOS3 among five candidate genes and performed functional assays specific to NOS3. Therefore the paper neither reports an association of ABCF1 with disease nor provides mechanistic data about ABCF1.

Limitations and scope of this conclusion: absence of ABCF1 from this paper is not proof that ABCF1 cannot be involved in any Mendelian disease — it only means this study does not provide supporting evidence for ABCF1. Additionally, the study itself has limitations (small single-family study, a relatively low reported LOD score of 1.1953, and linkage region spanning 215 MB containing many genes), so negative inference about other genes from this study is weak. To evaluate the claim about ABCF1, one would need studies that explicitly analyze ABCF1 variation in Mendelian disease cohorts, segregation evidence in pedigrees, functional assays of ABCF1 variants, or curated disease-gene databases; this paper does none of those.


[Read Paper](https://www.semanticscholar.org/paper/a952dbd9a3ebf1304cd3c460f802b84ef3bb424b)


## Search Queries Used

- ABCF1 pathogenic variant ClinVar OMIM Mendelian disease case report familial segregation exome sequencing de novo autosomal recessive autosomal dominant inherited disorder

- ABCF1 ATP binding cassette subfamily F member 1 function translation initiation immune response knockout mouse zebrafish developmental phenotype model organism embryo lethal

- ABCF1 loss of function intolerance pLI gnomAD constraint score haploinsufficiency probability rare variant burden study exome aggregation

- ABCF1 gene disease association monogenic rare disorder linkage analysis family based whole genome sequencing candidate gene co segregation mutation spectrum congenital disorder


## Bibliometric Configuration

- Use Bibliometrics: True

- Author Impact Weight: 0.4

- Citation Impact Weight: 0.4

- Venue Impact Weight: 0.2
