## Supplementary material for "MENDELSEEK: An algorithm that predicts Mendelian Genes and elucidates what makes them special": ABCF2.txt

# Claim: The gene ABCF2 is likely associated with at least one Mendelian disease.

**Status**: processed

**Overall Rating**: -1

**Explanation**:

No relevant papers were found that support or refute this claim.


## Relevant Papers


## Other Reviewed Papers


### The Burden of Candidate Pathogenic Variants for Kidney and Genitourinary Disorders Emerging From Exome Sequencing

**Why Not Relevant**: The provided paper content does not mention the gene ABCF2, does not report variants in ABCF2, and contains no mechanistic data (e.g., molecular function, loss-of-function evidence, inheritance patterns) specific to ABCF2. The paper focuses on genes associated with Mendelian kidney and genitourinary disorders and describes a curated list of 625 genes (and that the full list is available on the laboratory website), but the excerpted text here does not include that list or any per-gene entries. Consequently, there is no direct evidence in the provided content to support or refute the claim that ABCF2 is likely associated with at least one Mendelian disease. To evaluate the claim further, one would need the paper's full supplementary gene list (or the laboratory website resource referenced) or other studies that specifically report ABCF2 in OMIM/ClinVar/HGMD entries or provide mechanistic/segregation/functional data linking ABCF2 to a Mendelian phenotype.


[Read Paper](https://www.semanticscholar.org/paper/a75e297208c04992568e607d19aa9b9aaebd09d5)


### Autosomal Dominant Intellectual Development Disorder-6 (MRD6) Without Seizures Linked to a De Novo Mutation in the grin2b Gene Revealed by Exome Sequencing: A Case Report of a Moroccan Child

**Why Not Relevant**: The manuscript text provided exclusively describes a Mendelian association and mechanistic context for the grin2b (GRIN2B) gene and a specific GRIN2B nonsense variant causing MRD6. The paper does not mention ABCF2, report any ABCF2 variants, nor discuss functional, genetic, or clinical links between ABCF2 and Mendelian disease. Because the claim concerns ABCF2, and this paper contains no direct data, variant reports, mechanistic pathways, or literature citations related to ABCF2, it provides no evidence either supporting or refuting the claim that ABCF2 is likely associated with at least one Mendelian disease. From the absence of any ABCF2 content one cannot infer association by analogy or extrapolation—establishing a gene-disease relationship requires direct genetic/clinical evidence (case reports, segregation, population/variant data, functional validation) or at minimum a demonstrated mechanistic link to a known disease gene, none of which are present here. To evaluate the claim about ABCF2, sources explicitly mentioning ABCF2 (OMIM entries, ClinVar/ClinGen assertions, peer-reviewed case reports, or functional studies) should be sought.


[Read Paper](https://www.semanticscholar.org/paper/67d756e0cc302e5d46fdda9df34403f1f2de9a5a)


### P-540 Strategic implementation of PGT-M for multiple genes post qhole exome sequencing in couples with familial history: a comprehensive study

**Why Not Relevant**: The paper does not mention the gene ABCF2 at all and provides no gene-level lists or examples that include ABCF2. While the study uses WES and identifies pathogenic/likely pathogenic variants in multiple families, without explicit reporting of which genes were implicated there is no direct or mechanistic evidence in this text to support or refute the claim that ABCF2 is associated with at least one Mendelian disease. In other words, although the methods and results indicate the study was capable of detecting gene-disease associations, the absence of ABCF2 in the reported findings (or any table/list of implicated genes) means this paper cannot be used as evidence for the claim. Additionally, cohort-specific factors (high consanguinity, selection of couples referred for known familial disorders) limit generalizability and preclude inferring associations for genes not named in the report.


[Read Paper](https://www.semanticscholar.org/paper/1b721e406f9d31ac5983b146c5b9aa03d58fc49c)


## Search Queries Used

- ABCF2 gene pathogenic variant Mendelian disorder case report exome sequencing trio segregation ClinVar OMIM

- ABCF2 ATP binding cassette subfamily F protein function translation regulation ribosome interaction knockout mouse zebrafish CRISPR functional study

- ABCF2 loss of function constraint gnomAD pLI LOEUF haploinsufficiency intolerance population frequency rare variant burden sequencing study

- ABCF2 monogenic disease association autosomal recessive autosomal dominant de novo mutation familial segregation linkage study systematic review DECIPHER HGMD ClinVar rare disease database


## Bibliometric Configuration

- Use Bibliometrics: True

- Author Impact Weight: 0.4

- Citation Impact Weight: 0.4

- Venue Impact Weight: 0.2
