## Supplementary material for "MENDELSEEK: An algorithm that predicts Mendelian Genes and elucidates what makes them special": ACACB.txt

# Claim: The gene ACACB is likely associated with at least one Mendelian disease.

**Status**: processed

**Overall Rating**: 0

**Explanation**:

No relevant papers were found for this claim.


## Relevant Papers


## Search Queries Used

- ACACB ACACB gene ACC2 acetyl CoA carboxylase beta case report biallelic variant Mendelian disease

- ACACB ClinVar OMIM pathogenic variant exome sequencing inborn error of metabolism congenital metabolic disorder

- ACACB acetyl CoA carboxylase beta ACC2 fatty acid metabolism lipogenesis mitochondrial function knockout mouse zebrafish loss of function phenotype

- ACACB loss of function intolerance gnomAD pLI human knockout homozygous loss of function carriers population sequencing constraint Mendelian gene
