## Supplementary material for "MENDELSEEK: An algorithm that predicts Mendelian Genes and elucidates what makes them special": ACOX3.txt

# Claim: The gene ACOX3 is likely associated with at least one Mendelian disease.

**Status**: processed

**Overall Rating**: -1

**Explanation**:

No relevant papers were found that support or refute this claim.


## Relevant Papers


## Other Reviewed Papers


### Peroxisomes in Biology and Medicine

**Why Not Relevant**: The material supplied (paper title and brief description) focuses on peroxisomes and the nonspecific lipid transfer protein in rat and human liver and on phenotypic properties of preneoplastic and neoplastic hepatic lesions. It does not contain any verbatim sentences, paragraphs, figure captions, methods, results, or discussion text that mention the gene ACOX3, ACOX3 protein function, ACOX3 variants, patient phenotypes, pedigrees, or any Mendelian disease association. Therefore this paper, as provided, contains no direct evidence (no case reports, no genetic association, no linkage, no pathogenic variants) supporting or refuting the claim that ACOX3 is likely associated with at least one Mendelian disease.

Mechanistic relevance: while the paper’s focus on peroxisomes is biologically connected to ACOX3 (ACOX3 encodes a peroxisomal acyl-CoA oxidase in human/animal metabolism), the supplied content does not describe ACOX3 specifically or provide mechanistic data (e.g., alteration of ACOX3 expression, loss-of-function experiments, or biochemical defects tied to disease phenotypes). Thus any mechanistic link is speculative from the title alone and not supported by extracted evidence from this paper.

Limitations that reduce relevance:
- No textual content from the paper body was provided to allow verbatim excerpt extraction or assessment of methods/results.
- The title/subject matter is general (peroxisomes and lipid transfer proteins in liver lesions) and does not imply investigation of genetic disease associations or of the ACOX3 gene specifically.
- Without full text, figures, or supplementary data, it is impossible to determine whether ACOX3 was mentioned anywhere in the paper or whether there are incidental data relevant to Mendelian disease.

Recommendation: to evaluate the claim properly, provide the paper’s full text (so specific sentences/paragraphs can be quoted and evaluated) or consult primary genetics resources (OMIM, ClinVar, PubMed gene-centric reviews) for documented associations between ACOX3 and Mendelian disorders. Given the current inputs, the paper does not supply usable direct or mechanistic evidence regarding ACOX3’s involvement in Mendelian disease.


[Read Paper](https://www.semanticscholar.org/paper/9d15d2379ce5c3e03684056d711f086f8408552a)


### A phytol-enriched diet induces changes in fatty acid metabolism in mice both via PPARalpha-dependent and -independent pathways.

**Why Not Relevant**: The paper provides mechanistic and physiological evidence that peroxisomal fatty acid metabolism is regulated by PPARalpha and that perturbations of branched-chain fatty acid handling are relevant to Mendelian disorders (e.g., Refsum disease, peroxisome biogenesis disorders). However, the paper does not mention the gene ACOX3 by name, does not present any genetic or clinical data linking ACOX3 to disease, and contains no human genetic association or mutation data. All evidence is physiological (mouse, diet/knockout experiments) and pathway-level; thus there is no direct evidence that ACOX3 is associated with a Mendelian disease. Given the absence of any mention of ACOX3 or direct genetic/disease association data, the content is only weakly relevant (mechanistic plausibility only) to the claim.


[Read Paper](https://www.semanticscholar.org/paper/882890e34bb36ac7d61c16bde08f769b6547ea75)


### Very-Long-Chain and Branched-Chain Fatty Acyl-CoAs are High Affinity Ligands for the Peroxisome Proliferator-Activated Receptor α (PPARα)

**Why Not Relevant**: This paper excerpt does not provide direct evidence that the specific gene ACOX3 is associated with any Mendelian disease. The text discusses peroxisome proliferator-activated receptors (PPARα), the biochemistry of very-long-chain and branched-chain fatty acids, and peroxisomal β-oxidation enzymes in general, and it refers to 'AOX' (palmitoyl-CoA oxidase) in a non-paralog-specific way. While it (a) establishes that peroxisomal β-oxidation enzymes can cause Mendelian disorders when defective and (b) shows that disruption of an acyl-CoA oxidase in mice produces disease-relevant biochemical phenotypes, none of the quoted material names ACOX3, reports human genetic variation, segregation, case studies, linkage, or functional human-mutation data, or otherwise links ACOX3 specifically to a Mendelian phenotype. Further, the only in vivo genetic data are from a mouse 'AOX gene' disruption without clarity about which acyl-CoA oxidase paralog was targeted, limiting applicability to ACOX3. For these reasons the excerpt is mechanistically relevant in the abstract (it makes it plausible that peroxisomal oxidases can be Mendelian disease genes) but provides no direct or specific evidence for the claim about ACOX3; therefore overall relevance to the claim is very low.


[Read Paper](https://www.semanticscholar.org/paper/5e4b225496d17fd9d9aa433e2e474af42b7ce766)


### Resistance to erucic acid as a selectable marker for peroxisomal activity: Isolation of revertants of an infantile Refsum disease cell line

**Why Not Relevant**: The paper content provided does not mention the gene ACOX3, any variants in ACOX3, patient phenotypes, segregation data, clinical cases, or statistical associations. It only reports that certain peroxisome-related metabolic deficiencies and organelle paucity are partially corrected in selected clones and that peroxisomal activities (VLCFA β-oxidation and dihydroxyacetone phosphate acyltransferase activity) are re-established. While these findings are mechanistically relevant to peroxisomal function, they neither identify nor implicate ACOX3 specifically, nor do they provide evidence of a Mendelian disease association. Because the content lacks any direct genetic or clinical data linking ACOX3 to disease, its relevance to the specific claim is very low.


[Read Paper](https://www.semanticscholar.org/paper/3855ee673f0339c18594d95b4790cc1797970953)


### A phytol-enriched diet induces changes in fatty acid metabolism in mice both via PPARα-dependent and -independent pathways Published, JLR Papers in Press, January 16, 2005. DOI 10.1194/jlr.M400337-JLR200

**Why Not Relevant**: The provided paper excerpt does not mention the gene ACOX3, nor does it present any human genetic or clinical data linking any specific acyl-CoA oxidase gene to Mendelian disease. The evidence is mechanistic and pathway-level (mouse and in vitro studies of PPARα regulation and induction of peroxisomal enzymes) and does not include gene-specific variant, segregation, case, or population data required to support that ACOX3 is associated with a Mendelian disorder. Additionally, the experimental system is a mouse phytol-feeding model and regulatory responses in mice do not by themselves establish that variation in a given human gene causes Mendelian disease. For these reasons the paper provides mechanistic context only and no direct evidence that ACOX3 is likely associated with at least one Mendelian disease.


[Read Paper](https://www.semanticscholar.org/paper/48843dc93fec5194ad1538d021a2f1888d4ff6bd)


### NGS based studies on primary immunodeficiencies (PIDs) : causative gene identification, tool development and application

**Why Not Relevant**: This paper provides no direct or mechanistic evidence that ACOX3 is associated with any Mendelian disease. The provided content describes sequencing studies focused on primary immunodeficiency diseases (PIDs), reports specific genes implicated (e.g., LRBA, ATM, DOCK8, PMS2, TNFRSF18, PIK3CG, LILRB1, EPHB2, TXNIP, CD5, NLRP5, RAG1/RAG2 combinations, etc.), and summarizes analytical methods and diagnostic yield. Nowhere in the text is ACOX3 mentioned, nor are there mechanistic discussions (pathways, protein functions, biochemical assays) that involve ACOX3. Therefore: - There is no direct evidence in this paper supporting that ACOX3 is causal for a Mendelian disease. - There is no mechanistic evidence in this paper linking ACOX3 to disease processes (the paper focuses on immune-related genes and DNA repair/methylation effects on immunity). Important caveats: absence of mention in this single paper does not disprove any association between ACOX3 and Mendelian disease elsewhere. The study is concentrated on PIDs; ACOX3 (a peroxisomal acyl-CoA oxidase involved in fatty acid metabolism) would not be expected to appear unless it contributed to the immunological phenotypes studied or was present among variants in this cohort and flagged by the authors. Additionally, although WES was performed on many samples (and could detect variants in ACOX3), the authors only report genes they found to be implicated; ACOX3 is not among reported hits. Conclusion: relevance of this paper to the claim "ACOX3 is likely associated with at least one Mendelian disease" is essentially nil — the paper contains neither supporting nor mechanistic evidence for that claim.


[Read Paper](https://www.semanticscholar.org/paper/7786e43fbaf68904111f286747f0cf8968c2dd1e)


## Search Queries Used

- ACOX3 mutation pathogenic variant Mendelian monogenic inherited disease patient case report homozygous compound heterozygous loss of function deficiency ClinVar OMIM

- ACOX3 acyl CoA oxidase 3 pristanoyl CoA oxidase peroxisomal fatty acid oxidation pristanic acid metabolism peroxisomal disorder Zellweger Refsum disease metabolic phenotype

- ACOX3 rare variants exome sequencing genome sequencing cohort study gnomAD pLI LOEUF loss of function intolerance variant interpretation population database ClinVar summary

- ACOX3 knockout mouse zebrafish knockdown null mutation phenotype neurological liver developmental metabolic peroxisome functional study biochemical assay


## Bibliometric Configuration

- Use Bibliometrics: True

- Author Impact Weight: 0.4

- Citation Impact Weight: 0.4

- Venue Impact Weight: 0.2
