## Supplementary material for "MENDELSEEK: An algorithm that predicts Mendelian Genes and elucidates what makes them special": ACSS3.txt

# Claim: The gene ACSS3 is likely associated with at least one Mendelian disease.

**Status**: processed

**Overall Rating**: -1

**Explanation**:

No relevant papers were found that support or refute this claim.


## Relevant Papers


## Other Reviewed Papers


### How rare bone diseases have informed our knowledge of complex diseases.

**Why Not Relevant**: The provided paper content is a single general sentence: "Rare bone disorders in partnership with GWAS have brought us to the brink of a new era of personalized genomic medicine in which the prevention and management of complex diseases will be driven by the molecular understanding of each individuals contributing genetic risks for disease." This sentence contains no mention of the gene ACSS3, no presentation of experimental data, no patient or family observations, no statistical associations, no mechanistic pathways involving ACSS3, and no discussion of Mendelian disease. Therefore it provides neither direct evidence (e.g., a reported pathogenic variant in ACSS3 segregating with disease, case series, cohort association statistics) nor mechanistic evidence (e.g., biochemical function of ACSS3 linked to disease biology) to support or refute the claim that ACSS3 is likely associated with at least one Mendelian disease. Because the excerpt lacks methods, results, sample sizes, effect estimates, variant descriptions, or contextual details, it is insufficient to evaluate the claim. To assess the claim properly would require papers or database entries that specifically mention ACSS3 (e.g., clinical genetics reports, segregation studies, functional assays, ClinVar/OMIM entries, or GWAS/rare-variant analyses implicating ACSS3).


[Read Paper](https://www.semanticscholar.org/paper/1a29d961e2262c650360f47b647eb86ff5ed8bd4)


### Novel homozygous mutation in the SPATA7 gene causes autosomal recessive retinal degeneration in a consanguineous German family

**Why Not Relevant**: This paper is entirely focused on SPATA7 and its role in inherited retinal degeneration (retinitis pigmentosa and Leber congenital amaurosis). The text repeatedly documents identification of a SPATA7 homozygous variant in affected siblings, the established role of SPATA7 in Mendelian retinal disease, and a mechanistic explanation for SPATA7-related photoreceptor degeneration. Nowhere in the provided content is the gene ACSS3 mentioned, nor are there any data (genetic, phenotypic, or mechanistic) that address ACSS3. Although the study used a capture panel of 286 IRD genes, the manuscript content as provided reports findings only for SPATA7; there is no indication that ACSS3 was observed, analyzed, or implicated. Therefore the paper provides no direct evidence (no variants, co-segregation, case reports) and no mechanistic evidence linking ACSS3 to any Mendelian disease. Absence of mention in this paper is not proof that ACSS3 is not disease-associated, but it does mean this paper cannot be used to support the claim that ACSS3 is likely associated with at least one Mendelian disease. To evaluate the claim about ACSS3, a targeted literature search (OMIM, ClinVar, HGMD, PubMed) or examination of datasets listing gene-disease associations would be required.


[Read Paper](https://www.semanticscholar.org/paper/da1c2b59c82c6a0ce58e2b73a11a2be36b8e497c)


### Second cite

**Why Not Relevant**: The provided paper excerpt contains no mention of the gene ACSS3 (no symbol, no synonyms, and no sequence/variant data relating to ACSS3). The content exclusively discusses filaggrin (FLG), ichthyosis vulgaris (IV), atopic dermatitis (AD), and the identification of loss-of-function mutations in FLG (e.g. p.R510X, c.2282del4, c.3702delG) as a Mendelian cause and as a risk factor for a common disease. Therefore there is neither direct evidence (no genetic, clinical, or variant data linking ACSS3 to any Mendelian phenotype) nor mechanistic evidence (no functional, biochemical, or pathway information connecting ACSS3 to disease) in the excerpt provided. 

Limitations and caveats: (1) This assessment is limited to the text you supplied. If ACSS3 appears elsewhere in the full article outside this excerpt, that content was not available for review. (2) The paper does describe general principles and methods for discovering Mendelian disease genes (linkage, candidate gene screening, etc.), which are broadly applicable to gene discovery, but such methodological discussion does not constitute evidence that ACSS3 is associated with any Mendelian disease. (3) To evaluate the claim about ACSS3, targeted sources should be searched (e.g., full-text literature, OMIM, ClinVar, HGMD, gnomAD, PubMed) for documented pathogenic variants, reported Mendelian phenotypes, or functional studies implicating ACSS3.


[Read Paper](https://www.semanticscholar.org/paper/3cea4ccbda90825a0e49ea30b074f26e45de0875)


### Estimating the Population-Based Prevalence of Congenital Thrombotic Thrombocytopenic Purpura Using Large-Scale Sequencing Data

**Why Not Relevant**: This paper does not provide any evidence (direct or mechanistic) regarding the gene ACSS3. The manuscript exclusively studies ADAMTS13 and congenital thrombotic thrombocytopenic purpura (cTTP). Representative verbatim sentences from the provided content demonstrate the focus on ADAMTS13 and cTTP: "Thrombotic thrombocytopenic purpura (TTP) is a life-threatening disorder caused by severe ADAMTS13 deficiency." "Because of ADAMTS13 defects, an abnormal amount of ultra-large von Willebrand factor multimers is concentrated, resulting in the formation of platelet-rich microthrombi in the circulation." "We identified a total number of 2212 variants within ADAMTS13 among 282,912 alleles analyzed in the gnomAD population." "We then collected high-quality data from gnomAD population including 141,456 subjects with different ethnicities..." The paper's methods, results, and conclusions all refer to ADAMTS13 variant discovery, classification, and prevalence estimation; there is no mention of ACSS3, no functional or mechanistic data relevant to ACSS3, and no discussion of Mendelian associations for any gene other than ADAMTS13. Therefore the article is not relevant to assessing whether ACSS3 is likely associated with at least one Mendelian disease. No direct evidence (e.g., reported pathogenic variants, segregation, case reports) or mechanistic evidence (e.g., biochemical function tied to a disease phenotype) concerning ACSS3 is present in the provided content, so it cannot be used to support or refute the claim about ACSS3.


[Read Paper](https://www.semanticscholar.org/paper/e84a94fe315b28d008fc40fb3d0c7dda2e3ecab8)


### TTC7A Variants Previously Described to Cause Enteropathy Are Observed on a Single Haplotype and Appear Non-pathogenic in Pediatric Inflammatory Bowel Disease Patients

**Why Not Relevant**: The provided paper content concerns variants in the TTC7A gene and evaluates whether TTC7A variation contributes to polygenic inflammatory bowel disease (IBD), referencing compound heterozygote variants described by Lawless et al. The text makes no mention of the gene ACSS3, any ACSS3 variants, phenotypes associated with ACSS3, nor any mechanistic pathways involving ACSS3. Because the claim specifically concerns whether ACSS3 is likely associated with at least one Mendelian disease, and the paper content solely discusses TTC7A and IBD, there is no direct evidence (no reported ACSS3 variants, cases, segregation, pathogenicity assertions, or statistical associations) and no mechanistic evidence (no biochemical, cellular, or pathway data related to ACSS3) relevant to the claim. Therefore this paper provides no support for or against the claim. No verbatim excerpts about ACSS3 can be supplied because the gene is not mentioned in the provided text. Limitations: if additional sections of the paper not included in the provided content mention ACSS3, that material was not available for review; based on the given content alone, relevance to ACSS3 and Mendelian disease is effectively zero.


[Read Paper](https://www.semanticscholar.org/paper/3818a966b5d6e8b0eda65fdef043ac0727fa429c)


### Understanding Genetic Epidemiology: The Potential Benefits and Challenges of Genetics for Improving Human Health

**Why Not Relevant**: This paper is a high-level narrative review of the field of genetic epidemiology and does not mention ACSS3, provide any gene-level association data, nor discuss mechanisms specific to ACSS3. The text discusses general methods (linkage analysis, candidate gene studies, GWAS), gives examples of well-known Mendelian disorders (e.g., Tay-Sachs, Huntington’s, cystic fibrosis) and complex-trait findings (e.g., FTO, VKORC1, CYP2C9, CYP2C19), and addresses topics such as missing heritability and gene–environment interaction. None of the sentences or paragraphs in the provided content report associations, segregation, pathogenic variants, functional studies, knockout phenotypes, or mechanistic pathways for ACSS3. Therefore the paper provides neither direct evidence (no reported ACSS3 variants, case series, linkage or association results implicating ACSS3 in a Mendelian disease) nor mechanistic evidence (no discussion of ACSS3 function, biochemical pathways, or disease models) to support or refute the claim.

Limitations relevant to this assessment: (1) The manuscript is a general review (not a primary association or functional study), so gene-specific claims would only appear if the authors chose to highlight that gene; (2) The provided content appears to be from 2013, so later literature (post-2013) could contain evidence about ACSS3 not captured here. To evaluate the claim that "ACSS3 is likely associated with at least one Mendelian disease," consult gene-disease databases (OMIM, ClinVar, HGMD), case reports and family segregation studies in PubMed, population variant databases (gnomAD), and functional/animal-model literature for ACSS3. Those sources are required because this paper does not contain relevant data.


[Read Paper](https://www.semanticscholar.org/paper/1ea9ce6d8741214d75e3aa3f5dae45e0e6129c75)


### Two homozygous KRT85 mutations in a Chinese patient with pure hair and nail ectodermal dysplasia

**Why Not Relevant**: The paper reports a clinical and genetic study of pure hair and nail ectodermal dysplasia (PHNED) and identifies two homozygous mutations in KRT85 as the causal variants in the reported patient(s). There is no mention of the gene ACSS3 anywhere in the provided content. Although the authors used whole-exome sequencing (which would in principle detect variants in ACSS3), they report and discuss only KRT85 variants and mechanisms. Therefore this paper provides no direct evidence that ACSS3 is associated with any Mendelian disease. Limitations of this negative relevance: absence of ACSS3 in this single-study report does not constitute evidence that ACSS3 cannot be involved in Mendelian disease in other contexts; additionally, the paper does not present a systematic negative report of all genes (it focuses on KRT85), so lack of mention is not a definitive exclusion of ACSS3 involvement in other studies or datasets.


[Read Paper](https://www.semanticscholar.org/paper/751d0a591f939dcbbc93bea9a724918f104bce03)


### Contribution of rare transmitted and de novo variants among 2 , 871 1 congenital heart disease probands

**Why Not Relevant**: The provided paper content does not mention the gene ACSS3, either in the main text excerpts supplied or in the tables and statistics shown. The manuscript excerpts focus on congenital heart disease (CHD) cohort characteristics, overall findings about de novo and transmitted variants, and enrichment analyses for known CHD genes, but no sentences or paragraphs in the provided content reference ACSS3 or provide data (variant calls, enrichment, segregation, functional evidence) about that gene. Therefore the paper content as supplied provides neither direct evidence (e.g., reported pathogenic variants in ACSS3 segregating with a Mendelian phenotype) nor mechanistic evidence (e.g., functional studies or pathway links implicating ACSS3) relevant to the claim that ACSS3 is likely associated with a Mendelian disease. 

Limitations of this assessment: (1) Only the excerpted portions of the paper were provided. The full paper (including Supplementary Excel files and supplementary tables mentioned) might list many genes by name; ACSS3 could appear there. Because those supplementary tables are not included here, I cannot evaluate whether ACSS3 is reported elsewhere in the full article. (2) Absence of mention in the supplied text is not evidence that ACSS3 is not implicated in the study — it only means the supplied content contains no evidence for or against the claim. 

Suggested next steps to evaluate the claim: (a) Search the full paper and its supplementary materials (especially gene lists and Supplementary Excel files S1–S4 and S2 referenced) for "ACSS3"; (b) Query curated clinical resources (OMIM, ClinVar), population constraint metrics (gnomAD pLI/observed/expected, missense Z), and recent literature for ACSS3 associations or reported pathogenic variants; (c) Look for independent studies reporting segregating rare variants, functional validation (animal or cellular models), or replication in other cohorts to support a Mendelian disease link.


[Read Paper](https://www.semanticscholar.org/paper/e5830d9e95469f22518560007746202e62db26a0)


### Type : Invited Commentary Title : “ PLCD 1 and pilar cysts ”

**Why Not Relevant**: The provided paper content exclusively discusses PLCD1 (encoding PLCδ1) and its role in familial trichilemmal (pilar) cysts and, in some families, leukonychia. The text contains no mention of the gene ACSS3, no genetic data about ACSS3, and no mechanistic discussion that involves ACSS3. Key topics in the content include: identification of PLCD1 germline and somatic variants in affected families, functional assays showing reduced PLCδ1-dependent channel activation and reduced DAG generation, PLCD1 knockout mouse phenotypes, and a proposed monoallelic "two-hit" mechanism for cyst formation. Because ACSS3 is not discussed at all, there is neither direct evidence (genetic association, segregation, case reports) nor mechanistic evidence (biochemical pathway links, animal model data) in this paper content to support or refute the claim that ACSS3 is likely associated with at least one Mendelian disease. The absence of any mention of ACSS3 in the content means this paper is not informative for the claim. Limitations: it remains possible that ACSS3 is discussed elsewhere (other sections of the journal, other papers), but based strictly on the supplied content, no evidence regarding ACSS3 exists here.


[Read Paper](https://www.semanticscholar.org/paper/946fec15195f9adeac37ed32fc967a7d54d7e0bc)


### Regulation/ Gene Therapy Initial findings from whole exome sequencing of pediatric morphea lesional on of and in

**Why Not Relevant**: The provided paper content does not mention the gene ACSS3 anywhere, nor does it present data (genetic, transcriptomic, functional, or clinical) that implicates ACSS3 in any Mendelian disorder. The text discusses a variety of genes and diseases (e.g., MIF, PTPN22, NLRP1, KRT10, KRT1, SMARCAD1, ENPP1, DSG1) and describes WES, expression, and functional studies for those genes, but there are no sentences or paragraphs referring to ACSS3, ACSS3 variants, segregation with phenotype, pathogenic mechanisms involving ACSS3, or any assay results for ACSS3. Therefore there is no direct evidence (no reported ACSS3 variants associated with disease) and no mechanistic evidence (no described biochemical/pathway links, expression changes, or model systems implicating ACSS3) in this content to support or refute the claim that ACSS3 is likely associated with at least one Mendelian disease.

Limitations and implications: The absence of ACSS3 in this document neither proves nor disproves the claim; it only indicates that this specific paper does not provide relevant evidence. To evaluate the claim further, one should search disease-gene databases (OMIM, ClinVar), population and constraint data (gnomAD), and the broader literature (PubMed) for reported ACSS3 pathogenic variants, segregation data, de novo occurrences, functional studies, or clinical case reports implicating ACSS3 in a Mendelian phenotype.


[Read Paper](https://www.semanticscholar.org/paper/4e7717fce1922f46fbc50da6aaf5d727391b9e75)


### Acetyl-CoA Synthetase Short Chain Family Member-1 (ACSS1) acetylation status is an important regulator of adipose tissue thermogenic function

**Why Not Relevant**: The abstract exclusively studies ACSS1 (Acetyl-CoA Synthetase Short Chain Family Member-1) and contains no mention of ACSS3. All mechanistic and experimental evidence pertains to ACSS1 post-translational modification and its physiological effects in engineered mice; there are no human genetics data, Mendelian inheritance analyses, patient phenotypes, or references to ACSS3. Therefore the paper provides no direct or indirect evidence that ACSS3 is associated with any Mendelian disease. Additional limitations: the content is an abstract (not peer-reviewed within the journal), the experiments are in mice and focus on a specific post-translational modification rather than germline sequence variants, and there is no discussion of disease linkage or human variant pathogenicity that could be extrapolated to ACSS3.


[Read Paper](https://www.semanticscholar.org/paper/5b14a57e7dba99ba79b9de1c7386bcc34340b5dd)


## Search Queries Used

- ACSS3 mutation case report Mendelian disease exome sequencing homozygous variant compound heterozygous frameshift nonsense missense pathogenic ClinVar OMIM

- ACSS3 candidate gene rare disease gene discovery gene association monogenic genetic disorder autosomal recessive autosomal dominant pedigree segregation linkage study ACSF3 misannotation

- ACSS3 acyl CoA synthetase short chain member 3 acetate metabolism acetyl CoA synthetase mitochondrial function biochemical pathway fatty acid metabolism mouse knockout zebrafish phenotype functional assay

- ACSS3 gnomAD gene constraint loss of function intolerance pLI observed expected allele frequency population variation rare variant burden test collapsing analysis cohort study variant interpretation


## Bibliometric Configuration

- Use Bibliometrics: True

- Author Impact Weight: 0.4

- Citation Impact Weight: 0.4

- Venue Impact Weight: 0.2
