## Supplementary material for "MENDELSEEK: An algorithm that predicts Mendelian Genes and elucidates what makes them special": ACTL7B.txt

# Claim: The gene ACTL7B is likely associated with at least one Mendelian disease.

**Status**: processed

**Overall Rating**: -1

**Explanation**:

No relevant papers were found that support or refute this claim.


## Relevant Papers


## Other Reviewed Papers


### Rare genetic disorders: Beyond whole‐exome sequencing

**Why Not Relevant**: The provided paper content is a general review of next-generation sequencing technologies (WES, WGS, targeted panels, SNP arrays) and their roles in diagnosing rare genetic disorders. It does not mention the gene ACTL7B, any variants in ACTL7B, any patients or pedigrees, nor any functional or statistical data linking ACTL7B to a Mendelian phenotype. Therefore there is no direct evidence (case reports, segregation, variant pathogenicity, association statistics) and no mechanistic evidence (functional studies, molecular pathways, animal models) in this text that bears on the claim that ACTL7B is likely associated with at least one Mendelian disease.

Summary of why this content is not relevant:
- No occurrence of the gene name ACTL7B or any synonyms.
- No description of specific variants, their frequencies, or clinical phenotypes.
- No patient data, family segregation, linkage analysis, or cohort studies.
- No functional assays (in vitro, in vivo) or mechanistic pathways involving ACTL7B.
- The statements are high-level about the capabilities and limits of sequencing technologies (e.g., WES misses deep intronic variants, WGS can reveal structural variants), which are general methodological points and not evidence about any particular gene.

What would be required from a paper to support the claim:
- Verbatim reporting of pathogenic or likely pathogenic variants in ACTL7B in affected individuals with consistent phenotypes, ideally with segregation in families.
- Functional data showing that ACTL7B variants impair gene/protein function in a way that explains the phenotype (cellular assays, biochemical data, model organisms).
- Statistical evidence from case-control or cohort studies linking ACTL7B variants to disease, or multiple independent case reports.
- Consideration of allele frequency consistent with a Mendelian disorder and appropriate interpretation under ACMG/AMP guidelines.

Because none of those elements appear in the supplied text, the paper content cannot be used to support or refute the claim about ACTL7B.


[Read Paper](https://www.semanticscholar.org/paper/bd477d33a409ab5609c86cac0512c51ce998818a)


### Meta-analysis of whole-exome sequencing data from two independent cohorts finds no evidence for rare variant enrichment in Parkinson disease associated loci

**Why Not Relevant**: The paper studies rare variant enrichment within Parkinson disease (PD) genome-wide association loci using exome sequencing and replication genotyping. It does not mention the gene ACTL7B, does not test for associations with Mendelian diseases in general, nor report any analyses or results concerning ACTL7B. Therefore it provides no direct evidence supporting or refuting the claim that ACTL7B is likely associated with at least one Mendelian disease. The study's negative findings are restricted to genes in PD-associated loci in the analyzed datasets; absence of mention of ACTL7B in this PD-focused analysis is not evidence about ACTL7B's involvement in Mendelian disease. Additionally, the authors note limited power and the need for larger datasets, further limiting any indirect inference about genes not discussed.


[Read Paper](https://www.semanticscholar.org/paper/1e6b26572571d2090f6bbee2e33303523b8ae62b)


### Puzzling Out the Genetic Architecture of Endometriosis: Whole-Exome Sequencing and Novel Candidate Gene Identification in a Deeply Clinically Characterised Cohort

**Why Not Relevant**: The paper focuses on whole-exome sequencing of 80 endometriosis patients and reports rare damaging variants in genes previously associated with endometriosis and a few novel candidate genes. ACTL7B is not mentioned in the provided text or in the listed genes. The study design (targeting EM-associated genes and EM patients), small sample size for rare Mendelian discovery, and the absence of family-based segregation or functional validation mean this paper does not provide direct or mechanistic evidence for or against an association of ACTL7B with any Mendelian disease. Therefore its relevance to the claim is negligible.


[Read Paper](https://www.semanticscholar.org/paper/ebced5041637e2b958bf308e7d015f8aa88d727b)


### Association between NOTCH3 gene and Parkinson’s disease based on whole-exome sequencing

**Why Not Relevant**: The paper exclusively investigates variants in the NOTCH3 gene and their association with Parkinson’s disease in a Chinese cohort. The claim concerns ACTL7B and its likelihood of being associated with at least one Mendelian disease. The provided paper content contains no mention of ACTL7B, no data on variants in ACTL7B, and no mechanistic discussion that involves ACTL7B. Because the study's focus, methods, results, and conclusions all pertain to a different gene (NOTCH3) and phenotype (PD), none of its content provides direct or mechanistic evidence relevant to the ACTL7B–Mendelian disease claim. Absence of evidence in this paper is not evidence against the claim; assessing ACTL7B would require targeted genetic, clinical, or functional studies, or curated databases (e.g., OMIM, ClinVar) that specifically include ACTL7B.


[Read Paper](https://www.semanticscholar.org/paper/cbfdb180f6599898450c68d792c3f917150a7e1d)


### Importance of genetic studies in the diagnosis of glaucoma

**Why Not Relevant**: This paper focuses on genetic causes of glaucoma and reports pathogenic variants (and variants of uncertain significance) in genes such as MYOC, CYP1B1, FOXC1, COL2A1, SIX6, CARD10, MFN1, OPTC, OPTN, and WDR36. The provided text contains no mention of the gene ACTL7B, no association analyses involving ACTL7B, and no mechanistic discussion linking ACTL7B to any Mendelian phenotype. Therefore the paper provides no direct or mechanistic evidence to support or refute the claim that ACTL7B is likely associated with at least one Mendelian disease. Note that absence of mention in this glaucoma-focused cohort does not prove that ACTL7B is not associated with a Mendelian disease in other studies or phenotypes. To evaluate the claim further, consult dedicated genetic-disease resources (OMIM, ClinVar, HGMD), perform a PubMed search for ACTL7B + disease terms, and check large-scale sequencing databases (gnomAD, Decipher) and Mendelian gene curation resources.


[Read Paper](https://www.semanticscholar.org/paper/b36bb29dcafb8c0781b8b6d863a836b22e5a8fef)


### Genetic basis of Parkinson's disease and related disorders

**Why Not Relevant**: This paper focuses on genetic studies of Parkinson disease (familial and sporadic) and describes gene discoveries/associations (LRRK2, PRKRA, PARK2, SNCA, MAPT, etc.), genome-wide SNP and CNV analyses in PD cohorts, and replication attempts for putative PD genes. The gene ACTL7B is not mentioned anywhere in the provided text. Therefore the paper provides no direct evidence supporting or refuting an association between ACTL7B and any Mendelian disease. The only relevant information is indirect: the paper lists genes it implicates or tests for PD and describes methods (GWAS, autozygosity mapping, CNV detection) and their limits. Absence of ACTL7B from these reported findings is weak, indirect evidence that this study did not identify ACTL7B as a Mendelian disease gene in the contexts examined (mostly PD). Important limitations: (1) absence of mention in this PD-focused paper is not evidence that ACTL7B is not associated with any Mendelian disease elsewhere; (2) GWAS and SNP-array CNV methods used are best suited to common variants and large structural changes and are underpowered to discover rare, high-penetrance Mendelian variants or single-nucleotide variants unless targeted or sequenced; (3) the cohorts analyzed are PD-focused and may not include phenotypes relevant to other Mendelian diseases possibly caused by ACTL7B. Given these factors, the paper is essentially non-informative with respect to the claim that ACTL7B is likely associated with at least one Mendelian disease.


[Read Paper](https://www.semanticscholar.org/paper/b59ed2e51a73f5ce5958c18560d750b40e4034b6)


### Whole‐exome sequencing analyses in a Saudi Ischemic Stroke Cohort reveal association signals, and shows polygenic risk scores are related to Modified Rankin Scale Risk

**Why Not Relevant**: The provided paper content does not mention the gene ACTL7B, does not report gene-level associations for specific genes, and does not discuss Mendelian diseases. It describes gene-burden testing in a stroke cohort and an association between aggregate rare-variant burden in stroke genes and clinical outcome (mRS). Because the claim concerns whether ACTL7B is likely associated with at least one Mendelian disease, and the excerpt contains no data, named results, mechanistic discussion, or negative evidence regarding ACTL7B, the paper content is not relevant to supporting or refuting the claim. Additionally, without a full list of tested genes or gene-level results, one cannot infer from the excerpt whether ACTL7B was tested, observed, or excluded. Therefore the evidence in this excerpt is insufficient to evaluate the claim.


[Read Paper](https://www.semanticscholar.org/paper/ef1e3cc537505def74437c2048ef722316d024a6)


### British Society for Matrix Biology Autumn 2020 Meeting: “Basement Membranes in Health and Disease”

**Why Not Relevant**: The provided paper content does not mention the gene ACTL7B (no occurrence of the gene symbol or alternate names). There are no patient data, no Mendelian segregation analyses, no variant reports, no gene-disease association statistics, and no experimental perturbations (knockout, knockdown, overexpression) of ACTL7B described. The manuscript focuses on extracellular matrix biology (examples: giantin/GOLGB1 knockout effects on collagen processing, integrin α1β1 function, collagen VII and GBM components, and genes such as LAMB2, COL4A3/4/5, Col5a2), but none of these sections link mechanistically or experimentally to ACTL7B. Although the paper discusses Mendelian ECM diseases (for example, dystrophic epidermolysis bullosa and Alport syndrome) and mechanisms of collagen and basement membrane biology, that content does not provide direct or inferential mechanistic evidence implicating ACTL7B in any Mendelian disease. Therefore the paper provides no relevant direct evidence and no mechanistic evidence supporting the claim that ACTL7B is likely associated with at least one Mendelian disease. Limitations: It remains possible that ACTL7B could be implicated through pathways discussed in the paper, but such an implication would be speculative because the authors did not measure, manipulate, or report on ACTL7B; there is no sequence/variant/phenotype data tied to ACTL7B in the provided text. To establish relevance, one would need explicit mentions of ACTL7B (or synonyms), genetic association data, patient phenotypes, segregation evidence, or experimental perturbations showing disease-relevant phenotypes.


[Read Paper](https://www.semanticscholar.org/paper/e27ef31faa9280438adb116ad5e8faaf9838ff15)


## Search Queries Used

- ACTL7B actin like 7B variant pathogenic Mendelian disease monogenic case report family segregation autosomal recessive autosomal dominant

- ACTL7B exome sequencing undiagnosed rare disease cohort gene burden rare variant association trio whole exome sequencing clinical sequencing study

- ACTL7B function actin like 7B knockout mouse phenotype zebrafish knockdown cell model cytoskeleton spermatogenesis cilia functional characterization

- ACTL7B ClinVar OMIM gnomAD pLI loss of function intolerance constraint gene disease validity review systematic review meta analysis


## Bibliometric Configuration

- Use Bibliometrics: True

- Author Impact Weight: 0.4

- Citation Impact Weight: 0.4

- Venue Impact Weight: 0.2
