## Supplementary material for "MENDELSEEK: An algorithm that predicts Mendelian Genes and elucidates what makes them special": ACVR1C.txt

# Claim: The gene ACVR1C is likely associated with at least one Mendelian disease.

**Status**: processed

**Overall Rating**: -1

**Explanation**:

No relevant papers were found that support or refute this claim.


## Relevant Papers


## Other Reviewed Papers


### The Burden of Candidate Pathogenic Variants for Kidney and Genitourinary Disorders Emerging From Exome Sequencing

**Why Not Relevant**: The provided paper excerpt addresses identification and curation of genes associated with Mendelian kidney and genitourinary disorders and the prevalence of candidate pathogenic variants in a large control cohort. However, the excerpt does not mention the gene ACVR1C by name, nor does it present gene-specific variant data, mechanistic discussion, or case reports for ACVR1C. Because of that, the paper content supplied gives no direct evidence for or against the claim that ACVR1C is likely associated with at least one Mendelian disease. Additional reasons the excerpt is not informative for this claim: (1) the curated gene list (625 genes) is referenced but not shown, so one cannot tell whether ACVR1C is included; (2) the study focus is on kidney and genitourinary disorders—if ACVR1C is associated with a Mendelian disease outside those organ systems it may not have been considered; (3) no variant-level results, segregation, functional assays, or prior-disease reports specific to ACVR1C are provided in the text supplied. Recommended next actions to evaluate the claim: query authoritative gene–disease databases (OMIM, ClinVar, Orphanet, HGMD) for ACVR1C; search the primary literature for terms like “ACVR1C mutation,” “ACVR1C Mendelian,” or full gene name (activin A receptor type 1C / ALK7) combined with “familial,” “segregation,” or “pathogenic”; and review ClinVar/HGMD entries and any reported functional evidence (loss-of-function, gain-of-function, segregation, de novo occurrence) to assess whether ACVR1C has an established Mendelian association. Based on the content provided, no firm conclusion about ACVR1C can be drawn from this paper excerpt.


[Read Paper](https://www.semanticscholar.org/paper/a75e297208c04992568e607d19aa9b9aaebd09d5)


### Germline Variant Interpretation in Children with Severe Sepsis

**Why Not Relevant**: The supplied paper content does not mention ACVR1C or provide any gene-level data that would allow assessment of whether ACVR1C is associated with a Mendelian disease. The only sentence provided is: "The authors report that 200/330 of children admitted to the PICU with severe sepsis who underwent exome sequencing and had a clinically significant genetic variant in a gene(s) associated with a known inborn error of immunity (IEI) that may have contributed to the patient’s clinical presentation." This statement is broad and summarizes aggregate findings (number of patients with variants in one or more IEI-associated genes) but: (1) it does not list the specific genes or variants found, (2) it does not mention ACVR1C, (3) it does not provide variant-level pathogenicity, segregation, or functional evidence linking any particular gene to a Mendelian phenotype, and (4) an association with an "IEI-associated gene(s)" in this cohort is not sufficient by itself to establish that a specific gene (such as ACVR1C) is causally tied to a Mendelian disease. Because no direct or mechanistic evidence concerning ACVR1C appears in the provided content, the paper (as supplied) is not relevant to the claim that "ACVR1C is likely associated with at least one Mendelian disease." To evaluate that claim would require either (a) the paper naming ACVR1C among the implicated genes with supporting variant and clinical data, or (b) other primary evidence (e.g., case reports, segregation, functional studies, or database annotations) specifically addressing ACVR1C. None of those are present in the excerpt provided.


[Read Paper](https://www.semanticscholar.org/paper/8148b944a985ef1eb6ab7a15ba1b7990c783459a)


### The simplest explanation does not have to be preferred: co-occurrence of pathogenic variants in cancer-predisposing genes

**Why Not Relevant**: The provided paper content contains a single sentence fragment about reviewing the literature and UK 10,000 Genomes Project data for MINAS and cancer phenotypes, and whether multiple cancer susceptibility genes (CSGs) act additively or synergistically. It does not mention the gene ACVR1C, any specific genes, variants, Mendelian diseases, segregation data, case reports, functional assays, or any other data that could be used as direct or mechanistic evidence linking ACVR1C to a Mendelian disease. Therefore there is no direct evidence (no association statements, no variant-disease correlations, no statistical results) and no mechanistic evidence (no pathway, protein function, or pathogenic mechanism) in the provided text relevant to the claim. Additional limitations: the text is fragmentary and appears to be an introductory or methods-summary sentence rather than results or discussion; without the rest of the paper it is impossible to determine whether ACVR1C is addressed elsewhere. To support the claim that ACVR1C is likely associated with at least one Mendelian disease, the paper would need to provide explicit mentions of ACVR1C with supporting data (e.g., rare pathogenic variants segregating in families, statistical enrichment in cases vs controls, functional validation, or citation to established databases such as OMIM). None of those elements are present in the supplied content, so relevance to the claim is effectively zero.


[Read Paper](https://www.semanticscholar.org/paper/a1372a664bd2c33a0bda638975a8b26d112457fc)


### Case report: Preimplantation genetic testing for infantile GM1 gangliosidosis

**Why Not Relevant**: The paper exclusively addresses GM1 gangliosidosis caused by pathogenic variants in the GLB1 gene and describes diagnostic, genetic, and reproductive-management aspects specific to GLB1 (including variant details, PGT-M development for GLB1, and embryo genotypes). There is no mention of the gene ACVR1C, no ACVR1C variants, no phenotypes attributed to ACVR1C, and no mechanistic discussion linking ACVR1C to Mendelian disease. Therefore this paper provides no direct or mechanistic evidence supporting or refuting the claim that ACVR1C is likely associated with at least one Mendelian disease. Additionally, the study is a single-family case report focused on GLB1, so even indirect inference across genes would be unsupported. For these reasons the relevance to the ACVR1C claim is essentially zero.


[Read Paper](https://www.semanticscholar.org/paper/ce981bc89c565bb4d97baff441baaf8695595448)


## Search Queries Used

- ACVR1C ALK7 activin A receptor type 1C mutation pathogenic variant Mendelian disease case report family segregation autosomal dominant autosomal recessive

- ACVR1C ClinVar OMIM HGMD pathogenic variant germline monogenic hereditary disease clinical report

- ACVR1C ALK7 signaling SMAD2 SMAD3 kinase domain mutation developmental disorder congenital malformation mouse knockout phenotype human phenotype functional study

- ACVR1C whole exome sequencing whole genome sequencing rare disease cohort trio de novo loss of function rare variant segregation population constraint gnomAD pLI


## Bibliometric Configuration

- Use Bibliometrics: True

- Author Impact Weight: 0.4

- Citation Impact Weight: 0.4

- Venue Impact Weight: 0.2
