## Supplementary material for "MENDELSEEK: An algorithm that predicts Mendelian Genes and elucidates what makes them special": ADAM11.txt

# Claim: The gene ADAM11 is likely associated with at least one Mendelian disease.

**Status**: processed

**Overall Rating**: -1

**Explanation**:

No relevant papers were found that support or refute this claim.


## Relevant Papers


## Other Reviewed Papers


### Case report: A founder UGDH variant associated with developmental epileptic encephalopathy in Saudi Arabia

**Why Not Relevant**: The provided paper content exclusively describes a case study implicating the UGDH gene in a congenital disorder of glycosylation and gives mechanistic and population context for UGDH variants. There is no mention of ADAM11 anywhere in the supplied text, nor any data, discussion, or analysis that would directly or indirectly attribute pathogenicity to ADAM11. Therefore the paper does not provide evidence for or against the claim that ADAM11 is associated with a Mendelian disease.


[Read Paper](https://www.semanticscholar.org/paper/3f260fe2ca29d1be06660849ea4e56a0f7a2e9e1)


### Phenotypic and molecular reanalysis of a cohort of patients with monogenic diabetes reveals a case of partial lipodystrophy due to the A8344G mutation in the mitochondrial DNA

**Why Not Relevant**: The paper describes a single clinical case of familial partial lipodystrophy-like presentation that was ultimately attributed to a mitochondrial variant (m.A8344G). It focuses on LMNA and PPARG as the nuclear genes most commonly implicated in FPLD and reports sequencing of a 48-gene nuclear panel plus the mitochondrial genome. The text contains no mention of ADAM11, no experimental data (genetic, functional, segregation) involving ADAM11, and no mechanistic discussion linking ADAM11 to disease. Therefore the paper provides no direct evidence supporting the claim that ADAM11 is likely associated with at least one Mendelian disease. It also provides no mechanistic evidence that would increase or decrease the plausibility of ADAM11 as a Mendelian disease gene. Relevant limitations to the negative inference: absence of ADAM11 in this paper does not prove ADAM11 is not associated with Mendelian disease (the study was not designed to assess ADAM11, the 48-gene panel is not fully enumerated so ADAM11 may not have been assessed, and the paper reports a single case with a clear mitochondrial finding). Given these points, relevance to the ADAM11 claim is essentially zero.


[Read Paper](https://www.semanticscholar.org/paper/b84ebce0f78f814af19d16356bda255e32515b9d)


### Meningoencephalitis in a novel mutation in MNGIE (mitochondrial neurogastrointestinal encephalomyopathy) ending a familial diagnostic odyssey: A case series report

**Why Not Relevant**: This paper describes a familial case series of MNGIE and identifies a novel TYMP (thymidine phosphorylase) variant as causative in the reported family, plus a literature/database survey of MNGIE and MNGIE-like cases implicating genes such as POLG1, RRM2B, LIG3, RRM1, MTTV1, and MT-RNR1. Nowhere in the provided text is ADAM11 mentioned, nor are any data (genetic, mechanistic, functional, or epidemiologic) presented that link ADAM11 to MNGIE or to any other Mendelian disease. Therefore the paper provides no direct evidence supporting the claim that ADAM11 is likely associated with at least one Mendelian disease. It also provides no mechanistic evidence (e.g., pathway, protein interaction, expression, functional assay) implicating ADAM11. The only possible bearing is negative (i.e., the paper implicates other specific genes and not ADAM11), which does not constitute evidence for or against ADAM11 in the broader literature. In summary: relevance to the claim is effectively zero because the manuscript's genetic findings and mechanistic discussion concern TYMP and specified mitochondrial genes, and do not include ADAM11 or data that would plausibly implicate ADAM11 in Mendelian disease.


[Read Paper](https://www.semanticscholar.org/paper/27a86c38d19bd5a7eeb789892804f65161a6376e)


### TARGETED SEQUENCING REVEAL TWO PATHOGENIC MUTATIONS IN LDLR AND APOE GENES IN PATIENT WITH FAMILY HYPERCHOLESTEROLEMIA: CASE REPORT

**Why Not Relevant**: The paper reports a targeted genetic study of a single familial hypercholesterolemia (FH) case and identifies likely pathogenic variants in LDLR and APOE. ADAM11 is not mentioned anywhere in the provided content. Therefore the paper provides no direct evidence for or against an association between ADAM11 and any Mendelian disease. Important limitations that explain this lack of relevance: (1) the study used a targeted cardiovascular gene panel (TruSight Cardio) rather than whole exome/genome sequencing, so genes not on the panel would not be assayed or reported; (2) the sample is a single patient (with parents), so the study is not designed to discover novel gene-disease associations beyond the FH-relevant genes; and (3) the authors themselves recommend whole exome sequencing to identify other genes, indicating this work was not intended to comprehensively assess all possible Mendelian genes. Consequently, absence of ADAM11 in the report is not evidence that ADAM11 is not associated with Mendelian disease — it only shows this paper does not address the claim.


[Read Paper](https://www.semanticscholar.org/paper/04808c76b3962e7f9dc6e3ebec4ddbb8d15d0607)


### #1761 Whole-genome sequencing reveals contribution of rare and common variation to structural kidney and urinary tract malformations

**Why Not Relevant**: The paper does not mention ADAM11 or any data directly implicating or excluding ADAM11 as a Mendelian disease gene. The study focuses on structural kidney and urinary tract malformations and reports the genes and signals found in that context (e.g., HNF1B, AUTS2, ARHGAP5, ZNF879). Because ADAM11 is not reported among identified or statistically enriched genes, this paper provides at best weak, indirect evidence against ADAM11 being a Mendelian cause for the specific phenotypes studied. However, absence of mention in this cohort is not strong evidence that ADAM11 is not associated with any Mendelian disease: (1) the study’s phenotype focus may not include disorders caused by ADAM11, (2) very rare Mendelian associations can be missed given the diagnostic yield (4.3%) and cohort size, (3) analytic filters and case ascertainment could omit relevant ADAM11 variants, and (4) lack of statistical enrichment here does not exclude single-family or extremely rare pathogenic variants that would require different evidence (segregation, functional studies). Therefore the paper is not directly relevant to concluding that ADAM11 is likely associated with a Mendelian disease.


[Read Paper](https://www.semanticscholar.org/paper/490f0cc0aac67ac259732ccd40d6cf231466b691)


## Search Queries Used

- ADAM11 mutation variant monogenic Mendelian disease case report familial exome sequencing genome sequencing clinical phenotype congenital familial

- ADAM11 loss of function constraint gnomAD pLI observed expected o e o e haploinsufficiency population genetics ClinVar allele frequency rare variants

- ADAM11 knockout mouse phenotype loss of function model neurodevelopment synapse adhesion extracellular matrix metalloproteinase like non catalytic ADAM family functional study

- ADAM11 ClinVar OMIM DECIPHER Mendelian gene review ADAM family monogenic disease gene discovery diagnostic exome sequencing cohort replication study


## Bibliometric Configuration

- Use Bibliometrics: True

- Author Impact Weight: 0.4

- Citation Impact Weight: 0.4

- Venue Impact Weight: 0.2
