## Supplementary material for "MENDELSEEK: An algorithm that predicts Mendelian Genes and elucidates what makes them special": ADCY2.txt

# Claim: The gene ADCY2 is likely associated with at least one Mendelian disease.

**Status**: processed

**Overall Rating**: -1

**Explanation**:

No relevant papers were found that support or refute this claim.


## Relevant Papers


## Other Reviewed Papers


### Bi-allelic variants in TSPOAP1, encoding the active zone protein RIMBP1, cause autosomal recessive dystonia

**Why Not Relevant**: The submitted paper exclusively reports variants in TSPOAP1 (RIMBP1) and presents clinical, mouse-model, and in vitro mechanistic evidence linking TSPOAP1 dysfunction to autosomal recessive dystonia. The paper contains no mention of ADCY2, no data on ADCY2 variants, and no discussion connecting ADCY2 to the reported phenotypes or mechanisms. Therefore it provides no direct or mechanistic evidence for the claim that ADCY2 is likely associated with a Mendelian disease.


[Read Paper](https://www.semanticscholar.org/paper/49eb714efa47e65d21efcee0825d875c24a89950)


### Editorial: entering the age of whole-exome sequencing in rheumatic diseases: novel insights into disease pathogenicity.

**Why Not Relevant**: I reviewed the provided paper content in full. The text is an editorial discussing general principles and practical interpretation of whole-exome sequencing for Mendelian disease discovery (e.g., modes of inheritance, variant filtering strategies). It contains no mention of the gene name ADCY2 (or any ADCY2 synonyms), no reported variants in ADCY2, no case reports, no family segregation data, and no experimental or clinical results linking ADCY2 to any Mendelian phenotype. Thus the paper provides neither direct evidence (no observations, variants, or statistical associations involving ADCY2) nor mechanistic evidence (no functional studies, pathways, or biological plausibility statements about ADCY2) relevant to the claim that ADCY2 is likely associated with at least one Mendelian disease. The editorial content could be informative about how one would identify such a gene (for example, by describing exome sequencing and variant-filtering strategies), but it does not itself provide any gene-specific data that would support or refute the claim. Therefore relevance is 0.0.


[Read Paper](https://www.semanticscholar.org/paper/074f60e6d3662d0b3a5326ac719f8776de7a8c70)


### Case report: A novel de novo deletion mutation of DYRK1A is associated with intellectual developmental disorder, autosomal dominant 7

**Why Not Relevant**: The provided paper is entirely focused on DYRK1A (Intellectual developmental disorder 7/DYRK1A syndrome) and reports a de novo pathogenic deletion in DYRK1A in a single patient. Nowhere in the supplied text is ADCY2 mentioned, nor are there genetic, clinical, functional, mechanistic, population, or epidemiological data pertaining to ADCY2. Consequently, this paper provides no direct evidence that ADCY2 is associated with any Mendelian disease, nor does it provide mechanistic evidence that would make such an association plausible or implausible. The absence of ADCY2 discussion means the study is irrelevant to the claim. (Note: absence of evidence in this paper is not evidence that ADCY2 is not disease-associated — it simply means this paper cannot be used to evaluate the claim.)


[Read Paper](https://www.semanticscholar.org/paper/98eb2bb2850f75cb23ca3954609b1a07001ab461)


### Complex PKD1 Genetics in Early-Onset Cystic Kidney Disease

**Why Not Relevant**: The provided paper content does not mention ADCY2, nor does it present any clinical, genetic, or mechanistic data concerning ADCY2. The manuscript excerpt exclusively discusses PKD1 variants in pediatric-onset cystic kidney disease and briefly lists other cystic-kidney related genes (GANAB, DNAJB11, HNF1B, IFT140, PKHD1, ALG8, ALG9, DZIP1L, COL4A3/4/5) as possible contributors; ADCY2 is not included. The content therefore provides no direct evidence (no case reports, variant associations, segregation data, or statistical associations) linking ADCY2 to a Mendelian disease, nor any mechanistic evidence (biochemical pathways, functional studies, animal models) implicating ADCY2 in monogenic pathology. Additionally, the methodological and contextual discussion (phase determination, interpretation of missense variants, linkage disequilibrium, and ADPKD polycystin quantity hypotheses) is general to PKD1 and variant interpretation and does not provide transferrable evidence to support ADCY2 involvement in Mendelian disease. For these reasons the paper is not relevant to the claim that ADCY2 is likely associated with at least one Mendelian disease. To assess that claim, literature or data that explicitly reports pathogenic ADCY2 variants with segregation or functional validation, population-level enrichment, or mechanistic links to a Mendelian phenotype would be required.


[Read Paper](https://www.semanticscholar.org/paper/93aa5634e2918da0adf9d4d1273325fd11dae0af)


### Case Report: A novel de novo variant of COL1A1 in fetal genetic osteogenesis imperfecta

**Why Not Relevant**: The paper's content exclusively concerns a novel de novo COL1A1 variant (c.1309G>A, p.Gly437Ser) identified in a single fetus with osteogenesis imperfecta and discusses mechanistic reasons why a glycine substitution in the collagen triple helix is likely pathogenic. There is no mention of the gene ADCY2 anywhere in the provided text, nor any data, analysis, or discussion relevant to ADCY2. Therefore this paper provides no direct genetic association evidence (case reports, linkage, segregation, or statistical association) nor mechanistic evidence (biochemical pathway, functional assays, or predicted functional impact) linking ADCY2 to a Mendelian disease. Additional limitations relevant to the paper itself (single-case report, reliance on in silico prediction rather than functional validation) are moot with respect to ADCY2 because the gene is not addressed. For these reasons the relevance of this paper to the claim that "ADCY2 is likely associated with at least one Mendelian disease" is effectively zero.


[Read Paper](https://www.semanticscholar.org/paper/1caeaf6717a5af526be5f1f0dcef41ef4c377699)


### Identification and characterization of novel glomerulus-associated genes and proteins

**Why Not Relevant**: The paper studies adenylate cyclase type I (Adcy1) in mouse and other glomerular genes (dendrin, Crb2). It does not mention ADCY2 (adenylate cyclase type 2) or present human genetic evidence (segregation, pathogenic variants, or clinical reports) linking ADCY2 to a Mendelian disease. While the paper provides mechanistic context showing that adenylate cyclase family members produce cAMP and that loss of Adcy1 in mouse can create susceptibility to proteinuria under stress, this is indirect: different gene (ADCY1 vs ADCY2), different species (mouse), and the observed phenotype is stress-dependent/susceptibility rather than a clear Mendelian, fully penetrant disorder. No human mutation data, linkage, or disease association for ADCY2 are provided. For these reasons the paper provides negligible direct evidence for the claim that ADCY2 is likely associated with at least one Mendelian disease, though it offers weak mechanistic plausibility that other adenylate cyclase family members could affect kidney-related phenotypes.


[Read Paper](https://www.semanticscholar.org/paper/56b41bb5fe12fcfbcad36932804f3170b64a296a)


### Case report: A de novo Non-sense SOX9 mutation (p.Q417*) located in transactivation domain is Responsible for Campomelic Dysplasia

**Why Not Relevant**: This paper provides no evidence for or against the claim that ADCY2 is associated with a Mendelian disease. The manuscript exclusively discusses a novel heterozygous nonsense variant in SOX9 (c.1249C > T; p.Q417*) identified in a fetus with campomelic dysplasia (CD), and interprets pathogenicity in terms of SOX9 haploinsufficiency/reduced transactivation. There is no mention of ADCY2, adenylyl cyclase genes, or any data (genetic, functional, or mechanistic) related to ADCY2. Although whole-exome sequencing (WES) was used to find the causal variant, the authors report only the SOX9 result; the absence of any ADCY2 finding in the paper is not equivalent to negative evidence for ADCY2 involvement, it is simply not addressed. Therefore this paper is irrelevant to the claim. Limitations: one cannot infer from this paper that ADCY2 is not involved in Mendelian disease (no negative screen or report of ADCY2), but equally there is no supportive direct or mechanistic evidence presented linking ADCY2 to any Mendelian phenotype.


[Read Paper](https://www.semanticscholar.org/paper/bcc43a4127c8fa8d83c22e39fd3a13cbfa12fe1a)


### Case Report: Charcot-marie-tooth disease caused by a de novo MORC2 gene mutation - novel insights into pathogenicity and treatment

**Why Not Relevant**: This paper does not provide any direct or mechanistic evidence regarding the gene ADCY2. The study reports a single patient with Charcot-Marie-Tooth disease and identifies a de novo heterozygous missense mutation in MORC2 (NM_001303256.3: c.1199A>G, NP_001290186.1: p.Gln400Arg), and discusses clinical findings, diagnosis of CMT type 2Z, and potential treatment. ADCY2 is not mentioned anywhere in the provided content, and no data (positive association, negative finding, functional assay, segregation analysis, or database annotation) related to ADCY2 are presented. As such, there is neither direct evidence (observational or genetic) nor mechanistic evidence (pathway, functional studies, or molecular interactions) in this paper that supports or refutes the claim that ADCY2 is likely associated with at least one Mendelian disease. Limitations that further prevent any inference about ADCY2 from this paper: (1) the study is a single-case report focused on MORC2; (2) the authors report only the disease-causing variant they identified (MORC2) and do not report negative results for other genes; (3) no population-level, segregation, functional, or replication data relevant to ADCY2 are provided. If you want evidence about ADCY2 specifically, I recommend searching gene-disease databases (OMIM, ClinVar, Orphanet), variant repositories (HGMD, ClinVar), population frequency resources (gnomAD), and PubMed for publications linking ADCY2 to Mendelian phenotypes.


[Read Paper](https://www.semanticscholar.org/paper/f3b7a2400b32386ec81caae9078245d16ad29850)


### Calcium and cAMP homeostasis determine network organisation of respiratory pre-Bötzinger neurons in Mecp2 null mice in vitro

**Why Not Relevant**: The paper investigates cAMP signaling and adenylate cyclase activity generically in a Mecp2 knockout mouse model of Rett syndrome, providing mechanistic evidence that AC activity and cAMP homeostasis are altered in that Mendelian disease model. However, it does not mention ADCY2 (the ADCY2 gene or its protein product) by name, does not present genetic evidence (variants, segregation, linkage, de novo mutations, or patient phenotypes) implicating ADCY2 in any Mendelian disorder, and uses pharmacologic and imaging approaches that are not isoform-specific. Therefore the paper provides only indirect, mechanistic context (that adenylate cyclases as a class affect cAMP in a Mendelian disease model) but no direct or isoform-specific evidence supporting the claim that ADCY2 is likely associated with at least one Mendelian disease.


[Read Paper](https://www.semanticscholar.org/paper/39de7835d5d3d35c8c20d3cd7a53c9c03d166ba0)


## Search Queries Used

- ADCY2 pathogenic variant Mendelian disease case report exome sequencing whole exome sequencing de novo mutation

- ADCY2 monogenic disease autosomal dominant autosomal recessive loss of function missense nonsense frameshift pathogenic variant

- adenylate cyclase 2 ADCY2 cAMP signaling neuronal development synaptic function knockout mouse phenotype functional study

- ADCY2 ClinVar OMIM gnomAD pLI loss of function intolerance population variation no disease association review


## Bibliometric Configuration

- Use Bibliometrics: True

- Author Impact Weight: 0.4

- Citation Impact Weight: 0.4

- Venue Impact Weight: 0.2
