## Supplementary material for "MENDELSEEK: An algorithm that predicts Mendelian Genes and elucidates what makes them special": ADIPOR1.txt

# Claim: The gene ADIPOR1 is likely associated with at least one Mendelian disease.

**Status**: processed

**Overall Rating**: -1

**Explanation**:

No relevant papers were found that support or refute this claim.


## Relevant Papers


## Other Reviewed Papers


### Beyond the exome: utility of long-read whole genome sequencing in exome-negative autosomal recessive diseases

**Why Not Relevant**: The single provided sentence (“A path forward is proposed for the implementation of lrWGS sequencing in the setting of autosomal recessive diseases in a way that maximizes its utility.”) contains no mention of the gene ADIPOR1, no mention of any specific Mendelian disease, no data, and no mechanistic discussion linking any gene to disease. Therefore the paper content as provided supplies neither direct evidence (e.g., genetic association, case reports, segregation data, variant pathogenicity, or statistical associations) nor mechanistic evidence (e.g., functional studies, pathways, protein function changes) relevant to the claim that ADIPOR1 is likely associated with at least one Mendelian disease.

Limitations and reasons for irrelevance:
- No gene-level information: The excerpt does not name ADIPOR1 or any other gene.
- No disease-level information: It generically references 'autosomal recessive diseases' without specifying any phenotype or diagnosis.
- No primary data or methods: The sentence describes a proposed implementation path for lrWGS but provides no experimental results, case series, or analyses that could link a gene to disease.
- No mechanistic detail: There is no discussion of molecular mechanisms, protein function, pathogenic variants, or biological pathways that could support plausibility for ADIPOR1 involvement in Mendelian disease.
- Insufficient context/length: Only a single, high-level sentence was provided; even if the full paper discussed ADIPOR1 elsewhere, that content is not present here and cannot be evaluated.

Because of these points, the provided content cannot be used to support or refute the claim that ADIPOR1 is likely associated with at least one Mendelian disease.


[Read Paper](https://www.semanticscholar.org/paper/97e2370b00b1c292463a0e0f0eb6ee73b3066cbf)


### Precision Medicine in Internal Medicine

**Why Not Relevant**: The provided paper content is a general review of genetic testing methods, variant interpretation, and the state of gene–disease discovery. It contains no mention of ADIPOR1, no gene-specific data, no case reports, cohort analyses, linkage studies, functional experiments, or ClinVar/OMIM entries for ADIPOR1. Therefore it supplies only indirect, general context (e.g., that exome sequencing can detect novel gene–disease associations and that ~18% of genes were known Mendelian disease genes as of 2019). Because the claim concerns a specific gene's likely association with at least one Mendelian disease, and the paper offers no direct or gene-specific mechanistic evidence, its direct relevance to the claim is negligible.


[Read Paper](https://www.semanticscholar.org/paper/c7199e8a5226f8d9d55777da0e15e91797bb459a)


### Voretigene neparvovec for inherited retinal dystrophy due to RPE65 mutations: a scoping review of eligibility and treatment challenges from clinical trials to real practice

**Why Not Relevant**: The single provided sentence does not mention ADIPOR1, any specific gene, genetic variants, inheritance patterns, Mendelian disease, case reports, cohort genetics, linkage/association analyses, functional studies, or other data that could link a gene to a monogenic disorder. It only states a clinical/therapeutic observation about retinal degeneration and the importance of photoreceptor preservation for potential functional rescue. There is no direct evidence (e.g., pathogenic variants in ADIPOR1 causing disease) and no mechanistic evidence (e.g., ADIPOR1 expression/function in photoreceptors or molecular pathways leading to retinal degeneration) present in the excerpt. Because of these gaps the excerpt neither supports nor refutes the claim that ADIPOR1 is likely associated with at least one Mendelian disease. Limitations: only a single sentence was provided; if the full paper contains additional sections (methods, results, tables) that discuss ADIPOR1 or genetic evidence, those were not supplied and could change the relevance assessment. As given, the content is clinical context only and not informative for evaluating the genetic claim.


[Read Paper](https://www.semanticscholar.org/paper/e12434709606d93a377e141d4b1ca43a6ed74c1a)


### Whole exome sequencing identifies novel compound heterozygous pathogenic variants in the MYO15A gene leading to autosomal recessive non-syndromic hearing loss

**Why Not Relevant**: The provided paper content contains only a single, general summary sentence about the successful use of whole-exome sequencing to identify the molecular basis of ARNSHL in a patient. It does not mention the gene ADIPOR1, any specific variants, inheritance related to ADIPOR1, functional studies, or mechanistic pathways involving ADIPOR1. Therefore the content provides no direct or mechanistic evidence relevant to the claim that ADIPOR1 is likely associated with at least one Mendelian disease. Because the available text lacks any gene-level or disease-association detail, its relevance to the claim is effectively zero. Additionally, the snippet lacks experimental details (sample, methods, results, statistics) preventing any inference about other genes that might have been implicated in the study.


[Read Paper](https://www.semanticscholar.org/paper/13c5a6e28a8861e38b0ec26b82dc90c2ea6f4df0)


### Gene augmentation of LCA5-associated Leber congenital amaurosis ameliorates bulge region defects of the photoreceptor ciliary axoneme

**Why Not Relevant**: This paper does not provide any direct or mechanistic evidence related to the gene ADIPOR1 or its association with Mendelian disease. The content exclusively discusses Leber congenital amaurosis type 5 (LCA5), the protein lebercilin encoded by LCA5, and associated proteins RP1, IFT81 and IFT88, as well as LCA5-deficient mouse phenotypes and AAV-based LCA5 gene augmentation. There is no mention of ADIPOR1, adiponectin receptors, ADIPOR1-related phenotypes, or any genetic association data concerning ADIPOR1. Therefore: 

- Direct evidence: none. The paper contains no genetic association, patient data, variant pathogenicity analysis, linkage, segregation, or case reports concerning ADIPOR1.
- Mechanistic evidence: none. The molecular mechanisms described (lebercilin localization at the photoreceptor outer segment bulge, effects on axoneme, RP1 and IFT proteins, membrane disc formation) are specific to LCA5-related biology and do not implicate ADIPOR1 or provide a plausible mechanistic link to ADIPOR1-associated disease.
- Limitations and interpretive note: absence of ADIPOR1 in this paper means it cannot be used to support or refute the claim that ADIPOR1 is likely associated with a Mendelian disease. Absence of evidence in a single paper is not evidence of absence overall, but for the purpose of evaluating this paper's contribution to the claim, its relevance is effectively zero.

Because the paper contains no ADIPOR1-relevant sentences or paragraphs, no verbatim excerpts are provided and no mechanistic or direct evidence can be extracted from this text to address the claim.


[Read Paper](https://www.semanticscholar.org/paper/4d9f2bf75c6414edf990d585775747e3a75defee)


### RPE65 mutations in Leber congenital amaurosis, early-onset severe retinal dystrophy, and retinitis pigmentosa from a tertiary eye care center in India

**Why Not Relevant**: This paper does not provide any direct or mechanistic evidence regarding the gene ADIPOR1. The manuscript excerpt exclusively concerns RPE65 mutations in inherited retinal diseases (LCA, EOSRD, RP), methods used to screen RPE65 or an IRD gene panel, and the frequency/phenotype of RPE65 variants in the studied cohort. The provided text contains no mention of ADIPOR1, no reported ADIPOR1 variants, no functional or mechanistic data relevant to ADIPOR1, and no segregation or statistical analyses involving ADIPOR1. Although the Methods note a "targeted NGS gene panel for IRDs covering 260+ genes," the excerpt does not list which genes were included nor report any findings for ADIPOR1; therefore nothing in the text can be used to support or refute the claim that ADIPOR1 is likely associated with at least one Mendelian disease. Important caveats: absence of ADIPOR1 in this excerpt is not evidence that ADIPOR1 is not associated with Mendelian disease generally — it only means this particular paper and the provided content do not address ADIPOR1. To evaluate the claim one would need a paper that (a) reports pathogenic ADIPOR1 variants segregating with a Mendelian phenotype, (b) demonstrates functional impact consistent with disease mechanism, or (c) shows statistical enrichment of rare damaging ADIPOR1 variants in affected individuals versus controls.


[Read Paper](https://www.semanticscholar.org/paper/fcba678325dd2502e7b7123d521ff853e1ef9080)


### Identification of a novel compound heterozygous SMG9 variants in a Chinese family with heart and brain malformation syndrome using whole exome sequencing

**Why Not Relevant**: The provided paper content discusses SMG9 only and contains no mention of ADIPOR1 or any data, analyses, mechanisms, clinical observations, variants, or conclusions relating to ADIPOR1. Example sentences from the paper: "This report is the first description of a non-consanguineous Chinese pedigree with novel compound heterozygous variants in the SMG9 gene, and the molecular confirmation of the patient expands the genetic spectrum ofSMG9-deficiency syndrome, andThe manifestation of SMG 9- deficiency syndrome in the patient provides additional clinical information regarding this syndrome." Because the claim to be evaluated is specifically about the gene ADIPOR1 being likely associated with at least one Mendelian disease, and because the paper addresses a different gene (SMG9) and SMG9-deficiency syndrome, there is no direct evidence (e.g., variant-disease co-segregation, functional assays, case reports) nor mechanistic evidence (e.g., protein function, pathway overlap, molecular consequence of ADIPOR1 variants) in this paper that bears on ADIPOR1. Limitations and caveats: absence of any mention of ADIPOR1 in this paper means it cannot be used to support or refute the claim about ADIPOR1; however, lack of relevance of this single paper does not imply anything about the true association between ADIPOR1 and Mendelian disease in the broader literature. To evaluate the claim about ADIPOR1, papers that explicitly report ADIPOR1 variants in patients, functional studies of ADIPOR1 variant effects, segregation analyses, or curated disease-gene resources (e.g., OMIM, ClinVar, GeneReviews) should be examined instead.


[Read Paper](https://www.semanticscholar.org/paper/f6121a84131030977a3f1e5d9ffbf6f78963a53e)


### The simplest explanation does not have to be preferred: co-occurrence of pathogenic variants in cancer-predisposing genes

**Why Not Relevant**: The provided paper content makes no mention of ADIPOR1, ADIPOR1 variants, or any phenotype directly attributable to ADIPOR1. The excerpt discusses the frequency and cancer phenotypes associated with MINAS, review of recent literature, and analysis of UK 10,000 Genomes Project data for participants with tumor predisposition syndromes or multiple primary tumors, and whether effects are additive or synergistic among cancer susceptibility genes (CSGs). None of these sentences provide direct evidence that ADIPOR1 is associated with a Mendelian disease, nor do they provide mechanistic information involving ADIPOR1. Therefore, based on the supplied content, the paper does not provide evidence for or against the claim.

Limitations and caveats: The assessment is restricted to the single excerpt you provided. It is possible the full paper contains sections, tables, supplemental materials, or gene lists that mention ADIPOR1, but there is no indication of that in the text you supplied. To determine whether ADIPOR1 is associated with a Mendelian disease, a focused search of databases such as OMIM, ClinVar, and PubMed for ADIPOR1 and Mendelian phenotypes (or review of the full paper text and its gene lists/supplementary data) would be required.


[Read Paper](https://www.semanticscholar.org/paper/a1372a664bd2c33a0bda638975a8b26d112457fc)


### Sex-specific attenuation of photoreceptor degeneration by reserpine in a rhodopsin P23H rat model of autosomal dominant retinitis pigmentosa

**Why Not Relevant**: The provided paper content contains no mention of the gene ADIPOR1, no genetic association analyses, no variant reports, and no discussion of Mendelian inheritance for ADIPOR1. The excerpt focuses on (1) the therapeutic testing of reserpine in rodent models of inherited retinal degeneration (including the rhodopsin P23H rat and rd16 mouse that models CEP290-related disease), (2) functional outcomes (scotopic/a-wave/b-wave, contrast threshold, outer nuclear layer thickness), and (3) retinal transcriptome results describing sex-specific upregulation of phototransduction, proteostasis-related, and stress-response genes. None of these statements provide direct evidence that ADIPOR1 is associated with any Mendelian disease, nor do they name ADIPOR1 as part of the transcriptomic changes reported.

Because the paper excerpt does not (a) report ADIPOR1 sequence variants in patients or model organisms, (b) present segregation or case reports linking ADIPOR1 to a Mendelian phenotype, or (c) mechanistically implicate ADIPOR1 by name in disease pathways discussed, it yields neither direct nor mechanistic evidence for the claim. Absence of mention in this excerpt is not evidence that ADIPOR1 is not disease-associated; it only indicates this paper does not address the claim. To evaluate the claim further would require searching genetic-disease databases (e.g., OMIM, ClinVar), or literature specifically reporting ADIPOR1 variants and segregation/functional studies linking those variants to a Mendelian disorder.


[Read Paper](https://www.semanticscholar.org/paper/c209e1906261951db9f95224140e299c81985953)


### Probing the sub-cellular mechanisms of LCA5-Leber Congenital Amaurosis and associated gene therapy with expansion microscopy

**Why Not Relevant**: The provided paper content exclusively addresses Leber Congenital Amaurosis type 5 (LCA5) and the protein lebercilin, including mechanistic studies and gene augmentation therapy in mice. There is no mention of ADIPOR1, its sequence, phenotypes, patient genetics, model organism data, or any experimental or clinical evidence linking ADIPOR1 to disease. Therefore the paper provides no direct evidence that ADIPOR1 is associated with any Mendelian disease, nor does it provide mechanistic evidence that would plausibly connect ADIPOR1 to the conditions studied. Because the gene and disease focus differ (LCA5 versus ADIPOR1), and no inferential data regarding ADIPOR1 are present, the relevance of this paper to the claim is effectively zero.


[Read Paper](https://www.semanticscholar.org/paper/8ec94716d741a071af24f3ab3b17f3ccc87d0f19)


## Search Queries Used

- ADIPOR1 adiponectin receptor 1 Mendelian disease monogenic OMIM ClinVar pathogenic variant mutation inherited disease

- ADIPOR1 retinal dystrophy retinitis pigmentosa Leber congenital amaurosis inherited retinal disease photoreceptor degeneration mutation

- ADIPOR1 whole exome sequencing whole genome sequencing family segregation autosomal recessive compound heterozygous rare variant novel mutation

- ADIPOR1 knockout mouse zebrafish CRISPR photoreceptor degeneration retinal degeneration lipid metabolism ceramidase activity functional characterization disease model association study


## Bibliometric Configuration

- Use Bibliometrics: True

- Author Impact Weight: 0.4

- Citation Impact Weight: 0.4

- Venue Impact Weight: 0.2
