## Supplementary material for "MENDELSEEK: An algorithm that predicts Mendelian Genes and elucidates what makes them special": ADIPOR2.txt

# Claim: The gene ADIPOR2 is likely associated with at least one Mendelian disease.

**Status**: processed

**Overall Rating**: -1

**Explanation**:

No relevant papers were found that support or refute this claim.


## Relevant Papers


## Other Reviewed Papers


### Chronic increase of circulating galanin levels induces obesity and marked alterations in lipid metabolism similar to metabolic syndrome

**Why Not Relevant**: Verbatim sentence from paper content: "The hypothesis that increased circulating GAL levels contribute to the development of metabolic syndrome is supported, as chronically elevated GAL may regulate body weight, metabolic rate, and lipid and carbohydrate metabolism through a mechanism that is independent of feeding regulation." 

Why this is not relevant to the claim that ADIPOR2 is likely associated with at least one Mendelian disease:
- The provided paper content discusses GAL (galanin) and its possible role in metabolic syndrome. It does not mention ADIPOR2, adiponectin receptor 2, or any other gene.
- There is no genetic data, association analysis, variant reporting, segregation, case reports, or clinical descriptions that would constitute direct evidence linking ADIPOR2 to a Mendelian disorder.
- There is no mechanistic description connecting GAL signaling to ADIPOR2 function, regulation, or pathogenic variation. Therefore no plausible mechanistic pathway linking GAL-related findings in this paper to ADIPOR2 can be extracted from the text provided.
- The excerpt is a single summary sentence lacking methodological details (study design, subjects, species, assays, statistical results), so even for GAL-related conclusions the evidence quality cannot be assessed from this text alone.

Given the absence of any mention of ADIPOR2 or genetic/Mendelian evidence, this paper content provides no direct or mechanistic support for the claim and is not relevant.


[Read Paper](https://www.semanticscholar.org/paper/90b66b7c25c2b84a95acb71b98b1181d24487759)


### Myocardial Rev-erb–Mediated Diurnal Metabolic Rhythm and Obesity Paradox

**Why Not Relevant**: The provided paper text does not mention the gene ADIPOR2, adiponectin receptors, or any human Mendelian disease. The study focuses on cardiac-specific Rev-erbα/β knockout in mice, circadian regulation of myocardial metabolism, the obesity paradox in heart failure, and temporal patterns of Rev-erb expression in human dilated cardiomyopathy. There is no direct evidence (e.g., pathogenic ADIPOR2 variants, segregation data, case reports, or human genetic association with a Mendelian phenotype) and no mechanistic discussion linking ADIPOR2 to disease in the excerpt. Because the evidence required to support the claim (that ADIPOR2 is likely associated with at least one Mendelian disease) would need explicit genetic or clinical data implicating ADIPOR2, and none is present here, this paper is not relevant to the claim. Limitations: absence of any mention of ADIPOR2 or related signaling in the text prevents drawing direct or mechanistic inferences; the experimental system is primarily mouse knockout of Rev-erb genes and multi-omics analyses centered on those factors, so findings cannot be repurposed to support ADIPOR2 involvement in Mendelian disease without additional data.


[Read Paper](https://www.semanticscholar.org/paper/6db796e33b100dd7b87b1c39a91cc1ce4d5c2cf9)


### SIGMAR1 mutation associated with autosomal recessive Silver-like syndrome

**Why Not Relevant**: This paper does not provide evidence for or against the claim that ADIPOR2 is likely associated with at least one Mendelian disease. The provided content exclusively describes a study of SIGMAR1 (sigma nonopioid intracellular receptor–1) mutations in a patient with distal hereditary motor neuropathy (dHMN) and lower limb spasticity and a subsequent screen of additional dHMN patients for SIGMAR1 mutations. There is no mention of ADIPOR2, no data on variants in ADIPOR2, and no mechanistic discussion that implicates ADIPOR2 in the described phenotype. Therefore: 

- Direct evidence: None. The study identifies a homozygous missense variant (c.194T>A, p.Leu65Gln) in SIGMAR1 as the probable causative mutation and explicitly frames conclusions around SIGMAR1; no direct association between ADIPOR2 and any Mendelian disease is reported. 

- Mechanistic evidence: None relevant to ADIPOR2. Any mechanistic discussion in the paper (e.g., about SIGMAR1 pathogenicity, evolutionary conservation, in silico analyses, or phenotype similarities) applies to SIGMAR1 and cannot be extrapolated to support ADIPOR2 without additional data linking the two genes or shared pathways. 

- Study scope and limitations relevant to relevance assessment: The investigators used whole-exome sequencing but focused their reporting and follow-up analyses on SIGMAR1; absence of ADIPOR2 in the report does not constitute evidence that ADIPOR2 was examined and found negative. However, because the paper does not present any results, discussion, or mechanism involving ADIPOR2, it cannot be used to support the claim that ADIPOR2 is associated with a Mendelian disease. 

Given the complete lack of mention or data regarding ADIPOR2 in the supplied paper content, relevance to the claim is effectively zero.


[Read Paper](https://www.semanticscholar.org/paper/5a01bb902e68119cb07ea819e75cd17a3419146c)


### Liver Reptin/RUVBL2 controls glucose and lipid metabolism with opposite actions on mTORC1 and mTORC2 signalling

**Why Not Relevant**: This paper describes experiments on the AAA+ ATPase Reptin (Repin) using an inducible liver-specific Reptin knockout mouse model and reports effects on body/fat mass, glycaemia, lipidaemia, hepatic mTOR protein abundance and mTORC1/mTORC2 signalling, and the metabolic consequences in high-fat-diet insulin resistance. It does not mention ADIPOR2, adiponectin receptors, human genetic variation, Mendelian disease, or any clinical genetic case reports. There is therefore no direct evidence in this paper that ADIPOR2 is associated with any Mendelian disease. Mechanistically, although both the paper and ADIPOR2 (an adiponectin receptor involved in metabolic signalling) are related to metabolism, the paper provides no mechanistic data linking ADIPOR2 to disease (no expression, interaction, perturbation, or variant data for ADIPOR2). The study is in a mouse model focused on Reptin and hepatic mTOR signalling, so its findings cannot be used to infer ADIPOR2 Mendelian associations. Key limitations preventing relevance: no mention of ADIPOR2; no human genetics or Mendelian phenotype data; experimental system is a gene knockout in mice for a different gene. Because of these points, the paper provides no support for the claim that ADIPOR2 is likely associated with at least one Mendelian disease.


[Read Paper](https://www.semanticscholar.org/paper/c007d073ae4cfeadc0fa5742fb415d088995ab65)


### Precision Medicine in Internal Medicine

**Why Not Relevant**: This paper is a general review of clinical genetic testing strategies, variant interpretation, and the pace and challenges of gene discovery. It does not mention ADIPOR2, any ADIPOR2 variants, any ADIPOR2-related phenotypes, segregation analyses, functional studies of ADIPOR2, case reports, cohort analyses, or database entries specific to ADIPOR2. Therefore it contains no direct evidence supporting or refuting the claim that ADIPOR2 is likely associated with at least one Mendelian disease. The only potentially relevant material is high-level context: (1) that ES can discover gene-disease associations for genes not previously implicated, and (2) that only a minority of genes were linked to Mendelian disorders as of the cited date. These points are procedural or contextual and cannot be used to conclude that ADIPOR2 is or is not a Mendelian disease gene. To assess the claim about ADIPOR2 specifically would require gene-level evidence (e.g., OMIM/ClinVar/ClinGen entries for ADIPOR2, published case reports with segregating pathogenic variants, functional assays showing pathogenic mechanisms, or burden/variant-enrichment studies), none of which are provided in this paper.


[Read Paper](https://www.semanticscholar.org/paper/c7199e8a5226f8d9d55777da0e15e91797bb459a)


### Sphingosine‐1‐phosphate receptor 2: A novel bile acid receptor and regulator of hepatic lipid metabolism?

**Why Not Relevant**: The provided paper content makes no mention of the gene ADIPOR2 (adiponectin receptor 2), nor does it present genetic association data, clinical case reports, knockout phenotypes, or mechanistic experiments involving ADIPOR2. The manuscript focuses on bile-acid signaling pathways (FXR, TGR5), sphingosine-1-phosphate signaling (S1PR2, SphK2), and downstream epigenetic regulation (HDAC1/2) in hepatic lipid metabolism. Because ADIPOR2 is neither discussed nor experimentally examined, there is no direct evidence (genetic, clinical, or functional) in this paper to support or refute the claim that ADIPOR2 is associated with a Mendelian disease. While both ADIPOR2 and the pathways studied relate broadly to lipid metabolism, mechanistic similarity alone—without specific mention, experiments, or genetic/clinical data concerning ADIPOR2—cannot serve as evidence that ADIPOR2 is linked to any Mendelian condition. Therefore the paper is not relevant to the claim.


[Read Paper](https://www.semanticscholar.org/paper/3e20bf93d9b9428475aef29213e06a6efd112ea2)


### NGS based studies on primary immunodeficiencies (PIDs) : causative gene identification, tool development and application

**Why Not Relevant**: The paper does not mention ADIPOR2 anywhere in the provided content and provides no direct genetic, clinical, or mechanistic data linking ADIPOR2 to a Mendelian disease. The study focuses on primary immunodeficiency disorders (PIDs) and reports implicated genes and candidates within that domain (e.g., LRBA, ATM, DOCK8, PMS2, TNFRSF18, PIK3CG, LILRB1, EPHB2, TXNIP, CD5, NLRP5). Because ADIPOR2 is not named among the genes studied, found, or proposed, this paper provides no direct supportive or refutative evidence regarding the claim that ADIPOR2 is likely associated with at least one Mendelian disease. Important caveats: absence of evidence in this paper is not evidence of absence — ADIPOR2 could be associated with Mendelian disease outside the PID context or in other datasets not covered here. To evaluate the claim further, consult gene-disease databases (OMIM, ClinVar), targeted literature searches (PubMed), and large variant/phenotype resources (gnomAD, DECIPHER, HGMD) for any reported pathogenic variants or clinical case reports implicating ADIPOR2.


[Read Paper](https://www.semanticscholar.org/paper/7786e43fbaf68904111f286747f0cf8968c2dd1e)


### [Clinical and genetic analysis of three children with Menkes disease due to variants of ATP7A gene].

**Why Not Relevant**: The paper exclusively reports clinical and genetic findings for three children with Menkes disease and identifies pathogenic variants in the ATP7A gene (including a c.3294+1G>A splice-site variant and a c.77266650_77267178del CNV affecting ATP7A exons 8–9). Nowhere in the provided content is the gene ADIPOR2 mentioned, evaluated, or discussed; database searches and pathogenicity assessments described were applied to the ATP7A variants. There is no direct genetic evidence (variants, segregation, population frequency, or database annotations) relating ADIPOR2 to Mendelian disease in this paper, nor is there any mechanistic discussion that would plausibly link ATP7A findings to ADIPOR2. Therefore this paper provides no supporting or refuting evidence regarding the claim that ADIPOR2 is associated with at least one Mendelian disease. Limitations: absence of mention in this paper does not prove ADIPOR2 has no Mendelian associations elsewhere; it only shows this particular study does not address ADIPOR2.


[Read Paper](https://www.semanticscholar.org/paper/2c7af41359206fc3b0a3fce382c29206021a28c3)


### ATP10B and the Risk for Early‐Onset Parkinson's Disease

**Why Not Relevant**: The supplied paper text exclusively studies ATP10B and its relationship to early-onset Parkinson's disease. There is no mention of the gene ADIPOR2 (adiponectin receptor 2), no reported variants or phenotypes related to ADIPOR2, and no mechanistic discussion that connects ATP10B to ADIPOR2. Therefore the paper contains neither direct evidence (genetic association, reported pathogenic variants, segregation with Mendelian disease) nor mechanistic evidence (pathway links, functional data) that would support or refute the claim that ADIPOR2 is likely associated with at least one Mendelian disease. Note that absence of any mention of ADIPOR2 in this study does not constitute evidence that ADIPOR2 is not associated with a Mendelian disorder; it simply means this paper cannot inform that claim. To assess the claim about ADIPOR2, one should consult resources and studies that specifically examine ADIPOR2 variants and clinical phenotypes (for example, OMIM, ClinVar, locus-specific literature, or population/genetic studies reporting segregation or loss-of-function variants).


[Read Paper](https://www.semanticscholar.org/paper/3cba22be13141234d6b8d4367347be59fcaa70d6)


### Novel variants of NEK9 associated with neonatal arthrogryposis: Two case reports and a literature review

**Why Not Relevant**: This paper exclusively investigates pathogenic variants in NEK9 and their association with neonatal arthrogryposis and other NEK9-related phenotypes. The provided content (Objective, Methods, Results, Conclusion) repeatedly and specifically names NEK9 as the gene of interest and lists identified NEK9 variants; ADIPOR2 is not mentioned anywhere in the supplied text. Therefore the paper contains no direct evidence (variant associations, case reports, segregation, or statistical association) nor any mechanistic evidence (biochemical function, pathways, animal models) relevant to ADIPOR2 and Mendelian disease. Possible caveat: only the excerpted content was provided; if ADIPOR2 appeared elsewhere in the full paper (very unlikely given the clear NEK9 focus), that was not made available. However, based on the supplied content, relevance to the claim that ADIPOR2 is likely associated with at least one Mendelian disease is effectively zero.


[Read Paper](https://www.semanticscholar.org/paper/a69677b41b5f628b50b661893ae39d1b61fa7955)


### OA30 Genetic analysis of whole exome sequencing in a cohort of children with refractory JIA reveals genetic risk factors for rare juvenile diseases

**Why Not Relevant**: This paper does not provide direct evidence for or against the claim that ADIPOR2 is likely associated with a Mendelian disease. ADIPOR2 is not mentioned anywhere in the provided content; the study reports prioritized variants and illustrative examples in other genes (ADAR, ATP7B, MVK, HLA-DRB1) found in a cohort of 99 children with refractory juvenile idiopathic arthritis. Thus there is no direct association, variant report, mechanistic discussion, or functional data pertaining to ADIPOR2 in this paper.

Relevant limitations to interpreting this negative result: (1) Absence of ADIPOR2 in this study is not evidence that ADIPOR2 has no Mendelian associations generally — the cohort is limited (n=99) and specific to refractory JIA; a Mendelian ADIPOR2 disease could be ultra-rare, tissue-specific, or not represented in this disease cohort. (2) The authors applied allele frequency and in-silico score filters (excluded AF >1%, excluded CADD <15, required REVEL >0.7), which could miss genuine pathogenic variants that fail those thresholds. (3) Exome capture and coverage variability can lead to missing variants in some genes. (4) The authors prioritized genes from specific panels (primary immunodeficiency PanelApp, arthritis-associated genes, paediatric monogenic list), so if ADIPOR2 is not on those lists it might not have been evaluated or highlighted. Therefore, while this paper provides no supporting evidence for an ADIPOR2 Mendelian association, it cannot be used as strong evidence against such an association.


[Read Paper](https://www.semanticscholar.org/paper/5ceb90bf884c2beab8d50226ab822bc8d21c2783)


### The UK10K project identifies rare variants in health and disease

**Why Not Relevant**: The provided paper content does not mention the gene ADIPOR2 or report any variant- or gene-level associations involving ADIPOR2. The excerpts show that the UK10K project and its exome arm were well suited to discover Mendelian-disease genes generally (e.g., by detecting homozygous and compound heterozygous predicted loss-of-function variants and associating some with phenotypic outliers), but there is no direct evidence (no variants, statistical associations, or mechanistic data) linking ADIPOR2 to any Mendelian disease in the text supplied. Because the claim is specifically about ADIPOR2, these general methodological and resource-level statements are only weak, indirect support for plausibility and do not constitute evidence that ADIPOR2 is associated with a Mendelian disease.


[Read Paper](https://www.semanticscholar.org/paper/439b20628573147c514d93e0feb5140812cf6f34)


### Contribution of rare transmitted and de novo variants among 2 , 871 1 congenital heart disease probands

**Why Not Relevant**: The supplied paper content does not mention ADIPOR2 anywhere in the provided text. The study is focused on congenital heart disease (CHD) and reports aggregated analyses of de novo and transmitted damaging variants across curated CHD gene sets, but the provided gene lists and tables (e.g., curated sets of 212 human CHD genes, 61 mouse CHD genes, and summary counts of damaging recessive genotypes in known CHD genes) do not include ADIPOR2 in the excerpts shown. Therefore there is no direct evidence in this excerpt that ADIPOR2 is associated with any Mendelian disease. Mechanistic evidence is also absent: the paper discusses general categories of variants (LoF, damaging missense), statistical enrichment approaches, and results for CHD-related genes, but it provides no molecular, functional, or variant-level data concerning ADIPOR2. Limitations: (1) Only a portion of the paper was provided; it is possible (though unlikely given the CHD focus and curated gene lists) that ADIPOR2 appears elsewhere in the full paper or supplementary materials. (2) Even if ADIPOR2 had been sequenced in this cohort, absence of mention here means no reported significant association in this study. Recommended next steps to evaluate the claim: search the full text and supplementary files of this paper for "ADIPOR2"; search databases that curate Mendelian gene–disease relationships (OMIM, ClinVar, ClinGen) and population/trait association catalogs for ADIPOR2; and review functional studies of ADIPOR2 for mechanistic links to Mendelian phenotypes.


[Read Paper](https://www.semanticscholar.org/paper/e5830d9e95469f22518560007746202e62db26a0)


### SUN-653 Bypassing Skeletal Muscle Lipid Handling Deficiencies as a Therapy for Metabolic Disease

**Why Not Relevant**: The provided paper content (abstract) focuses entirely on KLF15 function in skeletal muscle, mouse metabolic phenotypes (obesity, insulin resistance, NAFLD), defects in muscle lipid handling (including changes in carnitine shuttle gene expression, long-chain acylcarnitines, Cd36 expression), and the effect of a short-chain fatty acid (SCFA)-rich diet. It does not mention ADIPOR2 (or 'adiponectin receptor 2') anywhere in the provided text, does not present any human genetic or Mendelian-disease data, and does not discuss associations between specific genes and Mendelian disorders. Therefore there is neither direct evidence (genetic association, pathogenic variants, clinical case reports) nor mechanistic evidence in this content that links ADIPOR2 to any Mendelian disease. Limitations: only the abstract was provided; it is possible (though unlikely based on the abstract focus) that the full paper might mention ADIPOR2 or relevant human genetics in other sections. To properly rule out any connection from this paper one would need to search the full text for 'ADIPOR2', 'adiponectin receptor 2', or human genetic/Mendelian disease terms. Based solely on the supplied content, relevance to the claim is nil.


[Read Paper](https://www.semanticscholar.org/paper/06a20fe12cf81d686afd98ce94821bc4dd888e2a)


## Search Queries Used

- ADIPOR2 adiponectin receptor 2 pathogenic variant Mendelian disease case report family segregation whole exome sequencing

- ADIPOR2 rare variant loss of function homozygous compound heterozygous monogenic disorder rare disease cohort exome sequencing burden analysis

- ADIPOR2 knockout mouse phenotype metabolic disease insulin resistance lipid metabolism ceramidase activity functional study human phenotype

- ADIPOR2 ClinVar OMIM gnomAD HGMD ClinGen gene disease validity pathogenic variant review


## Bibliometric Configuration

- Use Bibliometrics: True

- Author Impact Weight: 0.4

- Citation Impact Weight: 0.4

- Venue Impact Weight: 0.2
