## Supplementary material for "MENDELSEEK: An algorithm that predicts Mendelian Genes and elucidates what makes them special": AFDN.txt

# Claim: The gene AFDN is likely associated with at least one Mendelian disease.

**Status**: processed

**Overall Rating**: -1

**Explanation**:

No relevant papers were found that support or refute this claim.


## Relevant Papers


## Other Reviewed Papers


### Exome/Genome Sequencing in Undiagnosed Syndromes

**Why Not Relevant**: The provided paper content is a high-level overview of next-generation sequencing (NGS) applications (ES/GS) to undiagnosed rare/ultrarare Mendelian diseases and the technological and clinical challenges/opportunities in the field. It contains no mention of the gene AFDN, no case reports, no cohort analyses, no variant-level data, no functional experiments, and no statements linking AFDN to any Mendelian phenotype. Therefore it provides neither direct evidence (e.g., patient-level genotype–phenotype co-segregation, case series, statistical association) nor mechanistic evidence (e.g., molecular/biochemical pathway data, animal models, functional assays) that would support or refute the claim that AFDN is likely associated with at least one Mendelian disease.

Specific reasons: 
- No mention of the gene name “AFDN” (or synonyms) appears in the provided text. 
- The content is descriptive of technologies and broad challenges/opportunities; it does not present experimental methods, results, or conclusions about any particular gene. 
- There are no verbatim sentences or paragraphs in the supplied content that provide gene-level, variant-level, clinical, or mechanistic evidence relevant to AFDN. 

Because of these absences, relevance to the claim is effectively zero based on the supplied excerpt. To evaluate the claim about AFDN, one would need gene-specific sources such as: clinical case reports or series linking AFDN variants to a phenotype with segregation data, population-genetics evidence (e.g., constraint metrics, case–control burden), functional studies demonstrating variant impact on protein function or cellular pathways, animal models recapitulating disease features, or authoritative database entries (OMIM, ClinVar) noting pathogenic variants in AFDN. None of these are present in the provided text.


[Read Paper](https://www.semanticscholar.org/paper/2d9790fd8ef924917e3cd41d133ee99fe80fa0b1)


### Rare genetic disorders: Beyond whole‐exome sequencing

**Why Not Relevant**: The provided paper content is a general review of next-generation sequencing technologies (WES/WGS) and their role in diagnosing rare genetic disorders. It does not mention the gene AFDN, any specific genes, any variants, patient cases, association studies, linkage analyses, segregation data, functional experiments, or clinical reports that would support or refute an association between AFDN and a Mendelian disease. While the text states that WGS can reveal disease-causing variations and mechanisms in some cases, that is a methodological and contextual statement about technology utility, not evidence about any particular gene. Therefore there is no direct evidence (no reported variants, no statistical associations, no clinical cases) and no mechanistic evidence (no molecular/functional data implicating AFDN) in the excerpt. Limitations relevant to the claim: (1) absence of AFDN mention means no gene-level data are available here; (2) the paper's scope is technology and diagnostic yield, not gene discovery for AFDN; (3) general statements about WGS/WES applicability cannot be extrapolated to conclude that any specific gene—including AFDN—is likely associated with a Mendelian disease without supporting gene-specific data. Because of these points, the content is not relevant to assessing the claim that AFDN is likely associated with at least one Mendelian disease.


[Read Paper](https://www.semanticscholar.org/paper/bd477d33a409ab5609c86cac0512c51ce998818a)


### Diagnosing missed cases of spinal muscular atrophy in genome, exome, and panel sequencing datasets

**Why Not Relevant**: The paper's content exclusively addresses spinal muscular atrophy (SMA), the SMN1 and SMN2 genes, and the development and validation of a diagnostic tool (SMA Finder) for detecting SMN1/SMN2-related molecular causes of SMA from sequencing data. It contains no mention of the gene AFDN, no data on AFDN sequence variants, no genetic linkage or segregation analyses, no case reports implicating AFDN in Mendelian disease, and no functional/mechanistic experiments involving AFDN. Thus there is neither direct evidence (e.g., pathogenic AFDN variants observed in affected individuals) nor mechanistic evidence (e.g., biological pathways, protein function, or model organism phenotypes implicating AFDN in disease) relevant to the claim. While the paper demonstrates that exome/genome/targeted sequencing and analytic pipelines can detect disease-causing variants in genes located in segmental duplications (as with SMN1), that methodological point does not provide evidence that AFDN is associated with any Mendelian disease—only that similar technical approaches could, in principle, be used to study other genes. Therefore the paper does not support or refute the claim that AFDN is likely associated with at least one Mendelian disease.


[Read Paper](https://www.semanticscholar.org/paper/1cdee5f04f0788789a3f8b0cf937b1f62156e200)


### Exome sequencing identifies HELB as a novel susceptibility gene for non-mucinous, non-high-grade-serous epithelial ovarian cancer

**Why Not Relevant**: The paper is a genome-wide exome study of rare loss-of-function variants in relation to epithelial ovarian cancer risk. The study reports specific genes (BRCA1, BRCA2, BRIP1, RAD51C, RAD51D, MSH6, PALB2, HELB, OR2T35, NBN, MYO1A) associated with ovarian cancer in this data set and provides mechanistic linkage for HELB. The gene AFDN is not mentioned anywhere in the provided text. Therefore the paper provides no direct evidence that AFDN is associated with any Mendelian disease, nor any mechanistic data implicating AFDN. However, absence of mention in this ovarian-cancer-focused dataset is not definitive evidence that AFDN has no Mendelian-disease associations: the study tests only rare LOF variants (MAF < 0.1%) for association with ovarian cancer, so it would not detect (a) associations mediated by missense or regulatory variants in AFDN, (b) associations with phenotypes other than ovarian cancer, or (c) associations for which the study lacked variant observations or power. Given these points, the relevance of this paper to the claim about AFDN is very low.


[Read Paper](https://www.semanticscholar.org/paper/3ff2757719c7a067b7206590903a72ee90de73b7)


## Search Queries Used

- AFDN afadin AF6 Mendelian disease ClinVar OMIM case report pathogenic variant AFDN mutation monogenic disorder

- AFDN afadin AF6 missense nonsense frameshift loss of function de novo variant congenital developmental disorder intellectual disability congenital heart disease phenotype

- afadin AFDN adherens junction nectin Rap1 actin cytoskeleton cardiac development neural development knockout mouse zebrafish phenotype developmental defects

- AFDN rare variant burden exome sequencing genome sequencing gene discovery Mendelian gene panel unsolved rare disease cohort


## Bibliometric Configuration

- Use Bibliometrics: True

- Author Impact Weight: 0.4

- Citation Impact Weight: 0.4

- Venue Impact Weight: 0.2
