## Supplementary material for "MENDELSEEK: An algorithm that predicts Mendelian Genes and elucidates what makes them special": AGPAT1.txt

# Claim: The gene AGPAT1 is likely associated with at least one Mendelian disease.

**Status**: processed

**Overall Rating**: 3

**Explanation**:

Supporting evidence.
The provided paper shows that genetic loss of Agpat1 in mice produces a spectrum of severe, pleiotropic phenotypes: perinatal lethality for many homozygotes, low body weight and hypoglycemia, reduced adipose percentage (but not frank generalized lipodystrophy), impaired spermatogenesis and gonadal hormone abnormalities, abnormal hippocampal neuron development and audiogenic seizures. The manuscript also emphasizes the biochemical role of AGPAT enzymes in converting lysophosphatidic acid to phosphatidic acid, a precursor for major phospholipids and diacylglycerol, and notes that AGPAT1 and AGPAT2 are highly homologous and co-expressed in adipocytes. Importantly, the paper recalls that genetic defects in AGPAT2 cause congenital generalized lipodystrophy in humans, demonstrating that perturbation of this enzyme family is already established to cause a Mendelian disease. Together, the severe, multi-system mouse knockout phenotype combined with (1) the central biochemical role of AGPAT1 in phospholipid triacylglycerol biosynthesis and (2) precedent for AGPAT family members causing Mendelian disease, provide substantive biological plausibility that pathogenic AGPAT1 variants could underlie a human Mendelian disorder (likely affecting metabolism, reproduction, or neurologic development).

Caveats or contradictory evidence.
The excerpts do not provide any direct human genetic evidence linking AGPAT1 variants to a Mendelian disease. Mouse phenotypes do not always translate to observed human Mendelian disorders: (a) some genes when disrupted in mouse are embryonic/perinatal lethal and therefore may be underrepresented in human clinical sequencing cohorts, (b) human redundancy or compensatory pathways may differ from mouse, and (c) pathogenic mechanisms may differ (e.g., haploinsufficiency vs recessive loss of function). The paper also notes that Agpat1-/- mice do not show generalized lipodystrophy despite decreased fat percentage, indicating non-identical phenotypes between Agpat1 loss and AGPAT2-associated lipodystrophy. Thus, while the mouse data are compelling for biological importance, they do not establish that AGPAT1 is observed as a Mendelian disease gene in humans. Without direct human variant-disease co-segregation, population variant enrichment, or clinical case reports, the claim that AGPAT1 is "likely associated" with at least one Mendelian disease remains inferential rather than demonstrated.

Analysis of potential underlying mechanisms, deeper relationships, or second-order implications.
AGPAT1 catalyzes a key acyltransferase step in phosphatidic acid production; disruption will alter membrane phospholipid composition and signaling lipid pools (phosphatidic acid and downstream diacylglycerol), which plausibly affects multiple organ systems sensitive to membrane composition and lipid signaling (adipocytes, liver glucose metabolism, neurons, and germ cells). Given AGPAT2 already causes congenital generalized lipodystrophy when mutated, a reasonable mechanistic hypothesis is that AGPAT1 loss-of-function could produce a recessive metabolic syndrome with neurodevelopmental and reproductive manifestations, or alternatively embryonic/perinatal lethality if compensation is insufficient in humans. Modes of pathogenicity could include autosomal recessive loss-of-function (most likely, given mouse knockout data), dominant-negative effects, or hypomorphic alleles causing a milder, later-onset phenotype. The lack of compensation by AGPAT1 for AGPAT2 deficiency (and vice versa) in adipocytes suggests non-redundant roles that could yield distinct human phenotypes depending on which paralog is mutated.

Assessment and recommended rating.
The evidence supports strong biological plausibility: a critical biochemical role, severe multi-organ mouse knockout phenotypes, and a paralog (AGPAT2) already established as a Mendelian disease gene. However, the available evidence lacks any direct human genetic association (no case reports, no segregation data, no population genetics signals provided). Because the claim is that AGPAT1 is "likely associated" with at least one Mendelian disease, I weigh the absence of direct human data heavily. Plausibility is high, but absence of human evidence prevents concluding that an association is likely rather than merely possible. Therefore the best characterization given the provided information is that evidence is mixed: there is strong functional support but no human genetic corroboration.


**Final Reasoning**:

I reviewed the logic and evidence: (1) functional mouse knockout data are strong and show multi-system disease phenotypes; (2) the biochemical role of AGPAT1 and its paralogy to AGPAT2 (a known Mendelian gene) further increase plausibility; but (3) there is no presented human genetic evidence linking AGPAT1 variants to a Mendelian disorder. The claim wording—"is likely associated with at least one Mendelian disease"—implies a greater-than-even chance of an existing human Mendelian association. Given the lack of direct human data, I cannot justify a "Likely True" or stronger rating. Conversely, the strong functional evidence argues against ruling the claim false. The correct balance is therefore "Mixed Evidence": supportive functional and paralogous-disease evidence exists, but no direct human genetic confirmation is provided.


## Relevant Papers


### Metabolic, Reproductive, and Neurologic Abnormalities in Agpat1-Null Mice

**Authors**: Anil K. Agarwal (H-index: 2), A. Garg (H-index: 79)

**Relevance**: 0.8

**Bibliometric Impact**: 0.4656


**Excerpts**:

- Defects in the biosynthesis of phospholipids and neutral lipids are associated with cell membrane dysfunction, disrupted energy metabolism, and diseases including lipodystrophy.

- In these pathways, the 1-acylglycerol-3-phosphate O-acyltransferase (AGPAT) enzymes transfer a fatty acid to the sn-2 carbon of sn-1-acylglycerol-3-phosphate (lysophosphatidic acid) to form sn-1, 2-acylglycerol-3-phosphate [phosphatidic acid (PA)]. PA is a precursor for key phospholipids and diacylglycerol.

- AGPAT1 and AGPAT2 are highly homologous isoenzymes that are both expressed in adipocytes.

- Genetic defects in AGPAT2 cause congenital generalized lipodystrophy, indicating that AGPAT1 cannot compensate for loss of AGPAT2 in adipocytes.

- To further explore the physiology of AGPAT1, we characterized a loss-of-function mouse model (Agpat1-/-).

- The majority of Agpat1-/- mice died before weaning and had low body weight and low plasma glucose levels, independent of plasma insulin and glucagon levels, with reduced percentage of body fat but not generalized lipodystrophy.

- These mice also had decreased hepatic messenger RNA expression of Igf-1 and Foxo1, suggesting a decrease in gluconeogenesis.

- In male mice, sperm development was impaired, with a late meiotic arrest near the onset of round spermatid production, and gonadotropins were elevated.

- Female mice showed oligoanovulation yet retained responsiveness to gonadotropins.

- Agpat1-/- mice also demonstrated abnormal hippocampal neuron development and developed audiogenic seizures.

- In summary, Agpat1-/- mice developed widespread disturbances of metabolism, sperm development, and neurologic function resulting from disrupted phospholipid homeostasis.

- AGPAT1 appears to serve important functions in the physiology of multiple organ systems.

- The Agpat1-deficient mouse provides an important model in which to study the contribution of phospholipid and triacylglycerol synthesis to physiology and diseases.


**Explanations**:

- This sentence links defects in phospholipid/neutral lipid biosynthesis to human disease (lipodystrophy), establishing clinical relevance of the pathway in which AGPAT1 functions. This is mechanistic/contextual evidence that perturbing the pathway can produce Mendelian-like disease phenotypes, but it does not implicate AGPAT1 specifically in a human Mendelian disorder.

- This describes the biochemical function of AGPAT enzymes (including AGPAT1) in producing phosphatidic acid (PA), a precursor for key lipids. Mechanistic evidence: it explains how loss of AGPAT1 activity could perturb lipid biosynthesis and thereby cause cellular and organismal dysfunction. Strength: defines a clear molecular role. Limitation: enzyme function alone does not prove that human AGPAT1 variants cause Mendelian disease.

- This statement indicates that AGPAT1 and AGPAT2 are closely related and co-expressed in adipocytes. Mechanistic relevance: similarity to AGPAT2 (which is known to cause disease when mutated) increases plausibility that AGPAT1 perturbation could have disease consequences. Limitation: expression/homology do not equal causation; redundancy or tissue-specific roles may differ between isoenzymes and species.

- This is an important translational sentence: it provides direct human genetic evidence that mutations in the closely related gene AGPAT2 cause a Mendelian disease (congenital generalized lipodystrophy). It also states that AGPAT1 cannot compensate for AGPAT2 loss in adipocytes, implying non-redundant roles in at least some tissues. This strengthens plausibility that perturbation of an AGPAT family member can cause Mendelian disease. Limitation: AGPAT2 causation does not prove AGPAT1 causation; the sentence actually emphasizes AGPAT1's inability to replace AGPAT2 in adipocytes, which could mean AGPAT1 loss has different phenotypic consequences.

- This describes the experimental approach: generation and characterization of an Agpat1 loss-of-function (knockout) mouse. Relevance: provides a controlled in vivo system to observe phenotypic consequences of AGPAT1 deficiency. This is experimental (indirect) evidence relevant to assessing whether AGPAT1 loss can produce monogenic-like phenotypes. Limitation: mouse loss-of-function phenotypes are not direct proof of a human Mendelian disease.

- This reports major organismal phenotypes in Agpat1-/- mice (perinatal lethality, low weight, hypoglycemia, reduced fat but not generalized lipodystrophy). This is direct experimental evidence that AGPAT1 loss produces severe, multi-system phenotypes in mice. It supports the plausibility that AGPAT1 loss-of-function could cause a Mendelian disease in humans. Limitations: species differences, absence of corresponding human cases in this paper, and the note that generalized lipodystrophy was not observed (so phenotype differs from AGPAT2-associated disease). Also, statistical details (n, significance) are not present in the excerpt.

- This sentence links AGPAT1 deficiency to decreased hepatic Igf-1 and Foxo1 mRNA, suggesting reduced gluconeogenesis. This is mechanistic evidence connecting AGPAT1 loss to specific metabolic pathways that can account for hypoglycemia. Strength: provides a plausible biochemical/physiological pathway from gene loss to phenotype. Limitation: mRNA changes are correlative; causality and relevance to human physiology require further validation.

- This documents a specific reproductive phenotype (male infertility due to late meiotic arrest and elevated gonadotropins). This is experimental phenotype evidence indicating that AGPAT1 loss disrupts sperm development, strengthening the case that AGPAT1 has essential, non-redundant functions in specific tissues. Limitation: again, this is in mice and may not directly translate to human Mendelian disease without human genetic corroboration.

- This documents a female reproductive phenotype (oligoanovulation with preserved gonadotropin responsiveness). This broadens the organ systems affected by AGPAT1 deficiency and supports pleiotropic effects consistent with a monogenic syndrome. Limitation: mouse-specific data; details about penetrance and variability are not provided here.

- This reports neurologic phenotypes (abnormal hippocampal neuron development and audiogenic seizures) in knockout mice. Mechanistic relevance: links disrupted phospholipid homeostasis to neural development and function, offering a path by which AGPAT1 variants could cause neurologic Mendelian phenotypes. Limitation: audiogenic seizures in mice may not correspond to a human clinical syndrome, and the data remain preclinical.

- This summary sentence explicitly ties the observed multi-system phenotypes to disrupted phospholipid homeostasis caused by Agpat1 knockout. It provides mechanistic synthesis that AGPAT1 deficiency has pleiotropic effects across organ systems. Strength: concise mechanistic interpretation tying gene loss to physiological outcomes. Limitation: synthesis based on mouse model; human relevance is inferred but not demonstrated.

- This interpretive sentence highlights that AGPAT1 has important physiological roles across organ systems. Relevance: supports the biological plausibility that AGPAT1 variants could produce Mendelian disease phenotypes. Limitation: statement is inferential and not direct human genetic evidence.

- This concluding sentence positions the Agpat1-deficient mouse as a model to study contributions of phospholipid and triacylglycerol synthesis to physiology and disease. Relevance: underscores that AGPAT1 loss produces a definable syndrome in mice, making it a candidate gene for Mendelian disease discovery in humans. Limitation: the paper presents animal-model evidence; human genetic association/segregation data would be required to establish AGPAT1 as causative for a Mendelian disease.


[Read Paper](https://www.semanticscholar.org/paper/e39eaa8843eb74b6ff405a0d91a434acddc2885f)


## Other Reviewed Papers


### Enzymes of Lipid Metabolism II

**Why Not Relevant**: The provided paper content focuses on the mechanism of interaction of pancreatic phospholipase A2 with negatively charged substrates and properties of membrane-bound phospholipases A2. It contains no mention of AGPAT1 (1-acylglycerol-3-phosphate O-acyltransferase 1), no genetic or clinical data, no reported variants, and no discussion of Mendelian phenotypes. AGPAT1 is an acyltransferase (involved in the biosynthesis of phosphatidic acid), which is a distinct enzymatic activity from phospholipase A2 (which hydrolyzes fatty acyl chains). Thus (a) there is no direct evidence in this paper that AGPAT1 is associated with any Mendelian disease, and (b) there is no mechanistic pathway described that plausibly links the PLA2 substrate-interaction findings in this paper to pathogenic variation in AGPAT1. Without genetic association data, clinical case reports, functional studies specifically on AGPAT1 variants, or mechanistic experiments connecting PLA2 activity to AGPAT1 dysfunction and disease phenotypes, one cannot infer that AGPAT1 is likely associated with a Mendelian disease from this paper. Therefore the paper is not relevant to the claim.


[Read Paper](https://www.semanticscholar.org/paper/cc49b10be6a1411a1c6c30b7f1de1ecf4b22e17d)


### The Burden of Candidate Pathogenic Variants for Kidney and Genitourinary Disorders Emerging From Exome Sequencing

**Why Not Relevant**: The provided paper excerpt describes methods and general findings about identifying and curating genes associated with Mendelian kidney and genitourinary disorders, and it discusses database-driven and population-frequency considerations for variant pathogenicity. However, the excerpt does not mention AGPAT1 by name, nor does it provide the curated gene list or per-gene results that would allow confirmation that AGPAT1 is associated with a Mendelian disease. Because there is no direct statement, data, or mechanistic discussion in the supplied text specifically about AGPAT1, the paper content as provided cannot be used as evidence for or against the claim that AGPAT1 is likely associated with at least one Mendelian disease. The only potential indirect relevance is that AGPAT1 could be included among the 625 curated genes, but that cannot be ascertained from the material supplied here; therefore the relevance score is very low.


[Read Paper](https://www.semanticscholar.org/paper/a75e297208c04992568e607d19aa9b9aaebd09d5)


### Prevalence and phenotypic impact of rare potentially damaging variants in autism spectrum disorder

**Why Not Relevant**: The provided paper content is a single high-level summary sentence about the overall yield of potentially damaging variation in an ASD epidemiological sample and does not mention AGPAT1, any individual gene, specific variants, Mendelian disease associations, segregation, functional studies, or mechanistic pathways. The claim concerns whether AGPAT1 is likely associated with at least one Mendelian disease, which requires gene-specific evidence such as documented pathogenic variants, reported disease associations (e.g., OMIM entries), segregation in families, case reports, population-level enrichment analyses for damaging alleles, or functional/mechanistic data linking AGPAT1 perturbation to a Mendelian phenotype. None of these types of evidence are present in the supplied content. While the sentence implies that high-throughput sequencing finds many potentially damaging variants in ASD cohorts, that general statement cannot be mapped to AGPAT1 specifically; AGPAT1 could be absent from the cohort, present with benign variation, or present but not causative. Therefore the content is not relevant to assessing the claim and provides no direct or mechanistic support or refutation.


[Read Paper](https://www.semanticscholar.org/paper/fa3391717bcabc5b0a0eb7766e6489d8da5069b4)


### Expanding the phenotype of a recurrent de novo variant in PACS1 causing intellectual disability

**Why Not Relevant**: This paper exclusively reports and discusses a recurrent de novo missense variant in PACS1 and the resulting Mendelian syndrome MRD17. AGPAT1 is not mentioned anywhere in the provided content; there are no genetic, mechanistic, epidemiological, or experimental data related to AGPAT1. Therefore the paper provides no direct or mechanistic evidence to support or refute the claim that AGPAT1 is likely associated with at least one Mendelian disease. Absence of mention of AGPAT1 in this PACS1-focused study does not constitute evidence against AGPAT1's involvement in Mendelian disease; it only means this paper is irrelevant to the AGPAT1 claim.


[Read Paper](https://www.semanticscholar.org/paper/e460d89d850693b4fb1e3df12702be0dfb8d7f5b)


### A single center experience with publicly funded clinical exome sequencing for neurodevelopmental disorders or multiple congenital anomalies

**Why Not Relevant**: The provided paper content consists of a single summary sentence about the general utility of clinical exome sequencing in a diagnostic setting for GDD/ID and/or MCA. It contains no mention of the gene AGPAT1, no reported variants, no disease names linked to specific genes, and no mechanistic information that could connect AGPAT1 to a Mendelian disease. Because the content lacks any gene-specific data, case reports, cohort analyses, segregation studies, functional assays, or mechanistic descriptions, it cannot support or refute the claim that AGPAT1 is likely associated with at least one Mendelian disease. To evaluate that claim, primary data such as case reports with pathogenic AGPAT1 variants, cohort enrichment analyses, segregation in families, or functional studies demonstrating a disease mechanism for AGPAT1 would be required. None of those data are present in the excerpt provided.


[Read Paper](https://www.semanticscholar.org/paper/8ec4c0c03c573ad4019a6ee63b9197179a32b173)


### Precision Medicine in Internal Medicine

**Why Not Relevant**: The provided paper is a general review of genetic testing strategies, variant interpretation, and the state of gene–disease discovery as of the date cited. It does not mention AGPAT1, report variants in AGPAT1, present segregation or functional data involving AGPAT1, or cite any clinical cases linking AGPAT1 to a Mendelian phenotype. Therefore it contains no direct evidence supporting or refuting the claim that AGPAT1 is likely associated with at least one Mendelian disease. The included excerpts are general background that bear on the broader plausibility of discovering new Mendelian genes via exome sequencing, but they are insufficient to draw any conclusion about AGPAT1 specifically. To evaluate the claim about AGPAT1 would require gene-specific evidence (case reports, ClinVar/OMIM/HGMD entries, segregation analyses, functional studies, or cohort association data), none of which are present in this paper. Additionally, the OMIM statistic is dated (1 February 2019) and the landscape of gene–disease associations may have changed since then; however, even if updated, that kind of high-level statistic would still not constitute direct evidence for AGPAT1.


[Read Paper](https://www.semanticscholar.org/paper/c7199e8a5226f8d9d55777da0e15e91797bb459a)


### From case studies to community knowledge base: MSeqDR provides a platform for the curation and genomic analysis of mitochondrial diseases

**Why Not Relevant**: This paper does not provide evidence for or against the claim that AGPAT1 is likely associated with at least one Mendelian disease. The manuscript is an overview/editorial describing mitochondrial disease heterogeneity and introducing the MSeqDR community resource; it contains no mention of the gene AGPAT1, no case reports, no variant data, no functional assays, and no mechanistic discussion linking AGPAT1 to human disease. Although the paper states that MSeqDR curates >1360 nuclear and mitochondrial genes and synchronizes with ClinVar (which could, in principle, contain entries for many genes), the article gives no gene-level listings or specific variant–phenotype associations that would allow assessment of AGPAT1. Therefore, based on the content provided, there is no direct evidence (patient, genetic, statistical) nor mechanistic evidence (biochemical pathway, functional assay) in this paper to support or refute the claim. To evaluate the claim about AGPAT1, primary literature reporting patient cases, curated database entries listing AGPAT1 pathogenic variants, or functional studies of AGPAT1 would be required.


[Read Paper](https://www.semanticscholar.org/paper/6f85c595adaca6963c30fea73bc37266fa78be56)


### Whole-Exome Sequencing in Adults With Chronic Kidney Disease

**Why Not Relevant**: The provided paper excerpt does not mention AGPAT1 by name nor present any data, results, or discussion specifically about AGPAT1. The text describes study design, sequencing methods, gene-list curation, variant interpretation procedures, and general limitations of exome sequencing, which are relevant to how Mendelian disease associations would be discovered and validated but do not provide direct or mechanistic evidence linking AGPAT1 to any Mendelian disorder. Because no AGPAT1-specific evidence (presence/absence on the curated lists, variants detected, functional studies, or clinical correlations) appears in the supplied content, the paper as presented cannot be used to support or refute the claim that AGPAT1 is likely associated with at least one Mendelian disease. Note also that exome capture blind spots and lack of disclosure of the specific 287-gene list in this excerpt mean that even an absence of AGPAT1 in reported results here would be inconclusive: (a) AGPAT1 might not have been included in the curated gene list, (b) it might have been included but not reported in this excerpt, or (c) it might have had regions poorly captured by the exome kit and thus been missed. For these reasons the relevance score is very low.


[Read Paper](https://www.semanticscholar.org/paper/a29bd57964f68ac5f8e1abcafe1112ba488f53f5)


### The simplest explanation does not have to be preferred: co-occurrence of pathogenic variants in cancer-predisposing genes

**Why Not Relevant**: The provided text does not mention AGPAT1, Mendelian disease, or any gene-specific findings. It is a single sentence fragment describing that the paper reviews the frequency and cancer phenotypes associated with "MINAS" using recent literature and UK 10,000 Genomes Project data from participants with tumor predisposition syndromes or multiple primary tumors, and assesses whether effects of cancer susceptibility genes (CSGs) are additive or synergistic. This content (1) contains no reference to AGPAT1, (2) contains no direct genetic evidence (e.g., pathogenic variants, segregation in families, case reports) linking any gene to a Mendelian disorder, (3) provides no mechanistic data (e.g., functional assays, molecular pathways) that would support plausibility for AGPAT1 causing a Mendelian disease, and (4) includes no methods, results, statistics, or conclusions about gene-disease associations. Therefore the excerpted content provides neither direct nor mechanistic evidence for or against the claim that AGPAT1 is likely associated with at least one Mendelian disease. 

Limitations and caveats: (a) only a single sentence (fragment) of the paper was provided, so it is possible that other parts of the full paper discuss AGPAT1 or relevant evidence; that material was not available for review. (b) The sentence is general and focused on cancer predisposition and interactions among cancer susceptibility genes (CSGs), which is a different question than establishing that a specific gene (AGPAT1) causes a Mendelian disorder. To evaluate the claim properly would require full-text search for AGPAT1, any reported pathogenic variants, segregation/penetrance data, functional studies, or citations to other Mendelian-disease-focused literature, none of which are present in the provided content.


[Read Paper](https://www.semanticscholar.org/paper/a1372a664bd2c33a0bda638975a8b26d112457fc)


### Identification of novel disease-causing variants in rare diseases using trio exome sequencing

**Why Not Relevant**: The paper does not mention AGPAT1 anywhere in the provided content and thus contains no direct genetic, clinical, or mechanistic evidence linking AGPAT1 to a Mendelian disease. The study reports findings for several other genes discovered by trio exome sequencing in patients with specific congenital phenotypes, but AGPAT1 is absent from the reported results and discussion. Important limitations: (1) The absence of AGPAT1 in this small, phenotype-selected cohort does not disprove an association in other cohorts or phenotypes. (2) Exome sequencing can miss certain variant types and may have coverage gaps, so a non-detection here is not definitive. (3) The paper’s focus is on a limited set of clinical presentations (startle disease, hypomyelinating leukodystrophy, microcephaly/pontocerebellar hypoplasia, syndromic intellectual disability, developmental delay/hypotonia), so it would not detect AGPAT1 associations that cause different phenotypes. For these reasons, relevance to the claim is minimal (relevance score 0.02). Recommended next steps: search databases and literature specifically for AGPAT1 (OMIM, ClinVar, PubMed) or papers reporting functional/mechanistic evidence linking AGPAT1 to human Mendelian phenotypes.


[Read Paper](https://www.semanticscholar.org/paper/8ae081628e0ab2aec44f8fe9990f230956fb40ba)


## Search Queries Used

- AGPAT1 mutation human Mendelian disease monogenic disorder AGPAT1 pathogenic variant case report ClinVar OMIM

- AGPAT1 rare variants exome sequencing cohort trio sequencing developmental delay intellectual disability epilepsy congenital anomalies AGPAT1 de novo variants

- AGPAT1 1 acylglycerol 3 phosphate O acyltransferase 1 function phosphatidic acid biosynthesis lipid metabolism membrane biogenesis mechanism disease model

- AGPAT1 knockout mouse phenotype zebrafish morpholino loss of function phenotype embryonic lethal AGPAT1 null gnomAD loss of function constraint pLI gene constraint


## Usage Statistics

- Prompt Tokens: 0

- Completion Tokens: 0

- Total Tokens: 0

- Estimated Cost: $0.0533


## Bibliometric Configuration

- Use Bibliometrics: True

- Author Impact Weight: 0.4

- Citation Impact Weight: 0.4

- Venue Impact Weight: 0.2
