## Supplementary material for "MENDELSEEK: An algorithm that predicts Mendelian Genes and elucidates what makes them special": AKR1B10.txt

# Claim: The gene PTK2B is likely associated with at least one Mendelian disease.

**Status**: processed

**Overall Rating**: -1

**Explanation**:

No relevant papers were found that support or refute this claim.


## Relevant Papers


## Other Reviewed Papers


### Precision Medicine in Internal Medicine

**Why Not Relevant**: The provided paper content is a general review of genetic testing methods, the capabilities and limits of exome/genome sequencing, and the overall landscape of gene–disease discovery. It contains no mention of the gene PTK2B, no case reports, no variant-level data, no ClinVar/OMIM/OMIA entries specific to PTK2B, no segregation analyses, and no functional experiments involving PTK2B. Therefore it provides no direct evidence for or against the claim that PTK2B is likely associated with at least one Mendelian disease. The content does offer mechanistic context (how ES/GS can discover novel gene–disease links and the interpretive challenges), which increases general plausibility that new gene–disease relationships can be found, but such context is insufficient to support the specific claim about PTK2B. To evaluate the claim about PTK2B, one would need PTK2B-specific data such as: published case series linking PTK2B variants to a consistent phenotype, segregation evidence in families, de novo enrichment statistics, ClinVar/OMIM entries, experimental functional validation demonstrating pathogenic mechanisms, or population-based variant burden analyses.


[Read Paper](https://www.semanticscholar.org/paper/c7199e8a5226f8d9d55777da0e15e91797bb459a)


### Analysis of De Novo Mutations in Sporadic Cardiomyopathies Emphasizes Their Clinical Relevance and Points to Novel Candidate Genes

**Why Not Relevant**: The paper does not mention the gene PTK2B anywhere in the provided content. All reported de novo variants and candidate genes are different (TTN, DSP, SCN5A, TNNC1, TPM1, CRYAB, MYH7, TRIB3, SLC2A6, APOBEC family, UNC45A). The study focuses on cardiomyopathy patients (12 probands) and reports genes identified in that specific cohort; absence of PTK2B from these results is an implicit negative but is not informative evidence regarding whether PTK2B is or is not associated with any Mendelian disease in the broader literature. Key limitations that make this paper irrelevant to the PTK2B claim: small sample size, unclear gene panel composition (it is not stated whether PTK2B was included among tested/covered genes), limited phenotype focus (cardiomyopathies) which may not overlap with phenotypes caused by PTK2B variation, and the fact that absence of mention is not a systematic negative finding. To evaluate the claim that PTK2B is likely associated with at least one Mendelian disease, one should consult resources that aggregate gene–disease associations (e.g., OMIM, ClinVar, ClinGen gene curation, peer-reviewed reports specifically implicating PTK2B, and population/segregation evidence). This paper provides neither direct genetic association data nor mechanistic data for PTK2B, so its relevance to the claim is effectively zero.


[Read Paper](https://www.semanticscholar.org/paper/8b13172512a7f7359fcd75f7804754dfe9c36e05)


### De novo variation in ARID1B gene causes Coffin-Siris syndrome 1 in a Chinese family with excessive early-onset high myopia

**Why Not Relevant**: The provided paper content exclusively describes findings for the ARID1B gene (a de novo heterozygous frameshift insertion variant causing Coffin–Siris syndrome with excessive early-onset high myopia). There is no mention of PTK2B, no data, no variant reports, and no mechanistic discussion involving PTK2B. Therefore, based on the content supplied, the paper provides no direct or mechanistic evidence to support or refute the claim that PTK2B is likely associated with at least one Mendelian disease. It remains possible (but unsupported by the provided excerpt) that the full paper might discuss other genes; however, with only the supplied text, no connection to PTK2B can be established. Consequently the relevance score is very low.


[Read Paper](https://www.semanticscholar.org/paper/a0fe18dd082eeec9079cc284babcd59eb224375f)


### Whole-Exome Sequencing Identified a Novel DYRK1A Variant in a Patient With Intellectual Developmental Disorder, Autosomal Dominant 7

**Why Not Relevant**: This paper provides no direct or mechanistic evidence relating PTK2B to any Mendelian disease. The manuscript is a single-case report of intellectual developmental disorder, autosomal dominant 7 (MRD7) attributed to a novel heterozygous frameshift duplication in DYRK1A (c.848dup, p.(Asn283LysfsTer6)). There is no mention of the gene PTK2B, no data (genetic, functional, or clinical) implicating PTK2B, nor any discussion of pathways or interactions that would link PTK2B to the phenotype. Specific reasons for assigning relevance = 0.0: 1) Gene focus: All reported molecular findings and causal attribution are to DYRK1A; the pathogenic variant identified and validated by WES and Sanger sequencing is in DYRK1A. 2) No PTK2B data: The paper does not report sequencing results, variants, expression, protein studies, or case observations involving PTK2B. 3) No mechanistic linkage: There are no mechanistic experiments or cited literature in this paper connecting DYRK1A to PTK2B, nor any discussion that would provide plausibility for PTK2B causing a Mendelian disorder in this context. 4) Scope and sample size: The study is a single-patient diagnostic report; even if PTK2B had been mentioned, the sample size would limit generalizability. Caveat: The absence of PTK2B in this report does not disprove that PTK2B may be associated with a Mendelian disease elsewhere. It only means this paper does not provide supporting or refuting evidence. For establishing PTK2B as likely associated with a Mendelian disease, one would need papers that (a) report segregating pathogenic variants in PTK2B in affected families or multiple unrelated cases, (b) provide functional validation that variants alter PTK2B function in a disease-relevant way, or (c) present strong statistical enrichment of rare damaging PTK2B variants in cases versus controls. None of those elements are present in the provided content.


[Read Paper](https://www.semanticscholar.org/paper/62be40f3f7ee1a9d79fa945981f01aee7ba7049d)


### Modelling the evolutionarily conserved MANF/CDNF protein family in Drosophila melanogaster

**Why Not Relevant**: I found no mention of PTK2B (or its synonyms, e.g. Pyk2, FAK2) in the provided paper content. The manuscript focuses on the MANF/CDNF family (mesencephalic astrocyte-derived neurotrophic factor and cerebral dopamine neurotrophic factor), characterisation of the Drosophila orthologue DmManf, structural-functional analyses of MANF domains and motifs (for example the CXXC motif and ER retention signal), phenotypes of DmManf loss and point mutations, and genetic interactions between DmManf and genes encoding ER chaperones (e.g. GRP78), unfolded protein response components, and genes involved in ubiquinone synthesis and mitochondria. Those topics and quoted passages in the provided text do not provide direct human genetic evidence (familial segregation, pathogenic variants, linkage/association statistics, or clinical case reports) tying PTK2B to any Mendelian disease. Mechanistically, while the paper reports interactions of MANF with ER-chaperone pathways and mitochondrial-related genes (which are broad cellular processes), it does not describe any biochemical, genetic, or functional link to PTK2B or to protein-tyrosine-kinase signalling pathways that would allow an inference that PTK2B is implicated in a Mendelian disorder. The absence of any PTK2B mention means this paper neither supports nor refutes the claim that PTK2B is likely associated with at least one Mendelian disease. Note that absence of evidence in this single manuscript is not evidence of absence in the literature overall; to evaluate the claim about PTK2B you should consult human genetics databases (OMIM, ClinVar, HGMD), population-variant resources (gnomAD), and targeted PubMed searches for PTK2B and mendelian / familial / monogenic / pathogenic variants.


[Read Paper](https://www.semanticscholar.org/paper/ed79568891ddea6ebad39d87ffa8790295111571)


### Exome Survey and Candidate Gene Re-Sequencing Identifies Novel Exstrophy Candidate Genes and Implicates LZTR1 in Disease Formation

**Why Not Relevant**: The provided paper content does not mention PTK2B at all. The study focuses on exome analysis in cloacal exstrophy and related BEEC phenotypes and reports candidate genes (NR1H2, GKAP1, AKR1B10, CLSTN3, NDST4, PLEKHB1, SVEP1, LZTR1, SLC7A4). There is no direct genetic, statistical, or mechanistic data relating PTK2B to Mendelian disease in the text. While omission of PTK2B in the gene list may be taken as limited negative evidence in this specific cohort, absence of mention does not constitute a systematic search for or formal exclusion of PTK2B as a Mendelian gene. The study is phenotype-restricted (BEEC/CE) and reports rare/private variants in a small sample set; it neither evaluates PTK2B nor provides mechanistic data linking PTK2B to disease. Therefore the paper is not relevant to supporting the claim that PTK2B is likely associated with at least one Mendelian disease.


[Read Paper](https://www.semanticscholar.org/paper/7063988a5d8b29057b5aadc11328d505afac734d)


### The application of whole-exome sequencing in the early diagnosis of rare genetic diseases in children: a study from Southeastern China

**Why Not Relevant**: The provided paper content contains no mention of the gene PTK2B, no gene-specific variant data, and no mechanistic discussion related to PTK2B. The excerpt reports cohort-level whole-exome sequencing diagnostic statistics (overall diagnostic rate, comparison of Trio-WES vs singleton-WES, phenotype-specific diagnostic rates, proportions of inheritance modes, and proportion of de novo variants) but gives no gene names, no variant-level results, and no functional or segregation evidence that could support or refute an association between PTK2B and a Mendelian disease. Therefore, based on the supplied text alone, this paper provides no direct evidence (no reported pathogenic or likely pathogenic PTK2B variants, no case reports implicating PTK2B) nor mechanistic evidence (no experimental assays, animal models, or pathway analyses involving PTK2B) relevant to the claim.

Limitations / caveats: The assessment is limited to the excerpt you provided. It is possible that the full paper contains tables, supplementary materials, or unquoted sections that list implicated genes (including PTK2B) or describe functional follow-up; those sections were not supplied here. To properly evaluate the claim that PTK2B is likely associated with at least one Mendelian disease, one would need: (a) explicit reporting of PTK2B variants classified as pathogenic/likely pathogenic in affected individuals, with genotype–phenotype correlation and segregation data; and/or (b) mechanistic evidence (functional assays, animal models, molecular pathway perturbation) linking PTK2B disruption to disease phenotype. None of these are present in the provided text.


[Read Paper](https://www.semanticscholar.org/paper/ba9aa0637319a74b3dfdae65099a70edd6c60ebf)


## Search Queries Used

- PTK2B PYK2 pathogenic variant ClinVar OMIM Mendelian monogenic human disease case report family segregation

- PTK2B de novo variant exome sequencing trio rare disease diagnostic exome novel candidate gene autosomal dominant autosomal recessive

- PTK2B protein tyrosine kinase 2 beta knockout mouse phenotype zebrafish knockdown neuronal development synaptic function cell migration focal adhesion kinase functional study

- PTK2B gnomAD loss of function intolerance pLI constraint haploinsufficiency PolyPhen SIFT CADD pathogenicity prediction Alzheimer's disease GWAS eQTL


## Bibliometric Configuration

- Use Bibliometrics: True

- Author Impact Weight: 0.4

- Citation Impact Weight: 0.4

- Venue Impact Weight: 0.2
