## Supplementary material for "MENDELSEEK: An algorithm that predicts Mendelian Genes and elucidates what makes them special": AOC1.txt

# Claim: The gene AOC1 is likely associated with at least one Mendelian disease.

**Status**: processed

**Overall Rating**: 0

**Explanation**:

Supporting evidence:
The provided paper documents that non-coding variants, particularly those affecting upstream open reading frames (uORFs) in 5'UTRs, can cause Mendelian disorders and that systematic annotation of translation initiation sites in OMIM-listed disease genes can reveal such pathogenic variants. The authors manually annotated ~4.7k TISs related to uORFs in disease-associated genes, validated translation for a set of genes experimentally, and identified uORF-disrupting variants present in HGMD and ClinVar that could underlie Mendelian disease. The paper also reports development of a machine-learning approach to predict TISs in other human genes. These findings support the general principle that previously unrecognized non-coding variants in disease genes can be discovered by the methods described, and that expanding annotation can reveal new gene–disease links.

Caveats or contradictory evidence:
The excerpts do not mention AOC1 specifically. There is no direct evidence in the provided text that AOC1 was included among the OMIM genes annotated, that any uORF-affecting or other variants in AOC1 were identified in HGMD/ClinVar as pathogenic, or that AOC1 has been linked to any Mendelian phenotype in the datasets the authors analyzed. Thus, while the paper demonstrates a general approach and gives several gene-specific examples (ETFDH, PAX9, MAST1, HTT, TTN, GLI2, COL2A1, ZIC2), AOC1 is not among the validated examples cited in the excerpts. There is also no counter-evidence presented that would argue against AOC1 being associated with a Mendelian disease; the absence of mention is simply absence of evidence.

Analysis of potential underlying mechanisms and implications:
Mechanistically, the paper's logic is that disruption of regulatory elements in 5'UTRs (uORFs/TISs) can alter translation of the canonical coding sequence, producing loss- or gain-of-function effects that manifest as Mendelian phenotypes. If AOC1 is a gene with an annotated 5'UTR and contains uORFs or alternative TISs, it is mechanistically plausible that disruptive variants in those elements could contribute to disease. The paper's machine-learning approach could, in principle, be applied to AOC1 to predict TISs and flag potential pathogenic non-coding variants, but none of the provided excerpts state that this was done for AOC1. Therefore, while the method increases the general plausibility that additional genes (potentially including AOC1) could be linked to Mendelian disease via non-coding variants, there is no direct demonstration for AOC1.

Which rating is most appropriate:
Given the absence of any direct mention, data, or identified pathogenic variants for AOC1 in the provided excerpts, the evidence does not support the specific claim that AOC1 is likely associated with at least one Mendelian disease. The paper provides a general methodological and conceptual framework that could uncover such associations for many genes, but it does not present AOC1 as an example or report pathogenic variants in AOC1. Therefore, based solely on the provided evidence, the appropriate assessment is that there is no evidence for the claim in these materials.


**Final Reasoning**:

The provided paper supports the general concept that non-coding variants (uORF/TIS-disrupting) can cause Mendelian disease and that systematic annotation can reveal gene–disease links. However, the excerpts contain no direct data or statements implicating AOC1 specifically. No pathogenic AOC1 variants, no AOC1 TIS/uORF annotations, and no listing of AOC1 among the validated examples are provided. Because the claim is specific to AOC1 and the supplied evidence does not mention or analyze AOC1, the correct conclusion based on these materials is that there is no evidence to support or refute the claim. Accordingly, the rating is "No Evidence."


## Relevant Papers


### Annotation of uORFs in the OMIM genes allows to reveal pathogenic variants in 5′UTRs

**Authors**: A. Filatova (H-index: 8), M. Skoblov (H-index: 5)

**Relevance**: 0.18

**Bibliometric Impact**: 0.21600000000000003


**Excerpts**:

- An increasing number of studies emphasize the role of non-coding variants in the development of hereditary diseases.

- It was previously shown that variants in 5′-untranslated regions (5′UTRs) can lead to hereditary diseases due to disruption of upstream open reading frames (uORFs).

- Here, we performed a manual annotation of upstream translation initiation sites (TISs) in human disease-associated genes from the OMIM database and revealed ∼4.7 thousand of TISs related to uORFs.

- Using a luciferase assay, we experimentally validated the translation of uORFs in the ETFDH, PAX9, MAST1, HTT, TTN,GLI2 and COL2A1 genes, as well as existence of N-terminal CDS extension in the ZIC2 gene.

- We revealed the variants from the HGMD and ClinVar databases that disrupt uORFs and thereby could lead to Mendelian disorders.

- We also showed that the distribution of uORFs-affecting variants differs between pathogenic and population variants.

- Finally, drawing on manually curated data, we developed a machine-learning algorithm that allows us to predict the TISs in other human genes.


**Explanations**:

- This sentence provides general context that non-coding variation (the class that includes uORF-affecting variants) is implicated in hereditary diseases. It is mechanistic/ contextual evidence relevant to the plausibility that non-coding changes in a given gene (such as AOC1) could underlie Mendelian disease. Strength: states a widely supported concept. Limitation: no gene-specific data and no direct evidence about AOC1.

- This sentence states the mechanistic link specifically: 5′UTR variants can cause hereditary disease by disrupting uORFs. It strengthens biological plausibility that uORF disruption in any gene (potentially including AOC1) could produce Mendelian phenotypes. Strength: points to a concrete, experimentally supported mechanism. Limitation: still general — does not document that AOC1 has such variants or that they are pathogenic.

- This describes the scope of their annotation effort (manual TIS annotation in OMIM genes) and quantifies the number of TISs found. Mechanistic/relevance: indicates the authors systematically searched disease-associated genes for upstream TISs/uORFs, which is the dataset and approach that could detect uORF-related pathogenicity in genes like AOC1 if AOC1 were included in OMIM or their set. Strength: shows comprehensive dataset; Limitation: abstract does not list all genes analyzed and does not state whether AOC1 was among them.

- This sentence gives experimental validation in specific genes (ETFDH, PAX9, MAST1, HTT, TTN, GLI2, COL2A1, ZIC2). It is direct experimental evidence that uORFs or N-terminal extensions are real and can be functional in some disease genes. Relevance to AOC1: indirect — demonstrates the approach works for some genes but does not include AOC1. Strength: experimental validation increases confidence in the method; Limitation: AOC1 is not among the validated genes, so no direct evidence linking AOC1 to Mendelian disease here.

- This statement claims they identified variants in HGMD and ClinVar that disrupt uORFs and thereby could lead to Mendelian disorders. This is the closest assertion to disease association: it indicates that database-listed (often disease-cited) variants that affect uORFs exist. Relevance: indirect — if AOC1 had such variants in those databases, the paper's dataset/method could implicate it, but the abstract does not say AOC1 specifically appears. Strength: uses curated disease variant databases (HGMD, ClinVar); Limitation: 'could lead' indicates candidate status, not necessarily proven causality, and the abstract lacks gene-level detail and statistical confirmation.

- This indicates a statistical or distributional observation that uORF-affecting variants differ between pathogenic and population variants. Mechanistic/relevance: supports that uORF disruption is enriched among pathogenic variants and thus is a plausible disease mechanism for genes with such variants. Strength: comparative analysis supports non-random association; Limitation: no numerical/statistical details in the abstract and no gene-specific claims about AOC1.

- This describes a predictive tool to find TISs in other genes. Relevance: methodologically relevant because it could be applied to AOC1 to predict whether upstream TISs/uORFs exist and whether variants might affect them. Strength: offers a path to generate gene-specific hypotheses for AOC1. Limitation: prediction does not equal demonstrated pathogenic association; the abstract gives no performance metrics here.


[Read Paper](https://www.semanticscholar.org/paper/5d45c0cdf801505df45fc93a466a2a4101a12871)


## Other Reviewed Papers


### Incomplete penetrance and variable expressivity in monogenic diabetes; a challenge but also an opportunity

**Why Not Relevant**: The provided content is a single scope sentence from a review describing that it will address penetrance and expressivity variation in "MFD" and related factors. It does not mention the gene AOC1, any specific Mendelian disease, pathogenic variants, mechanistic pathways, clinical cases, or experimental results. Therefore, based on the supplied text alone, there is no relevant direct or mechanistic evidence to evaluate the claim. If you can provide additional excerpts or the full text of the paper, I can re-evaluate for any AOC1-related content.


[Read Paper](https://www.semanticscholar.org/paper/79fbedd297d3b7288b734021b4187afec402f5a3)


### The simplest explanation does not have to be preferred: co-occurrence of pathogenic variants in cancer-predisposing genes

**Why Not Relevant**: The supplied paper content does not mention the gene AOC1, Mendelian disease, or any specific gene-level associations. It is a single-sentence fragment describing a study approach: reviewing literature and checking UK 10,000 Genomes Project data for cancer phenotypes and whether effects of cancer susceptibility genes (CSGs) are additive or synergistic. No specific genes, variants, segregation data, case reports, functional assays, or diagnostic/clinical classifications (e.g., OMIM, ClinVar) are provided. Therefore there is no direct evidence supporting or refuting the claim that AOC1 is associated with a Mendelian disease, nor is there mechanistic evidence pertaining to AOC1. Additional reasons the content is not relevant: (1) it lacks gene names or results, (2) it lacks phenotype–genotype correlations, (3) it provides no statistical or experimental details, and (4) it is a methodological/overview statement rather than data or conclusions about specific genes. To evaluate the claim about AOC1 would require excerpts that name AOC1 and present genetic, clinical, or functional evidence; those are absent here.


[Read Paper](https://www.semanticscholar.org/paper/a1372a664bd2c33a0bda638975a8b26d112457fc)


### Case report: Preimplantation genetic testing for infantile GM1 gangliosidosis

**Why Not Relevant**: This paper describes a case of GM1 gangliosidosis caused by pathogenic variants in the GLB1 gene (including a frameshift c.699delG and a missense c.809A>C), the development of a PGT-M assay for those GLB1 variants, and clinical outcomes for the family and embryos. Nowhere in the provided content is the gene AOC1 mentioned, nor are there any data, analyses, or mechanistic discussions that implicate AOC1 in GM1 gangliosidosis or any other Mendelian disorder. The genetic findings and mechanistic discussion are specific to GLB1 (β-galactosidase deficiency) and to preimplantation/prenatal testing for GLB1 variants; the methods and results (next-generation sequencing identifying GLB1 variants, STR haplotyping around GLB1, PGT-M focusing on GLB1) therefore do not provide either direct evidence (e.g., AOC1 variants segregating with disease) or mechanistic evidence (e.g., biochemical interaction, pathway overlap, or functional studies) linking AOC1 to a Mendelian disease. Because the paper’s scope and data are confined to GLB1 and GM1 gangliosidosis, it neither supports nor refutes the claim that AOC1 is likely associated with at least one Mendelian disease. To evaluate that claim would require papers reporting AOC1 variants segregating with a Mendelian phenotype, functional studies of AOC1 disruption, or genetic database/clinical reports implicating AOC1 in disease — none of which are present in the provided content.


[Read Paper](https://www.semanticscholar.org/paper/ce981bc89c565bb4d97baff441baaf8695595448)


## Search Queries Used

- AOC1 pathogenic variant Mendelian disease monogenic disorder ClinVar OMIM

- AOC1 mutation case report whole exome sequencing congenital histamine intolerance diamine oxidase deficiency

- amine oxidase copper containing 1 diamine oxidase DAO enzyme deficiency mechanism loss of function autosomal recessive

- AOC1 polymorphism association study population genetic association migraine irritable bowel syndrome histamine intolerance systematic review meta analysis lack of association


## Usage Statistics

- Prompt Tokens: 0

- Completion Tokens: 0

- Total Tokens: 0

- Estimated Cost: $0.0165


## Bibliometric Configuration

- Use Bibliometrics: True

- Author Impact Weight: 0.4

- Citation Impact Weight: 0.4

- Venue Impact Weight: 0.2
