## Supplementary material for "MENDELSEEK: An algorithm that predicts Mendelian Genes and elucidates what makes them special": AOC2.txt

# Claim: The gene AOC2 is likely associated with at least one Mendelian disease.

**Status**: processed

**Overall Rating**: -1

**Explanation**:

No relevant papers were found that support or refute this claim.


## Relevant Papers


## Other Reviewed Papers


### De novo SOX10 nonsense mutation in a patient with Kallmann syndrome and hearing loss

**Why Not Relevant**: The provided paper content exclusively concerns the gene SOX10 and its role in Kallmann syndrome (KS), mentioning screening of patients and identification of novel loss-of-function SOX10 mutations. There is no mention of AOC2, no data, no mechanistic discussion, and no contextual information that links SOX10 findings to AOC2. Because the content contains no references to AOC2 or to any evidence that could be extrapolated to AOC2, it provides no direct or mechanistic support for the claim that AOC2 is likely associated with at least one Mendelian disease. Any assertion about AOC2 based on this content would be speculative and unsupported by the paper excerpt provided. Additionally, the excerpt is a single sentence with limited context, so even inference about related genes or pathways is not possible from this text alone.


[Read Paper](https://www.semanticscholar.org/paper/19652785549f1c4b8c8204cbaaf76d5fd9f8d90a)


### Identification of novel sarcoma risk genes using a two-stage genome wide DNA sequencing strategy in cancer cluster families and population case and control cohorts

**Why Not Relevant**: The provided paper content does not mention AOC2 or any data about Mendelian disease associations. The single sentence given reads: "Although neither of these two genes have been previously associated with sarcoma, ABCB5 has been shown to share clinical drug resistance associations with melanoma and leukaemia and C16orf96 shares regulatory elements with genes that are involved with TNF-alpha mediated apoptosis in a p53/TP53-dependent manner." This sentence refers only to ABCB5 and C16orf96 and describes (a) lack of prior sarcoma association, (b) clinical drug-resistance associations for ABCB5 with melanoma and leukaemia, and (c) shared regulatory elements for C16orf96 with genes involved in TNF-alpha mediated, p53/TP53-dependent apoptosis. None of these statements mention AOC2, report associations with Mendelian diseases, or provide mechanistic links that would plausibly connect AOC2 to a Mendelian disorder. In addition, the excerpt provides no original data, statistical results, experimental methods, patient/cohort information, or genetic evidence (e.g., segregation, de novo variants, pathogenic variant functional assays) that would be required to assess a Mendelian disease association. Therefore, based solely on the provided content, there is no direct or mechanistic evidence supporting the claim that AOC2 is likely associated with at least one Mendelian disease. Note: if the full paper contains additional sections not provided here, those sections were not available for review and may contain relevant information; the relevance assessment above applies only to the excerpt supplied.


[Read Paper](https://www.semanticscholar.org/paper/da5be3abeab13d421d0beee322fa9880f6ff3a72)


### Whole Exome Sequence Analysis Provides Novel Insights into the Genetic Framework of Childhood-Onset Pulmonary Arterial Hypertension

**Why Not Relevant**: The provided paper content does not mention AOC2, report variants in AOC2, present functional or mechanistic data involving AOC2, or otherwise address AOC2’s role in disease. The study focuses on childhood-onset pulmonary arterial hypertension and identifies risk alleles in other genes (ATP13A3, BMP10, PDGFD). Because the paper neither includes nor discusses AOC2, it provides no direct or mechanistic evidence for the claim that AOC2 is associated with at least one Mendelian disease. Additional reasons this paper cannot inform the claim: the cohort is small (n = 18), the analysis was limited to 26 candidate genes (the set is not provided here), and absence of a gene in a disease-focused study does not constitute evidence against that gene’s involvement in other Mendelian conditions. Therefore relevance to the claim is effectively zero and independent evidence (other genetic studies, ClinVar/OMIM records, functional studies) would be required to evaluate AOC2’s association with Mendelian disease.


[Read Paper](https://www.semanticscholar.org/paper/f2ee290311f5ab460b053ff2815ce4bd29d0af89)


### Identifying genomic variant associated with long QT syndrome type 2 in an ecuadorian mestizo individual: a case report

**Why Not Relevant**: This paper provides no direct or mechanistic evidence relevant to the claim that AOC2 is likely associated with at least one Mendelian disease. The report is a single-case clinical report of Long QT Syndrome with explicit identification of a pathogenic KCNH2 variant and two VUS in AKAP9 and TTN; AOC2 is not mentioned anywhere in the provided content. Because AOC2 is neither reported as having a variant in this patient nor discussed mechanistically, the paper cannot support the claim that AOC2 is a Mendelian disease gene. Further, the absence of AOC2 in the report is not reliable negative evidence: the text does not specify the breadth of sequencing (targeted panel vs exome/genome), reporting thresholds, filtering criteria, or whether no AOC2 variants were observed or simply not reported. The study is a single-case report (n=1) focused on LQTS and known LQTS-associated genes; it is not a gene discovery, burden, segregation, or functional study that could establish or refute a Mendelian role for AOC2. Therefore relevance to the AOC2 claim is effectively zero.


[Read Paper](https://www.semanticscholar.org/paper/aa65df486c4f1720d86feb42ddc72ed25ab57431)


### The First Case Report of a Homozygous Consensus Acceptor Splice Variant in the NUP214 Gene Associated With Fetal Hydrops and Arthrogryposis Multiplex

**Why Not Relevant**: The provided paper content exclusively discusses the NUP214 gene, its role in nucleocytoplasmic transport, and a reported homozygous splice-site variant (c.46-2A>G) in NUP214 associated with a fetal phenotype and with previously reported susceptibility to acute infection-induced encephalopathy type 9 (ILAE9). There is no mention of the gene AOC2 (nor any obvious synonyms) anywhere in the supplied text. Because the excerpt contains no references to AOC2, it provides neither direct evidence (e.g., patient variants, segregation, case reports, statistical associations) nor mechanistic evidence (e.g., biochemical function, pathways, protein interactions) linking AOC2 to any Mendelian disease. 

Limitations: This conclusion is limited to the text you provided. If the full paper (outside the supplied content) includes additional sections or supplementary material that mention AOC2, those were not available for review here. To evaluate the claim about AOC2, a targeted search of the full paper, the wider literature, and databases such as OMIM, ClinVar, HGMD, and PubMed would be needed.


[Read Paper](https://www.semanticscholar.org/paper/c45e23cc4c0d742b80b747141c024f5f66a2f05a)


### Case report: Preimplantation genetic testing for infantile GM1 gangliosidosis

**Why Not Relevant**: The provided paper content exclusively investigates GM1 gangliosidosis caused by mutations in the GLB1 gene. It describes clinical findings, identification of two GLB1 variants by next-generation sequencing, and development of PGT-M methods targeted to GLB1 and nearby STR markers. The text contains no mention of the gene AOC2, no variant, clinical, genetic, functional, or mechanistic data relating to AOC2, and no discussion linking AOC2 to GM1 gangliosidosis or any other Mendelian disorder. Therefore this paper provides no direct or mechanistic evidence supporting the claim that AOC2 is likely associated with at least one Mendelian disease. Note: absence of evidence in this single paper is not evidence of absence overall; it only indicates that this paper is irrelevant to assessing AOC2’s disease associations.


[Read Paper](https://www.semanticscholar.org/paper/ce981bc89c565bb4d97baff441baaf8695595448)


### Autosomal Dominant Intellectual Development Disorder-6 (MRD6) Without Seizures Linked to a De Novo Mutation in the grin2b Gene Revealed by Exome Sequencing: A Case Report of a Moroccan Child

**Why Not Relevant**: The provided paper content exclusively discusses a GRIN2B (grin2b) truncating mutation causing autosomal dominant intellectual development disorder-6 (MRD6). There is no mention of the gene AOC2, no data (genetic, functional, epidemiologic) about AOC2, nor any discussion that would relate GRIN2B findings to AOC2. Therefore this paper provides no direct or mechanistic evidence supporting or refuting the claim that AOC2 is likely associated with at least one Mendelian disease. Absence of AOC2 in the text means the paper cannot be used as evidence for the claim; it neither confirms nor denies any AOC2–disease association. Additional literature specifically addressing AOC2 would be required to evaluate the claim.


[Read Paper](https://www.semanticscholar.org/paper/67d756e0cc302e5d46fdda9df34403f1f2de9a5a)


### Contribution of rare transmitted and de novo variants among 2 , 871 1 congenital heart disease probands

**Why Not Relevant**: The provided paper content is a large exome-sequencing study focused specifically on congenital heart disease (CHD) and the identification of CHD-related genes through analysis of de novo and transmitted rare variants. The text does not mention AOC2 anywhere in the provided excerpts; there is no gene-level listing or specific result implicating AOC2. Therefore the paper provides no direct evidence that AOC2 is associated with any Mendelian disease. Indirectly, the study demonstrates an approach (WES of a disease cohort, variant calling and enrichment testing) that can identify Mendelian disease genes for CHD; absence of AOC2 from the reported significant genes implies it was not implicated in this CHD cohort. Limitations relevant to interpreting absence of AOC2 in this paper: (1) the cohort and analyses are CHD-focused — genes causing Mendelian diseases without CHD manifestations would not be expected to be found; (2) clinically ascertained CNV and aneuploidy cases were excluded, so structural variant–mediated associations would be missed; (3) whole-exome sequencing does not capture non-coding disease-causing variation; (4) extremely rare or population-specific AOC2 pathogenic variants might not be present in this cohort; (5) coverage gaps or variant-calling filters could miss some AOC2 variants. Given no mention of AOC2 and these methodological constraints, the paper does not provide support for the claim that AOC2 is likely associated with at least one Mendelian disease.


[Read Paper](https://www.semanticscholar.org/paper/e5830d9e95469f22518560007746202e62db26a0)


### Explorer CDKL 5 variants : improving our understanding of a rare neurological disorder

**Why Not Relevant**: This paper exclusively studies the gene CDKL5 and CDKL5-related variants and makes no mention of AOC2. All reported data, methods, results, and interpretations refer to CDKL5 (e.g., population data from ExAC for CDKL5, clustering of pathogenic missense variants in the catalytic domain of CDKL5, isoform-specific effects for CDKL5), and there are no experiments, analyses, case reports, or mechanistic discussions involving AOC2. Therefore the paper provides neither direct evidence (cases, segregation, pathogenic variants) nor mechanistic evidence (biochemical function, molecular pathways, functional assays) relevant to the claim that AOC2 is likely associated with a Mendelian disease. The study’s approaches (use of ExAC/1000G, variant reclassification, minigene splicing assays) could in principle be applied to evaluate AOC2 in other work, but this paper does not contain data or statements that support or refute any disease association for AOC2.


[Read Paper](https://www.semanticscholar.org/paper/a959fe9191159fa8a508c918a1e4e7dcf60553fd)


### 251 Expanding the genetic and clinical spectrum of the nono-associated x-linked intellectual disability syndrome with left ventricular non-compaction

**Why Not Relevant**: This paper describes a clinical and molecular case study implicating the X-linked gene NONO in a patient with developmental delay and cardiomyopathy. The provided text repeatedly names and discusses NONO (e.g., identification of an intronic NONO deletion, follow-up splicing studies, reclassification of a novel NONO variant as pathogenic) and the mechanistic evidence pertains to intron readthrough, use of an alternative donor site, frameshift, and nonsense-mediated decay for NONO transcripts. Nowhere in the provided content is the gene AOC2 mentioned, nor are there any data, variants, experiments, family segregation results, or mechanistic analyses that involve AOC2. Therefore the paper provides no direct evidence supporting or refuting the claim that AOC2 is likely associated with a Mendelian disease. Methodologically, although the study uses exome sequencing and RNA studies (techniques that could detect AOC2 variants in principle), the absence of any report or discussion of AOC2 in this case means there is no applicable information about AOC2’s disease association. Finally, the lack of AOC2 data in a single-case NONO study cannot be interpreted as negative evidence against AOC2’s involvement in Mendelian disease (absence of evidence is not evidence of absence), but it does mean this paper is irrelevant to evaluating the claim about AOC2.


[Read Paper](https://www.semanticscholar.org/paper/87d24f3f290951a33b8326893afc41835c74f8d8)


### 252 Germline transmission of pathogenic braf: expanding the clinical phenotype of cfc syndrome

**Why Not Relevant**: The paper excerpt concerns identification, functional validation, segregation, and reclassification of a pathogenic variant in the NONO gene in a single proband. There is no mention of the gene AOC2 anywhere in the provided content, no data (genetic, functional, or epidemiologic) related to AOC2, and no discussion drawing links between NONO and AOC2. Therefore the manuscript provides no direct or mechanistic evidence relevant to the claim that AOC2 is likely associated with at least one Mendelian disease. The available evidence instead supports NONO as disease-associated; it cannot be used to infer anything about AOC2 without additional data or explicit analysis connecting the two genes. Because of this complete absence of AOC2-related information, relevance to the claim is effectively zero.


[Read Paper](https://www.semanticscholar.org/paper/008f9c79db82794e3eaa341c0b8b751e8265b2cf)


### THU0480 Candidate Gene Discovery in Autoimmunity by Using Extreme Phenotypes and Whole Exome Capture

**Why Not Relevant**: This paper does not mention AOC2 at any point in the provided content. The study focuses on discovery of genes associated with multiple autoimmune syndrome (MAS) and reports candidate genes PKD1, ALPK2, and OBSCN; no results, functional data, variant calls, enrichment statistics, or mechanistic discussion concerning AOC2 are present. Therefore the paper provides no direct genetic, clinical, or mechanistic evidence that AOC2 is associated with any Mendelian disease. Additionally, the study's small sample size (12 sporadic cases and 24 familial individuals), its specific phenotype focus (MAS), and the filtration/analysis choices used mean that even an absence of AOC2 among reported candidates cannot be interpreted as strong evidence refuting AOC2 involvement in Mendelian disease more generally. For these reasons the paper is essentially irrelevant to the specific claim that AOC2 is likely associated with at least one Mendelian disease.


[Read Paper](https://www.semanticscholar.org/paper/907125dee95e1c0a04e7b7952a79d70875ca3463)


## Search Queries Used

- AOC2 gene Mendelian disease variant ClinVar OMIM case report

- AOC2 de novo variant exome sequencing trio rare Mendelian disease loss of function nonsense frameshift segregation

- AOC2 amine oxidase copper containing 2 enzyme function expression retina eye development knockout mouse phenotype

- AOC2 novel candidate gene Mendelian gene discovery rare variant burden unsolved exome cohorts family segregation study


## Bibliometric Configuration

- Use Bibliometrics: True

- Author Impact Weight: 0.4

- Citation Impact Weight: 0.4

- Venue Impact Weight: 0.2
