## Supplementary material for "MENDELSEEK: An algorithm that predicts Mendelian Genes and elucidates what makes them special": AOC3.txt

# Claim: The gene AOC3 is likely associated with at least one Mendelian disease.

**Status**: processed

**Overall Rating**: -1

**Explanation**:

No relevant papers were found that support or refute this claim.


## Relevant Papers


## Other Reviewed Papers


### mGAP: the macaque genotype and phenotype resource, a framework for accessing and interpreting macaque variant data, and identifying new models of human disease

**Why Not Relevant**: The provided paper content is a single, descriptive statement about a macaque variant resource and contains no mention of the human gene AOC3, no data on associations with Mendelian diseases, and no mechanistic information linking AOC3 to disease. The exact provided sentence is: "The macaque Genotype And Phenotype (mGAP) resource is the first public website providing searchable, annotated macaque variant data, and presents unprecedented opportunity to investigate potential genetic associations with currently characterized disease models, and to uncover new macaque models based on parallels with human risk alleles." This sentence only describes the existence and potential utility of the mGAP resource; it does not provide direct evidence (e.g., association statistics, case reports, segregation data) that AOC3 is associated with any Mendelian disease, nor does it provide mechanistic evidence (e.g., functional studies, pathogenic variants, biochemical pathways) implicating AOC3.

Detailed reasons for non-relevance and limitations:
- No direct evidence: The content contains no variant-level data, no phenotype-genotype correlations, and no named genes (AOC3 is not mentioned). Therefore it cannot support or refute the claim that AOC3 is likely associated with a Mendelian disease.
- No mechanistic evidence: The passage gives no information about gene function, pathogenic mechanisms, or biological pathways involving AOC3.
- No methods or results: There are no experimental methods, sample descriptions, statistical analyses, or results to assess evidence quality, effect sizes, or significance.
- Species and generalizability caveat: The statement concerns macaque variant resources. Even if mGAP contained AOC3 variants, cross-species differences in gene function, variant effects, and phenotype manifestation limit direct inference to human Mendelian disease without additional evidence.

What would be needed to evaluate the claim properly:
- Direct human evidence: entries in OMIM, ClinVar, HGMD, or peer-reviewed case reports showing pathogenic AOC3 variants segregating with Mendelian phenotypes, with supporting functional data.
- Mechanistic data: functional studies showing how specific AOC3 variants disrupt gene/protein function in a way that plausibly causes disease (e.g., loss-of-function, dominant-negative effects), including biochemical, cellular, or animal model evidence.
- Statistical evidence: case-control or family-based analyses with sufficient sample size and appropriate controls, and clear pathogenicity classifications following guidelines (e.g., ACMG criteria).
- If leveraging mGAP: explicit mGAP entries showing macaque AOC3 variants that model known human pathogenic variants and phenotype concordance, plus cross-species functional validation.

Conclusion: Based solely on the provided paper content, there is no evidence — direct or mechanistic — to support or refute the claim that AOC3 is likely associated with at least one Mendelian disease. To assess the claim, consult human genetic databases (OMIM, ClinVar, PubMed) and/or detailed mGAP variant records and functional studies if available.


[Read Paper](https://www.semanticscholar.org/paper/f47691ff666dd65548f1282e606d696eae56524d)


### Vascular adhesion protein 1 in nonalcoholic steatohepatitis: A novel biomarker?

**Why Not Relevant**: The paper provides functional, biochemical, clinical-association, and animal-model evidence that the AOC3 gene product (VAP-1) participates in liver inflammation and fibrosis and that its inhibition alters disease phenotypes in models and correlates with NASH severity in patients. However, it contains no genetic evidence that mutations in AOC3 cause a Mendelian (single-gene, monogenic) disorder: there are no reports of pathogenic germline variants, familial segregation analyses, linkage/association to rare high-penetrance alleles, or descriptions of discrete inherited clinical syndromes attributable to AOC3. The associations and knockout/antibody perturbation data address complex disease mechanisms and therapeutic modulation rather than Mendelian causality. Therefore, while the paper strengthens biological plausibility that AOC3 is disease-relevant, it does not provide direct or sufficient evidence to support the claim that AOC3 is likely associated with at least one Mendelian disease.


[Read Paper](https://www.semanticscholar.org/paper/f15197ba71ba7c2bde49d938ed9899e915562821)


### MSeqDR Quick‐Mitome (QM): Combining Phenotype‐Guided Variant Interpretation and Machine Learning Classifiers to Aid Primary Mitochondrial Disease Genetic Diagnosis

**Why Not Relevant**: The paper describes a bioinformatics/clinical‑research platform (MSeqDR Quick‑Mitome) and a variant pathogenicity classifier for diagnosing primary mitochondrial diseases. It does not mention the gene AOC3, present any AOC3 variant data, report case(s) implicating AOC3 in a Mendelian disorder, or present mechanistic/functional experiments concerning AOC3. The only potential relevance is indirect: the pipeline annotates variants using databases (OMIM, ClinVar) that might themselves contain AOC3 associations, and the classifier could in principle score AOC3 variants if they were included in an analysed dataset. However, the paper presents no direct or gene‑specific evidence for AOC3; therefore it does not support the claim that AOC3 is likely associated with at least one Mendelian disease. Consequently, the paper's relevance to the claim is negligible.


[Read Paper](https://www.semanticscholar.org/paper/d12bb508040b75453f2a34e64f096b0cc1f15b01)


### Generalized Analysis of Open Genetic Databases Reveals New Associations with Migraine

**Why Not Relevant**: The provided paper content does not mention the gene AOC3 by name nor provide any gene-level results that could be evaluated for Mendelian disease association. While the study interrogated ClinVar and selected variants annotated as "pathogenic" or "likely pathogenic" (which could indicate Mendelian disease associations), and while it groups identified variants into broad functional categories including "enzymes" (AOC3 encodes an enzyme), none of the excerpts list specific genes or show that AOC3 variants are pathogenic, associated with a Mendelian phenotype, or cited in ClinVar as disease-causing. Therefore there is no direct evidence in the supplied text that AOC3 is associated with any Mendelian disease. Mechanistically, the paper's approach (using ClinVar pathogenic annotations and GWAS associations) could detect Mendelian gene associations, but without gene-level reporting in the provided content, the claim about AOC3 cannot be supported or refuted. Given the absence of any explicit mention of AOC3 or Mendelian diagnoses linked to it in the supplied text, the relevance of this paper to the specific claim is very low.


[Read Paper](https://www.semanticscholar.org/paper/124525d37c8455ef131d488299564098b1b8761a)


### British Society for Matrix Biology Autumn 2020 Meeting: “Basement Membranes in Health and Disease”

**Why Not Relevant**: This paper content contains no direct or indirect evidence relating to the gene AOC3 (also known as VAP-1 or semicarbazide-sensitive amine oxidase). I searched the provided text for the gene name AOC3 and common synonyms (VAP-1, SSAO) and found no mentions. The manuscript excerpts focus on extracellular matrix biology, collagen biosynthesis and processing, integrin α1β1 function, basement membrane components (for example LAMB2, COL4A3, COL4A4, COL4A5, COL7A1, Col5a2), giantin knockout models, and genetic modifiers of collagen IV disease. While these topics concern Mendelian collagen and basement-membrane diseases (the text explicitly mentions known Mendelian disorders such as Pierson syndrome, Alport syndrome, thin basement membrane nephropathy, and dystrophic epidermolysis bullosa), none of the described experiments, results, or discussions involve AOC3, its protein product, pathogenic variants, clinical phenotypes, or human genetic association data.

Because the paper provides neither (a) mention of AOC3 nor (b) experimental, clinical, or genetic data that could plausibly be extrapolated to support an association between AOC3 and a Mendelian disease, it does not provide evidence for or against the claim that "AOC3 is likely associated with at least one Mendelian disease." Mechanistic pathways discussed (collagen processing, integrin-mediated regulation of growth factor signalling, basement membrane composition and remodeling) do not implicate AOC3 in the text, so no mechanistic link can be inferred from this document.

Limitations and suggested next steps: To evaluate the claim about AOC3, one should instead consult human genetics resources and primary literature that explicitly cover AOC3, for example OMIM, ClinVar, HGMD, gnomAD constraint metrics (pLI, LOEUF), human case reports, mouse knockout phenotypes for Aoc3, and PubMed searches for ‘‘AOC3 mutation’’ or ‘‘VAP-1 Mendelian’’. The current document does not supply the necessary data.


[Read Paper](https://www.semanticscholar.org/paper/e27ef31faa9280438adb116ad5e8faaf9838ff15)


## Search Queries Used

- AOC3 amine oxidase copper containing 3 VAP 1 vascular adhesion protein 1 mutation pathogenic variant Mendelian disease case report patient monogenic

- AOC3 ClinVar OMIM Orphanet pathogenic variant clinical annotation gene disease association monogenic

- AOC3 knockout mouse phenotype VAP 1 vascular adhesion protein 1 knockout leukocyte trafficking endothelial adhesion inflammation fibrosis metabolic phenotype

- AOC3 loss of function variant gnomAD pLI constraint intolerance loss of function depletion exome sequencing rare variant association monogenic disease discovery


## Bibliometric Configuration

- Use Bibliometrics: True

- Author Impact Weight: 0.4

- Citation Impact Weight: 0.4

- Venue Impact Weight: 0.2
