## Supplementary material for "MENDELSEEK: An algorithm that predicts Mendelian Genes and elucidates what makes them special": APEX1.txt

# Claim: The gene APEX1 is likely associated with at least one Mendelian disease.

**Status**: processed

**Overall Rating**: -1

**Explanation**:

No relevant papers were found that support or refute this claim.


## Relevant Papers


## Other Reviewed Papers


### ClinGen Myeloid Malignancy Variant Curation Expert Panel recommendations for germline RUNX1 variants.

**Why Not Relevant**: The provided paper content focuses on development and testing of gene- and disease-specific ACMG/AMP variant curation criteria by the ClinGen Myeloid Malignancy Variant Curation Expert Panel, with an explicit and repeated focus on RUNX1 and inherited myeloid malignancies. There is no mention of the gene APEX1, no data on APEX1 variants, and no mechanistic discussion involving APEX1. Because the paper does not address APEX1 at all, it provides neither direct evidence (e.g., observed pathogenic APEX1 variants segregating with Mendelian disease) nor mechanistic evidence (e.g., functional studies linking APEX1 perturbation to a Mendelian phenotype) relevant to the claim. Absence of evidence in this paper does not imply evidence of absence for APEX1 associations overall; to evaluate the claim one must consult other sources (OMIM, ClinVar, PubMed functional/genetic studies) that specifically report on APEX1.


[Read Paper](https://www.semanticscholar.org/paper/d2a2dd2ebfb5a63969c41c03899f8f978fb18c6a)


### Evidence-based recommendations for gene-specific ACMG/AMP variant classification from the ClinGen ENIGMA BRCA1 and BRCA2 Variant Curation Expert Panel

**Why Not Relevant**: The provided paper content exclusively describes the ENIGMA research consortium's work on developing and applying variant classification methods for the BRCA1 and BRCA2 genes, including adaptation of ACMG/AMP criteria, statistical calibration of evidence strength, and pilot testing on 40 BRCA1/2 variants. The text contains no mention of the gene APEX1, no data linking APEX1 to any Mendelian phenotype, and no mechanistic discussion involving APEX1. Therefore it provides neither direct evidence (e.g., genetic association, case reports, segregation data, pathogenic variants) nor mechanistic evidence (e.g., protein function, pathway disruption, molecular consequences) that would support or refute the claim that APEX1 is likely associated with at least one Mendelian disease. While the paper discusses general methods for gene-specific variant interpretation that could, in principle, be applied to APEX1, methodological descriptions alone do not constitute evidence that APEX1 is associated with a Mendelian disorder. In summary, because the content is focused on BRCA1/BRCA2 variant curation and contains no references or data about APEX1 or relevant Mendelian phenotypes, its relevance to the claim is effectively zero.


[Read Paper](https://www.semanticscholar.org/paper/83345c8cc8964ac610df160320ccc9a2a1724ac7)


### Re-evaluation of a Fibrillin-1 Gene Variant of Uncertain Significance Using the ClinGen Guidelines

**Why Not Relevant**: The provided paper content exclusively addresses fibrillin-1 (FBN1) and its role in Marfan syndrome (MFS), including re-evaluation of FBN1 variants, application of ClinGen FBN1 VCEP criteria, and conclusions about the clinical importance of FBN1 pathogenic variants. There is no mention of the gene APEX1, no genetic association data for APEX1, and no mechanistic discussion linking APEX1 to Mendelian disease. Therefore this paper provides no direct or mechanistic evidence supporting or refuting the claim that APEX1 is likely associated with at least one Mendelian disease. The absence of APEX1 in this study does not inform APEX1's disease relevance; it merely indicates that this publication is not a source of evidence regarding APEX1.


[Read Paper](https://www.semanticscholar.org/paper/bf6815f1239963b018387318ae7e6603ebb3e512)


## Search Queries Used

- APEX1 APE1 apurinic endonuclease 1 pathogenic variant mutation case report Mendelian disease patient family segregation

- APEX1 ClinVar OMIM ClinGen gene disease association curation pathogenic variants review

- APEX1 apurinic endonuclease 1 base excision repair DNA repair loss of function variant human phenotype monogenic disease mechanism REF1 redox effector factor 1

- APEX1 knockout mouse zebrafish phenotype embryonic lethal haploinsufficiency whole exome sequencing undiagnosed disease rare variant segregation autosomal recessive autosomal dominant


## Bibliometric Configuration

- Use Bibliometrics: True

- Author Impact Weight: 0.4

- Citation Impact Weight: 0.4

- Venue Impact Weight: 0.2
