## Supplementary material for "MENDELSEEK: An algorithm that predicts Mendelian Genes and elucidates what makes them special": APOD.txt

# Claim: The gene APOD is likely associated with at least one Mendelian disease.

**Status**: processed

**Overall Rating**: -1

**Explanation**:

No relevant papers were found that support or refute this claim.


## Relevant Papers


## Other Reviewed Papers


### Precision Medicine in Internal Medicine

**Why Not Relevant**: The provided paper content is a general review of genetic testing strategies, variant interpretation, and resources (ACMG guidelines, ClinVar, OMIM) and does not mention the APOD gene or present any gene-specific data. There are no experimental results, case reports, segregation analyses, functional studies, or variant classifications pertaining to APOD in the text. Therefore the paper provides only indirect, general context about how one would discover or evaluate gene–disease relationships (e.g., via exome sequencing or ClinVar) but contains no direct or gene-specific mechanistic evidence to support or refute the claim that APOD is likely associated with at least one Mendelian disease. For a determination about APOD specifically, one would need gene-focused evidence such as reported pathogenic variants in APOD with segregation and phenotype data, functional studies, or authoritative database entries linking APOD to a Mendelian disorder.


[Read Paper](https://www.semanticscholar.org/paper/c7199e8a5226f8d9d55777da0e15e91797bb459a)


### Whole-Exome Sequencing in Adults With Chronic Kidney Disease

**Why Not Relevant**: The supplied paper content does not mention the gene APOD anywhere in the provided text, nor does it report results or findings that directly involve APOD. The paper describes methods (WES), curated gene lists for nephropathy and hypertension (287 OMIM genes), and general analytic and technical limitations, but it gives no data, variant calls, mechanistic discussion, or clinical observations linking APOD to any Mendelian disease. Without APOD being named in the gene lists or results, there is neither direct evidence (observed pathogenic variants, case reports, segregation) nor mechanistic evidence (functional perturbation, pathway implicating APOD in a Mendelian phenotype) in the provided content. Therefore the paper, as supplied, is not relevant to supporting or refuting the claim that APOD is likely associated with at least one Mendelian disease.


[Read Paper](https://www.semanticscholar.org/paper/a29bd57964f68ac5f8e1abcafe1112ba488f53f5)


## Search Queries Used

- APOD apolipoprotein D mutation case report family segregation monogenic Mendelian disease clinical exome sequencing OMIM ClinVar pathogenic likely pathogenic variant

- apolipoprotein D APOD function lipid transport neurodegeneration oxidative stress myelin peripheral neuropathy hereditary neuropathy mouse model knockout apod null functional assay phenotype

- APOD loss of function intolerance pLI gnomAD truncating variant rare variant burden exome sequencing study ClinVar APOD pathogenic likely pathogenic variant frequency

- APOD not associated Mendelian disease no evidence apod benign variant annotation pseudogene gene annotation error GWAS common variant association expression quantitative trait locus eQTL


## Bibliometric Configuration

- Use Bibliometrics: True

- Author Impact Weight: 0.4

- Citation Impact Weight: 0.4

- Venue Impact Weight: 0.2
