## Supplementary material for "MENDELSEEK: An algorithm that predicts Mendelian Genes and elucidates what makes them special": ARHGAP6.txt

# Claim: The gene ARHGAP6 is likely associated with at least one Mendelian disease.

**Status**: processed

**Overall Rating**: -1

**Explanation**:

No relevant papers were found that support or refute this claim.


## Relevant Papers


## Other Reviewed Papers


### In vivo Perturb-Seq reveals neuronal and glial abnormalities associated with autism risk genes

**Why Not Relevant**: The provided paper content does not mention the gene ARHGAP6 anywhere in the quoted text and does not provide a gene-by-gene list of the 35 targeted ASD/ND risk genes, so there is no direct evidence in this content that ARHGAP6 was studied. The study design (pooled in utero CRISPR frameshift perturbations analyzed by single-cell RNA-seq in mouse cortex) assesses cell-intrinsic transcriptional effects and gene modules relevant to autism risk, but it does not provide human clinical genetic evidence (segregation, pathogenic variant enrichment, or replication in human families) required to establish that a gene is associated with a Mendelian disease. Even if ARHGAP6 were included among the 35 genes (which cannot be determined from the supplied text), functional perturbation phenotypes in mouse cortex would be mechanistic/functional evidence at best and insufficient alone to demonstrate that ARHGAP6 is a Mendelian-disease gene. Therefore the paper content supplied is not relevant to the claim that ARHGAP6 is likely associated with at least one Mendelian disease.


[Read Paper](https://www.semanticscholar.org/paper/ccebffce65b9879271b25238c491376d9c2ad3cb)


### Disruption of PHF21A causes syndromic intellectual disability with craniofacial anomalies, epilepsy, hypotonia, and neurobehavioral problems including autism

**Why Not Relevant**: The provided paper content discusses deleterious nonsense, frameshift, and missense mutations in PHF21A and their association with autism spectrum disorder, epilepsy, hypotonia, neurobehavioral problems, tapering fingers, clinodactyly, syndactyly, intellectual disability, and craniofacial anomalies. It contains no mention of ARHGAP6, no data on ARHGAP6 variants, no mechanistic links between PHF21A and ARHGAP6, and no discussion of Mendelian disease associations for ARHGAP6. Therefore the content provides neither direct evidence nor mechanistic evidence supporting or refuting the claim that ARHGAP6 is likely associated with at least one Mendelian disease. Because the single provided sentence is exclusively about PHF21A, it cannot be used to draw conclusions about ARHGAP6; any inference would require additional literature or data not present in the supplied content.


[Read Paper](https://www.semanticscholar.org/paper/87c951a7e4271d50548982a10337a6ca2a1ee1fc)


### The Burden of Candidate Pathogenic Variants for Kidney and Genitourinary Disorders Emerging From Exome Sequencing

**Why Not Relevant**: The provided paper content does not mention the gene ARHGAP6 anywhere, nor does it present gene-specific data or analysis that would allow assessment of ARHGAP6's association with a Mendelian disease. The article describes (a) methods for compiling a curated list of genes associated with Mendelian kidney and genitourinary disorders using OMIM and Orphanet, (b) cohort sequencing and variant-filtering pipelines, and (c) general observations about the prevalence of candidate pathogenic variants in a healthy cohort. Although these methods could in principle have included ARHGAP6 if it were annotated in OMIM/Orphanet and relevant to the phenotypes studied, the excerpt does not provide the curated gene list or any locus-specific results, variant counts, prior pathogenic reports, or mechanistic data for ARHGAP6. Therefore there is no direct evidence (e.g., reported pathogenic variants, segregation data, case reports) nor mechanistic evidence (e.g., functional impact, loss-of-function intolerance for that gene) in this content to support or refute the claim that ARHGAP6 is likely associated with at least one Mendelian disease. To evaluate the claim, one would need at minimum: (1) explicit inclusion of ARHGAP6 in the curated disease-gene list with supporting citations, (2) reported pathogenic variants (with descriptions and frequencies), or (3) mechanistic/functional data linking ARHGAP6 perturbation to a Mendelian phenotype. None of these are present in the provided text.


[Read Paper](https://www.semanticscholar.org/paper/a75e297208c04992568e607d19aa9b9aaebd09d5)


### NR4A2 and Dystonia with Dopa Responsiveness

**Why Not Relevant**: This paper focuses on NR4A2 and other dopamine-related genes (GCH1, TH, SPR) as causes of dopa-responsive and neurodevelopmental disorders. The provided text includes multiple direct statements and mechanistic discussion implicating NR4A2 in a novel monogenic (Mendelian) disease, including identification of a de novo NR4A2 variant by trio whole-exome sequencing and clinical response to levodopa. Nowhere in the provided content is the gene ARHGAP6 mentioned, discussed, or analyzed. Therefore the paper provides no direct genetic evidence (e.g., pathogenic variants, segregation, case reports) implicating ARHGAP6 in a Mendelian disorder, nor any mechanistic data (expression, function, pathway links) connecting ARHGAP6 to the phenotypes studied. Because of this absence, the paper is not relevant to the claim that ARHGAP6 is likely associated with at least one Mendelian disease. Note that absence of evidence in this single paper does not prove ARHGAP6 has no Mendelian associations; it only means this paper does not provide supporting evidence. To evaluate the claim about ARHGAP6, a targeted search of additional sources (OMIM, ClinVar, HGMD, PubMed, gnomAD constraint metrics, and case reports) would be required.


[Read Paper](https://www.semanticscholar.org/paper/0bc0da6e272dc700624e99bd3693b40c9787e03f)


### Mutation of ARHGEF 18 causes adult-onset retinal degeneration and underscores a key role of apicobasal polarity proteins in the maintenance of human photoreceptors

**Why Not Relevant**: The paper presents genetic, functional, phenotypic, and animal-model evidence that biallelic mutations in ARHGEF18 cause an inherited retinal dystrophy (a Mendelian disorder). However, the claim asks about ARHGAP6. The manuscript does not mention ARHGAP6 at all, nor does it present any ARHGAP6-specific genetic, functional, or clinical data. While ARHGEF18 operates in the RHOA signaling pathway (and thus provides mechanistic precedent that perturbation of RHO-pathway components can cause Mendelian retinal disease), that mechanistic overlap does not constitute direct evidence that ARHGAP6 is associated with a Mendelian disease. Additional, gene-specific evidence (e.g., human variants in ARHGAP6 segregating with disease, functional studies of ARHGAP6 variants, or animal models of Arhgap6 perturbation producing a similar phenotype) would be required to support the claim. For these reasons the paper is only indirectly relevant (pathway-level plausibility) and provides no direct evidence about ARHGAP6 itself.


[Read Paper](https://www.semanticscholar.org/paper/2f25ada77419b74d382941e0874d907d3b8b936e)


### Contribution of rare transmitted and de novo variants among 2 , 871 1 congenital heart disease probands

**Why Not Relevant**: The provided paper excerpt does not mention ARHGAP6, nor does it present any data (variant counts, de novo or transmitted genotypes, enrichment results, or mechanistic descriptions) tied to ARHGAP6. The manuscript focuses on exome sequencing of a large congenital heart disease (CHD) cohort and reports specific genes (e.g., GDF1, MYH6, FLT4) and aggregate gene-set results, but ARHGAP6 is not named in the supplied text. Therefore there is no direct evidence (observations of pathogenic variants, statistical association, or case reports) linking ARHGAP6 to any Mendelian disease in the provided content. Mechanistic evidence is also absent: no functional studies, pathway analyses, or gene-specific discussions of ARHGAP6 appear in the excerpt. Caveats: (1) the supplied content appears to be a portion of the full paper (main text and some table fragments); ARHGAP6 might hypothetically appear elsewhere (full text, other sections, figures, or supplementary tables/excel files) that were not provided. (2) Absence of mention in this excerpt does not prove ARHGAP6 is not associated with a Mendelian disease — it only means this paper excerpt provides no supporting evidence. Recommended next steps to evaluate the claim: search the full published paper (including all supplements and Excel tables) for ARHGAP6; check gene-disease resources such as OMIM, ClinVar, HGMD, and gnomAD/ExAC constraint metrics (pLI, LOEUF); and search the literature for case reports or functional studies implicating ARHGAP6 in Mendelian phenotypes.


[Read Paper](https://www.semanticscholar.org/paper/e5830d9e95469f22518560007746202e62db26a0)


### ID: TMG033 Clinical Utility of a Large Exome-Based Panel in the Genetic Diagnosis of Autism Spectrum Disorders and Intellectual Disabilities

**Why Not Relevant**: The provided paper content does not mention ARHGAP6 anywhere and contains no data (no variants, no case reports, no functional studies) pertaining to ARHGAP6. The text focuses on an ASD/ID diagnostic exome panel (describing capture, sequencing, and diagnostic yield) and on de novo missense mutations in DDB1 (with references to CUL4 and PHIP1). There is no gene-level list included in the excerpt that would allow inference that ARHGAP6 is among the tested genes. Therefore there is neither direct evidence (case reports, pathogenic variants, segregation data) nor mechanistic evidence (biological function, pathways, model systems) in this document that would support or refute the claim that ARHGAP6 is likely associated with a Mendelian disease. Limitations and caveats: (1) This is a set of abstracts/summary content rather than a comprehensive gene-disease review — absence of ARHGAP6 here does not prove absence of an association in the broader literature. (2) The panel described contains 1,905 genes but the specific gene list is not shown in the excerpt; ARHGAP6 could conceivably be on that panel but there is no evidence presented about variants in it. Suggested next steps to evaluate the claim: search databases and resources that curate gene-disease relationships (OMIM, ClinVar, ClinGen Gene-Disease Validity, DECIPHER), review PubMed for ARHGAP6 case reports or functional studies, and check variant frequency/constraint metrics in gnomAD and any reported pathogenic variants in locus-specific databases.


[Read Paper](https://www.semanticscholar.org/paper/f080f99ce279441d5248790319ba7f2085ed5ce2)


## Search Queries Used

- ARHGAP6 ARHGAP 6 Rho GTPase activating protein 6 Mendelian disease monogenic inherited disorder case report family segregation clinical phenotype

- ARHGAP6 mutation pathogenic variant loss of function nonsense frameshift missense de novo germline exome sequencing whole exome sequencing trio study sequencing panel

- ARHGAP6 RhoGAP6 Rho GTPase activating protein 6 knockout mouse zebrafish morpholino CRISPR functional assay cytoskeleton actin regulation neuronal development phenotype

- ARHGAP6 gene disease association OMIM ClinVar gene curation gene disease validity systematic review meta analysis rare disease database


## Bibliometric Configuration

- Use Bibliometrics: True

- Author Impact Weight: 0.4

- Citation Impact Weight: 0.4

- Venue Impact Weight: 0.2
