## Supplementary material for "MENDELSEEK: An algorithm that predicts Mendelian Genes and elucidates what makes them special": ARID5B.txt

# Claim: The gene ARID5B is likely associated with at least one Mendelian disease.

**Status**: processed

**Overall Rating**: -1

**Explanation**:

No relevant papers were found that support or refute this claim.


## Relevant Papers


## Other Reviewed Papers


### De novo mutations in autosomal recessive congenital malformations

**Why Not Relevant**: The provided paper content does not mention ARID5B or provide any gene-level data that could be attributed to ARID5B. The single sentence given describes a study design and a high-level result: that, among nine terminated pregnancies with severe fetal malformation, trio-based exome sequencing identified two unrelated cases in which an autosomal recessive disorder was caused by the combination of one de novo and one inherited variant. This statement contains no gene names, no variant descriptions, no segregation data, no functional or mechanistic evidence specific to ARID5B, and no statistical measures related to any particular gene. Therefore it supplies no direct evidence that ARID5B is associated with a Mendelian disease.

Mechanistic relevance is equally absent: while the sentence describes a general genetic mechanism (compound genotype of one de novo plus one inherited variant causing an autosomal recessive disorder), this is a general principle and cannot be used to infer involvement of ARID5B without explicit mention of ARID5B variants, pathogenicity assessments, segregation, or functional validation. Establishing that ARID5B is likely associated with at least one Mendelian disease would require identification of (a) pathogenic or likely pathogenic variants in ARID5B in affected individuals, (b) segregation with disease in families or recurrence in unrelated affected individuals, and ideally (c) supportive functional/mechanistic data — none of which are present in the supplied excerpt.

Limitations: the excerpt is a single summary sentence and may not reflect the full paper; ARID5B could appear elsewhere in the full text, supplementary material, tables, or figures. However, based solely on the provided content, there is no evidence for or against the claim about ARID5B. To properly evaluate the claim, the full paper (or any sections listing identified genes/variants and their annotations) would be needed.


[Read Paper](https://www.semanticscholar.org/paper/a2ba92abb5d06fff00c2b0d81291ff611c496f38)


### Molecular mechanisms of ARID5B-mediated genetic susceptibility to acute lymphoblastic leukemia.

**Why Not Relevant**: This paper provides strong evidence that common, non-coding variants at the ARID5B locus are associated with risk of childhood acute lymphoblastic leukemia and gives mechanistic insight into how a top risk allele (rs7090445) alters transcription factor binding and ARID5B regulation. However, the claim being evaluated is that ARID5B is likely associated with at least one Mendelian disease (i.e., a monogenic disorder with Mendelian inheritance). The study focuses on common-variant, polygenic susceptibility to a complex disease and uses an additive case–control association framework; it explicitly reports 54 common, non-coding risk variants and does not report rare, high-penetrance coding or loss-of-function variants, familial segregation, de novo mutations, or inheritance patterns consistent with a Mendelian disorder. Mechanistic experiments demonstrate regulatory effects in ALL cells but do not demonstrate that germline ARID5B mutations cause a Mendelian syndrome. Therefore, while the paper supports ARID5B as a locus influencing complex-trait disease risk and hematologic traits at the population level, it does not provide direct or robust evidence that ARID5B is associated with a Mendelian disease. Key limitations preventing relevance to a Mendelian claim: (1) focus on common non-coding variants with population-level effects, (2) absence of reported rare pathogenic variants with segregation data, and (3) analytical design (additive association testing) that is not targeted to detect Mendelian inheritance.


[Read Paper](https://www.semanticscholar.org/paper/860ca99eecc0f48bb862ccea0f0847f672e66371)


### De novo mutation of NAXE (APOAIBP)-related early-onset progressive encephalopathy with brain edema and/or leukoencephalopathy-1: A case report

**Why Not Relevant**: The paper exclusively investigates a severe neurometabolic disorder (PEBEL1) caused by NAXE (APOA1BP) biallelic variants. All reported genetic findings, clinical correlation, and conclusions center on NAXE; ARID5B is not mentioned anywhere in the provided content. Therefore the paper provides no direct genetic, clinical, or mechanistic evidence either supporting or refuting the claim that ARID5B is likely associated with at least one Mendelian disease. Absence of ARID5B discussion means this paper cannot be used as evidence for the claim. Note that lack of mention is not proof that ARID5B is not disease-associated, only that this manuscript offers no relevant data.


[Read Paper](https://www.semanticscholar.org/paper/5a593d0a440a88cd34e0a92d055a797c68cbc328)


### Case Report: The Monogenic Familial Steroid-Resistant Nephrotic Syndrome Caused by a Novel Missense Mutation of NPHS2 Gene A593C in a Chinese Family

**Why Not Relevant**: The paper exclusively reports on NPHS2 pathogenic variation causing autosomal recessive steroid-resistant nephrotic syndrome in a single consanguineous family and provides clinical, pathological, and mechanistic information specific to NPHS2/podocin. There is no mention of ARID5B anywhere in the provided content, nor are there data, analyses, or mechanistic discussions that implicate ARID5B in Mendelian disease. The trio whole-exome sequencing result identifies a causative NPHS2 variant and explicitly states no other SRNS-related gene variants were determined, but the text does not specify screening results for ARID5B or make any inference about ARID5B’s disease association. Therefore this paper does not provide direct evidence (no reported ARID5B variants co-segregating with disease) nor mechanistic evidence (no functional data, pathways, or phenotypes connected to ARID5B) to support the claim that ARID5B is likely associated with a Mendelian disease. Note that absence of mention in this paper does not disprove ARID5B involvement elsewhere; it only means this paper is not informative for the claim.


[Read Paper](https://www.semanticscholar.org/paper/26ba0a4e7e5097751e78e184e30cde938546d66d)


### SPG9A with the new occurrence of an ALDH18A1 mutation in a CMT1A family with PMP22 duplication: case report

**Why Not Relevant**: The provided paper content discusses a de novo mutation or gonadal mosaicism of ALDH18A1 in an SPG9A family and asks for further study of amino acid levels to understand P5CS deficiency. It does not mention ARID5B, any variants in ARID5B, phenotypes associated with ARID5B, or mechanistic pathways involving ARID5B. Therefore the content supplies no direct or mechanistic evidence for the claim that ARID5B is likely associated with at least one Mendelian disease. Caveat: this conclusion is based solely on the single sentence provided; if the full paper contains additional sections that mention ARID5B, those were not available for review here.


[Read Paper](https://www.semanticscholar.org/paper/c52899f94d0ef2b838ffdba1e3484424d8c18855)


### Germline mosaicism in a collagen VI‐related myopathy family: A cause of autosomal recessive inheritance

**Why Not Relevant**: The provided paper text focuses entirely on germline mosaicism and an autosomal recessive collagen VI-related myopathy caused by variants in the COL6A1 gene (with additional contextual mention of COL6A2 and COL6A3 and other examples such as COL4A4). There is no mention of the gene ARID5B, no data or analyses concerning ARID5B, and no mechanistic discussion that links ARID5B to Mendelian disease. Key topics in the text include: description of paternal germline mosaicism of a COL6A1 c.1057-2A>G variant, recurrence of Ullrich congenital muscular dystrophy in siblings, trio whole-exome sequencing results for COL6A1 variants (c.525delT and c.1057-2A>G), and implications of germline mosaicism for recurrence risk and carrier screening. Because the gene of interest in the claim (ARID5B) does not appear anywhere in the provided content, there is neither direct evidence (e.g., case reports, variant-disease segregation, statistical association) nor mechanistic evidence (e.g., functional studies, pathway connections) to support or refute the claim that ARID5B is likely associated with at least one Mendelian disease.

Consequently, no verbatim sentences or paragraphs from the provided text can be extracted that would support or refute the ARID5B–Mendelian disease claim. While the paper illustrates general genetic concepts (germline mosaicism, autosomal recessive inheritance, use of trio WES, and variant segregation in COL6A1), these generalities cannot be used to infer an association for a different, unmentioned gene. Inferring a Mendelian-disease association for ARID5B would require data specifically about ARID5B (clinical cases with segregating variants, functional/biochemical evidence, statistical burden tests, or literature linking ARID5B mutations to disease phenotypes), none of which are present in the provided content. Therefore the relevance of this paper to the ARID5B claim is effectively zero.


[Read Paper](https://www.semanticscholar.org/paper/4572f635bb354974678a75579201d9eedc2c40ea)


### Genome-Wide Association Study of Acute Lymphoblastic Leukemia in Hispanic/Latino Children Identifies a Putatively Novel Risk Locus at Chromosome 5q31.1

**Why Not Relevant**: This paper provides strong evidence that ARID5B harbors common genetic variants associated with susceptibility to childhood acute lymphoblastic leukemia (ALL) from GWAS. However, the claim is about ARID5B being "likely associated with at least one Mendelian disease," which implies a monogenic, typically rare and highly penetrant disorder caused by pathogenic variants in ARID5B. The study design and content do not address rare, high-penetrance variants or familial segregation required to support a Mendelian relationship: (1) the GWAS explicitly excluded variants with MAF <0.01, so it would not detect the rare variants that commonly underlie Mendelian disorders; (2) the associations reported are common-variant, modest-effect size (e.g., ORs ~1.3), characteristic of complex disease risk rather than Mendelian causation; and (3) there is no discussion of ARID5B loss-of-function variants, clinical case reports, segregation analyses, or functional experiments demonstrating monogenic disease mechanisms. Therefore, while the paper is directly relevant to ARID5B's role in complex-trait ALL susceptibility, it provides no direct or strong mechanistic evidence that ARID5B is the cause of any Mendelian disease. For these reasons the paper is of low relevance (0.05) to the specific claim.


[Read Paper](https://www.semanticscholar.org/paper/5a8adfd8ffe6d7b75534956e6b7f9f975ab38b05)


### Abstract 4516: Rare high-penetrance and common low-penetrance variants associated with risk of pediatric acute lymphoblastic leukemia

**Why Not Relevant**: The paper reports that ARID5B is a replicated locus for common (low-penetrance) germline association with pediatric ALL (a GWAS result), but it does not present evidence that ARID5B harbors rare pathogenic/likely pathogenic variants that would indicate a Mendelian (high-penetrance, monogenic) disease. The study explicitly screened 60 autosomal dominant cancer predisposition genes for P/LP variants and reports seven genes with enriched rare pathogenic variation; ARID5B is not among them. Therefore the paper provides relevant evidence that ARID5B is associated with ALL risk in a polygenic/common-variant sense, but no direct or mechanistic evidence that ARID5B is linked to any Mendelian disease. For these reasons the paper is of low relevance to the claim that ARID5B is likely associated with at least one Mendelian disease.


[Read Paper](https://www.semanticscholar.org/paper/7339a2bb7c17c1deb61734c523a33a392f1f57cb)


### Abstract A079: The impact of Indigenous American ancestry on the risk of acute lymphoblastic leukemia and the implications for future study designs

**Why Not Relevant**: The abstract addresses population-level enrichment and effect-size analyses of known ALL risk alleles across Indigenous American ancestry in Latino populations. It focuses on common-variant associations with a complex disease (acute lymphoblastic leukemia) and on methodological implications for GWAS in admixed populations. The text does not mention the gene ARID5B by name, nor does it present data about rare, high-penetrance variants, family segregation, loss-of-function alleles, or clinical syndromes—lines of evidence required to support that a gene is causative for a Mendelian disease. Therefore this paper provides no direct evidence that ARID5B is associated with a Mendelian disorder. Caveat: this is an abstract-level summary of a study of complex-trait risk; absence of ARID5B in the abstract does not prove that the gene has or lacks Mendelian associations elsewhere in the literature, but the provided content contains no information to support the claim.


[Read Paper](https://www.semanticscholar.org/paper/0e7c56504dda51f4bfc325d1e6fa653c1c359085)


### Abstract 2002: A genome-wide association study identifies novel sepsis risk loci in children with Down syndrome-associated acute lymphoblastic leukemia: A report from the Children’s Oncology Group

**Why Not Relevant**: The paper provides no direct or mechanistic evidence relevant to the claim that ARID5B is likely associated with at least one Mendelian disease. The document is a GWAS abstract focused on common-variant associations for sepsis risk in children with Down syndrome-associated ALL. Key reasons the paper is not relevant to the ARID5B–Mendelian disease claim: (1) ARID5B is not mentioned anywhere in the content; none of the reported loci or genes (TMEM163, ZFAND3, APOBEC3A, etc.) is ARID5B. (2) The study analyzed autosomal common variants with MAF >1% and excluded chromosome 21; Mendelian disease associations are typically driven by rare, high-penetrance variants which this study design would not detect. (3) The phenotype under study is treatment-related sepsis in a specific clinical cohort (DS-ALL), not familial segregation, rare variant discovery, or functional assays to establish causality for a Mendelian condition. (4) This is an abstract reporting preliminary GWAS associations with a small cohort (N=264) and describes confirmatory studies as ongoing; abstracts typically lack the depth (rare-variant analysis, segregation, in vitro/in vivo functional follow-up) required to support a claim about Mendelian disease causality. For these reasons, relevance to the ARID5B–Mendelian disease claim is effectively nil.


[Read Paper](https://www.semanticscholar.org/paper/91c47d0044654ab5322f858bb86bf8257714436c)


## Search Queries Used

- ARID5B germline mutation Mendelian disease monogenic autosomal dominant autosomal recessive de novo case report clinical exome sequencing

- ARID5B loss of function knockout mouse phenotype Arid5b null embryonic lethal developmental delay transcription factor chromatin remodeling functional study

- ARID5B ClinVar OMIM DECIPHER gnomAD pLI loss of function intolerance constraint pathogenic variants clinical significance

- ARID5B GWAS childhood acute lymphoblastic leukemia common variant risk association population study genetic association comparison monogenic evidence pleiotropy


## Bibliometric Configuration

- Use Bibliometrics: True

- Author Impact Weight: 0.4

- Citation Impact Weight: 0.4

- Venue Impact Weight: 0.2
