## Supplementary material for "MENDELSEEK: An algorithm that predicts Mendelian Genes and elucidates what makes them special": ASAH2.txt

# Claim: The gene ASAH2 is likely associated with at least one Mendelian disease.

**Status**: processed

**Overall Rating**: -1

**Explanation**:

No relevant papers were found that support or refute this claim.


## Relevant Papers


## Other Reviewed Papers


### The Burden of Candidate Pathogenic Variants for Kidney and Genitourinary Disorders Emerging From Exome Sequencing

**Why Not Relevant**: The provided paper content does not mention the gene ASAH2 by name, does not present any gene-level association results for ASAH2, nor does it discuss mechanistic evidence (molecular function, disease-causing variants, inheritance patterns) specific to ASAH2. The paper describes methods for identifying and curating genes associated with Mendelian kidney and genitourinary disorders, references databases (OMIM, Orphanet, ClinVar, HGMD), and notes a curated list of 625 genes accessible on a laboratory website or in appendices. However, without the actual curated gene list or any per-gene results in the supplied text, there is no direct evidence (case reports, variant associations, statistical signals) or mechanistic evidence provided here that would support or refute the claim that "ASAH2 is likely associated with at least one Mendelian disease." To evaluate that claim based on this study, one would need access to the supplementary gene list or the cited databases/appendices to see whether ASAH2 is included and, if so, what evidence (OMIM entry, ClinVar/HGMD pathogenic variants, inheritance mode, functional data) supports the association. Absent that, the relevance of this paper to the ASAH2 claim is minimal.


[Read Paper](https://www.semanticscholar.org/paper/a75e297208c04992568e607d19aa9b9aaebd09d5)


### Biallelic sequence and structural variants in RAX2 are a novel cause for autosomal recessive inherited retinal disease

**Why Not Relevant**: The provided paper content refers exclusively to the gene RAX2 and its role in recessive inherited retinal disease (IRD). The claim to be evaluated concerns the gene ASAH2 and whether it is likely associated with at least one Mendelian disease. The excerpt contains no mention of ASAH2, no experimental results, no mechanistic discussion, and no contextual information that links RAX2 findings to ASAH2. Based solely on the supplied content, there is no basis to support or refute the claim about ASAH2. If the full paper (not provided here) includes separate sections that mention ASAH2, those were not available for review; the relevance assessment above applies only to the text supplied.


[Read Paper](https://www.semanticscholar.org/paper/08e79066c138449120587d9b5111b8d9f8a01f2a)


### Biallelic variants in Plexin B2 (PLXNB2) cause amelogenesis imperfecta, hearing loss and intellectual disability

**Why Not Relevant**: The provided paper content focuses entirely on plexin family genes and specifically identifies PLXNB2 (plexin B2) as a Mendelian disease gene. There is no mention of ASAH2 anywhere in the supplied text, and no data (genetic, expression, functional, mechanistic, or segregation) concerning ASAH2 are presented. Therefore this paper provides no direct or mechanistic evidence that ASAH2 is associated with any Mendelian disease. Furthermore, evidence that establishes PLXNB2 as a disease gene (biallelic pathogenic variants segregating with disease, expression in relevant cell types, phenotype overlap with knockout animals) cannot be extrapolated to a different gene (ASAH2) without independent data. Note: assessment is limited to the content you provided; if the full paper contains sections not included here that mention ASAH2, those were not available for review.


[Read Paper](https://www.semanticscholar.org/paper/611d1262ba83537ba6cee109d4704fb0ab033e85)


### Distinct neurological phenotypes associated with biallelic loss of NOTCH3 function: evidence for recessive inheritance.

**Why Not Relevant**: The provided paper content does not mention ASAH2 anywhere. The single sentence only reports that the studied early-onset leukoencephalopathy appears to follow an autosomal recessive inheritance pattern and contrasts this with CADASIL's dominant gain-of-function mechanism. There is no gene name, no variant data, no segregation analysis tied to ASAH2, and no biochemical or cellular mechanism involving ASAH2 presented. Because the claim concerns whether ASAH2 is likely associated with at least one Mendelian disease, and the excerpt contains no information about ASAH2 (nor any evidence that could be mapped to ASAH2), the content is not relevant to the claim. Any inference that ASAH2 is implicated would be speculative and unsupported by the provided text. Additionally, mentioning CADASIL refers to a different disease mechanism and different gene(s), which does not bear on ASAH2 without explicit linkage. Therefore relevance is effectively zero.


[Read Paper](https://www.semanticscholar.org/paper/95bc213ad299f037a07a15d426f8cba70ff00aa2)


### Exome sequence analysis identifies a homozygous, pathogenic, frameshift variant in the MAN2B1 gene underlying clinical variant of α-mannosidosis

**Why Not Relevant**: The paper exclusively investigates α-mannosidosis and reports pathogenic, segregating biallelic variants in MAN2B1 as the cause in the studied families. The gene ASAH2 is not mentioned anywhere in the provided text, and no data, variants, phenotypes, or mechanistic analyses involving ASAH2 are presented. Therefore the paper provides no direct genetic evidence (no observed ASAH2 variants segregating with disease), nor any mechanistic evidence (no discussion of ASAH2 protein function, pathways, or plausible disease mechanisms) to support or refute the claim that ASAH2 is likely associated with at least one Mendelian disease. For these reasons the relevance of this paper to the claim is 0.0.


[Read Paper](https://www.semanticscholar.org/paper/455efa329ae9a2ace4b63d745696b455810b8e46)


### Biallelic loss-of-function variants in RAX2, encoding a homeobox-containing Rax transcription factor, cause autosomal recessive inherited retinal disease

**Why Not Relevant**: The paper content provided is entirely focused on the gene RAX2 and its role in autosomal recessive retinitis pigmentosa and contains no mention of ASAH2. Example verbatim sentences from the supplied text that demonstrate this focus include:

"RAX2 encodes a member of the homeobox-containing Rax family of transcription factors, which play a pivotal role in late retinogenesis in vertebrate species by regulating the spatial expression of photoreceptor-specific genes." 

"So far, only four monoallelic RAX2 variants have been tentatively implicated in autosomal dominant cone-dominated retinal disease." 

"In summary, our findings support a role for RAX2 as a novel disease gene for autosomal recessive RP." 

Because ASAH2 is not mentioned anywhere in the provided content, the paper provides no direct evidence (no genetic association data, no variant descriptions, no case reports) and no mechanistic evidence (no functional studies, pathways, or homology-based arguments) relevant to the claim that "ASAH2 is likely associated with at least one Mendelian disease." One cannot infer an association for ASAH2 from data on RAX2 because they are distinct genes with different functions and the paper does not report any genetic, clinical, or biochemical link between them. 

To evaluate the claim about ASAH2, one would need papers or database entries that directly report (a) pathogenic or likely pathogenic variants in ASAH2 segregating with Mendelian phenotypes, (b) functional studies showing a disease-causing mechanism for ASAH2 variants, or (c) curated entries in clinical genetics resources (e.g., OMIM, ClinVar) documenting ASAH2 as a disease gene. None of these are present in the provided text, so relevance to the ASAH2 claim is effectively zero.


[Read Paper](https://www.semanticscholar.org/paper/3f6d8a99a6cad3e50453c4497c622646c7beabb3)


### Mapping genetic variations to three-dimensional protein structures to enhance variant interpretation: a proposed framework

**Why Not Relevant**: The provided paper content is a workshop/report and field-review focused on methods, resources, and a proposed framework for integrating genetic variants with three-dimensional protein structures. The excerpt contains general descriptions of variant interpretation challenges, databases, and tools, and high-level mechanistic rationale for why structural mapping can help interpret variants. However, the excerpt contains no mention of the gene ASAH2, no data (case reports, cohort analyses, segregation studies, population frequency analyses, functional assays) concerning ASAH2 variants, and no statements asserting any association between ASAH2 and any Mendelian disease. Consequently, there is no direct evidence (such as statistically significant association, pathogenic variant descriptions, or clinical case reports) nor any ASAH2-specific mechanistic evidence presented in this paper content to support the claim that ASAH2 is likely associated with at least one Mendelian disease. 

Limitations of this assessment: absence of ASAH2 in this paper excerpt does not prove ASAH2 has no Mendelian associations; it only shows this particular paper does not provide supporting evidence. To evaluate the claim one would need gene-specific sources: clinical variant databases (e.g., ClinVar entries for ASAH2), primary genetic studies, case reports, or functional studies demonstrating pathogenic ASAH2 variants and segregation with disease. The paper does describe general methods that could be applied to ASAH2 in future work, but no ASAH2-specific results are included in the provided content.


[Read Paper](https://www.semanticscholar.org/paper/ae68aa934d5a5ad3d9009bb8811fa8d5fc4dbb1c)


## Search Queries Used

- ASAH2 variant Mendelian disease case report exome sequencing family segregation biallelic autosomal recessive autosomal dominant loss of function

- ASAH2 ClinVar OMIM HGMD review systematic review gene disease association clinical curation pathogenic variant annotation

- ASAH2 neutral ceramidase N acylsphingosine amidohydrolase 2 sphingolipid metabolism ceramide hydrolysis functional study knockout mouse zebrafish cell model phenotype neurodegeneration

- ASAH2 rare variant burden exome sequencing cohort Mendelian disease gene discovery linkage analysis family based study gene panel loss of function constraint ASAH1 paralog differentiation


## Bibliometric Configuration

- Use Bibliometrics: True

- Author Impact Weight: 0.4

- Citation Impact Weight: 0.4

- Venue Impact Weight: 0.2
