## Supplementary material for "MENDELSEEK: An algorithm that predicts Mendelian Genes and elucidates what makes them special": BACE1.txt

# Claim: The gene BACE1 is likely associated with at least one Mendelian disease.

**Status**: processed

**Overall Rating**: -1

**Explanation**:

No relevant papers were found that support or refute this claim.


## Relevant Papers


## Other Reviewed Papers


### Precision Medicine in Internal Medicine

**Why Not Relevant**: This paper is a general review of clinical genetic testing strategies, variant types, and interpretation frameworks. It contains no empirical data, case reports, gene-specific analyses, or database queries that mention BACE1 or evaluate variants in BACE1. Thus it provides neither direct evidence for nor direct evidence against the claim that BACE1 is associated with at least one Mendelian disease. The paper's content is relevant only at a methodological and conceptual level (e.g., describing how exome sequencing could discover gene–disease associations, defining truncating variants and loss-of-function, and noting that only ~18% of protein-coding genes were linked to Mendelian disorders as of 2019). Those points affect plausibility and context but do not substitute for gene-specific association data. Given the absence of any BACE1-specific information in the provided text, the paper cannot be used to support the claim; it only informs what kinds of evidence would be needed and why some genes remain unassociated in the literature/databases. Therefore the relevance score is very low (<0.1).


[Read Paper](https://www.semanticscholar.org/paper/c7199e8a5226f8d9d55777da0e15e91797bb459a)


### A Novel CLN6 Variant Associated With Juvenile Neuronal Ceroid Lipofuscinosis in Patients With Absence of Visual Loss as a Presenting Feature

**Why Not Relevant**: This paper focuses entirely on CLN6 and its role in neuronal ceroid lipofuscinoses (Batten disease). Multiple sentences explicitly state that mutations in CLN6 cause juvenile/late-infantile/adult-onset NCL and describe CLN6's cellular role and the specific CLN6 variants identified in affected families. The paper contains no mention of BACE1, no genetic or functional data concerning BACE1, and no discussion that would link BACE1 to any Mendelian phenotype. Therefore it provides no direct genetic evidence (variant identification, segregation, or case reports) nor mechanistic evidence (functional perturbation, pathway connection, or model organism data) that BACE1 is associated with a Mendelian disease. The absence of any mention of BACE1 in this study does not prove BACE1 is not associated with Mendelian disease, but based on this paper alone there is no support for the claim. To evaluate the claim about BACE1 would require papers that specifically report pathogenic BACE1 variants segregating with disease, functional validation, or authoritative gene-disease curation resources.


[Read Paper](https://www.semanticscholar.org/paper/541fb134c7e3f8dfbe116f47402985171d2b765b)


### New variant in deficiency of interleukin‐36 receptor antagonist syndrome (DITRA)

**Why Not Relevant**: This paper focuses exclusively on DITRA syndrome and pathogenic variants in the IL36RN gene. It contains no mention of the gene BACE1, no reported BACE1 variants, and no mechanistic discussion connecting BACE1 to any Mendelian phenotype. Although the authors performed mass sequencing on a panel of 4,427 genes (so BACE1 could conceivably have been included on the panel), the paper reports only IL36RN findings; absence of reporting of BACE1 data in this manuscript means there is no evidence here to support the claim that BACE1 is likely associated with a Mendelian disease. Note the logical limitation: a single paper not addressing a gene provides no evidence either for or against that gene's disease association — it is simply silent. Therefore, the correct assessment is that this paper is not relevant to the claim about BACE1.


[Read Paper](https://www.semanticscholar.org/paper/8ddda6aca3434df8829aee423d53ce69e59841dc)


### Pontocerebellar Hypoplasia Type 9: A New Case with a Novel Mutation and Review of Literature

**Why Not Relevant**: The paper exclusively investigates Pontocerebellar hypoplasia type 9 and identifies AMPD2 loss-of-function variants (including a novel c.1471G > A, p.Gly491Arg) as causal in the presented patient. All mechanistic and functional data concern AMPD2 (expression pattern, role in purine biosynthesis, enzyme activity reduction). The manuscript contains no mention of BACE1, no genetic, functional, or clinical data involving BACE1, and therefore provides no direct or mechanistic evidence to support or refute the claim that BACE1 is associated with at least one Mendelian disease. Absence of evidence in this paper does not disprove the claim, but the paper is irrelevant to establishing BACE1 as a Mendelian disease gene.


[Read Paper](https://www.semanticscholar.org/paper/9bda312f04b150ba923988a961bc394307daca34)


### Discovery of a Novel Shared Variant Among RTEL1 Gene and RTEL1-TNFRSF6B lncRNA at Chromosome 20q13.33 in Familial Progressive Myoclonus Epilepsy

**Why Not Relevant**: This paper does not mention BACE1 anywhere in the provided content. The study focuses on progressive myoclonus epilepsy (PME) in two siblings and reports candidate variants in KCNH8, RTEL1, RTEL1-TNFRSF6B, and SLC22A17 identified by WES and computational analyses. Although WES was performed (which could in principle detect variants in BACE1), the authors did not report any BACE1 variants or discuss BACE1-related mechanisms or diseases. Therefore the paper provides no direct evidence for or against the claim that BACE1 is likely associated with at least one Mendelian disease. Important limitations: absence of BACE1 discussion in this PME-focused, small-family study does not constitute evidence that BACE1 is not associated with any Mendelian disease elsewhere. To evaluate the claim about BACE1 you would need papers that specifically report pathogenic BACE1 variants segregating with Mendelian phenotypes, functional validation, or population-level burden analyses—none of which are present in this manuscript.


[Read Paper](https://www.semanticscholar.org/paper/c62784939aad39b807ef0d0a75076891bc28ca41)


### Beyond APOL 1 : Genetic Inroads into Understanding Population Disparities in Diabetic Kidney Disease

**Why Not Relevant**: This paper does not mention BACE1, nor does it present any genetic, clinical, experimental, or mechanistic data concerning BACE1. The content focuses on APOL1, CUBN, LRP2 (megalin), AMN, and related loci in the context of kidney disease and albuminuria, and it explicitly describes Mendelian disorders linked to CUBN, AMN, and LRP2. Because there is no discussion, analysis, or data about BACE1 in the provided text, there is no direct or mechanistic evidence in this paper to support or refute the claim that BACE1 is likely associated with at least one Mendelian disease. Therefore the paper's relevance to the specific claim about BACE1 is effectively zero.


[Read Paper](https://www.semanticscholar.org/paper/52d3f3490311ed466b2327ad26b88d847ffd485b)


## Search Queries Used

- BACE1 beta secretase 1 Mendelian disease monogenic disorder OMIM ClinVar gene disease association

- BACE1 mutation variant pathogenic loss of function biallelic homozygous de novo case report patient exome sequencing segregation

- BACE1 knockout mouse phenotype human phenocopy neurodevelopment myelination synaptic function APP processing amyloid beta mechanism single gene disorder

- BACE1 Alzheimer disease complex trait rare variant burden study genome wide association review meta analysis Mendelian versus complex inheritance


## Bibliometric Configuration

- Use Bibliometrics: True

- Author Impact Weight: 0.4

- Citation Impact Weight: 0.4

- Venue Impact Weight: 0.2
