## Supplementary material for "MENDELSEEK: An algorithm that predicts Mendelian Genes and elucidates what makes them special": BAIAP3.txt

# Claim: The gene BAIAP3 is likely associated with at least one Mendelian disease.

**Status**: processed

**Overall Rating**: -1

**Explanation**:

No relevant papers were found that support or refute this claim.


## Relevant Papers


## Other Reviewed Papers


### Needle in a Haystack: The Common Can Inform the Rare in Restless Legs Syndrome

**Why Not Relevant**: The supplied paper excerpt does not mention the gene BAIAP3, any variants in BAIAP3, nor any experimental or statistical results pertaining to BAIAP3. The content discusses general issues in restless legs syndrome (RLS) genetics, comparisons between GWAS and rare-variant (Mendelian) approaches, and an example (LRRK2) of a known Mendelian cause for a different neurological disease. While these general statements describe how one would identify Mendelian causes and the rationale for rare-variant burden testing, they provide no direct or mechanistic evidence about BAIAP3 specifically. Therefore, based on the provided text, there is no evidence to support or refute the claim that BAIAP3 is likely associated with at least one Mendelian disease. Note: if additional sections of the paper (methods, results, supplementary tables) that explicitly reference BAIAP3 are made available, I can re-evaluate and extract relevant verbatim excerpts and their implications.


[Read Paper](https://www.semanticscholar.org/paper/a32e565d62c1422ae2d6a6aa316fa63f0c5070f1)


### TTC7A Variants Previously Described to Cause Enteropathy Are Observed on a Single Haplotype and Appear Non-pathogenic in Pediatric Inflammatory Bowel Disease Patients

**Why Not Relevant**: The provided paper content refers exclusively to variants in the Tetratricopeptide Repeat Domain 7A (TTC7A) gene and evaluates TTC7A variation in relation to polygenic inflammatory bowel disease (IBD). There is no mention of BAIAP3, no reported variants in BAIAP3, and no mechanistic discussion linking TTC7A to BAIAP3. Therefore the excerpted content supplies no direct evidence (no case reports, statistical associations, or segregation data) nor mechanistic evidence (no pathway, functional, or interaction data) supporting or refuting the claim that BAIAP3 is likely associated with a Mendelian disease. Limitation: only a single-sentence summary of the study was provided; if the full article contains additional sections not included here that mention BAIAP3, those were not available for review. Based on the provided content alone, the paper is not relevant to the claim about BAIAP3.


[Read Paper](https://www.semanticscholar.org/paper/3818a966b5d6e8b0eda65fdef043ac0727fa429c)


### HOXA11 is another monogenic cause of congenital anomalies of the kidney and urinary tract

**Why Not Relevant**: The provided paper content does not mention BAIAP3 or any synonyms, and therefore provides no direct or mechanistic evidence bearing on the claim that BAIAP3 is likely associated with a Mendelian disease. The only sentence supplied is: "Adding HOXA11 to the list of monogenic causes of CAKUT may hopefully encourage physicians and researchers to look for HOxA11 variants in selected cases for prompt genetic diagnosis and genetic counselling, as well as to study the molecular pathogenesis of HOX a11-related disease in humans withCAKUT." This sentence concerns HOXA11 and congenital anomalies of the kidney and urinary tract (CAKUT), not BAIAP3. There are no data, variant descriptions, functional experiments, mechanistic pathways, statistical results, or conclusions in the provided content that mention BAIAP3 or permit inference about BAIAP3's involvement in Mendelian disease. Limitations: my assessment is limited to the text you provided; if the full paper contains other sections not included here that mention BAIAP3, those were not available for review. Based on the supplied content alone, relevance to the BAIAP3–Mendelian disease claim is effectively nil.


[Read Paper](https://www.semanticscholar.org/paper/db3d83a93884ccd5d66fca19ab8728ccabfc9ffc)


### OP0309 WES ANALYSIS OF RARE FAMILIES POINTS TO A PATHOGENIC ROLE FOR THE cGAS/STING-TYPE I IFN AXIS IN SSc

**Why Not Relevant**: The provided paper content does not mention the gene BAIAP3 anywhere, nor does it provide any gene list or data that would allow identification of BAIAP3 as one of the candidate genes. Although the paper states that "Mutations in two of our candidates cause Mendelian diseases," it does not name those genes in the provided excerpts; the named genes with functional evidence are TMEM173/STING, NT5E/CD73 and PRKD2, none of which are BAIAP3. The study focuses on familial systemic sclerosis and implicates cGAS/STING–type I IFN pathway genes; there is no direct or mechanistic evidence in the supplied text linking BAIAP3 to any Mendelian disease. Because of the absence of any mention or data about BAIAP3 in the provided content, the paper is not relevant (relevance score 0.02) to the claim that BAIAP3 is likely associated with at least one Mendelian disease.


[Read Paper](https://www.semanticscholar.org/paper/a5d1937aaf7ad488b3d5d4df3648ee2ca8966c0d)


### The UK10K project identifies rare variants in health and disease

**Why Not Relevant**: The provided paper content (UK10K project) describes a large-scale sequencing resource and reports broad findings (discovery of novel variants, identification of many genes carrying predicted loss-of-function alleles, and inclusion of rare-disease cohorts). However, the excerpt contains no mention of the gene BAIAP3, no gene-specific association results, and no mechanistic data linking BAIAP3 to any Mendelian phenotype. Therefore there is no direct evidence in this text that BAIAP3 is associated with a Mendelian disease. It is possible BAIAP3 could be among the genes with LoF variants detected in the full dataset or supplementary materials, but that cannot be inferred from the supplied excerpt. Absence of mention here is not definitive evidence against the claim; it only means this paper extract neither supports nor refutes the claim about BAIAP3. To evaluate the claim further would require checking gene-level results/supplementary tables from this study or searching dedicated clinical and variant databases (OMIM, ClinVar, HGMD), gnomAD constraint metrics (pLI), and the primary literature for reports of pathogenic variants in BAIAP3.


[Read Paper](https://www.semanticscholar.org/paper/439b20628573147c514d93e0feb5140812cf6f34)


## Search Queries Used

- BAIAP3 BAI1 associated protein 3 Mendelian disease monogenic rare disease case report exome sequencing segregation family study

- BAIAP3 ClinVar OMIM HGMD ClinGen gnomAD loss of function intolerance pLI constraint variant pathogenicity

- BAIAP3 functional study knockout mouse phenotype zebrafish knockdown neuronal dense core vesicle exocytosis secretory pathway BAI1 associated protein 3

- BAIAP3 de novo mutation developmental disorders Deciphering Developmental Disorders DDD exome sequencing trios cohort intellectual disability epilepsy autism gene discovery population frequency no association


## Bibliometric Configuration

- Use Bibliometrics: True

- Author Impact Weight: 0.4

- Citation Impact Weight: 0.4

- Venue Impact Weight: 0.2
