## Supplementary material for "MENDELSEEK: An algorithm that predicts Mendelian Genes and elucidates what makes them special": BDNF.txt

# Claim: The gene PPARGC1A is likely associated with at least one Mendelian disease.

**Status**: processed

**Overall Rating**: 0

**Explanation**:

Supporting evidence:
The provided paper documents robust biological roles for PPARGC1A (PGC-1α). It is expressed in high-oxidative tissues (brain, muscle) and acts as a transcriptional coactivator regulating many genes in response to metabolic cues (exercise, fasting). Tissue-specific deletion in mice produces clear phenotypes: AgRP-neuron deletion alters feeding, fat mass, locomotion and thermoregulation; muscle-specific deletion leads to loss of mitochondrial function, impaired calcium homeostasis, increased apoptosis, premature sarcopenia, and altered responses to ER stress. These functional data show that loss of PGC-1α activity can cause substantial organismal and tissue-specific dysfunction in model systems, establishing biological plausibility that severe disruption could produce a disease phenotype.

Caveats or contradictory evidence:
None of the excerpts report human genetic evidence linking PPARGC1A to a Mendelian disorder. The paper describes mouse knockouts and cell models, metabolic regulation, and age-associated pathology, but does not present human pedigrees, segregation analyses, pathogenic variants, or clinical descriptions consistent with a Mendelian inheritance pattern. Animal-model phenotypes and mechanistic insights do not by themselves demonstrate that PPARGC1A causes a Mendelian disease in humans. The distinction between a gene being biologically important and a gene being a known cause of a Mendelian disorder is critical; the provided evidence does not cross that threshold.

Analysis of mechanisms and second-order implications:
Mechanistically, PPARGC1A influences mitochondrial biogenesis, calcium handling, and transcriptional programs for oxidative metabolism. Loss-of-function variants in a gene with these roles could plausibly produce monogenic phenotypes (e.g., mitochondrial myopathy-like presentations, neuroendocrine/metabolic syndromes) if variants arise that severely disrupt protein function and segregate in families. However, plausibility does not equal documented causality. Many genes essential for cellular metabolism have no reported Mendelian human disease either because loss is embryonic-lethal, because pathogenic variants are rare and underascertained, or because dysfunction typically contributes as a risk factor for complex diseases rather than producing a clear Mendelian syndrome. The provided data do not address which of those possibilities applies to PPARGC1A in humans.

Which rating is most appropriate:
Given the evidence provided, the strongest conclusion is that there is no direct evidence in these excerpts that PPARGC1A is associated with any human Mendelian disease. The paper supplies supporting functional and animal-model data that make such an association biologically plausible, but it contains no human genetic or clinical evidence necessary to support the claim. Therefore the appropriate assessment is that there is no evidence (in the supplied material) to support the claim that PPARGC1A is likely associated with at least one Mendelian disease.


**Final Reasoning**:

I reviewed the logical steps: the claim asserts PPARGC1A is likely associated with at least one Mendelian disease. The provided paper offers mechanistic and mouse-model evidence that PGC-1α is important for metabolic and muscle physiology, which increases biological plausibility that severe disruption could cause disease. However, the definition of a gene being 'associated with a Mendelian disease' requires human genetic evidence (segregating pathogenic variants, clinical descriptions, database/OMIM entries, or equivalent). The paper excerpts contain no such data. Therefore the correct judgement based on the supplied evidence is that there is no evidence to support the claim. I find no internal inconsistency in this reasoning and reaffirm the rating below.


## Relevant Papers


### Neuronal control of energy balance and modulation of muscle aging by the transcriptional coactivator PGC-1α

**Authors**: Jonathan F. Gill (H-index: 7)

**Relevance**: 0.15

**Bibliometric Impact**: 0.05600000000000001


**Excerpts**:

- Cellular metabolic adaptations play a central role in the body's response to environmental changes and external stimuli and allow the maintenance of a proper energy balance. Transcriptional activators enable the integration of incoming signals and sensing of altered energy levels. Dysregulation of such metabolic pathways is a common mechanism of various tissue dysfunctions contributing to different diseases. A key player in cellular metabolism is the transcriptional coregulator Peroxisome proliferator-activated receptor coactivator 1 alpha (PGC-1α). PGC-1α is expressed in tissues with high basal oxidative capacity such as brain and muscle and regulates the expression of a plethora of genes in response to various external cues, including exercise and fasting. This makes PGC-1α an extremely powerful metabolic sensor and a potential therapeutic target in metabolic diseases.

- To determine the role of PGC-1α in central metabolic sensing and brain regulation of energy balance, we deleted the coactivator specifically in murine AgRP and POMC neurons located in the arcuate nucleus. The ablation of PGC-1α in POMC neurons did not reveal any major phenotype. Conversely, absence of PGC-1α in AgRP cells drove an increase in fat mass coupled with a reduction in locomotor activity and body temperature. Mice lacking the transcriptional coactivator in AgRP neurons exhibited a blunted leptin response and reduced food intake in fed and fasted conditions.

- Mechanistically, we demonstrated that fasting-induced AgRP expression is blunted in those mice and in an immortalized AgRP cell line, thereby leading to reduced feeding response upon fasting.

- To determine the role of PGC-1α in muscle aging, we used muscle-specific PGC-1α deletion and overexpression mouse models. We confirmed that PGC-1α preserves mitochondrial function and protein expression, as well as muscle endurance during aging. More importantly, we identified a novel function for PGC-1α in muscle, involving the regulation of calcium homeostasis through the control of genes responsible for mitochondrial calcium uptake and mitochondrial association with the sarcoplasmic reticulum.

- We suspect that PGC-1α-mediated amelioration of calcium metabolism protected the old muscle against ER stress and prevented tubular aggregate formation, which all contributed to reducing muscle apoptosis.

- Finally, we used muscle cells treated with ceramide or thapsigargin to demonstrate that PGC-1α inhibited apoptosis initiated by mitochondrial or calcium homeostasis impairments.

- We also showed that PGC-1α muscle deletion led to pre-mature sarcopenia.

- This work therefore improves our understanding of the biological processes regulated by PGC-1α and established the transcriptional regulator as a promising target for therapeutic approaches in metabolic and age-associated muscle diseases.


**Explanations**:

- This opening paragraph establishes that PGC-1α (encoded by PPARGC1A) is a central regulator of metabolism, is expressed in brain and muscle, and influences many downstream genes. Relation to claim: mechanistic plausibility. Strength: shows the gene has broad, important biological roles in tissues where monogenic defects might plausibly cause disease. Limitations: purely descriptive and functional, no human genetic or Mendelian disease data; general involvement in metabolism alone does not demonstrate a Mendelian disease association.

- These sentences describe conditional knockout experiments in mice (AgRP and POMC neurons) and report organismal phenotypes (increased fat mass, reduced locomotion and temperature, blunted leptin response). Relation to claim: mechanistic and functional evidence that loss of PGC-1α function causes physiological perturbations in mammals. Strength: in vivo loss-of-function in specific neuronal populations produces measurable systemic phenotypes, supporting biological relevance of the gene. Limitations: mouse conditional deletions are not equivalent to naturally occurring germline human mutations, the phenotypes are tissue- and model-specific, and no evidence is presented linking PPARGC1A variants to human Mendelian disease or demonstrating dominant/recessive inheritance patterns.

- This sentence provides a specific mechanistic observation that fasting-induced AgRP expression is blunted in PGC-1α-deficient mice and in an immortalized AgRP cell line, connecting the molecular effect (reduced AgRP expression) to the feeding phenotype. Relation to claim: mechanistic evidence. Strength: links gene perturbation to a defined molecular pathway (AgRP regulation) and cellular model confirmation. Limitations: still mechanistic and experimental (mouse and cell line); it does not demonstrate that human variants in PPARGC1A cause Mendelian disorders, nor does it quantify effect sizes or penetrance.

- This multi-sentence excerpt documents use of muscle-specific deletion and overexpression mouse models and reports that PGC-1α preserves mitochondrial function and muscle endurance, and regulates calcium-homeostasis genes linked to mitochondrial calcium uptake and SR association. Relation to claim: mechanistic evidence that loss or dysregulation of PPARGC1A affects core cellular processes (mitochondrial function, calcium handling) that, if severely disrupted, could underlie monogenic disease phenotypes. Strength: identifies concrete molecular processes and target gene sets under PGC-1α control, increasing biological plausibility for disease causation. Limitations: evidence is from engineered mouse models and does not include human genetic linkage, segregation, or case reports required to establish a Mendelian association.

- This sentence posits that improved calcium metabolism via PGC-1α protects aged muscle from ER stress and tubular aggregate formation, reducing apoptosis. Relation to claim: mechanistic evidence. Strength: ties PGC-1α action to cellular stress responses and pathology-relevant structures (tubular aggregates), suggesting possible mechanisms by which PPARGC1A dysfunction could cause disease. Limitations: phrasing is speculative ('We suspect'); evidence presented is inferential rather than direct demonstration, and again derived from animal/cellular models without human genetic correlation.

- This sentence reports that PGC-1α inhibits apoptosis in muscle cells subjected to mitochondrial or calcium stressors (ceramide or thapsigargin). Relation to claim: mechanistic evidence that PGC-1α activity can determine cell survival under pathological stresses, which could be relevant to disease phenotypes. Strength: experimental cellular demonstration of protective effect under defined insults. Limitations: in vitro cell treatments are artificial stress models; protective effect does not demonstrate that germline PPARGC1A variants cause Mendelian disease in humans.

- This short sentence reports that muscle-specific deletion of PGC-1α led to premature sarcopenia in mice. Relation to claim: functional/phenotypic evidence that loss of gene function produces a defined disease-like phenotype in an animal model. Strength: a clear organismal phenotype (sarcopenia) linked to gene deletion increases plausibility that severe perturbation of PPARGC1A could cause disease. Limitations: sarcopenia in an engineered mouse model is not direct proof of a human Mendelian disorder caused by PPARGC1A variants; no human genetic data, inheritance pattern, penetrance, or variant pathogenicity information is provided.

- This concluding sentence frames PPARGC1A/PGC-1α as a promising therapeutic target for metabolic and age-associated muscle diseases. Relation to claim: contextual, suggesting clinical relevance and that modulation of the gene affects disease-related phenotypes. Strength: summarizes translational potential based on mechanistic and animal-model data. Limitations: this is a translational statement and does not cite human genetic evidence of Mendelian disease association; therapeutic target status does not imply known monogenic disease causation.


[Read Paper](https://www.semanticscholar.org/paper/09b35b51b593e1cc4d26ebb005e358bd62ce91d9)


## Other Reviewed Papers


### ClinGen’s RASopathy Expert Panel Consensus Methods for Variant Interpretation

**Why Not Relevant**: The provided paper content discusses RAS-specific ACMG-AMP specifications and classification of RASopathy-associated variants within the Ras/MAPK pathway: "RAS-specific ACMG-AMP specifications optimized the utility of available clinical evidence and Ras/MAPK pathway–specific characteristics to consistently classify RASopathy-associated variants, highlighting how grouping genes by shared features promotes rapid multigenic variant assessment without sacrificing specificity and accuracy." This text makes no mention of the gene PPARGC1A, any evidence linking PPARGC1A to Mendelian disease, nor any mechanisms involving PPARGC1A. The content is explicitly about RAS/MAPK pathway genes and variant classification frameworks (RASopathies), which are distinct from PPARGC1A's known biology. Therefore, based on the provided content, there is no direct evidence (no observation, association, or variant-level data) nor mechanistic evidence (no pathway links, functional data, or plausible mechanism) supporting the claim that PPARGC1A is associated with a Mendelian disease. Limitations: only a single sentence (or brief excerpt) was supplied from the paper; if the full paper contains additional sections not provided here, it is possible (though unlikely given the excerpt's focused topic) that PPARGC1A is discussed elsewhere. However, within the supplied content, relevance to the claim is effectively zero.


[Read Paper](https://www.semanticscholar.org/paper/54e98adccb563b01deee283d5a738e8135c7a8d6)


### Effect of bezafibrate treatment on late-onset mitochondrial myopathy in mice.

**Why Not Relevant**: This paper provides mechanistic and therapeutic evidence that altering PGC1-alpha (the protein encoded by PPARGC1A) activity can modify the course of genetically caused mitochondrial myopathies in mouse models. However, it contains no data showing that germline variants in PPARGC1A cause a Mendelian disease in humans or segregate with disease in families. The experiments are primarily overexpression or pharmacologic modulation in mice and involve diseases caused by mutations in other genes (e.g., TWNK/Twinkle, COX10, SURF1). Therefore, while the work supports biological plausibility that PPARGC1A function impacts mitochondrial disease phenotypes (mechanistic evidence), it does not provide direct genetic evidence that PPARGC1A is itself associated with any Mendelian disorder. Given the lack of human genetic, segregation, or pathogenic-variant data for PPARGC1A in this paper, its relevance to the specific claim (PPARGC1A is likely associated with at least one Mendelian disease) is minimal.


[Read Paper](https://www.semanticscholar.org/paper/0a7a7ae5104e2ae0202ac788e0aff8e654c46da4)


### Muscle-generated BDNF (brain derived neurotrophic factor) maintains mitochondrial quality control in female mice

**Why Not Relevant**: The paper does not provide evidence that PPARGC1A is associated with any Mendelian disease. PPARGC1A appears only as an abbreviation/defined gene name within the abbreviation list; the experimental content and conclusions focus on BDNF-driven mitochondrial fission/mitophagy pathways in myotubes and muscle (mouse models and diet interventions). There are no human genetic data, no descriptions of pathogenic variants, no family segregation or inheritance information, and no statements that link PPARGC1A causally to a Mendelian phenotype. Experimental systems are cell culture and muscle-specific Bdnf knockout mice on high-fat diet, which are not appropriate to infer human Mendelian gene–disease relationships. For these reasons the paper is not relevant evidence for or against the claim that PPARGC1A is likely associated with at least one Mendelian disease.


[Read Paper](https://www.semanticscholar.org/paper/9e585440620d36206cd3035435b23a9a34b297df)


### Contractile activity attenuates autophagy suppression and reverses mitochondrial defects in skeletal muscle cells

**Why Not Relevant**: The paper provides mechanistic and expression-based evidence that PPARGC1A (PGC-1α) is a regulator of mitochondrial and lysosomal biogenesis and that its expression is modulated by chronic contractile activity; it also links pathway changes to autophagy/mitophagy markers and ROS-dependent mechanisms. However, it contains no human genetic data, no descriptions of pathogenic germline variants in PPARGC1A, no family segregation or case reports, and no statement that mutations in PPARGC1A cause a Mendelian disease. All evidence is experimental/physiological (muscle cell contractile activity, autophagy blockade, biochemical assays). Therefore the paper does not provide direct evidence that PPARGC1A is associated with any Mendelian disorder; at best it offers mechanistic plausibility that perturbation of the PPARGC1A-related pathway could contribute to disease phenotypes. Given the absence of genetic/clinical causation data, relevance to the specific claim (that PPARGC1A is likely associated with at least one Mendelian disease) is very low.


[Read Paper](https://www.semanticscholar.org/paper/a675f05aea7a6da3a936f0d89dc8f4339800aca5)


### Clingen Coagulation Factor Deficiency Variant Curation Expert Panel: Meeting the Need for Recommendations to Curate Variants in the Coagulation Factor Genes

**Why Not Relevant**: This paper is not relevant to the claim that PPARGC1A is likely associated with at least one Mendelian disease. The text is focused on ClinGen's Coagulation Factor Deficiency Variant Curation Expert Panel (CFD-VCEP) and on gene-specific variant curation for coagulation factor genes, specifically F8 and F9 (Hemophilia A and B). The paper provides no mention of PPARGC1A, no association studies, no clinical case reports, and no mechanistic data concerning PPARGC1A. Therefore it offers neither direct evidence (genetic association, case segregation, variant pathogenicity) nor mechanistic evidence (biological pathways, protein function linked to a Mendelian phenotype) relevant to the claim. Caveat: the absence of PPARGC1A in this paper does not disprove any possible PPARGC1A–disease associations reported elsewhere; it only indicates that this paper does not provide supportive or refuting information about PPARGC1A.


[Read Paper](https://www.semanticscholar.org/paper/9ea8d4ce19622ae1c6b0cdfd40848c93199f6280)


## Search Queries Used

- PPARGC1A mutation causes Mendelian disease case report family study PPARGC1A PGC1A PGC 1 alpha single gene disorder

- PPARGC1A pathogenic variant ClinVar OMIM gene curation review PPARGC1A disease association pathogenicity

- PPARGC1A PGC 1 alpha mitochondrial disease mitochondrial myopathy neurodegeneration muscle metabolic disorder mechanism

- PPARGC1A exome sequencing rare variant segregation linkage autosomal dominant autosomal recessive loss of function haploinsufficiency


## Usage Statistics

- Prompt Tokens: 0

- Completion Tokens: 0

- Total Tokens: 0

- Estimated Cost: $0.0369


## Bibliometric Configuration

- Use Bibliometrics: True

- Author Impact Weight: 0.4

- Citation Impact Weight: 0.4

- Venue Impact Weight: 0.2
