## Supplementary material for "MENDELSEEK: An algorithm that predicts Mendelian Genes and elucidates what makes them special": BHMT.txt

# Claim: The gene BHMT is likely associated with at least one Mendelian disease.

**Status**: processed

**Overall Rating**: 0

**Explanation**:

Supporting evidence.
The provided papers establish that BHMT (betaine-homocysteine S-methyltransferase) plays a physiologically important role in homocysteine remethylation and hepatic one-carbon/betaine metabolism. The first paper reports that in a mouse model of classical cystathionine β-synthase (CBS)–deficient homocystinuria (a well-characterized Mendelian metabolic disorder), BHMT mRNA, protein and enzyme activity were repressed by ~75%, hepatic betaine was depleted, and modulation of BHMT expression (e.g., by taurine or cysteine treatment) altered the therapeutic efficacy of betaine. The second paper describes methods for indirect measurement of BHMT activity and documents metabolite changes (e.g., N-dimethylglycine) that reflect BHMT pathway flux in human metabolic disease. Together these studies show BHMT is biochemically and functionally relevant to inherited disorders of sulfur/one-carbon metabolism and that its activity/expression changes in disease contexts.

Caveats or contradictory evidence.
None of the provided excerpts present evidence that germline variants in the BHMT gene cause a Mendelian disorder. The first study reports BHMT repression as a downstream effect of CBS deficiency (i.e., BHMT expression is altered secondary to another gene’s loss of function), not that BHMT itself is mutated. The second paper documents altered metabolites consistent with BHMT pathway perturbation in some metabolic conditions but does not link those perturbations to BHMT gene mutations. There are no genetic linkage, segregation, case reports, or loss-of-function mutation data presented that would indicate BHMT is causally implicated as a disease gene in a monogenic syndrome. Thus the evidence supports involvement of the BHMT enzyme in disease physiology but not that BHMT variants produce a Mendelian disease.

Analysis of potential underlying mechanisms and implications.
Mechanistically, BHMT catalyzes homocysteine methylation using betaine as methyl donor, a route complementary to methionine synthase (MTR)–mediated remethylation. If BHMT function is reduced (by repression, cofactor depletion, or enzymatic inhibition), one would expect perturbations in homocysteine/methionine/trimethylamine-N-oxide/DMG metabolism that could exacerbate homocystinuria-like phenotypes. This biochemical plausibility makes BHMT a reasonable candidate gene to consider in unexplained hyperhomocysteinemia or related metabolic phenotypes; however, biochemical plausibility alone does not prove that naturally occurring germline BHMT variants cause a Mendelian disorder. Secondary repression of BHMT in CBS deficiency illustrates that altered BHMT activity can be a downstream modifier of phenotype rather than the primary genetic cause. Without genetic evidence (e.g., pathogenic variants segregating with disease, functional studies of variants, or OMIM/ClinVar entries implicating BHMT), one cannot infer a Mendelian disease association from these metabolic/expressional data alone.

Which rating is most appropriate and why.
The preponderance of evidence in the provided excerpts supports BHMT's biochemical role and its altered expression/activity in at least one Mendelian metabolic disease (CBS-deficient homocystinuria), but there is no evidence that BHMT gene variants themselves cause a Mendelian disorder. Because the claim as worded—"The gene BHMT is likely associated with at least one Mendelian disease"—most naturally reads as an assertion that BHMT is genetically implicated in (i.e., causative of) at least one Mendelian disease, the provided papers do not supply that kind of evidence. Therefore the correct assessment based on the supplied literature is that there is no direct genetic evidence in these excerpts to support the claim. The appropriate rating is "No Evidence."


**Final Reasoning**:

I reviewed the excerpts for genetic evidence linking BHMT itself to a Mendelian disorder. The studies demonstrate that BHMT activity/expression is biologically important and is altered in a known Mendelian disease (CBS-deficient homocystinuria), and they provide biochemical/functional context. However, none of the excerpts show BHMT gene mutations causing a Mendelian disease (no case reports, segregation data, pathogenic variants, or curated disease-gene annotations). Thus, while BHMT is clearly disease-relevant biochemically, the provided evidence does not support the specific claim that BHMT is likely associated (in the genetic/causal sense) with at least one Mendelian disease. I therefore assign the rating "No Evidence."


## Relevant Papers


### Taurine alleviates repression of betaine‐homocysteine S‐methyltransferase and significantly improves the efficacy of long‐term betaine treatment in a mouse model of cystathionine β‐synthase–deficient homocystinuria

**Authors**: K. Maclean (H-index: 31), S. Stabler (H-index: 72)

**Relevance**: 0.25

**Bibliometric Impact**: 0.54096


**Excerpts**:

- Classical cystathionine β‐synthase‐deficient homocystinuria (HCU) is a life‐threatening inborn error of sulfur metabolism.

- Betaine treatment efficacy diminishes significantly over time due to impairment of betaine‐Hcy S‐methyltransferase (BHMT) function.

- Using a betaine‐responsive preclinical mouse model of HCU, we observed that this condition induces a 75% repression of BHMT mRNA, protein and enzyme activity, and significant depletion of hepatic betaine levels.

- BHMT repression was proportional to plasma Hcy levels but was not observed in mouse models of homocystinuria due to either methylenetetrahydrofolate reductase or Met synthase deficiency.

- Both Met supplementation and chemically induced glutathione depletion exacerbated hepatic BHMT repression in HCU mice but not wild‐type (WT) controls.

- Conversely, cysteine treatment normalized hepatic BHMT expression in HCU mice but had no effect in WT control animals.

- Taurine treatment induced BHMT expression in HCU mice by 5‐fold and restored maximal lowering of Hcy levels during long‐term betaine treatment with a concomitant normalization of inflammatory cytokine expression and a significantly improved coagulative phenotype.

- Collectively, our findings indicate that adjuvantial taurine treatment has the potential to significantly improve clinical outcomes in HCU.


**Explanations**:

- Context: This sentence defines the disease model studied (HCU), which is a Mendelian (inborn) metabolic disorder. Relevance: indirect/contextual. Strength/limitations: It establishes that the paper studies a genetically determined metabolic disease (HCU) but does not implicate BHMT as the causal gene for any Mendelian disease; rather it frames the biological context in which BHMT function is examined.

- Direct mechanistic assertion: This sentence links reduced BHMT function to loss of long‑term efficacy of betaine therapy. Relevance: mechanistic support that BHMT activity affects a Mendelian disease phenotype (treatment response) but not evidence that BHMT variation causes a Mendelian disease. Strength/limitations: Strong for implicating BHMT in disease pathophysiology (secondary impairment), but it does not demonstrate that germline BHMT mutations produce a Mendelian disorder in humans or mice.

- Mechanistic and quantitative finding: Reports large (75%) reductions in BHMT mRNA, protein, and enzyme activity in a mouse HCU model, plus depletion of hepatic betaine. Relevance: mechanistic evidence that BHMT function is substantially altered in the context of a Mendelian disease (CBS‑deficient HCU). Strength/limitations: Provides strong experimental evidence of BHMT dysregulation in disease state in mice, but this is secondary dysregulation in a model of CBS deficiency (not primary BHMT mutation). Also limited by being preclinical (mouse) and lacking human genetic data linking BHMT variants to Mendelian disease.

- Specificity evidence: States that BHMT repression correlated with plasma homocysteine and was absent in other mouse models of homocystinuria (MTHFR or Met synthase deficiency). Relevance: mechanistic — suggests BHMT repression is specifically associated with the CBS‑deficient HCU context and with hyperhomocysteinemia levels. Strength/limitations: Strengthens plausibility that BHMT activity is relevant to the disease mechanism in this specific genetic form of HCU, but still does not show that BHMT itself is a disease‑causing gene.

- Experimental manipulation result: Shows that Met supplementation and glutathione depletion worsened BHMT repression in HCU mice but not WT controls. Relevance: mechanistic — indicates environmental/metabolic modifiers affect BHMT expression in the disease model. Strength/limitations: Supports that BHMT can be modulated by metabolic state in the HCU context; does not provide evidence for BHMT as a Mendelian disease gene.

- Therapeutic mechanistic result: Cysteine normalized BHMT expression in HCU mice but had no effect in WT. Relevance: mechanistic — again supports that BHMT expression is responsive to thiol status in the disease model. Strength/limitations: Suggests potential therapeutic routes to restore BHMT function in disease, but not evidence of causative BHMT mutations.

- Therapeutic/phenotypic outcome: Demonstrates that taurine induced BHMT expression 5‑fold and restored maximal lowering of Hcy with improved inflammatory and coagulation phenotype. Relevance: mechanistic and phenotypic support that modulation of BHMT expression/function materially affects disease phenotype in this Mendelian disorder model. Strength/limitations: Strong functional evidence that BHMT activity matters to disease severity and treatment outcomes in the mouse model, but this remains secondary modulation rather than demonstration that BHMT germline variants are causal for a Mendelian disease.

- Interpretation: Authors propose taurine as an adjunct to improve outcomes in HCU. Relevance: contextual/concluding—supports the idea that BHMT activity is clinically meaningful in this Mendelian disease model. Strength/limitations: Reinforces biological importance of BHMT in disease context, but does not change the key limitation: no evidence presented that BHMT mutations themselves cause a Mendelian disease.


[Read Paper](https://www.semanticscholar.org/paper/09a238c710606dfc9ce1cc99a1366828fa8522c8)


### Indirect measurement of betaine--homocysteine methyltransferase activity using 1H nmr spectroscopy.

**Authors**: S. Burns (H-index: 11), R. Iles (H-index: 31)

**Relevance**: 0.25

**Bibliometric Impact**: 0.30806060606060603


**Excerpts**:

- The (re)methylation of homocysteine to methionine is an essential process for continuation of the cycle whereby Sadenosylmethionine acts as the major methyl donor for biosynthesis.

- There are two enzymes capable of mediating the homocysteine to m e t h i o n i n e c o n v e r s i o n : N 5 methyltetrahydrofolate-homocysteine methyltransferase with the methyl group provided by N5-methyltetrahydrofolate o r betaine-homocysteine methyl transferase (BHMT), the methyl group being donated by betaine leaving N-dimethylglycine (DMG) as product.

- We have investigated patients with the disorder multiple acyl CoA dehydrogenase deficiency ("glutaric aciduria type 11"; [2]) .

- The enzymes DMG dehydrogenase and sarcosine dehydrogenase are also ETF(-QO) dependent and whereas sarcosine excretion has been reported in such patients no investigation of DMG metabolism has been carried out.

- The spectrum from the patient shows several abnormal resonances. Signals from adipic, suberic , sebacic, glutaric and isovaleric acids are present. At 2.93 and 3.73 ppm are large signals from DMG and at 2.74 ppm a small signal from sarcosine.

- The excretion rate of DMG expressed as creatinine equivalents was 1.5. mol/mol.

- Excretion of DMG in healthy neonates is usually less than 0.3 mol/mol creatinine and sarcosine < 0.1 (Davies and Iles, unpublished).

- Perhaps surprisingly betaine excretion was within the normal range for a child of this age [41.


**Explanations**:

- Mechanistic background: This sentence establishes the biological importance of the homocysteine → methionine remethylation cycle and the role of S-adenosylmethionine as the principal methyl donor. Relevance to the claim: it provides context that perturbations in this cycle could plausibly produce disease phenotypes, but it does not mention BHMT specifically nor provide direct evidence linking BHMT to Mendelian disease. Limitation: purely background context; no genetic or clinical linkage to BHMT.

- Mechanistic evidence: This sentence explicitly identifies BHMT (betaine-homocysteine methyl transferase) as one of two enzymes that remethylate homocysteine, and states its biochemical product (DMG). Relevance to the claim: shows that BHMT participates in a medically important pathway and that its activity produces a measurable metabolite (DMG), supporting mechanistic plausibility that BHMT dysfunction could have metabolic consequences. Limitation: descriptive biochemical role only; no data on mutations, disease phenotypes, or human genetic linkage.

- Context / study population: This states that the investigators studied patients with multiple acyl-CoA dehydrogenase deficiency (glutaric aciduria type II). Relevance to the claim: indicates the clinical disorder under study is caused by defects in electron transfer flavoprotein or ETF-ubiquinone reductase, not BHMT. This weakens any inference that observed metabolic changes implicate BHMT as the primary causal gene for the patients' disease. Limitation: does not discuss BHMT variants or genetic testing.

- Mechanistic clarification: This indicates that DMG dehydrogenase and sarcosine dehydrogenase are ETF(-QO) dependent and that sarcosine excretion has been reported in such patients, while DMG metabolism had not been previously investigated. Relevance to the claim: provides a mechanistic explanation that accumulation of DMG (a BHMT product) can result from downstream oxidation defects (ETF-related), independent of BHMT function. This argues against interpreting DMG accumulation as evidence of BHMT gene pathogenicity. Limitation: doesn't present genetic data; inference depends on biochemical pathway knowledge.

- Direct metabolite observations: These sentences report the patient's NMR spectrum showing large signals from DMG and a small signal from sarcosine alongside other abnormal metabolites. Relevance to the claim: they document elevated DMG (product of BHMT) in a patient, which is consistent with altered flux through the BHMT→DMG→DMG-dehydrogenase pathway. However, this is not direct evidence that BHMT is mutated—elevation can be explained by downstream (ETF-dependent) enzyme failure as discussed elsewhere in the paper. Limitation: single-patient biochemical data; no genotype data; other metabolic defects could explain findings.

- Quantitative result: Reports the excretion rate of DMG in the patient as 1.5. mol/mol (creatinine equivalents). Relevance to the claim: shows markedly elevated DMG consistent with altered one-carbon/betaine metabolism, which could arise from altered BHMT flux but is equally or more plausibly due to impaired DMG catabolism. Limitation: format/units as reported are odd ("1.5. mol/mol"), sample size appears to be one neonate, and no genetic analysis of BHMT is presented.

- Comparative context: States normal neonatal DMG and sarcosine excretion ranges (DMG < 0.3 mol/mol creatinine; sarcosine < 0.1). Relevance to the claim: underscores that the patient's DMG excretion is substantially elevated relative to normal, confirming a metabolic perturbation. Limitation: still biochemical only; does not implicate BHMT gene mutations; reference cited as unpublished for normals.

- Additional metabolic observation: Notes that betaine excretion was within the normal range for age. Relevance to the claim: if betaine (the BHMT methyl donor) is normal, elevated DMG is less likely to be explained by increased substrate availability and more likely by downstream block in DMG oxidation — again arguing that BHMT gene dysfunction is not the necessary explanation. Limitation: single observation; does not exclude BHMT mutations but makes them less likely as the primary cause in this case.


[Read Paper](https://www.semanticscholar.org/paper/aa1738a25af7aeacd4c90284a21743fca5eae2d9)


## Other Reviewed Papers


### Exome sequencing of 20,979 individuals with epilepsy reveals shared and distinct ultra-rare genetic risk across disorder subtypes.

**Why Not Relevant**: The provided paper content is a high-level summary of a whole-exome sequencing study of epilepsy ("The largest whole-exome sequencing study of epilepsy to date, with more than 54,000 human exomes, is presented, comprising 20,979 deeply phenotyped patients from multiple genetic ancestry groups with diverse epilepsy subtypes and 33,444 controls, to investigate rare variants that confer disease risk."). Nowhere in the supplied text is the gene BHMT mentioned, nor are there any gene-specific results, variant lists, burden tests, mechanistic descriptions, or statements about genes being causal for Mendelian diseases. Therefore the content as provided contains no direct evidence (no variant associations, no statistical signals, no reported pathogenic variants) nor any mechanistic evidence (no functional data, pathways, or biological rationale) that would support or refute the claim that BHMT is likely associated with at least one Mendelian disease. 

Limitations that prevent relevance: (1) absence of any mention of BHMT in the provided excerpt; (2) lack of gene-level results, tables, or supplementary data in the provided content; (3) the summary nature of the text means even if BHMT were analyzed in the underlying study, no information about its association, effect sizes, significance, or mechanistic links is present here. It is possible that the full paper or its supplementary materials include BHMT-specific results, but those were not provided. To evaluate the claim, one would need gene-level association results, counts of rare predicted pathogenic variants in cases vs controls for BHMT, statistical significance (p-values, multiple-test correction), and any functional/mechanistic follow-up. Without such information in the supplied content, relevance to the BHMT–Mendelian disease claim is effectively zero.


[Read Paper](https://www.semanticscholar.org/paper/e67705489483e441dd2fd670d246565b27d3fd99)


### Whole exome sequencing identifies a novel mutation in ASPM and ultra-rare mutation in CDK5RAP2 causing Primary microcephaly in consanguineous Pakistani families

**Why Not Relevant**: The paper focuses exclusively on primary microcephaly (MCPH) and reports novel pathogenic variants in ASPM and CDK5RAP2 identified by whole-exome sequencing of two consanguineous Pakistani families. BHMT is not mentioned anywhere in the provided content (background, results, or conclusions), there are no mechanistic data linking BHMT to MCPH or to any Mendelian phenotype in this study, and no BHMT variants are reported. Because the paper neither investigates nor reports any findings about BHMT, it provides no direct or mechanistic evidence supporting or refuting the claim that BHMT is associated with a Mendelian disease. Note that while whole-exome sequencing was used (so, in principle, variants in BHMT could have been detected), the study's clinical focus (MCPH) and the small number of families mean the absence of BHMT findings here cannot be taken as meaningful evidence for or against BHMT causing Mendelian disease in general.


[Read Paper](https://www.semanticscholar.org/paper/82abfa052c72a2becdb46bed35c8854bbf631047)


### Biallelic variants identified in 36 Pakistani families and trios with autism spectrum disorder

**Why Not Relevant**: The provided paper content does not mention BHMT, nor does it present gene-level associations, variant lists, functional or clinical evidence, or statistical tests involving BHMT. The single sentence only states that the study reports genetic variability in ASD in Pakistan and a general enrichment for biallelic variants compared with outbreeding populations. While enrichment for biallelic variants is a general mechanistic factor that could increase the occurrence of recessive Mendelian diseases in a population, this is insufficient to implicate a specific gene. Therefore the paper content as provided offers no direct or specific mechanistic evidence linking BHMT to any Mendelian disease. To evaluate the claim about BHMT, one would need explicit mentions of BHMT (e.g., variants observed, segregation with disease, functional impact, statistical association), none of which are present in the excerpt. Because of that lack of gene-level information, the relevance to the BHMT claim is effectively negligible.


[Read Paper](https://www.semanticscholar.org/paper/17be98b475c43e8e296ed0d13d54250c91e9c42d)


### Deficiency of betaine-homocysteine methyltransferase activates glucose-6-phosphate dehydrogenase (G6PD) by decreasing arginine methylation of G6PD in hepatocellular carcinogenesis.

**Why Not Relevant**: The paper content provided addresses a mechanistic role for BHMT deficiency in hepatocarcinogenesis and potential therapeutic implications for HCC, but contains no information about germline variants, inheritance, familial segregation, penetrance, or clinical descriptions typical of Mendelian diseases. Mendelian disease associations require evidence such as identification of pathogenic germline variants, cosegregation in families, or reproducible genotype–phenotype links; none of these are present in the provided text. Therefore the content is minimally relevant to the specific claim that BHMT is likely associated with at least one Mendelian disease.


[Read Paper](https://www.semanticscholar.org/paper/8a1d460f1dee2e2c6475f2e7c1360a8e00aa83e9)


### Identification of novel variations in three cases with rare inherited neuromuscular disorder

**Why Not Relevant**: The provided paper content does not mention the gene BHMT anywhere, nor does it present data, analyses, or mechanistic discussion linking BHMT to any Mendelian disease. The study reports diagnostic variants in COL6A3, SPAST, and DMD in three pediatric IND cases and performs mechanistic work on a SPAST variant; none of these findings involve BHMT. Therefore this paper provides no direct evidence supporting or refuting the claim that BHMT is likely associated with at least one Mendelian disease. Note: absence of BHMT in this paper is not evidence that BHMT is not disease-associated; it only means this paper is not relevant to the claim. Additionally, the small sample size (three cases) and focus on IND limit any broader inferences about genes outside the study findings.


[Read Paper](https://www.semanticscholar.org/paper/e7130ad3c690f27033363e14a51ceb2eef859a74)


### Application of whole exome sequencing in the diagnosis of muscular disorders: a study of Taiwanese pediatric patients

**Why Not Relevant**: The paper does not mention the gene BHMT anywhere in the provided content, nor does it report findings (variants, pathogenicity assessments, mechanistic data, or disease associations) concerning BHMT. The study focuses on using whole exome sequencing to diagnose pediatric muscular dystrophies and congenital myopathies in a Taiwanese cohort and reports pathogenic variants across 28 genes relevant to those phenotypes; absence of any BHMT reference means there is no direct or mechanistic evidence in this paper supporting or refuting the claim that BHMT is likely associated with at least one Mendelian disease. Limitations relevant to this assessment: (1) absence of BHMT in this cohort or in the list of 28 genes does not prove BHMT has no Mendelian disease associations in other contexts or populations; (2) the study's clinical scope (muscle disorders) may not include phenotypes caused by BHMT variants even if such associations exist elsewhere; (3) WES and variant interpretation approaches may miss certain variant types (e.g., deep intronic, structural variants) or assign variants as uncertain that later turn out pathogenic. Given the complete lack of mention or data on BHMT in the paper content provided, the relevance to the BHMT-Mendelian-disease claim is essentially zero.


[Read Paper](https://www.semanticscholar.org/paper/455877818cc9f770df354051d974f7bb09689fcc)


## Search Queries Used

- BHMT betaine homocysteine methyltransferase pathogenic variant case report Mendelian disease biallelic mutation autosomal recessive

- BHMT BHMT2 whole exome sequencing rare disease candidate gene biallelic variants inherited disorder diagnostic sequencing study

- betaine homocysteine methyltransferase knockout mouse enzyme activity homocysteine methylation methionine cycle deficiency phenotype

- BHMT hyperhomocysteinemia homocystinuria inborn error of metabolism remethylation pathway MTHFR CBS differential diagnosis association study


## Usage Statistics

- Prompt Tokens: 0

- Completion Tokens: 0

- Total Tokens: 0

- Estimated Cost: $0.0395


## Bibliometric Configuration

- Use Bibliometrics: True

- Author Impact Weight: 0.4

- Citation Impact Weight: 0.4

- Venue Impact Weight: 0.2
