## Supplementary material for "MENDELSEEK: An algorithm that predicts Mendelian Genes and elucidates what makes them special": BRINP3.txt

# Claim: The gene BRINP3 is likely associated with at least one Mendelian disease.

**Status**: processed

**Overall Rating**: -1

**Explanation**:

No relevant papers were found that support or refute this claim.


## Relevant Papers


## Other Reviewed Papers


### Non-invasive prenatal sequencing for multiple Mendelian monogenic disorders using circulating cell-free fetal DNA

**Why Not Relevant**: The provided paper content is a single summary sentence that does not mention BRINP3, does not list the 30 genes targeted, and provides no gene-specific, variant-specific, clinical, segregation, or functional data. Because the claim is specifically about BRINP3 being likely associated with at least one Mendelian disease, the absence of any mention or data regarding BRINP3 means the paper content cannot support or refute the claim. The only possible (very weak) indirect relevance would be if BRINP3 is one of the unnamed 30 genes; however, that cannot be determined from the provided text. Additionally, even inclusion of BRINP3 as a target in a diagnostic assay would not alone establish that BRINP3 causes a Mendelian disease without accompanying clinical-genetic evidence. Given these substantial gaps, the paper content is not relevant to evaluating the claim about BRINP3.


[Read Paper](https://www.semanticscholar.org/paper/965b5d12657d5a39be70c31cf4cb3eb1617f5e96)


### Large-scale rare variant burden testing in Parkinson's disease.

**Why Not Relevant**: The paper focuses on rare-variant association analysis in Parkinson's disease and reports specific genes implicated in PD (GBA1, LRRK2) and several candidate PD risk genes (B3GNT3, AUNIP, ADH5, TUBA1B, OR1G1, CAPN10, TREML1). BRINP3 is not mentioned anywhere in the provided text. Therefore the paper provides no direct evidence that BRINP3 is associated with any Mendelian disease. The only indirect relevance is that the study's methods can detect monogenic (Mendelian) contributors to PD (as shown by LRRK2), which implies that if BRINP3 were a Mendelian PD gene present in these cohorts it might have been detected — but absence from the reported results is not strong evidence against BRINP3 causing a Mendelian disease in general, because (a) the analysis is limited to PD phenotypes and to the specific cohorts and allele frequency thresholds used; (b) the study may lack power for extremely rare or population-specific Mendelian variants; and (c) Mendelian associations for BRINP3 could involve phenotypes other than PD. Given no mention of BRINP3 and the PD-specific focus, the paper is essentially non-informative regarding the claim that BRINP3 is likely associated with at least one Mendelian disease, so overall relevance is very low.


[Read Paper](https://www.semanticscholar.org/paper/02f62832259be068f6aa4d92684745337ffa1c55)


### Rare genetic disorders: Beyond whole‐exome sequencing

**Why Not Relevant**: The provided paper content is a general review of next-generation sequencing technologies (WES/WGS) and their role in diagnosing rare genetic disorders. It contains no mention of the gene BRINP3, no patient cases, no variant data, and no statements linking BRINP3 to any Mendelian disease. The text discusses methodological issues (e.g., WES captures coding regions, WGS can detect structural and deep intronic variants) and the general diagnostic yield of sequencing strategies, but these are generic methodological points and not gene-specific evidence. Therefore, within the supplied content there is neither direct evidence (e.g., variant segregation, case reports, association statistics) nor mechanistic evidence (e.g., functional studies, biological pathways implicating BRINP3) bearing on the claim that BRINP3 is likely associated with at least one Mendelian disease. If you can provide sections of the paper that explicitly mention BRINP3 (e.g., results, tables, supplementary material) I can re-evaluate; based on the current excerpt, the claim cannot be supported or refuted.


[Read Paper](https://www.semanticscholar.org/paper/bd477d33a409ab5609c86cac0512c51ce998818a)


### Chromatin remodeling dysfunction extends the etiological spectrum of schizophrenia: a case report

**Why Not Relevant**: The provided paper content does not mention BRINP3, any specific gene, variants, or genetic data. It contains a single general statement: "This case underlines that schizophrenia may be, sometimes, underpinned by a Mendelian disease and addresses the question of systematic genetic investigations in the presence of warning signs such as a childhood onset of the schizophrenia or a resistant epilepsy." That sentence only asserts that some cases of schizophrenia can have a Mendelian basis and advocates genetic investigation when warning signs are present. It provides no direct evidence (no gene name, no variant, no segregation, no case series, no statistics) and no mechanistic evidence (no functional data, pathways, or molecular biology) linking BRINP3 to any Mendelian disease. Therefore the paper content as provided is irrelevant to the claim that BRINP3 is likely associated with at least one Mendelian disease. Limitations: the excerpt is very short and appears to be a general clinical/concluding remark (likely from a case report), so even if the full paper contains gene-level data elsewhere, that information is not present in the supplied text. Without explicit mention of BRINP3 or presentation of gene-specific genetic or mechanistic data, one cannot infer any support for the claim.


[Read Paper](https://www.semanticscholar.org/paper/29f6d9d6036caf286740be939452f1be2a4114b2)


### Genome-Wide Association Study-Guided Exome Rare Variant Burden Analysis Identifies IL1R1 and CD3E as Potential Autoimmunity Risk Genes for Celiac Disease

**Why Not Relevant**: The paper does not mention BRINP3 anywhere in the provided content and therefore provides no direct evidence that BRINP3 is associated with any Mendelian disease. The authors explicitly state they found "the absence of rare variants showing classical Mendelian inheritance in both families," and they prioritized other genes (IL1R1 and CD3E) based on their interactome and computational analyses. The study focuses on celiac disease in two Arab families, uses WES combined with GWAS loci and network analyses, and identifies candidate genes relevant to CeD pathogenesis—none of which include BRINP3. Limitations that bear on generalizability: the negative finding about Mendelian inheritance applies only to these two families and the study's detection filters; failure to detect BRINP3 here does not conclusively rule out BRINP3 involvement in Mendelian disease in other cohorts or via other study designs. However, based on the provided paper content, there is no direct or mechanistic evidence supporting the claim that BRINP3 is likely associated with at least one Mendelian disease.


[Read Paper](https://www.semanticscholar.org/paper/109c43874acfa1bf79ede793f0e10e4f8aa2ad9e)


### Case report: Novel compound heterozygous IL1RN mutations as the likely cause of a lethal form of deficiency of interleukin-1 receptor antagonist

**Why Not Relevant**: The provided paper does not mention BRINP3 and instead reports genetic and molecular evidence implicating IL1RN (interleukin-1 receptor antagonist) variants as the cause of a Mendelian autoinflammatory disease (DIRA) in the described family. All quoted content focuses on IL1RN: identification of two novel heterozygous IL1RN variants in the parents, decreased IL1RN mRNA production, truncated IL1RN transcripts, autosomal recessive segregation, and the conclusion that biallelic loss-of-function IL1RN variants cause DIRA. Because BRINP3 is not discussed, there is no direct evidence (variant-disease co-segregation, case reports, statistical association) or mechanistic evidence (effect of variants on gene/protein function, relevant pathways) in this paper relating BRINP3 to any Mendelian disease. Therefore this paper is not relevant to the claim that BRINP3 is likely associated with at least one Mendelian disease. If you want evidence for or against BRINP3 specifically, I can search literature/databases (PubMed, OMIM, ClinVar, gnomAD) and report any reported Mendelian associations or functional data for BRINP3.


[Read Paper](https://www.semanticscholar.org/paper/73a143f33c96c5392dc8b513a69039e0e391b92c)


### HOXA11 is another monogenic cause of congenital anomalies of the kidney and urinary tract—Reply

**Why Not Relevant**: The provided paper content discusses HXA11 and its insufficient evidence as a monogenic cause of CAKUT; it contains no mention of BRINP3, no patient or variant data, no functional assays, and no mechanistic information regarding BRINP3. Because the excerpt concerns a different gene and merely states a general standard for asserting monogenic causality (the need for additional patients and mutation characterization), it cannot be used to support or refute the claim that BRINP3 is likely associated with at least one Mendelian disease. To evaluate the claim about BRINP3 would require direct references to BRINP3 (clinical reports, segregation data, variant pathogenicity assessments, functional studies, or mechanistic pathways implicating BRINP3 in Mendelian phenotypes), none of which are present in the supplied content. Therefore the content is not relevant to the claim.


[Read Paper](https://www.semanticscholar.org/paper/1fdff7edb4655416163ed78c76a6471bc026c45d)


### An efficient design for whole genome trio sequencing identifies key variants in rare neurological disorder cases

**Why Not Relevant**: This paper does not provide evidence for or against the claim that BRINP3 is likely associated with at least one Mendelian disease. The study reports results from nine trios and explicitly names the genes found to be causal (CFAP52, DYNC1H1, FANCE, TCF4, and TOP3A). BRINP3 is not mentioned anywhere in the provided content. Because the dataset is small (nine trios), focused on neurological presentations in a specific underserved population, and used a sequencing design that could miss some variant classes or inherited variants (reduced parental coverage; CNV scanning applied inconsistently), the absence of BRINP3 findings in this paper is weak negative information: it neither supports nor meaningfully refutes a possible Mendelian association of BRINP3. In short, lack of mention = lack of relevant evidence, not disproval. Therefore overall relevance to the BRINP3 claim is negligible.


[Read Paper](https://www.semanticscholar.org/paper/3145118487852f8c0abaaebed17ba28b22502f54)


### Autosomal Dominant Intellectual Development Disorder-6 (MRD6) Without Seizures Linked to a De Novo Mutation in the grin2b Gene Revealed by Exome Sequencing: A Case Report of a Moroccan Child

**Why Not Relevant**: The paper exclusively discusses GRIN2B (presented as "grin2b") and a specific GRIN2B (grin2b) nonsense variant causing autosomal dominant intellectual development disorder-6 (MRD6). It provides detailed genetic, segregation, population-frequency, and ACMG-classification evidence for GRIN2B's role in a Mendelian disorder. However, the content contains no mention of BRINP3, no data on BRINP3 variants, no mechanistic pathways involving BRINP3, nor any comparative or interaction analyses that would connect BRINP3 to disease. Therefore the paper provides no direct or mechanistic evidence supporting the claim that BRINP3 is likely associated with at least one Mendelian disease.


[Read Paper](https://www.semanticscholar.org/paper/67d756e0cc302e5d46fdda9df34403f1f2de9a5a)


## Search Queries Used

- BRINP3 FAM5C pathogenic variant mutation Mendelian disease case report family segregation monogenic

- BRINP3 FAM5C ClinVar OMIM Decipher ClinGen GeneReviews reported pathogenic variants

- BRINP3 FAM5C whole exome sequencing whole genome sequencing trio study undiagnosed rare disease rare variant burden

- BRINP3 FAM5C expression function mouse knockout phenotype loss of function mechanism protein interaction pathway neural cardiac inflammation gnomAD pLI constraint


## Bibliometric Configuration

- Use Bibliometrics: True

- Author Impact Weight: 0.4

- Citation Impact Weight: 0.4

- Venue Impact Weight: 0.2
