## Supplementary material for "MENDELSEEK: An algorithm that predicts Mendelian Genes and elucidates what makes them special": BSN.txt

# Claim: The gene BSN is likely associated with at least one Mendelian disease.

**Status**: processed

**Overall Rating**: -1

**Explanation**:

No relevant papers were found that support or refute this claim.


## Relevant Papers


## Other Reviewed Papers


### Missense and truncating variants in CHD5 in a dominant neurodevelopmental disorder with intellectual disability, behavioral disturbances, and epilepsy

**Why Not Relevant**: The supplied paper content exclusively addresses CHD5 and its association with a variable neurodevelopmental syndrome. The claim concerns the gene BSN. The excerpt contains no mention of BSN, no data relating BSN to disease, and no mechanistic discussion that would plausibly connect findings about CHD5 to BSN. Without any mention or analysis of BSN, the paper content cannot provide direct or mechanistic evidence for or against the claim. Therefore relevance is effectively zero.


[Read Paper](https://www.semanticscholar.org/paper/09776c77585578bdb13befb2e2f389db53a18d89)


### De novo variants underlying monogenic syndromes with intellectual disability in a neurodevelopmental cohort from India

**Why Not Relevant**: The provided content contains no mention of the gene BSN, no gene-level variant data, no segregation or recurrence information, and no functional/mechanistic data related to BSN. Therefore the excerpt cannot support or refute the claim that BSN is likely associated with a Mendelian disease. To evaluate the claim one would need explicit reporting of BSN variants (e.g., de novo or segregating rare variants in affected individuals), phenotypic descriptions of carriers, statistical analyses or gene-level enrichment, and ideally functional evidence demonstrating an impact of BSN variation. None of these elements are present in the supplied text. If you can provide the full paper or the sections listing identified genes/variants, I can re-assess specifically for BSN.


[Read Paper](https://www.semanticscholar.org/paper/7bea82aa96284164e54212c2da9f191a36a23ada)


### Case report: A novel de novo deletion mutation of DYRK1A is associated with intellectual developmental disorder, autosomal dominant 7

**Why Not Relevant**: The supplied paper content focuses entirely on DYRK1A (Intellectual developmental disorder 7) and reports a de novo heterozygous deletion in DYRK1A as the pathogenic finding. Nowhere in the provided text is the gene BSN mentioned, nor are there mechanistic data, genetic findings, patient observations, segregation analyses, functional studies, or literature citations that implicate BSN in any Mendelian disease. Therefore this paper provides no direct evidence for or against the claim that BSN is likely associated with at least one Mendelian disease. Absence of mention in this single paper does not disprove an association elsewhere; it only means this paper is irrelevant to evaluating BSN. To evaluate the claim about BSN, one should instead consult sources that catalog gene-disease associations (OMIM, ClinVar, GenCC), search PubMed for BSN case reports or functional studies, and look for segregation, multiple unrelated affected individuals, or experimental knock-out/knock-in models implicating BSN.


[Read Paper](https://www.semanticscholar.org/paper/98eb2bb2850f75cb23ca3954609b1a07001ab461)


### Case report: Expanding the phenotype of FOXP1-related intellectual disability syndrome and hyperkinetic movement disorder in differential diagnosis with epileptic seizures

**Why Not Relevant**: This paper does not provide any direct or mechanistic evidence relating to the gene BSN. The study focuses on a single individual with a de novo splice site variant in FOXP1 (and an additional de novo CHD7 variant) and discusses clinical features and management of FOXP1-related intellectual disability. There is no mention of BSN, no BSN variants reported, no functional assays involving BSN, no segregation data, and no mechanistic discussion implicating BSN in Mendelian disease. Therefore the paper is not relevant to the claim that BSN is likely associated with at least one Mendelian disease. If you need evidence for or against BSN's involvement in Mendelian disease, relevant data would include reported pathogenic or likely pathogenic BSN variants in affected individuals (with segregation or de novo occurrence), functional studies showing a deleterious effect of BSN variants, population constraint metrics, animal-model phenotypes, and entries in databases such as OMIM, ClinVar, DECIPHER, or peer-reviewed case series; none of these are present in the provided paper content.


[Read Paper](https://www.semanticscholar.org/paper/3d7a33a707a435016dbc5fac3ff92c45624031a4)


### Burden re-analysis of neurodevelopmental disorder cohorts for prioritization of candidate genes.

**Why Not Relevant**: The provided content is a single, general methodological/summary sentence that does not mention BSN or any specific gene, does not present results, effect sizes, statistical significance, variant-level or segregation data, or functional/mechanistic findings. Because the excerpt contains only a high-level description of approaches used to discover novel NDD genes and no gene-specific evidence, it is insufficient to support or refute the claim that BSN is likely associated with at least one Mendelian disease. To evaluate the claim, one would need verbatim sentences or paragraphs from the paper that report BSN-specific findings (e.g., enrichment statistics, de novo or segregating variants in BSN, phenotype co-segregation, functional assays, or clinical case descriptions) along with methodological details (sample size, filtering thresholds, p-values, confidence intervals) and any limitations discussed by the authors.


[Read Paper](https://www.semanticscholar.org/paper/0a7e56a69b7001fbe8ee5a57ca3d63c5bc7951cc)


### Expanding the Spectrum of Autosomal Dominant ATP6V1A-Related Disease: Case Report and Literature Review

**Why Not Relevant**: This paper describes a clinical and genetic case concerning the ATP6V1A gene (a de novo heterozygous pathogenic variant c.82G>A, p.Val28Met) and the clinical spectrum of ATP6V1A-related neurodevelopmental disorders. There is no mention of the BSN gene, no data about BSN variants, and no mechanistic discussion linking ATP6V1A findings to BSN. Therefore the paper provides no direct evidence that BSN is associated with any Mendelian disease, nor does it provide mechanistic evidence that would support or refute such an association. The absence of BSN in the text means this paper cannot be used to evaluate the claim about BSN. If you want evidence for or against BSN being implicated in Mendelian disease, I can perform a literature search or analyze papers that explicitly study BSN (Bassoon) variants, their segregation in families, functional studies, or case reports.


[Read Paper](https://www.semanticscholar.org/paper/5a349716f192d755ac18fcd68585ae76bdea9c72)


### Autosomal dominant Ullrich congenital muscular dystrophy due to a de novo mutation in COL6A3 gene. A case report

**Why Not Relevant**: The provided paper content exclusively discusses collagen VI genes and a pathogenic variant in COL6A3 causing Ullrich congenital muscular dystrophy (UCMD). There is no mention of the BSN gene, no genetic data, mechanistic discussion, or clinical association involving BSN, and therefore the paper provides no direct or mechanistic evidence supporting the claim that BSN is associated with a Mendelian disease. The absence of BSN-related data in this paper does not disprove the claim overall — it simply means this paper is not relevant to evaluating BSN's disease association.


[Read Paper](https://www.semanticscholar.org/paper/2db22d53df5b68aee90bf24ad1489b2fa725984c)


### Case Report: Concurrent de novo pathogenic variants in the LMNA gene as a cause of sporadic partial lipodystrophy

**Why Not Relevant**: This paper does not mention the gene BSN and therefore provides no direct or mechanistic evidence bearing on the claim that BSN is likely associated with at least one Mendelian disease. The study focuses on germline de novo variants in LMNA in a sporadic case of partial lipodystrophy, for example: "Two heterozygous missense de novo pathogenic variants in cis within the LMNA gene were found in the proband: p.Y481H and p.K486N (NP_733821.1)." and "In a case of sporadic partial lipodystrophy, we describe two concurrent de novo pathogenic variants within the same gene (LMNA) as a novel pathogenic mechanism." Because BSN is not discussed, there is neither direct evidence (e.g., reported pathogenic BSN variants segregating with disease) nor mechanistic evidence (e.g., functional studies, protein modeling, or pathway links implicating BSN) in this paper. Limitation: the authors performed exome-wide analyses under multiple mendelian inheritance models but report only the LMNA findings; absence of any mention of BSN in the text means one cannot infer a negative result for BSN from this report (i.e., lack of reported BSN variants here is not strong evidence that BSN is not associated with Mendelian disease), but equally there is no supportive evidence to link BSN to a Mendelian disorder in this manuscript.


[Read Paper](https://www.semanticscholar.org/paper/0ed819a1194503a68324b1b37c5d69996acc257d)


### Missense and truncating variants in CHD5 in a dominant neurodevelopmental disorder with intellectual disability, behavioral disturbances, and epilepsy

**Why Not Relevant**: The provided paper content discusses heterozygous CHD5 variants and a neurodevelopmental syndrome; it contains no mention of the gene BSN, no clinical or genetic data linking BSN to disease, and no mechanistic information that would plausibly implicate BSN. Because the excerpt pertains to a different gene (CHD5) and no link between CHD5 and BSN is provided, the content cannot be used as direct or mechanistic evidence for the claim that BSN is likely associated with at least one Mendelian disease. To evaluate the claim about BSN would require papers reporting BSN variants in affected individuals with segregation/functional data, case series, or mechanistic studies implicating BSN in disease—none of which are present in the supplied text.


[Read Paper](https://www.semanticscholar.org/paper/59f28a970d4de4d91a257f01572ae31b32f28c4d)


## Search Queries Used

- BSN mutation pathogenic variant Mendelian disease case report de novo autosomal recessive autosomal dominant

- BSN loss of function intolerance pLI gnomAD ClinVar OMIM haploinsufficiency gene disease association

- Bassoon presynaptic cytomatrix BSN synaptic transmission epilepsy neurodegeneration mouse knockout seizure neuropathology

- exome sequencing BSN de novo variant neurodevelopmental disorder epilepsy intellectual disability cohort rare variant burden


## Bibliometric Configuration

- Use Bibliometrics: True

- Author Impact Weight: 0.4

- Citation Impact Weight: 0.4

- Venue Impact Weight: 0.2
