## Supplementary material for "MENDELSEEK: An algorithm that predicts Mendelian Genes and elucidates what makes them special": BTRC.txt

# Claim: The gene BTRC is likely associated with at least one Mendelian disease.

**Status**: processed

**Overall Rating**: -1

**Explanation**:

No relevant papers were found that support or refute this claim.


## Relevant Papers


## Other Reviewed Papers


### De novo mutations in epileptic encephalopathies

**Why Not Relevant**: The provided paper content does not mention the gene BTRC anywhere in the text, the lists of genes with de novo mutations, or the genes highlighted as intolerant or implicated in epileptic encephalopathies. The study reports exome sequencing of 264 trios and confirms 329 de novo mutations, and it names specific genes (for example: "Other genes with de novo mutations in this cohort include CACNA1A, CHD2, FLNA, GABRA1, GRIN1, GRIN2B, HNRNPU, IQSEC2, MTOR and NEDD4L."), but BTRC is not among them. Therefore this paper provides no direct evidence (no observed mutations, no statistical association) that BTRC is associated with any Mendelian disease. 

Mechanistic evidence for BTRC (e.g., functional studies, pathway links, intolerance scores for BTRC, or observed de novo or inherited pathogenic variants) is not provided in the excerpt; the paper only provides a general framework that could be applied to candidate genes but does not apply it to BTRC. Limitations relevant to using this paper to evaluate BTRC: absence of BTRC in this study is not evidence that BTRC is not disease-associated (negative data), because (1) the study size (264 trios) has limited power to detect associations for many genes, (2) the study focuses on de novo mutations in epileptic encephalopathies and would not detect inherited Mendelian associations or associations with other phenotypes, and (3) exome sequencing has incomplete sensitivity for some variant types (e.g., deep intronic variants, structural variants). In summary, relevance is effectively zero because the paper contains neither direct nor mechanistic evidence about BTRC; any inference about BTRC would require additional data not present in this paper.


[Read Paper](https://www.semanticscholar.org/paper/2b6ce4a1dfc9c8722254184f4ddbde209b364626)


### De novo mutation screening in childhood-onset cerebellar atrophy identifies gain-of-function mutations in the CACNA1G calcium channel gene

**Why Not Relevant**: The provided paper content focuses on early-onset cerebellar atrophy and specifically investigates CACNA1G, reporting de novo CACNA1G mutations and detailed functional (gain-of-function) evidence for Cav3.1 channel defects. There is no mention of BTRC, no data on variants in BTRC, and no mechanistic discussion involving BTRC protein function. Therefore this paper provides neither direct genetic association evidence nor mechanistic evidence linking BTRC to any Mendelian disease. Absence of BTRC in the provided content does not disprove a possible association elsewhere, but based on the text given, no support or refutation regarding BTRC can be drawn. If you want evidence about BTRC specifically, I recommend querying gene-disease resources (OMIM, ClinVar, HGMD) or searching the literature for studies that explicitly mention BTRC.


[Read Paper](https://www.semanticscholar.org/paper/0b264bac49270d7d896e6d7fd3a1b3db7daec8f2)


### Expanding the phenotype of a recurrent de novo variant in PACS1 causing intellectual disability

**Why Not Relevant**: The provided paper content exclusively discusses PACS1 (a trans-Golgi membrane traffic regulator) and a recurrent de novo PACS1 missense variant (p.Arg203Trp) causing the intellectual disability syndrome MRD17. There is no mention of the gene BTRC anywhere in the excerpt, nor is there any comparative, pathway, or mechanistic discussion linking PACS1 (or the reported findings) to BTRC. Consequently, the paper provides no direct genetic evidence (e.g., variants in BTRC in affected individuals), no mechanistic evidence (e.g., functional assays implicating BTRC, shared pathways connecting PACS1 and BTRC), and no epidemiological/clinical evidence relevant to the claim that 'BTRC is likely associated with at least one Mendelian disease.' Without any mention or analysis of BTRC, one cannot infer association from these PACS1-specific data. Therefore relevance is effectively zero. If you want evidence about BTRC, please provide papers mentioning BTRC or allow me to search the literature for BTRC-specific genetic or functional studies.


[Read Paper](https://www.semanticscholar.org/paper/e460d89d850693b4fb1e3df12702be0dfb8d7f5b)


### De novo variation in ARID1B gene causes Coffin-Siris syndrome 1 in a Chinese family with excessive early-onset high myopia

**Why Not Relevant**: The provided paper content exclusively discusses ARID1B and its role in Coffin–Siris syndrome (CSS) and early-onset high myopia (eoHM). There is no mention of the gene BTRC, no experimental results, case reports, genetic linkage/association analyses, variant interpretations, or mechanistic descriptions involving BTRC. Because the claim specifically concerns BTRC being likely associated with at least one Mendelian disease, and this paper provides no data or mechanistic rationale involving BTRC, the paper is not relevant to the claim. Note that lack of relevance here does not imply that BTRC is not disease-associated in other studies; it only reflects that this particular paper contains no evidence bearing on BTRC.


[Read Paper](https://www.semanticscholar.org/paper/a0fe18dd082eeec9079cc284babcd59eb224375f)


### Genetic characterization of Schuurs-Hoeijmakers syndrome in a moroccan individual with heterozygote PACS1 mutation

**Why Not Relevant**: The supplied paper content contains a single sentence: "Functional studies indicate the crucial role of PACS1 in craniofacial development and neurodevelopmental processes, with potential implications for autism spectrum disorders (ASD)." This sentence discusses PACS1 only and makes no mention of BTRC, Mendelian disease associations, pathogenic variants, clinical phenotypes, family segregation, or functional data relating to BTRC. Therefore the paper content provides neither direct evidence (e.g., reported pathogenic BTRC variants or clinical diagnoses linked to BTRC) nor mechanistic evidence (e.g., molecular pathways, protein interactions, loss-/gain-of-function experiments involving BTRC) relevant to the claim that "BTRC is likely associated with at least one Mendelian disease." Because the content is about a different gene (PACS1) and lacks any information about BTRC, the relevance to the claim is effectively zero. To evaluate the claim about BTRC one would need sources that explicitly discuss BTRC (for example OMIM/ClinVar entries, case reports showing co-segregation of BTRC variants with Mendelian phenotypes, or functional studies demonstrating that BTRC variants cause a disease mechanism).


[Read Paper](https://www.semanticscholar.org/paper/23aec955d8d80465837e00480f370abc21574484)


## Search Queries Used

- BTRC beta TrCP1 beta transducin repeat containing E3 ubiquitin ligase pathogenic variant Mendelian disease OMIM ClinVar case report

- BTRC de novo mutation exome sequencing intellectual disability developmental disorders early onset epilepsy congenital anomaly cohort study rare variant

- BTRC loss of function beta catenin Wnt signaling developmental defect mouse knockout phenotype human disease mechanism congenital malformation craniofacial limb

- BTRC gene disease association ClinGen gnomAD pLI loss of function intolerance haploinsufficiency gene curation Mendelian gene list review


## Bibliometric Configuration

- Use Bibliometrics: True

- Author Impact Weight: 0.4

- Citation Impact Weight: 0.4

- Venue Impact Weight: 0.2
