## Supplementary material for "MENDELSEEK: An algorithm that predicts Mendelian Genes and elucidates what makes them special": CACNG3.txt

# Claim: The gene CACNG3 is likely associated with at least one Mendelian disease.

**Status**: processed

**Overall Rating**: 2

**Explanation**:

Supporting evidence.
Several genetic studies report linkage and association signals implicating CACNG3 in childhood absence epilepsy (CAE). Linkage analyses in cohorts of CAE families yielded positive HLOD scores for markers encompassing CACNG3 (e.g., HLOD=3.54, α=0.62) and transmission disequilibrium was observed for SNPs in a ~35 kb high-LD region around the 5'UTR and exon 1. Re-sequencing identified many noncoding variants and four variants predicted computationally to affect exonic splicing. Functional inference from the gene family is also suggestive: CACNG3 belongs to the CACNG (stargazin-like) family, and loss-of-function mutations in the related gene Cacng2 produce recessive epilepsy/ataxia phenotypes in mice, showing that gamma subunits can produce clear monogenic phenotypes in vivo.

Caveats or contradictory evidence.
None of the cited human studies identified clear coding (protein-altering) mutations in CACNG3 that segregate with disease in a Mendelian fashion; re-sequencing of coding exons in affected individuals failed to find putative causal variants. The positive genetic signals are consistent with CACNG3 being a susceptibility locus in a subset of patients rather than a locus for a classical Mendelian disorder — CAE is an idiopathic generalized epilepsy with complex inheritance and locus heterogeneity. The reported variants are predominantly noncoding, and the splice-impact predictions are in silico only; there is no published evidence here of a variant with demonstrated loss/gain-of-function, segregation consistent with dominant or recessive inheritance, de novo occurrence, or functional validation showing pathogenicity. Evidence from Cacng2 is from a different paralog and from mouse models; paralogous gene disease in animals increases plausibility but is not direct proof for CACNG3 causing a human Mendelian disease.

Analysis of mechanisms and deeper implications.
Biologically, CACNG3 encodes an AMPA receptor trafficking regulator / gamma subunit that can influence synaptic transmission and possibly voltage-dependent calcium channel function. Disruptive mutations (loss-of-function or dominant-negative) could plausibly produce a severe neuronal phenotype causing epilepsy. The presence of noncoding/splice-region variation raises the possibility of regulatory or splicing defects causing disease, but establishing a Mendelian relationship would require (a) identification of rare, highly penetrant variants, (b) consistent segregation in families or recurrent de novo events, and (c) functional demonstration of pathogenic mechanism. The observed linkage and TDT findings suggest CACNG3 contributes to risk in some families, but the high locus heterogeneity and absence of coding pathogenic variants point to a polygenic / susceptibility role rather than monogenic causation based on current data.

Which rating is most appropriate.
Taken together, the preponderance of evidence indicates CACNG3 is implicated as a susceptibility locus for CAE in subsets of patients, but there is insufficient direct evidence to support that CACNG3 is associated with a Mendelian (monogenic) disease in humans. The lack of identified segregating coding mutations, absence of functional validation of predicted splice variants, and the complex inheritance pattern of CAE argue against concluding a likely Mendelian association. Paralog-based and animal-model evidence raise plausibility but do not offset the absence of human monogenic evidence. Therefore the most appropriate rating is "Likely False": it is unlikely, based on the provided evidence, that CACNG3 is associated with at least one Mendelian disease in humans (though it remains a plausible susceptibility gene and warrants further rare-variant/functional studies).


**Final Reasoning**:

I reviewed the linkage and association data showing CACNG3 as a susceptibility locus for childhood absence epilepsy, the absence of identified coding causal variants in resequencing efforts, the presence of only predicted (in silico) splice-affecting variants, and the fact that the genetic signals are consistent with locus heterogeneity and complex inheritance. Although mechanistic plausibility exists (gene function in synaptic regulation; disease in a paralogous gene in mice), there is no direct human evidence of rare, highly penetrant variants with Mendelian segregation or functional validation of pathogenicity in CACNG3. Given this balance — positive evidence for susceptibility but no evidence of monogenic causation — the claim that CACNG3 is likely associated with at least one Mendelian disease is not supported by the provided literature. I therefore assign the rating "Likely False."


## Relevant Papers


### Linkage and association analysis of CACNG3 in childhood absence epilepsy

**Authors**: Kate V. Everett (H-index: 14), M. Gardiner (H-index: 16)

**Relevance**: 0.35

**Bibliometric Impact**: 0.26103529411764703


**Excerpts**:

- Evidence supports CACNG3 as a susceptibility locus in a subset of CAE patients,

- although four variants are predicted to affect exonic splicing.


**Explanations**:

- This sentence provides direct evidence that CACNG3 is associated with a human disease phenotype (childhood absence epilepsy, CAE) by identifying it as a "susceptibility locus" in a subset of patients. Strength: it is a direct statement of an association with a clinical phenotype. Limitations: the phrasing "susceptibility locus" and "subset" imply an association or risk factor rather than a proven monogenic (Mendelian) cause. The sentence gives no information on segregation with disease in families, effect sizes, allele frequencies, statistical significance, inheritance pattern (dominant, recessive, de novo), or functional validation — all of which are necessary to support a claim of a Mendelian disease association. Therefore this is relevant evidence for association but weak/insufficient to demonstrate a Mendelian causal relationship.

- This sentence offers mechanistic plausibility by reporting that four variants are predicted to affect exonic splicing, which could plausibly alter protein function or expression and thereby contribute to disease. Strength: it points to potentially functional variants that could underlie pathogenicity. Limitations: the statement refers to in silico predictions only ("predicted"), with no provided functional assays, no demonstration that these variants are rare/pathogenic, and no family segregation or phenotype correlation presented here. Predicted splicing effects increase biological plausibility but do not by themselves establish pathogenicity or Mendelian inheritance.


[Read Paper](https://www.semanticscholar.org/paper/301b8571e7094fc3cb73d9546b2263b276337de4)


### The alpha-amino-3-hydroxyl-5-methyl-4-isoxazolepropionate receptor trafficking regulator "stargazin" is related to the claudin family of proteins by Its ability to mediate cell-cell adhesion.

**Authors**: M. Price (H-index: 16), D. Burgess (H-index: 22)

**Relevance**: 0.25

**Bibliometric Impact**: 0.29246315789473687


**Excerpts**:

- Mutations in the Cacng2 gene encoding the neuronal transmembrane protein stargazin result in recessively inherited epilepsy and ataxia in "stargazer" mice.

- Functional studies suggest a dual role for stargazin, both as a modulatory gamma subunit for voltage-dependent calcium channels and as a regulator of post-synaptic membrane targeting for alpha-amino-3-hydroxyl-5-methyl-4-isoxazolepropionate (AMPA)-type glutamate receptors.

- Co-immunoprecipitation experiments demonstrate that stargazin can bind proteins of either complex in vivo, but it remains unclear whether it can associate with both complexes simultaneously.

- Cacng2 is one of eight closely related genes (Cacng1-8) encoding proteins with four transmembrane segments, cytoplasmic termini, and molecular masses between 25 and 44 kDa.

- This group of Cacng genes constitutes only one branch of a larger monophyletic assembly dominated by over 20 genes encoding proteins known as claudins.

- Because stargazin is structurally similar to claudins, we hypothesized that it might also have retained claudin-like functions inherited from a common ancestor.

- Here, we report that expression of stargazin in mouse L-fibroblasts results in cell aggregation comparable with that produced by claudins, and present evidence that the interaction is heterotypic and calcium dependent.

- The data suggest that the cell adhesion function of stargazin preceded its current role in neurons as a regulator of either voltage-dependent calcium channels or AMPA receptors.

- We speculate these complexes may have co-opted the established presence of stargazin at sites of close cell-cell contact to facilitate their own evolving intercellular signaling functions.


**Explanations**:

- Relation to claim: This sentence documents a Mendelian (recessive) disease phenotype caused by mutations in a CACNG family member (Cacng2) in mice. Type of evidence: direct (for Cacng2 in mice). Strengths/limitations: Strong direct genetic-disease association, but it is in mouse Cacng2, not human CACNG3. Thus it supports the plausibility that gamma subunit family members can cause Mendelian disease, but does not provide direct evidence that CACNG3 is disease-causing in humans. Also, no human clinical data or CACNG3-specific mutations are reported here.

- Relation to claim: Describes molecular functions of stargazin relevant to neuronal physiology (calcium channel modulation and AMPA receptor trafficking). Type of evidence: mechanistic. Strengths/limitations: Mechanistic roles indicate how perturbation of a CACNG-family protein could lead to neurological disease, strengthening biological plausibility for related paralogs (including CACNG3). Limitation: These functions are described for Cacng2/stargazin; the excerpt does not show that CACNG3 shares the same activities in vivo or in human neurons.

- Relation to claim: Shows biochemical interactions of stargazin with proteins of the two complexes (VDCCs and AMPA receptor complexes). Type of evidence: mechanistic. Strengths/limitations: Supports the notion that perturbing such interactions could cause disease phenotypes; however, it addresses Cacng2/stargazin specifically and notes uncertainty about simultaneous association with both complexes. It does not provide data about CACNG3 binding partners or disease-causing variants.

- Relation to claim: States that Cacng2 is part of an eight-gene family (Cacng1-8), which implicitly includes Cacng3 as a closely related paralog. Type of evidence: contextual/mechanistic. Strengths/limitations: This supports the inference that disease mechanisms observed for Cacng2 might be relevant across the family, giving indirect support for CACNG3 candidacy. Limitation: Paralogy alone does not establish that CACNG3 is disease-causing; functional divergence among paralogs is common.

- Relation to claim: Places the Cacng gene subgroup into a larger evolutionary context with claudins. Type of evidence: mechanistic/evolutionary context. Strengths/limitations: Indicates conserved structural features and possible conserved functions (e.g., cell adhesion), which could imply shared vulnerability to pathogenic variants. Limitation: Evolutionary relationship does not equate to disease association for any specific member like CACNG3.

- Relation to claim: Authors hypothesize functional conservation with claudins due to structural similarity. Type of evidence: mechanistic/hypothesis. Strengths/limitations: Offers a mechanistic rationale (retained claudin-like functions) for how CACNG-family proteins might be important for cell physiology and thus disease-sensitive. Limitation: This is a stated hypothesis; it is not direct evidence of disease causation for any CACNG gene, and the text refers to stargazin specifically.

- Relation to claim: Demonstrates an experimentally observed cellular function (cell aggregation) for stargazin and provides mechanistic detail (heterotypic, calcium-dependent interaction). Type of evidence: mechanistic/functional. Strengths/limitations: Functional assays show stargazin influences cell adhesion, providing a mechanistic pathway by which mutations could perturb tissue/neuronal architecture and cause disease. Limitation: Experiments were performed with stargazin (Cacng2) in mouse fibroblasts; transferability to CACNG3, to human cells, and to in vivo pathogenesis is unproven in this paper.

- Relation to claim: Interprets the functional data in an evolutionary and mechanistic framework, proposing that cell adhesion preceded neuronal roles. Type of evidence: mechanistic/speculative. Strengths/limitations: Helps explain how disruption of these proteins could have pleiotropic effects relevant to Mendelian phenotypes. Limitation: Speculative and not direct evidence for CACNG3 involvement in human Mendelian disease.

- Relation to claim: Authors speculate that stargazin's localization to cell contacts was co-opted by other complexes; this is mechanistic/speculative context. Type of evidence: mechanistic/speculation. Strengths/limitations: Gives a plausible mechanism for how disruption of gamma-subunit/claudin-like proteins might impact intercellular signaling and lead to disease phenotypes. Limitation: Speculation about stargazin (Cacng2) only; no empirical data on CACNG3 or human Mendelian disease.


[Read Paper](https://www.semanticscholar.org/paper/46269e4dae96698d39affd5bbd7b02a220e3d003)


### Linkage analysis between childhood absence epilepsy and genes encoding GABAA and GABAB receptors, voltage-dependent calcium channels, and the ECA1 region on chromosome 8q.

**Authors**: R. Robinson (H-index: 15), R. Gardiner (H-index: 35)

**Relevance**: 0.7

**Bibliometric Impact**: 0.30025454545454544


**Excerpts**:

- Childhood absence epilepsy (CAE) is an idiopathic generalised epilepsy (IGE) characterised by onset of typical absence seizures in otherwise normal children of school age.

- Available evidence suggests that mutations in genes encoding GABA receptors or brain expressed voltage-dependent calcium channels (VDCCs) may underlie CAE.

- Seventeen VDCC subunit genes, ten GABA(A)R subunit genes, two GABA(B) receptor genes and the ECA1 locus on 8q24 were investigated using 35 microsatellite loci.

- Assuming locus homogeneity, all loci gave statistically significant negative LOD scores, excluding these genes as major loci in the majority of these families.

- Positive HLOD scores assuming locus heterogeneity were observed for CACNG3 on chromosome 16p12-p13.1 and the GABRA5, GABRB3, GABRG3 cluster on chromosome 15q11-q13.

- Association studies are required to determine whether these loci are the site of susceptibility alleles in a subset of patients with CAE.


**Explanations**:

- Context: This sentence defines the phenotype under study (CAE) and shows the paper is investigating a familial epilepsy phenotype. Relevance to claim: indirect/contextual. Strength/limitations: establishes the clinical phenotype but does not provide evidence about CACNG3 or Mendelian inheritance by itself.

- Mechanistic plausibility: This sentence states the biological rationale linking VDCCs (which include CACNG3-encoded subunits) and GABA receptor genes to CAE. Relevance to claim: mechanistic. Strength/limitations: it supports biological plausibility that a VDCC subunit gene could contribute to epilepsy, but it is a general hypothesis statement, not experimental evidence that CACNG3 causes a Mendelian disease.

- Methods/scope: This sentence describes the set of genes and markers tested (including VDCC subunit genes) and the linkage approach. Relevance to claim: methodological/direct. Strength/limitations: shows CACNG3 was within the tested gene set (because VDCC subunits were tested), so the study could provide direct genetic linkage evidence. It does not by itself provide results or prove association.

- Negative-result context: This sentence reports that under an assumption of locus homogeneity, all loci were excluded as major loci in the majority of families. Relevance to claim: direct evidence that weakens a simple, single-locus Mendelian interpretation. Strength/limitations: strong for excluding these genes as major, homogeneous causes across the cohort, but does not rule out locus heterogeneity or a role in a subset of families.

- Primary linkage finding: This sentence reports positive HLOD scores for CACNG3 (chr16p12-p13.1) when heterogeneity is allowed, and similarly for a GABA receptor cluster. Relevance to claim: direct evidence supporting that CACNG3 cosegregates with CAE in a subset of families (i.e., compatible with being a Mendelian locus in some pedigrees). Strength/limitations: this is suggestive linkage evidence (positive HLOD under heterogeneity), which increases plausibility that CACNG3 is associated with CAE in at least some families. Limitations include that linkage (especially heterogeneous HLOD) is not proof of causative mutation; no mutation screening, functional validation, or association/replication data are reported here. Also CAE can be genetically complex, so linkage in a subset does not by itself establish CACNG3 as a Mendelian disease gene.

- Caveat/future work: This sentence explicitly states that association studies are required to determine whether these loci harbour susceptibility alleles in a subset of patients. Relevance to claim: limitation/next steps. Strength/limitations: the authors acknowledge that their linkage results are not definitive proof of disease-causing alleles and that further genetic (association/mutation) and functional studies are needed to confirm CACNG3 as a disease gene.


[Read Paper](https://www.semanticscholar.org/paper/b15ab7f59514c387709d258fee16cc81eaa794eb)


### Linkage and association analysis of CACNG 3 in childhood absence epilepsy

**Authors**: Kate V. Everett (H-index: 14), M. Gardiner (H-index: 16)

**Relevance**: 0.4

**Bibliometric Impact**: 0.120225


**Excerpts**:

- Childhood absence epilepsy (CAE) is an idiopathic generalised epilepsy characterised by absence seizures manifested by transitory loss of awareness with 2.5-4Hz spike-wave complexes on ictal EEG.

- A genetic component to aetiology is established but the mechanism of inheritance and the genes involved are not fully defined.

- Available evidence suggests that genes encoding brain expressed voltage-gated calcium channels, including CACNG3 on chromosome 16p12-p13.1, may represent susceptibility loci for CAE.

- Assuming locus heterogeneity, a significant HLOD score (HLOD=3.54, α=0.62) was obtained for markers encompassing CACNG3 in 65 nuclear families with a proband with CAE. The maximum NPL score was 2.87 (p<0.002).

- Re-sequencing of the coding exons in 59 patients did not identify any putative causal variants.

- A linkage disequilibrium (LD) map of CACNG3 was constructed using 23 single nucleotide polymorphisms (SNPs). Transmission disequilibrium was sought using individual SNPs and SNP-based haplotypes with the pedigree disequilibrium test in 217 CAE trios and the 65 nuclear pedigrees.

- Evidence for transmission disequilibrium (p0.01) was found for SNPs within a ~35kb region of high LD encompassing the 5'UTR, exon 1 and part of intron 1 of CACNG3.

- Seventy-two variants were identified: 45 upstream; two 5'UTR; and 25 intronic SNPs. No coding sequence variants were identified, although four variants are predicted to affect exonic splicing.

- This evidence supports CACNG3 as a susceptibility locus in a subset of CAE patients.


**Explanations**:

- Context-setting: describes the clinical phenotype (CAE) under study. This is relevant context because the paper investigates genetic involvement in this specific epilepsy phenotype. It is not direct or mechanistic evidence regarding CACNG3 causing a Mendelian disease, but frames the trait under consideration.

- Context and limitation: the sentence explicitly states that while a genetic component exists, inheritance mechanisms and causal genes are not fully defined. This weakens a direct inference that any single gene (including CACNG3) is established as causing a Mendelian disorder, since the authors acknowledge incomplete understanding of inheritance.

- Direct association/supporting statement: the authors state that prior evidence implicates brain-expressed voltage-gated calcium channel genes, including CACNG3, as susceptibility loci for CAE. This is direct text asserting relevance of CACNG3 to CAE susceptibility, but the phrasing "susceptibility loci" implies risk/modifier effects rather than confirmed Mendelian causation.

- Direct genetic evidence (linkage): these sentences report statistically significant linkage metrics for markers around CACNG3 in 65 nuclear families (HLOD=3.54, α=0.62) and a strong NPL (2.87, p<0.002). This is relatively strong evidence that variants near CACNG3 co-segregate with disease in a subset of families (note α=0.62 indicates locus heterogeneity—only ~62% of families are estimated to be linked). While supporting a role for CACNG3 in familial CAE risk, linkage alone does not prove a Mendelian, monogenic cause without identification of segregating causal variants.

- Negative result limiting Mendelian causation: re-sequencing of coding exons in 59 patients failed to find putative causal coding variants. This weakens the claim that CACNG3 causes a Mendelian disease via coding (protein-altering) mutations in these cases; it suggests either non-coding regulatory/splicing variants, rare variants not captured, or locus is a susceptibility locus in a complex manner.

- Study design and sample size context: describes LD mapping using 23 SNPs and a transmission-disequilibrium approach in 217 trios plus the 65 pedigrees. This provides methodological context showing both family-based linkage and TDT/association analyses were performed; it supports the credibility of the association findings but does not by itself establish Mendelian causality.

- Association evidence in a specific non-coding interval: the reported transmission disequilibrium (p0.01) in a ~35kb high-LD region spanning 5'UTR, exon 1 and part of intron 1 implicates non-coding sequence near CACNG3. This is direct association evidence that variants in regulatory or first-exon/intron sequence may be associated with CAE. Such non-coding association supports a susceptibility role but is not definitive proof of a Mendelian, monogenic disease-causing gene.

- Mechanistic possibility and limitation: this reports identification of 72 variants (mostly non-coding) and explicitly states no coding sequence variants were found, but four variants are predicted to affect exonic splicing. This provides mechanistic hypotheses (splicing alteration) that could plausibly affect CACNG3 function; however, prediction alone is not functional proof, and absence of coding mutations limits straightforward classification as a Mendelian disease gene.

- Authors' conclusion: the paper concludes that the evidence supports CACNG3 as a susceptibility locus in a subset of CAE patients. This is a direct statement supporting association/susceptibility but does not claim CACNG3 is definitively a Mendelian disease gene. It emphasizes subset-specific association, consistent with locus heterogeneity and complex inheritance.


[Read Paper](https://www.semanticscholar.org/paper/e6839b4e6023224ebe877b32dca50e1e0319341b)


## Other Reviewed Papers


### Focal epilepsy resulting from a de novo SCN1A mutation.

**Why Not Relevant**: This paper does not provide evidence for or against the claim that CACNG3 is likely associated with at least one Mendelian disease. The text exclusively describes a de novo heterozygous missense mutation in SCN1A (c.5311A>T: I1771F) identified in a single patient with early-onset, fever-associated focal and generalized seizures. There is no mention of CACNG3, no genetic data, biochemical or functional assays, and no mechanistic discussion linking SCN1A to CACNG3. Because the paper focuses on a different gene (SCN1A) and a single-case clinical report, it does not contain direct genetic evidence (e.g., variants, segregation, population enrichment) nor mechanistic evidence (e.g., interaction, pathway perturbation, animal models) relevant to CACNG3. Limitations preventing any inference about CACNG3 from this paper include: (1) absence of any text or data about CACNG3; (2) single-patient case report design (no cohort, no segregation, no statistical evidence); and (3) no functional experiments that could plausibly implicate other genes via pathway perturbation. Therefore the paper is not relevant to the claim about CACNG3.


[Read Paper](https://www.semanticscholar.org/paper/f5c47694412e85dc5749b0bd5e945ac045ea65eb)


### A pathogenic variant in RAB32 causes autosomal dominant Parkinson’s disease and activates LRRK2 kinase

**Why Not Relevant**: The paper investigates genetic variation and functional effects of RAB GTPases in familial Parkinson disease, identifies RAB32 c.213C>G (Ser71Arg) as a pathogenic variant, and presents mechanistic data on LRRK2 activation. The study explicitly targeted RAB family genes (61 RAB GTPases) and reports results for RAB32; it contains no mention, data, analysis, association statistics, segregation, mechanistic links, or discussion regarding the gene CACNG3. Therefore the paper provides no direct genetic evidence, no segregation data, and no mechanistic evidence relevant to the claim that CACNG3 is likely associated with at least one Mendelian disease. The relevance is effectively zero because the subject genes, experimental focus, and reported results do not include CACNG3. Note that lack of mention here is not disproof of the claim generally—it only means this paper does not inform the claim.


[Read Paper](https://www.semanticscholar.org/paper/a36d28e828ede7c87c650af5f5117982b9749e7b)


### Novel LSS variants in alopecia and intellectual disability syndrome: New case report and clinical spectrum of LSS‐related rare disease traits

**Why Not Relevant**: The provided paper content exclusively discusses the gene LSS and its association with several Mendelian phenotypes (congenital cataract type 44, autosomal recessive hypotrichosis type 14, and alopecia–intellectual disability syndrome type 4). Nowhere in the provided text is the gene CACNG3 mentioned, nor are there mechanistic data, variant data, or phenotypic descriptions that implicate CACNG3. Because the claim concerns CACNG3 specifically, the paper provides neither direct evidence (variant–phenotype associations for CACNG3) nor mechanistic evidence (biological pathways, functional studies, or shared molecular mechanisms implicating CACNG3) to support or refute the claim. In short: absence of any mention or data on CACNG3 in this paper means it is not relevant to the claim. Caveat: this conclusion is limited to the provided content. If the full paper contains additional sections or supplementary data not included here that mention CACNG3, those were not available for review. If you want evidence about CACNG3 and Mendelian disease, I can search the literature/databases (OMIM, ClinVar, PubMed) for CACNG3-specific reports and provide extracted supporting or refuting evidence.


[Read Paper](https://www.semanticscholar.org/paper/fe90a5bb55278bc80588741eed5def6cf4f1aa82)


### GABRB1-related early onset developmental and epileptic encephalopathy: clinical trajectory and novel de novo mutation.

**Why Not Relevant**: This paper exclusively reports on DEE45 caused by pathogenic variants in GABRB1 and provides clinical and genetic data specific to GABRB1 (including a novel de novo GABRB1 mutation). The content contains no mention of CACNG3, no data on CACNG3 sequence variation, no patient phenotypes attributed to CACNG3, and no mechanistic discussion linking CACNG3 to Mendelian disease. Therefore the paper provides no direct or mechanistic evidence for or against the claim that CACNG3 is likely associated with at least one Mendelian disease. Note: absence of CACNG3 in this paper does not disprove the claim; it only means this paper is irrelevant to evaluating that claim. To assess the claim, literature that specifically investigates CACNG3 (genetic studies, case reports, functional assays, or gene-disease curation resources) should be examined.


[Read Paper](https://www.semanticscholar.org/paper/126396673b708f9178dc56b3859fd7963cfa91ea)


### Whole exome sequencing identified a homozygous novel variant in DOP1A gene in the Pakistan family with neurodevelopmental disabilities: case report and literature review

**Why Not Relevant**: This paper describes a novel homozygous missense variant in DOP1A (DOP1A(NM_001385863.1):c.2561A>G) and proposes DOP1A as a potential gene for neurodevelopmental disorders related to disrupted myelinogenesis. Nowhere in the provided content is the gene CACNG3 mentioned, nor are any variants, functional data, mechanistic pathways, co-expression, or clinical associations involving CACNG3 reported. Therefore the paper provides neither direct genetic evidence (e.g., variants, segregation, case reports) nor mechanistic evidence (e.g., biochemical pathways, functional assays) linking CACNG3 to any Mendelian disease. Note: the absence of evidence for CACNG3 in this paper does not imply CACNG3 is not associated with disease; it only means this paper is irrelevant to evaluating the claim. To assess the claim about CACNG3, one should consult gene-disease databases (OMIM, ClinVar), the primary literature for case reports or functional studies on CACNG3, and large-scale sequence/variant databases (gnomAD, DECIPHER).


[Read Paper](https://www.semanticscholar.org/paper/16845c10ba324247707bc427c38d7f3eb44d5ec1)


### RNA sequencing reveals a complete picture of a homozygous missense variant in a patient with VPS13D movement disorder: a case report and review of the literature

**Why Not Relevant**: The provided paper content discusses a single case involving VPS13D, describing RNA-seq functional characterization of a homozygous missense variant of uncertain significance in VPS13D, its impact on splicing, and a diagnosis of a VPS13D movement disorder. There is no mention of the gene CACNG3, no clinical or genetic data linking CACNG3 to disease, and no mechanistic discussion involving CACNG3. Because the content does not address CACNG3 in any way, it provides no direct or mechanistic evidence for or against the claim that CACNG3 is likely associated with at least one Mendelian disease. Additional reasons for irrelevance: (1) the methods/results relate to a different gene (VPS13D); (2) variant type, functional assay (RNA-seq confirming splicing), and diagnosis are all specific to VPS13D; (3) no comparative or cross-gene inference is presented that would allow extrapolation to CACNG3. To evaluate the claim about CACNG3, primary literature, case reports, or functional studies directly involving CACNG3 would be required.


[Read Paper](https://www.semanticscholar.org/paper/0655bdb1e80b122fdb5b84f07ee4efd77c63b603)


### Case Report: human early embryonic arrest in a consanguineous Chinese family caused by a novel missense variant of PADI6.

**Why Not Relevant**: The provided paper exclusively investigates PADI6 and reports a novel recessive PADI6 missense variant underlying female infertility by early embryonic arrest. Nowhere in the supplied text is CACNG3 mentioned, nor are any data, analyses, or mechanistic discussions related to CACNG3 included. Therefore the paper provides no direct genetic evidence (variants, segregation, case reports) nor mechanistic evidence (biological pathway links, functional studies) that would support or refute the claim that CACNG3 is likely associated with a Mendelian disease. While the study demonstrates the use of whole-exome sequencing and variant interpretation methods that can identify Mendelian disease genes, those methodological points do not constitute evidence about CACNG3 specifically. For these reasons the relevance of this paper to the claim about CACNG3 is effectively zero.


[Read Paper](https://www.semanticscholar.org/paper/3bc31c0fecd83bf6aae9393b6c538fa9129a5f57)


## Search Queries Used

- CACNG3 variant pathogenic Mendelian disease human case report family segregation exome sequencing

- CACNG3 epilepsy seizure childhood absence epilepsy genetic association linkage mutation de novo

- CACNG3 voltage dependent calcium channel gamma subunit 3 TARP gamma 3 AMPA receptor trafficking synaptic function knockout mouse seizure phenotype functional study

- CACNG3 ClinVar gnomAD loss of function constraint pLI missense constraint gene disease association review replication negative study


## Usage Statistics

- Prompt Tokens: 0

- Completion Tokens: 0

- Total Tokens: 0

- Estimated Cost: $0.0484


## Bibliometric Configuration

- Use Bibliometrics: True

- Author Impact Weight: 0.4

- Citation Impact Weight: 0.4

- Venue Impact Weight: 0.2
