## Supplementary material for "MENDELSEEK: An algorithm that predicts Mendelian Genes and elucidates what makes them special": CAMSAP2.txt

# Claim: The gene CAMSAP2 is likely associated with at least one Mendelian disease.

**Status**: processed

**Overall Rating**: -1

**Explanation**:

No relevant papers were found that support or refute this claim.


## Relevant Papers


## Other Reviewed Papers


### Precision Medicine in Internal Medicine

**Why Not Relevant**: The provided paper is a general review of genetic testing strategies, exome/genome sequencing, and challenges in variant interpretation. It contains no mention of the gene CAMSAP2, no genetic, clinical, or functional data concerning CAMSAP2 variants, and no analyses (case reports, cohort studies, segregation analyses, functional assays, or statistical associations) that would support or refute an association between CAMSAP2 and any Mendelian disease. The excerpts extracted are background and mechanistic context that explain how gene-disease associations are discovered and why many genes remain unassigned, but they do not constitute evidence about CAMSAP2 specifically. Therefore the paper provides essentially no direct or gene-specific mechanistic evidence relevant to the claim that CAMSAP2 is likely associated with at least one Mendelian disease.


[Read Paper](https://www.semanticscholar.org/paper/c7199e8a5226f8d9d55777da0e15e91797bb459a)


### Clinical and Genetic Characteristics of Patients with Unexplained Intellectual Disability/Developmental Delay without Epilepsy

**Why Not Relevant**: The paper content provided does not mention the gene CAMSAP2 anywhere in the Introduction, Methods, Results, or Conclusion excerpts. The study reports identified pathogenic/likely pathogenic variants and lists eight novel variants in other genes, but CAMSAP2 is not among them. Therefore, the paper provides no direct positive evidence that CAMSAP2 is associated with a Mendelian disease. The absence of CAMSAP2 in this small, disorder-specific cohort and in the reported findings should not be taken as strong evidence against CAMSAP2 involvement in Mendelian disease generally, because limitations include the cohort size (38 cases), the specific phenotypic focus (ID/DD/ASD), possible gaps in sequencing coverage or variant filtering for CAMSAP2, and the authors' choice to report only certain findings. Overall, relevance to the claim is negligible because CAMSAP2 is not discussed or implicated in the paper.


[Read Paper](https://www.semanticscholar.org/paper/e5289119c3a34a33fd81c6fcc009413eb467af22)


### Implications of a De Novo Variant in the SOX12 Gene in a Patient with Generalized Epilepsy, Intellectual Disability, and Childhood Emotional Behavioral Disorders

**Why Not Relevant**: Relevance is effectively zero because the provided paper content exclusively addresses SOX family genes, specifically a novel de novo SOX12 variant, its predicted pathogenicity, and mechanistic context within the HMG domain. CAMSAP2 is not mentioned anywhere in the provided text. There is no experimental data, genetic association, clinical observation, mechanistic discussion, or literature citation relating to CAMSAP2. Absence of CAMSAP2 discussion in this paper means it provides neither direct evidence (genetic/clinical association) nor mechanistic evidence (biological pathway overlap or functional similarity) to support or refute the claim that CAMSAP2 is likely associated with at least one Mendelian disease. Note that absence of evidence in this single SOX12-focused study is not evidence that CAMSAP2 is or is not associated with a Mendelian disease in other studies; it simply means this paper is not relevant to the claim.


[Read Paper](https://www.semanticscholar.org/paper/5cf652470efa4406f72843aea8389f4e398a99f8)


### The simplest explanation does not have to be preferred: co-occurrence of pathogenic variants in cancer-predisposing genes

**Why Not Relevant**: The provided paper content does not mention CAMSAP2 or any evidence linking that gene to Mendelian disease. The single sentence discusses frequency and cancer phenotypes associated with MINAS, review of recent literature, and checking UK 10,000 Genomes Project data for participants with tumor predisposition syndromes or multiple primary tumors, and whether effects of Cancer Susceptibility Genes (CSGs) are additive or synergistic. There is no direct evidence (case reports, segregation, pathogenic variants, statistical association) referencing CAMSAP2, and no mechanistic description connecting CAMSAP2 function to a Mendelian disorder. Methodological details and results that might contain gene-level findings are absent from the excerpt, so no inference about CAMSAP2 can be made. To evaluate the claim that CAMSAP2 is likely associated with at least one Mendelian disease would require gene-specific data (e.g., reported pathogenic variants, segregation in pedigrees, functional studies, population allele frequencies, constraint metrics) which are not present in this content. Recommended next steps: search gene-disease databases (OMIM, ClinVar, HGMD), PubMed for CAMSAP2 case reports or functional studies, and population/constraint resources (gnomAD) to assess plausibility.


[Read Paper](https://www.semanticscholar.org/paper/a1372a664bd2c33a0bda638975a8b26d112457fc)


### TTC7A Variants Previously Described to Cause Enteropathy Are Observed on a Single Haplotype and Appear Non-pathogenic in Pediatric Inflammatory Bowel Disease Patients

**Why Not Relevant**: The provided content exclusively describes variants in TTC7A and assessment of TTC7A in polygenic IBD: "This study describes the two variants observed in trans by Lawless et al. of compound heterozygote variants in the Tetratricopeptide Repeat Domain 7A gene, and assesses the evidence for variation in TTC7A contributing to polygenic IBD." There is no mention of CAMSAP2, no genetic association data for CAMSAP2, no phenotypes linked to CAMSAP2, and no mechanistic discussion involving CAMSAP2 in the text you supplied. Therefore this paper content provides no direct evidence (no observations, association statistics, segregation, or case reports) and no mechanistic evidence (no functional studies, pathways, or interactions) relevant to the claim that CAMSAP2 is likely associated with at least one Mendelian disease. Caveat: this assessment is limited to the exact content you provided. If the full paper contains sections not included here that mention CAMSAP2, those were not available for review and could alter the relevance assessment.


[Read Paper](https://www.semanticscholar.org/paper/3818a966b5d6e8b0eda65fdef043ac0727fa429c)


### Germline Variant Interpretation in Children with Severe Sepsis

**Why Not Relevant**: The provided paper content does not mention CAMSAP2 at all, nor does it include a gene list, variant-level data, or mechanistic descriptions that could link CAMSAP2 to a Mendelian disease. The single sentence supplied only reports the number of patients with clinically significant variants in genes associated with inborn errors of immunity; it therefore cannot be used to support or refute the claim about CAMSAP2. Without the full paper (including the list of genes/variants identified, phenotypic correlations, statistical analyses, and functional or segregation data), one cannot determine whether CAMSAP2 was observed or implicated. To assess the claim properly would require the full methods and results (gene lists, variant annotations, inheritance/segregation data, functional studies) from the study or other sources that explicitly report CAMSAP2 in Mendelian disease contexts.


[Read Paper](https://www.semanticscholar.org/paper/8148b944a985ef1eb6ab7a15ba1b7990c783459a)


### Molecular and phenotypical findings of a novel de novo SYNGAP1 gene variant in an 11-year-old Iranian boy with intellectual disability.

**Why Not Relevant**: The paper's content exclusively discusses SYNGAP1: its role in intellectual developmental disorder type 5, the identification of a de novo nonsense variant in SYNGAP1, and clinical implications for SYNGAP1-related phenotypes. There is no mention of the gene CAMSAP2, no experimental results, case reports, population analyses, nor mechanistic discussion that involve CAMSAP2 or molecular pathways linking SYNGAP1 to CAMSAP2. Therefore this paper provides no direct evidence (no reported CAMSAP2 variants or phenotypes) and no mechanistic evidence (no pathway or functional data involving CAMSAP2) to support or refute the claim that CAMSAP2 is likely associated with a Mendelian disease. Because the gene investigated is different and there are no cross-references or data relevant to CAMSAP2, the paper is not relevant to the claim.


[Read Paper](https://www.semanticscholar.org/paper/3f4b2eee9f7ad16af6b2e8b2a6417e1d72f20aed)


## Search Queries Used

- CAMSAP2 pathogenic variant Mendelian disease monogenic human germline pathogenic ClinVar case report segregation

- CAMSAP2 de novo variant neurodevelopmental disorder intellectual disability autism developmental delay epilepsy exome sequencing trio study

- CAMSAP2 microtubule minus end binding neuronal polarity axon dendrite development neuron migration mouse knockout zebrafish model congenital brain malformation

- CAMSAP family CAMSAP1 CAMSAP3 CAMSAP2 rare disease exome sequencing cohort Mendelian gene discovery gene burden pLI constraint gnomAD population genetics


## Bibliometric Configuration

- Use Bibliometrics: True

- Author Impact Weight: 0.4

- Citation Impact Weight: 0.4

- Venue Impact Weight: 0.2
