## Supplementary material for "MENDELSEEK: An algorithm that predicts Mendelian Genes and elucidates what makes them special": CCDC6.txt

# Claim: The gene CCDC6 is likely associated with at least one Mendelian disease.

**Status**: processed

**Overall Rating**: 0

**Explanation**:

Supporting evidence:
The provided paper documents CCDC6 as a recurrent fusion partner in papillary thyroid carcinoma (PTC), reporting CCDC6-RET gene fusions (alongside other RET and NTRK1 fusions). That establishes a clear, reproducible role for CCDC6 in oncogenic somatic rearrangements contributing to PTC tumorigenesis. This is evidence of CCDC6 involvement in human disease processes.

Caveats or contradictory evidence:
The excerpts explicitly describe CCDC6 involvement as part of somatic gene fusions in tumors. There is no mention in the provided material of germline CCDC6 variants, familial segregation, or any inherited (Mendelian) disorder associated with CCDC6. Somatic driver alterations in cancer (e.g., fusion genes) are mechanistically and clinically distinct from germline mutations that produce Mendelian inheritance patterns. The paper even separates findings of somatic alterations from the three germline variants it reports (ATM, MET, RET), none of which are CCDC6. Thus the evidence does not support — and does not provide examples of — CCDC6 acting as a Mendelian disease gene.

Analysis of mechanisms and deeper relationships:
CCDC6 is known (from the paper excerpt) to form fusion oncoproteins (CCDC6-RET) that produce constitutive kinase activity driving tumorigenesis; that is a somatic, gain-of-function oncogenic mechanism. Mendelian disease associations typically require germline variants that segregate with disease in families and often operate via loss-of-function, dominant-negative, or specific gain-of-function mechanisms present in the germline. The presence of frequent somatic fusions suggests that CCDC6 has relevance to disease biology, but it does not imply germline Mendelian causation. It remains possible, in principle, that rare germline CCDC6 variants could cause inherited disease, but the provided evidence neither demonstrates nor suggests such a relationship.

Assessment and rating rationale:
Based on the provided paper excerpts, there is no evidence that CCDC6 is associated with any Mendelian (inherited) disease. The paper provides strong evidence for somatic, cancer-related involvement of CCDC6 (fusion with RET) but no germline or familial data linking CCDC6 to Mendelian inheritance. Because the available evidence does not support the specific claim (and only documents somatic oncogenic roles), the appropriate assessment is that there is no evidence in the provided material to support the claim that "CCDC6 is likely associated with at least one Mendelian disease." Therefore the correct rating is "No Evidence."


**Final Reasoning**:

The single provided paper documents CCDC6 as a somatic fusion partner (CCDC6-RET) in papillary thyroid carcinoma, which is disease-relevant but explicitly somatic. The excerpts contain no report of germline CCDC6 variants, no familial segregation data, and no statement that CCDC6 causes an inherited disorder. Somatic oncogenic fusions do not by themselves establish a Mendelian disease association. Given the absence of any germline/Mendelian evidence in the provided material, the claim that CCDC6 is likely associated with at least one Mendelian disease is unsupported; therefore the most accurate rating with the supplied evidence is "No Evidence."


## Relevant Papers


### Genomic characterization of papillary thyroid carcinoma in Chinese Shandong population.

**Authors**: Xungeng Li (H-index: 4), Ling Li (H-index: 3)

**Relevance**: 0.15

**Bibliometric Impact**: 0.20840000000000003


**Excerpts**:

- Somatic and germline mutations were identified including 38 genes associated with tumor development.

- Sequencing data were analyzed to call tumor specific single nucleotide variants (SNV), small insertions and deletions (InDels), copy number alterations (CNA), gene fusions and rearrangements.

- Gene fusions, involving the RET and NTRK1 genes, were identified in three PTCs (2.4%), including NTRK1-TPM3, NTRK1-SQSTM1 and CCDC6-RET.

- And the three fusions patients were all BRAF V600E-negative cases.

- We also found that three PTC patients carried germline variants of uncertain significance, including ATM (p.L857P), MET (p.L211W) and RET (p.C515R), which are heterozygous mutations.


**Explanations**:

- This sentence indicates the study explicitly assessed both somatic and germline mutations across a panel of 38 genes. Relevance: indirect. Strength: shows germline testing was performed in the cohort. Limitation: it does not specify whether CCDC6 was among the 38 genes assessed for germline variation, nor does it report any germline CCDC6 findings.

- This sentence documents that the sequencing pipeline called gene fusions and rearrangements in addition to SNVs and CNAs. Relevance: indirect/mechanistic. Strength: establishes that fusion events (which could include CCDC6 rearrangements) were actively sought and reported. Limitation: does not by itself say which fusions were found or whether any germline structural variants were assessed.

- This is the key direct excerpt mentioning CCDC6: it reports a CCDC6-RET fusion among somatic gene fusions in papillary thyroid carcinoma samples. Relevance: mechanistic but not Mendelian. Strength: provides direct evidence that CCDC6 participates in somatic oncogenic rearrangements (CCDC6-RET fusion) in PTC, supporting a role for CCDC6 in tumorigenesis through fusion-driven activation of RET. Limitation: this is a somatic tumor fusion and therefore does not demonstrate that CCDC6 is associated with any Mendelian (inherited) disease; it does not report germline CCDC6 pathogenic variants nor familial segregation data.

- This sentence provides context that the fusion-positive tumors were BRAF V600E-negative, indicating mutual exclusivity patterns often seen in PTC. Relevance: mechanistic/contextual. Strength: helps interpret the biological role of the CCDC6-RET fusion within tumor molecular subtypes. Limitation: still tumor-contextual and does not address germline inheritance or Mendelian disease association.

- This sentence shows the study identified germline variants of uncertain significance in three patients, none of which were in CCDC6. Relevance: indirect/evidence against (in this cohort). Strength: demonstrates that germline analysis was done and lists specific germline findings; implies no germline CCDC6 variants were observed among reported germline findings. Limitation: small cohort (127 patients), the panel covered only 38 genes (it is not explicitly stated that CCDC6 was included among those for germline interpretation), and the absence of observed germline CCDC6 variants in this tumor cohort is weak evidence against a Mendelian association generally.


[Read Paper](https://www.semanticscholar.org/paper/1bb48ac5b5a75f65e680cdc4451047885b6ac0c8)


## Other Reviewed Papers


### The Burden of Candidate Pathogenic Variants for Kidney and Genitourinary Disorders Emerging From Exome Sequencing

**Why Not Relevant**: The provided paper content does not mention the gene CCDC6 anywhere in the excerpts or the surrounding text. The study focuses specifically on identification, curation, and annotation of genes associated with Mendelian kidney and genitourinary disorders and reports a final curated list of 625 genes; however, the provided content does not include the full gene list or any gene-level results beyond describing the curation process and cohort. Consequently, there is no direct evidence (variant observations, statistical association, case reports) or mechanistic evidence (functional studies, molecular pathways) in the supplied text that bears on whether CCDC6 is associated with any Mendelian disease. Limitations that prevent assessing the claim from this paper content: (1) absence of the CCDC6 name or any CCDC6-specific data in the provided excerpts; (2) the paper is scoped to kidney/genitourinary Mendelian disorders—if CCDC6 is associated with a Mendelian disease outside those categories it would not necessarily be included or discussed; (3) the full curated gene list (not included here) might or might not contain CCDC6, but without that list or explicit mention, no determination can be made. Recommendation: to evaluate the claim about CCDC6, consult gene-centric resources such as OMIM (entry for CCDC6), ClinVar, HGMD, GeneReviews, or the full supplementary gene list referenced by this paper (the laboratory web site), and examine case reports, segregation data, loss-of-function constraint metrics, and functional studies specific to CCDC6.


[Read Paper](https://www.semanticscholar.org/paper/a75e297208c04992568e607d19aa9b9aaebd09d5)


### Familial Whole Exome Sequencing Study of 30 Families With Early-Onset High Myopia

**Why Not Relevant**: The paper does not mention the gene symbol CCDC6 anywhere in the provided content. The gene lists and specific mutated genes reported (including CCDC111, SLC39A5, P4HA2, CPSF1, GRM6, etc.) do not include CCDC6. Therefore there is no direct supporting or refuting evidence in this paper regarding whether CCDC6 is associated with any Mendelian disease. The absence of CCDC6 in this cohort is weak negative evidence at best (it may simply not have been mutated in these 30 families or filtered out), and cohort- and phenotype-specific studies cannot definitively determine a gene's Mendelian status across all diseases. Additionally, the presence of CCDC111 in the text should not be interpreted as CCDC6—these are distinct gene symbols. Because the paper provides no direct mention, data, mechanistic discussion, or functional experiments concerning CCDC6, its relevance to the claim is effectively zero.


[Read Paper](https://www.semanticscholar.org/paper/8a51e1e2b13da3e9075f4d54e506e745280ab56b)


### Precision Medicine in Internal Medicine

**Why Not Relevant**: This paper is a general review of genetic testing strategies, variant interpretation, and the current scope of known Mendelian genes; it does not mention the gene CCDC6, does not present any data (genetic, clinical, segregation, or functional) related to CCDC6, and provides no mechanistic discussion that links CCDC6 to any Mendelian phenotype. The only statement in the provided text that even approaches gene-level context is a general statistic: "As of 1 February 2019, the Online Mendelian Inheritance in Man database indicated that out of 20000 protein-coding genes, only 3652 (approximately 18%) had been found to cause known single-gene disorders." That sentence provides background context about the overall fraction of genes with established Mendelian associations but does not provide any evidence for or against an association of CCDC6 with a Mendelian disease.

Because the paper contains no mention of CCDC6, there is no direct evidence (case reports, variant co-segregation, pathogenic variants, statistical association) and no mechanistic evidence (gene function, pathogenic mechanism, animal models, molecular pathways) in this text to support or refute the claim that "CCDC6 is likely associated with at least one Mendelian disease." Absence of mention in a general review does not constitute evidence that CCDC6 is or is not associated; it only indicates that the paper did not address that gene. The coverage date (text cites OMIM as of 1 February 2019) also matters: associations discovered after that date would not be discussed here.

Recommended next steps to evaluate the claim (not provided in this paper): search OMIM, ClinVar, HGMD, PubMed, and primary literature for "CCDC6" + "disease" or specific phenotype terms; check gnomAD for constraint metrics and allele frequencies; look for case reports, segregation studies, and functional experiments (cellular assays, animal models) implicating CCDC6; and review any expert curation (ClinGen gene-disease validity) entries for CCDC6. These sources would provide the direct and mechanistic evidence needed to assess the claim.


[Read Paper](https://www.semanticscholar.org/paper/c7199e8a5226f8d9d55777da0e15e91797bb459a)


### Whole-Exome Sequencing in Adults With Chronic Kidney Disease

**Why Not Relevant**: This paper excerpt does not mention the gene CCDC6 anywhere. The study focuses on using whole-exome sequencing to identify Mendelian causes of chronic kidney disease and hypertension and describes their gene lists (287 nephropathy OMIM genes and the ACMG 59 actionable genes), sequencing methods, coverage, variant annotation, and limitations of exome capture. While these procedural details are useful to assess whether the study could have detected pathogenic variants in a given gene, the provided text gives no gene-level results, no table or list of genes found to be associated with disease, and no explicit inclusion or exclusion of CCDC6 from their analyses. Therefore the excerpt provides no direct evidence that CCDC6 is associated (or not associated) with any Mendelian disease. Additionally, WES capture gaps and the study's kidney-focused gene list limit the ability to infer anything about CCDC6 from this paper fragment. For these reasons, the manuscript content supplied is not relevant to establishing the claim that CCDC6 is likely associated with at least one Mendelian disease.


[Read Paper](https://www.semanticscholar.org/paper/a29bd57964f68ac5f8e1abcafe1112ba488f53f5)


### Abstract 12295: Utilization of Whole-Exome Sequencing to Identify Causative Mutations in Familial Congenital Heart Disease

**Why Not Relevant**: The paper does not mention the gene CCDC6 anywhere in the provided content. The study focused on identifying pathogenic variants in familial congenital heart disease using a candidate list of 69 CHD genes and reports causative variants in MYH11, GATA4, and TLL1. Because CCDC6 is neither reported among the identified variants nor discussed, there is no direct evidence in this paper supporting an association between CCDC6 and any Mendelian disease. Furthermore, the paper's approach prioritized variants matching a predefined CHD gene list; if CCDC6 was not included on that list or if CCDC6-related phenotypes were outside the study's scope (CHD), the study would not have reported CCDC6 even if variants were present in the WES data. Therefore this paper provides no direct or mechanistic evidence for CCDC6 being associated with a Mendelian disease. Limitations that prevent concluding more strongly: small sample size (9 families), disease focus limited to CHD, and the use of a 69-gene candidate list which may have excluded other Mendelian disease genes including CCDC6.


[Read Paper](https://www.semanticscholar.org/paper/493c804828ca2e05804dd3041a93194ef222b680)


### P-540 Strategic implementation of PGT-M for multiple genes post qhole exome sequencing in couples with familial history: a comprehensive study

**Why Not Relevant**: The provided paper excerpt describes a retrospective WES study of couples referred for familial genetic disorders, reporting overall diagnostic yield, sequencing methods, and outcomes of PGT-M. Nowhere in the text is the gene name CCDC6 mentioned, nor are per-family gene lists, specific gene-disease associations, or any variant-level data that would allow assessment of CCDC6's involvement in a Mendelian disease. Therefore the paper provides no direct evidence that CCDC6 is associated with any Mendelian disease. It offers only general mechanistic context showing that WES can detect Mendelian disease genes and that the study design and sequencing depth were adequate to find coding pathogenic variants if present. Because the absence of any mention or data on CCDC6 prevents either support or refutation of the claim, the relevance to the specific claim that "CCDC6 is likely associated with at least one Mendelian disease" is effectively zero.


[Read Paper](https://www.semanticscholar.org/paper/1b721e406f9d31ac5983b146c5b9aa03d58fc49c)


### RF01 | PMON167 Isolated Cushing's Disease Associated With Rare Germline SDHx Variants

**Why Not Relevant**: This paper focuses on germline loss-of-function variants in the SDHx genes (SDHA, SDHB, SDHC, SDHD) as a rare cause of pituitary tumors and Cushing's disease. The provided text repeatedly names SDHx genes and other genes tested in tumors (USP8, USP48, BRAF, DICER1, TSC2) but contains no mention of the gene CCDC6. Therefore it provides no direct evidence for or against an association between CCDC6 and any Mendelian disease. Specific reasons why this paper is not relevant to the claim: 

- The abstract/introduction and results exclusively report variants and expression changes in SDHx subunits and a few other genes (USP8, USP48, BRAF, DICER1, TSC2). CCDC6 is not mentioned in any of the provided text.
- Although whole-exome sequencing (WES) was performed on many samples, the authors report on SDHx findings and a small set of tumor genes; absence of a reported CCDC6 result does not mean WES found no CCDC6 variants—more likely CCDC6 was not a focus or not reported. Thus there is no direct negative evidence either.
- The study cohort (primarily pediatric Cushing's disease patients) and the variant-focused analysis on SDHx limit any inference about other genes' disease associations; even if WES had data on CCDC6, the paper does not present it or discuss mechanisms linking CCDC6 to disease.

Limitations to this conclusion: absence of CCDC6 in the reported results cannot be interpreted as evidence that CCDC6 is not associated with Mendelian disease; it only means this paper does not address the claim. To evaluate the claim about CCDC6, one must consult studies that specifically examine CCDC6 variants, their segregation with disease, functional/ mechanistic data implicating CCDC6 in Mendelian inheritance, or curated resources (ClinVar, OMIM, gene-disease curation by ClinGen).


[Read Paper](https://www.semanticscholar.org/paper/369b45c751701de632cf46e834d78a2cc8297079)


### Mapping genetic variations to three-dimensional protein structures to enhance variant interpretation: a proposed framework

**Why Not Relevant**: The provided paper content is a workshop overview and methods-focused review about integrating genetic variants with 3D protein structures. The excerpt contains no mention of the gene CCDC6, no gene-specific data, no variant-level evidence, no disease associations, and no family segregation or Mendelian-disease analyses. Examples cited in the text (e.g., TRIM11, VHL) are other genes and illustrate methods rather than providing evidence about CCDC6. The paper is not a primary association study and does not present experimental or clinical data linking any specific gene to a Mendelian disorder in the provided content. Therefore, based on the excerpt supplied, there is no direct evidence for or against the claim that CCDC6 is likely associated with at least one Mendelian disease. Limitations: this assessment is limited to the supplied excerpt; the full paper or supplemental materials (not included here) might mention CCDC6, but no such content appears in the text provided. For a definitive evaluation, search the full text and relevant variant/disease databases (ClinVar, OMIM, HGMD, gnomAD) or primary literature for CCDC6-specific evidence.


[Read Paper](https://www.semanticscholar.org/paper/ae68aa934d5a5ad3d9009bb8811fa8d5fc4dbb1c)


### Abstract 2545: Identification of germline pathogenic alleles in 143 cancer predisposing genes in Mexican patients with hereditary breast cancer by massive parallel sequencing

**Why Not Relevant**: The abstract does not mention CCDC6 anywhere among the genes with pathogenic variants or VUS. It reports sequencing of 143 genes but does not provide a complete gene list, so we cannot confirm whether CCDC6 was included in the panel. The content therefore offers only weak, indirect negative evidence (absence from reported results) and provides no direct genetic association, mechanistic data, functional studies, or clinical-case evidence linking CCDC6 to any Mendelian disease. The cohort is small (69 patients) and restricted to Mexican hereditary breast cancer cases and to coding regions and splice sites; negative or missing evidence in this specific study is neither sufficient to support nor to refute the broader claim that "CCDC6 is likely associated with at least one Mendelian disease." Consequently, this paper is not a relevant source of evidence for the claim.


[Read Paper](https://www.semanticscholar.org/paper/f81c0918b71e08be5a70376c8526d47e8d0fc04f)


## Search Queries Used

- CCDC6 germline mutation Mendelian disease familial segregation case report whole exome sequencing monogenic inherited pathogenic variant

- CCDC6 ClinVar OMIM HGMD pathogenic variant clinical significance gene disease association germline annotation

- CCDC6 function DNA damage response genomic stability apoptosis haploinsufficiency loss of function mouse knockout phenotype developmental defect

- CCDC6 RET fusion papillary thyroid carcinoma somatic fusion oncoprotein constitutional germline somatic distinction


## Usage Statistics

- Prompt Tokens: 0

- Completion Tokens: 0

- Total Tokens: 0

- Estimated Cost: $0.0540


## Bibliometric Configuration

- Use Bibliometrics: True

- Author Impact Weight: 0.4

- Citation Impact Weight: 0.4

- Venue Impact Weight: 0.2
