## Supplementary material for "MENDELSEEK: An algorithm that predicts Mendelian Genes and elucidates what makes them special": CCL7.txt

# Claim: The gene CCL7 is likely associated with at least one Mendelian disease.

**Status**: processed

**Overall Rating**: -1

**Explanation**:

No relevant papers were found that support or refute this claim.


## Relevant Papers


## Other Reviewed Papers


### STAT3 mutations in the hyper-IgE syndrome.

**Why Not Relevant**: Relevance is very low because the paper exclusively reports findings implicating STAT3 in hyper-IgE (Job's) syndrome and does not mention the gene CCL7 (also known as MCP-3). The only chemokine explicitly named is "monocyte chemoattractant protein 1," which corresponds to CCL2 (MCP-1), not CCL7. Therefore the paper contains no direct genetic association, mechanistic data, expression evidence, or case reports implicating CCL7 in any Mendelian disease. Absence of evidence in this paper is not proof that CCL7 is not associated with a Mendelian disease, but this paper provides no support for the claim. Additional literature specifically examining CCL7/CCL7 variation, segregation in families, or functional effects would be required to evaluate the claim.


[Read Paper](https://www.semanticscholar.org/paper/c1d2bbab355088e1570dc5a942519f6b8ea31124)


### The Burden of Candidate Pathogenic Variants for Kidney and Genitourinary Disorders Emerging From Exome Sequencing

**Why Not Relevant**: The provided paper content does not mention the gene CCL7 nor provide any gene-level data or specific variant evidence for CCL7. The excerpt describes the study's goals, cohort, general methods for identifying and curating genes associated with Mendelian kidney and genitourinary disorders, and the size of the final curated gene list (625 genes), but it does not enumerate those genes or present any results for individual genes. Therefore there is no direct evidence (e.g., reports of pathogenic variants, genotype–phenotype correlations, inheritance mode, or case descriptions) nor mechanistic evidence (e.g., functional studies, molecular pathways implicating CCL7 in Mendelian disease) for or against the claim in the provided text. Key limitations preventing assessment include: (1) absence of the gene name CCL7 in the excerpt; (2) lack of gene-by-gene results or tabulated supplementary lists in the supplied content; and (3) no presentation of variant-level data, case reports, or functional assays relating to CCL7. If the full paper or its supplementary materials (the laboratory website, Appendix files, or the curated gene list referenced) were available, CCL7 might appear there; that possibility cannot be evaluated from the provided content. To adjudicate the claim one should examine the paper's full gene list/supplementary tables and external resources (OMIM, ClinVar, HGMD, or primary case reports) for: reported pathogenic CCL7 variants, inheritance pattern, segregation or de novo occurrence, population frequency incompatible with benign variation, and any functional evidence supporting pathogenicity.


[Read Paper](https://www.semanticscholar.org/paper/a75e297208c04992568e607d19aa9b9aaebd09d5)


### The simplest explanation does not have to be preferred: co-occurrence of pathogenic variants in cancer-predisposing genes

**Why Not Relevant**: The provided paper content comprises a single sentence describing reviewing literature and checking the UK 10,000 Genomes Project for cancer phenotypes and whether cancer-susceptibility genes act additively or synergistically. It contains no mention of the gene CCL7, no data, results, variants, or mechanistic discussion pertaining to CCL7, and no statements about Mendelian disease associations. Because the excerpt does not name or describe CCL7 or provide any genetic, clinical, or mechanistic evidence relevant to the claim, it cannot support or refute the claim that CCL7 is likely associated with at least one Mendelian disease. Therefore relevance is effectively zero.


[Read Paper](https://www.semanticscholar.org/paper/a1372a664bd2c33a0bda638975a8b26d112457fc)


### Primary immune deficiencies − improving our understanding of their role in immunological disease

**Why Not Relevant**: The provided paper content is a meeting report/classification update summarizing known primary immunodeficiency syndromes and listing specific genes added to various classification tables. The excerpted text names multiple immune-related genes (DNA ligase IV, CD3δ, AID, UNG, CD40, CD40L, ICOS, Rac2, β actin, MBP, MASP2, etc.) but contains no mention of the chemokine gene CCL7 nor any discussion of chemokines as Mendelian disease genes. Therefore the document, as provided, offers no direct or mechanistic evidence linking CCL7 to a Mendelian disease. Important caveats: (1) absence of mention in this excerpt is not proof that CCL7 has no Mendelian disease associations elsewhere in the literature; (2) the report is a high-level classification/review rather than a primary study establishing novel gene–disease causation; (3) only the supplied text was available for analysis—other parts of the full report (not provided) could conceivably mention CCL7, but there is no evidence of that here. Given these points, the relevance of this paper excerpt to the claim is negligible.


[Read Paper](https://www.semanticscholar.org/paper/c02ac4a2463f6f1b2f1a688fc0261b7f0e4471eb)


## Search Queries Used

- CCL7 MCP3 SCYA7 germline variant pathogenic mutation Mendelian monogenic disease case report OMIM ClinVar

- CCL7 chemokine monocyte chemoattractant protein 3 primary immunodeficiency autoinflammatory congenital immune disorder single gene familial mutation

- CCL7 knockout mouse phenotype CCR2 signaling chemokine pathway loss of function gain of function functional study model organism developmental defect inflammation

- CCL7 Mendelian disease review systematic review gene disease validity ClinGen DECIPHER ClinVar gene curation rare disease gene list


## Bibliometric Configuration

- Use Bibliometrics: True

- Author Impact Weight: 0.4

- Citation Impact Weight: 0.4

- Venue Impact Weight: 0.2
