## Supplementary material for "MENDELSEEK: An algorithm that predicts Mendelian Genes and elucidates what makes them special": CD38.txt

# Claim: The gene CD38 is likely associated with at least one Mendelian disease.

**Status**: processed

**Overall Rating**: -1

**Explanation**:

No relevant papers were found that support or refute this claim.


## Relevant Papers


## Other Reviewed Papers


### An open resource of structural variation for medical and population genetics

**Why Not Relevant**: The paper provides population-scale evidence that structural variants can disrupt or duplicate protein-coding genes and that such events contribute substantially to predicted loss-of-function variation and clinically reportable findings. These points give mechanistic plausibility that gene-disrupting variants (including SVs) can cause Mendelian disease. However, the provided text contains no mention of the gene CD38, no gene-level analyses or reported associations linking CD38 to any Mendelian phenotype, and the authors explicitly state that sample sizes are currently underpowered to estimate SV constraint at the level of individual genes. Because of the absence of any CD38-specific data (genetic, clinical, or statistical) and the stated limitation on per-gene inference, this paper does not provide direct evidence that CD38 is associated with at least one Mendelian disease. It only offers general, indirect mechanistic support that genes affected by SVs can be disease-causing, which is insufficient to establish the specific claim about CD38.


[Read Paper](https://www.semanticscholar.org/paper/d96c5f9215341f85056df52a914c403b5435de5b)


### A data set of variants derived from 1455 clinical and research exomes is efficient in variant prioritization for early‐onset monogenic disorders in Indians

**Why Not Relevant**: The provided paper content does not mention the gene CD38, nor does it present any gene-specific associations, tables, or named lists that would allow identification of CD38 as being linked to a Mendelian disease. The excerpts show that the study generated a large Indian exome variant resource, identified additional genes/human knockouts, and listed carrier frequencies for many recessive disorders, but none of these statements include gene names or phenotypic associations in the provided text. Therefore there is no direct evidence (no variant, genotype-phenotype link, segregation, statistical association, or case report) and no mechanistic evidence (no functional data, pathway analysis, or biological rationale) presented here that links CD38 to a Mendelian disease. It remains possible that CD38 could be among the "additional 19 genes" or included in the carrier-frequency listings in the full paper or supplements, but based solely on the supplied content, that cannot be determined. To evaluate the claim further would require the full paper (main text and supplementary tables) or external databases/literature that list gene-to-disease associations (e.g., OMIM, ClinVar, HGMD) to see whether CD38 is named and what evidence supports any Mendelian association.


[Read Paper](https://www.semanticscholar.org/paper/5580b42d884e2d4384f3dc66f68623ee1184b0f0)


### Whole-Exome Sequencing Reveals a Rare Missense Variant in SLC16A9 in a Pedigree with Early-Onset Gout

**Why Not Relevant**: The paper's entire genetic analysis and reported findings concern SLC16A9 and early-onset gout. There is no mention of the gene CD38, no data, no mechanistic discussion, and no experimental or cosegregation evidence involving CD38. Therefore this paper provides no direct evidence that CD38 is associated with any Mendelian disease. Absence of CD38 in this paper does not disprove the claim in general; it merely means this paper is irrelevant to evaluating CD38. To assess the claim about CD38 would require literature that specifically reports CD38 mutations, cosegregation with Mendelian phenotypes, functional/mechanistic studies implicating CD38 in monogenic disease, or curated disease-gene databases (e.g., OMIM, ClinVar) listing CD38-associated Mendelian disorders.


[Read Paper](https://www.semanticscholar.org/paper/69ca3cf7fa46cb144ca6c07c8b8add1acfa1382b)


### Predictors of the utility of clinical exome sequencing as a first-tier genetic test in patients with Mendelian phenotypes: results from a referral center study on 603 consecutive cases

**Why Not Relevant**: The provided paper content contains no mention of the gene CD38, no gene-level results, no variant counts for specific genes, and no mechanistic discussion that could link CD38 to any Mendelian phenotype. The single sentence supplied reports an aggregate diagnostic yield (40%) from clinical exome sequencing (CES) across a cohort with various Mendelian phenotypes and refers to predictors of CES utility, but it provides no information about which genes were implicated, which specific diagnoses were made, or any functional evidence for particular gene–disease associations. From that sentence alone one cannot (a) identify whether CD38 variants were observed in the cohort, (b) determine whether any observed CD38 variants were classified as pathogenic or likely pathogenic, or (c) evaluate segregation or functional data that would support a Mendelian association. Therefore this paper content does not provide direct evidence for or against the claim that "CD38 is likely associated with at least one Mendelian disease," nor does it provide mechanistic evidence to assess plausibility. To evaluate the claim would require gene-level results (counts of pathogenic/likely pathogenic CD38 variants), phenotypic descriptions of affected individuals, segregation or de novo status, functional/biochemical evidence for pathogenicity, or citations to prior studies linking CD38 to Mendelian disorders—none of which are present in the supplied text. Additionally, because only a single summary sentence was provided, methodological details (cohort composition, sequencing/variant-calling/interpretation pipelines, criteria for diagnostic classification) that could affect the possibility of detecting CD38 associations are absent, further preventing any inference about CD38 from this content.


[Read Paper](https://www.semanticscholar.org/paper/3e4076a587328718af716949d2e787a464220fa6)


### Case report: Exploring autosomal recessive woolly hair: genetic and scanning electron microscopic perspectives on a Japanese patient

**Why Not Relevant**: The paper focuses on the genetic causes and morphological features of autosomal recessive woolly hair, reporting pathogenic variants in LIPH (and noting LPAR6 and KRT25 as known causes). It does not mention the gene CD38, investigate CD38 variants, or address any phenotypes or Mendelian diseases linked to CD38. Therefore the paper provides no direct or mechanistic evidence for or against the claim that CD38 is associated with at least one Mendelian disease. Note that absence of any mention of CD38 in this study is not proof that CD38 is not associated with a Mendelian disease elsewhere; it simply means this paper is not relevant to the claim.


[Read Paper](https://www.semanticscholar.org/paper/8796d4a078e46956fc51bba8512fa7901da1182d)


### Case report: A novel FBXW7 gene variant causes global developmental delay

**Why Not Relevant**: The provided paper content exclusively describes a clinical case and genetic findings for FBXW7 (including objective, trio-WES results, and conclusions). There is no mention of the gene CD38, no functional/mechanistic discussion involving CD38, and no data that could be interpreted as evidence for or against an association between CD38 and any Mendelian disease. Because the paper does not address CD38 at all, it provides neither direct evidence (observational/genetic association) nor mechanistic evidence (pathway/function) relevant to the claim. Additionally, the study is a single-case report focused on FBXW7, so even indirect inference about other genes would be unjustified. To evaluate the claim about CD38, one should instead consult primary genetic-disease databases (e.g., OMIM, ClinVar, HGMD) and literature that specifically report pathogenic variants or Mendelian phenotypes linked to CD38.


[Read Paper](https://www.semanticscholar.org/paper/bb0b7ce4ae459771c428a6c9d1dbc93aecc6bcdc)


### A Rare Pathogenic Variant in the SERPINF1 Gene in Association with Early-Onset Severe Presentation of Autosomal Recessive Type of Osteogenesis Imperfecta VI: A Case Report

**Why Not Relevant**: The paper exclusively reports a case study of osteogenesis imperfecta type VI caused by a homozygous truncating variant in SERPINF1, with clinical descriptions and genetic confirmation in a single family. There is no mention of the gene CD38 anywhere in the provided text, no experimental data, no genetic association, and no mechanistic discussion related to CD38. Therefore the paper provides no direct evidence supporting or refuting the claim that CD38 is associated with at least one Mendelian disease. The absence of CD38 in this focused report cannot be interpreted as evidence against CD38's involvement in Mendelian disease generally; it simply indicates that this study did not examine CD38. Given the complete lack of content about CD38, the relevance of this paper to the claim is essentially zero.


[Read Paper](https://www.semanticscholar.org/paper/227249d6153b838015091ffee1a5bf887778cf71)


### Urofacial (Ochoa) syndrome with a founder pathogenic variant in the HPSE2 gene: a case report and mutation origin

**Why Not Relevant**: The provided paper content does not mention the gene CD38 anywhere and instead reports a pathogenic variant in HPSE2 associated with urofacial syndrome (UFS). Therefore it contains no direct or mechanistic evidence linking CD38 to any Mendelian disease. From a methodological and inference standpoint, one cannot infer an association of CD38 from a report that exclusively describes HPSE2: there is no reported genetic, biochemical, functional, or epidemiological connection between CD38 and the described findings. Absence of mention in this excerpt is not proof that CD38 is not involved in Mendelian disease, but for the purposes of assessing whether this paper provides supporting or refuting evidence for the claim, it provides none. To support the claim about CD38, one would need explicit evidence such as: identification of pathogenic CD38 variants segregating with disease in families, functional assays showing that CD38 variants cause relevant molecular defects, animal models recapitulating a Mendelian phenotype with CD38 disruption, or curated database/clinical reports (e.g., OMIM, ClinVar) documenting CD38 as causal for a Mendelian disorder. None of those types of evidence are present in the supplied text.


[Read Paper](https://www.semanticscholar.org/paper/aed48658f491795f6058cfadb58f6ceb0243fe11)


### Clinical utility of Whole Exome Sequencing for rare Mendelian disorders: phenotypic-driven strategy for a high diagnostic yield and identification of 48 novel variants

**Why Not Relevant**: The provided paper excerpt contains only high-level background, methods, cohort size, diagnostic yield, and aggregate counts of pathogenic variants, but it contains no mention of the gene CD38, no gene- or variant-level results, and no mechanistic discussion specific to CD38. Therefore it provides no direct evidence that CD38 is associated with any Mendelian disease, nor mechanistic data that would support or refute such an association. To evaluate the claim about CD38 specifically, the full paper's results, tables, or supplemental material listing identified genes/variants would need to be examined for any CD38 findings. Because the excerpt contains no CD38-related information, its relevance to the claim is minimal.


[Read Paper](https://www.semanticscholar.org/paper/5617e3ed3e23ea4eb7551539966ff2c7dce5de77)


### Whole-genome sequencing reveals contribution of rare and common variation to structural kidney and urinary tract malformations

**Why Not Relevant**: The provided paper excerpt contains no mention of the gene CD38 and provides no direct or mechanistic evidence linking CD38 to any Mendelian disease. The paper focuses on congenital anomalies of the kidneys and urinary tract (CAKUT), reporting overall diagnostic yield for monogenic causes, an association at a single common-variant marker (rs117473527), estimates of heritability attributed to common variation, and polygenic risk scores. Example verbatim sentences that illustrate the paper's focus (and absence of CD38) include: "Congenital anomalies of the kidneys and urinary tract (CAKUT) are the commonest cause of kidney failure in children and young adults with over 50 monogenic causes identified, largely in cohorts enriched for familial, syndromic, or consanguineous disease." and "Exome-wide rare variant and genome-wide common variant (minor allele frequency [≤] 0.1%) testing was performed in a subset of 813 patients and 25,205 ancestry-matched controls with significant association detected at rs117473527 (P=3.93x10-8; OR 3.17; 95% CI 2.10-4.78; MAF 0.02)." Neither these sentences nor any other sentence in the provided content names CD38 or describes functional or clinical evidence regarding CD38. Therefore, based solely on the supplied content, there is no support for the claim that "The gene CD38 is likely associated with at least one Mendelian disease." Caveat: absence of any mention of CD38 in this excerpt is not definitive proof that the full paper (including supplementary tables) contains no relevant information. If you can provide the full text or supplementary material, or allow a literature search, I can re-evaluate for any CD38-specific evidence.


[Read Paper](https://www.semanticscholar.org/paper/e961066c4ba5e47abd023a9e57f780a8a68c2c2c)


## Search Queries Used

- CD38 pathogenic variant ClinVar OMIM case report patient mutation

- CD38 Mendelian disease monogenic rare variant exome sequencing family segregation

- CD38 deficiency human loss of function NADase NAD glycohydrolase oxytocin autism social behavior neurodevelopmental disorder

- CD38 gene constraint gnomAD pLI ClinGen dosage sensitivity population genetics variant pathogenicity


## Bibliometric Configuration

- Use Bibliometrics: True

- Author Impact Weight: 0.4

- Citation Impact Weight: 0.4

- Venue Impact Weight: 0.2
