## Supplementary material for "MENDELSEEK: An algorithm that predicts Mendelian Genes and elucidates what makes them special": CDH8.txt

# Claim: The gene CDH8 is likely associated with at least one Mendelian disease.

**Status**: processed

**Overall Rating**: -1

**Explanation**:

No relevant papers were found that support or refute this claim.


## Relevant Papers


## Other Reviewed Papers


### Precision Medicine in Internal Medicine

**Why Not Relevant**: The provided paper content is a general review of genetic testing strategies, exome/genome sequencing, and variant interpretation. It does not mention the gene CDH8, report any CDH8 variants, provide clinical cases, segregation analyses, functional assays, statistical association tests, or mechanistic studies specific to CDH8. Therefore it contains no direct evidence (case reports, cohort studies, ClinVar/OMIM entries, or functional data) that CDH8 is associated with any Mendelian disease. The included excerpts offer only general contextual or mechanistic background about how gene-disease associations are discovered and the current state of gene annotation (e.g., that many genes are not yet linked to Mendelian disorders). Because none of this material is specific to CDH8, the document cannot be used to support or refute the claim that CDH8 is likely associated with at least one Mendelian disease. To evaluate the claim properly would require gene-specific sources such as OMIM/ClinVar entries for CDH8, published case reports or cohort studies showing pathogenic CDH8 variants segregating with disease, or functional evidence demonstrating a disease mechanism linked to CDH8.


[Read Paper](https://www.semanticscholar.org/paper/c7199e8a5226f8d9d55777da0e15e91797bb459a)


### Potential genetic causes of miscarriage in euploid pregnancies: a systematic review.

**Why Not Relevant**: The reviewed paper is a systematic review focused on monogenic causes of euploid pregnancy loss (≤20 weeks) and summarizes genes and mechanisms implicated in pregnancy loss. The paper lists specific genes (e.g., CHRNA1, DYNC2H1, RYR1) and discusses exome sequencing and CNV studies, but the content provided does not mention CDH8 anywhere. Therefore the paper provides no direct evidence that CDH8 is associated with a Mendelian disease. It also provides limited indirect evidence: because the review targeted genes causing early lethal phenotypes and excluded genes not lethal in utero, the absence of CDH8 might reflect phenotype selection (CDH8-associated disease, if any, might not present as early pregnancy lethality) or limitations of the literature search (date cutoff May 2018, database scope). Absence of mention in this review is not definitive evidence against CDH8 being a Mendelian disease gene, but it is also not supportive. Given the lack of any explicit data, variants, mechanism, or discussion about CDH8 in the paper content provided, the relevance of this paper to the claim that "CDH8 is likely associated with at least one Mendelian disease" is negligible.


[Read Paper](https://www.semanticscholar.org/paper/d03fb624d6bf2a0d59a8128a2d593f64934a07f4)


### Identifying genomic variant associated with long QT syndrome type 2 in an ecuadorian mestizo individual: a case report

**Why Not Relevant**: The paper does not mention CDH8 anywhere and provides no experimental, genetic, clinical, or mechanistic data pertaining to CDH8. It is a single-case report focused on Long QT syndrome type 2 and reports variants in KCNH2, AKAP9, and TTN only. Therefore it supplies neither direct evidence (e.g., pathogenic CDH8 variants segregating with disease) nor mechanistic evidence (e.g., functional studies implicating CDH8 in disease pathways) relevant to the claim that CDH8 is likely associated with at least one Mendelian disease. The absence of CDH8 findings in this narrow LQTS case report cannot be taken as evidence against CDH8 involvement in other Mendelian diseases; likewise, it offers no support for the claim. Given these points, the paper is not relevant to assessing the claim.


[Read Paper](https://www.semanticscholar.org/paper/aa65df486c4f1720d86feb42ddc72ed25ab57431)


### Novel, heterozygous, de novo pathogenic variant (c.4963delA: p.Thr1656Glnfs*42) of the NF1 gene in a Chinese family with neurofibromatosis type 1

**Why Not Relevant**: The provided paper content contains a single sentence: "This NF1 novel variant probably underlays the NF1 pathogenesis in this pedigree, which would help for the diagnosis, genetic counseling, and clinical management of this disorder." This sentence refers specifically to a novel variant in the NF1 gene and to NF1 pathogenesis. It does not mention the gene CDH8, does not present data or analysis concerning CDH8, and provides no mechanistic link between CDH8 and any Mendelian disease. While NF1 (neurofibromin 1) is itself a gene associated with a Mendelian disorder (neurofibromatosis type 1), that is unrelated to the claim about CDH8. Because there is no mention of CDH8, no genetic association, and no mechanistic discussion connecting CDH8 to disease in the provided text, this paper content provides no direct or mechanistic evidence for or against the claim that CDH8 is likely associated with at least one Mendelian disease. Additional or different source material that explicitly addresses CDH8 would be required to evaluate the claim.


[Read Paper](https://www.semanticscholar.org/paper/d060a6d0f5b9e4fcf8b566368457c6650fbc09f4)


### Prenatal ultrasound findings of X-linked congenital cataracts: case report and description of a novel variant.

**Why Not Relevant**: The paper's content focuses on prenatal diagnosis of congenital cataracts and reports a pathogenic variant in the OCRL gene associated with X-linked disorders (Dent disease 2, Lowe's syndrome). There is no mention of the gene CDH8 (no occurrences of the symbol/name 'CDH8' or any discussion of cadherin-8 or related loci) in the provided text. Therefore the paper contains neither direct evidence (e.g., variants, segregation, case reports, association statistics) nor mechanistic evidence (e.g., functional studies, pathway links) linking CDH8 to any Mendelian disease. Note that absence of CDH8 in this paper does not disprove an association in other literature, but this paper does not contribute supporting evidence toward the claim.


[Read Paper](https://www.semanticscholar.org/paper/058d8e6037d294c050f1e435bf38ad6e48b453db)


### New Pathogenic Homozygous Variant in (DGOUK) Gene Cause Vital Progressive Liver Failure in a Neonate: Case Report

**Why Not Relevant**: The provided content entirely concerns mitochondrial DNA depletion syndromes (MDS) and reports a homozygous variant in DGUOK (OMIM:601465) in an infant case. There is no mention of the gene CDH8, no data, no variant descriptions, no mechanistic discussion, and no clinical or genetic evidence relating CDH8 to any Mendelian disease. Therefore the paper content as provided is not relevant to the claim that "The gene CDH8 is likely associated with at least one Mendelian disease." Caveats: this assessment is limited to the supplied excerpt. If CDH8 appears elsewhere in the full paper (outside the provided text), that could change relevance; however, based on the content given, there is no supporting or mechanistic evidence about CDH8.


[Read Paper](https://www.semanticscholar.org/paper/74a649e26c83fe77b6b076668da3b563912accff)


### Case report: Preimplantation genetic testing for infantile GM1 gangliosidosis

**Why Not Relevant**: This paper focuses exclusively on GM1 gangliosidosis caused by mutations in the GLB1 gene, reports clinical and molecular findings for that gene, and describes preimplantation and prenatal testing for GLB1 variants. The text contains no mention of CDH8, no genetic data, segregation analysis, functional assays, or mechanistic discussion relating to CDH8. Therefore it provides neither direct evidence (case reports, variant associations, segregation) nor mechanistic evidence (biochemical pathway links, protein function perturbation) relevant to the claim that CDH8 is likely associated with a Mendelian disease. Absence of any reference to CDH8 in this paper means the paper is not informative with respect to the claim. Caveat: a single paper's lack of mention of CDH8 does not prove CDH8 is not disease-associated; it only means this paper does not supply supporting or refuting evidence. To assess the claim about CDH8, one should consult other sources (OMIM, ClinVar, gene-disease curation resources, and the primary literature) that specifically analyze CDH8 variants, segregation, and functional impact.


[Read Paper](https://www.semanticscholar.org/paper/ce981bc89c565bb4d97baff441baaf8695595448)


### Autosomal Dominant Intellectual Development Disorder-6 (MRD6) Without Seizures Linked to a De Novo Mutation in the grin2b Gene Revealed by Exome Sequencing: A Case Report of a Moroccan Child

**Why Not Relevant**: This paper does not provide evidence for or against the claim that the gene CDH8 is likely associated with at least one Mendelian disease. The entire provided content discusses GRIN2B (written as 'grin2b' in the text) and an autosomal dominant intellectual developmental disorder-6 (MRD6) caused by a GRIN2B nonsense variant (c.3912C>G, p.Tyr1304Ter). Specific points from the content that show the paper's focus include: the disorder name (MRD6), the gene discussed (GRIN2B/grin2b), the identified variant (c.3912C>G (p.Tyr1304Ter)), OMIM entry (613970), and the statement that the variant is considered pathogenic according to ACMG criteria. Nowhere in the provided text is CDH8 mentioned, nor are there mechanistic data, genetic segregation, case reports, population data, functional studies, or literature citations that link CDH8 to any Mendelian phenotype. Therefore: (1) there is no direct evidence in this paper supporting an association between CDH8 and a Mendelian disease; (2) there is no mechanistic evidence in this paper relevant to CDH8; and (3) the absence of any mention of CDH8 means the paper cannot be used to substantiate the claim. Note that absence of evidence in this paper is not evidence that CDH8 is not associated with a Mendelian disease; it only means this paper is irrelevant to the claim. To evaluate the claim about CDH8 one would need to search databases and literature sources specific to CDH8 (e.g., OMIM, ClinVar, PubMed, HGMD, gnomAD) or find case reports/functional studies directly addressing CDH8 variants and phenotypes.


[Read Paper](https://www.semanticscholar.org/paper/67d756e0cc302e5d46fdda9df34403f1f2de9a5a)


### A New De Novo Missense Variant of the TET3 Gene in a Patient with Epilepsy and Macrocephaly

**Why Not Relevant**: This paper provides no direct or mechanistic evidence relating to the gene CDH8. The supplied content exclusively discusses TET3 (Beck–Fahrner syndrome) and a de novo TET3 variant in a single patient with neurodevelopmental delay and epilepsy. No mention of CDH8 appears in the text, and there are no data, variant reports, segregation analyses, functional assays, or mechanistic discussions concerning CDH8. 

Relevant points from the provided content that demonstrate the focus on TET3 rather than CDH8 include statements such as: the association of pathogenic variants of the TET3 gene with Beck–Fahrner syndrome, the report that only 28 patients are reported with pathogenic variants of TET3, and the description of a de novo heterozygous TET3 variant (c.2867G>A p.(Arg956Gln)) identified by trio-based exome sequencing. These confirm the paper’s subject is TET3 and not CDH8. 

Because the claim specifically asks whether CDH8 is likely associated with at least one Mendelian disease, and the paper contains no mention or data on CDH8 (no variants, no phenotype correlation, no mechanistic experiments), the paper is not relevant. 

Limitations and notes: although this paper illustrates the types of evidence that can support a gene–disease association (e.g., de novo variants, absence from population databases, bioinformatic pathogenicity predictions, phenotypic description), those are generic methodological points only. They do not provide any empirical or mechanistic evidence for CDH8. To evaluate the claim about CDH8 one would need papers reporting CDH8 variants segregating with disease, recurrent pathogenic variants in multiple unrelated affected individuals, functional studies showing perturbation of CDH8 leads to disease-relevant biology, or animal models recapitulating phenotype. None of those are present in the provided content.


[Read Paper](https://www.semanticscholar.org/paper/46aff7c276c98934ac50dac8ca16dcaf6dacc5aa)


### Detection of chromosomal abnormalities, copy number variations (cnvs), and mutations causing

**Why Not Relevant**: The provided paper content is a general introductory overview of bone and mineral disorders, emphasizing that some are monogenic (Mendelian) while others are polygenic, and outlining clinical approaches to diagnosis and the use of molecular genetic tests. The text does not mention CDH8, any specific genes, variants, patient cases, segregation analyses, statistical results, functional assays, animal models, or mechanistic pathways involving CDH8. Therefore it contains no direct evidence (e.g., reported pathogenic CDH8 variants, disease co-segregation, case reports, association statistics) and no mechanistic evidence (e.g., biochemical function of CDH8 relevant to bone disease, pathway perturbation, or experimental perturbation demonstrating pathogenicity) to support or refute the claim that CDH8 is likely associated with at least one Mendelian disease. To evaluate the claim one would need gene-specific evidence such as: (1) curated entries in Mendelian disease resources (OMIM) linking CDH8 to a named disorder, (2) pathogenic or likely pathogenic CDH8 variants in ClinVar with supporting evidence, (3) peer-reviewed case reports or cohort studies showing segregation of CDH8 variants with disease, (4) functional experiments (cellular or animal) demonstrating that CDH8 loss or mutation reproduces relevant disease phenotypes, and/or (5) statistical evidence from rare-variant burden tests in appropriately powered cohorts. Because none of these elements are present in the supplied text, the relevance to the specific claim is effectively nil.


[Read Paper](https://www.semanticscholar.org/paper/8174cf54a2a6499063ac67c52b1427a8ec56b7b8)


## Search Queries Used

- CDH8 pathogenic variant Mendelian disease case report OMIM ClinVar

- CDH8 variant exome sequencing trio sequencing family segregation autosomal dominant autosomal recessive de novo

- CDH8 cadherin 8 loss of function knockout mouse model neuronal development cell adhesion synapse autism intellectual disability functional assay

- cadherin genes Mendelian disease review systematic review monogenic neurodevelopmental disorders copy number variation structural variant CDH8


## Bibliometric Configuration

- Use Bibliometrics: True

- Author Impact Weight: 0.4

- Citation Impact Weight: 0.4

- Venue Impact Weight: 0.2
