## Supplementary material for "MENDELSEEK: An algorithm that predicts Mendelian Genes and elucidates what makes them special": CEP76.txt

# Claim: The gene CEP76 is likely associated with at least one Mendelian disease.

**Status**: processed

**Overall Rating**: -1

**Explanation**:

No relevant papers were found that support or refute this claim.


## Relevant Papers


## Other Reviewed Papers


### Autosomal Dominant Intellectual Development Disorder-6 (MRD6) Without Seizures Linked to a De Novo Mutation in the grin2b Gene Revealed by Exome Sequencing: A Case Report of a Moroccan Child

**Why Not Relevant**: The paper content exclusively discusses a pathogenic GRIN2B (grin2b) stop-gain variant causing MRD6 and contains no mention of the gene CEP76 or of any data (genetic, functional, clinical) concerning CEP76. Therefore there is no direct evidence (case reports, segregation, variant data) linking CEP76 to any Mendelian disease in this text. Mechanistic evidence in the paper (nonsense mutation, de novo occurrence, ACMG classification) applies to GRIN2B and its role in neurodevelopment; those methods and lines of evidence demonstrate how one would establish a gene-disease association but do not constitute evidence for CEP76. Additionally, the paper does not present negative data about CEP76 (e.g., sequencing showing no CEP76 variants), so it neither supports nor formally refutes a CEP76-disease association. To evaluate the claim about CEP76, one would need papers or databases that specifically report CEP76 variants, segregation, functional studies, or curated gene-disease entries (e.g., OMIM, ClinVar, GeneReviews) for CEP76. Because the supplied content contains zero information about CEP76, its relevance to the claim is effectively nil.


[Read Paper](https://www.semanticscholar.org/paper/67d756e0cc302e5d46fdda9df34403f1f2de9a5a)


### Case report: A de novo Non-sense SOX9 mutation (p.Q417*) located in transactivation domain is Responsible for Campomelic Dysplasia

**Why Not Relevant**: This paper provides no evidence relevant to the claim that CEP76 is likely associated with at least one Mendelian disease. The provided content exclusively concerns Campomelic dysplasia (CD) and pathogenic variants of the SOX9 gene (including identification of a novel SOX9 nonsense variant c.1249C > T; p.Q417*). There is no mention of the gene CEP76, no experimental data involving CEP76, and no mechanistic discussion that links CEP76 to CD or any other Mendelian disorder. The study design (whole-exome sequencing of a fetus with CD, co-segregation analysis, bioinformatics prediction of SOX9 pathogenicity) and the conclusions all pertain to SOX9 haploinsufficiency and its transactivation activity; therefore, based on the provided text, this paper neither supports nor refutes any association between CEP76 and Mendelian disease. If additional sections of the full paper (not provided here) mention CEP76, those would need to be reviewed separately, but on the basis of the supplied content there is no relevant evidence.


[Read Paper](https://www.semanticscholar.org/paper/bcc43a4127c8fa8d83c22e39fd3a13cbfa12fe1a)


## Search Queries Used

- CEP76 mutation Mendelian disease case report exome sequencing family segregation de novo

- CEP76 ClinVar OMIM HGMD gnomAD constraint pLI loss intolerance pathogenic variant gene disease association

- CEP76 centrosomal protein 76 centriole duplication centrosome dysfunction ciliopathy microcephaly model mouse zebrafish cell line functional assay

- biallelic CEP76 loss of function variants autosomal recessive developmental disorder intellectual disability microcephaly whole exome sequencing cohort case series genotype phenotype correlation


## Bibliometric Configuration

- Use Bibliometrics: True

- Author Impact Weight: 0.4

- Citation Impact Weight: 0.4

- Venue Impact Weight: 0.2
