## Supplementary material for "MENDELSEEK: An algorithm that predicts Mendelian Genes and elucidates what makes them special": CEP89.txt

# Claim: The gene CEP89 is likely associated with at least one Mendelian disease.

**Status**: processed

**Overall Rating**: -1

**Explanation**:

No relevant papers were found that support or refute this claim.


## Relevant Papers


## Other Reviewed Papers


### Autosomal dominant familial dyskinesia and facial myokymia: single exome sequencing identifies a mutation in adenylyl cyclase 5.

**Why Not Relevant**: The paper exclusively investigates and implicates ADCY5 as the likely causal gene for familial dyskinesia with facial myokymia. There is no mention of the gene CEP89 anywhere in the provided content—no variant data, no co-segregation analysis, no expression or functional data relating to CEP89. Therefore this paper provides no direct evidence that CEP89 is associated with any Mendelian disease, nor does it provide mechanistic information that could be used to infer such an association for CEP89. Because the study's results and mechanistic discussion are specific to ADCY5, the content is not relevant to evaluating the claim about CEP89.


[Read Paper](https://www.semanticscholar.org/paper/efe6d6b532446cd5bbe0285d34d6deb9d4e4c726)


### The Burden of Candidate Pathogenic Variants for Kidney and Genitourinary Disorders Emerging From Exome Sequencing

**Why Not Relevant**: The provided paper content does not mention the gene CEP89 by name and contains no gene-level results or statements about CEP89. The excerpts describe methods for assembling a curated set of genes associated with Mendelian kidney and genitourinary disorders, cohort characteristics, and variant-filtering pipelines, but they do not present the curated gene list or any per-gene findings. Because there is no direct mention, summary, or data about CEP89 in the supplied text, the paper content cannot be used to support or refute the claim that CEP89 is likely associated with at least one Mendelian disease. It is possible CEP89 appears in the full gene list or in sections of the paper not included here, but based only on the provided content, there is insufficient information to evaluate the claim.


[Read Paper](https://www.semanticscholar.org/paper/a75e297208c04992568e607d19aa9b9aaebd09d5)


### Whole exome sequencing of intracranial aneurysm.

**Why Not Relevant**: The provided paper excerpt contains no mention of the gene CEP89, no data about variants in CEP89, and no genotype–phenotype segregation, functional, or mechanistic evidence linking CEP89 to any Mendelian disease. The excerpt is background and methods describing intracranial aneurysm (IA) familial recruitment and that whole exome sequencing was performed on selected families, but it does not include any results, gene lists, variant calls, or functional follow-up. Therefore there is no direct evidence (no reported CEP89 variants segregating with disease) and no mechanistic evidence (no discussion of CEP89 function, pathogenic mechanisms, or model systems) in the provided text to support or refute the claim. Limitations: the excerpt is truncated before results are reported, so it is possible the full paper (beyond the text provided) could contain relevant findings involving CEP89. To evaluate the claim properly, the sections reporting exome sequencing results, variant annotation, segregation analyses, and any functional assays or mechanistic discussion would be required. Without those, the excerpt cannot be used as evidence for CEP89 being associated with a Mendelian disease.


[Read Paper](https://www.semanticscholar.org/paper/3267b24524b6f935d4c9b5a0ba19b7d6846323e0)


### Rare Variants Analysis of Lysosomal Related Genes in Early-Onset and Familial Parkinson’s Disease in a Chinese Cohort

**Why Not Relevant**: The paper focuses specifically on lysosomal-related genes and Parkinson's disease in a Chinese cohort and reports findings for a limited set of genes (e.g., GBA, MAN2B1, SCARB2, MCOLN1, LYST, VPS16, VPS13C). The provided content (background, objective, methods, results, conclusion) contains no mention of the gene CEP89. Although whole-exome sequencing was performed, the authors report analyses restricted to 67 candidate lysosomal related genes; the paper does not present data implicating CEP89, nor does it describe mechanistic links involving CEP89. Therefore this paper provides no direct evidence that CEP89 is associated with any Mendelian disease. Important caveats: absence of CEP89 in the reported text does not prove CEP89 is not associated with a Mendelian disease — it only shows this particular study did not present such evidence. Additionally, because the methods indicate analysis focused on a predefined lysosomal gene list, even if WES data could in principle include CEP89, the reported analysis did not evaluate or report CEP89, so no inferences about CEP89 can be drawn from this paper. For evaluation of the claim about CEP89 one should consult studies/databases explicitly interrogating CEP89 (e.g., OMIM, ClinVar, locus-specific studies, or broader exome/genome-wide association analyses).


[Read Paper](https://www.semanticscholar.org/paper/1fd8997c5cf5d425181a4a955480e02f8ec3ed9c)


### PRIORITIZATION OF RESULTS FROM WHOLE EXOME SEQUENCING IN FAMILIAL INTRACRANIAL ANEURYSM

**Why Not Relevant**: The provided paper content describes a WES-based ranking system and reports counts of prioritized variants/genes in familial intracranial aneurysms, but it does not mention the gene CEP89 anywhere in the text supplied. There are no gene-specific results, lists, or statements tying CEP89 to any Mendelian disease. Therefore the paper provides no direct or indirect evidence that CEP89 is associated with at least one Mendelian disease. The study's methods (family WES, Mendelian-filtering) are relevant in principle to discovering Mendelian genes, but without CEP89 being named or described, the content cannot support or refute the claim.


[Read Paper](https://www.semanticscholar.org/paper/20b239904a9f7b1a9669e076a054804e126d963b)


### Identification by Exome Sequencing of Predisposing Variants in Familial Cases of Autoinflammatory Recurrent Fevers

**Why Not Relevant**: The paper does not mention CEP89 anywhere in the provided content. It focuses on exome sequencing of five familial autoinflammatory (AID) cases and reports candidate genes (FCRL6, PKN1, STAB1, PTDGR, VCAM1). Therefore the paper provides no direct evidence that CEP89 is associated with a Mendelian disease. The absence of CEP89 from the authors' candidate list is only weak, indirect negative evidence: the study is small (five families), phenotype-limited (periodic fever/autoinflammatory syndromes), and subject to specific variant filtering/prioritization pipelines, so CEP89 could still be disease-associated in other cohorts, phenotypes, or if analyzed with different methods. Because the manuscript neither reports variants in CEP89 nor discusses its function or mechanism, it cannot be used to support the claim that CEP89 is likely associated with a Mendelian disease; likewise it does not provide strong evidence against the claim. Consequently relevance to the claim is minimal.


[Read Paper](https://www.semanticscholar.org/paper/02404af0a77274d73bc9a97d762f58b6530e879c)


## Search Queries Used

- CEP89 centrosomal protein 89 C8orf41 biallelic loss of function missense splice site pathogenic variant exome sequencing case report Mendelian disease

- CEP89 ClinVar OMIM gnomAD gene constraint review systematic review meta analysis pathogenic annotation Mendelian disease

- CEP89 centrosomal protein 89 mitochondria oxidative phosphorylation respiratory chain cilia centriole functional study knockdown knockout mouse zebrafish cell line

- whole exome sequencing rare disease cohort CEP89 candidate gene autosomal recessive autosomal dominant segregation familial study undiagnosed Mendelian


## Bibliometric Configuration

- Use Bibliometrics: True

- Author Impact Weight: 0.4

- Citation Impact Weight: 0.4

- Venue Impact Weight: 0.2
