## Supplementary material for "MENDELSEEK: An algorithm that predicts Mendelian Genes and elucidates what makes them special": CHD6.txt

# Claim: The gene CHD6 is likely associated with at least one Mendelian disease.

**Status**: processed

**Overall Rating**: 0

**Explanation**:

Supporting evidence:
The provided paper describes the CHD (Chromodomain-Helicase-DNA binding) family as evolutionarily conserved proteins with roles in chromatin remodeling and DNA repair, and highlights that members of this family (e.g., CHD4) are essential for development (homozygous null embryos are lethal in mouse). These points establish that CHD-family proteins are biologically important and that disruption of family members can produce severe, often developmental, phenotypes — a general rationale for why mutation of a CHD-family gene could underlie a Mendelian disorder.

Caveats or contradictory evidence:
The excerpt does not mention CHD6 specifically at all; it focuses on the CHD family broadly and on CHD4 in particular. No experimental, genetic, clinical, or epidemiological data are provided that link CHD6 to any human Mendelian disease. The knockout/phenotype data cited refer to CHD4, not CHD6, so they cannot be taken as direct evidence for CHD6 disease association. Family-level functional importance does not by itself demonstrate that every paralog causes Mendelian disease when mutated; paralogs often differ in expression patterns, redundancy, and tolerance to variation.

Analysis of mechanisms and implications:
Mechanistically, CHD proteins (including CHD6 if it shares canonical domains) could plausibly cause disease when disrupted, because chromatin remodelers are critical for gene regulation during development and neurological function. Loss- or gain-of-function of such proteins has a well-established precedent producing Mendelian syndromes for other CHD paralogs (e.g., CHD7). However, whether CHD6 specifically behaves like those paralogs depends on its expression pattern, functional redundancy with other CHDs, tolerance to variation, and whether pathogenic variants occur in humans. None of those gene-specific attributes are provided in the excerpt.

Which rating is most appropriate:
Given the absence of any direct mention of CHD6 being associated with a Mendelian disease in the provided paper excerpt, and that the evidence is limited to general family-level plausibility and data on CHD4, the correct assessment based on the supplied evidence is that there is no evidence supporting the claim. The family-level arguments make the claim biologically plausible, but plausibility alone is not evidence of an actual Mendelian association for CHD6.


**Final Reasoning**:

I reviewed the provided excerpt and compared what it actually supports versus the claim. The paper establishes that CHD-family proteins play essential roles in chromatin remodeling and that disruption of at least one family member (CHD4) causes strong developmental phenotypes in mouse. This gives a biologically plausible basis to suspect other CHD genes might be disease-associated. However, the excerpt contains no data, case reports, genetic associations, or functional studies specific to CHD6. The claim is that CHD6 is likely associated with at least one Mendelian disease; the supplied evidence does not demonstrate any such association, nor does it cite human genetic findings for CHD6. Therefore, based strictly on the provided evidence, the appropriate rating is "No Evidence." If additional papers showing CHD6 pathogenic variants or clinical syndromes were supplied, the rating could be revised accordingly.


## Relevant Papers


### Spring 5-2005 Chromatin Remodeling in Mice

**Authors**: S. J. Dolgner (H-index: 6)

**Relevance**: 0.25

**Bibliometric Impact**: 0.048


**Excerpts**:

- The Chromodomain-HelicaseDNA binding domain gene family (CHD) is evolutionarily conserved and has distinct structural motifs that indicate a role in chromatin remodeling and DNA repair.

- The CHD proteins have both helicase activity, allowing the winding and unwinding of DNA, and an effect on histone acetylation through their role of the Nucleosome Remodeling and Histone Deacetylation (NuRD) complex.

- At the core of each complex is a helicase-like subunit that belongs to the SWI2/SNF2 family of proteins. Based on the relative homologies of these subunits, three major subfamilies have emerged: the SWI2/SNF2 family, the ISWI family, and the Mi-2/CHD family (4,5).

- In addition to the ATPase subunit CHD4, the NuRD complex also contains the histone deacetylases HDAC1 and HDAC2, the histone binding proteins RbAp46 and RbAp48, a methyl-CpG-binding domain-containing protein, MBD3, and a member of the metastasis-associated protein family, possibly leading to specialization of the complex (10,11).

- The largest component of the NuRD, CHD4, will be discussed in great detail in the next section, but is a multi-domain protein with two PHD Zn-finger domains, two chromodomains, the SWl2/SNF2-related ATPase domain, an additional helicase domain, and other domains whose functions are as yet unknown (12).

- We are using Chd4 knockout mice and cell lines to look further into the function of the protein. Preliminary data shows that homozygous null embryos are lethal, but the definite day of lethality has yet to be determined. In addition, cell line experiments show that cells heterozygous for Chd4 grow faster than wild type cells. Analysis of gene expression in mouse embryos shows gene expression in brain precursors, dermal precursors, and the dorsal aorta.

- Another large component of the NuRD is the histone deacetylases. These enzymes lead to the deacetylation of previously acetylated histones, causing the histone-bound DNA to become less accessible for transcription (13).


**Explanations**:

- This sentence identifies the CHD gene family as evolutionarily conserved and linked to chromatin remodeling and DNA repair. Relation to claim: mechanistic plausibility. Strength: establishes the family-level function that could make members disease-relevant. Limitation: it mentions the CHD family generally, not CHD6 specifically; conservation and structural motifs do not by themselves demonstrate Mendelian disease association.

- This sentence describes core biochemical activities of CHD proteins (helicase activity and influence on histone acetylation via NuRD). Relation to claim: mechanistic evidence that CHD proteins perform fundamental nuclear functions whose disruption could plausibly produce Mendelian phenotypes. Strength: links CHD proteins to essential chromatin processes. Limitation: still family-level and mechanistic, not direct evidence that CHD6 variants cause Mendelian disease.

- These sentences classify CHD (Mi-2/CHD) as a major subfamily of ATP-dependent chromatin remodelers. Relation to claim: mechanistic/evolutionary context indicating CHD proteins are part of essential remodeling machinery. Strength: situates CHD proteins among other key remodelers, supporting plausibility that disruption could have strong phenotypic consequences. Limitation: descriptive taxonomy; no CHD6 mention and no disease data.

- This sentence lists NuRD complex components and explicitly names CHD4 as the ATPase subunit within NuRD. Relation to claim: mechanistic/contextual evidence that certain CHD family members (CHD4) are integral to multi-protein complexes tied to transcriptional regulation. Strength: shows that CHD proteins function within complexes with other essential regulators, increasing the chance that their disruption is deleterious. Limitation: mentions CHD4 specifically, not CHD6; presence in complex is not direct evidence of CHD6's involvement in Mendelian disease.

- This sentence outlines domain architecture of CHD4 and emphasizes unknown functions of some domains. Relation to claim: mechanistic—demonstrates that CHD proteins (here CHD4) are multi-domain and plausibly essential. Strength: detailed structural context for why loss-of-function could have severe effects. Limitation: again CHD4-centric; CHD6 may differ in domain architecture and function, so inference is indirect.

- These sentences report preliminary functional data from Chd4 knockouts: homozygous null embryos are lethal; heterozygous cells grow faster; expression in specific embryonic tissues. Relation to claim: mechanistic/phenotypic evidence that perturbation of a CHD family member in vivo causes major developmental consequences, supporting the general plausibility that other CHD genes (potentially CHD6) could be Mendelian disease genes. Strength: in vivo evidence of essentiality for a CHD protein (Chd4) in mice. Limitations: (1) data are about Chd4, not CHD6; (2) described as preliminary without detailed methodology or statistical measures; (3) species differences (mouse vs human) and differences between paralogs reduce direct applicability to CHD6 Mendelian disease association.

- This sentence explains how histone deacetylases in NuRD repress transcription via deacetylation. Relation to claim: mechanistic—shows pathway (NuRD-mediated deacetylation) by which CHD family members influence gene expression and thus organismal phenotype. Strength: clarifies functional consequences of NuRD/CHD activity, supporting plausibility that disruption could produce disease. Limitation: general mechanistic statement about HDAC function; does not mention CHD6 or provide disease association evidence.


[Read Paper](https://www.semanticscholar.org/paper/dda8b6698e047480a4e73aec82cae41f60f9cdc3)


## Other Reviewed Papers


### DOES A DIAGNOSIS OF ASD HELP US TO HELP A PERSON WITH INTELLECTUAL DISABILITIES ?

**Why Not Relevant**: This paper content does not mention CHD6 and therefore provides no direct or mechanistic evidence bearing on the claim that CHD6 is associated with a Mendelian disease. The included material discusses caregiver well-being and ASD screening instruments, and a separate abstract focused on KCND1 as a candidate gene. Representative verbatim sentences from the paper illustrate the gene focus is KCND1 and related family members, not CHD6:

"We could identify and confirm by Sanger sequencing a novel missense mutation of the KCND1 gene in both affected half-brothers." 

"KCND1 is one of three members of the KCND/Kv4 family of voltagegated potassium channels." 

"Considering the identification of a sequence variant in KCND1 in two half brothers with moderate ID and autism spectrum disorder and the correlation of KCND2 and KCND3 with ID and autism we speculate that KCND1 might be a new candidate gene for intellectual disability." 

Other parts of the document concern caregiving for adults with autism and screening tools (DiBAS-R and ACL) and do not contain genetic evidence relevant to CHD6. Because CHD6 is not mentioned, there is no experimental, clinical, segregation, population-frequency, functional, or mechanistic data in this paper that would support or refute an association between CHD6 and any Mendelian disease.


[Read Paper](https://www.semanticscholar.org/paper/6d51deaa491eed78a7b02217a5218e42aa1bba39)


### Chromatin Remodeling Enzyme CHD7 Negatively Regulate Hematopoietic Stem Cell Function

**Why Not Relevant**: This paper exclusively studies CHD7 (Chromodomain Helicase DNA binding protein 7). The key disease-related sentence states that "Autosomal dominant CHD7 mutations are associated with CHARGE syndrome," which is direct evidence that CHD7 is a Mendelian disease gene. However, the claim to be evaluated is about CHD6. The manuscript content provided contains no mention of CHD6, no experiments on CHD6, and no data that directly implicate CHD6 in any phenotype or human disease. While CHD7 data illustrate that a member of the CHD family can cause Mendelian disease, similarity within a gene family (shared domain architecture or functions) is insufficient to infer that another paralog (CHD6) is likely a Mendelian disease gene without specific genetic, clinical, or functional evidence for CHD6. Therefore the paper is not relevant to the claim that CHD6 is likely associated with at least one Mendelian disease, aside from a very weak, indirect inference based on family-level biology. For these reasons the document provides negligible direct or mechanistic evidence for CHD6, and relevance is set very low.


[Read Paper](https://www.semanticscholar.org/paper/3363937a34556ffd276623ffa10187d9b7acaba8)


### Case report: A de novo Non-sense SOX9 mutation (p.Q417*) located in transactivation domain is Responsible for Campomelic Dysplasia

**Why Not Relevant**: The paper exclusively studies Campomelic dysplasia and reports a novel pathogenic SOX9 nonsense variant (p.Q417*) as the causal lesion in the affected fetus. There is no mention of the gene CHD6 anywhere in the provided text, nor any data, analysis, or discussion concerning CHD6. Therefore this paper provides no direct evidence for or against the claim that CHD6 is associated with a Mendelian disease. The study's narrow focus (single-family case, emphasis on SOX9) and lack of reporting about other genes means its contents are irrelevant to assessing CHD6. Note: absence of CHD6 discussion in this paper is not evidence that CHD6 has no Mendelian associations—only that this paper does not address that question.


[Read Paper](https://www.semanticscholar.org/paper/bcc43a4127c8fa8d83c22e39fd3a13cbfa12fe1a)


### Autosomal Dominant Intellectual Development Disorder-6 (MRD6) Without Seizures Linked to a De Novo Mutation in the grin2b Gene Revealed by Exome Sequencing: A Case Report of a Moroccan Child

**Why Not Relevant**: The paper content exclusively discusses GRIN2B (referred to as 'grin2b') and its association with MRD6 (OMIM 613970), including a specific pathogenic variant identified by exome sequencing. There is no mention of CHD6, CHD6 variants, CHD6 biology, or any data linking CHD6 to Mendelian disease. Therefore the paper provides no direct or mechanistic evidence relevant to the claim that CHD6 is likely associated with at least one Mendelian disease. The strong evidence in this paper supports GRIN2B as Mendelian-disease-associated, but gene-specific findings cannot be extrapolated to CHD6 without independent genetic, functional, or epidemiological data concerning CHD6.


[Read Paper](https://www.semanticscholar.org/paper/67d756e0cc302e5d46fdda9df34403f1f2de9a5a)


### Abstracts of papers presented at the 24th Genetics Society ’ s Mammalian Genetics and Development Workshop held at the Institute of Child Health, University College London on 21st November 2013

**Why Not Relevant**: The provided paper content does not mention CHD6 anywhere. The only CHD-family gene discussed is CHD7, with evidence that CHD7 haploinsufficiency causes CHARGE syndrome and that Chd7 ablation yields cardiac/vascular defects in mice. While this shows that a CHD-family chromatin remodeller can be a Mendelian disease gene (an indirect, mechanistic analogy), there is no direct genetic, clinical, functional, or mechanistic data specific to CHD6 in the text. Therefore the content does not provide direct support or refutation for the claim about CHD6. Inferring CHD6 involvement in Mendelian disease from CHD7 alone would be speculative without sequence/functional similarity data, variant evidence, patient phenotypes, or experimental perturbation of CHD6.


[Read Paper](https://www.semanticscholar.org/paper/b9217a59c8a6c141a43120d206471e3c06d88029)


### Chromatin Remodeling in Mice

**Why Not Relevant**: The provided paper content does not mention CHD6 anywhere and contains no genetic, clinical, or human pedigree data linking CHD6 to any Mendelian disorder. The manuscript focuses on CHD family biology and primarily on CHD4 (mouse knockouts and biochemical composition of the NuRD complex). While family-level and CHD4-specific mechanistic and experimental findings increase the plausibility that other CHD paralogs (including CHD6) could be disease-relevant, such inference is indirect and speculative. Concluding that CHD6 is likely associated with at least one Mendelian disease requires direct evidence (e.g., human pathogenic variants in CHD6 segregating with disease, case reports, population genetics indicating constrained variation, or functional studies of CHD6 variants), none of which are present in the text provided. Therefore the document provides mechanistic plausibility but no direct evidence linking CHD6 to Mendelian disease.


[Read Paper](https://www.semanticscholar.org/paper/88c919cd8208fa94066cb77fd32ca8fb7304cea9)


## Search Queries Used

- CHD6 mutation Mendelian disease case report family segregation exome sequencing de novo pathogenic variant

- CHD6 ClinVar OMIM gnomAD pLI loss of function constraint ClinGen haploinsufficiency

- Chromodomain helicase DNA binding protein 6 CHD6 chromatin remodeling ATP dependent helicase DNA repair transcriptional regulation functional study knockout mouse zebrafish developmental phenotype

- rare disease exome sequencing developmental disorder cohort candidate gene discovery CHD6 rare variant burden association intellectual disability growth delay craniofacial anomalies


## Usage Statistics

- Prompt Tokens: 0

- Completion Tokens: 0

- Total Tokens: 0

- Estimated Cost: $0.0415


## Bibliometric Configuration

- Use Bibliometrics: True

- Author Impact Weight: 0.4

- Citation Impact Weight: 0.4

- Venue Impact Weight: 0.2
