## Supplementary material for "MENDELSEEK: An algorithm that predicts Mendelian Genes and elucidates what makes them special": CHMP5.txt

# Claim: The gene CHMP5 is likely associated with at least one Mendelian disease.

**Status**: processed

**Overall Rating**: -1

**Explanation**:

No relevant papers were found that support or refute this claim.


## Relevant Papers


## Other Reviewed Papers


### Precision Medicine in Internal Medicine

**Why Not Relevant**: The paper content is a general review of genetic testing methods, variant interpretation, and the state of gene discovery; it does not mention CHMP5 by name, report observations of CHMP5 variants in affected individuals, provide segregation, functional, or case–control data, or describe mechanistic roles specific to CHMP5. Therefore the content offers only indirect, high-level support for the plausibility that an uncharacterized gene (such as CHMP5) might be discovered to cause a Mendelian disease in the future, but provides no direct evidence for or against the claim that CHMP5 is likely associated with at least one Mendelian disease. Given the absence of CHMP5-specific data, the paper is of very low relevance to the claim.


[Read Paper](https://www.semanticscholar.org/paper/c7199e8a5226f8d9d55777da0e15e91797bb459a)


### The Burden of Candidate Pathogenic Variants for Kidney and Genitourinary Disorders Emerging From Exome Sequencing

**Why Not Relevant**: Relevance is near zero because the provided paper content does not mention CHMP5 by name, does not present gene-level results or a public listing of the 625 curated genes within the excerpt, and contains no variant-level findings, mechanistic data, or prior reports specifically about CHMP5. The text describes the methods used to assemble and analyze a gene set for Mendelian kidney and genitourinary disorders and the cohort and variant-calling pipelines, but without the actual gene list or results one cannot determine whether CHMP5 is among the genes studied or whether any pathogenic variants in CHMP5 were observed. Therefore this paper excerpt provides no direct evidence (no reported association, no identified pathogenic variants) and no mechanistic evidence (no functional or biological information about CHMP5) to support or refute the claim that CHMP5 is likely associated with at least one Mendelian disease.


[Read Paper](https://www.semanticscholar.org/paper/a75e297208c04992568e607d19aa9b9aaebd09d5)


### POTENTIAL ROLE OF CERAMIDE METABOLISM IN LEWY BODY DISEASE

**Why Not Relevant**: The provided paper text contains no mention of the gene CHMP5 and provides neither direct genetic evidence (e.g. mutation descriptions, case reports, linkage, association, or Mendelian segregation) nor any mechanistic discussion specifically about CHMP5. The paper is a review focused on genes involved in ceramide metabolism and Lewy body diseases (e.g. GBA, PANK2, PLA2G6, NPC1, SPTLC1, ATP13A2), but none of the quoted passages refer to CHMP5 or to ESCRT/CHMP family function. Therefore this paper does not provide direct or specific mechanistic evidence that CHMP5 is associated with any Mendelian disease. While the review discusses lysosomal and ceramide-related pathways (which are biological processes that, in principle, could overlap with functions of endosomal sorting proteins such as CHMP5), such a generic overlap is speculative and is not supported by any text in the paper. Consequently the paper is not relevant to evaluating the claim that CHMP5 is likely associated with at least one Mendelian disease.


[Read Paper](https://www.semanticscholar.org/paper/a4b993dd7d1c93be882577eaa01aedc644f9aa45)


### Success and Pitfalls of Genetic Testing in Undiagnosed Diseases: Whole Exome Sequencing and Beyond

**Why Not Relevant**: The provided paper content does not mention CHMP5 in any capacity. It focuses on trio WES in a neurodevelopmental disorder cohort and describes findings for MEIS2, NIPBL, and unspecified genes of the chromatin-remodeling complex, with use of methylation signatures to evaluate variant pathogenicity. Because there are no data, case reports, variant descriptions, segregation analyses, functional assays, or mechanistic discussion involving CHMP5, this paper cannot support or refute the claim that CHMP5 is likely associated with a Mendelian disease. The paper's negative informational value is due to lack of coverage rather than evidence against CHMP5; to evaluate the claim would require literature or data that specifically report CHMP5 variants in affected individuals, segregation/penetrance data, functional studies showing pathogenic mechanisms, or population/constraint metrics implicating CHMP5. Additionally, this study's NDD-specific cohort means even a CHMP5 association to other Mendelian diseases could be missed. Therefore relevance is effectively zero for the specific claim.


[Read Paper](https://www.semanticscholar.org/paper/9c76f2886b2c4e59e399c5b986c6cf0018a2cdce)


### Novel CAD gene mutations in a boy with developmental and epileptic encephalopathy 50 with dramatic response to uridine therapy: a case report and a review of the literature

**Why Not Relevant**: The provided paper content refers exclusively to DEE50 (developmental and epileptic encephalopathy 50) and the CAD gene: "DEE 50 has early onset, refractory seizures, even status epilepticus leading to death, with favorable response to treatment with oral uridine, and the spectrum of CAD gene mutations is expanded." There is no mention of the gene CHMP5, no experimental data, case reports, genetic analyses, mechanistic discussion, or clinical associations involving CHMP5, and no evidence tying CHMP5 to Mendelian disease in this text. The described mechanism and therapeutic response (uridine) are specific to CAD (a gene involved in pyrimidine biosynthesis) and are mechanistically unrelated to CHMP5 (which encodes an ESCRT-III–associated protein involved in endosomal sorting and membrane remodeling). Because the excerpt does not mention CHMP5 and provides no direct or plausible mechanistic link to it, it cannot support or refute the claim that "CHMP5 is likely associated with at least one Mendelian disease." Additionally, the provided content is a single summary sentence without methods, results, or broader context, so even if CHMP5 had been mentioned, the snippet would be insufficient to establish a genetic-disease association. For evidence relevant to CHMP5 and Mendelian disease, literature specifically reporting CHMP5 variants in affected individuals, segregation, functional studies, or curated disease-gene resources (e.g., OMIM, ClinVar) should be consulted.


[Read Paper](https://www.semanticscholar.org/paper/b71aa97844d7f200b34377932dc0f0684e402757)


### How Many Genes Does It Take?

**Why Not Relevant**: The provided paper excerpt does not mention the gene CHMP5, any variants in CHMP5, or any phenotypes linked to CHMP5. The content discusses general concepts of autism spectrum disorder (ASD) genetics, consanguinity, approaches such as whole-exome sequencing (WES) and homozygosity mapping, and challenges in pinpointing causal loci in ASD families. None of these sentences provides direct evidence (e.g., reported CHMP5 pathogenic variants, segregation with disease, case reports) or mechanistic evidence (e.g., functional studies, pathways involving CHMP5) that would support or refute the claim that CHMP5 is likely associated with at least one Mendelian disease. 

Limitations and caveats: (1) The assessment is limited to the text you provided. It is possible that CHMP5 is discussed elsewhere in the full paper or in its supplementary materials; that information is not present in the excerpt and therefore cannot be evaluated here. (2) The excerpt does describe methods (WES, homozygosity mapping) that are appropriate for identifying Mendelian disease genes in consanguineous families, which is indirect and non-specific methodological context but does not constitute evidence about CHMP5 itself. 

Recommended next steps to evaluate the claim: search the full text and supplementary files of this paper for 'CHMP5'; query gene-disease databases and literature (OMIM, ClinVar, HGMD, PubMed) for CHMP5 pathogenic/likely pathogenic variants or reported Mendelian phenotypes; check population and constraint metrics (gnomAD, pLI/LOEUF) and any functional studies linking CHMP5 to disease mechanisms.


[Read Paper](https://www.semanticscholar.org/paper/324ffb74aa62f505b23c196e77da64af410d0121)


### Case report: Novel variants cause developmental and epileptic encephalopathy in three unrelated families from Mali

**Why Not Relevant**: This paper does not provide direct or mechanistic evidence regarding the gene CHMP5. The study reports whole exome sequencing results from three Malian families with developmental and epileptic encephalopathies and identifies variants in GRIN1, SYNJ1, and RARS2; CHMP5 is not mentioned. Absence of CHMP5 in this small, disease-specific cohort is not meaningful evidence for or against CHMP5's association with any Mendelian disease because: (1) the sample size is very small (three families, five patients), (2) the phenotypic focus is limited to DEEs and therefore would only detect CHMP5 involvement if it caused a DEE phenotype in these families, (3) the paper does not present systematic negative screening or population-level burden analyses for CHMP5, and (4) absence of evidence in one small study does not refute associations found elsewhere. Therefore the paper is not relevant to evaluating the claim that CHMP5 is likely associated with at least one Mendelian disease.


[Read Paper](https://www.semanticscholar.org/paper/a1ff4c7ea62aa361dd7f5bcff42679847e065e0e)


### Case Report: A novel de novo variant of COL1A1 in fetal genetic osteogenesis imperfecta

**Why Not Relevant**: The paper exclusively investigates and reports a novel de novo variant in COL1A1 as the likely cause of fetal osteogenesis imperfecta. There is no mention of CHMP5 anywhere in the provided content (no variants, no analyses, no discussion). Because the study's data, results, mechanistic interpretations, and conclusions all concern COL1A1 and collagen-related pathology, they do not provide either direct or mechanistic evidence that CHMP5 is associated with any Mendelian disease. While trio whole-exome sequencing was performed (which would likely detect coding variants in CHMP5 if present and prioritized), the authors do not report negative findings for CHMP5, nor do they present any data bearing on CHMP5. Therefore this paper is not relevant to the claim that CHMP5 is likely associated with at least one Mendelian disease.


[Read Paper](https://www.semanticscholar.org/paper/1caeaf6717a5af526be5f1f0dcef41ef4c377699)


### Clinical utility of Whole Exome Sequencing for rare Mendelian disorders: phenotypic-driven strategy for a high diagnostic yield and identification of 48 novel variants

**Why Not Relevant**: The paper content provided is a general summary of a clinical WES diagnostic study (162 cases, 52.5% diagnostic yield, 94 pathogenic variants characterized, 48 novel). Nowhere in the provided text is the gene CHMP5 mentioned, nor are any gene-specific results, variant identifiers, case descriptions, segregation analyses, or functional/mechanistic studies reported. Because the claim concerns a specific gene–disease association (CHMP5 → at least one Mendelian disease), evidence required would be: identification of pathogenic CHMP5 variants in affected individuals with segregation, recurrence in unrelated families, functional validation showing pathogenic mechanism, or prior literature linking CHMP5 to disease. None of those elements are present in the provided excerpt. Although the study reports novel pathogenic variants, without a gene-level breakdown or supplementary data listing those variants, one cannot attribute any of them to CHMP5. Therefore the provided paper content neither supports nor refutes the claim about CHMP5; it is non-informative with respect to that gene. To evaluate the claim, one would need the paper's full results/variant tables or other literature that explicitly report CHMP5 variants linked to disease.


[Read Paper](https://www.semanticscholar.org/paper/5617e3ed3e23ea4eb7551539966ff2c7dce5de77)


## Search Queries Used

- CHMP5 mutation Mendelian disease monogenic inherited disorder biallelic homozygous compound heterozygous de novo autosomal recessive autosomal dominant pathogenic variant

- CHMP5 whole exome sequencing trio undiagnosed rare disease clinical sequencing study patient cohort case report novel variant CHMP5

- CHMP5 knockout mouse phenotype ESCRT III charged multivesicular body protein 5 loss of function endosomal sorting autophagy neurodevelopment embryonic lethality model organism

- CHMP5 ClinVar OMIM ClinGen gnomAD gene disease association pathogenic likely pathogenic variant gene constraint pLI review genetic database


## Bibliometric Configuration

- Use Bibliometrics: True

- Author Impact Weight: 0.4

- Citation Impact Weight: 0.4

- Venue Impact Weight: 0.2
