## Supplementary material for "MENDELSEEK: An algorithm that predicts Mendelian Genes and elucidates what makes them special": CHRFAM7A.txt

# Claim: The gene CHRFAM7A is likely associated with at least one Mendelian disease.

**Status**: processed

**Overall Rating**: -1

**Explanation**:

No relevant papers were found that support or refute this claim.


## Relevant Papers


## Other Reviewed Papers


### High rate of disease-related copy number variations in childhood onset schizophrenia

**Why Not Relevant**: The provided paper content does not mention CHRFAM7A nor provide any data or specific discussion linking a particular gene to a Mendelian disease. The only sentence supplied is: "Clinically, broad CNV-based population screening is needed to assess their overall clinical burden and well-known pleiotropic effects of these CNVs suggesting shared abnormalities early in brain development are supported." This is a general statement about copy-number variants (CNVs) and their pleiotropic effects and does not (a) name CHRFAM7A, (b) present association evidence between CHRFAM7A and any Mendelian disorder, or (c) describe a mechanism implicating CHRFAM7A in a Mendelian disease pathway. Limitations: the excerpt is a single, high-level recommendation without methods, results, statistical data, or gene-level analysis; thus it cannot serve as direct or mechanistic evidence for the claim that CHRFAM7A is associated with at least one Mendelian disease. Because no relevant sentences or paragraphs about CHRFAM7A or Mendelian disease are present, no verbatim excerpts supporting or refuting the claim can be extracted.


[Read Paper](https://www.semanticscholar.org/paper/ae3f8f245ca6fc9edb069492dd091cf3878817ca)


### A pathogenic variant in RAB32 causes autosomal dominant Parkinson’s disease and activates LRRK2 kinase

**Why Not Relevant**: Relevance to the claim is effectively zero because the paper exclusively studies RAB GTPases, specifically identifying RAB32 c.213C>G (Ser71Arg) as co-segregating with autosomal dominant Parkinson disease and providing mechanistic data linking RAB32 to LRRK2 kinase activation. The paper does not mention CHRFAM7A, does not present any genetic, clinical, or mechanistic data about CHRFAM7A, and the methods indicate the gene set examined was RAB GTPases (61 genes). CHRFAM7A is a distinct gene (a human-specific partially duplicated fusion of CHRNA7) and would not be captured by analyses restricted to RAB family members unless specifically included; no such inclusion or incidental findings are reported in the provided text. Therefore this paper provides no direct or mechanistic evidence that CHRFAM7A is associated with any Mendelian disease.


[Read Paper](https://www.semanticscholar.org/paper/a36d28e828ede7c87c650af5f5117982b9749e7b)


### GM2 Gangliosidosis: Whole-Exome Sequencing Reveals Novel Homozygous Pathogenic Variant within the HEXA Gene in Iranian Family

**Why Not Relevant**: The provided content does not mention CHRFAM7A and therefore provides no direct or mechanistic evidence bearing on the claim. The single supplied sentence states: "The findings of this study offer additional support for the genetic heterogeneity of GM2 gangliosidosis and broaden the mutational gene spectrum of this inherited metabolic disorder by finding a novel HEXA pathogenic variant that had not been reported in these patients." This sentence concerns a novel pathogenic variant in HEXA and the genetic heterogeneity of GM2 gangliosidosis; it neither names nor describes CHRFAM7A, nor provides mechanistic pathways that could link HEXA to CHRFAM7A. Thus, from the supplied text there is neither direct evidence (e.g., identification of CHRFAM7A variants in Mendelian disease) nor mechanistic evidence (e.g., functional data, pathways, or interactions implicating CHRFAM7A) to support the claim.

Limitations: only a single sentence of the paper was provided. It is possible (though not shown here) that other parts of the full paper might mention CHRFAM7A; based solely on the supplied content, no inference can be made that CHRFAM7A is associated with any Mendelian disease. To properly evaluate the claim against this paper, a search of the full text for "CHRFAM7A" (and synonyms) or provision of additional excerpts would be necessary.


[Read Paper](https://www.semanticscholar.org/paper/6445f6e9e902634c0054e72cb027560096e8bd9e)


### Autosomal Dominant Intellectual Development Disorder-6 (MRD6) Without Seizures Linked to a De Novo Mutation in the grin2b Gene Revealed by Exome Sequencing: A Case Report of a Moroccan Child

**Why Not Relevant**: The paper exclusively reports a pathogenic, de novo nonsense variant in the GRIN2B (grin2b) gene causing autosomal dominant intellectual developmental disorder-6 (MRD6). There is no mention of the gene CHRFAM7A, no experimental or clinical data concerning CHRFAM7A, and no mechanistic discussion connecting GRIN2B findings to CHRFAM7A. Therefore this paper provides no direct evidence supporting or refuting the claim that CHRFAM7A is associated with any Mendelian disease. Limitation: absence of CHRFAM7A data in this paper does not prove CHRFAM7A is not disease-associated elsewhere; it only means this paper cannot be used as evidence for the claim.


[Read Paper](https://www.semanticscholar.org/paper/67d756e0cc302e5d46fdda9df34403f1f2de9a5a)


## Search Queries Used

- CHRFAM7A pathogenic variant Mendelian disease case report family segregation exome sequencing single gene disorder

- 15 q13 3 deletion CHRNA7 CHRFAM7A copy number variation CNV intellectual disability epilepsy neurodevelopmental disorder familial

- CHRFAM7A alpha 7 nicotinic receptor dominant negative function electrophysiology receptor assembly functional assay cell model induced pluripotent stem cell iPSC

- CHRFAM7A review ClinVar OMIM pathogenic variant population frequency loss of function pseudogene gene conversion human specific chimera monogenic


## Bibliometric Configuration

- Use Bibliometrics: True

- Author Impact Weight: 0.4

- Citation Impact Weight: 0.4

- Venue Impact Weight: 0.2
