## Supplementary material for "MENDELSEEK: An algorithm that predicts Mendelian Genes and elucidates what makes them special": CHTF8.txt

# Claim: The gene CHTF8 is likely associated with at least one Mendelian disease.

**Status**: processed

**Overall Rating**: -1

**Explanation**:

No relevant papers were found that support or refute this claim.


## Relevant Papers


## Other Reviewed Papers


### Genetic and phenotypic analysis of 101 patients with developmental delay or intellectual disability using whole‐exome sequencing

**Why Not Relevant**: The provided paper content does not mention the gene CHTF8 or any specific gene names. The text is a summary of whole-exome sequencing results in 101 Japanese patients with unexplained developmental delay/intellectual disability and reports aggregate findings (diagnostic yield, number of pathogenic variants, CNVs, and phenotype associations such as autism spectrum disorder and external ear anomalies). There are no verbatim sentences or paragraphs that name CHTF8, describe variants in CHTF8, or provide mechanistic information linking CHTF8 to a Mendelian disease. Therefore the paper content as provided contains no direct evidence (no reported pathogenic variants in CHTF8) and no mechanistic evidence (no functional, pathway, expression, or model organism data) to support or refute the claim that CHTF8 is likely associated with at least one Mendelian disease. Limitations relevant to this assessment: only an abstract-like excerpt was provided (no full methods, results, tables, or supplement), so it is possible CHTF8 could be mentioned elsewhere in the full paper; however, based solely on the supplied content there is no basis to evaluate the claim.


[Read Paper](https://www.semanticscholar.org/paper/c4a6b545539bc97495aa04ca6401c2b071178038)


### Generation and characterization of a novel gne Knockout Model in Zebrafish

**Why Not Relevant**: The provided paper excerpt exclusively studies the GNE gene (gne in zebrafish) and GNE Myopathy. There is no mention of the gene CHTF8, no named differentially expressed genes listed that could include CHTF8, nor any analysis, mutation data, pedigree, clinical correlation, or functional assays involving CHTF8. Therefore the paper provides neither direct evidence (genetic/clinical association of CHTF8 with a Mendelian phenotype) nor mechanistic evidence (pathway, expression, interaction data implicating CHTF8) relevant to the claim. The RNA-seq and differential-expression statement could hypothetically have contained information about other genes, but the excerpt does not report those gene identities; absent that data, no inference about CHTF8 can be drawn from this paper.


[Read Paper](https://www.semanticscholar.org/paper/360cdbcf8312091705578648d44e17a6c6c30cf2)


### SETD2 — linking stem cell survival and transformation

**Why Not Relevant**: The provided paper content discusses SETD2/Setd2 and its role in hematopoietic stem cell balance and in producing cell phenotypes consistent with myelodysplastic syndromes and acute myeloid leukemia. It makes no mention of the gene CHTF8, germline variants, familial segregation, or any Mendelian inheritance patterns. Therefore the content cannot support or refute the claim that CHTF8 is likely associated with at least one Mendelian disease. Additionally, the described phenotypes (MDS/AML) are commonly driven by somatic alterations and do not by themselves indicate a Mendelian (inherited) disease mechanism. Because the excerpt lacks any reference to CHTF8 or relevant germline evidence, relevance to the claim is effectively zero.


[Read Paper](https://www.semanticscholar.org/paper/6dcf99e24dd277ee03e893ba43001ebf3803a2f3)


### Non-Penetrance for Ocular Phenotype in Two Individuals Carrying Heterozygous Loss-of-Function ZEB1 Alleles

**Why Not Relevant**: The paper exclusively investigates the gene ZEB1 and its relationship to posterior polymorphous corneal dystrophy type 3 (PPCD3). None of the provided text mentions CHTF8, nor does it describe pathways, interactions, or data that would implicate or evaluate CHTF8. All quoted material is explicitly about ZEB1 (variants, cohort interrogation, population frequency, pLI score, and clinical phenotyping). Therefore the paper provides no direct genetic, phenotypic, population, or mechanistic evidence relevant to the claim that CHTF8 is likely associated with a Mendelian disease. Because there is no mention of CHTF8 and no data that could reasonably be extrapolated to CHTF8 (no overlap of gene names, no described paralogy or functional link), the relevance to the claim is effectively zero.


[Read Paper](https://www.semanticscholar.org/paper/26a1f359f1dc11c600a2a34ebaff763872b72050)


### Germline heterozygous exons 8–11 pathogenic BARD1 gene deletion reported for the first time in a family with suspicion of a hereditary colorectal cancer syndrome: more than an incidental finding?

**Why Not Relevant**: The provided paper content discusses a BARD1 deletion in a family with familial colorectal cancer type X (FCCTX) and explicitly addresses Lynch-syndrome clinical criteria and DNA mismatch repair status. There is no mention of the gene CHTF8, no description of variants, segregation data, functional assays, statistical association, or mechanistic links involving CHTF8. Because the content neither names nor implicates CHTF8, and offers no transferable mechanistic pathway connecting BARD1 findings to CHTF8, it cannot support or refute the claim that CHTF8 is likely associated with at least one Mendelian disease. To evaluate that claim, sources that explicitly discuss CHTF8 (variants, patient phenotypes, segregation, functional impact, or population association data) would be required.


[Read Paper](https://www.semanticscholar.org/paper/884a95b299f7238bf2a4a70eaeec5e815a99798c)


### Implications of a De Novo Variant in the SOX12 Gene in a Patient with Generalized Epilepsy, Intellectual Disability, and Childhood Emotional Behavioral Disorders

**Why Not Relevant**: The provided paper content concerns the SOX12 gene and its possible association with a neurodevelopmental phenotype. It repeatedly names SOX12 (and other SOX family genes) and describes a de novo variant in SOX12; it does not mention CHTF8 anywhere. Example verbatim sentences from the text showing the focus on SOX12: "Whole exome sequencing (WES) in patients presenting with generalized epilepsy, intellectual disability, and childhood emotional behavioral disorder, uncovered a de novo variation within SOX12 gene." and "The aim of this study is to establish a correlation between the SOX12 gene and the symptoms diagnosed in the patient." Because the claim is specifically about CHTF8 being likely associated with at least one Mendelian disease, and the paper contains no mention of CHTF8, no genetic data, no phenotypic correlation, and no mechanistic discussion pertaining to CHTF8, this paper provides no direct or mechanistic evidence relevant to the claim. 

Additional context on why the paper cannot be used to support the CHTF8 claim: (1) All described variants, functional-domain discussion (the HMG domain), sequence alignments, in silico pathogenicity predictions, and structural predictions are explicitly about SOX12 (and comparisons to SOX4/SOX11), not CHTF8. (2) The paper appears to be a single-case WES report focusing on a candidate novel gene-disease association (SOX12), so even for SOX12 the evidence is limited (single patient, in silico prediction, structural modeling), but this is irrelevant to CHTF8 because no data or discussion about CHTF8 are present. (3) There is no experimental, clinical, or computational link between SOX12 and CHTF8 presented that could plausibly allow inference that CHTF8 is disease-associated. 

Recommendation: To evaluate the claim about CHTF8 one should obtain literature or database evidence directly mentioning CHTF8 (e.g., OMIM, ClinVar, published case series, functional studies demonstrating pathogenic variants), since this SOX12-focused paper does not contribute any supporting or refuting evidence regarding CHTF8.


[Read Paper](https://www.semanticscholar.org/paper/5cf652470efa4406f72843aea8389f4e398a99f8)


### Identifying genomic variant associated with long QT syndrome type 2 in an ecuadorian mestizo individual: a case report

**Why Not Relevant**: This paper is a single-patient case report of Long QT Syndrome type 2 and describes genetic findings limited to a pathogenic variant in KCNH2 and two VUS in AKAP9 and TTN. The manuscript text provided contains no mention of the gene CHTF8, provides no genetic variants in CHTF8, and provides no mechanistic discussion involving CHTF8. Therefore the paper offers neither direct evidence (identified pathogenic variants, segregation data, case series) nor mechanistic evidence (functional studies, pathway links) supporting an association between CHTF8 and any Mendelian disease. Limitations and caveats: (1) The absence of CHTF8 in the reported results does not prove that the authors tested CHTF8 or that CHTF8 is not associated with disease generally — it only shows this case report did not implicate CHTF8. (2) The study is a single-case report focused on cardiac arrhythmia genes; even if CHTF8 had been tested and was negative, one negative case would not be strong evidence against an association. (3) The excerpt does not provide the full methods (gene panel composition, exome/genome scope), so it is unknown whether CHTF8 was covered by the sequencing assay used. Overall, because the paper does not mention or analyze CHTF8, it is not relevant to the claim that CHTF8 is likely associated with at least one Mendelian disease.


[Read Paper](https://www.semanticscholar.org/paper/aa65df486c4f1720d86feb42ddc72ed25ab57431)


### A de novo variant in ZBTB18 gene caused autosomal dominant non-syndromic intellectual disability 22 syndrome: A case report and literature review

**Why Not Relevant**: This paper does not provide any evidence for or against the claim about CHTF8. The report exclusively discusses a case of autosomal dominant non-syndromic intellectual disability 22 attributable to the ZBTB18 gene and describes a de novo pathogenic ZBTB18 variant; it does not mention CHTF8, any variant in CHTF8, nor any mechanistic connection between ZBTB18 and CHTF8. Representative verbatim sentences from the paper (showing its focus on ZBTB18) include:

"Rationale: Autosomal dominant non-syndromic intellectual disability 22 is a rare genetic disorder caused by the ZBTB18 gene." 

"Clinical whole exome sequencing revealed a de novo pathogenic variant in the ZBTB18 gene (c.1207delC, p. Arg403Alafs*60), which is a previously unreported site." 

"This variant causes the premature termination of peptide chain synthesis, leading to incomplete polypeptide chains." 

Additionally, the paper is a single-case report (one proband) describing clinical features, a novel ZBTB18 frameshift variant, and diagnostic considerations. Because CHTF8 is not mentioned anywhere in the provided text, there is: (1) no direct genetic association evidence linking CHTF8 to a Mendelian phenotype, (2) no mechanistic data implicating CHTF8 in the disease process described, and (3) no methodological or experimental data concerning CHTF8. Therefore the content is not relevant to evaluating the claim that "the gene CHTF8 is likely associated with at least one Mendelian disease."


[Read Paper](https://www.semanticscholar.org/paper/09bd993a331155206894ef8f2ad7afb75e164664)


### Normal and Pathological Development of Melanocytes metastasis of Targeted Knockout of β-Catenin in Adult Melanocyte Stem Cells Using a Mouse Line, Results in Disrupted Stem Cell Renewal and Defects.

**Why Not Relevant**: I found no mention of the gene symbol “CHTF8” (or an obvious synonym) anywhere in the provided paper content. The manuscript excerpts focus on melanoma biology and specific genes/pathways including STAT3, MITF, CEBP family members, TFEB/TFE3/TFEC, NRAS, BRAF, TDG, SMARCA4 (Brg1), BPTF, SOX10 and others, and on experimental approaches such as conditional mouse models, whole-genome expression profiling, ATAC-seq, 3D melanoma models, and preclinical drug sensitivity assays. The paper presents functional and mechanistic data connecting STAT3 → CEBP → repression of MITF and describes roles of other transcription factors and chromatin regulators in melanoma; it does not present genetic linkage, segregation, exome/genome sequencing, case reports, population genetics, ClinVar annotation, or other human genetic evidence that would support or refute an association of CHTF8 with a Mendelian disease. Because there is no mention of CHTF8 and no human genetic analyses relevant to Mendelian disease in the provided text, this paper provides no direct evidence and no mechanistic evidence bearing on the claim that “CHTF8 is likely associated with at least one Mendelian disease.”

Limitations: Absence of any mention of CHTF8 in this paper does not prove that CHTF8 is not associated with a Mendelian disease; it only means this paper does not provide supporting or contradicting data. To evaluate the claim one would need to consult human genetics resources (e.g., OMIM, ClinVar), segregation/linkage/exome studies, functional follow-up of rare variants, or case reports that specifically analyze CHTF8.


[Read Paper](https://www.semanticscholar.org/paper/85186af7892ccfa2b03d2e652220a9d99f875cb4)


### Whole‐exome sequencing identified five novel de novo variants in patients with unexplained intellectual disability

**Why Not Relevant**: The provided paper content consists only of background and the stated aim: "Intellectual disability (ID) represents a neurodevelopmental disorder, which is characterized by marked defects in the intellectual function and adaptive behavior, with an onset during the developmental period. ID is mainly caused by genetic factors, and it is extremely genetically heterogeneous. This study aims to identify the genetic cause of ID using trio‐WES analysis." There is no mention of the gene CHTF8, no results, no variant calls, no genotype–phenotype correlations, and no mechanistic data related to CHTF8. Therefore the excerpted content provides neither direct evidence (e.g., identification of pathogenic CHTF8 variants in affected individuals) nor mechanistic evidence (e.g., functional studies implicating CHTF8 in pathways relevant to Mendelian intellectual disability) to support or refute the claim.

Additional points relevant to interpretation:
- Methodological plausibility: trio whole-exome sequencing (trio-WES) is an appropriate method to discover Mendelian disease genes, so the overall study design could in principle generate evidence about CHTF8 if variants in that gene were observed and validated. However, no such data are present in the provided content.
- Possible incompleteness: because only a short excerpt (background and aim) was supplied, it is possible that the full paper contains relevant results (variant tables, case descriptions, functional follow-up) that mention CHTF8. With only the provided text, no evaluation of association between CHTF8 and any Mendelian disease can be made.

Conclusion: Based on the provided content, relevance to the claim is effectively zero because the gene CHTF8 is not discussed and no supporting or refuting evidence is presented. To assess the claim further, the full paper (results, tables, supplementary data) should be reviewed for any mention of CHTF8 variants or functional studies.


[Read Paper](https://www.semanticscholar.org/paper/dbdce024d78711ac3298e2df8e77679219149398)


### FRI387 Reproductive Capacity Of Znhit3 Knockout Mouse Model

**Why Not Relevant**: The provided paper content exclusively discusses the gene ZNHIT3 (also called TRIP-3) — its domains, involvement in pre-rRNA processing, previously reported human disease associations (biallelic variants with PEHO; heterozygous C-terminal variants with MRKH), and a Znhit3 knockout mouse study showing absence of homozygous pups suggesting embryonic lethality. There is no mention of CHTF8 anywhere in the abstract, introduction, methods, results, or conclusion text provided. The paper therefore contains no direct evidence (no human genetic data, no variant-disease associations, no case reports) implicating CHTF8 in Mendelian disease, nor any mechanistic data (no expression, interaction, pathway, functional studies, or homology analysis) linking CHTF8 to the described phenotypes. Because the content does not address CHTF8 at all, it cannot be used to support or refute the claim that CHTF8 is likely associated with at least one Mendelian disease. Note that absence of evidence in this paper is not evidence that CHTF8 is not disease-associated; it only means this paper is irrelevant to that claim. If you would like, I can search for and analyze publications or databases specifically about CHTF8 (e.g., ClinVar, OMIM, PubMed) to assess its Mendelian disease associations.


[Read Paper](https://www.semanticscholar.org/paper/31dabd0957ffec78267540944decdfe71d088dec)


### Case report: Preimplantation genetic testing for infantile GM1 gangliosidosis

**Why Not Relevant**: The paper exclusively addresses GM1 gangliosidosis and genetic findings in the GLB1 gene; there is no mention of the gene CHTF8, no variant data, no functional/mechanistic discussion, and no clinical associations presented for CHTF8. Therefore the paper provides no direct or mechanistic evidence for or against the claim that CHTF8 is associated with a Mendelian disease. Given the complete lack of CHTF8-related content, relevance to the claim is effectively zero.


[Read Paper](https://www.semanticscholar.org/paper/ce981bc89c565bb4d97baff441baaf8695595448)


### Autosomal Dominant Intellectual Development Disorder-6 (MRD6) Without Seizures Linked to a De Novo Mutation in the grin2b Gene Revealed by Exome Sequencing: A Case Report of a Moroccan Child

**Why Not Relevant**: The provided paper content exclusively discusses a pathogenic variant in the GRIN2B (grin2b) gene causing autosomal dominant intellectual development disorder-6 (MRD6). Example sentences from the text include: “Autosomal dominant intellectual development disorder-6 (MRD6) arises from a grin2b gene mutation, inducing neurodevelopmental issues.” and “Exome sequencing identified a nonsense (STOP) mutation c.3912C>G (p.Tyr1304Ter) in the grin2b gene in the heterozygous state known to be present in MRD6 (Online Mendelian Inheritance in Man (OMIM) 613970).” Nowhere in the provided content is the gene CHTF8 mentioned, nor are there any data, results, mechanistic descriptions, segregation analyses, or variant annotations that concern CHTF8. Therefore the paper provides no direct evidence (variant-disease association, segregation, case reports) nor mechanistic evidence (biological function, pathway involvement, model organism data) relevant to assessing whether CHTF8 is associated with a Mendelian disease. Limitations that reinforce this conclusion: the study is a single-case report focusing on identifying the causal variant for that patient (GRIN2B); although exome sequencing was performed, the authors report and interpret only the GRIN2B finding and explicitly characterize it as de novo and pathogenic—there is no mention of CHTF8 variants being observed, assessed, or ruled out. Given these facts, the paper is not relevant to the claim that CHTF8 is likely associated with at least one Mendelian disease.


[Read Paper](https://www.semanticscholar.org/paper/67d756e0cc302e5d46fdda9df34403f1f2de9a5a)


### Regulation of human heterochromatin protein 1 binding protein 3 (HP1BP3, also known as HP1BP74) in response to the DNA damage

**Why Not Relevant**: The paper's content exclusively addresses HP1BP3 (also written HP1B3/HP1-BP74), its interactions with FBXW7, regulation by ATM/ATR kinases, and functional consequences in cellular and cancer models. There is no mention of the gene CHTF8, no human genetic studies, no variant-disease segregation, no case reports, and no population or clinical data that would support an association between CHTF8 and any Mendelian disorder. All quoted material pertains to mechanistic cell and molecular biology of HP1BP3 and related pathways; such mechanistic findings about a different gene/protein do not constitute evidence that CHTF8 is associated with a Mendelian disease. Additionally, the experiments are largely biochemical and cell-line based (including CRISPR knockout in HCT116 cells), which cannot establish Mendelian inheritance or clinical disease causation in humans. For these reasons the paper provides no relevant direct or mechanistic evidence supporting the claim that CHTF8 is likely associated with at least one Mendelian disease.


[Read Paper](https://www.semanticscholar.org/paper/ff87086173716320d111684d19aadfcdf3cd0f2e)


## Search Queries Used

- CHTF8 gene pathogenic variant Mendelian disease case report family segregation OMIM ClinVar

- CHTF8 de novo variant trio exome sequencing unexplained developmental disorder neurodevelopmental intellectual disability rare variant burden

- CHTF8 protein function chromatin DNA repair development mouse knockout zebrafish CRISPR model phenotype

- CHTF8 loss of function intolerance haploinsufficiency gnomAD allele frequency pLI constraint score


## Bibliometric Configuration

- Use Bibliometrics: True

- Author Impact Weight: 0.4

- Citation Impact Weight: 0.4

- Venue Impact Weight: 0.2
