## Supplementary material for "MENDELSEEK: An algorithm that predicts Mendelian Genes and elucidates what makes them special": CIDEA.txt

# Claim: The gene CIDEA is likely associated with at least one Mendelian disease.

**Status**: processed

**Overall Rating**: -1

**Explanation**:

No relevant papers were found that support or refute this claim.


## Relevant Papers


## Other Reviewed Papers


### Genetic variants of frizzled‐4 gene in familial exudative vitreoretinopathy and advanced retinopathy of prematurity

**Why Not Relevant**: The provided paper excerpt focuses entirely on familial exudative vitreoretinopathy (FEVR), retinopathy of prematurity (ROP), and the genes NDP (ND), FZD4 and LRP5. There is no mention of the gene CIDEA, no genetic, clinical, or mechanistic data regarding CIDEA, and no discussion that could be interpreted as linking CIDEA to any Mendelian disease. Therefore this paper does not provide direct or mechanistic evidence supporting or refuting the claim that CIDEA is likely associated with at least one Mendelian disease. To evaluate the claim about CIDEA, one should consult resources and studies that specifically examine CIDEA (e.g., OMIM, ClinVar, HGMD, gnomAD constraint metrics, and PubMed-indexed genetic studies).


[Read Paper](https://www.semanticscholar.org/paper/0fd831a89ef36cbab38e7be7cb449ea6dfdeb0f7)


### The Burden of Candidate Pathogenic Variants for Kidney and Genitourinary Disorders Emerging From Exome Sequencing

**Why Not Relevant**: The provided paper content does not mention the gene CIDEA by name, nor does it provide gene-specific results or listings that would indicate whether CIDEA is included in the curated gene list or observed to harbor candidate pathogenic variants. The manuscript excerpt describes methods, curation of a gene set focused on kidney and genitourinary disorders, cohort characteristics, and variant-annotation pipelines (ClinVar, HGMD, criteria for candidate pathogenic variants), which are relevant as background and as the methodological framework that could detect gene–disease associations. However, without explicit reporting of CIDEA among the curated genes or presentation of variant findings in CIDEA, there is no direct evidence in this text to support or refute the claim that CIDEA is likely associated with at least one Mendelian disease. Any inference that CIDEA might be assessed by this pipeline is speculative: while the methods would be capable of identifying prior database reports or rare loss-of-function variants in CIDEA, the paper content provided does not supply the necessary gene-level data or results. Therefore the document is not meaningfully informative regarding the specific claim about CIDEA.


[Read Paper](https://www.semanticscholar.org/paper/a75e297208c04992568e607d19aa9b9aaebd09d5)


### Rapid genetic diagnosis in single‐gene movement disorders

**Why Not Relevant**: This article is a commentary about the use of exome sequencing in diagnosing genetically heterogeneous movement disorders and gives examples of genes (e.g. VPS13A, LRRK2, PLA2G6, VCP, ATXN2, ATXN3, EIF4G1, VPS35, TGM6, DHODH, DNAH5) and discusses methodological strengths and limitations. It does not mention the gene CIDEA, does not present any genetic, clinical, or mechanistic data related to CIDEA, and provides no evidence for or against an association between CIDEA and any Mendelian disease. Because there are no sentences or paragraphs in the provided content that reference CIDEA (or provide data that could be interpreted as evidence about CIDEA), the paper offers no direct or mechanistic support for the claim. Further, the paper is a general commentary rather than primary experimental evidence linking specific genes to disease; even its gene examples are presented as established or illustrative cases rather than new data about other genes. Absence of mention in this single paper does not prove CIDEA is not associated with a Mendelian disease, but it does mean this paper provides no relevant evidence to support the claim that CIDEA is likely associated with at least one Mendelian disease.


[Read Paper](https://www.semanticscholar.org/paper/3a1319f498ab2152bd07137e625cb9c821bfbfef)


### Exome-wide Analysis of De Novo and Rare Genetic Variants in Patients With Brain Arteriovenous Malformation

**Why Not Relevant**: Relevance is near zero because the paper focuses on germline variants and pathways underlying brain arteriovenous malformation (bAVM) and specifically reports findings implicating the ANGPTL8 regulatory pathway and the gene SLC19A3. The provided text contains no mention of the gene CIDEA, no data on CIDEA variants, and no mechanistic discussion linking CIDEA to bAVM or any other Mendelian disease. Therefore this paper provides no direct or indirect evidence supporting the claim that CIDEA is likely associated with at least one Mendelian disease. Limitations: absence of mention is not definitive proof that CIDEA has no Mendelian associations elsewhere, but within this paper there is simply no relevant information to support the claim.


[Read Paper](https://www.semanticscholar.org/paper/17051168a0ad2ef848d7692de1976646487e0c2b)


### A retrospective analysis of phosphatase catalytic subunit gene variants in patients with rare disorders identifies novel candidate neurodevelopmental disease genes

**Why Not Relevant**: The paper focuses specifically on phosphatase catalytic subunit genes (a defined set of 189 genes) and reports novel candidate phosphatase disease genes (PTPRD, PTPRG, PTPRT, PTPRU, PTPRZ1, MTMR3, GAK, TPTE2, PTPN18). CIDEA is not mentioned in the paper excerpts provided and is not among the listed candidate genes. Therefore the paper provides no direct genetic evidence (no variants, case matches, or statistical association) that CIDEA is associated with a Mendelian disease. Mechanistically, the paper discusses phosphorylation/dephosphorylation and phosphatase function; unless CIDEA were a phosphatase catalytic subunit (which it is not within the study scope), those mechanisms do not establish a link to CIDEA. Absence of mention in this phosphatase-focused dataset and candidate list means this paper is not a source of evidence for the claim. Note that absence of evidence in this paper is not evidence of absence overall: CIDEA could be implicated by other studies not covered here. Given the lack of any mention or data on CIDEA in the provided content, the paper's relevance to the claim is negligible.


[Read Paper](https://www.semanticscholar.org/paper/d0dfa58ee328453fa43927d0500f97d3a8f2dd39)


### P-742 Universal genome-wide haplotyping-based preimplantation genetic testing: a systematic seven-year experience at a single center

**Why Not Relevant**: The provided paper text contains no mention of the gene CIDEA, no gene-specific data, and no discussion of any gene–disease association involving CIDEA. The study is a retrospective analysis of universal preimplantation genetic testing (PGT) outcomes and reports aggregated information (e.g., numbers of couples, PGT cycles, proportions of AD/AR/XL indications, and that >200 genes were encompassed across various PGT-M indications), but it does not list those genes nor present variant, segregation, case report, or functional data for any specific gene other than mentioning BRCA1/BRCA2 by name. Establishing that a gene is “likely associated with at least one Mendelian disease” requires direct evidence such as reported pathogenic variants segregating with a Mendelian phenotype, case series, or mechanistic/functional data linking gene disruption to a disease phenotype. None of those types of evidence for CIDEA appear in this text. The only potentially related statement—that “Various PGT-M indications encompassed >200 genes in total”—is insufficient to infer that CIDEA is among them, and even if CIDEA were included in the unspecified list, the paper’s focus on PGT outcomes (implantation rates, biopsy types, embryo ranking) would not provide the necessary genetic or phenotypic evidence to support a gene–disease association. Therefore the paper content is not relevant for supporting or refuting the claim about CIDEA and Mendelian disease.


[Read Paper](https://www.semanticscholar.org/paper/ba73b220921052593f3ba2b10af90f6f05bed83d)


### 1684: PATHOGENIC VARIATION ASSOCIATED WITH MONOGENIC DISORDERS IN A SEVERE PEDIATRIC SEPSIS COHORT

**Why Not Relevant**: The provided paper content does not mention the gene CIDEA, nor provide any variant-level data, mechanistic discussion, or disease associations for CIDEA. The manuscript excerpt describes whole exome sequencing in 45 children with severe sepsis, screening a defined set of 96 genes and reporting pathogenic or likely pathogenic variants linked to specific monogenic disorders (for example, atypical hemolytic uremic syndrome, autoimmune lymphoproliferative syndrome, cryopyrin-associated periodic syndromes, etc.). However, none of the reported disorders, variant descriptions, or sentences in the provided text refer to CIDEA. Therefore the paper content as provided offers no direct evidence for or against the claim that CIDEA is likely associated with at least one Mendelian disease.

Limitations and caveats: It is possible that CIDEA was included among the 96 genes screened but not mentioned in this short excerpt because no pathogenic/likely pathogenic variants in CIDEA were found, or because results for CIDEA appeared only in full-text tables or supplements that were not provided here. The absence of mention in the supplied text is absence of evidence, not evidence of absence. To properly evaluate the claim regarding CIDEA, one would need the complete paper (including methods' gene list, full variant tables, or supplements) or independent sources (OMIM, ClinVar, HGMD, peer-reviewed reports) that specifically address CIDEA and Mendelian disease associations.


[Read Paper](https://www.semanticscholar.org/paper/cdd32cc853ff52097298692a62266f79b44b8c53)


### #4317 CLINICAL DIVERSITY OF STEROID-RESISTANT NEPHROTIC SYNDROME CAUSED BY TRPC6 MUTATIONS

**Why Not Relevant**: The paper focuses on genetic causes of steroid-resistant nephrotic syndrome (SRNS) in a cohort of 39 Japanese patients and reports pathogenic variants in known SRNS-related genes (e.g., TRPC6, NUP107, PLCE1, COL4A3, COL4A5). The gene CIDEA is not mentioned anywhere in the provided text, and no direct or mechanistic evidence linking CIDEA to any Mendelian disease is presented. While the methods (exome/panel sequencing and pathogenic-variant filtering) could have detected coding pathogenic variants, the study is phenotype-specific (SRNS), so the absence of CIDEA findings here only indicates that CIDEA was not implicated in SRNS in this cohort. It does not constitute evidence that CIDEA is or is not associated with any Mendelian disease in general. Therefore the paper provides no direct support for the claim and only very weak indirect (absence) evidence confined to this specific disease context and sample. Key limitations: small, disease-specific cohort; possible panel/exome coverage or filtering choices that could miss or exclude relevant variants; and absence of broader phenotypic or population-level investigation of CIDEA.


[Read Paper](https://www.semanticscholar.org/paper/c2747aaff3d6f0c6e29493cdff73cf48fd818d22)


### Abstract 1452: Analyses of whole exome sequencing (WES) data of hereditary lung cancer families identify germline copy number variations (CNVs) in multiple genes

**Why Not Relevant**: This paper does not provide evidence for or against an association between CIDEA and any Mendelian disease. The study focuses on germline copy-number variations (deletions) identified by whole-exome sequencing in hereditary lung cancer families and explicitly lists the genes in which CNVs were found (KIR2DL1, DMBT1, SIBPB1, LILRA6, HLA-DRB5). CIDEA is not mentioned anywhere in the provided content. Therefore there is no direct evidence (no reported pathogenic variants, segregation data, or statistical association for CIDEA) and no mechanistic evidence (no functional data, pathway links, or variant effect discussion specific to CIDEA) in this paper. Additional important caveats: the content is an abstract (preliminary data), CNV-focused (so it would not detect many other variant types that could implicate CIDEA), the sample is limited to hereditary lung cancer families (so negative findings here do not generalize to other diseases or populations), and validation was still ongoing. For these reasons the paper is essentially irrelevant to the claim that CIDEA is likely associated with at least one Mendelian disease.


[Read Paper](https://www.semanticscholar.org/paper/44bdd9bc1f77714a2ed2ff2f1307cc81e9c6f37d)


## Search Queries Used

- CIDEA mutation human Mendelian disease monogenic disorder single gene inherited familial case report patient autosomal dominant autosomal recessive X linked

- CIDEA ClinVar OMIM pathogenic variant ClinVar record gnomAD pLI loss of function intolerance constraint CIDEA gene annotation

- CIDEA function adipocyte lipid droplet thermogenesis energy homeostasis apoptosis Cidea knockout mouse phenotype lipodystrophy insulin resistance congenital metabolic disease

- whole exome sequencing rare variant burden de novo mutation trio sequencing families undiagnosed Mendelian disorders candidate gene CIDEA rare variants association


## Bibliometric Configuration

- Use Bibliometrics: True

- Author Impact Weight: 0.4

- Citation Impact Weight: 0.4

- Venue Impact Weight: 0.2
