## Supplementary material for "MENDELSEEK: An algorithm that predicts Mendelian Genes and elucidates what makes them special": CKMT1A.txt

# Claim: The gene CKMT1A is likely associated with at least one Mendelian disease.

**Status**: processed

**Overall Rating**: -1

**Explanation**:

No relevant papers were found that support or refute this claim.


## Relevant Papers


## Other Reviewed Papers


### Segregation of ATP10B variants in families with autosomal recessive parkinsonism

**Why Not Relevant**: The provided paper content does not mention CKMT1A, nor does it present any data or mechanistic discussion relating to Mendelian disease association for CKMT1A. The single sentence given states: "This study indicates that biallelic ATP10B missense variants increase the risk of PD, and provides evidence for a cis location of the two variants, both inherited from the mother, who was unaffected at age 58, suggesting that they cannot be causal for PD." That sentence refers specifically to ATP10B and Parkinson's disease (PD), discusses cis configuration of two variants and maternal inheritance with lack of maternal phenotype, and concludes these variants are unlikely causal for PD. There is no mention of CKMT1A, no evidence about segregation, pathogenicity, loss-of-function, gain-of-function, population frequency, or clinical phenotypes relevant to CKMT1A, and no mechanistic pathway linking ATP10B to CKMT1A or to a Mendelian disorder. Therefore this paper provides no direct or mechanistic evidence for or against the claim that CKMT1A is associated with at least one Mendelian disease. Limitations of this assessment: only the supplied sentence was available; if the paper contains other sections not provided here that mention CKMT1A, those were not evaluated.


[Read Paper](https://www.semanticscholar.org/paper/3010f2ddb5d6726ee0fe09b3dc43368ec04f224f)


### Non-invasive prenatal sequencing for multiple Mendelian monogenic disorders using circulating cell-free fetal DNA

**Why Not Relevant**: The provided paper content is a single summary sentence describing a prenatal test that detects mutations across 30 genes associated with dominant monogenic diseases and its reported high accuracy. The content does not (a) name CKMT1A, (b) provide the list of the 30 genes tested, (c) present any variant-level associations between CKMT1A and disease, or (d) give mechanistic or clinical evidence linking CKMT1A to a Mendelian disorder. Because the claim concerns the association of CKMT1A with at least one Mendelian disease, and the paper content gives no gene-level or variant-level information, no phenotype description, nor any mechanistic insight regarding CKMT1A, the content is insufficient to support or refute the claim. It remains possible (but unverifiable from this content) that CKMT1A could be among the 30 genes; however, without explicit naming or data, no inference about CKMT1A's disease association can be drawn. To evaluate the claim one would need the paper's full gene list, reported associations for each gene, variant pathogenicity evidence, cohort phenotypes, and any functional or segregation data relevant to CKMT1A.


[Read Paper](https://www.semanticscholar.org/paper/965b5d12657d5a39be70c31cf4cb3eb1617f5e96)


### Identifying genomic variant associated with long QT syndrome type 2 in an ecuadorian mestizo individual: a case report

**Why Not Relevant**: This paper provides no direct or mechanistic evidence relating to the gene CKMT1A. The report is a single-case clinical/genetic case report of Long QT Syndrome type 2 and explicitly discusses variants in KCNH2, AKAP9, and TTN. Relevant sentences from the provided content identify the causal gene and detected variants (for example: “...a pathogenic variant in the KCNH2 gene p.(Ala614Val) (ClinVar ID: VCV000029777.14) associated with LQTS type 2, and two variants of uncertain significance in the AKAP9 p.(Arg1654GlyfsTer23) (rs779447911), and TTN p. (Arg34653Cys) (ClinVar ID: VCV001475968.4) genes were identified.”), but CKMT1A is not mentioned anywhere in the supplied text. Therefore the paper neither supports nor refutes the claim that CKMT1A is likely associated with a Mendelian disease.

Additional points relevant to interpreting this absence:
- Lack of mention is not evidence against an association: the paper focuses on a specific clinical diagnosis (LQTS) and the genes known to be associated with that phenotype; it does not represent a systematic survey of all Mendelian disease genes. Thus absence of CKMT1A in this report does not argue that CKMT1A is not associated with any Mendelian disease.
- Scope and sample size limit utility: this is a single-case report (one patient), so even if CKMT1A had relevance to broad Mendelian disease genetics, this study would be unlikely to detect or address it unless the gene were directly implicated in the patient’s phenotype.
- No mechanistic data: the paper includes no functional, expression, segregation, or biochemical data that could be extrapolated to CKMT1A.

Conclusion: relevance to the claim is effectively zero because CKMT1A is not discussed and no evidence (direct or mechanistic) about CKMT1A’s disease association is present in the paper.


[Read Paper](https://www.semanticscholar.org/paper/aa65df486c4f1720d86feb42ddc72ed25ab57431)


### Case report: Preimplantation genetic testing for infantile GM1 gangliosidosis

**Why Not Relevant**: The paper exclusively reports a case of GM1 gangliosidosis caused by mutations in the GLB1 gene, including identification of GLB1 variants, development of a PGT-M testing system for GLB1, and embryo/fetal testing outcomes. CKMT1A is not mentioned anywhere in the provided content, no experiments or analyses involve CKMT1A, and no mechanistic discussion links GLB1 biology to CKMT1A. Therefore this paper provides no direct or mechanistic evidence supporting or refuting the claim that CKMT1A is likely associated with at least one Mendelian disease. The absence of CKMT1A in this study does not constitute evidence against an association in other studies, but it does mean this paper is irrelevant to assessing that claim.


[Read Paper](https://www.semanticscholar.org/paper/ce981bc89c565bb4d97baff441baaf8695595448)


## Search Queries Used

- CKMT1A mutation pathogenic variant Mendelian monogenic disease OMIM ClinVar case report

- CKMT1A mitochondrial creatine kinase function loss of function missense variant enzyme activity energy metabolism myopathy cardiomyopathy neurodegeneration functional study

- CKMT1A variants gnomAD allele frequency ClinVar loss of function constraint exome sequencing rare disease cohort trio family segregation autosomal recessive biallelic

- mitochondrial creatine kinase CKMT1A CKMT1B CKMT2 hereditary myopathy cardiomyopathy mitochondrial disease inborn error creatine kinase deficiency review systematic review


## Bibliometric Configuration

- Use Bibliometrics: True

- Author Impact Weight: 0.4

- Citation Impact Weight: 0.4

- Venue Impact Weight: 0.2
