## Supplementary material for "MENDELSEEK: An algorithm that predicts Mendelian Genes and elucidates what makes them special": CLASP2.txt

# Claim: The gene CLASP2 is likely associated with at least one Mendelian disease.

**Status**: processed

**Overall Rating**: -1

**Explanation**:

No relevant papers were found that support or refute this claim.


## Relevant Papers


## Other Reviewed Papers


### ClinGen’s RASopathy Expert Panel Consensus Methods for Variant Interpretation

**Why Not Relevant**: The provided paper content does not mention CLASP2 by name, does not present gene-specific variant or clinical evidence, and offers only a general statement about pathway-focused ACMG-AMP specification for RASopathy-associated variants. There are no mechanistic details linking CLASP2 to the Ras/MAPK pathway or to any Mendelian phenotype in the excerpt. Because the claim concerns association of a specific gene (CLASP2) with Mendelian disease, and the excerpt provides neither direct genetic/clinical evidence nor mechanistic links for CLASP2, the content is not relevant to supporting or refuting the claim. It is possible the full paper (not provided here) mentions additional genes, but based solely on the supplied text there is no basis to evaluate CLASP2’s disease association.


[Read Paper](https://www.semanticscholar.org/paper/54e98adccb563b01deee283d5a738e8135c7a8d6)


### The Burden of Candidate Pathogenic Variants for Kidney and Genitourinary Disorders Emerging From Exome Sequencing

**Why Not Relevant**: The provided paper excerpt does not mention the gene CLASP2 anywhere and contains no variant-level, case-level, or experimental data about CLASP2. The content is methodological and describes how a curated list of 625 genes associated with Mendelian kidney and genitourinary disorders was assembled and annotated, but the excerpt does not include the gene list itself (only a statement that the list is available on the laboratory website). Therefore, there is no direct evidence in this text that CLASP2 is associated with a Mendelian disease, nor any mechanistic discussion specific to CLASP2. To evaluate the claim that CLASP2 is likely associated with at least one Mendelian disease would require (a) checking the curated gene list referenced (laboratory website or supplementary materials) to see if CLASP2 is included; (b) searching primary gene–disease resources such as OMIM, ClinVar, HGMD, or peer-reviewed case reports for documented CLASP2 pathogenic variants and phenotype associations; or (c) reviewing experimental/mechanistic studies implicating CLASP2 in a Mendelian phenotype. Because none of those data are present in the provided excerpt, the excerpt is not relevant for supporting or refuting the claim.


[Read Paper](https://www.semanticscholar.org/paper/a75e297208c04992568e607d19aa9b9aaebd09d5)


### Familial focal segmental glomerulosclerosis associated with an ACTN4 mutation and paternal germline mosaicism.

**Why Not Relevant**: The paper's content exclusively discusses ACTN4 (and to a lesser extent NPHS1) in relation to focal segmental glomerulosclerosis (FSGS1) and reports a novel ACTN4 mutation and germline mosaicism. There is no mention of CLASP2, no experimental data, clinical observations, or mechanistic discussion involving CLASP2, and therefore the paper provides no direct or mechanistic evidence to support or refute the claim that CLASP2 is associated with a Mendelian disease. The absence of CLASP2 discussion in this study means it is not relevant to the claim; however, absence in this paper alone is not evidence that CLASP2 is not associated with Mendelian disease—only that this specific paper does not address CLASP2.


[Read Paper](https://www.semanticscholar.org/paper/16a08fd6ebb15730e4e5a44956e57231edeef62a)


### Diagnostic yield of patients with undiagnosed intellectual disability, global developmental delay and multiples congenital anomalies using karyotype, microarray analysis, whole exome sequencing from Central Brazil

**Why Not Relevant**: The provided paper content does not mention the gene CLASP2 anywhere, so it gives no direct evidence that CLASP2 is associated (or not associated) with any Mendelian disease. While the study used an exome-based ID gene panel of 1,252 genes and identified pathogenic variants in a subset of 19 trios, the text does not list gene-level results or describe mechanistic findings for specific genes. Therefore, one cannot infer from these excerpts whether CLASP2 was included on the panel, whether any CLASP2 variants were observed, or whether such variants segregated with disease or had functional validation. Additional reasons the paper cannot support the claim here: (1) the exome-panel sample size for sequencing (19 trios) is small, limiting power to detect rare associations; (2) absence of reported gene names or variant-level data prevents assessment of statistical or practical significance for any specific gene; (3) the study design and reported summaries do not include functional/mechanistic experiments that would support causality for a specific gene. To evaluate the claim about CLASP2, one would need either (a) explicit reporting of CLASP2 variants classified as pathogenic/likely pathogenic with segregation and phenotype data, (b) functional evidence linking CLASP2 disruption to disease-relevant mechanisms, or (c) independent genetic evidence (multiple unrelated families or case-control enrichment) — none of which are present in the provided text.


[Read Paper](https://www.semanticscholar.org/paper/23767daf16f48d9a91172aa0fc624f07b869a9c8)


### The G534E variant in HABP2 is not associated with increased risk of familial nonmedullary thyroid cancer in Brazilian Kindreds

**Why Not Relevant**: This paper investigates the role of a specific HABP2 germline variant (G534E) in familial nonmedullary thyroid cancer (FNMTC). It repeatedly names HABP2 as the gene of interest and reports genotyping and negative cosegregation results for that variant. The paper contains no mention of the gene CLASP2, no data, and no mechanistic discussion involving CLASP2. Therefore it provides no direct or mechanistic evidence bearing on the claim that CLASP2 is likely associated with at least one Mendelian disease. The study's relevance is effectively zero: its findings are limited to HABP2 in FNMTC kindreds (Brazilian cohort of 16 families, n=82 genotyped), and nothing in the methods, results, or discussion addresses CLASP2. Consequently, this paper cannot be used to support or refute the CLASP2–Mendelian disease claim.


[Read Paper](https://www.semanticscholar.org/paper/5c7ac0ba6467035bb1d39eb311512e46953c6e35)


### Germline APC Mutation and Familial Barrett Esophagus: Causal or Coincidence?

**Why Not Relevant**: This paper does not provide any direct or mechanistic evidence regarding the gene CLASP2. The text discusses familial Barrett esophagus, esophageal adenocarcinoma, and familial adenomatous polyposis/attenuated FAP (APC gene), and it describes Wnt signaling and β-catenin–related mechanisms. Nowhere in the provided content is CLASP2 named or discussed. Therefore there is no basis in this paper to support or refute the claim that CLASP2 is associated with a Mendelian disease. Specific points: (1) The genetic locus and gene emphasized throughout is APC; all genetic/mechanistic discussion centers on APC, β-catenin, and Wnt pathway alterations, not on CLASP2. (2) The paper is largely descriptive and speculative with respect to links between AFAP/FAP and familial Barrett esophagus (case report context, suggestions of possible linkage), and it cites segregation analysis suggesting rare autosomal-dominant susceptibility alleles for familial Barrett esophagus, but it does not identify any such genes. Thus even mechanistic plausibility for CLASP2 cannot be inferred from this paper. (3) Study types and limitations (case report, family observations, discussion of genotype–phenotype correlations for APC) further limit generalizability, and there is no experimental, genetic, linkage, sequencing, expression, or functional data pertaining to CLASP2. Because the required verbatim evidence for or against CLASP2 is absent, the paper is not relevant to the claim.


[Read Paper](https://www.semanticscholar.org/paper/3d4cf4dffb9bfae5c0410596be5f39acbb54d989)


### Homozygous Mutations in Thyroid Peroxidase (TPO) in Hypothyroidism with Intellectual Disability, Developmental Delay, and Hearing and Ocular Anomalies in Two Families: Severe Manifestation of Untreated TPO-deficiency Poses a Diagnostic Dilemma

**Why Not Relevant**: The paper focuses on congenital hypothyroidism (CH) and identifies homozygous pathogenic variants in the TPO gene as the genetic cause in the reported families. Nowhere in the provided content is the gene CLASP2 mentioned, studied, or implicated. The molecular analyses reported (SNP-based homozygosity mapping, linkage analysis, exome sequencing) led to TPO variant discovery, not CLASP2. Therefore this paper provides no direct evidence that CLASP2 is associated with any Mendelian disease, nor does it present mechanistic data implicating CLASP2. Absence of mention in this study is not definitive proof that CLASP2 has no Mendelian associations elsewhere, but for the purposes of evaluating CLASP2 specifically, this paper is not relevant.


[Read Paper](https://www.semanticscholar.org/paper/348b669fb02f61a677545917eed0c87312b1c29e)


### Genetic Analysis in a Familial Case With High Bone Mineral Density Suggests Additive Effects at Two Loci

**Why Not Relevant**: The paper does not mention the gene CLASP2 anywhere in the provided content. All identified candidate genes discussed are VAV3 and ADGRE5, with mechanistic and mouse-model evidence supporting their role in high bone mineral density in this family. The methods (whole-exome sequencing of three affected and three unaffected relatives) could in principle detect coding variants in CLASP2 if they existed, but because CLASP2 is not referenced in the results, discussion, or conclusions, this paper provides no direct genetic association, mechanistic data, or discussion relevant to the claim that CLASP2 is likely associated with at least one Mendelian disease. Limitations to this conclusion: absence of evidence in this paper is not evidence of absence—this single family study focused on other genes and may not have reported negative findings for every gene; also, if CLASP2 harbors noncoding variants or structural variants not detected by the reported analyses, those would not appear in these excerpts. Nonetheless, based on the content provided, relevance of this paper to the CLASP2–Mendelian disease claim is effectively zero.


[Read Paper](https://www.semanticscholar.org/paper/de079d062b38ceaec92ab3987723acef25f28dff)


### Contribution of variants across the allelic frequency spectrum to cystic kidney disease

**Why Not Relevant**: The paper presents a comprehensive whole-genome analysis of genetic contributors to cystic kidney disease (CyKD) and lists genes significantly associated with that phenotype (PKD1, PKD2, DNAJB11, COL4A3, PKHD1, IFT140, HNF1B locus). The paper does not mention CLASP2 at any point, nor does it present mechanistic data or associations linking CLASP2 to any Mendelian disease. Therefore this manuscript provides no direct evidence supporting that CLASP2 is associated with a Mendelian disease. It does provide indirect negative evidence for CLASP2 specifically causing CyKD in the studied cohorts (i.e., CLASP2 was not among identified or emergent gene-level associations), but that absence is limited to this phenotype and the samples analyzed and cannot exclude CLASP2 involvement in other Mendelian disorders. Key limitations: the study is disease-specific (CyKD), so its negative findings do not generalize to all Mendelian diseases; lack of mention of CLASP2 is not definitive proof of lack of any Mendelian association; and the analysis sensitivity depends on variant frequency, effect size, inheritance mode, and phenotypic ascertainment. Because the paper contains no information about CLASP2, its relevance to the claim is negligible.


[Read Paper](https://www.semanticscholar.org/paper/ac802a1b5b377860f361a8b5d89ff182d78b02b3)


### Novel SYNGAP1 Variant in an Adult Individual Affected by Intellectual Disability and Epilepsy: A Cold Case Solved through Whole-Exome Sequencing

**Why Not Relevant**: The paper's content, as provided, does not mention the gene CLASP2 at all. All molecular findings and interpretations concern a novel de novo variant in SYNGAP1 and the proband's phenotype consistent with SYNGAP1-related disease. Therefore the paper provides no direct evidence (variant, segregation, case reports) nor mechanistic evidence (functional studies, pathways implicating CLASP2) supporting the claim that CLASP2 is associated with a Mendelian disease. Because CLASP2 is not discussed, cited, or analyzed in this report, its relevance to the claim is effectively zero. Limitations: conclusion based solely on the provided text; if the full paper contains unprovided sections mentioning CLASP2, they were not available for analysis.


[Read Paper](https://www.semanticscholar.org/paper/e2f8d50ff5458adc25bc78983e545080ac7d5e62)


### The Scope of Mendelian Cardiomyopathy Genes

**Why Not Relevant**: The provided content does not mention CLASP2, any specific gene, any specific disease, Mendelian inheritance, pathogenic variants, patient data, experimental results, or any mechanistic details linking a particular gene to a Mendelian phenotype. It contains only a general recommendation that genes of primary cardiomyopathies (CMs) have broad effects and that molecular genetic studies are needed. Because the claim concerns a specific gene (CLASP2) and its likely association with at least one Mendelian disease, the absence of any specific data, observations, or mechanistic description about CLASP2 or Mendelian disorders makes the content effectively irrelevant for supporting or refuting the claim. The low relevance score reflects this lack of specific, direct, or even gene-level mechanistic evidence.


[Read Paper](https://www.semanticscholar.org/paper/3c511ca1645f8807664714166ae870af33d40bbc)


### The Scope of Mendelian Cardiomyopaties Genes

**Why Not Relevant**: The provided excerpt is a general review about the scope of genes implicated in Mendelian cardiomyopathies and states that 75 genes have pathogenic/probably pathogenic variants according to Simple ClinVar. However, the excerpt does not mention the gene CLASP2 anywhere, nor does it provide a list of the 75 genes or any gene-level data that would allow determining whether CLASP2 is among them. Therefore the paper content as provided gives no direct evidence (no case reports, variants, ClinVar/OMIM entries, or functional studies) that CLASP2 is associated with any Mendelian disease. It only provides indirect, high-level context about how CM genes commonly have broad tissue expression and diverse pathway involvement, which is insufficient to support the specific claim. Additionally, because the full gene list and any supplementary tables are not present in the excerpt, it is possible (but unresolvable from this content) that CLASP2 might be included elsewhere in the full paper; without that, no gene-specific inference can be made.


[Read Paper](https://www.semanticscholar.org/paper/14f6566784da01feb6aaf58462ae97bc7a8a6dbe)


### A novel AUTS2 variant in a patient with global developmental delay and intellectual disability

**Why Not Relevant**: The provided paper content exclusively concerns AUTS2 (also spelled AUST2 in one sentence) and reports a novel de novo loss-of-function variant in AUTS2 in a single case. There is no mention of the gene CLASP2 anywhere in the text, no genetic, clinical, or functional data about CLASP2, and no discussion of pathways, interactions, or mechanisms that link AUTS2 to CLASP2. Therefore the paper does not provide direct evidence that CLASP2 is associated with any Mendelian disease. Furthermore, although both AUTS2 and CLASP2 may be involved in neurodevelopmental processes in broader literature, nothing in this paper establishes such a connection. Given the absence of CLASP2-specific data (variants, segregation, functional studies, or mechanistic links), this paper is not relevant to supporting or refuting the claim that CLASP2 is likely associated with at least one Mendelian disease.


[Read Paper](https://www.semanticscholar.org/paper/f5e84e7bb3b42239a53cc7e5c457342cb38c6277)


## Search Queries Used

- CLASP2 mutation case report familial germline variant monogenic Mendelian disease

- CLASP2 exome sequencing rare variant intellectual disability developmental delay epilepsy congenital anomaly

- cytoplasmic linker associated protein 2 CLASP2 microtubule plus end tracking neuronal migration axon guidance cortical development knockout mouse phenotype

- CLASP2 pathogenic variants ClinVar OMIM ClinGen review gene disease association Mendelian evidence


## Bibliometric Configuration

- Use Bibliometrics: True

- Author Impact Weight: 0.4

- Citation Impact Weight: 0.4

- Venue Impact Weight: 0.2
