## Supplementary material for "MENDELSEEK: An algorithm that predicts Mendelian Genes and elucidates what makes them special": CMC4.txt

# Claim: The gene CMC4 is likely associated with at least one Mendelian disease.

**Status**: processed

**Overall Rating**: -1

**Explanation**:

No relevant papers were found that support or refute this claim.


## Relevant Papers


## Other Reviewed Papers


### mGAP: the macaque genotype and phenotype resource, a framework for accessing and interpreting macaque variant data, and identifying new models of human disease

**Why Not Relevant**: The provided paper content contains a single general statement about the mGAP resource: it is a public website with searchable, annotated macaque variant data that can be used to investigate genetic associations and identify macaque models based on parallels with human risk alleles. The text does not mention the gene CMC4, any specific variants, any disease phenotypes, nor any mechanistic data tying CMC4 to Mendelian disease. Therefore it provides no direct evidence (no reported associations, statistics, or case/control results) and no mechanistic evidence (no functional studies, pathways, or variant effects) to support or refute the claim that CMC4 is likely associated with at least one Mendelian disease. The only relevant point is that mGAP as a resource could be used in future to investigate such associations, but that is a statement about potential utility rather than evidence. Limitations: absence of gene name, absence of variant-phenotype links, absence of experimental details or statistical results; therefore the paper content as provided cannot be used to evaluate the claim.


[Read Paper](https://www.semanticscholar.org/paper/f47691ff666dd65548f1282e606d696eae56524d)


### Molecular findings among patients referred for clinical whole-exome sequencing: Editorial comment

**Why Not Relevant**: The provided paper text is a general report of clinical whole-exome sequencing in a 2000-patient cohort describing methods, cohort composition, overall diagnostic yield (504/2000 = 25.2%), variant counts and types, and modes of inheritance. Nowhere in the supplied content is the gene name CMC4 mentioned, nor are gene-level results, tables, or specific gene–disease associations included that would allow assessment of whether CMC4 is associated with a Mendelian disease. Although the study demonstrates that whole-exome sequencing can and did identify many disease-causing variants (and that many causative alleles were novel), the excerpt does not provide any direct evidence (variant, segregation, or phenotype association) or mechanistic data specifically linking CMC4 to disease. Therefore this paper, as provided, cannot be used to support or refute the claim that CMC4 is likely associated with at least one Mendelian disease. To evaluate the claim would require gene-specific data absent here (e.g., explicit listing of variants in CMC4 with pathogenicity assessments, segregation or de novo status, phenotypic correlation, functional studies, or references to prior disease reports).


[Read Paper](https://www.semanticscholar.org/paper/a122c53f80c050b861b70f5e57fbede67e0b7263)


### Mapping of breakpoints in balanced chromosomal translocations by shallow whole-genome sequencing points to EFNA5, BAHD1 and PPP2R5E as novel candidates for genes causing human Mendelian disorders

**Why Not Relevant**: The paper does not mention the gene CMC4 anywhere in the provided content. The study reports disrupted genes identified in nine probands with de novo balanced chromosomal translocations and discusses EFNA5, EBF3, LARGE, PPP2R5E, TXNDC5, ZNF423, NIPBL, BAHD1, RET, TRPS1 and SLC4A10. Although the paper describes methods (SGMPS and follow-up) and a prioritization strategy (pLI, DOMINO) that could in principle be applied to CMC4 in other datasets, it provides no direct or mechanistic evidence for or against an association between CMC4 and any Mendelian disease. The absence of CMC4 from the reported disrupted genes is not definitive evidence that CMC4 is not a Mendelian disease gene because (a) the study sample is small and limited to a specific class of variation (balanced translocations), (b) disease-causing variants can be of other types (SNVs, indels, CNVs) or occur in other cohorts, and (c) absence from a shortlist in one study does not refute prior or subsequent findings implicating a gene. Therefore the paper is not relevant to supporting the claim that "CMC4 is likely associated with at least one Mendelian disease."


[Read Paper](https://www.semanticscholar.org/paper/a5000f2e70b2dd6268015bdc7bfb9a566f21011e)


### Abstract 4147050: Genetic Insights into Arrhythmogenic Mitral Valve Prolapse – A Case-Control Study from UK Biobank and NIH All of Us

**Why Not Relevant**: The provided paper (abstract) does not mention the gene CMC4 anywhere in its hypothesis, methods, results, or conclusions. The study focused on Mendelian cardiomyopathy and arrhythmia genes as captured by clinical gene panels and ClinVar ≥2-star P/LP entries; it reports TTN as the most frequently implicated gene. Because CMC4 is neither named nor discussed, there is no direct evidence in this paper supporting or refuting an association between CMC4 and any Mendelian disease. Additional reasons this paper cannot inform the claim: (1) the gene panels used are not enumerated here, so CMC4 may not have been tested; (2) the analysis restricted pathogenic calls to ClinVar entries with ≥2-star curation, which would exclude novel or not-yet-curated CMC4 associations; and (3) the cohort focus (arrhythmogenic mitral valve prolapse and cardiomyopathy/arrhythmia genes) may not be the phenotype space where a CMC4 Mendelian disease would manifest. Therefore, relevance to the claim about CMC4 is effectively nil.


[Read Paper](https://www.semanticscholar.org/paper/2d898418a62e7c0d81f7271ae5e41274c04380e7)


### Contribution of rare transmitted and de novo variants among 2 , 871 1 congenital heart disease probands

**Why Not Relevant**: The provided paper content does not mention the gene CMC4 anywhere in the text excerpt. I found no verbatim sentences or paragraphs referring to CMC4, no variant, phenotype, mechanistic description, or statistical result that involves CMC4. The paper focuses on exome sequencing of congenital heart disease (CHD) probands, reporting aggregate results (e.g., counts of damaging recessive genotypes, genes implicated such as GDF1, MYH6, FLT4, and inferred numbers of contributing genes), but none of the quoted sections or tables in the provided content lists or discusses CMC4. Therefore there is no direct evidence in this excerpt that CMC4 is associated with any Mendelian disease, nor is there mechanistic evidence specific to CMC4.

Limitations and next steps: it is possible that CMC4 could appear in parts of the full paper not included in the provided excerpt (for example, in Supplementary Excel files S2–S4 or in gene lists referenced but not printed here). To evaluate CMC4 specifically, one should search the full text and all supplementary materials (especially the curated gene lists and Supplementary Excel files) for "CMC4" and, if present, extract the relevant variant, case, and statistical details. Based solely on the supplied content, relevance to the claim is effectively zero because CMC4 is not present or discussed.


[Read Paper](https://www.semanticscholar.org/paper/e5830d9e95469f22518560007746202e62db26a0)


### Cardiovascular Research Summit & Poster Fair Focus on Cardiac Arrhythmias

**Why Not Relevant**: I searched the supplied text for any mention of the gene symbol "CMC4" (or obvious synonyms) and found none. The provided content consists of multiple abstracts and excerpted paragraphs focused on cardiac proteomics, heart regeneration enhancers, myosin light chain isoforms, cardiac troponin proteomics, the 9p21.3 CAD locus, iPSC-derived vascular smooth muscle cells, and mitochondrial KATP assays (e.g., references to leptin b, MLC-1/2, cTn, SUR2A-55, ROMK, CCDC51, and the 9p21.3 locus). None of these sections contains human genetic association data, descriptions of variants, segregation/pedigree analyses, case reports of Mendelian phenotypes, citations to ClinVar/OMIM, or functional genetic experiments that implicate CMC4 in disease.

Therefore, the document provides no direct evidence (no reports of pathogenic variants, affected individuals, segregation, statistical association with Mendelian phenotypes) that CMC4 is associated with any Mendelian disease. It also provides no mechanistic evidence relevant to CMC4 (no functional assays, expression or perturbation experiments, or pathway analyses involving CMC4) that would support or refute the claim. The methods used in the cited studies (top-down and bottom-up proteomics, enhancer dissection in zebrafish, native mass spectrometry, iPSC differentiation, mitochondrial assays) are not targeted approaches for establishing gene–disease causality for a specific human Mendelian disorder; they do not include exome/genome sequencing, variant pathogenicity assessment, or familial segregation analysis.

Limitations that further reduce relevance: the excerpts include non-human models (swine, zebrafish, mouse) and small sample sizes for proteomic characterization, and focus on protein expression/biochemistry rather than genetic causality. Even where human tissue or cells are used (e.g., human cardiac tissues for troponin enrichment or iPSC lines for VSMC differentiation), the studies do not report variant discovery or Mendelian phenotype correlation.

Conclusion and recommendation: Relevance to the claim is effectively zero based on the supplied content. To evaluate whether CMC4 is likely causative for a Mendelian disease, one would need human genetic evidence such as rare (typically protein-disrupting) variants in affected individuals with segregation in families, statistical enrichment of rare variants in cases vs controls, functional assays showing variant impact consistent with disease mechanism, and/or clinical case reports linking CMC4 variants to a reproducible phenotype. None of those types of evidence appear in the provided text.


[Read Paper](https://www.semanticscholar.org/paper/bf0d25ec049e578867a2deb11b6f667f938787dd)


### MOL CLASS OF 2022 SENIOR THESIS ABSTRACTS Identification and characterization of positive regulators governing hepatitis C virus host tropism

**Why Not Relevant**: I found no direct or mechanistic evidence relevant to the claim that "CMC4 is likely associated with at least one Mendelian disease." The submitted paper content is a concatenation of multiple unrelated project/abstract summaries (HCV host factors, RNA m6A methylation, lung morphogenesis/WNT, developmental kurly mutants, Prdm1 regulation, bacterial defense systems, RTK biosensors, cerebellar tracing, etc.) and contains no mention of the gene name CMC4 (or variants such as "CMC-4", "C M C 4"), nor any discussion of Mendelian disease, monogenic inheritance, autosomal dominant/recessive inheritance, pathogenic variants, segregation, clinical phenotypes, human genetic linkage, exome/genome sequencing in affected families, or other standard lines of evidence used to implicate a gene in Mendelian disease.

To illustrate the topical focus of the provided content, here are verbatim sentences from the text (exact wording preserved) that show the document's unrelated subject matter:

- "Hepatitis C virus (HCV) continues to affect millions of lives worldwide." 

- "It is presently unknown which specific transcript(s) – being subject to epitranscriptomic regulation – are responsible for the observed phenotype." 

- "A better understanding of lung morphogenesis is key to improving current treatment options and will pave the way for future tissue engineering applications." 

- "By modulating canonical WNT using pharmacological inhibitors and activators, we show that canonical WNT signaling plays an important role in the differentiation and localization of airway smooth muscle around the epithelium." 

- "Analysis of the PADS Arsenal in 62,164 bacteria uncovered a high prevalence of defense system genes and a tendency for these genes to co-occur in the same genome, which is partially dependent on genome length." 

- "Receptor tyrosine kinases are important for studying cell signaling pathways." 

Because none of the quoted material — nor any other portion of the provided text — addresses CMC4, genetic variation in CMC4, clinical phenotypes linked to CMC4, family segregation data, functional assays of CMC4 variants, population-level enrichment of rare variants in cases versus controls, or other standard genetic/functional evidence used to implicate a gene in Mendelian disease, the document provides no support for or against the claim.

Limitations and caveats: the assessment above is limited to the exact text you supplied. If there is another section of the paper (not included here), a supplementary file, clinical case report, or separate manuscript that discusses CMC4 and human disease, that material would need to be provided to evaluate the claim. As provided, the content is insufficient and irrelevant to the claim that CMC4 is likely associated with a Mendelian disease.


[Read Paper](https://www.semanticscholar.org/paper/9571d247a1af65ff123ad6a20084e2ce86f82e58)


## Search Queries Used

- CMC4 gene human Mendelian disease association ClinVar OMIM case report patient phenotype

- CMC4 pathogenic variant loss of function missense nonsense frameshift splice site de novo inherited exome sequencing trio rare disease study

- CMC4 protein function mitochondrial localization subcellular localization expression tissue distribution knockout mouse zebrafish CRISPR functional study cellular assay

- CMC4 candidate gene gene prioritization rare variant burden test Mendelian gene discovery copy number variant CNV structural variant regulatory variant eQTL GTEx neighboring genes locus


## Bibliometric Configuration

- Use Bibliometrics: True

- Author Impact Weight: 0.4

- Citation Impact Weight: 0.4

- Venue Impact Weight: 0.2
