## Supplementary material for "MENDELSEEK: An algorithm that predicts Mendelian Genes and elucidates what makes them special": CNTN5.txt

# Claim: The gene CNTN5 is likely associated with at least one Mendelian disease.

**Status**: processed

**Overall Rating**: -1

**Explanation**:

No relevant papers were found that support or refute this claim.


## Relevant Papers


## Other Reviewed Papers


### Molecular findings among patients referred for clinical whole-exome sequencing.

**Why Not Relevant**: The supplied paper content (an observational study of 2000 clinical whole-exome sequencing cases) does not mention the gene CNTN5 anywhere in the provided text. The excerpts provided are aggregate results (diagnostic yield, proportions of inheritance modes, numbers of incidental findings) and do not include gene-level findings or mechanistic discussion of specific genes. Therefore there is no direct evidence in the provided content that CNTN5 is associated with any Mendelian disease, nor is there any mechanistic evidence concerning CNTN5. 

Specific reasons for the relevance score of 0.0:
- No occurrence of the string "CNTN5" or any obvious synonym is present in the supplied text.
- The paper text shown reports cohort-level outcomes (e.g., "A molecular diagnosis was reported for 504 patients (25.2%)...", "The Mendelian disease patterns of the 527 molecular diagnoses included 280 (53.1%) autosomal dominant...") but does not list the genes responsible for those diagnoses in the provided excerpt.
- Mechanistic discussion required to support plausibility (protein function, pathogenic variant types, disease mechanism) is absent for any individual gene in the provided material.

Limitations and next steps:
- This assessment is based only on the supplied excerpt. The full paper (main text, tables, and especially supplementary materials) may include a gene-level list or case reports that mention CNTN5. If you need a definitive determination, obtain and search the full text and supplements (including gene lists or Table Sx) for "CNTN5" or review databases such as OMIM, ClinVar, HGMD, or gene-disease curation resources (ClinGen) to see whether CNTN5 has an established Mendelian disease association.
- Even if CNTN5 appeared in the full paper, one should evaluate whether the reported variants meet standards for pathogenicity (segregation, functional evidence, recurrence) before concluding a Mendelian association.


[Read Paper](https://www.semanticscholar.org/paper/887bd08e76f0b826324d90f481b823ee9c4aef30)


### The Burden of Candidate Pathogenic Variants for Kidney and Genitourinary Disorders Emerging From Exome Sequencing

**Why Not Relevant**: The provided paper content focuses on identifying and curating genes associated with Mendelian kidney and genitourinary disorders and on assessing the prevalence of candidate pathogenic variants in a curated set of 625 such genes in a cohort of healthy adults. The excerpts and methods describe the gene curation process (OMIM/Orphanet), the final gene set size, and variant-annotation pipelines (ClinVar/HGMD), but the supplied text does not mention CNTN5 or present the full gene list or gene-level results. Therefore, the paper as provided contains no direct evidence that CNTN5 is associated with any Mendelian disease. Mechanistically, the paper's approach (curating disease-specific genes, prioritizing loss-of-function and previously reported pathogenic variants, and filtering by population frequency and inheritance mode) is relevant for assessing gene–disease associations in general, but because CNTN5 is not discussed in the provided content and the 625-gene list is not included here, no direct or mechanistic evidence about CNTN5 can be extracted. Caveats: CNTN5 could conceivably appear in the full 625-gene list or in appendices or on the referenced laboratory website (not included in the excerpt). To determine whether CNTN5 is likely associated with a Mendelian disease, one should directly query disease-gene resources (OMIM, ClinVar, HGMD, GeneReviews) or the full supplementary materials of this study rather than rely on the provided excerpt.


[Read Paper](https://www.semanticscholar.org/paper/a75e297208c04992568e607d19aa9b9aaebd09d5)


### Navigating MARRVEL, a Web-Based Tool that Integrates Human Genomics and Model Organism Genetics Information.

**Why Not Relevant**: The provided paper content describes MARRVEL, a protocol and tool for aggregating human genetic and model organism data to help evaluate whether gene variants may be pathogenic. It does not present any original data, analyses, or conclusions specific to CNTN5. There are no mentions of CNTN5, no reported variants, segregation data, statistical associations, clinical case descriptions, or functional experiments pertaining to CNTN5. Therefore the paper does not provide direct evidence that CNTN5 is associated with a Mendelian disease. At best, it is indirectly relevant because it describes resources and workflows that one could use to investigate CNTN5, but it does not itself support or refute the claim. Given the absence of CNTN5-specific content, the relevance to the claim is minimal.


[Read Paper](https://www.semanticscholar.org/paper/d4ce2c7d342a0fcd2325b51b035d147439ceecb7)


### Whole-Exome Sequencing in Adults With Chronic Kidney Disease

**Why Not Relevant**: The provided paper excerpt does not mention the gene CNTN5 anywhere, nor does it report findings (variants, segregation, phenotype correlations, functional assays, or case reports) implicating CNTN5 in any Mendelian disorder. The manuscript focuses on whole-exome sequencing in patients referred for chronic kidney disease and hypertension, the creation and use of a prioritized list of 287 nephropathy-related OMIM genes, and methodological details including regions not captured by the exome kit. While these details are useful for assessing whether an exome study could detect a disease association for a given gene, they do not constitute direct or mechanistic evidence that CNTN5 is associated with a Mendelian disease. The methodological excerpts do indicate potential blind spots (uncaptured exons, excluded genes such as PKD1) and that the authors also reviewed other OMIM and ACMG genes; however, without explicit mention of CNTN5 (in the gene list, results, tables, or text), the paper cannot be used to support the claim that CNTN5 is likely associated with at least one Mendelian disease. Therefore relevance to the claim is very low.


[Read Paper](https://www.semanticscholar.org/paper/a29bd57964f68ac5f8e1abcafe1112ba488f53f5)


### SHANK 3 overexpression causes manic-like behavior with unique pharmacogenetic properties

**Why Not Relevant**: This paper focuses on SHANK3 — its overexpression, duplications, and associated neuropsychiatric phenotypes in humans and mice. The provided content includes no mention of the gene CNTN5 (Contactin 5) and contains no data, observations, mechanistic discussion, or citations that concern CNTN5. Therefore it provides no direct evidence (genetic, clinical, or experimental) that CNTN5 is associated with any Mendelian disease, nor does it provide mechanistic evidence that would make such an association plausible by analogy. Because there is no mention or data about CNTN5, there are no relevant sentences or paragraphs to extract as supporting or refuting the claim. To evaluate the claim about CNTN5, one would need to examine other sources (e.g., primary genetic studies, ClinVar/OMIM entries, case reports of CNTN5 variants with segregation, functional studies of CNTN5 variants) — this SHANK3-focused paper cannot inform that assessment.


[Read Paper](https://www.semanticscholar.org/paper/b7cc563b034a688f8a7031a93315f2330b226fb1)


### Hereditary platelet function disorder from RASGRP2 gene mutations encoding CalDAG-GEFI identified by whole-exome sequencing in a Korean woman with severe bleeding

**Why Not Relevant**: This paper does not provide any evidence for or against the claim that CNTN5 is likely associated with a Mendelian disease. The manuscript focuses entirely on an inherited platelet function disorder caused by compound heterozygous mutations in RASGRP2, with clinical, genetic, and in silico evidence supporting RASGRP2 as the causal gene in the reported patient. Nowhere in the provided text is CNTN5 mentioned, nor are there any data (genetic, functional, or segregation) regarding CNTN5. The authors report whole-exome sequencing results that identified two heterozygous mutations in RASGRP2 and state that 72 genes associated with platelet disorders were screened, but they do not list those genes nor report any findings in CNTN5. Therefore: (1) there is no direct evidence linking CNTN5 to any Mendelian disease in this paper (no variants, no segregation, no phenotype correlation); (2) there is no mechanistic discussion involving CNTN5 that could plausibly tie it to the platelet disorder or any other Mendelian phenotype; and (3) absence of mention in this paper does not constitute evidence that CNTN5 is not associated with a Mendelian disease, it only means this paper provides no support for the claim. Because the paper contains no relevant sentences or data about CNTN5, excerpts and excerpt-specific explanations are not provided.


[Read Paper](https://www.semanticscholar.org/paper/d1297a1df15e312560df2f812ee1eaa0395296cd)


### Voluntary Running Improves Behavioral and Structural Abnormalities in a Mouse Model of CDKL5 Deficiency Disorder

**Why Not Relevant**: The provided paper content exclusively concerns CDKL5 deficiency disorder (CDD), experiments in Cdkl5 knockout mice, and the effects of voluntary running (with associated changes such as increased BDNF). There is no mention of the gene CNTN5, no human genetic data (e.g., segregation, de novo or rare variant enrichment), no variant pathogenicity evidence, and no functional assays involving CNTN5. Therefore this paper provides no direct or mechanistic evidence that CNTN5 is associated with any Mendelian disease. 

What would constitute relevant evidence (but is absent here): identification of pathogenic CNTN5 variants segregating with a Mendelian phenotype in families, statistically significant enrichment of rare damaging CNTN5 variants in affected individuals versus controls, functional assays showing that CNTN5 variants disrupt protein function in a way that plausibly causes disease, or clinical descriptions linking CNTN5 variants to a consistent phenotype. The present study's focus on a different gene (CDKL5), a mouse model, and an exercise intervention means its findings cannot be extrapolated to infer CNTN5 involvement in Mendelian disease. Additionally, species/model-specific findings for CDKL5 do not provide mechanistic parallels sufficient to implicate CNTN5.


[Read Paper](https://www.semanticscholar.org/paper/67caa6808fe3b4a3eed0e1a97e2b8946c3745591)


### The multiple faces of TOR1A: different inheritance, different phenotype.

**Why Not Relevant**: The supplied paper content does not mention CNTN5 at all and contains no data, experiments, variants, pedigrees, or mechanistic discussion specific to CNTN5. The excerpt discusses general concepts about Mendelian (monogenic) disorders and gives TOR1A as an illustrative example (DYT1-dystonia and homozygous TOR1A variants), but provides no direct evidence (no association, no variant-level data, no phenotype mapping) nor mechanistic evidence (no protein interaction, expression, functional assays, pathway membership) linking CNTN5 to any Mendelian disease. Therefore the paper content as provided cannot support or refute the claim that CNTN5 is likely associated with at least one Mendelian disease. Limitations and caveats: (1) absence of CNTN5 in this excerpt is not proof that the full paper (if longer than the provided excerpt) never mentions CNTN5 — the conclusion applies only to the provided content; (2) general statements about how mutations in a single gene can produce different inheritance patterns (as illustrated with TOR1A) are not gene-specific and cannot be used as evidence for CNTN5 without additional data; (3) no methodological details, statistical results, or experimental conditions relevant to CNTN5 are present here. Given these points, relevance to the claim is effectively zero based on the supplied text.


[Read Paper](https://www.semanticscholar.org/paper/b74b91e0829b14562a5293323e09bb3633156aab)


### Leveraging genetic diversity to understand monogenic Parkinson's disease's landscape in AfrAbia.

**Why Not Relevant**: This paper does not mention CNTN5 anywhere in the provided content and provides no data (genetic association, segregation, pathogenic variants, functional studies, or case reports) that link CNTN5 to any Mendelian disease. The article is a review/synthesis focused on monogenic forms of Parkinson's disease and explicitly lists well-established causal genes (SNCA, LRRK2, VPS35, PRKN, PINK1, and DJ) but none of these is CNTN5; no excerpt names CNTN5 or presents evidence about it. Therefore there is no direct evidence for or against the claim in this paper. Mechanistic evidence is also absent: the text does not describe CNTN5's molecular function, pathogenic mechanisms, animal models, or variant pathogenicity that could plausibly associate CNTN5 with a Mendelian disorder. Limitations: absence of mention in this paper is not evidence that CNTN5 is not associated with a Mendelian disease — it only shows that this particular review of Parkinson's disease genetics did not implicate CNTN5. To evaluate the claim that "CNTN5 is likely associated with at least one Mendelian disease," one would need papers reporting (a) rare segregating variants in families producing Mendelian inheritance with phenotype concordance, (b) statistically significant enrichment of rare damaging variants in cases vs controls, or (c) robust functional evidence that specific CNTN5 variants cause a disease phenotype. None of those types of evidence appear in the provided content, so this paper cannot be used to support the claim.


[Read Paper](https://www.semanticscholar.org/paper/c9efcd86f6f8889b27a7a94010795c933fc97e0c)


### Autism spectrum disorder and intellectual disability in an inherited 2q14.3 micro‐deletion involving CNTNAP5

**Why Not Relevant**: The paper exclusively discusses CNTNAP5 (Contactin-Associated Protein 5) and case reports of deletions affecting that gene. The claim concerns CNTN5 (Contactin 5), a distinct gene (different gene symbol and family). The provided content contains no mention of CNTN5, no data on CNTN5 sequence, function, variants, pedigrees, or mechanistic links between CNTNAP5 and CNTN5. Because the subject gene differs, the paper does not provide evidence for or against the claim that CNTN5 is likely associated with at least one Mendelian disease. Any inference would require additional data explicitly about CNTN5 or validated functional/genetic links between the two genes, which are not present in the text.


[Read Paper](https://www.semanticscholar.org/paper/55486bde9e53f788062ecc1b17957f53ee155e76)


### An adult male with SHANK2 variant with epilepsy and obsessive‐compulsive disorder: Expanding the shankopathy phenotypic spectrum

**Why Not Relevant**: This paper does not provide evidence for or against the claim that CNTN5 is likely associated with a Mendelian disease. The manuscript exclusively discusses SHANK-related disorders (SHANK1, SHANK2, SHANK3), reports a de novo frameshift variant in SHANK2, phenotypic features of a SHANK2-affected individual, and mechanistic observations related to SHANK2 (including synaptic transmission differences and mouse models). There is no mention of CNTN5, the contactin gene family, CNTN5 variants in patients, segregation data, functional studies of CNTN5, nor any pathway or interaction data linking CNTN5 to SHANK genes in this text. Because neither direct clinical/genetic evidence (e.g., pathogenic CNTN5 variants segregating with disease, de novo CNTN5 variants in affected individuals, or case reports) nor mechanistic evidence (e.g., functional assays, animal models, protein interaction data implicating CNTN5 in a Mendelian phenotype) are present, the paper is irrelevant to the claim. No verbatim excerpts can be provided that relate to CNTN5 because the gene is not discussed in the provided content. If you would like evidence about CNTN5, I can search for or analyze papers, databases (OMIM, ClinVar, gnomAD constraint metrics), or case reports specifically mentioning CNTN5.


[Read Paper](https://www.semanticscholar.org/paper/a07cc3110dc2f48548f0a08d78036fcfc8278ab4)


### Heterozygosity for neurodevelopmental disorder-associated TRIO variants yields distinct deficits in behavior, neuronal development, and synaptic transmission in mice

**Why Not Relevant**: The provided paper does not mention CNTN5 and contains no data, analysis, or discussion about CNTN5 or any diseases directly attributed to CNTN5. Instead, the paper is focused on the TRIO gene and TRIO variants. For example, the paper opens with: "Genetic variants in TRIO are associated with neurodevelopmental disorders (NDDs) including schizophrenia (SCZ), autism spectrum disorder (ASD) and intellectual disability." It further specifies the variants and mechanisms studied: "Here, we investigate how heterozygosity for NDD-associated Trio variants – +/K1431M (ASD), +/K1918X (SCZ), and +/M2145T (bipolar disorder, BPD) – impact mouse behavior, brain development, and synapse structure and function." The paper describes TRIO GEF domain activities and Rac1/Tiam1-related mechanisms (e.g., "The TRIO K1431M GEF1 domain has impaired ability to promote GTP exchange on Rac1, but +/K1431M mice exhibit increased Rac1 activity, associated with increased levels of the Rac1 GEF Tiam1."), and reports on mouse phenotypes and synaptic physiology. None of this provides direct evidence that CNTN5 is associated with any Mendelian disease, nor mechanistic evidence linking CNTN5 to Mendelian disorders. Because there is no mention of CNTN5 and no transferable mechanistic link presented in the paper, it cannot be used to support or refute the claim that "CNTN5 is likely associated with at least one Mendelian disease." Any inference about CNTN5 from this TRIO-focused study would be speculative and unsupported by the paper's content.


[Read Paper](https://www.semanticscholar.org/paper/0d2ecf7372a56b7b47786a8c6fa3ddd784e94ee0)


### Mosaic RAI1 variant in a Smith–Magenis syndrome patient with total anomalous pulmonary venous return

**Why Not Relevant**: The provided paper excerpt exclusively addresses Smith–Magenis syndrome (SMS) and genetic findings for the RAI1 gene (including a de novo mosaic truncating RAI1 variant). There is no mention of CNTN5 anywhere in the text excerpt, no data (genetic, clinical, or functional) concerning CNTN5, and no mechanistic discussion that connects RAI1 to CNTN5. The evidence presented (high-coverage trio exome sequencing identifying a mosaic RAI1 c.94C>T p.Q32* variant present in 32% of reads, clinical phenotype consistent with SMS, and discussion of 17p11.2 deletions and RAI1 haploinsufficiency) is strong for implicating RAI1 in a Mendelian disorder but is entirely unrelated to the claim about CNTN5. Because the paper does not evaluate or mention CNTN5, it provides neither direct genetic evidence (e.g., pathogenic variants, segregation, case reports) nor mechanistic evidence (e.g., functional studies, pathway linkage) supporting or refuting an association between CNTN5 and any Mendelian disease. Therefore the relevance to the claim that “CNTN5 is likely associated with at least one Mendelian disease” is effectively zero. If you want an evidence-based assessment of CNTN5 specifically, I can search and analyze publications, ClinVar/OMIM entries, and functional studies that directly address CNTN5 variants and disease associations.


[Read Paper](https://www.semanticscholar.org/paper/c0698b67528bb20f9dc24552db3e063eaef186d0)


### Identifying Rare Genetic Variation In Obsessive-Compulsive Disorder

**Why Not Relevant**: This paper does not provide evidence for or against the claim that CNTN5 is likely associated with at least one Mendelian disease. The content is a thesis abstract on rare genetic variation in obsessive-compulsive disorder (OCD) that reports analyses of post-zygotic single-nucleotide variants (PZVs) and copy-number variants (CNVs) in OCD and control trios. Nowhere in the provided text is the gene symbol CNTN5 mentioned, nor are any specific genes (by name) reported as being implicated by the observed CNVs or PZVs. The paper gives summary statistics (rates of PZVs per base pair, rate ratios, p-values; proportions of children with rare deletions or duplications) and general statements about using CNVs to identify OCD risk genes in the future, but it does not report gene-level findings, variant-level evidence, mechanistic data related to CNTN5, or any diagnostic/Mendelian disease associations. Therefore there is no direct evidence (no variants in CNTN5 reported, no disease association described) and no mechanistic evidence (no functional or pathway data involving CNTN5) in this paper to support the claim.

Limitations and caveats relevant to this assessment:
- Absence of mention of CNTN5 in this document does not prove CNTN5 has no Mendelian associations; it only shows this particular paper does not address CNTN5. 
- The paper's finding of an increased rate of de novo deletions in OCD cases suggests that gene-level analyses of CNV contents could implicate specific genes, but the authors explicitly state further work is needed and do not present such gene-level results here. 

If you want evidence about CNTN5 specifically, I recommend querying gene-disease databases (OMIM, ClinVar, Orphanet), searching peer-reviewed literature for CNTN5-associated variants, and checking large aggregate variant resources (gnomAD, ClinGen) for curated disease relationships.


[Read Paper](https://www.semanticscholar.org/paper/6836f39aa067f957e1211be9fdfcd02b590e97cc)


### #4317 CLINICAL DIVERSITY OF STEROID-RESISTANT NEPHROTIC SYNDROME CAUSED BY TRPC6 MUTATIONS

**Why Not Relevant**: The paper focuses on steroid-resistant nephrotic syndrome (SRNS) and reports pathogenic variants in SRNS-related genes (e.g., TRPC6, NUP107, PLCE1, COL4A3, COL4A5) identified in a cohort of 39 Japanese SRNS patients. CNTN5 is not mentioned anywhere in the provided content. The absence of CNTN5 in this disease-specific cohort and gene list does not constitute evidence that CNTN5 is not associated with any Mendelian disease; it only shows that CNTN5 was not implicated in this particular SRNS study. Additional reasons this paper is not relevant to the claim include: (1) narrow disease focus (SRNS) — CNTN5 might cause a different Mendelian disorder outside SRNS; (2) cohort size and ascertainment — a small, specific cohort may not include individuals with CNTN5-related disease; (3) methodological filtering — the authors reported only variants meeting ACMG/ClinVar pathogenic criteria, so novel CNTN5 pathogenic variants could have been excluded; (4) no functional or mechanistic discussion of CNTN5 is present. Because the paper provides neither direct evidence (reported pathogenic CNTN5 variants) nor mechanistic arguments regarding CNTN5 function or disease linkage, its relevance to the claim that CNTN5 is likely associated with at least one Mendelian disease is negligible.


[Read Paper](https://www.semanticscholar.org/paper/c2747aaff3d6f0c6e29493cdff73cf48fd818d22)


### #3320 GENETIC TESTING FOR MOLECULAR DIAGNOSIS OF NEPHROLITHIASIS AND NEPHROCALCINOSIS IN ADULT PATIENTS: A SINGLE-CENTER COHORT

**Why Not Relevant**: The provided paper focuses on genetic causes of nephrolithiasis/nephrocalcinosis and reports results from a targeted 45-gene nephrolithiasis panel. The paper does not mention CNTN5 anywhere in the supplied text, nor does it list CNTN5 among genes with pathogenic, likely pathogenic, or VUS findings. Therefore the paper provides no direct evidence that CNTN5 is associated with any Mendelian disease. The only indirect pieces of information relevant to assessing CNTN5 are (1) that a 45-gene panel was used (but the exact gene list is not shown), and (2) the list of genes where pathogenic variants were detected—CNTN5 is absent from that list. Limitations preventing a stronger conclusion: (a) the paper does not provide the full list of the 45 genes included in the panel, so CNTN5 may not have been assayed; (b) the cohort is small (35 patients) and focused on kidney stone phenotypes, so even if CNTN5 is a Mendelian disease gene for a non-kidney phenotype it would be unlikely to appear here; (c) absence of evidence in this study is not evidence of absence—this single disease-focused cohort cannot meaningfully support or refute a broader claim about CNTN5 being a Mendelian disease gene. To evaluate the claim properly, one would need studies that (a) specifically assay CNTN5 for rare damaging variants in multiple affected individuals with a consistent phenotype, (b) provide segregation or functional evidence, or (c) authoritative gene-disease curation resources (e.g., ClinGen) that evaluate CNTN5 for Mendelian disease associations. Because none of those are present here, I set relevance very low.


[Read Paper](https://www.semanticscholar.org/paper/2097cfd6ee6b8857428d8e8895413812313205a9)


## Search Queries Used

- CNTN5 contactin 5 mutation case report Mendelian disease monogenic autosomal dominant autosomal recessive clinical phenotype

- CNTN5 ClinVar OMIM pathogenic likely pathogenic loss of function rare variant exome sequencing clinical genetics database annotation

- CNTN5 contactin 5 knockout mouse model phenotype neuronal development synapse behavior autism intellectual disability model organism

- CNTN5 de novo variant trio exome autism spectrum disorder developmental delay copy number variation deletion duplication contiguous gene syndrome contactin family


## Bibliometric Configuration

- Use Bibliometrics: True

- Author Impact Weight: 0.4

- Citation Impact Weight: 0.4

- Venue Impact Weight: 0.2
