## Supplementary material for "MENDELSEEK: An algorithm that predicts Mendelian Genes and elucidates what makes them special": COL6A5.txt

# Claim: The gene COL6A5 is likely associated with at least one Mendelian disease.

**Status**: processed

**Overall Rating**: 3

**Explanation**:

Supporting evidence.
The provided paper documents that mutations in collagen VI genes cause Mendelian muscle diseases (Bethlem myopathy and Ullrich congenital muscular dystrophy) and that loss of Col6a1 in mice produces a myopathic phenotype with clear pathophysiological mechanisms (mitochondrial dysfunction, apoptosis, autophagy defects). Collagen VI is described as a major extracellular matrix protein composed of distinct alpha chains encoded by separate genes, and the authors show that lack of one collagen VI chain (Col6a1) produces a robust disease phenotype in mice that mirrors human collagen VI–related myopathies. These points establish that genes encoding collagen VI chains can be bona fide Mendelian disease genes and that disruption of collagen VI has strong mechanistic plausibility for causing inherited disease.

Caveats or contradictory evidence.
The paper excerpts refer specifically to α1, α2 and α3(VI) chains (encoded by COL6A1, COL6A2, COL6A3) and experimental work is on Col6a1 null mice. The excerpts do not mention COL6A5 at all, nor provide human genetic data linking COL6A5 variants to Mendelian disorders. Thus, while the collagen VI family as a whole includes disease-associated members, there is no direct evidence in the provided material that COL6A5 is mutated in, or responsible for, any Mendelian disease. The paper’s relevance to COL6A5 is therefore indirect and limited.

Analysis of potential underlying mechanisms, deeper relationships, or second-order implications.
Because collagen VI is a multi-subunit protein and disease-causing mutations have been established in specific subunits (COL6A1–3), it is biologically plausible that other subunit genes could, if they encode functional chains incorporated into the collagen VI network or affect assembly, likewise cause disease when disrupted. However, subunit-specific expression patterns, redundancy, and whether a given alpha chain is incorporated into functionally essential heterotrimers (or has tissue-specific roles) determine whether loss-of-function or dominant-negative variants would produce a Mendelian phenotype. Without data on COL6A5 expression, protein incorporation into collagen VI assemblies, or human/animal phenotypes from COL6A5 perturbation, mechanistic plausibility alone is insufficient to conclude COL6A5 is a Mendelian disease gene.

Which rating is most appropriate.
The evidence supports the general principle that collagen VI genes can be Mendelian disease genes, but the provided paper contains no direct evidence implicating COL6A5 specifically. Given the plausible link by family homology but absence of direct genetic, clinical, or functional data for COL6A5 in the excerpts, the appropriate conclusion is that evidence is equivocal: there is a plausible basis for the claim but no direct support. Therefore the best-fitting rating is "Mixed Evidence."


**Final Reasoning**:

I reviewed the logic and evidence: the paper robustly supports that mutations in some collagen VI genes (notably COL6A1–3) cause Mendelian muscular disorders and that loss of collagen VI function produces disease-relevant cellular defects. However, the excerpts do not mention COL6A5, present no human genetic associations for COL6A5, and show no experimental perturbation of COL6A5. Homology to disease-causing family members makes COL6A5 a plausible candidate, but plausibility is not evidence of causation. No contradictory evidence was presented, only a lack of direct evidence. Balancing the indirect supportive context with the absence of any direct link to COL6A5, the correct assessment is that the evidence is mixed/insufficient to confidently assert that COL6A5 is associated with a Mendelian disease. I therefore reaffirm the rating "Mixed Evidence."


## Relevant Papers


### Extracellular collagen type VI has prosurvival and autophagy instructive properties in mouse embryonic fibroblasts

**Authors**: S. Castagnaro (H-index: 9)

**Relevance**: 0.2

**Bibliometric Impact**: 0.072


**Excerpts**:

- Collagen VI is a major protein of the extracellular matrix with a broad distribution in many tissues, including skeletal muscle and connective tissues.

- It is composed of three distinct alpha chains, α1, α2 and α3(VI), ecoded by separate genes.

- Mutations of collagen VI genes in humans cause several muscle diseases, such as Bethlem myopathy and Ullrich congenital muscular dystrophy.

- Collagen VI null (Col6a1–/–) mice display a myopathic phenotype characterized by mitochondrial dysfunction, spontaneous apoptosis and autophagic impairments in myofibers.

- To assess how lack of collagen VI impacts on fibroblast functions, I generated stable mouse embryonic fibroblast (MEF) lines from wild type and Col6a1–/– mice and showed that collagen VI is necessary for autophagy regulation and has prosurvival properties in fibroblasts.

- Col6a1–/– MEFs displayed accumulation of LC3 both basally and following autophagy induction.

- I found that in Col6a1–/– MEFs the mTORC1 downstream targets, such as 4E-BP1 and S6, are persistently activated under nutrient depletion stimuli, leading to autophagy inhibition in starving cells.

- Furthermore, Col6a1–/– MEFs presented a general energy imbalance, leading to over-activation of the AMP-activated protein kinase.

- These signalling defects lead to massive accumulation of autophagosomes inside Col6a1–/– fibroblasts, due to a compromised autophagosome-lysosome fusion in association with the presence of enlarged lysosomes and LAMP-2 protein depletion.

- These lysosomal defects are also associated with aberrant localization and activity of TFEB, a master transcription factor for lysosome biogenesis and autophagy regulation.

- This phenotype was specifically rescued by culturing cells onto purified collagen VI provided as an adhesive substrate .

- Mitochondria of Col6a1–/– MEFs exhibited increased fragmented morphology, associated with Parkin translocation and defective mitophagy.

- These findings show that fibroblasts play a relevant role in the development of the pathophysiological defects of collagen VI null mice, a finding that provide a thus far undisclosed and valuable information for the diagnosis and therapy of inherited diseases associated with collagen VI gene mutations.

- Moreover, they reveal for the first time a direct effect of collagen VI on the regulation of autophagy and associated mechanisms in this cell type.


**Explanations**:

- This sentence establishes that collagen VI is widely expressed in tissues relevant to Mendelian disorders (skeletal muscle, connective tissues). Relation to claim: mechanistic/biological plausibility. Strengths: highlights target tissues where pathogenic mutations could cause disease. Limitations: does not mention specific genes (e.g., COL6A5) or genetic association evidence.

- This sentence states the composition of collagen VI as three alpha chains (α1, α2, α3) encoded by separate genes. Relation to claim: contextual/mechanistic. Strengths: clarifies which collagen VI chains are discussed in the paper. Limitations: as written, it implies three chains (α1–α3); it does not mention α4–α6 or COL6A5, and thus provides no direct support for COL6A5 involvement. (Note: the excerpt contains the original misspelling "ecoded".)

- This sentence gives direct human-disease context: mutations in collagen VI genes cause Bethlem myopathy and Ullrich congenital muscular dystrophy. Relation to claim: partial direct support for the general proposition that collagen VI genes can be Mendelian disease genes. Strengths: explicit link between collagen VI gene mutations and inherited muscle diseases. Limitations: the sentence does not specify which collagen VI genes (e.g., COL6A1/2/3 versus COL6A5) are implicated, so it is not direct evidence for COL6A5 specifically.

- This describes the phenotype of Col6a1 knockout mice, showing that loss of a collagen VI gene causes a myopathic phenotype with clear cellular defects. Relation to claim: mechanistic and organismal evidence that loss of collagen VI function causes disease phenotypes in vivo. Strengths: in vivo functional evidence connecting a collagen VI gene knockout to pathology. Limitations: this is mouse Col6a1, not human COL6A5; species and gene differences limit direct inference to COL6A5 as a Mendelian disease gene.

- This sentence reports an experimental system (wild type vs Col6a1–/– MEFs) and the conclusion that collagen VI is necessary for autophagy regulation and has prosurvival properties in fibroblasts. Relation to claim: mechanistic evidence that collagen VI function is critical for cell physiology and that its disruption can produce cellular pathology relevant to inherited disease. Strengths: direct cellular functional data. Limitations: focuses on Col6a1 in mouse fibroblasts; no data on COL6A5 sequence variation or genetic disease association in humans.

- This reports accumulation of LC3 in Col6a1–/– MEFs, indicating altered autophagy. Relation to claim: mechanistic evidence (autophagy impairment) that loss of a collagen VI gene disrupts cellular homeostasis, supporting plausibility that mutations could be pathogenic. Strengths: molecular marker (LC3) measured; direct cellular phenotype. Limitations: again, Col6a1 in mouse cells; not evidence for COL6A5 mutations in human Mendelian disease.

- This provides mechanistic detail that mTORC1 downstream targets are persistently activated in Col6a1–/– MEFs under nutrient depletion, leading to autophagy inhibition. Relation to claim: mechanistic pathway evidence linking collagen VI deficiency to disrupted nutrient-sensing and autophagy, which can be disease-relevant. Strengths: identifies specific, conserved signaling components (mTORC1, 4E-BP1, S6) affected by collagen VI loss. Limitations: pertains to Col6a1 knockout cells; no genetic association to COL6A5 is provided.

- This states energy imbalance and AMPK over-activation in Col6a1–/– MEFs. Relation to claim: mechanistic—provides another pathway by which collagen VI deficiency leads to cellular dysfunction that could underlie disease. Strengths: adds to mechanistic plausibility. Limitations: Col6a1 mouse model only; not evidence for COL6A5 involvement in human Mendelian disease.

- This links signaling defects to autophagosome accumulation, compromised autophagosome-lysosome fusion, enlarged lysosomes and LAMP-2 depletion in Col6a1–/– fibroblasts. Relation to claim: mechanistic—describes downstream cellular defects that could mediate pathology when collagen VI is deficient. Strengths: identifies lysosomal/autophagic defects with specific markers. Limitations: experimental system is Col6a1 knockout mouse fibroblasts; does not demonstrate that COL6A5 variants cause similar defects in humans.

- This ties lysosomal defects to aberrant TFEB localization and activity. Relation to claim: mechanistic—TFEB is a master regulator of lysosome biogenesis, so this finding strengthens the biological plausibility that collagen VI disruption can produce broad lysosomal/autophagy pathology relevant to disease. Strengths: connects collagen VI loss to transcriptional regulation of lysosome/autophagy pathways. Limitations: again, Col6a1 mouse-cell data; no direct human genetic evidence for COL6A5.

- This reports mitochondrial fragmentation with Parkin translocation and defective mitophagy in Col6a1–/– MEFs. Relation to claim: mechanistic—shows mitochondrial quality-control defects resulting from collagen VI loss, a plausible contributor to inherited myopathies. Strengths: provides another concrete cellular mechanism linking collagen VI deficiency to cell death and disease-relevant pathways. Limitations: model is Col6a1 knockout in mouse fibroblasts; no mention of COL6A5 sequencing or human disease causality.

- This sentence explicitly connects the fibroblast findings in collagen VI null mice to "inherited diseases associated with collagen VI gene mutations" and suggests diagnostic/therapeutic relevance. Relation to claim: indirect support—authors assert that their findings inform inherited collagen VI gene disorders. Strengths: authors draw a link to human inherited diseases caused by collagen VI gene mutations. Limitations: this is a general statement about collagen VI genes; it does not provide specific evidence implicating COL6A5 as a Mendelian disease gene.

- This concluding sentence claims a "direct effect of collagen VI on the regulation of autophagy" in fibroblasts. Relation to claim: mechanistic evidence reinforcing that collagen VI dysfunction can produce cellular phenotypes plausibly underlying Mendelian disease. Strengths: summarizes a novel mechanistic role for collagen VI. Limitations: demonstrated for Col6a1 loss in mouse cells; does not provide direct human genetic association data for COL6A5.


[Read Paper](https://www.semanticscholar.org/paper/1bb82cec187370aacf23e4b5489c1c819df6d76c)


## Other Reviewed Papers


### Collagen type VI myopathies.

**Why Not Relevant**: The provided paper content states: "This review highlights the four recognized clinical phenotypes of collagen VI related - myopathies; Ullrich congenital muscular dystrophy, Bethlem myopathy, autosomal dominant limb-girdle muscular Dystrophy phenotype and autosomal recessive myosclerosis, and discusses the diagnostic criteria of these disorders." However, it does not mention the gene COL6A5, report any mutations, describe any patient genotypes, or present mechanistic data specific to COL6A5. While the content establishes that disorders are related to "collagen VI," it gives no gene-level attribution or evidence linking COL6A5 to any Mendelian disease. Therefore the paper content as provided gives neither direct evidence (e.g., mutation–disease associations, segregation, case reports) nor mechanistic evidence (e.g., expression, functional studies, protein interactions) that would support or refute the claim that COL6A5 is likely associated with at least one Mendelian disease. Limitations: the excerpt is brief and high-level, addresses collagen VI-related phenotypes in general, and lacks any gene names, experimental methods, statistical results, or contextual detail needed to evaluate the COL6A5-specific claim. Because of this absence of COL6A5-specific information, the content is not relevant to the claim.


[Read Paper](https://www.semanticscholar.org/paper/08dd5078e4a3f27334e70d3f927530203d1a340b)


### Causative variant profile of collagen VI-related dystrophy in Japan

**Why Not Relevant**: The provided content does not mention COL6A5 or present any variant, genotype–phenotype, segregation, functional, or mechanistic data specific to COL6A5. It only states that causative variant profiles were reported for a set of collagen VI-related dystrophy cases, without naming the genes involved. Because the claim concerns whether COL6A5 is likely associated with at least one Mendelian disease, and the excerpt contains no reference to COL6A5 nor to evidence that would allow inference about that specific gene, the paper content as provided is effectively non-informative with respect to the claim.


[Read Paper](https://www.semanticscholar.org/paper/e861ebbba9a5af8d478119d3ad09e04258ae36ef)


### Novel LSS variants in alopecia and intellectual disability syndrome: New case report and clinical spectrum of LSS‐related rare disease traits

**Why Not Relevant**: The provided paper content exclusively discusses the gene LSS, its biallelic pathogenic variants, and LSS-associated Mendelian phenotypes (congenital cataract type 44, autosomal recessive hypotrichosis type 14, and APMR4). There is no mention of COL6A5, no data concerning variants in COL6A5, nor any mechanistic linkage described between LSS and COL6A5. Therefore this paper provides no direct evidence that COL6A5 is associated with any Mendelian disease, nor mechanistic evidence that would make such an association plausible. To evaluate the claim about COL6A5, sources that explicitly study COL6A5 (e.g., gene-disease curation resources such as OMIM, ClinGen, ClinVar, PubMed reports of COL6A5 variants with segregation/functional data) should be consulted. The absence of COL6A5 from this study is the primary reason for irrelevance.


[Read Paper](https://www.semanticscholar.org/paper/fe90a5bb55278bc80588741eed5def6cf4f1aa82)


### Ullrich Congenital Dystrophy and Bethlem Myopathy: Current Knowledge on the Clinical Spectrum, Pathogenesis, and Future Therapeutic Avenues of Collagen VI Related Muscular Dystrophies

**Why Not Relevant**: The paper is a 2009 review focused on collagen VI–related muscular dystrophies and explicitly identifies the three α-chains and encoding genes as COL6A1, COL6A2, and COL6A3. It provides no mention, data, or discussion of COL6A5. Therefore it contains no direct evidence that COL6A5 is associated with any Mendelian disease. The content offers mechanistic context for how mutations in collagen VI genes (the three genes listed) cause Mendelian disorders, but this is indirect with respect to COL6A5. Absence of COL6A5 in this review is weak negative/neutral evidence (absence of evidence is not evidence of absence), and the review's publication date (2009) means it may not reflect later gene discoveries or associations.


[Read Paper](https://www.semanticscholar.org/paper/be09fb8cf7d0dddb65055b30d558dd47d2f2c873)


### [A rare variant in the TTR gene (p.E112K) is associated with systemic amyloidosis and a new symptom - skin hyperemia in response to ethanol intake: family segregation analysis, literature review, and a clinical case. Case report].

**Why Not Relevant**: This paper provides no direct or mechanistic evidence relevant to the claim that COL6A5 is likely associated with a Mendelian disease. The manuscript content is exclusively about transthyretin (TTR) and transthyretin amyloidosis. For example, the paper states: "Transthyretin amyloidosis (ATTR-amyloidosis) is a systemic disorder associated with extracellular deposition in the tissues and organs of amyloid fibrils, transthyretin-containing insoluble protein-polysaccharide complexes." It also states: "To date, more than 140 variants in the TTR gene have been identified; however, most of them are described in single patients and do not have clear evidence of pathogenicity." And: "This article presents a clinical case of ATTRv-amyloidosis associated with a rare pathogenic variant in the TTR gene and a newly described skin symptom." These verbatim sentences demonstrate the paper's focus on TTR gene variants and ATTR-amyloidosis. Nowhere in the provided content is COL6A5 (or the collagen VI gene family) mentioned, nor are there any data, analyses, or mechanistic descriptions that involve COL6A5, its variants, protein function, pathogenic mechanisms, segregation with disease, functional assays, population frequency data, or clinical phenotypes relevant to collagen VI-related disorders. Because the claim concerns COL6A5 and this paper deals only with TTR, there is no basis in this document to support or refute the claim. Limitations and caveats: absence of evidence in this single paper is not evidence of absence for COL6A5 associations generally — it simply means this paper is not a relevant source. To evaluate the claim about COL6A5 one should consult dedicated genetic-disease resources (OMIM, ClinVar, HGMD), primary literature on COL6A5, and large-scale sequencing/phenotype databases, none of which are provided or cited in the supplied text.


[Read Paper](https://www.semanticscholar.org/paper/2567df8ee078bbbd995e907e0465d6337ffeeb26)


### Co-Inheritance of Pathogenic Variants in PKD1 and PKD2 Genes Determined by Parental Segregation and De Novo Origin: A Case Report

**Why Not Relevant**: The provided paper content exclusively discusses autosomal dominant polycystic kidney disease (ADPKD) and variants in PKD1 and PKD2 genes as determinants of phenotypic variability. There is no mention of COL6A5, collagen VI genes, or any data linking COL6A5 to a Mendelian disease. Therefore the excerpt contains neither direct evidence (genetic association, segregation, case reports, statistics) nor mechanistic evidence (biological pathways, functional studies) relevant to the claim that COL6A5 is likely associated with at least one Mendelian disease. 

Limitations and caveats: This assessment is based solely on the content you provided. If the full paper contains additional sections, supplementary data, or other mentions of COL6A5 that were not included here, those could change the relevance. However, within the supplied text there are no sentences or paragraphs that support or refute the claim, so relevance is effectively zero.


[Read Paper](https://www.semanticscholar.org/paper/6bc1fefe3972092c6b64b12c8c0733ffcbf974ec)


### Frequency and phenotype associations of rare variants in five monogenic cerebral small vessel disease genes in 200,000 UK Biobank participants with whole exome sequencing data

**Why Not Relevant**: The provided paper content does not mention COL6A5 anywhere and explicitly states which genes were investigated (CTSA, TREX1, HTRA1, COL4A1/2). Because COL6A5 was not part of the systematic review, ClinVar search, or association analyses reported, the paper provides no direct evidence (positive or negative) about COL6A5 being associated with any Mendelian disease. Mechanistic evidence is also absent: the paper focuses on cerebral small vessel disease genes (including COL4A1/2, members of the type IV collagen family) and their penetrance/associations in UK Biobank, whereas COL6A5 is a different collagen family (collagen VI alpha-5) with distinct biology. General findings about penetrance, overestimated pathogenicity, and limits of population cohort analyses do not constitute evidence for COL6A5; at best they are methodological context about how population-based studies may under-detect gene-phenotype associations. Therefore the content is essentially non-informative with respect to the claim that COL6A5 is likely associated with at least one Mendelian disease.


[Read Paper](https://www.semanticscholar.org/paper/1f0de3de50f290aae016ea35325c46c15e4af3e8)


### Case report: A novel mutation of the CAPN3 gene in a Chinese family with limb-girdle muscular dystrophy type 2A

**Why Not Relevant**: The paper exclusively studies LGMD2A/R1 and genetic variation in CAPN3. There is no mention of the gene COL6A5, no data on COL6A5 sequence variants, expression, function, or pathological mechanisms, and no discussion that would link COL6A5 to the phenotype under study. Therefore the paper provides no direct genetic evidence or mechanistic rationale supporting the claim that COL6A5 is likely associated with at least one Mendelian disease. Note: absence of any COL6A5 data in this paper does not disprove that COL6A5 could be involved in a Mendelian disease elsewhere; it simply means this paper is not relevant to that claim.


[Read Paper](https://www.semanticscholar.org/paper/e358d75a5baa346a5fb9b7b5f9b716f439e505a1)


### Exome-by-phenome-wide rare variant gene burden association with electronic health record phenotypes

**Why Not Relevant**: The provided paper content (abstract) does not mention COL6A5 anywhere in the methods, results, or conclusions. The study describes an exome-by-phenome PheWAS approach that detected 106 gene-burden associations (and 12 robust associations), and it lists specific genes identified (e.g., TTN, MYBPC3, CFTR, BRCA1/2, TGM6). Because COL6A5 is not listed among the discovered or validated genes in the text supplied, the paper provides no direct evidence that COL6A5 is associated with a Mendelian disease. The methodological descriptions (WES in N=11,451, pLOF MAF ≤ 0.1% collapsing, complementary missense analyses, replication in UKBB) are relevant context showing that the study was designed to detect Mendelian-like gene associations driven by rare pLOF variants. However, absence of COL6A5 in the reported findings does not constitute proof that COL6A5 is not associated with any Mendelian disease: limitations include modest discovery cohort size for very rare disorders, reliance on EHR-derived phenotypes that may not capture all Mendelian presentations, focus on pLOF (which would miss non–loss-of-function mechanisms), allele frequency/annotation thresholds that may exclude some pathogenic variants, and potential ancestry/phenotype differences between PMBB and UKBB. In summary, the paper does not provide supportive evidence for the claim (no direct mention or positive association for COL6A5), nor does it provide gene-specific mechanistic data for COL6A5; therefore its relevance to the claim is very low.


[Read Paper](https://www.semanticscholar.org/paper/4f7e7da3430a98395e1a8212d10130271c5eb764)


### Phenotypic Variability in Camurati–Engelmann Disease: A Case Report of a Family with the c.653G>A Pathogenic Variant in the TGFB1 Gene

**Why Not Relevant**: Relevance is near zero because the paper exclusively reports on Camurati–Engelmann Disease (CED) caused by heterozygous mutations in TGFB1 and describes a TGFB1 missense variant (c.653G>A, p.Arg218Cys) segregating with disease in a family. There is no mention of the gene COL6A5, no genetic data relating COL6A5 to the phenotype, and no mechanistic discussion linking COL6A5 to TGFB1, bone metabolism, or CED. Therefore the paper provides neither direct evidence (no observations, variants, segregation, linkage, or association data for COL6A5) nor mechanistic evidence (no biochemical, expression, pathway, or interaction data implicating COL6A5) relevant to the claim that COL6A5 is likely associated with at least one Mendelian disease. Absence of evidence in this paper is not proof that COL6A5 is not disease-associated in other studies, but this article cannot be used to support the claim. If you want, I can (a) search the literature, ClinVar, OMIM, and gene-disease databases for COL6A5 associations, or (b) evaluate other papers for direct or mechanistic evidence regarding COL6A5 and Mendelian disease.


[Read Paper](https://www.semanticscholar.org/paper/2d74d8dc9fa5f2b70a1dd80faea19ca419b8631c)


### Whole‐exome sequencing analyses in a Saudi Ischemic Stroke Cohort reveal association signals, and shows polygenic risk scores are related to Modified Rankin Scale Risk

**Why Not Relevant**: This paper excerpt does not provide evidence relevant to the claim that COL6A5 is likely associated with at least one Mendelian disease. The provided sentence reports a general result of gene-burden testing in a Saudi stroke cohort and an association between cumulative rare-variant burden in stroke genes and modified Rankin Scale (mRS) outcome. It (1) does not mention COL6A5 by name, (2) gives no gene-level results, (3) provides no variant-level, segregation, case-report, or functional/mechanistic data for COL6A5, and (4) focuses on stroke-related burden and clinical stratification rather than on identifying Mendelian disease genes. Absence of mention of COL6A5 in this excerpt is not evidence that COL6A5 is not Mendelian; it only means this paper excerpt does not address the claim. To assess the claim one would need gene-specific evidence such as case reports showing segregation of COL6A5 variants with disease, ClinVar/OMIM annotations, statistically significant enrichment of rare (likely pathogenic) variants in cases versus controls for COL6A5, functional assays demonstrating a pathogenic mechanism compatible with Mendelian inheritance, or animal models. None of those data are present in the excerpt, so the relevance to the COL6A5–Mendelian disease claim is negligible.


[Read Paper](https://www.semanticscholar.org/paper/ef1e3cc537505def74437c2048ef722316d024a6)


### Extracellular Matrix Perturbation in Down Syndrome Pathology

**Why Not Relevant**: The paper focuses on collagen type VI chains α1, α2 and α3 (COL6A1, COL6A2, COL6A3) and their perturbation in Down syndrome and extracellular matrix/hyaluronan biology. It contains no mention of COL6A5, no genetic linkage or mutation data implicating COL6A5 in any disorder, and no experimental assays targeting COL6A5 (protein, RNA, or genetic). Therefore it provides no direct evidence that COL6A5 is associated with any Mendelian disease, nor mechanistic evidence specific to COL6A5. The data concern trisomy 21 (a chromosomal aneuploidy) effects on COL6A1/2 and HA, which are not informative for evaluating whether COL6A5 causes Mendelian (single-gene) disease.


[Read Paper](https://www.semanticscholar.org/paper/ff0d99b10386f268e9f38f93fcff21a1c0cd0382)


### Pathogenetic Mechanisms of Collagen Type VI-Related Muscular Dystrophies

**Why Not Relevant**: The provided paper content does not mention COL6A5 at all. The study focuses on collagen type VI-related muscular dystrophies and on molecular pathways (e.g., TGFβ1) and transcriptomic/splicing changes in muscle biopsies from patients with established COL6RD, which are described as caused by mutations in the genes encoding the three α-chains of collagen VI. The methods reported (histology, microarray, RNA-Seq expression and splicing analyses) are aimed at characterizing disease mechanisms and expression changes rather than identifying or validating novel Mendelian gene-disease associations. There is no direct genetic evidence (e.g., variant identification, segregation, functional validation) implicating COL6A5 in a Mendelian disorder, nor is there any mention of altered COL6A5 expression or splicing in the excerpts provided. Therefore the paper provides no direct support for the claim, and only very weak indirect context (it studies collagen VI-related disease generally) that is insufficient to conclude that COL6A5 is likely associated with at least one Mendelian disease. Absence of mention in this paper is not proof that COL6A5 is not disease-associated, but this paper does not provide affirmative evidence for the claim.


[Read Paper](https://www.semanticscholar.org/paper/9eaef469b1a5b9c1bcd1b235888e06808ac02124)


## Search Queries Used

- COL6A5 mutation pathogenic Mendelian disease case report family segregation clinical phenotype

- COL6A5 variant exome sequencing rare variant burden gene discovery monogenic disease cohort trio

- COL6A5 collagen type VI alpha 5 chain extracellular matrix assembly interaction COL6A1 COL6A2 COL6A3 functional study expression tissue distribution muscle skin fibroblast

- collagen type VI review COL6A5 disease association Bethlem myopathy Ullrich congenital myopathy muscular dystrophy monogenic review


## Usage Statistics

- Prompt Tokens: 0

- Completion Tokens: 0

- Total Tokens: 0

- Estimated Cost: $0.0657


## Bibliometric Configuration

- Use Bibliometrics: True

- Author Impact Weight: 0.4

- Citation Impact Weight: 0.4

- Venue Impact Weight: 0.2
