## Supplementary material for "MENDELSEEK: An algorithm that predicts Mendelian Genes and elucidates what makes them special": COX7A2L.txt

# Claim: The gene COX7A2L is likely associated with at least one Mendelian disease.

**Status**: processed

**Overall Rating**: -1

**Explanation**:

No relevant papers were found that support or refute this claim.


## Relevant Papers


## Other Reviewed Papers


### Functional analysis of the APAF-1 binding protein FAM96A

**Why Not Relevant**: This paper provides no relevant evidence for the claim that COX7A2L is likely associated with at least one Mendelian disease. The manuscript excerpt exclusively discusses the gene/protein FAM96A (APAF-1 binding protein), its role in intrinsic apoptosis, interactions with Ciao1, effects on tumorigenesis (including GIST and other cancers), copy number losses, expression changes, and a conditional FAM96A knockout mouse. There is no mention of COX7A2L, no genetic-variant, segregation, case-report, cohort, or population-association data pertaining to COX7A2L, and no discussion of Mendelian disease phenotypes or inheritance patterns related to COX7A2L. The types of evidence required to support an association between a specific gene and a Mendelian disease (e.g., pathogenic variants in affected individuals, segregation in families, de novo occurrence, loss-of-function intolerance, functional rescue / loss-of-function modeling that reproduces a human phenotype) are absent for COX7A2L in this text. Although the paper includes mechanistic data (apoptosis pathways, Fe-S biogenesis, tumor suppressor behavior) and an animal knockout model for FAM96A, these findings are specific to FAM96A and cannot be extrapolated to COX7A2L without independent evidence. Limitations: because the provided content does not mention COX7A2L at all, neither direct evidence nor mechanistic plausibility relevant to COX7A2L can be extracted from this paper. Therefore the paper is not relevant to assessing the claim about COX7A2L and Mendelian disease.


[Read Paper](https://www.semanticscholar.org/paper/9884f9a50521510db057cc9d83f7245bc6da5c04)


## Search Queries Used

- COX7A2L SCAF1 cytochrome c oxidase subunit 7A2 like Mendelian disease monogenic hereditary single gene case report exome sequencing ClinVar OMIM pathogenic

- COX7A2L SCAF1 mitochondrial supercomplex assembly respiratory chain complex IV deficiency mitochondrial disease cardiomyopathy encephalopathy mitochondrial myopathy mechanism

- COX7A2L knockout mouse loss of function phenotype zebrafish Drosophila cell model respiratory chain assembly phenotype developmental phenotype

- COX7A2L variants population sequencing gnomAD constraint loss of function intolerance pLI gene burden analysis rare disease cohort monogenic candidate gene


## Bibliometric Configuration

- Use Bibliometrics: True

- Author Impact Weight: 0.4

- Citation Impact Weight: 0.4

- Venue Impact Weight: 0.2
