## Supplementary material for "MENDELSEEK: An algorithm that predicts Mendelian Genes and elucidates what makes them special": CRHBP.txt

# Claim: The gene TBC1D32 is likely associated with at least one Mendelian disease.

**Status**: processed

**Overall Rating**: -1

**Explanation**:

No relevant papers were found that support or refute this claim.


## Relevant Papers


## Other Reviewed Papers


### A whole exome sequencing study to identify rare variants in multiplex families with alcohol use disorder

**Why Not Relevant**: Relevance is nil because the provided paper content does not mention the gene TBC1D32 at all and therefore provides no direct evidence for or against an association between TBC1D32 and any Mendelian disease. The study focuses on rare variants in families multiplex for alcohol use disorder (a complex trait), reporting lists of candidate genes (e.g., CDSN, CHRNA9, IFT43, TLR6, SELENBP1, GMPPB, ZNF514, OXGR1, DIEXF, TMX4, MTBP, PON2, CRHBP, ANKRD46, AGTRAP, PPA1) but TBC1D32 is absent. Additional reasons the paper cannot support the claim: (1) the phenotype studied is AUD, not Mendelian disorders; (2) the sample of probands is small (N=53), limiting power to detect rare Mendelian-effect variants genome-wide; (3) reported findings are candidate-level and described as follow-up priorities rather than confirmed disease-causing genes. Therefore, this paper neither provides mechanistic evidence implicating TBC1D32 nor direct genetic association evidence for TBC1D32 with any Mendelian disease.


[Read Paper](https://www.semanticscholar.org/paper/f8b51fc93e484d1aa91efb50bcbee3b173b3392c)


### Novel LSS variants in alopecia and intellectual disability syndrome: New case report and clinical spectrum of LSS‐related rare disease traits

**Why Not Relevant**: The provided paper content exclusively discusses pathogenic biallelic variants in the gene LSS and their association with multiple Mendelian phenotypes (congenital cataract type 44, autosomal recessive hypotrichosis type 14, and alopecia‐intellectual disability syndrome type 4 (APMR4)). It describes trio exome sequencing findings in a patient with novel compound heterozygous LSS variants, phenotypic features, and genotype–phenotype correlations for LSS. Nowhere in the supplied text is the gene TBC1D32 mentioned, nor are there mechanistic data, interaction data, or other evidence linking TBC1D32 to the discussed phenotypes. Therefore, this paper provides no direct evidence that TBC1D32 is associated with any Mendelian disease, nor does it provide mechanistic support for such an association. Note: absence of evidence in this paper does not prove absence of an association overall, but based solely on the supplied content there is no relevant evidence for the claim.


[Read Paper](https://www.semanticscholar.org/paper/fe90a5bb55278bc80588741eed5def6cf4f1aa82)


### Exome‐wide assessment of isolated biliary atresia: A report from the National Birth Defects Prevention Study using child–parent trios and a case–control design to identify novel rare variants

**Why Not Relevant**: The provided paper content focuses exclusively on biliary atresia (BA) and reports genes (PKD1L1, NOTCH2, IFRD2) implicated or worthy of further study in BA or related gastrointestinal conditions. The text does not mention TBC1D32 anywhere, nor does it present data or mechanistic discussion linking TBC1D32 to BA or any other Mendelian disease. Therefore the paper does not provide direct or mechanistic evidence relevant to the claim that TBC1D32 is likely associated with at least one Mendelian disease. Additional reasons for non-relevance: the study cohort and analyses are disease-specific (BA) and relatively small for discovering new Mendelian gene associations outside that phenotype, and the authors explicitly note complexity and diagnostic difficulty in BA, which further limits extrapolation to other gene–disease claims.


[Read Paper](https://www.semanticscholar.org/paper/e936442497b14b6fd0f27d7a747cd2dc7cb2eade)


### A Big Catch for Germ Cell Tumour Research

**Why Not Relevant**: This paper does not mention the gene TBC1D32 anywhere in the provided content. All direct human genetic and mechanistic evidence in the paper pertains to LRRC50 (lrrc50) and other genes (e.g., BMPR1B, DND1). While the paper provides examples and mechanistic rationale that ciliary genes can underlie Mendelian predisposition to seminoma (which offers indirect plausibility that other ciliary genes might also be disease-associated), it supplies no direct genetic, functional, or epidemiologic evidence linking TBC1D32 to any Mendelian disease. Therefore the article is only tangentially relevant: it supports the general concept that disruption of ciliary proteins can cause heritable disease phenotypes, but it does not provide any evidence specific to TBC1D32. Because the claim concerns association of TBC1D32 with a Mendelian disease, the absence of any mention or data on TBC1D32 makes the paper non-informative for confirming or refuting the claim.


[Read Paper](https://www.semanticscholar.org/paper/025daf36bbd4370c5b91188705cdfb1dee0e191c)


### Whole-exome sequencing of pathogenic genes in a family with congenital heart disease: A case report

**Why Not Relevant**: The provided paper content does not mention the gene TBC1D32 at any point. The study reports whole-exome sequencing findings in a single Chinese family with congenital heart disease and identifies variants in AMER1 and KCNE1 (and two additional unspecified variants). Because TBC1D32 is neither reported among the discovered variants nor discussed mechanistically, this paper provides no direct genetic, clinical, or mechanistic evidence supporting or refuting the claim that TBC1D32 is associated with at least one Mendelian disease. Absence of mention in this paper is not evidence that TBC1D32 is unrelated to Mendelian disease more broadly, but it does mean this paper cannot be used to support the claim. Additional studies that specifically report variants in TBC1D32 segregating with disease, functional validation, or curated disease-gene associations would be required to evaluate the claim.


[Read Paper](https://www.semanticscholar.org/paper/2afe5ddc8ce353c6339a8bb041306ea8d115518f)


### Whole exome sequencing identifies risk variants associated with intracranial epidermoid cyst deterioration: A case report

**Why Not Relevant**: The paper does not mention the gene TBC1D32 anywhere in the provided text. The whole-exome sequencing results and the study conclusions specifically report pathogenic variants in GJB2 and TLR2 only. Consequently, there is no direct genetic association reported between TBC1D32 and any Mendelian disease in this paper, nor is there mechanistic discussion relevant to TBC1D32. Additionally, this is a single-case report: even if TBC1D32 had been absent from the findings, the study's scope (one proband, focused on GJB2 and TLR2) would limit generalizability. Because the paper contains no data, analyses, or discussion of TBC1D32, its relevance to the claim that TBC1D32 is likely associated with at least one Mendelian disease is effectively zero.


[Read Paper](https://www.semanticscholar.org/paper/3e9a22f9edbb69594a1d1f950d5f730041afc821)


### Exome sequencing in over 63,000 patients in TIMI trials uncovers pathogenic cardiomyopathy variant carriers with high risk for heart failure and cardiovascular death

**Why Not Relevant**: The paper focuses specifically on rare pathogenic/likely pathogenic variants associated with dilated, hypertrophic, and arrhythmogenic right ventricular cardiomyopathies identified within five cardiovascular clinical trials. The text provides study objectives, methods, and results centered on known cardiomyopathy genes (for example, TTN) and clinical outcomes (HHF, CV death). Nowhere in the provided content is the gene TBC1D32 mentioned, nor are there mechanistic data, variant reports, or curated classifications pertaining to TBC1D32. Therefore this paper does not provide direct evidence for or against the claim that TBC1D32 is likely associated with at least one Mendelian disease. Important caveat: absence of mention in this cardiomyopathy-focused study does not constitute evidence that TBC1D32 is not associated with a Mendelian disease; it only shows that this paper did not assess or report on that gene. Assessing the claim would require literature and database reviews (ClinVar, OMIM, gene-disease catalogs) or primary studies that explicitly examine TBC1D32.


[Read Paper](https://www.semanticscholar.org/paper/f9e3adf19cb2bfdb8171744405eb1112137a3bb9)


### Descriptive analysis of MC4R gene variants associated with obesity listed on ClinVar

**Why Not Relevant**: The provided paper focuses exclusively on MC4R gene variants (objectives, methods, results and conclusions all explicitly concern MC4R). There is no mention of TBC1D32, no data about TBC1D32 variants, no mechanistic discussion involving TBC1D32, and no broader genomic or interactome analysis that would implicate TBC1D32. Therefore this paper provides no direct evidence that TBC1D32 is associated with any Mendelian disease, nor does it provide mechanistic pathways linking TBC1D32 to disease. Absence of mention in this MC4R-focused study is not disproof that TBC1D32 is disease-associated, but it means this paper cannot be used to support the claim.


[Read Paper](https://www.semanticscholar.org/paper/6bae624c1c06216b5458a99ef3b5e0a4b2d4cd16)


## Search Queries Used

- TBC1D32 variants exome sequencing Mendelian disease family study case report rare disease

- TBC1D32 ClinVar OMIM DECIPHER pathogenic likely pathogenic variant gene disease association

- TBC1D32 ciliogenesis hedgehog signaling primary cilium knockout mouse zebrafish developmental phenotype functional study

- TBC1D32 gnomAD population frequency loss of function constraint pLI missense z score congenital malformation ciliopathy phenotype


## Bibliometric Configuration

- Use Bibliometrics: True

- Author Impact Weight: 0.4

- Citation Impact Weight: 0.4

- Venue Impact Weight: 0.2
