## Supplementary material for "MENDELSEEK: An algorithm that predicts Mendelian Genes and elucidates what makes them special": CRTAC1.txt

# Claim: The gene CRTAC1 is likely associated with at least one Mendelian disease.

**Status**: processed

**Overall Rating**: -1

**Explanation**:

No relevant papers were found that support or refute this claim.


## Relevant Papers


## Other Reviewed Papers


### Whole exome sequencing analyses reveal gene–microbiota interactions in the context of IBD

**Why Not Relevant**: The provided paper content does not mention CRTAC1 at all and focuses on host genetic effects on the gut microbiome in IBD. The study reports mbQTLs in IL17REL, MYRF, SEC16A, WDR78 and rare PTV/CNV effects in CYP2D6, GPR151 and CD160, with interaction analyses confirming TNFSF15 findings—none of which are CRTAC1. Therefore the paper provides no direct evidence that CRTAC1 is associated with any Mendelian disease. Weak indirect information can be drawn from the study design (whole exome sequencing was performed), which indicates that coding variants could in principle have been detected; however, the study's aims (microbiome associations in IBD), cohort composition (IBD patients and population controls), statistical focuses (mbQTLs, PTV/CNV burdens with specific MAF thresholds), and lack of family-based or Mendelian-phenotype analyses mean it is not an appropriate dataset to evaluate whether CRTAC1 causes a Mendelian disorder. Consequently, the absence of CRTAC1 in the reported findings is not strong evidence against the claim. Key limitations that prevent drawing conclusions about CRTAC1 from this paper: (1) study outcome is microbiome traits in IBD, not Mendelian disease phenotypes; (2) sample size and allele-frequency-focused analyses may lack power to detect ultra-rare pathogenic variants typical of Mendelian diseases; (3) no family segregation or clinical-genetic diagnostic workup was performed; (4) exact exome coverage for CRTAC1 and variant-level QC/annotation details are not provided. Given these points, the paper is not relevant for supporting or refuting the claim that CRTAC1 is likely associated with at least one Mendelian disease.


[Read Paper](https://www.semanticscholar.org/paper/7cb9697cbccc67657a0950080ec81f182f541636)


### Multiple epiphyseal dysplasia and related disorders: Molecular genetics, disease mechanisms, and therapeutic avenues

**Why Not Relevant**: The provided paper content does not mention the gene CRTAC1 anywhere, so it contains no direct evidence that CRTAC1 is associated with a Mendelian disease. The text provides only high-level background about rare genetic skeletal diseases and multiple epiphyseal dysplasia (MED), noting that MED is caused by mutations in genes encoding cartilage extracellular matrix proteins, enzymes, and transporter proteins, and discussing ER stress as a unifying disease mechanism. While these general statements describe plausible mechanistic pathways (e.g., misfolded cartilage proteins causing ER stress) that could, in principle, apply to any cartilage-expressed gene, including CRTAC1, the paper presents no data, examples, variant descriptions, segregation analyses, functional experiments, or explicit mention of CRTAC1 to support or refute an association. Therefore the evidence in this paper for the claim is effectively absent. Limitations: because the excerpt is background-only and lacks gene-specific data, one cannot infer CRTAC1 involvement from the mechanistic discussion alone; establishing a likely Mendelian association would require gene-specific genetic evidence (pathogenic variants segregating with disease, de novo events, case-control enrichment), functional validation (demonstrating variant impact on protein function or relevant cellular phenotype), or curated clinical-genetic resources, none of which are present in the provided content. The excerpt pages are not available, so no page references can be given.


[Read Paper](https://www.semanticscholar.org/paper/a9be94d6f40e19547fb65c1ce011541d3f3c2926)


### Meta-Analysis and Systematic Review of the Genomics of Mucosal Melanoma

**Why Not Relevant**: This paper reports a comprehensive collated analysis of somatic genomic alterations in mucosal melanoma tumor samples. The claim concerns whether the gene CRTAC1 is likely associated with at least one Mendelian (germline, heritable) disease. The manuscript's focus is tumor somatic mutation, copy-number, and structural variation across cohorts of melanoma samples, not germline variant segregation, pedigree analysis, or clinical descriptions of inherited disorders. The provided text lists significantly mutated somatic driver genes in this cancer type and pathways implicated in tumorigenesis; CRTAC1 is not mentioned. Absence of CRTAC1 in a somatic tumor-driver study does not meaningfully support or refute CRTAC1 having a Mendelian-disease association because (1) Mendelian disease evidence requires germline, familial, or case-control data and functional segregation information, and (2) the targeted panels used may not have included CRTAC1 or the gene may not be relevant to this cancer type. Therefore, the paper is not a relevant source of direct or mechanistic evidence for the claim. For these reasons I rate relevance as 0.02 (near zero but non-zero because the paper does present gene lists and sequencing approaches that indirectly inform what was or was not found in this tumor-focused dataset).


[Read Paper](https://www.semanticscholar.org/paper/c6f8faea99cebc1523bedefa8d01b289d63e804c)


### The pathogenesis of osteoclast diseases: Some knowns, but still many unknowns

**Why Not Relevant**: This paper does not mention CRTAC1 (no symbol, gene name, or product) anywhere in the provided content. The manuscript focuses on genes known to cause osteoclast-related Mendelian disorders (examples given include TCIRG1, CLCN7, OSTM1, PLEKHM1, CA2, TNFRSF11A, TNFRSF11B, SQSTM1, VCP, LRP5, CTSK, etc.) and on mechanistic pathways relevant to those genes (acidification/vesicular trafficking in osteoclasts and perturbation of proteasomal degradation). Because CRTAC1 is not discussed, the paper provides neither direct evidence (no reports of mutations, genotype–phenotype associations, pedigree or case reports) nor mechanistic evidence (no description of CRTAC1 function, expression, or pathway involvement that could plausibly link it to a Mendelian disease). 

Limitations to this conclusion: absence of mention in this paper is not proof that CRTAC1 is not associated with any Mendelian disease. This is a single review (dated 2007) focused on osteoclast disorders; CRTAC1 could be associated with Mendelian disease outside the scope of this review or discovered after its publication. To evaluate CRTAC1’s association with Mendelian disease, a targeted search of primary literature, clinical databases (e.g., OMIM, ClinVar), and more recent reviews is required.


[Read Paper](https://www.semanticscholar.org/paper/72869b7d23fa551f350748c543749060964e9b02)


### The ups and downs of parting COMPany Poly‐aspartic acid tract instability: a novel expansion–contraction mutation

**Why Not Relevant**: The provided paper text does not mention the gene CRTAC1 anywhere and contains no data (genetic, clinical, or mechanistic) that links CRTAC1 to any Mendelian disease. The content focuses on the class of dynamic (trinucleotide repeat expansion/contraction) mutations and gives examples involving genes such as FMR1, the myotonin kinase gene, and COMP (with discussion of pseudoachondroplasia and multiple epiphyseal dysplasia). Although the paper discusses mechanisms by which repeat expansions cause Mendelian disorders (e.g., instability of trinucleotide repeats, location of repeats in 5'UTR/exon/intron/3'UTR, and an example of a contraction–expansion mutation in COMP), none of these mechanisms or examples involve CRTAC1. Therefore there is neither direct evidence (observations, case reports, genetic linkage/segregation, variant pathogenicity) nor mechanistic evidence (molecular function, relevant mutation types in CRTAC1, animal models) in this paper to support or refute the claim that CRTAC1 is associated with a Mendelian disease.

Limitations and caveats: absence of evidence in this single paper does not prove CRTAC1 is not associated with a Mendelian disease; it only shows this paper is not a source of relevant evidence. To assess the claim, searches of gene-disease databases (OMIM, ClinVar, HGMD), PubMed for CRTAC1 case reports, and population/constraint metrics (gnomAD) or functional studies would be required.


[Read Paper](https://www.semanticscholar.org/paper/c4d8424428568c274ac27f6e6bf8fcbfaa912906)


### Genetic copy number variants, cognition and psychosis: a meta-analysis and a family study

**Why Not Relevant**: The provided paper content discusses associations of high-risk CNVs with schizophrenia, other neurodevelopmental disorders, and cognitive impairments—topics relevant to complex, often multifactorial disease risk. The claim concerns CRTAC1 being likely associated with at least one Mendelian disease (a monogenic, typically highly penetrant disorder). The excerpt contains no reference to CRTAC1, no gene-specific findings, no mutation/variant-level evidence, and no discussion of Mendelian inheritance or monogenic disease causation. Although CNVs can sometimes cause Mendelian disorders if they delete or duplicate a single critical gene, the excerpt does not indicate which genes are involved in the CNVs studied, provides no mechanistic evidence tying CRTAC1 to disease, and gives no segregation or variant pathogenicity data required to support a Mendelian-disease association. Therefore the content cannot be used to support or refute the claim about CRTAC1; additional gene-level, variant-level, and inheritance/segregation evidence would be necessary to evaluate the claim.


[Read Paper](https://www.semanticscholar.org/paper/0b161d85a225172a0757d14026cbc4e0631eaf49)


### Uncovering recessive alleles in rare Mendelian disorders by genome sequencing of 174 individuals with monoallelic pathogenic variants.

**Why Not Relevant**: The provided paper content does not mention CRTAC1, any specific gene, any variant in CRTAC1, any phenotype attributable to CRTAC1, or any Mendelian disease. It is a single general statement about genome sequencing improving diagnostic yield when structural and intronic variants are interpreted. Because the claim concerns a specific gene–disease association (CRTAC1 and a Mendelian disease), evidence relevant to that claim would need to include: observed pathogenic or likely pathogenic variants in CRTAC1 segregating with disease, statistical enrichment of rare damaging CRTAC1 variants in affected individuals vs controls, functional/mechanistic data linking CRTAC1 perturbation to disease biology, clinical case reports, or curated database entries (e.g., OMIM) implicating CRTAC1 in a Mendelian disorder. None of those elements are present. The sentence is actionable only as a general rationale for broader sequencing analyses and does not provide direct or gene-specific mechanistic evidence. Consequently, it is insufficient to support or refute the claim, so the relevance is very low.


[Read Paper](https://www.semanticscholar.org/paper/5ee57a7874147eec316d4452073c0a496149a855)


### A novel compound heterozygous variant of ECEL1 induced joint dysfunction and cartilage degradation: a case report and literature review

**Why Not Relevant**: This paper does not mention CRTAC1 and provides no direct or mechanistic evidence linking CRTAC1 to any Mendelian disease. The study exclusively describes a case of distal arthrogryposis type 5D (DA5D) associated with ECEL1. Relevant verbatim sentences from the paper illustrating its focus on ECEL1 are:

"Background Distal arthrogryposis type 5D (DA5D) represents a subtype of distal arthrogryposis (DA) characterized by congenital joint contractures in the distal extremities. DA5D is inherited in a rare autosomal recessive manner and is associated with the ECEL1 gene." 

and

"Whole-exome sequencing of the patient’s DNA revealed a compound heterozygous mutation of c.2152-15C>A and c.110_155del in ECEL1." 

These sentences show the paper's findings concern ECEL1 mutations causing DA5D. No sentences or paragraphs in the provided content reference CRTAC1, no data (genetic, clinical, or mechanistic) connect CRTAC1 to the phenotype described, and no discussion suggests CRTAC1 as a candidate gene. Therefore the paper offers neither direct evidence (e.g., variants in CRTAC1 segregating with disease) nor mechanistic evidence (e.g., functional studies, pathway links) relevant to the claim that CRTAC1 is likely associated with a Mendelian disease. Additional limitations of the paper (single-case report, predictions from in silico tools) are irrelevant to CRTAC1 because CRTAC1 is not addressed at all in the manuscript.


[Read Paper](https://www.semanticscholar.org/paper/aa42762668cf8099d0e49c48118c5bafd8799879)


### Novel de novo heterozygous CACNA1A gene variant in generalised dystonia: a case report

**Why Not Relevant**: The provided paper discusses a single-case report of dystonia associated with a variant in CACNA1A and makes no mention of the gene CRTAC1 anywhere in the text. Therefore it contains no direct genetic evidence (variants, segregation, functional studies) implicating CRTAC1 in any Mendelian phenotype, nor any mechanistic discussion that would plausibly connect CRTAC1 to the disease described. The paper's scope is limited to CACNA1A in dystonia (single patient, de novo likely-pathogenic variant), so it neither supports nor refutes the claim about CRTAC1. Because CRTAC1 is not discussed, there are no experimental conditions, methodologies, results, or mechanistic data in this paper that affect the plausibility of CRTAC1 causing a Mendelian disease.


[Read Paper](https://www.semanticscholar.org/paper/7f35be37862ec1c423e0d5f5879486b9d93cc5c0)


### Studying the Molecular Mechanism of Post-Translational Modification of Lamin A in Hutchinson-Gilford Progeria Syndrome

**Why Not Relevant**: This paper does not mention CRTAC1 at all. The content exclusively discusses Hutchinson-Gilford Progeria Syndrome (HGPS) and the LMNA gene (lamin A and C), including a point mutation in exon 11 of LMNA that leads to progerin production, and therapeutic approaches such as farnesyltransferase inhibitors. There are no data, statements, mechanistic descriptions, variant analyses, patient reports, or literature citations that link CRTAC1 to any Mendelian disease. Therefore the paper provides neither direct evidence (e.g., reported pathogenic variants, segregation in families, patients with CRTAC1 mutations) nor mechanistic evidence (e.g., biochemical function, pathway involvement relevant to disease) supporting the claim that CRTAC1 is likely associated with a Mendelian disease. Limitation: absence of evidence in this single paper does not prove CRTAC1 is not associated with a Mendelian disease, but based on this paper alone there is no supportive information and no basis for inferring such an association.


[Read Paper](https://www.semanticscholar.org/paper/51db901b1bb3ae32a4025123f20e72eb07e1a2d8)


### Autosomal Dominant Intellectual Development Disorder-6 (MRD6) Without Seizures Linked to a De Novo Mutation in the grin2b Gene Revealed by Exome Sequencing: A Case Report of a Moroccan Child

**Why Not Relevant**: The provided paper text exclusively describes a Mendelian disorder caused by variants in the GRIN2B (grin2b) gene (Autosomal dominant intellectual development disorder-6, MRD6). It reports a specific nonsense variant c.3912C>G (p.Tyr1304Ter) in GRIN2B, describes its de novo status confirmed by Sanger sequencing, notes absence from gnomAD, and applies ACMG pathogenicity criteria (PVS1, PS2, PM2, PP3, PP5). There is no mention of CRTAC1, no CRTAC1 variants, no mechanistic discussion involving CRTAC1, and no clinical or functional data related to CRTAC1. Therefore this paper provides no direct evidence (no observed CRTAC1-disease co-segregation, no case reports) and no mechanistic evidence (no functional studies, no protein or pathway links) relevant to the claim that CRTAC1 is likely associated with a Mendelian disease. Absence of CRTAC1 discussion in this paper does not disprove any association, but it means this paper cannot be used to support or refute the claim. Relevant evidence for CRTAC1 would require text showing CRTAC1 variants segregating with disease, OMIM entry or ClinVar classifications implicating CRTAC1, functional assays demonstrating pathogenic mechanisms, or clear mechanistic links to known disease pathways — none of which appear in the supplied content.


[Read Paper](https://www.semanticscholar.org/paper/67d756e0cc302e5d46fdda9df34403f1f2de9a5a)


### Case report: A de novo Non-sense SOX9 mutation (p.Q417*) located in transactivation domain is Responsible for Campomelic Dysplasia

**Why Not Relevant**: The paper exclusively reports a novel heterozygous nonsense variant in SOX9 (c.1249C > T; p.Q417*) as the pathogenic lesion in a fetus with campomelic dysplasia and discusses SOX9 transactivation as a mechanistic basis for disease. There is no mention of CRTAC1 anywhere in the provided text, no reported variants in CRTAC1, and no functional or genetic data bearing on CRTAC1. Therefore this paper provides no direct genetic, clinical, or mechanistic evidence that CRTAC1 is associated with any Mendelian disease. Limitations: absence of CRTAC1 from this paper does not prove CRTAC1 is not a Mendelian disease gene, but it means this paper cannot be used as supporting evidence for the claim. Additionally, the study reports a single family/fetus and relies on bioinformatics prediction (and a statement about transactivation) rather than extensive functional validation, but these limitations concern SOX9 evidence and do not create any link to CRTAC1.


[Read Paper](https://www.semanticscholar.org/paper/bcc43a4127c8fa8d83c22e39fd3a13cbfa12fe1a)


### THE JAPANESE TERATOLOGY SOCIETY 62ND ANNUAL MEETING

**Why Not Relevant**: I found no direct or mechanistic evidence in the provided paper content that links the gene CRTAC1 to any Mendelian disease. The text contains many development- and gene-related passages (e.g., about Sonic hedgehog (Shh), Satb2, CRISPR/Cas9 genome editing, Il2rg in an X-SCID example, chemical teratogenesis in zebrafish, LIF/ACTH/CRH signaling, and Galectin-9), but the gene name CRTAC1 does not appear anywhere in the supplied excerpts. There are no human genetic association data, no case reports, no family segregation or linkage analyses, no reports of pathogenic variants, no functional studies of CRTAC1 loss- or gain-of-function, and no animal models or RNA/protein expression data involving CRTAC1 described in the provided text. Mechanistic discussion in the paper focuses on processes such as interkinetic nuclear migration, Shh signaling, Wnt signaling in cleft palate models, and maternal–fetal LIF–ACTH signaling; none of these passages mention CRTAC1 or provide a mechanistic pathway that implicates CRTAC1 in Mendelian disease.

Limitations and caveats: this conclusion is strictly based on the text you supplied. If the full paper contains additional sections, figures, supplementary materials, or gene lists that were not included here (for example an RNA-seq table or mention of CRTAC1 elsewhere), those could potentially provide relevant evidence. Based only on the provided content, however, there is no support for the claim that CRTAC1 is likely associated with at least one Mendelian disease. To evaluate that claim would require one or more of the following in the paper: explicit mention of CRTAC1 with (a) identification of rare segregating variants in patients with a Mendelian phenotype, (b) experimental functional evidence that CRTAC1 variants cause loss/gain of function consistent with disease, (c) animal models with CRTAC1 disruption recapitulating a Mendelian phenotype, or (d) statistically significant burden tests/association analyses implicating CRTAC1 in a Mendelian disorder. None of those elements are present in the provided excerpts.


[Read Paper](https://www.semanticscholar.org/paper/162811312bde34697100a4eaab4c8748157e2132)


### Limb development, evolution, and regeneration and repair: Part two

**Why Not Relevant**: The provided paper content does not mention CRTAC1 at all, nor does it provide any human genetic data, variant/phenotype correlations, linkage/segregation analyses, case reports, or experiments that test CRTAC1 function. The text is an overview of articles in a special issue on limb development and evolution and discusses genes such as Grem1, Gdf5, Pax3/Pax7, tbx2/3, drumstick, Keratins, and others, but there is no reference to CRTAC1 or to any Mendelian disease associations. Because there is no direct mention of CRTAC1, there is no direct evidence (no observational, genetic, or experimental data) in this paper to support or refute the claim that CRTAC1 is likely associated with a Mendelian disease. Mechanistic inference is also impossible from this source: while the issue addresses cartilage and limb development mechanisms (for example, Gdf5’s role in phalangeal cartilages), none of those mechanistic discussions include CRTAC1 or provide data that would allow linking CRTAC1 to Mendelian inheritance or to a human phenotype. Limitations: absence of evidence in this paper does not prove absence of an association overall; it only means this paper provides no relevant evidence. Recommended next steps: search human genetics resources (OMIM, ClinVar, HGMD), population/variant databases (gnomAD), and primary literature for CRTAC1 functional studies, human case reports, or segregation analyses to evaluate any potential Mendelian disease association.


[Read Paper](https://www.semanticscholar.org/paper/2293fd491e26309be00dcd6e7437c99f7e5d517b)


### Exome copy number variant detection, analysis and classification in a large cohort of families with undiagnosed rare genetic disease

**Why Not Relevant**: The provided paper content discusses CNV detection from exome sequencing and reports cohort-level results (number and sizes of CNVs, diagnostic yield, and classification counts), but it does not mention CRTAC1 anywhere in the supplied text. Therefore the paper content as given contains no direct evidence (no reported variants, cases, or statistical associations) linking CRTAC1 to any Mendelian disease. It provides only general, mechanistic, and methodological context (that CNVs can cause rare disease and that exome CNV calling can identify pathogenic CNVs). Because the claim is gene-specific (CRTAC1 associated with at least one Mendelian disease), and the excerpted paper content does not report gene-level findings or list affected genes, it cannot be used to support or refute the claim. Additional data—such as a gene-level table, variant annotations, case reports, or specific mention of CRTAC1—would be required to evaluate the claim. Note also that absence of mention in this paper is not evidence that CRTAC1 is not associated with a Mendelian disease; it only means this paper does not provide relevant information either way. Finally, even if the paper had considered CRTAC1 and found no CNVs, exome CNV calling may not detect all pathogenic variant types (e.g., single-nucleotide variants, small indels, regulatory variants), so a negative result would not conclusively refute a gene–disease association.


[Read Paper](https://www.semanticscholar.org/paper/2b8162a945544b579cfc6f8ac2e7a50d528bfcde)


### British Society for Matrix Biology Autumn 2020 Meeting: “Basement Membranes in Health and Disease”

**Why Not Relevant**: The provided paper content does not mention CRTAC1 (by that symbol or any obvious synonym) anywhere in the text. The excerpts above show the paper focuses on giantin, collagen biosynthesis/processing, integrin-mediated regulation of ECM, and specific ECM genes (e.g., LAMB2 and COL4A3/4/5) that are known to cause Mendelian kidney diseases. However, there is no experimental, genetic, clinical, expression, or mechanistic data concerning CRTAC1 in this document. Because the claim concerns CRTAC1 being likely associated with at least one Mendelian disease, and because the paper contains no reference to CRTAC1 (no mutation reports, no human genetics, no knockout/knockdown of CRTAC1, no expression or functional studies involving CRTAC1), this paper provides neither direct evidence nor mechanistic support for or against the claim. Any inference that CRTAC1 is disease-associated based on this paper would be speculative and unsupported by the content provided. Therefore relevance to the claim is effectively zero.


[Read Paper](https://www.semanticscholar.org/paper/e27ef31faa9280438adb116ad5e8faaf9838ff15)


## Search Queries Used

- CRTAC1 variant mutation Mendelian disease case report family segregation de novo autosomal recessive autosomal dominant exome sequencing

- cartilage acidic protein 1 CRTAC1 monogenic rare disease skeletal cartilage joint disorder clinical phenotype mutation

- CRTAC1 loss of function knockout mouse zebrafish CRISPR functional study expression cartilage development molecular mechanism

- exome sequencing rare disease cohort gene burden CRTAC1 copy number variant CNV systematic review meta analysis


## Bibliometric Configuration

- Use Bibliometrics: True

- Author Impact Weight: 0.4

- Citation Impact Weight: 0.4

- Venue Impact Weight: 0.2
