## Supplementary material for "MENDELSEEK: An algorithm that predicts Mendelian Genes and elucidates what makes them special": CWC22.txt

# Claim: The gene CWC22 is likely associated with at least one Mendelian disease.

**Status**: processed

**Overall Rating**: -1

**Explanation**:

No relevant papers were found that support or refute this claim.


## Relevant Papers


## Other Reviewed Papers


### Spliceosome malfunction causes neurodevelopmental disorders with overlapping features

**Why Not Relevant**: The provided paper content does not mention CWC22 anywhere. All genetic, functional, cellular, and organismal evidence reported concerns U2AF2, PRPF19, and RBFOX1 (and their orthologs/effects). Therefore the paper supplies no direct genetic association, segregation, variant, functional assay, or model-organism result involving CWC22 that would support or refute the claim that CWC22 is likely associated with at least one Mendelian disease. While the study demonstrates that spliceosome components can be Mendelian disease genes (a mechanistic rationale that might make CWC22 plausibly relevant a priori because it is also a spliceosome-associated factor), this is indirect and insufficient: evidence for other spliceosomal genes cannot be treated as evidence for CWC22 without CWC22-specific genetic or experimental data (e.g., observed pathogenic variants, segregation, de novo occurrence, functional disruption). In short, relevance is very low because the paper provides no CWC22-specific data or statements; any inference about CWC22 would require independent genetic or functional evidence not present in this manuscript.


[Read Paper](https://www.semanticscholar.org/paper/98d90a681d5535485a604cb4883121852358e976)


### Case Report: A Rare Case of Benign Recurrent Intrahepatic Cholestasis-Type 1 With a Novel Heterozygous Pathogenic Variant of ATP8B1

**Why Not Relevant**: This paper does not provide evidence for or against the claim that the gene CWC22 is likely associated with at least one Mendelian disease. The entire content provided concerns benign recurrent intrahepatic cholestasis type 1 (BRIC1) and a novel heterozygous pathogenic variant in ATP8B1 (c.1429+2T>G). The text contains no mention of CWC22, no genetic data or mechanistic discussion involving CWC22, and no phenotypes linked to CWC22. Therefore it is not relevant to assessing whether CWC22 is associated with a Mendelian disorder.

Specific reasons:
- No occurrence of the gene name CWC22 or synonyms in the provided content; the only gene discussed is ATP8B1.
- The paper's genetic, histopathologic, and therapeutic observations all concern ATP8B1-related BRIC1 and bile transport mechanisms (e.g., bile salt export pump expression), which are mechanistically unrelated to CWC22 based on the provided text.
- The study appears to be a single-patient case report describing a novel ATP8B1 splice-site variant and clinical response to rifampicin; even for ATP8B1 this is limited evidence (single case, heterozygous variant in an autosomal recessive disorder), so it would not bear on independent claims about an unrelated gene.

Because the paper does not mention or analyze CWC22 in any way, it cannot be used as direct or mechanistic evidence for the claim that CWC22 is associated with a Mendelian disease.


[Read Paper](https://www.semanticscholar.org/paper/1f94ba6cb68f2fecb4b3b5b7d19d5241863cb9ce)


### Biallelic splicing variants in the nucleolar 60S assembly factor RBM28 cause the ribosomopathy ANE syndrome

**Why Not Relevant**: Relevance is zero because the provided paper content exclusively discusses RBM28 and its role in ANE syndrome; the gene CWC22 is not mentioned anywhere in the excerpt. The paper presents clinical, genetic, in silico, cell-based splicing, and yeast-model functional evidence linking RBM28 variants to a Mendelian ribosomopathy. None of the text provides direct association data, mechanistic links, or any commentary about CWC22, nor does it present cross-gene analyses that would allow inference about CWC22. Therefore this paper does not provide evidence for or against the claim that CWC22 is likely associated with at least one Mendelian disease.


[Read Paper](https://www.semanticscholar.org/paper/a60618e9deb270c19c972d700fb2814ffbd8337c)


### Clinical and Genetic Characteristics of Patients with Unexplained Intellectual Disability/Developmental Delay without Epilepsy

**Why Not Relevant**: The paper does not mention the gene CWC22 anywhere in the provided content. It reports diagnostic yields and lists specific genes and novel variants identified in a cohort of 38 patients with unexplained ID/DD and/or ASD, none of which are CWC22. While the methods used (CMA, CES, WES) are capable of detecting many classes of pathogenic variants that could implicate a Mendelian disease gene like CWC22, absence of CWC22 findings in this small, phenotype-selected cohort is weak negative evidence and cannot be used to conclude that CWC22 is or is not associated with a Mendelian disease. Additional reasons the paper is not relevant: (1) the study size is small (38 subjects), so power to detect rare CWC22-associated cases is limited; (2) WES/CES can miss noncoding or certain structural variants that might implicate CWC22; (3) the clinical phenotype focus (ID/DD/ASD) may not capture other phenotypes linked to CWC22 if any. Therefore, the paper neither provides direct supporting evidence nor mechanistic data specifically about CWC22, and its absence from the reported findings is not sufficient to support or refute the claim.


[Read Paper](https://www.semanticscholar.org/paper/e5289119c3a34a33fd81c6fcc009413eb467af22)


### Clinical case report: mosaic ANK3 pathogenic variant in a patient with autism spectrum disorder and neurodevelopmental delay

**Why Not Relevant**: The provided paper content exclusively discusses ankyrin proteins and ANK3 (Ankyrin 3)—its roles in the CNS, and a reported de novo mosaic nonsense variant in ANK3 associated with neurodevelopmental phenotypes. The text does not mention the gene CWC22, nor does it provide any data, discussion, or mechanistic information linking CWC22 to disease. Therefore there is no direct evidence (e.g., genetic, phenotypic, segregation, case reports) nor mechanistic evidence (e.g., biochemical function, pathway involvement, animal models) in this paper that bears on the claim that CWC22 is likely associated with at least one Mendelian disease. Because the gene of interest (CWC22) is not present in the paper, the content cannot support or refute the claim.


[Read Paper](https://www.semanticscholar.org/paper/8d26866252dcf55922aa6ce59017540fbbc29c46)


### Trio-whole exome sequencing reveals the importance of de novo variants in children with intellectual disability and developmental delay

**Why Not Relevant**: The provided paper content contains only a single general sentence about the diagnostic yield of trio-WES for DD/ID and does not mention CWC22 or any data (genetic, clinical, statistical, or functional) pertaining to that gene. There are no case reports, variant annotations, segregation analyses, functional assays, population frequency data, or mechanistic descriptions linking CWC22 to a disease phenotype. While the sentence implies that trio-WES can detect Mendelian causes of DD/ID (and thus could be used to discover gene-disease associations), it does not provide any evidence specific to CWC22. Because relevance to the claim requires gene-specific direct or mechanistic evidence, this content is not relevant (relevance score 0.0). To evaluate the claim about CWC22 one would need excerpts that report variants in CWC22 in affected individuals, statistical enrichment, segregation in families, functional impact of variants on gene/protein function, or other mechanistic data—none of which are present in the provided text.


[Read Paper](https://www.semanticscholar.org/paper/370a01dc9cf9e59a3df214c33c43cd7787d2426e)


### in an model. SAM68 is a physiological regulator of SMN2 splicing in spinal muscular atrophy

**Why Not Relevant**: The provided paper content focuses exclusively on spinal muscular atrophy (SMA), the SMN1/SMN2 genes, and the role of splicing regulators (notably SAM68, hnRNP A1, and U2AF65) in SMN2 exon 7 inclusion. There is no mention of the gene CWC22, no human genetic association data, no variant pathogenicity analyses, no case reports, and no discussion of Mendelian disease linkage for any gene other than SMN1/SMN2. The mechanistic data concern SAM68-mediated regulation of SMN2 splicing in an SMA mouse model; these results cannot be used to infer an association between CWC22 and any Mendelian disease. Because the paper contains no direct or indirect information about CWC22, it does not provide evidence supporting or refuting the claim that CWC22 is likely associated with at least one Mendelian disease.


[Read Paper](https://www.semanticscholar.org/paper/2d30ef51010150845d8d0fabba563e1a870129f9)


### RhoGAPp190: A potential player in tbph-mediated neurodegeneration in Drosophila

**Why Not Relevant**: The paper focuses on TDP-43/tbph function in Drosophila and the identification and characterization of RhoGAPp190 as a potential tbph target. There is no mention of the human gene CWC22, nor any discussion of Mendelian diseases or gene-disease associations relevant to CWC22. All quoted material pertains to TDP-43 biology, the fly ortholog tbph, and RhoGAPp190. Therefore the paper provides no direct evidence that CWC22 is associated with any Mendelian disease, nor mechanistic links involving CWC22. Because the content neither names nor implicates CWC22, its relevance to the claim is essentially nil.


[Read Paper](https://www.semanticscholar.org/paper/4d22772f6c9360e20ac0e21f1195bacf7880a6bf)


### A novel pathogenic variant of DNMT3A associated with craniosynostosis: a case report of Heyn–Sproul–Jackson syndrome

**Why Not Relevant**: The provided paper content exclusively discusses pathogenic variants in DNMT3A and their association with Tatton-Brown-Rahman syndrome and Heyn-Sproul-Jackson syndrome (HESJAS). There is no mention of the gene CWC22, no variant data reported for CWC22, and no mechanistic discussion that would link DNMT3A findings to CWC22 (for example via pathway analyses, genetic interactions, or co-segregation). Because the manuscript is a case report/series focused on DNMT3A and its clinical manifestations, it provides neither direct genetic evidence (variants, segregation, case series) nor mechanistic evidence (biochemical, cellular, or pathway data) that would support an association between CWC22 and a Mendelian disease. Additionally, the evidence in the paper is a single case report of a novel DNMT3A variant, which limits generalizability even for DNMT3A itself and does not permit inference to unrelated genes. Therefore, this paper does not provide relevant evidence for the claim that CWC22 is likely associated with at least one Mendelian disease.


[Read Paper](https://www.semanticscholar.org/paper/4593a650bdab490666074d3d6811588ba54430b1)


### A de novo PAK1 likely pathogenic variant and a de novo terminal 1q microdeletion in a Chinese girl with global developmental delay, severe intellectual disability, and seizures

**Why Not Relevant**: The provided paper content discusses a novel PAK1 variant associated with a terminal 1q microdeletion in a patient with neurodevelopmental disorder and states that the main clinical features may ascribe to the pathogenic variant in PAK1. There is no mention of the gene CWC22, no data about CWC22's involvement in the terminal 1q microdeletion, and no mechanistic or clinical evidence linking CWC22 to any Mendelian disease in the supplied text. Therefore the content neither supports nor refutes the claim that CWC22 is likely associated with at least one Mendelian disease.

Limitations and caveats: (1) The supplied content is a single summary sentence and may be incomplete. The full paper (including main text, results, tables, figures, and supplements) could mention other genes in the deleted interval — possibly including CWC22 — but that information is not present in the excerpt provided. (2) Because no explicit sentences or paragraphs about CWC22 are available in the provided content, no direct or mechanistic evidence can be extracted. To evaluate the claim properly, please provide the full text or the sections listing genes included in the terminal 1q deletion, variant tables, or any discussion of CWC22.


[Read Paper](https://www.semanticscholar.org/paper/8ac30d308f85987b8cf4644520ee11940136ebaa)


### KCNB1 frameshift variant caused inherited intellectual disability, developmental delay, and seizure.

**Why Not Relevant**: The provided paper content does not mention the gene CWC22 and therefore contains no direct or mechanistic evidence relevant to the claim that CWC22 is likely associated with at least one Mendelian disease. Instead, the paper concerns a different gene, KCNB1. Verbatim sentences from the supplied text illustrating this focus include: "Potassium voltage-gated channel subfamily B member 1 (KCNB1) encodes Kv2.1 potassium channel." and "We performed whole exome sequence analysis (WES) and identified a heterozygous frameshift mutation c.522delA in exon 1 of KCNB1 (NM_004975.4) predicting a premature stop codon p.Lys174Asnfs*20 in the proband." Because the gene discussed (KCNB1) is different from the gene in the claim (CWC22), the paper provides neither direct evidence (observations of CWC22 variants segregating with disease) nor mechanistic evidence (molecular or physiological mechanisms linking CWC22 dysfunction to a Mendelian phenotype) relevant to the claim. Note also that absence of CWC22 in this excerpt does not prove CWC22 is or is not associated with Mendelian disease; it only shows this particular paper is not a source of evidence for that claim. To evaluate the claim about CWC22, literature and database searches (e.g., PubMed, OMIM, ClinVar, gnomAD, HGMD) for CWC22-specific reports would be required.


[Read Paper](https://www.semanticscholar.org/paper/eb672d4472edfa39f564bbfe046982296e47094c)


### SAM 68 is a physiological regulator of SMN 2 splicing in spinal muscular atrophy

**Why Not Relevant**: The provided paper content does not mention the gene CWC22, nor does it contain any genetic-association data, case reports, linkage analyses, variant pathogenicity assessments, or other evidence that would support or refute an association between CWC22 and a Mendelian disease. Instead, the paper focuses on spinal muscular atrophy (SMA), the SMN1/SMN2 genes, and the role of the splicing factor SAM68 in modulating SMN2 exon 7 inclusion in an SMA mouse model. Representative verbatim sentences from the text that illustrate the paper's focus are: "Spinal muscular atrophy (SMA) is the leading genetic cause of infant mortality, with an incidence of approximately one in 6,000–10,000 newborns (Arnold and Burghes, 2013; Nurputra et al., 2013)."; "We found that knockout of the splicing factor SAM68 partially rescued body weight and viability of SMAΔ7 mice."; and "Mechanistically, SAM68 binds to SMN2 pre-mRNA, favoring recruitment of the splicing repressor hnRNP A1 and interfering with that of U2AF65 at the 3′ splice site of exon 7." These quotes show the study's mechanistic and phenotypic focus is on SMN2 splicing and SAM68 function in SMA models, not on CWC22.

Limitations relevant to the claim assessment: (1) No mention or data about CWC22 in the entire provided text, so there is no direct evidence for or against CWC22 being associated with any Mendelian disease. (2) The experimental system described is an SMA mouse model and concerns modulation of SMN2 splicing by SAM68; even if a splicing-related mechanism were generalizable, absence of any direct data on CWC22 means no mechanistic support can be inferred. (3) The paper addresses a specific gene-disease relationship (SMN1/SMN2 and SMA) and a regulatory factor (SAM68); extrapolating from these findings to implicate an unrelated gene (CWC22) in Mendelian disease would be speculative and unsupported by the text. For these reasons the relevance of this paper to the claim that "CWC22 is likely associated with at least one Mendelian disease" is effectively zero.


[Read Paper](https://www.semanticscholar.org/paper/8758722656d2e1f8f5bb8d001ea103fbd78a1cd9)


## Search Queries Used

- CWC22 pathogenic variant Mendelian disorder case report

- CWC22 rare variants exome sequencing de novo intellectual disability developmental delay cohort

- CWC22 pre mRNA splicing factor loss of function haploinsufficiency splicing defect cellular model

- CWC22 ClinVar OMIM gnomAD pLI loss of function intolerance population genetics


## Bibliometric Configuration

- Use Bibliometrics: True

- Author Impact Weight: 0.4

- Citation Impact Weight: 0.4

- Venue Impact Weight: 0.2
