## Supplementary material for "MENDELSEEK: An algorithm that predicts Mendelian Genes and elucidates what makes them special": DAZ3.txt

# Claim: The gene DAZ3 is likely associated with at least one Mendelian disease.

**Status**: processed

**Overall Rating**: 3

**Explanation**:

Supporting evidence.
The provided papers consistently link the AZFc region of the Y chromosome, which contains the DAZ gene cluster, to spermatogenic failure (azoospermia and severe oligozoospermia). Paper excerpts state that “the association of the complete AZFc deletion with spermatogenic failure is well established,” and that Y chromosome microdeletions (including AZFc deletions) account for ~10–15% of idiopathic azoospermia/severe oligozoospermia. The DAZ genes (DAZ1–DAZ4) are repeatedly mentioned as components of the AZFc locus; one excerpt notes that DAZ3 can be identified using a gene copy–specific STS amplification, and another describes how partial AZFc duplications/deletions change the number of DAZ copies and can be risk factors for male infertility. These points support the biological plausibility that loss or alteration of DAZ copies (including DAZ3) can contribute to a human reproductive phenotype that could be Mendelian in a Y-linked fashion.

Caveats or contradictory evidence.
Despite the strong association between AZFc deletions and spermatogenic failure, the excerpts also emphasize important uncertainties. Multiple papers state that AZF deletions remove several genes simultaneously and that “it is currently unknown which of these individual genes or any combination thereof play an essential role,” so causal attribution to a single DAZ copy (e.g., DAZ3) is unresolved. Partial AZFc deletions and duplications have been observed in both infertile and fertile men, indicating incomplete penetrance or context dependence (Y-chromosomal background or other modifiers). Another excerpt highlights that, after years of study, only a handful of genetic alterations have been clearly shown to cause spermatogenic failure, and identification of additional monogenic causes has been difficult. Thus, while DAZ cluster defects are implicated, direct evidence that DAZ3 specifically causes a Mendelian disease is lacking in these excerpts.

Analysis of potential mechanisms and deeper relationships.
Mechanistically, DAZ family genes encode RNA-binding proteins implicated in germ cell development; loss of copies could plausibly impair spermatogenesis. Because the DAZ genes reside on the male-specific region of the Y chromosome, any single-copy loss in a hemizygous male could produce a Y-linked (male-limited) Mendelian phenotype if that copy were necessary and sufficient. However, the DAZ genes occur in multiple nearly identical copies (four DAZ copies in some chromosomes), so functional redundancy among copies complicates single-gene causal inference: loss of one copy may be compensated by others, while deletion of the entire cluster or specific combinations may be required to produce phenotype. The observation that partial deletions/duplications have variable effects on fertility suggests dosage sensitivity combined with modifying background factors rather than a straightforward single-copy Mendelian effect for DAZ3 alone. Finally, ascertainment and methodological limitations (e.g., deletions removing several genes, difficulty assigning copy-specific effects) reduce the strength of single-gene claims.

Which rating is most appropriate.
The preponderance of evidence shows a well-supported association between AZFc/DAZ cluster abnormalities and spermatogenic failure, but the evidence does not demonstrate that DAZ3 by itself is a Mendelian disease gene. The provided excerpts explicitly note that it is unknown which individual genes within deleted regions are essential and that partial deletions are sometimes present in fertile men. Given the plausible mechanism and the clear involvement of the DAZ locus in a clinically relevant phenotype, DAZ3 is a candidate, but the specific claim that DAZ3 is “likely associated with at least one Mendelian disease” goes beyond what these data establish. Therefore the most defensible conclusion based on the supplied evidence is Mixed Evidence: there is plausible and suggestive support but insufficient direct, copy-specific genetic proof to confidently rate the claim as Likely True or Highly Supported.


**Final Reasoning**:

I reviewed the logic and citations in the essay: the AZFc region (including DAZ genes) is clearly implicated in spermatogenic failure, which is a clinically recognized phenotype often resulting from Y deletions. However, the supplied excerpts explicitly state that which individual genes in AZF deletions are causative is unknown, and partial DAZ deletions/duplications can be found in fertile men, indicating incomplete penetrance or context dependence. Because the claim asserts DAZ3 is "likely associated with at least one Mendelian disease," and the provided evidence does not give copy-specific causal proof or demonstrated segregation of DAZ3 variants with disease, it would be inappropriate to call the claim Likely True or Highly Supported. Conversely, there is nontrivial supporting evidence (biological plausibility, locus-level association), so dismissing the claim entirely would also be incorrect. Therefore the rating "Mixed Evidence" best reflects that DAZ3 is a plausible candidate but lacks direct, conclusive, copy-specific Mendelian evidence in the provided literature.


## Relevant Papers


### Genetic Causes of Male Infertility

**Authors**: D. Plaseska‐Karanfilska (H-index: 24), Z. Moneva (H-index: 1)

**Relevance**: 0.35

**Bibliometric Impact**: 0.18620000000000003


**Excerpts**:

- Deletions most frequently involve the AZFc region, less frequently the AZFb region, and only rarely the AZFa region.

- While the association of the complete AZFc deletion with spermatogenic failure is well established, the role of partial AZFc deletions on spermatogenesis and male infertility is still controversial.

- Duplications can occur on a chromosome with a partial AZFc deletion and generate a chromosome with four DAZ genes, but lacking some sequence tagged site (STS) markers [10,11].

- Recently, an AZFc partial duplication has been shown to be a risk factor for male infertility in Taiwan [12].

- Y chromosome microdeletions represent the etiological factor of 10.0–15.0% of idiopathic azoospermia and severe oligozoospermia [3].

- Y chromosome microdeletions are found almost exclusively in patients with azoospermia or severe oligozoospermia [5].

- Partial AZFc deletions as well as AZFc duplications were present in both infertile and fertile men. They may represent a risk factor for male infertility when present on certain Y chromosomal backgrounds.

- The analysis of polymorphisms in genes involved in spermatogenesis represents one of the most exciting areas of research in genetics of male infertility [16].


**Explanations**:

- This sentence establishes that AZFc is the Y-chromosome region most frequently affected by deletions. Relevance to the claim: indirect mechanistic context. Strength: indicates target genomic region where DAZ genes (including DAZ3) reside are commonly deleted in infertile men. Limitation: does not mention individual DAZ gene copies (DAZ3) or attribute disease specifically to any single DAZ gene.

- This sentence states that complete AZFc deletions are clearly associated with spermatogenic failure, while partial deletions are less certain. Relevance: mechanistic and epidemiological support that loss of material in AZFc causes a Mendelian-like phenotype (spermatogenic failure). Strength: asserts a well-established association for complete AZFc deletions. Limitation: does not identify which gene(s) within AZFc (e.g., DAZ3) are causative; partial-deletion uncertainty weakens inference to a single-gene Mendelian disease.

- This sentence explicitly names DAZ genes as being affected by AZFc structural changes and describes duplication generating four DAZ genes. Relevance: mechanistic evidence linking DAZ gene copy-number changes to structural variants in AZFc. Strength: directly implicates DAZ gene family as subject to deletions/duplications that alter copy number, a plausible mechanism for affecting spermatogenesis. Limitation: refers to DAZ genes generically (DAZ1–4) and does not single out DAZ3; no demonstration that alteration of DAZ3 alone causes a Mendelian disease.

- This reports that an AZFc partial duplication has been associated with increased risk of male infertility in a specific population. Relevance: epidemiological support that structural variation in the AZFc region (which includes DAZ genes) can be a risk factor—mechanistic/evidence link. Strength: cites population-level association. Limitation: is population-specific (Taiwan) and addresses partial duplications (not necessarily DAZ3-specific) and reports a risk factor rather than definitive Mendelian causation.

- This provides the fraction of idiopathic azoospermia and severe oligozoospermia cases attributable to Y chromosome microdeletions. Relevance: direct epidemiological evidence that Y-chromosome deletions, which include AZFc and DAZ genes, are common genetic causes of severe spermatogenic phenotypes. Strength: quantifies etiologic contribution (10–15%). Limitation: microdeletions typically remove multiple genes; does not demonstrate that DAZ3 specifically is necessary and sufficient for a Mendelian disease.

- This reinforces that Y microdeletions are observed predominantly in patients with severe spermatogenic phenotypes. Relevance: supports that deletions in regions containing DAZ genes are strongly associated with disease phenotypes. Strength: specificity of phenotype association. Limitation: same as above—association is with regions/multiple genes, not with a single DAZ3 gene; microdeletions are structural (CNV) variants rather than single-gene point mutations classically described in Mendelian disease annotations.

- This sentence notes that both partial AZFc deletions and duplications occur in fertile and infertile men and may act as risk factors depending on Y-chromosomal background. Relevance: provides nuance about penetrance and genetic background—mechanistic/epidemiologic context. Strength: highlights that AZFc/DAZ alterations can be modifiers/risk factors. Limitation: the presence in fertile men indicates incomplete penetrance and argues against a simple highly penetrant Mendelian relationship for any single DAZ copy like DAZ3.

- This general sentence emphasizes study of polymorphisms in spermatogenesis genes as an active area. Relevance: contextual mechanistic relevance—genetic variation in spermatogenesis genes (including DAZ family members) is a plausible cause of infertility. Strength: supports biological plausibility. Limitation: it is a general statement about research direction and not evidence for DAZ3 specifically causing a Mendelian disease.


[Read Paper](https://www.semanticscholar.org/paper/2204f207379ecabbbacd75c73cedc5e71ae9b40b)


### P-510 Global methylation status of human oocytes in ART Genetics: male P-511 Study of meiotic segregation and location of chromosome breakpoints in rare Robertsonian translocations

**Authors**: Y. Belaid (H-index: 1), F. Pellestor (H-index: 32)

**Relevance**: 0.25

**Bibliometric Impact**: 0.132


**Excerpts**:

- These data demonstrate that patients with an abnormal karyotype are found to have severe oligospermia or azoospermia.

- Chromosomal analysis, therefore, should be offered to patients with severe oligospermia (<5 million/ml) or sperm morphology of <5%, thus reducing the cost implications and patient anxiety.

- Our study analyses gr/gr, b1/b3 and b2/b3 deletions by biplex STS PCR (sY1161, sY1197, sY1191, sY1291, sY1206, sY1201), DAZ1, DAZ2, DAZ4 copies identification by SNV analysis using DHPLC with specific primers.

- DAZ3 is identified using a gene copy specific STS amplification.

- Although we were not yet able to identify UTY gene mutations in infertile men associated with some specific pathology, we can assume that the DBY gene is most likely the major AZFa gene causing the SCO syndrome as observed in men with AZFa deletions, whereas USP9Y gene deletions are expected to cause only a post-meiotic male germ line defect.


**Explanations**:

- Relation to claim: Indirect clinical evidence linking chromosomal abnormalities to a Mendelian-like disease phenotype (severe oligospermia/azoospermia). Strength/limitations: This is observational clinical evidence that chromosomal abnormalities (including Y-chromosome structural changes) are associated with male infertility. It does not mention DAZ3 specifically, nor does it demonstrate that a mutation in DAZ3 causes a Mendelian disease; it supports plausibility that Y-linked genes can underlie severe spermatogenic disorders.

- Relation to claim: Contextual clinical recommendation showing a diagnostic relationship between chromosomal/Y abnormalities and severe spermatogenic phenotypes. Strength/limitations: Reinforces that genetic testing is used because chromosomal/Y deletions have clinical impact. Still no direct statement that DAZ3 mutations cause a Mendelian disease; this supports the clinical relevance of detecting Y-linked gene copy changes but does not prove causality for DAZ3.

- Relation to claim: Methodological evidence that the study specifically assays known Y-chromosome deletions and DAZ gene copies (DAZ1, DAZ2, DAZ4). Strength/limitations: Demonstrates the authors target DAZ gene family members and known deletion types implicated in spermatogenic failure. However, it lists methods and targets but does not report an association between DAZ3 and disease in the quoted text; DAZ3 is not enumerated among DAZ1/2/4 in this sentence.

- Relation to claim: Direct procedural statement that DAZ3 copy is identified using a gene copy-specific STS amplification. Strength/limitations: This is the only explicit mention of DAZ3 in the provided text and shows the gene was assayed. It does not state any association between DAZ3 mutation/deletion and a Mendelian disease phenotype (e.g., azoospermia), nor provide results or statistical associations for DAZ3 specifically.

- Relation to claim: Mechanistic/contextual evidence that Y-chromosome gene deletions (e.g., DBY, USP9Y) can cause defined clinical syndromes (SCO syndrome or post-meiotic defects), illustrating precedent that single Y-linked genes can underlie spermatogenic disorders. Strength/limitations: This supports biological plausibility that a Y-linked gene such as DAZ3 could be causative of a Mendelian phenotype, but it does not provide direct evidence for DAZ3 itself. It also reflects inference rather than direct demonstration and notes that UTY mutations were not identified in that analysis.


[Read Paper](https://www.semanticscholar.org/paper/f0f90e413b46d6ef747e904e8799651c3552b5a9)


### Methodological issues in the study of the genetics of spermatogenic failure

**Authors**: L. Visser (H-index: 3), S. Repping (H-index: 58)

**Relevance**: 0.2

**Bibliometric Impact**: 0.244


**Excerpts**:

- Second, five recurrent Y-chromosome deletions, the so-called AZFa, P5/proximal P1 (AZFb), P5/distal P1, b2/b4 (AZFc) and gr/gr deletions, are associated with spermatogenic failure (Reijo et al. 1995, Vogt et al. 1996, Repping et al. 2002, 2003, Visser et al. 2009). These deletions all remove multiple genes that are potentially involved in normal human spermatogenesis. It is currently unknown which of these individual genes or any combination thereof play an essential role, and in which dosage (Noordam & Repping 2006).

- Spermatogenesis is governed by the parallel and serial actions of thousands of genes (Sha et al. 2002, Schultz et al. 2003, Schlecht et al. 2004, Ellis et al. 2007, Zamudio et al. 2008).

- In theory, alterations in any of these genes or their expression may cause spermatogenic failure. In reality, only a handful of genetic alterations have clearly been shown to cause spermatogenic failure (Nuti & Krausz 2008).

- Third, a number of monogenic disorders, listed in the Online Mendelian Inheritance in Man database, are associated with spermatogenic failure. Among these are Kallmann syndrome and Noonan syndrome. In monogenic disorders, spermatogenic failure results from hypogonadotrophic hypogonadism, cryptorchidism, delayed puberty or male pseudohermaphroditism and ambiguous genitalia (Hardelin & Dode 2008).

- Despite years of research, only few genetic factors have clearly been shown to cause spermatogenic failure, and the identification of additional genetic causes or risk factors has proven to be extremely difficult.

- First, many previous studies have included men based on their (reported) fertility, instead of their semen quality.


**Explanations**:

- Relation to claim: mechanistic/indirect. This passage states that recurrent Y-chromosome deletions (including AZFc) are associated with spermatogenic failure and that these deletions remove multiple genes potentially involved in spermatogenesis, but that it is unknown which individual genes are essential or in which dosage. Strength: establishes that genes within AZFc can be causally linked to a Mendelian-type phenotype (spermatogenic failure) when deleted. Limitation: the excerpt does not name DAZ3 specifically, and explicitly states that the causal contribution of individual genes within these deletions is unknown — therefore it is indirect evidence at best for DAZ3 being a Mendelian disease gene.

- Relation to claim: mechanistic. This sentence provides biological plausibility: spermatogenesis depends on thousands of genes, so single-gene defects could theoretically produce Mendelian spermatogenic phenotypes. Strength: supports plausibility that a specific gene (such as DAZ3) could be involved. Limitation: purely general and does not identify DAZ3 or give disease-association data.

- Relation to claim: direct/interpretive. These sentences temper expectations: while many genes could be involved, only a few genetic alterations have been clearly shown to cause spermatogenic failure. Strength: emphasizes the high evidentiary bar for assigning causality. Limitation: does not address DAZ3 directly, but implies that evidence for many candidate genes remains unproven, weakening an unsupported assertion that DAZ3 is a Mendelian disease gene.

- Relation to claim: contextual/direct. This passage documents that some monogenic (Mendelian) disorders are associated with spermatogenic failure and gives examples (Kallmann, Noonan). Strength: shows that Mendelian disorders can manifest with spermatogenic failure and are catalogued in OMIM. Limitation: the described monogenic disorders are systemic syndromes causing spermatogenic failure via endocrine or developmental mechanisms; the excerpt does not indicate that DAZ3 is among OMIM-listed Mendelian disease genes.

- Relation to claim: contextual/interpretive. This sentence reiterates that despite extensive research, only few genetic causes have been clearly proven, and finding additional causal genes is difficult. Strength: realistic assessment of current knowledge; limitation: again not specific to DAZ3 and highlights that candidate-gene claims require strong evidence which is not present in this paper.

- Relation to claim: methodological/caveat. This sentence points out a common methodological flaw in genetic association studies (case ascertainment based on reported fertility rather than measured semen parameters). Strength: explains why published associations (potentially including Y-linked gene associations) may be unreliable. Limitation: does not directly mention DAZ3, but implies that any weak association reported elsewhere would need careful re-evaluation.


[Read Paper](https://www.semanticscholar.org/paper/198a680ee854c51c9699f4ab8c091cd3e9e60e85)


## Search Queries Used

- DAZ3 variant mutation deletion DAZ3 gene Y chromosome AZFc azoospermia non obstructive azoospermia male infertility Mendelian disease association

- DAZ3 protein function RNA binding spermatogenesis germ cell development DAZ family DAZ1 DAZ2 DAZL functional study knockout model redundancy paralog compensation

- Y chromosome microdeletion AZFc DAZ genes copy number variation DAZ3 association spermatogenic failure case control study systematic review meta analysis

- DAZ3 copy number polymorphism benign variation lack of association normal fertility incomplete penetrance oligogenic multifactorial causes diagnostic sensitivity specificity


## Usage Statistics

- Prompt Tokens: 0

- Completion Tokens: 0

- Total Tokens: 0

- Estimated Cost: $0.0238


## Bibliometric Configuration

- Use Bibliometrics: True

- Author Impact Weight: 0.4

- Citation Impact Weight: 0.4

- Venue Impact Weight: 0.2
