## Supplementary material for "MENDELSEEK: An algorithm that predicts Mendelian Genes and elucidates what makes them special": DDIT3.txt

# Claim: The gene DDIT3 is likely associated with at least one Mendelian disease.

**Status**: processed

**Overall Rating**: 2

**Explanation**:

Supporting evidence:
The provided papers consistently report that DDIT3 (CHOP) is an important mediator of endoplasmic reticulum (ER) stress and that its expression or activity is associated with pathophysiological outcomes. For example, Taniguchi & Yoshida review ER stress pathways and note the PERK–ATF4–CHOP pathway is proapoptotic in some kidney diseases. Havighorst et al. report that CHOP (Ddit3) expression in deer mice predicts later-life metabolic pathology (elevated plasma cholesterol and hepatic steatosis). Germani & Zito show increased CHOP in a mouse model of RyR1-related pathology and that loss of CHOP did not rescue the phenotype. These findings support a role for DDIT3 in disease processes and in modulating susceptibility or severity in animal and cellular models.

Caveats or contradictory evidence:
None of the provided excerpts (or the studies they’re taken from) present evidence that germline variants in DDIT3 cause a Mendelian (monogenic, inherited) human disease. The studies describe CHOP as a mediator or biomarker of ER stress in complex or induced disease models or as part of somatically acquired oncogenic events in the literature (e.g., known somatic FUS–DDIT3 fusions in myxoid liposarcoma, which are cancer-specific, not inherited germline causes). The deer mouse and mouse-model papers describe expression-level associations, inducibility, or loss-of-function experiments in animals that affect disease phenotypes, but these do not demonstrate that pathogenic germline mutations in human DDIT3 produce a Mendelian syndrome with segregation in families.

Analysis of potential underlying mechanisms and implications:
Mechanistically, DDIT3 encodes CHOP, a transcription factor induced by the unfolded protein response that promotes apoptosis under unresolved ER stress. This positions DDIT3 as a downstream effector in pathways where other genes (e.g., EIF2AK3/PERK) when mutated can produce Mendelian disorders (for example, EIF2AK3 mutations cause Wolcott–Rallison syndrome). However, being a critical pathway effector does not by itself imply that germline mutations in DDIT3 produce a heritable monogenic disorder. Many genes that participate in stress-response pathways influence disease risk or progression without being primary causes of Mendelian conditions. The available data show DDIT3’s expression modulates disease processes, but do not document co-segregating, high-penetrance germline variants in human families.

Which rating is most appropriate and why:
Given the absence of any evidence in the provided papers for germline, segregating pathogenic variants in DDIT3 causing a Mendelian disease, and given the body of work that instead implicates DDIT3 as a mediator of ER stress responses and sometimes as a somatically altered oncogene in cancers, the claim that "DDIT3 is likely associated with at least one Mendelian disease" is not supported by the presented evidence. It is possible (though currently unsupported) that rare germline DDIT3 variants could be discovered as causes of monogenic disease in the future, but that is speculative. The balance of evidence indicates that DDIT3 is involved in disease biology and risk modulation but not established as a Mendelian disease gene. Therefore the most appropriate rating is "Likely False."


**Final Reasoning**:

I reviewed the provided excerpts and incorporated domain knowledge. The studies supplied show DDIT3/CHOP functions in ER stress, predicts or modulates disease phenotypes in animal models, and is implicated in somatic oncogenic fusions — none of which constitute evidence of a germline, segregating cause of a Mendelian disease. Other ER stress pathway genes are known Mendelian disease genes, which could lead to confusion, but that indirect relationship does not establish DDIT3 itself as a Mendelian gene. No paper provided reports family segregation, pathogenic germline variants, or a named Mendelian syndrome attributable to DDIT3. On that basis the claim is not supported by the evidence and is best rated "Likely False." I find no flaw in this logic given the materials; if new genetic evidence (e.g., multiple unrelated families with segregating DDIT3 pathogenic variants and consistent phenotype) appears, the assessment should be revisited.


## Relevant Papers


### Endoplasmic reticulum stress in kidney function and disease

**Authors**: M. Taniguchi (H-index: 9), H. Yoshida (H-index: 42)

**Relevance**: 0.18

**Bibliometric Impact**: 0.32911111111111113


**Excerpts**:

- Recent findingsER stress is evoked in various kidney diseases, including diabetic nephropathy, renal fibrosis, inflammation or osmolar contrast-induced renal injury, ischemia-reperfusion, genetic mutations of renal proteins, proteinuria and cyclosporine A treatment.

- In some cases, chemical chaperones, such as 4-phenylbutyrate and taurodeoxycholic acid, relieve the symptoms, indicating that ER stress-induced apoptosis of renal cells is one of the major causes of certain kidney diseases.

- Actually, the ER stress response provides protection against some kidney diseases, although the PERK-ATF4-CHOP pathway of the ER stress response is proapoptotic in some kidney diseases.

- SummaryBecause ER stress is a major cause of some kidney diseases, the ER stress response and autophagy, which deal with unfolded proteins that accumulate in the ER, are promising therapeutic targets in acute and chronic kidney diseases.


**Explanations**:

- This sentence documents that ER stress is observed in multiple kidney disease contexts, including cases caused by "genetic mutations of renal proteins." It is mechanistic/contextual evidence that ER stress pathways (which include DDIT3/CHOP) are active in human disease processes. Strength: establishes biological contexts where ER stress participates in disease. Limitation: it does not mention DDIT3 or CHOP explicitly by gene name, nor does it provide any genetic evidence linking variants in DDIT3 to a Mendelian (monogenic) disorder.

- This sentence reports that reducing ER stress with chemical chaperones can relieve symptoms, "indicating that ER stress-induced apoptosis of renal cells is one of the major causes of certain kidney diseases." This provides functional/mechanistic evidence that ER stress-mediated pathways (to which DDIT3/CHOP contributes) can drive disease phenotypes and be therapeutically targeted. Strength: functional intervention supports causality of ER stress in disease pathology. Limitation: it is therapeutic/mechanistic evidence from interventions, not genetic evidence tying DDIT3 variants to a Mendelian disease; method details (model systems, species, sample sizes, statistics) are not provided here.

- This sentence specifically names the PERK-ATF4-CHOP pathway as "proapoptotic in some kidney diseases." CHOP is the protein product of the DDIT3 gene (DDIT3 encodes CHOP/GADD153), so this is direct mechanistic evidence that the DDIT3-encoded protein participates in disease-relevant apoptosis pathways. Strength: directly links the CHOP node of the ER stress response to disease-relevant cell death, which increases biological plausibility that perturbation of DDIT3 could affect disease. Limitation: the statement describes pathway activity and pathogenic mechanism but does not present genetic linkage, variant pathogenicity, segregation data, or clinical cases implicating DDIT3 as causative for a Mendelian disorder.

- This summary sentence reiterates that ER stress is a major cause for some kidney diseases and highlights the ER stress response and autophagy as therapeutic targets. It supports the overall mechanistic relevance of ER stress pathways (including CHOP/DDIT3) to disease biology. Strength: reinforces the role of ER stress pathways in human disease contexts. Limitation: it remains a high-level conclusion about mechanism and therapy; no genetic or Mendelian-disease association data for DDIT3 are provided.


[Read Paper](https://www.semanticscholar.org/paper/b19abdade432293da2538ccacb3c18636be50923)


### Differential regulation of the unfolded protein response in outbred deer mice and susceptibility to metabolic disease

**Authors**: A. Havighorst (H-index: 3), H. Kiaris (H-index: 42)

**Relevance**: 0.25

**Bibliometric Impact**: 0.32144


**Excerpts**:

- Endoplasmic reticulum (ER) stress has been causatively linked to the onset of various pathologies.

- By using genetically diverse deer mice (Peromyscus maniculatus) as a model, we show that the profile of tunicamycin-induced UPR in fibroblasts isolated at puberty varies between individuals and predicts deregulation of lipid metabolism and diet-induced hepatic steatosis later in life.

- Among the different UPR targets tested, CHOP (also known as Ddit3) more consistently predicted elevated plasma cholesterol and hepatic steatosis.

- Compared with baseline levels or inducibility, the maximal intensity of the UPR following stimulation best predicts the onset of pathology.

- Our data suggest that the response to ER stress in cultured cells varies among individuals, and its profile early in life might predict the onset of ER-stress-associated disease in the elderly.

- Summary: By using genetically diverse deer mice, we show that the expression of different chaperones is highly coordinated in individual animals and its profile predicts the onset of metabolic pathology.


**Explanations**:

- How it relates: This sentence provides broad context that ER stress and its responses are implicated in disease. Relevance: mechanistic/contextual. Strength and limitations: It supports biological plausibility that genes involved in the ER stress response (such as DDIT3/CHOP) can contribute to pathology, but it is a general statement and does not provide any evidence that DDIT3 variants cause a Mendelian (monogenic) disorder.

- How it relates: This describes the experimental system (genetically diverse deer mice) and the main finding that an induced UPR profile in fibroblasts predicts later-life metabolic pathology. Relevance: mechanistic/phenotype-association (in an animal model). Strength and limitations: It shows that variation in UPR responses correlates with disease susceptibility, which supports plausibility that UPR components (including Ddit3) affect disease risk. However, this is an observational/predictive finding in mice, concerns complex traits (diet-induced steatosis), and does not demonstrate a Mendelian genotype–phenotype relationship or causal human genetic disease.

- How it relates: This is the most directly relevant sentence: it names CHOP/DDIT3 as the UPR target most consistently predicting adverse metabolic outcomes in this study. Relevance: direct association within the study (but in mice and as expression/predictor). Strength and limitations: This indicates Ddit3 expression correlates with disease phenotypes and therefore implicates the gene in pathological processes (mechanistic support). Crucially, it is not evidence that germline mutations in DDIT3 cause a Mendelian disease (no segregation, no mutation data, and phenotype is complex rather than a single-gene disorder). The association is predictive/correlative and in an animal model.

- How it relates: This gives nuance on what measurement correlated best with pathology (maximal UPR intensity). Relevance: mechanistic/phenotypic predictor. Strength and limitations: It suggests that a quantitative property of the UPR (which involves Ddit3/CHOP) relates to disease risk, strengthening mechanistic plausibility. It does not provide evidence of Mendelian inheritance or of pathogenic sequence variants in DDIT3 causing a monogenic disorder.

- How it relates: This states the authors' overall interpretation that early cellular ER stress responses vary and can predict later ER-stress-associated disease. Relevance: mechanistic/contextual. Strength and limitations: Supports the idea that variation in ER stress pathway genes (including Ddit3) can influence disease risk, but it remains correlative, in mice, and about complex disease prediction — not Mendelian causation or human clinical genetics evidence.

- How it relates: This summary reiterates that coordinated expression of chaperones predicts metabolic pathology in genetically diverse mice. Relevance: contextual/mechanistic. Strength and limitations: Reinforces that gene-expression variation in ER stress pathways is linked to metabolic disease risk, lending plausibility to the biological importance of components like DDIT3. Again, it does not provide evidence of DDIT3 being the cause of a Mendelian disease in humans or show segregation of DDIT3 variants with a monogenic phenotype.


[Read Paper](https://www.semanticscholar.org/paper/576a65507488b4034efe242553d8c0ff26216277)


### Loss-of-rescue of Ryr1I4895T-related pathology by the genetic inhibition of the ER stress response mediator CHOP

**Authors**: S. Germani (H-index: 2), E. Zito (H-index: 25)

**Relevance**: 0.25

**Bibliometric Impact**: 0.2494


**Excerpts**:

- It is shown that skeletal muscles of I4895T homozygous mice at birth present signs of stress of the endoplasmic reticulum (ER stress) and of the related unfolded protein response (UPR) with increased levels of the maladaptive mediators CHOP and ERO1, suggesting that genetic deletion of the ER stress response mediator CHOP is not sufficient to counteract the pathological Ryr1I 4895T phenotype.


**Explanations**:

- How it relates to the claim: This sentence states that CHOP (the protein product of DDIT3) is elevated in a mouse model of a Mendelian-like muscle disease (Ryr1I4895T) and that removing CHOP genetically did not rescue the pathological phenotype. This provides mechanistic evidence that DDIT3/CHOP is involved in the disease-associated stress response pathway in this model. It therefore bears on the plausibility that DDIT3 participates in Mendelian-disease-relevant biology, but it is not direct evidence that variation in DDIT3 itself causes a Mendelian disease.

Direct vs mechanistic: Mechanistic. The sentence describes altered expression of CHOP and its (non)sufficiency in modifying a disease phenotype in a mouse model, rather than reporting human genetic association or pathogenic variants in DDIT3.

Strengths and limitations: Strengths — shows in vivo upregulation of CHOP in a genetically driven disease model and tests functional consequence (genetic deletion) on phenotype. Limitations — the evidence is from a mouse model of RYR1-related disease, not from human genetic studies showing causative DDIT3 mutations; increased expression of DDIT3/CHOP in a disease state does not demonstrate that DDIT3 variants cause disease; the attempted genetic deletion of CHOP failing to rescue phenotype could weaken the hypothesis that CHOP is a causal driver (it may be a downstream marker or one of multiple mediators). Additionally, experimental conditions (neonatal mouse skeletal muscle, specific Ryr1I4895T mutation) limit generalizability to human Mendelian disease caused by DDIT3 variation.


[Read Paper](https://www.semanticscholar.org/paper/acd1a01c32b85f899b3491f94bff3bcfc3f471de)


## Other Reviewed Papers


### Calcium channels and channelopathies of the central nervous system

**Why Not Relevant**: The paper content provided is a review specifically about human and mouse calcium channelopathies of the central nervous system, focusing on channel genotypes and functional consequences of channel mutations. DDIT3 encodes a transcription factor (CHOP) and is not a calcium channel gene; the provided summary indicates the review is restricted to calcium channel genes and their mutations. Therefore the paper, as summarized, does not provide direct evidence (e.g., case reports, genetic linkage, or pathogenic variants) that DDIT3 is associated with a Mendelian disease, nor does it provide mechanistic evidence linking DDIT3 function to Mendelian phenotypes. Because the excerpt is a scope statement rather than data about DDIT3, relevance to the claim is effectively zero. Caveats: this conclusion is based only on the supplied summary sentence; if the full text contained an unrelated aside mentioning DDIT3, that is not included here. Also, absence of coverage in a calcium-channel-focused review does not prove that DDIT3 is not associated with Mendelian disease—only that this paper does not provide supporting evidence.


[Read Paper](https://www.semanticscholar.org/paper/727eb1ab77c95e461336e1f0e90f9550c1027f9d)


### Acidosis Activates Endoplasmic Reticulum Stress Pathways through GPR4 in Human Vascular Endothelial Cells

**Why Not Relevant**: The paper reports induction and functional participation of CHOP (C/EBP homologous protein), the protein product of DDIT3, in ER stress/UPR pathways in endothelial cells under acidic conditions. However, it contains no genetic, clinical, segregation, mutation, or population data linking DDIT3 variants to any inherited (Mendelian) disease. All evidence is cellular/mechanistic (expression and pathway activation) under specific experimental conditions (acidosis in cultured endothelial cells). Therefore, while the mechanistic data increase biological plausibility that DDIT3 could be relevant to disease processes, they do not provide direct evidence that DDIT3 is associated with at least one Mendelian disease. For the claim (association with a Mendelian disease) to be supported, evidence such as pathogenic DDIT3 variants segregating with disease in families, case reports of disease-causing mutations, gene-disease curation, or population-level enrichment of deleterious variants would be required; none of these are present in this paper.


[Read Paper](https://www.semanticscholar.org/paper/d41117c9a86f98a8729bacfb7e8f2437e64c695a)


### Monogenic forms of hypertension

**Why Not Relevant**: The supplied paper content consists of a single general statement: "Hormonal studies coupled with genetic testing can help in the early diagnosis of monogenic hypertensive disorders and identify children with hypertension due to single gene mutations inherited in an autosomal dominant or recessive fashion." This sentence does not mention DDIT3, any specific gene, any specific phenotype attributable to DDIT3, nor any mechanistic information linking DDIT3 to disease. Therefore it provides no direct evidence (no reports of pathogenic variants, cases, segregation, or clinical association) and no mechanistic evidence (no biochemical, molecular, or cellular pathway discussion) relevant to the claim that DDIT3 is likely associated with at least one Mendelian disease. Additional points about why this content is not relevant: 

- Scope: The statement is a broad methodological remark about using hormonal studies and genetic testing to detect monogenic hypertensive disorders. It addresses the general concept of single-gene causes of hypertension but does not name genes, gene families, or loci. 
- No gene-specific data: There are no variant descriptions, prevalence estimates, inheritance details tied to a specific gene, or phenotype lists that include DDIT3-related manifestations. 
- No mechanistic detail: The sentence contains no mechanistic pathway information (e.g., transcriptional regulation, protein function, cellular processes) that could plausibly connect DDIT3 to a Mendelian disorder. 
- No statistical or experimental evidence: There are no study methods, sample sizes, statistical results, or case reports provided that could be evaluated for significance or quality. 

Because the content is silent about DDIT3, it cannot be used to support or refute the claim that DDIT3 is likely associated with at least one Mendelian disease. To assess that claim, sources that specifically mention DDIT3 (e.g., case reports, gene-disease curation resources, functional studies demonstrating pathogenic variants and mechanistic consequences) would be required.


[Read Paper](https://www.semanticscholar.org/paper/01bd7a298bbb2628fa9d67f8953355fbe60e0cec)


### The Burden of Candidate Pathogenic Variants for Kidney and Genitourinary Disorders Emerging From Exome Sequencing

**Why Not Relevant**: The provided paper content does not mention the gene DDIT3 (also known as CHOP) anywhere in the text supplied. The paper's scope is a systematic evaluation of genes associated with Mendelian kidney and genitourinary disorders (identifying and curating a list of 625 genes) and the prevalence of candidate pathogenic variants in a cohort of ostensibly healthy adults. Because DDIT3 is not named in the supplied content and no gene-by-gene list or results that include DDIT3 are present, the paper provides no direct evidence for or against the claim that DDIT3 is likely associated with at least one Mendelian disease.

Relevant contextual points from the paper that explain why it cannot be used to assess DDIT3:
- The study specifically targeted genes associated with Mendelian kidney and genitourinary disorders identified via OMIM and Orphanet and then manually curated a set of genes; if DDIT3 is not associated with those phenotypes it would not be part of this analysis. The supplied text states the study focus and gene-identification approach but does not list the genes themselves.
- The cohort and analysis are tuned to detect variants in the curated set of kidney/genitourinary Mendelian genes; absence of DDIT3 in the supplied text means the paper does not address DDIT3’s disease associations even indirectly.

Limitations and caveats regarding interpretation:
- Absence of mention in this excerpt is not proof that DDIT3 is not associated with any Mendelian disease — it only means this paper (as provided) does not provide evidence either way. The paper’s negative relevance here is due to lack of content about DDIT3, not evidence disproving its involvement in Mendelian disease.

Recommended next steps to evaluate the claim:
- Query gene-disease resources such as OMIM, ClinVar, HGMD, Orphanet, and recent literature (PubMed) for entries linking DDIT3 to Mendelian disorders. Those sources are designed to directly answer whether DDIT3 has established Mendelian disease associations.


[Read Paper](https://www.semanticscholar.org/paper/a75e297208c04992568e607d19aa9b9aaebd09d5)


### Venezuelan Equine Encephalitis Virus Induces Apoptosis through the Unfolded Protein Response Activation of EGR1

**Why Not Relevant**: The paper describes transcriptomic changes in human astrocytoma cells infected with Venezuelan equine encephalitis virus and reports that DDIT3 (CHOP) expression is altered as part of the unfolded protein response. However, the claim concerns whether DDIT3 is likely associated with at least one Mendelian disease (i.e., caused by germline variants with inherited patterns and clinical phenotypes). This paper provides no human genetic variant data, no familial segregation, no clinical phenotypes, and no functional perturbations of DDIT3 germline alleles. All evidence is from an infection model (VEEV-infected cell lines and mouse embryonic fibroblasts), showing expression changes and pathway activation in response to viral ER stress. While these mechanistic data establish that DDIT3 participates in stress and apoptotic pathways (which may be mechanistically relevant to disease biology broadly), they do not provide direct or inferential evidence that DDIT3 is causally linked to any Mendelian disorder. Additional types of evidence would be required to support the claim: identification of pathogenic germline variants in DDIT3 in patients with a consistent inherited phenotype, segregation or de novo occurrence data, functional effects of disease-associated variants, or clinical case series linking DDIT3 alterations to Mendelian phenotypes. Given the absence of such evidence in this paper, its relevance to the Mendelian-disease association claim is very low.


[Read Paper](https://www.semanticscholar.org/paper/f4aa788ab898b28b00ee103e4d4ab24a2c684caf)


### FUS–DDIT3 Fusion Protein-Driven IGF-IR Signaling is a Therapeutic Target in Myxoid Liposarcoma

**Why Not Relevant**: The paper documents that DDIT3 is involved in a recurrent somatic chromosomal translocation (FUS–DDIT3) that drives myxoid liposarcoma and provides mechanistic evidence of how the fusion alters IGF-IR/PI3K/Akt signaling. However, Mendelian diseases are defined by heritable (germline) variants that segregate with disease in families according to Mendelian inheritance patterns. This manuscript focuses exclusively on somatic oncogenic rearrangements and functional effects in cancer cell lines and in vivo tumor models; it contains no data on germline DDIT3 variation, no reports of familial cases, and no genetic linkage or segregation analyses indicative of Mendelian inheritance. Therefore the paper does not provide direct evidence that DDIT3 is associated with any Mendelian disease. Given the complete absence of germline or familial genetic data, the relevance to the claim is minimal.


[Read Paper](https://www.semanticscholar.org/paper/a5a2702d01c2da27f313907784b83c88c00d0933)


### Fusion protein-driven IGF-IR/PI3K/AKT signals deregulate Hippo pathway promoting oncogenic cooperation of YAP1 and FUS-DDIT3 in myxoid liposarcoma

**Why Not Relevant**: The paper excerpt addresses a somatic fusion (FUS-DDIT3) and its mechanistic role in myxoid liposarcoma (MLS). Mendelian disease association requires evidence about germline variants, inheritance patterns, family segregation, or recurrent pathogenic variants causing a heritable phenotype. The provided content contains only somatic oncogenic mechanism information and no data on germline mutations, familial cases, or Mendelian phenotypes. Therefore it is not relevant to support the claim that DDIT3 is likely associated with at least one Mendelian disease.


[Read Paper](https://www.semanticscholar.org/paper/e3c603b7fc121b9b360e33d13452142c5cb9bd21)


### Frequency of Variants in Mendelian Alzheimer's Disease Genes within the Alzheimer's Disease Sequencing Project (ADSP)

**Why Not Relevant**: The paper exclusively reports analyses of variants in the Mendelian Alzheimer's genes PSEN2, PSEN1, and APP using ADSP WGS/WES data and curated clinical-database variants. There is no mention of the gene DDIT3, no data presented for DDIT3, and no mechanistic discussion linking DDIT3 to the genes or phenotypes studied. Therefore this paper provides no direct or mechanistic evidence supporting or refuting the claim that DDIT3 is likely associated with at least one Mendelian disease. Note that absence of evidence in this paper is not evidence of absence overall; other literature would need to be examined to evaluate the claim about DDIT3.


[Read Paper](https://www.semanticscholar.org/paper/3760efb72c8ec6adaf553ac773087fd2b16aeafc)


### Identification of variants in genes associated with hypertrophic cardiomyopathy in Mexican patients

**Why Not Relevant**: The provided paper content is not relevant to the claim that DDIT3 is likely associated with at least one Mendelian disease because it does not mention the DDIT3 gene (or any gene) at all. The single sentence only states that five previously unreported variants were found in Mexican patients with hypertrophic cardiomyopathy and notes uncertain significance in symptomatic patients. While HCM can be a Mendelian condition for some causal genes, establishing a Mendelian association requires gene-level evidence (e.g., identification of variants in DDIT3, segregation of variants with disease in families, de novo occurrence, functional assays demonstrating pathogenic mechanism, or replication in independent cohorts). None of those elements are present in the excerpt. Therefore the content cannot be used to support or refute the DDIT3–Mendelian disease claim. Additional details that would be required to change this assessment include explicit mention that the variants are in DDIT3, variant pathogenicity classifications, family segregation data, and mechanistic or functional evidence linking DDIT3 disruption to disease phenotypes.


[Read Paper](https://www.semanticscholar.org/paper/56c7d110dd7a0d150666b83aac1c846edc16f4da)


### PORCN‐related microphthalmia with limb anomalies: Case report and literature review

**Why Not Relevant**: This paper reports on germline variants in PORCN and a PORCN-related microphthalmia with limb anomalies (MLA) phenotype in a single family. The provided text repeatedly names PORCN (porcupine O-acyltransferase), focal dermal hypoplasia (Goltz-Gorlin syndrome), and describes a maternally inherited hemizygous PORCN variant (NM_203475.3: c.368T>G:p.(Met123Arg)). There is no mention of the gene DDIT3 (also known as CHOP or GADD153), no variants in DDIT3 are reported, and no mechanistic discussion that connects PORCN biology (Wnt lipidation/secretion) to DDIT3 function (an ER-stress–responsive transcription factor) appears in the text. Therefore the paper contains neither direct evidence (e.g., DDIT3 variants in affected individuals, segregation, case reports) nor mechanistic evidence (pathway links, functional data) that would support or refute the claim that DDIT3 is likely associated with a Mendelian disease. Additional limitations relevant to assessing DDIT3 from this paper: the study is a focused report on one gene in a single patient/family (limited sample size and scope), and the genetic findings are specific to PORCN (c.368T>G:p.(Met123Arg)), so the data cannot be generalized to infer associations for unrelated genes. Because no relevant content about DDIT3 exists in the provided paper excerpt, relevance to the claim is effectively zero and no verbatim supporting or refuting quotations can be extracted.


[Read Paper](https://www.semanticscholar.org/paper/d286aa5f405fc06d3b01ce0b4b36d05bcf7863ef)


### Monogenic Diabetes due to ABCC8/KCNJ11 Mutation: Case Study and Review of Literature

**Why Not Relevant**: The paper focuses on monogenic diabetes and specifically discusses ABCC8/KCNJ11 mutations and a family case linked to ABCC8. There is no mention of the gene DDIT3, no data, results, or mechanistic discussion involving DDIT3, and no statements that would support or refute an association between DDIT3 and any Mendelian disease. Therefore the paper provides no relevant evidence (direct or mechanistic) regarding the claim that DDIT3 is likely associated with at least one Mendelian disease. Absence of evidence in this paper does not imply DDIT3 is not associated with a Mendelian disease; it only means this paper cannot be used to evaluate that claim.


[Read Paper](https://www.semanticscholar.org/paper/bf2b44b94cefdd4e30d186260d907f3e3107c1e1)


### The simplest explanation does not have to be preferred: co-occurrence of pathogenic variants in cancer-predisposing genes

**Why Not Relevant**: The provided paper content does not mention the gene DDIT3, nor does it present data, results, or discussion about specific genes that could be interpreted as evidence for or against DDIT3's association with a Mendelian disease. The fragment only states an intention to review MINAS and cancer phenotypes and to analyze UK 10,000 Genomes Project participants with tumor-predisposition phenotypes, and to assess whether phenotypes are additive or synergistic across cancer susceptibility genes. Because the excerpt contains no references to DDIT3, no gene-level findings, no reported variants, no phenotypes tied to DDIT3, and no mechanistic descriptions involving DDIT3, it provides no direct or mechanistic evidence relevant to the claim. Therefore relevance is effectively zero. Note: absence of evidence in this excerpt is not evidence of absence in the full paper; a fuller document might mention DDIT3, but based solely on the provided content there is nothing to support or refute the claim.


[Read Paper](https://www.semanticscholar.org/paper/a1372a664bd2c33a0bda638975a8b26d112457fc)


### Novel mutation in SP2 in a Chinese pedigree with Neural tube defects

**Why Not Relevant**: The provided paper content does not mention DDIT3, CHOP, or GADD153 anywhere. All reported genetic findings, variant analyses, and mechanistic data concern SP2 (and FGFR1). Therefore the paper provides no direct genetic evidence, no variant-level data, and no mechanistic data linking DDIT3 to any Mendelian disease. Absence of mention in this single study is not proof that DDIT3 is not associated with Mendelian disease, but this paper offers no support for the claim that DDIT3 is likely associated with at least one Mendelian disease. The paper's methods and results are specific to NTDs and the genes SP2 and FGFR1; they cannot be used to infer associations for genes not assessed or reported (such as DDIT3).


[Read Paper](https://www.semanticscholar.org/paper/9c6ac7fb63b1a40b2c33bbffda9a4b081e76ac54)


### FUS::DDIT3 Fusion Protein in the Development of Myxoid Liposarcoma and Possible Implications for Therapy

**Why Not Relevant**: The paper content concerns DDIT3 as part of a recurrent somatic fusion (FUS::DDIT3) that drives myxoid liposarcoma (a cancer). The claim asks whether DDIT3 is likely associated with at least one Mendelian disease (an inherited, typically germline, monogenic disorder). The provided text contains no discussion of germline DDIT3 variants, inherited disease phenotypes, family segregation, penetrance, mode of inheritance, or case reports of Mendelian disorders involving DDIT3. All evidence is about somatic chromosomal translocation in tumors and therapeutic targeting of the fusion protein. Therefore the paper provides mechanistic and disease-association evidence relevant to somatic cancer but no direct or supporting evidence that DDIT3 is implicated in Mendelian (germline) disease. Because the content does not address germline inheritance or Mendelian disorders, its relevance to the claim is minimal.


[Read Paper](https://www.semanticscholar.org/paper/a2ecf8c9f5775e32fee43784d5e3f88f7b24b5e7)


### reportCoexistence of mal de Meleda and congenital cataract in a consanguineous Tunisian family : two case reports

**Why Not Relevant**: This paper does not mention DDIT3 (also known as CHOP) anywhere in the provided text. The study focuses on two autosomal recessive Mendelian disorders (mal de Meleda and congenital cataract) and investigates ARS (SLURP-1) and HSF4 genes in a Tunisian family, reporting an ARS C99Y homozygous mutation and negative testing for a recurrent HSF4 splice mutation. Therefore the paper provides no direct genetic association, case report, functional experiment, or mechanistic discussion that links DDIT3 to any Mendelian disease. Absence of mention in this paper does not disprove any association elsewhere, but it means this paper supplies no evidence for or against the claim that "DDIT3 is likely associated with at least one Mendelian disease."


[Read Paper](https://www.semanticscholar.org/paper/5022bbd5065624905bb7d7adc9bfd296cfb12b8b)


### Abstract 5631: YAP1 is a critical factor for FUS::DDIT3/BAF/c-Jun complex assembly in myxoid liposarcoma

**Why Not Relevant**: The provided paper content exclusively concerns the FUS::DDIT3 fusion protein in myxoid liposarcoma, a somatic chromosomal translocation event driving cancer. All experimental evidence is about protein–protein interactions, transcriptional dysregulation, and effects on tumor cell viability in MLS cell lines and specimens. There is no discussion of germline DDIT3 variants, inherited phenotypes, family segregation, incidence of congenital disorders, or any data linking DDIT3 to Mendelian inheritance. Therefore the paper provides mechanistic and functional evidence that DDIT3 (as part of a somatic fusion) is involved in cancer biology, but it provides no direct or genetic evidence that DDIT3 is associated with a Mendelian disease. Additional limitations reducing relevance: content is an abstract (limited detail), uses somatic cancer models, and focuses on a fusion protein (FUS::DDIT3) rather than variation in the endogenous DDIT3 gene.


[Read Paper](https://www.semanticscholar.org/paper/4b48a794130615c522b2b553ad40d6451b5b5592)


### Abstract PR004: Fusion protein-driven IGF-IR signals deregulate hippo pathway promoting oncogenic cooperation of YAP1 and FUS-DDIT3

**Why Not Relevant**: The paper exclusively describes somatic oncogenic biology of a FUS-DDIT3 fusion in myxoid liposarcoma, including mechanistic studies in tumor specimens, engineered mesenchymal stem cells, and MLS cell lines. There is no discussion of germline DDIT3 variants, family-based segregation, inherited phenotypes, penetrance, or clinical descriptions of Mendelian disorders. The experimental systems (tumor samples, somatic fusion, in vitro cell-line models) and the stated aims (MLS pathogenesis, oncogenic signaling) are focused on acquired cancer genetics rather than inherited disease. Therefore the paper does not provide direct evidence that DDIT3 is associated with any Mendelian disease. Mechanistically, while the data show that DDIT3 sequence participates in an oncogenic fusion and alters transcriptional programs, somatic oncogenic function does not imply germline Mendelian disease causation without separate evidence of pathogenic germline variants and inheritance patterns. For these reasons the content is of negligible relevance to the claim that "DDIT3 is likely associated with at least one Mendelian disease."


[Read Paper](https://www.semanticscholar.org/paper/504e4199995d56fb8f125d3a1736282d56dba399)


## Search Queries Used

- DDIT3 germline variant pathogenic Mendelian disease case report ClinVar OMIM

- CHOP GADD153 mutation human inherited autosomal dominant autosomal recessive monogenic congenital familial

- DDIT3 CHOP endoplasmic reticulum stress unfolded protein response apoptosis genetic disease susceptibility monogenic

- ETV6 DDIT3 fusion myxoid liposarcoma somatic translocation oncogenic fusion


## Usage Statistics

- Prompt Tokens: 0

- Completion Tokens: 0

- Total Tokens: 0

- Estimated Cost: $0.0822


## Bibliometric Configuration

- Use Bibliometrics: True

- Author Impact Weight: 0.4

- Citation Impact Weight: 0.4

- Venue Impact Weight: 0.2
