## Supplementary material for "MENDELSEEK: An algorithm that predicts Mendelian Genes and elucidates what makes them special": DDX43.txt

# Claim: The gene DDX43 is likely associated with at least one Mendelian disease.

**Status**: processed

**Overall Rating**: -1

**Explanation**:

No relevant papers were found that support or refute this claim.


## Relevant Papers


## Other Reviewed Papers


### The Burden of Candidate Pathogenic Variants for Kidney and Genitourinary Disorders Emerging From Exome Sequencing

**Why Not Relevant**: The provided paper content does not mention the gene DDX43, nor does it provide any data, case reports, variant annotations, or functional/mechanistic evidence specifically referring to DDX43. The manuscript excerpt focuses on incidental findings from exome/genome sequencing, methods for curation of genes associated with Mendelian kidney and genitourinary disorders, and general variant-filtering pipelines. Although it describes that a curated list of 625 genes was produced and that variant databases (HGMD, ClinVar) and OMIM/Orphanet were used, the actual gene list is not included in the provided content; therefore it is impossible from this excerpt to determine whether DDX43 was included among those genes or whether any variants in DDX43 were observed, reported, or functionally characterized. Because there is no direct mention or mechanistic discussion of DDX43 in the supplied text, the paper content as presented provides no direct or mechanistic evidence for or against the claim that DDX43 is likely associated with at least one Mendelian disease. To evaluate that claim would require either (a) the curated gene list or supplemental materials showing DDX43 inclusion and associated evidence, (b) variant-level reports (ClinVar/HGMD/OMIM entries) concerning DDX43 from this or other studies, or (c) case reports/segregation and functional studies directly implicating DDX43 in a Mendelian phenotype.


[Read Paper](https://www.semanticscholar.org/paper/a75e297208c04992568e607d19aa9b9aaebd09d5)


### TTC7A Variants Previously Described to Cause Enteropathy Are Observed on a Single Haplotype and Appear Non-pathogenic in Pediatric Inflammatory Bowel Disease Patients

**Why Not Relevant**: The provided paper content discusses variation in TTC7A and compound heterozygous variants observed in trans by Lawless et al., and explicitly assesses TTC7A's role in polygenic inflammatory bowel disease. Exact provided text: "This study describes the two variants observed in trans by Lawless et al. of compound heterozygote variants in the Tetratricopeptide Repeat Domain 7A gene, and assesses the evidence for variation in TTC7A contributing to polygenic IBD." There is no mention of the gene DDX43, no data or analysis concerning DDX43 variants, and no mechanistic discussion that links TTC7A to DDX43 or that would plausibly implicate DDX43 in Mendelian disease. Therefore, based on the supplied content, this paper provides no direct evidence (e.g., variant-disease co-segregation, case reports, statistical association) nor mechanistic evidence (e.g., functional studies, pathway links) supporting the claim that DDX43 is likely associated with at least one Mendelian disease. Limitations: the assessment is limited to the content you supplied; if the full paper contains additional sections not included here that mention DDX43, those were not available for review. Given only the provided text, relevance to the DDX43-Mendelian disease claim is effectively zero.


[Read Paper](https://www.semanticscholar.org/paper/3818a966b5d6e8b0eda65fdef043ac0727fa429c)


### Letter regarding “ SCN9A variant in a family of mixed breed dogs with congenital insensitivity to pain”—Navigating the pathogenicity of candidate gene mutations: Spotlight on paralog Nav genes

**Why Not Relevant**: The provided paper content exclusively discusses a variant in the SCN9A gene (Nav1.7) in dogs, its identification by whole-genome sequencing, and the authors' evaluation of its pathogenicity. There is no mention of the gene DDX43, no experimental data involving DDX43, no clinical cases linked to DDX43, and no mechanistic pathways involving DDX43 described anywhere in the text. Because the claim concerns whether DDX43 is likely associated with at least one Mendelian disease, and this document contains no information about DDX43, it provides neither direct evidence (no association data, no cases, no segregation, no functional assays) nor mechanistic evidence (no biochemical, cellular, or physiological data involving DDX43) relevant to that claim. The paper does include general discussion about standards for gene–disease interpretation (e.g., the need for family segregation, functional analysis, population frequency data), which could inform how one would evaluate evidence for any gene, including DDX43; however, those methodological remarks are generic and do not constitute evidence that DDX43 is associated with a Mendelian disease. Therefore the content is not relevant to the claim.


[Read Paper](https://www.semanticscholar.org/paper/39dfc4195578b71ad782115a3a552b9b9e2dae03)


### Autosomal Dominant Intellectual Development Disorder-6 (MRD6) Without Seizures Linked to a De Novo Mutation in the grin2b Gene Revealed by Exome Sequencing: A Case Report of a Moroccan Child

**Why Not Relevant**: The paper's entire presented content pertains to a GRIN2B (grin2b) nonsense variant causing MRD6 in a single patient and to the clinical and interpretive evidence supporting that association (de novo status, absence from gnomAD, ACMG pathogenic classification, and predicted deleterious effect). Nowhere in the provided text is DDX43 mentioned, discussed, or evaluated. Therefore there is no direct evidence (case data, genetic association, segregation) nor mechanistic evidence (functional studies, pathway links) in this paper that bears on whether DDX43 is associated with any Mendelian disease. The study design (single-case exome report) also would not be a suitable source to establish or refute gene-disease associations for genes not examined in the report. In short, relevance to the claim about DDX43 is effectively zero because the paper addresses a different gene and disorder entirely; absence of DDX43 in this report does not inform DDX43's disease associations in the literature at large.


[Read Paper](https://www.semanticscholar.org/paper/67d756e0cc302e5d46fdda9df34403f1f2de9a5a)


### Explorer CDKL 5 variants : improving our understanding of a rare neurological disorder

**Why Not Relevant**: This paper is focused entirely on the CDKL5 gene and CDKL5 deficiency (a rare X-linked neurologic disorder). The provided content discusses CDKL5 variant curation, ExAC population data for CDKL5, pathogenicity assignment criteria, minigene splicing assays for CDKL5 exon 14, and conclusions about CDKL5 isoforms and catalytic domain. The text contains no mention of the gene DDX43, its variants, expression, functional studies, clinical case reports, or any other information that would bear on whether DDX43 is associated with a Mendelian disease. Therefore it provides no direct evidence for or against the claim about DDX43. 

Note on indirect/methodological relevance: The paper describes general approaches (use of large-scale population databases such as ExAC, computational prediction tools, variant classification criteria, and minigene splicing assays) that are transferable methods for assessing whether any gene’s variants are likely pathogenic. However, methodological transferability is not evidence that DDX43 itself is associated with a Mendelian disease. If you want, I can (1) search the literature and databases (OMIM, ClinVar, gnomAD/ExAC, PubMed) for evidence linking DDX43 to Mendelian disease and summarize direct and mechanistic evidence, or (2) apply the same analytical framework to any available DDX43 variant data you provide.


[Read Paper](https://www.semanticscholar.org/paper/a959fe9191159fa8a508c918a1e4e7dcf60553fd)


## Search Queries Used

- DDX43 mutation Mendelian disease case report family segregation pathogenic variant ClinVar OMIM disease association

- DDX43 loss of function constraint gnomAD pLI observed expected LoF allele frequency population database variant pathogenicity

- DEAD box helicase 43 DDX43 HAGE knockout mouse zebrafish phenotype developmental disorder spermatogenesis infertility functional study RNA helicase mechanism

- DDX43 exome sequencing rare disease cohort trio undiagnosed disease program gene discovery germline variant hereditary cancer predisposition somatic mutation cancer testis antigen


## Bibliometric Configuration

- Use Bibliometrics: True

- Author Impact Weight: 0.4

- Citation Impact Weight: 0.4

- Venue Impact Weight: 0.2
