## Supplementary material for "MENDELSEEK: An algorithm that predicts Mendelian Genes and elucidates what makes them special": DHX40.txt

# Claim: The gene DHX40 is likely associated with at least one Mendelian disease.

**Status**: processed

**Overall Rating**: -1

**Explanation**:

No relevant papers were found that support or refute this claim.


## Relevant Papers


## Other Reviewed Papers


### Simultaneous identification and prioritization of variants in familial, de novo, and somatic genetic disorders with VariantMaster

**Why Not Relevant**: The provided paper content does not mention the gene DHX40, any variants in DHX40, or any results implicating specific genes in Mendelian diseases. The text focuses on the VariantMaster software: its algorithmic approach, integration with annotation tools, and general proof-of-concept applications (including an unspecified 'rare Mendelian genetic disease'). Because there is no gene-level data, no named disease–gene associations, and no variant lists or statistics relevant to DHX40, the paper provides no direct or gene-specific mechanistic evidence to support or refute the claim that DHX40 is likely associated with at least one Mendelian disease. It is possible (but unsubstantiated here) that DHX40 could have been among genes analyzed, but without explicit mention or data one cannot infer association. Therefore relevance is rated very low.


[Read Paper](https://www.semanticscholar.org/paper/a56e2dc7369c76b03ad5e9309093beb98a159719)


### Kindreds of dominantly inherited Parkinson's disease: Keys to the riddle

**Why Not Relevant**: This paper does not provide evidence for or against the specific claim that DHX40 is likely associated with at least one Mendelian disease. The text focuses on familial aggregation of Parkinson's disease, the general plausibility that single-gene (monogenic) causes exist for familial PD, and the value of linkage studies and pedigrees for gene discovery. Nowhere in the provided content is the gene name DHX40 (or any close variant) mentioned, nor are there any locus mapping results, sequencing data, statistical associations, or functional experiments that implicate any particular gene, let alone DHX40. The excerpts above only supply general mechanistic or methodological context for gene discovery in familial PD; they do not provide direct empirical support for DHX40 being causal in any Mendelian disease. Given the lack of any mention or data concerning DHX40, the paper is not relevant to the claim about that gene. Additional, specific evidence (e.g., linkage to the chromosomal location of DHX40, segregation of DHX40 variants with disease in pedigrees, case-control or population studies, and/or functional validation) would be required to evaluate the claim.


[Read Paper](https://www.semanticscholar.org/paper/b47c5b5c2c233ecbd785bd33340dff1c297bb8c2)


### Chloroplast RH3 DEAD-box RNA helicases in Zea mays and Arabidopsis thaliana function in splicing of specific group II introns and affect chloroplast ribosome biogenesis

**Why Not Relevant**: The paper studies plant (angiosperm) chloroplast DEAD-box RNA helicases, primarily AtRH3 in Arabidopsis and its maize co-orthologs (ZmRH3A/ZmRH3B). It contains no mention of the human gene DHX40, no human or clinical genetic data, and no discussion of Mendelian disease. The mechanisms described (group II intron splicing, assembly of the 50S plastid ribosomal particle, chloroplast biogenesis) are specific to plastids in plants and are not directly informative about human nuclear or mitochondrial RNA-processing pathways. Although DEAD-box family membership implies distant mechanistic relationships across taxa, the paper provides neither orthology analysis to human DHX40 nor any genetic or phenotypic evidence in humans or model systems that would support an association between DHX40 and a Mendelian disease. Therefore this paper is not relevant to assessing the claim that DHX40 is likely associated with at least one Mendelian disease.


[Read Paper](https://www.semanticscholar.org/paper/0d644ccf52b7899c032aad55ab06501d14bf3e40)


### A rare splice-site variant in cardiac troponin-T (TNNT2): The need for ancestral diversity in genomic reference datasets

**Why Not Relevant**: The provided paper text does not mention DHX40 and therefore contains no direct or mechanistic evidence relevant to the claim that "DHX40 is likely associated with at least one Mendelian disease." The paper instead focuses on a variant in TNNT2 (cardiac troponin T) and its population frequency, splice effect, and clinical classification. For example, the manuscript states: "We report a rare variant in the cardiac troponin T gene, TNNT2; NM_001001430.3: c.571-1G>A (rs483352835) identified via research-based whole exome sequencing in two unrelated probands of Oceanian ancestry with cardiac phenotypes." It also gives mechanistic detail specific to TNNT2: "The variant disrupts the canonical splice acceptor site, activating a cryptic acceptor and resulting in an in-frame deletion (p.Gln191del)." Finally, the authors conclude about that TNNT2 variant: "Based on these data, we classify this variant as benign, and conclude it is not a monogenic cause of disease." These statements make clear the gene, variant, mechanisms, and conclusions described all pertain to TNNT2, not DHX40. While the paper contains general points about variant interpretation and population representation that are broadly applicable to Mendelian gene discovery (e.g., higher VUS rates in non-European ancestries and the need for diverse reference datasets), those methodological points do not provide evidence for or against DHX40 being implicated in a Mendelian disease. Limitations/caveats: this assessment is based solely on the text you provided; if the full paper or supplementary materials (not included here) mention DHX40, those sections were not available for review. If you want an evidence-based evaluation of DHX40 specifically, I can search ClinVar, OMIM, PubMed, and gnomAD for reported pathogenic variants, disease associations, functional studies, and population frequency data for DHX40.


[Read Paper](https://www.semanticscholar.org/paper/4fed6e1e7cc6124f24c814ee86964f97c3ffaa39)


### Case Report: Identification of a rare nonsense mutation in the POC1A gene by NGS in a diabetes mellitus patient

**Why Not Relevant**: This paper presents a clinical case and molecular analysis attributing the patient's diabetes to a pathogenic truncating mutation in POC1A. The text supplied contains no mention of the gene DHX40, no reported variants in DHX40, and no experimental or mechanistic data involving DHX40. Although the study used trio whole-exome sequencing (a method that could detect coding variants in DHX40), the authors report only the POC1A finding and related clinical correlation; they do not report negative or positive results for DHX40. Therefore the paper provides no direct evidence supporting the claim that DHX40 is associated with a Mendelian disease. The absence of DHX40 in the report cannot be taken as strong evidence that DHX40 is not associated with disease (because a lack of reporting is not equivalent to a systematic negative result), but it does mean this paper offers no supportive or mechanistic evidence for the claim. Key limitations for using this paper as evidence regarding DHX40: (1) the study's aim, findings, and conclusions focus exclusively on POC1A; (2) no DHX40 data are presented; and (3) even though WES was performed, WES has known blind spots (structural variants, repeat expansions, deep intronic variants, low-level mosaicism) and the authors' filtering/reporting choices could omit unrelated variants. In sum, relevance to the claim is effectively zero because the paper does not address DHX40 at all.


[Read Paper](https://www.semanticscholar.org/paper/8f90a0a9b3c51d0406ae0337e2a4d8c2c0204107)


### A Novel Missense Mutation in the TGF-β-binding Protein-Like Domain 3 of FBN1 Causes Weill–Marchesani Syndrome with Intellectual Disability

**Why Not Relevant**: The paper focuses entirely on Weill–Marchesani syndrome (WMS) and reports a novel heterozygous FBN1 mutation causing autosomal dominant WMS in a single consanguineous family. The genes discussed and implicated in the study are FBN1 (dominant) and ADAMTS10/ADAMTS17/LTP2 (recessive). There is no mention of DHX40 anywhere in the provided text, and no experimental, genetic, segregation, functional, or mechanistic data concerning DHX40 are presented. Therefore this paper provides no direct evidence that DHX40 is associated with any Mendelian disease, nor does it provide mechanistic plausibility linking DHX40 to disease. Limitations of the paper (single family, focus on FBN1, WES targeted to proband with co-segregation of FBN1 variant) further confirm that it cannot inform the claim about DHX40. Because the content is entirely about other genes, relevance to the DHX40 claim is effectively zero.


[Read Paper](https://www.semanticscholar.org/paper/c998af9708683d9ea84961bc2527db3cfd0407ae)


### Adrenal function and MC1R gene analysis in a prepubertal girl with generalized hyperpigmentation: case report

**Why Not Relevant**: This paper provides no direct or indirect evidence regarding the gene DHX40. The manuscript focuses on a single clinical case of generalized hyperpigmentation and on sequencing and analysis of MC1R (and discussion of other pigmentation-related genes such as HERC2). There is no mention of DHX40 anywhere in the provided text, no testing of DHX40 (no sequencing, no variant discussion), and no mechanistic pathways or results that implicate DHX40 in pigmentation or any Mendelian disorder. Methodological scope is restricted to targeted analysis of MC1R (and discussion of other pigmentation genes), so absence of DHX40 is expected but does not provide evidence for or against an association. Key limitations: (1) single-patient case report (n=1), so even for genes examined the sample size is tiny; (2) genetic testing described was targeted to MC1R ("We did not find any mutations within the whole MC1R gene."), not exome or genome sequencing that could detect DHX40 variants; (3) the paper's conclusions pertain to pigmentation genes and adrenal/ACTH mechanisms (MC1R, MC2R, HERC2), not to DHX40. Therefore relevance to the claim that "DHX40 is likely associated with at least one Mendelian disease" is effectively zero based on this paper.


[Read Paper](https://www.semanticscholar.org/paper/cf0b637b0c02cc8b1ee679a9945b3290946babdd)


### reportCoexistence of mal de Meleda and congenital cataract in a consanguineous Tunisian family : two case reports

**Why Not Relevant**: This paper does not provide evidence supporting or refuting the claim that DHX40 is associated with a Mendelian disease. The study is a case report of a Tunisian family with Mal de Meleda and congenital cataract and presents targeted genetic testing of the ARS (SLURP-1) gene and a recurrent HSF4 splice mutation. The authors identify a homozygous C99Y mutation in ARS and report the absence of the recurrent HSF4 splice mutation. There is no mention of the DHX40 gene, no sequencing or analysis of DHX40, and no mechanistic discussion implicating DHX40. Because the study used candidate-gene/targeted testing rather than exome/genome-wide approaches, the absence of any DHX40 data reflects study design and scope rather than evidence about DHX40's role in Mendelian disease. Consequently, the paper is essentially irrelevant to the claim that DHX40 is likely associated with at least one Mendelian disease.


[Read Paper](https://www.semanticscholar.org/paper/5022bbd5065624905bb7d7adc9bfd296cfb12b8b)


### Novel de Novo Nonsense Variants in AGO3 and KHSRP: Insights into Global Developmental Delay and Autism Spectrum Disorders through Whole Genome Analysis

**Why Not Relevant**: The provided paper content focuses on a single case report identifying novel de novo nonsense variants in AGO3 and KHSRP and discusses those genes in relation to the patient's global developmental delay and autistic features. The text contains no mention of DHX40, no variants in DHX40, and no mechanistic data or discussion pertaining to DHX40. Therefore this paper provides no direct genetic evidence (no reported DHX40 variants, segregation, case series, or statistical association) and no mechanistic evidence (no functional studies, expression data, or pathway links) supporting or refuting an association between DHX40 and any Mendelian disease. Additional limitations relevant to interpreting the paper itself (single patient, candidate gene nomination without functional follow-up) further mean it cannot be repurposed to infer an association for an unrelated gene (DHX40). Consequently, the paper is not relevant to the claim that DHX40 is likely associated with at least one Mendelian disease.


[Read Paper](https://www.semanticscholar.org/paper/5aba59259e6535c76aa2fd80054cb774eb3d7452)


## Search Queries Used

- DHX40 Mendelian disease monogenic inherited autosomal dominant autosomal recessive familial congenital disorder case report pedigree segregation

- DHX40 variant pathogenic ClinVar OMIM gnomAD exome sequencing whole genome sequencing trio sequencing rare variant report clinical genetics

- DHX40 function RNA helicase DEAH box DExH box ATP dependent helicase RNA binding splicing ribonucleoprotein mouse knockout zebrafish Drosophila functional study

- DHX40 loss of function intolerance pLI constraint gnomAD population frequency rare variant burden exome cohort case control segregation linkage functional assay pathogenicity prediction


## Bibliometric Configuration

- Use Bibliometrics: True

- Author Impact Weight: 0.4

- Citation Impact Weight: 0.4

- Venue Impact Weight: 0.2
