## Supplementary material for "MENDELSEEK: An algorithm that predicts Mendelian Genes and elucidates what makes them special": DLG5.txt

# Claim: The gene DLG5 is likely associated with at least one Mendelian disease.

**Status**: processed

**Overall Rating**: -1

**Explanation**:

No relevant papers were found that support or refute this claim.


## Relevant Papers


## Other Reviewed Papers


### Genetic variation in DLG5 is associated with inflammatory bowel disease

**Why Not Relevant**: The paper excerpt reports an association between a DLG5 variant and Crohn disease and suggests gene-gene interaction with CARD15, but Crohn disease is characterized here as a polygenic disorder. The claim asks whether DLG5 is likely associated with at least one Mendelian disease (i.e., a monogenic disorder with Mendelian inheritance). The provided content contains no evidence of Mendelian (monogenic) causation: there are no family segregation data, no report of a highly penetrant pathogenic variant, no functional studies demonstrating a causal loss- or gain-of-function consistent with Mendelian inheritance, and no statistical or clinical details that would support classifying DLG5 as a Mendelian disease gene. Therefore the excerpt is not relevant to supporting the specific claim that DLG5 is likely associated with a Mendelian disease.


[Read Paper](https://www.semanticscholar.org/paper/59bd02011abaeac745eebcd7ffb3d68e37cdf586)


### Genome-wide association study of Crohn's disease in Koreans revealed three new susceptibility loci and common attributes of genetic susceptibility across ethnic populations

**Why Not Relevant**: The paper is a genome-wide association study of Crohn's disease in a Korean population, focused on identifying common-variant susceptibility loci for a complex, polygenic trait. The abstract (the provided content) names the loci found and investigated (e.g., TNFSF15, IL23R, MHC, RNASET2-FGFR1OP-CCR6, ATG16L2, FCHSD2) and does not mention the gene DLG5 or any Mendelian (monogenic) disease associations. Key reasons this paper is not relevant to the claim that "DLG5 is likely associated with at least one Mendelian disease":
- Absence of evidence: DLG5 is not reported among the replicated or novel loci; absence of DLG5 in this GWAS provides no positive support for the claim.
- Study design limitation: GWAS targets common variants influencing complex traits; it is not optimized to discover rare, highly penetrant variants that cause Mendelian diseases. Even if DLG5 were implicated in a monogenic disorder, this study would likely miss that association.
- Phenotype mismatch: The paper studies Crohn's disease (a complex IBD), not Mendelian disorders. Association of a gene with CD does not equate to association with a Mendelian disease.
Therefore, because the study does not mention DLG5 and is not designed to detect Mendelian disease genes, its relevance to assessing whether DLG5 is likely associated with a Mendelian disease is negligible. That is why relevance is scored very low (0.02) rather than absolute zero: the paper's negative/absence information and the methodological description provide limited indirect context but no substantive evidence for or against a DLG5–Mendelian disease link.


[Read Paper](https://www.semanticscholar.org/paper/195873c899176891e4215078291efa6bffaf0e1a)


### The Burden of Candidate Pathogenic Variants for Kidney and Genitourinary Disorders Emerging From Exome Sequencing

**Why Not Relevant**: The provided paper content does not mention the gene DLG5 anywhere in the quoted text, nor does it present gene-level results that would allow assessment of whether DLG5 is associated with a Mendelian disease. The excerpts describe methods (databases searched, curation workflow, cohort, and annotation pipelines) and note that a curated list of 625 genes was produced, but the specific gene list and gene-level findings are not included in the supplied content. Consequently, there is no direct evidence (e.g., reported pathogenic variants, statistical associations, segregation, case reports) or mechanistic evidence (e.g., functional studies, predicted loss-of-function intolerance specific to DLG5) in the provided text to support or refute the claim that DLG5 is likely associated with at least one Mendelian disease. If the full paper or the referenced laboratory web site (where the final gene list was said to be accessible) were available, it might be possible to determine whether DLG5 is included among the curated genes and whether any evidence is reported; however, based solely on the supplied content, the claim cannot be evaluated.


[Read Paper](https://www.semanticscholar.org/paper/a75e297208c04992568e607d19aa9b9aaebd09d5)


### Genetic Determinants of Pediatric Inflammatory Bowel Disease: Is Age of Onset Genetically Determined?

**Why Not Relevant**: The supplied paper content does not mention the gene DLG5 at all and contains no data—genetic, segregation, variant-level, or functional—pertaining to DLG5. The paper describes genome-wide association studies and analyses of polygenic risk and age-of-onset in inflammatory bowel disease, which are studies of common-variant contributions to a complex trait rather than investigations of high-penetrance, typically rare variants that would establish a gene as causing a Mendelian disease. Establishing a Mendelian disease association normally requires evidence such as rare damaging variants co-segregating with disease in families, demonstration of de novo or biallelic mutations in affected individuals with appropriate inheritance patterns, strong effect sizes, and often functional validation; none of these types of evidence are present in the provided content. Therefore this paper content provides no direct or mechanistic evidence for DLG5 being associated with a Mendelian disease. Additional limitations: the methods and results discussed focus on GWAS in pediatric versus adult cohorts and on discovery of small-effect loci (e.g., TNFRSF6 and PSMG1), negative/nominal associations for age-of-onset, and plans for larger GWAS meta-analyses — none of which are informative about Mendelian causality for any specific gene, including DLG5. Given the absence of any mention of DLG5 and the mismatch between study design (complex-trait GWAS) and the type of evidence required for Mendelian gene-disease associations, the relevance of this paper content to the claim is effectively zero.


[Read Paper](https://www.semanticscholar.org/paper/e6cbe09d51890193b9be408dc2552c45e0ced9b3)


### Precision Medicine in Internal Medicine

**Why Not Relevant**: The provided paper text is a general review of genetic testing strategies, exome/genome sequencing, and variant interpretation. It does not mention the gene DLG5 anywhere, nor does it present gene-specific data, case reports, linkage/segregation evidence, functional studies, or ClinVar/OMIM entries for DLG5. Therefore it contains no direct evidence (genetic, clinical, or functional) that DLG5 is associated with any Mendelian disease. The excerpts included convey only general context (e.g., how many genes currently have known Mendelian associations, what ES typically finds, and that ES can discover novel gene–disease links). Those contextual points modestly inform the background probability and mechanisms by which a gene might be discovered as disease-associated, but they do not provide specific support or refutation for the claim about DLG5. Because the paper lacks any DLG5-specific data, its relevance to the claim is minimal.


[Read Paper](https://www.semanticscholar.org/paper/c7199e8a5226f8d9d55777da0e15e91797bb459a)


### Familial Whole Exome Sequencing Study of 30 Families With Early-Onset High Myopia

**Why Not Relevant**: This paper does not provide evidence for or against the claim that DLG5 is likely associated with at least one Mendelian disease. The provided content (purpose, results, and conclusions) lists the genes detected and analyzed in this eoHM cohort (for example COL4A5, NYX, CACNA1F, CCDC111, SLC39A5, P4HA2, CPSF1, GRM6) and reports proportions of families with inherited retinal disease-associated genes, but does not mention DLG5 anywhere. Therefore there is no direct evidence (no reported DLG5 variants, segregation data, or phenotype linkage) and no mechanistic evidence concerning DLG5 in this paper. Limitations to this assessment: absence of DLG5 in the paper is only evidence that DLG5 was not observed/reported in this specific eoHM cohort and does not constitute evidence that DLG5 is unrelated to Mendelian disease in other cohorts or phenotypes. To evaluate the claim about DLG5, one would need studies that specifically report pathogenic DLG5 variants with segregation/functional evidence or mechanistic data linking DLG5 perturbation to a Mendelian phenotype.


[Read Paper](https://www.semanticscholar.org/paper/8a51e1e2b13da3e9075f4d54e506e745280ab56b)


### A pathogenic variant in RAB32 causes autosomal dominant Parkinson’s disease and activates LRRK2 kinase

**Why Not Relevant**: The provided paper content concerns RAB GTPases (specifically RAB32 Ser71Arg) and its association with autosomal dominant Parkinson disease and LRRK2 kinase activation. The text makes no mention of the gene DLG5, nor any findings, variants, segregation, mechanistic data, or clinical/pathological associations involving DLG5. Key sentences that demonstrate the paper's focus include: "To explore the role of RAB GTPases in PD, we undertook a comprehensive examination of their genetic variability in familial PD," and "We found RAB32 c.213C>G (Ser71Arg) to co-segregate with autosomal dominant parkinsonism in three multi-incident families." Because DLG5 is neither listed among the genes studied (the study targeted 61 RAB GTPases) nor discussed in any mechanistic or clinical context in the provided content, there is no direct evidence (genetic segregation, case reports, statistical association) nor mechanistic evidence in this paper that would support or refute the claim that "DLG5 is likely associated with at least one Mendelian disease." The absence of any mention of DLG5 in the methods, results, interpretation, or conclusions means this paper cannot be used to evaluate the claim. Note: this assessment is limited to the supplied paper content; if the full manuscript contains sections not provided here that mention DLG5, those were not available for review.


[Read Paper](https://www.semanticscholar.org/paper/a36d28e828ede7c87c650af5f5117982b9749e7b)


### OP11 Expanded genome-wide association study of Inflammatory Bowel Disease identifies 174 novel loci and directly implicates new genes in disease susceptibility

**Why Not Relevant**: The provided paper content does not mention DLG5 anywhere and therefore provides no direct evidence that DLG5 is associated with a Mendelian disease. The excerpts above only supply indirect, general evidence that GWAS loci can overlap genes implicated in Mendelian syndromes (in this study the authors list CARMIL2, DOCK8, G6PC3, HPS4, NCF1, and PIK3CD), and a mechanistic rationale that common non-coding variants can dysregulate Mendelian IBD genes. Because DLG5 is not named or discussed, the paper cannot be used to support or refute the specific claim about DLG5. If you need a direct assessment of DLG5, a search of papers or databases that specifically report Mendelian associations (e.g., OMIM, ClinVar, gene-specific case series or sequencing studies) would be required.


[Read Paper](https://www.semanticscholar.org/paper/9119bf9558131dc6529c1320f4b9127531312a86)


### Whole exome sequencing revealed a novel homozygous variant in the DGKE catalytic domain: a case report of familial hemolytic uremic syndrome

**Why Not Relevant**: The provided paper content discusses DGKE and its possible role in aHUS; it does not mention DLG5. DLG5 and DGKE are distinct genes with different sequences, functions, and disease associations. Therefore the content does not provide direct evidence for the claim that DLG5 is likely associated with at least one Mendelian disease, nor does it provide mechanistic evidence about DLG5. Additionally, the excerpt is speculative about DGKE ("It is suggested" / "might lead"), lacks experimental details, and contains no statistical or clinical data that could be repurposed to support a claim about DLG5. Because of the gene mismatch and absence of any reference to DLG5, the relevance to the claim is effectively zero.


[Read Paper](https://www.semanticscholar.org/paper/2691c9dc3e9f16f506828ae781284158953cd6a0)


### Screening Mutations of the Monogenic Syndromic High Myopia by Whole Exome Sequencing From MAGIC Project

**Why Not Relevant**: The paper focuses specifically on monogenic syndromic high myopia (msHM) and reports pathogenic variants in 84 genes (with examples provided) identified from a targeted panel of 298 msHM-related genes. Nowhere in the provided content is the gene DLG5 mentioned, nor are mechanistic data or gene lists including DLG5 presented. Therefore this paper provides no direct evidence that DLG5 is associated with any Mendelian disease, nor does it provide mechanistic evidence implicating DLG5. The only weakly relevant information is that a targeted panel of genes was assessed and certain genes were implicated; but without either an explicit mention of DLG5 in the panel or in the results, the study does not address the claim. Additional limitations: the excerpt does not provide the full 298-gene list or supplementary tables, so it is possible (but unproven here) that DLG5 was not included or was included but not found to be implicated. Given the lack of any mention of DLG5 or relevant mechanisms in the provided text, the paper's relevance to the claim is negligible.


[Read Paper](https://www.semanticscholar.org/paper/339423d81112e62718cf1ec21a06909ee07f9afe)


### THE JAPANESE TERATOLOGY SOCIETY 62ND ANNUAL MEETING

**Why Not Relevant**: The provided paper content does not mention the gene DLG5, nor does it present any genetic association data, case reports, sequencing results, loss-of-function or gain-of-function experiments, segregation analyses, or phenotype descriptions linked to DLG5 that would bear on its association with a Mendelian disease. The paper focuses on organ development, Shh, Satb2, Wnt signaling, LIF/CRH signaling, CRISPR genome-editing caveats (illustrated with Il2rg/X-SCID), and teratogenic models (including zebrafish). Because there is no direct discussion or mechanistic connection to DLG5 in the text, there is no evidence here to support or refute the claim that DLG5 is likely associated with at least one Mendelian disease.


[Read Paper](https://www.semanticscholar.org/paper/162811312bde34697100a4eaab4c8748157e2132)


## Search Queries Used

- DLG5 pathogenic variant Mendelian disease case report familial segregation whole exome sequencing monogenic

- DLG5 ClinVar OMIM HGMD pathogenic loss of function haploinsufficiency gene disease association variant classification

- DLG5 knockout mouse phenotype developmental defect epithelial polarity organ development congenital anomaly zebrafish functional study

- DLG5 association Crohn disease inflammatory bowel disease GWAS common variant association replication study complex disease versus monogenic explanation


## Bibliometric Configuration

- Use Bibliometrics: True

- Author Impact Weight: 0.4

- Citation Impact Weight: 0.4

- Venue Impact Weight: 0.2
