## Supplementary material for "MENDELSEEK: An algorithm that predicts Mendelian Genes and elucidates what makes them special": DNAH6.txt

# Claim: The gene DNAH6 is likely associated with at least one Mendelian disease.

**Status**: processed

**Overall Rating**: -1

**Explanation**:

No relevant papers were found that support or refute this claim.


## Relevant Papers


## Other Reviewed Papers


### The Burden of Candidate Pathogenic Variants for Kidney and Genitourinary Disorders Emerging From Exome Sequencing

**Why Not Relevant**: The provided paper content does not mention the gene DNAH6 anywhere in the excerpts supplied. The paper's focus is on assembling and curating a list of genes associated with Mendelian kidney and genitourinary disorders, annotating them, and assessing the prevalence of candidate pathogenic variants in a large control cohort. Although the methods describe using OMIM, Orphanet, ClinVar, HGMD, and a curated list of 625 genes, the actual gene list and any gene-specific results (including whether DNAH6 is included, excluded, or has pathogenic variants) are not provided in the supplied text. Therefore, no direct evidence (e.g., reported pathogenic variants in DNAH6, statistical association with a Mendelian phenotype, or mechanistic functional data implicating DNAH6) is present in the content you supplied. Because the claim specifically concerns DNAH6, and the paper content does not reference that gene or present gene-level findings, the paper as given cannot be used to support or refute the claim that DNAH6 is likely associated with at least one Mendelian disease.


[Read Paper](https://www.semanticscholar.org/paper/a75e297208c04992568e607d19aa9b9aaebd09d5)


### Contribution of rare transmitted and de novo variants among 2 , 871 1 congenital heart disease probands

**Why Not Relevant**: The supplied paper excerpt makes no mention of the gene DNAH6, does not report variants in DNAH6, and contains no mechanistic or statistical data linking DNAH6 to any Mendelian disease. The content focuses on exome sequencing in congenital heart disease (CHD) probands, analysis of de novo and recessive genotypes in known CHD genes, and enrichment statistics for curated CHD gene sets; none of the visible sentences, tables, or paragraphs reference DNAH6. Therefore the excerpt provides no direct or mechanistic evidence for or against the claim that DNAH6 is likely associated with a Mendelian disease. Caveat: only the provided excerpt was analyzed — DNAH6 could appear elsewhere in the full paper (for example in Supplementary Excel files or later sections not included here); absence from this excerpt does not rule out discussion elsewhere in the full publication or in other literature databases (OMIM, ClinVar, PubMed).


[Read Paper](https://www.semanticscholar.org/paper/e5830d9e95469f22518560007746202e62db26a0)


## Search Queries Used

- DNAH6 mutation biallelic autosomal recessive autosomal dominant de novo Mendelian case report family segregation primary ciliary dyskinesia PCD heterotaxy congenital heart disease

- DNAH6 dynein axonemal heavy chain 6 cilia flagella motility sperm motility male infertility MMAF functional study mouse knockout zebrafish knockdown electron microscopy axoneme

- DNAH6 exome sequencing genome sequencing rare variants loss of function predicted pathogenic ClinVar OMIM trio undiagnosed disease program Mendelian gene discovery burden analysis

- DNAH6 no association population frequency gnomAD allele frequency loss of function intolerance gene constraint pLI DNAH family DNAH5 DNAH11 genetic heterogeneity phenocopy


## Bibliometric Configuration

- Use Bibliometrics: True

- Author Impact Weight: 0.4

- Citation Impact Weight: 0.4

- Venue Impact Weight: 0.2
