## Supplementary material for "MENDELSEEK: An algorithm that predicts Mendelian Genes and elucidates what makes them special": DUS2.txt

# Claim: The gene DUS2 is likely associated with at least one Mendelian disease.

**Status**: processed

**Overall Rating**: -1

**Explanation**:

No relevant papers were found that support or refute this claim.


## Relevant Papers


## Other Reviewed Papers


### Identi ﬁ cation and functional characterization of de novo FOXP1 variants provides novel insights into the etiology of neurodevelopmental disorder

**Why Not Relevant**: The provided paper content exclusively discusses FOXP1: clinical cases, functional characterization of FOXP1 variants, and proposed mechanisms (loss of transcriptional repression and possible dominant-negative effects). There is no mention of the gene DUS2, nor any data, association tests, or mechanistic discussion relating to DUS2. Therefore this paper content provides no direct genetic or mechanistic evidence that DUS2 is associated with any Mendelian disease. Caveats: the analysis is limited to the text supplied here — if the full paper (outside the provided content) includes data on DUS2, that is not available for this review. Based on the provided excerpts alone, relevance to the claim about DUS2 is effectively zero.


[Read Paper](https://www.semanticscholar.org/paper/69811e3522216a7c9f789332730ac1ba8c2641db)


### De novo missense variants in LMBRD2 are associated with developmental and motor delays, brain structure abnormalities and dysmorphic features

**Why Not Relevant**: The provided paper describes LMBRD2 and de novo LMBRD2 variants; it does not mention the gene DUS2 anywhere in the supplied text and therefore provides no direct or mechanistic evidence regarding DUS2’s association with a Mendelian disease. For example, the paper states: "Objective To determine the potential disease association between variants in LMBRD2 and complex multisystem neurological and developmental delay phenotypes." and "Conclusion These findings indicate that rare de novo variants in LMBRD2 can lead to a previously unrecognised early-onset neurodevelopmental disorder." Both sentences explicitly focus on LMBRD2, not DUS2. The paper also notes that "LMBRD2 encodes an evolutionary ancient and widely expressed transmembrane protein with no known disease association, although two paralogues are involved in developmental and metabolic disorders," which again pertains to LMBRD2 and its paralogues rather than DUS2. Because DUS2 is not discussed, there is neither direct evidence (cases or variants in DUS2 linked to disease) nor mechanistic evidence (functional, biochemical, or model-organism data implicating DUS2) in this document. Limitations: absence of mention means nothing about DUS2’s true disease status—this paper cannot be used to support or refute the claim about DUS2. To evaluate the claim about DUS2, one should instead search gene-specific resources (OMIM, ClinVar, HGMD), literature for DUS2 human variants and phenotypes, functional studies of DUS2, and population constraint metrics (gnomAD), or perform targeted variant/segregation analyses.


[Read Paper](https://www.semanticscholar.org/paper/d35f5b055525b8e5fab475ae393f5abebfda34da)


### Genome Sequencing of Idiopathic Speech Delay

**Why Not Relevant**: This paper does not provide direct or mechanistic evidence relevant to the claim that DUS2 is likely associated with at least one Mendelian disease. The text mentions specific genes (SETD1A, SPTBN1, ARF3) found in a cohort of children with idiopathic speech delay and discusses pathogenic variants and candidate genes for speech-related disorders, but DUS2 is not mentioned anywhere in the provided content. Therefore there are no verbatim sentences or paragraphs in this paper that support or refute an association between DUS2 and a Mendelian disease. 

Detailed reasoning following the requested analysis guidelines:
- Direct evidence: None. The paper reports pathogenic variants in SETD1A and likely pathogenic variants in SPTBN1 and ARF3, but contains no data, case reports, variant calls, segregation, or clinical associations involving DUS2.
- Mechanistic evidence: None. There are no mechanistic descriptions (biochemical function, pathways, model systems, phenocopy experiments) related to DUS2 in the provided content.
- Methodology/results relevance: The study used genome sequencing in 23 probands (including parent trios for 10), a small cohort for discovery of high-penetrance variants in speech delay, and identified genes other than DUS2. Although the study design could in principle detect Mendelian associations for any gene (including DUS2) if variants were present, the absence of any mention or reported DUS2 variants means no information about DUS2 can be inferred from these results.
- Limitations and impact: The absence of DUS2 in the reported findings could be due to its not being mutated in this small sample, to filtering/prioritization choices, or simply because the gene was not implicated. Absence of evidence here is not evidence that DUS2 is not associated with a Mendelian disease, but it does mean this paper provides no affirmative support for the claim.

If you want evidence for or against the claim about DUS2, relevant materials would include: reports of rare pathogenic (e.g., loss-of-function) variants in DUS2 segregating with a Mendelian phenotype in families or recurrent de novo variants in affected individuals; functional/mechanistic studies showing how DUS2 loss or mutation produces a disease-relevant phenotype in cells or model organisms; or curated entries in disease databases (e.g., OMIM, ClinVar) linking DUS2 to a named Mendelian disorder. This paper provides none of those elements for DUS2.


[Read Paper](https://www.semanticscholar.org/paper/bc658050f962ece1b4738e7ad56cd11fc5364bb6)


## Search Queries Used

- DUS2 variant patient case report Mendelian disease DUS2 mutation dihydrouridine synthase 2 tRNA dihydrouridine synthase 2 monogenic autosomal recessive autosomal dominant familial

- DUS2 function tRNA modification dihydrouridine synthase 2 enzyme activity protein structure knockout mouse zebrafish drosophila yeast model translation mitochondrial translation neurodevelopment neurodegeneration

- DUS2 de novo rare variant exome sequencing cohort intellectual disability developmental delay neurodevelopmental disorders rare variant burden segregation loss of function variant

- ClinVar DUS2 OMIM DUS2 ClinGen DUS2 gnomAD pLI loss of function intolerance gene disease association clinical genetic testing panel DUS2


## Bibliometric Configuration

- Use Bibliometrics: True

- Author Impact Weight: 0.4

- Citation Impact Weight: 0.4

- Venue Impact Weight: 0.2
