## Supplementary material for "MENDELSEEK: An algorithm that predicts Mendelian Genes and elucidates what makes them special": ECSIT.txt

# Claim: The gene ECSIT is likely associated with at least one Mendelian disease.

**Status**: processed

**Overall Rating**: 3

**Explanation**:

Supporting evidence:
The provided excerpt shows that ECSIT (Evolutionarily Conserved Signaling Intermediate in Toll pathway) is functionally connected to ACAD9, a known complex I assembly factor: tissues were used to examine ECSIT and its interaction with ACAD9, and ECSIT levels were significantly reduced in the absence of ACAD9. The same excerpt emphasizes that ACAD9 has a dual role including complex I assembly and that ACAD9 deficiency is a recognized cause of isolated complex I deficiency in humans. Together, these points establish that ECSIT participates in the biology of mitochondrial complex I assembly and that perturbation of this assembly pathway (via ACAD9) leads to Mendelian disease.

Caveats or contradictory evidence:
The excerpt does not present any direct human genetic evidence linking ECSIT variants to a Mendelian phenotype: there are no reported pathogenic ECSIT variants, no patient genotype–phenotype data, no segregation analyses, and no functional rescue experiments implicating ECSIT mutations in human disease. The observation that ECSIT levels fall when ACAD9 is absent could reflect ECSIT being destabilized secondary to loss of an interacting partner rather than indicating that primary defects in ECSIT cause disease. The single provided paper and its excerpts are oriented toward ACAD9 function; ECSIT is mentioned in mechanistic context but not as an established disease gene. The provided metadata also indicates low bibliometric impact and low relevance score, suggesting sparse direct evidence in this dataset.

Analysis of potential mechanisms and implications:
ECSIT is described as a complex I assembly-related protein interacting with ACAD9. Many proteins involved in mitochondrial complex I assembly are known Mendelian disease genes: pathogenic variants in assembly factors frequently produce autosomal recessive mitochondrial encephalomyopathies or isolated complex I deficiency. Mechanistically, loss-of-function mutations in an essential assembly factor can impair complex I biogenesis and lead to multisystem mitochondrial disease, so disruption of ECSIT could plausibly produce a Mendelian mitochondrial disorder. However, functional involvement and plausibility do not substitute for genetic evidence: for a gene to be considered 'likely associated' with a Mendelian disease, one would expect at least case reports of pathogenic variants with supportive segregation and functional data. The current evidence establishes biological plausibility but not documented causal association.

Assessment and recommended rating:
Given the mechanistic link between ECSIT and a pathway in which other genes are established Mendelian disease genes, there is reasonable biological plausibility that ECSIT could be associated with a Mendelian disease. However, the provided evidence contains no direct human genetic data showing ECSIT variants causing disease. The balance of evidence is therefore not strongly supportive nor directly contradictory; it is suggestive but unproven. Accordingly, the most appropriate assessment based on the provided materials is 'Mixed Evidence'—plausible and biologically supported but lacking direct genetic/clinical confirmation in the supplied excerpts.


**Final Reasoning**:

I reviewed the provided excerpt and evaluated whether it supports the claim that ECSIT is likely associated with at least one Mendelian disease. The excerpt documents ECSIT's role in complex I assembly and its interaction with ACAD9, and notes that ACAD9 deficiency causes a Mendelian complex I deficiency. This establishes good biological plausibility: many complex I assembly factors are Mendelian disease genes, so ECSIT is a reasonable candidate. However, no direct human genetic evidence (pathogenic variants, affected patients, segregation, or functional rescue linking ECSIT mutation to disease) is provided. The observed reduction of ECSIT in ACAD9 deficiency could be a downstream effect rather than evidence that primary ECSIT defects cause disease. Because of this gap between plausible mechanism and lack of genetic/clinical confirmation in the supplied evidence, the correct classification is 'Mixed Evidence'—neither strongly supported nor contradicted by the materials provided.


## Relevant Papers


### Understanding ACAD9 function and the physiologic consequences of its deficiency

**Authors**: Andrew Sinsheimer (H-index: 1)

**Relevance**: 0.25

**Bibliometric Impact**: 0.008


**Excerpts**:

- Tissues from these animals were also used to examine a second complex I assembly factor, Evolutionarily Conserved Signaling Intermediate in Toll pathway (ECSIT), and its interaction with ACAD9.

- ECSIT levels were significantly reduced in the absence of ACAD9.

- In addition to its function in mitochondrial FAO, ACAD9 has been shown to have a second function as one of 14 factors responsible for assembly of complex I of the electron transport chain (ETC).

- ACAD9 deficiency leads to a combined defect in fatty acid oxidation and oxidative phosphorylation (OXPHOS) due to a duel role in the pathways.

- ACAD9 is the most common cause of isolated complex I deficiency in humans, underscoring the public health significance of these studies have relative to diagnosis and treatment.


**Explanations**:

- Mechanistic relevance: This sentence states that the authors examined ECSIT as a complex I assembly factor and its interaction with ACAD9. It provides mechanistic context linking ECSIT to a known Mendelian-disease-associated protein (ACAD9) involved in complex I assembly. Strength: directly ties ECSIT to the same biochemical pathway and experimental focus as a gene already disease-associated. Limitation: it does not provide genetic or clinical evidence that ECSIT itself causes Mendelian disease; it only establishes that ECSIT was studied in the ACAD9 models.

- Mechanistic relevance: This sentence reports that ECSIT protein levels fall when ACAD9 is absent, indicating a functional/biochemical dependence between the two proteins. Strength: supports biological plausibility that ECSIT participates in the same molecular machinery as ACAD9 (complex I assembly) and that perturbation of one affects the other. Limitation: decreased ECSIT levels in an ACAD9 knockout is indirect evidence — it could reflect secondary effects of ACAD9 loss rather than demonstrating that ECSIT mutations cause Mendelian disease. No human genetic or phenotype data for ECSIT are provided.

- Mechanistic/contextual relevance: This sentence establishes ACAD9 as one of the assembly factors for complex I. Strength: places ECSIT (noted elsewhere as a complex I assembly factor) in a disease-relevant pathway because defects in complex I assembly are known to cause Mendelian mitochondrial disorders. Limitation: this is background about ACAD9 and complex I assembly; it does not provide direct evidence linking ECSIT variation to human Mendelian disease.

- Contextual relevance: This describes the clinical consequence of ACAD9 deficiency (combined FAO and OXPHOS defect), underscoring that perturbation of proteins in this pathway can produce Mendelian phenotypes. Strength: supports the plausibility that other proteins in the same pathway (including ECSIT) could, in principle, be disease-causing. Limitation: still indirect — it documents ACAD9 pathogenicity but provides no direct ECSIT genetic/clinical data.

- Contextual relevance: This emphasizes that ACAD9 is a common genetic cause of isolated complex I deficiency in humans, reinforcing that components of the complex I assembly machinery are established Mendelian disease genes. Strength: bolsters mechanistic plausibility that ECSIT, as a complex I assembly factor interacting with ACAD9, could be implicated in Mendelian disease. Limitation: does not constitute direct evidence that ECSIT mutations cause disease; no variant, segregation, phenotype, or patient data are presented for ECSIT.


[Read Paper](https://www.semanticscholar.org/paper/fa6ca8afdae3473017629e03cb3b652bc8190a98)


## Other Reviewed Papers


### Novel De Novo KCND3 Mutation in a Japanese Patient with Intellectual Disability, Cerebellar Ataxia, Myoclonus, and Dystonia

**Why Not Relevant**: This paper content is not relevant to the claim that ECSIT is likely associated with at least one Mendelian disease. The supplied text describes a single case report of a 30-year-old Japanese man with intellectual disability, early-onset cerebellar ataxia, myoclonus, and dystonia, and states that a de novo novel missense mutation was revealed in SCA19/22. The content makes no mention of the gene ECSIT, provides no variants in ECSIT, no segregation data, no functional studies of ECSIT, and no mechanistic discussion relating ECSIT to disease. Therefore there is no direct evidence (no reported ECSIT variant linked to disease) and no mechanistic evidence (no molecular or pathway data implicating ECSIT) in this paper content to support or refute the claim.

Additional notes on why this is insufficient as evidence for the claim:
- A Mendelian disease association for a gene typically requires specific evidence such as pathogenic variants in that gene observed in affected individuals with appropriate segregation, recurrent observations across unrelated families, and/or functional demonstration that variants disrupt gene function in a disease-relevant manner. None of these elements are present for ECSIT in the provided text. 
- The paper instead implicates SCA19/22 (a spinocerebellar ataxia locus) and reports a novel missense mutation in that context; it therefore addresses different gene(s)/locus(es) and phenotype(s). 
- Because the provided content does not mention ECSIT at all, it cannot be used as supporting or opposing evidence regarding ECSIT’s involvement in Mendelian disease. 

If you can provide the full text or excerpts that explicitly mention ECSIT (variants, functional data, segregation, or mechanistic links), I can re-evaluate relevance and extract verbatim supporting or refuting evidence.


[Read Paper](https://www.semanticscholar.org/paper/b552f8e8d70e2896f399ad3236da2f8bf60efbe5)


### De novo mutations in autosomal recessive congenital malformations

**Why Not Relevant**: The provided paper content does not mention ECSIT, any variants in that gene, or any phenotypes linked to that gene. It only describes, in a single sentence, the use of trio-based exome sequencing in nine families and the detection of two cases where a recessive disorder was caused by a de novo plus an inherited variant. Because there is no specific information about ECSIT (no variant calls, no segregation or functional data, no mechanistic discussion involving ECSIT), the content cannot be used to support or refute the claim that ECSIT is associated with a Mendelian disease. To evaluate the claim, one would need explicit reporting of ECSIT variants segregating with disease and/or functional/mechanistic evidence implicating ECSIT in disease pathogenesis; none of those elements are present in the excerpt.


[Read Paper](https://www.semanticscholar.org/paper/a2ba92abb5d06fff00c2b0d81291ff611c496f38)


### A novel de novo nonsense mutation in SALL4 causing duane radial ray syndrome: a case report and expanding the phenotypic spectrum

**Why Not Relevant**: The single provided sentence refers to SALL4 and SALL4-related disorders and does not mention ECSIT. SALL4 and ECSIT are distinct genes with different known biology; therefore a statement that "The findings could extend the pathogenic mutations and phenotypic spectrum of SALL4 related disorders" provides no direct evidence that ECSIT is associated with any Mendelian disease, nor does it provide mechanistic evidence linking ECSIT to disease. In addition, only one short sentence was supplied from the paper, so there is no broader context, methods, results, or mechanistic discussion to evaluate. Because (1) ECSIT is not mentioned, (2) no mechanistic link between SALL4 and ECSIT is provided, and (3) insufficient text is available to search for any indirect evidence elsewhere in the paper, the content provided is not relevant to the claim that "ECSIT is likely associated with at least one Mendelian disease." To assess the claim we would need sentences or sections that explicitly mention ECSIT, describe pathogenic variants in ECSIT, association studies, segregation in families, functional assays, or mechanistic pathways implicating ECSIT in Mendelian phenotypes.


[Read Paper](https://www.semanticscholar.org/paper/bfc3fc258851307680f6d2e56bbc7a78bc1d3ab2)


### Missense Mutations in MAB21L1: Causation of Novel Autosomal Dominant Ocular BAMD Syndrome

**Why Not Relevant**: The provided paper content concerns the gene MAB21L1 and an autosomal dominant ocular syndrome (BAMD) caused by heterozygous MAB21L1 variants; it does not mention ECSIT or provide any data on that gene. Verbatim illustrative sentences from the paper (showing focus on MAB21L1) include:

"Purpose Biallelic MAB21L1 variants have been reported to cause autosomal recessive cerebellar, ocular, craniofacial, and genital syndrome (COFG), whereas only five heterozygous pathogenic variants have been suspected to cause autosomal dominant (AD) microphthalmia and aniridia in eight families. This study aimed to report an AD ocular syndrome (blepharophimosis plus anterior segment and macular dysgenesis [BAMD]) syndrome based on clinical and genetic findings from patients with monoallelic MAB21L1 pathogenic variants in our cohort and reported cases."

"Results Three heterozygous missense variants in MAB21L1, predicted to be damaging, were detected in 5 unrelated families, including c.152G>T in 2, c.152G>A in 2, and c.155T>G in one. All were absent from gnomAD. The variants were de novo in two families, transmitted from affected parents to offspring in two families, and with an unknown origin in the other family, demonstrating strong evidence of AD inheritance. All patients revealed similar BAMD phenotypes, including blepharophimosis, anterior segment dysgenesis, and macular dysgenesis."

"Conclusions Heterozygous pathogenic variants in MAB21L1 account for a new AD BAMD syndrome, which is completely different from COFG caused by homozygous variants in MAB21L1. Nucleotide c.152 is likely a mutation hot spot, and the encoded residue of p.Arg51 might be critical for MAB21L1."

Because ECSIT is not mentioned and no data (genetic, clinical, or mechanistic) concerning ECSIT are presented, this paper provides no direct evidence linking ECSIT to any Mendelian disease, nor any mechanistic evidence that would plausibly connect ECSIT to the phenotypes described. Therefore the paper is not relevant to the claim that "The gene ECSIT is likely associated with at least one Mendelian disease."


[Read Paper](https://www.semanticscholar.org/paper/372e5086b85699136a1393cc555e99e2818549cc)


### SPG9A with the new occurrence of an ALDH18A1 mutation in a CMT1A family with PMP22 duplication: case report

**Why Not Relevant**: The provided text makes no mention of the gene ECSIT or any data relating to ECSIT. It refers instead to ALDH18A1, P5CS deficiency, and an SPG9A family with a de novo mutation or possible gonadal mosaicism. There are no genetic, clinical, segregation, functional, or mechanistic data presented that link ECSIT to any Mendelian disease. The single sentence fragment supplied only indicates that further studies are required to clarify relationships between amino acid levels and clinical manifestations for P5CS (encoded by ALDH18A1) deficiency. Because the gene names and biological context do not match the claim, and because no direct or indirect evidence regarding ECSIT is present, this paper content is not relevant to evaluating whether ECSIT is likely associated with at least one Mendelian disease. Additional relevant information (e.g., explicit mention of ECSIT variants, segregation in families, functional assays, or disease associations) would be needed to support or refute the claim.


[Read Paper](https://www.semanticscholar.org/paper/c52899f94d0ef2b838ffdba1e3484424d8c18855)


## Search Queries Used

- ECSIT mutation Mendelian disease case report clinical exome biallelic de novo autosomal recessive autosomal dominant family segregation patient phenotype

- ECSIT mitochondrial complex I assembly oxidative phosphorylation mitochondrial disease cardiomyopathy encephalopathy functional study knockout mouse zebrafish patient fibroblast

- ECSIT NF kappa B Toll like receptor signaling innate immunity immunodeficiency autoinflammatory cytokine defect patient functional assay variant

- ECSIT loss of function intolerance gnomAD pLI constraint haploinsufficiency ClinVar gene disease association review Mendelian gene curation


## Usage Statistics

- Prompt Tokens: 0

- Completion Tokens: 0

- Total Tokens: 0

- Estimated Cost: $0.0216


## Bibliometric Configuration

- Use Bibliometrics: True

- Author Impact Weight: 0.4

- Citation Impact Weight: 0.4

- Venue Impact Weight: 0.2
