## Supplementary material for "MENDELSEEK: An algorithm that predicts Mendelian Genes and elucidates what makes them special": EHMT2.txt

# Claim: The gene EHMT2 is likely associated with at least one Mendelian disease.

**Status**: processed

**Overall Rating**: 0

**Explanation**:

Supporting evidence:
The provided excerpts show that EHMT2 (also known as G9a) is a bona fide histone lysine methyltransferase that catalyzes H3K9 dimethylation and participates in epigenetic regulation. In particular, the RNAi study demonstrates that reducing EHMT2 expression in mouse models and patient iPSC-derived cells leads to activation of maternally silenced Prader–Willi syndrome (PWS) genes (SNRPN, SNORD116) — roughly a twofold increase in expression after ~80–90% EHMT2 knockdown. This establishes a clear mechanistic role for EHMT2 in controlling imprinting at the PWS imprinting center and shows that perturbing EHMT2 function alters expression of disease-relevant genes.

Caveats or contradictory evidence:
Neither paper presents evidence that germline variants or mutations in EHMT2 cause a Mendelian disease in humans. The RNAi data use experimental knockdown (siRNA/shRNA) to change gene expression in model systems and patient-derived cells; such manipulations demonstrate function but do not show that natural EHMT2 alleles are pathogenic. The muscle paper explicitly states EHMT2 is dispensable for skeletal muscle development and regeneration in vivo, implying that loss of EHMT2 function does not necessarily produce an overt developmental phenotype in that tissue. There is no direct clinical genetic data, segregation analysis, case reports, or association of EHMT2 variants with monogenic disease provided in these excerpts.

Analysis of potential underlying mechanisms and implications:
Mechanistically, EHMT2’s role in H3K9me2-mediated silencing at the PWS imprinting center makes it plausible that alterations in EHMT2 activity could influence imprinting and thereby modify expression of imprinted loci. In theory, this could have phenotypic consequences and might contribute to imprinting-related disorders if disrupted in the appropriate context (germline mutation, imprinting establishment errors, or dominant-negative effects). However, experimental knockdown resulting in reactivation of maternally silenced genes is conceptually the opposite of how EHMT2 defects would need to act to produce classic PWS (which arises from loss of paternal expression). Moreover, tissue-specific dispensability (muscle) and lack of presented human genetic data reduce the likelihood that simple loss-of-function EHMT2 alleles cause a recognizable Mendelian syndrome, at least in the contexts tested.

Which rating is most appropriate and why:
Given the evidence provided, the most defensible conclusion is that there is no direct evidence in these papers that EHMT2 is associated with a Mendelian disease. The excerpts demonstrate a regulatory role relevant to a Mendelian disorder (PWS) but do not provide human genetic association data or examples of pathogenic EHMT2 variants segregating with disease. Therefore the claim that "EHMT2 is likely associated with at least one Mendelian disease" is not supported by the supplied evidence. Because the evidence is functional and mechanistic rather than genetic/clinical, and because one paper even shows tissue dispensability, the correct assessment based on the provided material is that there is no evidence for a Mendelian disease association in these excerpts.


**Final Reasoning**:

I reviewed the provided excerpts and logic carefully. The RNAi paper provides strong functional evidence that EHMT2 mediates epigenetic silencing of PWS-related genes and that reducing EHMT2 activity can reactivate those genes. However, that functional perturbation in experimental systems does not equate to evidence that germline variants in EHMT2 cause a Mendelian disorder. The muscle paper’s finding that EHMT2 is dispensable for skeletal muscle development further argues against a straightforward loss-of-function Mendelian phenotype affecting that tissue. There are no case reports, segregation data, human genetics analyses, or statements in the excerpts that link EHMT2 mutations to monogenic disease. Therefore the claim as stated is not supported by the provided papers. I find no direct evidence in the supplied excerpts for a Mendelian disease association and assign the rating "No Evidence."


## Relevant Papers


### The lysine methyltransferase Ehmt2/G9a is dispensable for skeletal muscle development and regeneration

**Authors**: Regan Zhang (H-index: 9), F. Rossi (H-index: 50)

**Relevance**: 0.15

**Bibliometric Impact**: 0.3376


**Excerpts**:

- Ehmt2 and H3K9me2 do not play significant roles in skeletal muscle development and regeneration in vivo and are not implicated in a number of biological processes relevant to cell fate control.


**Explanations**:

- Relation to claim: This sentence directly reports negative experimental findings about Ehmt2 (the mouse ortholog of human EHMT2) and the histone modification H3K9me2, stating they "do not play significant roles" in skeletal muscle development, regeneration, and several cell-fate–related biological processes. This constitutes evidence against EHMT2 being involved in Mendelian diseases that manifest via disrupted skeletal muscle development or regeneration or via the specific cell-fate processes studied. Type of evidence: mechanistic/functional (negative functional evidence) rather than human genetic or epidemiologic evidence. Strengths: The statement is a clear experimental conclusion about in vivo function (presumably based on the paper's experiments), which weakens the plausibility that EHMT2 causes Mendelian disease through those particular mechanisms or in skeletal muscle. Limitations/caveats: (1) The sentence reports a negative result in a particular biological context (skeletal muscle development and regeneration) and for "a number of biological processes relevant to cell fate control," but does not address other tissues, developmental stages, or molecular functions of EHMT2 that could underlie Mendelian disease. (2) It does not provide human genetic or clinical evidence of Mendelian inheritance or pathogenic variants; absence of functional effect in these assays does not exclude disease-causing roles in other contexts. (3) No experimental details, statistical measures, sample sizes, species or model specifics, or boundaries of the "number of biological processes" are provided in this excerpt, limiting assessment of generalizability. Overall assessment: relevant but limited. The paper provides mechanistic evidence weakening the claim for muscle- or cell-fate–related Mendelian disease mechanisms but does not directly test or refute a general association between EHMT2 and any Mendelian disease in humans.


[Read Paper](https://www.semanticscholar.org/paper/01cc7fa1f1473bf294ff90a4a96f531c9b50f857)


### RNAi Knockdown of EHMT2 in Maternal Expression of Prader–Willi Syndrome Genes

**Authors**: Violeta Zaric (H-index: 0), Ryan K Butler (H-index: 1)

**Relevance**: 0.35

**Bibliometric Impact**: 0.124


**Excerpts**:

- Background/objectives: Euchromatic histone lysine methyltransferase 2 (EHMT2, also known as G9a) is a mammalian histone methyltransferase that catalyzes the dimethylation of histone 3 lysine 9 (H3K9).

- On human chromosome 15, the parental-specific expression of Prader–Willi Syndrome (PWS)-related genes, such as SNRPN and SNORD116, are regulated through the genetic imprinting of the PWS imprinting center (PWS-IC).

- On the paternal allele, PWS genes are expressed whereas the epigenetic maternal silencing of PWS genes is controlled by the EHMT2-mediated methylation of H3K9 in PWS-IC.

- Here, we measured the effects of RNA interference of EHMT2 on the maternal expression of genes deficient in PWS in mouse model and patient iPSC-derived cells.

- We used small interfering RNA (siRNA) oligonucleotides and lentiviral short harpin RNA (shRNA) to reduce Ehtm2/EHMT2 expression in mouse Snord116 deletion primary neurons, PWS patient-derived induced pluripotent stem cell (iPSC) line and PWS iPSC-derived neurons.

- With an approximate reduction of 90% in EHMT2 mRNA and more than 80% of the EHMT2 protein, we demonstrated close to a 2-fold increase in the expression of maternal transcripts for SNRPN and SNORD116 in PWS iPSCs treated with siEHMT2 compared to PWS iPSC siControl.

- A similar increase in SNORD116 and SNRPN RNA expression was observed in PWS iPSC-derived neurons treated with shEHMT2.

- RNAi reduction in EHMT2 activates maternally silenced PWS genes.

- Further studies are needed to determine whether the increase is therapeutically relevant.

- This study confirms the role of EHMT2 in the epigenetic regulation of PWS genes.


**Explanations**:

- Mechanistic relevance: This sentence defines EHMT2's biochemical activity (H3K9 dimethylation). It provides mechanistic plausibility that EHMT2 can alter chromatin states and gene expression. Strength: clear molecular function relevant to epigenetic regulation. Limitation: does not by itself link EHMT2 to any human Mendelian disease.

- Contextual relevance: Establishes that SNRPN and SNORD116 (genes involved in Prader–Willi Syndrome) are regulated by imprinting at the PWS-IC on chromosome 15. Strength: links the biological system (PWS genes) that could be affected by EHMT2 activity. Limitation: descriptive of PWS locus, not of EHMT2 mutations causing disease.

- Direct mechanistic linkage: States that maternal silencing at the PWS-IC is controlled by EHMT2-mediated H3K9 methylation. This is the central mechanistic connection between EHMT2 activity and regulation of genes implicated in a Mendelian imprinting disorder (PWS). Strength: provides a specific epigenetic mechanism whereby EHMT2 affects disease-relevant loci. Limitation: mechanism shown as regulatory control, but does not demonstrate that genetic variants in EHMT2 cause PWS or another Mendelian disease in humans.

- Study aim and approach: Explains the experiment tested (RNAi of EHMT2) to assess effects on maternally silenced PWS genes in model systems. Relevance: shows the authors directly tested whether altering EHMT2 changes expression of Mendelian disease-related genes. Limitation: this is an experimental manipulation (knockdown) rather than observation of pathogenic human variants.

- Experimental methods: Describes the use of siRNA/shRNA in mouse primary neurons and patient-derived iPSCs/neurons to reduce EHMT2. Relevance: indicates the data derive from cellular knockdown models relevant to PWS. Limitation: knockdown magnitude and method may not reflect heterozygous germline loss-of-function or missense variants in humans; species and in vitro contexts affect generalizability.

- Result — effect size: Reports that ~90% mRNA and >80% protein reduction of EHMT2 produced ~2-fold increases in maternal SNRPN and SNORD116 transcripts in PWS iPSCs. Relevance: provides experimental evidence that reducing EHMT2 derepresses PWS-related genes, supporting a mechanistic role of EHMT2 in imprinting at a locus involved in a Mendelian disorder. Strength: quantitative effect reported. Limitations: no statistical values reported here (p-values, confidence intervals) in the excerpt; the perturbation is large (near-complete knockdown), which may not mirror typical human genetic variation; increased expression in PWS iPSCs (which already lack paternal expression) does not prove that EHMT2 variants cause PWS phenotypes in vivo.

- Replication in neuronal differentiation: States a similar increase was observed in iPSC-derived neurons treated with shEHMT2. Relevance: demonstrates the effect is reproducible in a differentiated cell type relevant to disease. Limitation: still an in vitro knockdown model; no evidence linking EHMT2 variants to Mendelian disease in patients.

- Conclusion about mechanism: Concise statement that RNAi reduction activates maternally silenced PWS genes. Relevance: directly supports mechanistic plausibility that loss of EHMT2 function could alter imprinting at disease-associated loci. Limitation: activation after experimental knockdown is not direct evidence that germline EHMT2 mutations cause a Mendelian disease.

- Caveat about clinical relevance: Authors explicitly state therapeutic relevance is undetermined. Relevance: indicates the study authors do not claim that EHMT2 perturbation equates to a Mendelian disease cause and acknowledge further work is needed. This weakens the strength of claiming EHMT2 is currently established as causative for any Mendelian disease based on these data.

- Overall interpretive statement: The study confirms EHMT2's role in epigenetic regulation of PWS genes. Relevance: supports mechanistic plausibility linking EHMT2 to pathways involved in a Mendelian imprinting disorder (PWS). Limitation: confirmation of regulatory role is not the same as genetic association of EHMT2 variants with a Mendelian disease; no human genetic evidence of causative EHMT2 variants is presented.


[Read Paper](https://www.semanticscholar.org/paper/5dfeb6c45f10fe20202435faa61c36668aa955ba)


## Other Reviewed Papers


### Non-invasive prenatal sequencing for multiple Mendelian monogenic disorders using circulating cell-free fetal DNA

**Why Not Relevant**: The paper content provided is a single summary sentence and does not mention EHMT2 by name. Therefore it provides no direct evidence that EHMT2 is associated with a Mendelian disease. The sentence only indicates that a 30-gene panel targeting genes "frequently associated with dominant monogenic diseases" was used; this is insufficient because (a) the specific gene list is not provided so we cannot determine whether EHMT2 is included, (b) even inclusion on a test panel would be only an indirect signal and would require citation or justification, and (c) there are no variant-level data, clinical correlations, segregation analyses, functional studies, or literature references presented here that would constitute mechanistic or causal evidence for EHMT2 involvement in a Mendelian disorder. To support the claim, the paper would need to explicitly name EHMT2 as one of the genes, report pathogenic or likely pathogenic EHMT2 variants in affected individuals with phenotypic correlation (or cite prior gene–disease associations), or present mechanistic/functional data linking EHMT2 dysfunction to disease. None of these are present in the provided content, so the relevance to the claim is minimal.


[Read Paper](https://www.semanticscholar.org/paper/965b5d12657d5a39be70c31cf4cb3eb1617f5e96)


### Targeting the histone methyltransferase G9a activates imprinted genes and improves survival of a mouse model of Prader–Willi syndrome

**Why Not Relevant**: The provided paper content contains only the single summary sentence: "This study provides the first proof of principle for an epigenetics-based therapy for PWS by displaying therapeutic effects in the PWS mouse model by improving the survival and the growth of m+/pΔS−U newborn pups." This sentence: (1) does not mention EHMT2 (also known as G9a) or any other specific gene; (2) does not present genetic-variant, segregation, case-report, cohort, or human clinical data that would bear on a Mendelian-disease association; and (3) gives no mechanistic details linking the described epigenetic therapy to EHMT2 activity or to causal genetic variation. While the study concerns epigenetic therapy in a mouse model of Prader–Willi syndrome (a genetic/imprinting disorder), that context alone is insufficient to support or refute the claim that "EHMT2 is likely associated with at least one Mendelian disease." There is no direct evidence (no mention of mutations, patient phenotypes, linkage/segregation, or pathogenic variants in EHMT2) and no mechanistic evidence (no description that the intervention targeted EHMT2 or that EHMT2 perturbation produced a Mendelian phenotype). Limitations preventing relevance: single-sentence content with no gene-level data, no experimental details, no species-specific gene attribution to EHMT2, and no discussion of Mendelian inheritance or human disease association. If you want a determination about EHMT2 specifically, I can (a) search the literature for human genetic studies implicating EHMT2 in Mendelian disorders, or (b) analyze full-text papers that mention EHMT2 and experimental evidence; please provide additional papers or allow me to search external databases.


[Read Paper](https://www.semanticscholar.org/paper/dc606f07e342146f96dc9af42533ea1b464ac04e)


### Implications of a De Novo Variant in the SOX12 Gene in a Patient with Generalized Epilepsy, Intellectual Disability, and Childhood Emotional Behavioral Disorders

**Why Not Relevant**: This paper excerpt does not mention EHMT2 at all and exclusively discusses SRY-box (SOX) family genes (SOX12, SOX4, SOX11) and a de novo variant found in SOX12. Example statements from the excerpt showing scope include: "SRY-box transcription factor (SOX) genes, a recently discovered gene family, play crucial roles in the regulation of neuronal stem cell proliferation and glial differentiation during nervous system development and neurogenesis." and "Whole exome sequencing (WES) in patients presenting with generalized epilepsy, intellectual disability, and childhood emotional behavioral disorder, uncovered a de novo variation within SOX12 gene." Because EHMT2 is not mentioned, the paper provides neither direct evidence (e.g., observed pathogenic variants in EHMT2 segregating with Mendelian disease) nor mechanistic evidence (e.g., functional data linking EHMT2 perturbation to disease mechanisms) relevant to the claim that "EHMT2 is likely associated with at least one Mendelian disease." The absence of EHMT2 discussion in this paper means it cannot be used to support or refute the claim. Note: lack of relevance here does not imply anything about the true association status of EHMT2 in other literature; it only reflects that this paper contains no information bearing on EHMT2.


[Read Paper](https://www.semanticscholar.org/paper/5cf652470efa4406f72843aea8389f4e398a99f8)


### A de novo frameshift variant in MED13 gene in a patient with autism spectrum disorder and magnetic resonance imaging abnormalities mimicking tuberous sclerosis

**Why Not Relevant**: The paper content provided exclusively concerns the MED13 gene and a de novo truncating variant found in a single patient; it describes clinical phenotypes and concludes that MED13-related disorders include brain abnormalities. There is no mention of EHMT2 anywhere in the provided text, no experimental or clinical data related to EHMT2, and no mechanistic discussion connecting MED13 to EHMT2 or shared pathways that would allow inference about EHMT2. Because the paper does not address EHMT2 directly or indirectly, it provides no evidence for or against the claim that EHMT2 is likely associated with at least one Mendelian disease. Additionally, the evidence presented is from a single case report about MED13, which would be limited in generalizability even for MED13 itself.


[Read Paper](https://www.semanticscholar.org/paper/c1a65b9ac26a735bddb776612339efbbc90ae89a)


### A de novo PAK1 likely pathogenic variant and a de novo terminal 1q microdeletion in a Chinese girl with global developmental delay, severe intellectual disability, and seizures

**Why Not Relevant**: The supplied paper content discusses identification of a novel PAK1 variant associated with a terminal 1q microdeletion in a patient with a neurodevelopmental disorder and proposes that the main clinical features may be ascribed to a pathogenic variant in PAK1. There is no mention of the gene EHMT2, no data concerning EHMT2 variants, and no mechanistic discussion implicating EHMT2 in Mendelian disease. Therefore this paper provides no direct evidence (no variant-phenotype associations, no segregation, no functional studies) and no mechanistic evidence (no biochemical, expression, or pathway data) relevant to the claim that EHMT2 is likely associated with at least one Mendelian disease.

Limitations and caveats: this assessment is based solely on the provided paper content. If the full text contains additional sections (supplementary tables, gene lists, or broader discussion) that were not included in the excerpt, those might mention EHMT2; however, given the paper’s stated focus on PAK1 and a terminal 1q microdeletion, it is unlikely to contain substantive evidence about EHMT2. For authoritative evaluation of the claim about EHMT2, consult dedicated genetic-disease resources (OMIM, ClinVar, HGMD) or publications explicitly studying EHMT2 variants and phenotypes.


[Read Paper](https://www.semanticscholar.org/paper/8ac30d308f85987b8cf4644520ee11940136ebaa)


### Genetic basis of Parkinson's disease and related disorders

**Why Not Relevant**: The manuscript focuses on genetic studies of Parkinson disease (familial and sporadic), reporting specific gene discoveries and association results (LRRK2, PRKRA, PARK2, SNCA, MAPT, etc.). EHMT2 is not mentioned anywhere in the provided text. Therefore the paper provides no direct evidence supporting that EHMT2 is associated with any Mendelian disease. The paper does demonstrate methods (haplotype mapping, autozygosity mapping, GWAS, CNV analysis) capable of detecting Mendelian disease genes in PD cohorts; the absence of EHMT2 in reported findings suggests it was not implicated in these PD-focused analyses. Important caveats: absence of EHMT2 in this PD-centered study is not definitive evidence that EHMT2 is not associated with any Mendelian disease — EHMT2 could be causal for Mendelian phenotypes outside the study’s scope, involve variant types not well captured by the study (very rare variants, non-coding regulatory changes, structural variants outside probe resolution), or occur in populations not sampled here. Given no mention of EHMT2, relevance to the claim is very low.


[Read Paper](https://www.semanticscholar.org/paper/b59ed2e51a73f5ce5958c18560d750b40e4034b6)


### NGS based studies on primary immunodeficiencies (PIDs) : causative gene identification, tool development and application

**Why Not Relevant**: This paper does not provide direct or specific evidence that EHMT2 is associated with any Mendelian disease. The study reports genes found to be causal or candidate genes in primary immunodeficiency cohorts (examples: LRBA, ATM, DOCK8, PMS2, TNFRSF18, PIK3CG, LILRB1, EPHB2, TXNIP, CD5, NLRP5, DNMT3B, ZBTB24) but EHMT2 is not mentioned anywhere in the supplied content. While the paper discusses methylation and epigenetic/dna-repair machinery as mechanistic contributors to disease, the methylation genes explicitly implicated are DNMT3B and ZBTB24 (and ATM for DNA repair). EHMT2 (a histone H3K9 methyltransferase) is neither listed among discovered causal genes nor discussed mechanistically in this excerpt. The study used WES (which could in principle detect EHMT2 variants) and TRS panels (219 PID genes), but the authors do not report any EHMT2 findings; moreover it is not specified whether EHMT2 was included in the TRS panel. Therefore, absence of EHMT2 in the reported results constitutes absence of evidence in this paper, not evidence of absence of any Mendelian association for EHMT2 in the wider literature. Given no direct mention, variant data, functional assays, or statistical association concerning EHMT2 are presented here, the content is not relevant for supporting the claim that "EHMT2 is likely associated with at least one Mendelian disease."


[Read Paper](https://www.semanticscholar.org/paper/7786e43fbaf68904111f286747f0cf8968c2dd1e)


### Selected Abstracts from the VIIth Biennial Conference of African Society for Immunodeficiencies (ASID): ASID VII Online, Khartoum, October 1-2, 2021

**Why Not Relevant**: The provided paper content (a short abstract for an oral presentation) contains no mention of the gene EHMT2, no gene names at all, no genetic data, no variant/segregation evidence, and no mechanistic discussion linking any specific gene to a Mendelian disease. The entire text is:

"This oral presentation will demonstrate a systematic approach on when to suspect and how to diagnose PID in infants and children laying stress on symptoms and signs suggestive of these disorders, as well as the basic and advanced laboratory tests that are essential for proper diagnosis."

Why this is not relevant to the claim that "EHMT2 is likely associated with at least one Mendelian disease":
- No direct evidence: The text contains no reports of EHMT2 variants, phenotypes, case reports, cohort studies, statistical associations, or clinical genetic diagnoses involving EHMT2. Therefore it provides no direct support or refutation for an EHMT2–Mendelian disease association.
- No mechanistic evidence: There is no molecular, cellular, or pathway-level discussion that implicates EHMT2 (e.g., changes in histone methylation, gene regulation by EHMT2, or functional consequences of EHMT2 disruption) that could make an EHMT2–disease link plausible.
- Context does not substitute for gene-specific evidence: Although primary immunodeficiencies (PIDs) can be Mendelian, the general topic of PID diagnosis does not permit inference about any particular gene. Inferring EHMT2 involvement solely because PIDs are sometimes Mendelian would be speculative and unsupported by the presented content.
- Insufficient methodological detail: The abstract is a brief description of an educational presentation, not an original study; it lacks methods, data, results, or references that could be examined for relevance.

Given these points, the document provides no usable evidence (direct or mechanistic) to support or contradict the claim about EHMT2 and Mendelian disease. To evaluate the claim, one would need primary research articles, case reports, genetic databases (e.g., OMIM, ClinVar), or functional studies that specifically mention EHMT2 and associated phenotypes.


[Read Paper](https://www.semanticscholar.org/paper/31c6c1d39776c2e71b78f01a64f293ac49a307d7)


### Molecular and phenotypical findings of a novel de novo SYNGAP1 gene variant in an 11-year-old Iranian boy with intellectual disability.

**Why Not Relevant**: The paper content provided does not mention EHMT2 anywhere and instead describes a study of a SYNGAP1 variant causing intellectual developmental disorder type 5. Relevant verbatim sentences from the provided content that demonstrate the paper's focus on SYNGAP1 include:

"OBJECTIVE Intellectual developmental disorder (IDD) type 5 is an autosomal dominant (AD) disorder and is characterized by intellectual disability (ID), psychomotor developmental delay, variable autism phenotypes, microcephaly, and seizure. IDD can be caused by mutations in the SYNGAP1 gene, which encodes a Ras GTPase-activating protein. This study revealed a novel de novo nonsense variant in SYNGAP1. The identification of such variants is essential for genetic counseling in patients and their families."

"RESULTS The de novo NM_006772.3(SYNGAP1):c.3685C>T variant was identified in an 11-year-old boy with severe intellectual disability, neurodevelopmental delay, speech disorder, ataxia, specific dysmorphic facial features, and aggressive behavior."

"CONCLUSION The current study findings expand the existing knowledge of variants in SYNGAP1 that have been previously associated with nonsyndromic intellectual disability and autism, extending the spectrum of phenotypes associated with this gene. The data have implications for genetic diagnosis and counseling in similar phenotypic presentations."

How this bears on the claim (that EHMT2 is likely associated with at least one Mendelian disease):
- No direct evidence: The paper reports a de novo SYNGAP1 nonsense variant in a patient and interprets that variant with clinical genetic guidelines. There is no mention of EHMT2, no EHMT2 variants, and no patient phenotypes linked to EHMT2. Therefore the paper provides no direct evidence supporting or refuting an association between EHMT2 and any Mendelian disease.
- No mechanistic evidence relevant to EHMT2: The mechanistic statements concern SYNGAP1 (not EHMT2) — e.g., SYNGAP1 encodes a Ras GTPase-activating protein. There are no mechanistic data, pathways, functional assays, or literature discussion in the provided text that involve EHMT2 or its biological functions. Thus the paper gives no mechanistic basis to infer EHMT2 involvement in Mendelian disease.
- Limitations and inability to generalize: Because the study is specific to SYNGAP1 (and apparently a single-case report confirmed by exome and Sanger sequencing), it cannot be used to support claims about other genes. Even strong evidence that SYNGAP1 causes a Mendelian disorder does not imply that EHMT2 does or does not cause one, unless the paper had explicit data linking the two genes (which it does not).

Conclusion: Relevance is effectively zero. The provided paper is unrelated to EHMT2 and therefore cannot be used to support or refute the claim that EHMT2 is likely associated with at least one Mendelian disease.


[Read Paper](https://www.semanticscholar.org/paper/3f4b2eee9f7ad16af6b2e8b2a6417e1d72f20aed)


### The lysine methyltransferase Ehmt2/G9a is dispensable for skeletal muscle development and regeneration

**Why Not Relevant**: The paper's content addresses functional roles of Ehmt2/H3K9me2 in skeletal muscle development, regeneration, and certain cell-fate processes in vivo. The claim concerns whether the human gene EHMT2 is likely associated with at least one Mendelian disease (a human genetic association). The provided sentence is a negative functional result in a specific experimental/contextual setting (mouse, muscle, selected processes) and does not include human genetic, clinical, segregation, variant pathogenicity, or population evidence that would directly establish or refute a Mendelian disease association. Because the paper lacks human genetic data, disease phenotyping in humans, or broader evidence linking EHMT2 variants to monogenic disorders, it is not directly relevant to adjudicating the claim. The negative functional finding is limited in scope and cannot exclude other mechanisms (e.g., tissue-specific disease, dominant-negative or gain-of-function variants, haploinsufficiency in humans) by which EHMT2 could be associated with Mendelian disease.


[Read Paper](https://www.semanticscholar.org/paper/41d031b152bd3bc3bf76906aea6e3dad9b6f52bb)


### Abstract LB-156: Delineating the pathological roles and molecular functions of histone methyltransferase G9a in hepatocellular carcinoma

**Why Not Relevant**: Relevance is very low for the claim that EHMT2 is likely associated with at least one Mendelian disease because this abstract reports somatic tumor biology of EHMT2 (G9a) in hepatocellular carcinoma. The paper documents somatic deregulation (up-regulation), somatic gene copy number gain, miRNA-mediated post-transcriptional regulation, effects of knockdown/knockout in tumor cell lines, pharmacologic inhibitor responses, and xenograft tumorigenicity—none of which are evidence of germline pathogenic variants, segregation in families, congenital phenotypes, or population genetic association typical of Mendelian disease causation. Additionally, this is a conference abstract (limited methodological detail and no reported sample sizes or germline analyses). To support the claim that EHMT2 is associated with a Mendelian disease would require evidence such as: (a) identification of rare, segregating germline variants in affected families with consistent phenotype(s); (b) de novo pathogenic variants in patients with congruent clinical features; (c) experimental models showing that specific germline variants recapitulate the inherited phenotype; or (d) population-level enrichment of rare damaging variants in cases versus controls. None of those types of data are present here. Therefore the content is mechanistically informative about somatic oncogenic roles of EHMT2 but does not provide direct or sufficient mechanistic evidence to support a Mendelian disease association.


[Read Paper](https://www.semanticscholar.org/paper/e40e12a8be15f6080f8663bfc28a017190e02a4c)


## Search Queries Used

- EHMT2 G9a KMT1C mutation pathogenic variant Mendelian disorder monogenic disease case report family study

- EHMT2 de novo variant exome sequencing trio neurodevelopmental disorder intellectual disability autism developmental delay

- euchromatic histone lysine methyltransferase 2 EHMT2 G9a H3K9 methylation epigenetic regulation mechanism mouse model knockout phenotype

- EHMT2 ClinVar OMIM ClinGen gnomAD loss of function intolerance population frequency constraint haploinsufficiency no association


## Usage Statistics

- Prompt Tokens: 0

- Completion Tokens: 0

- Total Tokens: 0

- Estimated Cost: $0.0592


## Bibliometric Configuration

- Use Bibliometrics: True

- Author Impact Weight: 0.4

- Citation Impact Weight: 0.4

- Venue Impact Weight: 0.2
