## Supplementary material for "MENDELSEEK: An algorithm that predicts Mendelian Genes and elucidates what makes them special": ENDOG.txt

# Claim: The gene ENDOG is likely associated with at least one Mendelian disease.

**Status**: processed

**Overall Rating**: -1

**Explanation**:

No relevant papers were found that support or refute this claim.


## Relevant Papers


## Other Reviewed Papers


### Chromatin remodeling dysfunction extends the etiological spectrum of schizophrenia: a case report

**Why Not Relevant**: The provided paper content (a single sentence) does not mention the gene ENDOG or any specific genes, variants, experimental results, pedigrees, or statistical associations. It only states: "This case underlines that schizophrenia may be, sometimes, underpinned by a Mendelian disease and addresses the question of systematic genetic investigations in the presence of warning signs such as a childhood onset of the schizophrenia or a resistant epilepsy." This is a general clinical observation and recommendation about performing genetic investigations in certain clinical contexts; it does not provide direct evidence (no variant, no segregation, no case-control data) nor mechanistic evidence (no functional data, no pathway description) linking ENDOG to any Mendelian disease. Key missing items that would be required to support the claim include: identification of ENDOG variants in affected individuals, segregation of such variants with disease in pedigrees, population frequency and pathogenicity assessments, functional assays demonstrating a damaging effect on ENDOG function, or statistical enrichment of ENDOG variants in cases versus controls. Because none of these elements appear in the supplied text, the content cannot be used to support or refute the claim that ENDOG is likely associated with at least one Mendelian disease. Note also that the provided snippet is short and may not reflect the full paper; if you can provide the full text or additional excerpts that mention ENDOG specifically, I can re-evaluate for direct and mechanistic evidence. 


[Read Paper](https://www.semanticscholar.org/paper/29f6d9d6036caf286740be939452f1be2a4114b2)


### HOXA11 is another monogenic cause of congenital anomalies of the kidney and urinary tract—Reply

**Why Not Relevant**: The provided paper content mentions only HXA11 and its insufficient evidence as a monogenic cause of CAKUT. It contains no mention of the gene ENDOG, no experiments, observations, mechanisms, or statistical associations related to ENDOG, and no discussion of Mendelian diseases in connection to ENDOG. Therefore the content is not relevant to evaluating the claim that ENDOG is likely associated with at least one Mendelian disease. Because the single excerpt addresses a different gene and condition, no inferences about ENDOG can be drawn from it. To assess the claim about ENDOG, papers that explicitly discuss ENDOG (genetic variants, segregation in families, functional studies, or disease associations) would be needed.


[Read Paper](https://www.semanticscholar.org/paper/1fdff7edb4655416163ed78c76a6471bc026c45d)


### Case Report: The Monogenic Familial Steroid-Resistant Nephrotic Syndrome Caused by a Novel Missense Mutation of NPHS2 Gene A593C in a Chinese Family

**Why Not Relevant**: The provided paper describes clinical and genetic findings in two siblings with steroid-resistant nephrotic syndrome caused by a homozygous missense variant in NPHS2 and discusses NPHS2-related pathology and diagnostic implications. The text contains no mention of ENDOG, no data on variants in ENDOG, and no mechanistic discussion involving ENDOG. Therefore the paper offers no direct or mechanistic evidence supporting or refuting the claim that ENDOG is associated with a Mendelian disease. Additional limitations: the study is disease- and gene-specific (SRNS and NPHS2), has a very small sample (two affected siblings), and explicitly states no other SRNS-related gene variants were determined — but this negative statement applies only to SRNS-related genes and does not imply anything about ENDOG's involvement in other Mendelian conditions. Given the complete absence of ENDOG-related content, relevance to the claim is effectively zero.


[Read Paper](https://www.semanticscholar.org/paper/26ba0a4e7e5097751e78e184e30cde938546d66d)


### HOXA11 is another monogenic cause of congenital anomalies of the kidney and urinary tract

**Why Not Relevant**: The provided paper content exclusively addresses HOXA11 and its potential role as a monogenic cause of CAKUT. There is no mention of the gene ENDOG, no presented genetic association studies, no variant data, no phenotype descriptions related to ENDOG, and no functional or mechanistic experiments involving ENDOG. Because the excerpt neither references ENDOG nor supplies any transferable mechanistic evidence that would implicate ENDOG in a Mendelian disease, the paper content is not relevant to the claim that ENDOG is likely associated with at least one Mendelian disease.


[Read Paper](https://www.semanticscholar.org/paper/db3d83a93884ccd5d66fca19ab8728ccabfc9ffc)


### Autosomal Dominant Intellectual Development Disorder-6 (MRD6) Without Seizures Linked to a De Novo Mutation in the grin2b Gene Revealed by Exome Sequencing: A Case Report of a Moroccan Child

**Why Not Relevant**: The paper describes a clinical case and genetic analysis implicating GRIN2B (a gene associated with autosomal dominant intellectual development disorder-6, MRD6). It provides variant-level evidence (a de novo nonsense variant c.3912C>G, p.Tyr1304Ter in GRIN2B), ACMG pathogenicity classification, and discussion of GRIN2B's role in neurodevelopment. Nowhere in the provided content is the gene ENDOG mentioned, nor are any data, analyses, or mechanistic pathways involving ENDOG discussed. There is no direct genetic, clinical, population, or mechanistic evidence linking ENDOG to a Mendelian disease in this paper. Therefore the paper is not relevant to the claim that ENDOG is likely associated with at least one Mendelian disease. Limitations that justify assigning relevance = 0.0: (1) absence of the gene name ENDOG or any synonyms/aliases in the text; (2) all genetic evidence pertains to a different gene (GRIN2B) with specific variant and phenotype associations; (3) no comparative analyses, gene lists, pathway links, or secondary findings are reported that might implicate ENDOG indirectly. Because none of the paper's content addresses ENDOG, it cannot provide supporting or refuting evidence for the user's claim.


[Read Paper](https://www.semanticscholar.org/paper/67d756e0cc302e5d46fdda9df34403f1f2de9a5a)


## Search Queries Used

- ENDOG mutation pathogenic variant Mendelian disease monogenic inherited case report family segregation homozygous heterozygous de novo

- ENDOG endonuclease G mitochondrial nuclease apoptosis mitochondrial DNA maintenance knockout mouse phenotype cardiomyopathy myopathy neurodegeneration infertility

- ENDOG gnomAD loss of function constraint pLI LoF observed expected loss of function variants homozygous frequency population exome sequencing cohort

- ENDOG ClinVar OMIM gene disease association review gene curation monogenic Mendelian autosomal recessive autosomal dominant


## Bibliometric Configuration

- Use Bibliometrics: True

- Author Impact Weight: 0.4

- Citation Impact Weight: 0.4

- Venue Impact Weight: 0.2
