## Supplementary material for "MENDELSEEK: An algorithm that predicts Mendelian Genes and elucidates what makes them special": ENGASE.txt

# Claim: The gene ENGASE is likely associated with at least one Mendelian disease.

**Status**: processed

**Overall Rating**: -1

**Explanation**:

No relevant papers were found that support or refute this claim.


## Relevant Papers


## Other Reviewed Papers


### Exome sequencing identifies a de novo mutation of CTNNB1 gene in a patient mainly presented with retinal detachment, lens and vitreous opacities, microcephaly, and developmental delay

**Why Not Relevant**: This paper is wholly focused on CTNNB1 (β-catenin) and a de novo loss-of-function mutation (c.1672C>T, p.Gln558X) found in a patient with developmental abnormalities and ophthalmic findings. The text discusses CTNNB1’s role in cell adhesion and Wnt signaling, prior CTNNB1 gain- and loss-of-function findings, and presents a case report and whole-exome sequencing result implicating CTNNB1. Nowhere in the provided content is the gene ENGASE mentioned, nor are there any data, analyses, or mechanistic discussions that involve ENGASE or its protein product. Therefore this paper provides neither direct evidence (variants in ENGASE segregating with disease, case reports, association statistics) nor mechanistic evidence (functional studies implicating ENGASE in pathways causing Mendelian phenotypes) relevant to the claim that ENGASE is likely associated with at least one Mendelian disease. The absence of any mention of ENGASE in this CTNNB1-centered report means the paper cannot be used to support or refute the claim about ENGASE; it is simply not applicable. Note that absence of evidence in this single paper does not constitute evidence that ENGASE is not disease-associated — it only indicates this paper provides no information relevant to that claim. For assessment of ENGASE, studies that (a) report pathogenic ENGASE variants in affected individuals with segregation or de novo status, (b) present case series or population-level association data, or (c) provide functional/mechanistic data showing how ENGASE perturbation causes Mendelian phenotypes are required.


[Read Paper](https://www.semanticscholar.org/paper/da848f914a19f0732ef95266b453087bda26d85e)


### De novo apparent loss-of-function mutations in PRR12 in three patients with intellectual disability and iris abnormalities

**Why Not Relevant**: The provided paper content contains a single sentence: "It is proposed that haploinsufficiency of PRR12 is associated with this novel multisystem neurodevelopmental disorder." This sentence refers specifically to PRR12 and to a proposed association between PRR12 haploinsufficiency and a neurodevelopmental disorder. There is no mention of ENGASE, no data, no methodological details, and no mechanistic discussion that links PRR12 to ENGASE. Therefore the content provides no direct evidence that ENGASE is associated with any Mendelian disease, nor any mechanistic evidence that would make such an association plausible. Furthermore, one cannot infer an association for ENGASE from an association reported for a different gene (PRR12) without additional data showing a relationship (e.g., genetic interaction, pathway overlap, co-segregation, shared phenotype with ENGASE variants, or population/clinical genetic evidence). The single-sentence excerpt also lacks information on study design, cohort size, statistical significance, variant types, inheritance pattern, functional validation, or phenotypic specifics; these omissions further preclude using this content to support or refute the claim about ENGASE. For these reasons relevance is effectively zero and no verbatim excerpts supporting or refuting the ENGASE claim are available in the provided content.


[Read Paper](https://www.semanticscholar.org/paper/47ae30fde0955d6a9334f598a39891d5ab1e7109)


### Biallelic loss of TRAPPC9 function links vesicle trafficking pathway to autosomal recessive intellectual disability

**Why Not Relevant**: The provided paper content does not mention ENGASE anywhere. The single sentence attributes the observed neurological findings to a loss-of-function variant in TRAPPC9 and proposes vesicle trafficking as the mechanism. Because there is no discussion, data, or analysis concerning ENGASE (no mention of variants, pathogenicity, segregation, functional assays, or clinical association for ENGASE), the paper provides no direct evidence for ENGASE being associated with a Mendelian disease. The content is only tangentially relevant insofar as it identifies an alternative gene (TRAPPC9) explaining the phenotype; however, absence of ENGASE in this report does not constitute evidence that ENGASE is not disease-associated. In summary, the paper neither supports nor provides direct mechanistic evidence for ENGASE involvement, so its relevance to the claim that ENGASE is likely associated with at least one Mendelian disease is negligible.


[Read Paper](https://www.semanticscholar.org/paper/2b0480686f56a16015b5c96581f5c10e1286f759)


### A murine model of hnRNPH2-related neurodevelopmental disorder recapitulates clinical features of human disease and reveals a mechanism for genetic compensation of HNRNPH2

**Why Not Relevant**: This paper describes mutations, mechanisms, and mouse models for HNRNPH2 and explicitly discusses Hnrnph2 and its paralog Hnrnph1. The content makes no mention of ENGASE, ENGASE variants, ENGASE-related phenotypes, or any experiments involving ENGASE. Because the claim concerns whether ENGASE is likely associated with at least one Mendelian disease, evidence relevant to that claim must implicate ENGASE directly (e.g., pathogenic variants in ENGASE segregating with disease, functional studies of ENGASE variants, patient phenotypes linked to ENGASE disruption, or mechanistic pathways specifically involving ENGASE). None of those types of data or statements appear in the provided text. While the paper does provide mechanistic concepts (e.g., toxic gain-of-function, genetic compensation by a paralog) that are broadly applicable to genetic disease biology, these general principles cannot be used to infer that ENGASE is associated with a Mendelian disorder without direct data on ENGASE. Therefore the paper provides no direct evidence and no specific mechanistic evidence relevant to the ENGASE claim. Limitations that lead to a relevance score of 0.0: (1) No occurrence of the gene name ENGASE in the text; (2) All experimental results, models, and conclusions concern HNRNPH2/Hnrnph2 and HNRNPH1/Hnrnph1; (3) No patient genotypes, segregation data, variant pathogenicity assessment, or functional assays are presented for ENGASE; (4) General mechanistic parallels (e.g., compensation by paralogs) are insufficient to support a gene-disease association for a different gene. Because of these points, the paper cannot be used to support or refute the claim that ENGASE is likely associated with at least one Mendelian disease.


[Read Paper](https://www.semanticscholar.org/paper/3216fe04785cf8d230007ebf3dedfd99dceedd8d)


### De novo apparent loss-of-function mutations in PRR12 in three patients with intellectual disability and iris abnormalities

**Why Not Relevant**: The provided paper content contains a single sentence: "It is proposed that haploinsufficiency of PRR12 is associated with this novel multisystem neurodevelopmental disorder." This sentence refers exclusively to PRR12 and makes a proposal about PRR12 haploinsufficiency; it does not mention the gene ENGASE, any variants in ENGASE, nor any clinical, genetic, or mechanistic data linking ENGASE to disease. Therefore the paper content gives neither direct evidence (no reported ENGASE variants segregating with disease, no case reports, no statistical association) nor mechanistic evidence (no biochemical pathway, expression, functional studies, or animal models) that would support the claim that ENGASE is associated with a Mendelian disease. Additional points limiting any inference from this content: (1) the statement is explicitly about a different gene (PRR12), so it cannot be applied to ENGASE without additional information linking the two genes (no such link is provided); (2) the sentence is phrased as a proposal, not as demonstrated evidence, so even for PRR12 the content is tentative; (3) there are no methodological details, results, variant descriptions, inheritance patterns, or statistical measures that could allow assessment of strength of evidence; and (4) absence of mention of ENGASE in this excerpt is not evidence that ENGASE is not associated with disease, only that this paper content provides no support. In sum, based solely on the provided text, the paper is not relevant to the claim that ENGASE is likely associated with at least one Mendelian disease.


[Read Paper](https://www.semanticscholar.org/paper/5b99961ae102b22a3dec205081dd94074df20cad)


## Search Queries Used

- ENGASE mutation human patient loss of function pathogenic clinical exome Mendelian disease intellectual disability developmental delay

- ENGASE knockout mouse endo beta N acetylglucosaminidase deficiency phenotype neurodevelopmental glycosylation pathway model organism

- ENGASE NGLY1 interaction genetic modifier rescue N glycanase 1 deglycosylation pathway ENGase inhibition NGLY1 deficiency patient cells

- ENGASE ClinVar OMIM gnomAD loss of function constraint pLI haploinsufficiency allele frequency rare variants Mendelian association


## Bibliometric Configuration

- Use Bibliometrics: True

- Author Impact Weight: 0.4

- Citation Impact Weight: 0.4

- Venue Impact Weight: 0.2
