## Supplementary material for "MENDELSEEK: An algorithm that predicts Mendelian Genes and elucidates what makes them special": ENO2.txt

# Claim: The gene ENO2 is likely associated with at least one Mendelian disease.

**Status**: processed

**Overall Rating**: -1

**Explanation**:

No relevant papers were found that support or refute this claim.


## Relevant Papers


## Other Reviewed Papers


### Prevalence and phenotypic impact of rare potentially damaging variants in autism spectrum disorder

**Why Not Relevant**: The provided paper content does not contain any mention of the gene ENO2, any specific genes, nor any discussion of Mendelian diseases. The only sentence supplied is: "In this epidemiological sample, rare pdSNV were more common than pdCNV and the combined yield of potentially damaging variation was substantial at 27%, providing compelling rationale for the use of high-throughout sequencing as part of routine clinical workup for ASD and support the development of precision medicine in ASD." This statement is a general summary about the frequency and combined yield of potentially damaging single-nucleotide variants (pdSNV) and copy-number variants (pdCNV) in an ASD sample and advocates sequencing as a clinical tool. It does not (a) identify ENO2, (b) report any association between ENO2 and disease, (c) describe a Mendelian phenotype or inheritance pattern, or (d) provide mechanistic data linking any particular gene to disease. Therefore there is no direct evidence in this content for or against the claim that ENO2 is likely associated with at least one Mendelian disease. At best, the sentence gives an indirect, very general rationale that high-throughput sequencing can identify potentially damaging variants in ASD cohorts; however, without gene-level results, variant annotation, segregation, functional validation, or clinical phenotyping, that general statement cannot be used to support the specific claim about ENO2. Key limitations: single-sentence excerpt, no gene names, no variant data, no clinical/segregation evidence, and no mechanistic descriptions relevant to ENO2.


[Read Paper](https://www.semanticscholar.org/paper/fa3391717bcabc5b0a0eb7766e6489d8da5069b4)


### ZMYND11 variants are a novel cause of centrotemporal and generalised epilepsies with neurodevelopmental disorder

**Why Not Relevant**: This paper discusses ZMYND11 and its role in chromosome 10p15.3 microdeletion syndrome and epilepsy. The provided content makes no mention of the gene ENO2, its variants, diseases associated with ENO2, or any mechanistic connection involving ENO2. Key points in the excerpt focus on: (a) ZMYND11 as the critical gene in 10p15.3 microdeletion syndrome, (b) the epilepsy phenotypes observed in individuals with pathogenic variants in ZMYND11, (c) that variants were mostly de novo and distributed across ZMYND11 with no precise genotype–phenotype correlation, and (d) a possible epileptogenic mechanism linked to interaction with histone H3.3. None of these items provide direct evidence that ENO2 is associated with a Mendelian disease, nor do they provide mechanistic evidence that would plausibly link ENO2 to such a disease. 

Methodological notes that further justify the low relevance: the genetic analyses described (gene panels or exome sequencing) are general approaches that could detect variants in many genes, but the paper reports only findings for ZMYND11 and does not report ENO2 findings or negative results for ENO2. The mechanistic detail given (interaction with histone H3.3) pertains to chromatin reading and is unrelated to the known biology of ENO2 (neuron-specific enolase, a glycolytic enzyme). 

Therefore, this paper does not provide direct evidence for or against the claim that ENO2 is likely associated with at least one Mendelian disease, nor does it provide mechanistic evidence relevant to ENO2. If you want an evidence-based assessment of ENO2 specifically, I can search and summarize gene-disease resources (e.g., OMIM, ClinVar, ClinGen curation, PubMed literature, gnomAD constraint metrics) for any reported Mendelian associations of ENO2.


[Read Paper](https://www.semanticscholar.org/paper/dfbe5519d7bbba01d84445ec593cc8506c0395d3)


### Detailed Clinical and Psychological Phenotype of the X-linked HNRNPH2-Related Neurodevelopmental Disorder

**Why Not Relevant**: The paper exclusively studies HNRNPH2 (an X-linked gene) and its associated neurodevelopmental disorder. There is no mention of the gene ENO2, no data on ENO2 variants, no mechanistic experiments involving ENO2, and no analysis that would permit inference about ENO2's role in Mendelian disease. Because the paper's whole content concerns a different gene, it cannot provide direct evidence supporting or refuting the claim that ENO2 is likely associated with at least one Mendelian disease. The absence of ENO2 data in this report is not evidence against ENO2 involvement (i.e., it is not a negative screen); it simply falls outside the scope needed to evaluate the claim.


[Read Paper](https://www.semanticscholar.org/paper/d13ac29fbc1ed2aa5c12352383e3f10f263cebb8)


### Case report: A novel FBXW7 gene variant causes global developmental delay

**Why Not Relevant**: This paper is not relevant to the claim that ENO2 is likely associated with at least one Mendelian disease. The entire provided content concerns a single case report implicating a de novo pathogenic variant in FBXW7 (c.1612C>T (p.Gln538*)) as the cause of a neurodevelopmental disorder. The text does not mention ENO2, does not provide any genetic, clinical, or mechanistic data about ENO2, and does not draw any connections between FBXW7 and ENO2. Therefore there is no direct evidence (e.g., ENO2 variants segregating with disease, case reports, cohort studies) nor mechanistic evidence (e.g., functional studies, biochemical pathways implicating ENO2 in disease processes) in this paper to support or to refute the claim. 

Limitations and caveats: the absence of ENO2 in this paper is not evidence that ENO2 is not associated with a Mendelian disease — it simply means this particular study provides no relevant data. This is a single-case exome-based diagnostic report focused on FBXW7; it cannot be used to assess ENO2’s disease associations. Additionally, because only an excerpt of the paper was provided, if the full article elsewhere contained any mention of ENO2 (unlikely given the provided objective, results, and conclusion), that text was not supplied here. 

Because relevance is < 0.1, no verbatim excerpts were extracted as evidence for or against the claim.


[Read Paper](https://www.semanticscholar.org/paper/bb0b7ce4ae459771c428a6c9d1dbc93aecc6bcdc)


### An adult male with SHANK2 variant with epilepsy and obsessive‐compulsive disorder: Expanding the shankopathy phenotypic spectrum

**Why Not Relevant**: This paper does not provide any direct or mechanistic evidence regarding the gene ENO2. The entire report discusses SHANK2 (and shankopathies) — for example, it describes a de novo heterozygous frameshift variant in SHANK2 (c.579del, p.Ser194Profs*31) and phenotypes in a single patient with ASD and epilepsy — but ENO2 (or enolase/neuron-specific enolase) is not mentioned anywhere in the provided text. Therefore there are no verbatim sentences or paragraphs in this paper that support or refute the claim that ENO2 is likely associated with a Mendelian disease. 

Because the paper focuses solely on SHANK2, any evidence it provides is relevant to SHANK2 gene-disease association (including statements about loss-of-function as a mechanism for SHANK2, mouse-model synaptic transmission differences, and the patient phenotype), but none of those findings can be used as evidence for ENO2. There is no mechanistic data linking ENO2 to the phenotypes, no reported ENO2 variants, and no population or functional data about ENO2 in this text. 

Limitations relevant to your claim: (1) absence of evidence in this paper is not evidence of absence for ENO2 — it simply was not studied here; (2) findings about SHANK2 cannot be extrapolated to ENO2 because the genes have different functions and disease mechanisms; (3) to assess ENO2 you would need targeted evidence such as reported pathogenic ENO2 variants in affected individuals, segregation data, functional studies, or curation in clinical databases (OMIM, ClinVar, ClinGen).


[Read Paper](https://www.semanticscholar.org/paper/a07cc3110dc2f48548f0a08d78036fcfc8278ab4)


### Case Report: Identification of a rare nonsense mutation in the POC1A gene by NGS in a diabetes mellitus patient

**Why Not Relevant**: The provided paper reports a case study of diabetes caused by a POC1A truncating mutation and describes trio whole-exome sequencing and variant verification, but it does not mention the gene ENO2 anywhere in the objective, methods, results, or conclusion. Therefore the paper provides no direct evidence (reports of ENO2 variants linked to disease) nor mechanistic evidence (functional data implicating ENO2 in Mendelian disease) relevant to the claim that ENO2 is likely associated with at least one Mendelian disease. Although the study used whole-exome sequencing (with stated high coverage), which in principle could detect ENO2 coding variants, the absence of any reported ENO2 findings means this paper cannot be used to support or refute the claim. Additional limitations: this is a single-patient report focused on POC1A, so even if ENO2 had been reported, a single case would provide limited evidence for a Mendelian association without segregation, functional validation, or recurrence in additional affected individuals.


[Read Paper](https://www.semanticscholar.org/paper/8f90a0a9b3c51d0406ae0337e2a4d8c2c0204107)


### TARGETED SEQUENCING REVEAL TWO PATHOGENIC MUTATIONS IN LDLR AND APOE GENES IN PATIENT WITH FAMILY HYPERCHOLESTEROLEMIA: CASE REPORT

**Why Not Relevant**: This paper does not provide any direct or mechanistic evidence regarding the gene ENO2. The content exclusively discusses familial hypercholesterolemia (FH) and genetic findings in LDLR, APOE and a few other genes (APOB, TTN, COL3A1, NOTCH1, LAMA4, MYPN, DMD). There is no mention of ENO2, no reported variants in ENO2, and no mechanistic discussion that would link ENO2 to FH or to any Mendelian disease. Key relevant sentences illustrating the paper's focus include: "In our study, we identified in FH patient likely pathogenic variant of LDLR and APOE genes.", "Sequencing of 96 genes and bioinformatics analysis identified 2 likely pathogenic variants - C526T:p.R176C in APOE and G295A:p.E99K in LDLR gene, 4 uncertain significance variants in APOB, TTN, COL3A1, NOTCH1 and 90 benign variants in APOB, TTN, LAMA4, MYPN, DMD.", and "In our study, we found the genetic mutations - two likely pathogenic genetic variants LDLR (G295A:p.E99K) and APOE (C526T:p.R176C) associated with FH in patient with clinical phenotype of FH using TruSight Cardio targeted panel of 96 genes." Because ENO2 is not discussed, there is neither supporting nor refuting evidence in this paper regarding the claim that "ENO2 is likely associated with at least one Mendelian disease." Limitations to this conclusion: absence of any mention of ENO2 here does not prove ENO2 has no Mendelian associations; the study used a targeted gene panel (TruSight Cardio / a 96-gene panel) and thus may simply have not assayed ENO2. To evaluate the claim about ENO2, one would need data from studies that specifically include ENO2 (e.g., whole exome/genome sequencing datasets, case-control studies of ENO2 variants, ClinVar/OMIM annotations, or functional/mechanistic experiments).


[Read Paper](https://www.semanticscholar.org/paper/04808c76b3962e7f9dc6e3ebec4ddbb8d15d0607)


### Whole exome sequencing identifies variants in the \(\textit{TNNI3}\) gene in vietnamese patient with restrictive cardiomyopathy - a case report

**Why Not Relevant**: The supplied paper does not provide any evidence (direct or mechanistic) relating to the gene ENO2. The study focuses on restrictive cardiomyopathy (RCM) and reports pathogenic variants in TNNI3. Relevant verbatim sentences from the paper that demonstrate the paper's focus (and absence of ENO2) include: "In this study, we conducted sequencing of the entire gene coding region (WES) in the patient and identified a compound heterozygote variants (c.289C>G, p.Arg97Gly and c.433C>T, p.Arg145Trp) in the TNNI3 gene." "These variants were inherited from the patient's father and mother, who were heterozygous variant carriers." "These variants were also identified as the pathogenic variants in the ClinVar database (accession number VCV001331910.2 and VCV000012426.28, respectively) and were the cause of the patient's disease."

Why relevance = 0.0: (1) The text contains no mention of ENO2, enolase, or any synonyms; therefore there is no direct evidence that ENO2 is associated with any Mendelian disease. (2) There is no mechanistic discussion linking ENO2 to RCM, to the pathogenic variants described, or to any Mendelian inheritance pattern. (3) The paper is a case-focused genetic diagnosis of RCM attributing disease causation to TNNI3 variants (single-family WES analysis), so its findings are specific to TNNI3 and are not generalizable to other genes not discussed.

Limitations and caveats: Because the provided content is limited to a single paper excerpt, absence of evidence in this text does not prove that ENO2 is not associated with a Mendelian disease — it only means this paper gives no support for that claim. To evaluate the claim about ENO2, one should search genetic-disease resources (e.g., OMIM, ClinVar, HGMD) and the broader literature for ENO2/ENO2-related pathogenic variants and mechanistic studies.


[Read Paper](https://www.semanticscholar.org/paper/83a11d71a34fe3ae5618faf4b4a593ecd29e0884)


## Search Queries Used

- ENO2 mutation pathogenic Mendelian disease case report ClinVar OMIM patient sequencing

- enolase 2 ENO2 rare variant exome sequencing cohort neurodevelopmental disorder intellectual disability epilepsy autism

- enolase 2 neuron specific enolase knockout mouse phenotype neuronal metabolism glycolysis neurodegeneration functional assay

- ENO2 gene constraint gnomAD pLI loss of function intolerance ClinGen gene disease validity systematic review monogenic association


## Bibliometric Configuration

- Use Bibliometrics: True

- Author Impact Weight: 0.4

- Citation Impact Weight: 0.4

- Venue Impact Weight: 0.2
