## Supplementary material for "MENDELSEEK: An algorithm that predicts Mendelian Genes and elucidates what makes them special": EZR.txt

# Claim: The gene EZR is likely associated with at least one Mendelian disease.

**Status**: processed

**Overall Rating**: -1

**Explanation**:

No relevant papers were found that support or refute this claim.


## Relevant Papers


## Other Reviewed Papers


### Precision Medicine in Internal Medicine

**Why Not Relevant**: The provided paper is a general review about clinical genetic testing, exome/genome sequencing, and variant interpretation. It contains general statements about the capabilities of ES/GS, variant frequencies, and the number of genes currently implicated in Mendelian disease, but it contains no mention of EZR, no data (genetic, segregation, functional, case reports, or statistical association) regarding EZR, and no mechanistic experiments specific to EZR. Therefore it provides no direct evidence that the gene EZR is associated with any Mendelian disease. The few excerpts above offer only indirect, general context (e.g., that many genes remain to be linked to disease and that ES can discover novel associations), which is insufficient to support the claim that EZR is likely associated with at least one Mendelian disease.


[Read Paper](https://www.semanticscholar.org/paper/c7199e8a5226f8d9d55777da0e15e91797bb459a)


### Simultaneous identification and prioritization of variants in familial, de novo, and somatic genetic disorders with VariantMaster

**Why Not Relevant**: The paper describes a variant-prioritization algorithm (VariantMaster) and demonstrates its application to discover de novo mutations and causative variants in at least one rare Mendelian disease. However, the provided content contains no mention of the gene EZR, no case reports, no variant-level data, no statistical associations, and no functional or mechanistic studies specific to EZR. Therefore it provides no direct evidence that EZR is associated with any Mendelian disease. The only indirect relevance is methodological: the tool described could, in principle, be used to identify causative EZR variants in future studies, but that is speculative and not evidence. For these reasons the paper's relevance to the claim that "EZR is likely associated with at least one Mendelian disease" is very low.


[Read Paper](https://www.semanticscholar.org/paper/a56e2dc7369c76b03ad5e9309093beb98a159719)


### The Burden of Candidate Pathogenic Variants for Kidney and Genitourinary Disorders Emerging From Exome Sequencing

**Why Not Relevant**: The provided paper excerpt describes methods and scope for identifying and curating genes associated with Mendelian kidney and genitourinary disorders and for analyzing variant prevalence in a large cohort. However, the excerpt does not mention the gene EZR by name, does not present a gene-level list or tables showing which genes were included in the curated 625-gene set, and does not provide any results or discussion specific to EZR. Therefore, there is no direct evidence in the excerpt that EZR is associated (or not associated) with any Mendelian disease. The study's design and dataset could in principle contain relevant information (e.g., EZR might be among the curated genes or might have variants observed), but without the gene list or gene-specific results, the paper excerpt is insufficient to evaluate the claim. Because of this absence of gene-level information, the excerpt provides only minimal, indirect relevance (methodological context) and cannot be used to support or refute the claim that "The gene EZR is likely associated with at least one Mendelian disease."


[Read Paper](https://www.semanticscholar.org/paper/a75e297208c04992568e607d19aa9b9aaebd09d5)


### Clingen Coagulation Factor Deficiency Variant Curation Expert Panel: Meeting the Need for Recommendations to Curate Variants in the Coagulation Factor Genes

**Why Not Relevant**: The provided paper content is focused on the development and specification of variant-interpretation guidelines (ClinGen CFD-VCEP) specifically for coagulation factor genes F8 and F9 and on curating variants in those genes. There is no mention of the gene EZR, no case reports, no genetic association data, and no functional/mechanistic experiments relating EZR to any Mendelian phenotype. Therefore the paper provides no direct or mechanistic evidence for or against the claim that EZR is associated with a Mendelian disease. The absence of discussion of EZR in this gene- and disease-specific guideline/methods paper is expected and does not constitute evidence about EZR generally. To evaluate the claim about EZR, one would need studies (case reports, family segregation, population-variant analyses, functional assays) that specifically assess EZR variants and link them to a Mendelian phenotype; none of these are present in the supplied text.


[Read Paper](https://www.semanticscholar.org/paper/9ea8d4ce19622ae1c6b0cdfd40848c93199f6280)


## Search Queries Used

- EZR ezrin germline mutation Mendelian disease monogenic autosomal dominant autosomal recessive family segregation case report

- EZR ezrin ClinVar OMIM ClinGen HGMD pathogenic variant clinical significance gene disease association

- EZR ezrin loss of function mutation actin cytoskeleton epithelial polarity microvillus formation knockout mouse developmental congenital phenotype

- EZR ezrin somatic mutation cancer carcinoma fusion oncogene somatic versus germline exome sequencing population frequency gnomAD no disease association


## Bibliometric Configuration

- Use Bibliometrics: True

- Author Impact Weight: 0.4

- Citation Impact Weight: 0.4

- Venue Impact Weight: 0.2
