## Supplementary material for "MENDELSEEK: An algorithm that predicts Mendelian Genes and elucidates what makes them special": F2R.txt

# Claim: The gene F2R is likely associated with at least one Mendelian disease.

**Status**: processed

**Overall Rating**: -1

**Explanation**:

No relevant papers were found that support or refute this claim.


## Relevant Papers


## Other Reviewed Papers


### The Burden of Candidate Pathogenic Variants for Kidney and Genitourinary Disorders Emerging From Exome Sequencing

**Why Not Relevant**: The provided paper content describes methods and a curated gene list for Mendelian kidney and genitourinary disorders, cohort characteristics, and variant-filtering strategies, but it does not mention the gene F2R, present any variant-level results for F2R, or discuss F2R in a mechanistic or disease-association context. Therefore there is no direct or specific mechanistic evidence in this excerpt that F2R is associated with any Mendelian disease. Because the gene lists and per-gene results are not included in the excerpt, one cannot infer F2R's status from the provided text; additional sections (full gene list or results) would be required to evaluate the claim.


[Read Paper](https://www.semanticscholar.org/paper/a75e297208c04992568e607d19aa9b9aaebd09d5)


### The Brain Hemostasis: Implications for Stroke Pathology and New Therapeutic Strategies

**Why Not Relevant**: The paper does not mention F2R, PAR1, or specific genetic variants of the thrombin receptor by name, nor does it present any human genetic, linkage, segregation, case report, or cohort data that would support an association of F2R with a Mendelian disease. All relevant passages are descriptive of hemostatic mechanisms and tissue-specific expression of coagulation regulators (e.g., TFPI, TM, EPCR, protease-activated receptors) and include animal-model data for thrombomodulin, but none provide direct genetic evidence (variants, inheritance patterns, pathogenicity, or documented Mendelian phenotypes) implicating F2R. Therefore the paper provides only peripheral mechanistic context (that thrombin signaling/PARs operate in endothelium) but no direct or substantive evidence that F2R is associated with any Mendelian disease.


[Read Paper](https://www.semanticscholar.org/paper/c49675c8f336dbc496be8edbb01cd94b01933f0a)


### Precision Medicine in Internal Medicine

**Why Not Relevant**: This paper is a general review about genetic testing strategies, exome/genome sequencing, variant interpretation, and the state of gene–disease discovery as of early 2019. It contains no mention of the gene F2R, no data on variants in F2R, and no discussion of any specific Mendelian disorders attributable to F2R. Therefore it provides no direct evidence supporting or refuting the specific claim that F2R is likely associated with at least one Mendelian disease. The included excerpts supply relevant methodological and contextual information (how gene–disease associations are discovered and judged), which informs how one would evaluate evidence for F2R, but they do not constitute evidence about F2R itself. Because the paper lacks any direct or gene-specific mechanistic data concerning F2R, its relevance to the claim is very low.


[Read Paper](https://www.semanticscholar.org/paper/c7199e8a5226f8d9d55777da0e15e91797bb459a)


## Search Queries Used

- F2R protease activated receptor 1 PAR1 thrombin receptor mutation Mendelian disease family segregation autosomal dominant autosomal recessive case report

- F2R ClinVar OMIM HGMD F2R gene pathogenic variant clinical significance molecular diagnosis database entry

- F2R loss of function knockout mouse protease activated receptor 1 thrombin receptor platelet function bleeding disorder vascular anomaly hereditary thrombosis functional study

- F2R rare variant exome sequencing family based linkage de novo mutation Mendelian inheritance severe early onset disease common variant GWAS thrombosis platelet aggregation risk allele


## Bibliometric Configuration

- Use Bibliometrics: True

- Author Impact Weight: 0.4

- Citation Impact Weight: 0.4

- Venue Impact Weight: 0.2
