## Supplementary material for "MENDELSEEK: An algorithm that predicts Mendelian Genes and elucidates what makes them special": FAM13A.txt

# Claim: The gene FAM13A is likely associated with at least one Mendelian disease.

**Status**: processed

**Overall Rating**: -1

**Explanation**:

No relevant papers were found that support or refute this claim.


## Relevant Papers


## Other Reviewed Papers


### Annotation of uORFs in the OMIM genes allows to reveal pathogenic variants in 5′UTRs

**Why Not Relevant**: The provided paper abstract discusses uORFs, their annotation in OMIM disease-associated genes, experimental validation in several named genes, and the presence of uORF-disrupting variants in HGMD/ClinVar that could cause Mendelian disorders. However, the abstract does not mention FAM13A anywhere, nor does it provide gene-specific results for FAM13A. Therefore the paper content supplied contains only general mechanistic and methodological context showing that uORF disruption can cause Mendelian disease, but it contains no direct evidence (experimental, database-reported, or statistical) linking FAM13A to any Mendelian disease. Because the claim is a gene-specific assertion and the paper content lacks any mention or data about FAM13A, the relevance to the claim is minimal.


[Read Paper](https://www.semanticscholar.org/paper/5d45c0cdf801505df45fc93a466a2a4101a12871)


### The Burden of Candidate Pathogenic Variants for Kidney and Genitourinary Disorders Emerging From Exome Sequencing

**Why Not Relevant**: This paper does not provide evidence for or against the claim that FAM13A is likely associated with at least one Mendelian disease. The provided content focuses specifically on identifying and curating genes linked to Mendelian kidney and genitourinary disorders using OMIM and Orphanet, producing a curated list of 625 genes that were annotated and analyzed for variant prevalence. The text excerpts supplied do not mention FAM13A by name, do not reproduce the curated gene list, nor present gene-level results that would allow determination of whether FAM13A is included or has evidence of Mendelian disease association. Furthermore, the study’s phenotypic focus (kidney and genitourinary disorders) and cohort design make it unlikely to address Mendelian associations for genes primarily implicated in other systems (for example, FAM13A has been implicated in pulmonary traits in other literature). While the authors indicate the curated list is available on a laboratory website, that external resource was not provided here; without access to the specific gene list or gene-level annotations/results, no direct or mechanistic evidence concerning FAM13A can be extracted from the supplied paper content. To assess the claim about FAM13A, one would need either (a) the paper’s supplementary gene list/annotations showing FAM13A is included with supporting evidence, (b) gene-level results or variant annotations implicating FAM13A in a Mendelian phenotype, or (c) independent sources (OMIM, ClinVar, peer-reviewed reports) that directly document a Mendelian disease association for FAM13A.


[Read Paper](https://www.semanticscholar.org/paper/a75e297208c04992568e607d19aa9b9aaebd09d5)


### A pathogenic variant in RAB32 causes autosomal dominant Parkinson’s disease and activates LRRK2 kinase

**Why Not Relevant**: The provided paper text discusses RAB GTPases, specifically identifying RAB32 c.213C>G (Ser71Arg) as co-segregating with autosomal dominant Parkinson disease and activating LRRK2 kinase. There is no mention of the gene FAM13A anywhere in the supplied content, nor any data, results, mechanisms, or discussion that link FAM13A to Mendelian disease. Therefore this paper provides no direct evidence for or against the claim that FAM13A is likely associated with at least one Mendelian disease. Additionally, there is no mechanistic information in the text that could plausibly be re-interpreted to implicate FAM13A (FAM13A is not a RAB GTPase and is not discussed). To evaluate the claim about FAM13A would require examining other literature and databases (e.g., OMIM, ClinVar, PubMed, gnomAD) for reports of pathogenic variants, segregation in families, functional studies, or mechanistic links to Mendelian phenotypes.


[Read Paper](https://www.semanticscholar.org/paper/a36d28e828ede7c87c650af5f5117982b9749e7b)


### Trio-based whole exome sequencing in patients with suspected sporadic inborn errors of immunity: A retrospective cohort study

**Why Not Relevant**: The provided paper content focuses on de novo variants discovered by trio-based whole exome sequencing in a cohort of 123 patients with suspected inborn errors of immunity (IEI). The genes explicitly discussed and functionally validated in this excerpt are IEI-relevant genes (for example NLRP3, RELA, and FBXW11) and other candidate genes (PSMB10, DDX1, KMT2C). The gene FAM13A is not mentioned anywhere in the provided text. Therefore this paper provides no direct genetic, clinical, or mechanistic evidence linking FAM13A to any Mendelian disease. Additionally, the study population is specialized (suspected sporadic IEI) and the analysis emphasized de novo coding and splice variants; absence of FAM13A findings in this specific cohort and variant class is weak negative evidence at best and does not permit a general conclusion about FAM13A's association with Mendelian disease. Because there is no mention or data on FAM13A in the supplied content, the paper is not relevant to supporting or refuting the claim that FAM13A is likely associated with at least one Mendelian disease.


[Read Paper](https://www.semanticscholar.org/paper/cdde2232daad22b7d92ec15faa38b9776f1e4c15)


### Novel LSS variants in alopecia and intellectual disability syndrome: New case report and clinical spectrum of LSS‐related rare disease traits

**Why Not Relevant**: The provided paper text concerns pathogenic biallelic variants in the gene LSS and their association with specific Mendelian traits (congenital cataract type 44, autosomal recessive hypotrichosis type 14, and alopecia‑intellectual disability syndrome type 4). There is no mention of the gene FAM13A, no data, no variants, and no mechanistic discussion linking FAM13A to disease in the excerpt. Therefore the paper provides neither direct evidence (variant‑to‑disease observations for FAM13A) nor mechanistic evidence (biochemical, genetic interaction, pathway, expression, or functional data connecting FAM13A to a Mendelian phenotype) relevant to the claim. Because the content focuses exclusively on LSS, it cannot be used to support or refute the claim that FAM13A is likely associated with at least one Mendelian disease. No verbatim sentences can be extracted to support the FAM13A claim, and no indirect links or caveats are present in the text that would allow inference about FAM13A. If you can provide a paper that mentions FAM13A or studies of that gene, I can perform a direct relevance and evidence assessment.


[Read Paper](https://www.semanticscholar.org/paper/fe90a5bb55278bc80588741eed5def6cf4f1aa82)


### Genetic Diagnosis of Rubinstein–Taybi Syndrome With Multiplex Ligation-Dependent Probe Amplification (MLPA) and Whole-Exome Sequencing (WES): Case Series With a Novel CREBBP Variant

**Why Not Relevant**: This paper focuses on Rubinstein–Taybi Syndrome (RSTS) and reports genetic findings involving CREBBP and EP300 (e.g., a large heterozygous deletion spanning CREBBP and three different CREBBP point/frameshift variants). The provided text contains no mention of FAM13A, no variants in FAM13A, and no mechanistic discussion implicating FAM13A in RSTS or any other Mendelian condition. Therefore the paper provides neither direct evidence (no observed FAM13A variants, no segregation, no case reports) nor mechanistic evidence (no pathway, functional or biochemical discussion involving FAM13A) to support or refute the claim that FAM13A is likely associated with at least one Mendelian disease. Limitations of the paper (small cohort of five patients, disease-specific focus) do not change this conclusion: absence of evidence for FAM13A in this disease-focused study means the paper is not informative about FAM13A’s disease associations. To assess the claim about FAM13A one should consult other sources (e.g., ClinVar, OMIM, ClinGen gene-disease validity curation, HGMD, and targeted case reports or large-scale sequencing studies) for variant, segregation, functional, and replication evidence regarding FAM13A.


[Read Paper](https://www.semanticscholar.org/paper/0ac8d8dae09dd7889f1b27fa9d2f397eb4565e16)


### [A rare variant in the TTR gene (p.E112K) is associated with systemic amyloidosis and a new symptom - skin hyperemia in response to ethanol intake: family segregation analysis, literature review, and a clinical case. Case report].

**Why Not Relevant**: The provided paper content exclusively discusses transthyretin amyloidosis (ATTR), pathogenic variants in the TTR gene, and clinical features associated with TTR mutations. There is no mention of the gene FAM13A, no data (genetic, clinical, linkage, segregation, case reports) involving FAM13A, and no mechanistic discussion connecting FAM13A to amyloid disease or any Mendelian phenotype. Therefore the paper gives no direct evidence (cases, variants, segregation, functional assays) nor mechanistic evidence (biological pathways, molecular interactions) supporting the claim that FAM13A is likely associated with at least one Mendelian disease. Additionally, because the article is a case report and literature review centered on TTR, its scope is narrow and not designed to survey or report associations for other genes. For these reasons the content is not relevant to the claim about FAM13A.


[Read Paper](https://www.semanticscholar.org/paper/2567df8ee078bbbd995e907e0465d6337ffeeb26)


### Co-Inheritance of Pathogenic Variants in PKD1 and PKD2 Genes Determined by Parental Segregation and De Novo Origin: A Case Report

**Why Not Relevant**: The provided paper content makes no mention of the gene FAM13A. All sentences in the supplied text refer to autosomal dominant polycystic kidney disease (ADPKD) and variants in PKD1 and PKD2, including a description of a family with a PKD2 nonsense variant and a novel PKD1 frameshift variant. Thus there is neither direct evidence (no reporting of variants, segregation, clinical association, or case reports involving FAM13A) nor mechanistic evidence (no functional, pathway, or interaction data implicating FAM13A) in the provided text that links FAM13A to any Mendelian disease. Limitations: this assessment is limited to the content you provided (which appears to be an abstract or summary focused on PKD1/PKD2). If the full paper or supplementary materials (not provided here) mention FAM13A, that content was not available for review. Based on the supplied text alone, the paper is irrelevant to the claim that FAM13A is likely associated with at least one Mendelian disease. For evaluation of that claim, consult primary sources that specifically report FAM13A variants, segregation data, functional studies, or curated databases (e.g., OMIM, ClinVar, HGMD).


[Read Paper](https://www.semanticscholar.org/paper/6bc1fefe3972092c6b64b12c8c0733ffcbf974ec)


### A rare Coffin-Siris syndrome induced by SOX11: a de novo nonsense variant of short stature

**Why Not Relevant**: The provided paper content contains a single statement: "The combination of clinical presentation and genetic testing results supports that the patient suffers from Coffin-Siris syndrome due to a genetic variant in the SOX11 gene, which expands the understanding of human gene variation." This statement concerns SOX11 as the implicated gene for Coffin-Siris syndrome and does not mention FAM13A. Therefore the paper provides no direct evidence (no genetic observations, segregation, case reports, or statistical association) that FAM13A is involved in any Mendelian disease. It also provides no mechanistic evidence (no functional studies, biochemical pathways, protein interactions, expression data, or model organism results) linking FAM13A to disease. Because the content is exclusively about a different gene (SOX11) and a clinical diagnosis (Coffin-Siris syndrome), it cannot be used to support or refute the claim that "FAM13A is likely associated with at least one Mendelian disease." To evaluate that claim requires literature or data showing FAM13A variants segregating with a Mendelian phenotype, pathogenic/likely pathogenic classifications, functional validation, or population genetics evidence; none of these are present in the supplied text. Additionally, the supplied content appears to describe a single patient case concerning SOX11, which would be of limited generalizability even if it were about FAM13A (single-case reports provide weak evidence without segregation or functional follow-up).


[Read Paper](https://www.semanticscholar.org/paper/4183ebf1ebe7bfefc2d4a82261d6c09b9fcf8a9f)


### A case report of PGAP2-related hyperphosphatasia with impaired intellectual development syndrome in a Chinese family and literature review

**Why Not Relevant**: This paper describes pathogenic variants in PGAP2 and other genes in the glycosyl phosphatidylinositol (GPI)-anchor synthesis pathway and reports two patients with PGAP2-related hyperphosphatasia with impaired intellectual development syndrome (HPMRS). The text provided makes no mention of the gene FAM13A, contains no genetic, clinical, or mechanistic data linking FAM13A to any Mendelian disorder, and presents no co‑segregation, variant, functional, or pathway evidence that would support or refute an association between FAM13A and Mendelian disease. Mechanistically, the paper focuses on GPI-anchor synthesis and PGAP2 function; there is no discussion of FAM13A in those pathways or any cross-talk that might plausibly implicate FAM13A. Therefore, this paper provides no direct evidence and no mechanistic evidence relevant to the claim that FAM13A is likely associated with at least one Mendelian disease. Note that absence of evidence in this single paper does not disprove the claim broadly, but this paper cannot be used to support it.


[Read Paper](https://www.semanticscholar.org/paper/63cba8ad2bff7348418f63782b7cc408f084157d)


### Pathogenic paralogous variants can be used to apply the ACMG PS1 and PM5 variant interpretation criteria

**Why Not Relevant**: The provided paper content describes a general methodological advance (paralog-aware extension of ACMG PS1/PM5 criteria) and reports aggregate results across many genes and gene families, but it contains no mention of the specific gene FAM13A, nor does it present gene-level lists or examples that identify FAM13A as having disease-associated variants or being reclassified as associated with a Mendelian disorder. Without explicit inclusion of FAM13A in the results or examples, the paper cannot be used as direct evidence that FAM13A is likely associated with at least one Mendelian disease. The content only implies that, if FAM13A had pathogenic paralogous variants or were in a family with disease-associated genes, the method might provide supporting evidence—but such inference requires checking the paper's full gene-level data or supplemental tables (not provided here). Because no FAM13A-specific data are present, the relevance to the claim is very low.


[Read Paper](https://www.semanticscholar.org/paper/0a921a972e8db185945c5510191b7ac52b42e325)


### Whole exome sequencing in the diagnosis of liver diseases

**Why Not Relevant**: The provided paper content does not mention the gene FAM13A anywhere and provides no gene-level results or lists that include it. The text describes a study of exome sequencing in liver disease cohorts, overall diagnostic yields, the number of genes examined (502) and that P/LP variants were found in 25 genes among the CLD cohort, but it does not identify those genes or provide any data, variants, phenotypes, or mechanistic discussion specific to FAM13A. Therefore there is no direct evidence (genetic association, reported pathogenic variants, case reports) nor mechanistic evidence (functional data, pathway links) in the supplied content to support or refute the claim that FAM13A is likely associated with at least one Mendelian disease.

Limitations and caveats: (1) The provided excerpt may be incomplete — the full paper or its supplementary materials might contain the gene list or case-level data that include FAM13A. (2) This assessment is strictly based on the supplied text; absence of mention in this excerpt is not definitive proof that the full article contains no relevant information. (3) The study focused on Mendelian liver disease genes; if FAM13A is not typically implicated in primary or secondary liver Mendelian disorders, it would be unlikely to appear, but that inference requires external data not present here.

Suggested next steps: request the paper's full text and supplementary tables (especially the list of the 25 genes with P/LP variants), or query curated resources (OMIM, ClinVar, HGMD) and the literature for FAM13A-specific Mendelian disease reports.


[Read Paper](https://www.semanticscholar.org/paper/b07ed494c5c173a3f7be94d5f6d5accc5255636b)


### OA30 Genetic analysis of whole exome sequencing in a cohort of children with refractory JIA reveals genetic risk factors for rare juvenile diseases

**Why Not Relevant**: The paper does not mention FAM13A anywhere in the provided content. All identified/prioritized genes discussed (ADAR, ATP7B, MVK, HLA-DRB1) are different from FAM13A. The methods state that variants were annotated only if they appeared in certain gene lists (primary immunodeficiency PanelApp, arthritis-associated genes, or a paediatric monogenic gene list), so if FAM13A is not on those lists it would not have been highlighted by this study. The paper demonstrates the study could detect and report known monogenic-disease-associated variants in other genes, but provides no direct evidence (no detected variants, no mention, no mechanistic discussion) that FAM13A is associated with any Mendelian disease. Additional limitations relevant to absence of evidence here: (1) cohort size is modest (99 individuals), so rare pathogenic variants in FAM13A could be missed by chance; (2) the authors applied variant frequency and pathogenicity filters (e.g., removing variants with allele frequency >1%, ClinVar benign/likely benign, and CADD PHRED <15 — note the text also states a REVEL score >0.7 in the filtering clause, which may be a typographical or interpretational issue), which could exclude some potentially relevant variants depending on parameter choices; (3) annotation was restricted to certain gene lists, so genes not on those lists would not be reported even if variants existed; and (4) the provided excerpt does not include a complete variant table or negative results listing, so absence of FAM13A mention here is not a formal proof that FAM13A was examined and found negative. In sum, this paper provides no direct or mechanistic evidence that FAM13A is associated with a Mendelian disease, and thus is not relevant to supporting the claim.


[Read Paper](https://www.semanticscholar.org/paper/5ceb90bf884c2beab8d50226ab822bc8d21c2783)


## Search Queries Used

- FAM13A pathogenic variant Mendelian disease case report family segregation

- FAM13A ClinVar OMIM HGMD monogenic disorder pathogenic annotation gene disease association

- FAM13A whole exome sequencing rare disease cohort de novo variant monogenic diagnosis

- FAM13A functional study knockout mouse phenotype loss of function gain of function Rho GTPase signaling COPD genome wide association


## Bibliometric Configuration

- Use Bibliometrics: True

- Author Impact Weight: 0.4

- Citation Impact Weight: 0.4

- Venue Impact Weight: 0.2
