## Supplementary material for "MENDELSEEK: An algorithm that predicts Mendelian Genes and elucidates what makes them special": FCN2.txt

# Claim: The gene FCN2 is likely associated with at least one Mendelian disease.

**Status**: processed

**Overall Rating**: -1

**Explanation**:

No relevant papers were found that support or refute this claim.


## Relevant Papers


## Other Reviewed Papers


### Annotation of uORFs in the OMIM genes allows to reveal pathogenic variants in 5′UTRs

**Why Not Relevant**: The provided paper abstract describes a general annotation and experimental-validation study of upstream open reading frames (uORFs) in disease-associated genes and reports that variants disrupting uORFs (from HGMD and ClinVar) can lead to Mendelian disorders. However, the abstract does not mention the gene FCN2 anywhere, and the list of experimentally validated genes (ETFDH, PAX9, MAST1, HTT, TTN,GLI2, COL2A1, ZIC2) does not include FCN2. Because there is no direct statement, list entry, variant report, or experimental result in the provided content tying FCN2 to a Mendelian disease, the paper as presented does not provide direct evidence for the claim. The paper does provide generalized mechanistic support (that uORF-disrupting variants can underlie Mendelian disease and that such variants are enriched among pathogenic calls), which makes it plausible that some genes studied could include FCN2, but plausibility or methodological applicability is not equivalent to evidence that FCN2 specifically is associated with at least one Mendelian disease. To support the claim about FCN2, one would need an explicit mention of FCN2 in the paper's gene lists, variant tables, or experimental validations (none are present in the supplied text), or a direct curated entry in OMIM/HGMD/ClinVar documented in the paper linking FCN2 to a Mendelian phenotype.


[Read Paper](https://www.semanticscholar.org/paper/5d45c0cdf801505df45fc93a466a2a4101a12871)


### Whole-Exome Sequencing Uncovers Novel Causative Variants and Additional Findings in Three Patients Affected by Glycogen Storage Disease Type VI and Fanconi−Bickel Syndrome

**Why Not Relevant**: The supplied paper content does not mention the gene FCN2 anywhere, nor does it report variants, mechanistic data, or clinical associations involving FCN2. The study focuses on glycogen storage disease (GSD) genes (examples given: PYGL, SLC2A2) and reports findings from whole-exome sequencing in three individuals with suspected liver GSDs. Although the authors note identification of eight pathogenic/likely pathogenic Mendelian-disease findings, the provided text does not list those genes and contains no statement implicating FCN2. Therefore the paper provides no direct or mechanistic evidence supporting or refuting the claim that FCN2 is likely associated with at least one Mendelian disease. Additional caveats: absence of mention in this small, disease-focused WES study is not strong evidence against FCN2 involvement in Mendelian disease generally (the study design targeted GSDs and had only three individuals), but from the provided content there is simply no information relevant to the claim.


[Read Paper](https://www.semanticscholar.org/paper/5cee85618cba2f1ecdcf2d93bcd128fa435df210)


### Molecular Genetic Study of a Large Inbred Pakistani Family Affected with Autosomal Recessive Congenital Ichthyosis Through Whole Exome Sequencing.

**Why Not Relevant**: The provided paper content focuses exclusively on a novel nonsense mutation in PNPLA1 causing autosomal recessive congenital ichthyoses (ARCI). All specific genetic, mechanistic, and contextual statements refer to PNPLA1 (e.g., "Analysis of the WES data identified a novel nonsense mutation, c.762C>G, in the PNPLA1 (patatin-like phospholipase domain containing 1) gene." and "The protein product of of this gene is involved in lipid organization during cornified cell envelope formation."). There is no mention of FCN2, ficolin-2, or any gene or protein related to FCN2, nor any discussion of diseases or phenotypes associated with FCN2. Therefore the paper provides neither direct evidence (no reported FCN2 variants or segregation with Mendelian disease) nor mechanistic evidence (no functional data or pathways involving FCN2) to support or refute the claim that "FCN2 is likely associated with at least one Mendelian disease." Note: absence of evidence in this single paper does not disprove the claim overall; it only means this paper is not informative for evaluating the association of FCN2 with Mendelian disease.


[Read Paper](https://www.semanticscholar.org/paper/06a186a63b447a0e978a698a6445035344530fda)


### Determining the pathogenicity of CFTR missense variants: Multiple comparisons of in silico predictors and variant annotation databases

**Why Not Relevant**: This paper analyzes pathogenicity prediction for missense variants in the CFTR gene and the performance of in silico predictors for classifying CFTR variants. The abstract and content focus exclusively on CFTR (cystic fibrosis transmembrane conductance regulator) and cystic fibrosis / CFTR-related disorders; there is no mention of the gene FCN2, no data or analyses involving FCN2 variants, and no mechanistic discussion that would connect CFTR findings to FCN2. Therefore the paper provides neither direct evidence (no reports of FCN2 variants causing Mendelian disease) nor mechanistic evidence (no biology, pathways, or functional data concerning FCN2) relevant to the claim that “FCN2 is likely associated with at least one Mendelian disease.”

Methodological reasons this paper cannot inform the claim: the study design is a gene-specific evaluation of variant pathogenicity predictors applied to CFTR missense variants; its results pertain to the reliability of predictors for CFTR and to clinical decision-making for CFTR variant interpretation. Those conclusions do not generalize to other genes such as FCN2 because variant effect, constraint, domain structure, and disease mechanisms differ between genes. Consequently, no inference about FCN2’s disease association can be drawn from this manuscript.

If you want evidence for or against an association between FCN2 and a Mendelian disease, consult gene-disease resources (OMIM, ClinGen, ClinVar, HGMD), primary case reports or family-based studies in PubMed, population-constraint metrics (gnomAD pLI, observed/expected), and functional or segregation studies. A targeted literature and database search for “FCN2” plus terms like “mutation,” “variant,” “Mendelian,” “familial,” “hereditary,” or specific phenotypes (for example immunodeficiency, if relevant) is the appropriate next step.


[Read Paper](https://www.semanticscholar.org/paper/80ac05d37d1e9b8b9d7ab0f8bd0a2fea507edd9c)


### Incomplete penetrance and variable expressivity in monogenic diabetes; a challenge but also an opportunity

**Why Not Relevant**: The single sentence provided from the paper—"This review will mainly address penetrance and expressivity variation in different types of MFD, factors that confer such variations and opportunities that come with such knowledge."—does not mention FCN2, any specific gene, any Mendelian disease linked to FCN2, or any mechanistic details related to FCN2. Therefore the excerpt contains no direct evidence (e.g., genetic association, case reports, segregation data) nor mechanistic evidence (e.g., biochemical function, pathogenic variants, animal models) that would support or refute the claim that FCN2 is likely associated with at least one Mendelian disease.

Additional limiting factors: only a single sentence from the review was provided, so it is possible that FCN2 is discussed elsewhere in the full paper; however, based on the supplied content alone there is no relevant information to evaluate the claim. To reach a verdict on the claim using this paper, the full text (or at least the sections discussing specific genes or disease examples) would be required. Alternatively, other sources (OMIM, ClinVar, PubMed articles describing FCN2 variants and clinical phenotypes) should be consulted.


[Read Paper](https://www.semanticscholar.org/paper/79fbedd297d3b7288b734021b4187afec402f5a3)


### [A rare variant in the TTR gene (p.E112K) is associated with systemic amyloidosis and a new symptom - skin hyperemia in response to ethanol intake: family segregation analysis, literature review, and a clinical case. Case report].

**Why Not Relevant**: The provided paper content discusses transthyretin amyloidosis (ATTR-amyloidosis) and variants of the TTR gene; it makes no mention of FCN2, ficolin-2, or any genes other than TTR. There is no direct evidence (case reports, genetic association, segregation, pathogenic variants) concerning FCN2, nor is there any mechanistic discussion (protein function, pathway involvement, loss/gain-of-function mechanisms) that could plausibly link FCN2 to a Mendelian disease. Therefore, based on the supplied text, this paper provides no support for or against the claim that FCN2 is likely associated with at least one Mendelian disease. Limitation: this assessment is strictly limited to the excerpt you provided; if the full paper contains other sections not included here that mention FCN2, those were not available for review and could change the relevance.


[Read Paper](https://www.semanticscholar.org/paper/2567df8ee078bbbd995e907e0465d6337ffeeb26)


### A case report of PGAP2-related hyperphosphatasia with impaired intellectual development syndrome in a Chinese family and literature review

**Why Not Relevant**: This paper does not provide evidence for or against the claim that FCN2 is likely associated with any Mendelian disease. The content and quoted sentences show the study focuses exclusively on PGAP2 and other GPI-anchor-synthesis genes (PIGA, PIGY, PIGO, PIGW, PGAP3) in relation to hyperphosphatasia with impaired intellectual development syndrome (HPMRS). Nowhere in the provided text is FCN2 (ficolin-2) mentioned, nor are there any genetic, clinical, mechanistic, or population-data links between FCN2 and the disorders discussed. Therefore the paper contains no direct evidence (no FCN2 variants, no segregation, no case reports) and no mechanistic evidence (no biochemical pathway involving FCN2 relevant to HPMRS) supporting the claim.

Limitations for assessing FCN2 from this paper: because the paper neither studies nor discusses FCN2, nothing in its methods, results, or conclusions can be extrapolated to FCN2 without additional data. To support the claim that FCN2 is associated with a Mendelian disease would require evidence such as: segregation of rare, likely damaging FCN2 variants with disease in families; recurrence of similar FCN2 variants in unrelated affected individuals; functional assays showing variant impact on ficolin-2 function; constraint/allele-frequency data consistent with pathogenicity; and/or an entry in curated resources (e.g., OMIM) linking FCN2 to a Mendelian phenotype. None of these appear in the provided paper.

Recommendation: to evaluate the FCN2–Mendelian disease claim, examine studies and databases that specifically report FCN2 variants, functional effects on ficolin-2 protein, familial segregation, case series, and clinical annotations (ClinVar, OMIM, gnomAD constraint metrics, PubMed reports). This paper on PGAP2/HPMRS does not contribute evidence for FCN2.


[Read Paper](https://www.semanticscholar.org/paper/63cba8ad2bff7348418f63782b7cc408f084157d)


### Two novel mutations in ALDH18A1 and SPG11 genes found by whole-exome sequencing in spastic paraplegia disease patients in Iran

**Why Not Relevant**: Relevance is effectively zero because the paper's entire content focuses on hereditary spastic paraplegia and reports likely pathogenic variants in ALDH18A1 and SPG11 in two cases. The manuscript contains no mention of the gene FCN2, no FCN2 variants, no phenotypes attributed to FCN2, and no mechanistic discussion involving FCN2. Although the methods state that whole-exome sequencing was performed (which in principle would capture FCN2 coding sequence), the authors report only variants in ALDH18A1 and SPG11 — the absence of any FCN2 findings in the reported results means this paper provides no direct or mechanistic evidence that FCN2 is associated with any Mendelian disease. Limitations: lack of mention is not definitive proof that FCN2 is not associated with disease (the study is small, focused on spastic paraplegia phenotypes, and the authors may have filtered or prioritized variants related to the presenting phenotype), but for the purpose of supporting the claim about FCN2, this paper supplies no relevant evidence.


[Read Paper](https://www.semanticscholar.org/paper/75a201bc47209dabc2b08a6a6888f29571c37a97)


### Autosomal Dominant Intellectual Development Disorder-6 (MRD6) Without Seizures Linked to a De Novo Mutation in the grin2b Gene Revealed by Exome Sequencing: A Case Report of a Moroccan Child

**Why Not Relevant**: The provided paper content exclusively concerns a pathogenic variant in the grin2b (GRIN2B) gene causing Autosomal dominant intellectual development disorder-6 (MRD6). It describes a de novo nonsense variant c.3912C>G (p.Tyr1304Ter) in GRIN2B, ACMG classification of that variant as pathogenic, and discusses GRIN2B's role in brain development and neurodevelopmental disease. Nowhere in the provided text is the gene FCN2 (or ficolin-2) mentioned, nor are any data, variants, mechanisms, or clinical associations described that involve FCN2. Therefore the paper provides no direct genetic, clinical, or mechanistic evidence bearing on the claim that FCN2 is associated with a Mendelian disease.

Specific reasons this content is not relevant to the claim:
- Gene mismatch: the entire manuscript excerpt deals with GRIN2B (grin2b) rather than FCN2; association evidence presented (de novo pathogenic variant, segregation, ACMG criteria) applies to GRIN2B only.
- No mechanistic connection to FCN2: the biological mechanisms discussed (neurodevelopmental effects, GRIN2B's role in nervous system communication) are specific to GRIN2B and cannot be used to infer FCN2 involvement because FCN2 encodes a lectin (ficolin-2) with distinct immune-related functions.
- No transferable variant or population data: the variant, its absence from gnomAD, and pathogenicity evidence are specific to the reported GRIN2B variant; there are no FCN2 variants, frequencies, segregation data, or functional assays reported.

Limitations on attempting to draw any inference: one cannot generalize from a well-supported gene-disease association for GRIN2B to another unrelated gene (FCN2) without direct data. Absent any mention of FCN2, the paper neither supports nor refutes the claim about FCN2 and therefore provides no usable evidence for the claim that FCN2 is likely associated with at least one Mendelian disease.


[Read Paper](https://www.semanticscholar.org/paper/67d756e0cc302e5d46fdda9df34403f1f2de9a5a)


### Camptodactyly-Arthropathy-Coxa Vara-Pericarditis Syndrome Caused by Truncating Mutations in the Prg4 Gene: Case Series and Literature Review

**Why Not Relevant**: The paper exclusively studies Camptodactyly Arthropathy Coxa-vara Pericarditis (CACP) syndrome and provides genetic, mechanistic, and clinical evidence implicating truncating mutations in the PRG4 gene as causal. The text contains no mention of the FCN2 gene, no data on FCN2 variants, and no mechanistic discussion involving FCN2. Therefore this paper provides no direct or mechanistic evidence for or against the claim that FCN2 is likely associated with at least one Mendelian disease. Absence of evidence in this paper does not prove FCN2 is not disease-associated, but it means this paper is irrelevant to evaluating the FCN2 claim.


[Read Paper](https://www.semanticscholar.org/paper/e87a78ef310a6c55e5787a7bb3c0ed0b048657a8)


### The application of whole-exome sequencing in the early diagnosis of rare genetic diseases in children: a study from Southeastern China

**Why Not Relevant**: This paper does not provide any direct or mechanistic evidence about the gene FCN2. The provided content summarizes a cohort whole-exome sequencing (WES) study (144 patients, 61 diagnosed) and reports aggregate diagnostic rates, phenotype groups, inheritance patterns (e.g., "Autosomal dominant disorders accounted for 70.49% (43/61) of positive cases" and "80.56% (29/36) of pathogenic variants were identified as de novo mutations through Trio-WES"), but it does not mention FCN2, ficolin-2, any variants in FCN2, nor any gene-level or variant-level results that would support or refute an association between FCN2 and a Mendelian disease. There are no mechanistic descriptions (e.g., functional assays, protein function perturbation, pathway involvement) pertaining to FCN2, no segregation data for FCN2 variants, and no case reports in which FCN2 variants are implicated. Therefore the paper provides neither supporting nor contradicting evidence regarding the claim that FCN2 is likely associated with at least one Mendelian disease.

Limitations and context: (1) The study design (WES of a diagnostic cohort) is appropriate for discovering gene-disease associations, but because this paper does not report gene-specific findings or list implicated genes, we cannot infer anything about FCN2 from the presented summary. (2) Absence of mention in this report is not evidence that FCN2 is not associated with Mendelian disease; it only means this paper does not address that gene. (3) To evaluate the claim further, one should examine variant-level databases and primary literature (e.g., ClinVar, OMIM, HGMD, PubMed) or look for case reports, segregation studies, functional assays, and population-frequency data specific to FCN2.


[Read Paper](https://www.semanticscholar.org/paper/ba9aa0637319a74b3dfdae65099a70edd6c60ebf)


### Pathogenic paralogous variants can be used to apply the ACMG PS1 and PM5 variant interpretation criteria

**Why Not Relevant**: The provided paper content describes a general methodological advance (para-PS1/PM5) for using pathogenic paralogous variants to extend ACMG PS1/PM5 criteria across many genes. It contains no mention of the gene FCN2, no gene-specific results, and no variant-level evidence linking FCN2 to any Mendelian phenotype. While the method could in principle be applied to FCN2 to help classify missense variants or to aggregate paralogous evidence, the paper as provided does not present any direct genetic, clinical, segregation, functional, or statistical data pertaining to FCN2. Therefore it does not provide direct or strong mechanistic evidence that FCN2 is likely associated with at least one Mendelian disease. Given the absence of any FCN2-specific content, the relevance to the specific claim is very low.


[Read Paper](https://www.semanticscholar.org/paper/0a921a972e8db185945c5510191b7ac52b42e325)


## Search Queries Used

- FCN2 ficolin 2 mutations pathogenic variants Mendelian disease case report family segregation

- FCN2 ClinVar OMIM pathogenic variant monogenic clinical annotation

- FCN2 ficolin 2 loss of function lectin complement pathway primary immunodeficiency recurrent infections congenital deficiency

- FCN2 rare variants whole exome sequencing trio rare disease autosomal recessive autosomal dominant segregation analysis


## Bibliometric Configuration

- Use Bibliometrics: True

- Author Impact Weight: 0.4

- Citation Impact Weight: 0.4

- Venue Impact Weight: 0.2
