## Supplementary material for "MENDELSEEK: An algorithm that predicts Mendelian Genes and elucidates what makes them special": FGFR1OP2.txt

# Claim: The gene FGFR1OP2 is likely associated with at least one Mendelian disease.

**Status**: processed

**Overall Rating**: -1

**Explanation**:

No relevant papers were found that support or refute this claim.


## Relevant Papers


## Other Reviewed Papers


### Developmental mechanisms of heart disease

**Why Not Relevant**: The supplied document is a table-of-contents style list of sections and chapter titles from a volume on cardiac development and clinical molecular cardiology. It contains no mention of the gene symbol "FGFR1OP2" (or obvious synonyms) and includes no experimental methods, results, case reports, variant descriptions, statistical associations, or mechanistic descriptions that are specific to FGFR1OP2. Therefore it provides neither direct evidence (e.g., genetic linkage, segregation, pathogenic variants, case series) nor mechanistic evidence (e.g., protein function, pathway involvement, animal models) that would support or refute the claim that FGFR1OP2 is likely associated with a Mendelian disease.

Specific reasons for assigning zero relevance:
- No direct mentions: The provided text contains only chapter and section titles (e.g., "Regulation of Early Heart Development by the Fibroblast Growth Factor-2 System", "Molecular Analysis of the inv Insertional Mutation", "Genetic Analysis of Hypertrophic Cardiomyopathy in Japan") but none of these headings mention FGFR1OP2, nor any variant identifiers, patient summaries, or gene-disease statements pertaining to FGFR1OP2.
- No mechanistic detail for FGFR1OP2: Although some chapter titles refer broadly to growth factor systems or molecular mechanisms in cardiac development, there are no excerpts describing FGFR1OP2 protein function, expression, interaction partners, loss- or gain-of-function phenotypes, or model-organism data for this specific gene.
- No methodological or statistical information: The content provided has no methods, sample sizes, segregation analyses, LOD scores, odds ratios, p-values, or other data that could be interpreted as evidence for a Mendelian association.
- Document type limitation: The supplied content appears to be a high-level index/TOC rather than chapter text. It is possible that full chapter texts (not provided) could discuss specific genes, but based solely on the material given there is nothing to extract or quote that bears on the FGFR1OP2–Mendelian disease claim.

Implications and recommended next steps:
- Because this document contains no relevant evidence, you should consult primary genetic resources for claims about FGFR1OP2: OMIM (Online Mendelian Inheritance in Man), ClinVar, HGMD (if available), gnomAD (for constraint metrics), PubMed (for gene-specific literature), and locus-specific databases. Search for gene synonyms (if any) and for reported pathogenic or likely pathogenic variants, segregation with disease, or animal-model phenotypes.
- If you can provide full-text chapters or specific excerpts from the volume that mention FGFR1OP2, I can re-analyze those for direct or mechanistic evidence. As provided, there is no text to extract or evaluate with respect to the claim.


[Read Paper](https://www.semanticscholar.org/paper/65e202db3d23610a48f150372b008c899147385f)


### Novel variant of KIF11 associated with MCLMR syndrome

**Why Not Relevant**: The paper exclusively investigates and reports a heterozygous, likely pathogenic, de novo frameshift variant in KIF11 as the cause of the patient's Mendelian syndrome. It details targeted sequencing of the 22 coding exons and exon–intron boundaries of KIF11 and provides mechanistic commentary (loss-of-function via nonsense-mediated decay/protein truncation) for the KIF11 variant. Nowhere in the provided text is FGFR1OP2 mentioned, nor are exome- or genome-wide analyses reported that could have identified FGFR1OP2 variants. Because the genetic testing was targeted to KIF11, the study cannot provide direct evidence for or against an association between FGFR1OP2 and any Mendelian disease. Additionally, the report is a single-case study without functional follow-up for the variant; even for KIF11 the evidence is limited to genetics and inference rather than functional validation. Therefore, this paper does not provide direct or mechanistic evidence relevant to the claim that FGFR1OP2 is likely associated with at least one Mendelian disease.


[Read Paper](https://www.semanticscholar.org/paper/598b1c7508865997c295d53a98d38407fbecde29)


### Neural expression and chromosomal mapping of Neu differentiation factor to 8 pl 2p 21 ( polypeptide growth factor / receptor tyouine kinase / erbB2 / nervous system )

**Why Not Relevant**: This paper describes the Neu differentiation factor (NDF/heregulin): its biochemical purification, cloning, embryonic expression pattern in mouse, and chromosomal localization of the NDF gene to human chromosome 8 (short arm, bands p12-p21). The text makes no mention of the gene FGFR1OP2 (no occurrences of that gene symbol or any obvious synonyms), provides no clinical case reports, no mutation data, no linkage/segregation analyses, and no population or association studies that would address Mendelian disease association. The experimental focus (in situ hybridization of NDF mRNA in mouse embryos, cloning of NDF cDNA, and hybridization of an NDF probe to human metaphase spreads) is on gene expression and chromosomal mapping of NDF, not on identifying or characterizing Mendelian disorders or genes other than NDF/heregulin. Mechanistically, the paper discusses receptor tyrosine kinases (Neu/ErbB-2 and EGF receptor) and NDF as a ligand, but that mechanistic content pertains to NDF/heregulin biology and is not linked to FGFR1OP2. Because the provided content does not mention FGFR1OP2 and contains no data that could be interpreted as evidence for or against FGFR1OP2 being associated with any Mendelian disease, it cannot be used to support or refute the claim. Limitations: although growth-factor signaling mechanisms described here are broadly relevant to many genes, without any explicit mention, mapping, functional assays, mutation screening, or clinical phenotypes related to FGFR1OP2, there is no direct or plausible mechanistic bridge in this paper to conclude anything about FGFR1OP2 and Mendelian disease.


[Read Paper](https://www.semanticscholar.org/paper/b26c856a1e72ac2ebf20c867aa0184d4a3882a55)


### Kiyokazu Kakugawa DYNAMIC TRANSLOCATION OF THEMIS FROM THE CYTOPLASM TO THE NUCLEUS IS ESSENTIAL FOR ITS FUNCTION IN T CELLS

**Why Not Relevant**: The supplied document is a collection of conference abstracts on immunology topics (Treg cells, THEMIS, LAT, Th17/exTh17, BAF complex and CD8+ T cells, etc.). There is no mention of the gene name FGFR1OP2 (or any obvious synonyms), no human genetic or Mendelian-disease phenotypes, no segregation or case reports, no population or association analyses, and no functional genetic experiments tying FGFR1OP2 to disease. Therefore the content provides neither direct evidence (e.g., co-segregation, pathogenic variants, case descriptions, statistical association) nor mechanistic evidence (e.g., loss-of-function experiments demonstrating a disease-relevant phenotype) that FGFR1OP2 is associated with a Mendelian disease. Limitations/caveats: (1) the provided excerpt may be only a subset of the full proceedings; if other sections of the proceedings or separate publications mention FGFR1OP2 they were not included here; (2) absence of evidence in this document is not evidence of absence overall — establishing the claim requires targeted genetic studies, clinical reports, or database searches (OMIM, ClinVar, HGMD, gnomAD constraint metrics). For evaluation of the claim, I recommend searching primary genetic-disease resources and the literature for FGFR1OP2/aliases, looking for family segregation studies, reported pathogenic variants, and experimental loss/gain-of-function evidence.


[Read Paper](https://www.semanticscholar.org/paper/876ac5e0bd7d88047c7f293c82c685335472a133)


### Discovery of a β-cell check on insulin action

**Why Not Relevant**: The provided paper content does not mention the gene FGFR1OP2 at all; it only discusses a gene labeled "Iir." The sentence reports functional (knockdown) and mechanistic findings for Iir (effects on β cell mass, glucose homeostasis, and interactions with INSR/IGF1R). To support the claim that FGFR1OP2 is likely associated with at least one Mendelian disease, one would expect direct human genetic evidence (pathogenic variants in FGFR1OP2 segregating with disease, recurrent de novo mutations, or strong genotype–phenotype correlations), or at least explicit functional data linking FGFR1OP2 perturbation to a Mendelian phenotype. None of that is present in the excerpt. Possible caveat: if "Iir" is an alternative name or alias for FGFR1OP2, the excerpt could be relevant, but the text does not state this, so we cannot assume equivalence. Additionally, the excerpt reports knockdown experiments (likely in model systems), which, even if on FGFR1OP2, would be mechanistic/functional evidence but would not by itself establish a Mendelian disease association without human genetic/clinical corroboration. For these reasons the content is essentially non‑relevant to the claim.


[Read Paper](https://www.semanticscholar.org/paper/29c05f83cf169d484bcc7e625f2bbffc6d0b8029)


### INVESTIGATION OF PEPTIDE LIGAND/RECEPTOR INTERACTIONS IN YEAST USING ADVANCEMENTS ON THE GROWTH FACTOR REGULATION OF STEROID RECEPTOR TRANSCRIPTIONAL ACTIVITY.

**Why Not Relevant**: The supplied paper content does not mention FGFR1OP2, nor does it present any human genetic, clinical, linkage, segregation, variant, or phenotype data that could support or refute an association between FGFR1OP2 and a Mendelian disease. The excerpts are methodological and biological abstracts (yeast two-hybrid, microscopy, prolactin biology, steroid receptor function) focused on protein interactions, imaging, and endocrine signaling—none provide direct evidence (e.g., pathogenic variants, segregation in families, case reports) or specific mechanistic pathways connecting FGFR1OP2 to Mendelian pathogenesis. Because there is no gene-specific content, no human genetic data, and no disease phenotype descriptions relating to FGFR1OP2, the document is irrelevant to the claim.


[Read Paper](https://www.semanticscholar.org/paper/e837a38d6f557dd7829c41f8a1b43cab771ff7ea)


## Search Queries Used

- FGFR1OP2 Mendelian disease monogenic inherited syndrome FGFR1 oncogene partner 2 FOP2 gene

- FGFR1OP2 pathogenic variant ClinVar OMIM case report exome sequencing family segregation de novo loss of function missense truncating frameshift

- Fibroblast growth factor receptor 1 oncogene partner 2 protein function expression developmental phenotype knockout mouse zebrafish cell biology localization functional study

- exome sequencing rare disease cohort gene discovery FGFR1OP2 gene burden rare variant association de novo mutation gnomAD constraint pLI loss of function intolerance


## Bibliometric Configuration

- Use Bibliometrics: True

- Author Impact Weight: 0.4

- Citation Impact Weight: 0.4

- Venue Impact Weight: 0.2
