## Supplementary material for "MENDELSEEK: An algorithm that predicts Mendelian Genes and elucidates what makes them special": FN3K.txt

# Claim: The gene FN3K is likely associated with at least one Mendelian disease.

**Status**: processed

**Overall Rating**: -1

**Explanation**:

No relevant papers were found that support or refute this claim.


## Relevant Papers


## Other Reviewed Papers


### High diagnostic rate of trio exome sequencing in consanguineous families with neurogenetic diseases

**Why Not Relevant**: The provided paper content is limited to the abstract and does not mention the gene FN3K by name, does not present variant-level data tied to specific gene symbols beyond aggregate counts, and does not provide mechanistic data that implicates FN3K specifically. Therefore the paper content as provided contains no direct evidence that FN3K is associated with a Mendelian disease. It is possible that the full article or its supplementary tables list the 119 known disease genes or the 27 novel genes and might include FN3K, but that information is not present in the excerpt given. To properly assess the claim one would need the full text and supplementary materials (gene/variant lists, tables, and any case descriptions) to see whether FN3K appears among implicated genes and to evaluate the evidence for pathogenicity (variant type, segregation, frequency, functional data). Given the absence of any mention of FN3K in the supplied content, the relevance of this paper to the specific claim is very low.


[Read Paper](https://www.semanticscholar.org/paper/4563b1f6db311c786c75738c544f25cd3e75932f)


### Implementation of Exome Sequencing in Prenatal Diagnostics: Chances and Challenges

**Why Not Relevant**: The provided paper text does not mention the gene FN3K, contains no data, variant calls, gene lists, or mechanistic discussion related to FN3K, and therefore provides no direct or mechanistic evidence for or against the claim that FN3K is associated with a Mendelian disease. The extract is a brief report of a one-year prenatal WES diagnostic series (28 fetus-parent trios) with overall diagnostic yield and operational details (turn-around time, inheritance modes of detected pathogenic variants). While it states that 7 cases (25%) had pathogenic or likely pathogenic variants with various inheritance patterns (autosomal recessive, de novo, dominant), it does not identify any of the causal genes or phenotypes, so one cannot infer FN3K involvement from this report. Important limitations preventing relevance: (1) absence of any mention of FN3K or specific gene-level results; (2) small cohort focused on diagnostic yield rather than gene discovery; (3) no mechanistic or functional data presented; (4) absence of variant details, segregation, or phenotype description tied to particular genes. Absence of evidence for FN3K in this paper is not evidence that FN3K is or is not associated with Mendelian disease. To evaluate the claim one would need sources that report FN3K variants in individuals with consistent, segregating phenotypes and/or functional work implicating FN3K in disease (e.g., ClinVar/OMIM entries, case series, functional assays, or animal models).


[Read Paper](https://www.semanticscholar.org/paper/f0b9c404eead58932c3e10e563291e1b0ae2f5a5)


### Identification of a novel de novo pathogenic variant in GFAP in an Iranian family with Alexander disease by whole-exome sequencing

**Why Not Relevant**: The provided paper content does not mention the gene FN3K, any variants in FN3K, nor any clinical or molecular data tying FN3K to Alexander disease or any other Mendelian disorder. The single sentence only states a general aim to expand the genetic spectrum of Iranian AxD patients; it contains no gene-specific findings, no mechanistic discussion, no statistical results, and no diagnostic evidence. Because association of FN3K with a Mendelian disease would require explicit gene-level evidence (e.g., identification of pathogenic variants, segregation data, functional assays, or mechanistic pathways), this excerpt is insufficient and irrelevant to assess the claim. If more of the paper (full text, methods, results, variant lists, or tables) were available, a re-evaluation could be performed; based on the current content, relevance to the FN3K claim is effectively nil.


[Read Paper](https://www.semanticscholar.org/paper/9b228b684c3371c832698bc6ddcbd35325a8af79)


### Chinese family with Blau syndrome: Mutated NOD2 allele transmitted from the father with de novo somatic and germ line mosaicism

**Why Not Relevant**: The paper exclusively reports on Blau syndrome and genetic findings related to the NOD2 gene (including a specific pathogenic variant, mosaicism across tissues, and diagnostic sequencing considerations). There is no mention of the gene FN3K, no sequence/variant data for FN3K, no genetic segregation or functional studies involving FN3K, and no mechanistic discussion that would link FN3K to any Mendelian phenotype. Because the document contains no information about FN3K, it provides neither supporting nor refuting evidence for the claim that "FN3K is likely associated with at least one Mendelian disease." Absence of discussion of FN3K in this paper means it cannot be used as evidence for that claim; to evaluate the claim one would need papers reporting FN3K variants segregating with disease, functional assays showing pathogenicity, or genetic/clinical case reports implicating FN3K in Mendelian inheritance.


[Read Paper](https://www.semanticscholar.org/paper/86abd598b42cc68307bbee0d238cdc840dab1ed3)


### A novel CHD7 variant in a chinese family with CHARGE syndrome

**Why Not Relevant**: The provided paper content contains only a single summary sentence about genetic testing facilitating prenatal diagnosis of CHARGE syndrome and promoting appropriate genetic counseling. It does not mention the gene FN3K, any specific genes or variants, segregation data, functional studies, case reports, population associations, or mechanistic pathways relevant to FN3K. Because none of the required types of evidence for associating a gene with a Mendelian disease (e.g., identification of pathogenic variants in affected individuals, co-segregation in families, functional validation, statistical enrichment, or described disease mechanism) are present in the provided text, the paper content is not relevant to the claim. Additional reasons for assigning relevance = 0.0: (1) absence of gene name FN3K or synonyms, (2) no variant, phenotype, or inheritance information tied to FN3K, (3) no experimental or clinical data to evaluate, and (4) the single sentence pertains to CHARGE syndrome broadly without linking it to FN3K. If you can provide the full paper text or other papers that mention FN3K, I can re-evaluate for direct and mechanistic evidence.


[Read Paper](https://www.semanticscholar.org/paper/53b970f2aca721c79ce44f35acdd1636c979ac75)


### Whole-exome sequencing of pathogenic genes in a family with congenital heart disease: A case report

**Why Not Relevant**: The paper focuses on identifying causative variants for congenital heart disease in a single Chinese family using whole-exome sequencing and reports variants in AMER1 and KCNE1 (and two unspecified variants). The gene FN3K is not mentioned anywhere in the provided content, and no results, mechanistic discussion, or variant reports involving FN3K are present. Therefore this paper provides no direct evidence supporting or refuting the claim that FN3K is likely associated with at least one Mendelian disease. Note: because whole-exome sequencing was performed, if FN3K had been implicated in this family the authors would likely have reported it; however, the absence of FN3K in this small, disease-specific study is not definitive evidence that FN3K has no Mendelian associations in other contexts. To evaluate the claim about FN3K, sources that specifically examine FN3K variants (e.g., disease gene catalogs like OMIM, ClinVar, gene-specific association studies, or larger sequencing cohorts) would be required.


[Read Paper](https://www.semanticscholar.org/paper/2afe5ddc8ce353c6339a8bb041306ea8d115518f)


### Germline mosaicism in a collagen VI‐related myopathy family: A cause of autosomal recessive inheritance

**Why Not Relevant**: This paper describes paternal germline mosaicism of pathogenic variants in the COL6A1 gene causing collagen VI-related myopathy (Ullrich congenital muscular dystrophy). The content repeatedly names COL6A1 and related collagen genes (COL6A1, COL6A2, COL6A3) and details specific COL6A1 sequence variants, inheritance, and clinical consequences. Nowhere in the provided text is the gene FN3K mentioned, nor are any data, variants, functional assays, patient phenotypes, or mechanistic discussions concerning FN3K presented. Therefore the paper provides no direct genetic, clinical, or mechanistic evidence for or against the claim that FN3K is likely associated with a Mendelian disease. At best, the paper discusses the general concept of germline mosaicism as a mechanism that can affect recurrence risk in Mendelian disorders; this is a generic concept that could apply broadly to many genes, but it is not specific evidence linking FN3K to any Mendelian phenotype. Because the paper's experimental subjects, variants, and conclusions are all about COL6A1, its relevance to the FN3K claim is effectively zero.


[Read Paper](https://www.semanticscholar.org/paper/4572f635bb354974678a75579201d9eedc2c40ea)


### Exome Sequencing Identifies a Novel SGCA Gene Mutation in an Iranian Family with Limb Girdle Muscular Dystrophy: A Case Report

**Why Not Relevant**: The provided paper content does not mention the gene FN3K anywhere and instead reports a novel homozygous SGCA mutation (p.A107fs:c.319-329del) causing autosomal recessive limb-girdle muscular dystrophy (LGMD) in a single case from an Iranian consanguineous family. The study uses whole exome sequencing followed by Sanger sequencing to identify and confirm the SGCA variant and concludes that the SGCA mutation is pathogenic for LGMD. Because there is no mention of FN3K, no genetic, clinical, or mechanistic data pertaining to FN3K are present in the text, so the paper provides neither direct evidence (e.g., FN3K variants segregating with disease) nor mechanistic evidence (e.g., functional studies, pathway links) to support or refute the claim that FN3K is associated with a Mendelian disease. Limitations/caveats: absence of evidence in this specific paper does not imply FN3K is not associated with any Mendelian disease overall — it only means this paper is irrelevant to that claim. To evaluate the claim about FN3K would require papers that explicitly study FN3K variants, segregation, functional impact, or disease cohorts involving FN3K.


[Read Paper](https://www.semanticscholar.org/paper/46a6274976aa6da24830f44d465c15762c6beb3f)


### TARGETED SEQUENCING REVEAL TWO PATHOGENIC MUTATIONS IN LDLR AND APOE GENES IN PATIENT WITH FAMILY HYPERCHOLESTEROLEMIA: CASE REPORT

**Why Not Relevant**: The provided paper focuses on familial hypercholesterolemia and reports likely pathogenic variants in LDLR and APOE identified by a targeted cardiology gene panel. The text contains no mention of FN3K, no data on FN3K variants, and no mechanistic discussion involving FN3K. Therefore the paper offers no direct evidence that FN3K is associated with any Mendelian disease. The methodological details (targeted panel of cardiovascular genes, and the authors' suggestion that whole exome sequencing could identify other genes) mean that FN3K could simply have been outside the study's scope; absence of FN3K in this report is not evidence of absence in the broader literature. In sum, relevance to the claim is very low because the study neither tests nor discusses FN3K; any assertion about FN3K's association with Mendelian disease cannot be supported or refuted from this paper's content.


[Read Paper](https://www.semanticscholar.org/paper/04808c76b3962e7f9dc6e3ebec4ddbb8d15d0607)


## Search Queries Used

- FN3K pathogenic variant Mendelian monogenic disease exome sequencing case report family study

- FN3K ClinVar OMIM Orphanet DECIPHER GeneReviews FN3K pathogenic annotation gene disease

- fructosamine 3 kinase FN3K knockout mouse phenotype loss of function deglycation glycated proteins diabetic complications congenital phenotype

- FN3K rare variant exome sequencing trio sequencing burden test family linkage autosomal recessive autosomal dominant de novo


## Bibliometric Configuration

- Use Bibliometrics: True

- Author Impact Weight: 0.4

- Citation Impact Weight: 0.4

- Venue Impact Weight: 0.2
