## Supplementary material for "MENDELSEEK: An algorithm that predicts Mendelian Genes and elucidates what makes them special": GART.txt

# Claim: The gene GART is likely associated with at least one Mendelian disease.

**Status**: processed

**Overall Rating**: -1

**Explanation**:

No relevant papers were found that support or refute this claim.


## Relevant Papers


## Other Reviewed Papers


### Frequency of Variants in Mendelian Alzheimer's Disease Genes within the Alzheimer's Disease Sequencing Project (ADSP)

**Why Not Relevant**: The provided paper content focuses exclusively on three Mendelian Alzheimer's disease genes (PSEN2, PSEN1, and APP) and reports analyses of variants only in those genes within the ADSP WGS/WES datasets. There is no mention of the gene GART, no data presented on GART variants, and no mechanistic discussion involving GART. Therefore the paper provides no direct evidence for or against the claim that GART is likely associated with at least one Mendelian disease. Note that absence of GART from this paper does not imply that GART is not associated with Mendelian disease—only that this study did not evaluate it. To assess the claim properly would require literature or database searches (ClinVar, OMIM, HGMD), gene-specific case reports, or broader exome/genome-wide analyses that include GART.


[Read Paper](https://www.semanticscholar.org/paper/3760efb72c8ec6adaf553ac773087fd2b16aeafc)


### Case Report: Identification of a rare nonsense mutation in the POC1A gene by NGS in a diabetes mellitus patient

**Why Not Relevant**: The paper exclusively investigates and reports a pathogenic truncating mutation in POC1A as the cause of the patient's diabetes. The text contains no mention of the gene GART, no reported variants in GART, and no mechanistic discussion linking POC1A findings to GART function. Although trio whole-exome sequencing with high coverage and a comprehensive variant interpretation pipeline was used (so GART coding variants could potentially have been detected), the manuscript does not report negative or positive findings for GART specifically. Therefore the paper provides no direct evidence that GART is associated with any Mendelian disease, nor does it provide mechanistic evidence that would increase the plausibility of such an association. Given the complete absence of GART-related content, the relevance of this paper to the claim that "The gene GART is likely associated with at least one Mendelian disease" is effectively zero.


[Read Paper](https://www.semanticscholar.org/paper/8f90a0a9b3c51d0406ae0337e2a4d8c2c0204107)


### Whole‐exome sequencing for genetic diagnosis of idiopathic liver injury in children

**Why Not Relevant**: The paper does not mention the gene GART anywhere in the provided content. The study focused on pediatric liver disease and performed WES with filtering against a manually curated panel of 380 OMIM hepatobiliary genes; the genes reported (ACOX2, PYGL, ABCB4, PHKA2, CDAN1, JAG1, PCK2, SLC27A5, VPS33B) do not include GART. Therefore this paper provides no direct evidence supporting or refuting the claim that GART is associated with at least one Mendelian disease. Additional important caveats: (1) absence of GART in the reported results is not evidence that GART is not disease-associated — it may not have been on the curated panel, may not have been mutated in this small cohort, or variants may have been filtered out; (2) the cohort is small (20 patients) and ascertained for liver phenotypes, so it would only detect GART associations relevant to those phenotypes and present in this sample. To evaluate the claim about GART, one would need studies that specifically report pathogenic variants in GART with segregation/phenotype data, OMIM entries implicating GART, or functional/mechanistic studies linking GART loss or alteration to Mendelian phenotypes.


[Read Paper](https://www.semanticscholar.org/paper/bc3354e33080c6dc662b37cec78639ac41c8e1b9)


### Whole exome sequencing identifies variants in the \(\textit{TNNI3}\) gene in vietnamese patient with restrictive cardiomyopathy - a case report

**Why Not Relevant**: The paper content provided focuses entirely on restrictive cardiomyopathy (RCM) and reports pathogenic variants in the TNNI3 gene as the cause of the patient's disease. There is no mention of the gene GART anywhere in the provided text, no reported variants in GART, and no discussion of mechanisms or pathways that would link GART to RCM or to any Mendelian disorder. The explicit identification of TNNI3 variants (including genomic changes, amino-acid changes, inheritance, and ClinVar accession numbers) shows the paper attributes causation to TNNI3, not GART. Because the paper neither names GART nor provides experimental, genetic, clinical, or mechanistic data about it, it provides no direct evidence supporting the claim that GART is associated with at least one Mendelian disease. It also provides no mechanistic evidence that would make the claim plausible (e.g., pathway connections, functional studies, or population/segregation data for GART). Absence of evidence in this single paper is not evidence of absence for GART generally, but for the purposes of evaluating this paper alone, it is not relevant to the claim. If you want evidence for or against GART being associated with a Mendelian disease, I recommend querying gene-disease resources (OMIM, ClinVar, HGMD), searching the primary literature for GART case reports or functional studies, and checking population databases (gnomAD) for constraint metrics and known pathogenic variants.


[Read Paper](https://www.semanticscholar.org/paper/83a11d71a34fe3ae5618faf4b4a593ecd29e0884)


## Search Queries Used

- GART pathogenic variant mutation Mendelian monogenic disease case report ClinVar OMIM whole exome sequencing whole genome sequencing rare disease

- GART phosphoribosylglycinamide formyltransferase phosphoribosylglycinamide synthetase trifunctional enzyme de novo purine biosynthesis purine metabolism enzyme deficiency developmental delay intellectual disability metabolic encephalopathy

- GART knockout mouse conditional knockout embryonic lethal zebrafish morpholino CRISPR functional study purinosome cell model loss of function phenotype

- GART loss of function intolerance pLI gnomAD constraint missense Z score population variation allele frequency pathogenic variant burden


## Bibliometric Configuration

- Use Bibliometrics: True

- Author Impact Weight: 0.4

- Citation Impact Weight: 0.4

- Venue Impact Weight: 0.2
