## Supplementary material for "MENDELSEEK: An algorithm that predicts Mendelian Genes and elucidates what makes them special": GAS1.txt

# Claim: The gene GAS1 is likely associated with at least one Mendelian disease.

**Status**: processed

**Overall Rating**: 4

**Explanation**:

Supporting evidence.
Multiple lines of evidence from the provided papers support an association between GAS1 and at least one Mendelian developmental disorder (holoprosencephaly, HPE, or HPE-like craniofacial malformations). In mouse genetics, targeted deletion of Gas1 produces midline craniofacial defects described as microform HPE (midfacial hypoplasia, incisor fusion, cleft palate and ear defects) and is associated with reduced Sonic hedgehog (Shh) signaling at a distance from the ligand source. The Gas1(-/-) phenotype is genetically interactive with Shh: loss of a single Shh allele in a Gas1(-/-) background significantly worsens midline craniofacial defects. This provides strong functional evidence that Gas1 is required for normal SHH pathway activity in craniofacial development and that its loss can produce an HPE-like phenotype in an experimental mammal.

Human genetic evidence in the provided excerpts is consistent with GAS1 being a candidate disease locus. The Seppälä paper notes human GAS1 maps to 9q21.3–q22, a region previously linked to nonsyndromic cleft palate and congenital deafness, and explicitly proposes GAS1 as a potential locus for several craniofacial malformations. The Ribeiro et al. report describes four Brazilian patients with GAS1 DNA sequence changes who presented a spectrum from classic HPE to HPE-like signs; in two patients the only identified change was in GAS1, while in two others GAS1 change co-occurred with SHH mutation. These human observations, combined with the mouse functional data and the biochemical interaction with SHH signaling, form a plausible chain of evidence that GAS1 variation can contribute to HPE-spectrum phenotypes in humans.

Caveats or contradictory evidence.
The human genetic evidence is limited in scale and detail in the provided excerpts. The Ribeiro paper describes "GAS1 DNA sequence change(s)" but the excerpts do not document whether those variants were definitively pathogenic (e.g., segregation in families, de novo occurrence, functional validation, population frequency, or in silico/pathogenicity classification). Two of the four patients also had SHH mutations, indicating possible oligogenic (multiple-hit) etiology rather than GAS1 being a sole Mendelian cause. The Seppälä mouse data show forebrain gross morphology remained intact in Gas1 nulls (microform HPE rather than full HPE), suggesting that GAS1 deficiency may produce milder or tissue-limited phenotypes and/or act as a modifier of other HPE genes rather than being a highly penetrant primary cause. HPE is genetically heterogeneous with variable expressivity and incomplete penetrance, and environmental modifiers can play roles; this complicates interpretation of small human series and mouse phenotypes.

Analysis of mechanisms, deeper relationships, and implications.
Biologically, GAS1 encodes a GPI-anchored membrane glycoprotein that modulates SHH signaling; the mouse data indicate Gas1 can potentiate hedgehog signaling in facial primordia. Since pathogenic variants in SHH pathway genes (SHH, PTCH1, GLI2, etc.) are established causes of Mendelian HPE, a gene that directly modulates SHH signaling has a clear mechanistic route to producing similar phenotypes. The genetic interaction data (Gas1 null exacerbating Shh haploinsufficiency) and co-occurrence of GAS1 and SHH variants in some patients argue for GAS1 functioning in the same developmental pathway and, depending on variant type and genetic background, acting either as a primary causal gene (in some instances) or as a modifier that influences penetrance/severity.

Which rating is most appropriate and why.
Weighing the evidence: the combination of (1) reproducible functional animal data showing an HPE-like phenotype with loss of Gas1, (2) mechanistic plausibility via direct modulation of SHH signaling, and (3) human case reports describing GAS1 sequence changes in patients with HPE-spectrum phenotypes (including some with GAS1 change alone) supports an association between GAS1 and at least one Mendelian developmental disorder. However, the human evidence as presented is limited in sample size and lacks definitive pathogenicity proof and broader genetic validation; GAS1 appears in the evidence sometimes as a modifier or contributor in an oligogenic context. Therefore the evidence is reasonably strong but not definitive. The best-fitting category is "Likely True": GAS1 is likely associated with at least one Mendelian disease (HPE or HPE-like craniofacial malformations), though larger human genetic series, rigorous variant pathogenicity work, and segregation/functional validation would be required to elevate this to "Highly Supported."


**Final Reasoning**:

I reviewed the logic and evidence and find it consistent: (a) strong mechanistic and functional support from mouse knockouts and pathway interaction with SHH; (b) human case series showing GAS1 variants in patients with HPE-like phenotypes, including instances where GAS1 was the only identified change; (c) caveats that human data are limited and some cases involve additional mutations (oligogenicity), and full pathogenic proof for human variants is not provided in the excerpts. Taking the totality of evidence, the claim that GAS1 is likely associated with at least one Mendelian disease (specifically HPE-spectrum disorders) is supported but not proven beyond doubt. Therefore "Likely True" is the appropriate rating.


## Relevant Papers


### Gas1 is a modifier for holoprosencephaly and genetically interacts with sonic hedgehog.

**Authors**: M. Seppala (H-index: 12), M. Cobourne (H-index: 44)

**Relevance**: 0.85

**Bibliometric Impact**: 0.40717647058823536


**Excerpts**:

- Holoprosencephaly (HPE) is a clinically heterogeneous developmental anomaly affecting the CNS and face, in which the embryonic forebrain fails to divide into distinct halves.

- Numerous genetic loci and environmental factors are implicated in HPE, but mutation in the sonic hedgehog (Shh) gene is an established cause in both humans and mice.

- As growth arrest-specific 1 (Gas1) encodes a membrane glycoprotein previously identified as a Shh antagonist in the somite, we analyzed the craniofacial phenotype of mice harboring a targeted Gas1 deletion.

- Gas1(-/-) mice exhibited microform HPE, including midfacial hypoplasia, premaxillary incisor fusion, and cleft palate, in addition to severe ear defects; however, gross integrity of the forebrain remained intact.

- These defects were associated with partial loss of Shh signaling in cells at a distance from the source of transcription, suggesting that Gas1 can potentiate hedgehog signaling in the early face.

- Loss of a single Shh allele in a Gas1(-/-) background significantly exacerbated the midline craniofacial phenotype, providing genetic evidence that Shh and Gas1 interact.

- As human GAS1 maps to chromosome 9q21.3-q22, a region previously associated with nonsyndromic cleft palate and congenital deafness, our results establish GAS1 as a potential locus for several human craniofacial malformations.


**Explanations**:

- Context sentence defining holoprosencephaly (HPE). This provides disease context relevant to Mendelian craniofacial disorders; it is background (neither direct nor mechanistic evidence for GAS1) but necessary to interpret the phenotype described later.

- States that SHH mutations are established causes of HPE in humans and mice. This provides mechanistic and comparative disease context: since GAS1 interacts with the SHH pathway, the established Mendelian role of SHH strengthens biological plausibility that GAS1 perturbation could produce Mendelian-like craniofacial defects. Limitation: this sentence does not provide direct evidence on GAS1 itself.

- Describes GAS1 as a membrane glycoprotein previously identified as a SHH antagonist in the somite and states the experimental approach (analysis of mice with targeted Gas1 deletion). This is mechanistic context and experimental setup, establishing that the subsequent findings derive from a Gas1 knockout mouse model. Limitation: use of a mouse model does not by itself prove a human Mendelian disease association.

- Reports the primary experimental finding: Gas1(-/-) mice show microform HPE features (midfacial hypoplasia, premaxillary incisor fusion, cleft palate) plus severe ear defects, while the gross forebrain remains intact. This is direct experimental evidence that loss of Gas1 causes craniofacial malformations in a mammalian model, supporting the plausibility that GAS1 mutations could cause Mendelian developmental disorders. Limitations: phenotype is a 'microform' HPE rather than full HPE; findings are in mice (model organism), and no human pathogenic variants or segregation data are presented here.

- Provides mechanistic evidence linking the Gas1-null phenotype to a partial loss of SHH signaling in cells distant from the transcription source, and suggests GAS1 can potentiate hedgehog signaling in the early face. This directly supports a biological mechanism by which GAS1 influences craniofacial development and thus strengthens plausibility for disease association. Limitation: mechanistic inference is based on the experimental system used (mouse embryos); degree and universality of this mechanism in humans are not established here.

- Describes a genetic interaction: loss of one Shh allele on a Gas1(-/-) background significantly worsens the midline craniofacial phenotype, providing genetic evidence that Shh and Gas1 interact. This is strong mechanistic/genetic evidence (epistasis/modifier effect) that GAS1 functions in the SHH pathway relevant to midline craniofacial development. Limitation: interaction observed in engineered mouse genotypes; it supports pathway involvement but does not demonstrate a human Mendelian GAS1 mutation causing disease.

- States that human GAS1 maps to 9q21.3-q22, a region previously associated with nonsyndromic cleft palate and congenital deafness, and concludes GAS1 is a potential locus for several human craniofacial malformations. This ties the mouse findings to human genetics and is the paper's claim that GAS1 is a candidate human disease locus. Strength: provides positional human-genetic context linking GAS1 to known craniofacial malformation loci. Limitations: mapping to a chromosomal region and being a plausible candidate is not equivalent to demonstrating causative Mendelian mutations in GAS1; no human mutation or segregation data are provided in the quoted content.


[Read Paper](https://www.semanticscholar.org/paper/ebe7bb065d08d5c83dbe2dcef44f38d09241d687)


### Holoprosencephaly and holoprosencephaly‐like phenotype and GAS1 DNA sequence changes: Report of four Brazilian patients

**Authors**: L. Ribeiro (H-index: 11), A. Richieri‐Costa (H-index: 34)

**Relevance**: 0.8

**Bibliometric Impact**: 0.3217714285714286


**Excerpts**:

- Holoprosencephaly (HPE) is genetically heterogeneous.

- Variable phenotypic manifestations within families with normal and affected patients have been attributed to the number and type of HPE gene mutations.

- Environmental agents may also contribute to the severity as well as the requirement of multiple hits.

- Main genes involved include SHH, GLI2, PTCH1, TGIF, ZIC2, TDGF1, SIX3; however, several other candidates have been proposed.

- Recently it was established that the human growth arrest specific gene 1 (GAS1) is a potential locus for several human craniofacial malformations.

- Here, we report on four Brazilian patients with GAS1 DNA sequence change who presented variable phenotypical manifestations ranging from classic HPE to HPE‐like signs.

- Two patients had single DNA sequence change in the GAS1 gene, while in other two, an additional mutation in the SHH gene was observed.

- Clinical manifestations presented by these patients suggest that GAS1 could be considered a candidate locus for one of the types of human HPE.


**Explanations**:

- How it relates: Provides disease context showing HPE is caused by multiple genes. Relevance: contextual/mechanistic. Strength: establishes that HPE has Mendelian genetic causes, so discovering additional genes (like GAS1) is plausible. Limitation: This sentence is background only and does not provide evidence specific to GAS1.

- How it relates: Explains that the number and nature of mutations in HPE genes influence phenotype. Relevance: mechanistic/contextual. Strength: supports the idea that variants in a gene can be causative or contributory to HPE. Limitation: General statement – does not present data on GAS1 variants or segregation.

- How it relates: States that non-genetic factors and multi-hit models contribute to HPE severity. Relevance: mechanistic. Strength: highlights biological plausibility for variable expressivity and gene–environment or gene–gene interactions involving candidate loci like GAS1. Limitation: Again, general; does not demonstrate GAS1-specific mechanisms or causation.

- How it relates: Lists well-established HPE genes and notes other candidates have been proposed. Relevance: contextual. Strength: Positions GAS1 in the broader gene-discovery landscape (i.e., not yet a main established gene). Limitation: Does not by itself support or refute GAS1 association; indicates GAS1 is among putative candidates.

- How it relates: Direct statement that GAS1 has been identified as a potential locus for craniofacial malformations. Relevance: direct evidence/interpretation by authors. Strength: Explicitly connects GAS1 to relevant human malformations; increases plausibility that GAS1 variants can cause or contribute to disease. Limitation: Uses the word "potential," indicating preliminary/uncertain status; no supporting data or strength of association given in this sentence.

- How it relates: Reports primary observational data: four patients with GAS1 DNA sequence changes and HPE or HPE-like phenotypes. Relevance: direct evidence. Strength: Provides actual cases showing co-occurrence of GAS1 variants and HPE phenotypes, which is key evidence toward association. Limitation: Small sample size; the excerpt does not report whether variants were rare, predicted damaging, segregated in families, or functionally validated—so causality is not established.

- How it relates: Details genotype findings among the four cases: two with GAS1-only changes and two with GAS1 plus SHH mutation. Relevance: direct and mechanistic. Strength: The presence of GAS1-only cases could support a causal role; the GAS1+SHH cases suggest possible gene–gene interaction or that GAS1 may act as a modifier. Limitation: Without segregation, functional studies, or population frequency data, these observations could reflect coincidental rare variation, and the presence of SHH mutations in some cases complicates causal attribution to GAS1 alone.

- How it relates: Authors' conclusion that GAS1 should be considered a candidate locus for a type of human HPE. Relevance: direct interpretive evidence. Strength: The authors synthesize their observations and prior information to propose GAS1 as a candidate Mendelian locus for HPE. Limitation: Phraseology "could be considered a candidate" signals that evidence is suggestive but not definitive; the excerpt does not convey evidence strength, statistical support, segregation data, or functional validation required to confirm GAS1 as a Mendelian disease gene.


[Read Paper](https://www.semanticscholar.org/paper/834ab90b7e72cdede629a9202030e0f5527f8ca3)


### THE JAPANESE TERATOLOGY SOCIETY 62ND ANNUAL MEETING

**Authors**: H. Sakata-Haga (H-index: 0), T. Mochizuki (H-index: 0)

**Relevance**: 0.2

**Bibliometric Impact**: 0.0


**Excerpts**:

- The Sonic hedgehog (Shh) signaling pathway plays a crucial role in craniofacial development, but the severe defects in early developmental stage in Shh-/- mice has made it difficult to understand the detailed molecular roles of this gene in craniofacial development.

- In this study, we investigated the mice in which Shh expression is specifically deleted in the forebrain (Sox1-Cre;Shhfl/fl) and the oropharyngeal epithelium (Isl1-Cre;Shhfl/fl).

- Sox1-Cre;Shhfl/fl mice demonstrated mild holoprosencephaly, whereas Isl1-Cre;Shhfl/fl mice exhibited essentially normal brain with relatively smaller cerebrum.

- In Sox1-Cre; Shhfl/fl mice, the bones and cartilages in proximity to the brain, as the cranial base and the orbital floor, were largely defected.

- In contrast, the secondary palate in Sox1-Cre;Shhfl/fl mice was normally formed, suggesting that Shh expression in the forebrain is crucial for the development of bones adjacent to the brain but not necessary to the palate development.

- In Isl1-Cre;Shhfl/fl mice, the distal portion of the upper jaw, the cranial base, and the palate were largely defected.

- Our RNA-seq analysis revealed that Satb2 (Special AT-rich sequence-binding protein 2) is a downstream target of Shh in the upper jaw formation.

- The development of genome editing tools, especially CRISPR/Cas9, an RNA-dependent nuclease has made it possible to perform direct knockout and site-specific knockin on fertilized eggs, enabling the generation of animal models of congenital diseases with arbitrary genetic abnormalities at an unprecedented speed.

- We have generated novel X-SCID lines using CRISPR/Cas9. One of the lines expressed Il2rg protein despite a 7-nucleotide deletion in Exon 2. This is an example of a genomic deletion not granting a phenotype, and suggests the need to confirm expression at the protein level in knockout lines.


**Explanations**:

- Mechanistic relevance: This sentence establishes that Shh signaling is critical for craniofacial development and notes severe phenotypes when Shh is entirely lost in mice. Relevance to the GAS1 claim: GAS1 is known (outside this paper) to modulate SHH signaling, so the centrality of SHH to a Mendelian craniofacial phenotype makes it biologically plausible that genes affecting SHH (like GAS1) could underlie Mendelian disease. Limitations: The paper does not mention GAS1 or provide any direct genetic or clinical evidence linking GAS1 to disease. This is mechanistic, not direct, evidence.

- Contextual/methodological relevance: This sentence describes conditional deletion strategies used to dissect tissue-specific roles of Shh. Relevance to GAS1: it shows how perturbation of SHH in particular tissues produces discrete developmental outcomes — supporting the idea that components/modulators of SHH signaling could produce Mendelian phenotypes when disrupted. Limitation: no data on GAS1 specifically; conditional Shh deletions are not equivalent to mutations in a SHH-modulating gene.

- Mechanistic evidence: Demonstrates that loss of Shh in certain contexts causes holoprosencephaly and other brain size defects. Relevance to GAS1: holoprosencephaly is a well-characterized Mendelian or oligogenic phenotype in humans that can result from disrupted SHH pathway activity; therefore, genes that alter SHH signaling could plausibly be disease genes. Limitation: This is indirect; the paper shows consequences of Shh loss, not of GAS1 alteration, and mouse conditional-knockout phenotypes do not prove human Mendelian association.

- Mechanistic evidence linking SHH perturbation to craniofacial skeletal defects. Relevance to GAS1: if GAS1 modulates SHH activity during cranial base/orbital floor development, variation in GAS1 could plausibly produce similar skeletal defects and hence Mendelian craniofacial disorders. Limitation: the excerpt is an observation about Shh loss; it does not measure or implicate GAS1, nor does it provide human genetic data or demonstrate causality for GAS1.

- Mechanistic/contextual: Shows that Shh's spatial expression matters for which craniofacial structures are affected. Relevance to GAS1: suggests that genes regulating SHH spatial/temporal signaling can have specific phenotypic consequences; again this provides mechanistic plausibility for a SHH-modulating gene being a Mendelian disease gene. Limitation: no mention of GAS1, no direct genetic association.

- Mechanistic evidence of tissue-specific defects following loss of Shh in a different domain. Relevance to GAS1: reinforces that altering SHH signaling in different embryonic compartments leads to definable congenital malformations — supporting the plausibility that a SHH-interacting gene could cause Mendelian craniofacial disease. Limitation: indirect inference only.

- Mechanistic and molecular relevance: identifies downstream targets of Shh (Satb2) implicated in jaw formation. Relevance to GAS1: indicates that SHH acts via transcriptional programs to pattern craniofacial structures; a SHH co-receptor or modulator (such as GAS1) could affect these downstream programs and thus underlie Mendelian phenotypes. Limitation: Satb2 is the named downstream target, not GAS1; no experimental link to GAS1 is provided.

- Methodological relevance: explains that CRISPR/Cas9 can rapidly generate animal models of congenital disease. Relevance to GAS1: indicates that genes suspected of causing Mendelian disease (e.g., hypothetical GAS1 variants) can be modeled to test pathogenicity. Limitation: this is methodological and does not provide evidence that GAS1 is associated with any Mendelian disease.

- Practical caveat about knockout interpretation: describes an instance where a genomic deletion did not abolish protein expression or phenotype. Relevance to GAS1: warns that not all genomic alterations produce expected phenotypes and that protein-level confirmation is necessary when modelling candidate Mendelian genes such as GAS1. Limitation: does not provide evidence for or against GAS1 involvement in disease; it is a general technical caveat.


[Read Paper](https://www.semanticscholar.org/paper/162811312bde34697100a4eaab4c8748157e2132)


## Other Reviewed Papers


### Precision Medicine in Internal Medicine

**Why Not Relevant**: This paper is a general review of genetic testing strategies, exome/genome sequencing, interpretation frameworks, and the state of gene–disease discovery. It contains no mention of GAS1, no experimental data, no case reports, no statistical associations, and no mechanistic studies specific to GAS1. Therefore it provides no direct evidence for or against the claim that GAS1 is associated with at least one Mendelian disease. The included excerpts only offer general context about how gene–disease associations are discovered and interpreted; they do not address GAS1 specifically. Because the claim concerns a specific gene–disease association, the paper is not relevant as direct evidence.


[Read Paper](https://www.semanticscholar.org/paper/c7199e8a5226f8d9d55777da0e15e91797bb459a)


### CDKN1C gene mutation causing familial Silver–Russell syndrome: A case report and review of literature

**Why Not Relevant**: This paper presents a familial case report and analysis of a CDKN1C missense mutation (p.Arg279His) causing a Silver-Russell syndrome (SRS) phenotype and discusses the relationship of CDKN1C mutations to IMAGe syndrome. The provided text does not mention the gene GAS1, GAS1 variants, or any diseases or mechanisms involving GAS1. Therefore the paper contains no direct genetic evidence (segregation, variant pathogenicity, case reports) nor any mechanistic evidence (protein interactions, pathways, functional assays) that would support or refute the claim that GAS1 is associated with a Mendelian disease. Specific reasons the paper is not relevant to the GAS1 claim: 

- Gene focus: All genetic findings reported are for CDKN1C (c.836G>A, p.Arg279His). No sentences in the provided content refer to GAS1.
- Diseases discussed: The clinical syndromes described are Silver-Russell syndrome and IMAGe syndrome, both discussed in relation to CDKN1C. There is no discussion of any phenotype or syndrome linked to GAS1.
- Mechanisms discussed: Mechanistic comments concern CDKN1C function, the PCNA domain, and how different missense mutations in that region relate to phenotype. No mechanistic pathway involving GAS1 is described.
- Evidence type: The paper is a family case report with exome sequencing for CDKN1C; even if strong for CDKN1C, it cannot be extrapolated to provide evidence about a different gene not mentioned in the study.

Given the complete absence of GAS1 in the paper content, this document does not provide any supporting or refuting evidence for the claim that GAS1 is likely associated with at least one Mendelian disease. To evaluate that claim one would need studies (case reports, case series, linkage/segregation analyses, functional assays, or curated database entries) that specifically report GAS1 variants segregating with a Mendelian phenotype or demonstrate mechanistic pathogenicity of GAS1 variants.


[Read Paper](https://www.semanticscholar.org/paper/40aebdbf1ccb88434e9e6c904faab1782e362cc9)


### A pediatric case of congenital neutropenia with SRP54 gene mutation in which monocytosis and gingival swelling were useful in differentiating from autoimmune neutropenia

**Why Not Relevant**: The paper exclusively discusses SRP54 (and, in comparison, ELANE) mutations causing congenital neutropenia and associated clinical/hematologic findings. The text contains specific genetic, phenotypic, and treatment-response information for SRP54 but contains no mention of the GAS1 gene, no data on GAS1 variants, and no mechanistic discussion involving GAS1. Therefore this paper provides no direct or mechanistic evidence for or against the claim that GAS1 is likely associated with at least one Mendelian disease. Absence of evidence in this paper is not evidence that GAS1 is unrelated to Mendelian disease in general, but for the purpose of evaluating the stated claim, this document is not relevant.


[Read Paper](https://www.semanticscholar.org/paper/fe60f7a6e60346243006c1f2e266d8dcb817e74b)


### Stepping on the GAS: a brake pedal for melanoma metastasis?

**Why Not Relevant**: The paper focuses on GAS1 in the context of melanoma metastasis, presenting evidence from somatic tumor samples, cancer cell lines, and mouse metastasis models. It documents altered expression, functional effects on metastasis when GAS1 is manipulated, and somatic deletions of the GAS1 locus in tumors. However, it contains no data on germline variants, family segregation, penetrance, inheritance patterns, clinical descriptions of an inherited syndrome, or case-control associations that would indicate GAS1 causes or is associated with a Mendelian disease. While the mechanistic link to SHH signaling provides biological plausibility that germline GAS1 perturbation could affect development, the paper does not present direct or epidemiologic evidence of Mendelian disease. Therefore the content is largely not relevant to the claim that GAS1 is likely associated with at least one Mendelian disease.


[Read Paper](https://www.semanticscholar.org/paper/d274ab000c1015a99f134c7a0726b352b2ce6bf8)


### Severe congenital neutropenia with mastoidectomy: A case report of HAX1

**Why Not Relevant**: The provided paper content concerns severe congenital neutropenia (Kostmann disease) and identifies ELANE, HAX1, G6PC3, and GFI1 as genes implicated in that Mendelian disorder; it gives a specific HAX1 homozygous mutation (pW44X) in a patient. Nowhere in the supplied text is the gene GAS1 mentioned, nor are there data, analyses, or mechanistic discussions linking GAS1 to any Mendelian phenotype. Therefore this paper does not provide evidence for or against the claim that GAS1 is associated with a Mendelian disease. Absence of mention in this disease-focused paper does not constitute evidence that GAS1 is not associated with any Mendelian disease elsewhere; it simply means this source is not relevant to evaluating the claim about GAS1.


[Read Paper](https://www.semanticscholar.org/paper/4286358f6cfe92498f9f4460ec9da5aaddd838f0)


### CHAPTER 12 Growth Faaors Regulating Neural Crest Cell Fate Decisions

**Why Not Relevant**: The provided paper content does not mention the gene GAS1 (no occurrences of the string "GAS1", "Gas1", or variants) and contains no discussion of specific genes being linked to Mendelian diseases. The text is a review-style discussion of neural crest cell plasticity and the roles of various growth factors (e.g., GDNF, ET3, SCF, BMP2, TGFβ, FGF2, Shh, Notch, NRG1) in lineage decisions. While it provides mechanistic background on signaling pathways (including sonic hedgehog signaling) that could be conceptually relevant to a gene like GAS1, there is no direct genetic, clinical, or experimental evidence in this excerpt that associates GAS1 with any Mendelian disorder. Therefore the paper content is largely non-relevant to the specific claim that "GAS1 is likely associated with at least one Mendelian disease."


[Read Paper](https://www.semanticscholar.org/paper/22aed51de8c7cbe6391b8e4ef640711ce2305d96)


## Search Queries Used

- GAS1 mutation holoprosencephaly case report growth arrest specific 1

- GAS1 pathogenic variant ClinVar OMIM segregation family de novo growth arrest specific 1 Mendelian disease

- GAS1 sonic hedgehog SHH signaling developmental biology mouse knockout phenotype embryo functional study growth arrest specific 1

- GAS1 population frequency gnomAD penetrance incomplete penetrance variable expressivity modifier gene review growth arrest specific 1


## Usage Statistics

- Prompt Tokens: 0

- Completion Tokens: 0

- Total Tokens: 0

- Estimated Cost: $0.0511


## Bibliometric Configuration

- Use Bibliometrics: True

- Author Impact Weight: 0.4

- Citation Impact Weight: 0.4

- Venue Impact Weight: 0.2
