## Supplementary material for "MENDELSEEK: An algorithm that predicts Mendelian Genes and elucidates what makes them special": GAS6.txt

# Claim: The gene GAS6 is likely associated with at least one Mendelian disease.

**Status**: processed

**Overall Rating**: -1

**Explanation**:

No relevant papers were found that support or refute this claim.


## Relevant Papers


## Other Reviewed Papers


### The Burden of Candidate Pathogenic Variants for Kidney and Genitourinary Disorders Emerging From Exome Sequencing

**Why Not Relevant**: The provided paper content describes methods for identifying and curating genes associated with Mendelian kidney and genitourinary disorders and for screening a large control cohort for candidate pathogenic variants. However, the excerpts and surrounding content do not mention the gene GAS6 by name, nor do they present gene-level results or tables that would indicate whether GAS6 is included among the curated 625 genes or whether it has variants classified as pathogenic. All included sentences are methodological or high-level results about the gene-curation and variant-annotation workflow. Because the claim concerns the likelihood that GAS6 is associated with at least one Mendelian disease, and the paper content supplied contains no direct or indirect evidence about GAS6 specifically (no explicit inclusion, variant counts, disease associations, or mechanistic data for GAS6), this document cannot be used to support or refute the claim. If the full paper or the supplementary gene list were available, one could check whether GAS6 appears among the curated genes or whether ClinVar/HGMD entries for GAS6 exist; absent that, relevance to the GAS6-specific claim is negligible.


[Read Paper](https://www.semanticscholar.org/paper/a75e297208c04992568e607d19aa9b9aaebd09d5)


### Whole-Exome Sequencing in Adults With Chronic Kidney Disease

**Why Not Relevant**: The provided paper content does not mention the gene GAS6, nor does it present results (positive or negative) about GAS6 variants or phenotypes. The study focuses on using whole-exome sequencing to identify Mendelian causes of chronic kidney disease and hypertensive disorders and describes a curated list of 287 OMIM 'nephropathy genes', sequencing coverage, variant-calling and annotation methods, and known capture blind spots. Because the gene-level results and Supplement Table 3 (the gene list) are not included in the provided text, there is no direct evidence here that GAS6 is included among the evaluated Mendelian genes, nor any mechanistic or clinical findings linking GAS6 to a Mendelian disease. The methodological excerpts show how the authors would detect Mendelian variants and also highlight limitations (uncaptured exons, filtering thresholds) that could lead to false negatives, but they do not supply any GAS6-specific data. Therefore this paper, as provided, cannot be used to support or refute the claim that GAS6 is likely associated with at least one Mendelian disease. Absence of mention in this excerpt is not definitive evidence that GAS6 is not a Mendelian disease gene; it only means this paper's available text provides no relevant evidence either way.


[Read Paper](https://www.semanticscholar.org/paper/a29bd57964f68ac5f8e1abcafe1112ba488f53f5)


### From case studies to community knowledge base: MSeqDR provides a platform for the curation and genomic analysis of mitochondrial diseases

**Why Not Relevant**: This paper is an overview of mitochondrial disease heterogeneity and describes the MSeqDR resource for collecting, curating, and analyzing mitochondrial disease genes and variants, with specific discussion of PMPCA as an example. The text makes no mention of the gene GAS6, provides no data (genetic association, case reports, variant pathogenicity, segregation, functional assays) linking GAS6 to any Mendelian phenotype, and discusses mechanisms and evidence only in the context of mitochondrial proteins and mitochondrial disease gene discovery. Therefore the paper provides no direct evidence supporting the claim that GAS6 is associated with a Mendelian disease, nor does it provide mechanistic evidence implicating GAS6. Absence of evidence in this single paper is not evidence that GAS6 is not associated with a Mendelian disease in the broader literature, but based on the provided content alone, there is no support for the claim.


[Read Paper](https://www.semanticscholar.org/paper/6f85c595adaca6963c30fea73bc37266fa78be56)


### Autosomal Dominant Intellectual Development Disorder-6 (MRD6) Without Seizures Linked to a De Novo Mutation in the grin2b Gene Revealed by Exome Sequencing: A Case Report of a Moroccan Child

**Why Not Relevant**: The paper exclusively discusses GRIN2B (grin2b) and a specific nonsense variant (c.3912C>G, p.Tyr1304Ter) causing MRD6 (OMIM 613970). There is no mention of the gene GAS6, no data (genetic, clinical, population, or functional) concerning GAS6, and no discussion of mechanistic links between GAS6 and any Mendelian phenotype. Therefore the paper provides no direct or mechanistic evidence supporting or refuting the claim that GAS6 is likely associated with at least one Mendelian disease. Absence of evidence in this paper does not disprove the claim generally, but it means this paper cannot be used to support the claim. Additional literature specifically addressing GAS6 would be required to evaluate the claim.


[Read Paper](https://www.semanticscholar.org/paper/67d756e0cc302e5d46fdda9df34403f1f2de9a5a)


### The V368i mutation in Twinkle does not segregate with adPEO

**Why Not Relevant**: The paper does not mention the gene GAS6, perform any experiments on GAS6, or provide genetic/functional evidence bearing on GAS6. All reported genetic findings and mechanistic discussion concern ANT1, C10ORF2 (Twinkle), POLG1, mtDNA deletions, and the Twinkle V368I variant. Therefore this paper provides no direct or mechanistic evidence to support the claim that GAS6 is likely associated with any Mendelian disease. Absence of a GAS6 discussion in this disease-focused study does not constitute evidence that GAS6 is not associated with Mendelian disease in other contexts; it merely indicates GAS6 was not studied or implicated here.


[Read Paper](https://www.semanticscholar.org/paper/4f4aef3668232d5c4630ee81bc3c87051593ff0a)


## Search Queries Used

- GAS6 mutation pathogenic Mendelian disease case report family segregation autosomal dominant autosomal recessive OMIM ClinVar

- GAS6 TAM receptor AXL MERTK TYRO3 interaction modifier gene retinitis pigmentosa retinal dystrophy Mendelian knockout mouse phenotype

- GAS6 rare variants exome sequencing rare disease cohort burden analysis loss of function constraint gnomAD pLI ClinVar population genetics

- GAS6 gene disease association review systematic review meta analysis ClinGen OMIM Orphanet HGMD gene curation Mendelian disease clinical genetics


## Bibliometric Configuration

- Use Bibliometrics: True

- Author Impact Weight: 0.4

- Citation Impact Weight: 0.4

- Venue Impact Weight: 0.2
