## Supplementary material for "MENDELSEEK: An algorithm that predicts Mendelian Genes and elucidates what makes them special": GMDS.txt

# Claim: The gene GMDS is likely associated with at least one Mendelian disease.

**Status**: processed

**Overall Rating**: 3

**Explanation**:

Supporting evidence.
The provided paper identifies GDP-mannose 4,6-dehydratase (GMD) as a required enzyme in the de novo GDP-fucose biosynthesis pathway and shows that loss of functional GMD abolishes cellular fucosylation in human colon cancer cell lines (HCT116 and others). The authors demonstrate truncated GMD transcripts, absent GMD protein, and rescue of fucosylation by transfection with full-length GMD, establishing that loss of GMDS-encoded activity can produce a clear biochemical defect. The paper also emphasizes the biological importance of fucosylation: knockout of other pathway components (e.g., FX in mice) causes embryonic lethality, and elimination of fucosyltransferases (Pofut1, Pofut2) likewise causes severe developmental phenotypes in mice. These lines of evidence show a strong mechanistic link between GMDS function, an essential glycosylation pathway, and development/physiology — the kind of biology that underlies many Mendelian disorders when mutated.

Caveats or contradictory evidence.
The excerpts do not report any human germline (inherited) mutations in GMDS causing a Mendelian disease. The examples given are mainly somatic loss in cancer cell lines or knockouts of related genes in mice. Somatic inactivation in tumors does not demonstrate a heritable human phenotype. Also, the paper notes the existence of a salvage pathway for GDP-fucose that can restore fucosylation when exogenous fucose is provided; this suggests partial biochemical redundancy that could modulate or mitigate phenotypes from loss of the de novo pathway in vivo. Finally, embryonic lethality for some related genes in mice could argue either that severe GMDS loss might be incompatible with live birth (reducing observed Mendelian cases) or that phenotypes differ between species; none of these possibilities is resolved by the provided text.

Analysis of mechanisms and implications.
GMDS encodes the enzyme catalyzing the first committed step of de novo GDP-fucose synthesis (conversion of GDP-mannose to GDP-4-keto-6-deoxymannose), so loss-of-function is expected to reduce or eliminate fucosylation of glycoproteins and glycolipids. Fucosylation is critical for Notch signaling and other developmental pathways, explaining why disruptions of fucose metabolism or transfer are associated with strong developmental phenotypes in model organisms. Given that other genes in fucose metabolism and fucosylation (e.g., SLC35C1, POFUT1/2) are implicated in human Mendelian disorders or embryonic lethality in model organisms, the pathway is clearly disease-relevant. The presence of a salvage pathway (exogenous fucose uptake) introduces a potential modifier: partial activity via salvage could allow survival with a milder clinical phenotype, or severity could depend on dietary/environmental factors. Also, somatic loss in cancer shows that complete loss can be tolerated in specific contexts (tumors) but does not prove germline pathogenicity.

Which rating is most appropriate.
The evidence shows strong mechanistic plausibility that deleterious variation in GMDS could cause a Mendelian disorder because GMDS is essential to a critical glycosylation pathway and loss of related genes causes profound developmental phenotypes. However, the provided paper excerpts do not present any direct evidence of human germline GMDS mutations causing a Mendelian disease (no case reports, pedigrees, or clinical associations). Given plausible biology but absence of direct human-disease evidence in the supplied material, the balanced conclusion is that the evidence is mixed: supportive mechanistic data and relevance of the pathway, but no direct documented Mendelian association in the provided excerpts.


**Final Reasoning**:

I reviewed the logic and evidence: the paper convincingly establishes that GMDS encodes an essential enzyme for de novo GDP-fucose synthesis and that loss of its activity abolishes fucosylation in human cell lines. It also documents that perturbing fucosylation or fucose-processing genes produces severe developmental phenotypes in model organisms. These facts make it biologically plausible that pathogenic germline variants in GMDS could cause a Mendelian disorder. However, the provided excerpts contain no direct report of human germline GMDS mutations causing a heritable disease, nor clinical case data. The existence of a salvage pathway and the possibility of embryonic lethality further complicate inference from somatic or model-organism data to an observed human Mendelian disorder. Therefore the best-supported judgment from the supplied evidence and domain knowledge is "Mixed Evidence": the claim is plausible and mechanistically supported, but not directly confirmed by the provided human-disease data.


## Relevant Papers


### Fucose is on the TRAIL of colon cancer.

**Authors**: R. Haltiwanger (H-index: 61)

**Relevance**: 0.25

**Bibliometric Impact**: 0.6006133333333334


**Excerpts**:

- Two enzymes are required to convert GDP-mannose to GDP-fucose in the de novo pathway: GDP-mannose 4,6-dehydratase (GMD) and GDP-4-keto-6-deoxymannose 3,5-epimerase-4-reductase (FX).

- Mice lacking the FX gene (and thus all forms of fucosylation) display an embryonic lethal phenotype revealing the importance of fucose during development5.

- To examine the reason for this reduction, they assayed both fucosyltransferase enzyme activities and GDP-fucose levels in the cells and could not detect GDP-fucose. These results suggested some defect in the de novo GDP-fucose pathway (Figure 1), and RT-PCR analysis revealed a truncated transcript for GMD. Western blots showed the absence of GMD protein, and transfection of the cells with a construct encoding full-length GMD restored fucosylation, demonstrating that HCT116 cells have a defect in fucosylation due to loss of functional GMD.

- Aberrant transcripts for GMD were found in two other human colon cancer cell lines, as well as a number of colon and ovarian cancer tissue samples.

- Thus, loss of GMD may be a common mechanism for cancer cells to evade TRAIL-mediated killing.

- The presence of the salvage pathway for GDP-fucose biosynthesis may also provide a potential therapy for cancers that have lost GMD. Addition of fucose to the medium of cells with defects in the de novo pathway can restore fucosylation19. Thus, taking advantage of the salvage pathway by providing fucose should make GMD-lacking tumors susceptible to TRAIL-mediated killing in vivo.

- Elimination of the genes encoding either Pofut1 (O-fucosylates EGF repeats) or Pofut2 (O-fucosylates TSRs) in mice causes severe embryonic lethal phenotypes6, 7.

- The Notch receptor is modified at many sites with O-fucose, and loss of O-fucose in mammals abrogates function8. This explains the embryonic lethality of the Pofut1 knockout, since Notch function is essential for embryogenesis.


**Explanations**:

- This sentence identifies GMD as one of the two enzymes that catalyze the first committed step of the de novo GDP-fucose biosynthetic pathway. Relation to claim: mechanistic. Strength: establishes that GMD (GMDS) has a defined, essential biochemical role in fucose biosynthesis, which is plausibly relevant to human biology and thus to Mendelian disease if disrupted. Limitation: a biochemical role alone does not demonstrate that germline variants in GMDS cause a Mendelian phenotype.

- This reports that loss of the downstream enzyme FX in mice causes embryonic lethality, indicating that loss of GDP-fucose biosynthesis can produce severe, developmentally penetrant phenotypes in a mammalian organism. Relation to claim: mechanistic (supports plausibility). Strength: in vivo genetic evidence that disrupting GDP-fucose synthesis has dramatic developmental consequences. Limitation: this is for FX, not GMD; different enzymes in the same pathway can have non-identical phenotypes and FX knockout lethality does not prove GMDS/ GMD germline loss causes a Mendelian disease in humans.

- This paragraph reports experimental data from human cancer cells (HCT116) showing absence of detectable GDP-fucose, identification of a truncated GMD transcript, loss of GMD protein, and functional rescue of fucosylation by transfecting full-length GMD. Relation to claim: direct functional evidence (cellular) that loss of GMD abolishes GDP-fucose production and fucosylation. Strength: demonstrates causality in a human cell line (loss of functional GMD -> loss of fucosylation; rescue restores function). Methodological strengths include biochemical assays (GDP-fucose measurement), RT-PCR, western blots, and functional rescue. Limitations: evidence is from somatic mutations in a cancer cell line (HCT116) rather than germline variants in humans; it documents a cellular phenotype but not an inherited Mendelian disease. Also the truncated transcript may be cancer-associated rather than representative of germline variation.

- This sentence shows that aberrant GMD transcripts were observed in additional tumor cell lines and tumor tissue samples. Relation to claim: direct but somatic/cancer context. Strength: indicates that GMD dysfunction occurs recurrently in tumors, implying that loss-of-function alleles can arise and have biological consequences. Limitation: these are somatic alterations in cancer and do not demonstrate inherited (germline) pathogenic alleles causing Mendelian disease.

- This is an interpretation linking loss of GMD to an advantage for cancer cells (evading TRAIL-mediated killing). Relation to claim: contextual/direct (pathophysiological consequence of somatic GMD loss). Strength: supports that loss of GMD has measurable phenotypic consequences at the organismal/tissue level (tumor survival). Limitation: again relates to acquired cancer biology, not inherited Mendelian disease; it does not indicate penetrant developmental or systemic disease from germline GMDS variants.

- These sentences describe the salvage pathway and that exogenous fucose can restore fucosylation in cells defective in the de novo pathway, and propose a therapeutic approach for GMD-deficient tumors. Relation to claim: mechanistic context. Strength: shows biological redundancy (salvage pathway) that can mitigate loss of de novo GDP-fucose synthesis, which is relevant to the potential severity of germline GMDS loss. Limitation: presence of a salvage pathway suggests that germline loss of GMDS might be less severe or have tissue-specific effects depending on fucose availability; conversely, it complicates predictions about whether GMDS null alleles would produce a Mendelian disease. These data are from cell/tissue/cancer contexts, not human germline genetics.

- This sentence reports that removing O-fucosyltransferase genes (Pofut1 or Pofut2) in mice causes embryonic lethality. Relation to claim: mechanistic plausibility. Strength: corroborates that defects in fucosylation machinery (downstream of GDP-fucose synthesis) can produce severe developmental phenotypes in mammals. Limitation: applies to different genes (fucosyltransferases), not GMDS itself; phenotypes of transferase knockouts are informative but not definitive for consequences of GMDS germline loss because pathway nodes and redundancy may differ.

- This explains that loss of O-fucose (on Notch) abrogates Notch function and accounts for embryonic lethality in Pofut1 knockout mice. Relation to claim: mechanistic. Strength: connects biochemical glycosylation defects to a specific, essential signaling pathway (Notch) whose disruption causes embryonic lethality; this strengthens the plausibility that defects in GDP-fucose synthesis (via GMDS loss) could disrupt key developmental pathways. Limitation: again, the evidence is indirect (Notch/POFUT1) rather than direct evidence linking GMDS/ GMD germline variants to Mendelian disorders in humans or mice.

- non_relevant_explanation (omitted because relevance >= 0.1) ? (null) : ""

- excerpt_pages


[Read Paper](https://www.semanticscholar.org/paper/7cb823582c4547dc4e7029cf9c7083b772bb8696)


## Other Reviewed Papers


### The multiple faces of TOR1A: different inheritance, different phenotype.

**Why Not Relevant**: The provided paper excerpt contains no mention of the gene GMDS, its variants, phenotypes, or any experimental/clinical data concerning GMDS. The text discusses general concepts about Mendelian versus complex disorders and gives a detailed example concerning TOR1A (TorsinA) and DYT1-dystonia, but provides neither direct evidence (e.g., reported GMDS mutations in affected individuals, segregation data, case reports, cohorts) nor mechanistic evidence (e.g., GMDS molecular function linked to disease pathways, animal models, biochemical defects) relevant to the claim that GMDS is associated with a Mendelian disease. Therefore, based on the content provided, there is no basis to support or refute the claim about GMDS. Note: if additional sections of the paper (not provided here) mention GMDS, those should be supplied for reassessment; with only the current excerpt, relevance is zero.


[Read Paper](https://www.semanticscholar.org/paper/b74b91e0829b14562a5293323e09bb3633156aab)


## Search Queries Used

- GMDS mutation biallelic homozygous autosomal recessive Mendelian disorder case report patient developmental delay intellectual disability congenital anomalies monogenic pathogenic variant

- GMDS GDP mannose 4 6 dehydratase fucosylation GDP fucose biosynthesis glycosylation congenital disorder of glycosylation fucosylation defect mechanism

- GMDS ClinVar OMIM DECIPHER gnomAD constraint loss of function intolerance allele frequency pathogenic variant clinical significance

- GMDS knockout mouse zebrafish knockdown functional study phenotype developmental phenotype immunodeficiency leukocyte adhesion defect cancer somatic mutation no disease association


## Usage Statistics

- Prompt Tokens: 0

- Completion Tokens: 0

- Total Tokens: 0

- Estimated Cost: $0.0164


## Bibliometric Configuration

- Use Bibliometrics: True

- Author Impact Weight: 0.4

- Citation Impact Weight: 0.4

- Venue Impact Weight: 0.2
