## Supplementary material for "MENDELSEEK: An algorithm that predicts Mendelian Genes and elucidates what makes them special": GMPS.txt

# Claim: The gene GMPS is likely associated with at least one Mendelian disease.

**Status**: processed

**Overall Rating**: -1

**Explanation**:

No relevant papers were found that support or refute this claim.


## Relevant Papers


## Other Reviewed Papers


### The Burden of Candidate Pathogenic Variants for Kidney and Genitourinary Disorders Emerging From Exome Sequencing

**Why Not Relevant**: The provided paper content does not mention the gene GMPS (or obvious synonyms such as GMP synthase) nor present any data, variant counts, case reports, or mechanistic discussion specific to GMPS. The excerpt focuses on methods for identifying and curating genes associated with Mendelian kidney and genitourinary disorders, cohort composition, variant-filtering criteria (e.g., MAF < 1%, protein-truncating variants, use of HGMD/ClinVar), and the general problem of misclassified pathogenic variants in databases. While these methods and criteria are relevant to how one would assess whether GMPS is associated with a Mendelian disease, they do not provide direct or mechanistic evidence about GMPS itself. Because no verbatim sentences or paragraphs in the supplied content address GMPS, there is no basis in this text to support or refute the claim that GMPS is likely associated with at least one Mendelian disease. To evaluate the claim, one would need either (a) the paper's full curated gene list (625 genes) to check whether GMPS was included, or (b) gene-specific evidence from OMIM/Orphanet/ClinVar/HGMD or case series describing pathogenic variants in GMPS and their inheritance/phenotype. Those data are not present in the provided excerpt.


[Read Paper](https://www.semanticscholar.org/paper/a75e297208c04992568e607d19aa9b9aaebd09d5)


### Whole-Exome Sequencing in Adults With Chronic Kidney Disease

**Why Not Relevant**: The provided paper excerpt does not mention the gene GMPS (guanosine monophosphate synthetase) anywhere in the text, nor does it present any results or variant-level data tying GMPS to a Mendelian disease. The content describes study rationale, methods (WES capture, variant annotation, curated gene lists including 287 OMIM 'nephropathy genes'), and analytical/validation procedures, but does not include the actual gene list (Supplement Table 3) or result tables showing which genes had pathogenic or likely pathogenic variants. Therefore, from the supplied text it is not possible to determine whether GMPS is included among the curated OMIM genes, whether any GMPS variants were detected, or whether GMPS has an established Mendelian disease association. The methodological excerpts do provide indirect, mechanistic context that whole-exome sequencing and curated OMIM lists can detect Mendelian gene–disease associations and that technical limitations (exon capture gaps) could lead to false negatives. However, because no direct mention or data concerning GMPS are present in the excerpt, the paper content as provided offers no evidence for or against the claim that GMPS is likely associated with at least one Mendelian disease. To evaluate the claim further, one would need either (a) the full Supplement Table 3 to see whether GMPS is listed among the OMIM nephropathy genes, (b) result tables or text describing detected pathogenic variants (including any in GMPS), or (c) external sources/databases (OMIM, ClinVar, HGMD, peer-reviewed publications) reporting GMPS disease associations.


[Read Paper](https://www.semanticscholar.org/paper/a29bd57964f68ac5f8e1abcafe1112ba488f53f5)


### From case studies to community knowledge base: MSeqDR provides a platform for the curation and genomic analysis of mitochondrial diseases

**Why Not Relevant**: The provided paper is a description of the MSeqDR resource and its role in aggregating, curating, and sharing mitochondrial disease genomic data. It does not mention the gene GMPS nor present any experimental, clinical, or genetic evidence linking GMPS to a Mendelian disease. All relevant excerpts are contextual about databases, curation, and the capacity to add novel disease genes (e.g., PMPCA), but none constitute direct or mechanistic evidence for GMPS involvement in a Mendelian disorder. Therefore the paper is minimally relevant to the claim. The absence of GMPS in this text neither supports nor refutes the claim; it simply does not address it. Because the developer instructions requested a detailed explanation when relevance < 0.1, this statement documents that gap: the paper's focus is on mitochondrial disease data infrastructure and curation processes rather than on presenting new gene–disease associations for GMPS.


[Read Paper](https://www.semanticscholar.org/paper/6f85c595adaca6963c30fea73bc37266fa78be56)


### Next-generation whole-exome sequencing contribution to identification of rare autosomal recessive diseases

**Why Not Relevant**: The paper excerpt contains only high-level background about rare diseases and a methodological statement that whole-exome sequencing has led to discovery of novel mutations in autosomal recessive disease genes. It does not mention the gene GMPS, does not report any variants, phenotype correlations, segregation data, functional assays, or statistical evidence linking GMPS to a Mendelian phenotype. Mechanistic details that could support plausibility for GMPS (e.g., biochemical function of GMPS, pathways, animal models, or cellular phenotypes) are absent. Because the excerpt lacks any direct or gene-specific mechanistic evidence, it cannot be used to support or refute the claim that GMPS is likely associated with at least one Mendelian disease. The only value of the excerpt is as general context: it indicates that WES is a technique that can discover Mendelian disease genes, but that general statement does not provide evidence about GMPS itself.


[Read Paper](https://www.semanticscholar.org/paper/01c6f2a79bacd3813dfcd91d21b2bf7bc68d7e62)


### Gene discovery and cellular modelling of rare autosomal recessive neurodevelopmental conditions

**Why Not Relevant**: This paper does not provide direct or mechanistic evidence relevant to the claim that GMPS is likely associated with at least one Mendelian disease. The manuscript excerpt describes a family-based whole exome sequencing study of neurodevelopmental disorders and reports pathogenic or candidate variants in HERC2, TUBA1A, KLHL7 and MAL, including functional characterization for HERC2, KLHL7 and MAL. Nowhere in the provided text is the gene GMPS mentioned, nor are any variants, functional studies, segregation data, or phenotype correlations relating to GMPS reported. Therefore there is no direct evidence (e.g., reported GMPS variants segregating with disease, statistical association, or case reports) and no mechanistic evidence (e.g., functional assays implicating GMPS in disease-relevant biology) in this paper supporting the claim. Limitations and caveats: absence of any mention of GMPS in this paper is not proof that GMPS is not associated with a Mendelian disease — it only shows that this particular study did not identify or study GMPS. The lack of evidence here cannot be used to refute the claim; it only indicates the paper is not a source of supporting evidence for GMPS-disease association. For a conclusion about GMPS one would need to examine other genetic studies, case reports, databases (e.g., ClinVar, OMIM), or functional studies specifically involving GMPS.


[Read Paper](https://www.semanticscholar.org/paper/1916e63deed84eba433676327e1d8b5a8d009cf7)


### The simplest explanation does not have to be preferred: co-occurrence of pathogenic variants in cancer-predisposing genes

**Why Not Relevant**: The provided paper content is a single sentence fragment discussing the frequency and cancer phenotypes associated with "MINAS," a review of recent literature, and checking UK 10,000 Genomes Project data in participants with tumor predisposition syndromes or multiple primary tumors, plus whether phenotypes are additive or synergistic. It contains no mention of the gene GMPS, no data or conclusions about GMPS, and no mechanistic discussion involving GMPS. Therefore it provides neither direct evidence (e.g., variant-disease associations, case reports, segregation, allele frequencies, pathogenicity assessments) nor mechanistic evidence (e.g., functional studies, biochemical pathways, animal models) relevant to the claim that GMPS is likely associated with at least one Mendelian disease. The fragment also lacks methodological or results details, sample sizes, statistical measures, or contextual identifiers (page numbers, figures, or tables) that could be used to infer relevance. If additional sections of the paper that explicitly mention GMPS are available, those should be provided for proper evaluation.


[Read Paper](https://www.semanticscholar.org/paper/a1372a664bd2c33a0bda638975a8b26d112457fc)


## Search Queries Used

- GMPS GMP synthetase mutation pathogenic variant Mendelian disease case report ClinVar OMIM

- GMPS whole exome sequencing whole genome sequencing rare disease gene discovery trio de novo autosomal recessive monogenic

- GMPS GMP synthetase knockout mouse zebrafish loss of function purine biosynthesis neurological developmental delay phenotype

- GMPS gene disease validity review ClinGen OMIM ClinVar gnomAD loss of function intolerance constraint pLI essential gene embryonic lethal


## Bibliometric Configuration

- Use Bibliometrics: True

- Author Impact Weight: 0.4

- Citation Impact Weight: 0.4

- Venue Impact Weight: 0.2
