## Supplementary material for "MENDELSEEK: An algorithm that predicts Mendelian Genes and elucidates what makes them special": GNAZ.txt

# Claim: The gene GNAZ is likely associated with at least one Mendelian disease.

**Status**: processed

**Overall Rating**: -1

**Explanation**:

No relevant papers were found that support or refute this claim.


## Relevant Papers


## Other Reviewed Papers


### Phenotypes of a female patient with novel de novo frameshift ARX variant identified by whole-exome sequencing: a case report

**Why Not Relevant**: The provided paper discusses a case report and genetic analysis of a novel de novo frameshift variant in the ARX gene and its phenotypic consequences. Nowhere in the provided content is the gene GNAZ mentioned, nor are there data, analyses, or mechanistic discussions that involve GNAZ. Therefore this paper provides no direct genetic evidence (e.g., reported pathogenic variants in GNAZ, segregation data, case series) nor mechanistic evidence (e.g., functional studies, protein interactions implicating GNAZ in Mendelian disease) relevant to the claim that GNAZ is likely associated with a Mendelian disorder. The absence of mention does not prove GNAZ is or is not disease-associated, but it means this paper cannot be used as supporting or refuting evidence for the claim. For evaluation of the claim about GNAZ, literature, database (OMIM, ClinVar), or experimental studies specifically involving GNAZ should be consulted.


[Read Paper](https://www.semanticscholar.org/paper/c809ee0f8458dfa0baefd3da4fbd76dfef205dc0)


### Combined exome and whole transcriptome sequencing identifies a de novo intronic SRCAP variant causing DEHMBA syndrome with severe sleep disorder.

**Why Not Relevant**: The provided paper content does not mention the gene GNAZ, nor does it present any data, results, or discussion that link GNAZ to a Mendelian disease. The single-sentence summary states: "This study demonstrated the efficacy of a combined ES-WTS approach in solving undiagnosed cases and speculated that sleep respiratory disorder may be an underdiagnosed complication of DEHMBA syndrome." This describes (1) a methodological outcome (efficacy of combined exome sequencing and whole-transcriptome sequencing) and (2) a clinical speculation about DEHMBA syndrome and sleep respiratory disorder. Neither element provides: identification of GNAZ variants, segregation data, functional assays, statistical association, nor mechanistic pathways implicating GNAZ in disease. Evidence that would be directly relevant to the claim (for example: detection of pathogenic GNAZ variants in affected individuals, segregation of such variants with disease, experimental demonstration that GNAZ alteration causes relevant pathology, or statistical enrichment analyses implicating GNAZ) is absent from the provided content. Additionally, because the excerpt is a short summary, it lacks methods, results, variant-level details, or gene lists that might permit indirect inference. While DEHMBA syndrome might itself be Mendelian, the summary does not name causal genes, so no inference connecting GNAZ to DEHMBA (or to any Mendelian phenotype) can be made. For these reasons the content is not relevant to assessing the claim that "The gene GNAZ is likely associated with at least one Mendelian disease."


[Read Paper](https://www.semanticscholar.org/paper/22ade7a766f116133e03729baaf0b3b717e6cd33)


### De novo HDAC3 variants leading to epigenetic machinery dysfunction are associated with a neurodevelopmental disorder

**Why Not Relevant**: The provided paper content exclusively addresses heterozygous de novo variants in the HDAC3 gene and their association with a novel neurodevelopmental syndrome. It describes specific HDAC3 missense variants (D93N, A110T, P201S, L266S, G267S, and R359C), functional assays (HDAC activity, western blots), proteomic/co-immunoprecipitation results (altered interactions with CoREST and NCoR), and altered nuclear localization for certain HDAC3 variants. Nowhere in the supplied content is the gene GNAZ mentioned, nor are GNAZ protein function, variants, or any data pertaining to GNAZ discussed. Because the claim concerns whether GNAZ is likely associated with at least one Mendelian disease, and this paper contains no direct observations, variant data, mechanistic links, literature citations, or incidental findings about GNAZ, it provides neither direct nor mechanistic evidence for or against the claim. In short: absence of any reference to GNAZ in the paper means the paper cannot inform the claim. If you want assessment of GNAZ specifically, I can (a) search for or analyze publications that mention GNAZ and disease associations, (b) examine human genetic resources (OMIM, ClinVar, gnomAD, DECIPHER) for evidence of pathogenic variants in GNAZ, or (c) evaluate mechanistic plausibility based on GNAZ biology—please indicate which you prefer.


[Read Paper](https://www.semanticscholar.org/paper/524b09f9e39f306b6871f816044577dccb53abd6)


### Deciphering the Non-Coding Genome in Autism Spectrum Disorders (ASD): A Study of De Novo and Rare Inherited Variants through Targeted Sequencing in Regulatory Regions

**Why Not Relevant**: The provided paper content does not mention GNAZ at all nor does it report associations between specific genes and Mendelian diseases. The study explicitly focused on non-coding regulatory variation in ASD and intentionally excluded cases with coding loss-of-function mutations and large CNVs (the types of variation that often reveal Mendelian gene–disease relationships). Although the paper describes mechanistic approaches (Sei, PLAC-Seq, ATAC-seq, implication of CTCF dysregulation) that can link regulatory variants to target genes, the excerpted text does not list gene-level results or any evidence that GNAZ is implicated. Therefore there is no direct or gene-specific mechanistic evidence in the provided content to support the claim that GNAZ is likely associated with at least one Mendelian disease. It remains possible that the full paper or its supplementary tables name GNAZ among implicated genes, but that information is not present here; without such gene-level data, the claim cannot be evaluated or supported from the supplied text.


[Read Paper](https://www.semanticscholar.org/paper/649aeab19738420d26b974bc80eaee323f848ea6)


## Search Queries Used

- GNAZ pathogenic variant Mendelian disease clinical case report G alpha z guanine nucleotide binding protein alpha z subunit ClinVar OMIM

- GNAZ de novo variant neurodevelopmental disorder whole exome sequencing trio study rare variant burden novel variants

- GNAZ knockout mouse phenotype functional study G alpha z GPCR signaling neuronal synaptic transmission behavior electrophysiology

- GNAZ ClinGen gene curation Mendelian gene list diagnostic gene panel gnomAD constraint pLI loss of function intolerance


## Bibliometric Configuration

- Use Bibliometrics: True

- Author Impact Weight: 0.4

- Citation Impact Weight: 0.4

- Venue Impact Weight: 0.2
