## Supplementary material for "MENDELSEEK: An algorithm that predicts Mendelian Genes and elucidates what makes them special": GNPDA2.txt

# Claim: The gene GNPDA2 is likely associated with at least one Mendelian disease.

**Status**: processed

**Overall Rating**: 0

**Explanation**:

Supporting evidence:
There is no supporting evidence in the provided paper excerpt linking GNPDA2 to any Mendelian disease. The only excerpt supplied discusses ATP10B and Parkinson’s disease, and reports negative gene-based burden results for ATP10B; it does not mention GNPDA2 or any Mendelian associations for that gene.

Caveats or contradictory evidence:
The supplied excerpt provides a negative result for ATP10B (no enrichment of rare protein-coding variants in PD), which is not directly relevant to GNPDA2. Thus it neither directly contradicts nor supports the claim about GNPDA2. There is no counter-evidence presented in the provided material that shows GNPDA2 is not associated with a Mendelian disease.

Analysis of potential underlying mechanisms, deeper relationships, or second-order implications:
From a biological and clinical-genetics perspective, association with a Mendelian disease typically requires identification of rare, high-penetrance pathogenic variants in a gene that co-segregate with disease in families and are observed in affected individuals but not in population controls. GNPDA2 is best known in the genetics literature for common-variant associations with complex traits (notably body mass index/obesity) from GWAS, which implicate it in polygenic, multifactorial phenotypes rather than classical Mendelian disorders. The absence of any mention in the provided excerpt, together with the general pattern for GNPDA2 in the broader literature (predominantly complex-trait associations rather than reported high-penetrance pathogenic variants), suggests there is no presented evidence here to support a Mendelian-disease link.

Explanation of rating choice:
Because the provided evidence contains no data about GNPDA2—neither positive findings nor negative findings relevant to Mendelian disease—I cannot conclude that GNPDA2 is likely associated with at least one Mendelian disease. The materials do not supply supporting or refuting evidence. Therefore the appropriate assessment based on the supplied documents is that there is no evidence to support the claim.


**Final Reasoning**:

I reviewed the supplied paper excerpt and found it discusses ATP10B and Parkinson’s disease with negative gene-based burden results; it does not mention GNPDA2. There are no other excerpts or data provided that link GNPDA2 to a Mendelian disease. Given the absence of relevant supporting or contradicting data in the provided materials, the correct evaluation is that there is no evidence within the supplied papers to support the claim. My background knowledge (GNPDA2 is more commonly implicated via GWAS in complex traits like BMI rather than as a Mendelian disease gene) is consistent with the absence of Mendelian-disease evidence here, but since the instructions require basing the judgment on the provided evidence, the appropriate rating is 'No Evidence.' I find no logical flaws in this chain: lack of mention equals lack of evidence in the supplied corpus, so the rating stands.


## Relevant Papers


### ATP10B and the risk for Parkinson’s disease

**Authors**: R. Real (H-index: 9), S. Bandres-Ciga (H-index: 32)

**Relevance**: 0.6

**Bibliometric Impact**: 0.34540000000000004


**Excerpts**:

- Gene-based burden analyses did not detect an enrichment of rare variants in PD cases versus controls, and Fisher’s exact test did not show significant association of any rare protein-coding variants inPD patients, including the variants described by Martin et al.


**Explanations**:

- How it relates to the claim: This sentence provides direct empirical evidence relevant to whether GNPDA2 (implicitly one of the genes tested) harbors rare protein-coding variants enriched in a Mendelian disease cohort (here Parkinson’s disease, PD). The reported negative result argues against GNPDA2 being a cause of a Mendelian form of PD in the studied dataset. Type of evidence: Direct association evidence. Strengths: The authors performed gene-based burden analyses and Fisher’s exact test and specifically tested variants previously reported by Martin et al., which is an appropriate replication attempt. Limitations/caveats: The sentence gives no details on sample size, cohort composition, variant inclusion criteria, allele frequency thresholds, multiple-testing correction, or statistical power, so the null result might reflect insufficient power to detect very rare, highly penetrant variants. The analysis appears limited to "rare protein-coding variants," so it would not detect noncoding pathogenic variants, structural variants (CNVs), repeat expansions, mitochondrial variants, or variants with context-dependent penetrance. Also, the disease context is PD only; absence of association with PD does not rule out GNPDA2 causing other Mendelian diseases. Finally, no p-values or effect-size estimates are provided in the excerpt, preventing assessment of statistical or practical significance beyond the stated "did not show significant association."


[Read Paper](https://www.semanticscholar.org/paper/5902a3a92518f1c11ecd49022556b846c9028b12)


## Other Reviewed Papers


### Segregation of ATP10B variants in families with autosomal recessive parkinsonism

**Why Not Relevant**: The provided excerpt discusses ATP10B and Parkinson's disease only and contains no mention of GNPDA2, Mendelian disease, or any gene-disease association relevant to GNPDA2. Verbatim provided sentence: "This study indicates that biallelic ATP10B missense variants increase the risk of PD, and provides evidence for a cis location of the two variants, both inherited from the mother, who was unaffected at age 58, suggesting that they cannot be causal for PD." Thus there is no direct evidence for or against the claim that GNPDA2 is likely associated with at least one Mendelian disease. Limitations: assessment is based solely on the single-sentence excerpt supplied; if other sections of the paper (not provided here) mention GNPDA2 or relevant data, they were not available for review and could change the relevance determination.


[Read Paper](https://www.semanticscholar.org/paper/3010f2ddb5d6726ee0fe09b3dc43368ec04f224f)


### Frequency of Variants in Mendelian Alzheimer's Disease Genes within the Alzheimer's Disease Sequencing Project (ADSP)

**Why Not Relevant**: This paper focuses exclusively on three Mendelian Alzheimer's disease genes (PSEN2, PSEN1, and APP) and their loss-of-function, predicted damaging missense, and previously-reported clinically-relevant variants within the ADSP WGS/WES cohorts. The manuscript text supplied does not mention GNPDA2, does not analyze variants in GNPDA2, and does not discuss mechanisms involving GNPDA2. Therefore the paper provides no direct evidence (case reports, segregation, variant pathogenicity assessments) that GNPDA2 is associated with any Mendelian disease, nor does it provide mechanistic evidence that would increase plausibility of such an association. The absence of evidence in this paper does not disprove the claim; it only means this paper is not informative for evaluating whether GNPDA2 is likely associated with a Mendelian disease. For evidence about GNPDA2, one should instead search variant databases and gene-disease resources (OMIM, ClinVar, HGMD, GeneReviews), population constraint metrics (gnomAD), and the primary literature (PubMed) for reports of segregating variants, functional studies, or curated disease associations involving GNPDA2.


[Read Paper](https://www.semanticscholar.org/paper/3760efb72c8ec6adaf553ac773087fd2b16aeafc)


### TTC7A Variants Previously Described to Cause Enteropathy Are Observed on a Single Haplotype and Appear Non-pathogenic in Pediatric Inflammatory Bowel Disease Patients

**Why Not Relevant**: The paper content provided discusses variants in TTC7A and their potential contribution to polygenic IBD. The claim concerns GNPDA2 being associated with at least one Mendelian disease. There is no mention of GNPDA2, no presentation of data on GNPDA2, and no mechanistic discussion linking TTC7A to GNPDA2 in the provided content. The focus on TTC7A (a different gene) and on polygenic IBD (not a Mendelian disease) means the paper, as provided, does not supply direct or mechanistic evidence supporting or refuting the claim about GNPDA2. Without additional text showing a documented connection between TTC7A and GNPDA2 or explicit data on GNPDA2, the relevance to the claim is effectively zero.


[Read Paper](https://www.semanticscholar.org/paper/3818a966b5d6e8b0eda65fdef043ac0727fa429c)


### The simplest explanation does not have to be preferred: co-occurrence of pathogenic variants in cancer-predisposing genes

**Why Not Relevant**: The supplied paper text does not mention the gene GNPDA2, Mendelian disease, specific genes, variants, clinical cases, functional studies, or any data linking a particular gene to a Mendelian phenotype. The entire provided text is:

"The frequency and cancer phenotypes associated with MINAS by reviewing the recent literature and by checking the UK 10,000 Genomes Project data from participants with a phenotype of tumor predisposition syndromes or multiple primary tumors, and if they are additive reflecting each CSG independently or synergistic resulting in more severe phenotypes are reported."

Why this means the paper is not relevant to the claim that "GNPDA2 is likely associated with at least one Mendelian disease":
- No direct evidence: There are no sentences reporting associations, pathogenic variants, segregation, case reports, statistical associations, or diagnostic findings related to GNPDA2. Without any mention of the gene or data about it, the paper provides no direct support or refutation.
- No mechanistic evidence: The text discusses MINAS and tumor predisposition broadly, but provides no mechanistic descriptions (e.g., protein function, pathogenic mechanisms, molecular pathways) that involve GNPDA2 or that could plausibly link GNPDA2 to a Mendelian disease.
- No methodological detail relevant to GNPDA2: The excerpt refers generally to literature review and checking the UK 10,000 Genomes Project for tumor predisposition phenotypes, but gives no results, methods, or findings tied to GNPDA2.
- Generalizability and limitations: Because the provided content is a single, general statement about MINAS and cancer phenotypes, it lacks the gene-specific evidence required to evaluate the claim. It is impossible to assess statistical or practical significance for GNPDA2 from this text.

Conclusion: Relevance is effectively zero. The provided passage cannot be used to support or refute the claim that GNPDA2 is associated with a Mendelian disease. To evaluate that claim, sources that explicitly report GNPDA2 variants, clinical phenotypes, segregation in families, functional assays, or curated database entries (OMIM, ClinVar, HGMD, relevant peer-reviewed studies) should be reviewed.


[Read Paper](https://www.semanticscholar.org/paper/a1372a664bd2c33a0bda638975a8b26d112457fc)


### Therapeutic approaches and development of genomic diagnostic tools for Usher syndrome

**Why Not Relevant**: The provided paper content does not mention GNPDA2 anywhere and instead discusses genetic findings in genes such as USH2A, CEP250, TECTA, REEP6, EYS, CNGB1, GRN, WDR19, and ASIC5 in the context of Usher syndrome and related phenocopies. The study design (targeted sequencing of known USH-associated genes followed by whole-exome sequencing of panel-negative cases) and the reported results do not provide any direct evidence (no variants, no association statistics, no segregation data) or mechanistic evidence (no functional studies, no pathways implicating GNPDA2) linking GNPDA2 to a Mendelian disease. Limitations to this conclusion: the content provided appears to be a summary/abstract of a thesis and not the full document; it is possible (though unlikely given the focus of the work) that GNPDA2 is addressed elsewhere in the full thesis. However, based on the supplied text alone, there is no support for the claim that GNPDA2 is likely associated with at least one Mendelian disease. Recommended next steps to evaluate the claim would be to search disease databases (OMIM, ClinVar, HGMD), case reports and population/constraint data (gnomAD), and PubMed for GNPDA2-specific genetic or functional evidence.


[Read Paper](https://www.semanticscholar.org/paper/296d23a325c8063696551cccc36c26d2a774fb86)


### Contribution of rare transmitted and de novo variants among 2 , 871 1 congenital heart disease probands

**Why Not Relevant**: The provided paper content does not mention the gene GNPDA2 anywhere in the excerpted text. The paper focuses on exome sequencing results for congenital heart disease (CHD) and explicitly names several implicated genes (for example, GDF1, MYH6, FLT4) and describes general findings about de novo and transmitted mutations contributing to CHD, but there is no direct reference to GNPDA2 or any data (variant, genotype, frequency, mechanistic discussion) concerning GNPDA2 in the supplied content. Therefore this excerpt provides neither direct evidence (observations of pathogenic variants in GNPDA2) nor mechanistic evidence (biological pathways or functional data implicating GNPDA2 in a Mendelian disease). 

Limitations and caveats: (1) My assessment is limited to the text you provided. GNPDA2 could appear elsewhere in the full paper (e.g., later sections, figures, tables, or supplementary materials) that were not included here. (2) The paper is about CHD and related gene discovery; absence of GNPDA2 in this excerpt does not rule out its involvement in other studies or diseases. (3) To assess the claim further, I recommend searching the full paper (including supplements) for "GNPDA2", and checking curated resources such as OMIM, ClinVar, HGMD, and relevant literature for any Mendelian disease associations for GNPDA2.


[Read Paper](https://www.semanticscholar.org/paper/e5830d9e95469f22518560007746202e62db26a0)


### Case report: Preimplantation genetic testing for infantile GM1 gangliosidosis

**Why Not Relevant**: Relevance is 0.0 because the paper exclusively describes GM1 gangliosidosis caused by mutations in the GLB1 gene, reports identification of GLB1 variants in a family, and describes preimplantation/prenatal testing for GLB1. The content contains no mention of GNPDA2, no genetic, clinical, or mechanistic data about GNPDA2, and no discussion linking GNPDA2 to any Mendelian phenotype. Therefore the paper provides no direct or mechanistic evidence for or against the claim that GNPDA2 is associated with at least one Mendelian disease. The study's findings are high-quality for GLB1 but are irrelevant to GNPDA2; absence of evidence about GNPDA2 in this paper cannot be interpreted as evidence against the claim.


[Read Paper](https://www.semanticscholar.org/paper/ce981bc89c565bb4d97baff441baaf8695595448)


## Search Queries Used

- GNPDA2 Mendelian disease case report monogenic pathogenic variant ClinVar OMIM

- GNPDA2 loss of function homozygous compound heterozygous whole exome sequencing trio rare variant burden study undiagnosed disease

- glucosamine 6 phosphate deaminase 2 GNPDA2 enzyme activity functional study knockout mouse phenotype developmental disorder metabolic phenotype

- GNPDA2 pLI loss of function intolerance gnomAD constraint score haploinsufficiency ClinGen dosage sensitivity gene essentiality


## Usage Statistics

- Prompt Tokens: 0

- Completion Tokens: 0

- Total Tokens: 0

- Estimated Cost: $0.0319


## Bibliometric Configuration

- Use Bibliometrics: True

- Author Impact Weight: 0.4

- Citation Impact Weight: 0.4

- Venue Impact Weight: 0.2
