## Supplementary material for "MENDELSEEK: An algorithm that predicts Mendelian Genes and elucidates what makes them special": GNPTG.txt

# Claim: The gene GNPTG is likely associated with at least one Mendelian disease.

**Status**: processed

**Overall Rating**: 5

**Explanation**:

Supporting evidence.
The provided primary review explicitly states that mutations in GNPTG cause mucolipidosis (ML) type III gamma (and related ML types), which are autosomal recessive lysosomal storage disorders. The GNPTG gene encodes the γ-subunit of N-acetylglucosamine-1-phosphotransferase (GlcNAc-1-phosphotransferase), a key enzyme for generation of mannose-6-phosphate targeting signals on lysosomal enzymes. Defective activity of this enzyme causes missorting of lysosomal enzymes and lysosomal accumulation of non-degradable macromolecules leading to characteristic clinical phenotypes (e.g., MLIII gamma). The review reports a substantial catalog of disease-associated variation (50 mutations in GNPTG reported, including novel variants) and a literature-wide set of affected individuals for MLII/MLIII, supporting that GNPTG harbors pathogenic variants that produce a Mendelian (autosomal recessive) disease.

Caveats or contradictory evidence.
A second paper documents an unusual phenomenon in two MLIII gamma siblings homozygous for nonsense GNPTG mutations: in vivo transcript-level ‘correction’ (mRNA editing) to wild-type sequence was observed, suggesting that transcript analysis (cDNA) may sometimes be misleading and that genomic DNA (gDNA) analysis is required for reliable genetic diagnosis. This finding complicates molecular diagnostics but does not contradict that GNPTG variants cause disease; instead it warns about potential false negatives if only RNA is assayed. No evidence in the excerpts refutes a causal role for GNPTG in a Mendelian disorder.

Analysis of mechanisms, deeper relationships, and implications.
The molecular mechanism is consistent with a Mendelian loss-of-function disease model. GNPTG encodes an obligate subunit of the GlcNAc-1-phosphotransferase complex required for mannose-6-phosphate tagging of lysosomal hydrolases. Biallelic loss-of-function (or other deleterious) variants would be expected to impair lysosomal enzyme targeting broadly, producing a lysosomal storage phenotype across many cell types — exactly what is observed in mucolipidosis. The autosomal recessive inheritance is mechanistically plausible because partial residual activity (e.g., heterozygotes) is typically sufficient to avoid severe phenotypes, while biallelic damaging variants lead to disease. The documented genotype–phenotype correlations noted in the review further strengthen causality. The transcript-editing observation has second-order implications for diagnostics and variant interpretation (necessitating gDNA testing and careful functional validation), but does not negate pathogenicity.

Which rating is most appropriate and why.
Taken together, the evidence is direct and robust: peer-reviewed literature explicitly attributes a Mendelian (autosomal recessive) lysosomal storage disorder (mucolipidosis III gamma) to mutations in GNPTG, with numerous reported pathogenic variants and mechanistic plausibility. The isolated caveat about transcript-level editing affects diagnostic practice but does not undermine the causal relationship. Therefore the appropriate rating is "Highly Supported."


**Final Reasoning**:

The core claim is that GNPTG is likely associated with at least one Mendelian disease. The reviewed literature explicitly states that mutations in GNPTG cause mucolipidosis III gamma, an autosomal recessive lysosomal storage disorder, and documents dozens of GNPTG variants reported in affected individuals. Mechanistically, loss of GNPTG function disrupts GlcNAc-1-phosphotransferase activity and lysosomal enzyme targeting, producing the expected disease phenotype. The only substantive caveat is that mRNA editing can sometimes mask pathogenic variants at the transcript level, which complicates molecular detection but does not refute causation. Given direct attribution in multiple publications, a clear mechanistic pathway, and reported pathogenic variants, the preponderance of evidence strongly supports the claim. I therefore reaffirm the rating "Highly Supported."


## Relevant Papers


### The lysosomal storage disorders mucolipidosis type II, type III alpha/beta, and type III gamma: Update on GNPTAB and GNPTG mutations

**Authors**: R. V. Velho (H-index: 16), S. Pohl (H-index: 21)

**Relevance**: 1.0

**Bibliometric Impact**: 0.29223999999999994


**Excerpts**:

- Mutations in the GNPTAB and GNPTG genes cause mucolipidosis (ML) type II, type III alpha/beta, and type III gamma, which are autosomal recessively inherited lysosomal storage disorders.

- GNPTAB and GNPTG encode the α/β‐precursor and the γ‐subunit of N‐acetylglucosamine (GlcNAc)‐1‐phosphotransferase, respectively, the key enzyme for the generation of mannose 6‐phosphate targeting signals on lysosomal enzymes.

- Defective GlcNAc‐1‐phosphotransferase results in missorting of lysosomal enzymes and accumulation of non‐degradable macromolecules in lysosomes, strongly impairing cellular function.

- MLII‐affected patients have coarse facial features, cessation of statural growth and neuromotor development, severe skeletal abnormalities, organomegaly, and cardiorespiratory insufficiency leading to death in early childhood.

- MLIII alpha/beta and MLIII gamma are attenuated forms of the disease.

- Since the identification of the GNPTAB and GNPTG genes, 564 individuals affected by MLII or MLIII have been described in the literature.

- In this report, we provide an overview on 258 and 50 mutations in GNPTAB and GNPTG, respectively, including 58 novel GNPTAB and seven novel GNPTG variants.

- Comprehensive functional studies of GNPTAB missense mutations did not only gain insights into the composition and function of the GlcNAc‐1‐phosphotransferase, but also helped to define genotype‐phenotype correlations to predict the clinical outcome in patients.


**Explanations**:

- Direct evidence: This sentence explicitly states that mutations in GNPTG cause mucolipidosis type III gamma and that these disorders are autosomal recessive. This is direct, clear evidence that GNPTG is associated with at least one Mendelian disease (MLIII gamma). Strength: unambiguous causal wording. Limitation: the sentence is a summary statement from the paper; the excerpt itself does not present primary molecular or statistical data here, but asserts established causality.

- Mechanistic evidence: This sentence identifies the molecular product of GNPTG (the γ‐subunit of GlcNAc‐1‐phosphotransferase) and places it in the enzymatic complex responsible for generating mannose 6‐phosphate targeting signals. This provides mechanistic plausibility linking GNPTG dysfunction to lysosomal enzyme targeting defects. Strength: specifies the protein role. Limitation: while it explains the protein's normal function, it does not by itself demonstrate that particular variants disrupt that function (functional assays would be needed).

- Mechanistic evidence: This sentence describes the cellular consequence of defective GlcNAc‐1‐phosphotransferase (missorting of lysosomal enzymes and accumulation of macromolecules) and links enzymatic defect to impaired cellular function. This mechanism explains how mutations in GNPTG (as part of the enzyme) can produce a lysosomal storage disease phenotype, strengthening causal inference. Limitation: it is a general mechanistic description and does not specify experimental details or whether these specific outcomes were observed in GNPTG-mutant cells in this report.

- Context / phenotype: This sentence details the severe clinical phenotype of MLII, providing context for the types of clinical manifestations expected when the pathway is disrupted. Its relevance to the claim is contextual—illustrates the Mendelian disease phenotype associated with defects in the same pathway. Limitation: this sentence describes MLII phenotypes (often GNPTAB-related) rather than MLIII gamma specifically, so it is indirect with respect to GNPTG.

- Context / direct linkage: This statement clarifies that MLIII gamma is an attenuated (less severe) form of the disease spectrum and, by naming MLIII gamma, connects the clinical subtype that the GNPTG gene is implicated in. Strength: supports that GNPTG-related disease exists within a recognized clinical category. Limitation: it does not by itself provide molecular evidence, but supports phenotype classification.

- Context / epidemiology: This sentence provides the number of affected individuals described in the literature since gene identification (564 individuals with MLII or MLIII), supporting that these are documented, diagnosed Mendelian conditions observed in multiple patients. Strength: shows breadth of clinical/variant observation. Limitation: aggregated number across GNPTAB and GNPTG; does not disaggregate how many of these are GNPTG-specific.

- Supportive context and caveat about functional data: This sentence reports that comprehensive functional studies were performed for GNPTAB missense mutations and that these studies helped define genotype–phenotype correlations. Strength: demonstrates that functional and genotype–phenotype work exists for the pathway, supporting mechanistic understanding and clinical prediction for related genes. Limitation: this excerpt specifically refers to GNPTAB missense mutations (α/β subunit) rather than GNPTG, so direct functional data for GNPTG in this sentence are not described; the depth of functional characterization for GNPTG variants is not specified here.


[Read Paper](https://www.semanticscholar.org/paper/db6447e58caf7f424316ed2870b9fabe2e63282b)


### Enigmatic in vivo GlcNAc-1-phosphotransferase (GNPTG) transcript correction to wild type in two mucolipidosis III gamma siblings homozygous for nonsense mutations

**Authors**: R. V. Velho (H-index: 16), I. Schwartz (H-index: 34)

**Relevance**: 0.4

**Bibliometric Impact**: 0.3207


**Excerpts**:

- The results suggest that mRNA editing can be more frequent than expected in monogenic disorders and that GNPTG analysis should be performed on gDNA.


**Explanations**:

- How it relates: This sentence links GNPTG to the context of monogenic (Mendelian) disorders by stating that mRNA editing may be common in such disorders and explicitly recommending GNPTG analysis on genomic DNA. It therefore provides indirect evidence relevant to the claim that GNPTG is associated with at least one Mendelian disease — if GNPTG sequence analysis is clinically recommended in the context of monogenic disorders, that implies relevance of GNPTG to those disorders.

Type of evidence: Mostly indirect/inferential (procedural recommendation) with mechanistic implication (mRNA editing can confound mRNA-based detection of pathogenic variants).

Strengths: The sentence explicitly connects GNPTG to diagnostic practice in monogenic disorders and raises a plausible mechanism (mRNA editing) that could affect variant detection, which supports the idea that GNPTG is examined in the context of Mendelian disease.

Limitations/caveats: The excerpt is a single, summary statement with no supporting data, no mention of specific Mendelian disease(s), no demonstration of causative GNPTG variants, no sample size, no statistical measures, and no methodological details. The recommendation to perform GNPTG analysis on gDNA could reflect caution to avoid mRNA-editing artifacts rather than proof that GNPTG causes a Mendelian disease. Therefore this is suggestive but not direct proof of association.


[Read Paper](https://www.semanticscholar.org/paper/44a0bb4904d558b004e7407bb646efa18b597e34)


## Other Reviewed Papers


### The Burden of Candidate Pathogenic Variants for Kidney and Genitourinary Disorders Emerging From Exome Sequencing

**Why Not Relevant**: The provided paper content contains no mention of the gene symbol "GNPTG" nor any data, results, or discussion specific to that gene. The manuscript excerpt describes methods for identifying and curating genes associated with Mendelian kidney and genitourinary disorders (databases used, cohort, filtering pipelines) and states that a curated list of 625 genes is available on the laboratory website, but the excerpt does not include the gene names themselves. Because no verbatim sentences or paragraphs in the provided content refer to GNPTG, there is no direct evidence (variant observations, frequency data, or reported disease associations) nor any mechanistic evidence (functional data, pathogenic mechanism, or genotype–phenotype description) pertaining to GNPTG in this text. If GNPTG appears on the authors' curated gene list hosted externally, that information is not present in the supplied excerpt and therefore cannot be used here to support the claim. To evaluate the claim that GNPTG is associated with at least one Mendelian disease, one would need to consult gene-disease resources (for example, OMIM, ClinVar, HGMD) or the authors' full gene list (laboratory website) and any specific statements or data about GNPTG; none of these are provided in the excerpt.


[Read Paper](https://www.semanticscholar.org/paper/a75e297208c04992568e607d19aa9b9aaebd09d5)


### Variants of Uncertain Significance and “Missing Pathogenicity”

**Why Not Relevant**: The provided paper content does not mention the gene GNPTG anywhere, and it focuses on analysis of a set of medically actionable genes (with particular emphasis on 30 cardiac genes from the ACMG 59-gene set) and on variant-interpretation issues in a biobank cohort. Because GNPTG is not referenced, there is no direct evidence (e.g., reported pathogenic variants, segregation data, clinical phenotypes) nor mechanistic data (e.g., functional assays, pathway links) relating GNPTG to any Mendelian disease in this text. General statements about variant prevalence and interpretation in other genes do not constitute gene-specific evidence for GNPTG. Therefore the content is not relevant to assessing the claim that GNPTG is likely associated with at least one Mendelian disease.


[Read Paper](https://www.semanticscholar.org/paper/a2df451199c905611c0053a2f7d8a6a4dfecc14b)


### Confirmation of a Causal Role for SHQ1 Variants in Early Infantile‐Onset Recessive Dystonia

**Why Not Relevant**: The provided paper content discusses genetics of dystonia and implicates genes such as EIF2AK2, PRKRA, and SHQ1 in translational-control–related mechanisms and autosomal recessive early-onset dystonia. Nowhere in the supplied text is the gene GNPTG mentioned, nor are there any data, variants, mechanistic descriptions, or citations relating GNPTG to a Mendelian disease. Therefore the excerpt contains no direct evidence (e.g., reported pathogenic variants, segregation data, clinical descriptions tied to GNPTG) nor mechanistic evidence (e.g., biochemical pathways, functional assays) that would support or refute the claim that GNPTG is likely associated with at least one Mendelian disease. Limitations: this assessment is restricted to the text you supplied; if GNPTG appears elsewhere in the full paper (outside the provided content), that material was not available for review. To evaluate the claim further, consult the full article text and external resources (OMIM, ClinVar, gnomAD, primary genetic studies) for any GNPTG-disease associations.


[Read Paper](https://www.semanticscholar.org/paper/86973ad706273ccdf837552de1288ee84b4e5aab)


### Identification and characterization of 30 novel pathogenic variations in 69 unrelated Indian patients with Mucolipidosis Type II and Type III

**Why Not Relevant**: The provided paper content consists of a single summary sentence describing the scope of a study on Mucolipidosis II/III in an Indian population. It does not mention the gene GNPTG, does not present any mutation findings, does not describe mechanisms, and contains no experimental or statistical results. Because the claim concerns whether GNPTG is likely associated with at least one Mendelian disease, and the supplied content contains no direct or mechanistic information about GNPTG, the content is effectively non-informative with respect to the claim. Any assertion that GNPTG is associated with a Mendelian disease would require additional text from the paper (e.g., specific gene mutation findings, tables, results, or discussion) or external sources; such information is not present in the provided content, so relevance is rated very low.


[Read Paper](https://www.semanticscholar.org/paper/1aa038cb55e53fbbdacf6cd3aa4e55ac627070f9)


### Incidence of Germline ATM Variants in a Consecutive Clinical Cohort of CLL Patients

**Why Not Relevant**: This paper does not provide any direct or mechanistic evidence relevant to the claim that GNPTG is likely associated with at least one Mendelian disease. The entire text of the provided manuscript focuses on the ATM gene and its germline/missense variants in chronic lymphocytic leukemia (CLL) patients, the frequency of ATM variants across hematologic malignancies, and correlations between ATM variants and CLL clinical/genetic features (for example, 11q deletion and TP53 status). There is no mention of the gene GNPTG, no discussion of GNPTG-related biology, no data bearing on GNPTG variant frequencies, and no mechanistic pathways linking ATM (or any other genes discussed) to GNPTG. Because the paper contains no statements about GNPTG, no verbatim excerpts can be provided that would support or refute the claim. Any inference about GNPTG from this paper would be purely speculative and unsupported by the data or text presented. To evaluate the claim about GNPTG, one must consult other sources (e.g., OMIM, ClinVar, gene-specific literature, or population/genetic studies that include GNPTG).


[Read Paper](https://www.semanticscholar.org/paper/146d680685af39f2208198ed081c4ae39fd4b22f)


### Identification and characterization of 30 novel pathogenic variations in 69 unrelated Indian patients with Mucolipidosis Type II and Type III

**Why Not Relevant**: The provided content is a single-summary sentence stating the study scope (Mucolipidosis II/III mutation spectrum in an Indian population) but contains no mention of specific genes (including GNPTG), no mutation findings, no mechanistic descriptions, and no data. While GNPTG is known from other sources to be associated with Mucolipidosis III (a Mendelian disorder), the excerpt supplied does not provide direct or mechanistic evidence relevant to the claim. Therefore the relevance of the provided paper content to the claim that 'GNPTG is likely associated with at least one Mendelian disease' is very low. To evaluate the claim properly would require the paper's specific results, named-gene mutation lists, pathogenicity assessments, or mechanistic discussion; none of that is present in the supplied text.


[Read Paper](https://www.semanticscholar.org/paper/7284417c4f8a4ed51ac613c7d6e955958184487d)


## Search Queries Used

- GNPTG mutations mucolipidosis III gamma case report clinical phenotype biallelic autosomal recessive GNPTG pathogenic variants

- N acetylglucosamine 1 phosphotransferase gamma subunit mannose 6 phosphate lysosomal enzyme trafficking GNPTG loss of function molecular mechanism functional assay

- GNPTG exome sequencing gene discovery Mendelian disease candidate gene pathogenic variants population frequency gnomAD variant of uncertain significance

- GNPTG OMIM mucolipidosis review systematic review genotype phenotype animal model knockout mouse gnptg lysosomal storage disease


## Usage Statistics

- Prompt Tokens: 0

- Completion Tokens: 0

- Total Tokens: 0

- Estimated Cost: $0.0307


## Bibliometric Configuration

- Use Bibliometrics: True

- Author Impact Weight: 0.4

- Citation Impact Weight: 0.4

- Venue Impact Weight: 0.2
