## Supplementary material for "MENDELSEEK: An algorithm that predicts Mendelian Genes and elucidates what makes them special": HAO1.txt

# Claim: The gene HAO1 is likely associated with at least one Mendelian disease.

**Status**: processed

**Overall Rating**: 4

**Explanation**:

Supporting evidence.
The provided paper excerpt explicitly describes a patient with "GO deficiency" (glycolate oxidase deficiency) and notes that the child has two unrelated autosomal recessive conditions. HAO1 encodes glycolate oxidase (GO), so a reported biochemical diagnosis of GO deficiency in a patient from consanguineous parents is consistent with a Mendelian (autosomal recessive) enzyme-deficiency disorder linked to HAO1. The co-occurrence of GO deficiency with other recessive disease(s) in a consanguineous family further increases the plausibility that GO deficiency in this case is due to a homozygous pathogenic variant rather than an acquired or multifactorial cause.

Caveats or contradictory evidence.
The excerpt is brief and does not present direct genetic evidence (e.g., identification of pathogenic HAO1 variants) in this patient; it reports a clinical/biochemical diagnosis of GO deficiency without showing the causative mutation(s). A single case report (or a single brief excerpt) provides limited evidence for a gene–disease relationship: it could reflect a rare biochemical abnormality that is not strictly Mendelian, a misattribution, or a secondary metabolic effect. The excerpt also mentions concurrent conditions (congenital hyperinsulinism and unexplained hyperoxaluria), which complicates phenotype attribution: some features could arise from other causes, and pleiotropy or blended phenotypes in consanguineous pedigrees can confound interpretation. Without broader genetic, segregation, or functional data, one cannot be certain that HAO1 variants are causative in a Mendelian sense.

Analysis of potential underlying mechanisms, deeper relationships, or second-order implications.
Biologically, HAO1 encodes glycolate oxidase, an enzyme that oxidizes glycolate to glyoxylate, a precursor of oxalate. Loss-of-function of GO would be expected to alter glycolate/glyoxylate/oxalate metabolism, producing biochemical signatures such as elevated glycolate and potentially altered oxalate levels; such a mechanism is compatible with an enzyme-deficiency Mendelian disorder inherited in an autosomal recessive fashion. This mechanistic link strengthens plausibility: the gene product has a clear, testable metabolic role tied to the reported biochemical phenotype. Also, consanguinity and the mention of "unrelated autosomal recessive conditions" in the same patient fit a model where a homozygous HAO1 variant could underlie GO deficiency. Second-order implications include that if HAO1 loss causes a Mendelian enzyme deficiency, then sequencing and functional assays should identify pathogenic loss-of-function variants and consistent biochemical profiles across patients, and HAO1 would be a potential target for therapies manipulating oxalate metabolism.

Explanation of chosen rating.
Given that the excerpt reports a clinically recognized GO deficiency—which corresponds directly to loss of function of the HAO1 product—and that the case is described in a consanguineous family with autosomal recessive conditions, the evidence supports a Mendelian association. However, the provided evidence is limited (no direct genetic confirmation or multiple independent reports in the excerpts). The biological mechanism linking HAO1 to glycolate oxidase activity and oxalate metabolism is well-established, which strongly favors a causal relationship if pathogenic variants are present. Balancing the direct mechanistic plausibility and the clinical report against the paucity of explicit genetic proof in the provided excerpt, the appropriate assessment is that HAO1 is likely associated with at least one Mendelian disease, while acknowledging the need for more direct genetic and segregation data to reach a higher level of support.


**Final Reasoning**:

The excerpt documents a patient with glycolate oxidase deficiency (GO deficiency) and explicitly frames this as an autosomal recessive condition occurring in a child of consanguineous parents, which is consistent with a Mendelian enzyme deficiency due to HAO1 disruption. The gene–product relationship (HAO1 encodes glycolate oxidase) provides a clear mechanistic rationale linking genotype to biochemical phenotype. The main limitation is that the excerpt does not include genetic confirmation (identified pathogenic HAO1 variants, segregation data, or functional validation). Given mechanistic plausibility plus a reported clinical case but limited direct genetic evidence in the materials provided, the best-supported rating is "Likely True": the balance of evidence and biological plausibility indicates HAO1 is likely associated with at least one Mendelian (autosomal recessive) disease, though stronger confirmation would require genetic and functional data from additional cases.


## Relevant Papers


### Glycolate oxidase deficiency in a patient with congenital hyperinsulinism and unexplained hyperoxaluria

**Authors**: O. Clifford-Mobley (H-index: 6), S. Senniappan (H-index: 20)

**Relevance**: 0.4

**Bibliometric Impact**: 0.24485714285714286


**Excerpts**:

- This baby girl was born at 39 weeks gestation to consanguineous Asian parents and has two unrelated autosomal recessive conditions, CHI and GO deficiency, and also hyperoxaluria of unknown aetiology.


**Explanations**:

- Relation to the claim: This sentence provides indirect mechanistic support for the claim that HAO1 is likely associated with at least one Mendelian disease. GO deficiency (glycolate oxidase deficiency) is named as one of the autosomal recessive conditions in the patient; because HAO1 encodes glycolate oxidase (GO), the clinical observation of a GO deficiency phenotype is mechanistically consistent with pathogenic variation in HAO1 causing a Mendelian (recessive) disease. Strength of evidence: mechanistic plausibility is present (phenotype consistent with loss of the GO enzyme). Limitations/caveats: the excerpt does not state that HAO1 was tested or found to be mutated in this patient, so there is no direct genetic evidence linking HAO1 to disease in this paper. The report is a single case, so no statistical evidence or segregation/functional data are provided. The presence of multiple autosomal recessive conditions in a consanguineous family complicates attribution of specific phenotypes to specific genes. The phrase "hyperoxaluria of unknown aetiology" further indicates incomplete understanding of the biochemical/genetic cause in this patient. Overall, the excerpt supports mechanistic plausibility but does not provide direct genetic proof that HAO1 is causal.


[Read Paper](https://www.semanticscholar.org/paper/a1d5ad11381ce7a1c519cadc8c5d3a64a870d13a)


## Search Queries Used

- HAO1 hydroxyacid oxidase 1 glycolate oxidase Mendelian disease case report biallelic homozygous compound heterozygous pathogenic variant ClinVar OMIM

- glycolate oxidase deficiency glycolic aciduria glycolate excretion HAO1 inborn error of metabolism autosomal recessive case report urinary glycolate oxalate

- HAO1 knockout mouse glycolate oxidase knockout rat phenotype oxalate glyoxylate metabolism primary hyperoxaluria therapeutic target antisense oligonucleotide RNA interference enzyme activity

- HAO1 loss of function variants gnomAD pLI human knockout homozygous loss of function population sequencing exome study ClinVar ClinGen Mendelian gene curation


## Usage Statistics

- Prompt Tokens: 0

- Completion Tokens: 0

- Total Tokens: 0

- Estimated Cost: $0.0097


## Bibliometric Configuration

- Use Bibliometrics: True

- Author Impact Weight: 0.4

- Citation Impact Weight: 0.4

- Venue Impact Weight: 0.2
