## Supplementary material for "MENDELSEEK: An algorithm that predicts Mendelian Genes and elucidates what makes them special": HAO2.txt

# Claim: The gene RBM14 is likely associated with at least one Mendelian disease.

**Status**: processed

**Overall Rating**: -1

**Explanation**:

No relevant papers were found that support or refute this claim.


## Relevant Papers


## Other Reviewed Papers


### Unmet Expectations: miR-34 Plays No Role in p53-Mediated Tumor Suppression In Vivo

**Why Not Relevant**: The provided paper content focuses entirely on p53 and the miR-34 family (miR-34a/b/c), their roles in p53-regulated pathways, and mouse knockout/functional studies of those miRNAs. There is no mention of the gene symbol RBM14, the name "RBM14", alternate spellings (e.g., "RBM-14"), or the protein name "RNA-binding motif protein 14" anywhere in the supplied text. The paper contains no human genetic data, no case reports of segregating variants, no gene-disease association analyses, and no discussion of Mendelian phenotypes or clinical genetics that could support or refute an association between RBM14 and any Mendelian disease. Methodologically, the work described is in vivo and in vitro functional analysis of miRNA and p53 biology in mice and cells, not a study of human Mendelian genetics; therefore it cannot provide direct genetic evidence (segregation, de novo occurrence, variant pathogenicity, population frequency evidence) or mechanistic evidence specifically implicating RBM14 in Mendelian disease. Absence of evidence in this excerpt does not demonstrate RBM14 is not disease-associated, but this paper does not contribute relevant evidence for the claim that RBM14 is likely associated with at least one Mendelian disease.


[Read Paper](https://www.semanticscholar.org/paper/64cede98bac79fad7a83cd1410c99f74f4e5583c)


### Detailed genetic and clinical analysis of a novel de novo variant in HPRT1: Case report of a female patient from Saudi Arabia with Lesch–Nyhan syndrome

**Why Not Relevant**: The paper exclusively studies HPRT1 and Lesch–Nyhan syndrome; RBM14 is not mentioned anywhere in the provided content. All genetic, mechanistic, and clinical evidence in the paper pertains to HPRT1 (including identification of a de novo frameshift variant and X-inactivation explaining disease manifestation in a female). Because the paper does not discuss, analyze, or provide data on RBM14, it offers no direct or mechanistic evidence that RBM14 is associated with any Mendelian disease. Therefore relevance to the claim about RBM14 is effectively zero.


[Read Paper](https://www.semanticscholar.org/paper/16daa186f81b3719f537f57e29a674411e64bb19)


### Case report: De novo mutation of a-galactosidase A in a female patient with end-stage renal disease: report of a case of late diagnosis of Anderson–Fabry disease

**Why Not Relevant**: The provided paper content is a case report focused entirely on Anderson–Fabry disease and pathogenic variants in the GLA (alpha-galactosidase A) gene. There is no mention of the gene RBM14, no analyses (genetic, functional, statistical, or mechanistic) involving RBM14, and no discussion that could be interpreted as linking RBM14 to any Mendelian phenotype. Therefore this paper provides no direct evidence (positive or negative) nor mechanistic rationale regarding the claim that RBM14 is likely associated with at least one Mendelian disease. Absence of mention in this single paper does not disprove an association; it merely means this document is irrelevant to evaluating the claim. To assess the claim about RBM14, one would need to consult other sources (e.g., genetic case reports implicating RBM14 variants, ClinVar/OMIM entries, cohort sequencing studies, functional assays, animal models, or constraint/variant burden analyses).


[Read Paper](https://www.semanticscholar.org/paper/a7c27f053151b6265c6c2b709b2f2085f8129f0e)


### Case Report: A de novo NLRP3 variant resulting in autoinflammatory disease in a Chinese newborn

**Why Not Relevant**: The provided paper content exclusively discusses a de novo variant in the NLRP3 gene and NLRP3-associated autoinflammatory disease (NLRP3-AID / CAPS). There is no mention of the gene RBM14 (nor any obvious synonyms or related proteins) in the background, results, or conclusions. Therefore the paper contains no direct evidence linking RBM14 to any Mendelian disease, nor does it provide mechanistic data that would plausibly implicate RBM14 by inference. Limitations: my judgment is based solely on the text supplied above; if the full paper contains other sections not included here (e.g., Supplementary Data, Results sections, or a gene panel listing) that mention RBM14, those were not available for review. Based on the provided content, however, this paper is not relevant to the claim that RBM14 is likely associated with at least one Mendelian disease.


[Read Paper](https://www.semanticscholar.org/paper/282bab9e5b483d880402d57adb139944db12b1a8)


### Tuberous sclerosis complex associated lymphangioleiomyomatosis caused by de novo mutation of TSC2 gene in Vietnam: A case report

**Why Not Relevant**: This paper does not provide any evidence that RBM14 is associated with a Mendelian disease. The manuscript describes a single patient with LAM/TSC-LAM and reports a de novo frameshift mutation in TSC2 (c.4504del (p.L1502Cfs*74)) as the likely genetic cause. RBM14 is not mentioned anywhere in the provided text, and no variants in RBM14 are reported or discussed. Mechanistically the paper links the phenotype to TSC1/TSC2 pathogenic variation—established Mendelian genes—rather than to RBM14. Limitations to note: the absence of RBM14 in this report is not proof that RBM14 cannot be involved in Mendelian disease elsewhere (absence of evidence is not evidence of absence). Also, this is a single-case report, so it cannot comprehensively rule out RBM14 involvement in other diseases or cohorts. However, for the specific claim that 'RBM14 is likely associated with at least one Mendelian disease,' this paper provides no supporting direct evidence, no mechanistic rationale involving RBM14, and thus is not relevant to substantiating that claim.


[Read Paper](https://www.semanticscholar.org/paper/6b7d5d519fb771d9bd7eb969a3b3847dcfdd8dad)


### Analysis of SAFB1 and SAFB2 Knockout Mice Reveals Non-Redundant Functions of the Two Estrogen Receptor Co-Repressor Paralogs.

**Why Not Relevant**: The paper exclusively studies the Scaffold Attachment Factor genes SAFB1 and SAFB2 in mouse models and their roles as estrogen receptor co-repressors and in mammary gland development. Nowhere in the provided content is the gene RBM14 mentioned, nor are there any human genetic data, pedigree analyses, variant pathogenicity assessments, or statements about Mendelian disease causation. Therefore the paper provides no direct evidence that RBM14 is associated with any Mendelian disease. Mechanistically, while the paper discusses RNA- and DNA-binding domains for SAFB proteins, this information pertains to SAFB1/2 and cannot be extrapolated to RBM14 without additional data showing functional or genetic linkage. Additional limitations that prevent relevance include: (1) experiments are in mouse knockouts of different genes, (2) phenotypes described are developmental and tissue-specific rather than human inherited disease descriptions, and (3) there is no genetic association or mutation data for RBM14. Because the paper does not mention RBM14 at all and contains no human Mendelian genetics evidence, its relevance to the claim is effectively zero.


[Read Paper](https://www.semanticscholar.org/paper/c0cfdff2a89eecaaeea6f7a39303451ac70c7bb7)


### Construction of phenotype and genotype database for Arabidopsis to contribute to research using diversity.

**Why Not Relevant**: I found no direct or indirect mention of the gene RBM14 in the supplied paper content. The claim asks whether RBM14 is likely associated with at least one Mendelian disease, which requires either (a) direct genetic/clinical evidence (human mutations in RBM14 segregating with a Mendelian phenotype, case reports, linkage/association in families) or (b) mechanistic evidence connecting RBM14 function to a well-established Mendelian disease pathway with supporting experiments. The provided text contains multiple topics (yeast replication origins and chromatin modifiers Abf1/Gcn5/Esa1, fission yeast Pol2-interacting factor and heterochromatin, gene body DNA methylation in Ciona, royalactin/Egfr in insects, Xist in mouse germ cells, NO/8-nitro-cGMP signaling, odorant-binding proteins in Drosophila, autophagy/TORC1/Atg proteins, Gtr1/Gtr2 interactions) but none mention RBM14 or a homolog by name, nor any human genetic or Mendelian-disease data. While some passages discuss transcription- or RNA-related factors (e.g., a Pol2-interacting factor), these are unnamed in the excerpt and cannot be attributed to RBM14 without sequence, orthology, or functional evidence. All experiments appear to be in non-human model organisms (yeast, Ciona, Drosophila, mouse in one Xist allele experiment), and no human clinical genetics or family segregation data are presented. Therefore the paper provides no direct evidence supporting or refuting an association between RBM14 and any Mendelian disease, and it provides no mechanistic evidence that implicates RBM14. Limitations: this assessment is based solely on the supplied text; if the full paper or supplemental materials (not provided here) include RBM14 or human genetic data, that material was not available for review. Given the absence of any mention or data about RBM14 in the provided content, the relevance to the claim is effectively zero.


[Read Paper](https://www.semanticscholar.org/paper/5b0fca04d97e6641aab5c5a51a09f80bfb63dc09)


### Expression of Huntingtons disease markers in eye tissues

**Why Not Relevant**: This document contains no mention of RBM14 and provides no data (genetic, experimental, clinical, or mechanistic) that links RBM14 to any Mendelian disease. The text focuses on Huntington disease (HTT), ATM, AAK1/SEL-5, BCAR1/p130Cas, a MommeD3 mutant (with its causative mutation not specified in this excerpt), and various developmental and cell-biological projects. There are no human genetic linkage/segregation analyses, no rare variant associations, no reported pathogenic RBM14 variants, no functional perturbation (knockout/knock-in) data of RBM14, and no mechanistic discussion implicating RBM14 in Mendelian inheritance or monogenic phenotypes. Because the paper neither names RBM14 nor presents any direct or mechanistic evidence relevant to the claim, its relevance to the claim “RBM14 is likely associated with at least one Mendelian disease” is effectively zero. Note: absence of evidence in this paper does not disprove the claim in general, but within the content provided there is no supportive or contradictory information concerning RBM14.


[Read Paper](https://www.semanticscholar.org/paper/20ae6edb20d81fe8df2617d06f8ba75e11d0ee30)


### 114 Involucrin deﬁciency results in decreased vitamin D receptor-mediated inﬂammation and Csnk1e isoform bias transcriptome reveals increased atherosclerosis markers and immune and barrier differences amongst subtypes

**Why Not Relevant**: The provided paper content does not mention RBM14, nor does it present human genetic-association, variant, clinical, or functional data for RBM14 that could support or refute a Mendelian-disease association. The manuscript excerpts focus on involucrin (IVL) biology and knockout mouse phenotypes, lncRNA expression in keratinocyte aging, SDR9C7 mutations in ichthyosis, antimicrobial peptides, transcriptomes of ichthyosis subtypes, and proteomics/lipidomics of photoaging. Genes explicitly named in the text include IVL, TSLP, VDR, Csnk1e, NORAD, MIAThad, PINT, PVT1, SDR9C7/Sdr9c7, FADS1/2, FAR2, FA2H, ELOVL3, HAO2, CLDN8, CLDN23, CDH10, SPRR1A/1B/2C/2G, PGLYRP3/4, LCE3D, VNN1, PCSK9, IL36A/B/G, HBEGF, SELE, ACE2, ApoE, Hmmr, and Mmp9. None of these sentences or paragraphs provide direct evidence (genetic linkage, segregation, pathogenic variants, case reports) or mechanistic evidence (gene function, molecular pathways implicating RBM14 in disease) relevant to the claim that RBM14 is associated with a Mendelian disease. Therefore, based on the supplied content, the paper is not relevant to evaluating the claim.


[Read Paper](https://www.semanticscholar.org/paper/6b2b0a699353892d34f2a9d1d5e1601df3f8b70c)


## Search Queries Used

- RBM14 mutation de novo variant patient case report mendelian disease

- RNA binding motif protein 14 RBM14 CoAA germline mutation hereditary disease familial syndrome

- RBM14 knockout mouse phenotype embryonic development loss of function haploinsufficiency functional study RNA binding transcription coactivator

- RBM14 gnomAD pLI loss of function intolerance constraint ClinVar pathogenic review Mendelian


## Bibliometric Configuration

- Use Bibliometrics: True

- Author Impact Weight: 0.4

- Citation Impact Weight: 0.4

- Venue Impact Weight: 0.2
