## Supplementary material for "MENDELSEEK: An algorithm that predicts Mendelian Genes and elucidates what makes them special": HBE1.txt

# Claim: The gene HBE1 is likely associated with at least one Mendelian disease.

**Status**: processed

**Overall Rating**: 0

**Explanation**:

Supporting evidence:
The provided excerpts discuss globin gene regulation (particularly gamma globin) and conditions that alter fetal hemoglobin (HPFH) or influence HbF levels in sickle cell disease. They note that NF-E1/YY1 represses embryonic (epsilon) and fetal (gamma) globin genes and describe a C→G substitution in the A gamma promoter that affects transcription factor binding and reactivates gamma expression. One excerpt mentions the epsilon gene in a list of genes assayed in a beta-globin cluster analysis. These points establish general biological context that globin-gene regulation and mutations can affect hemoglobin expression.

Caveats or contradictory evidence:
None of the excerpts present any direct data linking HBE1 (the epsilon globin gene) to a Mendelian disease. The first paper focuses on an A gamma (HBG1/HBG2) promoter variant and HPFH (hereditary persistence of fetal hemoglobin), not on HBE1 variants causing disease. The second paper discusses HbF levels in sickle cell disease and deletional HPFH in the beta-globin cluster; it does not attribute Mendelian disorders to HBE1. Thus the provided material contains no direct supportive evidence that HBE1 is associated with a Mendelian disorder, nor does it provide explicit contradictory evidence (e.g., statements that HBE1 has been shown to be benign).

Analysis of mechanisms and deeper relationships:
Biologically, globin genes frequently underlie Mendelian hemoglobinopathies (e.g., HBB mutations causing beta-thalassemia/sickle cell disease). It is therefore plausible in principle that variants in HBE1 could produce a Mendelian phenotype. However, HBE1 encodes an embryonic hemoglobin (epsilon) primarily expressed during early development; deleterious variants might have embryonic effects, be lethal, or have little clinical consequence in adults once expression ceases. The provided papers emphasize regulation of fetal (gamma) globin and HPFH mechanisms rather than pathogenic variants in embryonic genes. The mere fact that a gene is in the globin cluster and is a functional globin does not, by itself, constitute evidence that it is associated with a Mendelian disease.

Why the rating selected is most appropriate:
Given the contents of the supplied papers, there is no direct evidence supporting the claim that HBE1 is associated with at least one Mendelian disease. The excerpts do not report HBE1 mutations causing a hereditary disease, nor do they cite clinical phenotypes tied to HBE1 variation. While biological plausibility exists (globin genes often cause Mendelian hemoglobinopathies), the provided evidence does not demonstrate or document such an association for HBE1. Therefore the most accurate assessment based on the supplied material is that there is no evidence in these papers to support the claim.


**Final Reasoning**:

I re-read the essay for logical consistency. The papers provided focus on gamma globin regulation and HPFH and briefly reference the epsilon gene only in passing as part of assays; they do not present association data linking HBE1 to Mendelian disease. Although globin genes commonly underlie Mendelian disorders, and it is biologically plausible that HBE1 variants could be pathogenic, plausibility is not evidence. Because the instructions require basing the rating on the provided evidence, the appropriate conclusion is that the materials supply no evidence to support or refute the claim. Thus the rating 'No Evidence' accurately reflects the preponderance of the provided excerpts.


## Relevant Papers


### The -195C→G Substitution In Brazilian Hereditary Persistence of Fetal Hemoglobin Decreases NF-E1/YY1 Binding and Increases PAX1 Binding to the A Gamma Globin Promoter Region

**Authors**: F. M. Roversi (H-index: 9), F. Costa (H-index: 44)

**Relevance**: 0.15

**Bibliometric Impact**: 0.2120857142857143


**Excerpts**:

- In vitro primary human erythroblast cultures, derived from human CD34 + hematopoietic cells from 4 Brazilian ndHPFH type subjects and 4 control subjects, were proliferated and differentiated into late stage erythroblasts.

- NF-E1/YY1 is a transcription factor that represses embryonic (epsilon) and fetal (gamma) globin genes.

- Protein-DNA array and EMSA showed a decreased binding of NF-E1/YY1 in Brazilian ndHPFH nuclear extracts and ChIP analysis revealed diminished NF-E1/YY1 occupancy at the –195 A gamma globin promoter region of Brazilian ndHPFH.

- The consensus binding site for NF-E1/YY1 is a CCAN motif that is observed between the –195 and –192 position in the A gamma globin promoter region.

- The C→G substitution at –195 position may disrupt this DNA binding site, cause decreased NF-E1/YY1 interaction and probably allows the binding of PAX-1, a transcriptional activator with a paired box DNA-binding domain that has as a DNA binding core motif, the sequence TTCCGC.

- Our protein-DNA array and EMSA results showed an increased binding of PAX-1 in the Brazilian ndHPFH nuclear extracts and quantitative ChIP analysis with anti-PAX-1 antibody showed that PAX-1 binds to the –195 A gamma globin promoter region only in the presence of this C→G substitution.

- These results suggest that the –195 site (C→G) in the A gamma globin promoter region may decrease NF-E1/YY1 binding and increase PAX-1 binding in this DNA region, probably resulting in the reactivation of the A gamma globin gene.

- Hereditary persistence of fetal hemoglobin (HPFH) is a condition that prevents hemoglobin switching and the consequent silencing of the gamma globin genes, resulting in continued hemoglobin (Hb) F synthesis in adults.


**Explanations**:

- Methodological/contextual excerpt. This sentence describes sample source and scale (4 affected and 4 controls) and the in vitro erythroblast differentiation used to generate nuclear extracts. It is relevant because it frames the experimental basis for the molecular findings, but it does not provide any evidence that links HBE1 (epsilon) to a Mendelian disease. Limitation: small sample size and in vitro culture may not capture in vivo disease phenotypes.

- Mechanistic excerpt. This sentence explicitly names the embryonic (epsilon) globin among NF-E1/YY1 targets, directly connecting the transcription factor studied to HBE1 (epsilon). It provides mechanistic relevance because it indicates that the regulatory machinery investigated in this paper acts on HBE1. However, it does not present evidence that HBE1 itself is mutated in, causes, or is associated with any Mendelian disease—only that it is a regulatory target. Strength: direct statement of regulatory relationship. Limitation: statement of repression does not equate to disease association.

- Mechanistic/experimental excerpt. This describes observed changes in NF-E1/YY1 binding in the Brazilian ndHPFH samples using Protein-DNA array, EMSA, and ChIP. It shows that alterations in NF-E1/YY1 occupancy are associated with a pathological hemoglobin persistence phenotype (HPFH) related to gamma globin reactivation. Relevance to HBE1 is indirect: since NF-E1/YY1 also represses epsilon (HBE1), altered NF-E1/YY1 activity could conceivably affect epsilon expression, but the paper measures occupancy at the A gamma promoter, not at HBE1, and does not report effects on epsilon expression or disease caused by HBE1. Limitation: no direct measurement of HBE1 expression or HBE1 sequence variation; changes shown are at the gamma promoter.

- Mechanistic excerpt. This identifies the NF-E1/YY1 binding motif location in the A gamma promoter, providing molecular detail underlying the altered binding observed. It strengthens the mechanistic plausibility that promoter sequence changes can alter TF binding. Relevance to HBE1: indirect—describes motif at gamma promoter, not HBE1 promoter. Limitation: locus-specific detail for HBG (gamma) rather than HBE1.

- Mechanistic excerpt. This hypothesizes how the specific C→G mutation could disrupt NF-E1/YY1 binding and permit PAX-1 binding, with sequence-level detail. It demonstrates the authors' mechanistic model for gamma reactivation. Relevance to HBE1 is indirect—shows how TF binding at a promoter alters globin expression; since NF-E1/YY1 also represses epsilon, analogous promoter changes could affect HBE1, but the paper does not test this for HBE1. Limitation: speculative ('may disrupt', 'probably allows') and focused on A gamma promoter.

- Mechanistic/experimental excerpt. This provides experimental confirmation that PAX-1 binds the mutated A gamma promoter and that binding is mutation-dependent. It supports a mechanism for reactivation of gamma globin (HBG) in HPFH. Relevance to HBE1 is indirect and inferential: demonstrates that TF switching at globin promoters can change expression, but HBE1 is not directly assayed. Limitation: no data on HBE1 promoter occupancy or expression.

- Mechanistic/conclusion excerpt. The authors summarize their mechanistic interpretation linking the –195 mutation to altered TF binding and A gamma reactivation. It reinforces that the paper's findings are about gamma globin reactivation. Relevance to HBE1: conceptual (shared regulatory TF can act on epsilon), but there is no direct evidence here that HBE1 is involved in HPFH or any Mendelian disease. Limitation: conclusion specific to A gamma; HBE1 not experimentally addressed.

- Contextual excerpt. This defines HPFH and its effect on globin switching and adult HbF levels, framing the disease phenotype studied. It is relevant background but does not discuss HBE1 mutation or HBE1-linked Mendelian disease. Limitation: descriptive background only; does not provide data linking HBE1 to Mendelian disease.


[Read Paper](https://www.semanticscholar.org/paper/7b7e16b7112661cee0ec3f70b11f61b3238c5142)


### High Hemoglobin F in Sickle Cell Disease: Waning Protection with Age

**Authors**: Seda S. Tolu (H-index: 9), C. Minniti (H-index: 41)

**Relevance**: 0.15

**Bibliometric Impact**: 0.38016000000000005


**Excerpts**:

- Beta globin dosage genetic analysis was performed on a subset of patients (8/11) which used multiplex-polymerase chain amplification of fragments specific for portions of the epsilon, gammaG, gammaA, delta, and beta globin genes, which is commonly associated with deletional variant of HPFH (Quest Diagnostics, VA).

- SCD patients with HBS and deletional hereditary persistence of fetal hemoglobin (HPFH) have high levels of HbF (>30%) in a pancellular distribution, and absence of symptoms as suggested by prior studies.

- Available flow cytometry and genetic data revealed heterocellular distribution of HbF in 8 patients, and pancellular distribution in 2 patients: patient (#2) and patient (#10). Patient (#2) is positive for deletional HPFH with genetic confirmation of heterozygous positive deletion in the beta globin gene cluster.

- In our cohort, two patients with confirmed pancellular distribution of HbF, one of which with confirmed deletional HPFH, were not fully protected from the clinical complications of SCD, suggesting alternative pathogenic mechanisms for VOCs may be at play.


**Explanations**:

- Relation to claim: mechanistic/assay evidence. This sentence states that the diagnostic multiplex PCR assay targeted fragments specific to the epsilon, gammaG, gammaA, delta, and beta globin genes when assessing for deletional HPFH. Strength: it demonstrates that the epsilon gene region (commonly HBE1) is included in a clinical diagnostic panel used to detect deletions associated with HPFH, implying that structural variation involving the epsilon locus is considered relevant in detecting deletional variants of the beta-globin cluster. Limitations/caveats: the sentence does not name HBE1 explicitly, does not report any pathogenic variants in epsilon/HBE1, and does not demonstrate that variation in HBE1 causes a Mendelian disease by itself. It only shows that the assay interrogates that genomic region as part of a larger diagnostic target.

- Relation to claim: mechanistic/phenotype context. This sentence links deletional hereditary persistence of fetal hemoglobin (HPFH) — a Mendelian, genetically determined condition of the beta-globin cluster — to a recognizable phenotype (high HbF, pancellular distribution, typically asymptomatic). Strength: confirms HPFH is a Mendelian variant of the globin locus with clear clinical consequences and is caused by genetic alterations in the beta-globin gene cluster. Limitations/caveats: the sentence does not implicate any single gene (such as HBE1) within the cluster as causal; HPFH is often due to deletions spanning multiple globin genes or regulatory elements. Thus, it supports the plausibility that genes in the cluster can be involved in Mendelian traits, but does not specifically tie HBE1 to disease causation.

- Relation to claim: direct/diagnostic result for the cohort. These sentences report that genetic testing identified a heterozygous deletion in the beta globin gene cluster consistent with deletional HPFH in one patient, and that the genetic assay interrogated the epsilon gene region (see Methods excerpt). Strength: provides direct evidence that deletional events in the beta-globin cluster were detected in patient(s) in this cohort, confirming that cluster deletions are present in humans and are clinically detectable. Limitations/caveats: the report refers to a deletion in the beta globin gene cluster but does not specify which genes are included in the deletion (it does not state whether HBE1/epsilon is part of the deletion). Therefore it does not demonstrate that HBE1 variants are causative of a Mendelian disease on their own.

- Relation to claim: contextual/mechanistic limitation. This sentence shows that even when cluster deletions consistent with HPFH are present (a Mendelian condition affecting globin gene expression), the expected protective phenotype is not absolute in the context of sickle cell disease, implying phenotypic complexity for cluster variants. Strength: emphasizes that variants affecting the beta-globin cluster produce measurable clinical effects (i.e., HPFH is biologically meaningful). Limitation/caveat: it does not attribute disease causation to any particular gene within the cluster (such as HBE1) and instead highlights phenotypic variability and other pathogenic mechanisms; therefore it does not strengthen a specific link between HBE1 and a Mendelian disease.


[Read Paper](https://www.semanticscholar.org/paper/1034196c6e101fd31848074d92c0655c8fabc105)


## Other Reviewed Papers


### Targeted deletion of 5'HS2 of the murine beta-globin LCR reveals that it is not essential for proper regulation of the beta-globin locus.

**Why Not Relevant**: The paper reports mouse experimental work on the beta-globin locus control region and the effect of deleting or disrupting 5'HS2 (including insertional effects of a selectable marker) on expression of mouse beta-like globin genes (including the embryonic gene Ey) and on embryonic viability. It does not mention the human HBE1 gene, any human or mouse allelic variants of HBE1, clinical phenotypes attributable to mutations in HBE1, or any Mendelian inheritance patterns linking HBE1 to disease. The observed embryonic lethality and expression changes are either due to insertional artifacts (neo) or are specific to the regulatory element in the mouse locus; the authors explicitly show that removal of the selectable marker largely rescues expression. Therefore the manuscript provides mechanistic background about globin locus regulation but no direct evidence that HBE1 is associated with at least one Mendelian disease. Additional reasons relevance is very low: (1) species difference (mouse Ey versus human HBE1), (2) focus on regulatory element rather than coding gene mutations, (3) absence of clinical/genetic linkage data or variant-disease associations in the paper.


[Read Paper](https://www.semanticscholar.org/paper/e689d53cb00cbaf535a7a77cbaaf4b2899b15cbb)


### An "in-out" strategy using gene targeting and FLP recombinase for the functional dissection of complex DNA regulatory elements: analysis of the beta-globin locus control region.

**Why Not Relevant**: The paper focuses on functional analysis of the locus control region (LCR) regulating beta-like globin genes using a somatic hybrid cell model and recombination methods. It does not mention the HBE1 gene by name, provides no genetic association or variant-level data, and presents no clinical or phenotypic evidence linking any specific globin gene to Mendelian disease. All findings are mechanistic and methodological (effects of inserting/deleting a selectable marker on LCR function) in an engineered cell system. While the mechanistic result—that LCR integrity controls expression of beta-like globin genes—makes it biologically plausible that regulatory mutations in the beta-globin locus could cause globin-related Mendelian diseases, the paper contains no direct or specific evidence that HBE1 is associated with a Mendelian disease. Additional limitations reducing relevance: the experimental system is an artificial somatic hybrid cell line (murine erythroleukemia hybrids with human chromosome 11), the manipulations are engineered insertions/deletions (not naturally occurring variants), and no human genetic or clinical data are provided. For these reasons, the paper provides only very indirect mechanistic background at best and no direct evidence supporting the claim that HBE1 is associated with a Mendelian disease.


[Read Paper](https://www.semanticscholar.org/paper/688380ede95d89163f6ac5a79e110eb089ac97a5)


### The Burden of Candidate Pathogenic Variants for Kidney and Genitourinary Disorders Emerging From Exome Sequencing

**Why Not Relevant**: The supplied paper content does not mention the gene HBE1 (or synonymous terms such as "hemoglobin epsilon 1", "epsilon globin", or related hemoglobinopathies). The study is focused on genes associated with Mendelian kidney and genitourinary disorders (it describes methodologic steps: querying OMIM and Orphanet, manual curation to a 625-gene list, use of ClinVar/HGMD/ExAC, and analysis of a cohort of 7,974 ostensibly healthy adults). All excerpts in the provided text relate to the process for identifying and filtering variants in kidney/genitourinary disease genes and to the generalized problem of variant misclassification; none provide direct evidence (variant-level association, segregation, case reports) or mechanistic evidence (gene function, pathogenic mechanism, animal models, expression data) relevant to HBE1.

Limitations and implications:
- Absence of HBE1 in the provided content means this paper neither supports nor refutes the claim that "HBE1 is likely associated with at least one Mendelian disease." Lack of mention is not definitive proof either way; it only shows this paper does not supply relevant evidence.
- I do not have access to the appendices or the full curated gene list referenced (only the text excerpts you provided). If HBE1 appears in the paper's full gene list or appendices, that was not included in the supplied content and therefore cannot be evaluated here.

Recommended next steps to evaluate the claim directly:
- Query gene-disease resources (OMIM, ClinVar, HGMD, Orphanet) for HBE1 to look for documented Mendelian associations, reported pathogenic variants, and associated phenotypes.
- Search primary literature (PubMed) for case reports, segregation studies, functional assays, or animal model evidence linking HBE1 to Mendelian disease.
- Review population frequency data (gnomAD/ExAC) for HBE1 variants and any ClinVar classifications to assess plausibility and prior classifications.

Because the required verbatim excerpts relevant to HBE1 are absent from the provided document, no direct or mechanistic quotations or excerpt-specific explanations can be supplied from this paper.


[Read Paper](https://www.semanticscholar.org/paper/a75e297208c04992568e607d19aa9b9aaebd09d5)


### Developmental regulation of human gamma- and beta-globin genes in the absence of the locus control region

**Why Not Relevant**: The paper describes experiments on transgenic mice carrying human gamma- (Gγ and Aγ) and beta-globin transgenes and examines hereditary persistence of fetal hemoglobin (HPFH) promoter mutations (-202 Gγ and -117 Aγ). Nowhere in the provided text is the HBE1 gene (the human embryonic epsilon-globin gene) mentioned. The content therefore contains no direct evidence that HBE1 is associated with any Mendelian disease. Mechanistically, the paper shows that mutations in regulatory elements of globin genes (gamma and beta) can produce Mendelian-like phenotypes (HPFH) in a transgenic mouse model; this is a general statement about the globin gene cluster rather than evidence implicating HBE1 specifically. Limitations that further reduce relevance: (1) the study focuses on Gγ, Aγ and β-globin expression and specific promoter mutations associated with HPFH, not on epsilon (HBE1); (2) the work is in transgenic mice and pertains to developmental regulation and promoter effects in that context, so even mechanistic inferences about other globin genes are indirect; (3) without any mention of HBE1 sequence, function, mutation, or phenotype, there is no direct or strong mechanistic link to support the claim that HBE1 is likely associated with at least one Mendelian disease. For these reasons the paper is not relevant evidence for the claim.


[Read Paper](https://www.semanticscholar.org/paper/e03a4d8fa1f9ad5be1c207e040bbb1c9962e975e)


### Developmental regulation of human gamma- and beta-globin genes in the absence of the locus control region.

**Why Not Relevant**: The paper describes experiments on human gamma- and beta-globin transgenes (and specific HPFH promoter mutations) in mice, addressing developmental regulation and functional effects of promoter variants for G gamma and A gamma genes. It does not mention HBE1 (epsilon-globin) anywhere in the provided text, nor does it present data on HBE1 sequence, expression, variants, or disease association. Therefore the paper provides no direct or mechanistic evidence linking HBE1 to any Mendelian disease. The biological mechanisms discussed (cis-regulatory control of gamma/beta genes, effects of specific promoter mutations) are specific to those genes and cannot be assumed to apply to HBE1 without direct data. For these reasons the relevance to the claim that "HBE1 is likely associated with at least one Mendelian disease" is negligible.


[Read Paper](https://www.semanticscholar.org/paper/39a0125b65f1ef37a4d7f42a6f2e75f213bb731f)


### Precision Medicine in Internal Medicine

**Why Not Relevant**: The paper is a general overview of genetic testing strategies, variant interpretation, and the state of gene–disease discovery. It does not mention HBE1 or present any experimental, clinical, or database-specific evidence (e.g., case reports, segregation studies, ClinVar/OMIM entries) linking HBE1 to a Mendelian disease. The few statements in the paper that bear indirectly on the claim are high-level (percent of genes linked to Mendelian disorders; capabilities of exome sequencing; challenges in establishing associations) and therefore only provide contextual or methodological relevance. They do not provide direct or mechanistic evidence that HBE1 is associated with any Mendelian disorder. Because the paper contains no HBE1-specific data, it cannot be used to support the claim that HBE1 is likely associated with at least one Mendelian disease. Additional sources (gene-specific literature, OMIM/ClinVar entries, case reports, functional studies) would be required to evaluate the claim directly.


[Read Paper](https://www.semanticscholar.org/paper/c7199e8a5226f8d9d55777da0e15e91797bb459a)


### Activation of the beta-like globin genes in transgenic mice is dependent on the presence of the beta-locus control region.

**Why Not Relevant**: The paper investigates the regulatory requirement of the beta-globin LCR for expression of globin genes (epsilon/gamma/beta) using transgenic YACs in mice and tests how an HPFH-promoter mutation in gamma-globin behaves without the LCR. Although it provides mechanistic evidence that epsilon-globin (HBE1) expression depends on the LCR (i.e., removal of the LCR abolishes epsilon transcript detection), it contains no data linking HBE1 sequence variation or functional loss directly to any Mendelian disease in humans. There are no human genetics, clinical phenotype, inheritance pattern, or disease-association data presented. The experiments are performed in engineered mouse transgenics and probe regulatory-element dependence rather than allele-disease causality. Therefore the paper is only weakly relevant (mechanistic background) to the claim that HBE1 is likely associated with at least one Mendelian disease.


[Read Paper](https://www.semanticscholar.org/paper/7d237bbdb946a4501d8a617e0a63f14a83303d1e)


## Search Queries Used

- HBE1 HBE epsilon globin HBE1 mutation variant pathogenic germline monogenic Mendelian disease case report

- HBE1 epsilon globin hereditary persistence of fetal hemoglobin HPFH delta beta thalassemia hemoglobinopathy HBE1 associated disease

- beta globin locus regulation locus control region LCR HBE1 promoter enhancer deletion HPFH regulatory mutation embryonic globin expression

- HBE1 OMIM ClinVar Online Mendelian Inheritance in Man gene disease association review systematic review meta analysis monogenic gene


## Usage Statistics

- Prompt Tokens: 0

- Completion Tokens: 0

- Total Tokens: 0

- Estimated Cost: $0.0473


## Bibliometric Configuration

- Use Bibliometrics: True

- Author Impact Weight: 0.4

- Citation Impact Weight: 0.4

- Venue Impact Weight: 0.2
