## Supplementary material for "MENDELSEEK: An algorithm that predicts Mendelian Genes and elucidates what makes them special": HHEX.txt

# Claim: The gene HHEX is likely associated with at least one Mendelian disease.

**Status**: processed

**Overall Rating**: -1

**Explanation**:

No relevant papers were found that support or refute this claim.


## Relevant Papers


## Other Reviewed Papers


### Evaluating the Clinical Validity of Hypertrophic Cardiomyopathy Genes

**Why Not Relevant**: The paper focuses specifically on evaluating gene-disease associations for hypertrophic cardiomyopathy (HCM) and syndromic genes with left ventricular hypertrophy. The provided content lists and discusses specific HCM genes (e.g., MYBPC3, MYH7, TNNT2, etc.) and aggregate findings about 57 curated genes and ClinVar variants, but it does not mention the gene HHEX anywhere. Because HHEX is not named or described, there is no direct evidence (case reports, segregation data, variant pathogenicity, or functional/mechanistic studies) in this paper for or against HHEX being associated with any Mendelian disease. The paper's conclusions about the general quality of HCM gene associations likewise cannot be used to infer an association for HHEX. Therefore the paper is essentially not relevant to the specific claim that HHEX is likely associated with at least one Mendelian disease. Note that lack of mention in this paper does not imply HHEX is not associated with a Mendelian disease; it only means this paper provides no supporting or refuting evidence regarding HHEX.


[Read Paper](https://www.semanticscholar.org/paper/d7dda87eae6df767a7c390a8435f02a29fa32532)


### The Burden of Candidate Pathogenic Variants for Kidney and Genitourinary Disorders Emerging From Exome Sequencing

**Why Not Relevant**: The provided paper excerpt does not mention the gene HHEX anywhere, nor does it present gene-level data or variant-level observations for HHEX. The manuscript describes methods for identifying and curating genes associated with Mendelian kidney and genitourinary disorders (OMIM/Orphanet search, manual curation to a final list of 625 genes) and describes cohort sequencing and variant-filtering workflows, but the excerpt does not list the curated genes or any gene-specific results. Therefore there is no direct evidence (case reports, variant associations, statistical signals) or mechanistic discussion in the supplied text that would support or refute the claim that HHEX is likely associated with at least one Mendelian disease. 

Limitations preventing assessment from this paper content:
- No occurrence of the symbol/name "HHEX" or any synonym in the provided text. 
- The curated gene list is referenced but not included in the excerpt; without that list or the supplementary material, one cannot determine whether HHEX was included among the 625 genes. 
- No variant-level data, phenotype associations, inheritance information, functional studies, or mechanistic descriptions specific to HHEX are present in the excerpt. 

To evaluate the claim using this study, one would need access to the paper's full supplementary material (the curated gene list or Appendix tables) or explicit mentions of HHEX within the results or discussion. Alternatively, consult gene-disease resources (OMIM, ClinVar, HGMD) or primary studies that report HHEX variants and associated Mendelian phenotypes.


[Read Paper](https://www.semanticscholar.org/paper/a75e297208c04992568e607d19aa9b9aaebd09d5)


### Expanding the phenotype of a recurrent de novo variant in PACS1 causing intellectual disability

**Why Not Relevant**: This paper does not provide evidence for or against the claim that HHEX is associated with any Mendelian disease. The entire report focuses on a recurrent de novo missense variant in PACS1 (p.Arg203Trp) causing an intellectual disability syndrome (MRD17). There is no mention of the gene HHEX, no HHEX variants analyzed or reported, and no experiments or mechanistic discussions that implicate HHEX. Mechanistic comments in the paper relate specifically to PACS1 (e.g., its role as a trans-Golgi membrane traffic regulator, proximity of the variant to a CK2-binding motif, and zebrafish antisense experiments affecting SOX10-positive neural crest cells), which are not mechanistically connected to HHEX in this manuscript. Because HHEX is neither discussed nor investigated in the provided content, the paper supplies no direct genetic evidence (variant–phenotype cosegregation or case reports), no functional evidence, and no mechanistic rationale linking HHEX to a Mendelian disease. Therefore the relevance of this paper to the claim about HHEX is effectively zero.


[Read Paper](https://www.semanticscholar.org/paper/e460d89d850693b4fb1e3df12702be0dfb8d7f5b)


### Systematic use of phenotype evidence in clinical genetic testing reduces the frequency of variants of uncertain significance

**Why Not Relevant**: The provided paper content describes a general methodological framework for curating phenotype evidence and demonstrates that the approach improves variant interpretation accuracy. However, it contains no mention of the gene HHEX, no presentation of HHEX variants or HHEX-associated phenotypes, and no mechanistic discussion of HHEX protein function or pathogenic mechanisms. Therefore the paper provides no direct evidence (case reports, cohort data, segregation, functional assays) nor mechanistic evidence linking HHEX to any Mendelian disease. The method described could in principle be applied to HHEX if HHEX phenotypes were curated and predictive, but that is speculative and not contained in the text. Because the content lacks any HHEX-specific data or discussion, its relevance to the claim that HHEX is likely associated with at least one Mendelian disease is negligible.


[Read Paper](https://www.semanticscholar.org/paper/2620a6cde34a04b05159b44505c04aca71239387)


### Actionable Genotypes and Their Association with Life Span in Iceland.

**Why Not Relevant**: The provided paper content describes a study that curated and analyzed variants in the ACMG SF v3.0 gene list and reports aggregate and group-level findings (number of actionable genotypes, counts of genes, survival differences). However, the excerpted text does not mention the gene HHEX anywhere, nor does it provide a gene-level list or variant-level results that include HHEX. Because the claim concerns whether HHEX is likely associated with at least one Mendelian disease, evidence would require (a) explicit inclusion of HHEX among genes associated with Mendelian disease in this study, (b) identification of pathogenic/likely pathogenic variants in HHEX, or (c) mechanistic/phenotypic associations implicating HHEX. None of those are present in the supplied content. Therefore the paper content as provided is not informative regarding HHEX specifically. It is possible that the full paper (outside the supplied excerpt) lists individual genes or variant tables that include HHEX, but that information is not available here. Given the absence of any mention of HHEX in the provided text, the relevance of this content to the claim is negligible.


[Read Paper](https://www.semanticscholar.org/paper/56acca1792702d6478faca121435d7f8e2037b1b)


### Prenatal diagnosis of Norrie disease after whole exome sequencing of an affected proband during an ongoing pregnancy: a case report

**Why Not Relevant**: The provided paper content does not contain any information about the gene HHEX, nor does it mention any specific genes, variants, cases, phenotypes tied to HHEX, or any named Mendelian diseases. The only sentence in the provided content is: "Clinical polymorphism of hereditary ophthalmic pathology can severely complicate establishment of an exact diagnosis and make it time- and cost-consuming, and NGS appears to be the method-of-choice in complicated cases, and this could substantially hasten the establishment of a diagnosis and genetic risk estimation."

Why this does not support the claim:
- No direct evidence: The text contains no report of HHEX variants, no association statistics, no case reports, and no diagnostic findings that would link HHEX to any Mendelian disease. Therefore there is no direct supportive or refuting evidence in this excerpt.
- No mechanistic evidence: The sentence discusses next-generation sequencing (NGS) as a diagnostic tool in hereditary ophthalmic pathology but does not describe molecular mechanisms, functional effects, or biological pathways involving HHEX that would make an association plausible.
- Lack of methodology/results: The content is a general statement or conclusion about NGS utility and clinical polymorphism; it contains no experimental methods, sample sizes, analyses, or results that could be evaluated for quality, significance, or generalizability.

Limitations and caveats of the excerpt for the claim:
- The statement is high-level and generic; while it implies that NGS can be used to find genetic causes of hereditary ophthalmic diseases, it does not present any data, named genes, or findings.
- It is possible (but speculative) to infer that NGS studies could identify HHEX if HHEX were causal in some cases, but inference is not evidence. No specific linkage, segregation, pathogenicity assessment, or functional validation concerning HHEX is provided.

Conclusion: Relevance is effectively zero because the excerpt lacks any direct or mechanistic information about HHEX or its association with Mendelian disease. To evaluate the claim that "HHEX is likely associated with at least one Mendelian disease," one would need primary data (case reports, segregation, variant pathogenicity, functional studies) or authoritative database entries (OMIM, ClinVar) referencing HHEX — none of which are present in the provided content.


[Read Paper](https://www.semanticscholar.org/paper/f445f8140f38787d8a90a992bc8eb55919719ec8)


### Case Report: A Disease Phenotype of Rett Syndrome and Neurofibromatosis Resulting from A Bilocus Variant Combination

**Why Not Relevant**: The provided paper content does not mention HHEX, nor does it provide genetic, functional, or inheritance data that could be linked to HHEX. The single sentence only notes a clinical observation (a child with ASD later showing a Rett-like phenotype) and speculates about possible protein- or cellular-level relationships among 'multilocus variations'. That speculation is generic and could, in principle, apply to any gene set; without identification of the specific loci, variants, pathogenicity assessments, segregation information, or experimental mechanistic data, the excerpt offers no direct or gene-specific mechanistic evidence pertinent to HHEX. Therefore the content is essentially irrelevant to the claim that HHEX is likely associated with at least one Mendelian disease.


[Read Paper](https://www.semanticscholar.org/paper/993edabc0eeab1343cda0a7dbad248110f7e87ad)


### Whole exome sequencing in the diagnosis of liver diseases

**Why Not Relevant**: The provided paper content does not mention the gene HHEX by name and does not provide a gene-level list that would allow determination of whether HHEX was evaluated or found to harbor P/LP variants. The study focused on 502 genes linked to Mendelian liver phenotypes and reports that P/LP variants were found in 25 genes, but the abstracted content supplied here does not identify those genes. Therefore there is no direct evidence (observational association or identified pathogenic variants) nor mechanistic evidence (functional data or described pathways involving HHEX) in the provided text to support or refute the claim that HHEX is associated with at least one Mendelian disease. Additional information (the full gene list examined and the list of 25 genes with P/LP variants, or any supplementary tables) would be required to evaluate HHEX specifically.


[Read Paper](https://www.semanticscholar.org/paper/b07ed494c5c173a3f7be94d5f6d5accc5255636b)


### Case report: Preimplantation genetic testing for infantile GM1 gangliosidosis

**Why Not Relevant**: This paper focuses exclusively on GM1 gangliosidosis caused by pathogenic variants in the GLB1 gene (clinical description, sequencing identification of two GLB1 variants, and preimplantation genetic testing for GLB1). The text contains no mention of the HHEX gene, no HHEX variants, no mechanistic links between HHEX and GLB1 or the disease pathways discussed, and no analyses (genetic, functional, epidemiological) that would bear on whether HHEX is associated with any Mendelian disease. Therefore the paper provides no direct evidence supporting the claim that HHEX is likely associated with at least one Mendelian disease, nor mechanistic evidence strengthening such a claim. Note that absence of evidence in this paper is not evidence of absence—this case report simply does not address HHEX and so cannot be used to support the claim. Given the complete lack of HHEX-related content, relevance to the claim is effectively zero.


[Read Paper](https://www.semanticscholar.org/paper/ce981bc89c565bb4d97baff441baaf8695595448)


## Search Queries Used

- HHEX pathogenic variant HHEX mutation monogenic disorder Mendelian disease clinical case report exome sequencing ClinVar OMIM

- HHEX ClinVar OMIM gene disease association review systematic review gene curation pathogenicity evidence

- hematopoietically expressed homeobox HHEX knockout mouse embryonic development pancreas liver forebrain hematopoiesis phenotype human disease relevance

- developmental disorder exome sequencing HHEX de novo variant Decipher DDD congenital anomalies rare disease cohort HHEX association


## Bibliometric Configuration

- Use Bibliometrics: True

- Author Impact Weight: 0.4

- Citation Impact Weight: 0.4

- Venue Impact Weight: 0.2
