## Supplementary material for "MENDELSEEK: An algorithm that predicts Mendelian Genes and elucidates what makes them special": HOXA9.txt

# Claim: The gene HOXA9 is likely associated with at least one Mendelian disease.

**Status**: processed

**Overall Rating**: -1

**Explanation**:

No relevant papers were found that support or refute this claim.


## Relevant Papers


## Other Reviewed Papers


### Familial myeloid malignancies with germline TET2 mutation

**Why Not Relevant**: The provided paper content does not mention HOXA9 or any data linking HOXA9 to Mendelian disease. The only sentence provided is: "Analysis of Tet2-knockout mice, which are viable, fertile and develop grossly normally, demonstrated that TET2 functions as a tumor suppressor whose haploinsufficiency increases susceptibility to hematological diseases." This sentence describes work on Tet2/TET2 and its role as a tumor suppressor in the context of hematological disease susceptibility. It does not (a) mention HOXA9, (b) report any human Mendelian phenotypes, or (c) describe mechanistic connections between TET2 and HOXA9. Therefore the content provides no direct evidence that HOXA9 is associated with a Mendelian disease. Potential indirect relevance (for example, if HOXA9 were regulated by TET2 or acted in the same pathway) cannot be inferred from this sentence because no such relationships are described; drawing such inferences would require additional data or citations not present here. Limitations: only a single sentence was provided, and it concerns a different gene (TET2) in a mouse model; there is no genetic, clinical, or mechanistic information about HOXA9 in the provided content.


[Read Paper](https://www.semanticscholar.org/paper/dbc8335b2a72a0ea81565ad2f2cfaecb78be96e7)


### Precision Medicine in Internal Medicine

**Why Not Relevant**: The provided paper is a general review of clinical genetic testing strategies, exome/genome sequencing capabilities, variant interpretation challenges, and the state of gene–disease discovery as of 2019. It does not mention the HOXA9 gene, present any data (genetic, clinical, segregation, functional, or statistical) about HOXA9, nor make any gene-specific disease assertions. Therefore it contains only indirect, methodological, and contextual information relevant to how one would discover or evaluate a gene's Mendelian-disease association, but no direct or mechanistic evidence specific to HOXA9. Because the claim asks whether HOXA9 is likely associated with at least one Mendelian disease, and the paper provides no HOXA9-specific findings or evaluations, the paper is not informative for confirming or refuting that claim.


[Read Paper](https://www.semanticscholar.org/paper/c7199e8a5226f8d9d55777da0e15e91797bb459a)


### Congenital Heart Disease Entering a New Era of Human Genetics

**Why Not Relevant**: The provided paper excerpt discusses the genetics of congenital heart disease, de novo mutations, study design (exome sequencing of trios), and that 28 genes were implicated in CHD. Nowhere in the supplied text is the gene HOXA9 named or described. Therefore there is no direct evidence in this paper excerpt linking HOXA9 to any Mendelian disease. The content offers only general mechanistic and contextual support (that single-gene mutations and de novo variants in developmentally expressed genes can cause Mendelian CHD), which makes HOXA9's involvement biologically plausible but does not provide gene-specific evidence. Because the claim concerns association of HOXA9 with at least one Mendelian disease and the excerpt contains no mention of HOXA9 or data about it, the relevance of this paper to the claim is very low.


[Read Paper](https://www.semanticscholar.org/paper/353c3c35c0c635cc27f19a86a407f39e72d6b6d2)


### De Novo RPS20 Mutations in Diamond Blackfan Anemia

**Why Not Relevant**: The provided paper content contains no mention of the gene HOXA9. The study focuses on Diamond Blackfan anemia and genes encoding ribosomal proteins (e.g., RPS7, RPS10, RPS17, RPS19, RPS24, RPS26, RPS29, RPL5, RPL11, RPL15, RPL26, RPL31, RPL35A) and additionally references GATA1 and newly identified de novo variants in RPS20. Mechanistic discussion centers on ribosome assembly defects, altered rRNA processing, nucleolar stress and downstream p53 pathways. None of the text links HOXA9 to DBA or any Mendelian phenotype. Therefore this paper provides no direct evidence that HOXA9 is associated with any Mendelian disease, nor mechanistic evidence that would plausibly implicate HOXA9. Limitations: absence of HOXA9 in this single paper is not definitive evidence that HOXA9 is not disease-associated—lack of mention only means this paper does not inform the claim. To evaluate the claim about HOXA9, one should examine gene-disease databases (OMIM, ClinVar, HGMD), locus-specific literature, and large-scale gene-disease curation resources. The relevance score (0.0) reflects that the provided content offers no supporting or refuting evidence about HOXA9.


[Read Paper](https://www.semanticscholar.org/paper/a615979f9c2853ae21d1e572a50954ed02561c7a)


### NGS based studies on primary immunodeficiencies (PIDs) : causative gene identification, tool development and application

**Why Not Relevant**: I found no direct or mechanistic evidence in the provided paper content linking HOXA9 to any Mendelian disease. The text lists many genes identified or investigated in the context of primary immunodeficiency disorders (e.g., LRBA, ATM, DOCK8, PMS2, TNFRSF18, PIK3CG, LILRB1, EPHB2, TXNIP, CD5, NLRP5, RAG1, RAG2, DNMT3B, ZBTB24, etc.), but HOXA9 does not appear anywhere in the supplied content. The paper focuses on genetic causes and mechanisms of primary immunodeficiencies and the development of variant-prediction tools (VIPPID), and discusses DNA repair/methylation and T-cell repertoire effects for specific genes; none of these sections mention HOXA9 or provide data (variants, segregation, functional assays, or mechanistic rationale) implicating HOXA9 in a Mendelian condition. Therefore this document provides neither direct evidence (no reported HOXA9 variants, case reports, segregation, or pathogenicity data) nor mechanistic evidence (no discussed pathways involving HOXA9 relevant to disease in this cohort) to support the claim.

Limitations of this assessment: absence of mention in this document is not proof that HOXA9 is not associated with any Mendelian disease—only that this particular paper does not provide supporting evidence. To evaluate the claim further, one should search other primary literature and clinical databases (e.g., OMIM, ClinVar, HGMD) for HOXA9 disease associations, or seek sequencing studies that report HOXA9 variants with segregation and functional validation.


[Read Paper](https://www.semanticscholar.org/paper/7786e43fbaf68904111f286747f0cf8968c2dd1e)


## Search Queries Used

- HOXA9 germline mutation case report familial inherited monogenic Mendelian disease de novo variant human patients

- HOXA9 ClinVar OMIM DECIPHER ClinGen gnomAD variant pathogenicity clinical significance HOXA9 germline

- HOXA9 loss of function haploinsufficiency mouse knockout developmental phenotype hematopoiesis homeobox A9 germline functional study

- HOXA9 somatic mutation acute myeloid leukemia AML oncogenic overexpression translocation fusion somatic versus germline evidence


## Bibliometric Configuration

- Use Bibliometrics: True

- Author Impact Weight: 0.4

- Citation Impact Weight: 0.4

- Venue Impact Weight: 0.2
