## Supplementary material for "MENDELSEEK: An algorithm that predicts Mendelian Genes and elucidates what makes them special": HOXC5.txt

# Claim: The gene HOXC5 is likely associated with at least one Mendelian disease.

**Status**: processed

**Overall Rating**: -1

**Explanation**:

No relevant papers were found that support or refute this claim.


## Relevant Papers


## Other Reviewed Papers


### Non-alcoholic fatty liver disease in a pediatric patient with heterozygous familial hypobetalipoproteinemia due to a novel APOB variant: a case report and systematic literature review

**Why Not Relevant**: The paper exclusively investigates and reports a novel pathogenic variant in the APOB gene causing familial hypobetalipoproteinemia (FHBL). There is no mention of the gene HOXC5, no data (genetic, mechanistic, segregation, or literature review) about HOXC5, and no experimental or observational results that implicate or assess HOXC5. Therefore this paper provides no direct or mechanistic evidence for or against the claim that HOXC5 is associated with a Mendelian disease. Note that absence of any mention in this APOB-focused study is not evidence that HOXC5 is not associated with Mendelian disease; it only means this paper cannot be used to support the claim.


[Read Paper](https://www.semanticscholar.org/paper/1f7237acededf0b660a9f44ccc339ccc177fb97b)


### A pathogenic variant in RAB32 causes autosomal dominant Parkinson’s disease and activates LRRK2 kinase

**Why Not Relevant**: The paper exclusively investigates RAB GTPases (explicitly screening 61 RAB family genes) and identifies a specific RAB32 variant (c.213C>G, Ser71Arg) that co-segregates with autosomal dominant parkinsonism and activates LRRK2 in functional assays. Nowhere in the provided content is the gene HOXC5 mentioned, nor are homeobox genes or data on HOXC5 variants, segregation, or disease association reported. Therefore this paper provides no direct genetic evidence, mechanistic data, or clinical observations linking HOXC5 to any Mendelian disease. Because HOXC5 is a different gene family with distinct biological functions (homeobox transcription factor versus RAB GTPase), findings about RAB32 do not provide mechanistic plausibility for HOXC5 involvement. Given the absence of any mention or data on HOXC5 in the provided text, the relevance of this paper to the claim that "HOXC5 is likely associated with at least one Mendelian disease" is effectively zero.


[Read Paper](https://www.semanticscholar.org/paper/a36d28e828ede7c87c650af5f5117982b9749e7b)


### Implications of a De Novo Variant in the SOX12 Gene in a Patient with Generalized Epilepsy, Intellectual Disability, and Childhood Emotional Behavioral Disorders

**Why Not Relevant**: This paper does not provide any direct or mechanistic evidence relevant to the claim that HOXC5 is likely associated with at least one Mendelian disease. The text exclusively discusses SRY-box (SOX) family genes (SOX12, SOX4, SOX11), a de novo variation in SOX12 found by whole exome sequencing, and the possible pathogenicity of that SOX12 variant. There is no mention of the HOX gene family or the specific gene HOXC5, no variant data for HOXC5, and no mechanistic discussion that links SOXC genes to HOX genes or that would plausibly implicate HOXC5 in Mendelian disease. Because the paper contains no statements, data, or mechanistic pathways pertaining to HOXC5, it cannot be used to support or refute the claim. Limitations: absence of evidence in this paper is not evidence that HOXC5 is or is not associated with a Mendelian disorder—only that this particular manuscript does not address HOXC5. If you would like, I can search the literature, OMIM, ClinVar, and population/clinical variant databases for evidence about HOXC5 and Mendelian disease associations.


[Read Paper](https://www.semanticscholar.org/paper/5cf652470efa4406f72843aea8389f4e398a99f8)


### A case report of a patient with primary familial brain calcification with a PDGFRB genetic variant

**Why Not Relevant**: This paper concerns primary familial brain calcification (Fahr’s disease) and reports a PDGFRB variant (p.Arg919Gln) as a candidate variant in a single-family case. The paper names genes previously implicated in PFBC (PDGFB, PDGFRB, SLC20A2, XPR1, MYORG, and JAM2) and discusses pathogenicity predictions and the need for functional validation of the PDGFRB variant. Nowhere in the provided text is the gene HOXC5 mentioned, nor are there any data, analyses, mechanistic discussions, or clinical correlations linking HOXC5 to a Mendelian disease. Therefore this paper provides no direct evidence for or against the claim that HOXC5 is likely associated with at least one Mendelian disease. Note that absence of any mention of HOXC5 in this single paper is not evidence that HOXC5 is not associated with a Mendelian disease more broadly; it simply means this paper is not a relevant source to evaluate that claim. To assess the claim about HOXC5 one would need to examine other sources such as gene-disease databases (OMIM, ClinVar), peer-reviewed reports describing pathogenic HOXC5 variants co-segregating with disease, functional studies, or relevant population/segregation data.


[Read Paper](https://www.semanticscholar.org/paper/5cbb86c12218cdf91c19ae96c89901510a745009)


### A de novo frameshift variant in MED13 gene in a patient with autism spectrum disorder and magnetic resonance imaging abnormalities mimicking tuberous sclerosis

**Why Not Relevant**: Relevance is near zero because the paper content exclusively addresses MED13. There is no mention of the HOXC5 gene, no data (genetic, clinical, or mechanistic) concerning HOXC5, and no discussion of pathways or interactions that would link MED13 findings to HOXC5. Therefore this paper provides no direct evidence (variants, segregation, phenotype) nor mechanistic evidence (functional studies, pathway connections) supporting or refuting the claim that HOXC5 is likely associated with a Mendelian disease. The excerpts chosen are verbatim sentences from the provided text showing the paper's focus on MED13 to justify lack of relevance.


[Read Paper](https://www.semanticscholar.org/paper/c1a65b9ac26a735bddb776612339efbbc90ae89a)


### A de novo PAK1 likely pathogenic variant and a de novo terminal 1q microdeletion in a Chinese girl with global developmental delay, severe intellectual disability, and seizures

**Why Not Relevant**: The provided paper content does not mention the gene HOXC5 at all and instead concerns a novel PAK1 variant and a terminal 1q microdeletion in a patient with a neurodevelopmental disorder. Because there is no reference to HOXC5, no genetic association data, no mechanistic pathways involving HOXC5, and no discussion of HOXC5's genomic location or function in the excerpt, the paper provides no direct or mechanistic evidence for or against the claim that HOXC5 is likely associated with at least one Mendelian disease. Additional considerations that further reduce relevance: the paper is a single-patient report focused on PAK1 and chromosome 1q abnormalities (where HOXC5 is not implicated), and therefore even indirect inference is not supported. To evaluate the claim about HOXC5, sources that explicitly analyze HOXC5 variants, linkage/segregation in pedigrees, case reports associating HOXC5 disruption with Mendelian phenotypes, or functional studies of HOXC5 would be required; none are present in the provided content.


[Read Paper](https://www.semanticscholar.org/paper/8ac30d308f85987b8cf4644520ee11940136ebaa)


### A novel AUTS2 variant in a patient with global developmental delay and intellectual disability

**Why Not Relevant**: The provided paper text discusses a novel, de novo loss-of-function variant in the AUTS2 gene and the clinical characterization of AUTS2 (AUTS2 syndrome). The text makes no mention of the gene HOXC5, no comparison between AUTS2 and HOXC5, and provides no mechanistic pathway information that connects AUTS2 to HOXC5. Therefore it contains no direct evidence that HOXC5 is associated with any Mendelian disease, nor does it provide mechanistic evidence that would make such an association plausible. In addition, the study is a single-case report focused on AUTS2, so even indirect inference (for example by gene family, pathway, or positional overlap) is unsupported by the content provided. For these reasons the paper is not relevant to evaluating the claim that "HOXC5 is likely associated with at least one Mendelian disease."


[Read Paper](https://www.semanticscholar.org/paper/f5e84e7bb3b42239a53cc7e5c457342cb38c6277)


### Genotype first approach & familial segregation analysis help in the elucidation of disease-causing variant for fucosidosis

**Why Not Relevant**: The provided paper content exclusively concerns the FUCA1 gene and fucosidosis (a lysosomal storage disorder). There is no mention of HOXC5, no data, no experimental results, nor any mechanistic discussion relating HOXC5 to disease. Therefore the content does not provide direct evidence (variant associations, case reports, segregation, pathogenic variants) nor mechanistic evidence (biochemical function, developmental role, pathway perturbation) for HOXC5 being associated with a Mendelian disease. Because the paper neither treats HOXC5 nor discusses related pathways or phenotypes that could plausibly implicate HOXC5, it cannot support the claim. The assigned relevance score (0.01) reflects that while the document is clearly genetic/Mendelian-disease–focused, its content is entirely about a different gene (FUCA1); therefore it is effectively non‑relevant to the HOXC5 claim.


[Read Paper](https://www.semanticscholar.org/paper/5cf518959c6fdfb9a9e4906ff08b43ae36b8e31d)


## Search Queries Used

- HOXC5 pathogenic variant monogenic disorder Mendelian disease case report familial segregation

- HOXC5 de novo variant exome sequencing neurodevelopmental disorder intellectual disability autism developmental delay trio sequencing

- HOXC5 homeobox C5 HOX C5 mouse knockout zebrafish morpholino phenotype craniofacial limb axial patterning functional assay transcription factor

- HOXC5 OMIM ClinVar HGMD gene disease association curation review HOX gene family redundancy compensation lack of human disease


## Bibliometric Configuration

- Use Bibliometrics: True

- Author Impact Weight: 0.4

- Citation Impact Weight: 0.4

- Venue Impact Weight: 0.2
