## Supplementary material for "MENDELSEEK: An algorithm that predicts Mendelian Genes and elucidates what makes them special": HTR2C.txt

# Claim: The gene HTR2C is likely associated with at least one Mendelian disease.

**Status**: processed

**Overall Rating**: 3

**Explanation**:

Supporting evidence.
The provided paper reports a loss-of-function (LOF) mutation in the mouse Htr2c gene (Htr2cF327L/Y) that produces a clear, reproducible phenotype: hyperphagia, obesity and related metabolic dysfunction (hyperglycemia). The authors show reversal or mitigation of that phenotype by re-expressing functional 5-HT2CR specifically in hypothalamic POMC neurons, by pharmacologic activation of downstream melanocortin signaling (MC4R agonist), and by behavioral interventions. These experiments establish a causal link in mice between Htr2c LOF and a Mendelian-like phenotype (single-gene manipulation producing a consistent physiological outcome). The authors additionally state that MC4R agonists and lifestyle changes “might hold promise in counteracting … in individuals carrying rare variants in Htr2c,” implying a translational hypothesis that rare HTR2C variants in humans could produce similar phenotypes.

Caveats or contradictory evidence.
The evidence is entirely from a mouse model; no human genetic data are provided in the excerpt (no reported human cases, no segregation data, no OMIM/clinical annotation). Demonstration that a gene causes a phenotype in a model organism is necessary but not sufficient to establish that gene as causative for a Mendelian disease in humans. Differences in physiology, compensatory biology, variant-specific effects, and species-specific expression patterns can all lead to discordance between mouse knockout/LOF phenotypes and human monogenic disease. The excerpt does not present human pathogenic variants, de novo events, family segregation, or clinical reports that would meet usual criteria for assigning a Mendelian disease to HTR2C.

Analysis of potential mechanisms and implications.
Mechanistically, 5-HT2CR is expressed in hypothalamic POMC neurons and modulates melanocortin signaling, a pathway already implicated in human monogenic obesity (e.g., MC4R mutations). The mouse data show that Htr2c LOF produces hyperphagia and obesity that can be rescued by downstream activation, consistent with HTR2C functioning upstream in a pathway that is relevant to human energy homeostasis. Because HTR2C is X-linked, hemizygous LOF variants in males could, in principle, produce a penetrant phenotype analogous to the mouse model. Therefore, from a mechanistic and biological plausibility standpoint, HTR2C is a credible candidate gene for a Mendelian (likely X-linked) syndrome of severe early-onset hyperphagia/obesity. However, mechanistic plausibility and animal model causality do not substitute for direct human genetic evidence required to classify a gene as definitively associated with a Mendelian disease.

Which rating is most appropriate and why.
Given the strong, causal mouse-model evidence and plausible mechanism linking HTR2C LOF to a phenotype already known to be monogenic when other pathway components (e.g., MC4R) are mutated, there is biologic plausibility that HTR2C could underlie a Mendelian disorder in humans. However, the provided evidence contains no direct human genetic or clinical data demonstrating that HTR2C variants segregate with disease or produce a reproducible Mendelian phenotype in people. Therefore the correct assessment is not “Highly Supported” or “Likely True,” but also not “No Evidence” or “Contradicted.” The appropriate classification is that evidence is mixed: compelling preclinical support and plausible mechanism exist, but human Mendelian association has not been demonstrated in the provided material.


**Final Reasoning**:

I reviewed the logic and evidence: the single provided paper gives robust experimental proof that Htr2c LOF causes hyperphagia and obesity in mice and identifies a clear mechanistic pathway (5-HT2CR → POMC → MC4R). That creates plausible grounds for hypothesizing a human Mendelian condition due to HTR2C LOF, particularly given known monogenic obesity genes in the same pathway and the X-linked nature of HTR2C. However, the claim to be evaluated is specifically that HTR2C is "likely associated with at least one Mendelian disease." The evidence provided lacks any human genetic or clinical data (no documented pathogenic variants in patients, no familial segregation, no clinical reports), which are required to move from plausible candidate to likely human Mendelian gene. Balancing strong mechanistic and animal-model support against the absence of human genetic evidence in the provided materials, the best-fit rating is "Mixed Evidence." I reaffirm this rating: suggestive and plausible but not confirmed by human Mendelian data in the excerpts given.


## Relevant Papers


### Therapeutic strategies against metabolic imbalance in a male mouse model with 5-HT2CR loss-of-function.

**Authors**: Hailan Liu (H-index: 2), Yang He (H-index: 2)

**Relevance**: 0.75

**Bibliometric Impact**: 0.15599999999999997


**Excerpts**:

- The serotonin 2C receptor (5-HT2CR)-melanocortin pathway plays well-established roles in the regulation of feeding behavior and body weight homeostasis.

- Dysfunctions in this system, such as loss-of-function mutations in the Htr2c gene, can lead to hyperphagia and obesity.

- In this study, we aimed to investigate the potential therapeutic strategies for ameliorating hyperphagia, hyperglycemia and obesity associated with a loss-of-function mutation in the Htr2c gene (Htr2cF327L/Y).

- We demonstrated that re-expressing functional 5-HT2CR solely in hypothalamic pro-opiomelanocortin (POMC) neurons is sufficient to reduce food intake and body weight in Htr2cF327L/Y mice subjected to a high-fat diet.

- In addition, 5-HT2CR expression restores the responsiveness of POMC neurons to lorcaserin, a selective agonist for 5-HT2CR.

- Similarly, administration of melanotan II (MT II), an agonist of the melanocortin receptor 4 (MC4R), effectively suppresses feeding and weight gain in Htr2cF327L/Y mice.

- Strikingly, promoting wheel running activity in Htr2cF327L/Y mice results in a decrease in HFD consumption and improved glucose homeostasis.

- Together, our findings underscore the crucial role of the melanocortin system in alleviating hyperphagia and obesity related to dysfunctions of the 5-HT2CR, and further suggest that MC4R agonists and lifestyle interventions might hold promise in counteracting hyperphagia, hyperglycemia and obesity in individuals carrying rare variants in Htr2c gene.


**Explanations**:

- Mechanistic context: This sentence establishes the biological pathway (5-HT2CR–melanocortin) relevant to energy balance. It strengthens plausibility that perturbations of HTR2C can produce clinically relevant phenotypes (feeding, weight). Limitation: general statement; no experimental data or species specified in this sentence.

- Direct mechanistic/phenotypic link (inferred from the paper): This asserts that loss-of-function mutations in Htr2c produce hyperphagia and obesity, directly tying gene dysfunction to a clear phenotype. Strength: makes a causal connection. Limitation: the excerpt does not specify species or experimental details here (later text indicates a mouse model), so translation to human Mendelian disease remains inferential rather than demonstrated.

- Model specification: This sentence indicates the study examines a specific loss-of-function allele (Htr2cF327L/Y), i.e., a defined mutant model. Relevance: shows the authors used a single-gene mutation model to study the phenotype, which is the typical approach for Mendelian-gene inference. Limitation: it specifies an experimental mouse allele; it does not provide human genetic evidence or clinical cases.

- Mechanistic rescue/direct causal evidence in model organism: Re-expression of functional 5-HT2CR only in POMC neurons sufficed to reduce hyperphagia/body weight in the mutant mice on a high-fat diet. Strength: strong causal evidence in the model that loss of 5-HT2CR function causes the phenotype and that restoring the gene in a specific cell type rescues it—consistent with a monogenic effect. Limitations: this is an experimental mouse rescue under a high-fat diet; it demonstrates causality in mice but is not direct proof of a human Mendelian disease. Also, quantitative/statistical outcomes are not provided in the excerpt.

- Mechanistic detail: Shows that 5-HT2CR expression restores pharmacologic responsiveness of POMC neurons to a selective agonist, connecting molecular expression to cellular function. Relevance: strengthens mechanism by which HTR2C dysfunction leads to altered neuronal signaling and thus feeding behavior. Limitation: cellular pharmacology in the model; no human clinical data.

- Therapeutic/mechanistic evidence in the model: Demonstrates that downstream activation of MC4R (with MT II) suppresses feeding/weight gain in the Htr2c mutant mice, supporting that the melanocortin pathway downstream of 5-HT2CR mediates the phenotype. Relevance: supports mechanistic pathway linking HTR2C to an obesity phenotype and suggests potential interventions. Limitation: again in mice; not direct human genetic disease evidence.

- Phenotypic modulation by lifestyle in model: Shows that non-genetic intervention (exercise) can ameliorate aspects of the phenotype in mutant mice, indicating phenotype is modifiable. Relevance: underscores that the mutant-gene–associated phenotype is physiologically meaningful. Limitation: this does not bear directly on Mendelian causation in humans; it shows environmental modulation in the mouse model.

- Translational inference and context: The authors conclude the melanocortin system is crucial for alleviating hyperphagia/obesity related to 5-HT2CR dysfunction and explicitly suggest that MC4R agonists and lifestyle changes might help "individuals carrying rare variants in Htr2c gene." Relevance: the sentence explicitly connects the mouse findings to the possibility of human carriers of rare HTR2C variants having related metabolic phenotypes—thus supporting the plausibility that HTR2C variants could underlie a Mendelian-like disorder of hyperphagia/obesity. Limitation: this is an inference/implication rather than empirical human genetic evidence; no human pedigrees, segregation, or clinical diagnoses are reported in the provided text.


[Read Paper](https://www.semanticscholar.org/paper/3ba8839c20517102207d11cbddc84d5c10611d71)


## Other Reviewed Papers


### A de Novo ZMIZ1 Pathogenic Variant for Neurodevelopmental Disorder With Dysmorphic Facies and Distal Skeletal Anomalies

**Why Not Relevant**: The paper exclusively studies NEDDFSA caused by heterozygous variants in ZMIZ1. All presented background, methods, results, mechanistic analyses, and conclusions relate to ZMIZ1 and the reported de novo ZMIZ1 missense variant. There is no mention of HTR2C, no reported variants in HTR2C, and no mechanistic discussion involving HTR2C. Therefore the paper provides no direct evidence that HTR2C is associated with any Mendelian disease, nor does it provide mechanistic evidence relevant to HTR2C. Caveats: the authors performed whole-exome sequencing (WES), which could in principle detect HTR2C variants, but the manuscript excerpts provided do not list all WES findings (they note 20 variants were detected but only report the ZMIZ1 variant). Thus, absence of mention of HTR2C in the text is not a definitive negative screen for HTR2C association, but it does mean this paper contains no usable evidence for the claim that HTR2C is likely associated with a Mendelian disease.


[Read Paper](https://www.semanticscholar.org/paper/77be037b2243e6dec4d9cad4679927a8190d818f)


### A novel UBE2A mutation in a Chinese family with X-linked intellectual disability.

**Why Not Relevant**: This paper does not provide evidence for or against the claim that HTR2C is associated with a Mendelian disease. The manuscript exclusively reports on UBE2A and UBE2A deficiency syndrome. For example, the paper states verbatim: "BACKGROUND X-linked intellectual disability type Nascimento, also known as UBE2A deficiency syndrome, is an intellectual disability syndrome characterized by moderate to severe intellectual disability, dysmorphic facial features, speech impairment, genital anomalies and skin abnormalities. The syndrome is caused by mutations of the UBE2A gene, or larger deletions of Xq24 encompassing UBE2A." It also states: "RESULTS A novel UBE2A mutation (c.TAT245TGT, p.Tyr82Cys) was identified in our patient. The heterozygous missense UBE2A mutation was identified in his mother, but not in his father or sister." The paper describes methods ("Whole exome sequencing and Sanger sequencing were used to identify the disease-causing mutations in this intellectual disabilities and epilepsy patient") and concludes: "This study identified a novel UBE2A mutation in a patient with severe intellectual disability and seizures. Our findings expand the mutational spectrum of UBE2A gene." Nowhere in the provided content is HTR2C mentioned, nor are any data, mechanisms, or analyses concerning HTR2C presented. Therefore this paper neither supports nor refutes the claim about HTR2C. Limitations: absence of HTR2C in this paper does not imply HTR2C is unrelated to Mendelian disease in other studies; it only means this particular paper provides no relevant evidence for the claim.


[Read Paper](https://www.semanticscholar.org/paper/ae64d1417c91cc55d8d2a8bfed174a5ec89589e0)


### Case Report: De novo DDX3X mutation caused intellectual disability in a female with skewed X-chromosome inactivation on the mutant allele

**Why Not Relevant**: The paper's content is entirely focused on DDX3X and the role of skewed X-chromosome inactivation (XCI) in a female proband carrying a DDX3X:c.694_711dup18 de novo mutation. The text describes DDX3X-specific genetic findings, assays (PCR-Sanger SNP detection, AR assay, RNA sequencing, RT-PCR), allele-specific expression (70% normal vs. 30% mutated), and conclusions about XCI effects on DDX3X-related neurodevelopmental disorder. Nowhere in the provided content is the gene HTR2C mentioned, nor are there any data, analyses, or mechanistic discussions that link HTR2C to disease. Although the paper provides mechanistic insights about XCI and how XCI-escape may affect phenotypes for X-linked genes (which is a concept that could, in principle, be relevant to other X-linked genes), such general mechanistic discussion is insufficient to support the specific claim that HTR2C is likely associated with at least one Mendelian disease. In short, this paper provides no direct genetic, clinical, or mechanistic evidence pertaining to HTR2C; therefore it cannot be used to support or refute the claim about HTR2C's Mendelian disease associations.


[Read Paper](https://www.semanticscholar.org/paper/1b83fd16d7ea5994e1e071db326a487d9e833820)


### Danon disease in a Sardinian family: different aspects of the same mutation—a case report

**Why Not Relevant**: The provided paper content is not relevant to the claim that HTR2C is likely associated with at least one Mendelian disease. The abstract explicitly attributes the disorder under study to a different gene and disease: "Background Danon disease (DD) is a rare X-linked disorder due to mutations in the lysosome-associated membrane protein 2 gene." The paper focuses on Danon disease and LAMP2-related pathology (for example: "It is characterized by a clinical triad of hypertrophic cardiomyopathy, skeletal myopathy, and a variable degree of intellectual disability."). Nowhere in the supplied text is HTR2C, serotonin receptor 2C, 5-HT2C, or any alternate gene besides LAMP2 mentioned. Therefore there is no direct evidence (no reported mutations, case reports, linkage, segregation, or association) tying HTR2C to a Mendelian disease in this paper. There is also no mechanistic discussion linking HTR2C to the pathophysiology described for Danon disease (lysosome-associated membrane protein 2 dysfunction leading to cardiomyopathy and myopathy). Limitations: the assessment is based solely on the text excerpt you provided (the abstract and case-summary material). If the full paper included additional sections not provided here (e.g., a broader literature review or supplementary data), those were not available for review; however, given the explicit and narrow focus on LAMP2 and Danon disease in the supplied content, it is very unlikely that this paper provides evidence supporting an association between HTR2C and a Mendelian disorder.


[Read Paper](https://www.semanticscholar.org/paper/d9133abd84b99059f8be1c8c9d89b5228c7afa1d)


### Unveiling a Novel THOC2 Mutation's Role in X-linked Intellectual Disability

**Why Not Relevant**: The paper's entire genetic analysis and conclusions focus on THOC2 as the causative gene for X-linked intellectual disability in the reported family. The provided content contains no mention of the gene HTR2C, no functional or mechanistic data concerning HTR2C, and no discussion linking HTR2C to any Mendelian phenotype. Therefore this paper does not provide supporting or refuting evidence regarding the claim that HTR2C is likely associated with at least one Mendelian disease. Note that absence of evidence in this single THOC2-focused case report does not disprove HTR2C involvement in Mendelian disease elsewhere, but for the purpose of assessing this paper's relevance to the claim, relevance is effectively zero because HTR2C is not addressed.


[Read Paper](https://www.semanticscholar.org/paper/24e8651552c97edfac56156718bd864e005e292d)


### A de novo pathogenic variant in DHX30 gene in a fetus with isolated dysgenesis of the corpus callosum

**Why Not Relevant**: The provided paper content does not mention the gene HTR2C anywhere. It describes a prenatal case with a de novo missense pathogenic mutation c.2353 C > T in the DHX30 gene and discusses DHX30-associated neurodevelopmental disorder, imaging features, and the role of exome sequencing. Because the claim concerns HTR2C being likely associated with at least one Mendelian disease, and there is no direct or mechanistic discussion of HTR2C, this paper provides no direct evidence (no case reports, no variant data, no segregation, no functional studies) nor mechanistic evidence (no pathway, expression, or functional assays) relevant to HTR2C. Limitations: while the paper shows that exome sequencing can identify Mendelian causes of fetal brain anomalies (an indirect methodological point), that does not constitute evidence linking HTR2C to a Mendelian disease. Therefore the paper is not relevant to supporting or refuting the claim about HTR2C.


[Read Paper](https://www.semanticscholar.org/paper/185b90b03c94008f9d651d2f43e9c135a7a1b5aa)


### A Novel Neuroimaging Phenotype in the X-Linked Intellectual Disability with a Missense Mutation of CNKSR2 Gene

**Why Not Relevant**: The paper focuses exclusively on a missense pathogenic variant in the CNKSR2 gene in a single case of X-linked syndromic intellectual disability (MRXSHG). The text contains no mention of HTR2C, no experiments involving HTR2C, and no mechanistic discussion linking CNKSR2 to HTR2C. Therefore the paper provides no direct or mechanistic evidence that HTR2C is associated with any Mendelian disease. Additional reasons: (1) the genetic finding described is specific to CNKSR2 (hemizygous variant; maternal carrier), (2) the report is a single-case clinical report and would not by itself establish broad gene–disease associations even if HTR2C were discussed, and (3) there are no functional, genetic, or clinical data in this paper pertinent to HTR2C. Because the content does not address HTR2C at all, its relevance to the claim is essentially nil.


[Read Paper](https://www.semanticscholar.org/paper/dbf16043597ced631e613919449fd6c42f1ab18e)


### Ablation of MARK 4 , an AMPK-related Kinase , Leads to Insulin Hypersensitivity and Resistance to Diet-induced Obesity

**Why Not Relevant**: The provided paper content exclusively concerns MARK4 (Par-1d/MARK4), an AMPK-related kinase, and reports mouse knockout experiments and metabolic phenotypes (hyperphagia, hyperactivity, hypermetabolism, effects on brown adipose tissue, AKT phosphorylation, and AMPK activity). There is no mention of HTR2C (the serotonin receptor 2C gene), no discussion of human genetic variation, and no reference to Mendelian diseases or clinical genetic associations. Mechanistic pathways described (kinase signaling, AMPK, AKT, brown-fat activity) do not implicate HTR2C in this text. Therefore the paper content provides neither direct evidence (genetic association, patient phenotypes, linkage/segregation data) nor mechanistic evidence connecting HTR2C to any Mendelian disorder. Because the content does not address the gene in question or Mendelian disease concepts at all, its relevance to the claim is effectively zero.


[Read Paper](https://www.semanticscholar.org/paper/07fd2ee21a549dff06d6c71f43def6dd32725d93)


### c.1103T>C (p.Ile368Th) de novo Variant in Synaptotagmin 1 (SYT1) Gene is Pathogenic, Leading to an Ultra-Rare Neurodevelopmental Disorder: The Baker-Gordon Syndrome

**Why Not Relevant**: The paper excerpt exclusively concerns Baker-Gordon Syndrome and pathogenic variants in the SYT1 gene. There is no mention of HTR2C, no association studies, mutation reports, mechanistic links, or phenotypic descriptions involving HTR2C. Therefore this paper provides no direct or mechanistic evidence for or against the claim that HTR2C is likely associated with at least one Mendelian disease. Because the gene of interest (HTR2C) is not discussed or analyzed in any way in the provided content, relevance to the HTR2C claim is effectively zero. Limitations: assessment is based solely on the supplied abstract content; if other sections of the full paper (not provided) mention HTR2C, that content was not available for review.


[Read Paper](https://www.semanticscholar.org/paper/ee225e1b916fbf3fda1d1b5451fe2038e281b8c8)


### Abstract SY14-02: From disease genetics to diagnostics to functional biology: High-throughput proteomics and protein interaction network building guides discovery of novel disease genes in ciliopathies, cystic disease, and cancer

**Why Not Relevant**: The paper content contains no mentions of HTR2C (the serotonin 2C receptor), no genetic association data (linkage, mutation, sequencing) involving HTR2C, and no functional assays or clinical descriptions implicating HTR2C in any Mendelian disease. The mechanistic material about GPCR trafficking into cilia (TULP3/IFT-A) is generic to GPCRs and could, in principle, be relevant to GPCR family members, but the paper provides no evidence that HTR2C localizes to cilia, is affected by the described pathways, or has pathogenic variants causing a Mendelian syndrome. Therefore the paper provides at best speculative mechanistic plausibility for GPCRs broadly, but no direct or specific evidence supporting the claim that HTR2C is likely associated with at least one Mendelian disease. Given these points, the paper is not relevant to establishing HTR2C as a Mendelian disease gene.


[Read Paper](https://www.semanticscholar.org/paper/c4d4c5b6e9383a8226e18fbeef555bd9008870c8)


## Search Queries Used

- HTR2C mutation Mendelian disease case report X linked intellectual disability obesity

- serotonin receptor 2C HTR2C gene monogenic disease review ClinVar OMIM

- HTR2C loss of function knockout mouse hyperphagia obesity neurodevelopmental functional study

- HTR2C rare variant de novo pathogenic variant exome sequencing Mendelian neurodevelopmental disorder


## Usage Statistics

- Prompt Tokens: 0

- Completion Tokens: 0

- Total Tokens: 0

- Estimated Cost: $0.0541


## Bibliometric Configuration

- Use Bibliometrics: True

- Author Impact Weight: 0.4

- Citation Impact Weight: 0.4

- Venue Impact Weight: 0.2
