## Supplementary material for "MENDELSEEK: An algorithm that predicts Mendelian Genes and elucidates what makes them special": HTR3A.txt

# Claim: The gene HTR3A is likely associated with at least one Mendelian disease.

**Status**: processed

**Overall Rating**: -1

**Explanation**:

No relevant papers were found that support or refute this claim.


## Relevant Papers


## Other Reviewed Papers


### The Burden of Candidate Pathogenic Variants for Kidney and Genitourinary Disorders Emerging From Exome Sequencing

**Why Not Relevant**: The paper content provided focuses specifically on identification, curation, and analysis of genes associated with Mendelian kidney and genitourinary disorders (a curated list of 625 genes) and on assessing the prevalence of candidate pathogenic variants in those genes in a cohort of ostensibly healthy adults. The excerpts and methods make clear the scope is limited to kidney/genitourinary Mendelian genes; the provided text does not mention HTR3A, perform analyses on HTR3A, or present mechanistic or clinical evidence linking HTR3A to any Mendelian disease. Therefore this paper does not provide direct or mechanistic evidence relevant to the claim that HTR3A is likely associated with at least one Mendelian disease. Note that absence of mention in this disease-focused paper is not definitive proof that HTR3A has no Mendelian associations — it only means this study did not address that gene or relevant phenotypes.


[Read Paper](https://www.semanticscholar.org/paper/a75e297208c04992568e607d19aa9b9aaebd09d5)


### Whole-Exome Sequencing in Adults With Chronic Kidney Disease

**Why Not Relevant**: The provided paper text does not mention HTR3A, report variants in HTR3A, or describe any clinical or mechanistic evidence linking HTR3A to a Mendelian disease. The study focuses on diagnosing Mendelian forms of kidney disease and hypertension using whole-exome sequencing with a curated list of 287 nephropathy-related OMIM genes and review of other OMIM and ACMG actionable genes. While that analytic framework could in principle detect Mendelian associations in genes present in OMIM, the excerpted manuscript content contains no findings, statements, or mechanistic discussion about HTR3A. Methodological caveats in the paper (exome capture blind spots) mean a negative result could reflect technical limits, but no data are presented here about HTR3A capture status or observed variant frequencies. Therefore this paper, as excerpted, provides no direct or mechanistic evidence to support or refute the claim that HTR3A is likely associated with at least one Mendelian disease.


[Read Paper](https://www.semanticscholar.org/paper/a29bd57964f68ac5f8e1abcafe1112ba488f53f5)


### Annotation of uORFs in the OMIM genes allows to reveal pathogenic variants in 5′UTRs

**Why Not Relevant**: The provided content is only the paper abstract and does not mention the gene HTR3A anywhere. The abstract describes a broad study: manual annotation of upstream translation initiation sites (TISs) in human disease-associated genes from the OMIM database, experimental validation of uORFs in several named genes (ETFDH, PAX9, MAST1, HTT, TTN, GLI2, COL2A1, ZIC2), and identification of variants from HGMD and ClinVar that disrupt uORFs and "thereby could lead to Mendelian disorders." None of these sentences cite or implicate HTR3A specifically, and no gene-level list or supplementary data are provided in the excerpt that could be used to infer whether HTR3A was included among the OMIM genes analyzed or among the genes with uORF-disrupting variants.

Because the abstract does not name HTR3A, there is neither direct evidence (e.g., a statement that HTR3A has pathogenic variants or is implicated in a Mendelian disease) nor mechanistic evidence specific to HTR3A (e.g., demonstration that a uORF in HTR3A is disrupted by pathogenic variants) available in the provided content. It is possible that the full text or supplementary materials include HTR3A in their OMIM gene list or variant tables, but that cannot be determined from the abstract alone. Therefore, based solely on the provided content, the paper does not provide usable evidence for or against the claim that HTR3A is likely associated with at least one Mendelian disease.

If you want a definitive assessment using this paper, please provide the full text or the supplementary tables (gene lists / variant lists) so I can check whether HTR3A is included and extract any relevant verbatim sentences and mechanistic details. Alternatively, I can search other literature or variant databases (OMIM, ClinVar, HGMD) for HTR3A-specific evidence if you would like.


[Read Paper](https://www.semanticscholar.org/paper/5d45c0cdf801505df45fc93a466a2a4101a12871)


### From case studies to community knowledge base: MSeqDR provides a platform for the curation and genomic analysis of mitochondrial diseases

**Why Not Relevant**: The paper is a descriptive report of the MSeqDR resource and the broader context of mitochondrial disease genetics. Nowhere in the provided text is the gene HTR3A mentioned, nor are there any patient cases, variant data, functional studies, or mechanistic discussions that involve HTR3A. The excerpts demonstrate the manuscript's focus on mitochondrial biology and genes curated in MSeqDR; although the resource synchronizes variant annotations from ClinVar and contains many nuclear genes linked to mitochondrial disease, the paper does not present any direct genetic association, case reports, or mechanistic evidence that would support the claim that HTR3A is likely associated with at least one Mendelian disease. Therefore, based on the content provided, the paper does not provide relevant evidence for or against the claim. Limitations: the paper describes a database that could, in other contexts, include many genes, but without explicit mention or listings of HTR3A or any data concerning it in this text, no inference about HTR3A’s association to Mendelian disease can be made from this paper alone.


[Read Paper](https://www.semanticscholar.org/paper/6f85c595adaca6963c30fea73bc37266fa78be56)


### Whole genome sequencing of 4,787 individuals identifies gene-based rare variants in age-related macular degeneration.

**Why Not Relevant**: This paper presents a whole-genome sequencing case-control study of age-related macular degeneration (AMD) and reports association signals and rare loss-of-function enrichment for specific genes (CFH, C2, C3, NRTN, CFI, ORMDL2). The paper text provided does not mention HTR3A anywhere and contains no direct genetic, clinical, or mechanistic data linking HTR3A to any Mendelian disease. Therefore it provides neither direct evidence (e.g., mutation-disease co-segregation, pathogenic variants in HTR3A, case reports, or association with a Mendelian phenotype) nor mechanistic evidence (e.g., functional studies, pathway connections relevant to Mendelian disease) supporting the claim that HTR3A is likely associated with a Mendelian disorder.

Limitations relevant to interpreting absence of HTR3A in this paper:
- Disease scope: The study focuses on AMD; lack of HTR3A findings here does not address HTR3A associations with other diseases or Mendelian disorders. Absence of mention in an AMD GWAS is not evidence that HTR3A is not involved in any Mendelian disease.
- Study design and power: The study analyzes common and rare variants in an AMD case-control cohort (2394 cases, 2393 controls) and uses ExAC for gene-based tests. It is not designed to discover Mendelian, highly penetrant disease genes across the genome or to evaluate segregation in families, which are typically required to establish Mendelian disease associations.
- Reported gene list: The paper explicitly reports gene-based signals in CFH, C2, C3, NRTN and enrichment of rare loss-of-function variants in CFH, CFI, and ORMDL2 (these findings are specific to AMD in this cohort). The absence of HTR3A from those reported genes only indicates it was not implicated in this AMD analysis, not that HTR3A is unassociated with Mendelian disease broadly.

Conclusion: Relevance of this paper to the claim that HTR3A is likely associated with at least one Mendelian disease is effectively zero because HTR3A is not discussed and no data in the paper bear on Mendelian associations involving HTR3A. To evaluate the claim, one should consult sources that catalog Mendelian disease genes and pathogenic variants (for example OMIM, ClinVar, ClinGen, or peer-reviewed family-based studies demonstrating segregation and pathogenicity of HTR3A variants).


[Read Paper](https://www.semanticscholar.org/paper/30bbd57cc4dd8bb9b445226f1ebfeb2aac667f53)


### Genome-Wide Association Study-Guided Exome Rare Variant Burden Analysis Identifies IL1R1 and CD3E as Potential Autoimmunity Risk Genes for Celiac Disease

**Why Not Relevant**: The provided paper focuses on identifying candidate genes for celiac disease (CeD) using whole-exome sequencing in two Arab families and network-based integration with GWAS loci. It explicitly reports "the absence of rare variants showing classical Mendelian inheritance in both families," and prioritizes IL1R1 and CD3E as candidate genes. The manuscript text contains no mention of HTR3A, no variants in HTR3A, nor any mechanistic or statistical analysis relating HTR3A to Mendelian disease. Therefore the paper provides no direct or mechanistic evidence supporting or refuting the claim that HTR3A is likely associated with at least one Mendelian disease. The lack of mention does not disprove such an association in general; it only shows that this study did not implicate HTR3A in the CeD families analyzed. Limits of generalization include small sample size (two families), WES technical limits (noncoding regions, structural variants, mosaicism), and disease focus (CeD) rather than broad Mendelian-disease discovery.


[Read Paper](https://www.semanticscholar.org/paper/109c43874acfa1bf79ede793f0e10e4f8aa2ad9e)


### Quantifying variant contributions in cystic kidney disease using national-scale whole-genome sequencing

**Why Not Relevant**: This paper focuses specifically on genetic contributors to cystic kidney disease (CyKD) and reports signals in known CyKD genes such as COL4A3 and PKHD1. The provided text contains no mention of HTR3A, no analyses of HTR3A variants, and no mechanistic discussion implicating the HTR3A gene or its protein product in CyKD or any other Mendelian condition. Therefore the paper provides no direct evidence supporting the claim that HTR3A is likely associated with at least one Mendelian disease. The absence of HTR3A in this kidney-disease-focused study does not constitute evidence that HTR3A is not associated with any Mendelian disease (absence of evidence is not evidence of absence), but it means this paper cannot be used to support the claim. Additionally, the study's phenotype specificity (CyKD) and the gene-level focus restrict generalizability: even a comprehensive CyKD analysis would not be expected to identify Mendelian associations for genes unrelated to kidney cystic phenotypes. In short, relevance to the HTR3A claim is negligible because HTR3A is neither analyzed nor discussed in the manuscript.


[Read Paper](https://www.semanticscholar.org/paper/39307f4edc822aa5abc019e0ce2a01c4d3243c95)


### Phenome-Wide Association Study of Common Genetic Variants in the DPP4 Gene and GLP1R Gene and Kidney Outcomes of European Ancestry Individuals in the Million Veteran Program (MVP)

**Why Not Relevant**: The paper's content exclusively discusses FRAS1 as the gene implicated in familial rapid renal decline and diabetic nephropathy; HTR3A is not mentioned anywhere in the provided text. Therefore the paper contains no direct genetic, epidemiological, or mechanistic evidence for or against the claim that HTR3A is associated with a Mendelian disease. Absence of evidence in this paper is not evidence of absence overall (i.e., this single paper cannot disprove an association), but it provides no support for the claim and cannot be used to evaluate HTR3A's Mendelian-disease status.


[Read Paper](https://www.semanticscholar.org/paper/8825241774f4233adb410a77772ab21c48b2f425)


### Genome-Wide and Exome-Wide Association Study Identifies Genetic Underpinning of Comorbidity between Myocardial Infarction and Severe Mental Disorders

**Why Not Relevant**: The provided paper content does not mention HTR3A anywhere. The study focuses on common-variant GWAS, Mendelian randomization for complex-trait causality, and exome-wide association in the context of comorbidity between myocardial infarction and severe mental disorders. It reports specific genes (e.g., GIGYF2, KCNJ13, PCCB, STAG1, HLA-C, HLA-B, FURIN, FES, SMG6, MUC2) but not HTR3A. The paper does not investigate Mendelian (monogenic) disease associations, family segregation, clinical genetic diagnoses, or curated pathogenic variants that would be the standard evidence for labeling a gene as causing a Mendelian disease. Absence of HTR3A in this study is not definitive disproof of a Mendelian association, but it provides no direct or mechanistic support for the claim. Additionally, the study's case ascertainment (UK Biobank participants with MI and SMDs) and analytic focus on complex-trait genetics limit its ability to detect or evaluate rare, high-penetrance Mendelian variants; such detection typically requires different study designs (family-based studies, clinical sequencing of affected pedigrees, or databases of established Mendelian gene–disease relationships). Therefore the paper is not relevant evidence for the claim that HTR3A is likely associated with at least one Mendelian disease.


[Read Paper](https://www.semanticscholar.org/paper/741e95fe69342ed2ac63d3964f83c0a8d68cc061)


## Search Queries Used

- HTR3A 5 HT3A serotonin receptor 3A mutation monogenic Mendelian disorder case report family segregating variant

- HTR3A ClinVar OMIM HGMD pathogenic variant HTR3A Mendelian disease annotation review

- HTR3A variant functional study electrophysiology receptor function loss of function gain of function knockout mouse behavioral phenotype seizure anxiety

- HTR3A whole exome sequencing rare variant family based sequencing rare disease Mendelian gene discovery genome wide association study GWAS common variant psychiatric


## Bibliometric Configuration

- Use Bibliometrics: True

- Author Impact Weight: 0.4

- Citation Impact Weight: 0.4

- Venue Impact Weight: 0.2
