## Supplementary material for "MENDELSEEK: An algorithm that predicts Mendelian Genes and elucidates what makes them special": IDH3G.txt

# Claim: The gene IDH3G is likely associated with at least one Mendelian disease.

**Status**: processed

**Overall Rating**: -1

**Explanation**:

No relevant papers were found that support or refute this claim.


## Relevant Papers


## Other Reviewed Papers


### A recurrent rare intronic variant in CAPN3 alters mRNA splicing and causes autosomal recessive limb‐girdle muscular dystrophy‐1 in three Pakistani pedigrees

**Why Not Relevant**: The paper is entirely focused on biallelic mutations in CAPN3 causing autosomal recessive limb-girdle muscular dystrophy (LGMDR1). The provided text describes identification of a homozygous CAPN3 splice-site variant (NM_000070.3: c.379 + 3A>G; p.Asp128Glyfs*15), mRNA splicing studies demonstrating partial intron retention and introduction of a premature stop codon, reduced CAPN3 expression in patient fibroblasts, and genome-wide homozygosity mapping supporting a founder effect. Nowhere in the supplied content is the gene IDH3G mentioned, nor are any data, analyses, or mechanistic links presented that connect IDH3G to Mendelian disease. Although whole-exome sequencing was used in these families (and thus could in principle detect variants in many genes, including IDH3G), the manuscript reports only CAPN3 findings and provides no negative evidence (e.g., explicit absence of pathogenic IDH3G variants) that would bear on the claim. Therefore this paper provides no direct evidence supporting or refuting the claim that IDH3G is likely associated with at least one Mendelian disease. Limitations: absence of any mention of IDH3G in the text does not prove IDH3G is not disease-associated; it only indicates that this study did not identify or report any IDH3G-related findings. The study population and analyses are specific to CAPN3-associated LGMDR1 in consanguineous Pakistani families, so results are not generalizable to assessment of other genes.


[Read Paper](https://www.semanticscholar.org/paper/6eb80e582a041fb89b352d780fb73f7a8f1034de)


### Decreased calcium permeability caused by biallelic TRPV5 mutation leads to autosomal recessive renal calcium-wasting hypercalciuria

**Why Not Relevant**: The provided paper content pertains exclusively to TRPV5 and the conclusion that biallelic loss of TRPV5 causes a novel autosomal recessive renal Ca2+-wasting hypercalciuria (RCWH). The claim to evaluate concerns IDH3G being likely associated with at least one Mendelian disease. There is no mention of IDH3G in the provided text, no genetic, clinical, or mechanistic data relating IDH3G to disease, and no discussion of any genes other than TRPV5. Because the gene names and described biological mechanism (TRPV5-mediated renal calcium transport) are different from IDH3G (which encodes a subunit of isocitrate dehydrogenase in the TCA cycle), the excerpt offers neither direct evidence (e.g., variants, segregation, case reports) nor mechanistic evidence (shared pathway, protein interaction, functional consequences) to support or refute an association between IDH3G and a Mendelian disorder. Consequently, relevance to the claim is effectively zero based on the provided content. To evaluate the claim about IDH3G would require different evidence: reports of pathogenic IDH3G variants segregating with disease, case series, population burden analyses, functional assays showing variant impact on IDH3G function, or mechanistic links between IDH3G perturbation and a Mendelian phenotype.


[Read Paper](https://www.semanticscholar.org/paper/eed13d61e621b499f021b1f84abbc8811a578c04)


### A bird’s eye view on the use of whole exome sequencing in rare congenital ophthalmic diseases

**Why Not Relevant**: The provided paper content contains only a single sentence stating that WES expands the analytical portfolio for ocular dysgenesis and is useful for differential diagnosis in ASD. It does not mention IDH3G, any variants in IDH3G, patient phenotypes linked to IDH3G, segregation or inheritance data, functional studies, or mechanistic pathways involving IDH3G. Therefore there is no direct evidence in this text that IDH3G is associated with a Mendelian disease. Mechanistic evidence is also absent: the excerpt does not discuss metabolic pathways, isocitrate dehydrogenase function, mitochondrial biology, or any cellular processes that could link IDH3G to a disease phenotype. Because the content lacks gene-specific information and necessary evidence types (variant identification, co-segregation, functional validation, statistical association), it is insufficient to support or refute the claim that IDH3G is likely associated with at least one Mendelian disease.


[Read Paper](https://www.semanticscholar.org/paper/2dcaaa7df6f96f5bc69eede8888aebfdab24b8b7)


### Whole exome sequencing revealed a novel homozygous variant in the DGKE catalytic domain: a case report of familial hemolytic uremic syndrome

**Why Not Relevant**: The provided paper content discusses DGKE and a putative DGKE variant (p.Asn314Lys) with respect to aHUS. The claim to evaluate is about the gene IDH3G being likely associated with at least one Mendelian disease. Because the excerpt names a different gene (DGKE) and contains no mention of IDH3G, it provides no direct evidence for or against an association between IDH3G and any Mendelian disease. Mechanistically, while the excerpt illustrates a general principle (catalytic-domain variants can alter protein structure and potentially cause disease), that generic principle cannot be used to infer that IDH3G is disease-associated without additional specific evidence (e.g., pathogenic variants in IDH3G, functional data, segregation, population and clinical data). Additionally, the sentence is speculative and lacks experimental detail, statistical evidence, and contextual data that might allow cautious extrapolation. For these reasons the relevance of this paper content to the claim about IDH3G is negligible (relevance = 0.0).


[Read Paper](https://www.semanticscholar.org/paper/2691c9dc3e9f16f506828ae781284158953cd6a0)


### Familial juvenile polyposis syndrome with a de novo germline missense variant in BMPR1A gene: a case report

**Why Not Relevant**: The provided paper content does not mention the gene IDH3G anywhere and instead reports a de novo germline missense variant in BMPR1A in a family with juvenile polyposis. Because the text concerns a different gene (BMPR1A) and provides no mechanistic or associative information about IDH3G, it cannot support or refute the claim about IDH3G's association with any Mendelian disease. There is no direct evidence (no IDH3G variants, no phenotype linkage) and no mechanistic pathway connecting BMPR1A findings to IDH3G in the provided content. Therefore relevance to the claim is effectively zero. Additionally, the excerpt is a brief clinical summary without variant-level data, functional studies, segregation analysis, or population/penetrance statistics that might allow indirect inference about other genes.


[Read Paper](https://www.semanticscholar.org/paper/02b7c910bb70e933d6994ab70b31e03d7c422449)


### Novel mutation in the TGFBI gene in a Moroccan family with atypical corneal dystrophy: a case report

**Why Not Relevant**: The provided paper content discusses a novel mutation in the TGFBI gene associated with Thiel-Behnke corneal dystrophy and a genotype–phenotype correlation for that family. It contains no mention of the gene IDH3G, no data on IDH3G variants, no mechanistic discussion linking IDH3G to disease, and no genetic, statistical, or clinical evidence concerning IDH3G. Therefore the paper provides neither direct evidence (e.g., reported pathogenic variants, segregation, functional assays) nor mechanistic evidence (e.g., biochemical role, pathway disruption) that IDH3G is associated with any Mendelian disease. 

Limitations and context: the absence of IDH3G-related information in this paper does not disprove that IDH3G could be disease-associated in other studies, but this paper is irrelevant to the claim because it focuses exclusively on TGFBI. The single sentence provided lacks methodological details, sample size, pedigree data, variant nomenclature, functional assays, or any other elements that could be reinterpreted to support or refute the IDH3G–Mendelian disease claim.


[Read Paper](https://www.semanticscholar.org/paper/f901921467669d44be4f58c9350f9f390c9bf428)


### The Clinicopathologic and Genetic Characteristics of Autosomal Recessive Polycystic Kidney Disease Presenting in Adulthood

**Why Not Relevant**: The provided paper excerpt concerns autosomal recessive polycystic kidney disease (ARPKD) and reports biallelic pathogenic variants in the PKHD1 gene in several families. The text makes no mention of the gene IDH3G, nor any data, results, or mechanistic discussion involving IDH3G. Although the Methods note that whole exome sequencing (WES) was performed — a technique that could in principle detect variants in IDH3G — the Results and Background report only PKHD1 findings. Therefore this paper provides no direct evidence that IDH3G is associated with any Mendelian disease, and it provides no mechanistic evidence bearing on IDH3G. Limitations: the excerpt may be partial and other sections (e.g., Supplementary Data) could conceivably mention other genes, but based on the content provided there is no information relevant to the claim. Absence of evidence in this paper is not evidence that IDH3G is not disease-associated; it simply means this paper cannot be used to support or refute the claim.


[Read Paper](https://www.semanticscholar.org/paper/d7e39e24c54c37f51b033e406dfe95e686b2e857)


### Whole‐exome sequencing analyses in a Saudi Ischemic Stroke Cohort reveal association signals, and shows polygenic risk scores are related to Modified Rankin Scale Risk

**Why Not Relevant**: The supplied paper content does not mention the gene IDH3G, Mendelian disease associations, or any gene-specific findings that could be linked to IDH3G. The single provided sentence: "Gene burden testing in Saudi stroke populations reveals a number of statistically significant signals under different disease inheritance models and, interestingly, stroke subjects with mRS of 3 or lower had lower cumulative genetic risk from rare variants in stroke genes and therefore, determining the potential mRS cutoffs to use for clinical significance may allow risk stratification of this population." is a general summary about burden testing in stroke genes and mRS-based stratification and contains no direct evidence (no gene names, no variant-level data, no phenotype linking) nor mechanistic information about IDH3G. Therefore the excerpt provides neither direct nor mechanistic evidence for or against the claim that IDH3G is likely associated with at least one Mendelian disease.

Limitations and caveats: (1) Only a single short paragraph was provided; it is possible that other sections of the full paper (methods, results, supplementary tables) might list gene-level associations that include IDH3G. Those sections were not supplied, so this assessment is limited to the provided content. (2) The provided sentence refers broadly to "stroke genes" and "statistically significant signals" without specifying genes or variants, so it cannot be used to infer associations for any specific gene. (3) To evaluate the claim about IDH3G, one would need gene-specific association results, reported pathogenic variants, clinical segregation data, or independent entries from disease databases (e.g., OMIM, ClinVar, HGMD), none of which are present in the excerpt.

If you can provide the full paper or the specific results/tables listing significant genes or variants, I can re-assess for direct or mechanistic evidence regarding IDH3G. Alternatively, I can search external genetic-disease databases and literature for existing evidence linking IDH3G to Mendelian disease if you would like.


[Read Paper](https://www.semanticscholar.org/paper/ef1e3cc537505def74437c2048ef722316d024a6)


## Search Queries Used

- IDH3G mutation Mendelian disease case report familial segregation pathogenic variant

- IDH3G variant whole exome sequencing rare disease cohort monogenic biallelic autosomal recessive

- isocitrate dehydrogenase 3 gamma subunit IDH3G loss of function enzyme deficiency mitochondrial dysfunction Krebs cycle model organism knockout phenotype

- IDH3G gnomAD constraint pLI gene intolerance ClinVar OMIM gene disease association rare variant burden IDH3A IDH3B


## Bibliometric Configuration

- Use Bibliometrics: True

- Author Impact Weight: 0.4

- Citation Impact Weight: 0.4

- Venue Impact Weight: 0.2
