## Supplementary material for "MENDELSEEK: An algorithm that predicts Mendelian Genes and elucidates what makes them special": IER3.txt

# Claim: The gene IER3 is likely associated with at least one Mendelian disease.

**Status**: processed

**Overall Rating**: -1

**Explanation**:

No relevant papers were found that support or refute this claim.


## Relevant Papers


## Other Reviewed Papers


### The Burden of Candidate Pathogenic Variants for Kidney and Genitourinary Disorders Emerging From Exome Sequencing

**Why Not Relevant**: The provided paper content describes methods and scope for identifying and evaluating genes associated with Mendelian kidney and genitourinary disorders and the variant-filtering pipelines used, but it does not mention the gene IER3 by name, nor does it provide gene-level results that would indicate whether IER3 is associated with a Mendelian disease. All excerpts are methodological or describe the curated gene list generically (counts and annotation approach). Therefore, based solely on the text supplied, there is no direct or indirect evidence that IER3 is associated with at least one Mendelian disease. It remains possible that IER3 is included in the study's final list of 625 genes (the authors note the list is available on the laboratory website), but that information is not present in the provided content; without the explicit listing or reported results for IER3, the paper cannot be used as evidence for the claim.


[Read Paper](https://www.semanticscholar.org/paper/a75e297208c04992568e607d19aa9b9aaebd09d5)


### Disruption of PHF21A causes syndromic intellectual disability with craniofacial anomalies, epilepsy, hypotonia, and neurobehavioral problems including autism

**Why Not Relevant**: The provided content refers exclusively to PHF21A, not IER3. Example verbatim sentence from the content: "Deleterious nonsense, frameshift, and missense mutations disrupting the AT Hook domain and/or an intrinsically disordered region in PHF21A were found to be associated with autism spectrum disorder, epilepsy, hypotonia, neurobehavioral problems, tapering fingers, clinodactyly, and syndactyle, in addition to intellectual disability and craniofacial anomalies." This sentence documents associations between mutations in PHF21A and a range of neurodevelopmental and physical phenotypes, but it makes no mention of IER3 or any data implicating IER3 in Mendelian disease. Therefore the paper content provided offers neither direct evidence (no reported IER3 variants, phenotypes, segregation, or case reports) nor mechanistic evidence (no functional studies, pathways, or interactions involving IER3) to support or refute the claim that IER3 is likely associated with at least one Mendelian disease. Limitations: the assessment is based solely on the single sentence supplied; if the full paper includes other sections mentioning IER3, those were not provided and were not assessed. If you can supply the full text or sections that mention IER3, I can re-evaluate for direct and mechanistic evidence.


[Read Paper](https://www.semanticscholar.org/paper/87c951a7e4271d50548982a10337a6ca2a1ee1fc)


### How autophagy, a potential therapeutic target, regulates intestinal inflammation

**Why Not Relevant**: The provided paper is a review of autophagy in inflammatory bowel disease (IBD) and discusses specific autophagy-related genes (ATG16L1, IRGM, NOD2, LRRK2) and pathways relevant to intestinal homeostasis and IBD pathogenesis. The text contains no mention of the gene IER3, nor any discussion of Mendelian disease associations for IER3. Therefore the paper provides no direct genetic evidence (case reports, linkage, pathogenic variants) nor mechanistic evidence linking IER3 to any Mendelian disorder. Important caveats: absence of IER3 in this IBD/autophagy-focused review is not definitive proof that IER3 has no Mendelian disease associations elsewhere; it only indicates this paper does not address that claim. To evaluate the claim properly would require literature or database searches specifically for IER3 (e.g., OMIM, ClinVar, HGMD, gene-focused case reports) or functional studies tying IER3 mutations to Mendelian phenotypes.


[Read Paper](https://www.semanticscholar.org/paper/5b74665535de17e374e09d2f0d020311e834ac42)


### Endoplasmic reticulum stress response and transcriptional reprogramming

**Why Not Relevant**: This paper does not provide direct evidence that the gene IER3 is associated with any Mendelian disease. The content is an overview/collection focused on endoplasmic reticulum stress, unfolded protein response (UPR), and transcriptional reprogramming. The gene name IER3 does not appear anywhere in the supplied text. The paper discusses IRE1α (inositol-requiring 1α) and related mechanisms (XBP1 splicing, RIDD), but these are distinct from IER3 (Immediate Early Response 3) as commonly annotated in human genetics. One sentence in the text contains "IER1α RNase," which appears to be a typographic or nomenclature inconsistency relative to the earlier, correctly spelled "IRE1α"; this should not be interpreted as referring to IER3. Importantly, establishing a Mendelian disease association requires gene-specific genetic evidence (e.g., segregation in families, pathogenic variants, case reports, or statistically significant burden in affected individuals)—none of which is present in this text. The paper provides mechanistic context showing that ER stress pathways can contribute to disease broadly, but that is insufficient to infer that IER3 is likely causative of a Mendelian disorder. For these reasons the relevance to the specific claim is negligible.


[Read Paper](https://www.semanticscholar.org/paper/3ad2b1517e1a08cb2e655453d962feeddaae4a5a)


### Cases in Precision Medicine: When Patients Present With Direct-to-Consumer Genetic Test Results

**Why Not Relevant**: This paper is about direct-to-consumer (DTC) genetic testing (23andMe), its limitations, clinician guidance, and regulatory history. Although it names several specific Mendelian conditions (for example, Bloom syndrome, Gaucher disease type 1, cystic fibrosis, hereditary hemochromatosis) in the context of tests that 23andMe offers or that clinicians might encounter, the paper contains no mention of the gene IER3, no association studies, no case reports, no segregation or linkage data, and no functional/mechanistic experiments related to IER3. Therefore it provides neither direct evidence (genetic association, case-series, statistical association) nor mechanistic evidence (biochemical function, pathogenic variants causing disease phenotypes, animal models) that would support or refute the claim that IER3 is likely associated with at least one Mendelian disease. Methodological details relevant to gene-disease association (cohort description, sequencing/variant-calling pipelines, statistical tests, functional assays) are absent for IER3 specifically. Because the paper does not address IER3 in any way, its content cannot meaningfully inform the claim. 

What would be needed to assess the claim: evidence such as (1) segregation of rare pathogenic variants in IER3 with a consistent Mendelian phenotype in multiple families; (2) statistically significant enrichment of (likely) pathogenic IER3 variants in affected individuals versus controls; (3) curated clinical assertions (ClinVar) or an OMIM entry linking IER3 to a Mendelian disorder; and/or (4) functional data showing that disease-associated IER3 variants perturb gene/protein function in a manner consistent with the phenotype (cellular assays, animal models). None of these evidence types appear in the provided paper content.


[Read Paper](https://www.semanticscholar.org/paper/b8f37dd6a7da030eade004848c55b6838eab4965)


### Patients with coronary heart disease, dilated cardiomyopathy and idiopathic ventricular tachycardia share overlapping patterns of pathogenic variation in cardiac risk genes

**Why Not Relevant**: Relevance is set to 0.0 because the supplied paper content contains no direct mention of the gene IER3, no reported variants in IER3, and no mechanistic discussion connecting IER3 to a Mendelian disease. The paper reports results from a targeted cardiac-gene panel and names multiple genes found to carry pathogenic or likely pathogenic variants (e.g., LAMA2, MYBPC3, TTN, GAA, KCNQ1, DSG2, MYH6) but IER3 is not listed in the provided text. Important caveats: (1) The absence of IER3 in the supplied excerpts does not prove that IER3 was not included in the 96-gene panel — the full gene list and full variant tables are not provided here. (2) Even if IER3 were included but yielded no notable variants in this 92-patient cardiac cohort, that would be limited negative evidence only for Mendelian associations relevant to ventricular tachycardia or cardiomyopathy in this population; it would not rule out IER3 involvement in other Mendelian diseases or in other cohorts. (3) Establishing that a gene is "likely associated with at least one Mendelian disease" generally requires gene-level evidence such as multiple unrelated pathogenic variants with consistent phenotypes, segregation in families, functional/biochemical data, or curation in databases (ClinVar, OMIM). None of these IER3-specific data are present in the provided text. Therefore, based on the content supplied, this paper does not provide evidence supporting or refuting the claim regarding IER3.


[Read Paper](https://www.semanticscholar.org/paper/3f3b535d385c9b16f6a599120d890be36b90f4b6)


### Deoxynivalenol-Induced Spleen Toxicity in Mice: Inflammation, Endoplasmic Reticulum Stress, Macrophage Polarization, and the Dysregulation of LncRNA Expression

**Why Not Relevant**: This paper does not provide evidence relevant to the claim that IER3 is likely associated with at least one Mendelian disease. The manuscript focuses on deoxynivalenol (DON) toxicity in mouse spleen, reporting inflammation, macrophage M1 polarization, ER stress (PERK-eIF2α-ATF4), apoptosis (caspase-12, caspase-3), effects of the ER stress inhibitor 4-Phenylbutyric acid, and broad transcriptome changes (numbers of differentially expressed lncRNAs and mRNAs and enriched immune pathways). The provided text contains no mention of IER3 (the gene name), no analysis of specific genes linked to human Mendelian disorders, no genetic variant, segregation, case-control, or clinical phenotype data, and no functional follow-up that would demonstrate causality of rare variants in disease. Differential expression in a mouse toxin exposure model does not by itself establish a gene’s involvement in Mendelian disease; establishing such an association requires human genetic evidence (rare pathogenic variants segregating with disease, de novo variants, multiple unrelated affected individuals, or well-controlled functional studies demonstrating that disease-causing variants alter gene function in a manner consistent with the phenotype).

Relevant verbatim context from the paper (showing scope and why it is not relevant):
"The spleen is a primary target of deoxynivalenol (DON) toxicity, but its underlying molecular mechanisms remain unclear. This study investigates the effects of DON on inflammation, splenic macrophage polarization, endoplasmic reticulum (ER) stress, and transcriptome changes (mRNA and lncRNAs) in mouse spleen." 
"Transcriptome analysis identified 1968 differentially expressed (DE) lncRNAs and 2664 DE mRNAs in mouse spleen following DON exposure. Functional enrichment analysis indicated that the upregulated genes were involved in pathways associated with immunity, including Th17 cell differentiation, TNF signaling, and IL-17 signaling, while downregulated mRNAs were linked to cell survival and growth pathways." 
"This study provides new insights into the mechanisms underlying DON toxicity and its effects on the immune system."

Why these statements matter for relevance: they demonstrate the study design (toxin exposure in mouse spleen, transcriptome-wide differential expression, pathway enrichment) and show that the work is not examining Mendelian disease genetics or specific gene–disease causality. Even if IER3 (or its mouse ortholog) were among the DE mRNAs (the paper does not provide a gene list in the provided text), that would at most be mechanistic evidence of stress-responsive expression, not evidence that variants in IER3 cause a Mendelian disorder.

Limitations preventing use of this paper to support the claim:
- No mention of IER3 by name in the provided content.
- No human genetic data, no variant analyses, no familial segregation, no case reports, and no disease association statistics.
- Mouse toxin exposure and differential expression do not equate to Mendelian disease causation.
- Pathway enrichment and ER stress/apoptosis findings are mechanistic for DON toxicity but do not link a given gene to heritable monogenic disease.

Conclusion and recommended next steps to evaluate the claim: this paper is not relevant to establishing whether IER3 is associated with a Mendelian disease. To assess the claim, consult human genetics resources (OMIM, ClinVar, HGMD), population sequencing databases (gnomAD), and published human genetic studies reporting pathogenic/likely pathogenic variants in IER3 with segregation or case series. Functional studies that test human IER3 variants for loss- or gain-of-function consistent with a disease phenotype would also be needed.


[Read Paper](https://www.semanticscholar.org/paper/34f96364586f4c785253362d7fc89ad36d7cfdba)


### Differences in Expression of IQSEC2 Transcript Isoforms in Male and Female Cases with Loss of Function Variants and Neurodevelopmental Disorder

**Why Not Relevant**: The provided paper text exclusively discusses the gene IQSEC2: its pathogenic hemizygous or heterozygous mutations, clinical consequences (XLID1), case reports with de novo IQSEC2 variants, and mechanistic evidence (an intronic variant creating an alternative splice site). There is no mention of the gene IER3 anywhere in the text. Therefore the paper provides no direct evidence (no reported variants, cases, or association) and no mechanistic evidence (no functional data, splicing or expression effects, or pathway links) related to IER3. Because the genes are distinct, evidence about IQSEC2 cannot be taken as evidence for IER3 without independent data demonstrating a specific relationship (e.g., genetic linkage, protein interaction, shared pathway with demonstrated causality, or homology with established relevance). In short, this paper does not support or refute the claim that IER3 is associated with a Mendelian disease; it is simply not relevant to IER3.


[Read Paper](https://www.semanticscholar.org/paper/d14a9429bb9a71e15141ff7330f8bd5e699cf638)


### The multiple faces of TOR1A: different inheritance, different phenotype.

**Why Not Relevant**: The provided paper excerpt contains general background about Mendelian versus complex disorders, discussion of variable expressivity and incomplete penetrance, and a specific example involving TOR1A. Nowhere in the text is the gene IER3 mentioned, nor are there any experimental data, case reports, sequencing results, segregation analyses, functional assays, or mechanistic studies concerning IER3. Therefore there is no direct evidence that IER3 is associated with any Mendelian disease in this content. The excerpt only offers conceptual and analogical context (showing how other genes can be linked to Mendelian disorders), which affords minimal indirect support for the *possibility* that any given gene might be disease-associated but provides no specific evidence for IER3. Because of the complete absence of IER3-specific information, the relevance to the claim that “IER3 is likely associated with at least one Mendelian disease” is very low.


[Read Paper](https://www.semanticscholar.org/paper/b74b91e0829b14562a5293323e09bb3633156aab)


### Clingen Coagulation Factor Deficiency Variant Curation Expert Panel: Meeting the Need for Recommendations to Curate Variants in the Coagulation Factor Genes

**Why Not Relevant**: This paper does not provide evidence for or against the claim that IER3 is associated with a Mendelian disease. The content exclusively describes an effort to specify ACMG/AMP variant-interpretation guidelines for coagulation factor genes, and repeatedly and explicitly names F8 and F9 as the genes under consideration (e.g., “The first two genes under the purview of CFD-VCEP are F8 (OMIM: 300841) and F9 (OMIM: 300746).” and “Pathogenic variants in the F8 and F9 genes resulting in the loss of protein function cause Hemophilia A and B, respectively.”). There is no mention of the gene IER3 anywhere in the provided text, no reported variants in IER3, no clinical cases involving IER3, and no mechanistic discussion that links IER3 to coagulation or to any Mendelian phenotype. The paper is a guideline/consensus document focused on variant curation workflows and gene-specific specification of ACMG/AMP criteria for coagulation-factor genes; it contains no experimental data, case reports, or literature review relevant to IER3. Because direct evidence (case reports, segregation, pathogenic variants) and mechanistic evidence (biological role, pathway links) about IER3 are entirely absent from the manuscript content provided, this source cannot be used to support or refute the claim. To evaluate the claim about IER3 and Mendelian disease association, one would need to consult other sources (e.g., OMIM, ClinVar, peer-reviewed case reports, functional studies) that explicitly mention IER3 and any disease associations. The absence of IER3 in this coagulation-focused guideline paper is a fundamental limitation preventing any inference regarding IER3’s involvement in Mendelian disease.


[Read Paper](https://www.semanticscholar.org/paper/9ea8d4ce19622ae1c6b0cdfd40848c93199f6280)


### NGS based studies on primary immunodeficiencies (PIDs) : causative gene identification, tool development and application

**Why Not Relevant**: The paper focuses on genetic findings in primary immunodeficiency disorders (PIDs) using WES/TRS and reports specific genes and candidate novel PID genes. Nowhere in the provided text is the gene name IER3 mentioned, nor are variants in IER3, pathogenic mechanisms involving IER3, or clinical cases linked to IER3 discussed. Therefore this paper provides no direct evidence that IER3 is associated with any Mendelian disease, nor mechanistic evidence supporting such an association. Caveats: absence of mention in this PID-focused cohort is not definitive disproof that IER3 can be associated with a Mendelian disease—IER3 could be implicated in other disease classes, in cohorts not studied here, or missed due to filtering, limited sample size for extremely rare disorders, or phenotype-driven inclusion criteria. To evaluate the claim further, examination of disease databases (OMIM, ClinVar), population variant resources (gnomAD), and the broader literature (PubMed) for IER3-specific reports is recommended.


[Read Paper](https://www.semanticscholar.org/paper/7786e43fbaf68904111f286747f0cf8968c2dd1e)


### Mapping genetic variations to three-dimensional protein structures to enhance variant interpretation: a proposed framework

**Why Not Relevant**: The provided paper is a methods/workshop/review-style article about integrating genetic variant data with 3D protein structures and tools for variant interpretation. It does not mention the gene IER3, any variants in IER3, or any Mendelian disease associations. Therefore it contains no direct evidence supporting or refuting the claim that "IER3 is likely associated with at least one Mendelian disease." The excerpts above show mechanistic and methodological context relevant to how one would evaluate such a claim, but they do not constitute gene-specific or disease-specific evidence. Because the paper contains no IER3-specific data, the relevance to the specific claim is very low.


[Read Paper](https://www.semanticscholar.org/paper/ae68aa934d5a5ad3d9009bb8811fa8d5fc4dbb1c)


### Travelling to Rome : inflammation, endoplasmic reticulum stress and angiogenesis during atherosclerotic plaque development

**Why Not Relevant**: This paper does not provide evidence for or against the claim that IER3 is likely associated with at least one Mendelian disease. The provided dissertation text focuses on atherosclerosis, immune cells (iNKT cells, CD1d-expressing APCs), angiogenesis, endothelial dysfunction, ER stress, the unfolded protein response (UPR), and T-cadherin. There is no mention of the gene IER3, no human genetic data, no descriptions of pathogenic variants, no family segregation or de novo mutation evidence, and no statements linking any gene to a Mendelian phenotype. Representative verbatim sentences from the paper showing the topics covered (verbatim from the provided content) include: "Cardiovascular diseases are the leading cause of death worldwide followed by cancer." "Unfolded protein response (UPR) is the front line of defense during ER stress, aiming to re-establish cellular homeostasis and rescue the cell from apoptosis." "During the development of an atherosclerotic plaque, lesions are infiltrated by inflammatory cells and professional antigen presenting cells (APCs)." "Upregulation of T-cadherin expression on EC during ER stress attenuates the activation of the proapoptotic PERK (PKR (double-stranded RNA-activated protein kinase)-like ER kinase) branch of the UPR cascade and thereby protects EC from ER stress-induced apoptosis."

How this bears on the claim:
- Direct evidence: None. The paper contains no direct mention of IER3, no mutation data, no case reports, no linkage/segregation analyses, and no population genetic evidence linking IER3 to a Mendelian disorder.
- Mechanistic evidence: Limited/indirect at best. The paper discusses ER stress, UPR, apoptosis, and inflammatory pathways (mechanisms that some genes implicated in Mendelian diseases may also affect). However, without any explicit discussion of IER3, shared pathway involvement cannot be used to infer that IER3 is causally or likely associated with a Mendelian disease. Mechanistic plausibility alone (i.e., that a gene participates in pathways relevant to disease biology) is insufficient to establish a Mendelian gene–disease relationship; one requires genetic evidence (segregation, recurrent pathogenic variants, de novo events, strong case-control enrichment, functional validation of patient variants, etc.).

Limitations and recommended next steps:
- Because IER3 is not mentioned, this document cannot inform the claim. To evaluate the claim properly, search sources of genetic disease associations and variant curation such as OMIM, ClinVar, HGMD (if available), PubMed for "IER3 mutation" or "IER3 disease", and population/constraint databases (gnomAD) for intolerance metrics. Look for case reports, cosegregation studies, de novo variant reports in trios, and functional studies of patient variants to support a Mendelian association.


[Read Paper](https://www.semanticscholar.org/paper/085d9af306c2c5651cfcf0975fb03d4f4c2494e3)


### ADAR1 plays a protective role in proximal tubular cells under high glucose conditions by attenuating the PI3K/AKT/mTOR signaling pathway

**Why Not Relevant**: This paper does not provide any evidence — direct or mechanistic — relevant to the claim that IER3 is likely associated with at least one Mendelian disease. The content supplied concerns ADAR1 expression and function in diabetic db/db mice and high-glucose-treated HK-2 cells, describing experiments on ADAR1 overexpression and inhibition and effects on endoplasmic reticulum stress, apoptosis, and the PI3K/AKT/mTOR pathway. Nowhere in the provided text is the gene IER3 mentioned, nor are there any genetic-association data, pedigree analyses, reports of pathogenic variants, case reports, or references to Mendelian inheritance or Mendelian disease databases (e.g., OMIM). The experimental approaches described (mouse model expression analysis, HK-2 cell high-glucose injury, plasmid overexpression, pharmacologic inhibition) are molecular/functional and do not constitute evidence for a gene–Mendelian disease association in humans. Because there is no mention of IER3, no mechanistic pathway tying IER3 to disease is presented, and no human genetic data are reported, the paper provides no support for or against the claim. Limitations that further reduce any indirect relevance: (1) species and model-specific results (db/db mice, HK-2 cells) without human genetic correlation; (2) absence of variant-level or inheritance data required to establish Mendelian associations; and (3) focus on a different gene (ADAR1) and signaling pathways (PI3K/AKT/mTOR), so any extrapolation to IER3 would be purely speculative and unsupported by the text. Because relevance is below 0.1, no verbatim excerpts mentioning IER3 or Mendelian disease are available from the provided paper content.


[Read Paper](https://www.semanticscholar.org/paper/32764f989ab0d4678a0b6744b7b594c87a15f792)


### Consolidating the Role of Mutated ATP2B2 in Neurodevelopmental and Cerebellar Pathologies

**Why Not Relevant**: The provided paper content discusses ATP2B genes (specifically ATP2B2 encoding PMCA2) and their association with a human neurodevelopmental syndrome. The abstract's text refers exclusively to ATP2B/PMCA2 (e.g., "ATP2B2 (encoding PMCA2)", "ATP2B2-related condition", "ATP2B2 variants") and does not mention the gene IER3 anywhere. Because IER3 is not named or discussed, there is no direct evidence (genetic, clinical, or statistical) relating IER3 to any Mendelian disease in this paper, nor is there any mechanistic discussion that could plausibly connect findings about ATP2B2/PMCA2 to IER3. Findings that ATP2B2 is implicated in a Mendelian neurodevelopmental syndrome cannot be extrapolated to support the claim about IER3 without additional data. Therefore this paper does not provide relevant direct or mechanistic evidence for or against the claim that IER3 is likely associated with at least one Mendelian disease.


[Read Paper](https://www.semanticscholar.org/paper/18af3e328d760f2e7be9187299c0243a7d49beec)


## Search Queries Used

- IER3 IEX1 immediate early response 3 mutation variant pathogenic Mendelian monogenic disease case report family exome sequencing

- IER3 ClinVar OMIM HGMD gene disease association pathogenic variant clinical significance IER3 gene

- IER3 IEX1 apoptosis cell survival endoplasmic reticulum stress immune regulation inflammation pathway mechanism human disease phenotype

- IER3 loss of function truncating variant haploinsufficiency knockout mouse phenotype developmental disorder trio exome sequencing de novo neurodevelopmental intellectual disability epilepsy


## Bibliometric Configuration

- Use Bibliometrics: True

- Author Impact Weight: 0.4

- Citation Impact Weight: 0.4

- Venue Impact Weight: 0.2
