## Supplementary material for "MENDELSEEK: An algorithm that predicts Mendelian Genes and elucidates what makes them special": INHBB.txt

# Claim: The gene INHBB is likely associated with at least one Mendelian disease.

**Status**: processed

**Overall Rating**: -1

**Explanation**:

No relevant papers were found that support or refute this claim.


## Relevant Papers


## Other Reviewed Papers


### Rare Variants in Tissue Inhibitor of Metalloproteinase 2 as a Risk Factor for Schizophrenia: Evidence From Familial and Cohort Analysis

**Why Not Relevant**: Relevance is near zero because the paper focuses on TIMP2 (Tissue Inhibitor of Metalloproteinase 2) and its possible role in schizophrenia. The gene named in the claim, INHBB, is not mentioned in the provided content. Therefore this paper provides no direct genetic, mechanistic, or epidemiological evidence linking INHBB to any Mendelian disease. All extracted excerpts pertain to TIMP2 and SZ; none support or refute associations between INHBB and Mendelian disorders. Limitations: absence of any mention of INHBB in the text makes it impossible to draw any inference about INHBB from this paper.


[Read Paper](https://www.semanticscholar.org/paper/01f2a0354c9dab947d005e4c30f7a22f1f502c2b)


### Genome-wide DNA methylation profiling and exome sequencing resolved a long-time misdiagnosed case

**Why Not Relevant**: The provided paper content contains only a single, general sentence about the usefulness of combining genomic and epigenomic strategies with phenotype analysis for diagnosing rare genetic disorders. It does not mention the gene INHBB, any specific variants, patient cases, segregation or inheritance data, pathogenicity assessments, functional studies, or mechanistic pathways involving INHBB. Because none of the required types of evidence (direct association data, mechanistic plausibility specific to INHBB, or case/variant-level information) are present, the content cannot be used to support or refute the claim that INHBB is likely associated with at least one Mendelian disease.


[Read Paper](https://www.semanticscholar.org/paper/88a966ac3495d9dcafa55433a8d3b45cfbe85576)


### Chung–Jansen syndrome can mimic Cornelia de Lange syndrome: Another player among chromatinopathies?

**Why Not Relevant**: The paper exclusively discusses a familial CdLS-like presentation and the identification of a likely pathogenic variant in PHIP (c.40G > A (p.Glu14Lys)) associated with Chung-Jansen syndrome. There is no mention of the gene INHBB in the provided content, no experimental data, clinical observations, mechanistic discussion, or genetic association involving INHBB, and therefore no direct or mechanistic evidence in this paper that bears on whether INHBB is associated with any Mendelian disease. Because the study's findings pertain to a different gene (PHIP) and disorder, the paper does not provide support for or against the claim about INHBB. Additionally, the evidence here is limited to a small familial report for PHIP and cannot be extrapolated to infer associations for unrelated genes such as INHBB.


[Read Paper](https://www.semanticscholar.org/paper/6b47ca6e79752c41120ad9aab71071ea8a96ad1b)


### Phenotypic Differences in Multiple Osteochondromas in Monozygotic Twins: A Case Report

**Why Not Relevant**: The paper focuses entirely on hereditary multiple exostoses (MO/HME) and presents genetic evidence implicating EXT1 and EXT2 as causal genes (including a specific EXT1 frameshift mutation). There is no mention of the gene INHBB anywhere in the provided text, no experiments or analyses involving INHBB, and no mechanistic discussion linking INHBB to MO or any other Mendelian condition. Therefore this paper provides no direct or mechanistic evidence that INHBB is associated with any Mendelian disease. Absence of INHBB in this disease-focused study does not prove INHBB is not associated with any Mendelian disorder, but it means this paper is irrelevant to supporting the claim that INHBB is likely associated with at least one Mendelian disease.


[Read Paper](https://www.semanticscholar.org/paper/6e18bac0a7ae14f6064092c786e37feb59c64d13)


### Autosomal Dominant Intellectual Development Disorder-6 (MRD6) Without Seizures Linked to a De Novo Mutation in the grin2b Gene Revealed by Exome Sequencing: A Case Report of a Moroccan Child

**Why Not Relevant**: Relevance is near zero because the paper exclusively discusses GRIN2B (grin2b) and a GRIN2B-associated Mendelian disorder (MRD6). The text contains multiple direct and mechanistic pieces of evidence that GRIN2B mutations cause an autosomal dominant intellectual developmental disorder (specific pathogenic variant, de novo segregation, absence from population databases, ACMG classification, predicted deleterious effect), but it contains no mention of the gene INHBB, no variants in INHBB, no clinical associations involving INHBB, and no mechanistic links (e.g., shared pathways, protein interactions, or homology) connecting INHBB to the phenotype or to GRIN2B. Therefore this paper provides no direct or indirect evidence supporting the claim that INHBB is associated with any Mendelian disease. To support the claim about INHBB one would need explicit data (genetic variants in INHBB segregating with disease, case reports, functional studies, population absence/enrichment data, or authoritative database annotation), none of which are present in this text.


[Read Paper](https://www.semanticscholar.org/paper/67d756e0cc302e5d46fdda9df34403f1f2de9a5a)


### Case report: Preimplantation genetic testing for infantile GM1 gangliosidosis

**Why Not Relevant**: This paper provides no direct or mechanistic evidence regarding the gene INHBB. The study exclusively concerns GM1 gangliosidosis and pathogenic variants in the GLB1 gene. Relevant verbatim sentences from the paper that demonstrate this focus include: "Ganglioside-monosialic acid (GM1) gangliosidosis (ICD-10: E75.1; OMIM: 230500, 230600, 230650) is a rare autosomal recessive hereditary disease, lysosomal storage disorder caused by mutations in the GLB1 gene that lead to the absence or insufficiency of β-galactosidase." and "The analysis revealed two variants in the heterozygous state: a frameshift variant c.699delG (rs1452318343, ClinVar ID 928700) in exon 6 and a missense variant c.809A>C (rs371546950, ClinVar ID 198727) in exon 8 of the GLB1 gene." There is no mention of INHBB (inhibin beta B) or any experiments, observations, or mechanistic discussion involving INHBB in the provided text. Therefore, this paper neither supports nor refutes the claim that "INHBB is likely associated with at least one Mendelian disease." Absence of evidence in this paper does not constitute evidence against the claim in general; it only means this paper is irrelevant to evaluating INHBB's disease associations.


[Read Paper](https://www.semanticscholar.org/paper/ce981bc89c565bb4d97baff441baaf8695595448)


### #1049 Genetic characteristics of autosomal dominant polycystic kidney disease in a Greek population

**Why Not Relevant**: This paper does not provide direct evidence for or against the claim that INHBB is likely associated with at least one Mendelian disease. The study focuses on genetic causes of autosomal dominant polycystic kidney disease (ADPKD) and reports pathogenic or likely pathogenic variants primarily in PKD1 and PKD2, with a few variants in SEC61B and SLC3A1. INHBB is not mentioned anywhere in the provided content. While the methods note sequencing of over 600 clinically relevant genes, the paper does not specify whether INHBB was included on the sequencing panel or whether any INHBB variants were absent or present; therefore absence of mention is not strong negative evidence. Important limitations that reduce relevance: (1) the cohort is ascertained for ADPKD, so only genes plausibly causing that phenotype would be expected to be identified; INHBB could be associated with a different phenotype and would not be captured here; (2) even with tNGS/WES/WGS, some variant types or noncoding pathogenic variants can be missed; (3) the sample size and ethnic homogeneity limit the ability to detect rare Mendelian associations for genes unrelated to the studied phenotype. Given these factors, the paper provides at best weak, indirect circumstantial evidence (absence of reported INHBB findings in an ADPKD cohort) and no mechanistic data linking INHBB to Mendelian disease, so its relevance to the claim is minimal.


[Read Paper](https://www.semanticscholar.org/paper/809bc9272642701e2600d56210354f9dc489b442)


### Identification of novel disease-causing variants in rare diseases using trio exome sequencing

**Why Not Relevant**: The provided paper content does not mention INHBB at all and therefore provides no direct evidence for or against the claim that INHBB is likely associated with at least one Mendelian disease. The paper describes trio exome sequencing applied to several cohorts and reports variants in specific genes; example verbatim sentences from the text show the genes investigated and the findings (no occurrence of INHBB):

"The use of trio exome next-generation sequencing (NGS), an analysis of the entire complement of coding exons within an affected index patient and unaffected parents, is increasingly being utilised in the analysis of patients with congenital diseases." 

"This revealed a single case of allelic drop-out of a known GLRA1 p.Y307C variant in a novel de novo pattern of inheritance, three cases where known or expected pathogenic variants had been identified in genes known to cause a seizure phenotype (DOLK and DMD), and an association with variants in ATRN, a null mutant of which is associated with a seizure phenotype in Mus musculus." 

"Second, whole exome analysis of three individual cases with rare diseases was performed; a case of hypomyelinating leukodystrophy with a homozygous NKX6-2 p.L54Qfs variant, a case of microcephaly and pontocerebellar hypoplasia with a MED14 de novo p.R1162C variant, and a case of syndromic intellectual disability with hypotonia and a homozygous DRG1 p.Y295* variant." 

"Third, novel de novo GRIN1 p.M818R and GRIN2B p.M824V variants were identified in further cases with developmental delay and hypotonia, adding to recent publications expanding the phenotype of variants within the NMDA receptor subunits." 

Why this means the paper is not relevant to the INHBB claim:
- Absence of mention: INHBB is not named in any of the reported findings or gene lists in the provided content, so there is no direct report of INHBB variants causing disease in this work.
- No mechanistic connection provided: The mechanistic descriptions and candidate genes relate to glycinergic synapses, seizure phenotypes, hypomyelination, NMDA receptor subunits, etc.; none implicate INHBB or pathways directly connected to INHBB function.
- Study scope and limitations: Trio/whole-exome sequencing could detect causal variants in INHBB if such variants were present in the studied individuals and relevant to their phenotypes, but the study samples are small (cohort of eight trios plus three individual exomes and additional cases) and focused on specific congenital neurological phenotypes. Therefore absence of a reported INHBB finding here does not constitute evidence that INHBB is not associated with any Mendelian disease. The paper simply does not address INHBB.

Conclusion: Relevance is essentially zero because the paper provides neither direct nor mechanistic evidence regarding INHBB. To evaluate the claim about INHBB, one would need studies that specifically investigate INHBB variation in affected individuals, report segregating rare variants, functional assays, or mechanistic links to a Mendelian phenotype — none of which are present in the provided content.


[Read Paper](https://www.semanticscholar.org/paper/8ae081628e0ab2aec44f8fe9990f230956fb40ba)


### Abstract 9929: Novel Mutations Identified by Whole Exome Sequencing in Families With Congenital Heart Disease

**Why Not Relevant**: The provided paper content does not mention the gene INHBB at all. The study focuses on using whole-exome sequencing to identify novel variants segregating with familial congenital heart disease (CHD) and reports aggregate counts and examples (e.g., Eph-family interacting proteins), but the text supplied does not list INHBB among the genes found nor present any data, segregation, functional, or prior-disease associations for INHBB. Therefore there is no direct evidence in this paper that INHBB is associated with any Mendelian disease. Further, the study context reduces the likelihood of supporting the claim even indirectly: CHD is described as a complex, multifactorial disorder, the sample of sequenced individuals is small (12 affected, 7 unaffected across 5 families), identified variants are characterized primarily by in silico prediction (SIFT, PolyPhen, MutationTaster) without reported functional validation, and the authors explicitly call for replication and candidate-gene resequencing. These limitations mean that, even if INHBB had appeared among the 69 genes (which is not shown in the provided text), this study alone would be insufficient to establish INHBB as causative for a Mendelian disease. Given the absence of any mention or data on INHBB in the supplied content, the paper provides no support for the claim.


[Read Paper](https://www.semanticscholar.org/paper/cd733afc1d20916deebb33a5523de1cc7d9d2d01)


## Search Queries Used

- INHBB mutation Mendelian disease monogenic disorder case report family segregation OMIM ClinVar

- INHBB rare variant association exome sequencing whole exome sequencing genome sequencing trio sequencing developmental disorder congenital anomaly familial segregation de novo

- INHBB inhibin beta B activin beta B function TGF beta SMAD knockout mouse phenotype zebrafish morpholino fertility gonadal development reproductive failure

- INHBB loss of function intolerance pLI gnomAD constraint allele frequency ClinVar pathogenic review monogenic gene candidate autosomal dominant autosomal recessive


## Bibliometric Configuration

- Use Bibliometrics: True

- Author Impact Weight: 0.4

- Citation Impact Weight: 0.4

- Venue Impact Weight: 0.2
