## Supplementary material for "MENDELSEEK: An algorithm that predicts Mendelian Genes and elucidates what makes them special": INTS9.txt

# Claim: The gene INTS9 is likely associated with at least one Mendelian disease.

**Status**: processed

**Overall Rating**: -1

**Explanation**:

No relevant papers were found that support or refute this claim.


## Relevant Papers


## Other Reviewed Papers


### Expanding the phenotype of a recurrent de novo variant in PACS1 causing intellectual disability

**Why Not Relevant**: The provided paper focuses entirely on PACS1 as the gene implicated in a recurrent de novo missense mutation causing an autosomal-dominant intellectual disability syndrome (MRD17). The text contains multiple direct and mechanistic statements about PACS1 (genetic finding, molecular function, expression pattern, and a mechanistic hypothesis involving the CK2-binding motif). Nowhere in the provided content is the gene INTS9 mentioned, nor are any data presented (genetic, functional, or epidemiological) that could be plausibly interpreted as evidence linking INTS9 to a Mendelian disease. Therefore this paper provides no direct evidence for the claim, no mechanistic pathway implicating INTS9, and no indirect evidence that would meaningfully change the prior probability that INTS9 is associated with a Mendelian condition. The absence of mention of INTS9 in a report about a different causal gene does not constitute negative evidence against INTS9; it simply is uninformative with respect to the claim.


[Read Paper](https://www.semanticscholar.org/paper/e460d89d850693b4fb1e3df12702be0dfb8d7f5b)


### Duplication 8q12: confirmation of a novel recognizable phenotype with duane retraction syndrome and developmental delay

**Why Not Relevant**: The provided paper content contains a single locus-level sentence: "It is observed that patients with 8q12 duplication share a common recognizable phenotype characterized by DRS, developmental delay and facial features, which strongly suggest that this entity may define a novel syndrome." This sentence does not mention the gene INTS9 or any other gene by name, does not report variant-level or family segregation data, and gives no mechanistic or functional data linking a specific gene to the phenotype. Therefore it provides no direct evidence that INTS9 is associated with a Mendelian disease. 

Detailed reasons: 
- No gene-level mention: The text refers only to a chromosomal region (8q12 duplication) and a shared phenotype (DRS, developmental delay, facial features). It does not identify which genes within 8q12 are duplicated, nor does it single out INTS9. Without explicit naming or mapping, one cannot infer that INTS9 is involved. 
- Locus-level vs gene-level evidence: A duplication at 8q12 may encompass many genes; locus-level phenotype correlation does not establish causality for any single gene within the interval. Assigning the phenotype to INTS9 would require additional evidence (e.g., focal variants in INTS9, minimal critical region mapping narrowing to INTS9, segregation of INTS9 variants with disease, or functional perturbation of INTS9 producing the phenotype). None of these are present. 
- No mechanistic data: The sentence provides no mechanistic hypothesis (e.g., haploinsufficiency, gain-of-function, dominant-negative) involving INTS9, nor functional assays, expression data, or model organism results. 
- No Mendelian evidence: There is no information about inheritance patterns, de novo occurrences, family segregation, or recurrence consistent with a Mendelian trait attributable to a specific gene. 
- Generalizability and boundaries: The statement suggests a novel syndrome associated with an 8q12 duplication, but without breakpoint coordinates or gene content, it is impossible to generalize or attribute the syndrome to INTS9 specifically. 

What would be needed to support the claim: demonstration that INTS9 is within the minimal duplicated region; variant-level evidence (rare pathogenic variants in INTS9) segregating with disease in families; de novo damaging variants in INTS9 in multiple unrelated affected individuals; functional data showing that perturbation of INTS9 recapitulates key features; or high-resolution mapping narrowing the critical interval to include only INTS9. None of these appear in the provided text, so the paper content as given is not relevant to the claim that INTS9 is likely associated with at least one Mendelian disease.


[Read Paper](https://www.semanticscholar.org/paper/d58c242dcf0888de78b139ad3ea03c456767685a)


### A recessive syndrome of intellectual disability, moderate overgrowth, and renal dysplasia predisposing to Wilms tumor is caused by a mutation in FIBP gene

**Why Not Relevant**: The provided paper does not mention INTS9 at all and instead identifies and studies FIBP as the gene implicated in a recessive overgrowth syndrome. All primary genetic, mechanistic, and contextual statements in the paper refer to FIBP (including mapping, variant description, predicted mechanistic effect, and prior reports). Therefore the paper provides no direct genetic or mechanistic evidence that INTS9 is associated with any Mendelian disease. Limitations: absence of mention of INTS9 in the excerpt means we cannot infer anything about INTS9 from this study; unless additional, unprovided sections of the paper discuss INTS9 (which is not indicated here), this paper is irrelevant to the claim that INTS9 is likely associated with a Mendelian disorder.


[Read Paper](https://www.semanticscholar.org/paper/53849cb0c70575cf51bd83ce3aec9cbf5bc51943)


### Clinical and genetic characteristics and prenatal diagnosis of patients presented GDD/ID with rare monogenic causes

**Why Not Relevant**: The provided paper content contains a single general sentence: "Patients presented with GDD/ID caused by rare single gene variants are characterized by early onset, relatively severe symptoms and great clinical variability and genetic heterogeneity." This statement does not mention the gene INTS9, report any variants in INTS9, present case(s), segregation data, functional assays, population/constraint metrics, or any mechanistic information linking INTS9 to a Mendelian phenotype. It is a high-level background remark about the clinical features and heterogeneity of GDD/ID caused by rare single-gene variants and therefore cannot serve as direct evidence that INTS9 is associated with a Mendelian disease. Mechanistically, the sentence provides no information about INTS9 function, the effect of variants, or biological pathways that would make an INTS9–disease link plausible. Methodological and result-oriented elements required to evaluate a gene–disease association (e.g., number of affected individuals, inheritance pattern, statistical enrichment, functional validation, or replication) are entirely absent. For these reasons the content is not relevant to the claim that "INTS9 is likely associated with at least one Mendelian disease." To assess that claim one would need excerpts reporting INTS9-specific variants in affected individuals with segregation or de novo occurrence, functional evidence of pathogenicity, population frequency/constraint data, or inclusion in curated disease resources (e.g., OMIM), none of which are present in the supplied text.


[Read Paper](https://www.semanticscholar.org/paper/87ded00e6def971d09456b16f6ca88fa38fc15cb)


### Novel LSS variants in alopecia and intellectual disability syndrome: New case report and clinical spectrum of LSS‐related rare disease traits

**Why Not Relevant**: This paper reports pathogenic biallelic variants in the gene LSS and describes clinical phenotypes (including congenital cataract, hypotrichosis, alopecia–intellectual disability syndrome, and expanded neurologic features) found in a proband and in a review of reported LSS cases. The provided text makes no mention of the gene INTS9, the Integrator complex, or any genetic, biochemical, or clinical link between INTS9 and human Mendelian disease. Therefore it provides neither direct evidence (no statements that INTS9 is mutated in any patients or associated with disease) nor mechanistic evidence (no discussion of interactions, pathways, protein complexes, or functional data connecting INTS9 to LSS or to phenotypes). Limitations and caveats: it is possible that INTS9 could be connected to LSS or to similar phenotypes in other literature, but this paper provides no information to support or refute that possibility. To evaluate the claim about INTS9 you should consult other sources (e.g., primary literature on INTS9, OMIM, ClinVar, HGMD, gnomAD constraint metrics, or functional studies of INTS9) because this LSS-focused paper is not relevant.


[Read Paper](https://www.semanticscholar.org/paper/fe90a5bb55278bc80588741eed5def6cf4f1aa82)


### De Novo p.Asp3368Gly Variant of Dystrophin Gene Associated with X-Linked Dilated Cardiomyopathy and Skeletal Myopathy: Clinical Features and In Silico Analysis

**Why Not Relevant**: The provided paper content focuses exclusively on a pathogenic variant in the dystrophin (DMD) gene identified by whole exome sequencing and confirmed by Sanger sequencing, and on the clinical consequences (dilated cardiomyopathy and skeletal muscle disease) in the proband and her son. There is no mention of the gene INTS9 anywhere in the text provided. Therefore the paper contains neither direct evidence (e.g., identification of pathogenic variants in INTS9 segregating with disease) nor mechanistic evidence (e.g., functional studies of INTS9, pathway links to disease phenotypes) that would support or refute the claim that INTS9 is likely associated with at least one Mendelian disease. Although WES was used in the study (so INTS9 could in principle have been assayed), the absence of any reported INTS9 findings in the provided text means no positive or negative inference about INTS9 can be made from this paper. In summary: relevance to the INTS9–Mendelian disease claim is effectively zero because the study reports findings only about DMD and provides no data, results, or discussion concerning INTS9.


[Read Paper](https://www.semanticscholar.org/paper/d71f314236ef259f325f8caa52d9474e9a03cb85)


### Novel Compound Heterozygous Splice-Site Variants in TPM3 Revealed by RNA Sequencing in a Patient with an Unusual Form of Nemaline Myopathy: A Case Report

**Why Not Relevant**: The paper exclusively investigates TPM3 as the disease-causing gene in a patient with nemaline myopathy. All reported genetic findings, mechanistic RNA and protein evidence, methods, and conclusions pertain to TPM3. There is no mention of INTS9, no variants reported in INTS9, and no mechanistic discussion involving INTS9. Although the study used broad genetic methods (WES/WGS) capable of detecting variants in many genes, the absence of any data or discussion about INTS9 means this paper provides no direct evidence that INTS9 is associated with any Mendelian disease, nor does it provide mechanistic evidence supporting such an association. Therefore the paper is not relevant to the claim that INTS9 is likely associated with at least one Mendelian disease.


[Read Paper](https://www.semanticscholar.org/paper/1be01b71ad7221e3e9cc9367fa3b86b00423f3f2)


### Novel PHOX2B germline mutation in childhood medulloblastoma: a case report

**Why Not Relevant**: The provided paper content exclusively discusses a de novo PHOX2B germline mutation as a potential cause of medulloblastoma and suggests familial germline screening. There is no mention of the gene INTS9, no genetic association data concerning INTS9, no mechanistic pathway involving INTS9, and no discussion of Mendelian disease associations for INTS9. Because the supplied text contains no reference to INTS9 or evidence (direct or mechanistic) bearing on its involvement in Mendelian disease, it cannot be used to support or refute the claim that INTS9 is likely associated with at least one Mendelian disease. Any inference about INTS9 from this paper would be speculative and unsupported. To evaluate the claim about INTS9, sources that explicitly report variants, segregation, functional impact, or mechanistic links involving INTS9 and Mendelian phenotypes are required.


[Read Paper](https://www.semanticscholar.org/paper/6089dea8e271ef8eec8c0b36a3965719a4fe554e)


### Novel PHOX2B germline mutation in childhood medulloblastoma: a case report

**Why Not Relevant**: The provided paper content discusses a de novo PHOX2B germline mutation in a patient with medulloblastoma and recommends familial germline variant screening. It does not mention INTS9, does not present data (genetic, clinical, or mechanistic) about INTS9, and therefore supplies no direct or mechanistic evidence for or against an association between INTS9 and any Mendelian disease. The lack of relevance arises from a gene mismatch: the reported genetic finding concerns PHOX2B, not INTS9. Caveat: this assessment is limited to the excerpt supplied; if other sections of the full paper (not provided here) discuss INTS9, those sections would need to be reviewed. Based on the supplied content alone, relevance to the claim is effectively zero.


[Read Paper](https://www.semanticscholar.org/paper/a8c43338873b285e085862dbe1dcc72cd8b0df14)


### Case report: Neonatal-onset inflammatory bowel disease due to novel compound heterozygous mutations in DUOX2

**Why Not Relevant**: The paper exclusively studies DUOX2 and its role in Very Early Onset Inflammatory Bowel Disease, reporting compound heterozygous DUOX2 variants as a monogenic cause in the presented patient. There is no mention of the gene INTS9, no genetic, clinical, or mechanistic data concerning INTS9, and no discussion that would link INTS9 to the phenotype or to Mendelian disease in general. Therefore this paper provides no direct or mechanistic evidence supporting the claim that INTS9 is likely associated with at least one Mendelian disease.


[Read Paper](https://www.semanticscholar.org/paper/78a31d77d8707f934b79756b02d086d433a871c3)


### Delineating the overlapping roles of the single-stranded DNA binding proteins Ssb1 and Ssb2 in the maintenance of genomic stability and intestinal homeostasis

**Why Not Relevant**: The paper provides mechanistic and functional evidence that Integrator activity (and the Integrator catalytic components IntS9/IntS11) is critical for snRNA processing, splicing and cellular viability in mouse/cell models. However, it contains no direct genetic or clinical evidence that variants in INTS9 cause a Mendelian disorder: there are no human patient data, no family segregation or phenotype descriptions attributable to INTS9 mutations, and no experiments directly perturbing INTS9 (the experiments perturb Ssb1/2). The experimental systems are mouse models and cultured MEFs (including SV40-immortalised cells) with inducible Ssb1/2 double knockouts; conditions and perturbations differ from germline heterozygous/homozygous human variants leading to Mendelian disease. Therefore while the paper offers mechanistic plausibility that INTS9 is biologically important (supporting the idea it could be disease-relevant), it does not provide direct evidence that INTS9 is associated with any Mendelian disease.


[Read Paper](https://www.semanticscholar.org/paper/d1af8501b91df5b68859e730f4da4bb3875d14aa)


## Search Queries Used

- INTS9 variant Mendelian disease case report whole exome sequencing de novo biallelic germline pathogenic

- INTS9 gene association monogenic disease rare disease clinical phenotype intellectual disability developmental delay congenital anomaly genotype phenotype

- Integrator complex subunit 9 INTS9 function RNA polymerase II snRNA processing transcriptional regulation loss of function knockout mouse zebrafish functional assay

- INTS9 ClinVar pathogenic gnomAD loss of function intolerance pLI LOEUF constraint haploinsufficiency gene essentiality population variation


## Bibliometric Configuration

- Use Bibliometrics: True

- Author Impact Weight: 0.4

- Citation Impact Weight: 0.4

- Venue Impact Weight: 0.2
