## Supplementary material for "MENDELSEEK: An algorithm that predicts Mendelian Genes and elucidates what makes them special": ITIH4.txt

# Claim: The gene ITIH4 is likely associated with at least one Mendelian disease.

**Status**: processed

**Overall Rating**: -1

**Explanation**:

No relevant papers were found that support or refute this claim.


## Relevant Papers


## Other Reviewed Papers


### Frequent expression loss of Inter-alpha-trypsin inhibitor heavy chain (ITIH) genes in multiple human solid tumors: A systematic expression analysis

**Why Not Relevant**: The claim concerns whether the gene ITIH4 is likely associated with at least one Mendelian disease (i.e., a disease caused by germline variation in a single gene with Mendelian inheritance). The provided paper content reports somatic downregulation of ITIH family genes in solid tumors (breast, colon, lung). Somatic expression changes in cancers do not constitute evidence of a germline, Mendelian disease association. The excerpt does not name ITIH4 specifically, does not discuss germline variants, inheritance, segregation, penetrance, or any clinical Mendelian phenotype, and contains no statistical or methodological detail. Thus the content is not relevant to establishing or refuting a Mendelian association for ITIH4.


[Read Paper](https://www.semanticscholar.org/paper/5d14d48e9730a3bacebe89113369cd48e7c59c8a)


### Exome sequencing of Finnish isolates enhances rare-variant association power

**Why Not Relevant**: The provided paper content contains only a single summary sentence describing that exome sequencing in Finnish populations found 26 deleterious alleles associated with 64 quantitative traits and that these show geographical clustering similar to Finnish Mendelian disease mutations. It does not mention ITIH4 or any specific genes, nor does it present data linking any identified allele to a clinically defined Mendelian disease. The claim under consideration is gene-specific (that ITIH4 is likely associated with at least one Mendelian disease). To evaluate that claim one would need explicit evidence such as: (a) ITIH4 being listed among the deleterious alleles discovered; (b) an association of ITIH4 variants with a Mendelian phenotype (i.e., fully penetrant monogenic disease, segregation in families, or clinical diagnosis meeting Mendelian criteria); or (c) mechanistic/functional data connecting ITIH4 disruption to disease pathogenesis. None of these elements are present in the supplied content. The observed associations in the sentence are with quantitative traits (which may reflect polygenic or complex disease influences) rather than bona fide Mendelian disorders, and population-level clustering alone cannot attribute Mendelian causality to a specific gene. Therefore the paper content as provided is insufficient to support or refute the claim about ITIH4, yielding a relevance score near zero.


[Read Paper](https://www.semanticscholar.org/paper/dba3e4f4a553f447c8aa37a6d4f5ff8b17f0021e)


### Renal inter-alpha-trypsin inhibitor heavy chain 3 increases in calcium oxalate stone-forming patients.

**Why Not Relevant**: The provided content contains a single observational sentence about protein expression/localization in stone formers and a reference to "inter-alpha-trypsin inhibitor components" being linked to human stone disease. This does not meet the types of evidence required to support the claim that ITIH4 is likely associated with at least one Mendelian disease because: (1) ITIH4 is not mentioned by name—the sentence refers to "heavy chain 3," which would more plausibly refer to ITIH3, not ITIH4; (2) the content reports protein expression/localization, not genetic variation, inheritance patterns, segregation, or pathogenicity data that would indicate a Mendelian disorder; (3) stone disease (kidney stones) is typically multifactorial and not classically a Mendelian disease, and no familial or genetic analysis is presented; (4) there are no methods, sample sizes, statistics, or experimental conditions provided to assess the robustness or generalizability of the observation. For these reasons, the excerpt is insufficient to support the claim and relevance is very low.


[Read Paper](https://www.semanticscholar.org/paper/b0907025183725f48ef03fb735430d4590d6eda6)


### Statistical Guidance for Experimental Design and Data Analysis of Mutation Detection in Rare Monogenic Mendelian Diseases by Exome Sequencing

**Why Not Relevant**: This paper does not provide any direct or mechanistic evidence about the gene ITIH4. The manuscript excerpt is methodological and statistical in scope: it presents a statistical modeling framework for power calculations in exome sequencing studies and discusses general properties of gene discovery under various inheritance models and locus heterogeneity. It contains no experimental data, gene lists, variant observations, genotype–phenotype correlations, functional assays, or any mention of ITIH4. Therefore it cannot support or refute the claim that "ITIH4 is likely associated with at least one Mendelian disease." 

Representative verbatim sentences from the excerpt that demonstrate the paper's scope (included to justify the non-relevance):
- "To address these questions, we present a statistical modeling framework to calculate the power, the probability of identifying truly disease-causing genes, under various inheritance models and experimental conditions, providing guidance for both proper experimental design and data analysis." 
- "Based on our model, we found that the exome sequencing approach is well-powered for mutation detection in recessive, but not dominant, Mendelian diseases with high locus heterogeneity." 

Why these statements imply non-relevance to the ITIH4 claim:
- The quoted sentences show the paper's focus is on statistical power and study design, not on reporting specific gene associations. No gene-level results are presented in the provided content. 
- Because the paper lacks gene-specific data, it provides no direct evidence (e.g., observed pathogenic variants in ITIH4 segregating with disease) and no mechanistic evidence (e.g., functional impact of ITIH4 disruption, pathways implicating ITIH4 in Mendelian phenotypes) that would be necessary to evaluate whether ITIH4 is associated with a Mendelian disease. 

Possible indirect / contextual relevance (insufficient to change relevance score):
- The paper's conclusions about the power of exome sequencing imply that, under certain study designs and effect sizes (for example, a gene responsible for ~5% of cases in a recessive disorder with 200 unrelated patients), exome sequencing could discover disease genes. That is a general statement about discoverability, not evidence that ITIH4 is associated with disease. 

Limitations that prevent using this paper to evaluate the ITIH4 claim:
- No mention of ITIH4 or any specific genes in the provided content.
- No case series, cohort data, segregation analyses, variant pathogenicity assessments, or functional experiments related to ITIH4.
- The paper's statistical results cannot be mapped to ITIH4 without independent data showing effect size, inheritance model, allele frequency, or observed variants in affected individuals. 

Conclusion: Relevance = 0.0 because the paper contains no relevant evidence (direct or mechanistic) pertaining to ITIH4 and therefore cannot be used to support or refute the claim that ITIH4 is likely associated with at least one Mendelian disease.


[Read Paper](https://www.semanticscholar.org/paper/4855f23094ced564082f0bdbccf7dc2d23429706)


### Thrombin Cleavage of Inter-α-inhibitor Heavy Chain 1 Regulates Leukocyte Binding to an Inflammatory Hyaluronan Matrix*♦

**Why Not Relevant**: This paper does not provide evidence that ITIH4 is associated with any Mendelian disease. The experimental focus is on extracellular matrix biology and the proteolytic cleavage of heavy chain 1 (HC1) by thrombin. HC1 corresponds to the gene ITIH1 (not ITIH4). The manuscript contains mechanistic and functional data about HC-HA complexes, thrombin cleavage of HC1, and leukocyte adhesion, but it contains no genetic data, no discussion of ITIH4, no variant or inheritance analyses, and no clinical phenotypes consistent with Mendelian disorders. Therefore the content is not relevant to the claim that ITIH4 is likely associated with at least one Mendelian disease.


[Read Paper](https://www.semanticscholar.org/paper/2a7391171b00293afe95486ec8685d44d6648187)


### Genetic Sequencing of Pediatric Patients Identifies Mutations in Monogenic Inflammatory Bowel Disease Genes that Translate to Distinct Clinical Phenotypes

**Why Not Relevant**: The paper's abstract describes a whole-exome sequencing study of an unselected pediatric IBD cohort focused on identifying pathogenic or likely pathogenic variants in 68 monogenic IBD genes. The abstract lists specific genes that were implicated (TRIM22, WAS, NOD2) and others that did not meet monogenic criteria (ADA, FERMT1, LRBA). Nowhere in the provided text is the gene ITIH4 mentioned, discussed, or listed among the genes evaluated. Because the study is disease-focused (monogenic causes of IBD) and the abstract neither names ITIH4 nor provides the full list of 68 genes, there is no direct evidence in this paper excerpt that ITIH4 is associated with any Mendelian disease. Absence of mention in this abstract cannot be interpreted as evidence against ITIH4's involvement in Mendelian disease for several reasons: (1) the paper may not have included ITIH4 among the 68 genes, (2) even if ITIH4 were included, a negative result or non-mention would not be reported in the abstract unless it met study thresholds, and (3) this cohort is limited to pediatric IBD patients, so associations with Mendelian diseases outside of IBD would not be detected. Additionally, methodological details (e.g., full gene list, variant-filtering criteria, statistical power for individual genes, phenotypic breadth) are not provided in the excerpt, preventing any reliable inference about ITIH4. For these reasons the paper content provided is essentially non-informative with respect to the claim that ITIH4 is likely associated with at least one Mendelian disease.


[Read Paper](https://www.semanticscholar.org/paper/06f782a67bd23b9faef27200da3457378695fd73)


### A novel STAT1 loss-of-function mutation associated with Mendelian susceptibility to mycobacterial disease

**Why Not Relevant**: Relevance is scored 0.0 because the paper's content exclusively addresses a STAT1 loss-of-function variant causing Mendelian susceptibility to mycobacterial disease (MSMD). The document does not mention ITIH4, present any variants in ITIH4, evaluate ITIH4 expression or function, or propose mechanistic links between ITIH4 and MSMD or any other Mendelian disorder. The provided excerpts demonstrate the paper's focus on STAT1 (identification of a STAT1 variant by whole-exome sequencing, functional assays showing impaired STAT1 activation, and interpretation that STAT1 LOF causes MSMD). Therefore this paper provides no direct evidence (no variant calls, no genetic association, no case reports) and no mechanistic evidence (no pathway, expression, interaction, or functional data) that would support the claim that ITIH4 is likely associated with a Mendelian disease. Caveats: absence of any mention of ITIH4 in this paper is not proof that ITIH4 has no Mendelian associations elsewhere; it only means this particular paper does not contribute evidence for the claim. To evaluate the claim further, one should search human genetics databases (e.g., OMIM, ClinVar, HGMD), population variant databases (gnomAD), and the wider literature for ITIH4 variants, segregation studies, functional assays, or case reports linking ITIH4 to Mendelian phenotypes.


[Read Paper](https://www.semanticscholar.org/paper/9cec67e638d5279f19cbaf5c3800b2b0f9d44ecf)


### Whole-exome sequencing in familial type 2 diabetes identifies an atypical missense variant in the RyR2 gene

**Why Not Relevant**: The paper's content exclusively reports findings about a missense variant in the RyR2 gene and its relationship to type 2 diabetes and CPVT-related pathogenic hotspots. There is no mention of ITIH4 anywhere in the provided text, no data (genetic, functional, or epidemiologic) about ITIH4, and no discussion of Mendelian diseases linked to ITIH4. Therefore this paper provides no direct or mechanistic evidence for or against the claim that ITIH4 is likely associated with at least one Mendelian disease. Relevance is effectively zero because the gene of interest (ITIH4) is absent from the reported analyses and conclusions.


[Read Paper](https://www.semanticscholar.org/paper/21d68d0282d328f9f7038ea8caa048ae43248fbb)


### Identifying genomic variant associated with long QT syndrome type 2 in an ecuadorian mestizo individual: a case report

**Why Not Relevant**: The paper is a case report of Long QT Syndrome type 2 and reports sequencing results implicating KCNH2 (pathogenic) and variants of uncertain significance in AKAP9 and TTN. The manuscript contains no mention of the gene ITIH4, no data (genetic, functional, epidemiologic) about ITIH4, and no discussion that would support or refute an association between ITIH4 and any Mendelian disease. Therefore it provides no direct evidence linking ITIH4 to a Mendelian disorder, nor mechanistic evidence about ITIH4. The absence of ITIH4 in this focused LQTS genetic analysis does not constitute comprehensive negative evidence about ITIH4 generally; it simply means this study is not informative regarding the claim. Given this, relevance to the claim is effectively zero.


[Read Paper](https://www.semanticscholar.org/paper/aa65df486c4f1720d86feb42ddc72ed25ab57431)


### Frequent expression loss of Inter-alpha-trypsin inhibitor heavy chain ( ITIH ) genes in multiple human solid tumors : A systematic expression analysis

**Why Not Relevant**: The paper focuses on expression analysis of ITIH family genes in human solid tumors (cancer biology), reporting family-level biochemical roles and tumor-associated downregulation. It does not present any data on germline variants, segregation in families, case reports of monogenic disease, mutation screens, linkage analyses, or functional studies of variants that would be required to support a claim that ITIH4 is associated with a Mendelian disease. ITIH4 is only mentioned as a member of the ITIH gene family; it is not specifically analyzed (the detailed follow-up focused on ITIH2). Therefore the content provides mechanistic/functional context relevant to general biology of ITIH genes but offers no direct or sufficient mechanistic genetic evidence to support that ITIH4 is likely associated with at least one Mendelian disease.


[Read Paper](https://www.semanticscholar.org/paper/a2306d0452209babadf15251ef2a726658d44f4f)


### CNVoyant a machine learning framework for accurate and explainable copy number variant classification

**Why Not Relevant**: The supplied paper content is a single sentence describing the CNVoyant tool's performance and features: "Compared to alternative approaches, CNVoyant shows marked improvements in precision-recall and ROC AUC metrics for binary pathogenic classifications while going one step further, offering multi-classification of clinical significance and corresponding SHAP explainability plots." This sentence does not mention the gene ITIH4, any Mendelian disease, variant-level or gene-level associations, clinical cases, cohort analyses, functional experiments, or any data linking ITIH4 to disease. Therefore there is no direct evidence (no reported association, statistics, or cases) and no mechanistic evidence (no biological pathway, functional assay, or mechanistic rationale) in the provided content that ITIH4 is associated with a Mendelian disease. Limitations/caveats: only the single supplied sentence was available for assessment; if the full paper contains sections not provided here that discuss genes, variant findings, or disease associations (including ITIH4), those were not supplied and could change this conclusion. Based solely on the provided content, relevance to the claim is effectively zero.


[Read Paper](https://www.semanticscholar.org/paper/56f08cb7ff8b47dcf67c8982001cb74c98b5d841)


### CNVoyant: A Highly Performant and Explainable Multi-Classifier Machine Learning Approach for Determining the Clinical Significance of Copy Number Variants

**Why Not Relevant**: The provided paper content does not mention the gene ITIH4, does not report any gene-level associations or gene-centric results, and presents only a description of a machine-learning framework for classifying CNVs. While the method uses features such as "disease-gene annotations" and "dosage sensitivity," which are mechanistically relevant to identifying genes involved in Mendelian disease, the excerpt contains no direct data, examples, or findings about ITIH4 specifically. There are no quoted results (e.g., variant counts, pathogenic classifications, statistical associations) tied to ITIH4, nor any case reports, tables, or supplementary notes in the provided text that would allow assessment of ITIH4's association with a Mendelian disease. Because the content is methodological and high-level, it cannot support or refute the claim that "ITIH4 is likely associated with at least one Mendelian disease." To evaluate that claim, one would need gene-specific evidence such as ClinVar or DECIPHER entries linking ITIH4 variants to Mendelian phenotypes, reported pathogenic variants in ITIH4, dosage-sensitivity annotations specific to ITIH4, or curated gene-disease associations (e.g., OMIM entries). None of those are present in the supplied excerpt, so the relevance to the claim is negligible.


[Read Paper](https://www.semanticscholar.org/paper/6db12c5a0a30823e95cd0b4de7760931a0dc67a6)


### Comprehensive in Vitro Characterization of CXCR4WHIM variants to Decipher Genotype-Phenotype Correlations in WHIM Syndrome

**Why Not Relevant**: The paper exclusively studies WHIM syndrome and functional effects of CXCR4 gain-of-function mutations. The text supplied contains no mention of the gene ITIH4, no experimental data involving ITIH4, and no mechanistic discussion linking ITIH4 to CXCR4, WHIM syndrome, or any other Mendelian disease. Therefore this paper does not provide direct evidence that ITIH4 is associated with any Mendelian disease, nor does it provide mechanistic evidence that would make such an association plausible. Absence of evidence in this single paper does not refute the claim in general, but it means this paper is not informative for assessing whether ITIH4 is likely associated with a Mendelian disease. No relevant pages were provided.


[Read Paper](https://www.semanticscholar.org/paper/140b73b1f5755688518e300347c90d981dfe06c3)


## Search Queries Used

- ITIH4 mutation rare variant exome sequencing Mendelian disease monogenic

- ITIH4 ClinVar OMIM Decipher pathogenic variant clinical significance

- Inter alpha trypsin inhibitor heavy chain H4 hyaluronan extracellular matrix protease inhibitor inflammation connective tissue disease

- ITIH4 loss of function intolerance gnomAD pLI constraint exome sequencing trio developmental disorder novel candidate gene


## Bibliometric Configuration

- Use Bibliometrics: True

- Author Impact Weight: 0.4

- Citation Impact Weight: 0.4

- Venue Impact Weight: 0.2
