## Supplementary material for "MENDELSEEK: An algorithm that predicts Mendelian Genes and elucidates what makes them special": K1109_HUMAN.txt

# Claim: The gene K1109_HUMAN is likely associated with at least one Mendelian disease.

**Status**: processed

**Overall Rating**: 0

**Explanation**:

Supporting evidence:
None of the provided paper excerpts mention K1109_HUMAN (or KIAA1109) or present genetic, clinical, or functional data linking that specific gene/protein to a Mendelian disease. The first paper documents pathogenic biallelic variants in LSS causing several Mendelian conditions (congenital cataract type 44, autosomal recessive hypotrichosis type 14, and alopecia–intellectual disability syndrome type 4). The second paper describes disease phenotypes resulting from NARFL deficiency and links NARFL variants/polymorphisms to susceptibility for several conditions. While both papers provide clear examples of genes (LSS, NARFL) implicated in Mendelian or Mendelian-like disorders, neither provides any evidence regarding K1109_HUMAN.

Caveats or contradictory evidence:
There is no explicit contradictory evidence about K1109_HUMAN in the excerpts—only absence of mention. A potential caveat is naming ambiguity: if K1109_HUMAN was intended as an alternate or truncated label for another gene discussed (for example, LSS_HUMAN or NARFL_HUMAN), then the excerpts would be relevant. However, the gene symbols and protein names in the excerpts (LSS, NARFL) do not map to K1109_HUMAN; K1109_HUMAN likely refers to KIAA1109 (a distinct gene/protein). Because no data in the excerpts link KIAA1109/K1109_HUMAN to Mendelian disease, we cannot infer association. Another caveat is that absence of evidence in these two papers does not prove absence of any Mendelian association for K1109_HUMAN elsewhere in the literature; it only means the provided evidence does not support the claim.

Analysis of potential underlying mechanisms, deeper relationships, or second-order implications:
Establishing that a gene is “associated with at least one Mendelian disease” typically requires demonstration of pathogenic variants with appropriate inheritance and phenotype correlation (e.g., segregation in families, recurrence in unrelated affected individuals, functional evidence). The provided excerpts include examples of this standard for LSS and functional knockout models for NARFL, illustrating the kinds of data one would expect to support a Mendelian association. No analogous data (human pathogenic variants, segregation, case reports, or functional rescue/knockout data specific to K1109_HUMAN) are present. Mechanistically, neither LSS (sterol biosynthesis enzyme) nor NARFL (CIA pathway-related protein) is related to the putative KIAA1109 protein, so mechanistic inference across these genes is not justified from the excerpts.

Explanation of the rating decision:
Given that neither excerpt mentions K1109_HUMAN or supplies any genetic, clinical, or experimental evidence linking that gene to a Mendelian disorder, the appropriate conclusion is that there is no evidence in the provided material to support the claim. The two papers do support that some genes (LSS, NARFL) are implicated in Mendelian or Mendelian-like conditions, but that does not bear on K1109_HUMAN specifically. Therefore the claim that K1109_HUMAN is likely associated with at least one Mendelian disease is unsupported by the provided evidence; the correct rating is "No Evidence."


**Final Reasoning**:

I reviewed the supplied excerpts carefully. The first paper attributes Mendelian disease causation to LSS variants; the second attributes disease phenotypes to NARFL deficiency. Neither paper mentions K1109_HUMAN (likely KIAA1109) or provides any human genetic or functional data relevant to that gene. Because the assessment task is to evaluate the claim based on the provided evidence, and that evidence contains no support for K1109_HUMAN's association with a Mendelian disease, the rational conclusion is that there is no evidence in these materials to support the claim. I considered possible naming confusion as a caveat, but the gene names in the excerpts do not map to K1109_HUMAN, so conflation is not justified. Therefore I confirm the rating "No Evidence."


## Relevant Papers


### Novel LSS variants in alopecia and intellectual disability syndrome: New case report and clinical spectrum of LSS‐related rare disease traits

**Authors**: H. Elbendary (H-index: 8), M. Zaki (H-index: 46)

**Relevance**: 0.9

**Bibliometric Impact**: 0.35840000000000005


**Excerpts**:

- Pathogenic biallelic variants in LSS are associated with three Mendelian rare disease traits including congenital cataract type 44, autosomal recessive hypotrichosis type 14, and alopecia‐intellectual disability syndrome type 4 (APMR4).

- We performed trio research exome sequencing on a family with a four‐year‐old male with global developmental delay, epilepsy and striking alopecia, and identified novel compound heterozygous LSS splice site (c.14+2T>C) and missense (c.1357 G>A; p.V453L) variant alleles.

- Rare features associated with APMR4 such as cryptorchidism, micropenis, mild cortical brain atrophy and thin corpus callosum were detected.

- Previously unreported APMR4 findings including cerebellar involvement in the form of unsteady ataxic gait, small vermis with prominent folia, were noted. A review of all reported variants to date in 29 families with LSS‐related phenotypes showed an emerging genotype–phenotype correlation. Our report potentially expands LSS‐related phenotypic spectrum and highlights the importance of performing brain imaging in LSS‐related conditions.


**Explanations**:

- Direct relevance: This sentence explicitly states that pathogenic biallelic variants in the gene LSS are associated with three named Mendelian rare disease traits, i.e. it asserts LSS as causally linked to Mendelian disorders. Strengths: clear, concise summary linking LSS to Mendelian diseases. Limitations/caveats: it is a summary statement and does not present the underlying data in this sentence; applicability to the claim depends on whether the queried identifier (K1109_HUMAN) refers to the LSS protein/gene — the paper uses the gene symbol LSS, so mapping between K1109_HUMAN and LSS must be confirmed externally.

- Direct evidence: This sentence reports a specific family-based (trio) exome sequencing result identifying compound heterozygous LSS variants (one splice-site and one missense) in a patient with relevant clinical features. Strengths: case-level genetic evidence (biallelic variants in LSS) in a proband with compatible phenotype strengthens a causal association. Limitations/caveats: single-family report without functional validation described here; causality inferred from genotype–phenotype concordance but no biochemical/functional assays are presented in this excerpt; potential for coincidental findings or contributions from other variants cannot be excluded without broader analysis.

- Contextual/direct evidence: These observed clinical features are described as being associated with APMR4, the LSS-related syndrome. Strengths: clinical phenotype matches previously described syndrome manifestations, supporting that the identified LSS variants likely underlie a Mendelian phenotype in this patient. Limitations/caveats: phenotypic observations alone do not prove causation; sample-size limitation (one family here) and no statistical assessment provided in this sentence.

- Mechanistic/contextual/direct evidence: The first sentence documents additional, previously unreported phenotypic features (cerebellar involvement) expanding the clinical spectrum; the second sentence reports a review across 29 families showing an "emerging genotype–phenotype correlation," which supports a reproducible relationship between LSS variants and disease phenotypes; the third emphasizes that this report expands the phenotypic spectrum and clinical recommendations. Strengths: aggregation across 29 families and noted genotype–phenotype correlation increase the weight of evidence for LSS as a Mendelian disease gene and suggest reproducibility and some genotype–phenotype structure. Limitations/caveats: "emerging" implies the correlation is not yet definitive; no mechanistic biochemical or cellular data are provided here to demonstrate how specific variants affect LSS function and cause disease; review/sample ascertainment biases and heterogeneity in case reports may affect conclusions; statistical strength (e.g., segregation metrics, LOD scores, burden tests) is not provided in these sentences.


[Read Paper](https://www.semanticscholar.org/paper/fe90a5bb55278bc80588741eed5def6cf4f1aa82)


### The Mechanism of Vascular Endothelial Dysfunction Induced by Ferroptosis Mediated by NARFL Knockout

**Authors**: H. Hu (H-index: 0), S. Shi (H-index: 0)

**Relevance**: 0.85

**Bibliometric Impact**: 0.1


**Excerpts**:

- In our previous study, we identified a novel homozygous mutation in NARFL that led to decreased expression in a consanguineous family with diffuse pulmonary arteriovenous malformations (DPAVMs) secondary to pulmonary hypertension.

- Narfl deletion in zebrafish resulted in larvae lethality, vascular malformation with abnormal blood flow, and abnormal blood-brain barrier (BBB) structur.

- Fluorescence probe detection showed increased iron, enhanced oxidative stress, and lipid peroxidation in response to narfl deficiency, which could be partially alleviated by the use of Ferrostatin-1.

- In HPMEC cells, NARFL deficiency resulted in decreased proliferation, abnormal mitochondrial morphology, oxidative stress, decreased mitochondrial respiration, decreased tube formation ability and enhanced permeability.

- In mice, Ciao3 deletion resulted in embryonic death, vascular dysplasia, impaired differentiation of endothelial progenitor cells, and abnormalities in the expression of ferroptosis-related proteins.

- NARFL polymorphisms were associated with susceptibility to pulmonary hypertension, epilepsy, and neurodegenerative diseases.

- NARFL knockout disrupts its interaction with CIA system-related proteins, leading to decreased aconitase activity, increased IRP1 activity, endothelial cell ferroptosis pathway abnormalities, enhanced ferroptosis and oxidative stress, and ultimately vascular endothelial dysfunction.

- This dysfunction is responsible for the death of embryos in narfl-/- zebrafish and Ciao3-/- mice, as well as the susceptibility to pulmonary hypertension, epilepsy, and neurodegenerative diseases.


**Explanations**:

- Direct human genetic evidence supportive of a Mendelian (recessive) association: a novel homozygous mutation was observed in a consanguineous family with a clearly defined, rare clinical phenotype (DPAVMs with secondary pulmonary hypertension). Strengths: homozygosity in a consanguineous family and decreased expression (functional effect) are consistent with a recessive pathogenic variant. Limitations: this sentence references "our previous study" without presenting segregation statistics, variant annotation, number of affected relatives, or criteria used to assign pathogenicity in this abstract; the full paper would need variant-level details, segregation data, and population/functional evidence to definitively establish a Mendelian causal relationship.

- Model-organism evidence showing a strong loss-of-function phenotype: narfl deletion causes larval lethality and specific vascular/BBB malformations in zebrafish. Relation to claim: supports that loss of this gene's function produces severe, developmentally penetrant phenotypes, increasing plausibility that human mutations could produce Mendelian disease. Limitations: zebrafish homolog effects may differ from human disease manifestations and complete knockout lethality does not by itself prove that the human observed homozygous variant is causal for the reported human phenotype (dose, allelic series, and species differences matter). Note: the sentence contains a typographical error ("structur") copied verbatim from the source.

- Mechanistic cellular/biochemical evidence: narfl deficiency increases cellular iron, oxidative stress, and lipid peroxidation that can be partially rescued by a ferroptosis inhibitor (Ferrostatin-1). Relation to claim: provides a plausible mechanism (iron handling → oxidative stress → cell death) by which NARFL dysfunction could produce tissue pathology consistent with disease. Limitations: fluorescence probe readouts are indirect measures; extent of rescue is reported as partial and quantitative effect sizes, replicates, and statistical significance are not provided here.

- Cellular-level mechanistic evidence in human-derived endothelial cells: NARFL deficiency impairs proliferation, mitochondrial morphology and function, angiogenic (tube formation) ability, and barrier integrity. Relation to claim: connects gene dysfunction to endothelial cell pathology relevant to the vascular phenotype observed in the human family (DPAVMs). Limitations: in vitro HPMEC knockouts model cell-autonomous effects but may not capture organismal complexity; dose dependence, rescue experiments, and whether the human variant recapitulates these specific defects are not detailed here.

- Human genetic association evidence: reported associations between NARFL polymorphisms and susceptibility to pulmonary hypertension, epilepsy, and neurodegenerative diseases. Relation to claim: supports broader clinical relevance of variation at this locus to human disease risk. Limitations: association with susceptibility (polymorphisms) is not equivalent to a Mendelian, high-penetrance causal allele; association studies can reflect linkage disequilibrium, population stratification, or small effect sizes and do not by themselves establish Mendelian inheritance.

- Mechanistic summary linking molecular consequence to pathology: knockout disrupts interactions with CIA (cytosolic iron-sulfur assembly) proteins, reduces aconitase activity and increases IRP1 activity, producing ferroptosis/oxidative stress and endothelial dysfunction. Relation to claim: provides a coherent mechanistic pathway from gene loss to cellular dysfunction and organ-level pathology, strengthening biological plausibility that pathogenic variants in this gene could cause Mendelian disease affecting vasculature and neural tissue. Limitations: mechanistic claims are based on a combination of biochemical and model-system results; the abstract does not present quantitative effect sizes, statistical measures, or evidence directly tying these molecular changes to the specific human pedigree's variant.

- Integrative conclusion linking model phenotypes to human disease susceptibility and embryonic lethality: the authors attribute embryonic death in knockout models and human disease susceptibility to NARFL deficiency. Relation to claim: synthesizes human genetics and cross-species functional data to argue that NARFL disruption can cause severe, likely Mendelian, phenotypes. Limitations: this is a conclusion statement summarizing the study's interpretation; as with prior points, direct variant-level evidence (e.g., segregation, population frequency, functional complementation/rescue by wild-type human allele) is not included in the abstract and would be required for definitive Mendelian classification.


[Read Paper](https://www.semanticscholar.org/paper/29202721282d1a05b8664dbd5e0ba7c589e11f58)


## Other Reviewed Papers


### Wild-type and A315T mutant TDP-43 exert differential neurotoxicity in a Drosophila model of ALS.

**Why Not Relevant**: The provided paper discusses the RNA-binding protein TDP-43 (gene TARDBP) and an A315T missense mutation linked to amyotrophic lateral sclerosis (ALS), using Drosophila overexpression models to study neurotoxicity and mechanisms. It does not mention the gene identifier or name K1109_HUMAN anywhere in the text, nor does it provide data, variants, or mechanistic links involving K1109_HUMAN. Therefore there is no direct or mechanistic evidence in this paper that K1109_HUMAN is associated with any Mendelian disease. 

Relevant specifics from the paper that demonstrate the mismatch: the text repeatedly refers to TDP-43, the A315T TDP-43 missense mutation, and ALS-related phenotypes (neuronal loss, locomotor dysfunction, neuromuscular junction architecture, proteasome/HSP70/apoptosis modulation). These pertain to TARDBP biology and ALS but cannot be extrapolated to support a claim about a different gene (K1109_HUMAN) without additional information linking K1109_HUMAN to TARDBP or to ALS. 

Limitations and caveats: if K1109_HUMAN were an alias, isoform name, or database identifier for TARDBP (which is not indicated in the provided content), one would need explicit mapping evidence (e.g., UniProt entry, HGNC symbol) to treat findings about TDP-43 as evidence for K1109_HUMAN. Because the paper text supplied contains no such mapping or mention, no inference can be made. If you can provide the sequence, alias, or database identifier mapping showing that K1109_HUMAN refers to TARDBP (or another gene discussed), I can re-evaluate relevance and extract supporting excerpts accordingly.


[Read Paper](https://www.semanticscholar.org/paper/d76f9d89bd0fb65e3a117266ec0b0e22760b6ba9)


### The alliance of genome resources: transforming comparative genomics

**Why Not Relevant**: The provided paper content contains a single sentence: "The modular infrastructure developed by the Alliance allows the resource to serve as an extensible “knowledge commons” capable of expanding to accommodate additional model organisms." This sentence makes no mention of the gene K1109_HUMAN, any specific gene, Mendelian disease, phenotype, genetic association, patient data, variant evidence, functional studies, or mechanisms linking genes to disease. It instead refers to software/infrastructure and the potential to include more model organisms. There is therefore no direct evidence (no association data, no statistical results) and no mechanistic evidence (no molecular function, pathway, or phenotypic mechanism) relevant to the claim that K1109_HUMAN is likely associated with at least one Mendelian disease. Limitations of the provided content: it is a single descriptive sentence about infrastructure with no experimental methods, results, or gene-specific information; it cannot be used to support or refute the claim. While the infrastructure description implies that the resource could in the future contain data linking genes to disease, that is speculative and not evidence in this text.


[Read Paper](https://www.semanticscholar.org/paper/b6ed706f5f8cc682b853c888e79df4c7fb0eb2c7)


### Correlation between Phenotype and Genotype in CTNNB1 Syndrome: A Systematic Review of the Literature

**Why Not Relevant**: Relevance is effectively zero because the paper exclusively discusses CTNNB1 Syndrome and genotype–phenotype correlations for CTNNB1 variants. The provided content contains no mention of the gene identifier or symbol 'K1109_HUMAN' (nor obvious synonyms or identifiers for KIAA1109 or other possible mappings). Therefore the paper provides neither direct evidence (observational or genetic data linking K1109_HUMAN to a Mendelian disease) nor mechanistic evidence (functional studies or pathway links) relevant to the claim. To evaluate the claim about K1109_HUMAN, sources that explicitly mention that gene (e.g., gene-specific case reports, OMIM, ClinVar, or functional studies) should be consulted. Limitations: if 'K1109_HUMAN' is an alternate/uncommon alias for CTNNB1 (which is unlikely given the text), that mapping is not stated in the paper content provided; absent any explicit identifier mapping, no inference can be made from this CTNNB1-focused paper to support the claim about K1109_HUMAN.


[Read Paper](https://www.semanticscholar.org/paper/1128b67e6e3a19366fa824315324d962ca2223f3)


### Precision Medicine in Internal Medicine

**Why Not Relevant**: The paper is a general review about clinical genetic testing methods, variant interpretation frameworks, and the current state of gene-disease discovery. It contains no gene-specific data or statements about K1109_HUMAN, no reported variants in that gene, no segregation, case series, functional studies, nor database entries explicitly naming K1109_HUMAN. Therefore it does not provide direct evidence that K1109_HUMAN is associated with any Mendelian disease. The included excerpts only offer background context on how such associations might be discovered and interpreted, and on the fact that, as of 1 February 2019, only ~18% of protein-coding genes were known to cause single-gene disorders. Because there is no mention of K1109_HUMAN anywhere in the provided content, the paper cannot be used to support the specific claim.


[Read Paper](https://www.semanticscholar.org/paper/c7199e8a5226f8d9d55777da0e15e91797bb459a)


### The genotype–phenotype correlations of the CACNA1A-related neurodevelopmental disorders: a small case series and literature reviews

**Why Not Relevant**: The paper exclusively studies CACNA1A and CACNA1A-related neurodevelopmental disorders (GDD/ID, epilepsy, ASD). The claim concerns a different gene identifier, K1109_HUMAN, which does not appear anywhere in the provided paper content. Because the gene names/identifiers do not match and no mapping between K1109_HUMAN and CACNA1A is supplied, this paper provides no direct or mechanistic evidence that K1109_HUMAN is associated with any Mendelian disease. The paper does, however, provide clear evidence that CACNA1A is associated with Mendelian neurodevelopmental disorders — but that is not evidence for K1109_HUMAN. If you intended a different gene (for example, a specific UniProt entry or a KIAA gene), please clarify the exact gene symbol/accession so I can re-evaluate relevant literature.


[Read Paper](https://www.semanticscholar.org/paper/79f5fcb007791190e8376552b51c85a1f327b965)


### The simplest explanation does not have to be preferred: co-occurrence of pathogenic variants in cancer-predisposing genes

**Why Not Relevant**: The provided paper content does not mention the gene K1109_HUMAN, any specific gene identifier, nor any Mendelian disease. The single supplied sentence discusses: frequency and cancer phenotypes associated with MINAS, a review of recent literature, checking the UK 10,000 Genomes Project for participants with tumor predisposition syndromes or multiple primary tumors, and whether effects are additive or synergistic for cancer susceptibility genes (CSGs). None of these elements provide direct evidence that K1109_HUMAN is associated with a Mendelian disease, nor do they provide mechanistic evidence linking that specific gene to Mendelian inheritance of disease. Specifically:

- No occurrence of the string "K1109_HUMAN" or any obvious synonyms/identifiers for that gene is present in the excerpt.
- There are no variant, segregation, functional, biochemical, or inheritance data reported that could connect any gene to a Mendelian disorder.
- The text addresses cancer phenotypes and analyses in aggregate (MINAS, CSGs, UK 10,000 Genomes Project) but contains no gene-level findings, no pathogenicity claims, and no mechanistic descriptions (e.g., protein function, disrupted pathway, loss/gain of function) applicable to K1109_HUMAN.
- The excerpt is a brief high-level methods/aims-style sentence; it lacks results, data, or conclusions that could be evaluated for statistical or practical significance regarding any gene-disease association.

Because of the absence of any mention of K1109_HUMAN or any Mendelian disease evidence in the supplied text, relevance to the claim is effectively zero. If you can provide the full paper text or a section that names K1109_HUMAN (or its gene symbol/aliases, UniProt accession, OMIM ID, etc.), I can repeat the analysis and extract verbatim supporting or refuting sentences and evaluate mechanistic evidence and study limitations. Alternatively, performing a targeted search of the full paper for the gene identifier would be necessary to determine whether the unprovided portions contain relevant evidence.


[Read Paper](https://www.semanticscholar.org/paper/a1372a664bd2c33a0bda638975a8b26d112457fc)


### A frameshift mutation in the SCNN1B gene in a family with Liddle syndrome: A case report and systematic review

**Why Not Relevant**: The provided paper text exclusively describes Liddle syndrome caused by mutations in SCNN1A, SCNN1B, or SCNN1G, and reports a specific frameshift mutation in SCNN1B (c.1806dupG, p.Pro603Alafs*5) in a family with early-onset hypertension and hypokalemia. The gene identifier K1109_HUMAN does not appear anywhere in the supplied content, and no data, variants, co-segregation analyses, functional studies, or mechanistic discussion are provided for K1109_HUMAN. Therefore this paper provides no direct evidence (no reported variants or genetic linkage) and no mechanistic evidence (no protein function, pathway, or interaction data) supporting an association between K1109_HUMAN and any Mendelian disease. 

Limitations and caveats: the conclusion that this paper is non-relevant to the claim is based solely on the excerpt you provided. If the full paper contains additional sections not included here that mention K1109_HUMAN, those were not available for review. To support the claim that K1109_HUMAN is associated with a Mendelian disease, one would need evidence such as: (a) segregation of rare damaging variants in K1109_HUMAN with disease in pedigrees, (b) de novo pathogenic variants in affected individuals, (c) statistically significant case-control enrichment of rare loss-of-function or deleterious variants, and/or (d) functional/mechanistic data showing that disruptive variants in K1109_HUMAN alter protein function in a way consistent with disease biology. None of those evidence types are present in the provided text.


[Read Paper](https://www.semanticscholar.org/paper/b93e9a7f64d9f9ebd09ef30cf3dd0b2fcc350596)


### Case report: A new de novo 6q21q22.1 interstitial deletion case in a girl with cerebellar vermis hypoplasia and developmental delay and literature review

**Why Not Relevant**: This paper does not provide evidence for or against the claim that K1109_HUMAN is associated with a Mendelian disease. The provided text describes a 7.9 Mb de novo interstitial deletion at 6q21q22.1 and discusses clinical phenotypes observed in the patient and literature review, but it does not mention the gene or protein identifier 'K1109_HUMAN' (or any specific gene names) anywhere in the supplied content. Because the paper excerpt does not list the genes included in the deleted interval, does not single out any individual gene as causal, and presents only a broad cytogenetic interval and associated phenotypes, there is neither direct evidence (e.g., mutation/segregation/functional data implicating K1109_HUMAN) nor mechanistic evidence (e.g., gene function consistent with the phenotype, dosage sensitivity, model organism data) relating K1109_HUMAN to a Mendelian disease.

Limitations that prevent using this paper to support the claim:
- No explicit mention of K1109_HUMAN or any gene-level findings in the provided text, so the presence or absence of that gene within the deleted coordinates cannot be ascertained from this excerpt.
- The reported deletion is large (7.9 Mb) and spans many genes; even if K1109_HUMAN were within the interval, the paper does not perform gene-level analyses (e.g., genotype–phenotype correlation, single-gene mutations, segregation, functional assays) that could implicate a specific gene as causative. Large multi-gene deletions provide limited evidence for attributing phenotype to any single gene without additional data.
- The study design (single-case report and literature review of overlapping deletions) is not sufficient to establish a Mendelian disease association for an individual gene without replication, allele-specific variants, or mechanistic follow-up.

Conclusion: Based on the supplied content, the paper is not relevant to assessing whether K1109_HUMAN is likely associated with a Mendelian disease. Additional sources that (a) explicitly name K1109_HUMAN/KIAA1109 in the deleted interval, (b) report pathogenic variants in that gene segregating with disease, or (c) provide functional evidence linking that gene to a phenotype would be required to evaluate the claim.


[Read Paper](https://www.semanticscholar.org/paper/2cf859729d4f79e0d8be861fc43ea509d83c6774)


### LGP2 Facilitates Bacterial Escape through Binding Peptidoglycan via EEK Motif and Suppressing NOD2-RIP2 Axis in Cyprinidae and Xenocyprididae Families.

**Why Not Relevant**: This paper does not provide relevant evidence for or against the claim that K1109_HUMAN is likely associated with at least one Mendelian disease. Reasons: (1) The manuscript focuses on LGP2 (CiLGP2) and NOD2 biology primarily in fish (Ctenopharyngodon idella and other Cyprinidae/Xenocyprididae), not on any human gene labeled K1109_HUMAN. There is no mention of the gene name K1109_HUMAN or any human-specific protein/UniProt identifier. (2) The study reports mechanistic immunology (competition of LGP2 with NOD2 for peptidoglycan binding, suppression of the NOD2-RIP2 axis) and motif/species-specific analyses (EEK motif), but contains no human genetic data: no sequence variants, no genotype–phenotype correlations, no family segregation or case reports, no linkage/association statistics, and no clinical phenotype descriptions that would support a Mendelian disease assignment. (3) Experimental subjects and constructs are fish proteins and recombinant proteins; insertion of the EEK motif into zebrafish and mouse LGP2 did not confer PGN binding activity, underscoring species-specific biology and limiting extrapolation to humans. (4) No statements in the paper claim or imply that LGP2 (in fish or other species) is causally linked to inherited human disease; thus there is no direct evidence (variant pathogenicity, loss/gain-of-function causing a heritable phenotype) or indirect, persuasive mechanistic evidence connecting K1109_HUMAN to a Mendelian disorder. Given these facts, the paper is not relevant to evaluating the claim about K1109_HUMAN and Mendelian disease association.


[Read Paper](https://www.semanticscholar.org/paper/c1bbff49678a28dafdef049d358befae689d2dc4)


### Whole-exome sequencing revealed a novel mutation of the ALMS1 gene in a Chinese family with Alström syndrome: a case report

**Why Not Relevant**: The provided paper content contains a single sentence: "Genetic testing indeed should be incorporated into the diagnosis of syndromic deafness, as it can help avoid misdiagnoses of AS, and expands the mutation spectrum of ALMS1." This sentence mentions ALMS1 and Alström syndrome (AS) and argues for genetic testing in syndromic deafness, but it makes no mention of K1109_HUMAN, presents no genetic association data, no variant-level evidence, no segregation or case reports, and no mechanistic data tying K1109_HUMAN to any Mendelian phenotype. Therefore the paper provides no direct evidence (no association, no statistical results, no pathogenic variants) and no mechanistic evidence (no functional studies, protein interaction data, or pathway links) relevant to the claim that K1109_HUMAN is likely associated with at least one Mendelian disease. Additional limitations: the content is a single declarative sentence with no methods, results, or supporting data; it addresses ALMS1 specifically, which is a different gene, so any inference about K1109_HUMAN would be unsupported. Because the paper does not mention K1109_HUMAN at all, its relevance to the claim is effectively zero.


[Read Paper](https://www.semanticscholar.org/paper/5c07910460140f13b29f4c63c75d324e4579ec67)


### Exome Sequencing Identifies a Novel SGCA Gene Mutation in an Iranian Family with Limb Girdle Muscular Dystrophy: A Case Report

**Why Not Relevant**: The paper reports a novel homozygous SGCA gene mutation causing autosomal recessive limb-girdle muscular dystrophy in a single affected individual from a consanguineous family. The content makes no mention of the gene or protein referred to as K1109_HUMAN (no sequence identifiers, aliases, or annotations linking K1109_HUMAN to SGCA or to this study). Therefore the paper provides no direct or mechanistic evidence supporting or refuting the claim that K1109_HUMAN is associated with at least one Mendelian disease. Additionally, the paper's evidence pertains specifically to SGCA and is a single-family case report (limited sample size, no functional validation in the provided text), so it cannot be extrapolated to other genes without explicit data linking them.


[Read Paper](https://www.semanticscholar.org/paper/46a6274976aa6da24830f44d465c15762c6beb3f)


### Autosomal Dominant Intellectual Development Disorder-6 (MRD6) Without Seizures Linked to a De Novo Mutation in the grin2b Gene Revealed by Exome Sequencing: A Case Report of a Moroccan Child

**Why Not Relevant**: This paper describes a pathogenic variant in the grin2b (GRIN2B) gene causing an autosomal dominant intellectual developmental disorder (MRD6). The text repeatedly names grin2b/GRIN2B (including the OMIM entry 613970), describes a de novo nonsense variant (c.3912C>G, p.Tyr1304Ter) confirmed by Sanger sequencing, notes absence from gnomAD, and applies ACMG criteria to classify the variant as pathogenic. Nowhere in the provided content is the gene identifier “K1109_HUMAN” (or obvious synonyms such as KIAA1109) mentioned, nor are any data presented about that gene. Therefore the paper provides no direct evidence that K1109_HUMAN is associated with any Mendelian disease. Mechanistically, the paper provides strong mechanistic and clinical evidence linking GRIN2B disruption to MRD6 (de novo truncating mutation, phenotype description, ACMG classification), but that mechanistic evidence is about a different gene and cannot be used to infer association for K1109_HUMAN without independent evidence that K1109_HUMAN is an alias for GRIN2B (which it is not).

Limitations and caveats relevant to this assessment:
- Identifier confusion: If “K1109_HUMAN” was intended to refer to a different gene (for example KIAA1109 or another UniProt entry), that mapping should be clarified. As presented, there is no textual or contextual link between K1109_HUMAN and the GRIN2B data in this paper. 
- Absence of negative evidence: The paper does not evaluate K1109_HUMAN at all (neither sequencing nor functional data), so it does not provide evidence against the claim either; it is simply irrelevant to K1109_HUMAN.

Suggested next steps to evaluate the claim about K1109_HUMAN:
- Confirm the correct gene identifier (UniProt entry name, HGNC symbol, or gene symbol). If you meant KIAA1109 or another gene, search the paper/texts explicitly for that name.
- Query resources for Mendelian-disease associations for the correct gene identifier: OMIM, ClinVar, HGMD, and UniProt disease annotations; look for curated disease associations and reported pathogenic variants.
- If you want, provide a paper or text that mentions K1109_HUMAN (or the intended gene identifier) and I will analyze that for direct and mechanistic evidence per your original instructions.


[Read Paper](https://www.semanticscholar.org/paper/67d756e0cc302e5d46fdda9df34403f1f2de9a5a)


### MO005 GENERATION OF NOVEL 3D CO-CULTURE SYSTEMS TO STUDY PODOCYTOPATHIES EX VIVO IN A PERSONALIZED MANNER PARAMETERS IN GLOMERULAR CELLS IN VITRO AND CONTRIBUTE TO PROTEINURIA IN A ZEBRAFISH MODEL *

**Why Not Relevant**: The provided paper content does not contain any mention of the gene name or identifier "K1109_HUMAN" (or a closely matching symbol). The text excerpts supplied discuss multiple kidney-related topics and gene/protein names including KMO, Afmid, Kynu, PKD1, PKD2, various TRP channels (TRPML1, TRPC1/3, TRPM3/4/7, TRPV4, TRPA1), STIM1/2, Orai1/2/3, and inflammatory/biomarker molecules (GDF-15, YKL-40, MMP9), but none of these corresponds to K1109_HUMAN. Key points supporting non-relevance: 

- No direct evidence: There are no sentences or paragraphs reporting genetic linkage, pathogenic variants, segregation, case reports, exome/genome sequencing, or clinical associations that involve K1109_HUMAN or an equivalent gene symbol. Therefore the paper provides no direct evidence for or against an association between K1109_HUMAN and any Mendelian disease. 

- No mechanistic evidence: The mechanistic experiments described (e.g., morpholino knockdown of Afmid/Kmo/Kynu in zebrafish, KMO inhibition in podocytes, characterization of calcium-coupled GPCRs and TRP channels, gadolinium toxicity assays in HK-2 cells) implicate other genes/proteins and pathways (kynurenine pathway, NAD metabolism, calcium signaling) and do not implicate K1109_HUMAN. 

- Study types and limitations: The studies are a mixture of in vitro cell culture assays, zebrafish morpholino knockdowns, pharmacologic inhibition, and descriptive expression profiling. Even if a gene related to K1109_HUMAN function were discussed, these experimental designs (without human genetic data showing segregation or variant pathogenicity) would not on their own establish an association with a Mendelian disease. 

- Scope limitation: Because the provided content appears to be multiple abstracts focusing on renal biology and toxicology, absence of K1109_HUMAN in this text only indicates that this particular paper/content is not relevant to the claim. It does not rule out evidence in other literature. 

Conclusion: Relevance is effectively zero for assessing whether K1109_HUMAN is likely associated with at least one Mendelian disease, because the paper contains neither direct genetic association data nor mechanistic discussion of that gene.


[Read Paper](https://www.semanticscholar.org/paper/0e15561c2c8df273d721bf1649bd4d7e486635ad)


## Search Queries Used

- K1109_HUMAN KIAA1109 Mendelian disease OMIM ClinVar single gene disorder monogenic

- KIAA1109 variant pathogenic mutation exome sequencing family segregation autosomal recessive autosomal dominant case report genetic disease

- KIAA1109 K1109_HUMAN protein function molecular pathway protein interaction expression mouse model zebrafish neurodevelopment

- KIAA1109 genotype phenotype review systematic review case series clinical phenotype developmental delay intellectual disability congenital malformation


## Usage Statistics

- Prompt Tokens: 0

- Completion Tokens: 0

- Total Tokens: 0

- Estimated Cost: $0.0655


## Bibliometric Configuration

- Use Bibliometrics: True

- Author Impact Weight: 0.4

- Citation Impact Weight: 0.4

- Venue Impact Weight: 0.2
