## Supplementary material for "MENDELSEEK: An algorithm that predicts Mendelian Genes and elucidates what makes them special": KCNA3.txt

# Claim: The gene KCNA3 is likely associated with at least one Mendelian disease.

**Status**: processed

**Overall Rating**: 0

**Explanation**:

Supporting evidence:
The provided review identifies KCNA3 (Kv1.3) as the principal voltage-gated potassium channel in leukocytes and states that "alterations in Kv1.3 are associated with multiple immune disorders." It notes that excessive channel activity correlates with autoimmune diseases and reduced currents associate with increased cancer prevalence and immunodeficiencies. The review also indicates that diseases arising from dysregulation of the channel are discussed and that Kv1.3 is a potential immune target in a variety of diseases. These statements establish a connection between KCNA3 function and pathological immune phenotypes, which at least prima facie links KCNA3 biology to human disease.

Caveats or contradictory evidence:
None of the provided excerpts explicitly report germline pathogenic variants in KCNA3 that cause a Mendelian (monogenic, inherited) disease. The language in the review refers to "alterations" and "dysregulations," which can encompass non‑genetic mechanisms (e.g., altered expression, autoantibodies, post‑translational modification, somatic changes, environmental modulation) or complex/ polygenic associations rather than clearly inherited Mendelian disorders. The excerpts do not cite cases of families with segregating KCNA3 mutations, nor do they name an OMIM or similar monogenic condition linked to KCNA3. The bibliometric/relevance indicators supplied are low, and only a single review excerpt is available, so direct primary genetic evidence (case reports, segregation studies, functional characterization of germline variants) is absent.

Analysis of potential underlying mechanisms and implications:
Biologically, the role of Kv1.3 in leukocyte Ca2+ signaling and cell activation makes it plausible that a deleterious germline variant severely disrupting channel function could produce an immunodeficiency phenotype, and that gain‑of‑function variants might predispose to immune hyperactivity or autoimmunity. Thus, mechanistic plausibility exists for KCNA3 to be involved in monogenic immune disease. However, plausibility alone is not evidence of an existing Mendelian association; many genes with plausible mechanisms have no confirmed monogenic diseases. Establishing a Mendelian association requires documented germline variants with appropriate segregation, recurrence, and functional validation—none of which are provided here.

Which rating is most appropriate:
Given the absence of explicit evidence in the provided paper excerpts for germline pathogenic KCNA3 variants causing a Mendelian disease, and the fact that the review's statements can be interpreted as referring to non‑Mendelian dysregulation, the available evidence does not support the claim that KCNA3 is likely associated with at least one Mendelian disease. Although the gene is plausibly linked to immune phenotypes (making a future Mendelian association possible), there is no direct supporting evidence in the supplied materials. Therefore the most appropriate assessment based on the provided evidence is that there is No Evidence to support the specific claim of a Mendelian disease association.


**Final Reasoning**:

I reviewed the provided review excerpt carefully. It links KCNA3 function to immune disorders and states that dysregulation is associated with various pathologies, which supports a general disease connection. However, the claim to be evaluated is specifically about association with at least one Mendelian disease (i.e., a monogenic, inherited disorder). The excerpt does not report germline pathogenic variants, family segregation, or a named Mendelian condition tied to KCNA3. The term "alterations" in a review context can mean many non‑Mendelian phenomena. Mechanistic plausibility exists for KCNA3 variants to cause monogenic immune disease, but plausibility is not evidence. Because no direct genetic evidence for a Mendelian association is present in the supplied material, the correct classification is "No Evidence." I find no flaws in this logic given the evidence provided; if additional papers (case reports, genetic studies, OMIM entries) were supplied that document germline KCNA3 pathogenic variants causing a segregating disease, the rating would warrant revision.


## Relevant Papers


### Kv1.3 in the spotlight for treating immune diseases

**Authors**: María Navarro-Pérez (H-index: 6), Antonio Felipe (H-index: 3)

**Relevance**: 0.25

**Bibliometric Impact**: 0.15839999999999999


**Excerpts**:

- Kv1.3 is the main voltage-gated potassium channel of leukocytes from both the innate and adaptive immune systems.

- Channel function is required for common processes such as Ca2+ signaling but also for cell-specific events.

- In this context, alterations in Kv1.3 are associated with multiple immune disorders.

- Excessive channel activity correlates with numerous autoimmune diseases, while reduced currents result in increased cancer prevalence and immunodeficiencies.

- This review offers a general view of the role of Kv1.3 in every type of leukocyte.

- Moreover, diseases stemming from dysregulations of the channel are detailed, as well as current advances in their therapeutic research.

- Kv1.3 arises as a potential immune target in a variety of diseases.


**Explanations**:

- How it relates: Establishes the gene product (Kv1.3) and its expression in leukocytes, linking KCNA3 to immune cell physiology. Type of evidence: mechanistic/contextual. Strengths/limitations: Strong contextual relevance because gene expression and protein identity are fundamental; however, this sentence does not provide evidence of KCNA3 variants causing Mendelian disease or any genetic association data.

- How it relates: Describes the functional role of the channel in key cellular processes (e.g., Ca2+ signaling), providing a plausible mechanism by which channel perturbation could produce disease phenotypes. Type of evidence: mechanistic. Strengths/limitations: Supports biological plausibility that KCNA3 dysfunction could have pathologic effects; does not constitute direct evidence of hereditary (Mendelian) disorders nor provide genetic/clinical data linking specific variants to disease.

- How it relates: States that alterations in Kv1.3 are 'associated with multiple immune disorders,' which is an explicit claim of disease association. Type of evidence: associative (likely derived from reviewed studies). Strengths/limitations: This is direct language indicating disease associations, but as presented in the abstract it is non-specific (no details on whether associations are genetic, acquired, sporadic, or Mendelian). The sentence does not document familial segregation, pathogenic variants, or inheritance patterns required to support a Mendelian disease claim.

- How it relates: Provides directionality of functional effects: excessive activity linked to autoimmune disease, reduced currents linked to cancer and immunodeficiencies—this ties channel functional changes to distinct disease phenotypes. Type of evidence: mechanistic/associative. Strengths/limitations: Useful for plausibility that KCNA3 dysfunction can cause clinically meaningful immune phenotypes. Limitation: 'Correlates' and 'result in' are broad terms here in a review abstract; no genetic evidence, variant-level data, prevalence, penetrance, or inheritance information are provided to support Mendelian causality.

- How it relates: Indicates the paper is a review covering Kv1.3 roles across leukocytes, signaling that the content synthesizes multiple lines of evidence relevant to function and disease. Type of evidence: contextual/meta-scientific. Strengths/limitations: Confirms the article is a synthesis (not primary genetic discovery), so it may summarize studies that could include genetic data, but the abstract provides no direct genetic or Mendelian evidence itself.

- How it relates: States the review details diseases stemming from channel dysregulation and therapeutic research, implying discussion of disease links. Type of evidence: contextual/associative. Strengths/limitations: Suggests that disease connections are discussed in the full text; however, the abstract gives no specifics about causative mutations, inheritance patterns, or cases of Mendelian disease attributable to KCNA3.

- How it relates: The expert opinion that Kv1.3 is a potential immune target supports the channel's clinical relevance in disease. Type of evidence: opinion/clinical relevance. Strengths/limitations: Reinforces practical significance but does not supply genetic evidence for KCNA3 causing Mendelian disease. As an opinion statement, it cannot substitute for genetic proof.


[Read Paper](https://www.semanticscholar.org/paper/1fb40c38e09ef56721b5304abaf4f3721773053e)


## Other Reviewed Papers


### PINK1 protein in normal human brain and Parkinson's disease.

**Why Not Relevant**: The paper focuses on the PINK1 gene and its role in Parkinson's disease (including the identification of PINK1 mutations causing autosomal recessive parkinsonism and mechanistic studies showing mitochondrial localization). There is no mention of KCNA3 anywhere in the provided text—no genetic data, no mutation reports, no functional or segregation studies, and no clinical descriptions related to KCNA3. Therefore this paper provides neither direct genetic evidence (e.g., pathogenic KCNA3 variants segregating with disease, linkage/association data, ClinVar/OMIM entries) nor mechanistic evidence (e.g., functional assays of KCNA3 variants, relevant expression/localization in disease tissue) to support the claim that KCNA3 is likely associated with a Mendelian disease. To support the claim about KCNA3 one would need a different set of evidence such as: (1) published reports of KCNA3 variants with clear segregation in families and clinical phenotypes consistent with Mendelian inheritance, (2) functional assays showing that those variants alter channel function in a way consistent with disease mechanism, and/or (3) database entries (OMIM, ClinVar) listing KCNA3 as causative for a Mendelian disorder. None of those data are present in this paper, so its relevance to the KCNA3 claim is effectively zero.


[Read Paper](https://www.semanticscholar.org/paper/268102fa8ad5cd5b017e9d023c4cd6f7cbb8cab3)


### Voltage-Gated Potassium Channel Kv1.3 as a Target in Therapy of Cancer

**Why Not Relevant**: The provided paper content is a review focused on the Kv1.3 potassium channel (protein product of KCNA3) as a molecular target in cancer therapy. It supplies background and mechanistic information (channel identity, ion conductance, expression, roles in proliferation and apoptosis, and effects of inhibitors) that increase biological plausibility that perturbing KCNA3 function could influence cell physiology. However, it contains no direct evidence that KCNA3 is associated with any Mendelian (inherited, monogenic) human disease. Specifically, there are no data on germline variants, familial segregation, case reports of inherited syndromes, linkage/association studies implicating KCNA3 in Mendelian disorders, nor animal models showing heritable phenotypes corresponding to human Mendelian disease. The paper's focus on cancer, pharmacologic inhibition, and cellular mechanisms means it is not designed to identify or demonstrate Mendelian genetic associations. Therefore, while mechanistic information is relevant background, it does not meet the criterion of providing direct evidence that KCNA3 is likely associated with at least one Mendelian disease.


[Read Paper](https://www.semanticscholar.org/paper/0bb52c1f10cbb7ae3fc48f36fe59c11a00895f87)


### Voltage-Gated Potassium Channels Kv1.3--Potentially New Molecular Target in Cancer Diagnostics and Therapy.

**Why Not Relevant**: The provided paper content is a review focusing on Kv1.3 channel biology, expression (including in cancer), subcellular localization, functional roles (membrane potential, proliferation, apoptosis), and potential as a therapeutic target. It contains mechanistic and disease-context information relevant to how KCNA3 (the gene encoding Kv1.3) could affect cell physiology and contribute to disease processes, particularly cancer. However, it contains no human genetic data: there are no reports of germline variants in KCNA3, no case reports or family pedigrees, no linkage or segregation analyses, no mention of pathogenic variants, and no explicit statement that mutations in KCNA3 cause any Mendelian disorder. Therefore the paper does not provide direct evidence supporting the claim that KCNA3 is associated with at least one Mendelian disease. The mechanistic information increases biological plausibility that KCNA3 dysfunction could produce disease phenotypes, but without genetic/clinical data this is speculative and insufficient to support the claim.


[Read Paper](https://www.semanticscholar.org/paper/33ab568f80dc705b858735c2f63c4006eb999adb)


### Kv1.3 K+ channel physiology assessed by genetic and pharmacological modulation.

**Why Not Relevant**: The provided paper excerpt is a functional/physiological review of Kv1.3 (KCNA3) focusing on pharmacological modulation and knockout animal models and their links to complex pathologies (autoimmune disease, obesity, cancer). It contains mechanistic and phenotypic context that makes KCNA3 plausible as biologically important, but it contains no human genetic evidence (no case reports, no segregation analyses, no pathogenic variants, no linkage or association data specific to Mendelian inheritance). Therefore it does not provide direct evidence that KCNA3 is associated with a Mendelian (single-gene, typically familial) disease. Functional knockout or pharmacological data alone are insufficient to establish a gene as causing a Mendelian disorder in humans.


[Read Paper](https://www.semanticscholar.org/paper/ee5fed6038adb9c3609312e23e907cbf69ba0b44)


### Therapeutic role of voltage-gated potassium channels in age-related neurodegenerative diseases

**Why Not Relevant**: The provided paper excerpt is a review focusing on the functional role of voltage-gated potassium channels and their involvement in neurodegenerative (largely complex) diseases, oxidative stress effects, and therapeutic targeting. It does not mention the gene KCNA3 or Kv1.3 explicitly, nor does it present any genetic data (mutation reports, segregation analyses, allele frequencies, or clinical phenotypes) that would indicate KCNA3 is causal for a Mendelian (monogenic) disease. The disorders discussed (Alzheimer Disease, Parkinson’s Disease, Huntington Disease, Spinocerebellar Ataxia) are mainly complex or have distinct genetic architectures; no sentences in the excerpt describe an inherited Mendelian syndrome attributable to a KCNA-family gene. Therefore, while the paper provides mechanistic plausibility that KV channel genes could be disease-relevant, it provides no direct evidence supporting the specific claim that KCNA3 is likely associated with at least one Mendelian disease.


[Read Paper](https://www.semanticscholar.org/paper/edb68f971664978aed396732eece5420fea0f1e2)


### Regulatory T‐cell function: When suppressor cells can't suppress

**Why Not Relevant**: The provided paper content discusses Treg cell phenotype and FoxP3 expression and their functional consequences in immune regulation. The claim concerns the gene KCNA3 and its likely association with a Mendelian disease. The excerpt contains no mention of KCNA3 (gene name, protein product, channel function, expression pattern, variants, or disease associations) and contains no genetic or clinical data relevant to Mendelian disease. It therefore supplies neither direct evidence (no genetic association, case reports, segregation data, or pathogenic variants described) nor mechanistic evidence linking KCNA3 to Mendelian disease (no pathway, functional perturbation, or downstream effect involving KCNA3). In addition, the excerpt lacks methodological detail, sample size, statistical analyses, or clinical context that might allow indirect inference. For these reasons the relevance score is 0.0: the paper content as provided is not relevant to assessing whether KCNA3 is associated with any Mendelian disease.


[Read Paper](https://www.semanticscholar.org/paper/77de946672a784cbb3ea556c2e269670e416143b)


### Expanding the spectrum of phenotypes for MPDZ: Report of four unrelated families and review of the literature

**Why Not Relevant**: The provided paper content exclusively discusses the gene MPDZ and its association with biallelic variants causing a heterogeneous Mendelian phenotype. There is no mention of the gene KCNA3 anywhere in the supplied text, nor are there any data, methods, results, or mechanistic discussions that pertain to KCNA3. Key statements from the paper refer specifically to MPDZ (for example: "MPDZ, a gene with diverse functions ... is associated with a spectrum of clinically heterogeneous phenotypes with biallelic perturbation" and "Using exome and genome sequencing, we identified ... variants in MPDZ"), demonstrating the study's focus on MPDZ rather than KCNA3. Because the paper contains no direct evidence (no reported KCNA3 variants, genotype-phenotype correlations, segregation, or case reports) and no mechanistic evidence (no functional studies, animal models, or biochemical data) related to KCNA3, it provides no support for the claim that KCNA3 is likely associated with at least one Mendelian disease. The only possible, very weakly inferential point is that the paper uses exome/genome sequencing and animal-model analyses as approaches to establish gene-disease links; however, that methodological information is generic and does not supply evidence about KCNA3 specifically. Therefore the relevance to the claim is effectively zero. If you want an evidence-based assessment of KCNA3 itself, I can search the literature/databases (ClinVar, OMIM, PubMed) for reported KCNA3 pathogenic variants, segregation evidence, functional studies, or clinical case reports that would bear on its Mendelian-disease status.


[Read Paper](https://www.semanticscholar.org/paper/6ab0ca0c7127a6fd16fc33ead203e54f32e60f02)


### A LGI1 Mutation (C.263 T>A P.Leu88*) Identified in a Chinese Patient with Epilepsy: A Case Study

**Why Not Relevant**: The paper reports a pathogenic LGI1 mutation causing familial epilepsy and notes that LGI1 regulates voltage-gated potassium channels. However, it does not mention KCNA3 anywhere, nor does it present genetic, clinical, or experimental evidence that variants in KCNA3 are associated with a Mendelian disease. The only connection to KCNA3 would be an indirect, speculative mechanistic inference because LGI1 influences potassium channel activity and KCNA3 encodes a voltage-gated potassium channel subunit. Such an inference is weak: demonstrating that a regulator of potassium channels causes disease (LGI1) does not establish that a particular potassium channel gene (KCNA3) is itself a Mendelian disease gene. Additionally, the paper is based on a single family/case report about LGI1, limiting generalizability to other genes. Therefore the content is not relevant evidence for the claim that KCNA3 is likely associated with at least one Mendelian disease.


[Read Paper](https://www.semanticscholar.org/paper/ebf243fd07263d4622f87673ab3433235af19f77)


### Evaluating the effectiveness of cascade clinical screening for familial glaucoma in rural Western Australia

**Why Not Relevant**: This paper does not provide evidence for or against the claim that KCNA3 is likely associated with at least one Mendelian disease. The manuscript focuses on clinical family cascade screening for primary open-angle glaucoma and on programmatic/epidemiologic issues; it does not mention KCNA3, report genetic testing results for KCNA3, describe variants in KCNA3, present segregation data, functional assays, linkage analyses, case reports, or epidemiologic associations involving KCNA3. The only gene explicitly referenced in a Mendelian context is MYOC (as an example of a Mendelian glaucoma gene), which illustrates that the authors consider Mendelian genetic screening relevant for glaucoma but does not implicate KCNA3. To support the claim about KCNA3 would require gene-specific evidence such as: (a) identification of pathogenic or likely pathogenic KCNA3 variants segregating with a Mendelian phenotype in families, (b) multiple unrelated cases with consistent phenotype and KCNA3 variants, (c) functional studies showing variant impact on KCNA3 protein function consistent with disease mechanism, or (d) inclusion of KCNA3 in curated disease-gene databases (e.g., OMIM, ClinGen) with asserted disease relationship. None of these elements are present in this paper. Additional limitations of the paper relative to the claim: small sample of index cases consenting to registry follow-up, study design focused on screening uptake and logistics (not gene discovery), and absence of any genetic-sequencing or variant-analysis methods or results. Given the complete absence of KCNA3 content, the relevance to the specific claim is effectively zero.


[Read Paper](https://www.semanticscholar.org/paper/f7d91140ebcf7eda3e66c9d4b71818477daa7ef0)


### Immune Phenotypic Characterization of a TRAIL-Knockout Mouse

**Why Not Relevant**: The provided paper content concerns immunological phenotyping of TRAIL (TNF-related apoptosis inducing ligand) knockout mice and the effects of TRAIL on T-cell proliferation and dendritic cell populations. Nowhere in the supplied text is the gene KCNA3 (or Kv1.3 potassium channels, potassium channel genes, or any human Mendelian disease) mentioned. The study is an experimental mouse immunology study, not a human genetic association study, variant segregation analysis, clinical case series, or OMIM-style annotation that would provide evidence for KCNA3 being associated with a Mendelian disease. Therefore this paper provides no direct evidence (no case reports, linkage/segregation, pathogenic variants, or statistical association) and no mechanistic evidence linking KCNA3 to a Mendelian disorder. Limitations: the assessment is based only on the content you provided (Simple Summary and Abstract lines). If KCNA3 evidence were present elsewhere in the full paper (methods, results, supplements) that was not included here, it is not available to evaluate. Given the provided text, relevance to the claim is effectively nil.


[Read Paper](https://www.semanticscholar.org/paper/785b498e879d97cdc3e6bd03eac066b3e4bb3360)


### Genetic Analysis in a Familial Case With High Bone Mineral Density Suggests Additive Effects at Two Loci

**Why Not Relevant**: This paper does not mention KCNA3 anywhere in the provided content. It reports whole-exome sequencing in a family segregating high bone mineral density and identifies VAV3 and ADGRE5 as candidate Mendelian contributors, with mechanistic support from osteoclast biology and mouse knockouts. Because KCNA3 is neither reported among identified variants nor discussed mechanistically, the paper provides no direct evidence that KCNA3 is associated with any Mendelian disease. Absence of mention in this single study is not definitive disproof of a KCNA3–Mendelian disease association (absence of evidence is not evidence of absence), but for the claim as stated, this paper supplies no supporting data and thus is irrelevant to establishing KCNA3 as likely associated with a Mendelian disease. Limitations: the study used exome sequencing in only six family members (three affected, three unaffected) from a single pedigree focusing on bone phenotype; the authors do not provide a comprehensive negative report listing all genes evaluated or explicitly stating that KCNA3 was evaluated and found negative, so we cannot conclude KCNA3 was definitively tested and excluded in this dataset.


[Read Paper](https://www.semanticscholar.org/paper/de079d062b38ceaec92ab3987723acef25f28dff)


### Targeted knockout of Kcna3 and its function in Jurkat T cells using the CRISPR/Cas9 system

**Why Not Relevant**: The paper provides experimental evidence that KCNA3 (Kcna3) is the gene encoding the Kv1.3 channel and that its disruption in Jurkat T cells alters channel currents, calcium signaling, and IL-2 secretion. These are mechanistic and functional data in an in vitro T cell line. However, the claim concerns whether KCNA3 is likely associated with at least one Mendelian disease (i.e., a human genetic disorder with Mendelian inheritance). This paper contains no data on human patients, familial segregation, pathogenic germline variants, genotype-phenotype correlations, population genetics, or clinical phenotypes; it only reports CRISPR/Cas9-mediated knockout experiments in an immortalized cell line. Therefore it does not provide direct evidence for or against a Mendelian disease association. At best, the functional data increase biological plausibility that KCNA3 variation could have physiological consequences, but that is insufficient to infer a Mendelian disease association without human genetic/clinical evidence. Because the study lacks human genetic or clinical data, its relevance to the specific claim is very low.


[Read Paper](https://www.semanticscholar.org/paper/44b12c120cc4b4a01a11635a5a7c74fb072065a5)


### 007 Upregulation of miR-941 in peripheral CD14+ monocytes enhances osteoclast activation and osteolysis in patients with psoriatic arthritis: a potential diagnostic biomarker and treatment target

**Why Not Relevant**: The provided paper content contains no mention of the gene symbol KCNA3, its common protein name Kv1.3, the terms "potassium channel", or any direct references to Mendelian disease associations. The abstracts focus on: miR-941 in CD14+ monocytes and osteoclast activation in psoriatic arthritis; eosinophil degranulation in DRESS; non-desmoglein autoantibodies (including Dsc3, M3AR, SPCA1/ATP2C1) in pemphigus vulgaris; IL-27/IL-15 signaling in contact hypersensitivity; and MMP-3 and alpha2-antiplasmin in systemic sclerosis. While some gene names appear in the text (for example, "ATP2C1" and "SPCA1"), none are KCNA3 nor do any excerpts link potassium channel function or KCNA3 variants to Mendelian diseases. Therefore there is neither direct evidence (e.g., variant–disease co-segregation, patient genotype data, or clinical descriptions of a KCNA3-linked Mendelian disorder) nor mechanistic evidence (e.g., functional assays of KCNA3/Kv1.3, pathway analyses implicating KCNA3) in the provided content.

Limitations and caveats: (1) The material appears to be a set of conference abstracts (limited scope and length); if other sections of the full papers not included here discuss KCNA3, those were not available for review. (2) Absence of evidence in these excerpts is not definitive evidence that KCNA3 has no Mendelian associations elsewhere in the literature; it only indicates no support for the claim within the provided text.


[Read Paper](https://www.semanticscholar.org/paper/34d76596313e6374fa5ce415741db2e42e3a1237)


## Search Queries Used

- KCNA3 mutation human case report familial Mendelian disease Kv1.3

- KCNA3 Kv1.3 review channelopathy voltage gated potassium channel Mendelian disease review

- KCNA3 knockout mouse phenotype immunology T cell Kv1.3 electrophysiology functional study

- KCNA3 de novo variant exome sequencing undiagnosed disease cohort rare variant pathogenic ClinVar loss of function gain of function


## Usage Statistics

- Prompt Tokens: 0

- Completion Tokens: 0

- Total Tokens: 0

- Estimated Cost: $0.0638


## Bibliometric Configuration

- Use Bibliometrics: True

- Author Impact Weight: 0.4

- Citation Impact Weight: 0.4

- Venue Impact Weight: 0.2
