## Supplementary material for "MENDELSEEK: An algorithm that predicts Mendelian Genes and elucidates what makes them special": KIF3A.txt

# Claim: The gene RENBP is likely associated with at least one Mendelian disease.

**Status**: processed

**Overall Rating**: -1

**Explanation**:

No relevant papers were found that support or refute this claim.


## Relevant Papers


## Other Reviewed Papers


### Modification of β-adrenoceptor signal transduction pathway by genetic manipulation and heart failure

**Why Not Relevant**: The provided paper content is a single sentence about regulation of β-adrenergic (β-AR) signal transduction in the failing heart: "Both local and circulating renin-angiotensin systems, sympathetic nervous system and endothelial cell function appears to regulate the status of β-AR signal transduction pathway in the failing heart." This sentence does not mention the gene RENBP, any genetic variants, inheritance patterns, clinical phenotypes, or Mendelian disease associations. While the sentence references the renin-angiotensin system (a physiological pathway in which renin participates), it provides no data, experimental results, or discussion linking RENBP (renin-binding protein gene) to disease, nor does it present mechanistic evidence about RENBP expression, function, pathogenic variants, or hereditary transmission. There is therefore no direct evidence (no genetic association, case reports, segregation analyses, or variant functional studies) nor usable mechanistic evidence specifically involving RENBP. Additionally, the content lacks methodology, results, statistical analysis, or clinical context needed to assess any gene-disease relationship. For these reasons the excerpt is not relevant to the claim that "The gene RENBP is likely associated with at least one Mendelian disease."


[Read Paper](https://www.semanticscholar.org/paper/37a281e9e4518171e09787f9b16f354a79d873db)


### Precision Medicine in Internal Medicine

**Why Not Relevant**: The provided paper content is a general review about genetic testing strategies, exome/genome sequencing, variant interpretation, and the state of gene–disease discovery as of early 2019. It does not mention the gene RENBP, present any variant-level data for RENBP, report any cohort, case, segregation, functional, or statistical evidence linking RENBP to a Mendelian phenotype, nor does it include mechanistic experimental data specific to RENBP. Therefore the paper provides only indirect, contextual information (about methods and the general landscape of gene discovery) that can inform plausibility but offers no direct evidence for or against the claim that RENBP is associated with at least one Mendelian disease. Key limitations: absence of RENBP-specific data, lack of case reports or segregation analyses, no functional assays or population genetics data for RENBP, and the snapshot date (2019) which may predate subsequent discoveries. Because of these gaps, this source cannot be used to support the claim beyond noting that many genes remain uncharacterized and that ES/GS could in principle discover novel gene–disease relationships.


[Read Paper](https://www.semanticscholar.org/paper/c7199e8a5226f8d9d55777da0e15e91797bb459a)


### Detailed genetic and clinical analysis of a novel de novo variant in HPRT1: Case report of a female patient from Saudi Arabia with Lesch–Nyhan syndrome

**Why Not Relevant**: The paper exclusively investigates a case of Lesch–Nyhan syndrome caused by a de novo frameshift variant in HPRT1 and associated X-inactivation and expression changes. The gene RENBP is not mentioned anywhere in the provided text, no variants in RENBP are reported, and no mechanistic discussion links RENBP to the disease phenotypes or to HPRT1-related pathways. Therefore this paper provides no direct genetic, clinical, or mechanistic evidence that RENBP is associated with any Mendelian disease. Limitations relevant to this assessment: absence of RENBP in the paper's content means the paper cannot be used as supporting evidence for the claim; however, lack of mention here does not negate evidence for RENBP that might exist in other studies not provided.


[Read Paper](https://www.semanticscholar.org/paper/16daa186f81b3719f537f57e29a674411e64bb19)


### Copy number variation associated with Kallmann syndrome: new genetics insights from genome-wide studies.

**Why Not Relevant**: The provided paper content does not mention RENBP, does not give genomic coordinates or gene-level annotation for the reported microdeletions, and contains only a single-sentence summary from one-family array analysis. Therefore it cannot provide direct or mechanistic evidence that RENBP is associated with a Mendelian disease. Additional information that would be needed to evaluate the claim includes: the chromosomal locations and gene content of the five CNVs, whether RENBP lies within any of them, segregation of any RENBP-impacting variant with disease in the family, functional evidence that RENBP disruption causes disease-relevant phenotypes, and replication in additional unrelated affected individuals or families. Because none of that information is present in the excerpt, the relevance to the claim is negligible.


[Read Paper](https://www.semanticscholar.org/paper/9456d3fa700e40348b9beab88065ed630ec194cc)


### A De novo Mutation in the COL1A1 Gene Leading to Severe Osteogenesis Imperfecta: Case Report and Review of the Literature

**Why Not Relevant**: This paper does not provide any direct or mechanistic evidence relevant to the claim that RENBP is likely associated with at least one Mendelian disease. The study describes a novel heterozygous COL1A1 (c.2174G > T/p.(G725V)) pathogenic variant causing severe osteogenesis imperfecta in a fetus, and discusses COL1A1 sequence context, pathogenicity classification, prenatal findings, and implications for genetic counseling. The provided text contains no mention of the RENBP gene, no reported RENBP variants, no phenotype descriptions mapped to RENBP, and no mechanistic links (biochemical pathways, protein interactions, or genetic evidence) connecting COL1A1 and RENBP. Therefore: 

- There is no direct evidence: the paper does not report any RENBP mutation, association, or Mendelian phenotype attributable to RENBP.
- There is no mechanistic evidence: the paper does not discuss RENBP, nor any pathways or interactions that would plausibly link COL1A1 mutations to RENBP function or suggest RENBP involvement in osteogenesis imperfecta or other Mendelian disorders.

Limitations of this conclusion: absence of RENBP discussion in this paper is not disproof that RENBP is associated with Mendelian disease; it only shows this particular paper is irrelevant to that claim. To evaluate the claim about RENBP, targeted data are needed (e.g., case reports of RENBP variants with segregating phenotypes, ClinVar/HGMD entries for RENBP, functional studies, or population genetics evidence). Recommended next steps are literature/database searches specifically for RENBP in ClinVar, HGMD, OMIM, PubMed, and gnomAD.


[Read Paper](https://www.semanticscholar.org/paper/da7d630add2f62f5b7bd8087fce91e30f87dbc38)


### Autosomal Dominant Intellectual Development Disorder-6 (MRD6) Without Seizures Linked to a De Novo Mutation in the grin2b Gene Revealed by Exome Sequencing: A Case Report of a Moroccan Child

**Why Not Relevant**: The paper content exclusively discusses GRIN2B (grin2b) and a specific GRIN2B nonsense variant causing autosomal dominant intellectual development disorder-6 (MRD6). There is no mention of RENBP, no data on RENBP variants, and no mechanistic discussion linking RENBP to disease in this text. Therefore the paper provides no direct or mechanistic evidence that RENBP is associated with any Mendelian disease and cannot be used to support or refute the claim about RENBP. Any inference regarding RENBP would require data or discussion not present in this paper.


[Read Paper](https://www.semanticscholar.org/paper/67d756e0cc302e5d46fdda9df34403f1f2de9a5a)


### STEM-18. RENIN-ANGIOTENSIN SYSTEM IDENTIFIED BY PHAGE DISPLAY SCREENING IS A NEW THERAPEUTIC TARGET FOR GLIOBLASTOMA AND REGULATES PROLIFERATION OF GLIOBLASTOMA STEM CELLS

**Why Not Relevant**: This paper text does not mention the gene RENBP (renin binding protein) nor present any human genetic, clinical, or pedigree evidence linking RENBP to a Mendelian disease. The manuscript excerpts focus on components of the renin–angiotensin system (renin, angiotensinogen, ACE), glioblastoma stem cells, transcription factors (e.g., TCF12, OLIG2, ID1), and pharmacologic manipulation of RAS in cancer models. While renin and ACE are discussed, there are no statements about RENBP sequence variation, loss-of-function or gain-of-function mutations, genotype–phenotype co-segregation, Mendelian inheritance patterns, case reports, population genetics, or clinical phenotypes that would constitute direct or persuasive mechanistic evidence for RENBP causing a Mendelian disease. Therefore the paper provides no direct evidence and no mechanistic genetic evidence relevant to the claim. Limitations and caveats: (1) RAS component expression in tumors (renin, angiotensinogen, ACE) and responses to RAS inhibitors are cancer biology observations and do not imply germline Mendelian disease associations for other RAS-related genes such as RENBP. (2) Absence of RENBP mention in this text means no material here can be quoted to support or refute the claim; assessing RENBP’s Mendelian disease associations would require querying human genetics resources (OMIM, ClinVar, HGMD) or primary genetic studies, which are outside the content of this paper. (3) Even if RENBP were functionally related to renin, mechanistic involvement in a pathway active in cancer does not by itself indicate a Mendelian (germline) disease association.


[Read Paper](https://www.semanticscholar.org/paper/c95004fdb440bce11a57417dd186eeeee0e31bc6)


### MICRORNA REGULATION OF INTRA-TUMOUR METABOLIC HETEROGENEITY IN GLIOBLASTOMA MULTIFORME

**Why Not Relevant**: The provided paper content contains no mention of the gene RENBP, no germline/Mendelian genetic analyses, and no data linking any variants to inherited disease. The abstracts focus on somatic cancer biology (glioblastoma multiforme), microRNA regulation, cell-line and tumour-sample assays, signalling receptors (AT1R, AT2R), Rho GTPase pathway components (DOCK4, DOCK9), and GPR158/microRNA interactions. Specific relevant points from the content that demonstrate lack of relevance include: (1) gene names discussed are AT1R, AT2R, DOCK4, DOCK9, GPR158 and microRNAs (mir-449a) — RENBP is not listed; (2) study designs are tumour-focused (surgical specimens, patient-derived cell lines, in vitro assays, animal models) and address somatic processes like proliferation, migration, angiogenesis and metabolic heterogeneity rather than inherited disease; (3) there are no reports of familial segregation, inherited variants, loss-of-function or gain-of-function germline mutations, clinical case reports of Mendelian phenotypes, or references to databases of Mendelian disorders (e.g., OMIM/ClinVar) that would support a claim of RENBP being linked to a Mendelian disease; and (4) the statistical and experimental results presented (e.g., P<0.001 for lactate production differences) pertain to tumour metabolic assays and are not genetic-association or inheritance data. Because none of the required types of evidence for the claim (direct variant–disease associations, inheritance patterns, family studies, or mechanistic germline pathogenicity data) are present in the provided content, the paper is not relevant to assessing whether RENBP is likely associated with at least one Mendelian disease.


[Read Paper](https://www.semanticscholar.org/paper/3ac674acb67328c84dd9e3b7d830a51bf11836ce)


### SSBP1: A Target Molecule for Prevention of Fibrosis

**Why Not Relevant**: This paper does not provide evidence relevant to the claim that RENBP is likely associated with at least one Mendelian disease. The text discusses cardiac fibrosis, cellular sources of extracellular matrix, profibrotic signaling (e.g., TGF-β, Nox/ROS), angiotensin II and the renin-angiotensin-aldosterone system (RAS) in general, and the mitochondrial single-stranded DNA-binding protein 1 (SSBP1). Nowhere in the provided content is the gene name RENBP, renin-binding protein, RENBP variants, patient phenotypes, family segregation, linkage, or any Mendelian inheritance patterns mentioned. Mentions of the “renin-angiotensin-aldosterone system (RAS)-related hormones” or of “angiotensin II (Ang II)” are general physiological references to the RAS pathway and do not constitute evidence implicating RENBP specifically. The discussion of SSBP1 is about a different gene/protein (mitochondrial single-stranded DNA-binding protein 1) and does not establish a connection to RENBP or to Mendelian disease causation. There is no genetic, clinical, or variant-level data, nor any mechanistic experiments involving RENBP in this excerpt. The only possible (but highly speculative) link is that RENBP, by name, might be functionally related to components of the renin-angiotensin system; however, this paper provides no data, citations, or arguments that connect RENBP to disease phenotypes, inheritance, or pathogenic mechanisms. Therefore the content is not relevant to evaluating the claim, and no direct or mechanistic evidence for or against the association of RENBP with a Mendelian disease can be extracted from this paper excerpt.


[Read Paper](https://www.semanticscholar.org/paper/347a80db1be4c2dd8b3932bf762582a3ca60e394)


### CBIO-22. SELECTIVE INHIBITION OF HDAC6 AFFECTS CILIOGENESIS AND PREVENTS GLIOBLASTOMA GROWTH

**Why Not Relevant**: The provided paper content does not mention the gene RENBP (renin binding protein) nor any Mendelian disease or germline genetic disorder. The text focuses on glioblastoma and other brain tumor biology (e.g., STAT3, SLUG, HDAC6, PON2, CK2, AT1R/AT2R, KIF3A, NFIA), experimental interventions in cancer cell lines, TCGA transcriptomic analyses, and pharmacologic inhibition—i.e., somatic tumor pathways and therapeutic studies. There are no sentences reporting germline variants, familial segregation, monogenic disease phenotypes, OMIM/ClinVar citations, or functional studies of RENBP relevant to inherited disease. Because none of the necessary direct or mechanistic evidence (germline mutations causing a Mendelian phenotype, cosegregation in families, pathogenic variant reports, or animal models demonstrating a monogenic disease mechanism) is present in the excerpt, it provides no support for or against the claim that RENBP is likely associated with at least one Mendelian disease. To evaluate that claim one would need literature or database evidence reporting pathogenic germline RENBP variants with Mendelian inheritance, clinical case series, or functional assays linking RENBP loss/gain-of-function to a heritable disease phenotype (e.g., OMIM, ClinVar, peer-reviewed genetic studies, or human pedigrees).


[Read Paper](https://www.semanticscholar.org/paper/f20ecb7bb4d1fa038b160c035d052805db1f9dd5)


### Contribution of rare transmitted and de novo variants among 2 , 871 1 congenital heart disease probands

**Why Not Relevant**: I found no mention of the gene RENBP in the provided paper content. The excerpt describes an exome-sequencing study of congenital heart disease (CHD) probands, methods for calling de novo and transmitted variants, and enrichment analyses for known CHD genes, but it does not name RENBP nor present any data, results, or mechanistic discussion involving RENBP. Specific points relevant to this determination: 

- The text repeatedly refers to curated CHD gene sets (e.g., "a set of 212 human CHD genes (H-CHD genes)", "human orthologs of 61 mouse CHD genes"), but the actual gene lists are referenced as being in Supplementary Excel files which were not provided in the content you supplied. Thus, while RENBP could theoretically appear in those supplements, there is no evidence for that in the quoted main text. 

- All reported gene-level results and examples (e.g., GDF1, MYH6, FLT4) are CHD-related genes; none are RENBP. No sentence or paragraph in the provided content asserts an association between RENBP and any Mendelian disease, nor does it describe a plausible mechanistic role for RENBP in CHD or other Mendelian disorders. 

- Because the supplied content lacks any direct mention or data concerning RENBP, there is no direct evidence supporting or refuting the claim that "RENBP is likely associated with at least one Mendelian disease." There is also no mechanistic evidence in the provided excerpt that would strengthen or weaken such a claim. 

Limitations: The full paper (including Supplementary Excel files and any unquoted sections) might contain gene lists or additional analyses that mention RENBP. My conclusion is strictly based on the provided excerpt; without the full main text and supplements I cannot definitively exclude the possibility that RENBP is discussed elsewhere in the full publication. Based on the supplied content alone, however, the paper provides no relevant evidence for the claim.


[Read Paper](https://www.semanticscholar.org/paper/e5830d9e95469f22518560007746202e62db26a0)


## Search Queries Used

- RENBP mutation Mendelian disease exome sequencing case report monogenic de novo inherited variant

- RENBP ClinVar OMIM ClinGen gnomAD HGMD pathogenic variant clinical significance RENBP entry

- RENBP renin binding protein function renin angiotensin system knockout mouse phenotype developmental expression functional study

- RENBP rare variant burden study gene based association rare disease loss of function missense de novo copy number variation CNV deletion duplication neighboring gene


## Bibliometric Configuration

- Use Bibliometrics: True

- Author Impact Weight: 0.4

- Citation Impact Weight: 0.4

- Venue Impact Weight: 0.2
