## Supplementary material for "MENDELSEEK: An algorithm that predicts Mendelian Genes and elucidates what makes them special": KIF20B.txt

# Claim: The gene KIF20B is likely associated with at least one Mendelian disease.

**Status**: processed

**Overall Rating**: -1

**Explanation**:

No relevant papers were found that support or refute this claim.


## Relevant Papers


## Other Reviewed Papers


### A systematic review of the validated monogenic causes of human male infertility: 2020 update and a discussion of emerging gene–disease relationships

**Why Not Relevant**: The provided paper content is a systematic review of genes implicated in male infertility and explains methods, scoring, and overall outcomes, but the supplied text does not mention KIF20B anywhere nor provide any gene-specific data about KIF20B. Therefore the content cannot directly support or refute the claim that KIF20B is associated with at least one Mendelian disease. The paper could be indirectly relevant if KIF20B appears in the authors' full gene lists or spreadsheets (hosted at imigc.org) or in parts of the review not included here, but based solely on the provided excerpts there is no evidence specific to KIF20B. Additional steps to evaluate the claim would be: (1) inspect the full review's gene lists/spreadsheets linked at imigc.org for any KIF20B entries and their clinical validity scores, (2) search primary literature for KIF20B disease associations, and (3) examine functional/segregation evidence for KIF20B variants in Mendelian pedigrees.


[Read Paper](https://www.semanticscholar.org/paper/60ce04d3c61b5475a3b5f2e99ed2865bdcb38828)


### Genetic Modifiers of Mendelian Monogenic Collagen IV Nephropathies in Humans and Mice

**Why Not Relevant**: The supplied paper content does not mention KIF20B or present any data, observations, or mechanistic discussion linking KIF20B to Mendelian disease. The text exclusively discusses familial hematuria, Alport syndrome, and pathogenic variants in the collagen-IV genes COL4A3, COL4A4, and COL4A5, and it refers generally to investigations of candidate genetic modifiers in patients and mouse models. There are no verbatim sentences or paragraphs in the provided excerpt that name KIF20B, describe variants in KIF20B, show segregation of KIF20B variants with disease, demonstrate functional data implicating KIF20B, or propose a mechanistic pathway by which KIF20B would cause or modify a Mendelian disorder.

Because of this, the paper content as provided offers neither direct evidence (e.g., identified pathogenic/likely pathogenic KIF20B variants segregating with disease, case reports, or cohort associations) nor mechanistic evidence (e.g., biochemical, cell biological, or animal-model data implicating KIF20B) to support the claim that KIF20B is likely associated with at least one Mendelian disease. Note that absence of KIF20B in the excerpt is not definitive proof that the full article contains no mention of KIF20B; however, based solely on the supplied text, there is no relevant evidence.

If you want to establish whether this paper (in full) contains relevant information about KIF20B, I recommend searching the full text for the string "KIF20B" (and common synonyms) or providing additional portions of the paper. If you want evidence for or against KIF20B being linked to Mendelian disease more broadly, I can search the literature/databases (ClinVar, OMIM, PubMed) for reported pathogenic variants, disease associations, segregation, and functional studies for KIF20B.


[Read Paper](https://www.semanticscholar.org/paper/9cedc7aad43e0bb7a16afb724682bc76ec9dd0f3)


### Case Report: A Novel De Novo Missense Mutation of the GRIA2 Gene in a Chinese Case of Neurodevelopmental Disorder With Language Impairment

**Why Not Relevant**: The paper exclusively reports a novel de novo missense mutation in the GRIA2 gene associated with NEDLIB and discusses GRIA2 mutation spectrum and diagnostic utility of WES. There is no mention of the gene KIF20B, no data on KIF20B variants, and no mechanistic discussion that would link GRIA2 findings to KIF20B. Therefore the paper provides no direct or mechanistic evidence for or against the claim that KIF20B is associated with a Mendelian disease. Note that absence of KIF20B in this paper does not imply KIF20B has no Mendelian associations in other literature; it only means this particular paper is irrelevant to that claim.


[Read Paper](https://www.semanticscholar.org/paper/97a575a51e3abe25d15ff9c46defdb2d51861b21)


### Autosomal Dominant Intellectual Development Disorder-6 (MRD6) Without Seizures Linked to a De Novo Mutation in the grin2b Gene Revealed by Exome Sequencing: A Case Report of a Moroccan Child

**Why Not Relevant**: The paper content exclusively describes a case of MRD6 caused by a GRIN2B (grin2b) nonsense variant and related diagnostic, segregation, population-database, and ACMG-pathogenicity observations. There is no mention of the gene KIF20B, no variants reported in KIF20B, no functional/mechanistic data involving KIF20B, and no discussion that could be interpreted as implicating KIF20B in Mendelian disease. Therefore this paper provides no direct or mechanistic evidence for or against the claim that KIF20B is likely associated with at least one Mendelian disease.

Limitations and further notes: absence of KIF20B discussion in this GRIN2B-focused case report does not constitute evidence that KIF20B is not associated with disease (absence of evidence is not evidence of absence). To evaluate the claim about KIF20B one would need papers that (a) report rare, segregating KIF20B variants in affected families or de novo KIF20B variants in cases with consistent phenotypes, (b) provide functional/biochemical evidence that perturbation of KIF20B alters relevant biological processes, and (c) show statistical enrichment of rare damaging KIF20B variants in cases versus controls or replicate findings across cohorts. This paper contains none of those elements.


[Read Paper](https://www.semanticscholar.org/paper/67d756e0cc302e5d46fdda9df34403f1f2de9a5a)


### c.1103T>C (p.Ile368Th) de novo Variant in Synaptotagmin 1 (SYT1) Gene is Pathogenic, Leading to an Ultra-Rare Neurodevelopmental Disorder: The Baker-Gordon Syndrome

**Why Not Relevant**: The paper exclusively discusses Baker-Gordon Syndrome (BAGOS) and genetic, mechanistic, and clinical findings related to the synaptotagmin 1 (SYT1) gene. There is no mention of KIF20B anywhere in the provided content (abstract and case-report summary). Therefore the paper provides no direct genetic association evidence, no mechanistic evidence, and no contextual information that would support or refute the claim that KIF20B is likely associated with at least one Mendelian disease. In short, the manuscript is focused on a different gene (SYT1) and a distinct Mendelian disorder, so it is not relevant to assessing associations for KIF20B.


[Read Paper](https://www.semanticscholar.org/paper/ee225e1b916fbf3fda1d1b5451fe2038e281b8c8)


### Tuberous sclerosis complex associated lymphangioleiomyomatosis caused by de novo mutation of TSC2 gene in Vietnam: A case report

**Why Not Relevant**: This paper focuses exclusively on a clinical case of pulmonary lymphangioleiomyomatosis associated with a de novo frameshift mutation in TSC2; there is no mention of KIF20B, no reported variants in KIF20B, and no mechanistic discussion that would implicate KIF20B in this disorder. Therefore the paper provides no direct or mechanistic evidence supporting the claim that KIF20B is likely associated with at least one Mendelian disease. Note that absence of evidence in this single paper is not evidence that KIF20B is unrelated to Mendelian disease broadly — it only means this publication does not address KIF20B. Additional studies (case series, cohort exome/genome analyses, functional assays implicating KIF20B) would be required to evaluate the claim.


[Read Paper](https://www.semanticscholar.org/paper/6b7d5d519fb771d9bd7eb969a3b3847dcfdd8dad)


### Leveraging Clinical, Functional, Molecular and Population Genetic Data Reveals Genotype Phenotype Association and Health Disparity in a Monogenic Disorder, CTX

**Why Not Relevant**: The paper exclusively studies Cerebrotendinous Xanthomatosis (CTX) and pathogenic variants in the gene CYP27A1. The provided text contains no mention of KIF20B, no data, analyses, or mechanistic discussion relating to KIF20B, and no general evidence that could reasonably be extrapolated to support or refute an association between KIF20B and any Mendelian disease. Therefore this paper provides no relevant direct or mechanistic evidence regarding the claim that KIF20B is likely associated with at least one Mendelian disease.


[Read Paper](https://www.semanticscholar.org/paper/ecd9a6ae7be2a66a942f75a505fc4b04dff8e862)


### A brief clinical genetics review: stepwise diagnostic processes of a monogenic disorder—hypertriglyceridemia

**Why Not Relevant**: The provided paper text is a general review about advances in genetics, education of physicians, GWAS, and polygenic risk scores, and does not mention the gene KIF20B, any specific genes, nor any Mendelian diseases. Example sentences from the text that illustrate its general scope include: "Thus, there is an urgent need to educate physicians and healthcare providers with basic knowledge and skills in genetics." and "In addition, some important considerations in delineating and incorporating clinical genetic testing in the diagnosis and management of a monogenic disorder are illustrated in a stepwise fashion." These sentences show the paper discusses broad concepts (education, clinical genetic testing, PRS) rather than presenting gene-specific association data, family segregation, case reports, variant pathogenicity evidence, or functional/mechanistic experiments relevant to KIF20B. Therefore it provides no direct evidence (e.g., documented pathogenic variants in KIF20B segregating with disease, case series, statistical association with Mendelian phenotypes) nor mechanistic evidence (e.g., functional assays, animal models demonstrating disease-causing effects of KIF20B perturbation) to support or refute the claim that "KIF20B is likely associated with at least one Mendelian disease."

What would be required in a paper to be relevant to the claim: specific data such as (a) clinical reports of individuals/families with rare, likely pathogenic KIF20B variants showing segregation with a consistent phenotype; (b) de novo KIF20B variants in affected probands with appropriate statistical enrichment; (c) functional studies showing that variants impair KIF20B function in a way that plausibly causes disease; or (d) population genetics metrics and case-control burden tests supporting a role in a monogenic disorder. The provided text contains none of these elements, so its relevance to the claim is effectively zero.


[Read Paper](https://www.semanticscholar.org/paper/50284e86441df1a72744d288d3696df5aff5a6bd)


## Search Queries Used

- KIF20B de novo mutation exome sequencing patient case report neurodevelopmental disorder

- KIF20B Mendelian disease monogenic gene association human genetic disease OMIM review

- KIF20B knockout mouse phenotype loss of function cytokinesis midbody abscission neuronal development neuronal migration

- KIF20B loss of function intolerance gnomAD pLI constraint ClinVar pathogenic variants rare variant burden study


## Bibliometric Configuration

- Use Bibliometrics: True

- Author Impact Weight: 0.4

- Citation Impact Weight: 0.4

- Venue Impact Weight: 0.2
