## Supplementary material for "MENDELSEEK: An algorithm that predicts Mendelian Genes and elucidates what makes them special": KIF21B.txt

# Claim: The gene KIF21B is likely associated with at least one Mendelian disease.

**Status**: processed

**Overall Rating**: -1

**Explanation**:

No relevant papers were found that support or refute this claim.


## Relevant Papers


## Other Reviewed Papers


### A systematic review of Mendelian randomization studies on multiple sclerosis

**Why Not Relevant**: The provided paper content is a systematic review of Mendelian randomization studies assessing causal effects of genetic and non-genetic exposures on multiple sclerosis risk, emphasizing modifiable exposures (for example, low vitamin D and high BMI). It does not mention the gene KIF21B, nor does it discuss Mendelian (monogenic) disease associations or single-gene pathogenic variants. Furthermore, the MR approach and GWAS summary-level data emphasized in this review are geared toward assessing causal relationships between exposures and complex traits at the population level, typically using common variant instruments; this framework is not suited to demonstrating that a specific gene is causally linked to a Mendelian disease caused by rare, highly penetrant variants. Therefore, based on the paper content provided, there is no direct evidence supporting or refuting the claim that KIF21B is likely associated with at least one Mendelian disease, and no mechanistic discussion in this paper pertains to KIF21B or to monogenic disease mechanisms. Any assessment of KIF21B and Mendelian disease would require different types of evidence (e.g., case reports, segregation studies, functional assays of rare variants, ClinVar/OMIM entries), which are not present in this review.


[Read Paper](https://www.semanticscholar.org/paper/866887030eaa2c14b4cd19f5ab1bed453223e101)


### Precision Medicine in Internal Medicine

**Why Not Relevant**: The paper is a general review about genetic testing strategies, exome/genome sequencing, variant interpretation, and the state of gene discovery as of early 2019. It does not mention the gene KIF21B, present any case reports, genetic association data, pathogenic variants, segregation analyses, functional/mechanistic studies, or other gene-specific evidence linking KIF21B to any Mendelian phenotype. Therefore it provides no direct evidence for or against the claim that KIF21B is associated with at least one Mendelian disease. The extracted sentences only offer general context about how gene-disease associations are discovered and the large number of genes not yet linked to Mendelian disorders, which is indirect and nonspecific to KIF21B.


[Read Paper](https://www.semanticscholar.org/paper/c7199e8a5226f8d9d55777da0e15e91797bb459a)


### Family Relations in the Genomic Era: Communicating about Intergenerational Transmission of Risk for Disability

**Why Not Relevant**: The provided paper content contains general background on genetic causes of developmental disabilities, Mendelian inheritance, OMIM counts, genetic susceptibility, and epigenetics, but it does not mention KIF21B, report any observational/genetic/clinical/segregation data, functional experiments, variant-level evidence, case reports, or statistics that link KIF21B to any Mendelian disorder. Therefore the content provides no direct evidence for or against the claim and only offers general mechanistic context that could apply to any gene. Because the claim concerns a specific gene (KIF21B), and the paper contains no KIF21B-specific information, the relevance to the claim is very low.


[Read Paper](https://www.semanticscholar.org/paper/a6ab963ce2ad3023249041b17f3dc6ff2b939705)


### Tubulin glutamylation: a skeleton key for neurodegenerative diseases

**Why Not Relevant**: The provided paper text does not mention KIF21B, nor does it present genetic, clinical, or pedigree data linking any kinesin gene (including KIF21B) to a Mendelian disorder. The content focuses on tubulin post-translational modifications, their enzymes (TTLLs and CCPs), and the regulatory protein CSAP, and discusses general effects of PTMs on microtubule-associated proteins and molecular motors. While this yields mechanistic context (that tubulin PTMs can modulate kinesin function), such general mechanistic statements cannot be used to infer that KIF21B is "likely associated with at least one Mendelian disease." There is no direct experimental, clinical, or population-genetic evidence in the excerpt to support or refute the claim. Any inference that KIF21B is disease-associated would therefore be speculative and unsupported by the paper's content. Additional data (e.g., reports of pathogenic variants in KIF21B segregating with disease, functional studies of KIF21B mutations causing cellular phenotypes, or clinical case series) would be required to evaluate the claim.


[Read Paper](https://www.semanticscholar.org/paper/3560acd61817be3066152e1e88ac2ffa0ff85303)


### Finding New Molecular Targets for Parkinson's Disease through Gene Hunting in Queensland Families with Inherited Parkinsonism

**Why Not Relevant**: The provided paper content does not mention KIF21B anywhere and therefore offers no direct evidence supporting or refuting the claim that KIF21B is likely associated with at least one Mendelian disease. The paper focuses on Parkinson's disease families from the Queensland Parkinson’s Project and reports known PD genes (VPS35, LRRK2, SNCA, PARK2) and several candidate novel genes (e.g., KCNJ15, SIPA1L1, FAM134B). The authors used whole exome sequencing, copy-number variation analysis, and other supportive data to prioritize candidates, which means that coding high-impact variants in KIF21B within these families could have been detected if present and prioritized; however, the absence of any KIF21B finding in the reported results does not constitute evidence against KIF21B being Mendelian elsewhere. Limitations relevant to interpreting absence: (1) the study is restricted to PD-affected families, so it would not detect Mendelian associations of KIF21B with non-PD diseases; (2) exome sequencing and the study's prioritization/filtering criteria can miss non-coding pathogenic variants, certain structural variants, low-coverage regions, or variants filtered out for frequency/annotation reasons; (3) negative reporting (not listing a gene) is not the same as a systematic negative result for that gene. For these reasons the paper is not relevant to the claim about KIF21B, and it provides neither direct nor mechanistic evidence that KIF21B is associated with a Mendelian disease.


[Read Paper](https://www.semanticscholar.org/paper/d73a572362bfaa0214e74416736dfd0ea17d8e33)


## Search Queries Used

- KIF21B Kinesin family member 21B mutation variant Mendelian monogenic inherited disease case report familial de novo

- KIF21B function kinesin microtubule motor axonal transport neuronal development synaptic function mouse model zebrafish model knockdown knockout phenotype

- KIF21B exome sequencing rare variant burden whole exome cohort intellectual disability epilepsy developmental delay gnomAD constraint pLI loss of function intolerance

- KIF21B genome wide association study GWAS multiple sclerosis autoimmune risk association review systematic review meta analysis Mendelian versus complex disease


## Bibliometric Configuration

- Use Bibliometrics: True

- Author Impact Weight: 0.4

- Citation Impact Weight: 0.4

- Venue Impact Weight: 0.2
