## Supplementary material for "MENDELSEEK: An algorithm that predicts Mendelian Genes and elucidates what makes them special": KIFAP3.txt

# Claim: The gene KIFAP3 is likely associated with at least one Mendelian disease.

**Status**: processed

**Overall Rating**: -1

**Explanation**:

No relevant papers were found that support or refute this claim.


## Relevant Papers


## Other Reviewed Papers


### A pathogenic variant in RAB32 causes autosomal dominant Parkinson’s disease and activates LRRK2 kinase

**Why Not Relevant**: The paper exclusively studies RAB GTPases, and its principal genetic and functional findings concern a RAB32 variant (c.213C>G, Ser71Arg) and its role in autosomal dominant parkinsonism, including co-segregation in families, case-control association, and functional activation of LRRK2. Nowhere in the provided content is KIFAP3 mentioned, nor are there data, analyses, or mechanistic links reported that involve KIFAP3. Therefore this paper provides no direct genetic evidence that KIFAP3 is associated with any Mendelian disease, nor does it provide mechanistic evidence that would plausibly implicate KIFAP3 by analogy or pathway connection. Absence of evidence in this paper does not disprove any possible association of KIFAP3 discovered elsewhere, but based on the provided content there is no support for the claim that KIFAP3 is likely associated with at least one Mendelian disease.


[Read Paper](https://www.semanticscholar.org/paper/a36d28e828ede7c87c650af5f5117982b9749e7b)


### Whole exome sequencing revealed a novel homozygous variant in the DGKE catalytic domain: a case report of familial hemolytic uremic syndrome

**Why Not Relevant**: The provided paper content does not provide any evidence relevant to the claim that KIFAP3 is likely associated with at least one Mendelian disease. The only sentence in the content is: "It is suggested that some variations in the catalytic domain of DGKE like p.Asn314Lys which can cause alterations in secondary and 3-D structure of protein, might lead to aHUS." This sentence (verbatim) refers to DGKE and to a possible link between DGKE catalytic-domain variants (example p.Asn314Lys) and atypical hemolytic uremic syndrome (aHUS). It does not mention KIFAP3, any variant of KIFAP3, or any phenotype or Mendelian disease associated with KIFAP3. Therefore there is no direct evidence (no reported KIFAP3 variants, no genotype–phenotype correlations, no segregation or case reports) in this content that supports or refutes the claim about KIFAP3.

Mechanistic relevance: the sentence presents a speculative mechanistic idea about DGKE (that catalytic-domain changes may alter secondary/tertiary structure and thereby contribute to disease). That is mechanistic content about DGKE only; there is no mechanistic link made between DGKE and KIFAP3, nor any shared pathway or functional evidence in the provided text that could plausibly be extrapolated to KIFAP3. Without explicit mention or data on KIFAP3, this DGKE-focused mechanistic statement cannot be taken as mechanistic evidence for KIFAP3 involvement in Mendelian disease.

Limitations and caveats in the provided content: (1) the text is a single speculative sentence using the phrase "It is suggested" and "might lead to", indicating hypothesis rather than demonstrated result; (2) no experimental methods, data, statistical evidence, case descriptions, or segregation analyses are provided; (3) the statement concerns a different gene (DGKE), so it does not address the KIFAP3 claim; (4) because of the absence of details, even the DGKE assertion would be weak evidence on its own.

Conclusion: relevance is effectively zero because the content does not mention KIFAP3 or provide data/mechanistic links applicable to KIFAP3. No excerpts supporting or refuting the KIFAP3 claim are available in the provided text.


[Read Paper](https://www.semanticscholar.org/paper/2691c9dc3e9f16f506828ae781284158953cd6a0)


### Contribution of rare transmitted and de novo variants among 2 , 871 1 congenital heart disease probands

**Why Not Relevant**: The provided paper excerpt does not mention KIFAP3 anywhere in the visible text. The content describes cohort size, overall findings (e.g., specific genes GDF1, MYH6, FLT4), the number of genes implicated at statistical thresholds, curated gene sets from OMIM and mouse screens, variant classes analyzed, and counts of damaging recessive genotypes. None of these quoted sentences or paragraphs provide direct evidence that KIFAP3 is associated with a Mendelian disease. It is possible that KIFAP3 could appear in supplementary materials (Supplementary Excel S2 or S3) or in other parts of the full paper not provided here; however, based solely on the supplied text, there is no mention, result, or mechanistic discussion regarding KIFAP3. Therefore the relevance to the claim that 'KIFAP3 is likely associated with at least one Mendelian disease' is negligible in this excerpt. To evaluate the claim properly, one would need to (1) search the paper's supplementary gene lists (Supplementary Excel S2/S3) for KIFAP3, (2) inspect other sections of the full text not included here for any mention of KIFAP3, or (3) consult other publications or databases (e.g., OMIM, ClinVar, HGMD, gnomAD) for evidence linking KIFAP3 to Mendelian phenotypes.


[Read Paper](https://www.semanticscholar.org/paper/e5830d9e95469f22518560007746202e62db26a0)


### Exome Sequencing of Consanguineous Pashtun Families With Familial Epilepsy Reveals Causative and Candidate Variants in TSEN54, MOCS2, and OPHN1

**Why Not Relevant**: The paper does not mention KIFAP3 anywhere in the abstract content provided. All reported gene-disease associations refer to other genes (TSEN54, MOCS2, OPHN1) identified in this small exome-sequencing study of four families with presumed Mendelian epilepsy. Therefore the paper provides no direct genetic association data, mechanistic evidence, or discussion concerning KIFAP3 and its role in Mendelian disease. The study's small sample size and narrow phenotype focus (epilepsy in a specific population) further limit any implicit negative inference about KIFAP3: lack of mention in this study is not evidence that KIFAP3 is not disease-associated. Because there is neither direct nor mechanistic evidence relating KIFAP3 to a Mendelian disease in this paper, the relevance to the claim is effectively zero.


[Read Paper](https://www.semanticscholar.org/paper/1a6092d22e5cb11c253d76dafbaa8214cf0e51ec)


## Search Queries Used

- KIFAP3 variant Mendelian disease case report exome sequencing familial segregation pathogenic variant

- KIFAP3 ClinVar OMIM gnomAD loss of function constraint clinical annotation pathogenic likely pathogenic KIFAP3 gene

- Kinesin associated protein 3 KAP3 KIFAP3 loss of function knockout mouse zebrafish neuronal axonal transport neurodegeneration mechanism

- KIFAP3 rare variant burden exome sequencing cohort case control inherited disease penetrance expressivity phenotypic heterogeneity


## Bibliometric Configuration

- Use Bibliometrics: True

- Author Impact Weight: 0.4

- Citation Impact Weight: 0.4

- Venue Impact Weight: 0.2
