## Supplementary material for "MENDELSEEK: An algorithm that predicts Mendelian Genes and elucidates what makes them special": KRT24.txt

# Claim: The gene KRT24 is likely associated with at least one Mendelian disease.

**Status**: processed

**Overall Rating**: -1

**Explanation**:

No relevant papers were found that support or refute this claim.


## Relevant Papers


## Other Reviewed Papers


### Whole-exome sequencing identifies a GREB1L variant in a three-generation family with Müllerian and renal agenesis: a novel candidate gene in Mayer-Rokitansky-Küster-Hauser (MRKH) syndrome. A case report.

**Why Not Relevant**: This paper does not provide evidence for or against the claim that KRT24 is associated with a Mendelian disease. The study focuses on Mayer–Rokitansky–Küster–Hauser (MRKH) syndrome and reports identification of a novel pathogenic c.705G>T missense variant in GREB1L (CADD = 33): “leading to the identification of a novel pathogenic (CADD = 33) c.705G>T missense variant in GREB1L, a gene recently identified as a novel cause of RA.” The text repeatedly centers on GREB1L and its role in renal and female genital tract development and suggests GREB1L as a candidate gene for MRKH syndrome: “we suggest GREB1L as a novel and promising candidate gene in the aetiology of MRKH syndrome.” Nowhere in the provided content is KRT24 mentioned, analyzed, or discussed. Therefore there is no direct evidence (genetic, segregation, phenotypic, or statistical) relating KRT24 to any Mendelian phenotype in this paper, nor is there mechanistic data concerning KRT24. The absence of any mention or analysis of KRT24 means this paper cannot be used to support the claim; it likewise does not actively refute the claim (absence of evidence in this single paper is not evidence of absence). To evaluate the claim about KRT24 would require consulting studies that specifically assay KRT24 (e.g., ClinVar, OMIM, HGMD, published WES/WGS studies, or functional studies), which are not present here.


[Read Paper](https://www.semanticscholar.org/paper/ab2ca8c9473a42256e3a154ff0eacac1a32cbe2b)


### Familial Whole Exome Sequencing Study of 30 Families With Early-Onset High Myopia

**Why Not Relevant**: This paper does not provide relevant evidence for or against the claim that KRT24 is likely associated with at least one Mendelian disease. The provided content (Purpose, Methods, Results, Conclusions) describes whole-exome sequencing in 30 families with early-onset high myopia (eoHM) and lists genes and variants identified in that cohort (examples given include COL4A5, NYX, CACNA1F, CCDC111, SLC39A5, P4HA2, CPSF1, GRM6). Nowhere in the supplied text is KRT24 mentioned, nor are any variants in KRT24 reported. Therefore there is no direct evidence (no reported KRT24 variants segregating with a Mendelian phenotype) and no mechanistic evidence (no discussion of KRT24 function, expression, or pathways relevant to a Mendelian disease) in this paper.

Limitations and additional notes relevant to interpreting this absence of evidence:
- Absence of KRT24 in the reported results does not prove that KRT24 is not associated with a Mendelian disease; it only means this study did not identify or report such an association in its eoHM cohort.
- The study used whole-exome sequencing, which in principle could detect coding variants in KRT24 if present and covered, but the provided text does not state coverage metrics or explicitly confirm that KRT24 was adequately captured in all samples; thus, a negative observation in this paper (non-reporting of KRT24) cannot be taken as definitive disproof.
- The study's focus is on eoHM and inherited retinal/ocular disease genes; if KRT24 is associated with a Mendelian disease unrelated to the ocular phenotypes they studied, this cohort would be an insensitive context to detect that association.

In summary: relevance is effectively zero because KRT24 is not mentioned and no supporting or refuting data or mechanistic discussion concerning KRT24 appears in the provided paper content.


[Read Paper](https://www.semanticscholar.org/paper/8a51e1e2b13da3e9075f4d54e506e745280ab56b)


### CNGB3 Missense Variant Causes Recessive Achromatopsia in Original Braunvieh Cattle

**Why Not Relevant**: The paper reports identification and mechanistic characterization of a CNGB3 missense variant causing recessive achromatopsia in Original Braunvieh cattle. The provided text repeatedly and exclusively implicates CNGB3 (including genotype, variant p.Asp251Asn, localization changes, and functional phenotype) and contains no mention of KRT24 or keratin genes. Therefore it provides no direct genetic, clinical, or mechanistic evidence for or against an association between KRT24 and any Mendelian disease. Because the paper's entire focus is a different gene (CNGB3) and phenotype, its relevance to the specific claim about KRT24 is effectively nil.


[Read Paper](https://www.semanticscholar.org/paper/0ce67fef63d36e0a2ccf00bc7cd5ea3970c7022b)


### Report of a novel missense TDP1 variant in a Pakistani family affected with an extremely rare disorder congenital spinocerebellar ataxia with axonal neuropathy type 1 (SCAN1)

**Why Not Relevant**: The provided paper content reports on a single patient from a consanguineous Pakistani family with autosomal recessive spinocerebellar ataxia with axonal neuropathy type 1 and states that the evidence supporting a pathogenic role for TDP1 in cerebellar ataxias with peripheral neuropathy is increased. The exact text provided is: "A female patient born to a consanguineous Pakistani family segregating autosomal recessive spinocerebellar ataxia with axonal neuropathy type 1 is reported and the body of evidence that supports the pathogenic role of TDP1 in cerebellar ataxias with peripheral neuropathy is increased." This content discusses TDP1 and a specific neurogenetic phenotype; it contains no mention of KRT24, no data on variants in KRT24, no segregation analysis involving KRT24, no functional/mechanistic data for KRT24, and no clinical association between KRT24 and any Mendelian disease. Therefore the paper provides neither direct evidence (e.g., pathogenic KRT24 variants segregating with disease) nor mechanistic evidence (e.g., disrupted KRT24 protein function causing a Mendelian phenotype) relevant to the claim that "KRT24 is likely associated with at least one Mendelian disease." Limitations and caveats: it is possible (but not indicated in the provided text) that other parts of the full paper or supplementary data mention additional genes; however, based solely on the supplied content, there is no basis to support or refute an association between KRT24 and Mendelian disease. To evaluate the claim about KRT24 one would need a paper that explicitly reports KRT24 variants with segregation, case-control enrichment, functional assays, or animal-model phenotypes linking KRT24 disruption to a Mendelian disorder.


[Read Paper](https://www.semanticscholar.org/paper/f89b8882ce3bcf1efc93731fa7a65ea676dccbfc)


### The simplest explanation does not have to be preferred: co-occurrence of pathogenic variants in cancer-predisposing genes

**Why Not Relevant**: The provided paper content contains a single run-on sentence that discusses reviewing literature and UK 10,000 Genomes Project data for cancer phenotypes associated with MINAS and whether multiple cancer-susceptibility gene (CSG) effects are additive or synergistic. It does not mention the gene KRT24 at all, nor does it present any variant-level data, case reports, segregation analyses, functional/mechanistic studies, or statistical associations that could connect KRT24 to a Mendelian disease. Because none of the required types of evidence (direct observational genetic evidence, mechanistic molecular evidence, or even a mention of KRT24) are present, the content cannot support or refute the claim. Additional, gene-specific data or sections of the paper that explicitly discuss KRT24 would be required to assess the claim.


[Read Paper](https://www.semanticscholar.org/paper/a1372a664bd2c33a0bda638975a8b26d112457fc)


### Whole exome sequencing identified a novel frameshift variant in the BHLHA9 in an Iranian family with mesoaxial synostotic syndactyly

**Why Not Relevant**: The paper exclusively studies Mesoaxial synostotic syndactyly with phalangeal reduction (MSSD) and reports a novel homozygous frameshift variant in the BHLHA9 gene as causal in the presented family. There is no mention of KRT24 anywhere in the provided content, no sequence, expression, functional, linkage, or segregation data for KRT24, and no mechanistic discussion connecting KRT24 to the phenotype. Therefore the paper provides no direct evidence that KRT24 is associated with any Mendelian disease. The paper's attribution of this particular Mendelian disorder to BHLHA9 is indirect negative evidence only for KRT24’s role in this specific phenotype but does not address whether KRT24 may be associated with other Mendelian conditions. Additional limitations: the study is a single-family report focused on one gene/phenotype, so absence of evidence for KRT24 here cannot be taken as evidence of absence across all Mendelian diseases.


[Read Paper](https://www.semanticscholar.org/paper/f1e9bde70ac150678e1315f6a78f2e105d282b76)


### A novel missense mutation in the CRYBA2 caused autosomal dominant presenile cataract in a Chinese family

**Why Not Relevant**: The paper content provided (only the title) concerns a mutation in CRYBA2 associated with presenile cataract. It contains no mention of KRT24, keratin family genes, or any data (genetic, clinical, or mechanistic) involving KRT24. Therefore it provides no direct or mechanistic evidence for or against the claim that KRT24 is associated with a Mendelian disease. With only the title available, there is no basis to infer an indirect connection (e.g., shared pathways or genetic linkage) to KRT24. Because the content neither names nor discusses KRT24, relevance to the KRT24–Mendelian disease claim is effectively zero.


[Read Paper](https://www.semanticscholar.org/paper/29794dd1f9055a399851fa2e6dfa586ebf74dc13)


### Unraveling genetic contributions to neurodevelopmental disorders and epilepsy in a pediatric cohort from a Mexican state

**Why Not Relevant**: The provided paper content does not mention the gene KRT24 anywhere, nor does it list the specific genes, variant calls, or gene-level findings implicated in the diagnosed Mendelian diseases. Although the study reports identification of Mendelian diagnoses and novel mutations using next-generation sequencing in a small pediatric cohort, the absence of any explicit reference to KRT24 means there is no direct evidence (e.g., pathogenic KRT24 variants, segregation, functional or mechanistic data) supporting or refuting the claim that KRT24 is likely associated with at least one Mendelian disease. Additionally, the small sample size (24 patients) and selective parental testing constrain the ability to generalize or detect rare gene-disease associations. Therefore the paper content as provided is not relevant to establishing an association between KRT24 and a Mendelian disease.


[Read Paper](https://www.semanticscholar.org/paper/9162aad99d110fbef19855b4be76684d86ddd148)


### Development of a mouse model for Hutchinson-Gilford progeria syndrome reveal defects in adult stem cell maintenance

**Why Not Relevant**: This paper focuses on Hutchinson-Gilford progeria syndrome caused by mutations in LMNA and on mouse models expressing mutant lamin A/progerin. The text contains no mention of the gene KRT24, no data on KRT24 variants, and no genetic association or clinical descriptions linking KRT24 to a Mendelian disease. Although the study targeted expression to keratin 5-expressing tissues (using a keratin-5 promoter) and examined skin phenotypes, that experimental choice is a tissue-targeting strategy and does not implicate any keratin gene (including KRT24) as causal. Mechanistic descriptions concern A-type lamins and the nuclear lamina rather than keratin-24 or the keratin gene family more broadly. Therefore the paper provides no direct or mechanistic evidence for the claim that KRT24 is likely associated with at least one Mendelian disease.


[Read Paper](https://www.semanticscholar.org/paper/a242c87f7c8b391e5fdee78c6106d095f2ae3c43)


## Search Queries Used

- KRT24 keratin 24 mutation pathogenic variant Mendelian disease monogenic inherited disorder

- KRT24 ClinVar OMIM gnomAD pLI loss of function intolerance pathogenic variant annotation

- keratin 24 KRT24 expression epidermis skin hair cornea intermediate filament function knockout mouse phenotype skin disease

- whole exome sequencing family segregation rare variant KRT24 novel Mendelian gene discovery autosomal dominant autosomal recessive


## Bibliometric Configuration

- Use Bibliometrics: True

- Author Impact Weight: 0.4

- Citation Impact Weight: 0.4

- Venue Impact Weight: 0.2
