## Supplementary material for "MENDELSEEK: An algorithm that predicts Mendelian Genes and elucidates what makes them special": LAP3.txt

# Claim: The gene LAP3 is likely associated with at least one Mendelian disease.

**Status**: processed

**Overall Rating**: -1

**Explanation**:

No relevant papers were found that support or refute this claim.


## Relevant Papers


## Other Reviewed Papers


### Homozygous NLRP1 gain-of-function mutation in siblings with a syndromic form of recurrent respiratory papillomatosis

**Why Not Relevant**: The provided paper content contains no mention of the gene symbol 'LAP3' (nor any obvious synonyms or descriptions of that gene), and all genetic and mechanistic discussion centers on NLRP1 and human papillomavirus–related juvenile-onset recurrent respiratory papillomatosis (JRRP). Key statements describe a private homozygous mutation in NLRP1, its gain-of-function effect on inflammasome activation, elevated IL-1β secretion, and that six NLRP1 GOF mutations underlie three allelic Mendelian diseases. Because the text exclusively addresses NLRP1 (not LAP3), there is neither direct evidence (genetic association, case reports, segregation, variant pathogenicity) nor mechanistic evidence (biochemical pathways, functional assays linking LAP3 to disease) in the provided content to support or refute the claim that LAP3 is likely associated with at least one Mendelian disease. Limitations/caveats: this assessment is limited to the supplied excerpt only; the full paper (main text, supplemental data, or other sections not provided here) could potentially mention LAP3, but based on the content given, no relevant information about LAP3 is present.


[Read Paper](https://www.semanticscholar.org/paper/44227dfbf79fd311c75515f48f4ca895e29eb370)


### Heterozygous Mutations in DNA Repair Genes and Hereditary Breast Cancer: A Question of Power

**Why Not Relevant**: The provided paper text does not mention the gene LAP3 or present any data, analysis, or mechanistic discussion involving LAP3. The manuscript excerpt focuses on exome sequencing in hereditary breast cancer families, with emphasis on genes involved in homologous recombination and DNA repair (notably BRCA1, BRCA2, FANCC, and BLM). Specific findings and counts reported concern FANCC and BLM (e.g., identification of heterozygous mutations in these genes in breast cancer families and their frequency in various cohorts), and the text discusses general challenges of detecting rare Mendelian risk alleles by sequencing. Because LAP3 is neither named nor discussed in the text, there is no direct evidence (no variant counts, segregation data, case-control comparisons, or reported pathogenic mutations) supporting or refuting an association between LAP3 and any Mendelian disease. Likewise, there is no mechanistic evidence in this excerpt linking LAP3 to disease pathways (the mechanistic focus here is on homologous recombination and Fanconi anemia / Bloom syndrome molecular overlap). Important caveats: absence of mention in this single paper does not prove that LAP3 is not associated with a Mendelian disease; it merely means this paper provides no relevant evidence. To evaluate the claim about LAP3, one would need to consult other publications, variant databases, or experimentally focused mechanistic studies that specifically include LAP3.


[Read Paper](https://www.semanticscholar.org/paper/f950f5349779a29089fa04d12059c03231f6edc4)


### PADI3, hair disorders and genomic investigation

**Why Not Relevant**: The provided paper content focuses on the genetics and molecular mechanisms of uncombable hair syndrome (UHS) and central centrifugal cicatricial alopecia (CCCA), specifically implicating PADI3, TGM3 and TCHH and discussing whole exome sequencing as a method. Nowhere in the supplied text is the gene symbol "LAP3" mentioned, nor are any data, variants, functional studies, or mechanistic pathways involving LAP3 presented. Therefore the paper provides no direct genetic evidence (no reported LAP3 variants segregating with disease) and no mechanistic evidence (no biochemical or cell biological data implicating LAP3) to support or refute the claim that "LAP3 is likely associated with at least one Mendelian disease." While the paper does discuss methods (WES) that in principle could detect exonic LAP3 variants, the absence of any LAP3-specific results or discussion means there is no evidentiary basis in this paper to evaluate the claim. Consequently relevance to the claim is effectively zero.


[Read Paper](https://www.semanticscholar.org/paper/049d93b6c8f29f096e562e667131b4835d1700e3)


## Search Queries Used

- LAP3 leucine aminopeptidase 3 mutation Mendelian disease case report family segregation exome sequencing human

- LAP3 pathogenic variant ClinVar OMIM gnomAD loss of function constraint LOEUF pLI

- LAP3 knockout mouse phenotype zebrafish functional study aminopeptidase activity loss of function developmental neurological phenotype

- rare LAP3 variants exome sequencing trios gene discovery Mendelian disease autosomal recessive biallelic rare variant burden paralog homology misannotation mapping error pseudogene


## Bibliometric Configuration

- Use Bibliometrics: True

- Author Impact Weight: 0.4

- Citation Impact Weight: 0.4

- Venue Impact Weight: 0.2
