## Supplementary material for "MENDELSEEK: An algorithm that predicts Mendelian Genes and elucidates what makes them special": LOXL2.txt

# Claim: The gene LOXL2 is likely associated with at least one Mendelian disease.

**Status**: processed

**Overall Rating**: -1

**Explanation**:

No relevant papers were found that support or refute this claim.


## Relevant Papers


## Other Reviewed Papers


### Prenatal Diagnosis of PPP2R1A-Related Neurodevelopmental Disorders Using Whole Exome Sequencing: Clinical Report and Review of Literature

**Why Not Relevant**: The paper exclusively studies PPP2R1A-related neurodevelopmental disorder and reports de novo pathogenic or likely pathogenic variants in PPP2R1A found prenatally in two fetuses. There is no mention of LOXL2 anywhere in the provided content, no data (genetic, clinical, or mechanistic) linking LOXL2 to disease, and no cross-references that could plausibly connect LOXL2 to PPP2R1A-related pathology. Because the paper's entire focus, results, and conclusions pertain to a different gene (PPP2R1A), it provides no direct evidence supporting or refuting the claim that LOXL2 is associated with at least one Mendelian disease. Additionally, there are no mechanistic discussions (pathways, protein interactions, functional assays) that could be used to infer an association for LOXL2. For these reasons the paper is not relevant to the LOXL2 claim (relevance = 0).


[Read Paper](https://www.semanticscholar.org/paper/7595f38ac95f91f2d03163020f3687301e6f88d0)


### Diagnostic yield of patients with undiagnosed intellectual disability, global developmental delay and multiples congenital anomalies using karyotype, microarray analysis, whole exome sequencing from Central Brazil

**Why Not Relevant**: The paper excerpt does not mention LOXL2 by name, does not provide a list of genes analyzed or the gene-level results, and contains no mechanistic information about LOXL2. All available sentences describe study design, cohort size, diagnostic yields, and general conclusions about methods; none provide direct or indirect evidence that LOXL2 is associated with any Mendelian disease. Therefore the content cannot be used to support or refute the claim that LOXL2 is likely associated with at least one Mendelian disease.


[Read Paper](https://www.semanticscholar.org/paper/23767daf16f48d9a91172aa0fc624f07b869a9c8)


### Trio-Based Whole-Exome Sequencing Identifies a De novo EFNB1 Mutation as a Genetic Cause in Female Infant With Brain Anomaly and Developmental Delay

**Why Not Relevant**: The paper exclusively studies EFNB1 in relation to craniofrontonasal syndrome and a single-case presentation of schizencephaly with global developmental delay. There is no mention of LOXL2, no experimental data, variant calls, phenotypic associations, functional assays, or mechanistic discussion involving LOXL2. Because the gene of interest (LOXL2) is not addressed at all, the paper provides no direct genetic or mechanistic evidence for or against the claim that LOXL2 is associated with a Mendelian disease. Additionally, the study is a single-case EFNB1 report (limited sample size and scope), further limiting any indirect inference about other genes.


[Read Paper](https://www.semanticscholar.org/paper/939ec66d3caac7b206eab2e4073583eff41bfa48)


### Expanding phenotype of MED13-associated syndrome presenting novel de novo missense variant in a patient with multiple congenital anomalies

**Why Not Relevant**: The provided paper content discusses a de novo missense variant in MED13 and proposes expansion of the MED13-associated phenotype. It contains no mention of the gene LOXL2, no LOXL2 variants, no patient data implicating LOXL2, and no mechanistic data (protein interaction, pathway, expression, functional assays) linking LOXL2 to disease. Because the paper's subject is MED13 and no LOXL2-related information is present in the supplied excerpt, it offers no evidence for or against the claim that LOXL2 is likely associated with at least one Mendelian disease. If the full manuscript includes additional material not provided here (e.g., supplemental tables listing other candidate genes including LOXL2, or pathway analyses linking MED13 and LOXL2), that material would need to be reviewed; based on the content supplied, relevance is effectively zero.


[Read Paper](https://www.semanticscholar.org/paper/9b2cc0996fdf5fb478e07dbf06be71de50a057ee)


### De novo HDAC3 variants leading to epigenetic machinery dysfunction are associated with a neurodevelopmental disorder

**Why Not Relevant**: This paper focuses exclusively on heterozygous de novo variants in HDAC3 and their association with a novel neurodevelopmental syndrome. The provided content contains no mention of the gene LOXL2, no data on LOXL2 variants, and no analyses (genetic, functional, or clinical) that implicate LOXL2 in any Mendelian disease. The mechanistic findings described (reduced histone deacetylation activity for specific HDAC3 variants; disrupted interactions with CoREST and NCoR complexes; impaired nuclear localization) are specific to HDAC3 and its protein interactions and therefore do not provide direct or mechanistic evidence linking LOXL2 to a Mendelian condition. Because the paper neither examines nor references LOXL2, it cannot be used to support or refute the claim that LOXL2 is likely associated with at least one Mendelian disease.


[Read Paper](https://www.semanticscholar.org/paper/524b09f9e39f306b6871f816044577dccb53abd6)


## Search Queries Used

- LOXL2 lysyl oxidase like 2 pathogenic variant ClinVar OMIM case report Mendelian disease autosomal dominant autosomal recessive

- LOXL2 de novo mutation whole exome sequencing trio congenital anomaly developmental disorder Mendelian

- LOXL2 loss of function haploinsufficiency gain of function collagen crosslinking extracellular matrix connective tissue disorder vascular malformation Ehlers Danlos fibrosis

- LOXL2 knockout mouse zebrafish functional study human phenotype developmental defects ex vivo collagen crosslinking enzymatic activity rare variant burden analysis exome sequencing cohort


## Bibliometric Configuration

- Use Bibliometrics: True

- Author Impact Weight: 0.4

- Citation Impact Weight: 0.4

- Venue Impact Weight: 0.2
