## Supplementary material for "MENDELSEEK: An algorithm that predicts Mendelian Genes and elucidates what makes them special": MAP3K4.txt

# Claim: The gene ELF3 is likely associated with at least one Mendelian disease.

**Status**: processed

**Overall Rating**: -1

**Explanation**:

No relevant papers were found that support or refute this claim.


## Relevant Papers


## Other Reviewed Papers


### Expanding the phenotype of a recurrent de novo variant in PACS1 causing intellectual disability

**Why Not Relevant**: The provided paper content exclusively discusses PACS1 (and PACS2) and reports a recurrent de novo missense variant in PACS1 causing an autosomal-dominant intellectual disability syndrome (MRD17). There is no mention of the gene ELF3 anywhere in the text, no data (genetic, clinical, or functional) concerning ELF3, and no discussion that would allow inference about ELF3's involvement in Mendelian disease. Therefore the paper provides no direct or mechanistic evidence for or against the claim that ELF3 is likely associated with at least one Mendelian disease.


[Read Paper](https://www.semanticscholar.org/paper/e460d89d850693b4fb1e3df12702be0dfb8d7f5b)


### A single center experience with publicly funded clinical exome sequencing for neurodevelopmental disorders or multiple congenital anomalies

**Why Not Relevant**: The provided paper content consists of a single sentence: "The findings support the utility of clinical ES in a real-world setting, as part of a publicly funded genetic workup for individuals with GDD/ID and/or MCA." This sentence (1) does not mention the gene ELF3, (2) contains no variant-level data, genotype–phenotype correlations, segregation analyses, recurrence evidence, or case descriptions involving ELF3, and (3) provides no mechanistic information (e.g., gene function, pathogenic mechanism, animal or cellular models) that would link ELF3 to any Mendelian phenotype. While the sentence states that clinical exome sequencing (ES) is useful for investigating individuals with GDD/ID (global developmental delay/intellectual disability) and/or MCA (multiple congenital anomalies), that general statement does not constitute evidence that ELF3 is associated with a Mendelian disease. To support the claim about ELF3 one would expect verbatim sentences or paragraphs reporting one or more of the following: identification of (likely) pathogenic ELF3 variants in affected individuals, segregation of ELF3 variants with disease in families, statistical enrichment of ELF3 variants in cases versus controls, functional assays demonstrating deleterious effects of ELF3 variants, or discussion of a plausible mechanistic pathway linking ELF3 disruption to a phenotype. None of those elements are present in the provided content. Therefore the content is not relevant to the claim that "The gene ELF3 is likely associated with at least one Mendelian disease." If more of the paper (full text, figures, variant tables, or case reports) is available, those sections should be supplied for a focused assessment of any evidence implicating ELF3.


[Read Paper](https://www.semanticscholar.org/paper/8ec4c0c03c573ad4019a6ee63b9197179a32b173)


### Exome-wide Analysis of De Novo and Rare Genetic Variants in Patients With Brain Arteriovenous Malformation

**Why Not Relevant**: The paper's content focuses on germline genetic contributors to brain arteriovenous malformation (bAVM) and reports pathway enrichment for ANGPTL8 and a disease-associated gene SLC19A3 identified by exome-wide rare variant association. The text provided does not mention ELF3 at all. Therefore this paper provides neither direct evidence (e.g., genetic association, pathogenic variants, cosegregation) nor mechanistic evidence (e.g., functional data, pathway linkage) that ELF3 is associated with any Mendelian disease. Limitations/caveats: (1) Absence of ELF3 in this bAVM-focused exome study is not proof that ELF3 is not associated with any Mendelian disease elsewhere—the study's phenotype focus, sample composition, and analysis scope mean it would only detect genes relevant to bAVM in the cohorts analyzed. (2) Exome sequencing may miss noncoding variants or certain structural variants; if ELF3-related Mendelian disease were caused by such variants or by variants in populations not represented here, this study would not detect them. (3) The study reports other gene findings (SLC19A3) and pathways (ANGPTL8) pertinent to bAVM, but none provide indirect mechanistic links to ELF3 in the provided text. Given the complete lack of mention or data about ELF3 in the paper content supplied, the paper is not relevant to assessing the claim that "ELF3 is likely associated with at least one Mendelian disease."


[Read Paper](https://www.semanticscholar.org/paper/17051168a0ad2ef848d7692de1976646487e0c2b)


### Murine models of ARVC: what have we learned and where do we go? Insight for therapeutics.

**Why Not Relevant**: The provided paper content focuses on arrhythmogenic right ventricular cardiomyopathy (ARVC) and genes encoding desmosomal proteins (for example DSP, plakoglobin, plakophilin 2, desmocollin 2, desmoglein 2). The text contains mechanistic and genetic evidence linking desmosomal genes (notably DSP) to Mendelian cardiac disorders (ARVC, Carvajal syndrome), but ELF3 is not mentioned anywhere in the provided content. Therefore this paper provides no direct genetic association or mechanistic evidence linking ELF3 to any Mendelian disease. Absence of mention in this paper is not definitive disproof that ELF3 is or is not associated with a Mendelian disorder in other literature, but based on this document alone there is no support for the claim. Excerpt page numbers were not provided in the content, so excerpt_pages are set to null.


[Read Paper](https://www.semanticscholar.org/paper/2a5f1f23b76c02c35f17bd8a4ffc485355f8cd2e)


### De novo AHDC1 Deletions Identified by Genome Sequencing in Two Individuals with Xia-Gibbs Syndrome

**Why Not Relevant**: This paper exclusively reports on AHDC1 and Xia-Gibbs syndrome (XGS). All described genetic findings, mechanistic interpretations, and clinical correlations concern heterozygous variants or deletions of AHDC1 and AHDC1 haploinsufficiency as the disease mechanism in XGS. The provided content contains no mention of ELF3, no data about variants in ELF3, no functional or mechanistic experiments involving ELF3, and no clinical or genetic evidence linking ELF3 to any Mendelian phenotype. Therefore the paper provides no direct evidence (variant-disease co-segregation, case reports, functional validation) nor mechanistic evidence (pathway links, protein interactions, expression/knockout phenotypes) relevant to the claim that ELF3 is likely associated with at least one Mendelian disease.

Limitations and caveats: the absence of any mention of ELF3 in this paper does not prove ELF3 is not associated with Mendelian disease overall — it only means this particular paper offers no supporting or refuting data. To evaluate the claim about ELF3, one should review studies, case reports, or databases (e.g., OMIM, ClinVar, HGMD, gene-disease curated resources) that specifically assess ELF3 variants and their clinical consequences. Recommended next steps: search the primary literature and clinical variant databases for "ELF3" combined with terms such as "mutation", "variant", "Mendelian", "syndrome", "heritable", "de novo", or specific disease phenotypes.


[Read Paper](https://www.semanticscholar.org/paper/8ab12fd3ec0c21caabba032cb023668720bf18f8)


### Distinct Treatment Outcomes between MVID Patient-Modeled Mice Suggest Epithelial Cell Maturation Is Influenced by MYO5B Functional State

**Why Not Relevant**: Relevance is near zero because the abstract exclusively addresses Microvillus inclusion disease (MVID) caused by mutations in Myosin Vb (MYO5B), the generation and analysis of MYO5B-mutant mouse models, and treatments targeting pathways altered by MYO5B loss. Nowhere in the provided abstract text is the gene ELF3 mentioned, nor are there any genetic association data, clinical reports, pedigree analyses, variant pathogenicity assessments, or statements implicating ELF3 in a Mendelian disorder. The methods (e.g., RNA-sequencing of enteroids from Myo5b-mutant mice) and results are specifically framed around MYO5B perturbations; they do not provide direct evidence (e.g., linkage, segregation, de novo or biallelic pathogenic variants) or mechanistic evidence linking ELF3 to a Mendelian disease. Limitations that justify this assessment: the content provided is an abstract only and may omit secondary findings, but given the explicit MYO5B focus in background, methods, results, and conclusions, there is no basis in this document to support the claim about ELF3. To evaluate the claim that ELF3 is likely associated with at least one Mendelian disease, one would need papers presenting genetic evidence (human pedigrees, pathogenic variants, case reports, ClinVar/HGMD entries) or functional studies directly implicating ELF3 variants in Mendelian phenotypes—none of which are present in this abstract.


[Read Paper](https://www.semanticscholar.org/paper/16b55b636bdea532fff1a51c4dc68be2d32bd01d)


### De novo HDAC3 variants leading to epigenetic machinery dysfunction are associated with a neurodevelopmental disorder

**Why Not Relevant**: The provided paper focuses exclusively on HDAC3 (histone deacetylase 3) and reports heterozygous de novo variants in HDAC3 that associate with a neurodevelopmental syndrome. There is no mention of the gene ELF3 anywhere in the supplied text, no sequence or variant data concerning ELF3, and no mechanistic discussion involving ELF3. Therefore the paper provides neither direct evidence (e.g., genetic variants in ELF3 segregating with disease) nor mechanistic evidence (e.g., functional assays, protein interactions, localization) relevant to the claim that ELF3 is likely associated with at least one Mendelian disease. While the paper contains both genetic and functional evidence tying HDAC3 to disease (as shown in the excerpts), those data are gene-specific. One cannot infer from HDAC3 pathogenic mechanisms or disease association that ELF3 is likewise implicated without independent evidence; gene functions, interactions, and modes of pathogenicity differ and are not interchangeable. In summary, because ELF3 is not discussed or analyzed in this paper, it does not support or refute the claim about ELF3 and thus is not relevant to the claim.


[Read Paper](https://www.semanticscholar.org/paper/524b09f9e39f306b6871f816044577dccb53abd6)


### THE JAPANESE TERATOLOGY SOCIETY 62ND ANNUAL MEETING

**Why Not Relevant**: The supplied paper content does not mention the gene ELF3, any variants in ELF3, nor any phenotypes or Mendelian disorders linked to ELF3. The text discusses organogenesis, interkinetic nuclear migration, Sonic hedgehog (Shh) signaling, Satb2 as a downstream target of Shh, chemical teratogenicity in zebrafish, Wnt signaling, leukemia inhibitory factor (LIF), CRH/Pomc signaling, Galectin-9, and a brief note about genome editing (with an example concerning Il2rg and X-SCID). There are no genetic association studies, family or case reports, linkage/segregation data, loss/gain-of-function variants, knockout phenotypes for ELF3, functional assays of ELF3, or clinical descriptions implicating ELF3 in any Mendelian disease. Mechanistic discussions in the paper concern other genes/pathways (Shh, Satb2, Wnt, LIF, CRH/Pomc) and do not provide a plausible mechanistic link to ELF3. Therefore the paper provides no direct evidence and no mechanistic evidence for the claim that ELF3 is associated with at least one Mendelian disease. Limitations: because ELF3 is not discussed at all in the provided content, no inferences about ELF3 can be drawn from this paper; a relevant assessment would require papers that explicitly study ELF3 (e.g., human genetic studies, model organism knockouts of ELF3, functional assays of ELF3 variants, or clinical case reports).


[Read Paper](https://www.semanticscholar.org/paper/162811312bde34697100a4eaab4c8748157e2132)


### Abstracts of papers presented at the 24th Genetics Society ’ s Mammalian Genetics and Development Workshop held at the Institute of Child Health, University College London on 21st November 2013

**Why Not Relevant**: I found no mention of ELF3 (or common synonyms such as ELF-3, ESE-1, ESE1, or 'E74-like factor 3') in the provided paper content. The text is a compendium of short abstracts describing work on many other genes and processes (e.g., Fgf10, CHD7, MAP3K4, HEATR2, ATMIN, Caph2) and their links to developmental processes or Mendelian disorders (for example, CHD7 haploinsufficiency is explicitly implicated in CHARGE syndrome), but none of these passages refers to ELF3. Because ELF3 is not present in the supplied content, there is neither direct evidence (genetic association, patient data, linkage, segregation, pathogenic variants) nor mechanistic evidence (biochemical function, pathways, model-organism phenotypes linking ELF3 to a Mendelian phenotype) to support or refute the claim that ELF3 is likely associated with at least one Mendelian disease. Limitations and caveats: (1) The absence of ELF3 in the excerpted content means this source cannot be used to evaluate the claim. (2) The supplied text appears to be a set of abstracts or summaries and may be truncated; ELF3 could appear elsewhere in the full document or in other publications. Recommended next steps: search primary genetic-disease resources (OMIM, ClinVar), PubMed, and recent reviews for 'ELF3' and synonyms to identify any reported Mendelian disease associations and supporting mechanistic or clinical evidence.


[Read Paper](https://www.semanticscholar.org/paper/b9217a59c8a6c141a43120d206471e3c06d88029)


## Search Queries Used

- ELF3 ESE1 ESX E74 like ETS transcription factor 3 germline variant pathogenic variant patient case report Mendelian disease

- ELF3 de novo variant exome sequencing trio sequencing developmental disorder congenital anomaly rare variant burden monogenic disease

- ELF3 knockout mouse phenotype loss of function haploinsufficiency mouse model epithelial differentiation congenital defect

- ELF3 gnomAD loss of function intolerance pLI ClinVar OMIM gene disease association monogenic Mendelian


## Bibliometric Configuration

- Use Bibliometrics: True

- Author Impact Weight: 0.4

- Citation Impact Weight: 0.4

- Venue Impact Weight: 0.2
