## Supplementary material for "MENDELSEEK: An algorithm that predicts Mendelian Genes and elucidates what makes them special": MCM7.txt

# Claim: The gene MCM7 is likely associated with at least one Mendelian disease.

**Status**: processed

**Overall Rating**: -1

**Explanation**:

No relevant papers were found that support or refute this claim.


## Relevant Papers


## Other Reviewed Papers


### Identifying genomic variant associated with long QT syndrome type 2 in an ecuadorian mestizo individual: a case report

**Why Not Relevant**: This paper is a case report of congenital Long QT Syndrome type 2 and describes a pathogenic variant in KCNH2 and variants of uncertain significance in AKAP9 and TTN. The provided text contains no mention of the gene MCM7, no MCM7 variants, and no mechanistic discussion involving MCM7. Therefore it provides neither direct evidence (no observed MCM7 pathogenic variants or clinical associations) nor mechanistic evidence (no discussion of MCM7 function, pathways, or plausible links to Mendelian disease) bearing on the claim that MCM7 is likely associated with at least one Mendelian disease. Absence of mention in this single LQTS case report does not disprove the claim (absence of evidence is not evidence of absence), but it does mean this paper neither supports nor refutes the claim. To evaluate the claim about MCM7, literature or database searches specifically targeting MCM7 (ClinVar, OMIM, HGMD, PubMed) or studies reporting Mendelian phenotypes linked to MCM7 would be required.


[Read Paper](https://www.semanticscholar.org/paper/aa65df486c4f1720d86feb42ddc72ed25ab57431)


### Prenatal ultrasound findings of X-linked congenital cataracts: case report and description of a novel variant.

**Why Not Relevant**: This paper does not provide evidence for or against the claim that MCM7 is likely associated with at least one Mendelian disease. The content of the paper describes a prenatal case of bilateral congenital cataracts and reports a pathogenic variant in exon 11 of the OCRL gene, implicating X-linked recessive disorders such as Dent disease 2 (OMIM: 300555) or Lowe's syndrome (OMIM: 309000). There is no mention of the gene MCM7, no data on MCM7 variants, and no mechanistic discussion linking MCM7 to congenital cataracts or to any Mendelian phenotype. Because the paper focuses solely on OCRL in a single prenatal case report (ultrasound findings, family history, and genetic testing of the fetus and family members), it contains neither direct genetic evidence (variants, segregation, frequency, pathogenicity data) nor mechanistic evidence (molecular function, pathways, animal or cellular models) relating MCM7 to Mendelian disease. Additionally, the study design (a single-case prenatal report) limits generalizability and would not support broad gene–disease inferences even if MCM7 had been mentioned. To evaluate the claim about MCM7, a literature search for studies reporting pathogenic MCM7 variants, segregation with disease, functional assays, or OMIM entries would be required; this paper provides no such information.


[Read Paper](https://www.semanticscholar.org/paper/058d8e6037d294c050f1e435bf38ad6e48b453db)


### Whole Exome Sequencing and Panel-Based Analysis in 176 Spanish Children with Neurodevelopmental Disorders: Focus on Autism Spectrum Disorder and/or Intellectual Disability/Global Developmental Delay

**Why Not Relevant**: This paper does not provide any direct or mechanistic evidence relevant to the claim that MCM7 is likely associated with at least one Mendelian disease. The provided content contains no mention of the gene MCM7, no gene-level results, no case reports of patients with variants in MCM7, and no functional/mechanistic data concerning MCM7. The paper reports aggregate diagnostic yields from whole exome sequencing (WES), the proportion of variants of uncertain significance (VUS), and the impact of parental testing, but no specific genes or specific Mendelian disease associations are described. Therefore, nothing in the paper allows assessment of whether MCM7 is associated with a Mendelian disease. To evaluate the claim one would need variant-level findings (e.g., pathogenic or likely pathogenic variants in MCM7), segregation data, phenotype descriptions linked to MCM7 variants, or functional/mechanistic studies implicating MCM7 in a disease process—none of which are present in the provided text. Additional limitations that prevent inference about MCM7 from this paper: (1) results are reported only as overall diagnostic yield and VUS frequency without gene lists; (2) no genotype–phenotype correlations or specific variant classifications by gene are given; (3) absence of any mechanistic discussion of particular genes; and (4) even if WES was used, the paper does not report which genes contributed to diagnoses. Because of these factors, the paper is not informative for assessing the claim about MCM7.


[Read Paper](https://www.semanticscholar.org/paper/c6545402f04cad1b5fa80d4e00f55b18237edb1b)


### Implications of a De Novo Variant in the SOX12 Gene in a Patient with Generalized Epilepsy, Intellectual Disability, and Childhood Emotional Behavioral Disorders

**Why Not Relevant**: The provided paper content exclusively addresses the SOX gene family and, specifically, a de novo variant in SOX12. There is no mention of the gene MCM7, no data (genetic, clinical, or mechanistic) about MCM7, and no discussion that would allow inference about MCM7's involvement in Mendelian disease. Because the content neither addresses nor references MCM7 directly or indirectly, it cannot provide supporting or refuting evidence for the claim that MCM7 is likely associated with at least one Mendelian disease. Absence of mention in this SOX-focused study is not evidence about MCM7; to evaluate the claim one would need studies that explicitly analyze MCM7 (case reports, genetic association/Mendelian segregation evidence, functional studies of pathogenic variants, or curated clinical-database entries).


[Read Paper](https://www.semanticscholar.org/paper/5cf652470efa4406f72843aea8389f4e398a99f8)


### New Pathogenic Homozygous Variant in (DGOUK) Gene Cause Vital Progressive Liver Failure in a Neonate: Case Report

**Why Not Relevant**: The provided paper content contains no mention of the gene MCM7. The text focuses on mitochondrial DNA depletion syndromes (MDS) and reports a case implicating DGUOK (OMIM:601465). Although the paper discusses Mendelian inheritance and mechanistic aspects of MDS (mtDNA depletion and impaired respiratory chain synthesis), none of the excerpts reference MCM7, nor do they provide mechanistic pathways involving MCM7. Therefore, based on the supplied content, there is no direct or mechanistic evidence that MCM7 is associated with any Mendelian disease. Caveat: this assessment is limited to the supplied excerpt; the full paper might mention other genes such as MCM7 elsewhere, but that is not present in the text provided.


[Read Paper](https://www.semanticscholar.org/paper/74a649e26c83fe77b6b076668da3b563912accff)


### Case report: Preimplantation genetic testing for infantile GM1 gangliosidosis

**Why Not Relevant**: The provided paper exclusively studies GM1 gangliosidosis and genetic variants in the GLB1 gene; it contains no mention of the gene MCM7, no association analyses involving MCM7, no functional/mechanistic data about MCM7, and no clinical cases attributed to MCM7. Therefore it provides no direct evidence that MCM7 is associated with any Mendelian disease, nor mechanistic evidence that would increase or decrease the plausibility of such an association. Note: the absence of MCM7 in this paper does not constitute evidence that MCM7 is not associated with Mendelian disease in other literature; it only means this paper is irrelevant to evaluating that claim.


[Read Paper](https://www.semanticscholar.org/paper/ce981bc89c565bb4d97baff441baaf8695595448)


### A de novo PAK1 likely pathogenic variant and a de novo terminal 1q microdeletion in a Chinese girl with global developmental delay, severe intellectual disability, and seizures

**Why Not Relevant**: The provided content focuses solely on a novel PAK1 variant and an associated terminal 1q microdeletion in a patient with a neurodevelopmental disorder, and explicitly attributes the clinical features to PAK1. There is no mention of the gene MCM7, no data (genetic, functional, segregation, or mechanistic) concerning MCM7, and no gene list or coordinates for the microdeletion that would allow determination of whether MCM7 lies within the deleted interval. Therefore this paper content does not provide direct or mechanistic evidence that MCM7 is associated with any Mendelian disease. The absence of MCM7 in the provided text cannot be taken as definitive evidence against MCM7 involvement, because the excerpt is brief and may omit gene-level details; however, based on the content supplied, relevance to the claim is negligible.


[Read Paper](https://www.semanticscholar.org/paper/8ac30d308f85987b8cf4644520ee11940136ebaa)


### A novel AUTS2 variant in a patient with global developmental delay and intellectual disability

**Why Not Relevant**: The paper exclusively studies the AUTS2 gene and reports a de novo loss-of-function variant in AUTS2 associated with a neurodevelopmental disorder. There is no mention of the gene MCM7, no experimental data, no genetic association, and no mechanistic discussion involving MCM7. Therefore this paper provides no direct or mechanistic evidence for or against the claim that MCM7 is likely associated with at least one Mendelian disease. Absence of evidence in this paper is not evidence of absence—this single AUTS2-focused report cannot be used to infer anything about MCM7. Because none of the content addresses MCM7, relevance to the claim is essentially zero.


[Read Paper](https://www.semanticscholar.org/paper/f5e84e7bb3b42239a53cc7e5c457342cb38c6277)


## Search Queries Used

- MCM7 pathogenic variant Mendelian disease case report OMIM ClinVar

- MCM7 de novo variant whole exome sequencing neurodevelopmental disorder developmental delay intellectual disability

- MCM7 loss of function haploinsufficiency DNA replication minichromosome maintenance complex knockout mouse embryonic phenotype congenital malformation

- MCM7 gnomAD loss of function constraint pLI intolerance MCM family germline mutation cancer predisposition monogenic disorder review


## Bibliometric Configuration

- Use Bibliometrics: True

- Author Impact Weight: 0.4

- Citation Impact Weight: 0.4

- Venue Impact Weight: 0.2
