## Supplementary material for "MENDELSEEK: An algorithm that predicts Mendelian Genes and elucidates what makes them special": MLLT1.txt

# Claim: The gene MLLT1 is likely associated with at least one Mendelian disease.

**Status**: processed

**Overall Rating**: -1

**Explanation**:

No relevant papers were found that support or refute this claim.


## Relevant Papers


## Other Reviewed Papers


### Diagnostic yield of patients with undiagnosed intellectual disability, global developmental delay and multiples congenital anomalies using karyotype, microarray analysis, whole exome sequencing from Central Brazil

**Why Not Relevant**: The paper does not mention the gene MLLT1 by name and provides no gene-level results that would allow assessment of an association between MLLT1 and any Mendelian disease. All gene-related information in the manuscript excerpt is aggregate or methodological (size of the gene panel, number of trios, overall diagnostic yield and variant classifications), without listing which genes carried pathogenic, likely pathogenic, or VUS findings. Therefore there is neither direct evidence (e.g., identification of pathogenic MLLT1 variants segregating with disease) nor mechanistic evidence (e.g., functional studies implicating MLLT1 in a disease pathway) in this paper to support or refute the claim. To evaluate the claim about MLLT1 one would need explicit reporting of variants in MLLT1 with clinical and segregation data, recurrence across unrelated cases, and ideally functional validation demonstrating a plausible disease mechanism. Because those items are absent here, the relevance of this paper to the claim is effectively zero.


[Read Paper](https://www.semanticscholar.org/paper/23767daf16f48d9a91172aa0fc624f07b869a9c8)


### Whole-Exome Sequencing in Adults With Chronic Kidney Disease

**Why Not Relevant**: The provided excerpt contains no mention of the gene symbol "MLLT1" and presents no data—genetic, clinical, or mechanistic—linking MLLT1 to any Mendelian disease. The text describes a study of whole-exome sequencing (WES) applied to adults with chronic kidney disease, the methods used for sequencing and variant interpretation, and the authors' creation of a curated list of 287 OMIM genes implicated in Mendelian kidney disorders (referred to as "nephropathy genes"). However, the excerpt does not include the full gene list (Supplement Table 3), the study's variant/diagnosis results, or any supplementary material where MLLT1 might appear. Therefore, from the material provided here one cannot (a) find direct evidence that MLLT1 is reported as disease-associated in this study, nor (b) evaluate any mechanistic arguments linking MLLT1 to a Mendelian disorder.

Limitations and caveats relevant to this assessment:
- Absence of mention in the excerpt is not evidence that the full paper or its supplements do not mention MLLT1. The gene could appear in Supplementary Table 3 (the curated gene list) or in result tables not included in the excerpt. Without those sections, we cannot determine whether the authors assessed MLLT1 or observed variants in it.
- The paper's focus is on nephropathy genes and Mendelian causes of kidney disease; if MLLT1 is not a known kidney disease gene, this study may not be an appropriate source to evaluate MLLT1's broader Mendelian disease associations.
- The study does describe methods (WES with confirmation by Sanger sequencing, use of ClinVar/OMIM for pathogenicity assessment) that could detect Mendelian disease genes in principle, but no evidence specific to MLLT1 is provided in the excerpt.

Recommended next steps to evaluate the claim properly:
1. Search the full paper and all supplementary files (especially Supplement Table 3 and the results/tables) for the string "MLLT1." 2. If MLLT1 is listed among genes or reported with variants, extract the exact sentences, the variant details, ACMG classifications, and any clinical correlation (phenotype) provided. 3. Independently check clinical databases (OMIM, ClinVar) and the literature for MLLT1-associated Mendelian phenotypes. Until those steps are done, the supplied excerpt neither supports nor refutes the claim that "MLLT1 is likely associated with at least one Mendelian disease."


[Read Paper](https://www.semanticscholar.org/paper/a29bd57964f68ac5f8e1abcafe1112ba488f53f5)


### The simplest explanation does not have to be preferred: co-occurrence of pathogenic variants in cancer-predisposing genes

**Why Not Relevant**: The single provided sentence discusses reviewing the literature and checking UK 10,000 Genomes Project data for MINAS (tumor predisposition syndromes or multiple primary tumors) and whether effects of cancer susceptibility genes (CSGs) are additive or synergistic: "The frequency and cancer phenotypes associated with MINAS by reviewing the recent literature and by checking the UK 10,000 Genomes Project data from participants with a phenotype of tumor predisposition syndromes or multiple primary tumors, and if they are additive reflecting each CSG independently or synergistic resulting in more severe phenotypes are reported."

This excerpt contains no mention of the gene MLLT1, no data or results about specific genes, and no mechanistic descriptions linking any gene (including MLLT1) to Mendelian disease. Therefore, based solely on the provided content, the paper gives no direct evidence supporting or refuting the claim that MLLT1 is likely associated with at least one Mendelian disease. It also provides no mechanistic evidence (e.g., protein function, pathogenic variants, segregation, functional assays) that would bear on the plausibility of such an association.

Limitations and caveats: (1) Only a single sentence/fragment of the paper was provided. It is possible that other parts of the full paper discuss specific genes including MLLT1; that information was not available here. (2) The excerpt describes scope and data sources (literature review and UK 10k Genomes participants with tumor-predisposition phenotypes) but contains no methods, results, or gene-level findings, so no inference about MLLT1 can be made. (3) Because no experimental conditions, sample sizes, statistical results, or variant-level information are provided, no assessment of evidence strength, statistical or practical significance, or generalizability can be performed.

If you can provide additional text (results, tables, or gene lists) from the paper or other papers that mention MLLT1, I can re-evaluate and extract relevant verbatim passages and assess direct and mechanistic evidence for an association with a Mendelian disease.


[Read Paper](https://www.semanticscholar.org/paper/a1372a664bd2c33a0bda638975a8b26d112457fc)


### A novel de novo hemizygous ARHGEF9 mutation associated with severe intellectual disability and epilepsy: a case report

**Why Not Relevant**: The supplied paper content exclusively concerns the gene ARHGEF9 and a novel hemizygous loss-of-function deletion (c.656_c.669delACTTCTTTGAGGCC (p. His219Leu fs*9)) identified in a boy with severe intellectual disability, epilepsy, and developmental regression. The text includes statements such as “ARHGEF9 encodes collybistin, a brain-specific guanosine diphosphate-guanosine-5′-triphosphate exchange factor…” and “Here, we report a boy with clinical symptoms of severe intellectual disability, epilepsy, and developmental delay and regression. Trio exome sequencing ... identified a novel hemizygous deletion ... in exon 5 of ARHGEF9.” Nowhere in the provided content is the gene MLLT1 mentioned, nor are there descriptions of biological pathways, protein interactions, shared phenotypes, or genetic loci that link ARHGEF9 to MLLT1. Because the paper neither discusses MLLT1 nor provides mechanistic or epidemiological data that could plausibly be used to infer an association between MLLT1 and a Mendelian disease, it provides no direct or mechanistic evidence relevant to the claim that “MLLT1 is likely associated with at least one Mendelian disease.”

Limitations and caveats on this assessment:
- Absence of evidence in this paper is not evidence of absence: the paper’s lack of mention of MLLT1 does not prove MLLT1 is not associated with a Mendelian disease. It only means this paper does not inform that question.
- No indirect inference is possible from the presented ARHGEF9 data because there is no information linking ARHGEF9 and MLLT1 (no shared pathway, interaction, co-expression, or locus information provided).

Recommendation: to evaluate the claim about MLLT1, consult primary sources that directly address MLLT1 (OMIM, ClinVar, HGMD, locus-specific studies, case reports, or gene-disease curation resources such as ClinGen), or search the literature for studies explicitly reporting pathogenic variants in MLLT1 or functional data implicating it in Mendelian phenotypes.


[Read Paper](https://www.semanticscholar.org/paper/988bf7595fc677ddeb7de11713ff3345556d6a96)


### De novo HDAC3 variants leading to epigenetic machinery dysfunction are associated with a neurodevelopmental disorder

**Why Not Relevant**: The paper exclusively studies HDAC3: its de novo variants, functional assays, protein interactions, and clinical correlations in a small cohort. The gene MLLT1 is not mentioned anywhere in the provided content. Therefore this paper provides no direct genetic, clinical, or mechanistic evidence supporting or refuting an association between MLLT1 and any Mendelian disease. Findings about HDAC3 cannot be assumed to apply to MLLT1 because they are distinct genes with different sequences, functions, and interaction partners. Consequently, the relevance of this paper to the claim that MLLT1 is likely associated with at least one Mendelian disease is effectively zero.


[Read Paper](https://www.semanticscholar.org/paper/524b09f9e39f306b6871f816044577dccb53abd6)


### Novel De Novo Heterozygous Variants in the SON Gene Causing ZTTK Syndrome: A Case Report of Two Patients and Review of Neurological Findings

**Why Not Relevant**: The provided paper content discusses ZTTK syndrome caused by mutations in the SON gene (located at 21q22.11), describing clinical features and reporting new pathogenic variants in SON identified via microarray analysis and whole-exome sequencing. There is no mention of the gene MLLT1 anywhere in the text, nor any discussion of genes other than SON. Therefore the paper provides no direct evidence (no case reports, variant associations, or statistical associations) linking MLLT1 to any Mendelian disease. It also provides no mechanistic evidence (no functional studies, pathway analyses, protein interactions, or regulatory relationships) that would plausibly connect MLLT1 to the disease processes described. Because the necessary information to support or refute the claim about MLLT1 is entirely absent, relevance to the claim is essentially zero. Limitations: the excerpt is limited to a report on SON and ZTTK syndrome; it is possible that other sections of the full article outside the provided content could mention additional genes, but based solely on the supplied text there is no basis to evaluate MLLT1. If you want an assessment of MLLT1 specifically, I can search the literature and extract relevant evidence linking MLLT1 to Mendelian disease(s).


[Read Paper](https://www.semanticscholar.org/paper/5667c04c1f6b809971401be55530db53c77310e6)


## Search Queries Used

- MLLT1 germline variant Mendelian disease monogenic syndrome OMIM ClinVar pathogenic case report familial segregation

- MLLT1 mutation de novo developmental disorder intellectual disability congenital anomaly exome sequencing trio cohort case series

- MLLT1 ENL YEATS domain chromatin regulator loss of function knockout mouse model embryonic development cellular differentiation mechanistic study

- MLLT1 gnomAD constraint loss of function intolerance pLI haploinsufficiency ClinGen variant frequency population database variant interpretation


## Bibliometric Configuration

- Use Bibliometrics: True

- Author Impact Weight: 0.4

- Citation Impact Weight: 0.4

- Venue Impact Weight: 0.2
