## Supplementary material for "MENDELSEEK: An algorithm that predicts Mendelian Genes and elucidates what makes them special": MLLT3.txt

# Claim: The gene MLLT3 is likely associated with at least one Mendelian disease.

**Status**: processed

**Overall Rating**: -1

**Explanation**:

No relevant papers were found that support or refute this claim.


## Relevant Papers


## Other Reviewed Papers


### Clinical whole-exome sequencing for the diagnosis of mendelian disorders.

**Why Not Relevant**: The provided paper excerpt contains no mention of the gene MLLT3 and provides no gene-level findings that could be used to support or refute an association between MLLT3 and a Mendelian disease. The text is a high-level summary of a clinical whole-exome sequencing (WES) cohort (first 250 probands) reporting overall diagnostic yield (25%), counts of mutated alleles (86 in 62 patients), and inheritance-mode summaries (33 autosomal dominant, 16 autosomal recessive, 9 X-linked; percentages of de novo events). None of these sentences refer to MLLT3, describe variants in MLLT3, provide segregation or functional evidence for MLLT3, or discuss biological mechanisms involving MLLT3. Therefore there is no direct evidence (variant reports, segregation, recurrence, case descriptions) and no mechanistic evidence (functional assays, pathway analyses, model organisms) in the supplied content relevant to the claim.

Limitations and caveats: (1) My assessment is limited to the exact paper content you provided. The full paper (or its supplementary tables) might list specific genes and variants identified, and MLLT3 could conceivably appear there; that information is not present in the excerpt. (2) This excerpt is a summary/abstract-like passage and therefore may omit gene-level detail that could be elsewhere in the manuscript or supplemental data. Because none of those details are included here, I cannot evaluate MLLT3 specifically from this text.

Recommended next steps to evaluate the claim: search the full paper and its supplementary materials for "MLLT3" or check variant/gene tables; search curated resources (OMIM, ClinVar, ClinGen, HGMD) and primary literature (PubMed) for MLLT3 disease associations; query databases of rare variants and constraint metrics (gnomAD, DECIPHER) and Matchmaker Exchange for reported Mendelian cases involving MLLT3; and look for functional studies demonstrating pathogenic mechanisms (loss/gain of function, animal models) that would support causality.

Because the supplied text contains no MLLT3-specific data or mechanistic discussion, it cannot be used to support or refute the claim that "MLLT3 is likely associated with at least one Mendelian disease."


[Read Paper](https://www.semanticscholar.org/paper/73921a10d537f4f1013f1627227301a056facc73)


### Congenital immunodeficiency in an individual with Wiedemann–Steiner syndrome due to a novel missense mutation in KMT2A

**Why Not Relevant**: The paper exclusively discusses Wiedemann–Steiner Syndrome and implicates KMT2A heterozygous mutations (including a de novo KMT2A missense mutation) as the molecular cause in the reported case. There is no mention of MLLT3, no genetic variants reported in MLLT3, no association analyses, and no mechanistic discussion involving MLLT3. Therefore the paper provides no direct or mechanistic evidence supporting or refuting the claim that MLLT3 is likely associated with at least one Mendelian disease. Because the gene discussed (KMT2A) is different from the gene in the claim (MLLT3), the content is not relevant to assessing MLLT3’s Mendelian disease association.


[Read Paper](https://www.semanticscholar.org/paper/2a32dfef52c1aed451b77191d9c569f0aae4050e)


### Further delineation of the rare GDACCF (global developmental delay, absent or hypoplastic corpus callosum, dysmorphic facies syndrome): genotype and phenotype of 22 patients with ZNF148 mutations

**Why Not Relevant**: The paper exclusively investigates pathogenic variants in ZNF148 and describes clinical and molecular genetic features of individuals with ZNF148 mutations. The provided content contains no mention of the gene MLLT3, no data, no case reports, and no mechanistic experiments concerning MLLT3. Because the paper does not study or reference MLLT3 in any way, it cannot provide direct evidence that MLLT3 is associated with a Mendelian disease, nor can it provide mechanistic evidence that would strengthen such a claim by analogy. Therefore the paper is not relevant to the claim that MLLT3 is likely associated with at least one Mendelian disease.


[Read Paper](https://www.semanticscholar.org/paper/db6f85fe2efebebb6838530baf54ac26288fd0b5)


### First Report of a de novo 10q23.31q23.33 Microdeletion: Obesity, Intellectual Disability and Microcephaly

**Why Not Relevant**: The paper reports a pathogenic 3.1-Mb deletion at 10q23.31–10q23.33 and attributes the patient's phenotype to haploinsufficiency of genes within that deleted interval (GPR120, KIF11, EXOC6, CYP26A1, CYP26C1, and LGI1). The gene MLLT3 is not mentioned anywhere in the provided text, and none of the candidate genes listed are MLLT3. Therefore the paper contains no direct genetic, clinical, or mechanistic evidence linking MLLT3 to any Mendelian disease. The study is a single-case CNV report (limiting generalizability), and its findings are confined to the genes within the reported deletion; this further limits any inference about genes not present in that interval, including MLLT3.


[Read Paper](https://www.semanticscholar.org/paper/31000c6c6083b249dcc5a5449ca4c1c34cf06243)


### Exome sequencing reveals neurodevelopmental genes in simplex consanguineous Iranian families with syndromic autism

**Why Not Relevant**: The provided paper content contains a single statement: "The discovery of variants in two muscular dystrophy genes, DMD and CHKB, indicates that some genes transcend the boundary of apparently unrelated clinical categories, thereby establishing a novel connection between ASD and muscular dystrophy." This sentence (1) mentions only DMD and CHKB, (2) draws a broad conceptual conclusion about genes crossing clinical-category boundaries and a connection between ASD and muscular dystrophy, and (3) contains no mention of the gene MLLT3, no data about MLLT3 variants, no clinical phenotypes attributable to MLLT3, and no mechanistic evidence linking MLLT3 to any Mendelian disorder. Therefore the content offers neither direct evidence (e.g., variant identification, segregation, functional studies, case descriptions) nor mechanistic evidence (e.g., pathway involvement, loss-of-function intolerance, model organism phenotypes) that would support or refute the claim that MLLT3 is likely associated with at least one Mendelian disease. The general observation that "some genes transcend the boundary of apparently unrelated clinical categories" is a conceptual point but is non-specific and cannot be used to infer an association for a particular gene (MLLT3) without gene-specific data. In short, because MLLT3 is not mentioned and no relevant data or mechanisms about it are provided, the paper content is not relevant to the claim.


[Read Paper](https://www.semanticscholar.org/paper/9cb23120953e32cc9faf4f47f59f270690fe7487)


### Germline mosaicism in a collagen VI‐related myopathy family: A cause of autosomal recessive inheritance

**Why Not Relevant**: The paper exclusively reports on COL6A1 (and related COL6A2/COL6A3) variants and paternal germline mosaicism causing an autosomal recessive collagen VI-related myopathy (Ullrich congenital muscular dystrophy) in a family. The text contains no mention of MLLT3, no genetic, clinical, or mechanistic data concerning MLLT3, and no analyses (e.g., variant calls, segregation, functional studies) relevant to MLLT3. Therefore this paper provides no direct or mechanistic evidence for or against the claim that MLLT3 is associated with a Mendelian disease. Absence of evidence in this paper does not imply evidence of absence elsewhere; it simply means this paper is irrelevant to the MLLT3 claim.


[Read Paper](https://www.semanticscholar.org/paper/4572f635bb354974678a75579201d9eedc2c40ea)


### X-Linked Intellectual Disability with NEXMIF Gene Mutation and Developmental Delay with GNAO1 Gene Mutation: Case Report

**Why Not Relevant**: The provided paper content does not mention the gene MLLT3 anywhere. It discusses X-linked intellectual disability (XLID) and specifically describes variants in the NEXMIF (KIAA2022) and GNAO1 genes in a single affected child, along with the clinical phenotype. Because MLLT3 is not referenced, there is no direct evidence (e.g., reported pathogenic variants, segregation, case series) nor any mechanistic evidence (e.g., functional studies, pathway links) in this paper that would support or refute the claim that MLLT3 is likely associated with a Mendelian disease. The only genes named are NEXMIF and GNAO1, and all quoted clinical and genetic findings in the text relate to those genes; therefore no verbatim excerpts can be provided that are relevant to MLLT3. In short, the paper is not relevant to the claim because it contains no information about MLLT3, offers no mechanistic pathways involving MLLT3, and provides no data that could be extrapolated to assess MLLT3’s involvement in Mendelian disease.


[Read Paper](https://www.semanticscholar.org/paper/f313ed42d3ded706a55d1a60bbf8fbecf55a57c2)


## Search Queries Used

- MLLT3 germline variant Mendelian disease monogenic pathogenic variant autosomal dominant autosomal recessive de novo human exome sequencing

- MLLT3 mutation developmental delay intellectual disability congenital anomaly dysmorphism neurodevelopmental disorder epilepsy clinical case report

- MLLT3 loss of function haploinsufficiency knockout mouse zebrafish model hematopoiesis neuronal development phenocopy germline mutation functional study

- MLLT3 ClinVar OMIM HGMD ClinGen gene disease association pathogenic evidence gnomAD pLI loss of function intolerance somatic fusion leukemia MLL translocation AF9


## Bibliometric Configuration

- Use Bibliometrics: True

- Author Impact Weight: 0.4

- Citation Impact Weight: 0.4

- Venue Impact Weight: 0.2
