## Supplementary material for "MENDELSEEK: An algorithm that predicts Mendelian Genes and elucidates what makes them special": MMS19.txt

# Claim: The gene MMS19 is likely associated with at least one Mendelian disease.

**Status**: processed

**Overall Rating**: 3

**Explanation**:

Supporting evidence.
The provided papers collectively establish MMS19 as an essential component of the cytosolic iron–sulfur (Fe–S) cluster assembly (CIA) targeting complex and document its direct role in maturation of multiple nuclear DNA metabolism proteins. Vashisht & Wohlschlegel and the review-style excerpts state that MMS19 (with CIAO1 and FAM96B) is required for Fe–S insertion into extramitochondrial proteins including the XPD helicase, and that XPD must acquire its Fe–S cluster from the CIA targeting complex before assembling into TFIIH. The CIA pathway papers explicitly list MMS19 as interacting with numerous Fe–S client proteins that are themselves linked to nuclear DNA integrity—DNA polymerases (e.g., POLD1), helicases involved in DNA repair (XPD, FANCJ), and telomere maintenance factors (RTEL1). Functional/structural work (Kassube et al.) further shows MMS19 has a conserved structural role in client binding and Fe–S delivery. Phenotypic consequences of MMS19 dysfunction are also reported: cells with mutated MMS19 are hypersensitive to DNA damage and have altered telomere length, and mms19 null alleles are lethal in some organisms. Because many of the MMS19 client proteins are known causes of Mendelian disorders when mutated (e.g., XPD mutations cause xeroderma pigmentosum / trichothiodystrophy-related syndromes; POLD1/RTEL1/FANCJ mutations cause Mendelian syndromes), perturbation of MMS19 function could plausibly produce Mendelian phenotypes either by directly impairing those clients or by causing independent genetic disease.

Caveats or contradictory evidence.
None of the provided excerpts report a direct, documented association between human MMS19 variants and a named Mendelian disease. The evidence is mechanistic and correlative: MMS19 is required for assembly of disease-relevant proteins, and MMS19 perturbation yields cellular phenotypes (DNA damage sensitivity, telomere changes), but no clinical case series, segregating family data, or clinical genetic findings linking MMS19 mutations to specific inherited disorders are presented. Additionally, the observation that mms19 null alleles are lethal in some species raises the possibility that complete loss-of-function in humans might be embryonic lethal, which would reduce the chance of observing a classical inherited syndrome in living patients; conversely, partial loss-of-function or hypomorphic alleles might have subtler or late-onset phenotypes (e.g., cancer predisposition) rather than clear-cut Mendelian traits. The plant study shows species differences (Arabidopsis mms19 mutants viable), underscoring that organismal consequences of MMS19 disruption can vary and cannot be directly extrapolated to human Mendelian disease without human genetic evidence.

Analysis of potential underlying mechanisms, deeper relationships, and implications.
Mechanistically, MMS19 functions as a scaffold/docking and specificity factor in the CIA targeting complex, mediating Fe–S delivery to a set of nuclear clients that are critical for DNA replication, repair, telomere maintenance, and transcription. Loss or dysfunction of MMS19 could therefore produce multi-system cellular defects by undermining the function of many Fe–S–containing client proteins. Two broad outcomes are plausible: (1) severe, widespread LOF in MMS19 might be incompatible with development (embryonic lethality) or produce pleiotropic syndromes difficult to ascribe to a single Mendelian label; (2) partial LOF or hypomorphic variants could phenocopy individual client gene defects or produce cancer/aging predisposition phenotypes. Given that multiple client proteins are themselves established Mendelian disease genes, MMS19 dysfunction could act as an upstream cause of similar phenotypes (genetic epistasis). However, the directionality matters: some client proteins may accept Fe–S clusters from redundant pathways or have differing sensitivities to MMS19 impairment, producing heterogeneous outcomes. Also, phenotypes may be recessive if complete loss is required, or dominant-negative if specific alleles interfere with complex assembly.

Which rating is most appropriate based on the relative strength of the evidence.
The weight of the provided evidence supports high biological plausibility that MMS19 perturbation can cause human disease because of its central role in maturing multiple proteins whose defects are Mendelian. However, the excerpts do not provide direct human genetic or clinical evidence linking MMS19 variants to a named inherited disorder. The lethality of null alleles in model organisms further complicates expectations for observable Mendelian syndromes in humans. Given strong mechanistic plausibility but absence of documented human Mendelian associations in the supplied material, the most appropriate classification is "Mixed Evidence": the claim is plausible and supported mechanistically, but not directly demonstrated by the provided papers.


**Final Reasoning**:

I reviewed the provided excerpts for direct and indirect evidence that MMS19 is associated with at least one Mendelian disease. The literature supplied gives strong mechanistic support—MMS19 is a required CIA targeting complex component that inserts Fe–S clusters into many nuclear proteins (XPD, POLD1, FANCJ, RTEL1) that are themselves Mendelian disease genes; MMS19 dysfunction produces cellular DNA repair defects and organismal lethality in some species. These points make an association with human inherited disease biologically plausible: MMS19 loss could phenocopy or contribute to Mendelian disorders affecting DNA repair, replication, or telomere maintenance. However, the excerpts contain no human genetic or clinical reports of MMS19 mutations segregating with an inherited phenotype; they do not cite patient data, ClinVar/OMIM links, or familial segregation. The possibility that complete loss is embryonic lethal in humans further reduces the likelihood of observing a classical Mendelian syndrome due to null alleles, although hypomorphic alleles could produce disease. Balancing strong mechanistic plausibility against the lack of direct human genetic evidence provided, the correct judgment is that the evidence is mixed: suggestive but not definitive. Therefore I assign the rating "Mixed Evidence."


## Relevant Papers


### The Association of the Xeroderma Pigmentosum Group D DNA Helicase (XPD) with Transcription Factor IIH Is Regulated by the Cytosolic Iron-Sulfur Cluster Assembly Pathway*

**Authors**: A. Vashisht (H-index: 46), J. Wohlschlegel (H-index: 72)

**Relevance**: 0.4

**Bibliometric Impact**: 0.5617777777777778


**Excerpts**:

- Significance: The XPD-CIA targeting complex interaction is required for TFIIH assembly.

- Xeroderma pigmentosum group D (XPD) helicase is a component of the transcription factor IIH (TFIIH) transcription complex and plays essential roles in transcription and nucleotide excision repair.

- We recently identified a cytoplasmic Fe-S cluster assembly (CIA) targeting complex composed of MMS19, CIAO1, and FAM96B that is required for the biogenesis of extramitochondrial Fe-S proteins including XPD.

- Multiple lines of evidence indicate that this process occurs in a stepwise fashion in which XPD acquires a Fe-S cluster from the CIA targeting complex before assembling into TFIIH.

- First, XPD was found to associate in a mutually exclusive fashion with either TFIIH or the CIA targeting complex.

- Second, disrupting Fe-S cluster assembly on XPD by either 1) depleting cellular iron levels or 2) utilizing XPD mutants defective in either Fe-S cluster or CIA targeting complex binding blocks Fe-S cluster assembly and prevents XPD incorporation into TFIIH.

- Finally, subcellular fractionation studies indicate that the association of XPD with the CIA targeting complex occurs in the cytoplasm, whereas its association with TFIIH occurs largely in the nucleus where TFIIH functions.

- Conclusion: XPD maturation is a stepwise process in which XPD acquires its Fe-S cluster before binding TFIIH.


**Explanations**:

- This sentence states that the interaction between XPD and the CIA targeting complex is necessary for assembly of TFIIH. It is mechanistic evidence linking the CIA targeting complex (which includes MMS19) to the functional maturation of TFIIH via XPD. Strength: establishes functional consequence of the interaction. Limitation: it does not provide any genetic or clinical data tying MMS19 variants to human disease.

- This sentence places XPD in a critical DNA repair and transcriptional complex. It is mechanistic/contextual: showing that perturbation of XPD (and thus its biogenesis) can be expected to affect processes that, when defective, cause Mendelian disorders (e.g., nucleotide excision repair defects). Strength: explains biological importance of XPD. Limitation: the sentence does not mention MMS19 or provide direct evidence that MMS19 alterations cause disease.

- This sentence explicitly names MMS19 as a component of the cytoplasmic CIA targeting complex required for biogenesis of extramitochondrial Fe-S proteins, including XPD. This is direct mechanistic evidence that MMS19 participates in the maturation of XPD, a protein with established links to Mendelian disease when mutated. Strength: directly implicates MMS19 in the biochemical pathway that generates a known disease-related protein. Limitation: this is biochemical/cellular evidence of function; it does not present genetic association, patient data, or phenotypes caused by MMS19 variants.

- This sentence describes the sequential mechanism by which XPD acquires its Fe-S cluster from the CIA targeting complex prior to TFIIH assembly. This is mechanistic evidence showing MMS19-containing complex acts upstream of TFIIH incorporation of XPD. Strength: supports plausibility that defects in MMS19 could perturb XPD function. Limitation: mechanistic plausibility does not prove that MMS19 variants cause Mendelian disease in humans.

- This sentence provides a specific mechanistic observation that XPD alternates association with the CIA complex (cytoplasmic) or TFIIH (nuclear), implying a discrete maturation step in which MMS19-containing complex participates. This strengthens plausibility that MMS19 dysfunction would yield immature XPD. Limitation: observational cell biology—no direct phenotype/genetic evidence.

- This sentence details experimental manipulations that prevent Fe-S assembly on XPD and thereby block its incorporation into TFIIH. It provides mechanistic evidence that disrupting Fe-S assembly (the process in which MMS19 participates) functionally disables XPD integration into TFIIH. Strength: demonstrates functional consequences of impaired Fe-S assembly. Limitation: the interventions described (iron depletion, XPD mutants) are not MMS19 mutations and do not directly show MMS19 genetic causation of disease.

- This sentence localizes the MMS19-related process to the cytoplasm for CIA interaction and the nucleus for TFIIH association, supporting the stepwise model of XPD maturation. It is mechanistic and supports the idea that loss of MMS19 function could specifically block maturation before nuclear incorporation and activity of XPD. Limitation: again, cellular localization/mechanism does not equal clinical genetic evidence.

- This conclusion restates the stepwise maturation model: XPD acquires Fe-S cluster before binding TFIIH, implying that MMS19 (as a CIA targeting complex component) is upstream of TFIIH function. Mechanistic strength: reinforces causal chain from MMS19 → XPD maturation → TFIIH assembly/function. Limitation: the study does not present data on MMS19 variants in patients or phenotypic consequences in organisms that would constitute direct evidence of MMS19 as a Mendelian disease gene.


[Read Paper](https://www.semanticscholar.org/paper/2f2c74adb161b5d9e60fbe3d0bd4dc9638e991af)


### The Cytosolic Iron-Sulfur Cluster Assembly Protein MMS19 Regulates Transcriptional Gene Silencing, DNA Repair, and Flowering Time in Arabidopsis

**Authors**: Yong-Feng Han (H-index: 6), Xin‐Jian He (H-index: 35)

**Relevance**: 0.25

**Bibliometric Impact**: 0.28502222222222223


**Excerpts**:

- MMS19 is an essential component of the cytoplasmic iron-sulfur (Fe-S) cluster assembly complex in fungi and mammals; the mms19 null mutant alleles are lethal.

- Our study demonstrates that MMS19/MET18 in Arabidopsis thaliana interacts with the cytoplasmic Fe-S cluster assembly complex but is not an essential component of the complex.

- We find that MMS19 also interacts with the catalytic subunits of DNA polymerases, which have been demonstrated to be involved in transcriptional gene silencing (TGS), DNA repair, and flowering time regulation.

- Our results indicate that MMS19 has a similar biological function, suggesting a functional link between MMS19 and DNA polymerases.

- In the mms19 null mutant, the assembly of Fe-S clusters on the catalytic subunit of DNA polymerase α is reduced but not blocked, which is consistent with the viability of the mutant.

- Our study suggests that MMS19 assists the assembly of Fe-S clusters on DNA polymerases in the cytosol, thereby facilitating transcriptional gene silencing, DNA repair, and flowering time control.


**Explanations**:

- How it relates: This sentence provides cross‑species context indicating that MMS19 is essential in fungi and mammals and that null alleles are lethal. Relevance type: mechanistic/functional inference rather than direct clinical evidence. Strengths: establishes that MMS19 has an essential cellular function in mammals, which increases plausibility that perturbation could impact organismal viability or health. Limitations/caveats: lethality of null alleles in mammals does not demonstrate the existence of a Mendelian disease in humans; embryonic or perinatal lethality would reduce the chance of observing a classical Mendelian disorder in living patients. The statement is not supported here by human genetic data or clinical phenotypes, and the species statement ("mammals") is broad without specific experimental or clinical details.

- How it relates: This sentence reports the study's finding in Arabidopsis that MMS19 interacts with the cytosolic Fe‑S assembly complex but is not essential in that plant. Relevance type: mechanistic/functional evidence. Strengths: provides experimental data showing a biochemical interaction with the Fe‑S assembly machinery, supporting a conserved role in Fe‑S cluster biology. Limitations/caveats: this is plant data (Arabidopsis); the lack of essentiality in plants argues that complete loss of MMS19 function can be viable in some organisms, weakening a straightforward inference that human MMS19 loss‑of‑function would produce a Mendelian disease phenotype. Cross‑species differences limit direct applicability to humans.

- How it relates: This sentence shows MMS19 physically interacts with DNA polymerase catalytic subunits, and links DNA polymerases to TGS, DNA repair and developmental regulation. Relevance type: mechanistic. Strengths: provides a plausible pathway whereby MMS19 dysfunction could impair DNA replication/repair and epigenetic regulation—mechanisms known to underlie multiple Mendelian disorders. Limitations/caveats: interaction evidence alone does not demonstrate that human MMS19 variants cause disease; the described downstream processes are implicated in disease broadly, but no data are presented showing MMS19 variants produce pathogenic phenotypes in animals or humans.

- How it relates: This states the authors infer MMS19 has similar biological function to DNA polymerases and posits a functional link. Relevance type: mechanistic. Strengths: supports a functional role for MMS19 in processes (DNA replication/repair) that, when disrupted, can cause Mendelian disorders, thereby increasing biological plausibility. Limitations/caveats: the sentence expresses an interpretation of functional similarity rather than direct evidence of pathogenicity or of a disease association in humans.

- How it relates: This reports an experimental result in the plant mms19 null mutant: Fe‑S cluster assembly onto DNA polymerase α is reduced but not eliminated, correlating with mutant viability. Relevance type: mechanistic with experimental result. Strengths: shows partial impairment of a critical biochemical modification (Fe‑S insertion into polymerase) that could plausibly compromise polymerase function and genome maintenance. Limitations/caveats: result is in Arabidopsis and shows only reduction (not absence) of Fe‑S assembly; partial function may permit viability and milder phenotypes. There is no evidence here that similar partial deficits in mammals cause Mendelian disease, and species differences may alter phenotypic outcome.

- How it relates: This summarizes the proposed mechanism: MMS19 assists Fe‑S assembly on DNA polymerases in the cytosol, facilitating TGS, DNA repair and flowering time control. Relevance type: mechanistic. Strengths: provides a clear mechanistic hypothesis linking MMS19 to essential cellular processes (DNA repair/replication and gene silencing) that are plausible routes to Mendelian disease if disrupted. Limitations/caveats: this is a mechanistic proposal derived from plant data; it does not provide direct human genetic or clinical evidence linking MMS19 mutations to a Mendelian disorder. Also, the functional outcomes described include plant‑specific phenotype (flowering time), reducing direct translational relevance to human disease.


[Read Paper](https://www.semanticscholar.org/paper/7fcc36be8ef4eaa82d2ebd239bb9bce6eaf2607e)


### Cytosolic iron-sulfur cluster assembly

**Relevance**: 0.7

**Bibliometric Impact**: 0.0


**Excerpts**:

- A complex of four proteins termed CIA targeting complex is involved in the delivery of the NUBP1-NUBP2-bound Fe/S cluster to specific target apoproteins.

- NARFL (IOP1, Nar1 in yeast) is similar in se-quence to iron-only hydrogenases and binds two [4Fe-4S] clusters. These cofactors are essential for biogenesis but their molecular function is unknown.

- The WD40 domain protein CIAO1 (Cia1 in yeast) forms a propeller-like structure which may serve as a docking site for the other members of the CIA targeting complex.

- The role of FAM96B (YHR122W, CIA2 in yeast) and MMS19 is still unclear, but they show some specificity for the delivery of Fe/S clusters into certain target apoproteins.

- They directly interact with nu-merous Fe/S target proteins including many proteins involved in nuclear DNA integrity and maintenance such as DNA polymerases (POLD1), helicases of the DNA repair pathway (XPD or FANCJ) or of telomere maintenance (RTEL1), and primases.

- Cells with mutated MMS19 are hypersensitive to DNA damage and have elongated telomeres therefore these target proteins are relevant to various cancer-related diseases and aging.


**Explanations**:

- This sentence provides mechanistic context: MMS19 is part of a CIA targeting complex that delivers Fe/S clusters to apoproteins. Relevance to the claim: mechanistic support because participation in an essential protein-assembly pathway implies that MMS19 dysfunction could impact proteins whose defects cause Mendelian disorders. Limitation: this is mechanistic/contextual information only and does not provide direct genetic or clinical evidence linking MMS19 variants to a Mendelian disease.

- This describes an adjacent complex member (NARFL) binding Fe/S clusters and notes that cofactor function is essential but incompletely understood. Relevance: mechanistic background that the CIA pathway handles essential cofactors, strengthening plausibility that disruption (including via MMS19) could have phenotypic consequences. Limitation: indirect; describes another protein (NARFL) and states unknowns rather than providing evidence for disease association.

- This sentence further defines structural features of another CIA component (CIAO1) that may organize the complex. Relevance: mechanistic—helps explain how the CIA complex (including MMS19) might interact with client proteins; supports plausibility that MMS19 perturbation can have downstream effects. Limitation: structural/biochemical inference only; no direct disease association reported.

- This sentence states explicitly that MMS19's role is unclear but that MMS19 shows specificity for delivery of Fe/S clusters to certain apoproteins. Relevance: mechanistic evidence directly involving MMS19, indicating it has a specialized, nonredundant function in Fe/S cluster delivery. Strength: implicates MMS19 in a specific molecular function that could, if disrupted, produce cellular defects. Limitation: still mechanistic and descriptive; does not provide genetic evidence of Mendelian disease due to MMS19 mutations.

- This sentence identifies direct interactions between the CIA complex (including MMS19) and numerous Fe/S target proteins, specifically listing DNA maintenance proteins (POLD1, XPD/FANCJ, RTEL1, primases). Relevance: strong mechanistic evidence linking MMS19 to proteins whose mutations are known causes of Mendelian disorders (for example, mutations in XPD/ERCC2 and RTEL1 are established Mendelian disease genes). This increases plausibility that MMS19 dysfunction could produce Mendelian phenotypes via failure to mature these client proteins. Limitations: interaction does not equal causation—no data here showing MMS19 mutations cause the same clinical phenotypes, nor is there genetic evidence presented. The degree and context of the interactions (species, experimental system) are not given in this excerpt.

- This sentence reports cellular phenotypes: cells with mutated MMS19 are hypersensitive to DNA damage and have elongated telomeres, and it infers relevance to cancer-related diseases and aging. Relevance: functional (phenotypic) evidence that MMS19 loss affects cellular processes commonly disrupted in Mendelian DNA-repair/telomere maintenance disorders. Strength: provides phenotype-level evidence that MMS19 perturbation has biologically significant consequences. Limitations: these are cellular phenotypes (unspecified system), not clinical phenotypes in humans; no evidence is provided that germline MMS19 variants segregate with Mendelian disease in families, nor are statistics, sample sizes, or organismal/clinical data given. Thus this supports plausibility but is not direct proof of MMS19 causing a Mendelian disorder.


[Read Paper](https://www.semanticscholar.org/paper/854190977d78886c6d2f2e343bd6eeb3bc174a6f)


### Fe-S protein assembly involves bipartite client binding and conformational flexibility in the CIA targeting complex

**Authors**: S. Kassube (H-index: 5), N. Thomae (H-index: 1)

**Relevance**: 0.35

**Bibliometric Impact**: 0.144


**Excerpts**:

- The cytosolic iron sulfur (Fe-S) assembly (CIA) pathway is required for the insertion of Fe-S clusters into cytosolic and nuclear client proteins, including many DNA metabolism proteins.

- The molecular mechanisms of client protein recognition and Fe-S cluster transfer remain unknown.

- Here we report crystal structures of the CIA targeting complex and cryo-EM reconstructions of the complex bound either to the DNA replication factor primase or the DNA helicase DNA2.

- The structures, combined with biochemical, biophysical and yeast complementation assays, reveal an evolutionarily conserved, bipartite client binding mode facilitated by the structural flexibility of the MMS19 subunit.

- The primase Fe-S cluster is located ∼70  away from the catalytic cysteine in the CIA targeting complex, pointing to a conformationally dynamic mechanism of Fe-S cluster transfer.

- Altogether, our studies suggest a model for Fe-S cluster insertion and thus provide a mechanistic framework to understand the biogenesis of critical DNA replication and repair factors.


**Explanations**:

- This sentence establishes the biological role of the CIA pathway and explicitly states that it inserts Fe-S clusters into nuclear and cytosolic client proteins, including many DNA metabolism proteins. Relation to claim: mechanistic relevance. Strength: it shows MMS19 acts within a pathway that modifies proteins critical for genome maintenance, which is a plausible route by which MMS19 dysfunction could produce Mendelian disease phenotypes (e.g., DNA repair/replication defects). Limitation: this sentence names the pathway and its targets but does not mention MMS19 specifically nor provide any human genetic or phenotypic data linking MMS19 variants to Mendelian disease.

- This sentence notes a knowledge gap about how client proteins are recognized and how Fe-S clusters are transferred. Relation to claim: contextual mechanistic relevance. Strength: highlights that this study addresses mechanistic questions whose answers could inform how MMS19 dysfunction might cause disease. Limitation: it does not provide evidence for MMS19 causing disease; it only frames the mechanistic unknowns.

- This sentence describes the core experimental approaches (crystal structures and cryo-EM) and that the complexes were reconstructed with DNA replication factors (primase and DNA2). Relation to claim: mechanistic relevance. Strength: indicates the paper provides molecular-resolution structural data showing how the CIA targeting complex interacts with essential DNA replication/repair proteins — supporting plausibility that MMS19 perturbation could impair those processes. Limitation: structural/biophysical data alone do not constitute evidence of MMS19 being disease-causing in humans; no patient/variant/phenotype data are presented here.

- This sentence explicitly implicates MMS19 by name and states that the MMS19 subunit’s structural flexibility facilitates an evolutionarily conserved, bipartite client binding mode, supported by biochemical, biophysical and yeast complementation assays. Relation to claim: direct mechanistic evidence linking MMS19 function to the biogenesis of DNA metabolism proteins. Strength: shows MMS19 has a conserved, essential mechanistic role and that functional assays (including yeast complementation) were used, which strengthens the argument that MMS19 activity is important for cellular viability and function — a necessary precondition for being a Mendelian disease gene. Limitation: while yeast complementation suggests functional importance and conservation, it does not provide human genetic evidence (no human mutations, patient phenotypes, segregation, or pathogenicity data). Yeast models may not capture all human-specific effects or phenotypes; experimental conditions (in vitro, structural, yeast) limit direct inference to human Mendelian disease.

- This sentence reports a structural detail that the primase Fe-S cluster is positioned ~70 Å away from the catalytic cysteine, implying a conformationally dynamic transfer mechanism. Relation to claim: mechanistic relevance. Strength: supports a plausible, detailed mechanism for how MMS19-containing complexes deliver Fe-S clusters to essential DNA enzymes; such mechanistic detail strengthens plausibility that MMS19 defects could disrupt DNA replication/repair. Limitation: again, this is mechanistic/structural inference without accompanying human genetic or phenotypic evidence linking MMS19 mutations to Mendelian disease.

- This concluding sentence summarizes that the study proposes a model for Fe-S cluster insertion and provides a mechanistic framework for the biogenesis of critical DNA replication and repair factors. Relation to claim: mechanistic relevance and contextual support. Strength: reinforces that MMS19 is central to assembling/targeting Fe-S clusters into proteins critical for genome maintenance — a biological function where loss-of-function could plausibly cause Mendelian disorders. Limitation: it remains an inference from mechanism to disease risk; the paper does not present direct genetic or clinical evidence linking MMS19 variants to Mendelian disease.


[Read Paper](https://www.semanticscholar.org/paper/95b983bf3dcb3bf59289a72c5ff1789a78f1c42c)


## Other Reviewed Papers


### The Deciphering Developmental Disorders (DDD) study

**Why Not Relevant**: The supplied paper content is exclusively a general background on the challenges of diagnosing paediatric neurodevelopmental disorders and the value of genomic technologies. It contains no mention of the gene MMS19, no case reports or cohort data, no genetic variants, no segregation or de novo observations, and no functional or mechanistic experiments relating MMS19 to disease. Therefore there is no direct evidence (no observed pathogenic variants, no clinical associations) and no mechanistic evidence (no biochemical, cellular, or organismal studies implicating MMS19) in the provided text to support or refute the claim that MMS19 is associated with a Mendelian disease. Because the necessary data (methods, results, variant descriptions, phenotype correlations, functional assays) are absent, the content is irrelevant to the specific claim. To evaluate the claim would require excerpts such as: (a) identification of MMS19 variants in affected individuals with phenotype descriptions and segregation or de novo status; (b) statistical evidence of enrichment of damaging MMS19 variants in cases versus controls; and/or (c) functional studies demonstrating a pathogenic mechanism linking MMS19 perturbation to disease phenotypes. None of these are present in the provided text.


[Read Paper](https://www.semanticscholar.org/paper/640a3d8aeaa4d543a29b584c42eb3ccce085a17a)


### Structural insights into Fe–S protein biogenesis by the CIA targeting complex

**Why Not Relevant**: The provided paper content is focused on structural and mechanistic characterization of the CIA targeting complex and the MMS19 subunit (Cryo-EM reconstructions and Fe–S cluster transfer mechanism). It does not mention clinical phenotypes, genetic variants, Mendelian inheritance, patient data, or any disease association. While mechanistic data increases biological plausibility that perturbation of MMS19 could have pathological consequences, there is no direct or indirect genetic/clinical evidence presented here that links MMS19 to a Mendelian disorder. Because the claim concerns disease association (a clinical/genetic question) and the paper content provides only biochemical/structural mechanistic information, the relevance to the claim is very low. To support the claim one would need explicit genetic or clinical evidence (e.g., pathogenic MMS19 variants segregating with disease, case series, population genetics analyses, or clinical database annotations), none of which are present in the excerpt.


[Read Paper](https://www.semanticscholar.org/paper/6c9f49113aa230d4f8af52ba3e98c5645ef05317)


### Expanding the phenotype of a recurrent de novo variant in PACS1 causing intellectual disability

**Why Not Relevant**: The provided paper excerpt makes no mention of the gene MMS19. The manuscript describes a recurrent de novo missense variant in PACS1 (c.[607C>T], p.[Arg203Trp]) as causative for an autosomal dominant intellectual disability syndrome (MRD17), and discusses PACS1 function, expression, proximity of the variant to a CK2-binding motif, antisense zebrafish experiments, and DECIPHER microdeletion data. All described genetic, mechanistic, and clinical evidence pertains to PACS1 (and briefly mentions PACS2), not MMS19. Therefore there is neither direct genetic evidence (e.g., variants in MMS19 segregating with disease) nor mechanistic evidence (e.g., molecular function of MMS19 connected to a disease process) in this paper to support or refute the claim that MMS19 is likely associated with a Mendelian disease. Limitations: this assessment is limited to the text you provided; if other sections of the full paper (not included here) mention MMS19, they were not available for review. To evaluate the claim further, one should search primary literature, ClinVar, OMIM, and aggregated sequencing datasets for MMS19 variants with segregation or functional data linking MMS19 to Mendelian phenotypes.


[Read Paper](https://www.semanticscholar.org/paper/e460d89d850693b4fb1e3df12702be0dfb8d7f5b)


### Identifying genomic variant associated with long QT syndrome type 2 in an ecuadorian mestizo individual: a case report

**Why Not Relevant**: The paper is a case report focused on congenital Long QT Syndrome (LQTS) and reports pathogenic and uncertain variants in genes directly related to LQTS (KCNH2, AKAP9, TTN). The gene MMS19 is not mentioned anywhere in the provided content. Therefore the paper provides no direct evidence (no reported variants, no genotype-phenotype correlations, no functional or mechanistic data) relating MMS19 to any Mendelian disease. Absence of mention in a study whose scope is a specific cardiac disorder does not constitute evidence for or against MMS19 involvement in Mendelian disease generally. Additionally, the study design (single-case report centered on LQTS) and the gene panel/sequencing context limit its ability to address unrelated genes. For these reasons relevance to the claim is effectively zero.


[Read Paper](https://www.semanticscholar.org/paper/aa65df486c4f1720d86feb42ddc72ed25ab57431)


### Case report: Preimplantation genetic testing for infantile GM1 gangliosidosis

**Why Not Relevant**: The submitted paper content exclusively addresses GM1 gangliosidosis and pathogenic variants in the GLB1 gene, including clinical presentation, next-generation sequencing results identifying GLB1 variants, and reproductive/PGT-M outcomes based on those GLB1 findings. The text contains no mention of MMS19, no mechanistic discussion linking MMS19 to pathology, and no data (genetic, functional, population, or clinical) about MMS19. Therefore this paper provides no direct evidence supporting or refuting the claim that MMS19 is likely associated with at least one Mendelian disease. Absence of any mention of MMS19 in a paper focused on a different gene does not constitute evidence that MMS19 is or is not disease-associated; it only means this paper is irrelevant to evaluating the MMS19 claim. To assess MMS19's association with Mendelian disease, sources that specifically report MMS19 variants in patients, functional studies of MMS19 perturbation, or curated database entries (e.g., OMIM, ClinVar, HGMD) should be consulted.


[Read Paper](https://www.semanticscholar.org/paper/ce981bc89c565bb4d97baff441baaf8695595448)


### Autosomal Dominant Intellectual Development Disorder-6 (MRD6) Without Seizures Linked to a De Novo Mutation in the grin2b Gene Revealed by Exome Sequencing: A Case Report of a Moroccan Child

**Why Not Relevant**: The provided paper content exclusively discusses a pathogenic GRIN2B (grin2b) nonsense variant (c.3912C>G, p.Tyr1304Ter) causing Autosomal dominant intellectual development disorder-6 (MRD6) in a single patient. There is no mention of the gene MMS19, no data on MMS19 variants, and no mechanistic discussion that involves MMS19. Therefore the paper provides neither direct evidence (reported MMS19 variants segregating with disease, functional assays of MMS19, clinical descriptions tied to MMS19) nor mechanistic evidence (pathways implicating MMS19 in the phenotypes described) relevant to the claim that MMS19 is likely associated with at least one Mendelian disease.

Limitations and caveats: this conclusion is based solely on the supplied paper text. It is possible (but not indicated in the provided excerpt) that other sections, supplements, or referenced datasets from the full article could mention MMS19; those were not provided and therefore not evaluable here. Absence of evidence in this excerpt is not evidence that MMS19 is or is not disease-associated in other studies, only that this paper's content does not bear on the claim.


[Read Paper](https://www.semanticscholar.org/paper/67d756e0cc302e5d46fdda9df34403f1f2de9a5a)


### A study to evaluate the effectiveness of planned teaching programme on knowledge regarding developmental delay among mothers of preschool children admitted at selected hospital Belgaum, Karnataka

**Why Not Relevant**: This paper is primarily a small-scale pre-experimental educational intervention study assessing mothers' knowledge about developmental delay, with a literature review that briefly summarizes large-scale genetic studies identifying genes (e.g., PURA, SETBP1, ZMYND11) associated with neurodevelopmental disorders. Nowhere in the provided content is the gene MMS19 mentioned, nor are there any experimental data, case reports, variant analyses, segregation data, functional assays, or mechanistic studies related to MMS19. Therefore the paper contains no direct evidence (genetic, clinical, or mechanistic) supporting or refuting the claim that MMS19 is associated with a Mendelian disease. The included genetic-study excerpts are contextually relevant only insofar as they show that gene discovery methods can identify Mendelian causes of developmental disorders, but they do not implicate MMS19. Because the core claim concerns MMS19 specifically and this paper provides no information about that gene, the relevance to the claim is negligible.


[Read Paper](https://www.semanticscholar.org/paper/43f49246ed0fa454b8d9a7bc989fe4186ed7dd1a)


## Search Queries Used

- MMS19 mutation pathogenic variant Mendelian disease case report family segregation ClinVar OMIM

- MMS19 whole exome sequencing rare variant developmental disorder intellectual disability gene burden study DDD Decipher trio sequencing diagnostic cohort

- MMS19 iron sulfur assembly CIA complex Fe S cluster biogenesis cytosolic iron sulfur assembly DNA repair transcription helicase XPD Fe S protein maturation functional study

- MMS19 ClinGen gene disease validity review gene curation gnomAD constraint pLI loss of function tolerance missense Z population frequency no Mendelian disease association


## Usage Statistics

- Prompt Tokens: 0

- Completion Tokens: 0

- Total Tokens: 0

- Estimated Cost: $0.0458


## Bibliometric Configuration

- Use Bibliometrics: True

- Author Impact Weight: 0.4

- Citation Impact Weight: 0.4

- Venue Impact Weight: 0.2
