## Supplementary material for "MENDELSEEK: An algorithm that predicts Mendelian Genes and elucidates what makes them special": MT3.txt

# Claim: The gene MT3 is likely associated with at least one Mendelian disease.

**Status**: processed

**Overall Rating**: -1

**Explanation**:

No relevant papers were found that support or refute this claim.


## Relevant Papers


## Other Reviewed Papers


### Exome‐Wide Analyses in Paroxysmal Kinesigenic Dyskinesia Confirm TMEM151A as a Novel Causative Gene

**Why Not Relevant**: The provided paper text discusses genetic causes of paroxysmal kinesigenic dyskinesia (PKD), focusing on PRRT2 and TMEM151A. The gene symbol discussed throughout is TMEM151A (and PRRT2); the content does not mention MT3 (no occurrences of the symbol "MT3" or the name "metallothionein 3"). Key sentences such as "Our observations further emphasized TMEM151A is a bona fide PKD causative gene." and the description of identified TMEM151A variants demonstrate that the study's evidence pertains exclusively to TMEM151A as a disease-associated gene. Because MT3 is not referenced anywhere in the excerpt, there is neither direct evidence (observational/genetic association, segregation, variant data) nor mechanistic evidence (biochemical pathway, functional studies, model organisms) bearing on the claim that "MT3 is likely associated with at least one Mendelian disease." Furthermore, no sequence homology, functional analogy, or co-expression data linking TMEM151A (the gene studied) to MT3 are provided that would allow a justified inference about MT3 from the TMEM151A results. Therefore the paper content is not relevant to evaluating the claim about MT3.


[Read Paper](https://www.semanticscholar.org/paper/3b2758a6a23714ef7aea60c7f0cad73c823d2788)


### Biallelic loss-of-function in NRAP is a cause of recessive dilated cardiomyopathy.

**Why Not Relevant**: This paper investigates the role of the NRAP gene in autosomal recessive dilated cardiomyopathy (DCM). The entire provided content discusses NRAP variants, their enrichment in DCM cases, family segregation, and estimated contribution of biallelic NRAP loss-of-function variants to DCM. There is no mention of the gene MT3, no experimental data, segregation, population-frequency analysis, pathogenic-variant descriptions, or mechanistic discussion related to MT3. Therefore the paper provides no direct evidence (e.g., cases, co-segregation, variant pathogenicity) nor mechanistic evidence (e.g., protein function, pathways, animal models) that would support or refute the claim that MT3 is associated with a Mendelian disease. Limitations: absence of MT3 in this single paper does not disprove any MT3–disease association reported elsewhere; it only means this paper is not informative for the claim. If you want, I can perform a literature search or analyze other papers specifically about MT3 for evidence of Mendelian disease association.


[Read Paper](https://www.semanticscholar.org/paper/74bbb02db2eb68ef9d9eeef09e62e2cd907c7428)


### Case report: Novel NUS1 variant in a Chinese patient with tremors and intellectual disability

**Why Not Relevant**: The paper reports a case and genetic analysis implicating NUS1 (nuclear undecaprenyl pyrophosphate synthase 1) variants in a congenital glycosylation disorder with neurological features. The text contains no mention of the gene MT3, no MT3 variants, and no mechanistic discussion involving MT3. Therefore there is neither direct evidence (e.g., MT3 pathogenic variants in affected individuals) nor mechanistic evidence (e.g., shared pathways, functional studies linking MT3 to the reported phenotype) supporting the claim that MT3 is associated with a Mendelian disease. Because the study is specific to NUS1 and does not address MT3 in any way, its relevance to the MT3–Mendelian disease claim is effectively zero. Limitations that reinforce non-relevance: single-gene focus (NUS1), single-case report context, and no cross-gene comparative analyses or databases queried for MT3.


[Read Paper](https://www.semanticscholar.org/paper/875bea84e261d364738e752da6cd1ef9bbc26e60)


### Correspondence on “Clinical, neuropathological, and genetic characterization of STUB1 variants in cerebellar ataxias: a frequent cause of predominant cognitive impairment” by Roux et al.

**Why Not Relevant**: The provided paper content discusses heterozygous variants in the gene STUB1 in an ataxia patient cohort and explicitly remarks that the functional relevance of certain STUB1 variants is unclear. It does not mention the gene MT3, does not present data or mechanisms related to MT3, and does not provide any evidence (direct or mechanistic) linking MT3 to a Mendelian disease. Therefore the content is not relevant to the claim that MT3 is likely associated with at least one Mendelian disease.


[Read Paper](https://www.semanticscholar.org/paper/d133829fd8bd83f58a0e216c6bf9a5c26eaeadd6)


### Screening Mutations of the Monogenic Syndromic High Myopia by Whole Exome Sequencing From MAGIC Project

**Why Not Relevant**: This paper provides no direct evidence for or against the claim that MT3 is associated with at least one Mendelian disease. The provided text does not mention the gene MT3 anywhere; it reports findings for a set of msHM-related genes (examples listed) and summarizes diagnostic yield in a cohort with syndromic high myopia. Limitations that prevent using this study as evidence for MT3 association: (1) the study is restricted to monogenic syndromic high myopia, so genes causing Mendelian diseases without an msHM phenotype would not be detected or reported; (2) the excerpt does not include the full list of the 298 genes in the targeted panel nor the full list of the 84 implicated genes, so it is unknown whether MT3 was assayed but not found, assayed and found but not reported in the excerpt, or not included in the panel at all; (3) absence of mention in this paper is not proof that MT3 is not associated with any Mendelian disease. Therefore the study is not relevant evidence to support or refute the claim about MT3.


[Read Paper](https://www.semanticscholar.org/paper/339423d81112e62718cf1ec21a06909ee07f9afe)


### Whole-exome sequencing analysis in a case of primary congenital glaucoma due to the partial uniparental isodisomy

**Why Not Relevant**: The provided paper content exclusively discusses a pathogenic variant in the CYP1B1 gene identified in a single case of primary congenital glaucoma (including trio WES and Sanger confirmation). There is no mention of the gene MT3, no reported variants in MT3, and no mechanistic discussion linking MT3 to any Mendelian phenotype. While the authors note 'secondary/incidental findings' and the use of whole-exome sequencing (which in principle could detect MT3 variants), the content does not present any data, observations, or discussion about MT3. Therefore this paper provides no direct evidence that MT3 is associated with a Mendelian disease, nor mechanistic evidence supporting such an association. Limitations: the absence of any MT3 information in the excerpt means we cannot rule out that the full paper or raw WES data might contain unreported MT3 findings, but based on the supplied content there is nothing to support the claim.


[Read Paper](https://www.semanticscholar.org/paper/cbda5e9c6201514cf19b5b27e453c30a53060acd)


### Case report: A novel FBXW7 gene variant causes global developmental delay

**Why Not Relevant**: This paper focuses entirely on a case of neurodevelopmental disorder caused by a de novo pathogenic variant in FBXW7; MT3 is not mentioned anywhere in the provided content. All presented objective, methods, results, and conclusion statements concern FBXW7 and the patient's phenotype. Because the paper does not report any data, observations, or mechanistic discussion related to the MT3 gene, it provides no direct evidence (variant-disease association, segregation, case reports) nor mechanistic evidence (functional studies, pathway links) that would support or refute the claim that MT3 is associated with at least one Mendelian disease. Therefore the relevance to the MT3 claim is essentially zero. Note: absence of MT3 in this paper is not evidence that MT3 is or is not disease-associated; it only means this paper cannot inform that claim.


[Read Paper](https://www.semanticscholar.org/paper/bb0b7ce4ae459771c428a6c9d1dbc93aecc6bcdc)


### Understanding Genetic Epidemiology: The Potential Benefits and Challenges of Genetics for Improving Human Health

**Why Not Relevant**: This paper is a high-level review of the history, methods, and broad accomplishments of genetic epidemiology and does not mention the gene MT3 (metallothionein 3) or any data linking MT3 to Mendelian disease. The text discusses general methods used to discover genes for monogenic disorders (e.g., linkage analysis) and gives examples of genes and disorders (Tay-Sachs, Huntington’s, cystic fibrosis, PKU, VKORC1, CYP2C9, CYP2C19, FTO, GPT, ACP) but provides no direct genetic, clinical, or functional evidence concerning MT3. Therefore it contains neither direct evidence (e.g., pathogenic MT3 variants segregating with a Mendelian phenotype, case reports, cohort studies, ClinVar/OMIM annotations) nor mechanistic evidence (e.g., functional data, pathways, or molecular mechanisms that would plausibly link MT3 dysfunction to a Mendelian disorder) relevant to the claim.

Why relevance is effectively zero:
- No mention of MT3 by name anywhere in the provided text, so there are no verbatim excerpts that support or refute the claim. 
- The paper’s examples of monogenic disease gene discoveries and discussions of methods do not provide gene-specific evidence for MT3. 
- Absence of MT3 in a general review is not evidence that MT3 is or is not associated with a Mendelian disease; it only means this paper does not address the question.

What would constitute relevant evidence (and is absent here):
- Direct evidence: reports of pathogenic MT3 variants in multiple affected individuals/families with consistent Mendelian inheritance and segregation, statistical association of rare MT3 variants with a Mendelian phenotype, or authoritative database entries (OMIM, ClinVar) linking MT3 to a named Mendelian disorder. 
- Mechanistic evidence: experimental demonstrations that MT3 loss- or gain-of-function produces the disease phenotype in model systems, or clear molecular pathways showing how MT3 perturbation leads to the disease symptoms.

Given these points, this paper cannot be used to support or refute the claim that MT3 is likely associated with at least one Mendelian disease. To evaluate the claim, consult gene-focused resources (OMIM, ClinVar, HGMD), primary genetic studies (family linkage/exome/genome sequencing), or functional studies directly examining MT3 variants and phenotypes.


[Read Paper](https://www.semanticscholar.org/paper/1ea9ce6d8741214d75e3aa3f5dae45e0e6129c75)


### A novel mutation in the ATP7B gene causing hepatolenticular degeneration in a Chinese family: A case report

**Why Not Relevant**: The paper exclusively discusses ATP7B and Wilson disease; it identifies and characterizes a novel ATP7B variant and provides mechanistic and clinical context for ATP7B-related hepatolenticular degeneration. There is no mention of the gene MT3, no data (genetic, functional, segregation, or epidemiologic) regarding MT3, and no discussion connecting metallothioneins or MT3 to the disease phenotype. Although whole-exome sequencing was performed (which could, in principle, detect MT3 variants), the authors do not report any findings related to MT3. Therefore this paper provides no direct or mechanistic evidence supporting or refuting the claim that MT3 is associated with a Mendelian disease.


[Read Paper](https://www.semanticscholar.org/paper/9d4b911621bf3cc38aca3619e915780869df23d8)


## Search Queries Used

- MT3 mutation pathogenic variant Mendelian disease case report whole exome sequencing trio monogenic

- metallothionein 3 MT3 gene genetic disorder monogenic rare disease OMIM ClinVar HGMD review

- metallothionein 3 MT3 function metal homeostasis zinc copper neuronal growth inhibitory factor neurodegeneration knockout mouse phenotype loss of function study

- MT3 rare variants exome sequencing cohort burden test family segregation linkage study autosomal dominant autosomal recessive penetrance expressivity population frequency gnomAD


## Bibliometric Configuration

- Use Bibliometrics: True

- Author Impact Weight: 0.4

- Citation Impact Weight: 0.4

- Venue Impact Weight: 0.2
