## Supplementary material for "MENDELSEEK: An algorithm that predicts Mendelian Genes and elucidates what makes them special": MTCP1.txt

# Claim: The gene MTCP1 is likely associated with at least one Mendelian disease.

**Status**: processed

**Overall Rating**: -1

**Explanation**:

No relevant papers were found that support or refute this claim.


## Relevant Papers


## Other Reviewed Papers


### Whole-Exome Sequencing in Adults With Chronic Kidney Disease

**Why Not Relevant**: The provided excerpt contains no mention of the gene MTCP1 (no exact string match and no contextual discussion of MTCP1 or known synonyms). The manuscript fragment describes a whole-exome sequencing study of adults with chronic kidney disease, methods for sequencing and variant interpretation, and the construction/usage of a 287-gene nephropathy OMIM list and ACMG actionable gene review, but it does not report any results or discussion that name MTCP1 or link it to any Mendelian disorder. Therefore, there is no direct evidence in this content that MTCP1 is associated with a Mendelian disease, nor is there mechanistic discussion implicating MTCP1 in disease pathways.

Limitations and caveats:
- The assessment is limited to the text you provided. The full paper (including Results, Discussion, and Supplementary Tables 1–4 and Supplement Table 3 mentioned in the excerpt) might contain gene lists, variant tables, or statements that mention MTCP1. Those portions were not included here, so absence of MTCP1 in this excerpt is not definitive evidence that the full paper lacks any information about MTCP1.
- The excerpt focuses on nephropathy and hypertension genes; MTCP1 may not be a kidney-related gene and thus might not be expected to appear in the nephropathy-focused gene list used by this study.

Recommended next steps if you want a definitive answer from this paper:
1) Inspect the full text, especially the Results and Discussion sections and all Supplementary Tables (particularly the nephropathy gene list in Supplement Table 3 and variant lists), for any mention of MTCP1.
2) Search the paper PDF (and its supplements) for the exact string "MTCP1" and common synonyms/aliases.
3) If the paper does not mention MTCP1, consult primary databases (OMIM, ClinVar, HGMD) and the literature for MTCP1 to evaluate any reported Mendelian associations outside this paper.


[Read Paper](https://www.semanticscholar.org/paper/a29bd57964f68ac5f8e1abcafe1112ba488f53f5)


### The simplest explanation does not have to be preferred: co-occurrence of pathogenic variants in cancer-predisposing genes

**Why Not Relevant**: The supplied paper content does not provide any evidence for or against the claim that MTCP1 is likely associated with at least one Mendelian disease. The single sentence refers only to reviewing the literature and UK 10,000 Genomes Project data concerning MINAS, cancer phenotypes, and cancer susceptibility genes (CSGs) in participants with tumor predisposition syndromes or multiple primary tumors. It does not mention the gene MTCP1, any specific gene-level findings, variant-level data, case reports, segregation analyses, population frequencies, functional studies, or mechanistic pathways involving MTCP1. Therefore there is no direct evidence (no reported association, statistical results, or clinical cases implicating MTCP1) and no mechanistic evidence (no molecular function, pathway perturbation, or experimental validation linking MTCP1 to disease) in the provided content.

Additional limitations that prevent drawing any inference about MTCP1 from the provided text:
- The excerpt is a single, general methodological/introductory sentence rather than results or data: no methods, no sample counts, no statistical analyses, and no specific genes named.
- MINAS and CSG are generically referenced; without specific gene names or variants, one cannot map those statements to MTCP1.
- No experimental conditions, patient phenotypes linked to MTCP1, or functional assays are described, so mechanistic plausibility cannot be assessed.
- The lack of verbatim mention of MTCP1 means there is no way to extract supporting or refuting quotations per the requested guidelines.

Recommendation: to evaluate the claim about MTCP1 being associated with a Mendelian disease, consult primary sources that specifically report MTCP1 variants or disease phenotypes (for example: OMIM, ClinVar, HGMD, gene-specific case reports, segregation studies, functional assays, or cohort sequencing studies that list MTCP1 results).


[Read Paper](https://www.semanticscholar.org/paper/a1372a664bd2c33a0bda638975a8b26d112457fc)


### Role of gdf11 in retinal neurogenesis

**Why Not Relevant**: I found no direct or mechanistic evidence in the provided paper content relating to the gene MTCP1 or to any Mendelian disease associated with MTCP1. The provided text consists of meeting abstracts and short reports discussing other genes (for example, FGFR1, E2F1, GDF11, TCL1, AKT isoforms) and developmental processes in mouse models. The word "MTCP1" does not appear in the text, nor are there discussions of human genetic disease associations, Mendelian inheritance, pathogenic variants, patient phenotypes, linkage/segregation studies, loss-of-function/gain-of-function mutations, or functional assays relevant to MTCP1. Examples of the genes/topics actually present include: "Fgfr1", "E2F1", "Gdf11", "TCL1", and "AKT1/AKT2/AKT3". Because the paper content neither mentions MTCP1 nor provides experimental or clinical data that could be extrapolated mechanistically to MTCP1, it cannot support or refute the claim that "MTCP1 is likely associated with at least one Mendelian disease." Additional reasons why this content is not informative for the claim: (1) the material is primarily developmental biology abstracts rather than human genetic studies; (2) there are no genetic linkage, variant pathogenicity, case reports, or animal models involving MTCP1 presented; and (3) mechanistic pathways described (e.g., FGFR1 in radial glia proliferation, GDF11 in retinal neurogenesis, TCL1 interactions with AKT) are gene-specific and do not include MTCP1 as a component. To evaluate the claim about MTCP1, one would need literature that explicitly reports MTCP1 variants in affected individuals, segregation with disease, functional studies of MTCP1 mutations, or well-supported mechanistic links between MTCP1 dysfunction and a Mendelian phenotype. The provided content does not contain such information.


[Read Paper](https://www.semanticscholar.org/paper/fc915b0500e7436128291f85c160e278a03191b9)


## Search Queries Used

- MTCP1 pathogenic variant germline mutation Mendelian monogenic hereditary disease OMIM ClinVar ClinGen case report

- MTCP1 translocation T cell prolymphocytic leukemia somatic oncogene TCL1 family germline predisposition inherited cancer risk

- MTCP1 function AKT interaction TCL1 family proto oncogene signaling pathway functional study mouse knockout zebrafish loss of function developmental phenotype

- MTCP1 monogenic disorder X chromosome X linked germline variant rare disease exome sequencing diagnostic case series systematic review meta analysis Human Gene Mutation Database HGMD gnomAD population frequency


## Bibliometric Configuration

- Use Bibliometrics: True

- Author Impact Weight: 0.4

- Citation Impact Weight: 0.4

- Venue Impact Weight: 0.2
