## Supplementary material for "MENDELSEEK: An algorithm that predicts Mendelian Genes and elucidates what makes them special": MX1.txt

# Claim: The gene MX1 is likely associated with at least one Mendelian disease.

**Status**: processed

**Overall Rating**: -1

**Explanation**:

No relevant papers were found that support or refute this claim.


## Relevant Papers


## Other Reviewed Papers


### Loss of Function Mutation in ELF4 Causes Autoinflammatory and Immunodeficiency Disease in Human

**Why Not Relevant**: The provided paper content contains no mention of the gene MX1, no genetic testing or variant data, and no mechanistic data linking any gene to the described phenotype. The single-sentence summary states: "A pediatric patient suffering from recurrent viral and bacterial respiratory infection, refractory oral ulcer, constipation, and arthritis is described, and the second report of DEX is presented." This describes a clinical case and the reporting of a disorder abbreviated as DEX, but it does not: (1) name MX1 or any other gene, (2) report a sequence variant, genotype, segregation, or inheritance pattern, (3) provide functional assays or molecular mechanism implicating MX1, or (4) present statistical or family-based evidence consistent with a Mendelian gene-disease relationship. Because association of a gene with a Mendelian disease requires genetic and/or genetic-mechanistic evidence (e.g., pathogenic variant(s) segregating with disease, de novo occurrence, loss/gain-of-function demonstrated in vitro or in vivo, or recurrent variants in multiple unrelated affected individuals), the absence of any such information in the supplied content makes this paper irrelevant to the claim that "MX1 is likely associated with at least one Mendelian disease." Additional limitations that further reduce relevance: the report appears to be a single-case clinical description (insufficient sample size for establishing Mendelian association), and the abbreviation DEX is not expanded or linked to a gene in the provided text. Therefore, based on the supplied content alone, this paper provides no direct or mechanistic evidence for or against the claim.


[Read Paper](https://www.semanticscholar.org/paper/168cdf9d79bf512e3e598508966ba1eeff2e7177)


## Search Queries Used

- MX1 myxovirus resistance 1 Mendelian disease OMIM ClinVar pathogenic variant

- MX1 mutation human case report whole exome sequencing monogenic severe viral infection familial influenza

- MX1 MxA interferon induced GTPase loss of function knockout mouse phenotype viral susceptibility innate immunity human relevance

- MX1 variant frequency gnomAD ClinVar population genetics rare variant burden penetrance polymorphism confounding


## Bibliometric Configuration

- Use Bibliometrics: True

- Author Impact Weight: 0.4

- Citation Impact Weight: 0.4

- Venue Impact Weight: 0.2
