## Supplementary material for "MENDELSEEK: An algorithm that predicts Mendelian Genes and elucidates what makes them special": MX2.txt

# Claim: The gene MX2 is likely associated with at least one Mendelian disease.

**Status**: processed

**Overall Rating**: -1

**Explanation**:

No relevant papers were found that support or refute this claim.


## Relevant Papers


## Other Reviewed Papers


### Polygenic risk scores for low-density lipoprotein cholesterol and familial hypercholesterolemia

**Why Not Relevant**: The provided paper content contains no mention of the gene MX2 or of any specific Mendelian disease. The single sentence provided reads: "In conclusion, mutation negative CDFH patients demonstrated significantly higher PRSLDLC than controls, and PR SLDLC may have little additional effect on LDL-C and CAD among FH patients, while genome-wide association study summary statistic from the BioBank Japan Project could not detect any association between PRS and CAD." This sentence discusses (1) polygenic risk scores (PRS) and PRSLDLC, (2) LDL-C and coronary artery disease (CAD) in familial hypercholesterolemia (FH) patients, and (3) a GWAS summary statistic from the BioBank Japan Project. None of these elements address MX2, rare/monogenic variant analyses, segregation with disease, functional consequences of MX2 variants, or any Mendelian phenotype attribution. 

Why this is not relevant to the claim that "MX2 is likely associated with at least one Mendelian disease":
- No direct evidence: There are no data, results, or statements in the excerpt relating to MX2, its variants, or any disease association. Therefore there is no direct support or refutation of the claim. 
- No mechanistic evidence: The excerpt focuses on PRS (polygenic effects) and GWAS summary statistics, which examine common-variant contributions to complex traits, not rare high-penetrance variants or gene-level mechanisms characteristic of Mendelian disorders. 
- Study design mismatch: GWAS/PRS approaches are not designed to identify rare, highly penetrant Mendelian gene–disease relationships; absence of a PRS association with CAD does not inform whether a specific gene (MX2) causes a Mendelian disease. 
- Absence-of-evidence caveat: The lack of mention of MX2 in this excerpt cannot be interpreted as evidence that MX2 is not associated with a Mendelian disease. It simply indicates the paper (or at least the provided excerpt) did not examine that question. 

Additional limitations that prevent using this excerpt to evaluate the MX2-Mendelian-disease claim:
- The excerpt is a single concluding sentence without methods, sample sizes, variant-level analyses, or gene lists. 
- The focus is on polygenic risk and common-variant GWAS summary statistics (BioBank Japan), which do not capture the rare variant, segregation, or functional data needed to establish Mendelian gene–disease associations. 

Conclusion: Relevance to the claim is effectively zero based on the provided content. To assess whether MX2 is likely associated with a Mendelian disease would require different evidence types (e.g., rare-variant sequencing in pedigrees, segregation analysis, case reports of pathogenic MX2 variants, functional studies showing loss/gain-of-function consistent with disease), none of which are present here.


[Read Paper](https://www.semanticscholar.org/paper/6d972a1c19dc1e3425eeec805fea36b152d12c38)


### Tumoural soft tissue calcification in Down syndrome: association with heterozygous germline SAMD9 mutation and hyperactive type I interferon signaling.

**Why Not Relevant**: The paper is a case report of a person with trisomy 21 and mixed connective tissue disease describing upregulation of interferon signalling and IFN-stimulated genes, and noting upregulation of an IFN-induced MX protein in affected tissue. However, it contains no genetic data implicating MX2 (no reported MX2 variants, no segregation/penetrance information, no association statistics, no functional studies of MX2 variants), and does not claim or demonstrate that MX2 causes a Mendelian disease. The evidence is limited to expression and pathway activation in an acquired (trisomy 21) context; this supports mechanistic plausibility that MX-family proteins participate in pathology but does not provide direct or genetic mechanistic evidence that MX2 is a Mendelian disease gene. Therefore the paper provides negligible direct evidence for the claim.


[Read Paper](https://www.semanticscholar.org/paper/c66134ab91385fac6ab93328ad6722ebe236de51)


### Clonal Hematopoiesis as a pitfall in germline variant interpretation in the context of Mendelian disorders.

**Why Not Relevant**: The provided paper content describes a general finding that 58 genes show both somatic involvement in age-related clonal hematopoiesis and germline associations with Mendelian disorders, and it explains the study methods and interpretation pitfalls. However, the excerpted content does not include a list of the 58 genes or any mention of MX2 by name. Therefore there is no direct evidence in the supplied text that MX2 is associated with any Mendelian disease. The statements are mechanistically and contextually relevant (they show that such dual associations exist and could plausibly include MX2), but without the specific gene list, variant-level data, or mention of MX2, one cannot conclude from this paper content that MX2 is likely associated with a Mendelian disease. For that conclusion one would need the authors' gene list, supplementary tables, or explicit statements naming MX2.


[Read Paper](https://www.semanticscholar.org/paper/c9e08cffcbac82013969d352ac230110cb51287c)


### Abstract TMP1: Multi-omic Biomarker Development In A Mendelian Neurovascular Disease, Cavernous Angioma

**Why Not Relevant**: The supplied paper content does not provide direct or mechanistic evidence linking the gene MX2 to any Mendelian disease. The text focuses on Cavernous Angioma (familial cases attributed to three CCM genes) and on circulating biomarkers (plasma microRNAs, metabolites, and proteins) and their diagnostic models. There is no mention of MX2, no genetic association testing of MX2, no description of MX2 variants, nor any functional/mechanistic experiments involving MX2. Therefore this paper cannot be used to support the claim that MX2 is likely associated with at least one Mendelian disease. Note that absence of evidence in this paper is not evidence of absence overall: determining whether MX2 is associated with a Mendelian disease would require genetic studies, case reports, linkage/segregation analyses, or functional data not present here.


[Read Paper](https://www.semanticscholar.org/paper/b813d23fbb16d1dc4f7d24379b599d1675b5e77f)


### Contribution of variants across the allelic frequency spectrum to cystic kidney disease

**Why Not Relevant**: The paper focuses exclusively on genetic contributors to cystic kidney disease (CyKD). The gene MX2 is not mentioned anywhere in the provided content, and no direct or mechanistic evidence linking MX2 to a Mendelian disease is presented. The study used whole-genome sequencing and large-scale, hypothesis-free analyses across variant classes and also performed a large meta-analysis, but its negative findings pertain to CyKD specifically. Therefore this paper provides only indirect/negative evidence regarding MX2 (i.e., MX2 was not identified as associated with CyKD in this cohort). Absence of evidence in this phenotype-focused study does not rule out the possibility that MX2 is associated with a different Mendelian disease or with Mendelian disease mechanisms not well captured by the study (for example, ultra-rare private variants, family-specific segregation, non-CyKD phenotypes, or variant types with limited power in the analyses). Given MX2 is not reported or discussed in the paper, the content is not directly relevant to the claim that "MX2 is likely associated with at least one Mendelian disease."


[Read Paper](https://www.semanticscholar.org/paper/ac802a1b5b377860f361a8b5d89ff182d78b02b3)


## Search Queries Used

- MX2 mutation germline familial monogenic Mendelian disease association

- MX2 MxB myxovirus resistance 2 variants missense nonsense frameshift splice site pathogenic germline

- MX2 antiviral GTPase HIV restriction capsid interaction nucleocytoplasmic transport interferon stimulated gene functional study human immune deficiency susceptibility

- MX2 gnomAD loss of function constraint pLI population frequency ClinVar pathogenic variant OMIM exome sequencing


## Bibliometric Configuration

- Use Bibliometrics: True

- Author Impact Weight: 0.4

- Citation Impact Weight: 0.4

- Venue Impact Weight: 0.2
