## Supplementary material for "MENDELSEEK: An algorithm that predicts Mendelian Genes and elucidates what makes them special": MYCBP2.txt

# Claim: The gene MYCBP2 is likely associated with at least one Mendelian disease.

**Status**: processed

**Overall Rating**: -1

**Explanation**:

No relevant papers were found that support or refute this claim.


## Relevant Papers


## Other Reviewed Papers


### Extending the phenotype associated with the CSNK2A1‐related Okur–Chung syndrome—A clinical study of 11 individuals

**Why Not Relevant**: The provided paper text discusses variants in the CSNK2A1 gene (encoding the Protein Kinase CK2 alpha subunit) and their relationship to a specific neurodevelopmental syndrome (Okur–Chung neurodevelopmental syndrome). The claim concerns the gene MYCBP2, but the paper contains no mention of MYCBP2, no variant data for MYCBP2, and no mechanistic discussion linking CSNK2A1 to MYCBP2. Therefore it provides neither direct evidence (e.g., pathogenic MYCBP2 variants segregating with disease, de novo MYCBP2 variants, or case reports) nor mechanistic evidence (e.g., biochemical pathways implicating MYCBP2 in disease) relevant to the claim. Because the study is explicitly focused on CSNK2A1, its findings cannot be used to support or refute an association between MYCBP2 and any Mendelian disease. Limitations that lead to this conclusion: the paper's scope is limited to a different gene (CSNK2A1); absence of mention of MYCBP2 in the text means no data, statistics, or mechanistic descriptions involving MYCBP2 are available to evaluate the claim; and no indirect links or cross-references to MYCBP2 are provided that could permit cautious inference.


[Read Paper](https://www.semanticscholar.org/paper/5c58295d43e575146836115eee7d7d7ea1b0f9dd)


### DOES A DIAGNOSIS OF ASD HELP US TO HELP A PERSON WITH INTELLECTUAL DISABILITIES ?

**Why Not Relevant**: The provided paper content does not mention MYCBP2 anywhere. The genetic material in this content concerns KCND1 (and related KCND2/KCND3) as candidate genes for X-linked intellectual disability/autism spectrum disorder, and other parts concern caregiving and ASD screening instruments. No direct genetic association, linkage, mutation, functional data, case reports, or mechanistic discussion regarding MYCBP2 are present. Therefore this paper provides no direct or mechanistic evidence supporting or refuting the claim that MYCBP2 is likely associated with at least one Mendelian disease. Absence of evidence in this paper does not disprove the claim overall; it simply means this source is not informative for MYCBP2.


[Read Paper](https://www.semanticscholar.org/paper/6d51deaa491eed78a7b02217a5218e42aa1bba39)


### Identifying Rare Genetic Variation In Obsessive-Compulsive Disorder

**Why Not Relevant**: The provided paper text does not mention MYCBP2, nor does it present any variant-, gene-, or disease-level data that could be mapped to MYCBP2. The manuscript excerpt focuses on rates of post-zygotic single-nucleotide variants (PZVs) and copy-number variants (CNVs) in obsessive-compulsive disorder (OCD) probands versus controls, reporting summary statistics (e.g., rates and p-values) and a higher rate of de novo deletions in cases versus controls. However, no gene names, lists of implicated loci, variant-level results, or descriptions of Mendelian disease associations are provided. Therefore it contains neither direct evidence (e.g., pathogenic variants in MYCBP2, segregation with disease, case reports) nor mechanistic evidence (e.g., functional studies, pathway links) relevant to the claim that MYCBP2 is likely associated with at least one Mendelian disease.

Limitations and implications of this absence:
- Absence of evidence in this paper is not evidence that MYCBP2 is or is not associated with Mendelian disease; it only means this paper does not address that claim. 
- The study design (exome sequencing of trios for mosaic SNVs and CNV burden analyses in OCD) could in principle generate gene-level findings, but none are reported in the provided text, so no inference about MYCBP2 can be drawn from these excerpts. 
- To evaluate the claim about MYCBP2 one would need variant-level data and clinical/segregation evidence (e.g., reported pathogenic variants in MYCBP2 in affected individuals, replication in independent families, functional validation) or authoritative clinical genetics resources (e.g., OMIM, ClinVar, gene-disease validity curation).


[Read Paper](https://www.semanticscholar.org/paper/6836f39aa067f957e1211be9fdfcd02b590e97cc)


## Search Queries Used

- MYCBP2 MYC binding protein 2 PAM pathogenic variant de novo variant Mendelian disease monogenic autosomal dominant autosomal recessive intellectual disability epileptic encephalopathy neurodevelopmental disorder case report

- MYCBP2 ClinVar OMIM ClinGen Decipher gnomAD pLI loss of function intolerance haploinsufficiency gene disease validity

- MYCBP2 PAM E3 ubiquitin ligase ubiquitination neuronal development axon degeneration synaptic growth Highwire RPM 1 Drosophila mouse knockout phenotype neurodevelopment

- exome sequencing de novo MYCBP2 developmental disorders DDD Decipher autism intellectual disability trio study rare variant burden gene based association


## Bibliometric Configuration

- Use Bibliometrics: True

- Author Impact Weight: 0.4

- Citation Impact Weight: 0.4

- Venue Impact Weight: 0.2
