## Supplementary material for "MENDELSEEK: An algorithm that predicts Mendelian Genes and elucidates what makes them special": NCKIPSD.txt

# Claim: The gene NCKIPSD is likely associated with at least one Mendelian disease.

**Status**: processed

**Overall Rating**: 0

**Explanation**:

No relevant papers were found for this claim.


## Relevant Papers


## Search Queries Used

- NCKIPSD mutation Mendelian disease case report patient pathogenic variant NCK interacting protein with SH3 domain

- NCKIPSD ClinVar OMIM HGMD DECIPHER pathogenic variant monogenic disorder NCK interacting protein with SH3 domain

- NCKIPSD protein function NCK interacting protein with SH3 domain actin cytoskeleton signaling mouse knockout zebrafish knockdown model

- NCKIPSD loss of function intolerance pLI gnomAD constraint haploinsufficiency human gene damage index Mendelian disease association
