## Supplementary material for "MENDELSEEK: An algorithm that predicts Mendelian Genes and elucidates what makes them special": NEFM.txt

# Claim: The gene NEFM is likely associated with at least one Mendelian disease.

**Status**: processed

**Overall Rating**: 3

**Explanation**:

Supporting evidence.
The two provided papers establish neurofilament proteins as biologically plausible contributors to Mendelian neurological disease. The first paper reports that mutations in neurofilaments have been linked to some forms of Charcot–Marie–Tooth (CMT) disease, and demonstrates experimentally that CMT-associated neurofilament mutants disrupt filament assembly, axonal transport, and mitochondrial localization in neurons—mechanisms plausibly sufficient to cause a genetic neuropathy. The second paper explicitly considered both NF-L (NEFL) and NF-M (NEFM) as positional candidate genes in a CMT locus, and highlights neurofilament proteins as likely targets in axonal pathology and as logical candidate genes for other CMT loci. Together, these findings provide mechanistic and positional plausibility that perturbation of neurofilament genes can produce Mendelian neuropathies, and they identify NEFM as a reasonable candidate for such a role.

Caveats or contradictory evidence.
Neither excerpt provides direct evidence that NEFM (the gene encoding neurofilament medium) actually harbors pathogenic variants causing a Mendelian disease. In the second paper, the authors screened both NF-L and NF-M coding regions but report identification of a disease-associated missense mutation only in NF-L (NEFL); no NEFM mutation is reported in the provided excerpts. Thus the data show that at least one neurofilament gene (NEFL) is causally linked to a Mendelian form of CMT, but do not demonstrate that NEFM is. The cited papers therefore supply indirect or candidate-level support at best; they do not document a confirmed pathogenic NEFM variant or segregation evidence tying NEFM to a Mendelian phenotype. Additional limitations include the possibility that NEFM involvement could be noncoding, regulatory, or rare and missed by the screens described, but that is speculative and not shown by the excerpts.

Analysis of potential underlying mechanisms, deeper relationships, or second-order implications.
Neurofilaments are multisubunit intermediate filaments composed of light (NEFL), medium (NEFM), and heavy (NEFH) subunits; proper assembly requires stoichiometric interactions among these proteins. A pathogenic variant in one subunit can disrupt filament assembly or axonal transport, producing a dominant-negative effect or a loss-of-function phenotype that manifests as a neuropathy. The observation that NEFL mutations cause CMT lends biological credence to the idea that NEFM mutations could have similar effects, since NEFM participates directly in the same structural complexes and cellular processes. Moreover, functional perturbation of NEFM might have variable phenotypic manifestations depending on mutation type (dominant-negative vs recessive loss) and cellular context. However, mechanistic plausibility alone does not establish causation; empirical genetic evidence (segregation, recurrence, functional corroboration for specific NEFM variants) would be required to confirm NEFM as a Mendelian disease gene.

Explanation of rating.
On balance, the provided evidence supports NEFM as a biologically plausible candidate for Mendelian neurological disease (especially CMT), but it does not include direct genetic proof of NEFM causing a Mendelian disorder. The strongest empirical result shown is a NEFL mutation causing CMT; NEFM is discussed as a candidate and was screened, but no NEFM causal variant is reported in the excerpts. Given plausible mechanism and positional candidacy but absence of documented pathogenic NEFM variants in the provided material, the appropriate assessment is that evidence is mixed: there is substantive indirect support but no direct confirmation in these papers. Therefore the claim that NEFM is likely associated with at least one Mendelian disease is supported to a degree (plausible and worthy of further study) but not established by the supplied excerpts. The best fit rating is "Mixed Evidence."


**Final Reasoning**:

I reviewed the logical chain linking NEFM to Mendelian disease in the provided excerpts. The arguments in favor are (1) neurofilament mutations cause CMT and disrupt axonal biology; (2) NEFM (NF-M) is a positional and biological candidate in CMT loci and was explicitly considered by investigators. These points justify considering NEFM as a plausible Mendelian disease gene. The counterpoint is that the concrete genetic finding reported in the excerpts is in NEFL (NF-L), not NEFM, and no NEFM pathogenic variant is documented. That makes the support indirect rather than confirmatory. No stronger contradictory evidence (e.g., evidence refuting NEFM involvement) is presented—only absence of reported NEFM mutations. Therefore the most accurate classification given the supplied evidence is "Mixed Evidence": biologically plausible and candidate-supported but not directly demonstrated in these papers. I find no internal inconsistency in this reasoning and affirm the rating.


## Relevant Papers


### Charcot-Marie-Tooth disease neurofilament mutations disrupt neurofilament assembly and axonal transport.

**Authors**: J. Brownlees (H-index: 21), Christopher C. J. Miller (H-index: 49)

**Relevance**: 0.25

**Bibliometric Impact**: 0.42401818181818174


**Excerpts**:

- Charcot-Marie-Tooth disease (CMT) is the most common inherited disorder of the peripheral nervous system, and mutations in neurofilaments have been linked to some forms of CMT.

- Neurofilaments are the major intermediate filaments of neurones, but the mechanisms by which the CMT mutations induce disease are not known.

- Here, we demonstrate that CMT mutant neurofilaments disrupt both neurofilament assembly and axonal transport of neurofilaments in cultured mammalian cells and neurones.

- We also show that CMT mutant neurofilaments perturb the localization of mitochondria in neurones.

- Accummulations of neurofilaments are a pathological feature of several neurodegenerative diseases, including amyotrophic lateral sclerosis (ALS), Alzheimer's disease, Parkinson's disease, dementia with Lewy bodies, and diabetic neuropathy.

- Our results demonstrate that aberrant neurofilament assembly and transport can induce neurological disease, and further implicate defective neurofilament metabolism in the pathogenesis of human neurodegenerative diseases.


**Explanations**:

- How it relates: Directly relevant background linking mutations in neurofilaments to Charcot–Marie–Tooth disease, which is an inherited (i.e. Mendelian) disorder. This sentence provides direct disease-level evidence that mutations in neurofilament genes can cause Mendelian neuropathy. Strengths: explicit statement connecting neurofilament mutations to an inherited human disease. Limitations: the sentence does not specify which neurofilament genes are involved (NEFM is not named here), so it is indirect with respect to the specific claim about NEFM; no supporting data or genomic evidence are provided in this quoted sentence.

- How it relates: Context about the biological role of neurofilaments and an explicit statement that the mechanisms of pathogenicity are not fully understood. This is important for assessing whether mechanistic work in the paper could plausibly implicate a particular neurofilament gene in Mendelian disease. Strengths: frames the remaining mechanistic findings as addressing an acknowledged knowledge gap. Limitations: it emphasizes uncertainty—the paper is not claiming full mechanistic resolution—so this weakens any definitive inference that a specific neurofilament gene (NEFM) is established as a Mendelian disease gene.

- How it relates: Mechanistic evidence showing that disease-associated neurofilament mutants disrupt filament assembly and axonal transport in cultured cells and neurons. Strengths: provides plausible cellular mechanisms (assembly/transport defects) by which neurofilament gene mutations can produce neuronal pathology, supporting the biological plausibility that a mutation in any neurofilament gene (including NEFM) could be pathogenic in a Mendelian fashion. Limitations: the work is experimental (cell culture and neuronal models) rather than human genetic evidence; the excerpt does not indicate which neurofilament proteins (specific genes) were mutated in these experiments, and in vitro results may not fully recapitulate in vivo human disease.

- How it relates: Additional mechanistic evidence that mutant neurofilaments alter mitochondrial localization in neurons, suggesting downstream cellular dysfunction that could underlie neuropathology. Strengths: expands the mechanistic pathways (beyond assembly/transport) by which neurofilament mutations may cause neuronal disease, increasing plausibility for neurofilament gene involvement in Mendelian disorders. Limitations: again, the experiments are model-based and the excerpt lacks specification of gene identity (NEFM not named), quantitative effect sizes, or demonstration in human patients.

- How it relates: Contextualizes neurofilament pathology as a feature of multiple neurodegenerative diseases, implying that neurofilament dysfunction has broad pathological consequences. Strengths: broadens the clinical relevance of neurofilament abnormalities and supports the idea that neurofilament gene mutations can have pathogenic outcomes. Limitations: the listed diseases are mostly sporadic neurodegenerative conditions, not necessarily Mendelian, and this sentence does not provide gene-specific (NEFM) genetic evidence linking NEFM to a Mendelian disease.

- How it relates: The authors' concluding statement that aberrant neurofilament assembly and transport can induce neurological disease summarizes both direct and mechanistic findings and argues that defective neurofilament metabolism contributes to human disease pathogenesis. Strengths: synthesizes mechanistic and disease-relevant data, strengthening plausibility that mutations in neurofilament genes can be causal for neurological disorders. Limitations: it remains a general conclusion about neurofilament genes as a class; it does not provide gene-specific genetic evidence that NEFM is associated with a Mendelian disease, nor does it present human genetic linkage/segregation or variant pathogenicity data for NEFM.


[Read Paper](https://www.semanticscholar.org/paper/165c9052e95ca5d877b21bb930c06368f2ef5e7a)


### Axonal Charcot-Marie-Tooth disease and the neurofilament light gene (NF-L)

**Authors**: J. Lupski (H-index: 141)

**Relevance**: 0.35

**Bibliometric Impact**: 0.5803333333333334


**Excerpts**:

- Historical recombinants further narrowed the locus to a 16-cM interval that contained two excellent positional candidate genes: NF-L, which encodes the neurofilament light proteins, and NF-M, which encodes the neurofilament medium proteins.

- The entire coding regions were screened for both genes. An A→C transversion mutation predicted to result in a Gln333Pro missense mutation was identified in the first exon of NF-L.

- Neurofilaments are important for the structure and function of axons and may be necessary for axonal transport, regeneration, and longevity.

- The finding of an NF-L mutation associated with CMT2 supports a role for neurofilament proteins in this disease and also delineates a potential final common pathway for different forms of CMT1.

- As aptly pointed out by Mersiyanova et al. (2000 [in this issue]), neurofilament proteins are involved in the pathogenesis of several other neurological disorders, including giant axonal neuropathy, amyotrophic lateral sclerosis (ALS), and Parkinson and Alzheimer diseases.

- Furthermore, since neurofilament proteins apparently are key targets in axonal pathology, their corresponding genes—as well as genes for proteins that interact with or regulate neurofilaments—should be considered as candidate genes for other CMT loci.


**Explanations**:

- How it relates: This sentence identifies NF-M (the gene commonly referred to as NEFM) explicitly as an "excellent positional candidate" within the linked interval for CMT2 in the studied family. Type: indirect (positional) evidence. Strengths: NF-M/NEFM was in the linked interval, which is a standard initial criterion for disease-gene candidacy. Limitations/caveats: Being a positional candidate does not constitute demonstration of causality; the statement merely records NF-M's presence in the locus and its candidacy, not any mutation or segregation data implicating NF-M/NEFM.

- How it relates: These sentences report that both NF-L and NF-M coding regions were screened but that a causal coding mutation was found in NF-L (not NF-M) in this family. Type: direct genetic evidence (negative for NF-M in this family; positive for NF-L). Strengths: The authors explicitly screened NF-M coding sequence in the linked interval, and a clear disease-segregating NF-L mutation was identified, strengthening the conclusion that NF-L (not NF-M) explains the phenotype in this pedigree. Limitations/caveats: The negative result for NF-M refers only to coding-region screening in this single family; it does not exclude NEFM (NF-M) as a Mendelian disease gene in other families, nor does it exclude noncoding/regulatory variants or structural variants affecting NF-M that the screening could have missed.

- How it relates: Provides mechanistic context by describing the biological role of neurofilaments in axonal structure and transport—functions plausibly relevant to peripheral neuropathies. Type: mechanistic evidence. Strengths: Establishes biological plausibility that perturbation of neurofilament proteins (including NF-M/NEFM) could cause axonopathy and thus a Mendelian neuropathy. Limitations/caveats: This is general cell-biology context, not direct evidence that NEFM mutations cause human Mendelian disease; plausibility does not equal proof.

- How it relates: Shows that a bona fide neurofilament gene (NF-L) has a disease-causing mutation in CMT2, supporting the idea that neurofilament genes can be Mendelian disease genes. Type: indirect but strong mechanistic/genetic evidence. Strengths: Demonstration that mutation of one neurofilament gene causes a Mendelian neuropathy increases the prior probability that related neurofilament genes (like NEFM) could also be disease-causing. Limitations/caveats: This is evidence for the family and gene studied (NF-L) and only increases plausibility for NEFM; it is not direct evidence that NEFM itself is mutated in Mendelian disease.

- How it relates: Places neurofilament dysfunction in the broader landscape of neurological Mendelian and complex disorders, implying that neurofilament genes can be disease-relevant. Type: mechanistic/contextual evidence. Strengths: The association of neurofilament pathology with multiple neurological diseases supports the biological relevance of neurofilaments to disease. Limitations/caveats: The listed disorders are heterogeneous and many are not classic single-gene Mendelian conditions with established NEFM mutations; the sentence does not claim NF-M/NEFM mutations cause those diseases.

- How it relates: The authors explicitly recommend considering neurofilament genes and interacting regulators as candidate genes for other CMT loci—this is an explicit, reasoned argument that NF-M/NEFM should be evaluated as a potential Mendelian disease gene. Type: mechanistic/candidate-gene recommendation. Strengths: This statement synthesizes the genetic finding (NF-L mutation) and the mechanistic role of neurofilaments to justify testing NEFM in other CMT/axonopathy loci. Limitations/caveats: This is an expert recommendation rather than empirical proof; it supports targeted investigation but is not evidence of an existing NEFM-disease association.


[Read Paper](https://www.semanticscholar.org/paper/13ecb5f62e8dd6941596c05d354654e8ee0aee27)


## Other Reviewed Papers


### Axonal Charcot‐Marie‐Tooth neuropathy concurrent with distal and proximal weakness by translational elongation of the 3′ UTR in NEFH

**Why Not Relevant**: The provided paper content exclusively discusses NEFH (neurofilament heavy chain) and reports NEFH frameshift/stop-loss mutations and a 3' UTR translational-elongation mechanism that cause axonal neuropathy (CMT2CC) and are associated with ALS susceptibility. There is no mention of NEFM (the neurofilament medium chain gene), no data, no variants, and no mechanistic experiments involving NEFM. Because the claim concerns NEFM being likely associated with at least one Mendelian disease, evidence must directly involve NEFM (genetic, clinical, or mechanistic). Findings about NEFH cannot be taken as direct evidence for NEFM: although NEFH and NEFM are paralogous members of the neurofilament family, paralogy alone does not establish shared pathogenic variants, identical mutational mechanisms (e.g., presence of a cryptic amyloidogenic element in the 3' UTR), or Mendelian disease association. Inferring NEFM pathogenicity from NEFH would be speculative without sequence-specific, genetic, or functional data for NEFM. Therefore the paper is not relevant to the claim that NEFM is associated with a Mendelian disease.


[Read Paper](https://www.semanticscholar.org/paper/805aa8a1d0b4f145ed9ffa1fad539406a083195f)


### Heterogeneity in biomarkers, mitogenome and genetic disorders of the Arab population with special emphasis on large-scale whole-exome sequencing

**Why Not Relevant**: The provided paper content does not mention NEFM (neurofilament, medium polypeptide) or any synonyms/aliases for that gene, nor does it present data, analyses, or mechanistic discussion that bear on whether NEFM is associated with a Mendelian disease. The text is a high-level narrative summary of genomic findings in Arab populations (e.g., counts of novel variants, carrier frequencies for intellectual disability, identification of C12orf57 variants, and mitogenome variation) but contains no direct evidence (genetic association, segregation, pathogenic variants, case reports) or mechanistic evidence (functional studies, protein dysfunction, pathway involvement) linking NEFM to any Mendelian disorder. Therefore this paper provides neither supporting nor refuting evidence for the claim.

Limitations and caveats: absence of any mention of NEFM in this excerpt is not proof that NEFM is not associated with a Mendelian disease — it only shows that this particular review excerpt does not address NEFM. Narrative reviews may omit many genes outside their chosen examples (here C12orf57, intellectual disability, retinal dystrophy, mtDNA variation). To evaluate the claim further would require targeted searches of primary literature and curated disease-gene databases (e.g., OMIM, ClinVar, HGMD, PubMed) for NEFM-specific clinical reports, segregation studies, functional assays, or authoritative gene-disease validity assessments.


[Read Paper](https://www.semanticscholar.org/paper/b1efa17ab35bc6fcf1f887ba557baea309003a1d)


### A rare splice-site variant in cardiac troponin-T (TNNT2): The need for ancestral diversity in genomic reference datasets

**Why Not Relevant**: This paper does not provide any direct or mechanistic evidence for or against the claim that NEFM is likely associated with at least one Mendelian disease. The manuscript is entirely focused on a variant in the cardiac troponin T gene (TNNT2) and issues of ancestry and variant interpretation. Representative verbatim sentences from the provided content make the scope explicit: 

"We report a rare variant in the cardiac troponin T gene, TNNT2; NM_001001430.3: c.571-1G>A (rs483352835) identified via research-based whole exome sequencing in two unrelated probands of Oceanian ancestry with cardiac phenotypes." 

"Based on these data, we classify this variant as benign, and conclude it is not a monogenic cause of disease." 

Other sentences emphasize population-frequency and ancestry interpretation issues rather than data about NEFM: 

"The underrepresentation of different ancestry groups in large genomic datasets creates difficulties in interpreting the pathogenicity of monogenic variants." 

None of the text provided mentions NEFM, neurofilament medium chain, or any variants or phenotypes related to NEFM. There are no experimental data, case reports, segregation analyses, functional studies, mechanistic descriptions, or variant annotations concerning NEFM in the excerpt. 

Possible indirect or contextual relevance: the paper discusses general principles of variant interpretation, population frequency, and misclassification risk in underrepresented ancestries. Those topics are broadly relevant to how one would evaluate claims about gene-disease associations (including for NEFM) but do not supply any evidence about NEFM itself. 

Given the absence of any reference to NEFM or relevant data/mechanisms in the provided text, the relevance to the specific claim is effectively zero.


[Read Paper](https://www.semanticscholar.org/paper/4fed6e1e7cc6124f24c814ee86964f97c3ffaa39)


### Sural biopsy to detect the axonal cytoskeleton defects in KIF5A-related Charcot-Marie-Tooth disease type 2.

**Why Not Relevant**: The paper does not mention the NEFM gene, nor does it present any genetic evidence (variants, segregation, functional assays) implicating NEFM in disease. All genetic findings reported relate to KIF5A (p.E755K) and the reported pathology (neurofilament accumulation) is presented as secondary to defective axonal transport due to a KIF5A mutation. While the observed neurofilament pathology is mechanistically relevant to proteins encoded by NEFM (since NEFM encodes a neurofilament subunit), that is indirect: the study shows neurofilament disorganization as a downstream consequence of KIF5A dysfunction rather than primary disease caused by NEFM variants. Therefore this paper provides mechanistic context that neurofilament biology can be involved in neuropathology, but it provides no direct or genetic evidence that NEFM is associated with any Mendelian disease. Given the absence of NEFM-specific data, the relevance to the claim that "NEFM is likely associated with at least one Mendelian disease" is minimal.


[Read Paper](https://www.semanticscholar.org/paper/968c8f9dd01e691a0701c5d8e59400a58bc290f2)


### First case report of Charcot–Marie–Tooth disease type 2CC with a frameshift mutation of NEFH gene in Greece

**Why Not Relevant**: The provided content does not name NEFM or present any genetic, segregation, functional, or mechanistic data tying NEFM to a Mendelian disorder. It only generically states that electrophysiology plus genetic testing can diagnose CMT2CC and distinguish polyneuropathies. Without explicit mention of NEFM or presentation of evidence (variants, pathogenicity, inheritance, functional mechanism), the passage provides no direct or mechanistic support for the claim that NEFM is likely associated with at least one Mendelian disease. Additionally, the content is a single sentence without experimental detail, so it cannot establish association.


[Read Paper](https://www.semanticscholar.org/paper/44538be6f2817cbdbff19f080193da8734dc90a9)


### PSAP-genomic-regions: a method leveraging population data to prioritize coding and non-coding variants in whole genome sequencing for rare disease diagnosis

**Why Not Relevant**: This paper is a methods paper describing PSAP-genomic-regions, a prioritization tool for coding and non-coding variants in whole-genome sequencing. The provided content contains no mention of the gene NEFM, no data on variants in NEFM, and no experimental or clinical observations linking NEFM to any Mendelian phenotype. The paper reports evaluations on simulated datasets (ClinVar pathogenic variants inserted into population exomes/genomes) and on real cases (Cerebral Small Vessel Disease and male infertility), but no gene-specific results or lists are given in the provided text that would allow inference about NEFM. Mechanistic evidence (e.g., functional studies showing that NEFM mutations alter protein function leading to disease) and direct genetic evidence (e.g., segregation of NEFM variants with a Mendelian phenotype, case-control enrichment, or ClinVar entries for NEFM mutations) are absent from the excerpt. While the method described could, in principle, be used to prioritize pathogenic variants in NEFM if such variants existed in an analyzed genome, that is hypothetical and not evidence that NEFM is associated with any Mendelian disease. Therefore this paper provides no direct or mechanistic evidence for or against the claim that NEFM is likely associated with at least one Mendelian disease. If you want evidence about NEFM specifically, I recommend: (1) searching ClinVar and OMIM for NEFM entries, (2) searching PubMed for NEFM + disease/variant/mutation, and (3) examining case reports or functional studies that test NEFM variant pathogenicity.


[Read Paper](https://www.semanticscholar.org/paper/a2a1cfe7eb702f28e5bb9211607b7e7dc280bc7f)


### Rare Variant Genetic Association Study for Transplant-Associated Thrombotic Microangiopathy (TA-TMA) Via Whole Exome Sequencing

**Why Not Relevant**: The paper does not provide evidence for or against the claim that NEFM is associated with a Mendelian disease. NEFM (neurofilament medium polypeptide) is not mentioned in the manuscript, and the authors explicitly restricted their gene-level analyses to five a priori selected pathways and the genes within them (complement regulation, VWF and coagulation, VWF clearance, ADAMTS13-related proteins, and angiopoietin/endothelial activation). Their reported positive finding was in LRP1 within the VWF clearance pathway, and they report negative findings in the complement pathway. Although whole exome sequencing was performed, the study’s analytical focus and filtering criteria mean NEFM was not evaluated or reported here; therefore this paper provides no direct or mechanistic evidence regarding NEFM’s involvement in Mendelian disease. Additionally, the modest sample size (91 cases, 93 controls) limits power to detect rare Mendelian associations even for genes that were analyzed. In sum, relevance to the NEFM-Mendelian disease claim is negligible because NEFM is neither analyzed nor discussed in the study.


[Read Paper](https://www.semanticscholar.org/paper/a28f55f5071fa816fb5c466832a38b225dfaef02)


## Search Queries Used

- NEFM mutation familial segregation Mendelian disease case report neurofilament medium polypeptide hereditary

- NEFM variant rare variant exome sequencing whole exome sequencing whole genome sequencing pathogenicity ClinVar gnomAD population frequency neurofilament medium

- NEFM neurofilament medium axonal neuropathy Charcot Marie Tooth peripheral neuropathy amyotrophic lateral sclerosis Alzheimer's disease neurodegeneration mutation

- neurofilament medium NEFM function axonal transport intermediate filament assembly interaction with NEFL NEFH mechanism functional study review systematic review meta analysis Mendelian


## Usage Statistics

- Prompt Tokens: 0

- Completion Tokens: 0

- Total Tokens: 0

- Estimated Cost: $0.0441


## Bibliometric Configuration

- Use Bibliometrics: True

- Author Impact Weight: 0.4

- Citation Impact Weight: 0.4

- Venue Impact Weight: 0.2
