## Supplementary material for "MENDELSEEK: An algorithm that predicts Mendelian Genes and elucidates what makes them special": NFATC4.txt

# Claim: The gene NFATC4 is likely associated with at least one Mendelian disease.

**Status**: processed

**Overall Rating**: -1

**Explanation**:

No relevant papers were found that support or refute this claim.


## Relevant Papers


## Other Reviewed Papers


### Mutation-dependent recessive inheritance of NPHS2-associated steroid-resistant nephrotic syndrome

**Why Not Relevant**: The provided paper content discusses the NPHS2 gene and the p.Arg229Gln podocin variant (altered heterodimerization and mislocalization) and states that disease results only when this allele co-occurs with certain 3′ NPHS2 mutations. It makes no mention of NFATC4, does not present linkage, mutation, functional, or clinical evidence for NFATC4, nor does it describe pathways that would plausibly connect NPHS2 findings to NFATC4. Because the content addresses a different gene (NPHS2/podocin) and provides no direct or mechanistic information about NFATC4, it is not relevant to the claim that NFATC4 is likely associated with a Mendelian disease.


[Read Paper](https://www.semanticscholar.org/paper/8cd90c3b1f5497421ccb70b43eb0b8a848ddd4f8)


### The Burden of Candidate Pathogenic Variants for Kidney and Genitourinary Disorders Emerging From Exome Sequencing

**Why Not Relevant**: The provided paper excerpt describes methods for identifying and curating genes associated with Mendelian kidney and genitourinary disorders and for screening a large control cohort for candidate pathogenic variants. However, the excerpt does not mention the gene NFATC4 by name, nor does it provide the curated list of 625 genes or gene-level results. Therefore the paper content as provided cannot directly support or refute the claim that "NFATC4 is likely associated with at least one Mendelian disease." At best, the paper establishes a framework whereby NFATC4 would be captured if it were listed in OMIM/Orphanet and retained after curation, but absent the actual gene list or explicit statements about NFATC4, no gene-specific conclusion can be drawn. To evaluate the claim one would need either (a) the paper's full supplementary gene list (to see if NFATC4 is included and with what evidence), (b) direct statements or results about NFATC4 in the text, or (c) independent primary sources (OMIM, ClinVar, HGMD, or gene-specific literature) documenting NFATC4 associations with Mendelian phenotypes. Because the excerpt lacks any NFATC4-specific data, I assign a low relevance score (<0.1).


[Read Paper](https://www.semanticscholar.org/paper/a75e297208c04992568e607d19aa9b9aaebd09d5)


### Novel variants identified with next-generation sequencing in Polish patients with cone-rod dystrophy

**Why Not Relevant**: Relevance is very low (0.02) because the paper focuses specifically on identifying genes and variants underlying cone-rod dystrophy (CRD) in 18 Polish families and reports implicated genes (RPGR-ORF15, PROM1, PRPH2, ATF6, ABCA4, etc.). The paper does not mention NFATC4 at all, provides no mechanistic data involving NFATC4, and does not present any direct evidence for or against NFATC4 being associated with any Mendelian disease. The methodological design (targeted IRD gene panel plus selective WES) means NFATC4 could have been omitted from the panel or missed in families not subjected to WES; furthermore, absence of NFATC4 findings in a small, phenotype-restricted cohort is weak negative evidence and does not refute NFATC4 association with other Mendelian diseases. Therefore this paper neither supports nor meaningfully refutes the claim that NFATC4 is likely associated with at least one Mendelian disease.


[Read Paper](https://www.semanticscholar.org/paper/91e01bc532b442d01b6280069be5ef8f62922c5f)


### Expanding the phenotype of a recurrent de novo variant in PACS1 causing intellectual disability

**Why Not Relevant**: The paper exclusively reports and discusses a recurrent de novo PACS1 missense variant causing an intellectual disability syndrome (MRD17). It provides clinical, genetic, and mechanistic evidence related to PACS1 (including functional data in zebrafish and hypotheses about altered CK2-motif binding) but contains no mention, data, analysis, or discussion of the gene NFATC4, its variants, or any pathway linking NFATC4 to human Mendelian disease. Absence of mention in this PACS1-focused study means it supplies no direct or mechanistic evidence for or against the claim that NFATC4 is associated with a Mendelian disease. This lack of information—not evidence of absence of effect—drives the relevance rating of 0.0.


[Read Paper](https://www.semanticscholar.org/paper/e460d89d850693b4fb1e3df12702be0dfb8d7f5b)


### Mutations and mechanism: how PINK1 may contribute to risk of sporadic Parkinson's disease.

**Why Not Relevant**: The provided paper content is a short scientific commentary focused on PINK1 and other established Parkinson's disease genes and mechanisms. The text explicitly names SNCA, LRRK2, VPS35, PINK1, PARK7/DJ-1, PARK2/PARKIN, PLA2G6, ATP13A2, and FBXO7 as monogenic Parkinson's disease genes and discusses the relationship between monogenic and sporadic Parkinson's disease. Nowhere in the supplied content is the gene NFATC4 mentioned, nor are any pathways, functional data, genetic associations, or mechanistic discussions involving NFATC4 presented. Therefore this paper provides neither direct evidence (e.g., mutation-disease segregation, case reports, linkage, or association) nor mechanistic evidence (e.g., functional studies implicating NFATC4 in disease biology) supporting the claim that NFATC4 is associated with a Mendelian disease.

Limitations of this assessment: absence of NFATC4 in this commentary does not prove NFATC4 has no Mendelian associations in the broader literature. It only demonstrates that this particular paper does not contain relevant evidence. To evaluate the claim about NFATC4 one must consult primary genetic-disease resources (OMIM, ClinVar), gene-focused studies, or other publications that directly examine NFATC4 sequence variation and disease segregation or functional impact.


[Read Paper](https://www.semanticscholar.org/paper/99a394b75e80548eaff92176d5e13d2748cf9bc6)


### Unravelling the genetic causes of multiple malformation syndromes: A whole exome sequencing study of the Cypriot population

**Why Not Relevant**: The paper does not mention NFATC4 anywhere in the provided content. It reports the genes in which pathogenic or likely pathogenic variants were found (PCNT, UBE3A, KAT6A, SPR, POMGNT1, PIEZO2, PXDN, KDM6A, PHIP, HECW2, TFAP2A, CNOT3, AGTPBP1, GAMT) and provides study design, yield and conclusions about WES utility. Because NFATC4 is neither reported among identified genes nor discussed mechanistically, the paper provides no direct evidence that NFATC4 is associated with a Mendelian disease. The absence of NFATC4 findings in this specific cohort is only very weak indirect evidence against an association; it could reflect limited cohort size, phenotype ascertainment (MMS cases only), population specificity (Greek-Cypriot), or simply that NFATC4-related disease is not present in this sample. Therefore the content is essentially not relevant to the claim that "NFATC4 is likely associated with at least one Mendelian disease."


[Read Paper](https://www.semanticscholar.org/paper/3fe1dcfb954c4d8ab6d06a10dee5a5940f27cdfa)


### Title Plant Homeo Domain Finger Protein 8 Regulates Mesodermal and Cardiac Differentiation of Embryonic Stem Cells Through Mediating the Histone Demethylation of pmaip 1

**Why Not Relevant**: This paper does not provide evidence for or against the claim that NFATC4 is likely associated with at least one Mendelian disease. The manuscript focuses entirely on the histone demethylase PHF8 and its regulation of the pro-apoptotic gene pmaip1 during differentiation of mouse embryonic stem cells (mESCs). There is no mention of NFATC4, no analysis of NFATC4 expression or function, no human genetic or clinical data, and no discussion of Mendelian disease associations. Representative verbatim sentences from the provided paper text that illustrate its focus (and the absence of any NFATC4 content) are reproduced here exactly as in the source to show context and scope: 

"Histone demethylases have emerged as key regulators of biological processes. The H3K9me2 demethylase plant homeo domain finger protein 8(PHF8), for example, is involved in neuronal differentiation, but its potential function in the differentiation of embryonic stem cells (ESCs) to cardiomyocytes is poorly understood." 

"Using a phf8 knockout (ph8) model, we found that deletion of phf8 in ESCs did not affect self-renewal, proliferation or early ectodermal/endodermal differentiation, but it did promote the mesodermal lineage commitment with the enhanced cardiomyocyte differentiation." 

"Functionally, PHF8 promoted the loss of a repressive mark H3K9me2 from the transcription start site of a proapoptotic gene pmaip1 and activated its transcription." 

"These results reveal that the histone demethylase PHF8 regulates mesodermal lineage and cell fate decisions in differentiating mESCs through epigenetic control of the gene critical to programmed cell death pathways." 

Because NFATC4 is neither named nor studied in any experimental, mechanistic, genetic, or clinical context in this paper, the study cannot be used to support or refute an association between NFATC4 and any Mendelian disease. Additional types of evidence that would be required to support the claim (none of which are present here) include: (1) observation of NFATC4 mutations segregating with a Mendelian phenotype in families, (2) case-control or population genetic enrichment of rare damaging NFATC4 variants in affected individuals, (3) functional studies showing that NFATC4 variants alter protein function in a manner consistent with disease biology, or (4) clinical reports linking NFATC4 genetic variation to specific inherited disorders. The present study's methodological scope (mouse ESC differentiation, PHF8 knockout, epigenetic regulation of pmaip1) and its experimental conditions (in vitro differentiation, mouse model) limit its generalizability to human genetic disease and provide no mechanistic connection to NFATC4.


[Read Paper](https://www.semanticscholar.org/paper/c6e507e133e6719400902fa52351028cb65a295e)


### RLIM enhances BMP signalling mediated fetal lung development in mice

**Why Not Relevant**: The provided paper text discusses RLIM (also called RNF12), an X-linked E3 ubiquitin ligase, its role in regulating SMAD7 and BMP/TGFβ signalling, and phenotypic effects of Rlim knockout in mice. Nowhere in the supplied content is the gene NFATC4 mentioned, nor are any data, variants, phenotypes, or mechanistic pathways linking NFATC4 to Mendelian disease presented. Therefore the paper provides no direct evidence (no reports of NFATC4 variants segregating with Mendelian phenotypes, no clinical descriptions involving NFATC4) and no mechanistic evidence (no biochemical, regulatory, or pathway connections between RLIM/SMAD/BMP and NFATC4) relevant to the claim that NFATC4 is likely associated with at least one Mendelian disease.

Specific reasons for assigning relevance = 0.0:
- Absence of NFATC4: The entire text refers to RLIM/RNF12 and BMP pathway components (SMAD7, SMAD1/5/9, SMAD5) and to mouse Rlim knockout phenotypes; NFATC4 is not mentioned.
- No genetic association data for NFATC4: The paper reports human RLIM mutations and mouse Rlim knockout phenotypes, but provides no variant-level, segregation, case-series, or case-control data for NFATC4.
- No mechanistic linkage: The molecular mechanisms described concern ubiquitin-mediated regulation of SMAD proteins and downstream BMP signalling; there is no presentation of transcriptional, post-translational, or protein–protein interactions connecting NFATC4 to the pathways or phenotypes discussed.

Implications and recommended next steps to evaluate the claim properly:
- To assess whether NFATC4 is likely associated with a Mendelian disease, search primary resources that catalog gene-disease associations and human variants (e.g., OMIM, ClinVar, HGMD) and peer-reviewed human genetics literature (PubMed) for reports of pathogenic/likely pathogenic variants in NFATC4 with segregating phenotypes.
- Evaluate functional/mechanistic evidence in human or model systems that links NFATC4 perturbation to disease-relevant biology (e.g., loss-of-function experiments, dominant-negative effects, animal models) if such reports exist.

Because the supplied paper contains zero mention or data about NFATC4, no verbatim excerpts from it can be offered that bear on the claim.


[Read Paper](https://www.semanticscholar.org/paper/c3f852c711d11feb67caac86074b509f852e72af)


### Negative selection on human genes causing severe inborn errors depends on disease outcome and both the mode and mechanism of inheritance

**Why Not Relevant**: The provided paper content contains no mention of the specific gene NFATC4 and does not present gene-level results (i.e., CoNeS or negative selection scores) for NFATC4. The paper reports aggregate, population-level relationships between mode/mechanism/severity of Mendelian disorders and measures of negative selection across sets of disease genes. Those results are mechanistically relevant to assessing whether a gene with certain properties might be disease-associated, but they do not constitute direct evidence that NFATC4 is associated with any Mendelian disease. Without NFATC4 being named or its gene-specific CoNeS/selection metrics reported, the paper cannot support or refute the claim about NFATC4. Therefore the relevance to the specific claim is very low.


[Read Paper](https://www.semanticscholar.org/paper/1b29ee22a0cce5556f7265d657162fbd4c4f40c2)


### Ultra-rare de novo damaging coding variants are enriched in attention-deficit/hyperactivity disorder and identify risk genes

**Why Not Relevant**: The paper does not mention NFATC4 anywhere in the provided content and focuses exclusively on discovering ADHD risk genes through de novo and rare variant analyses in parent-child trios and case-control cohorts. The study's design and conclusions pertain to genetic contributions to ADHD (a complex neurodevelopmental disorder) and to identification of ADHD risk genes (e.g., KDM5B), not to demonstrating that any specific gene is causative for a Mendelian disease. Establishing that NFATC4 is 'likely associated with at least one Mendelian disease' requires different evidence types (e.g., multiple unrelated families with segregating, highly penetrant pathogenic variants in NFATC4; ClinVar/OMIM entries linking NFATC4 to a defined clinical syndrome; functional assays showing variants produce disease-relevant loss/gain of function). None of those are provided here. The modest trio sample size (152 trios) and the study's focus on ADHD phenotypes further limit the ability to infer Mendelian disease associations from these results. Therefore the paper provides no direct or mechanistic evidence supporting the claim about NFATC4; at best its negative result (no mention of NFATC4 among reported genes) is weak, indirect evidence and insufficient to support or refute the claim.


[Read Paper](https://www.semanticscholar.org/paper/8aac736fb881aba6c934da0e920b58ba657d373f)


### Abstract 17136: Irx3 and Irx5 Cooperatively Regulate Mouse Heart Development and Adult Cardiac Function

**Why Not Relevant**: This paper does not provide evidence for or against the claim that NFATC4 is likely associated with at least one Mendelian disease. The text exclusively discusses the Iroquois homeobox family members Irx3 and Irx5 and their roles in cardiac development and adult cardiac function. There is no mention of NFATC4, no data (genetic, clinical, linkage, or association) concerning NFATC4, and no mechanistic pathways described that connect Irx3/Irx5 with NFATC4. Therefore there is neither direct evidence (e.g., pathogenic variants in NFATC4 causing Mendelian phenotypes) nor mechanistic evidence (e.g., NFATC4 participating in the studied cardiac developmental pathways) present in this paper. The absence of NFATC4-related content in this manuscript means it is not informative for assessing NFATC4’s Mendelian disease status. Note that absence of evidence in this paper does not disprove the claim; it only indicates this paper is not relevant. To evaluate the claim would require sources that (a) report segregating NFATC4 variants in Mendelian pedigrees, (b) show de novo NFATC4 pathogenic variants in individuals with consistent phenotypes, or (c) provide functional data linking NFATC4 perturbation to a Mendelian phenotype.


[Read Paper](https://www.semanticscholar.org/paper/6630d29d31ebb00cbe22661265d5d20e80fc77d2)


### Investigating the role of Prox1 in cardiac and skeletal muscle development and disease

**Why Not Relevant**: The provided paper text discusses Prox1-driven regulation of contractile protein isoforms and resultant cardiomyopathy in mice; it does not mention NFATC4 or any NFAT family member, nor does it present human genetic association data, variant analyses, segregation studies, or clinical descriptions that would indicate a Mendelian disease linked to NFATC4. There is no direct evidence (no mention of the gene, variants, patients, or disease phenotype attributed to NFATC4) and no mechanistic evidence tying NFATC4 into the described Prox1-related pathways in the text supplied. Because the content does not contain any reference to NFATC4 or human Mendelian genetics, it cannot support or refute the claim that NFATC4 is likely associated with at least one Mendelian disease. To evaluate that claim, one would need to consult human genetics resources (OMIM, ClinVar, ClinGen), published human genetic studies, or functional studies explicitly involving NFATC4 and human disease phenotypes.


[Read Paper](https://www.semanticscholar.org/paper/e1cb8fd6d46f6f97860fa1420fdce6cc86bad537)


### Whole genome analysis of rare deleterious variants adds further evidence to BRSK2 and other risk genes in Autism Spectrum Disorder

**Why Not Relevant**: The paper's content focuses on rare variants identified in autism spectrum disorder cases and enumerates specific genes found to carry potentially damaging variants. NFATC4 is not mentioned in the excerpted content (nor in the list of genes reported), and no experimental, clinical, or mechanistic data involving NFATC4 are presented. Therefore this paper does not provide supporting or refuting evidence for the claim that NFATC4 is likely associated with a Mendelian disease. Note that a lack of mention in this study is not disproof: NFATC4 could be associated with Mendelian disease in other cohorts or via variant types not detectable or not present in this dataset (e.g., regulatory variants, rare structural changes, different phenotypes, or very small number of cases). Additionally, the paper's sample size and ascertainment (ASD-focused families) limit its ability to detect Mendelian disease associations for genes not relevant to ASD phenotypes.


[Read Paper](https://www.semanticscholar.org/paper/04930d2ae0bf3e76a2523ea2f11fc318ed3e3779)


### Study of the Effect of Calreticulin on Orai1 Function

**Why Not Relevant**: The paper discusses NFAT as a family of transcription factors and describes how calcium signaling, SOCE, calcineurin, and calreticulin affect NFAT activation and nuclear import. However, it does not mention NFATC4 by name, does not present any genetic analyses, variant data, segregation studies, clinical phenotypes, or human disease associations, and contains no evidence (statistical or case-based) linking NFATC4 to a Mendelian disorder. Therefore the content provides mechanistic context about NFAT signaling but no direct or specific evidence that the gene NFATC4 is associated with at least one Mendelian disease.


[Read Paper](https://www.semanticscholar.org/paper/6152eeb09ce6619057151dfc346465a9b164e3ab)


### Absent meibomian glands and cone dystrophy in ADULT syndrome: identification by whole exome sequencing of pathogenic variants in two causal genes TP63 and CNGB3

**Why Not Relevant**: The paper focuses on a clinical case (and family) with syndromic ectrodactyly and cone dystrophy and identifies pathogenic variants in TP63 and CNGB3 via Whole Exome Sequencing. NFATC4 is not mentioned anywhere in the provided content. Therefore the paper provides no direct evidence supporting or refuting the claim that NFATC4 is associated with a Mendelian disease. At best it offers weak, indirect evidence by omission (WES was performed but no NFATC4 variant was reported), which is insufficient to support or disprove a gene–disease association. Limitations: absence of mention in a single case report does not constitute evidence against NFATC4 involvement in Mendelian disease; WES detection and reporting depend on coverage, variant interpretation, and the authors' focus on variants matching the observed phenotypes. For these reasons relevance to the claim is effectively zero.


[Read Paper](https://www.semanticscholar.org/paper/1f9e7c1f5d2fb3721402c7e010cc91377c9bb2ee)


## Search Queries Used

- NFATC4 mutation pathogenic variant Mendelian disease monogenic disorder autosomal dominant autosomal recessive haploinsufficiency dominant negative

- NFATC4 ClinVar OMIM HGMD ClinGen review systematic review NFATC4 gene disease association

- NFATC4 transcription factor calcineurin pathway loss of function knockout mouse phenotype cardiac development neuronal development gene knockout study

- whole exome sequencing whole genome sequencing rare damaging variants NFATC4 de novo variant neurodevelopmental disorder congenital malformation case report family segregation study


## Bibliometric Configuration

- Use Bibliometrics: True

- Author Impact Weight: 0.4

- Citation Impact Weight: 0.4

- Venue Impact Weight: 0.2
