## Supplementary material for "MENDELSEEK: An algorithm that predicts Mendelian Genes and elucidates what makes them special": NKX3-1.txt

# Claim: The gene NKX3-1 is likely associated with at least one Mendelian disease.

**Status**: processed

**Overall Rating**: -1

**Explanation**:

No relevant papers were found that support or refute this claim.


## Relevant Papers


## Other Reviewed Papers


### The Burden of Candidate Pathogenic Variants for Kidney and Genitourinary Disorders Emerging From Exome Sequencing

**Why Not Relevant**: The provided paper content does not mention the gene NKX3-1 anywhere in the quoted text, nor does it provide a gene-by-gene list that would allow determination of whether NKX3-1 was identified as associated with a Mendelian disorder. The manuscript excerpts describe methods for identifying and curating genes associated with Mendelian kidney and genitourinary disorders (e.g., use of OMIM and Orphanet, curation to a final list of 625 genes), cohort characteristics, and variant-filtering pipelines, but they do not include the curated gene list or any specific results for individual genes. Therefore there is no direct evidence in the supplied content that NKX3-1 is associated with any Mendelian disease, nor is there mechanistic or variant-level data about NKX3-1 (such as reported pathogenic variants, inheritance mode, functional annotations, or frequency data) that could be used to infer such an association.

Limitations that prevent assessment from this paper content:
- No occurrence of the symbol or name "NKX3-1" in the provided text.
- The curated gene list (the final 625 genes) is referenced but not included in the provided excerpts or appendices, so it is impossible to tell whether NKX3-1 is among the genes the authors considered associated with Mendelian kidney or genitourinary disorders.
- No gene-specific results, variant counts, or mechanistic discussion for individual genes are present in the supplied content.

Possible indirect relevance (not sufficient as evidence): the paper documents a systematic pipeline (OMIM/Orphanet searches and manual curation) for identifying genes associated with Mendelian kidney/genitourinary disorders and analyzes variant prevalence in a large healthy-cohort exome dataset. If NKX3-1 had been included in their curated gene list and reported elsewhere in the full paper or appendices with associated pathogenic variants or relevant mechanistic notes, that material could be relevant. However, such information is not present in the supplied text, so no determination can be made from these excerpts about NKX3-1's association with any Mendelian disease.


[Read Paper](https://www.semanticscholar.org/paper/a75e297208c04992568e607d19aa9b9aaebd09d5)


### A commentary on the promise of whole-exome sequencing in medical genetics

**Why Not Relevant**: The provided paper content contains only a single general sentence describing whole-exome sequencing as a technique. It does not mention the gene NKX3-1, any phenotype, any patient or family data, variant findings, segregation analysis, functional studies, or clinical interpretation. Because there is no gene-specific or disease-specific information, there is no direct or substantive mechanistic evidence in this content to support or refute the claim that NKX3-1 is likely associated with at least one Mendelian disease. The only tenuous connection is methodological: WES is a tool that could be used to discover such an association, but the content gives no actual findings or analysis related to NKX3-1. For these reasons the relevance to the claim is minimal.


[Read Paper](https://www.semanticscholar.org/paper/daa71ffe97fb47c9e5836d58ac8ed461198deb7f)


### Whole-Exome Sequencing in Adults With Chronic Kidney Disease

**Why Not Relevant**: The supplied paper text does not mention the gene NKX3-1, nor does it present any data, results, or discussion that directly link NKX3-1 to a Mendelian disease. The study focuses on diagnosing Mendelian forms of kidney disease using a curated set of nephropathy genes and on reviewing OMIM and ACMG genes, but the actual gene lists (Supplement Table 3) and gene-level findings are not included in the provided content. Because NKX3-1 is not named anywhere in the text provided and no variant-level results for that gene are presented, the paper cannot be used to support or refute the claim that NKX3-1 is likely associated with at least one Mendelian disease. Additional information needed to assess the claim from this study would include the full Supplement Table 3 (to see if NKX3-1 was included), any gene-level results or tables, and any mention in the results or discussion sections. Given the absence of any direct mention or data on NKX3-1 in the supplied content, the relevance of this paper to the claim is negligible.


[Read Paper](https://www.semanticscholar.org/paper/a29bd57964f68ac5f8e1abcafe1112ba488f53f5)


### Germline and Somatic Whole-Exome Sequencing Identifies New Candidate Genes Involved in Familial Predisposition to Serrated Polyposis Syndrome

**Why Not Relevant**: The paper does not mention NKX3-1 anywhere in the provided content. Its aim and results are confined to identifying candidate germline predisposition genes for serrated polyposis syndrome/colorectal cancer, and the genes highlighted (ANXA10, ASXL1, CFTR, DOT1L, HIC1, INO80, KLF3, MCM3AP, MCM8, PDLIM2, POLD1, TP53BP1, WNK2 and WRN) do not include NKX3-1. While the study used whole-exome sequencing in familial SPS cases (so it could have detected coding germline variants in NKX3-1 if present and passing filters), the absence of NKX3-1 from the reported candidates is limited as evidence because: (1) the study is disease-specific (SPS/CRC) and a negative result here does not speak to NKX3-1 associations with other Mendelian diseases; (2) the sample size is modest (39 patients across 16 families), so rare Mendelian associations might not be represented; (3) whole-exome sequencing and filtering pipelines can miss certain variant types (e.g., noncoding regulatory variants, structural variants, low-coverage exons) or filter out true positives; and (4) the authors themselves characterize their candidate list as provisional pending functional validation and replication. For these reasons, the paper provides no substantive evidence for or against the claim that NKX3-1 is likely associated with at least one Mendelian disease.


[Read Paper](https://www.semanticscholar.org/paper/f986efea9a4a25f8c852540c4e0e82473d2812b8)


### Whole exome sequencing identified a novel POT1 variant as a candidate pathogenic allele underlying a Li–Fraumeni-like family

**Why Not Relevant**: The paper focuses on a Li-Fraumeni-like family and reports whole-exome sequencing results identifying a novel germline variant in POT1 as the likely causal allele. The abstract contains no mention of NKX3-1, no reported germline or somatic variants in NKX3-1, and no mechanistic discussion involving NKX3-1. Although the study used WES (which in principle could detect variants in NKX3-1), the abstract does not state that NKX3-1 was examined, found, or explicitly excluded. Therefore this paper provides essentially no direct or mechanistic evidence relevant to the claim that NKX3-1 is likely associated with at least one Mendelian disease. Absence of evidence in this single, small (13 individuals) family-based study does not meaningfully inform the broader question of NKX3-1 disease associations; lack of mention should not be interpreted as evidence against NKX3-1 involvement in Mendelian disease elsewhere.


[Read Paper](https://www.semanticscholar.org/paper/d370ec500a223fc8d190c6a3ba8acf529c381b55)


### The simplest explanation does not have to be preferred: co-occurrence of pathogenic variants in cancer-predisposing genes

**Why Not Relevant**: The provided text contains only a single summary-methods sentence describing that the paper reviews MINAS frequency/phenotypes and checks UK 10,000 Genomes Project data for tumor-predisposition participants and whether CSG effects are additive or synergistic. It does not mention NKX3-1, any specific gene-level associations, variant data, segregation evidence, OMIM links, or mechanistic pathways relevant to Mendelian disease. Therefore the excerpt provides no direct or mechanistic evidence for or against the claim that NKX3-1 is likely associated with at least one Mendelian disease. No conclusions about NKX3-1 can be drawn from this content; establishing relevance would require paper text that names NKX3-1 and reports variant-phenotype co-segregation, pathogenicity evidence, or mechanistic data.


[Read Paper](https://www.semanticscholar.org/paper/a1372a664bd2c33a0bda638975a8b26d112457fc)


### Familial Predisposition in Myeloproliferative Neoplasms: An NGS-Driven Analysis

**Why Not Relevant**: This paper does not provide evidence that NKX3-1 is likely associated with a Mendelian disease. NKX3-1 is not mentioned anywhere in the provided text, it does not appear among the listed genes found to be mutated in this familial MPN cohort, and the study used a targeted 70-gene panel (genes "selected based on the available evidence for Hereditary cancer syndromes"). Therefore: (1) there is no direct positive evidence (no reported NKX3-1 pathogenic or likely pathogenic germline variants); (2) there is no mechanistic discussion linking NKX3-1 to Mendelian disease in this paper. Important limitations affecting the ability of this paper to inform the claim: the study used a targeted gene panel so absence of NKX3-1 findings could reflect that NKX3-1 was not included on the panel rather than true absence of association; the cohort is modest in size (54 subjects) and specific to familial myeloid neoplasms, which limits generalizability to other Mendelian diseases; and the paper focused on genes related to DNA repair, telomere regulation, and known cancer predisposition genes rather than an unbiased genome-wide search for novel Mendelian genes. For these reasons the relevance of this paper to the claim about NKX3-1 is very low.


[Read Paper](https://www.semanticscholar.org/paper/4667b77e18b0caa52a8e17c16d764fc139df55a0)


### Somatic Mutation Screening in Adult Patients with Myeloid Malignancies Provides Insights into Likely Germline Variants

**Why Not Relevant**: This paper does not provide evidence that directly supports or refutes the claim that NKX3-1 is likely associated with at least one Mendelian disease. NKX3-1 is not mentioned anywhere in the provided text. The study focuses on germline-associated variants relevant to myeloid neoplasms (AML/MDS), and the genes explicitly reported as frequent findings (e.g., TP53, RUNX1, DDX41, CEBPA, NF1, BRCA1) do not include NKX3-1. Moreover, the authors used tumor-only somatic NGS on diagnostic bone marrow with limited confirmatory germline testing and evolving panels over time—conditions that both limit detection of germline variants and constrain the gene set surveyed. Therefore: (1) absence of NKX3-1 from this paper does not provide substantive negative evidence against its involvement in Mendelian disease; (2) the study design and stated limitations mean it was not well suited to discover or evaluate associations for genes outside established myeloid predisposition panels. To assess the claim about NKX3-1, one should consult resources specifically curated for Mendelian disease associations (e.g., OMIM, ClinVar, HGMD, gene-disease curation efforts such as ClinGen) and targeted literature describing NKX3-1 variants in familial disease or functional/mechanistic studies. Given the absence of any mention or analysis of NKX3-1 in this paper, its relevance to the claim is negligible.


[Read Paper](https://www.semanticscholar.org/paper/9e17b5c78f81cb7c08b9e3faf01919f391e308ae)


### Beyond APOL 1 : Genetic Inroads into Understanding Population Disparities in Diabetic Kidney Disease

**Why Not Relevant**: The provided paper excerpt does not mention the gene NKX3-1 anywhere. All explicit Mendelian disease statements relate to other genes (CUBN, AMN, LRP2) involved in kidney or multisystem syndromes. Thus the paper contains no direct genetic, clinical, mechanistic, or epidemiologic evidence linking NKX3-1 to any Mendelian disorder. The included sentences are useful only as background (examples of Mendelian gene–disease relationships and conceptual discussion about predictability between rare high‑penetrance variants and common variant associations). Because NKX3-1 is absent from the text, this source cannot be used to support the claim that "NKX3-1 is likely associated with at least one Mendelian disease." The relevance score is therefore very low; any assessment about NKX3-1 would require other sources that explicitly mention NKX3-1 mutations and phenotypes.


[Read Paper](https://www.semanticscholar.org/paper/52d3f3490311ed466b2327ad26b88d847ffd485b)


## Search Queries Used

- NKX3 1 NKX3.1 germline mutation variant pathogenic Mendelian disease monogenic hereditary syndrome familial case report OMIM ClinVar

- NKX3 1 NKX3.1 knockout mouse haploinsufficiency loss of function developmental phenotype prostate urinary tract malformation homeobox gene

- NKX3 1 NKX3.1 germline somatic mutation hereditary prostate cancer familial predisposition exome sequencing family study cancer predisposition gene

- NKX3 1 NKX3.1 rare disease exome sequencing undiagnosed disease Mendelian gene discovery candidate gene review meta analysis gene disease association


## Bibliometric Configuration

- Use Bibliometrics: True

- Author Impact Weight: 0.4

- Citation Impact Weight: 0.4

- Venue Impact Weight: 0.2
