## Supplementary material for "MENDELSEEK: An algorithm that predicts Mendelian Genes and elucidates what makes them special": NOTCH4.txt

# Claim: The gene LRCH1 is likely associated with at least one Mendelian disease.

**Status**: processed

**Overall Rating**: -1

**Explanation**:

No relevant papers were found that support or refute this claim.


## Relevant Papers


## Other Reviewed Papers


### Exome Sequencing of a Clinical Population for Autosomal Dominant Polycystic Kidney Disease.

**Why Not Relevant**: The paper is focused specifically on autosomal dominant polycystic kidney disease (ADPKD) and the contribution of PKD1, PKD2, and a defined set of other cystic-kidney-related genes (ALG8, ALG9, DNAJB11, GANAB, HNF1B, IFT140, SEC61B, PKHD1, PRKCSH, SEC63). LRCH1 is not mentioned anywhere in the provided text, was not among the genes analyzed or reported as associated, and no data (genetic, phenotypic, or mechanistic) about LRCH1 are provided. Therefore this study offers no direct evidence supporting or refuting the claim that LRCH1 is associated with a Mendelian disease. Note that absence of mention in this disease-focused study does not constitute evidence that LRCH1 is not a Mendelian disease gene in other contexts; it simply means this paper does not address the claim.


[Read Paper](https://www.semanticscholar.org/paper/62a9d9f9cdfbebdb1239b48dc88b23e2c4b4781c)


### Association of a rare NOTCH4 coding variant with systemic sclerosis: a family-based whole exome sequencing study

**Why Not Relevant**: The provided paper content contains a single sentence describing a family in which a rare, penetrant coding variant at an unnamed locus increases risk for systemic sclerosis compared to common non-coding variants. However, the sentence does not mention the gene LRCH1, does not report any specific variant, genotype-phenotype co-segregation data, diagnostic criteria for a Mendelian disorder, or statistical evidence tying LRCH1 to disease. The claim to evaluate is specifically that LRCH1 is likely associated with at least one Mendelian disease. Because the excerpt lacks any mention of LRCH1 or direct gene-level evidence (variant description, segregation, recurrence, functional validation tied to LRCH1), it cannot be used to support or refute the claim. The sentence only offers a general mechanistic observation about how rare coding variants can act in families, which is relevant background but insufficient to establish a LRCH1–Mendelian disease association. For a relevant assessment, the paper would need to explicitly identify LRCH1 (or a variant in LRCH1), present familial segregation or case series consistent with Mendelian inheritance, or provide functional evidence linking LRCH1 disruption to a Mendelian phenotype.


[Read Paper](https://www.semanticscholar.org/paper/c2397a9298b1470564bc85303532224d8bf43ab3)


### Sequencing individual genomes with recurrent genomic disorder deletions: an approach to characterize genes for autosomal recessive rare disease traits

**Why Not Relevant**: The provided paper content contains a single, general methodological statement: "The utility of NAHR recurrent deletions to enhance discovery in the challenging context of autosomal recessive (AR) traits and biallelic variation is demonstrated and NAHR events could potentially enable human haploid genetic screens as an approach to experimental inquiry into disease biology." This sentence does not mention LRCH1, any specific gene, any clinical cases, variant data, linkage/association statistics, functional assays, or phenotype descriptions. Therefore it provides no direct evidence that LRCH1 is associated with any Mendelian disease, nor mechanistic evidence specific to LRCH1. 

Although the sentence describes a general mechanism (NAHR recurrent deletions can aid discovery of autosomal recessive traits and could enable haploid genetic screens), that is a methodological or conceptual point about how to discover gene-disease relationships; it does not supply data, examples, or analyses linking LRCH1 to disease. In short, the content is relevant at the level of general strategy for gene discovery but contains zero specific evidence (direct or mechanistic) regarding LRCH1. 

Limitations that justify the relevance score of 0.0: 
- No mention of LRCH1 by name or by genomic coordinate. 
- No reported patient(s), variants, segregation, case-control, or population data implicating LRCH1. 
- No functional assays or mechanistic experiments performed on LRCH1. 
- No statistical associations, effect sizes, or confidence intervals relating LRCH1 to disease. 

To evaluate the claim that "LRCH1 is likely associated with at least one Mendelian disease," one would need evidence such as: documented pathogenic variants in LRCH1 segregating with a Mendelian phenotype, case reports linking biallelic or dominant variants to consistent clinical phenotypes, robust genetic association or linkage, or functional experiments showing a causative mechanism. None of these are present in the provided content.


[Read Paper](https://www.semanticscholar.org/paper/82356870534f47fd080d37bdf3980d65b074ecab)


### Whole-exome sequencing of pathogenic genes in a family with congenital heart disease: A case report

**Why Not Relevant**: The paper does not mention LRCH1 anywhere in the provided content. All reported sequence variants and candidate genes in this study are AMER1 and KCNE1 (and two unspecified variants that did not fit inheritance patterns). Therefore this paper provides no direct evidence that LRCH1 is associated with any Mendelian disease. Mechanistically, the paper offers no information about LRCH1 function, variant effects, segregation, or pathogenicity assessments that could support LRCH1's involvement. The study is a single-family CHD exome analysis and while it uses an appropriate method for Mendelian gene discovery, its negative relevance to LRCH1 is definitive in the context of this paper: absence of mention equals absence of supporting evidence. Note that the lack of LRCH1 evidence here does not disprove LRCH1 involvement in other studies or cohorts, but this paper contributes nothing for or against the specific claim regarding LRCH1.


[Read Paper](https://www.semanticscholar.org/paper/2afe5ddc8ce353c6339a8bb041306ea8d115518f)


### Whole exome sequencing identified a pathogenic nonsense mutation in LMNA in a family with a progressive cardiac conduction defect: A case report

**Why Not Relevant**: The paper describes identification of a nonsense mutation in the LMNA gene (c.1443C>A, p.Tyr481*) as a potential cause of progressive cardiac conduction defect (PCCD) in a Chinese family. The content repeatedly references LMNA as the implicated gene and does not mention LRCH1 at any point. Because the claim asks about LRCH1 being associated with at least one Mendelian disease, and this paper provides genetic, phenotypic, and in silico evidence only for LMNA, there is no direct evidence (no observed LRCH1 variants, no segregation data, no functional or mechanistic discussion of LRCH1) nor mechanistic evidence (no pathway, expression, interaction, or functional data involving LRCH1) to support or refute the claim. In short, the study is relevant to LMNA and PCCD but contains no information about LRCH1; therefore it cannot be used to assess whether LRCH1 is likely associated with a Mendelian disease.


[Read Paper](https://www.semanticscholar.org/paper/1bda863d40686845df5ee3f07e8920b140ce554e)


### A nonsense variant in KRT31 is associated with autosomal-dominant monilethrix.

**Why Not Relevant**: This paper does not provide evidence for or against the claim about LRCH1. The study focuses on monilethrix and identifies pathogenic variants in KRT31 (and references known genes KRT81, KRT83, KRT86, and DSG4). The text contains no mention of LRCH1, no data (genetic, statistical, or functional) involving LRCH1, and no mechanistic discussion that implicates LRCH1 in monogenic disease. Instead, the paper reports direct genetic and cellular evidence implicating KRT31: exome sequencing in affected individuals identified a nonsense variant c.1081G>T; p.(Glu361*) in KRT31, Sanger sequencing showed segregation in families, and cell-based assays (immunoblotting, immunofluorescence, RT-qPCR) were used to characterize the mutant keratin 31 protein. Because LRCH1 is not mentioned and no results concern LRCH1, this paper supplies no relevant direct or mechanistic evidence bearing on the claim that LRCH1 is likely associated with at least one Mendelian disease. Limitations: absence of evidence in this paper is not evidence of absence for LRCH1 overall, but for the purpose of evaluating LRCH1 specifically, this paper is irrelevant.


[Read Paper](https://www.semanticscholar.org/paper/70feee1879ff5c8a41c5624a6e6398f7d98ce27e)


### Exome Sequencing Identifies a Novel SGCA Gene Mutation in an Iranian Family with Limb Girdle Muscular Dystrophy: A Case Report

**Why Not Relevant**: This paper does not mention LRCH1 at all. The entire report describes a case of autosomal recessive limb-girdle muscular dystrophy in which whole exome sequencing identified a novel homozygous frameshift mutation in the SGCA gene (p.A107fs:c.319-329del) and concludes that the SGCA variant is pathogenic for LGMD. There is no direct evidence (no LRCH1 variant reported, no association analyses, no phenotypic correlation involving LRCH1) and no mechanistic discussion relating LRCH1 to disease. Therefore the paper provides neither supporting nor refuting evidence for the claim that LRCH1 is likely associated with a Mendelian disease. Note also that absence of mention is not evidence of absence: the paper does not perform a negative association study of LRCH1 or state that LRCH1 was examined and found normal. It simply does not address LRCH1, so it is irrelevant to the claim.


[Read Paper](https://www.semanticscholar.org/paper/46a6274976aa6da24830f44d465c15762c6beb3f)


## Search Queries Used

- LRCH1 mutation pathogenic variant case report family segregation exome sequencing Mendelian disease

- LRCH1 ClinVar OMIM DECIPHER gnomAD pLI loss of function intolerance constraint database annotation

- leucine rich repeat calponin homology domain containing 1 LRCH1 function protein interaction cytoskeleton immune cell migration knockout mouse zebrafish phenotype

- LRCH1 rare variant burden sequencing family based exome sequencing autosomal dominant autosomal recessive monogenic rare disease study


## Bibliometric Configuration

- Use Bibliometrics: True

- Author Impact Weight: 0.4

- Citation Impact Weight: 0.4

- Venue Impact Weight: 0.2
