## Supplementary material for "MENDELSEEK: An algorithm that predicts Mendelian Genes and elucidates what makes them special": NPR1.txt

# Claim: The gene NPR1 is likely associated with at least one Mendelian disease.

**Status**: processed

**Overall Rating**: -1

**Explanation**:

No relevant papers were found that support or refute this claim.


## Relevant Papers


## Other Reviewed Papers


### Pathogenesis of Hypertension

**Why Not Relevant**: The provided paper is a clinical review of hypertension pathophysiology and genetics. It discusses natriuretic peptides as part of blood-pressure regulation and summarizes that single-gene (Mendelian) causes of blood-pressure disorders exist, but it does not mention the NPR1 gene by name, present any genetic, clinical, linkage, mutation, or association data for NPR1, nor does it list NPR1 among the known Mendelian hypertension/hypotension genes. Therefore the paper provides only indirect mechanistic plausibility (natriuretic-peptide pathway importance) but no direct or specific evidence that NPR1 is associated with any Mendelian disease. Because the claim is about NPR1 being likely associated with at least one Mendelian disease, and the paper contains no gene-specific evidence for NPR1, the content is not relevant as direct support for the claim.


[Read Paper](https://www.semanticscholar.org/paper/70997d8bdccb350467a97b91fad1a779480ea1bb)


### Identification and characterization of two SERPINC1 mutations causing congenital antithrombin deficiency

**Why Not Relevant**: The provided paper content does not mention NPR1 at all and therefore provides no direct or mechanistic evidence for the claim that "The gene NPR1 is likely associated with at least one Mendelian disease." The only sentence supplied is: "The p.Asn460Thrfs*20 and p.Arg229* variants of SERPINC1 were responsible for the two hereditary AT deficiency pedigrees, which led to AT deficiency by different mechanisms, and the structure and function of the mutations were analyzed." This sentence refers to SERPINC1 and hereditary antithrombin (AT) deficiency, a Mendelian condition caused by SERPINC1 variants, but it does not relate to NPR1. Limitations and caveats: (1) the content provided is a single sentence about SERPINC1; it contains no mention of NPR1, its variants, clinical phenotypes, or mechanistic links to Mendelian disease. (2) Because only this excerpt was supplied, it is not possible to assess the remainder of the paper for any potential NPR1-relevant data — if NPR1 appears elsewhere in the full paper, that content was not provided. (3) To evaluate the claim about NPR1, one would need human genetic evidence (segregation in pedigrees, pathogenic variants reported in databases such as OMIM/ClinVar, case reports), functional/mechanistic studies demonstrating variant impact, or population genetics evidence; none of these are present in the supplied text. Recommended next steps: search for primary literature, OMIM/ClinVar entries, and functional studies specifically about NPR1 variants and human disease to properly assess the claim.


[Read Paper](https://www.semanticscholar.org/paper/136919dd91daf7190dadf5efa3b5051530061308)


### A cryptic splice-altering KCNQ1 variant in trans with R259L leading to Jervell and Lange-Nielsen syndrome

**Why Not Relevant**: The provided paper content focuses entirely on a clinical case of Jervell and Lange-Nielsen syndrome and characterization of a cryptic splice-altering variant in KCNQ1. There is no mention of the gene NPR1, no discussion of diseases related to NPR1, and no mechanistic data linking NPR1 to Mendelian inheritance presented in the supplied text. Consequently, the paper content as provided does not provide direct or mechanistic evidence for or against the claim that NPR1 is likely associated with at least one Mendelian disease. Caveats: this assessment is limited to the excerpt you supplied. If the full paper contains sections not provided here that discuss NPR1, those were not available for review; based on the available content, relevance to NPR1 is effectively nil.


[Read Paper](https://www.semanticscholar.org/paper/a4d6ea79ec9b18da106075d8c4c76a9c47c29eaf)


### Novel variant of KIF11 associated with MCLMR syndrome

**Why Not Relevant**: The paper is a case report focused on a novel heterozygous KIF11 frameshift variant causing a microcephaly-lymphedema-chorioretinal dysplasia (MCLMR/MLCRD) phenotype. The manuscript documents clinical findings, KIF11-targeted sequencing, identification of a de novo frameshift KIF11 variant, and plausible loss-of-function mechanism for KIF11. The gene NPR1 is not mentioned anywhere in the provided text, nor are any data, analyses, or mechanistic discussions related to NPR1 presented. Therefore this paper provides no direct evidence supporting or refuting the claim that NPR1 is likely associated with at least one Mendelian disease. Note: absence of discussion of NPR1 in a KIF11-focused case report is not evidence that NPR1 is not disease-associated; it only means this paper is not informative for the NPR1 claim. To evaluate the claim about NPR1, literature searches (OMIM, ClinVar, PubMed) or targeted genetic studies involving NPR1 would be required.


[Read Paper](https://www.semanticscholar.org/paper/598b1c7508865997c295d53a98d38407fbecde29)


### Complex PKD1 Genetics in Early-Onset Cystic Kidney Disease

**Why Not Relevant**: The provided paper content contains no mention of the gene NPR1 (nor of common synonyms such as NPRA, NPR-A, natriuretic peptide receptor A). The excerpt is exclusively focused on PKD1 and autosomal dominant polycystic kidney disease (ADPKD), variant phase (in cis vs in trans), interpretation of missense versus truncating variants, and the possible contribution of additional cystic-kidney genes or modifiers. Examples of the paper's topical focus include phrases such as "compound heterozygous PKD1 variants," "autosomal dominant polycystic kidney disease (ADPKD)", and "polycystin-1-pol ycystin-2" (all from the provided text). Because none of the sentences or paragraphs in the supplied content address NPR1, its variants, human phenotypes linked to NPR1, segregation data, functional assays, animal models, ClinVar/OMIM entries, or other evidence types used to implicate a gene in Mendelian disease, there is no direct evidence in this excerpt supporting or refuting the claim that "NPR1 is likely associated with at least one Mendelian disease."

The excerpt does contain general mechanistic discussion that is broadly relevant to how genes are evaluated for Mendelian disease causation (for example, distinguishing loss-of-function/truncating variants from missense/hypomorphic variants and discussing compound heterozygosity and digenic inheritance). However, these are generic points about variant interpretation and do not provide mechanistic evidence specific to NPR1. Therefore they do not strengthen or weaken the specific claim about NPR1.

Limitations: this assessment is strictly based on the supplied text. If NPR1-related data appear elsewhere in the full article (supplements, figures, or sections not included here), that content was not available for review. If you would like, I can (a) search literature/databases (OMIM, ClinVar, PubMed) for evidence linking NPR1 to Mendelian disease, or (b) re-check additional sections of the paper if you can provide them.


[Read Paper](https://www.semanticscholar.org/paper/93aa5634e2918da0adf9d4d1273325fd11dae0af)


## Search Queries Used

- NPR1 NPRA guanylate cyclase A mutation family segregation Mendelian monogenic case report trio exome sequencing

- NPR1 ClinVar OMIM pathogenic variant loss of function truncating nonsense frameshift missense rare variant clinical report

- NPR1 knockout mouse natriuretic peptide receptor A cGMP signalling functional study cardiac hypertrophy salt sensitive hypertension loss of function assay

- NPR1 genetic association familial hypertension hereditary cardiomyopathy review meta analysis linkage family based study natriuretic peptide receptor 1


## Bibliometric Configuration

- Use Bibliometrics: True

- Author Impact Weight: 0.4

- Citation Impact Weight: 0.4

- Venue Impact Weight: 0.2
