## Supplementary material for "MENDELSEEK: An algorithm that predicts Mendelian Genes and elucidates what makes them special": NPY.txt

# Claim: The gene NPY is likely associated with at least one Mendelian disease.

**Status**: processed

**Overall Rating**: -1

**Explanation**:

No relevant papers were found that support or refute this claim.


## Relevant Papers


## Other Reviewed Papers


### The Burden of Candidate Pathogenic Variants for Kidney and Genitourinary Disorders Emerging From Exome Sequencing

**Why Not Relevant**: The provided paper content does not mention the gene NPY nor present any data, results, or curated-gene lists (within the quoted text) that include or exclude NPY. The manuscript excerpt focuses on methods for identifying and curating genes associated with Mendelian kidney and genitourinary disorders, cohort characteristics, sequencing and variant-filtering pipelines, and high-level results about the prevalence of candidate pathogenic variants in that cohort, but contains no statements about NPY or any evidence linking NPY to a Mendelian disease. Because no direct or mechanistic evidence about NPY is present in the supplied text, this paper excerpt cannot be used to support or refute the claim that “NPY is likely associated with at least one Mendelian disease.”

Caveats: absence of any mention of NPY in the provided excerpt is not evidence that the full paper or its supplementary materials do or do not include NPY; the authors note that a full curated gene list is available on the laboratory website (not included here). To determine whether NPY is listed by these authors as associated with a Mendelian disease, one would need to consult the paper’s full gene list (supplementary materials or laboratory website) or primary gene-disease databases (OMIM, ClinVar, Orphanet, HGMD).


[Read Paper](https://www.semanticscholar.org/paper/a75e297208c04992568e607d19aa9b9aaebd09d5)


### mGAP: the macaque genotype and phenotype resource, a framework for accessing and interpreting macaque variant data, and identifying new models of human disease

**Why Not Relevant**: The provided paper content does not mention the gene NPY, nor does it present any data, statistical analyses, case reports, or mechanistic discussion linking NPY to any Mendelian disease. The single excerpt only describes a public macaque variant database (mGAP) and its potential to enable future investigations of genetic associations. While such a resource could facilitate discovery of gene–disease links in the future, the content contains no direct or indirect evidence that NPY is associated with at least one Mendelian disease. Therefore the paper content is not relevant to supporting or refuting the claim.


[Read Paper](https://www.semanticscholar.org/paper/f47691ff666dd65548f1282e606d696eae56524d)


### Simultaneous identification and prioritization of variants in familial, de novo, and somatic genetic disorders with VariantMaster

**Why Not Relevant**: The provided paper content describes a computational tool (VariantMaster) and its applications to identify causative variants in Mendelian diseases and somatic driver mutations in cancer, but it contains no mention of the specific gene NPY, nor any gene-level results or lists of implicated genes. Therefore there is no direct evidence (e.g., observed pathogenic NPY variants, statistical association, or clinical cases implicating NPY) nor mechanistic, gene-specific evidence connecting NPY to any Mendelian disease. The content only shows that the methodology could, in principle, detect such an association if NPY variants were present in the analyzed data, but that is speculative and does not support the claim. Because the paper provides no NPY-specific data, the relevance to the claim that "NPY is likely associated with at least one Mendelian disease" is negligible.


[Read Paper](https://www.semanticscholar.org/paper/a56e2dc7369c76b03ad5e9309093beb98a159719)


### Mc3 and Mc4 receptors: complementary role in weight control.

**Why Not Relevant**: The paper provides mechanistic context placing neuropeptide Y (NPY) within the leptin–hypothalamic regulatory circuit for appetite and energy homeostasis and gives examples of other genes in the same pathway (POMC, MC4R) that have known human genetic links to obesity. However, it contains no direct evidence that variation in the NPY gene causes a Mendelian disease: there are no reports of NPY mutations, no human genotype–phenotype segregation data, no case descriptions, and no functional genetic studies linking NPY variants to a Mendelian (single-gene, typically highly penetrant) disorder. Therefore the paper offers only indirect mechanistic plausibility (NPY is biologically relevant to appetite control) but no direct support for the claim that NPY is likely associated with at least one Mendelian disease. Given the absence of any NPY-specific genetic or clinical data in this paper, the relevance to the claim is very low.


[Read Paper](https://www.semanticscholar.org/paper/4c9c0cc72605199c05df6f7cab02717f69f6c169)


### Precision Medicine in Internal Medicine

**Why Not Relevant**: The provided paper is a general review of clinical genetic testing methods, variant interpretation, and the state of gene discovery. It does not mention the gene NPY anywhere in the supplied text, reports no data (genetic, clinical, or experimental) linking NPY to any Mendelian (single-gene) disorder, and supplies no mutation, segregation, functional, or epidemiologic evidence about NPY. Therefore the paper provides no direct evidence for or against the claim that NPY is associated with at least one Mendelian disease. The included sentences are contextual background about the fraction of genes currently linked to Mendelian diseases and about the capabilities and limitations of exome sequencing and variant interpretation; they can only speak to the general possibility that undiscovered associations (such as one involving NPY) might be found, not to any concrete association for NPY. Because no NPY-specific content, variants, phenotypes, segregation analyses, functional studies, or database annotations for NPY are presented, this paper cannot be used to evaluate the claim about NPY.


[Read Paper](https://www.semanticscholar.org/paper/c7199e8a5226f8d9d55777da0e15e91797bb459a)


### Adropin – a circulating factor in metabolic control or a drop in the ocean?

**Why Not Relevant**: The paper exclusively studies the ENHO gene product (adropin) and its role in metabolic physiology, using expression profiling and engineered knockout mice. There is no mention of the NPY gene, NPY protein, or any data connecting NPY to disease. The experimental evidence is animal-based (mouse knockouts and phenotype assays) demonstrating that loss of ENHO affects metabolic traits (fat mass, insulin resistance, glucose intolerance, dyslipidemia). These are complex metabolic phenotypes and do not constitute evidence that any gene (including NPY) is associated with a Mendelian disease in humans. The paper contains no human genetic linkage, segregation, case reports, variant pathogenicity assessments, or clinical descriptions of Mendelian inheritance for NPY or any other gene. Therefore the paper provides no direct or mechanistic evidence relevant to the claim that NPY is likely associated with at least one Mendelian disease.


[Read Paper](https://www.semanticscholar.org/paper/2322943d6c5826fe195c2fd207a44f3709e2aa23)


### Advancing long-read nanopore genome assembly and accurate variant calling for rare disease detection

**Why Not Relevant**: The paper does not mention the gene NPY anywhere in the provided content and provides no gene-specific results or lists that include NPY. All quoted material is either general context (diagnostic gap), methodological (cohort, coverage), or generic mechanistic/empirical findings about LRS's ability to detect variant classes and solve cases. While these mechanistic points explain how LRS could discover Mendelian associations for genes (including NPY) in principle, they do not provide direct evidence that NPY is associated with any Mendelian disease. Because there is no explicit observation, case report, variant, statistical association, or named-gene result involving NPY in the text supplied, this paper cannot be used as evidence for or against the claim that "The gene NPY is likely associated with at least one Mendelian disease." The absence of a gene list or named solved genes in the supplied excerpt is a key limitation: even though the study reports that LRS covered additional Mendelian disease genes and solved cases, without the specific gene identities one cannot evaluate NPY's involvement. Therefore the relevance to the specific claim is minimal.


[Read Paper](https://www.semanticscholar.org/paper/9ce7b266983316c2aa4955fbf3a61f90c3fd1785)


### Session Overview: Bone as an Endocrine Organ

**Why Not Relevant**: Relevance to the claim is very low because the provided text contains only physiological and mechanistic discussion of NPY's role in hypothalamic signaling, appetite regulation, and effects on bone in animal-model contexts. The paper content (a conference-session summary) does not present any genetic association data, case reports, familial segregation analyses, mutation descriptions, or clinical phenotypes that would indicate NPY is causally associated with a Mendelian disease. Mechanistic involvement of a gene in a physiological pathway does not by itself establish that pathogenic variants in that gene cause a Mendelian disorder. Additionally, the excerpts refer mainly to animal-model experiments and secondary summaries (conference talks) without methods, quantitative results, or evidence of heritable, monogenic disease, so they cannot be interpreted as direct evidence for the claim.


[Read Paper](https://www.semanticscholar.org/paper/2b15111783c0c7c05f26664bb17e22d5ceacfbd9)


### Genetic basis of Parkinson's disease and related disorders

**Why Not Relevant**: This paper does not mention the gene NPY anywhere in the provided text. The study is focused on Parkinson disease genetics (familial and sporadic PD), reporting discoveries and association results for LRRK2, PRKRA, SNCA, MAPT and other PD-relevant loci, and discusses methods (haplotype analysis, autozygosity mapping, GWAS, CNV detection) applied in PD cohorts. Because the claim concerns whether NPY is likely associated with at least one Mendelian disease, and the paper neither reports any data on NPY nor assesses Mendelian diseases beyond specific PD-related families/cohorts, the paper provides no direct evidence for or against the claim. The few relevant passages are only indirect: they list genes implicated in the authors' PD studies (and omit NPY), or describe methodological limitations (limited power, disease focus) that further reduce the ability to infer anything about NPY. Therefore the paper's relevance to the claim is negligible. Limitations of this assessment: absence of NPY in this PD-focused paper does not constitute evidence that NPY is not associated with some Mendelian disease elsewhere; establishing that would require literature searches or data specifically analyzing NPY in the context of Mendelian disorders.


[Read Paper](https://www.semanticscholar.org/paper/b59ed2e51a73f5ce5958c18560d750b40e4034b6)


### BONE AS IT IS KNOWN TO BE : A TAKER Coordinated Control of Bone Mass and Energy Metabolism : The Viewpoint

**Why Not Relevant**: This paper is a review of bone endocrinology that focuses on leptin, the sympathetic nervous system (Adrβ2), and CART as mediators linking central pathways to osteoblasts and bone mass regulation. The provided text contains mechanistic and physiological discussion of leptin and related mediators in mouse and comparative physiology, but it does not mention the gene NPY (neuropeptide Y) or any association between NPY and human Mendelian diseases. There are no experimental results, genetic association data, case reports, or literature citations in the excerpt that relate NPY to Mendelian conditions. Because the paper neither discusses NPY nor provides human genetic disease evidence, it contains no direct or indirect (mechanistic) evidence supporting or refuting the claim that NPY is likely associated with at least one Mendelian disease. Therefore the relevance to the claim is effectively zero. Limitations: the conclusion is based on the provided excerpt only; if the full paper contains sections not included here that mention NPY or Mendelian disease associations, those were not available for review.


[Read Paper](https://www.semanticscholar.org/paper/4e4965dc438bac9b5a2dde9ba9f95e4a2f8296c5)


### #1761 Whole-genome sequencing reveals contribution of rare and common variation to structural kidney and urinary tract malformations

**Why Not Relevant**: The provided paper studies genetic causes of structural kidney and urinary tract malformations using whole-genome sequencing in a cohort of affected individuals. The text does not mention the NPY gene anywhere, nor does it present any direct experimental or statistical evidence for or against NPY being associated with a Mendelian disease. The findings and gene-level associations reported (AUTS2, ARHGAP5, HNF1B, ZNF879) are specific to the studied phenotypes. Therefore the paper is only indirectly relevant: the absence of NPY among reported hits in this kidney/urinary tract cohort offers weak, phenotype-limited negative evidence, but it cannot be used to conclude whether NPY is or is not associated with any Mendelian disease in general. Additional studies targeting NPY, other phenotypes, or larger/more diverse cohorts would be required to evaluate the claim directly.


[Read Paper](https://www.semanticscholar.org/paper/490f0cc0aac67ac259732ccd40d6cf231466b691)


## Search Queries Used

- NPY neuropeptide Y mutation Mendelian disease case report familial monogenic disorder autosomal dominant autosomal recessive

- neuropeptide Y loss of function human phenotype deficiency knockout mouse model phenotype appetite feeding obesity energy homeostasis endocrine regulation receptor pathways

- NPY ClinVar pathogenic variant OMIM entry HGMD gene NPY disease association gene annotation

- NPY rare variant exome sequencing whole genome sequencing Mendelian cohort undiagnosed disease diagnostic yield de novo variant rare inherited variant burden


## Bibliometric Configuration

- Use Bibliometrics: True

- Author Impact Weight: 0.4

- Citation Impact Weight: 0.4

- Venue Impact Weight: 0.2
