## Supplementary material for "MENDELSEEK: An algorithm that predicts Mendelian Genes and elucidates what makes them special": NTF3.txt

# Claim: The gene NTF3 is likely associated with at least one Mendelian disease.

**Status**: processed

**Overall Rating**: -1

**Explanation**:

No relevant papers were found that support or refute this claim.


## Relevant Papers


## Other Reviewed Papers


### Phenotypic and biochemical analysis of an international cohort of individuals with variants in NAA10 and NAA15.

**Why Not Relevant**: The paper's content exclusively addresses genetic variation, clinical phenotypes, and biochemical mechanisms related to NAA10 and NAA15 (the NatA complex) and does not mention the gene NTF3 anywhere in the provided text. Therefore this paper provides no direct evidence (genetic association, case reports, segregation, or population data) nor mechanistic evidence linking NTF3 to any Mendelian disease. The absence of NTF3 in this study means it cannot be used to support or refute the claim that NTF3 is likely associated with at least one Mendelian disease. Note: absence of evidence in this paper is not evidence of absence overall; other literature would need to be searched to evaluate the claim about NTF3.


[Read Paper](https://www.semanticscholar.org/paper/704ec885f2ae72a140397706ab74dc438e8605b1)


### Expanding the mutational spectrum of monogenic hypogonadotropic hypogonadism: novel mutations in ANOS1 and FGFR1 genes

**Why Not Relevant**: The provided paper content contains only a single sentence referring to ANOS1 and FGFR1 as classical CHH (congenital hypogonadotropic hypogonadism) genes and notes newly identified variants in those genes: "Despite the fact that ANOS1 and FGFR1 are classical CHH genes and were thoroughly explored in several CHH cohorts the authors identified new, yet undescribed variants within their sequence, which support the genetic complexity of the disorder." This sentence does not mention NTF3, does not present data, mechanisms, or discussion about NTF3, nor does it provide any evidence (direct or mechanistic) linking NTF3 to a Mendelian disease. Therefore the excerpt is not relevant to the claim that "The gene NTF3 is likely associated with at least one Mendelian disease." 

Limitations and impact on evidence quality:
- Absence of mention: The content does not mention NTF3 at all, so no inference about NTF3 can be drawn from it.
- Scope: The sentence focuses on ANOS1 and FGFR1 in CHH cohorts; these are different genes and a different disorder context than the claim about NTF3.
- Insufficient data: No experimental methods, results, mechanistic descriptions, statistical evidence, or clinical case reports related to NTF3 are provided.

Conclusion: Based solely on the provided paper content, there is no direct or mechanistic evidence to support or refute the claim that NTF3 is associated with at least one Mendelian disease. To evaluate the claim, additional sources that explicitly discuss NTF3 (genetic studies, case reports, functional/mechanistic work, or curated disease-gene databases) would be required.


[Read Paper](https://www.semanticscholar.org/paper/dcceaee0bd8232b5abfc3d2d144738c98d42f76e)


### TTC7A Variants Previously Described to Cause Enteropathy Are Observed on a Single Haplotype and Appear Non-pathogenic in Pediatric Inflammatory Bowel Disease Patients

**Why Not Relevant**: The provided paper content focuses exclusively on TTC7A (Tetratricopeptide Repeat Domain 7A) and its variants in relation to polygenic inflammatory bowel disease, with mention of compound heterozygote variants observed in trans. The claim is about NTF3 and whether that gene is likely associated with at least one Mendelian disease. There is no mention of NTF3, no data or analyses concerning NTF3, and no mechanistic discussion linking NTF3 to Mendelian inheritance or disease in the provided content. While the excerpt references a Mendelian concept (compound heterozygosity in trans), this applies to TTC7A only and cannot be generalized to NTF3 without additional evidence. Therefore the paper content as provided is not relevant to evaluating the claim about NTF3. Additional reasons limiting relevance: (1) gene mismatch (TTC7A vs NTF3), (2) phenotype mismatch (polygenic IBD context rather than explicit Mendelian disease attribution), and (3) absence of mechanistic or statistical data about NTF3. Given these points, the relevance to the claim is effectively zero based on the supplied content.


[Read Paper](https://www.semanticscholar.org/paper/3818a966b5d6e8b0eda65fdef043ac0727fa429c)


### A case of neurodegenerative disorder caused by PLA2G6 compound heterozygous pathogenic variant and SCA17 pathogenic variant

**Why Not Relevant**: The provided paper content exclusively reports a case attributed to a pathogenic variant in PLA2G6 and contains no mention of the gene NTF3, any of its variants, phenotypes linked to NTF3, or mechanistic links between PLA2G6 and NTF3. There is therefore no direct or mechanistic evidence in this content that bears on the claim that NTF3 is associated with at least one Mendelian disease. Because the paper addresses a different gene (PLA2G6) and provides no data, discussion, or cross-reference to NTF3, it is not relevant to evaluating the claim about NTF3.


[Read Paper](https://www.semanticscholar.org/paper/aca969072f60778a8fc52afb27352aad4277cdf7)


### Congenital hypothyroidism and hearing loss without inner ear malformation: Think TPO

**Why Not Relevant**: The paper exclusively investigates TPO variants as the cause of thyroid dyshormonogenesis type 2A associated with hearing impairment. NTF3 is not mentioned anywhere in the provided text. The study's findings implicate TPO variants in the reported families and explicitly state that no other causal variants in genes associated with hearing impairment were identified in these cases; however, that negative statement applies only to this small, phenotype-specific cohort and does not constitute a systematic evaluation of NTF3 across Mendelian disorders. Exome sequencing may miss certain variant types and the sample size is small, so absence of evidence for NTF3 here is weak and not generalizable. Therefore the paper provides no substantive evidence for or against the broader claim that NTF3 is likely associated with at least one Mendelian disease.


[Read Paper](https://www.semanticscholar.org/paper/5bd4bf8d39ead9d9dbda04ab33519a204899cd57)


### Abstract. Kindler syndrome (KS) is a rare subtype of epider‐ molysis bullosa that is inherited in an autosomal recessive manner with mutations in FERMT1. A number of mutations

**Why Not Relevant**: This paper is not relevant to the claim that NTF3 is likely associated with at least one Mendelian disease. The manuscript exclusively addresses Kindler syndrome and variants in FERMT1 (kindlin-1). It reports a novel homozygous frameshift mutation c.1885_1901del (p.Val629fs) in FERMT1 in a single Chinese patient and summarizes previously reported pathogenic FERMT1 variants; it contains no mention of the gene NTF3, no genetic findings involving NTF3, and no mechanistic discussion linking NTF3 to any Mendelian phenotype. Although the study used a multi-gene panel of 541 loci, the paper only reports a FERMT1 pathogenic variant in the patient and does not report NTF3 as tested, mutated, or implicated. Therefore the article provides neither direct evidence (no reported NTF3 pathogenic variants or case associations) nor mechanistic evidence (no functional data, pathways, or biological links) supporting an association between NTF3 and a Mendelian disease. Caveats: absence of any mention of NTF3 in this paper does not disprove the claim — it only means this paper does not inform the claim. To evaluate the claim one needs papers that specifically report NTF3 mutations segregating with Mendelian phenotypes, functional studies of NTF3 loss/gain of function causing disease, or entries in curated disease-gene databases (e.g., OMIM, ClinGen) that implicate NTF3.


[Read Paper](https://www.semanticscholar.org/paper/852363faf04651740d0a2132e4b5ffca67b9ad9e)


### 1684: PATHOGENIC VARIATION ASSOCIATED WITH MONOGENIC DISORDERS IN A SEVERE PEDIATRIC SEPSIS COHORT

**Why Not Relevant**: The paper does not mention NTF3, nor does it present a gene list that would allow determination of whether NTF3 was examined. The study analyzed a defined panel of 96 genes and reports pathogenic variants found among those genes, but the reported results and conclusions enumerate specific disorders and genes identified (e.g., PIK3R1) without any reference to NTF3. Therefore the paper provides no direct evidence that NTF3 is associated with a Mendelian disease. Because absence of mention does not prove absence of association, this paper cannot be used as evidence for or against the claim: it is simply non-informative with respect to NTF3. Additional limitations reducing relevance: small cohort (N=45), targeted gene panel (not whole-genome reporting here), and lack of a full gene list in the excerpt provided.


[Read Paper](https://www.semanticscholar.org/paper/cdd32cc853ff52097298692a62266f79b44b8c53)


## Search Queries Used

- NTF3 mutation pathogenic variant Mendelian disease case report neurotrophin 3 autosomal dominant autosomal recessive monogenic

- neurotrophin 3 NTF3 knockout mouse phenotype sensory neuron development peripheral neuropathy TrkC NTRK3 mechanistic study

- NTF3 ClinVar OMIM gnomAD loss of function intolerance haploinsufficiency pLI gene constraint database annotation

- neurotrophin 3 de novo variant exome sequencing gene discovery intellectual disability autism congenital hearing loss hereditary neuropathy


## Bibliometric Configuration

- Use Bibliometrics: True

- Author Impact Weight: 0.4

- Citation Impact Weight: 0.4

- Venue Impact Weight: 0.2
