## Supplementary material for "MENDELSEEK: An algorithm that predicts Mendelian Genes and elucidates what makes them special": NTNG1.txt

# Claim: The gene NTNG1 is likely associated with at least one Mendelian disease.

**Status**: processed

**Overall Rating**: -1

**Explanation**:

No relevant papers were found that support or refute this claim.


## Relevant Papers


## Other Reviewed Papers


### Discovery and functional prioritization of Parkinson’s disease candidate genes from large-scale whole exome sequencing

**Why Not Relevant**: The provided paper content does not mention NTNG1 or provide any gene-level detail that would allow assessment of NTNG1's association with a Mendelian disease. The single sentence supplied only states a broad result—that 27 genes with homozygous or compound heterozygous loss-of-function variants were identified in PD cases and that several PD susceptibility candidates were nominated—without listing the genes, presenting segregation or family-based evidence, specifying whether identified variants cosegregate with a Mendelian phenotype, or describing any Mendelian disease diagnoses. Therefore the content cannot be used as direct evidence that NTNG1 is associated with a Mendelian disease. Furthermore, identifying homozygous/compound heterozygous LoF variants in cases is suggestive but not sufficient to establish a Mendelian disease association: establishing Mendelian causality generally requires cosegregation in families, strong statistical association and replication, functional validation demonstrating a penetrant effect, and clear phenotype matching. None of these items specific to NTNG1 are present in the supplied text, so relevance to the NTNG1 claim is negligible.


[Read Paper](https://www.semanticscholar.org/paper/489d796282e50673b7189ad7a170224d5ce73767)


### Effective variant filtering and expected candidate variant yield in studies of rare human disease

**Why Not Relevant**: The provided paper content contains only a single, general methodological sentence about variant filtering and expected candidate counts by inheritance mode. It includes no mention of the gene NTNG1, no reported variants in NTNG1, no clinical case reports or family segregation data, no functional or mechanistic studies involving NTNG1, and no statistical association results. Therefore, based on the supplied content, the paper offers no relevant evidence (neither direct nor mechanistic) bearing on whether NTNG1 is associated with any Mendelian disease. If additional portions of the paper exist that specifically discuss NTNG1 (e.g., variant lists, gene-level results, case descriptions, functional assays), those would need to be provided to reassess relevance. As is, the content is insufficient to evaluate the claim.


[Read Paper](https://www.semanticscholar.org/paper/8da5eda5547e3521df7aecc8c8e4b7380a7641fa)


### Exploring the Impact of Genetics in a Large Cohort of Moebius Patients by Trio Whole Exome Sequencing

**Why Not Relevant**: The provided paper content does not mention NTNG1 anywhere and provides no direct genetic, clinical, or mechanistic data about NTNG1. The study focuses on trio whole-exome sequencing (WES) in a cohort of 37 patients with Moebius syndrome (MBS) and MBS-like presentations, reporting that no de novo variants were found in REV3L and PLXND1 and that WES failed to identify mutations possibly associated with MBS. None of those statements support or refute an association between NTNG1 and any Mendelian disease. 

Specific reasons this paper is not relevant to the claim that "NTNG1 is likely associated with at least one Mendelian disease":
- Absence of any mention of NTNG1: The text does not list NTNG1 among candidate genes, findings, or discussion points, so there is no direct genetic evidence linking NTNG1 to MBS or to any other Mendelian condition in this paper.
- Scope limited to Moebius syndrome: The cohort and conclusions pertain specifically to MBS and MBS-like phenotypes. Even if the paper had negative findings for genes examined, that would not provide evidence about NTNG1's involvement in other Mendelian diseases.
- Negative/neutral WES findings do not imply anything about NTNG1: The authors report that WES "failed to identify mutations possibly associated with MBS," which is a negative result for causative genes in this cohort but does not exclude NTNG1 as a cause of other Mendelian disorders or even as an untested candidate for MBS. There is no indication that NTNG1 was included in the in silico candidate panel or was assessable in these analyses.
- Methodological limitations prevent strong inferences about genes not mentioned: WES can miss structural variants, deep intronic or regulatory variants, repeat expansions, mosaic variants, and regions with poor coverage. The study size (n=37) and focus on a specific phenotype limit generalizability to broader claims about gene–disease associations.

Because the paper provides no direct evidence (positive or negative) nor mechanistic discussion relevant to NTNG1, it cannot be used to support or refute the claim that NTNG1 is likely associated with at least one Mendelian disease. To evaluate that claim, one would need studies that explicitly analyze NTNG1 variants in patients with Mendelian phenotypes, functional studies linking NTNG1 perturbation to disease-relevant pathways, or curated clinical/variant-databases reporting pathogenic NTNG1 variants and associated phenotypes.


[Read Paper](https://www.semanticscholar.org/paper/226b561a7e847eca27fae7e2558a187686dd8f75)


### The Burden of Candidate Pathogenic Variants for Kidney and Genitourinary Disorders Emerging From Exome Sequencing

**Why Not Relevant**: This paper focuses on identifying and evaluating genes associated with Mendelian kidney and genitourinary disorders and on the prevalence and misclassification of purportedly pathogenic variants in a curated set of such genes. The provided content does not mention NTNG1 anywhere, does not show the full curated gene list, and does not report findings for genes outside the kidney/genitourinary scope. Therefore the paper fragment contains no direct evidence (positive or negative) about whether NTNG1 is associated with at least one Mendelian disease. The methodological passages included are relevant only as background (how the authors defined and curated genes and variants) and cannot be used to support the claim about NTNG1. Given the absence of any mention of NTNG1 or data pertaining to it, the relevance of this paper excerpt to the specific claim is negligible.


[Read Paper](https://www.semanticscholar.org/paper/a75e297208c04992568e607d19aa9b9aaebd09d5)


### Letter regarding “ SCN9A variant in a family of mixed breed dogs with congenital insensitivity to pain”—Navigating the pathogenicity of candidate gene mutations: Spotlight on paralog Nav genes

**Why Not Relevant**: This document does not provide any evidence for or against the claim that NTNG1 is associated with a Mendelian disease. The entire text discusses a different gene (SCN9A) and evaluation of a specific SCN9A variant in dogs. Example verbatim sentences from the provided content that show the paper's focus are: "We read the recent publication by Dr. Gutierrez-Quintana et al. titled “SCN9A variant in a family of mixed breed dogs with congenital insensitivity to pain” with great interest." and "In their study, the authors conducted whole-genome sequencing analysis of a dog with congenital insensitivity to pain and identified the pathogenic variant c.C2761T (p.R921C) in SCN9A, which encodes the alpha subunit of voltage-gated neuronal sodium channel Nav1.7 that is mainly expressed in nociceptive neurons." Nowhere in the provided text is NTNG1 mentioned (neither the gene symbol NTNG1 nor the name Netrin-G1 appear), nor are there data, mechanisms, functional studies, population frequencies, case reports, or family segregation analyses related to NTNG1. The paper instead critiques evidence for pathogenicity of a Nav1.7 (SCN9A) variant and discusses general issues of variant interpretation (e.g., in-silico prediction limits, ACMG-AMP guideline considerations), which are methodologically relevant to genetic variant assessment but do not provide evidence about NTNG1. Because no NTNG1-specific statements, data, or mechanisms are present in the supplied content, relevance to the claim is effectively zero.


[Read Paper](https://www.semanticscholar.org/paper/39dfc4195578b71ad782115a3a552b9b9e2dae03)


### Clingen Coagulation Factor Deficiency Variant Curation Expert Panel: Meeting the Need for Recommendations to Curate Variants in the Coagulation Factor Genes

**Why Not Relevant**: This paper does not provide evidence for or against the claim that NTNG1 is likely associated with at least one Mendelian disease. The content is focused on the development and specification of variant curation guidelines for coagulation factor genes (specifically F8 and F9) and on curation of variants in those genes. There is no mention of NTNG1, no presentation of genetic association data, no discussion of pathogenic variants in NTNG1, and no mechanistic data relating NTNG1 to any Mendelian phenotype. Because the paper's scope is limited to blood coagulation genes and methodological guideline specification, it cannot inform the NTNG1 claim. To evaluate the NTNG1 claim one would need primary genetic evidence (e.g., reported pathogenic variants in NTNG1 with segregation, de novo occurrence, or functional validation), database entries linking NTNG1 to Mendelian disorders (OMIM, ClinVar), case reports, or mechanistic studies implicating NTNG1 dysfunction in a Mendelian phenotype—none of which are present in this paper.


[Read Paper](https://www.semanticscholar.org/paper/9ea8d4ce19622ae1c6b0cdfd40848c93199f6280)


## Search Queries Used

- NTNG1 Netrin G1 pathogenic variant Mendelian disease monogenic mutation biallelic homozygous compound heterozygous family segregation case report

- NTNG1 ClinVar OMIM ClinGen HGMD ClinVar annotation pathogenic likely pathogenic clinical significance gene disease association review

- NTNG1 whole exome sequencing whole genome sequencing rare disease gene discovery trio sequencing de novo candidate gene autosomal recessive autosomal dominant large cohort

- NTNG1 Netrin G1 knockout mouse loss of function neuronal development synapse axon guidance intellectual disability epilepsy schizophrenia autism association case control functional characterization


## Bibliometric Configuration

- Use Bibliometrics: True

- Author Impact Weight: 0.4

- Citation Impact Weight: 0.4

- Venue Impact Weight: 0.2
