## Supplementary material for "MENDELSEEK: An algorithm that predicts Mendelian Genes and elucidates what makes them special": NUDT1.txt

# Claim: The gene NUDT1 is likely associated with at least one Mendelian disease.

**Status**: processed

**Overall Rating**: -1

**Explanation**:

No relevant papers were found that support or refute this claim.


## Relevant Papers


## Other Reviewed Papers


### ClinGen Myeloid Malignancy Variant Curation Expert Panel recommendations for germline RUNX1 variants.

**Why Not Relevant**: This paper focuses exclusively on developing and testing RUNX1-specific modifications to ACMG/AMP variant curation rules for inherited myeloid malignancies (familial platelet disorder with predisposition to acute myeloid leukemia). There is no mention of the gene NUDT1, no data on NUDT1 variants, no phenotype associations involving NUDT1, and no mechanistic discussion that would link RUNX1 findings to NUDT1. The methodological approaches described (gene-specific rule tailoring and variant curation workflows) are generally applicable and could be used in principle to evaluate NUDT1 in the future, but the paper provides no direct or mechanistic evidence to support or refute the claim that NUDT1 is associated with a Mendelian disease. Therefore the content is not relevant to the claim.


[Read Paper](https://www.semanticscholar.org/paper/d2a2dd2ebfb5a63969c41c03899f8f978fb18c6a)


### Re-evaluation of a Fibrillin-1 Gene Variant of Uncertain Significance Using the ClinGen Guidelines

**Why Not Relevant**: This paper is not relevant to the claim that NUDT1 is likely associated with at least one Mendelian disease. The study exclusively addresses FBN1 (fibrillin-1) variants in Marfan syndrome and contains no mention, data, or mechanistic discussion of NUDT1. Relevant verbatim sentences from the paper illustrating its focus: "Background Marfan syndrome (MFS) is caused by fibrillin-1 gene (FBN1) variants." "Using the Clinical Genome Resource (ClinGen) FBN1 variant curation expert panel (VCEP), we re-evaluated FBN1 germline variants reported as variants of uncertain significance (VUSs)." "Methods We re-evaluated 26 VUSs in FBN1 reported in 161 patients with MFS." "Results Four missense variants affecting cysteines (c.460T>C, c.1006T>C, c.5330G>C, and c.8020T>C) were reclassified as likely pathogenic and were assigned PM1_strong or PM1." "Conclusions ... The ClinGen FBN1 VCEP encompasses mutational hotspots and/or well-established critical functional domains and adjusts the criteria specifically for MFS; therefore, it is beneficial not only for identifying pathogenic FBN1 variants but also for distinguishing these variants from those that cause other connective tissue disorders with overlapping clinical features." Because the paper does not mention NUDT1, contains no variant or phenotype data for NUDT1, and discusses mechanisms and variant interpretation only in the context of FBN1 and Marfan syndrome, it provides neither direct evidence (observational/clinical genetic association) nor mechanistic evidence relevant to NUDT1. Therefore this paper cannot be used to support or refute the claim about NUDT1.


[Read Paper](https://www.semanticscholar.org/paper/bf6815f1239963b018387318ae7e6603ebb3e512)


## Search Queries Used

- NUDT1 MTH1 Nudix hydrolase 1 pathogenic variant Mendelian disease case report family segregation clinical phenotype

- NUDT1 ClinVar OMIM ClinGen germline mutation pathogenic review gene disease association

- NUDT1 loss of function knockout mouse zebrafish phenotype DNA repair 8 oxo dGTP oxidative stress neurodegeneration developmental disorder functional study

- NUDT1 exome sequencing rare variant burden monogenic disorder trio biallelic compound heterozygous gnomAD loss of function constraint pLI population genetics


## Bibliometric Configuration

- Use Bibliometrics: True

- Author Impact Weight: 0.4

- Citation Impact Weight: 0.4

- Venue Impact Weight: 0.2
