## Supplementary material for "MENDELSEEK: An algorithm that predicts Mendelian Genes and elucidates what makes them special": OPN4.txt

# Claim: The gene OPN4 is likely associated with at least one Mendelian disease.

**Status**: processed

**Overall Rating**: -1

**Explanation**:

No relevant papers were found that support or refute this claim.


## Relevant Papers


## Other Reviewed Papers


### The Burden of Candidate Pathogenic Variants for Kidney and Genitourinary Disorders Emerging From Exome Sequencing

**Why Not Relevant**: The paper is specifically focused on genes associated with Mendelian kidney and genitourinary disorders and on the problem of incidental and misclassified variant findings within that organ-system-specific gene set. The provided excerpts and the study description do not mention OPN4, nor do they analyze genes outside the kidney/genitourinary scope in a way that would provide direct or mechanistic evidence about OPN4. Therefore this paper does not provide relevant direct evidence that OPN4 is associated with at least one Mendelian disease. The absence of OPN4 from the provided text is not proof that OPN4 has no Mendelian associations; it only shows this study did not address that gene. To assess the claim about OPN4, one would need a paper that either lists OPN4 among curated Mendelian disease genes, reports pathogenic variants in OPN4 segregating with disease, or provides mechanistic evidence linking OPN4 dysfunction to a Mendelian phenotype. None of those types of evidence are present in the supplied content.


[Read Paper](https://www.semanticscholar.org/paper/a75e297208c04992568e607d19aa9b9aaebd09d5)


### P-540 Strategic implementation of PGT-M for multiple genes post qhole exome sequencing in couples with familial history: a comprehensive study

**Why Not Relevant**: This paper does not provide evidence for or against the claim that the gene OPN4 is likely associated with at least one Mendelian disease. The provided content describes a retrospective study using whole exome sequencing (WES) in couples with familial histories of genetic disorders, reporting overall diagnostic yield, consanguinity rates, and outcomes of PGT-M, but it contains no mention of the gene OPN4, no gene-level findings listing specific genes or variants (OPN4 is not named), and no mechanistic discussion involving OPN4. Specific points supporting non-relevance: (1) the text repeatedly summarizes cohort-level methods and results (e.g., number of families with reportable pathogenic variants, diagnostic confirmation rate) without listing the causal genes or variants; (2) there is no discussion of any disease linked to OPN4, nor any functional or mechanistic data about OPN4; (3) absence of OPN4 in the manuscript text means this paper neither supports nor refutes an association between OPN4 and a Mendelian disease. Limitations and caveats: (a) because raw data or a full gene list are not included in the provided excerpt, it is possible (but not evidenced here) that OPN4 variants were observed in the study cohort; the absence of mention in the text does not prove OPN4 is not associated with disease—only that this paper does not address that gene specifically; (b) the study’s focus and reporting (cohort-level yield, PGT-M outcomes) suggest it was not intended as a gene-specific association study for OPN4. Therefore, relevance to the specific claim is nil based on the provided content.


[Read Paper](https://www.semanticscholar.org/paper/1b721e406f9d31ac5983b146c5b9aa03d58fc49c)


### Whole-Exome Sequencing in Adults With Chronic Kidney Disease

**Why Not Relevant**: The provided paper text describes a whole-exome sequencing study of adults with chronic kidney disease that focused on a curated set of nephropathy and hypertension OMIM genes, methods for variant annotation and interpretation, and study logistics. Nowhere in the supplied content is the gene OPN4 mentioned, nor are any results, variant calls, or mechanistic discussions involving OPN4 presented. The only potentially related material is reference to Supplement Table 3 (the curated list of 287 OMIM nephropathy genes), which is not included in the provided excerpt; therefore it is not possible from this text to determine whether OPN4 was examined or implicated. Because the paper text contains no direct observation, result, or mechanistic pathway involving OPN4, it does not provide evidence for or against the claim that "OPN4 is likely associated with at least one Mendelian disease." Absence of evidence in this paper is not evidence of absence in the literature overall, but based solely on the provided content, there is effectively no relevant evidence to evaluate the claim.


[Read Paper](https://www.semanticscholar.org/paper/a29bd57964f68ac5f8e1abcafe1112ba488f53f5)


## Search Queries Used

- OPN4 human mutation Mendelian disease familial segregation case report whole exome sequencing family study OMIM ClinVar pathogenic

- melanopsin OPN4 mutation pupillary light reflex intrinsically photosensitive retinal ganglion cell non image forming vision circadian rhythm disorder sleep disorder non 24 congenital pupil anomaly

- OPN4 variants population frequency gnomAD allele frequency loss of function intolerance pLI constraint ClinVar variant interpretation pathogenic benign

- Opn4 knockout mouse melanopsin loss of function electrophysiology photoresponse behavioral circadian entrainment retinal ganglion cell phototransduction functional assay humanized variant


## Bibliometric Configuration

- Use Bibliometrics: True

- Author Impact Weight: 0.4

- Citation Impact Weight: 0.4

- Venue Impact Weight: 0.2
