## Supplementary material for "MENDELSEEK: An algorithm that predicts Mendelian Genes and elucidates what makes them special": OPRK1.txt

# Claim: The gene OPRK1 is likely associated with at least one Mendelian disease.

**Status**: processed

**Overall Rating**: -1

**Explanation**:

No relevant papers were found that support or refute this claim.


## Relevant Papers


## Other Reviewed Papers


### ClinGen Myeloid Malignancy Variant Curation Expert Panel recommendations for germline RUNX1 variants.

**Why Not Relevant**: The provided paper content exclusively addresses the development and application of gene- and disease-specific variant curation criteria for inherited myeloid malignancy genes, with explicit focus on RUNX1. There is no mention of OPRK1, no reported variants in OPRK1, no discussion of clinical cases implicating OPRK1 in Mendelian disease, and no mechanistic data relating OPRK1 to a Mendelian phenotype. Thus the paper provides no direct evidence supporting or refuting the claim that OPRK1 is associated with at least one Mendelian disease. The methodological approaches described (ACMG/AMP tailoring, allele frequency thresholds, functional assay thresholds) are broadly relevant to variant curation but are applied here only to RUNX1; without data or discussion of OPRK1, nothing in this paper changes the plausibility of the claim about OPRK1.


[Read Paper](https://www.semanticscholar.org/paper/d2a2dd2ebfb5a63969c41c03899f8f978fb18c6a)


### Trio-based whole exome sequencing in patients with suspected sporadic inborn errors of immunity: A retrospective cohort study

**Why Not Relevant**: Relevance is effectively zero because the provided paper content does not mention OPRK1 anywhere, nor does it present data, variants, functional assays, or discussion pertaining to OPRK1. The study focuses on de novo variants identified in a 123-patient cohort with suspected inborn errors of immunity and reports specific genes (NLRP3, RELA, PSMB10, DDX1, KMT2C, FBXW11) for which they found candidate or functionally supported variants. While the absence of OPRK1 among identified candidates is weak, indirect negative information within this IEI-focused cohort cannot be used to conclude that OPRK1 is or is not associated with a Mendelian disease: (a) the cohort is disease-specific (IEI) and relatively small (123 trios), so many Mendelian genes (especially those causing non-IEI phenotypes) would not be expected to appear; (b) the analysis targeted rare, non-synonymous de novo coding/splice-site variants, so inherited pathogenic variants, non-coding pathogenic variants, structural variants, or variants in other populations would be missed; (c) the paper reports functional follow-up only for FBXW11, not for other genes. Therefore this paper provides no direct or mechanistic evidence for or against the claim that OPRK1 is likely associated with at least one Mendelian disease.


[Read Paper](https://www.semanticscholar.org/paper/cdde2232daad22b7d92ec15faa38b9776f1e4c15)


### The Burden of Candidate Pathogenic Variants for Kidney and Genitourinary Disorders Emerging From Exome Sequencing

**Why Not Relevant**: The paper focuses on identifying and curating genes associated with Mendelian kidney and genitourinary disorders and on assessing the prevalence of candidate pathogenic variants in a healthy cohort. The provided content details the databases and methods used to assemble and annotate a domain-specific gene list (final n = 625 genes) and the variant curation pipeline. Nowhere in the supplied excerpts (or elsewhere in the provided content) is OPRK1 mentioned, nor are there any data, variant reports, or mechanistic discussions specific to OPRK1. Therefore the paper provides no direct evidence that OPRK1 is associated with any Mendelian disease. It also provides no mechanistic evidence regarding OPRK1 function or pathogenicity. Because the study is disease-domain–specific (kidney/genitourinary) and applies particular inclusion criteria, absence of OPRK1 in the presented material cannot be interpreted as positive or negative evidence for OPRK1 generally; it simply indicates the paper does not address the claim. For these reasons relevance to the claim is effectively zero.


[Read Paper](https://www.semanticscholar.org/paper/a75e297208c04992568e607d19aa9b9aaebd09d5)


### Exploring the Impact of Genetics in a Large Cohort of Moebius Patients by Trio Whole Exome Sequencing

**Why Not Relevant**: This paper is not relevant to the claim that OPRK1 is likely associated with at least one Mendelian disease. The manuscript excerpt discusses whole-exome sequencing (WES) in a cohort of 37 patients with Moebius syndrome (MBS), the absence of de novo variants in REV3L and PLXND1 in this cohort, and the failure of WES to identify mutations associated with MBS. The text does not mention OPRK1, does not report any variants in OPRK1, and provides no mechanistic data (biochemical, expression, pathway, or animal-model evidence) concerning OPRK1. Therefore there is neither direct evidence (observational or genetic association) nor mechanistic evidence in this paper that bears on whether OPRK1 is associated with any Mendelian disease.

Additional context and caveats: (1) The negative WES results reported here concern Moebius syndrome specifically and the inability to find common causative genes in this cohort; they do not constitute evidence that OPRK1 is not involved in other Mendelian diseases. (2) WES has known limitations (it can miss deep intronic/regulatory variants, copy-number or structural variants, repeat expansions, low-level mosaicism, and some GC-rich exons), so the absence of a reported finding in a WES study does not definitively rule out involvement of any gene in any disease. (3) Because OPRK1 is not mentioned, there is no excerpt to extract that would directly or mechanistically support or refute the claim. Consequently, relevance is effectively zero and no supporting verbatim sentences can be provided from this paper.


[Read Paper](https://www.semanticscholar.org/paper/226b561a7e847eca27fae7e2558a187686dd8f75)


### Clingen Coagulation Factor Deficiency Variant Curation Expert Panel: Meeting the Need for Recommendations to Curate Variants in the Coagulation Factor Genes

**Why Not Relevant**: The provided paper content focuses entirely on variant curation for coagulation factor genes (specifically F8 and F9) and the development/specification of ACMG/AMP guideline adaptations by the ClinGen CFD-VCEP. There is no mention of the gene OPRK1, no association studies, no variant lists including OPRK1, and no mechanistic discussion relevant to OPRK1. Therefore the paper provides no direct or mechanistic evidence for or against the claim that OPRK1 is likely associated with at least one Mendelian disease. The absence of any reference to OPRK1 or to non-coagulation genes means this source is not relevant to evaluating that claim.


[Read Paper](https://www.semanticscholar.org/paper/9ea8d4ce19622ae1c6b0cdfd40848c93199f6280)


### Descriptive analysis of MC4R gene variants associated with obesity listed on ClinVar

**Why Not Relevant**: The provided paper addresses the MC4R gene and its variants; it does not mention OPRK1 or present any data bearing on OPRK1. Examples from the text demonstrating its MC4R focus include: "Objectives The most recent version of ClinVar was utilized to filter variants of the MC4R gene based on location, condition, and clinical significance with the goal of obtaining benign and disease-associated variants of the MC4R gene." and "MC4R gene variants can lead to dysregulation of energy expenditure and appetite control, which prompted this study to delineate the distinctive features of MC4R gene variants submitted to the ClinVar repository regarding their association with obesity and related phenotypes." The results and conclusions likewise discuss pathogenic variants of MC4R (e.g., "A total of 103 pathogenic variants were observed.") and implications for monogenic obesity and diabetes related to MC4R. Because the paper exclusively analyses and discusses MC4R, it provides no direct evidence (no reported variants, phenotypes, clinical assertions, or mechanistic data) for or against the claim that OPRK1 is associated with a Mendelian disease. Limitations and implications: (1) Absence of mention of OPRK1 in this paper is not evidence that OPRK1 is not associated with a Mendelian disease; it only means this paper does not address the gene. (2) To evaluate the claim about OPRK1 one would need gene-specific evidence such as ClinVar/OMIM entries, published case reports linking OPRK1 variants to segregating Mendelian phenotypes, functional/biochemical studies of pathogenic variants, or curated gene–disease validity assessments. (3) No mechanistic connection between MC4R findings and OPRK1 is provided here; while both genes encode G-protein-coupled receptors, the paper does not discuss cross-gene mechanisms that would allow inference about OPRK1. In summary, relevance to the claim is effectively zero because the paper focuses entirely on MC4R and contains no data, statements, or mechanistic discussion about OPRK1.


[Read Paper](https://www.semanticscholar.org/paper/6bae624c1c06216b5458a99ef3b5e0a4b2d4cd16)


### Deciphering the Non-Coding Genome in Autism Spectrum Disorders (ASD): A Study of De Novo and Rare Inherited Variants through Targeted Sequencing in Regulatory Regions

**Why Not Relevant**: The provided paper content does not mention OPRK1 at any point and contains no direct evidence linking OPRK1 to any Mendelian disease. The study focuses on non-coding regulatory variation in ASD and explicitly excluded cases with coding loss-of-function variants and large CNVs (which are common sources of Mendelian gene discoveries). Mechanistic findings reported (e.g., global dysregulation of CTCF) are general and do not implicate OPRK1. Therefore there is neither direct genetic evidence (e.g., pathogenic coding variants in OPRK1 segregating with disease) nor mechanistic evidence in this text supporting the claim that OPRK1 is likely associated with a Mendelian disease. Limitations and caveats: absence of OPRK1 in the provided excerpt is not absolute proof that the full paper or its supplementary materials do not contain any mention of OPRK1; however, based on the supplied content, no support is present. To evaluate the claim further, consult primary disease-gene resources (OMIM, ClinVar, HGMD), PubMed for OPRK1 clinical genetics literature, and large-scale gene-disease aggregation efforts (e.g., ClinGen) for curated evidence of Mendelian associations.


[Read Paper](https://www.semanticscholar.org/paper/649aeab19738420d26b974bc80eaee323f848ea6)


## Search Queries Used

- OPRK1 KOR kappa opioid receptor mutation case report family segregation novel variant single gene disorder Mendelian disease

- OPRK1 ClinVar OMIM ClinGen pathogenic likely pathogenic variants clinical phenotype gene disease association OPRK1

- whole exome sequencing exome sequencing trio rare Mendelian disorders OPRK1 de novo variant candidate gene rare disease cohort

- OPRK1 kappa opioid receptor loss of function gain of function mutation knockout mouse zebrafish neurodevelopmental phenotype functional assay G protein coupled receptor signaling


## Bibliometric Configuration

- Use Bibliometrics: True

- Author Impact Weight: 0.4

- Citation Impact Weight: 0.4

- Venue Impact Weight: 0.2
