## Supplementary material for "MENDELSEEK: An algorithm that predicts Mendelian Genes and elucidates what makes them special": PCDH18.txt

# Claim: The gene PCDH18 is likely associated with at least one Mendelian disease.

**Status**: processed

**Overall Rating**: -1

**Explanation**:

No relevant papers were found that support or refute this claim.


## Relevant Papers


## Other Reviewed Papers


### The Burden of Candidate Pathogenic Variants for Kidney and Genitourinary Disorders Emerging From Exome Sequencing

**Why Not Relevant**: The provided paper content does not mention the gene PCDH18, does not list the curated gene set in the excerpt, and contains no data or statements that directly or mechanistically link PCDH18 to any Mendelian disease. The paper focuses on (1) methods for identifying and curating genes associated with Mendelian kidney and genitourinary disorders using OMIM and Orphanet, (2) variant-filtering pipelines and the prevalence of candidate pathogenic variants in a large cohort of ostensibly healthy adults, and (3) the problem of misclassification of pathogenic variants in public databases. Although the authors state that their final curated list of 625 genes was annotated and is accessible on a laboratory website, that list is not present in the provided content. Therefore: 

- There is no direct evidence in the excerpt that PCDH18 has been identified by the authors as disease-associated. 
- There is no mechanistic discussion (e.g., function of PCDH18, pathogenic variant types, inheritance mode, or model systems) relevant to PCDH18. 
- The methods described (OMIM/Orphanet search, manual curation, ExAC frequency filtering, ClinVar/HGMD annotation) are applicable approaches that could in principle be used to evaluate PCDH18, but the excerpt contains no results or statements about PCDH18 itself. 

Limitations that prevent supporting the claim from this paper: 
- Absence of the gene name PCDH18 anywhere in the provided text. 
- The curated gene list is referenced but not included in the provided content, so one cannot determine whether PCDH18 was included. 
- The paper’s scope is kidney and genitourinary Mendelian disorders; if PCDH18 is associated with a non-kidney phenotype, it would be outside the paper’s main focus and unlikely to be discussed. 

Conclusion: Based solely on the provided excerpt, this paper does not provide direct or mechanistic evidence that PCDH18 is likely associated with at least one Mendelian disease. If you want an evidence-based assessment of PCDH18, please provide the full paper (including the linked curated gene list) or other literature specifically mentioning PCDH18, and I will analyze those for direct associations, variant evidence, mechanistic plausibility, and study limitations.


[Read Paper](https://www.semanticscholar.org/paper/a75e297208c04992568e607d19aa9b9aaebd09d5)


### Case report of a novel homozygous splice site mutation in PLA2G6 gene causing infantile neuroaxonal dystrophy in a Sudanese family

**Why Not Relevant**: The provided paper content is insufficiently specific to support or refute the claim that PCDH18 is likely associated with at least one Mendelian disease. The only sentence given is: "This variant has one very strong (loss of function mutation) and three supporting evidences for its pathogenicity (segregation with the disease, multiple computational evidence and specific patients’ phenotype)." That sentence (1) does not name the gene (PCDH18 is not mentioned), (2) does not describe the variant (genomic coordinates, transcript, zygosity), (3) does not specify the inheritance pattern or number of affected individuals/families, (4) provides no study methods, sample sizes, statistical measures, or functional validation, and (5) does not state whether the reported pathogenicity demonstrates a causal gene–disease relationship versus a single pathogenic variant in an already established disease gene. While the sentence lists types of evidence that are potentially relevant (loss-of-function as a mechanistic class of variant, segregation as direct evidence, computational predictions, and phenotype match), those elements alone—presented without gene identification, detailed data, or context—are insufficient to establish that PCDH18 is associated with a Mendelian disorder. Important limitations: the statement appears to be a summary assessment (analogous to ACMG-style variant classification) but lacks provenance and specificity; a single reported pathogenic variant (even with segregation in one family) does not by itself prove a gene–disease relationship, which normally requires replication (multiple unrelated families), experimental functional evidence, clear inheritance consistency, and population frequency analysis. Therefore, based only on the supplied content, relevance to the claim about PCDH18 is very low. To properly evaluate the claim we would need the full paper text showing (at minimum) that the variant is in PCDH18, detailed segregation data across families, inheritance pattern, case/control or population frequency data, and any functional/mechanistic experiments linking loss of PCDH18 function to the disease phenotype.


[Read Paper](https://www.semanticscholar.org/paper/2230af79fdcb2bff652cfee20cf54fdf6d8f079f)


### Precision Medicine in Internal Medicine

**Why Not Relevant**: The provided paper is a general review of genetic testing strategies, exome/genome sequencing, and variant interpretation. It does not mention the gene PCDH18 by name, nor does it present any data (genetic, clinical, segregation, functional, or statistical) concerning PCDH18. Therefore the paper contains no direct evidence that PCDH18 is associated with any Mendelian disease, nor does it contain mechanistic data specific to PCDH18 (e.g., expression, protein function, loss-of-function intolerance, animal models, or pathogenic variants). The included excerpts are general context about how gene-disease associations are discovered and how many genes are currently known to cause Mendelian disorders; they may inform the plausibility that additional genes (potentially including PCDH18) could be discovered in the future, but they do not support or refute the specific claim about PCDH18. Because no gene-specific evidence is present, this paper is not useful for assessing whether PCDH18 is likely associated with at least one Mendelian disease.


[Read Paper](https://www.semanticscholar.org/paper/c7199e8a5226f8d9d55777da0e15e91797bb459a)


### A distinct APC pathogenic germline variant identified in a southern Thai family with familial adenomatous polyposis

**Why Not Relevant**: The provided content concerns a pathogenic variant in the APC gene and its segregation in a single family. The claim under consideration is about the gene PCDH18. The excerpt makes no mention of PCDH18, provides no data (genetic, clinical, segregation, functional, or mechanistic) relating to PCDH18, and therefore does not support or refute the claim that PCDH18 is likely associated with at least one Mendelian disease. APC and PCDH18 are distinct genes with different known biological roles (APC is a well-established tumor suppressor implicated in familial adenomatous polyposis; PCDH18 is a protocadherin), so evidence about APC cannot be extrapolated to PCDH18 without additional specific data. Note also that only a single sentence from the paper was provided; if the full paper includes other sections mentioning PCDH18, those were not made available for assessment.


[Read Paper](https://www.semanticscholar.org/paper/79d240917f7d527a68ee4003646f72690c4bd09e)


### Germline heterozygous exons 8–11 pathogenic BARD1 gene deletion reported for the first time in a family with suspicion of a hereditary colorectal cancer syndrome: more than an incidental finding?

**Why Not Relevant**: The provided paper content contains a single sentence: "This is the first published work in which this BARD1 deletion is detected in a family with familial colorectal cancer type X (FCCTX) syndrome, in which the clinical criteria for Lynch syndrome without alteration of the DNA mismatch repair (MMR) system are fulfilled." This sentence refers specifically to BARD1 and familial colorectal cancer type X (FCCTX) and makes no mention of the gene PCDH18, Mendelian inheritance of PCDH18 variants, segregation of PCDH18 variants with disease, pathogenic variants in PCDH18, mechanistic links between PCDH18 and disease biology, or any experimental or clinical data relevant to PCDH18. Therefore the paper content as provided gives no direct evidence for or against the claim that "PCDH18 is likely associated with at least one Mendelian disease," nor does it provide mechanistic evidence bearing on that claim. Additional limitations that reduce relevance: (1) only one sentence of content is provided (no methods, results, or discussion), (2) the gene discussed (BARD1) is different from the target gene in the claim (PCDH18), and (3) no data, statistical analyses, segregation information, functional assays, or cited literature relevant to PCDH18 are present. To evaluate the PCDH18 claim one would need studies that explicitly analyze PCDH18 (e.g., segregation in families, case-control or cohort studies, ClinVar/OMIM entries, functional/animal-model data, loss-of-function intolerance metrics), none of which are present in the provided content.


[Read Paper](https://www.semanticscholar.org/paper/884a95b299f7238bf2a4a70eaeec5e815a99798c)


### Autosomal Dominant Intellectual Development Disorder-6 (MRD6) Without Seizures Linked to a De Novo Mutation in the grin2b Gene Revealed by Exome Sequencing: A Case Report of a Moroccan Child

**Why Not Relevant**: The paper exclusively discusses a GRIN2B (grin2b) loss-of-function variant causing autosomal dominant intellectual development disorder-6 (MRD6). There is no mention of the gene PCDH18, no data (genetic, clinical, or mechanistic) concerning PCDH18, and no discussion of pathways or interactions that would plausibly link GRIN2B findings to PCDH18. Consequently, the paper provides no direct or mechanistic evidence for or against the claim that PCDH18 is likely associated with at least one Mendelian disease. Without any reference to PCDH18 or relevant connecting biology, the content cannot inform the claim.


[Read Paper](https://www.semanticscholar.org/paper/67d756e0cc302e5d46fdda9df34403f1f2de9a5a)


### A Novel Loss-of-Function Variant in COL4A3 in a Consanguineous Moroccan Family Displaying the Alport Syndrome with Variable Clinical Expression

**Why Not Relevant**: This paper provides no evidence supporting an association between PCDH18 and any Mendelian disease. The study is a case report of Alport syndrome in a consanguineous family and identifies a pathogenic homozygous nonsense variant in COL4A3 as the likely cause. The text repeatedly attributes the disease to COL4A3/COL4A4/COL4A5 (type IV collagen genes) and contains no mention of PCDH18, no reported variants in PCDH18, and no mechanistic discussion linking protocadherin-18 to the phenotype. Limitations to this negative assessment: the paper does not state the exact breadth of the next-generation sequencing (e.g., targeted COL4 panel vs exome/genome), so it is not certain whether PCDH18 was interrogated and found negative. Nevertheless, because the authors identify and classify a clearly plausible pathogenic COL4A3 variant that segregates with disease in the family, this report supplies no direct or mechanistic evidence that would support the claim that PCDH18 is likely associated with a Mendelian disorder. Absence of mention in this focused COL4A3-centered study is not proof that PCDH18 is uninvolved in any Mendelian disease broadly, but it does mean this paper is not a source of supporting evidence for the claim.


[Read Paper](https://www.semanticscholar.org/paper/d878160b7c91c8ae5e15426f4a2cb331ab38d3d3)


## Search Queries Used

- PCDH18 protocadherin 18 variant pathogenic Mendelian disease case report family segregation

- PCDH18 ClinVar OMIM HGMD gene disease association clinical significance PCDH18

- PCDH18 protocadherin 18 neuronal development synaptic adhesion knockout mouse phenotype neurodevelopment behavioral

- PCDH18 de novo variant copy number variant deletion exome sequencing intellectual disability autism epilepsy rare disease cohort gnomAD pLI LOEUF constraint


## Bibliometric Configuration

- Use Bibliometrics: True

- Author Impact Weight: 0.4

- Citation Impact Weight: 0.4

- Venue Impact Weight: 0.2
