## Supplementary material for "MENDELSEEK: An algorithm that predicts Mendelian Genes and elucidates what makes them special": PDE5A.txt

# Claim: The gene PDE5A is likely associated with at least one Mendelian disease.

**Status**: processed

**Overall Rating**: -1

**Explanation**:

No relevant papers were found that support or refute this claim.


## Relevant Papers


## Other Reviewed Papers


### The Burden of Candidate Pathogenic Variants for Kidney and Genitourinary Disorders Emerging From Exome Sequencing

**Why Not Relevant**: The provided paper content discusses methods and aggregate results for identifying and curating genes associated with Mendelian kidney and genitourinary disorders and highlights challenges with variant annotation and prevalence of purportedly pathogenic variants in control cohorts. However, the excerpt contains no mention of the specific gene PDE5A, no locus-specific variant data, and no direct clinical or mechanistic evidence linking PDE5A to any Mendelian disease. The paper's methods imply that if PDE5A were implicated in Mendelian kidney/genitourinary disease it might appear in their curated list, but the excerpt does not include the gene list or any PDE5A-related results. Therefore, while the paper is relevant in that it outlines a rigorous framework and warns about false-positive gene-disease annotations (which is useful background when evaluating any gene claim), it provides no direct or specific mechanistic evidence for or against the claim that PDE5A is associated with a Mendelian disease.


[Read Paper](https://www.semanticscholar.org/paper/a75e297208c04992568e607d19aa9b9aaebd09d5)


### An increased burden of rare exonic variants in NRXN1 microdeletion carriers is likely to enhance the penetrance for autism spectrum disorder

**Why Not Relevant**: The paper focuses specifically on Autism Spectrum Disorder and on NRXN1 intragenic deletions as a model of an ASD‑associated susceptibility variant. The methods and results (chromosomal microarray, whole‑exome sequencing, mtDNA deep sequencing) are applied to evaluate genetic modifiers of NRXN1 deletion penetrance; the text provided contains no mention of the gene PDE5A, no variant data for PDE5A, and no gene–disease association analyses that include PDE5A. Consequently, there is no direct evidence (no observations of pathogenic variants in PDE5A linked to disease in the study) nor mechanistic evidence (no functional or pathway data involving PDE5A) relevant to the claim that PDE5A is likely associated with at least one Mendelian disease. Because the absence of evidence in this paper is not evidence of absence, one cannot use this paper to support or refute the claim about PDE5A; therefore the paper is not relevant for evaluating that claim.


[Read Paper](https://www.semanticscholar.org/paper/bcd5750c86d7966d234b0e990ed6c7f1a7727279)


### A New Approach to Identify Pathogenic Mutations and Inherited Variants By Exome Sequencing – Using a Pair of Identical Twins with Monoclonal Lymphosis As Case Model

**Why Not Relevant**: The paper focuses on developing an annotation/ranking tool and applies it to whole-exome data from a pair of identical twins discordant for B-CLL/monoclonal lymphocytosis. The reported results list several genes implicated in CLL (IRF8, RUNX1, TET2, PLCB1, ELF4, MAP2K3, CHST15, CHD2, ABL2, MYH1, TNFAIP6, NOTCH4, TOPBP1, DNMT1, CHD7, SETD1B, etc.), but there is no mention of the gene PDE5A anywhere in the text. There are no experimental data, family segregation analyses, statistical associations, functional assays, or mechanistic discussions involving PDE5A. Therefore the paper provides neither direct evidence (e.g., reported pathogenic variants, segregation with disease, statistical association) nor mechanistic evidence (molecular function linked to a Mendelian phenotype) that PDE5A is associated with any Mendelian disease. This absence of evidence in this paper means its relevance to the claim is effectively zero; however, absence of mention here does not prove PDE5A has no Mendelian associations elsewhere—it only indicates this paper does not support the claim.


[Read Paper](https://www.semanticscholar.org/paper/155462d32af17033d5ac53072be832d7bb2a0381)


### The genetic architecture of schizophrenia and bipolar disorder: identity by descent and exome sequencing in a family-based population sample

**Why Not Relevant**: The provided paper content investigates rare sequence variants associated with schizophrenia and bipolar disorder in a closed population and lists candidate genes (GRM7, GRM8, NCAM1, NCAN) and implicated pathways (axon guidance, synaptic transmission). Nowhere in the supplied text is PDE5A mentioned, nor are there data, results, or mechanistic discussions that implicate PDE5A. Therefore this paper provides no direct evidence that PDE5A is associated with any Mendelian disease. Indirectly, the study's design (exome sequencing of 17 patients from a closed population) could have detected rare Mendelian variants if PDE5A were causal in this cohort, but the absence of PDE5A from the reported candidate genes and pathways means the paper does not support the claim. Limitations that further reduce relevance: the study focuses on complex psychiatric disorders rather than classical Mendelian phenotypes, the sequenced sample is small (17 patients), and candidate findings emphasize polygenic architectures; none of these produce evidence for PDE5A's involvement in Mendelian disease within this dataset.


[Read Paper](https://www.semanticscholar.org/paper/6afb1d6752f9c11ae2337df73aa8cc500c2ed480)


## Search Queries Used

- PDE5A phosphodiesterase 5A mutation case report Mendelian disorder monogenic disease pathogenic variant germline familial segregation

- PDE5A OMIM ClinVar ClinGen Orphanet HGMD PDE5A gene disease association Mendelian gene entry review

- whole exome sequencing PDE5A rare variants gene burden monogenic disorder familial inherited exome study case series rare damaging variants

- Pde5a knockout mouse phenotype loss of function phosphodiesterase 5A cGMP signaling pulmonary hypertension vascular development erectile dysfunction animal model


## Bibliometric Configuration

- Use Bibliometrics: True

- Author Impact Weight: 0.4

- Citation Impact Weight: 0.4

- Venue Impact Weight: 0.2
