## Supplementary material for "MENDELSEEK: An algorithm that predicts Mendelian Genes and elucidates what makes them special": PEG10.txt

# Claim: The gene PEG10 is likely associated with at least one Mendelian disease.

**Status**: processed

**Overall Rating**: -1

**Explanation**:

No relevant papers were found that support or refute this claim.


## Relevant Papers


## Other Reviewed Papers


### Genome sequencing for detection of pathogenic deep intronic variation: A clinical case report illustrating opportunities and challenges

**Why Not Relevant**: The provided paper content makes no mention of the gene PEG10 and instead exclusively describes variants in JAM3 and their relationship to a specific autosomal recessive disorder. Example verbatim sentences from the provided content (kept exactly as given) showing the paper's focus are:

"Variants in JAM3 have been reported in four families manifesting a severe autosomal recessive disorder characterized by hemorrhagic destruction of the brain, subependymal calcification, and cataracts." 

"We describe a 7‐year‐old male with a similar presentation found by research‐based quad genome sequencing to have two novel splicing variants in trans in JAM3, including one deep intronic variant (NM_032801.4: c.256+1260G>C) not detectable by standard exome sequencing." 

"Targeted sequencing of RNA isolated from transformed lymphoblastoid cell lines confirmed that each of the two variants has a deleterious effect on JAM3 mRNA splicing." 

"The role for genome sequencing as a clinical diagnostic test extends to those patients with phenotypes strongly suggestive of a specific Mendelian disorder, especially when the causal genetic variant(s) are not found by a more targeted approach." 

These sentences demonstrate that the study's subject, results, and mechanistic discussion (splicing effects of JAM3 variants and diagnostic implications of genome sequencing) concern JAM3 rather than PEG10. Therefore, the paper as provided supplies no direct evidence (no observation of PEG10 variants causing Mendelian disease) nor mechanistic evidence (no functional data, pathways, or experiments implicating PEG10) to support or refute the claim that "the gene PEG10 is likely associated with at least one Mendelian disease." 

Caveat: this conclusion applies only to the provided excerpt. If the full paper contains other sections not included here that mention PEG10, those were not available for review. To evaluate the claim further, I recommend searching the full text of this paper for "PEG10" or consulting genetic-disease databases and the broader literature for PEG10-specific evidence.


[Read Paper](https://www.semanticscholar.org/paper/3d3350f760ece7001258cb9733aec254e582d1f9)


### Biases in arginine codon usage correlate with genetic disease risk

**Why Not Relevant**: The provided content contains only a general conclusion about a genome-wide association between arginine codon usage patterns and an increased probability that a gene underlies single-gene (Mendelian) disorders. It does not mention the gene PEG10, does not report PEG10's codon usage, and provides no direct genetic, clinical, or experimental evidence linking PEG10 to any Mendelian disease. Thus the paper excerpt is only indirectly relevant: it establishes a rule that could make PEG10 a candidate only if PEG10 meets the specific codon-usage criterion (CGN arginine codons) and even then would only change prior probability rather than provide direct proof. Because the excerpt lacks PEG10-specific data or results, and because the association described is probabilistic rather than deterministic, the content is insufficient to support the claim that "PEG10 is likely associated with at least one Mendelian disease." To assess PEG10 specifically would require (a) reporting of PEG10's arginine codon usage, (b) statistical evidence that PEG10 lies within the enriched class and (c) direct genetic/clinical evidence (variants segregating with disease, functional pathogenicity data, etc.). None of these are present in the provided text, so relevance to the specific claim about PEG10 is very low.


[Read Paper](https://www.semanticscholar.org/paper/b5c65cb630e8425dd9853bce349269f6032bb648)


### Novel LSS variants in alopecia and intellectual disability syndrome: New case report and clinical spectrum of LSS‐related rare disease traits

**Why Not Relevant**: The provided paper content exclusively discusses pathogenic variants in the LSS gene and associated Mendelian phenotypes (e.g., congenital cataract type 44, autosomal recessive hypotrichosis type 14, alopecia‐intellectual disability syndrome type 4). There is no mention of PEG10, PEG10 variants, PEG10-associated phenotypes, or any mechanistic link between LSS and PEG10 in the text supplied. Example sentences from the content that demonstrate the paper's focus include: "Pathogenic biallelic variants in LSS are associated with three Mendelian rare disease traits including congenital cataract type 44, autosomal recessive hypotrichosis type 14, and alopecia‐intellectual disability syndrome type 4 (APMR4)." and "We performed trio research exome sequencing on a family with a four‐year‐old male with global developmental delay, epilepsy and striking alopecia, and identified novel compound heterozygous LSS splice site (c.14+2T>C) and missense (c.1357 G>A; p.V453L) variant alleles." Because PEG10 is not discussed and no data, results, or mechanistic statements pertaining to PEG10 are present, this paper provides no direct or mechanistic evidence for or against the claim that "The gene PEG10 is likely associated with at least one Mendelian disease."


[Read Paper](https://www.semanticscholar.org/paper/fe90a5bb55278bc80588741eed5def6cf4f1aa82)


### Case report: Discovery of a de novo FAM111B pathogenic variant in a patient with an APECED-like clinical phenotype

**Why Not Relevant**: The paper focuses on AIRE and FAM111B as causes of APECED and POIKTMP and reports a de novo pathogenic variant in FAM111B in the described patient. PEG10 is not mentioned anywhere in the provided text; no results, discussion, or mechanistic data concerning PEG10 are presented. Although exome sequencing was performed (so PEG10 variants could in principle have been assessed), the authors do not report any findings or analysis related to PEG10. Therefore the paper provides no direct or mechanistic evidence to support or refute the claim that PEG10 is associated with at least one Mendelian disease.


[Read Paper](https://www.semanticscholar.org/paper/6ed49c6ffda4dbf2de15474aa07c9326d5636a34)


### #5782 AN EXCEPTIONAL CASE OF A PATIENT WITH TWO RARE GENETIC VARIANTS OF AUTOSOMAL DOMINANT POLYCYSTIC KIDNEY DISEASE AND ALPORT SYNDROME

**Why Not Relevant**: The provided paper content does not mention PEG10 anywhere and contains no data (genetic, clinical, or mechanistic) about PEG10. The paper is a case report focused on Alport syndrome and autosomal dominant polycystic kidney disease (ADPKD) and explicitly discusses different genes. Examples of verbatim sentences from the text that show the genes studied and the clinical focus are:

"Alport syndrome and autosomal dominant polycystic kidney disease (ADPKD) belong to the most prevalent monogenic causes of chronic kidney disease (CKD) and both progress to end stage renal disease (ESRD)."

"ADPKD is mostly caused by mutations in PKD1and PKD2 genes but recently, the GANAB gene, that encodes the glucosidase IIa subunit, has been identified has a cause of a milder form of ADPKD."

"Genetic testing via next generation sequencing was performed revealing heterozygous mutations in COL4A3 (c.4421T>C p.(Leu1474Pro)), reported in ClinVar, and described in literature as an autosomal dominant variant present in Alport families."

"A mutation in the GANAB (c.2281A>G p.(Thr761Ala)) gene, classified as variants of unknown significance in was also detected, but this mutations is not reported in gnomAD or ClinVar."

How this bears on the claim: none of the above sentences (or any other text provided) address PEG10. There is no direct evidence (no reported PEG10 variants, no segregation or case reports, no statistical association) and no mechanistic evidence (no discussion of PEG10 function, imprinting, pathogenic mechanisms, or animal/functional studies) that would support or refute an association between PEG10 and any Mendelian disease. The absence of PEG10 in the report means it cannot be used as supporting evidence for the claim.

Limitations and caveats: the absence of any mention of PEG10 in this paper does not prove PEG10 is not associated with a Mendelian disease; it only means this particular paper provides no relevant evidence. It is possible other papers or datasets (not provided here) do include PEG10-related findings. Additionally, because the paper is a single-case report focused on specific kidney diseases and genes (COL4A3, PKD1/2, GANAB), it would not be expected to address PEG10 unless it were implicated in that case. Given the provided content alone, the relevance to the claim is effectively zero.


[Read Paper](https://www.semanticscholar.org/paper/0337e73775f38924bbc68438e6e09327983393d6)


### Thiamine-Responsive Megaloblastic Anemia Syndrome Combined with Thalassemia Trait: A Rare Association

**Why Not Relevant**: This paper provides no direct or mechanistic evidence relevant to the claim that PEG10 is likely associated with at least one Mendelian disease. The manuscript is a clinical case report of thiamine-responsive megaloblastic anemia (TRMA) that explicitly attributes the genetic cause of TRMA to mutations in SLC19A2 (second excerpt). The paper contains no mention of PEG10, no genetic data implicating PEG10, and no mechanistic discussion that would plausibly link PEG10 to the described phenotype. Additional important limitations that further reduce relevance: (1) this is a single-case report without molecular genetic confirmation (parents refused genetic diagnosis), so it does not even provide primary sequence data for SLC19A2 or any other gene; (2) the focus is clinical (presentation and thiamine response) rather than genetic discovery; and (3) the paper's mechanisms relate to thiamine transport via SLC19A2, which is unrelated to PEG10's known biology. For these reasons the paper cannot be used to support or refute the claim regarding PEG10 and Mendelian disease.


[Read Paper](https://www.semanticscholar.org/paper/10efb4633c1180941f25a09807f0b4048b07cea4)


### Explorer CDKL 5 variants : improving our understanding of a rare neurological disorder

**Why Not Relevant**: This paper focuses entirely on the CDKL5 gene and CDKL5 deficiency and contains no data, statements, or analyses about the gene PEG10. The objective, methods, results, and conclusions repeatedly refer to CDKL5 (e.g., discussion of pathogenic missense variants clustering in the catalytic domain of CDKL5, ExAC counts for CDKL5 variants, isoforms hCDKL5_1 and hCDKL5_5, and minigene splicing assays for CDKL5 exon 14). There is no mention of PEG10, no population-variant data for PEG10, no functional or mechanistic experiments involving PEG10, and no discussion of Mendelian diseases associated with PEG10. Therefore the paper provides neither direct evidence (case reports, segregation, variant pathogenicity data) nor mechanistic evidence (functional assays, protein/domain relevance, imprinting or expression patterns relevant to disease) relevant to the claim that "PEG10 is likely associated with at least one Mendelian disease." The analytical approaches used in the paper (use of ExAC, variant reclassification, minigene splicing assays) are methods that could, in principle, be applied to PEG10 in another study, but because this paper does not include PEG10 in its data or discussion, it cannot be used to support or refute the claim about PEG10.


[Read Paper](https://www.semanticscholar.org/paper/a959fe9191159fa8a508c918a1e4e7dcf60553fd)


### Leveraging Clinical, Functional, Molecular and Population Genetic Data Reveals Genotype Phenotype Association and Health Disparity in a Monogenic Disorder, CTX

**Why Not Relevant**: The provided paper exclusively studies Cerebrotendinous Xanthomatosis (CTX) and pathogenic variants in CYP27A1. There is no mention of PEG10, no data, analyses, or discussion linking PEG10 to disease, and no mechanistic statements involving PEG10. Therefore the paper does not provide direct evidence supporting or refuting the claim that PEG10 is likely associated with at least one Mendelian disease. Absence of PEG10 in this paper is not evidence that PEG10 is not associated with a Mendelian disorder; it only means this paper is irrelevant to the claim. To evaluate the claim about PEG10, a targeted literature search (genetic association studies, OMIM, ClinVar, gene-disease validity resources such as ClinGen, case reports, or functional studies of PEG10) would be required.


[Read Paper](https://www.semanticscholar.org/paper/ecd9a6ae7be2a66a942f75a505fc4b04dff8e862)


## Search Queries Used

- PEG10 pathogenic variant Mendelian disease exome sequencing case report clinical genetics

- PEG10 gene association rare disease monogenic syndrome OMIM ClinVar genetic diagnosis

- PEG10 imprinting paternal expression knock out mouse embryo lethality placenta development haploinsufficiency imprinting disorder functional study

- PEG10 germline variant population frequency gnomAD loss of function constraint pLI observed expected variants constraint metrics


## Bibliometric Configuration

- Use Bibliometrics: True

- Author Impact Weight: 0.4

- Citation Impact Weight: 0.4

- Venue Impact Weight: 0.2
