## Supplementary material for "MENDELSEEK: An algorithm that predicts Mendelian Genes and elucidates what makes them special": PELO.txt

# Claim: The gene HBS1L is likely associated with at least one Mendelian disease.

**Status**: processed

**Overall Rating**: -1

**Explanation**:

No relevant papers were found that support or refute this claim.


## Relevant Papers


## Other Reviewed Papers


### Endophenotype effect sizes support variant pathogenicity in monogenic disease susceptibility genes

**Why Not Relevant**: The provided paper content contains only a single, general methodological sentence and does not mention HBS1L, any specific variants, diseases, cohorts, statistical results, or mechanistic pathways. Therefore it supplies neither direct evidence (no case reports, association statistics, segregation data, or functional studies implicating HBS1L in Mendelian disease) nor gene-specific mechanistic evidence (no molecular or biological data about HBS1L). As a result, the content cannot meaningfully support or refute the claim that "The gene HBS1L is likely associated with at least one Mendelian disease." The statement only notes that population-based association approaches can be informative in general, which is potentially relevant as a methodological note for future studies of HBS1L, but it is insufficient as evidence regarding HBS1L itself.


[Read Paper](https://www.semanticscholar.org/paper/3ca53d5f1779ccd5c7db249a4a5782667f585773)


### Frequency and Phenotype Associations of Rare Variants in 5 Monogenic Cerebral Small Vessel Disease Genes in 200,000 UK Biobank Participants

**Why Not Relevant**: The paper does not mention HBS1L anywhere in the provided text. Its scope is limited to specific cerebral small vessel disease genes (CTSA, TREX1, HTRA1, and COL4A1/2) and analyzes putative pathogenic rare variants only in those genes. Because HBS1L is not among the genes studied, there is no direct genetic association data, no reported phenotypes, and no mechanistic discussion relevant to HBS1L in this paper. Therefore the paper provides no direct or mechanistic evidence to support or refute the claim that HBS1L is likely associated with at least one Mendelian disease. Note that absence of HBS1L from this study cannot be interpreted as evidence against its involvement in Mendelian disease; it simply lies outside the study's examined gene set and aims.


[Read Paper](https://www.semanticscholar.org/paper/003917f77c527c984dcd62b1b162615bc9383692)


### A second look at exome sequencing data: detecting mobile elements insertion in a rare disease cohort

**Why Not Relevant**: The provided paper content contains only a single general summary sentence about detecting pathological mobile element insertions (MEIs) in coding sequences and the impact on exome sequencing diagnostic yield. It does not mention HBS1L, any specific genes, any identified pathogenic variants, patient phenotypes, Mendelian disease names, or mechanistic links involving HBS1L. Without any gene-specific data or variant-level findings, the excerpt cannot provide direct evidence (e.g., identification of a pathogenic HBS1L variant segregating with disease) nor mechanistic evidence (e.g., functional impact of HBS1L disruption) supporting the claim. Because the necessary information to assess association of HBS1L with a Mendelian disease is absent, relevance to the claim is effectively zero.


[Read Paper](https://www.semanticscholar.org/paper/522d0a5dda91f0a8f91bd19d8234dd4127a2c918)


### A Case of PLACK Syndrome with a Novel Splicing Mutation in CAST: The Evidence for Loss-of-Function mechanism through mis-splicing.

**Why Not Relevant**: The paper exclusively reports a genetic and mechanistic investigation implicating CAST in a case of PLACK (Peeling Skin) syndrome. There is no mention of HBS1L, no data or analysis involving HBS1L, and no discussion connecting HBS1L to PLACK or to any Mendelian phenotype. The evidence presented is focused on a different gene (CAST), based on a single patient report with molecular follow-up (mRNA sequencing, segregation/expression analysis). Therefore the paper provides no direct or mechanistic evidence supporting or refuting the claim that HBS1L is likely associated with at least one Mendelian disease.


[Read Paper](https://www.semanticscholar.org/paper/07d7fed6c2e3ebabc373a6601645fe590d3233b1)


### A novel pathogenic variant in MRAP2 in an obese patient with successful outcome of bariatric surgery.

**Why Not Relevant**: The provided paper content exclusively addresses MRAP2 and its role in a monogenic form of severe obesity. There is no mention of HBS1L (or any synonyms) anywhere in the text; no genetic, clinical, functional, or mechanistic data are provided for HBS1L. Therefore the paper gives no direct evidence linking HBS1L to a Mendelian disease, nor does it present mechanistic information that could plausibly be extrapolated to implicate HBS1L. Additionally, the study is a single-patient case report focused on MRAP2 (with limited sample size and limited methodological detail in the excerpt), so even indirect inference by analogy to another gene would be unacceptable. For these reasons relevance to the claim that "HBS1L is likely associated with at least one Mendelian disease" is effectively nil.


[Read Paper](https://www.semanticscholar.org/paper/342b37b3321f360ad6b3b71aed1309973f2b69b0)


### Case report: Two sisters with a germline CHEK2 variant and distinct endocrine neoplasias

**Why Not Relevant**: The provided paper content contains no mention of HBS1L, directly or indirectly. All quoted sentences concern CHEK2 (CHK2), NF1, and the clinical utility of tumor sequencing in two case-report siblings with endocrine tumors. There is no genetic, clinical, mechanistic, statistical, or experimental data related to HBS1L in the text. Therefore the paper does not provide supporting or refuting evidence about HBS1L’s association with any Mendelian disease. Note that absence of mention in this paper does not constitute evidence that HBS1L is not disease-associated overall; it only means this paper is not a source of evidence for that claim.


[Read Paper](https://www.semanticscholar.org/paper/4e8a99f4716428876237e1c0c09dfd9a53b79634)


### Frequency and phenotype associations of rare variants in five monogenic cerebral small vessel disease genes in 200,000 UK Biobank participants with whole exome sequencing data

**Why Not Relevant**: The paper focuses exclusively on a set of cerebral small vessel disease (cSVD) genes (CTSA, TREX1, HTRA1, COL4A1/2) and analyzes variant frequency and phenotype associations for those genes in UK Biobank. HBS1L is not mentioned among the genes reviewed or analyzed, and no experimental, association, or mechanistic data about HBS1L are presented. Therefore this paper provides no direct evidence supporting or refuting the claim that HBS1L is likely associated with at least one Mendelian disease. Note that absence of HBS1L in this study does not imply HBS1L is not associated with Mendelian disease; it only indicates this publication does not address that gene. To evaluate the claim about HBS1L, primary literature, disease databases (e.g., OMIM, ClinVar), or functional studies that specifically examine HBS1L should be consulted.


[Read Paper](https://www.semanticscholar.org/paper/1f0de3de50f290aae016ea35325c46c15e4af3e8)


### Leveraging Clinical, Functional, Molecular and Population Genetic Data Reveals Genotype Phenotype Association and Health Disparity in a Monogenic Disorder, CTX

**Why Not Relevant**: The provided paper content is exclusively about cerebrotendinous xanthomatosis (CTX) and pathogenic variants in CYP27A1 (OMIM 213700). There is no mention of the gene HBS1L, no data, analyses, mechanistic discussion, or population-genetic results concerning HBS1L, and no statements that link HBS1L to CTX or any other Mendelian disorder. Key sentences in the excerpt (for example, the opening sentence that identifies the causal gene as CYP27A1) indicate the study's scope is limited to CYP27A1 genotype–phenotype associations and population frequencies of CYP27A1 alleles. Because the content contains no references to HBS1L and provides no experimental, clinical, or mechanistic evidence relevant to HBS1L, it cannot be used to support or refute the claim that "HBS1L is likely associated with at least one Mendelian disease." If you would like, I can (a) search the literature and curated databases (OMIM, ClinVar, HGMD, PubMed, gnomAD) for evidence linking HBS1L to Mendelian disease, or (b) re-check a larger portion of the paper if additional sections are available that were not included here.


[Read Paper](https://www.semanticscholar.org/paper/ecd9a6ae7be2a66a942f75a505fc4b04dff8e862)


### #5782 AN EXCEPTIONAL CASE OF A PATIENT WITH TWO RARE GENETIC VARIANTS OF AUTOSOMAL DOMINANT POLYCYSTIC KIDNEY DISEASE AND ALPORT SYNDROME

**Why Not Relevant**: This paper does not provide evidence for or against the claim that HBS1L is likely associated with at least one Mendelian disease. The study is a case report concerning Alport syndrome and ADPKD and focuses on genes COL4A3, PKD1, PKD2, and GANAB; HBS1L is not mentioned anywhere in the provided content. The excerpts above show which genes were considered and which variants were found, but none involve HBS1L. Limitations of using this paper to evaluate the claim: (1) absence of mention of HBS1L in a disease-specific case report is not evidence that HBS1L is unrelated to Mendelian disease overall; (2) a single case report targeting kidney phenotypes cannot comprehensively assess gene-disease relationships across other phenotypes; (3) the paper does not provide mechanistic data, segregation analysis, population variant data, functional assays, or case series regarding HBS1L. To evaluate the claim about HBS1L you would need direct evidence such as: pathogenic or likely pathogenic HBS1L variants reported in multiple unrelated affected individuals with concordant phenotypes and segregation data, functional studies demonstrating a deleterious effect consistent with disease mechanism, and/or curated entries in disease databases (OMIM, ClinVar) linking HBS1L to a named Mendelian disorder. This paper neither supplies such evidence nor discusses HBS1L, so its relevance to the claim is negligible.


[Read Paper](https://www.semanticscholar.org/paper/0337e73775f38924bbc68438e6e09327983393d6)


### Exome copy number variant detection, analysis and classification in a large cohort of families with undiagnosed rare genetic disease

**Why Not Relevant**: This paper's content discusses the detection and classification of copy number variants (CNVs) from exome sequencing and reports cohort-level diagnostic yield and numbers of pathogenic CNVs. It does not mention the gene HBS1L, does not present any gene-specific results or case reports implicating HBS1L, and provides no variant- or phenotype-level data that would allow assessment of HBS1L's association with Mendelian disease. Thus the paper provides only very indirect, general mechanistic support (that CNVs can cause Mendelian disease) but no direct or gene-specific evidence for HBS1L. Because the claim is specifically about HBS1L being likely associated with at least one Mendelian disease, the absence of any mention, data, or analysis related to HBS1L means this paper is not a relevant source for supporting or refuting the claim.


[Read Paper](https://www.semanticscholar.org/paper/2b8162a945544b579cfc6f8ac2e7a50d528bfcde)


### Clinical whole Exome Sequencing Reveals Novel Homozygous Missense Variant in the PMPCA Gene causing Autosomal Recessive Spinocerebellar Ataxia

**Why Not Relevant**: The paper exclusively investigates PMPCA (Peptidase, Mitochondrial Processing Subunit Alpha) and reports a novel homozygous PMPCA variant causing SCAR2 in a single patient. There is no mention of the gene HBS1L anywhere in the provided content, no genetic or clinical data involving HBS1L, and no mechanistic discussion linking PMPCA and HBS1L. Because the study's objective, results, and conclusions all pertain to PMPCA, this paper provides no direct genetic evidence that HBS1L is associated with any Mendelian disease, nor does it present mechanistic data that would plausibly implicate HBS1L. Additional limitations relevant to extrapolation: the report appears to be a single-case/family observation (limiting generalizability), focuses on a distinct molecular pathway (mitochondrial protein processing via PMPCA), and does not attempt gene-agnostic claims or broad genotype–phenotype correlations that would include HBS1L. Therefore the paper is not relevant to the claim that HBS1L is likely associated with at least one Mendelian disease.


[Read Paper](https://www.semanticscholar.org/paper/18d53a907d41dc419601512438adb5650b42476c)


### Whole-exome sequencing revealed a novel homozygous missense variant in OSGEP gene: a case report of Galloway–Mowat syndrome in Iran

**Why Not Relevant**: The provided paper content reports a case implicating OSGEP in Galloway–Mowat syndrome and does not mention HBS1L at all. Therefore the paper provides no direct genetic, clinical, or mechanistic evidence regarding whether HBS1L is associated with any Mendelian disease. Because the content is focused exclusively on OSGEP, it is irrelevant to the claim about HBS1L. Note that lack of relevance here does not imply HBS1L is or is not disease-associated in other studies; it only means this paper does not inform that question.


[Read Paper](https://www.semanticscholar.org/paper/ea9cceaa84e1b07a4276db70840e43b0ef851bb5)


## Search Queries Used

- HBS1L mutation patient case report Mendelian disease pathogenic variant loss of function

- HBS1L ClinVar OMIM gene disease association monogenic rare disease

- HBS1L function translation GTPase Pelota PELO ribosome rescue no go decay erythropoiesis fetal hemoglobin regulation HBS1L MYB intergenic GWAS modifier

- exome sequencing HBS1L homozygous variant recessive disease undiagnosed rare disease cohort


## Bibliometric Configuration

- Use Bibliometrics: True

- Author Impact Weight: 0.4

- Citation Impact Weight: 0.4

- Venue Impact Weight: 0.2
