## Supplementary material for "MENDELSEEK: An algorithm that predicts Mendelian Genes and elucidates what makes them special": PHF23.txt

# Claim: The gene PHF23 is likely associated with at least one Mendelian disease.

**Status**: processed

**Overall Rating**: -1

**Explanation**:

No relevant papers were found that support or refute this claim.


## Relevant Papers


## Other Reviewed Papers


### The Burden of Candidate Pathogenic Variants for Kidney and Genitourinary Disorders Emerging From Exome Sequencing

**Why Not Relevant**: The provided paper content does not mention the gene PHF23 by name, nor does it present gene-level results that would allow assessing whether PHF23 is associated with a Mendelian disease. The excerpts give methods, filtering criteria, cohort description, and general observations about variant databases and reclassification, but they do not list the 625 curated genes or report findings for individual genes. While the methodology described (database searches, ClinVar/HGMD reliance, LoF criteria, MAF filters) shows how the authors would have assessed a gene like PHF23, there is no direct or indirect data in the provided text identifying PHF23 as included in the curated gene list, having pathogenic variants, or being implicated mechanistically in any Mendelian phenotype. It is possible that PHF23 appears in the full supplemental gene list referenced (laboratory Web site or appendixes), but that information is not present in the supplied content. Therefore the supplied paper content is insufficient to support or refute the claim about PHF23.


[Read Paper](https://www.semanticscholar.org/paper/a75e297208c04992568e607d19aa9b9aaebd09d5)


### Genetic Diagnostic Evaluation of Trio-Based Whole Exome Sequencing Among Children With Diagnosed or Suspected Autism Spectrum Disorder

**Why Not Relevant**: The paper provides essentially no evidence for or against the claim that PHF23 is likely associated with at least one Mendelian disease. The study reports results from whole-exome sequencing of 80 simplex families with ASD or suspected ASD and explicitly lists seven genes identified as presumed causative (CHD8, AFF2, ADNP, POGZ, SHANK3, IL1RAPL1, and PTEN); PHF23 is not mentioned anywhere in the provided text. Therefore there is no direct evidence (no reported PHF23 variants, segregation, or disease association) and no mechanistic discussion involving PHF23 in this paper.

Additional important limitations explaining why the paper cannot inform the claim:
- Study scope: The cohort is limited to 80 simplex ASD families (Chinese children, CNV-negative), so negative findings for PHF23 only indicate that this study did not find PHF23-linked Mendelian disease in that specific cohort and phenotype, not that PHF23 is unrelated to Mendelian disease more broadly.
- Assay limitations: The study used whole-exome sequencing, which does not reliably capture noncoding variants, some structural variants, deep intronic variants, or repeat expansions that could implicate a gene in disease.
- Statistical/power limits: Sample size is modest for detecting rare gene-disease associations; failure to observe PHF23 variants could be due to low power rather than true absence of effect.
- Phenotype specificity: The study focused on ASD; if PHF23 is associated with a Mendelian disease with different clinical features, this study would not be expected to detect it.

In sum, relevance is near zero because PHF23 is not reported or discussed in the paper; the paper neither supports nor refutes the claim. To evaluate the claim about PHF23, one should examine other sources such as ClinVar, OMIM, gnomAD constraint metrics, relevant case reports or cohorts, and functional/mechanistic studies specifically addressing PHF23.


[Read Paper](https://www.semanticscholar.org/paper/e83590642a0ed51634cc31d0b28fac2e3ec75069)


### Parkinsonism, Intellectual Disability, and Catatonia in a Young Male With MECP2 Variant

**Why Not Relevant**: This paper focuses exclusively on MECP2 (Methyl CpG-binding protein 2) deficiency and its phenotypic spectrum in a male patient (Rett syndrome / MeCP2-related neurodevelopmental disorder and associated parkinsonian features). The text repeatedly and only names MECP2 as the gene of interest (for example, descriptions of the novel MECP2 c.503G>A (p.Arg168Gln) variant, discussion of MECP2 A140V, and statements that "MeCP2 deficiency accounts for 1.3% to 1.7% of males with cognitive impairment"). There is no mention of the gene PHF23 anywhere in the provided content, nor is there any discussion of pathways, mechanisms, variant evidence, segregation data, functional studies, or clinical phenotypes related to PHF23. Consequently: (1) there is no direct evidence in this paper that PHF23 is associated with any Mendelian disease, and (2) there is no mechanistic evidence tying PHF23 to disease-relevant biology presented here. The absence of PHF23 in this report means it cannot be used to support the claim that PHF23 is likely associated with at least one Mendelian disease. Note that absence of evidence in this single paper does not disprove an association overall; it only indicates that this paper provides no relevant data for evaluating the PHF23 claim. To evaluate the claim, one would need papers that specifically report PHF23 variants segregating with disease, functional studies of PHF23 perturbation, or credible statistical genetic evidence implicating PHF23 in Mendelian disorders.


[Read Paper](https://www.semanticscholar.org/paper/37bdfd82e4a0244261cf29dd7be95410ac155410)


### Connectome Analysis in an Individual with SETD1B-Related Neurodevelopmental Disorder and Epilepsy

**Why Not Relevant**: The provided paper content exclusively describes findings concerning SETD1B and a de novo missense variant c.5699A>G, p.(Tyr1900Cys) in SETD1B. There is no mention of the gene PHF23, nor of any data, results, discussion, or mechanistic analysis involving PHF23 in the supplied text. Specific elements from the supplied content that make it irrelevant to the claim include the abstract's focus: (1) the objective statement that causal variants in SETD1B have been associated with a neurodevelopmental phenotype, (2) the methods and results describing a single individual with a SETD1B variant and associated fMRI findings, and (3) the conclusion that expands the phenotype of SETD1B-related disorder. Because PHF23 is not named or discussed, there is neither direct genetic/clinical evidence nor mechanistic data in this paper excerpt that would support or refute the claim that PHF23 is likely associated with at least one Mendelian disease.

Limitations and caveats: (1) The supplied text notes that supplementary material is available on the journal website; I do not have that supplementary material here. It is possible (though seems unlikely given the abstract's clear focus) that PHF23 could be mentioned in supplementary data; without that material I cannot rule that out entirely. (2) The conclusion that this excerpt provides is strictly about SETD1B, so no extrapolation to PHF23 can be made from the provided content. 

Recommended next steps to evaluate the claim: search the full article (including supplements) for PHF23; perform a literature search (PubMed), and query disease-gene databases (OMIM, ClinVar, HGMD, DECIPHER, PanelApp) and large-scale population/constraint metrics (gnomAD, pLI/LOEUF) for PHF23 to find direct genetic or mechanistic evidence linking PHF23 to Mendelian disease.


[Read Paper](https://www.semanticscholar.org/paper/b31a3eb73aef8d82ab5fc80595d2b4bdb8d45def)


### The multiple faces of TOR1A: different inheritance, different phenotype.

**Why Not Relevant**: The provided paper content does not mention the gene PHF23 anywhere and contains no experimental, genetic, clinical, or epidemiological data regarding PHF23. All excerpts are general background about Mendelian genetics or a detailed example concerning a different gene (TOR1A). While the paper's discussion of how single genes can underlie Mendelian disorders (and how inheritance can vary by allele/zygosity) is mechanistically relevant in the abstract, it cannot be used as direct evidence that PHF23 is associated with any Mendelian disease. Therefore the content is largely irrelevant to the specific claim that PHF23 is likely associated with at least one Mendelian disease.


[Read Paper](https://www.semanticscholar.org/paper/b74b91e0829b14562a5293323e09bb3633156aab)


### Burden re-analysis of neurodevelopmental disorder cohorts for prioritization of candidate genes.

**Why Not Relevant**: The provided paper content is a single, high-level methodological summary about how the authors identify novel genes implicated in neurodevelopmental disorders. It contains no mention of the gene PHF23, no PHF23-specific variant or case data, no statistics (e.g., burden p-values or variant counts) for PHF23, and no mechanistic or functional data concerning PHF23. Therefore there is effectively no direct or mechanistic evidence in the supplied text to support or refute the claim that PHF23 is likely associated with at least one Mendelian disease. 

What would be required to evaluate the claim: explicit PHF23-focused evidence such as (1) observed de novo or segregating rare variants in PHF23 in multiple unrelated affected individuals with consistent phenotypes and appropriate statistical enrichment (burden or mutation rate analysis with p/q values), (2) segregation data showing Mendelian inheritance in families, (3) detailed clinical phenotypes linked to PHF23 variants, and/or (4) mechanistic/functional data (cellular assays, model organisms, expression patterns) demonstrating how PHF23 disruption leads to disease. None of these are present in the provided excerpt. 

Note: it is possible that the full paper (beyond the single sentence provided) includes PHF23-specific results in the results section, figures, tables, or supplementary materials. To make a reliable determination about PHF23 one would need those specific sections (gene hit lists, variant tables, case descriptions, statistical analyses, and functional follow-up).


[Read Paper](https://www.semanticscholar.org/paper/0a7e56a69b7001fbe8ee5a57ca3d63c5bc7951cc)


### Case Report: A Disease Phenotype of Rett Syndrome and Neurofibromatosis Resulting from A Bilocus Variant Combination

**Why Not Relevant**: The provided paper content does not mention PHF23 or present any gene-specific evidence. The single-sentence excerpt reports a clinical observation of overlapping ASD and Rett-like features and broadly suggests multilocus interactions at a protein or cellular level, but it gives no genetic, molecular, or statistical data tying PHF23 to disease. To support the claim that PHF23 is likely associated with a Mendelian disease the paper would need one or more of: explicit identification of pathogenic PHF23 variants in affected individuals, segregation of such variants with disease in families or recurrent de novo variants, functional studies showing that PHF23 disruption causes a plausible disease mechanism, or statistical enrichment of PHF23 variants in cases versus controls. None of these elements are present in the provided text, so the content is insufficient to support or refute the claim about PHF23.


[Read Paper](https://www.semanticscholar.org/paper/993edabc0eeab1343cda0a7dbad248110f7e87ad)


### Correspondence on “Pathogenic variants in USP7 cause a neurodevelopmental disorder with speech delays, altered behavior, and neurologic anomalies” by Fountain et al.

**Why Not Relevant**: The provided paper content exclusively discusses a de novo heterozygous variant in USP7 and clinical findings related to USP6 disorder (including fetal complications and cardiac involvement). There is no mention of PHF23, PHF23 variants, phenotypes attributable to PHF23, nor mechanistic pathways involving PHF23. Consequently the content is not relevant to the claim that PHF23 is likely associated with at least one Mendelian disease. If you can provide additional text from this paper or other papers that mention PHF23, I can re-evaluate for direct or mechanistic evidence. For independent assessment of the claim, consider searching databases such as OMIM, ClinVar, HGMD, and relevant peer-reviewed literature for PHF23 pathogenic variants, reported syndromes, functional studies, or segregation data.


[Read Paper](https://www.semanticscholar.org/paper/b3775e0fbd9d9b5506cd4a54ed1e5e2178151d54)


### De novo HDAC3 variants leading to epigenetic machinery dysfunction are associated with a neurodevelopmental disorder

**Why Not Relevant**: The provided paper content exclusively describes heterozygous de novo variants in HDAC3 and a novel neurodevelopmental syndrome linked to those HDAC3 variants. Nowhere in the supplied text is the gene PHF23 mentioned, nor are there any data, experiments, or mechanistic descriptions that implicate PHF23 (e.g., genetic variants in PHF23, PHF23 expression or localization changes, PHF23 protein interactions, or phenotypes in models related to PHF23). Because the claim concerns PHF23 being associated with at least one Mendelian disease, and the supplied content contains no reference to PHF23 or evidence that could be used to infer such an association, this paper excerpt provides no direct or mechanistic evidence for or against the claim.

Limitations and caveats:
- Assessment is limited to the supplied paper content. It is possible (but not indicated in the provided text) that the full paper contains additional sections, supplementary materials, or references that mention PHF23; those were not provided and therefore were not assessed. If you can supply the full text or search the full paper for "PHF23", I can re-check.
- Indirect inference (for example, if PHF23 were known from other literature to interact with HDAC3) cannot be made from this excerpt because no interaction data or network analyses including PHF23 are present here. Establishing PHF23 as a Mendelian disease gene would require (a) human genetic evidence (de novo or segregating pathogenic variants), (b) phenotypic correlation, and/or (c) compelling mechanistic data linking PHF23 perturbation to disease phenotypes; none of these are present in the supplied HDAC3-focused content.

Recommendation: To evaluate the claim about PHF23, please provide the specific paper(s) that mention PHF23 or allow a literature/database search (PubMed, OMIM, ClinVar, gnomAD, DECIPHER) for PHF23 (gene symbol) to identify any human genetic evidence or experimental mechanistic studies linking PHF23 to Mendelian disease.


[Read Paper](https://www.semanticscholar.org/paper/524b09f9e39f306b6871f816044577dccb53abd6)


### Novel de Novo Nonsense Variants in AGO3 and KHSRP: Insights into Global Developmental Delay and Autism Spectrum Disorders through Whole Genome Analysis

**Why Not Relevant**: The paper's entire provided content centers on a single patient and on de novo nonsense variants in AGO3 and KHSRP as potential contributors to global developmental delay and autistic features. Nowhere in the provided text is the gene PHF23 mentioned, discussed, or analyzed. There are no experimental data, case observations, literature citations, mechanistic proposals, or statistical analyses concerning PHF23. Therefore the paper provides no direct evidence (variant observations, case reports, cosegregation, functional assays) nor mechanistic evidence (biological function, pathway involvement, animal models) relevant to whether PHF23 is associated with a Mendelian disease. Given the absence of any PHF23 information, the document cannot be used to support or refute the claim about PHF23; its relevance is effectively nil. The small nonzero relevance score (0.02) reflects that the paper is about Mendelian/NDD context generally (background), but that general context does not meaningfully inform PHF23’s disease association.


[Read Paper](https://www.semanticscholar.org/paper/5aba59259e6535c76aa2fd80054cb774eb3d7452)


## Search Queries Used

- PHF23 PHD finger protein 23 gene pathogenic variant case report Mendelian disease monogenic disorder clinical phenotype

- PHF23 de novo variants exome sequencing trio cohort neurodevelopmental disorder intellectual disability autism rare disease ClinVar OMIM

- PHF23 PHD finger chromatin binding histone binding functional study mouse model zebrafish knockdown loss of function cellular phenotype transcriptional regulation

- PHF23 loss of function intolerance pLI gnomAD constraint haploinsufficiency essential gene haploinsufficiency score gene disease association review PHF family PHD finger genes Mendelian disease


## Bibliometric Configuration

- Use Bibliometrics: True

- Author Impact Weight: 0.4

- Citation Impact Weight: 0.4

- Venue Impact Weight: 0.2
