## Supplementary material for "MENDELSEEK: An algorithm that predicts Mendelian Genes and elucidates what makes them special": PIK3R4.txt

# Claim: The gene PIK3R4 is likely associated with at least one Mendelian disease.

**Status**: processed

**Overall Rating**: -1

**Explanation**:

No relevant papers were found that support or refute this claim.


## Relevant Papers


## Other Reviewed Papers


### Discovery and functional prioritization of Parkinson’s disease candidate genes from large-scale whole exome sequencing

**Why Not Relevant**: The provided paper content does not mention PIK3R4 at all; it only states that 27 genes with homozygous or compound heterozygous loss-of-function variants were identified in PD cases and that several PD susceptibility candidates were nominated. To assess the claim that PIK3R4 is likely associated with at least one Mendelian disease, one would need explicit mention of PIK3R4 among the identified genes or separate evidence showing: (a) PIK3R4 loss- or gain-of-function variants segregating with disease in families consistent with Mendelian inheritance, (b) pathogenic classification of variants in PIK3R4, (c) strong functional/mechanistic data linking PIK3R4 disruption to a Mendelian phenotype, or (d) replication in independent cohorts. None of those elements are present in the excerpt. Additionally, the study appears focused on PD susceptibility and does not by itself demonstrate that any of the identified genes cause classical Mendelian diseases; association in case cohorts (even with homozygous/compound heterozygous LoF variants) is not equivalent to establishing causality for a Mendelian disorder without family segregation, high penetrance, or mechanistic proof. Therefore the excerpt is insufficient to support or refute the claim regarding PIK3R4.


[Read Paper](https://www.semanticscholar.org/paper/489d796282e50673b7189ad7a170224d5ce73767)


### Abstract 18770: Elucidation of MRAS-Mediated Noonan Syndrome With Cardiac Hypertrophy

**Why Not Relevant**: The provided paper content exclusively reports discovery, modeling, and cohort screening implicating MRAS as a Noonan syndrome (Mendelian) gene and describes MRAS's interactions in the Ras/MAPK pathway. The text contains no mention of PIK3R4, no experiments or analyses involving PIK3R4, and no mechanistic discussion linking PIK3R4 to Noonan syndrome or any other Mendelian disease. Therefore the paper provides no direct or mechanistic evidence supporting the claim that PIK3R4 is likely associated with at least one Mendelian disease. Absence of evidence in this paper does not prove PIK3R4 is not associated with a Mendelian disease, but it does mean this paper cannot be used to support the claim.


[Read Paper](https://www.semanticscholar.org/paper/eed57cf29dcfb8c6e5fea6cfdd54b0603c83e163)


### Molecular findings among patients referred for clinical whole-exome sequencing: Editorial comment

**Why Not Relevant**: The provided paper content describes a large clinical whole-exome sequencing cohort, methods of interpretation and reporting, aggregate counts of variants, diagnostic yield, and a general conclusion that exome sequencing can detect rare and novel variants. However, the text contains no mention of the gene PIK3R4, no gene-specific results, and no case reports or analyses implicating PIK3R4 in any Mendelian phenotype. Therefore the paper provides no direct evidence that PIK3R4 is associated with a Mendelian disease. The excerpts chosen show mechanistic plausibility (that exome sequencing can discover novel gene-disease associations) but cannot substitute for direct gene-level evidence. Given the absence of any PIK3R4-specific data, the paper is not relevant for establishing that PIK3R4 is likely associated with at least one Mendelian disease.


[Read Paper](https://www.semanticscholar.org/paper/a122c53f80c050b861b70f5e57fbede67e0b7263)


### Precision Medicine in Internal Medicine

**Why Not Relevant**: The provided paper content is a general review of genetic testing approaches, exome/genome sequencing, variant interpretation, and the state of gene discovery as of 2019. It contains no mention of the gene PIK3R4, no case reports, cohort studies, variant-level data, segregation analyses, functional studies, or any other empirical evidence linking PIK3R4 to a Mendelian disorder. Therefore it does not provide direct evidence for or against the claim that PIK3R4 is likely associated with at least one Mendelian disease. The excerpts included are contextual and mechanistic (they describe how exome sequencing can discover gene–disease associations and the overall fraction of genes already linked to Mendelian disease) but do not address PIK3R4 specifically. Because the paper lacks gene-specific data, its relevance to the claim is minimal.


[Read Paper](https://www.semanticscholar.org/paper/c7199e8a5226f8d9d55777da0e15e91797bb459a)


### Biallelic variants identified in 36 Pakistani families and trios with autism spectrum disorder

**Why Not Relevant**: The provided paper content contains only the single sentence: "Information on the genetic variability associated with ASD in Pakistan is offered, and a marked enrichment for biallelic variants over that reported in outbreeding populations is demonstrated." This sentence is a high-level summary about autism spectrum disorder (ASD) genetics in a specific population and a general observation about enrichment for biallelic variants. It does not mention the gene PIK3R4, it does not report any gene-level associations, Mendelian disease links, variant lists, functional or mechanistic data relating to PIK3R4, nor any statistical evidence concerning that gene. Therefore, based on the content provided, the paper gives no direct evidence (no reported variants, no case reports, no segregation, no pathogenicity assertions) and no mechanistic evidence (no functional studies, pathway links, or model organism data) to support or refute the claim that PIK3R4 is likely associated with at least one Mendelian disease.

Limitations and caveats: (1) The assessment is limited to the single sentence you provided; the full paper, figures, tables, supplementary materials, or gene-specific lists might contain relevant data about PIK3R4 that is not present in this excerpt. (2) The sentence’s observation that biallelic variants are enriched in this population is a general point that increases the prior probability that recessive Mendelian disease genes may be discovered in such cohorts, but this is only an indirect, population-level consideration and does not constitute evidence specific to PIK3R4. (3) To properly evaluate the claim about PIK3R4, one would need the full text (especially methods, results, variant tables, and supplementary data), and external resources such as OMIM, ClinVar, gnomAD, and functional studies.

Recommended next steps to test the claim: search the full paper and its supplementary tables for any mentions of PIK3R4 or its variants; search ClinVar/OMIM for PIK3R4 disease associations; review published case reports or functional studies implicating PIK3R4 in Mendelian phenotypes.


[Read Paper](https://www.semanticscholar.org/paper/17be98b475c43e8e296ed0d13d54250c91e9c42d)


### Contribution of rare transmitted and de novo variants among 2 , 871 1 congenital heart disease probands

**Why Not Relevant**: I found no mention of the gene PIK3R4 in the provided paper content. The excerpt focuses on congenital heart disease (CHD) cohort sequencing, numbers of de novo and transmitted mutations, and specific genes (for example GDF1, MYH6, FLT4) and gene-set enrichment results, but it does not report any data, variants, mechanistic discussion, or gene lists that include PIK3R4. Therefore the paper content as provided gives neither direct evidence (no reported PIK3R4 variants, case counts, statistical tests, or clinical correlations) nor mechanistic evidence (no discussion of PIK3R4 function, pathways, or model systems) supporting an association between PIK3R4 and a Mendelian disease.

Limitations and caveats: 1) My judgment is based solely on the text you supplied. The full paper (including main text beyond this excerpt, figures, and supplementary files referenced here such as Supplementary Excel tables) might include additional gene lists or data that were not included in the excerpt and could mention PIK3R4. 2) The excerpt does describe methods (exome sequencing, analysis of de novo and recessive genotypes, enrichment tests) that are appropriate for discovering Mendelian disease genes, so the study design could in principle detect a PIK3R4 association if variants in that gene were observed and reported—but none are present in the supplied text.

Recommended next steps to determine whether PIK3R4 is linked to a Mendelian disease: search the full paper and all supplementary tables for PIK3R4; query disease and variant databases (OMIM, ClinVar, HGMD), large population datasets (gnomAD) for rare pathogenic variants and constraint metrics, and literature databases (PubMed) for case reports or functional studies implicating PIK3R4. If you want, I can (1) scan the full paper/supplementary material if you provide it, or (2) query public databases/literature and summarize evidence regarding PIK3R4 and Mendelian disease associations.


[Read Paper](https://www.semanticscholar.org/paper/e5830d9e95469f22518560007746202e62db26a0)


## Search Queries Used

- PIK3R4 VPS15 p150 mutation pathogenic variant Mendelian disease monogenic inherited disorder case report family segregation de novo

- PIK3R4 whole exome sequencing rare disease cohort de novo variants autosomal dominant autosomal recessive homozygous compound heterozygous loss of function

- PIK3R4 VPS15 autophagy endosomal trafficking VPS34 complex neuronal development neurodegeneration model mouse zebrafish human phenotype mechanism

- PIK3R4 ClinVar OMIM Decipher ClinGen haploinsufficiency loss of function intolerance pLI gene dosage copy number variant contiguous gene deletion review


## Bibliometric Configuration

- Use Bibliometrics: True

- Author Impact Weight: 0.4

- Citation Impact Weight: 0.4

- Venue Impact Weight: 0.2
