## Supplementary material for "MENDELSEEK: An algorithm that predicts Mendelian Genes and elucidates what makes them special": PLK1.txt

# Claim: The gene PLK1 is likely associated with at least one Mendelian disease.

**Status**: processed

**Overall Rating**: -1

**Explanation**:

No relevant papers were found that support or refute this claim.


## Relevant Papers


## Other Reviewed Papers


### Genetic and phenotypic analysis of 101 patients with developmental delay or intellectual disability using whole‐exome sequencing

**Why Not Relevant**: The provided paper content (abstract) does not mention the gene PLK1, any variants in PLK1, nor any mechanistic pathways explicitly involving PLK1. The text focuses on whole-exome sequencing in 101 Japanese patients with unexplained developmental delay/intellectual disability, diagnostic yield (54 cases, 53.5%), copy number variants, de novo variants, associations with autism spectrum disorder and external ear anomalies, and the importance of reanalysis and detection of CNVs and somatic variants. None of these sentences or paragraphs provide direct evidence (e.g., identification of pathogenic PLK1 variants in patients) or mechanistic evidence (e.g., PLK1's role in a disease pathway) that would support or refute the claim that PLK1 is likely associated with at least one Mendelian disease. Therefore, based on the supplied content, the paper is not relevant to the claim.

Limitations/caveat: analysis is restricted to the text you supplied. If the full paper (beyond this abstract) contains mention of PLK1 or relevant variant findings, that content was not available here. To evaluate PLK1 specifically, one should search the full paper text and supplementary materials for "PLK1" or reported gene lists, or consult other studies that explicitly report PLK1 pathogenic variants associated with Mendelian phenotypes.


[Read Paper](https://www.semanticscholar.org/paper/c4a6b545539bc97495aa04ca6401c2b071178038)


### Diagnostic yield of patients with undiagnosed intellectual disability, global developmental delay and multiples congenital anomalies using karyotype, microarray analysis, whole exome sequencing from Central Brazil

**Why Not Relevant**: The paper text provided does not mention PLK1 anywhere and contains no gene-level results or mechanistic discussion specific to PLK1. While the study used an ID gene panel of 1,252 genes and performed exome sequencing on 19 trios, the authors did not list the panel genes nor report findings for individual genes in the excerpt. Therefore: (1) there is no direct evidence (no reported PLK1 variants, co-segregation, case reports, or statistical association) to support that PLK1 is associated with a Mendelian disease; (2) there is no mechanistic evidence in this text linking PLK1 function to an ID phenotype; and (3) absence of evidence here cannot be taken as evidence of absence — the panel may have included PLK1 and either found no pathogenic variants in these 19 trios or simply did not report it in the provided excerpt. Additionally, the exome cohort is small (19 trios), limiting power to detect rare PLK1-associated Mendelian conditions. To assess the claim reliably would require explicit gene-level findings (variants in PLK1 with pathogenic classification and segregation/functional data) or independent literature reporting PLK1 mutations causing Mendelian disease.


[Read Paper](https://www.semanticscholar.org/paper/23767daf16f48d9a91172aa0fc624f07b869a9c8)


### Expanding phenotype of MED13-associated syndrome presenting novel de novo missense variant in a patient with multiple congenital anomalies

**Why Not Relevant**: The provided paper content exclusively describes a case involving a de novo missense variant in MED13 and proposes expansion of the MED13-associated phenotype. There is no mention of PLK1, PLK1 variants, PLK1-associated phenotypes, or mechanistic pathways linking MED13 to PLK1. Because the claim concerns PLK1 being associated with a Mendelian disease, and the paper does not discuss PLK1 in any capacity (no genetic, clinical, experimental, or pathway data involving PLK1), the paper provides no direct or mechanistic evidence for or against the claim. Therefore its relevance to the claim is effectively zero. If further evaluation is desired, papers that specifically study PLK1 variants, PLK1 knockouts, or clinical cohorts with PLK1 mutations (or that demonstrate MED13–PLK1 interactions with phenotypic consequences) would be needed to assess the claim.


[Read Paper](https://www.semanticscholar.org/paper/9b2cc0996fdf5fb478e07dbf06be71de50a057ee)


### De novo HDAC3 variants leading to epigenetic machinery dysfunction are associated with a neurodevelopmental disorder

**Why Not Relevant**: This paper focuses entirely on HDAC3 and reports heterozygous de novo missense variants in HDAC3 (D93N, A110T, P201S, L266S, G267S, and R359C) linked to a novel neurodevelopmental syndrome. The content describes HDAC3-specific functional assays (reduced deacetylation activity for L266S and G267S), disrupted interactions with CoREST and NCoR (A110T), and impaired nuclear localization for several HDAC3 variants. Nowhere in the provided text is the gene PLK1 mentioned, nor are PLK1-related pathways, proteins, or disease associations discussed. Therefore the paper provides neither direct evidence (no observations or reported variants in PLK1) nor mechanistic evidence (no biochemical, interaction, or pathway data involving PLK1) bearing on the claim that PLK1 is likely associated with at least one Mendelian disease. Absence of evidence in this paper does not disprove the claim overall; it simply means this paper is irrelevant to evaluating PLK1’s disease association. To assess the claim about PLK1, one should consult sources that specifically report PLK1 variants and phenotypes (e.g., OMIM, ClinVar, PubMed case reports, or population / constraint databases such as gnomAD).


[Read Paper](https://www.semanticscholar.org/paper/524b09f9e39f306b6871f816044577dccb53abd6)


### A novel AUTS2 variant in a patient with global developmental delay and intellectual disability

**Why Not Relevant**: The provided paper content concerns a novel heterozygous, likely loss-of-function variant in the AUTS2 gene and the resulting AUTS2 haploinsufficiency syndrome. There is no mention of the gene PLK1, no PLK1 variants, no experimental data involving PLK1, and no mechanistic discussion linking AUTS2 (or any other gene discussed) to PLK1. Therefore the paper contains no direct evidence that PLK1 is associated with any Mendelian disease, nor any mechanistic evidence that would support or refute such an association. Because the absence of any reference to PLK1 means neither supporting nor contradicting data are present, the paper is not relevant to the claim.


[Read Paper](https://www.semanticscholar.org/paper/f5e84e7bb3b42239a53cc7e5c457342cb38c6277)


## Search Queries Used

- PLK1 ClinVar OMIM HGMD ClinGen gene disease association germline pathogenic variant PLK1 Mendelian

- PLK1 de novo variant exome sequencing trio developmental disorder intellectual disability congenital anomaly DDD Deciphering Developmental Disorders

- Plk1 knockout mouse embryonic lethal heterozygous phenotype haploinsufficiency developmental defects centrosome mitotic regulation germline mutation model organism

- PLK1 germline predisposition hereditary cancer syndrome family segregation linkage gene panel systematic review ClinGen curation monogenic inherited


## Bibliometric Configuration

- Use Bibliometrics: True

- Author Impact Weight: 0.4

- Citation Impact Weight: 0.4

- Venue Impact Weight: 0.2
