## Supplementary material for "MENDELSEEK: An algorithm that predicts Mendelian Genes and elucidates what makes them special": PLRG1.txt

# Claim: The gene PLRG1 is likely associated with at least one Mendelian disease.

**Status**: processed

**Overall Rating**: 0

**Explanation**:

Supporting evidence:
The provided paper excerpt documents a molecular role for PLRG1: it interacts with hnRNP-M and CDC5L and is implicated in splice site choice. This establishes PLRG1 as a component of the spliceosome or splicing-regulatory machinery and indicates that perturbing its interaction domain alters alternative splicing of a reporter substrate. Such molecular data show PLRG1 is functionally important for RNA processing in cells.

Caveats or contradictory evidence:
The excerpt does not report any human genetic data, clinical phenotypes, segregating variants, linkage, or association with disease. There is no mention of mutations in PLRG1 causing a Mendelian disorder, nor presentation of phenotypic consequences in organisms or patients tied to PLRG1 loss- or gain-of-function. The paper describes an in vitro functional effect on splicing of a mini-gene; this does not constitute evidence that germline variants in PLRG1 cause a Mendelian disease in humans.

Analysis of potential underlying mechanisms, deeper relationships, or second-order implications:
Given PLRG1's role in core splicing machinery, it is biologically plausible that deleterious variants could lead to widespread splicing defects and, consequently, disease phenotypes. Many genes encoding spliceosomal components are intolerant to loss-of-function and some are implicated in human disorders. However, plausibility does not substitute for direct evidence. Demonstrating a Mendelian disease association requires genetic evidence (segregation in families, de novo mutations in affected individuals, population genetics consistent with pathogenicity) or strong experimental evidence linking variants to disease-relevant phenotypes in vivo.

Explanation of rating choice:
Because the only provided evidence is a molecular interaction and functional effect on a reporter substrate, and there is no genetic, clinical, or organismal phenotype data linking PLRG1 variants to a Mendelian disorder, the supplied materials do not support the claim that PLRG1 is likely associated with at least one Mendelian disease. The data make the hypothesis plausible but do not provide supporting evidence. Therefore the appropriate assessment based on the provided papers is that there is no evidence for a Mendelian disease association.


**Final Reasoning**:

I reviewed the logic and evidence: the excerpt demonstrates PLRG1's role in splicing, which increases biological plausibility that perturbation could cause disease. However, the claim is categorical (PLRG1 is likely associated with at least one Mendelian disease) and requires genetic/clinical evidence that is absent in the provided material. No contradictory genetic evidence was provided either (e.g., evidence that deleterious variants are tolerated), but absence of supportive data leads to the conclusion that there is no evidence in the supplied papers to support the claim. Thus the rating 'No Evidence' is appropriate.


## Relevant Papers


### Direct interaction between hnRNP‐M and CDC5L/PLRG1 proteins affects alternative splice site choice

**Authors**: D. Llères (H-index: 21), A. Lamond (H-index: 96)

**Relevance**: 0.25

**Bibliometric Impact**: 0.5618571428571428


**Excerpts**:

- Heterogeneous nuclear ribonucleoprotein‐M is an abundant nuclear protein that binds to pre‐mRNA and is a component of the spliceosome complex and an hnRNP‐M mutant lacking the CDC5L/PLRG1 interaction domain is unable to modulate alternative splicing of an adeno‐E1A mini‐gene substrate.


**Explanations**:

- This sentence is mechanistic evidence that ties PLRG1 (as part of a CDC5L/PLRG1 interaction) into the spliceosome machinery and into control of alternative splicing. It shows that disruption of the interaction domain (in hnRNP‑M) abolishes modulation of alternative splicing of a reporter (an adeno‑E1A mini‑gene), which supports biological plausibility that perturbing components or interactions involving PLRG1 could affect pre‑mRNA splicing. Strengths: it identifies PLRG1 as functionally connected to core splicing factors and to regulation of alternative splicing — a mechanism by which variants could produce pathogenic effects. Limitations/caveats: this is not direct evidence of PLRG1 causing any Mendelian disease. The experiment describes a mutation in hnRNP‑M that removes the CDC5L/PLRG1 interaction domain, not a mutation in PLRG1 itself. The assay appears to use an artificial reporter substrate (adeno‑E1A mini‑gene), so results may not generalize to endogenous transcripts or to organismal phenotypes. The sentence provides no genetic, clinical, or statistical data linking PLRG1 variants to disease, no information about species or cell type, and no experimental details (methods, sample size, effect sizes). Therefore the evidence increases plausibility but does not constitute direct support that PLRG1 is associated with a Mendelian disease.


[Read Paper](https://www.semanticscholar.org/paper/953532d95e26ee9c9963aaf8e278486f1f67dbb6)


## Other Reviewed Papers


### Implications of a De Novo Variant in the SOX12 Gene in a Patient with Generalized Epilepsy, Intellectual Disability, and Childhood Emotional Behavioral Disorders

**Why Not Relevant**: This paper does not provide evidence relevant to the claim that PLRG1 is likely associated with at least one Mendelian disease. The text exclusively discusses SOX family genes and a de novo variant in SOX12. For example, the paper states: "SRY-box transcription factor (SOX) genes, a recently discovered gene family, play crucial roles in the regulation of neuronal stem cell proliferation and glial differentiation during nervous system development and neurogenesis." and "Whole exome sequencing (WES) in patients presenting with generalized epilepsy, intellectual disability, and childhood emotional behavioral disorder, uncovered a de novo variation within SOX12 gene." The manuscript further focuses on SOX12-specific observations (e.g., HMG domain, conservation within SoxC, in silico pathogenicity predictions) and does not mention PLRG1, present any genetic data on PLRG1, or propose mechanistic links between SOX12 and PLRG1. Because there is no direct genetic evidence, no mechanistic pathway, and no discussion of PLRG1 in the paper, it cannot support or refute the claim about PLRG1’s association with a Mendelian disease. Note also that absence of discussion of PLRG1 in this SOX12-focused report is not negative evidence against PLRG1’s involvement in Mendelian disease—rather, it is simply unrelated. To evaluate the claim about PLRG1 would require papers that specifically present PLRG1 variants segregating with Mendelian phenotypes, functional studies of PLRG1 perturbation consistent with disease mechanisms, or curated clinical genetics database entries linking PLRG1 to a Mendelian disorder.


[Read Paper](https://www.semanticscholar.org/paper/5cf652470efa4406f72843aea8389f4e398a99f8)


### Elevated level of PLRG1 is critical for the proliferation and maintenance of genome stability of tumor cells

**Why Not Relevant**: The paper provides functional and mechanistic data about PLRG1 (spliceosomal role, involvement in pre-mRNA splicing and DNA repair, effects of experimental knockdown in cancer and normal cells), but it contains no direct human genetic evidence connecting PLRG1 variants to a Mendelian, inherited disease. Specifically, there are no reports of germline mutations in PLRG1, no case reports or pedigrees showing segregation of PLRG1 variants with disease, no population genetics or variant pathogenicity analyses, and no in vivo organismal phenotypes tied to inherited PLRG1 disruption. The experimental evidence is largely from somatic/cell-line knockdown or expression studies in cancer contexts; such perturbations demonstrate biological importance and mechanistic plausibility but do not establish that PLRG1 is causally associated with an inherited monogenic disorder. Therefore the content is only weakly relevant (mechanistic plausibility) and does not constitute direct evidence for the claim.


[Read Paper](https://www.semanticscholar.org/paper/b700736d5478e84916c9ff9f02f731e8b9f827c0)


### Expanding phenotype of MED13-associated syndrome presenting novel de novo missense variant in a patient with multiple congenital anomalies

**Why Not Relevant**: The provided paper content exclusively describes a de novo p.Pro835Ser missense variant in MED13 and proposes to expand the MED13-associated phenotype. There is no mention of the gene PLRG1, no reported PLRG1 variants, and no mechanistic discussion connecting MED13 to PLRG1. Consequently, the paper contains neither direct evidence (variants, segregation, phenotype attribution) nor mechanistic evidence (pathway interaction, functional studies) relevant to whether PLRG1 is associated with a Mendelian disease. Additionally, the study appears to be a single-case report focused on MED13, which further prevents extrapolation to other genes. Because the text contains no content about PLRG1, its relevance to the claim that "PLRG1 is likely associated with at least one Mendelian disease" is effectively zero.


[Read Paper](https://www.semanticscholar.org/paper/9b2cc0996fdf5fb478e07dbf06be71de50a057ee)


### De novo HDAC3 variants leading to epigenetic machinery dysfunction are associated with a neurodevelopmental disorder

**Why Not Relevant**: The paper content provided concerns heterozygous de novo variants in HDAC3 and a novel neurodevelopmental syndrome; it does not mention the gene PLRG1 anywhere. Example verbatim sentences from the paper that demonstrate its focus include: "Histone deacetylase 3 (HDAC3) is a crucial epigenetic modulator essential for brain development." and "This study establishes a novel link between heterozygous de novo variants in HDAC3 and a distinct neurodevelopmental syndrome, characterized by intellectual disability, developmental delays, and other variable manifestations such as musculoskeletal anomalies and congenital heart defects." All described genetic findings (D93N, A110T, P201S, L266S, G267S, and R359C), biochemical assays (HDAC activity, western blot), proteomic interactions (CoREST and NCoR), and cellular phenotypes (nuclear localization) explicitly refer to HDAC3. Because the manuscript contains no mention of PLRG1 and provides no data, observations, or discussion about PLRG1 sequence variants, phenotypes, functional assays, or mechanistic connections to disease, it provides neither direct evidence nor mechanistic evidence for the claim that "PLRG1 is likely associated with at least one Mendelian disease." Any inference that PLRG1 might be implicated would be purely speculative and not supported by the text. Limitations: although HDAC3-related mechanisms (epigenetic regulation, multi-protein complex interactions, nuclear localization) are described, there is no provided link between those mechanisms and PLRG1 (for example, shared pathways or physical interactions) in the paper content; therefore mechanistic plausibility for PLRG1 involvement cannot be assessed from this document.


[Read Paper](https://www.semanticscholar.org/paper/524b09f9e39f306b6871f816044577dccb53abd6)


### Combined exome and whole transcriptome sequencing identifies a de novo intronic SRCAP variant causing DEHMBA syndrome with severe sleep disorder.

**Why Not Relevant**: The provided paper content contains no mention of the gene PLRG1, no variant or genotype data, no patient or family segregation evidence, and no functional or mechanistic data linking PLRG1 to any phenotype. The single sentence provided ("This study demonstrated the efficacy of a combined ES-WTS approach in solving undiagnosed cases and speculated that sleep respiratory disorder may be an underdiagnosed complication of DEHMBA syndrome.") refers only to a diagnostic approach (ES-WTS), undiagnosed cases, and a possible complication of DEHMBA syndrome. None of this provides direct evidence that PLRG1 is associated with a Mendelian disease, nor does it provide mechanistic rationale implicating PLRG1. Limitations/caveats: (1) only a single short excerpt was supplied rather than the full paper — it is possible the full article contains discussion or data about PLRG1, but that content was not provided; (2) ES-WTS refers to exome sequencing and whole-transcriptome sequencing approaches that could in principle detect gene-disease associations, but the excerpt gives no results or gene names; and (3) DEHMBA syndrome is named but without gene attribution in the excerpt, so no inference about PLRG1 can be made. Based solely on the supplied content, the paper does not provide direct or mechanistic evidence supporting the claim that PLRG1 is likely associated with at least one Mendelian disease. For a thorough assessment, please provide the full text or sections that mention PLRG1, variant data, segregation, functional assays, or mechanistic discussion.


[Read Paper](https://www.semanticscholar.org/paper/22ade7a766f116133e03729baaf0b3b717e6cd33)


### Hypoparathyroidism in aicardi-goutieres syndrome (AGS) a descriptive case report

**Why Not Relevant**: The provided paper content focuses exclusively on Aicardi-Goutieres syndrome (AGS) and a case involving a SAMHD1 variant. It does not mention the gene PLRG1 anywhere in the text, nor does it present data, analyses, or mechanistic discussion that implicate PLRG1 in any Mendelian disease. Therefore the paper provides no direct evidence for the claim that PLRG1 is likely associated with at least one Mendelian disease. Absence of mention in this paper is not evidence that PLRG1 is not associated with a Mendelian disease—it simply means this paper is not relevant to evaluating that claim. Additionally, the paper is a single-case report focused on SAMHD1 and AGS; even if it cataloged known AGS genes, omission of PLRG1 would be a weak negative inference at best. Because the document contains no information about PLRG1, its relevance to the claim is effectively zero.


[Read Paper](https://www.semanticscholar.org/paper/ad1436aaef408b572435b20864c7a225565c4f40)


### Expanding the Phenotype of Congenital Glucocorticoid Deficiency: An Iranian Patient with Cholestasis due to Pathogenic Variants in the MC2R Gene

**Why Not Relevant**: The supplied paper text does not mention PLRG1 anywhere and therefore provides no direct evidence (genetic, clinical, or segregation data) nor mechanistic evidence linking PLRG1 to any Mendelian disease. Relevant verbatim sentences from the paper that demonstrate its focus on other genes include: "Familial glucocorticoid deficiency is caused by variants in the MC2R and MRAP genes." "We report an Iranian patient with congenital glucocorticoid deficiency and cholestasis due to pathogenic variants in the MC2R gene." "Two variants in the MC2R gene, c.560delT and c.676G > C, were detected and classified as pathogenic and likely pathogenic, respectively." Because PLRG1 is not mentioned, there are no experimental results, segregation analyses, phenotype descriptions, or mechanistic explanations in this paper that support or refute an association between PLRG1 and any Mendelian disorder.

Limitations and caveats: (1) Lack of mention in this paper is absence of evidence only for this specific report — it does not prove PLRG1 is not associated with Mendelian disease. (2) This paper addresses a distinct clinical/genetic question (MC2R/MRAP-related familial glucocorticoid deficiency) and thus is not expected to survey all genes; to evaluate the claim about PLRG1 one must examine other sources (OMIM, ClinVar, PubMed, gnomAD, gene-disease curation resources). Recommended next steps to assess the claim: search disease-gene databases (OMIM, ClinGen), ClinVar for pathogenic PLRG1 variants, PubMed for PLRG1 case reports or functional studies, and population constraint metrics (gnomAD) for PLRG1.


[Read Paper](https://www.semanticscholar.org/paper/421c6aec02ec3b76508151c8c693d2e40f197599)


### Phenotypic and molecular reanalysis of a cohort of patients with monogenic diabetes reveals a case of partial lipodystrophy due to the A8344G mutation in the mitochondrial DNA

**Why Not Relevant**: The provided paper content contains no mention of the gene PLRG1 and provides no direct or mechanistic evidence linking PLRG1 to any Mendelian disease. Instead, the report focuses on a single patient with a phenotype resembling familial partial lipodystrophy for whom no pathogenic variants were found in the classical nuclear genes (LMNA, PPARG) and who was ultimately found to carry a mitochondrial m.A8344G variant at high heteroplasmy. The findings therefore support a mitochondrial etiology for this patient’s disease and do not implicate nuclear genes such as PLRG1. Methodological caveats: the authors state they used a target panel of 48 nuclear genes related to monogenic diabetes but do not list those genes in the provided text, so it is not possible from this excerpt to confirm whether PLRG1 was included in the assay. Even if PLRG1 had been included and found wild-type in this single case, that would not constitute evidence for or against PLRG1 being associated with a Mendelian disease generally. Overall, because PLRG1 is not discussed and no data relating to it are presented, this paper is not relevant to the claim that PLRG1 is likely associated with at least one Mendelian disease.


[Read Paper](https://www.semanticscholar.org/paper/b84ebce0f78f814af19d16356bda255e32515b9d)


## Search Queries Used

- PLRG1 de novo variant developmental disorder exome sequencing trio ClinVar OMIM

- PLRG1 pathogenic variant case report monogenic syndrome familial segregation congenital phenotype

- PLRG1 pleiotropic regulator 1 spliceosome pre mRNA splicing CDC5L complex splicing defect disease mechanism

- PLRG1 gnomAD pLI loss of function intolerance constraint haploinsufficiency knockout mouse embryonic lethal zebrafish developmental phenotype


## Usage Statistics

- Prompt Tokens: 0

- Completion Tokens: 0

- Total Tokens: 0

- Estimated Cost: $0.0367


## Bibliometric Configuration

- Use Bibliometrics: True

- Author Impact Weight: 0.4

- Citation Impact Weight: 0.4

- Venue Impact Weight: 0.2
