## Supplementary material for "MENDELSEEK: An algorithm that predicts Mendelian Genes and elucidates what makes them special": PMCH.txt

# Claim: The gene PMCH is likely associated with at least one Mendelian disease.

**Status**: processed

**Overall Rating**: -1

**Explanation**:

No relevant papers were found that support or refute this claim.


## Relevant Papers


## Other Reviewed Papers


### Diagnostic clarity of exome sequencing following negative comprehensive panel testing in the neonatal intensive care unit

**Why Not Relevant**: The provided paper text does not mention the gene PMCH, does not report any variants in PMCH, and provides no functional or mechanistic data regarding PMCH. The study describes exome and panel sequencing of NICU patients and reports several molecular diagnoses (including mention of COL12A1 as a candidate), but PMCH is not listed among the identified genes nor discussed as a candidate. Therefore there is no direct evidence (no reported PMCH variants segregating with disease, no clinical descriptions tied to PMCH) and no mechanistic evidence (no expression, functional assay, pathway, or animal model data) in this paper that bears on the claim that PMCH is associated with a Mendelian disease. Limitations and context relevant to this negative assessment: (1) absence of evidence in this paper is not evidence of absence overall — this cohort is small (12 trios re-sequenced) and focused on NICU patients with specific phenotypes, so PMCH could be implicated in other cohorts or phenotypes not represented here; (2) the paper emphasizes that some disease genes discovered since the original panel were detectable only by later sequencing approaches, which illustrates that novel Mendelian genes continue to be found, but this general point does not implicate PMCH specifically; (3) to evaluate the claim about PMCH one would need to examine other sources (case reports, ClinVar/DECIPHER/OMIM entries, gene-disease curation resources, functional studies) because this paper provides no supporting or refuting data. Because the paper contains no relevant passages about PMCH, no verbatim excerpts or mechanistic descriptions could be extracted.


[Read Paper](https://www.semanticscholar.org/paper/62096c8bc6e246796ed40c8a0de20b582381c61f)


### Editorial: Hypothalamic Melanocortin Signaling in Cachexia.

**Why Not Relevant**: The provided paper discusses neuropeptides involved in appetite regulation and explicitly names melanin concentrating hormone (MCH) as one of several peptides that "may also contribute" to energy balance. However, it contains no mention of the PMCH gene by name, no genetic or clinical data (no mutations, no affected families, no case reports), and no statements linking MCH/PMCH to any Mendelian disorder. There is no direct human genetic evidence, no description of loss- or gain-of-function variants, no segregation or penetrance data, and no experimental demonstrations that PMCH variation causes a heritable disease. Therefore the paper does not provide direct evidence to support the claim that PMCH is likely associated with at least one Mendelian disease; at best it supplies a general mechanistic rationale (MCH participates in appetite regulation) without any genetic-disease linkage.


[Read Paper](https://www.semanticscholar.org/paper/71f76ed4c9c60174bfd5bf06e630436693a50e21)


### Frequency of Variants in Mendelian Alzheimer's Disease Genes within the Alzheimer's Disease Sequencing Project (ADSP)

**Why Not Relevant**: The paper exclusively studies variants in three established Mendelian Alzheimer's disease genes (PSEN2, PSEN1, APP) using ADSP WGS/WES data. The text provided contains no mention of the gene PMCH, no analysis of PMCH variants, and no mechanistic discussion that would link PMCH to a Mendelian disease. Therefore this paper provides no direct evidence (no data, no reported variants, no association statistics) nor mechanistic evidence (no functional or pathway discussion) relevant to the claim that PMCH is likely associated with at least one Mendelian disease. Important caveat: absence of evidence in this paper is not evidence of absence overall — the lack of PMCH data here simply means this source cannot be used to support or refute the claim about PMCH. To evaluate the claim about PMCH, one would need studies that explicitly analyze PMCH sequence variation, clinical case reports linking PMCH variants to Mendelian phenotypes, curated clinical-database entries (ClinVar/OMIM) implicating PMCH, or mechanistic functional studies showing pathogenic effects of PMCH variants.


[Read Paper](https://www.semanticscholar.org/paper/3760efb72c8ec6adaf553ac773087fd2b16aeafc)


### Comprehensive genetic screening of early-onset dementia patients in an Austrian cohort-suggesting new disease-contributing genes

**Why Not Relevant**: The provided paper content contains only a single sentence about uploading genetic and phenotypic data to the European Genome-phenome Archive to facilitate replication. It does not mention the PMCH gene, any variants in PMCH, any patient phenotypes attributable to PMCH, segregation or linkage data, association statistics, functional assays, or mechanistic pathways involving PMCH. Because none of the necessary elements to evaluate a gene-disease relationship (gene name, variant data, phenotype descriptions, inheritance patterns, statistical association, or functional mechanism) are present, the content cannot be used to support or refute the claim that PMCH is likely associated with at least one Mendelian disease. The only relevant point is that data sharing could enable future confirmation by independent groups, but that is prospective and not evidence of an existing association.


[Read Paper](https://www.semanticscholar.org/paper/75e46adadffa5677302c6111574f9b2f7a427a8c)


### Identifying genomic variant associated with long QT syndrome type 2 in an ecuadorian mestizo individual: a case report

**Why Not Relevant**: The paper does not mention the gene PMCH anywhere in the provided content. The study is a single-case report focused on congenital Long QT Syndrome type 2 and reports a pathogenic variant in KCNH2 and variants of uncertain significance in AKAP9 and TTN. There is no direct evidence (no variants reported in PMCH) nor mechanistic discussion linking PMCH to a Mendelian disease. Absence of PMCH in this paper does not constitute proof that PMCH is not associated with any Mendelian disease, but it does mean this paper provides no supporting or mechanistic evidence for the claim. Additional reasons for low relevance: the work's scope is LQTS genetics (different gene set), sample size is one individual, and methodological details about the breadth of the sequencing or variant reporting are not provided, so no inference about PMCH can be drawn from this study.


[Read Paper](https://www.semanticscholar.org/paper/aa65df486c4f1720d86feb42ddc72ed25ab57431)


### Evaluating High‐Confidence Genes in Conotruncal Cardiac Defects by Gene Burden Analyses

**Why Not Relevant**: This paper focuses on gene burden analyses in nonsyndromic conotruncal cardiac defects using a predefined high‑confidence congenital heart disease gene list and identifies specific CHD‑relevant genes (GATA6, NOTCH1, ANKRD11, DOCK6, NPHP4, STRA6, FLT4, TBX1). The text provided contains no mention of the gene PMCH, no analysis of PMCH variants, and no mechanistic discussion linking PMCH to Mendelian disease. Because PMCH is neither included in the described gene list nor reported among significant findings, the paper provides no direct or mechanistic evidence for the claim that PMCH is likely associated with at least one Mendelian disease. Absence of evidence in this phenotype‑focused study is not evidence that PMCH is not Mendelian, but it does mean this paper is not a relevant source to support the claim.


[Read Paper](https://www.semanticscholar.org/paper/74a8e360545d22e33b5e40474896c9f9c3b6f150)


### Case report: Preimplantation genetic testing for infantile GM1 gangliosidosis

**Why Not Relevant**: The paper exclusively describes GM1 gangliosidosis caused by mutations in the GLB1 gene, reports GLB1 sequence variants in a family, and outlines genetic testing and reproductive counseling related to GLB1. There is no mention of the PMCH gene, no association tests, no case reports, and no mechanistic discussion involving PMCH. Therefore this paper provides no direct evidence that PMCH is associated with any Mendelian disease nor mechanistic data that would support such a claim. Note: absence of evidence in this single paper does not disprove any association for PMCH in the broader literature, but within the content provided here there is nothing relevant to the claim.


[Read Paper](https://www.semanticscholar.org/paper/ce981bc89c565bb4d97baff441baaf8695595448)


### Autosomal Dominant Intellectual Development Disorder-6 (MRD6) Without Seizures Linked to a De Novo Mutation in the grin2b Gene Revealed by Exome Sequencing: A Case Report of a Moroccan Child

**Why Not Relevant**: The provided paper content exclusively discusses the gene grin2b (GRIN2B) and its role in Autosomal dominant intellectual development disorder-6 (MRD6). There is no mention of the gene PMCH, no data on PMCH variants, no mechanistic connection presented between PMCH and GRIN2B, and no inference or comparative analysis that would allow one to conclude anything about PMCH's association with Mendelian disease. Because the paper's observations, genetic findings (including a de novo nonsense variant c.3912C>G (p.Tyr1304Ter)), and mechanistic commentary all concern GRIN2B, they are not applicable evidence for the claim that PMCH is likely associated with at least one Mendelian disease. Absence of evidence in this paper is not evidence of absence for PMCH — but based on this document alone, there is no relevant direct or mechanistic support for the claim about PMCH.


[Read Paper](https://www.semanticscholar.org/paper/67d756e0cc302e5d46fdda9df34403f1f2de9a5a)


## Search Queries Used

- PMCH pathogenic variant ClinVar OMIM PMCH gene Mendelian disease case report pathogenic mutation

- exome sequencing PMCH rare variant gene burden monogenic disease diagnostic cohort rare disease panel PMCH

- PMCH loss of function knockout mouse human phenotype melanin concentrating hormone hypothalamic obesity hyperphagia neuroendocrine disorder

- MCHR1 MCHR2 PMCH signaling pathway mutations Mendelian neurodevelopmental disorder intellectual disability epilepsy congenital syndrome review


## Bibliometric Configuration

- Use Bibliometrics: True

- Author Impact Weight: 0.4

- Citation Impact Weight: 0.4

- Venue Impact Weight: 0.2
