## Supplementary material for "MENDELSEEK: An algorithm that predicts Mendelian Genes and elucidates what makes them special": PMM1.txt

# Claim: The gene PMM1 is likely associated with at least one Mendelian disease.

**Status**: processed

**Overall Rating**: -1

**Explanation**:

No relevant papers were found that support or refute this claim.


## Relevant Papers


## Other Reviewed Papers


### The Burden of Candidate Pathogenic Variants for Kidney and Genitourinary Disorders Emerging From Exome Sequencing

**Why Not Relevant**: The provided paper excerpt does not mention the gene PMM1 by name and does not include the curated gene list or gene-level results that would allow assessment of PMM1's association with any Mendelian disease. The paper focuses on identifying and curating genes associated with Mendelian kidney and genitourinary disorders using OMIM, Orphanet, ClinVar, and HGMD, and describes cohort and filtering methods, but none of the quoted material or surrounding text in the excerpt provides direct evidence (case reports, statistical association, functional data) or even a listing that includes PMM1. Because the claim concerns PMM1 specifically, and the excerpt lacks any explicit reference to PMM1 or to results that can be mapped to PMM1, this document excerpt cannot be used to support or refute the claim. To evaluate the claim, one would need either (a) the curated gene list from the study (to see if PMM1 is included) or (b) other sources (OMIM, ClinVar, HGMD, peer-reviewed reports) that specifically annotate PMM1 with a Mendelian disease association.


[Read Paper](https://www.semanticscholar.org/paper/a75e297208c04992568e607d19aa9b9aaebd09d5)


### Mutations in both SAMD9 and SLC19A2 genes caused complex phenotypes characterized by recurrent infection, dysphagia and profound deafness – a case report for dual diagnosis

**Why Not Relevant**: The provided paper content contains a single clinical statement: "It is reported that it is the first case of a patient with both MIRAGE syndrome and TRMA syndrome, and the patient died from sepsis and multiple organ failure before 1 year old." This text does not mention the gene PMM1, does not present any genetic findings implicating PMM1, and provides no mechanistic data linking PMM1 to disease. MIRAGE syndrome and TRMA syndrome are distinct clinical entities (associated with other genes), so the case report of their co-occurrence does not constitute evidence for or against PMM1 involvement. There are no experimental methods, variant data, biochemical assays, or pathway analyses in the provided content that could be interpreted as direct or mechanistic evidence relating PMM1 to a Mendelian disorder. For these reasons the paper content is not relevant to the claim that PMM1 is likely associated with at least one Mendelian disease.


[Read Paper](https://www.semanticscholar.org/paper/3849dfeacf877fd4bdf34f5f2e51c7e10de2faf9)


### Case Report: The Monogenic Familial Steroid-Resistant Nephrotic Syndrome Caused by a Novel Missense Mutation of NPHS2 Gene A593C in a Chinese Family

**Why Not Relevant**: The paper exclusively reports on NPHS2 (podocin) mutations causing autosomal recessive steroid-resistant nephrotic syndrome in a consanguineous family. The content provided contains no mention of the gene PMM1, no reported PMM1 variants, no mechanistic discussion relating PMM1 to disease, and no experimental or clinical data about PMM1. Therefore the paper provides no direct or mechanistic evidence supporting or refuting the claim that PMM1 is associated with at least one Mendelian disease. Additional limitations: the study is a small case report (two affected siblings) focused on a single gene (NPHS2); negative statements that 'no other SRNS-related gene variants ... were determined' do not specifically exclude PMM1 unless PMM1 is an SRNS gene and explicitly tested/reported, which is not stated. Because PMM1 is not addressed anywhere in the provided content, relevance to the claim is effectively zero.


[Read Paper](https://www.semanticscholar.org/paper/26ba0a4e7e5097751e78e184e30cde938546d66d)


### Comment on: Disease gene identification strategies for exome sequencing by Gilissen et al. 2012

**Why Not Relevant**: The provided paper content is a single general sentence about the complexity of diagnostic challenges and the need to combine approaches; it does not mention the gene PMM1, any Mendelian disease, genetic variants, inheritance patterns, functional/mechanistic studies, case reports, linkage/segregation evidence, population/variant frequency data, or any experimental methods or results. Therefore it contains no direct evidence (e.g., reported pathogenic variants, patient phenotypes, segregation) nor any mechanistic evidence (e.g., molecular function of PMM1, pathway perturbation, animal or cellular models) relevant to the claim that PMM1 is likely associated with at least one Mendelian disease. Because the single sentence contains no gene- or disease-specific information, there are no verbatim excerpts that support or refute the claim to extract. Limitations: the content is too short and too general to permit any inference about PMM1 or Mendelian disease associations; no methodology, data, or citations are provided. Recommended next steps to evaluate the claim: search gene-specific resources (OMIM, ClinVar, HGMD), PubMed for PMM1-related publications, and databases of gene-disease associations; verify that the gene symbol is correct and consider possible confusion with related genes (for example PMM2 is a known disease gene).


[Read Paper](https://www.semanticscholar.org/paper/18c97807528fb94ab3726d65c9a20f45a339b3de)


### Relevance of next generation sequencing (NGS) data re-analysis in the diagnosis of monogenic diseases leading to organ failure

**Why Not Relevant**: The provided paper content contains only a general statement about the value of periodic, systematic re-analysis of negative or inconclusive NGS reports: "It is indicated that periodic and systematic re-analysis of negative or inconclusive NGS data reports can lead to new variant identification or reclassification in a small but significant proportion of cases, with benefits for patients’ management." This sentence does not mention PMM1, any specific gene, any variant, any phenotype, segregation, functional data, case reports, cohort analyses, statistical association with a Mendelian phenotype, or mechanistic evidence linking PMM1 to disease. Therefore the paper content provides no direct evidence (case-level or variant-level) that PMM1 is associated with a Mendelian disease, and no mechanistic evidence (biochemical function, pathogenic mechanism, animal model, or cellular assay) that would support such an association. The statement is purely procedural/epidemiological about re-analysis workflows and cannot be used to support or refute the specific claim about PMM1. To evaluate the claim one would need excerpts reporting PMM1-specific findings such as: identified pathogenic or likely pathogenic variants in PMM1 with linked phenotypes, segregation in families, functional assay results showing loss/gain of function consistent with disease mechanism, or population/association statistics. Because none of those are present in the provided text, relevance to the claim is effectively zero.


[Read Paper](https://www.semanticscholar.org/paper/8f848728768f7aff5d12a97615800c9bdb992e10)


## Search Queries Used

- PMM1 variant patient case report de novo PMM1 mutation monogenic Mendelian disease clinical exome sequencing

- phosphomannomutase 1 PMM1 enzyme activity biochemical assay cellular glycosylation defect PMM1 knockout mouse model loss of function study

- PMM1 PMM2 paralog comparison phosphomannomutase PMM2 congenital disorder of glycosylation CDG functional redundancy differential diagnosis

- PMM1 loss of function variant population frequency gnomAD constraint pLI ClinVar pathogenic exome sequencing rare disease cohort gene burden study systematic review


## Bibliometric Configuration

- Use Bibliometrics: True

- Author Impact Weight: 0.4

- Citation Impact Weight: 0.4

- Venue Impact Weight: 0.2
