## Supplementary material for "MENDELSEEK: An algorithm that predicts Mendelian Genes and elucidates what makes them special": POLR2J.txt

# Claim: The gene POLR2J is likely associated with at least one Mendelian disease.

**Status**: processed

**Overall Rating**: -1

**Explanation**:

No relevant papers were found that support or refute this claim.


## Relevant Papers


## Other Reviewed Papers


### Unravelling undiagnosed rare disease cases by HiFi long-read genome sequencing

**Why Not Relevant**: Relevance is very low because the provided paper content does not mention POLR2J, does not report variants in POLR2J, and provides no mechanistic discussion involving POLR2J. The excerpts show which genes and findings the authors highlighted (e.g., MCF2/FGF13 fusion, PSMA3 deletion) and summarize diagnostic yield, but there is no direct or indirect evidence linking POLR2J to a Mendelian disease in the supplied text. Limitations: this conclusion is based solely on the supplied summary text; the full paper, supplementary tables, or raw variant lists (not provided here) could in principle contain POLR2J observations. To evaluate the claim more fully, one should search the full article, supplements, Solve-RD variant repositories, and disease databases (OMIM, ClinVar, HGMD) for POLR2J entries.


[Read Paper](https://www.semanticscholar.org/paper/c18381966e0231a841f688b515aad06ffabd74d7)


### A pathogenic variant in RAB32 causes autosomal dominant Parkinson’s disease and activates LRRK2 kinase

**Why Not Relevant**: This paper investigates RAB GTPases (specifically reporting RAB32 c.213C>G (Ser71Arg)) in familial Parkinson disease using whole-exome sequencing, segregation analysis, case-control genotyping, and functional assays. The provided text contains no mention of the gene POLR2J, no variants in POLR2J, and no clinical, genetic, or mechanistic data relating POLR2J to any Mendelian disorder. Therefore it provides no direct or mechanistic evidence for or against the claim that POLR2J is likely associated with at least one Mendelian disease. 

Notes on relevance of methods: Although the study design (co-segregation in multi-incident families, case-control association, shared haplotype, and functional activation assays) exemplifies the kinds of evidence that would support a gene-disease Mendelian association, those methods are applied here to RAB32 and not to POLR2J. In the absence of any textual evidence referring to POLR2J, no quotes can be extracted that would bear on the claim.


[Read Paper](https://www.semanticscholar.org/paper/a36d28e828ede7c87c650af5f5117982b9749e7b)


### Novel LSS variants in alopecia and intellectual disability syndrome: New case report and clinical spectrum of LSS‐related rare disease traits

**Why Not Relevant**: The paper's content is entirely focused on the gene LSS and its association with Mendelian disease phenotypes (including description of novel LSS variants and expanded clinical features). There is no mention of POLR2J, no genetic data concerning POLR2J, and no mechanistic discussion linking POLR2J to the reported phenotypes. Therefore this paper provides no direct or mechanistic evidence to support or refute the claim that POLR2J is likely associated with at least one Mendelian disease. In short, absence of any reference to POLR2J in the text means the paper is not relevant to the claim.


[Read Paper](https://www.semanticscholar.org/paper/fe90a5bb55278bc80588741eed5def6cf4f1aa82)


### Reanalysis of Trio Whole-Genome Sequencing Data Doubles the Yield in Autism Spectrum Disorder: De Novo Variants Present in Half

**Why Not Relevant**: The provided paper content does not mention POLR2J anywhere, nor does it provide a gene-level list or examples that would allow identification of POLR2J as implicated. The text is cohort-level and methodological, describing general findings about diagnostic yield, the importance of de novo variants, and that reanalysis implicates genes not previously associated with ASD. While these points describe plausible mechanisms (e.g., de novo, conserved coding variants implicating novel Mendelian genes), there is no direct or specific evidence in the excerpt tying POLR2J to a Mendelian disease. Therefore the content is only very weakly relevant: it establishes that the study design and results could identify novel disease genes, but it does not provide any gene-specific data or explicit association for POLR2J. Additional data (e.g., supplemental tables, gene lists, variant calls, or named examples) would be required to evaluate whether POLR2J is implicated.


[Read Paper](https://www.semanticscholar.org/paper/7c69c67de98e0a5eec703d15c87fdc71b3951364)


### RNA sequencing reveals a complete picture of a homozygous missense variant in a patient with VPS13D movement disorder: a case report and review of the literature

**Why Not Relevant**: The provided paper content discusses a case involving VPS13D—a different gene—and the use of RNA-seq to confirm pathogenic splicing of a VPS13D variant and diagnose a VPS13D movement disorder. There is no mention of POLR2J, no genetic, clinical, or functional data related to POLR2J, and no mechanistic link presented between VPS13D and POLR2J. Therefore the content provides no direct or mechanistic evidence bearing on the claim that POLR2J is likely associated with at least one Mendelian disease. To evaluate that claim one would need different evidence (e.g., reports of pathogenic POLR2J variants segregating with disease, population/constraint data, functional studies implicating POLR2J in disease-relevant biology, or curation in databases such as OMIM); none of those are present in the provided text.


[Read Paper](https://www.semanticscholar.org/paper/0655bdb1e80b122fdb5b84f07ee4efd77c63b603)


### Expanded carrier screening for inherited genetic disease using exome and genome sequencing

**Why Not Relevant**: The provided paper content does not mention the gene POLR2J, report any gene-specific findings, or provide any mechanistic information related to POLR2J. The study description is about assessing the feasibility of exome/genome sequencing for preconception carrier screening in couples and reports aggregate outcomes (e.g., 17 families identified as at-risk, 150 couples evaluated) and general limitations of ES/GS (ethical concerns, technical difficulties). Because no POLR2J-specific data, variant calls, inheritance information, phenotypes, or mechanistic discussion are present in the excerpt, there is no direct or mechanistic evidence in this content to support or refute the claim that POLR2J is likely associated with at least one Mendelian disease. Relevant limitations: absence of evidence in this excerpt is not evidence of absence—full paper or supplementary data might list specific genes/variants—but based solely on the provided text, no assessment of POLR2J can be made. Additional information required to evaluate the claim would include any mention of POLR2J variants detected, segregation or functional data, disease phenotype linkage, inheritance pattern, population frequency, and experimental or computational evidence of pathogenicity.


[Read Paper](https://www.semanticscholar.org/paper/1b853bde9e4b2d5fc6c52be6c5135c93f92d34ae)


### Exome sequence analysis identifies a homozygous, pathogenic, frameshift variant in the MAN2B1 gene underlying clinical variant of α-mannosidosis

**Why Not Relevant**: The manuscript exclusively investigates α-mannosidosis and reports identification, segregation, and structural modeling of a pathogenic MAN2B1 (NM_000528.4: c.2402dupG; p.S802fs*129) homozygous variant in two families. There is no mention of the gene POLR2J anywhere in the provided text, nor any data, analysis, or discussion that connects POLR2J to the phenotypes studied. Thus the paper provides no direct genetic, clinical, or mechanistic evidence for or against an association between POLR2J and any Mendelian disease. Absence of evidence in this paper does not disprove the claim about POLR2J, but it means this paper is not a relevant source to support the claim. Additional literature or data specifically addressing POLR2J would be required to evaluate the claim.


[Read Paper](https://www.semanticscholar.org/paper/455efa329ae9a2ace4b63d745696b455810b8e46)


### Deciphering the Non-Coding Genome in Autism Spectrum Disorders (ASD): A Study of De Novo and Rare Inherited Variants through Targeted Sequencing in Regulatory Regions

**Why Not Relevant**: The provided paper content contains no mention of the gene POLR2J, no POLR2J-specific variants, and no direct discussion of Mendelian disease associations for any specific gene. The study focuses on non-coding regulatory variation in ASD and explicitly excluded cases with WES-identified loss‑of‑function mutations and large CNVs, reducing its ability to detect canonical Mendelian gene–disease relationships. Although the paper describes mechanistic findings (e.g., CTCF dysregulation, PLAC‑Seq annotation) that could in principle implicate genes via regulatory links, none of the quoted sentences or paragraphs provide gene-level evidence implicating POLR2J. Therefore, based on the supplied content, there is insufficient direct or gene‑specific mechanistic evidence to support the claim that POLR2J is likely associated with at least one Mendelian disease.


[Read Paper](https://www.semanticscholar.org/paper/649aeab19738420d26b974bc80eaee323f848ea6)


## Search Queries Used

- POLR2J pathogenic variant Mendelian disease case report clinical exome family segregation POLR2J variant pathogenic

- exome sequencing genome sequencing rare disease cohort de novo variant trio POLR2J gene discovery rare inherited disorder

- POLR2J knockout mouse zebrafish knockdown loss of function RNA polymerase II subunit J transcription phenotype embryonic lethal haploinsufficiency

- RNA polymerase II subunit J POLR2J RPB11 POLR2J2 POLR2J3 Mendelian disease monogenic syndrome ClinVar OMIM variant interpretation pseudogene copy number


## Bibliometric Configuration

- Use Bibliometrics: True

- Author Impact Weight: 0.4

- Citation Impact Weight: 0.4

- Venue Impact Weight: 0.2
