## Supplementary material for "MENDELSEEK: An algorithm that predicts Mendelian Genes and elucidates what makes them special": POLR2K.txt

# Claim: The gene POLR2K is likely associated with at least one Mendelian disease.

**Status**: processed

**Overall Rating**: -1

**Explanation**:

No relevant papers were found that support or refute this claim.


## Relevant Papers


## Other Reviewed Papers


### Clinical and molecular characteristics of a novel rare de novo variant in PPP2CA in a patient with a developmental disorder, autism, and epilepsy

**Why Not Relevant**: The paper focuses on PP2A-related neurodevelopmental disorders and reports genetic, biochemical, and clinical evidence implicating PPP2R5D, PPP2R1A, and PPP2CA. POLR2K is not mentioned anywhere in the provided content. Therefore the paper provides no direct evidence for an association between POLR2K and any Mendelian disease, nor does it provide mechanistic data that would plausibly connect POLR2K to the PP2A-related disorders discussed. The absence of POLR2K from the paper only indicates that this study did not investigate that gene; absence of evidence in this single paper is not evidence of absence overall. To assess the claim about POLR2K, one would need literature or database searches specifically addressing POLR2K (e.g., case reports, cohort sequencing studies, ClinVar/OMIM entries, or functional studies). Given the paper's narrow focus on PP2A subunit genes and the lack of any mention, experimental data, or mechanistic rationale linking POLR2K to Mendelian disease, its relevance to the claim is negligible.


[Read Paper](https://www.semanticscholar.org/paper/69f0588223aa3a22b9c0ec3aa2960ef400867a5c)


### Case report: A novel de novo deletion mutation of DYRK1A is associated with intellectual developmental disorder, autosomal dominant 7

**Why Not Relevant**: The provided paper describes a case of Intellectual developmental disorder 7 (DYRK1A syndrome) caused by a de novo heterozygous deletion in the DYRK1A gene and makes no mention of POLR2K. Relevant sentences in the paper explicitly and repeatedly name DYRK1A as the causal gene (for example: "Using whole-exome sequencing, we identified a 9,424 bp de novo heterozygous deletion ... in DYRK1A, which is responsible for DYRK1A syndrome."), and the conclusions state the findings augment pathogenic variant data for DYRK1A. There is no direct evidence (no observations, variants, case reports, statistical associations, or experimental data) relating POLR2K to any Mendelian disease in the text provided. There is also no mechanistic discussion that connects DYRK1A biology to POLR2K or that would allow a plausible inference that POLR2K is implicated in Mendelian disease. Because the paper focuses solely on DYRK1A, it neither supports nor refutes the claim about POLR2K; it is simply not relevant. Additional limitations: the study is a single-case report concerning DYRK1A, so even for DYRK1A the evidence is limited to one proband and a classified pathogenic deletion; nevertheless, that is unrelated to POLR2K. If you want evidence about POLR2K specifically, I can search the literature or evaluate papers that explicitly mention POLR2K and Mendelian disease associations.


[Read Paper](https://www.semanticscholar.org/paper/98eb2bb2850f75cb23ca3954609b1a07001ab461)


### CAMK2D De Novo Missense Variant in Patient with Syndromic Neurodevelopmental Disorder: A Case Report

**Why Not Relevant**: This paper presents a single-case study identifying a de novo missense variant in CAMK2D (p.Arg275His) and discusses mechanistic and structural evidence implicating CAMK2D/CaMKIIδ in dilated cardiomyopathy with neurodevelopmental delay. The paper does not mention POLR2K anywhere, provides no data (genetic, functional, or clinical) about POLR2K, and does not evaluate POLR2K's association with Mendelian disease. Therefore it neither supports nor refutes the claim that POLR2K is likely associated with a Mendelian disease. The lack of mention of POLR2K means relevance to the claim is effectively nil; absence of evidence in this paper is not evidence against the claim—it's simply unaddressed here.


[Read Paper](https://www.semanticscholar.org/paper/fdc2eee435bd629ce54b9a5b129e1f46af6a6efe)


## Search Queries Used

- POLR2K pathogenic variant Mendelian disease patient case report de novo mutation developmental disorder

- POLR2K RNA polymerase II subunit K RPB10 loss of function haploinsufficiency transcription assembly knockout mouse embryonic lethality developmental phenotype

- POLR2K gnomAD pLI loss of function intolerance constraint rare variant burden exome sequencing de novo enrichment developmental disorder cohort

- RNA polymerase II subunit monogenic disease POLR2K POLR2A POLR2B POLR2C RPB10 Mendelian disorder gene discovery large scale exome sequencing Deciphering Developmental Disorders DDG2P


## Bibliometric Configuration

- Use Bibliometrics: True

- Author Impact Weight: 0.4

- Citation Impact Weight: 0.4

- Venue Impact Weight: 0.2
