## Supplementary material for "MENDELSEEK: An algorithm that predicts Mendelian Genes and elucidates what makes them special": POLR2L.txt

# Claim: The gene POLR2L is likely associated with at least one Mendelian disease.

**Status**: processed

**Overall Rating**: -1

**Explanation**:

No relevant papers were found that support or refute this claim.


## Relevant Papers


## Other Reviewed Papers


### Exome sequencing in obsessive–compulsive disorder reveals a burden of rare damaging coding variants

**Why Not Relevant**: The provided paper content is a single-sentence summary indicating that rare damaging coding variants contribute to OCD risk in a large exome sequencing dataset. It does not mention POLR2L, does not report any gene-level associations, and does not discuss Mendelian diseases or inheritance patterns. Because the claim concerns a specific gene's likely association with a Mendelian disease, evidence would require identification of POLR2L variants segregating with a Mendelian phenotype, statistically significant gene-level burdens consistent with dominant/recessive inheritance, or mechanistic data linking loss/gain of POLR2L function to a Mendelian disorder. None of these elements are present in the supplied content. Therefore the paper, as provided, does not provide relevant evidence for or against the claim. Note: absence of mention in this brief summary is not definitive evidence that the full paper contains no relevant data, but based on the supplied content alone, there is insufficient information to support the claim.


[Read Paper](https://www.semanticscholar.org/paper/34acd840408caf9db7479f8477105f2ee974323d)


### Annotation of uORFs in the OMIM genes allows to reveal pathogenic variants in 5′UTRs

**Why Not Relevant**: The paper describes a systematic annotation of upstream translation initiation sites (uORFs) in OMIM disease-associated genes, reports experimental validation for a set of genes (ETFDH, PAX9, MAST1, HTT, TTN, GLI2, COL2A1, ZIC2), and shows that uORF-disrupting variants in aggregate are enriched among pathogenic variants. However, the provided content (abstract) does not mention POLR2L by name, does not present any gene-specific results or variants for POLR2L, and does not report experimentally validated uORF effects or ClinVar/HGMD hits specifically in POLR2L. Therefore, while the paper provides mechanistic and cohort-level evidence that uORF disruption can cause Mendelian disease and tools to identify such variants, it does not provide direct evidence that POLR2L is associated with a Mendelian disease. The absence of any POLR2L-specific data or statements in the provided text is the reason relevance to the claim "POLR2L is likely associated with at least one Mendelian disease" is very low.


[Read Paper](https://www.semanticscholar.org/paper/5d45c0cdf801505df45fc93a466a2a4101a12871)


### Case report: Novel compound heterozygous IL1RN mutations as the likely cause of a lethal form of deficiency of interleukin-1 receptor antagonist

**Why Not Relevant**: This paper examines a family with a lethal neonatal-onset inflammatory disorder and identifies novel loss-of-function variants in IL1RN as the likely genetic cause. The text repeatedly names IL1RN (interleukin-1 receptor antagonist) and describes biallelic IL1RN LoF variants causing DIRA (deficiency of interleukin-1 receptor antagonist). There is no mention of the gene POLR2L, no data (genetic, transcriptomic, functional, or segregation) relating to POLR2L, and no mechanistic discussion that connects IL1RN findings to POLR2L function. Therefore the paper provides neither direct evidence (no observations of pathogenic POLR2L variants segregating with disease) nor mechanistic evidence (no biological rationale linking POLR2L perturbation to the described phenotype) supporting the claim that POLR2L is likely associated with a Mendelian disease. Key contextual sentences from the paper (illustrating its focus) include mention of IL1RN and DIRA, for example: "We identified two brothers... We conducted DNA and mRNA analyses... which identified two novel heterozygous IL1RN variants..." and "Biallelic LoF variants at the IL1RN gene cause the deficiency of interleukin-1 receptor antagonist (DIRA), a monogenic autoinflammatory disease..." Because POLR2L is not studied or discussed, the paper does not alter the evidence base for POLR2L and is not relevant to the claim.


[Read Paper](https://www.semanticscholar.org/paper/73a143f33c96c5392dc8b513a69039e0e391b92c)


### Variant annotation across homologous proteins (“Paralogue Annotation”) identifies disease-causing missense variants with high precision, and is widely applicable across protein families

**Why Not Relevant**: The paper does not mention POLR2L, nor does it present any data (variant observations, case reports, functional assays, or case-control associations) specific to POLR2L. All content relates to a general computational framework (Paralogue Annotation) and its aggregate performance across many genes (e.g., ClinVar benchmarking, TP53 and PPARG functional comparisons, and case-control tests for certain disorders). While the described method provides a mechanistic rationale for inferring pathogenicity in a gene from pathogenic variants in homologues—which could in principle be applied to POLR2L—no direct evidence, predictions, or results concerning POLR2L are provided. Therefore the paper is only indirectly relevant: it shows a method that might help establish whether POLR2L is disease-associated, but it does not itself provide evidence that POLR2L is associated with any Mendelian disease. To evaluate the claim for POLR2L specifically would require gene-focused data such as: reported pathogenic variants in POLR2L in ClinVar/OMIM/DECIPHER, segregation or de novo variants in affected individuals, functional assays demonstrating a deleterious effect relevant to disease mechanism, or Paralogue Annotation output applied to POLR2L and validated clinically.


[Read Paper](https://www.semanticscholar.org/paper/4f4ff71b890a0c69250f62e4e8170ac545a8968f)


### MSeqDR Quick‐Mitome (QM): Combining Phenotype‐Guided Variant Interpretation and Machine Learning Classifiers to Aid Primary Mitochondrial Disease Genetic Diagnosis

**Why Not Relevant**: This paper describes a web‑based platform (MSeqDR Quick‑Mitome, QM) and a machine‑learning classifier (MDV) for automated variant interpretation in primary mitochondrial disease. It is a methods/resource paper and does not report any gene‑specific Mendelian disease associations, does not mention the gene POLR2L anywhere in the provided content, and presents no patient or variant data linking POLR2L to disease. The paper therefore provides no direct evidence that POLR2L is associated with any Mendelian disease. Mechanistically, the paper focuses on variant annotation, ranking, and mtDNA‑specific analyses (haplogroups, heteroplasmy) and on classifier performance; it does not provide functional/mechanistic data about POLR2L (e.g., gene function, pathogenic variants, segregation, or model organism phenotypes) that would support or refute the claim. Limitations that prevent this paper from informing the claim: (1) absence of any mention of POLR2L or presentation of gene‑level association data; (2) no patient case reports or cohort analyses implicating POLR2L; (3) no experimental mechanistic work on POLR2L; and (4) the gene lists used to train or benchmark the classifier are not provided in the excerpt, so one cannot infer from classifier performance that POLR2L is implicated. The methods/tool described could potentially be used to analyze POLR2L variants in other datasets, but that is not evidence that POLR2L is associated with a Mendelian disease.


[Read Paper](https://www.semanticscholar.org/paper/d12bb508040b75453f2a34e64f096b0cc1f15b01)


### A novel mutation in the ATP7B gene causing hepatolenticular degeneration in a Chinese family: A case report

**Why Not Relevant**: The paper exclusively studies Wilson disease and reports a novel variant in ATP7B; it contains no mention of POLR2L, no genetic data or variant calls for POLR2L, and no mechanistic discussion involving POLR2L. Although whole-exome sequencing was performed (so POLR2L variants could in principle have been observed), the paper's text presents only ATP7B-focused results and interpretations. Therefore it provides no direct evidence (no reported POLR2L variants, cosegregation, case series, or statistical association) and no mechanistic evidence (no functional data, pathway links, or literature discussion) that would support or refute the claim that POLR2L is likely associated with a Mendelian disease. Additionally, this is a single-case report centered on ATP7B with limited generalizability beyond Wilson disease; its scope and reported findings do not address POLR2L. Because of the complete absence of POLR2L-related content, the paper is not relevant to evaluating the claim.


[Read Paper](https://www.semanticscholar.org/paper/9d4b911621bf3cc38aca3619e915780869df23d8)


### Autosomal Dominant Intellectual Development Disorder-6 (MRD6) Without Seizures Linked to a De Novo Mutation in the grin2b Gene Revealed by Exome Sequencing: A Case Report of a Moroccan Child

**Why Not Relevant**: The provided paper text exclusively addresses GRIN2B (grin2b) and its role in Autosomal dominant intellectual development disorder-6 (MRD6), including identification of a specific de novo nonsense variant (c.3912C>G, p.Tyr1304Ter), ACMG pathogenic classification, and biological implications for GRIN2B in brain development. There is no mention of the gene POLR2L, no data on POLR2L variants, and no mechanistic discussion that involves POLR2L. Because the claim concerns whether POLR2L is likely associated with at least one Mendelian disease, this paper provides no direct evidence (variant associations, segregation, case reports) nor mechanistic evidence (functional studies, pathway overlap implicating POLR2L) relating to POLR2L. Therefore the paper is not relevant to the claim.


[Read Paper](https://www.semanticscholar.org/paper/67d756e0cc302e5d46fdda9df34403f1f2de9a5a)


### Case Report: A Disease Phenotype of Rett Syndrome and Neurofibromatosis Resulting from A Bilocus Variant Combination

**Why Not Relevant**: The provided paper content does not mention POLR2L by name, nor does it present any genetic, segregation, variant-level, functional, or mechanistic data specific to POLR2L. The single available sentence only reports a clinical case with multilocus variations and a speculative comment about protein or cellular level relationships among the involved loci. Without explicit identification of POLR2L among the involved loci or any data concerning POLR2L (variants, functional studies, inheritance pattern, population frequency, in silico predictions, or experimental validation), the paper content cannot be used to support or refute the claim that POLR2L is likely associated with at least one Mendelian disease. To assess that claim, one would need excerpts demonstrating POLR2L involvement (e.g., identification of pathogenic variants in POLR2L segregating with disease, functional assays showing deleterious effects of POLR2L variants, or multiple unrelated patients with compatible phenotypes and POLR2L variants).


[Read Paper](https://www.semanticscholar.org/paper/993edabc0eeab1343cda0a7dbad248110f7e87ad)


### DYNA: Disease-Specific Language Model for Variant Pathogenicity

**Why Not Relevant**: The provided paper content describes a computational method (DYNA) for disease-specific fine-tuning of variant effect predictors and its evaluation on broad tasks (cardiovascular diseases and RNA splicing). Nowhere in the supplied text is POLR2L mentioned, nor are there gene-level results, variant-level findings, or any specific Mendelian disease associations reported for individual genes. Thus the content provides no direct evidence that POLR2L is associated with a Mendelian disease. Mechanistically, while the method could in principle be applied to variants in POLR2L to help assess pathogenicity in disease-specific contexts, no such application or result is presented here. Because of the absence of any mention of POLR2L or specific gene-disease associations, the relevance of this paper to the claim is essentially nil.


[Read Paper](https://www.semanticscholar.org/paper/ea7e73128d26759b0af8d083bb377daa366ba57a)


### De novo AHDC1 Deletions Identified by Genome Sequencing in Two Individuals with Xia-Gibbs Syndrome

**Why Not Relevant**: The provided paper content focuses entirely on AHDC1 and its role in Xia-Gibbs syndrome (XGS), including case reports of de novo deletions affecting AHDC1 and the conclusion that AHDC1 haploinsufficiency is a disease mechanism. There is no mention of POLR2L, no data, no variants, and no mechanistic discussion involving POLR2L. Therefore the paper provides no direct evidence (no reported POLR2L variants in patients, no association statistics, no segregation data) nor mechanistic evidence (no functional studies, pathways, or interactions implicating POLR2L) to support or refute the claim that POLR2L is likely associated with at least one Mendelian disease. Absence of evidence in this paper does not disprove the claim generally, but within this paper there is nothing relevant to POLR2L.


[Read Paper](https://www.semanticscholar.org/paper/8ab12fd3ec0c21caabba032cb023668720bf18f8)


### Novel de Novo Nonsense Variants in AGO3 and KHSRP: Insights into Global Developmental Delay and Autism Spectrum Disorders through Whole Genome Analysis

**Why Not Relevant**: The paper's content focuses on a single patient trio whole-genome analysis that identified novel de novo nonsense variants in AGO3 and KHSRP. POLR2L is not mentioned anywhere in the provided text, and there are no data, analyses, or mechanistic discussions concerning POLR2L. Therefore the paper provides no direct genetic, clinical, population, segregation, or functional evidence supporting or refuting an association between POLR2L and any Mendelian disease. Additional limitations that further reduce relevance: the study is a single-case report (limited generalizability), the two genes identified are explicitly stated as not currently associated with OMIM phenotypes (so even the paper's main genes lack established Mendelian links), and the authors call for further studies. Because none of the study's data concern POLR2L, the manuscript is not relevant to the claim that POLR2L is likely associated with at least one Mendelian disease.


[Read Paper](https://www.semanticscholar.org/paper/5aba59259e6535c76aa2fd80054cb774eb3d7452)


### De novo protein-coding gene variants in developmental stuttering

**Why Not Relevant**: The provided paper text does not mention POLR2L (or any obvious synonyms/aliases) and contains no data, analyses, or discussion relating to POLR2L or its role in Mendelian disease. The study focuses on de novo and rare variants in genes implicated in stuttering (examples given: SPTBN1, PRPF8, TRIO, ZBTB7A, FLT3, IREB2, and a set of twelve genes associated with monogenic forms of stuttering). Because POLR2L is not referenced, there is no direct genetic evidence in this paper linking POLR2L to any Mendelian disorder, nor any mechanistic discussion (e.g., functional assays, pathway analyses, expression data) that would support or refute such an association. 

Limitations and caveats: (1) My conclusion is limited to the content you provided. It is possible that POLR2L appears in other parts of the full article (methods supplement, extended tables, or supplementary materials) that were not included here. (2) Absence of evidence in this paper is not evidence of absence overall — other literature or databases (OMIM, ClinVar, HGMD, gnomAD constraint metrics, model organism studies) may contain relevant evidence about POLR2L and Mendelian disease. 

Recommended next steps to assess the claim: check OMIM and ClinVar for reported pathogenic/likely pathogenic POLR2L variants and associated phenotypes; review gnomAD constraint metrics (pLI, LOEUF) and population variant frequencies for POLR2L; search the literature for case reports or functional studies implicating POLR2L in human disease; and examine knockout/knockdown phenotypes in model organisms.


[Read Paper](https://www.semanticscholar.org/paper/8a0180d90cace5fafaf5b3b913882ae5a379c585)


## Search Queries Used

- POLR2L mutation variant pathogenic monogenic Mendelian disease case report family segregation human

- POLR2L ClinVar OMIM pathogenic variant clinical annotation gene disease association review

- RNA polymerase II subunit L POLR2L RPB10 knockout loss of function essential mouse zebrafish model phenotype developmental defect

- exome sequencing genome sequencing trio sequencing developmental disorder rare disease de novo POLR2L gene discovery burden analysis


## Bibliometric Configuration

- Use Bibliometrics: True

- Author Impact Weight: 0.4

- Citation Impact Weight: 0.4

- Venue Impact Weight: 0.2
