## Supplementary material for "MENDELSEEK: An algorithm that predicts Mendelian Genes and elucidates what makes them special": PRKCZ.txt

# Claim: The gene PRKCZ is likely associated with at least one Mendelian disease.

**Status**: processed

**Overall Rating**: -1

**Explanation**:

No relevant papers were found that support or refute this claim.


## Relevant Papers


## Other Reviewed Papers


### The 100,000 Genomes Pilot on Rare Disease Diagnosis in Healthcare − A Preliminary Report

**Why Not Relevant**: The paper excerpt provided is a high-level summary of a pilot genome-sequencing study and reports cohort-level outcomes (sample size, methods used, number of new genes discovered) but contains no gene-specific results or named gene lists. The claim concerns a specific gene, PRKCZ, and whether it is likely associated with at least one Mendelian disease. Because the excerpt does not mention PRKCZ, nor provide gene-level evidence, variant-level data, statistical association results, or mechanistic studies involving PRKCZ, it cannot provide direct support or refutation of the claim. The single sentence noting discovery of "3 new disease genes and 19 novel associations" is indirect and insufficient: without names, effect sizes, phenotype descriptions, statistical significance, or replication, one cannot infer that PRKCZ is among those discoveries. Therefore the content is of minimal relevance (score 0.03) — it indicates that novel Mendelian genes were found by the project (showing the study's capacity) but provides no evidence specific to PRKCZ. If you want a definitive assessment of PRKCZ, please provide the parts of the paper (or dataset/supplementary tables) listing gene-level findings, or I can search the literature/databases (OMIM, ClinVar, HGMD, PubMed) for evidence linking PRKCZ to Mendelian disease.


[Read Paper](https://www.semanticscholar.org/paper/2f56729cea5352bbce9b947f6bd4e560d82d58be)


### The Burden of Candidate Pathogenic Variants for Kidney and Genitourinary Disorders Emerging From Exome Sequencing

**Why Not Relevant**: The provided paper excerpt describes methods for identifying and curating genes associated with Mendelian kidney and genitourinary disorders, variant annotation pipelines, and the cohort used, but it does not mention the gene PRKCZ anywhere in the supplied text, nor does it present the full curated gene list within the excerpt. Therefore there is no direct evidence in this content that PRKCZ is associated with any Mendelian disease. The excerpt only indicates that a curated list of 625 genes exists and is available on the laboratory website; confirmation whether PRKCZ appears on that list or has evidence of disease association would require accessing the external gene list or other databases (OMIM, ClinVar, HGMD) not included in the provided content. Because the excerpt lacks any PRKCZ-specific statements, the relevance to the claim is very low.


[Read Paper](https://www.semanticscholar.org/paper/a75e297208c04992568e607d19aa9b9aaebd09d5)


### Unique Capabilities of Genome Sequencing for Rare Disease Diagnosis

**Why Not Relevant**: The provided paper content is an abstract describing the overall diagnostic yield and advantages of genome sequencing in a cohort of 744 undiagnosed individuals. The abstract summarizes methods, aggregate results (numbers of solved cases and variant types discovered), and conclusions about GS utility, but it does not mention any specific genes by name, including PRKCZ. Consequently, there is no direct evidence (case reports, identified pathogenic variants, segregation data, or statistical association) in the provided text to support or refute the claim that PRKCZ is associated with a Mendelian disease. Mechanistic information relevant to PRKCZ (e.g., functional studies, variant effects on PRKCZ protein function, pathways implicating PRKCZ in disease) is also absent. It is possible that the full article or its supplementary data include a gene-level list of solved cases or reported variants and that PRKCZ could appear there; that information is not present in the excerpt supplied. Therefore, based solely on the supplied content, the paper is not relevant to evaluating the claim about PRKCZ.


[Read Paper](https://www.semanticscholar.org/paper/14c14433e5800efa69d39173f95c49fac150cd46)


### Undiagnosed disease program in South Africa: Results from first 100 exomes

**Why Not Relevant**: The provided paper text does not mention PRKCZ by name, nor does it present a list of the 22 genes or the 25 novel variants referenced. Therefore there is no direct evidence (variant, segregation, phenotype match, or functional/ mechanistic data) linking PRKCZ to a Mendelian disease in this text. The passages describe study design, diagnostic yield, methods (including CNV confirmation), and that novel variants in 22 genes were found, but without gene-level identifiers we cannot ascertain whether PRKCZ is implicated. Mechanistic evidence (e.g., pathogenic variant effect on PRKCZ protein function, pathway disruption, or segregation data specific to PRKCZ) is absent. Given these gaps, the paper content as provided is insufficient to support or refute the claim that PRKCZ is likely associated with at least one Mendelian disease.


[Read Paper](https://www.semanticscholar.org/paper/f59152a7d31bc6a53d506efe5863a0ff91d185c0)


### Rare genetic disorders: Beyond whole‐exome sequencing

**Why Not Relevant**: The provided paper content is a general methodological/overview discussion of NGS technologies (WES and WGS) and their roles in diagnosing rare genetic disorders. It contains no mention of the gene PRKCZ, no case reports, cohort analyses, linkage studies, variant pathogenicity evidence, or functional/mechanistic data specific to PRKCZ. Therefore it provides no direct evidence that PRKCZ is associated with any Mendelian disease. While the paper discusses variant classes and sequencing approaches that are mechanistically relevant to discovering gene–disease associations in general (e.g., WGS detecting structural variants or deep intronic variants), that is generic context and does not bear on PRKCZ specifically. Given the complete absence of PRKCZ-specific data or discussion, the paper's relevance to the claim is effectively zero.


[Read Paper](https://www.semanticscholar.org/paper/bd477d33a409ab5609c86cac0512c51ce998818a)


### Uncovering recessive alleles in rare Mendelian disorders by genome sequencing of 174 individuals with monoallelic pathogenic variants.

**Why Not Relevant**: The provided paper content is a single general statement about using genome sequencing (GS) as a first-tier diagnostic test: "When performing GS as first-tier diagnostic test, including the interpretation of SVs and rare intronic variants in known recessive disease genes, the overall diagnostic yield of rare disease will increase." This sentence does not mention the gene PRKCZ, does not present any data, does not describe any specific Mendelian disease, nor does it provide mechanistic or association evidence linking PRKCZ to any disease. Therefore it provides no direct evidence (e.g., genetic association, segregation, pathogenic variants, case reports) nor mechanistic evidence (e.g., functional studies, pathogenic mechanism) relevant to the claim that PRKCZ is likely associated with at least one Mendelian disease. Limitations: the statement is a general methodological/predictive claim about GS improving diagnostic yield; it contains no gene-level or disease-level information, no experimental methods, no results, and no statistical measures that could be extrapolated to support or refute an association for PRKCZ. To evaluate the claim about PRKCZ, one would need papers reporting pathogenic variants in PRKCZ with segregation in families, recurrent de novo variants, case-control burden, functional assays demonstrating disruption consistent with disease mechanism, or inclusion of PRKCZ in curated disease-gene databases—none of which are present in the supplied content.


[Read Paper](https://www.semanticscholar.org/paper/5ee57a7874147eec316d4452073c0a496149a855)


## Search Queries Used

- PRKCZ protein kinase C zeta PKC zeta PRKCZ mutation PRKCZ variant pathogenic variant Mendelian disease monogenic single gene disorder case report family segregation OMIM ClinVar

- PRKCZ exome sequencing genome sequencing trio sequencing undiagnosed rare disease cohort rare variant burden copy number variant structural variant diagnostic gene panel single gene disorder

- PRKCZ knockout knockdown mouse zebrafish drosophila phenotype loss of function gain of function protein kinase C zeta developmental neurological immunological cardiac metabolic phenotype model organism functional study

- PRKCZ gene disease association review systematic review meta analysis ClinGen gene curation gene disease validity OMIM HGMD ClinVar Mendelian gene list diagnostic gene panel review


## Bibliometric Configuration

- Use Bibliometrics: True

- Author Impact Weight: 0.4

- Citation Impact Weight: 0.4

- Venue Impact Weight: 0.2
