## Supplementary material for "MENDELSEEK: An algorithm that predicts Mendelian Genes and elucidates what makes them special": PRM1.txt

# Claim: The gene PRM1 is likely associated with at least one Mendelian disease.

**Status**: processed

**Overall Rating**: -1

**Explanation**:

No relevant papers were found that support or refute this claim.


## Relevant Papers


## Other Reviewed Papers


### Translational aspects of novel findings in genetics of male infertility—status quo 2021 and diagnosis and epidemiology of winged scapula in breast cancer patients: a systematic review and meta-analysis

**Why Not Relevant**: The provided paper content is a brief introduction to a review on the genetics of male infertility and does not mention PRM1 anywhere. All excerpts are general context about the prevalence, established genetic causes (e.g., CFTR, karyotype, Y microdeletions), the difficulty of mapping monogenic causes for quantitative spermatogenic failure, and recommendations for broad sequencing approaches. None of the sentences contain direct evidence (e.g., variant-level data, case reports, segregation, functional assays) that PRM1 is associated with any Mendelian disease, nor do they describe mechanistic experiments implicating PRM1 in a disease process. Therefore the document provides only very indirect contextual support for the broad plausibility that some genes can underlie Mendelian male infertility, but it provides no specific or direct evidence for or against the claim that PRM1 is likely associated with at least one Mendelian disease.


[Read Paper](https://www.semanticscholar.org/paper/278e9d06a3ebb8141cb79f718ebdedc60c43cf9a)


## Search Queries Used

- PRM1 protamine 1 PRM1 human pathogenic variant ClinVar OMIM Mendelian disease monogenic infertility azoospermia oligozoospermia case report family segregation

- PRM1 protamine 1 knockout mouse spermatogenesis chromatin condensation loss of function infertility male germ cell functional study transgenic null mutant

- PRM1 gene loss of function intolerance pLI gnomAD rare variant frequency damaging variant burden monogenic disease association population genetics constraint

- protamine review male infertility monogenic genes PRM1 PRM2 systematic review meta analysis genetic testing diagnostic gene panel hereditary infertility clinical genetics


## Bibliometric Configuration

- Use Bibliometrics: True

- Author Impact Weight: 0.4

- Citation Impact Weight: 0.4

- Venue Impact Weight: 0.2
