## Supplementary material for "MENDELSEEK: An algorithm that predicts Mendelian Genes and elucidates what makes them special": PRMT5.txt

# Claim: The gene PRMT5 is likely associated with at least one Mendelian disease.

**Status**: processed

**Overall Rating**: -1

**Explanation**:

No relevant papers were found that support or refute this claim.


## Relevant Papers


## Other Reviewed Papers


### Germline activating MTOR mutation arising through gonadal mosaicism in two brothers with megalencephaly and neurodevelopmental abnormalities

**Why Not Relevant**: The provided paper content describes identification of an activating MTOR missense alteration of gonadal mosaic origin as the likely causative mutation in two brothers with neurological and developmental abnormalities. There is no mention of PRMT5, no data on PRMT5 sequence, expression, function, variants, or disease associations, and no mechanistic discussion involving PRMT5. Therefore the paper provides neither direct evidence (no reported PRMT5 variants in affected individuals, no segregation data, no clinical correlation) nor mechanistic evidence (no biochemical, cellular, or pathway data implicating PRMT5) supporting the claim that PRMT5 is associated with a Mendelian disease. Limitations: only a single-sentence summary of the paper was provided rather than the full text; however, that summary explicitly names MTOR as the implicated gene and contains no reference to PRMT5, so absent additional unseen text mentioning PRMT5 this source is not relevant to the claim.


[Read Paper](https://www.semanticscholar.org/paper/15eed4d5efb2abf2f4852f9f04205fe5d8d899ca)


### WWP1 germline variants are associated with normocephalic autism spectrum disorder

**Why Not Relevant**: The provided paper content addresses WWP1 variation and its role as a modifier for autism spectrum disorder (ASD) phenotypes in PTEN hamartoma tumor syndrome (PHTS) cases and references ~600 mainly American PTEN mutation–positive research subjects. It contains no mention of the gene PRMT5, no data on PRMT5 variants, no discussion of Mendelian diseases linked to PRMT5, and no mechanistic pathways involving PRMT5. Because the content is limited to a different gene (WWP1) and context (PTEN/PHTS/ASD), it cannot be used to support or refute the claim about PRMT5 and Mendelian disease association. Additionally, the provided text is a single, somewhat ungrammatical sentence with no methods, results, or detailed context, further preventing any indirect inference about PRMT5.


[Read Paper](https://www.semanticscholar.org/paper/908188af051f3d586880eb95875249c04f23819d)


### De Novo Germline Mutations in SEMA5A Associated With Infantile Spasms

**Why Not Relevant**: The paper's content exclusively discusses findings related to SEMA5A and CLTCL1 in the context of infantile spasm and neurodevelopmental disorders. There is no mention of PRMT5, no genetic, clinical, or mechanistic data involving PRMT5, and no negative statements about PRMT5. Therefore the paper provides no direct evidence for or against the claim that PRMT5 is associated with at least one Mendelian disease. Absence of mention in this single paper does not prove PRMT5 is not disease-associated overall, but this article cannot be used as supporting evidence for the claim.


[Read Paper](https://www.semanticscholar.org/paper/1e06a722f5eedcfe8ec19cb777aaa2048dd3a561)


### Case report: Discovery of a de novo FAM111B pathogenic variant in a patient with an APECED-like clinical phenotype

**Why Not Relevant**: The paper focuses on two specific Mendelian syndromes (APECED and POIKTMP) and the genes AIRE and FAM111B. There is no mention of the gene PRMT5, no reported variants in PRMT5, and no mechanistic discussion involving PRMT5. Although the study performed exome sequencing on a single patient, the absence of any PRMT5 data or discussion means the paper provides neither direct genetic evidence nor mechanistic rationale for or against an association between PRMT5 and a Mendelian disease. Therefore the paper is not relevant to the claim.


[Read Paper](https://www.semanticscholar.org/paper/6ed49c6ffda4dbf2de15474aa07c9326d5636a34)


### Case report and literature review: A de novo pathogenic missense variant in ACTN4 gene caused rapid progression to end-stage renal disease

**Why Not Relevant**: The provided paper content does not mention PRMT5 at all. The study is a case report and genetic analysis implicating ACTN4 in focal segmental glomerulosclerosis (FSGS) in a single patient and discusses ACTN4 variant pathogenicity. There is no experimental data, variant data, literature review, mechanistic discussion, or negative result pertaining to PRMT5. Therefore this paper provides no direct evidence that PRMT5 is associated with any Mendelian disease, nor mechanistic evidence for or against such an association. Absence of mention in this paper does not constitute proof that PRMT5 is not associated with Mendelian disease; it only means this paper is irrelevant to evaluating the PRMT5 claim.


[Read Paper](https://www.semanticscholar.org/paper/177a4961d13db90ae29f92f6efe63bbb6a26f181)


### A novel de novo hemizygous ARHGEF9 mutation associated with severe intellectual disability and epilepsy: a case report

**Why Not Relevant**: The paper exclusively addresses ARHGEF9: its function, a reported de novo loss-of-function variant, and a clinical case of X-linked intellectual disability. There is no mention of PRMT5 (the protein arginine methyltransferase 5 gene), no data on PRMT5 variants, no functional or pathway analyses involving PRMT5, and no comparative genomics or mechanism that would plausibly connect ARHGEF9 findings to PRMT5. Therefore this paper provides no direct evidence that PRMT5 is associated with any Mendelian disease, nor mechanistic evidence that would make such an association plausible. In general, demonstrating that one gene is causative for a Mendelian disorder (here ARHGEF9) does not imply causation for an unrelated gene (PRMT5) unless the paper provides explicit genetic, biochemical, or pathway connections — which it does not. For these reasons the content is not relevant to the claim that PRMT5 is likely associated with at least one Mendelian disease.


[Read Paper](https://www.semanticscholar.org/paper/988bf7595fc677ddeb7de11713ff3345556d6a96)


### Case Report: A novel de novo variant of COL1A1 in fetal genetic osteogenesis imperfecta

**Why Not Relevant**: The paper exclusively studies a novel de novo COL1A1 variant in a fetus with osteogenesis imperfecta. It contains no mention of the PRMT5 gene, no genetic association analyses involving PRMT5, no mechanistic experiments implicating PRMT5, and no discussion that links PRMT5 to the phenotype. Therefore this paper provides no direct or mechanistic evidence that PRMT5 is associated with any Mendelian disease, and no information from this study can be used to support the claim.


[Read Paper](https://www.semanticscholar.org/paper/1caeaf6717a5af526be5f1f0dcef41ef4c377699)


## Search Queries Used

- PRMT5 germline variant pathogenic Mendelian disease case report clinical sequencing de novo

- protein arginine methyltransferase 5 germline mutation neurodevelopmental disorder intellectual disability autism exome sequencing trio

- PRMT5 loss of function haploinsufficiency splicing histone methylation symmetric dimethylarginine mechanism mouse knockout model embryonic lethal

- PRMT5 ClinVar OMIM gnomAD pLI constraint intolerance to variation Mendelian gene curation review


## Bibliometric Configuration

- Use Bibliometrics: True

- Author Impact Weight: 0.4

- Citation Impact Weight: 0.4

- Venue Impact Weight: 0.2
