## Supplementary material for "MENDELSEEK: An algorithm that predicts Mendelian Genes and elucidates what makes them special": PRMT6.txt

# Claim: The gene ZFPM1 is likely associated with at least one Mendelian disease.

**Status**: processed

**Overall Rating**: 0

**Explanation**:

Supporting evidence:
The provided paper documents that a germline GATA1 mutation impairs erythroid and megakaryocytic differentiation and that this mutant shows decreased binding to a set of normal GATA1-interaction proteins, explicitly including the essential co‑factor FOG1 (encoded by ZFPM1). GATA1 is a well‑established master hematopoietic transcription factor, and germline GATA1 mutations are known causes of congenital anemias and thrombocytopenias. The paper therefore establishes a functional connection between a Mendelian disease gene (GATA1) and ZFPM1 at the level of protein–protein interaction and disease mechanism, and it highlights that perturbation of the GATA1–FOG1 interaction is relevant to human congenital anemia.

Caveats or contradictory evidence:
The excerpts do not present any direct evidence that germline or pathogenic variants in ZFPM1 (FOG1) cause a Mendelian disease in humans. The paper documents altered binding of mutant GATA1 to FOG1, but that is different from showing that ZFPM1 itself harbors disease‑causing variants. There is no mention of patients with ZFPM1 variants segregating with a Mendelian phenotype, no functional studies of ZFPM1 variants, and no clinical genetic evidence (e.g., linkage, de novo variants, cohort enrichment, ClinVar/OMIM entries) presented in the provided material that would establish ZFPM1 as a Mendelian disease gene. Functional interaction with a known disease protein increases biological plausibility but is insufficient by itself to claim ZFPM1 is associated with a Mendelian disorder.

Analysis of potential underlying mechanisms and implications:
Biologically, ZFPM1 (FOG1) is an essential cofactor for GATA1 transcriptional activity in erythropoiesis and megakaryopoiesis. Disruption of the GATA1–FOG1 complex could plausibly produce hematopoietic phenotypes similar to those caused by GATA1 mutations. Thus, mechanistically, loss‑of‑function or dominant‑negative variants in ZFPM1 could produce Mendelian hematologic phenotypes. However, plausibility does not equal demonstrated causation. Establishing a gene as a Mendelian disease gene requires human genetic evidence (segregation, recurrent independent pathogenic variants, de novo events, statistical enrichment in affected cohorts) and/or strong functional demonstration that specific ZFPM1 variants produce disease phenotypes consistent with inheritance. The provided paper supplies mechanistic relevance but lacks the human genetic or variant‑level functional data required to support the claim.

Explanation of rating decision:
Given that the only provided evidence links GATA1 disease biology to FOG1 via altered binding but does not present any instances of pathogenic ZFPM1 variants in humans or other direct genetic evidence of ZFPM1 causing Mendelian disease, the available evidence does not support the claim. While the interaction makes ZFPM1 a plausible candidate and warrants further genetic investigation, plausibility alone is not sufficient to assert that ZFPM1 is likely associated with at least one Mendelian disease. Therefore, based on the provided excerpts and the standards for assigning a gene as implicated in Mendelian disease, the appropriate judgment is that there is no direct supporting evidence in the supplied material that ZFPM1 is associated with a Mendelian disorder.


**Final Reasoning**:

The provided paper establishes that FOG1 (ZFPM1) is a key interactor of GATA1 and that perturbation of this interaction is implicated in congenital anemia due to mutant GATA1. This establishes biological plausibility that perturbing ZFPM1 function could have similar phenotypic consequences. However, the claim asserts that ZFPM1 is "likely associated with at least one Mendelian disease." To support that claim requires direct human genetic evidence (e.g., identified pathogenic ZFPM1 variants segregating with disease, de novo variants in affected individuals, multiple independent pathogenic alleles, or curated entries in disease databases). None of that is provided in the excerpts. The distinction between mechanistic plausibility and demonstrated genetic causation is crucial; the paper supports the former but not the latter. Therefore the proper assessment based on the supplied evidence is that there is no evidence presented that ZFPM1 is associated with a Mendelian disease.


## Relevant Papers


### Enhancement of PRMT6 binding to a novel germline GATA1 mutation associated with congenital anemia

**Authors**: Yingsi Lu (H-index: 4), Chengming Zhu (H-index: 1)

**Relevance**: 0.4

**Bibliometric Impact**: 0.18040000000000003


**Excerpts**:

- Mutations in the master hematopoietic transcription factor GATA1 are often associated with functional defects in erythropoiesis and megakaryopoiesis.

- To investigate the specific function and pathogenic mechanism of this mutant, in vitro mutant models of stable re-expression cells were generated.

- The mutation was subsequently validated to cause diminished transcriptional activity of GATA1 and defective differentiation of erythroid and megakaryocytes.

- Using proximity labeling and mass spectrometry, we identified selective alterations in the proximal protein networks of the mutant, revealing decreased binding to a set of normal GATA1-interaction proteins, including the essential co-factor FOG1.

- Taken together, our results provide molecular insights into erythropoiesis in which mutation leads to partial loss of GATA1 function, and the role of PRMT6 and its inhibitor MS023 in congenital anemia, highlighting PRMT6 binding as a negative factor of GATA1 transcriptional activity in aberrant hematopoiesis.


**Explanations**:

- How it relates: This sentence establishes that GATA1 mutations cause Mendelian hematologic phenotypes (congenital anemia / megakaryocyte defects). Relevance to claim: indirect — it defines the disease context in which GATA1 and its interactors operate. Strength/limitations: Directly shows that mutations in a transcription factor cause Mendelian disease, but it does not mention ZFPM1/FOG1 or provide evidence about ZFPM1 itself. This is contextual background rather than evidence for ZFPM1 causality.

- How it relates: Describes the experimental approach (in vitro re-expression models) used to probe the mutant GATA1's function. Relevance to claim: methodological context indicating that subsequent mechanistic findings come from cell models. Strength/limitations: Supports that mechanistic interactions (e.g., with cofactors) were experimentally tested, but also highlights a limitation — findings are from in vitro cellular models which may not fully capture in vivo Mendelian disease causation.

- How it relates: Demonstrates that the GATA1 mutation produces a pathogenic cellular phenotype (reduced transcriptional activity and defective differentiation). Relevance to claim: indirect — shows that perturbation of GATA1 function can produce the disease phenotype, which provides a rationale that partners of GATA1 (like ZFPM1/FOG1) could also be critical. Strength/limitations: Strong evidence that altered GATA1 function causes disease phenotypes in the model system; does not provide direct genetic evidence implicating ZFPM1 variants in Mendelian disease.

- How it relates: Mechanistic evidence that the GATA1 mutant shows decreased binding to several normal interacting proteins, explicitly naming FOG1 as an 'essential co-factor'. Relevance to claim: mechanistic support — if FOG1 (the protein mentioned) corresponds to ZFPM1, loss or disruption of this cofactor interaction is implicated in the disease mechanism, increasing biological plausibility that ZFPM1 dysfunction could cause similar disease. Strength/limitations: This is the most directly relevant excerpt for ZFPM1: it links disrupted GATA1–FOG1 interaction to disease phenotype in the model. However, it is not direct genetic evidence that variants in ZFPM1 cause Mendelian disease; it shows altered binding when GATA1 is mutated. It does not report ZFPM1 mutations, inheritance patterns, segregation, or patient phenotypes attributable to ZFPM1 variants.

- How it relates: Summarizes the study's overall molecular insights tying GATA1 loss-of-function and associated regulatory mechanisms (including recruitment of PRMT6) to congenital anemia. Relevance to claim: indirect/mechanistic — supports the concept that perturbation of the GATA1 regulatory complex yields Mendelian-like disease, implying that critical components of that complex (such as FOG1/ZFPM1) are plausible disease genes. Strength/limitations: Useful synthesis showing pathway-level causality but again lacks direct evidence (no ZFPM1 variants or patient data implicating ZFPM1). Also emphasizes other factors (e.g., PRMT6) whose involvement was experimentally shown, meaning multiple components are at play and causal inference for ZFPM1 alone is not established here.


[Read Paper](https://www.semanticscholar.org/paper/749ea7f74376cc19f06a2d6949ca5a59aa499c43)


## Other Reviewed Papers


### De novo and recessive forms of congenital heart disease have distinct genetic and phenotypic landscapes

**Why Not Relevant**: The provided paper content consists of a single-sentence summary describing broad findings about congenital heart disease genetics and cilia-related genes. It does not mention the gene ZFPM1, give gene-level association results, report variant or inheritance patterns for ZFPM1, nor describe mechanisms involving ZFPM1. Therefore the content cannot be used to support or refute the specific claim that ZFPM1 is likely associated with at least one Mendelian disease. Without additional text from the paper (e.g., gene lists, tables, methods, or variant-level results) or other sources that explicitly discuss ZFPM1, there is no direct or mechanistic evidence in the supplied content relevant to the claim.


[Read Paper](https://www.semanticscholar.org/paper/b541f81fb313e38766a6dfacf2fc7f3f6d96d72e)


### The Burden of Candidate Pathogenic Variants for Kidney and Genitourinary Disorders Emerging From Exome Sequencing

**Why Not Relevant**: The provided paper content describes methods and a curated gene set for investigating Mendelian kidney and genitourinary disorder genes, variant annotation practices, and cohort characteristics, but it does not mention the gene ZFPM1 explicitly nor present gene-level results that include ZFPM1. Because the excerpts do not list the curated genes or report findings for individual genes, there is no direct or specific mechanistic evidence in the provided text that ZFPM1 is associated with any Mendelian disease. It is possible that ZFPM1 appears in the study's full gene list or appendices, or that variant data for ZFPM1 exist elsewhere in the paper, but those sections were not included in the provided content. Therefore, based on the available text, we cannot support or refute the claim that ZFPM1 is likely associated with at least one Mendelian disease. The relevance is correspondingly very low.


[Read Paper](https://www.semanticscholar.org/paper/a75e297208c04992568e607d19aa9b9aaebd09d5)


### From case studies to community knowledge base: MSeqDR provides a platform for the curation and genomic analysis of mitochondrial diseases

**Why Not Relevant**: The provided paper content focuses on mitochondrial disease, the discovery and validation of mitochondrial disease genes (example: PMPCA), and on MSeqDR, a community resource that curates mitochondrial genes and variants. The text does not mention the gene ZFPM1 anywhere, nor does it present data, analyses, functional studies, statistics, or mechanistic discussion specific to ZFPM1. Therefore the paper provides no direct evidence (genetic association, case reports, segregation, functional validation, variant pathogenicity) that ZFPM1 is associated with a Mendelian disease. Indirectly, the paper describes resources and methods (massively parallel sequencing, curated databases like MSeqDR and ClinVar) that could be used to evaluate whether ZFPM1 is disease-associated, but that is not evidence that ZFPM1 is in fact associated with any Mendelian phenotype. Limitations: absence of mention in this paper is not proof that ZFPM1 lacks disease associations elsewhere; it may be outside the mitochondrial focus of this work or included in other resources not discussed here. Because the content contains no direct or mechanistic information about ZFPM1, its relevance to the claim is minimal.


[Read Paper](https://www.semanticscholar.org/paper/6f85c595adaca6963c30fea73bc37266fa78be56)


### Whole-Exome Sequencing in Adults With Chronic Kidney Disease

**Why Not Relevant**: The provided paper content does not mention the gene ZFPM1 (by name) nor present data, results, or discussion specific to ZFPM1. The text describes the authors' methods (WES in adults with CKD), their curated list of 287 nephropathy OMIM genes (referenced as Supplement Table 3), the review of other OMIM genes and ACMG actionable genes, and technical and interpretative procedures and limitations (including that some exons in 29 nephropathy genes were not captured). Because ZFPM1 is not named in the supplied content, there is no direct evidence in this excerpt that ZFPM1 is associated with any Mendelian disease. The methodological descriptions are relevant only insofar as they indicate that (a) the study could have detected pathogenic variants in Mendelian disease genes if those genes were included in the curated lists and adequately captured by the exome kit, and (b) technical blind spots and gene selection criteria (for example, exclusion of poorly captured genes like PKD1) limit the ability to infer absence of association. Therefore, relevance to the specific claim is very low: absence of mention here is not evidence against an association, and the paper as supplied cannot be used to support the claim that ZFPM1 is likely associated with at least one Mendelian disease.


[Read Paper](https://www.semanticscholar.org/paper/a29bd57964f68ac5f8e1abcafe1112ba488f53f5)


### Integrative modeling of transmitted and de novo variants identifies novel risk genes for congenital heart disease

**Why Not Relevant**: The provided paper content (abstract) describes a statistical method and its application to congenital heart disease trio data, reporting that 15 genes were identified as significant. However, the abstract does not list the identified genes or mention ZFPM1 anywhere. Therefore there is no direct evidence in the supplied text that ZFPM1 is associated with any Mendelian disease. The content is only indirectly relevant because the study's approach and results demonstrate that WES analyses can discover disease genes; this increases the plausibility that genes discovered by such methods (if ZFPM1 were among them) could be Mendelian disease genes, but the supplied text itself contains no gene-level data or mechanistic evidence specific to ZFPM1. Without additional sections of the paper (e.g., tables or the results section listing the 15 genes) or supplementary materials naming ZFPM1, the paper excerpt cannot be used to support or refute the claim about ZFPM1.


[Read Paper](https://www.semanticscholar.org/paper/b5cd3995050caef93f234179ebde44733849a57e)


### The simplest explanation does not have to be preferred: co-occurrence of pathogenic variants in cancer-predisposing genes

**Why Not Relevant**: The provided paper content does not mention the gene ZFPM1, Mendelian disease(s), or any gene-specific findings. The single sentence refers broadly to "MINAS," cancer phenotypes, and review/UK 10,000 Genomes Project data in participants with tumor predisposition syndromes or multiple primary tumors, but it contains no results, gene names, variant evidence, statistical data, or mechanistic descriptions that could be linked to ZFPM1. Therefore there is no direct evidence (e.g., pathogenic variants, segregation, case reports, enrichment statistics) and no mechanistic evidence (e.g., functional assays, biological pathways implicating ZFPM1) in the excerpt to support or refute the claim that ZFPM1 is likely associated with at least one Mendelian disease. Additional limitations: the text is a brief descriptive fragment (appears to be part of an abstract or background) and provides no methodological details, sample sizes, variant-level findings, or citations to follow up. Because the required gene-specific information is absent, the excerpt cannot be used to evaluate the claim and relevance is effectively zero.


[Read Paper](https://www.semanticscholar.org/paper/a1372a664bd2c33a0bda638975a8b26d112457fc)


### Case report: Preimplantation genetic testing for infantile GM1 gangliosidosis

**Why Not Relevant**: The paper exclusively addresses GM1 gangliosidosis caused by GLB1 mutations: it identifies specific GLB1 variants, describes sequencing and PGT-M methods targeted at GLB1, and reports embryo and pregnancy outcomes with respect to GLB1. The manuscript contains no mention of ZFPM1, no variants, experimental data, clinical correlations, functional studies, or mechanistic discussion involving ZFPM1. Therefore it provides no direct or mechanistic evidence supporting or refuting the claim that ZFPM1 is associated with a Mendelian disease. Absence of evidence in this paper is not evidence of absence overall, but this paper cannot be used to support the claim.


[Read Paper](https://www.semanticscholar.org/paper/ce981bc89c565bb4d97baff441baaf8695595448)


### Abstract 4143019: Fetuses with Congenital Heart Disease and Placental Malperfusion Have Adverse Outcomes and
 De Novo
 Variants in Key Placental Developmental Pathways

**Why Not Relevant**: The paper does not mention ZFPM1 at all. The study focuses on congenital heart disease fetuses and placental malperfusion and reports enrichment of de novo variants in particular pathways and specific genes (e.g., NOTCH1, NOTCH3, DLL4, SMAD6, VIM, TIE1, COL4A4, FN1, MMP2). Because ZFPM1 is neither listed among implicated genes nor discussed, the paper provides no direct evidence that ZFPM1 is associated with any Mendelian disease. Methodological limits further restrict negative inference: the authors filtered for rare, predicted deleterious exonic or putative splicing SNVs and small indels and emphasized de novo variants in this specific CHD/PMP cohort. Thus, absence of ZFPM1 in the reported results could reflect cohort composition, variant-type/inheritance-mode filtering, or reporting focus, rather than definitive evidence against ZFPM1 being a Mendelian gene. For these reasons the paper is minimally relevant to the claim.


[Read Paper](https://www.semanticscholar.org/paper/cbe436d1db7e626c0f3eef6e106c791f2d5adcef)


### Net fishing for de novo variants in congenital heart disease

**Why Not Relevant**: The supplied paper content does not mention ZFPM1 (also known as FOG1) at all. The text provided only states that the study: identified CHD candidate genes, proved the causal role of a rare missense GLYR1 de novo variant (DNV) in congenital heart disease (CHD), and developed/validated a pipeline to prioritize missense DNVs (and top-scoring DNVs in GT-PPI genes). There are no verbatim sentences or paragraphs in the provided content that refer to ZFPM1, any variant in ZFPM1, segregation or case reports involving ZFPM1, functional assays of ZFPM1, or any other mechanistic data linking ZFPM1 to a Mendelian phenotype.

Because of that absence, this paper excerpt provides neither direct evidence (human genetic cases, segregation, de novo/rare pathogenic variants, case-control enrichment) nor mechanistic evidence (functional assays, animal models, biochemical pathways implicating ZFPM1) relevant to the claim that "ZFPM1 is likely associated with at least one Mendelian disease."

Limitations and further steps:
- The provided content is a brief summary; it is possible the full paper (including main text, figures, and supplementary tables) contains a list of CHD candidate genes or top-ranked DNVs that might include ZFPM1. You should search the full article and its supplements (gene lists, tables S1–Sn) for "ZFPM1" or its synonyms (FOG1) to confirm.
- Absence of mention in this summary is not evidence against ZFPM1 being disease-associated; it is simply lack of evidence from this excerpt.

Recommended next actions to evaluate the claim properly:
1. Search the full paper (main text + supplements) for "ZFPM1"/"FOG1" and examine any reported variants, statistical associations, or functional follow-up. If present, extract verbatim sentences/paragraphs and re-run the relevance analysis.
2. Check curated clinical and genetic databases: OMIM (Online Mendelian Inheritance in Man), ClinVar, HGMD (if available), and PubMed for case reports linking ZFPM1 to Mendelian phenotypes.
3. Assess population constraint/variant frequency in gnomAD (pLI/LOEUF for loss-of-function intolerance, missense z-score) to see whether deleterious variation is depleted.
4. Look for experimental/mechanistic evidence: in vitro functional assays, animal knockout/knock-in phenotypes, or pathway analyses showing how ZFPM1 perturbation causes disease-relevant biology.
5. For strong inference of a Mendelian role, prioritize evidence types in this order: multiple unrelated human patients with rare damaging variants and consistent phenotype (segregation or de novo), replication in independent cohorts, supportive functional data, and responsive animal models.

If you can provide the full paper text or supplementary tables (or confirm whether ZFPM1 appears among the CHD candidate genes), I will extract verbatim quotes and evaluate direct and mechanistic evidence specifically for ZFPM1.


[Read Paper](https://www.semanticscholar.org/paper/367a95ffe2a23a2c63a57cb65ef5799a5344a8f7)


## Search Queries Used

- ZFPM1 FOG1 pathogenic variant causes Mendelian disease monogenic disorder OMIM ClinVar case report

- ZFPM1 FOG1 GATA1 interaction hematopoiesis erythropoiesis megakaryopoiesis congenital anemia thrombocytopenia congenital heart disease mechanism

- exome sequencing ZFPM1 rare variant de novo trio sequencing burden test congenital heart disease developmental disorder rare disease cohort

- ZFPM1 gene disease association review OMIM ClinVar HGMD PanelApp ClinGen gene curation Mendelian gene panel


## Usage Statistics

- Prompt Tokens: 0

- Completion Tokens: 0

- Total Tokens: 0

- Estimated Cost: $0.0487


## Bibliometric Configuration

- Use Bibliometrics: True

- Author Impact Weight: 0.4

- Citation Impact Weight: 0.4

- Venue Impact Weight: 0.2
