## Supplementary material for "MENDELSEEK: An algorithm that predicts Mendelian Genes and elucidates what makes them special": PSMD14.txt

# Claim: The gene PSMD14 is likely associated with at least one Mendelian disease.

**Status**: processed

**Overall Rating**: -1

**Explanation**:

No relevant papers were found that support or refute this claim.


## Relevant Papers


## Other Reviewed Papers


### Human gene and disease associations for clinical‐genomics and precision medicine research

**Why Not Relevant**: The abstract describes a large aggregated gene–SNP–disease database and lists source databases (including OMIM, HGMD, Orphanet) that catalog Mendelian disease associations, but it does not mention PSMD14 or any specific gene-level association. There is no direct evidence in the provided text that PSMD14 is associated with any Mendelian disease, nor any mechanistic data bearing on PSMD14. The paper therefore provides only indirect, speculative relevance (i.e., that the aggregated resource might contain such an association) but no substantive evidence to support or refute the claim that PSMD14 is likely associated with at least one Mendelian disease.


[Read Paper](https://www.semanticscholar.org/paper/bcd1e8a761d2915c1dedbd5a8ef0c314721d2bf6)


### Nearly Half of TP53 Germline Variants Predicted To Be Pathogenic in Patients With Osteosarcoma Are De Novo: A Report From the Children's Oncology Group.

**Why Not Relevant**: The paper focuses on recurrent de novo pathogenic/likely pathogenic variants in a cohort of pediatric osteosarcoma patients. The gene PSMD14 is not mentioned anywhere in the provided content. Therefore the study neither provides direct evidence that PSMD14 is associated with a Mendelian disease nor mechanistic data supporting such an association. At best the paper provides indirect, limited negative evidence that PSMD14 was not a recurrent de novo P/LP gene in this specific osteosarcoma cohort; this is insufficient to support or refute the broader claim that PSMD14 is likely associated with at least one Mendelian disease because: (1) the study is disease-specific (osteosarcoma) and may not capture Mendelian syndromes with other phenotypes; (2) the design prioritized recurrent de novo variants, so inherited pathogenic variants or non-recurrent de novo events in PSMD14 would be missed; (3) PSMD14 may not have been nominated for targeted sequencing if not seen in phase 1; and (4) absence in this dataset is not strong evidence against Mendelian associations observed in other populations or disease contexts.


[Read Paper](https://www.semanticscholar.org/paper/74181d48437b3bd252da941159b8872db80236e1)


### CNVoyant a machine learning framework for accurate and explainable copy number variant classification

**Why Not Relevant**: The provided paper content contains no information about the gene PSMD14, no mention of any Mendelian disease, no gene-disease association analyses, and no experimental or clinical evidence linking PSMD14 to a Mendelian disorder. The only sentence given is: "Compared to alternative approaches, CNVoyant shows marked improvements in precision-recall and ROC AUC metrics for binary pathogenic classifications while going one step further, offering multi-classification of clinical significance and corresponding SHAP explainability plots." This sentence describes the performance and features of a computational tool (CNVoyant) for classifying pathogenicity and providing explainability plots. It does not (a) report variant-level or gene-level association results, (b) present patient phenotypes or segregation data, (c) describe functional/mechanistic experiments involving PSMD14, or (d) cite prior literature linking PSMD14 to Mendelian disease. While such a tool could theoretically be used to analyze variants in PSMD14, that is speculative; the provided content contains no direct or mechanistic evidence relevant to the claim. Therefore the content is not relevant to assessing whether PSMD14 is likely associated with at least one Mendelian disease. Limitations: because only a single summary sentence from the paper was provided, it is possible (but not indicated) that other sections of the full paper could contain relevant analyses; this assessment is limited to the supplied content.


[Read Paper](https://www.semanticscholar.org/paper/56f08cb7ff8b47dcf67c8982001cb74c98b5d841)


### A de novo germline pathogenic BRCA1 variant identified following an osteosarcoma pangenomic molecular analysis.

**Why Not Relevant**: The provided paper content discusses a BRCA1 germline pathogenic variant and makes no mention of PSMD14, its variants, associated phenotypes, functional studies, or mechanistic pathways that could link PSMD14 to Mendelian disease. Because the content neither directly addresses PSMD14 nor provides mechanistic information applicable to PSMD14, it is not relevant to the claim that "PSMD14 is likely associated with at least one Mendelian disease." No supporting or refuting evidence for PSMD14 is present in the supplied text. If additional sections of the paper mention PSMD14, those should be supplied for re-evaluation.


[Read Paper](https://www.semanticscholar.org/paper/d14306369be885b24c7a9ee5c3e695c335ef108a)


## Search Queries Used

- PSMD14 germline mutation de novo inherited pathogenic variant whole exome sequencing whole genome sequencing case report Mendelian disease

- PSMD14 ClinVar OMIM DECIPHER pathogenic variant clinical database germline gene disease association

- PSMD14 POH1 RPN11 deubiquitinase JAMM metalloprotease proteasome 26 S loss of function haploinsufficiency neurodevelopmental disorder intellectual disability microcephaly functional study mouse model

- proteasome subunit gene Mendelian disorder monogenic proteasome associated autoinflammatory neurodevelopmental syndrome PSMD14 PSMD12 PSMB8 review cohort exome sequencing


## Bibliometric Configuration

- Use Bibliometrics: True

- Author Impact Weight: 0.4

- Citation Impact Weight: 0.4

- Venue Impact Weight: 0.2
