## Supplementary material for "MENDELSEEK: An algorithm that predicts Mendelian Genes and elucidates what makes them special": PTGDS.txt

# Claim: The gene PTGDS is likely associated with at least one Mendelian disease.

**Status**: processed

**Overall Rating**: -1

**Explanation**:

No relevant papers were found that support or refute this claim.


## Relevant Papers


## Other Reviewed Papers


### SI-1 Mechanism of seasonal time measurement in vertebrates: Japanese quail and mouse SI-2 Analysis of Gene Function using C. elegans KO strains and a mammalian cell line

**Why Not Relevant**: The paper does not mention PTGDS (the gene encoding lipocalin-type prostaglandin D synthase) anywhere. All prostaglandin-related statements refer specifically to hematopoietic PGD synthase (H-PGDS), which is encoded by a different gene (HPGDS). The reported data are mechanistic/biochemical and pharmacologic (expression changes in patient tissue, inhibitor development, and effects in mdx mice) rather than genetic linkage or mutation data that would indicate a Mendelian disease association. The Mendelian diseases discussed in the paper (e.g., Duchenne muscular dystrophy) are caused by mutations in other genes (dystrophin), and the H-PGDS observations relate to disease progression/response, not causation by germline mutations. Therefore the paper provides no direct or persuasive mechanistic genetic evidence that PTGDS is associated with a Mendelian disease.


[Read Paper](https://www.semanticscholar.org/paper/8fe1b176e9b61d46a702cbf027cb36d2a87f3069)


## Search Queries Used

- PTGDS mutation variant pathogenic Mendelian disease case report family segregation autosomal recessive autosomal dominant prostaglandin D synthase L PGDS beta trace protein

- PTGDS de novo variant exome sequencing cohort intellectual disability developmental delay rare disease gene discovery candidate gene prostaglandin D synthase L PGDS

- PTGDS knockout mouse phenotype loss of function mouse model neurodevelopment brain development neurological phenotype prostaglandin D synthase L PGDS

- PTGDS ClinVar OMIM ClinGen gnomAD constraint pLI loss of function intolerance pathogenic variant clinical significance prostaglandin D synthase beta trace protein


## Bibliometric Configuration

- Use Bibliometrics: True

- Author Impact Weight: 0.4

- Citation Impact Weight: 0.4

- Venue Impact Weight: 0.2
