## Supplementary material for "MENDELSEEK: An algorithm that predicts Mendelian Genes and elucidates what makes them special": PTMA.txt

# Claim: The gene PTMA is likely associated with at least one Mendelian disease.

**Status**: processed

**Overall Rating**: -1

**Explanation**:

No relevant papers were found that support or refute this claim.


## Relevant Papers


## Other Reviewed Papers


### The Burden of Candidate Pathogenic Variants for Kidney and Genitourinary Disorders Emerging From Exome Sequencing

**Why Not Relevant**: The provided paper content does not mention the gene PTMA (prothymosin alpha) anywhere, nor does it present data, analyses, or mechanistic discussion specific to PTMA. The paper focuses on identifying and curating genes associated with Mendelian kidney and genitourinary disorders (a curated list of 625 genes identified via OMIM and Orphanet), describing a cohort of 7,974 ostensibly healthy adults, and evaluating the prevalence and interpretation of candidate pathogenic variants in those kidney/genitourinary-associated genes. Because PTMA is not named in the methods, results, gene lists, or discussion in the supplied text, there is no direct evidence (variant associations, case reports, statistical enrichment) and no mechanistic evidence (biological function, pathogenic mechanism, inheritance patterns) in this paper that bears on the claim that PTMA is likely associated with at least one Mendelian disease. Limitations that lead to this conclusion: (1) no verbatim occurrence of 'PTMA' or its synonyms in the supplied content; (2) the study scope is explicitly limited to Mendelian kidney and genitourinary genes, so genes outside that domain would not be expected to be assessed here; (3) without inclusion of PTMA in the curated gene set or in any reported results, neither positive nor negative evidence about PTMA can be inferred from this paper. For assessment of the claim about PTMA, a targeted search of gene-disease resources (OMIM, ClinVar, HGMD), case reports, or functional studies specific to PTMA would be required.


[Read Paper](https://www.semanticscholar.org/paper/a75e297208c04992568e607d19aa9b9aaebd09d5)


### Precision Medicine in Internal Medicine

**Why Not Relevant**: The provided paper content does not mention PTMA or present any gene-specific data (variant observations, segregation, functional assays, case reports, or statistical association) for PTMA. All excerpts are general background about exome/genome sequencing, variant interpretation, and the proportion of genes currently known to cause Mendelian disorders. While that background is relevant to understanding how a gene such as PTMA could be implicated in a Mendelian disease (i.e., it provides mechanistic plausibility and context about discovery processes), it provides no direct evidence for or against the claim that PTMA is likely associated with at least one Mendelian disease. Therefore the overall relevance to the specific claim is very low.


[Read Paper](https://www.semanticscholar.org/paper/c7199e8a5226f8d9d55777da0e15e91797bb459a)


### Monogenic causation of pediatric nephrolithiasis

**Why Not Relevant**: The provided paper content discusses nephrolithiasis and the role of genetic (including monogenic/Mendelian) causes in pediatric kidney stone disease, but it does not mention the gene PTMA, any data on PTMA (genetic variants, association studies, functional assays, segregation in families), or any mechanistic link between PTMA and Mendelian disease. Therefore the paper provides no direct evidence for or against the claim that PTMA is associated with at least one Mendelian disease. The text only offers general background that some genes cause Mendelian nephrolithiasis, which is indirect and insufficient to support the specific claim about PTMA.


[Read Paper](https://www.semanticscholar.org/paper/ec3abc8d85ef0aeed9d6784c7263cfae9113aafd)


### The simplest explanation does not have to be preferred: co-occurrence of pathogenic variants in cancer-predisposing genes

**Why Not Relevant**: The provided paper content is a single sentence fragment about MINAS (multiple inherited neoplasia alleles syndrome), cancer phenotypes, and review/analysis of UK 10,000 Genomes Project participants with tumor predisposition syndromes or multiple primary tumors. It contains no mention of the gene PTMA, no data on PTMA variants, no association statistics, no clinical cases implicating PTMA in Mendelian disease, and no mechanistic discussion involving PTMA. Therefore it provides neither direct evidence (e.g., reported pathogenic PTMA variants segregating with disease or statistical association) nor mechanistic evidence (e.g., functional impairment of PTMA causing a Mendelian phenotype) relevant to the claim. Additional limitations of the provided content: it is fragmentary (appears to be part of an introduction or methods statement), contains no methods, results, or conclusions specific to any gene, and does not report sample sizes, variant calls, or phenotypic descriptions that could be used to infer relevance to PTMA. To evaluate the claim about PTMA would require different content (e.g., explicit mention of PTMA in cohort analyses, case reports, functional experiments, ClinVar/OMIM entries, or genetic linkage/segregation data).


[Read Paper](https://www.semanticscholar.org/paper/a1372a664bd2c33a0bda638975a8b26d112457fc)


### SAT0200 UNUSUAL SYSTEMIC LUPUS ERYTHEMATOSUS/SJOEGREN'S SYNDROME PHENOTYPE IN A PATIENT WITH A TNFAIP3 GENE MUTATION

**Why Not Relevant**: The provided paper content exclusively discusses mutations in the TNFAIP3 gene (A20) and their relationship to autoinflammatory/autoimmune phenotypes (Behçet-like disease, SLE/SS-like disease). There is no mention of the gene symbol PTMA, its protein product, or any experiments, observations, or mechanistic data involving PTMA. Therefore the paper provides no direct evidence (genetic association, segregation, case reports, or experimental perturbation) that PTMA is associated with any Mendelian disease, nor any mechanistic evidence (protein interaction, pathway involvement, functional studies) linking PTMA to a Mendelian phenotype. The paper does provide evidence that TNFAIP3 variants can cause monogenic disease, but that is about a different gene. Because the genes are distinct (PTMA encodes prothymosin alpha, whereas TNFAIP3 encodes A20) and no cross-reference, interaction, or mention of PTMA appears in the text, this source cannot be used to support the claim that PTMA is likely associated with at least one Mendelian disease. No excerpts are provided because there are no verbatim sentences in the paper content that refer to PTMA or relevant mechanistic links involving PTMA.


[Read Paper](https://www.semanticscholar.org/paper/f1a0805b2b2ba8a9383778f4ae60294f46a13406)


### An Encouraging Successful Result of Allogeneic Hematopoietic Stem Cell Transplantation in a Boy Both with Juvenile Myelomonocytic Leukemia and Osteogenesis Imperfecta : A Case Report

**Why Not Relevant**: The paper does not mention the gene PTMA anywhere in the provided content. The genetic findings and discussion are limited to NF1, PTPN11, COL1A1 (and COL1A2 as known OI genes in background). Because the claim concerns an association between PTMA and at least one Mendelian disease, relevant evidence would require the paper to report PTMA mutations, functional/mechanistic data linking PTMA to a Mendelian phenotype, or population/segregation data implicating PTMA. None of these appear in the text. The extracted sentences were provided to show the paper's actual genetic scope and to document that PTMA is not addressed here. Limitations: absence of any mention of PTMA in this single paper does not disprove an association between PTMA and Mendelian disease in the broader literature. To evaluate the claim one would need papers that explicitly report PTMA variants in affected individuals, segregation analyses, functional studies, or curated gene-disease assertions (e.g., ClinGen, OMIM).


[Read Paper](https://www.semanticscholar.org/paper/457df7a04fddf5dee2432ed70c9125ef795548fe)


### SP0120 Trap deficiency, autoimmunity and other mendelian variants of lupus

**Why Not Relevant**: The paper's content focuses on ACP5 (tartrate resistant acid phosphatase, TRAP), TREX1, early complement components, and the role of type I interferon in Mendelian disorders predisposing to SLE. The gene PTMA is not mentioned anywhere in the provided text. Therefore this paper provides no direct or mechanistic evidence that PTMA is associated with any Mendelian disease. Absence of mention in this paper is not proof that PTMA has no association with Mendelian disease (lack of evidence is not evidence of absence), but from the content provided one cannot support the claim that PTMA is likely associated with a Mendelian disease. To evaluate PTMA specifically would require papers that directly study PTMA mutations, inheritance patterns, functional effects of PTMA variants, or case reports linking PTMA variants to Mendelian phenotypes.


[Read Paper](https://www.semanticscholar.org/paper/4e2ecbbbe528197e2a316d3152bce0ef4fdd5ca1)


## Search Queries Used

- PTMA prothymosin alpha Mendelian disease monogenic single gene disorder case report patient mutation

- PTMA prothymosin alpha ClinVar OMIM HGMD pathogenic variant clinical significance gene disease association

- PTMA prothymosin alpha loss of function knockout mouse phenotype embryonic lethality apoptosis cell proliferation chromatin histone interaction

- PTMA prothymosin alpha exome sequencing undiagnosed disease rare variant burden de novo mutation gnomAD pLI loss of function intolerance


## Bibliometric Configuration

- Use Bibliometrics: True

- Author Impact Weight: 0.4

- Citation Impact Weight: 0.4

- Venue Impact Weight: 0.2
