## Supplementary material for "MENDELSEEK: An algorithm that predicts Mendelian Genes and elucidates what makes them special": PTPRD.txt

# Claim: The gene TRIM21 is likely associated with at least one Mendelian disease.

**Status**: processed

**Overall Rating**: -1

**Explanation**:

No relevant papers were found that support or refute this claim.


## Relevant Papers


## Other Reviewed Papers


### Simultaneous identification and prioritization of variants in familial, de novo, and somatic genetic disorders with VariantMaster

**Why Not Relevant**: The paper describes a computational tool (VariantMaster) for identifying causative variants in familial and sporadic genetic diseases and provides methodological details and proof-of-concept applications. However, the text contains no mention of the gene TRIM21, no gene-specific results, and no direct genetic or clinical evidence linking TRIM21 to any Mendelian disease. Consequently, it does not provide direct evidence to support or refute the claim that TRIM21 is likely associated with at least one Mendelian disease. At best the paper offers indirect mechanistic context showing that such associations can be discovered using the described methods, but the absence of TRIM21-specific data means the paper is not informative about the claim.


[Read Paper](https://www.semanticscholar.org/paper/a56e2dc7369c76b03ad5e9309093beb98a159719)


### Chromatin remodeling dysfunction extends the etiological spectrum of schizophrenia: a case report

**Why Not Relevant**: The supplied paper content (single sentence) does not mention TRIM21, contains no gene-specific data, and provides no experimental, genetic, or clinical evidence linking any particular gene to a Mendelian disease. The sentence only states a general point that schizophrenia can sometimes be underpinned by a Mendelian disease and recommends systematic genetic investigation when there are warning signs (e.g., childhood onset or resistant epilepsy). There are no patient data, no identified variants, no segregation analyses, no functional studies, and no statistical associations presented that could support or refute an association between TRIM21 and any Mendelian disorder. Because the necessary types of evidence (identification of TRIM21 variants in affected individuals, co-segregation in families, demonstration of pathogenicity or plausible mechanism, or population-level association statistics) are entirely absent from the provided content, the paper content is not relevant to the claim that TRIM21 is likely associated with at least one Mendelian disease. Limitations: only a single, general sentence was provided; without additional sections (methods, results, figures, tables, or specific gene mentions) no inference about TRIM21 can be made. To support the claim, the paper would need to present explicit TRIM21-related findings (e.g., pathogenic variants, segregation, functional impact) which are not present in the excerpt.


[Read Paper](https://www.semanticscholar.org/paper/29f6d9d6036caf286740be939452f1be2a4114b2)


### Mind the Gap between the Endothelium and E3 Ubiquitin Ligase: TRIM21 Is a Viable Therapeutic Target in Sepsis-induced Endothelial Dysfunction

**Why Not Relevant**: The paper contains mechanistic and experimental evidence that TRIM21 is an important E3 ubiquitin ligase involved in immune and endothelial responses, and documents clinical relevance via autoantibody recognition in autoimmune diseases. However, it does not present any human germline genetic evidence (e.g., segregating variants, linkage, de novo mutations, case reports of Mendelian inheritance, or loss-of-function intolerance data) that would support TRIM21 being associated with a Mendelian disease. All clinical associations described are autoimmune or acquired (autoantibodies, altered expression), and the experimental evidence comes from overexpression/deficiency models in cells or mice. Therefore, while the mechanistic data make it biologically plausible that TRIM21 variants could produce disease, this paper provides no direct genetic evidence for a Mendelian disorder attributable to TRIM21; consequently its relevance to the specific claim is very low.


[Read Paper](https://www.semanticscholar.org/paper/6b028a0ee60c764d5f5e5b78efc9eb9411c8f834)


### Undiagnosed disease program in South Africa: Results from first 100 exomes

**Why Not Relevant**: The provided paper excerpt does not mention TRIM21 by name, nor does it provide a list of the 22 genes with novel variants, variant annotations, or any mechanistic data referring to TRIM21. All available statements are high-level descriptions of cohort, methods, diagnostic yield, and general gene-discovery potential. Therefore, there is no direct evidence in the excerpt that TRIM21 is associated with any Mendelian disease. Although the study reports novel variants in multiple genes ("Twenty‐five novel variants in 22 genes are reported here"), because the gene identities are not included in the provided content it is impossible to determine from this excerpt whether TRIM21 is among them. In addition, no mechanistic experiments, functional validations, segregation data, or pathogenicity assessments specific to TRIM21 are presented. For these reasons the relevance to the claim that "TRIM21 is likely associated with at least one Mendelian disease" is negligible based on the supplied text. If the full paper's gene list or supplementary tables were available and included TRIM21 with evidence (segregation, pathogenic classification, functional data), relevance would increase; but that information is not present in the provided content.


[Read Paper](https://www.semanticscholar.org/paper/f59152a7d31bc6a53d506efe5863a0ff91d185c0)


### Regulatory T cell Itch reins in Th2 inflammation

**Why Not Relevant**: The paper excerpt exclusively discusses the E3 ubiquitin ligase Itch and its role in regulatory T cell biology and Th2 inflammation. It does not mention TRIM21, any other gene names, or any clinical/Mendelian disease associations. Because the claim concerns TRIM21 being likely associated with at least one Mendelian disease, and the provided content contains no reference to TRIM21 or evidence linking it to Mendelian disease, the paper content is not relevant to the claim.


[Read Paper](https://www.semanticscholar.org/paper/44403a906341fe3161e88d71ea36c2650602d746)


### Case report: Novel compound heterozygous IL1RN mutations as the likely cause of a lethal form of deficiency of interleukin-1 receptor antagonist

**Why Not Relevant**: The paper's content focuses exclusively on IL1RN and its involvement in a monogenic autoinflammatory disease (DIRA) in a single family. All reported genetic, transcriptomic, mechanistic, and clinical evidence pertains to IL1RN variants (specific intronic and intragenic deletion variants), decreased IL1RN mRNA, truncated IL1RN transcripts, autosomal recessive segregation, and the established link between biallelic IL1RN loss-of-function and DIRA. The text contains no mention of TRIM21, no experimental data on TRIM21, and no mechanistic discussion that would connect TRIM21 to the disease phenotype. Therefore this paper provides no direct or indirect evidence that TRIM21 is associated with any Mendelian disease, and is not relevant to the claim about TRIM21.


[Read Paper](https://www.semanticscholar.org/paper/73a143f33c96c5392dc8b513a69039e0e391b92c)


### HOXA11 is another monogenic cause of congenital anomalies of the kidney and urinary tract—Reply

**Why Not Relevant**: The provided paper content contains one sentence about the gene HXA11 and congenital anomalies of the kidney and urinary tract (CAKUT): "HXA11 is an example of a gene that does not yet have sufficient evidence supporting it as a monogenic cause of CAKUT in humans and it is believed that any generalizations regarding its direct causal role must await the description and characterization of mutations in additional patients." This sentence (1) concerns HXA11, not TRIM21; (2) addresses the current insufficiency of evidence for HXA11 as a monogenic cause of CAKUT; and (3) does not mention TRIM21, any Mendelian disease linked to TRIM21, nor any mechanistic data relevant to TRIM21. Therefore the content provides no direct evidence (no reported pathogenic TRIM21 variants, no segregation data, no clinical associations) and no mechanistic evidence (no functional studies, pathways, or biological rationale) bearing on the claim that "TRIM21 is likely associated with at least one Mendelian disease." Additional limitations: only a single sentence was provided, no figures, tables, results, or methods relevant to TRIM21 are present, and no page numbering or bibliographic context was supplied. To assess the claim about TRIM21 would require independent evidence such as reports of TRIM21 variants segregating with disease, statistically significant burden analyses, functional assays showing pathogenic mechanisms, or curated clinical-genetic databases—none of which are present in the supplied content.


[Read Paper](https://www.semanticscholar.org/paper/1fdff7edb4655416163ed78c76a6471bc026c45d)


### Experience of the first adult-focussed undiagnosed disease program in Australia (AHA-UDP): solving rare and puzzling genetic disorders is ageless

**Why Not Relevant**: The provided paper content is a single-sentence summary: "The utility of a UDP approach applying genome sequencing approaches in diagnosing adults with rare diseases who have had uninformative conventional genetic analysis is demonstrated, informing clinical management, recurrence risk, and recommendations for relatives." This sentence does not mention the gene TRIM21, does not report any gene-specific findings, and contains no data, associations, or mechanistic descriptions linking TRIM21 to any Mendelian disease. Therefore the excerpted content provides no direct evidence (no reported variants, case reports, cohort data, statistical associations, or clinical diagnoses involving TRIM21) and no mechanistic evidence (no functional studies, protein interactions, pathogenic mechanism, or genotype–phenotype description) relevant to the claim that TRIM21 is likely associated with at least one Mendelian disease. Limitations relevant to this assessment: (1) Only a single summary sentence was provided rather than the full paper text; it is possible that the full article contains gene-level results including TRIM21, but that information is not present in the content you supplied. (2) Because no gene-specific data appear in the provided content, I cannot evaluate evidence quality, methodology, statistical or practical significance, or experimental conditions with respect to TRIM21. If you can provide the full paper text or the specific sections (results, tables, supplementary data) that mention TRIM21, I can re-evaluate and extract relevant verbatim quotes and assess direct and mechanistic evidence.


[Read Paper](https://www.semanticscholar.org/paper/24eda8dfb581396db3979ef16ccb8e01634d084b)


### A retrospective analysis of phosphatase catalytic subunit gene variants in patients with rare disorders identifies novel candidate neurodevelopmental disease genes

**Why Not Relevant**: The provided paper text focuses specifically on a systematic survey of de novo variants in 189 genes encoding phosphatase catalytic subunits and reports candidate phosphatase disease genes (e.g., PTPRD, PTPRG, PTPRT, PTPRU, PTPRZ1, MTMR3, GAK, TPTE2, PTPN18). There is no mention of TRIM21, nor of E3 ubiquitin ligases, TRIM family genes, or any data linking TRIM21 to Mendelian disease in the supplied content. Therefore the paper provides no direct evidence for the claim that TRIM21 is likely associated with at least one Mendelian disease. Mechanistic evidence is also absent: the study concentrates on phosphorylation/dephosphorylation and phosphatase genes, which are a different functional class from TRIM21 (an E3 ubiquitin ligase), so there are no mechanistic pathways, experimental results, or phenotypic matches in the text that could be used to support or refute the TRIM21–Mendelian disease association. Limitations: If the full paper (beyond the supplied excerpt) contains sections not provided here, those could potentially mention TRIM21; my conclusion is limited to the supplied content only. Given the absence of any mention or data related to TRIM21 in the provided text, relevance to the claim is effectively zero.


[Read Paper](https://www.semanticscholar.org/paper/d0dfa58ee328453fa43927d0500f97d3a8f2dd39)


### Autosomal Dominant Intellectual Development Disorder-6 (MRD6) Without Seizures Linked to a De Novo Mutation in the grin2b Gene Revealed by Exome Sequencing: A Case Report of a Moroccan Child

**Why Not Relevant**: The paper exclusively discusses GRIN2B (referred to here as "grin2b") and a de novo nonsense variant (p.Tyr1304Ter) causing MRD6. There is no mention, data, analysis, or mechanism involving the gene TRIM21 anywhere in the provided text. Therefore the paper provides no direct evidence that TRIM21 is associated with any Mendelian disease, nor mechanistic evidence that would make such an association plausible. Absence of TRIM21 in this paper is not disproof of any TRIM21-disease link — it simply provides no information relevant to the claim. To evaluate the claim about TRIM21 one would need papers, clinical reports, or database entries (OMIM, ClinVar, HGMD, PubMed) that specifically address TRIM21 variants and their disease associations; those sources were not present in this document.


[Read Paper](https://www.semanticscholar.org/paper/67d756e0cc302e5d46fdda9df34403f1f2de9a5a)


### Exome copy number variant detection, analysis and classification in a large cohort of families with undiagnosed rare genetic disease

**Why Not Relevant**: The provided paper content describes methods and overall results of CNV detection from exome sequencing across a large, heterogeneous cohort and reports cohort-level yields and CNV characteristics. Nowhere in the supplied text is TRIM21 mentioned, nor are gene-level results or lists of affected genes provided. Because the claim concerns whether TRIM21 is likely associated with at least one Mendelian disease, evidence must be gene-specific (e.g., identification of pathogenic or likely pathogenic variants affecting TRIM21, segregation data, functional/mechanistic evidence implicating TRIM21 in disease). The supplied content only offers high-level, study-wide findings that CNV calling can identify causal variants across many genes. While this establishes general mechanistic plausibility (that CNVs can disrupt genes and cause Mendelian disease), it provides no direct or specific data about TRIM21. Therefore the relevance of this paper excerpt to the claim about TRIM21 is minimal.


[Read Paper](https://www.semanticscholar.org/paper/2b8162a945544b579cfc6f8ac2e7a50d528bfcde)


### The E3 Ubiquitin Ligase HUWE1 Is a Central Regulator of Podocyte Homeostasis

**Why Not Relevant**: The provided paper exclusively investigates Ercc1 (a nucleotide excision repair gene) using a podocyte-specific knockout mouse and examines pathways such as mTORC1 activation, inflammation, and the effects of calorie restriction and rapamycin. The text contains no mention of TRIM21 or any data (genetic association, mutation screening, functional assays, patient phenotypes, linkage/segregation, or case reports) relating TRIM21 to disease. Therefore the paper contains no direct evidence that TRIM21 is associated with any Mendelian disease, nor mechanistic data that would plausibly implicate TRIM21. Because the gene studied (ERCC1) and the disease mechanisms described are unrelated to TRIM21, no inference about TRIM21's Mendelian-disease associations can be drawn from this work.


[Read Paper](https://www.semanticscholar.org/paper/f538fe92dbad1862e814d66502d128f562629056)


### Shining a light on age-related adrenal cancer progression

**Why Not Relevant**: The provided content focuses on a mouse model of age-dependent adrenal cancer, myeloid immune cells, androgen effects, and senescent adrenal cortex cells with a SASP (Znrf3-cKO). There is no mention of the gene TRIM21, no discussion of human genetic variation, no report of Mendelian inheritance patterns, and no genetic association data. Thus the excerpt contains no direct or mechanistic evidence relevant to the claim that TRIM21 is likely associated with at least one Mendelian disease. Note that the assessment is limited to the supplied excerpt; to fully exclude relevance one would need to search the full paper text and genetic-disease databases (e.g., OMIM, ClinVar, HGMD) for TRIM21 associations.


[Read Paper](https://www.semanticscholar.org/paper/15946ef12c517778be92c49702bb563da756baaf)


### Signalling of E3 ubiquitin-protein ligases in the regulation of priming and tolerance of T cells

**Why Not Relevant**: This paper does not provide direct evidence supporting or refuting the claim that TRIM21 is likely associated with at least one Mendelian disease. The manuscript focuses on the roles of E3 ubiquitin-protein ligases in T cell anergy and priming, and specifically measures and discusses Cbl-b, Itch and Grail. The word "TRIM21" (or synonyms like Ro52) does not appear in the provided content, and there are no genetic, inheritance, mutation, patient, or clinical-disease association data presented that would link any particular E3 ligase (including TRIM21) to a Mendelian disorder. While several excerpts describe general mechanisms by which E3 ubiquitin ligases can alter immune cell function — a mechanistic class-level link that is shared by TRIM21 — such mechanistic similarity alone is insufficient to infer that TRIM21 is associated with a Mendelian disease. Additional required evidence would include segregation of TRIM21 variants with disease in families, identification of pathogenic (rare, deleterious) variants in TRIM21 in affected individuals, functional demonstration that those variants disrupt TRIM21 in a way consistent with disease mechanism, or epidemiological enrichment of damaging TRIM21 variants in patients versus controls. None of those data or analyses are present here. Other limitations reducing relevance: the experiments are immunological and expression-based (mouse transgenic and in vitro/in vivo T cell assays), not human genetic studies; findings are specific to other E3 ligases; and expression changes do not establish causality or Mendelian inheritance. For these reasons the paper provides at most very indirect mechanistic context about E3 ligases generally, but no specific or substantive evidence about TRIM21 and Mendelian disease.


[Read Paper](https://www.semanticscholar.org/paper/7c8892fe045390d07ff7f35a16364c13b542327f)


## Search Queries Used

- TRIM21 mutation Mendelian monogenic disease case report family segregation de novo biallelic heterozygous

- TRIM21 variant exome sequencing rare disease cohort undiagnosed disease program gene discovery candidate gene

- TRIM21 knockout mouse phenotype immune deficiency autoimmunity E3 ubiquitin ligase functional study inflammation

- TRIM21 Tripartite motif containing 21 Ro52 SSA1 ClinVar OMIM gnomAD pLI loss of function intolerance review gene disease association


## Bibliometric Configuration

- Use Bibliometrics: True

- Author Impact Weight: 0.4

- Citation Impact Weight: 0.4

- Venue Impact Weight: 0.2
