## Supplementary material for "MENDELSEEK: An algorithm that predicts Mendelian Genes and elucidates what makes them special": PTPRS.txt

# Claim: The gene PTPRS is likely associated with at least one Mendelian disease.

**Status**: processed

**Overall Rating**: -1

**Explanation**:

No relevant papers were found that support or refute this claim.


## Relevant Papers


## Other Reviewed Papers


### Expanding the phenotype of a recurrent de novo variant in PACS1 causing intellectual disability

**Why Not Relevant**: This paper discusses a recurrent de novo missense variant in PACS1 (c.[607C>T], p.[Arg203Trp]) causing an autosomal-dominant intellectual disability syndrome (MRD17). The text makes no mention of the gene PTPRS (no occurrences of the symbol PTPRS, its synonyms, or related protein tyrosine phosphatase family members). Therefore the paper contains no direct evidence that PTPRS is associated with any Mendelian disease, nor does it provide mechanistic data linking PACS1 findings to PTPRS. Because the gene discussed (PACS1) is distinct from PTPRS, findings about PACS1 cannot be used to support the claim about PTPRS. Note that absence of mention of PTPRS in this paper is not evidence that PTPRS is not associated with disease overall — it only means this paper does not provide relevant supporting or refuting evidence for the claim.


[Read Paper](https://www.semanticscholar.org/paper/e460d89d850693b4fb1e3df12702be0dfb8d7f5b)


### A novel NONO nonsense variant in a fetus with renal abnormalities

**Why Not Relevant**: The paper content exclusively describes a clinical case and genetic finding involving the NONO gene (a novel de novo hemizygous loss‑of‑function variant in NONO) and discusses the prenatal phenotype associated with NONO-related intellectual developmental disorder (OMIM#300967). There is no mention of PTPRS, no data (genetic, clinical, functional, or mechanistic) concerning PTPRS, and no discussion that could be used to infer an association between PTPRS and any Mendelian disease. Therefore the paper provides no direct or mechanistic evidence relevant to the claim that PTPRS is likely associated with at least one Mendelian disease.


[Read Paper](https://www.semanticscholar.org/paper/c188bc6b84c091f181b4802ec9293ca711cf7c62)


### articleDownregulation of genes with a function in axon outgrowth and synapse formation in motor neurones of the VEGFδ / δ mouse model of amyotrophic lateral sclerosis

**Why Not Relevant**: The provided paper content contains no mention of the gene PTPRS (Protein Tyrosine Phosphatase, Receptor Type S) and presents no data (direct or indirect) that links PTPRS to any Mendelian disease. The study focuses on VEGF (vascular endothelial growth factor), VEGFδ/δ mice, transcriptional changes in motor neurons, and pathways related to axonogenesis, synapse formation, and cholesterol synthesis. Although the authors performed microarray analysis (reporting numbers of differentially expressed genes at various disease stages), the excerpt does not include gene-level results or supplemental gene lists that could be searched for PTPRS. Therefore, based on the provided content, there is neither direct evidence (e.g., mutation–disease co-segregation, pathogenic variants, case reports) nor mechanistic evidence (e.g., functional experiments implicating PTPRS in disease processes) to support the claim that PTPRS is likely associated with at least one Mendelian disease.

Limitations relevant to this assessment:
- Absence of PTPRS in the text: No sentence or paragraph in the provided excerpt names PTPRS or describes its expression, mutation status, or functional role.
- Microarray possibility: The study performed microarray profiling and identified hundreds of differentially expressed genes, so it is possible (but not shown here) that PTPRS could appear in full gene lists or supplemental material; such material would need to be inspected to confirm presence or absence.
- Model and sample size: The experiments are in a mouse model (VEGF HRE deletion) with small sample groups (n=3 per genotype per time point in the excerpt), which limits generalizability to human Mendelian disease even if PTPRS were implicated in mouse transcriptional changes.

Suggested next steps to evaluate the claim properly:
1) Inspect the full paper (especially supplemental tables) for complete lists of differentially expressed genes to see whether PTPRS is reported.
2) Search databases of Mendelian disease associations (OMIM, ClinVar, HGMD, GeneReviews) and the literature for pathogenic variants or published pedigrees linking PTPRS to inherited disease.
3) If PTPRS appears in expression or functional data, assess whether there are human genetic data (segregation, de novo mutations, recurrent pathogenic variants) that meet criteria for Mendelian disease association.

Because the provided excerpt contains no mention of PTPRS and no relevant experimental or mechanistic information about that gene, the paper as given cannot be used to support the claim that PTPRS is likely associated with at least one Mendelian disease.


[Read Paper](https://www.semanticscholar.org/paper/2be2725a2198ee00f71fbcbb92587e717eb18434)


### De novo HDAC3 variants leading to epigenetic machinery dysfunction are associated with a neurodevelopmental disorder

**Why Not Relevant**: The provided paper text focuses entirely on HDAC3 and heterozygous de novo variants in HDAC3 as the cause of a novel neurodevelopmental syndrome. There is no mention of the gene PTPRS, no data concerning variants in PTPRS, and no comparative or cross-gene analyses that would link HDAC3 findings to PTPRS. Therefore the paper provides no direct evidence that PTPRS is associated with any Mendelian disease. Mechanistic findings described (reduced deacetylation activity, disrupted interactions with CoREST and NCoR, impaired nuclear localization) are specific to HDAC3 protein function and cannot, without additional data, be generalized or used as mechanistic support for pathogenicity of PTPRS. In short, because the manuscript does not discuss PTPRS at all, it is not relevant to the claim that PTPRS is likely associated with at least one Mendelian disease. If you would like, I can search for or analyze papers or databases that directly address PTPRS and Mendelian disease associations (e.g., OMIM, ClinVar, PubMed).


[Read Paper](https://www.semanticscholar.org/paper/524b09f9e39f306b6871f816044577dccb53abd6)


## Search Queries Used

- PTPRS PTP sigma receptor type S mutation Mendelian disease case report family segregation clinical exome sequencing autosomal dominant autosomal recessive

- PTPRS de novo variants exome sequencing trio intellectual disability developmental delay rare variant burden ClinVar OMIM DECIPHER

- PTPRS receptor type tyrosine phosphatase S neuronal development axon guidance synapse formation signaling pathway loss of function pathogenic mechanism neurodevelopmental disease

- PTPRS loss of function constraint gnomAD pLI gene constraint ClinGen gene curation Mendelian disease likelihood population variation copy number variation


## Bibliometric Configuration

- Use Bibliometrics: True

- Author Impact Weight: 0.4

- Citation Impact Weight: 0.4

- Venue Impact Weight: 0.2
