## Supplementary material for "MENDELSEEK: An algorithm that predicts Mendelian Genes and elucidates what makes them special": RICTOR.txt

# Claim: The gene RICTOR is likely associated with at least one Mendelian disease.

**Status**: processed

**Overall Rating**: -1

**Explanation**:

No relevant papers were found that support or refute this claim.


## Relevant Papers


## Other Reviewed Papers


### Impact of genetic insights into mendelian disease on cardiovascular clinical practice.

**Why Not Relevant**: The provided paper excerpt contains no mention of the gene RICTOR nor any synonyms or descriptions that clearly refer to RICTOR (for example, mTOR complex 2 component RICTOR). The text is a general review-style introduction about Mendelian cardiovascular diseases (inheritance patterns, clinical implications, and an introduction to hypertrophic cardiomyopathy) and contains no gene-specific data, case reports, variant evidence, segregation analyses, functional assays, or mechanistic discussion that connect RICTOR to any Mendelian disorder. Because there are no sentences or paragraphs that reference RICTOR, I cannot extract direct or mechanistic evidence linking RICTOR to a Mendelian disease from this content. Important limitations: (1) only an excerpt was provided — if the full paper contains sections not included here that discuss RICTOR, those were not available for review; (2) as supplied, the text is high-level clinical context and does not contain experimental, genetic, or molecular data that could support or refute the claim. To assess the claim properly would require primary evidence such as case reports, linkage/segregation studies, population/variant-disease association data, ClinVar/OMIM entries, or functional studies showing how RICTOR variants cause a Mendelian phenotype. None of those are present in the provided content.


[Read Paper](https://www.semanticscholar.org/paper/9540e5f30cfc13896da8382f41afc9854af87e6c)


### TTC7A Variants Previously Described to Cause Enteropathy Are Observed on a Single Haplotype and Appear Non-pathogenic in Pediatric Inflammatory Bowel Disease Patients

**Why Not Relevant**: The provided paper content discusses compound heterozygous variants in the Tetratricopeptide Repeat Domain 7A gene (TTC7A) and evaluates TTC7A variation in relation to polygenic inflammatory bowel disease. It does not mention the gene RICTOR, nor does it present any data, mechanistic discussion, or analyses that link TTC7A findings to RICTOR or to Mendelian disease status of RICTOR. Therefore there is no direct evidence (e.g., genetic association, segregation, pathogenic variants) nor mechanistic evidence (e.g., pathway interaction, functional assay) in this content that bears on the claim that RICTOR is likely associated with at least one Mendelian disease. Limitations: this judgment is based solely on the excerpt provided; if the full paper contains additional sections not included here that discuss RICTOR, those were not available for review. To evaluate the claim about RICTOR, one should examine studies that explicitly report RICTOR variants segregating with Mendelian phenotypes, functional follow-up on RICTOR perturbation, or curated clinical resources (e.g., OMIM, ClinVar, gene-disease curation databases).


[Read Paper](https://www.semanticscholar.org/paper/3818a966b5d6e8b0eda65fdef043ac0727fa429c)


### Clinical and Genetic Characteristics of Patients with Unexplained Intellectual Disability/Developmental Delay without Epilepsy

**Why Not Relevant**: This paper does not mention the gene RICTOR anywhere in the provided text. The study reports findings from a focused cohort of 38 patients with unexplained ID/DD and/or ASD and lists specific genes in which novel or pathogenic variants were identified (TUBA1A, TMEM63A, YY1AP1, ABCA13, USP9X, ANKRD17, GRIA4), but RICTOR is not among them. Therefore the paper provides no direct evidence that RICTOR is associated with a Mendelian disease, and it offers no mechanistic discussion implicating RICTOR in disease pathways. The absence of RICTOR in this small, phenotype-selected cohort is weak negative evidence at best: it neither supports nor robustly refutes the claim because (1) the sample size is small (38 individuals) and phenotypically restricted to neurodevelopmental presentations, (2) the study design and gene list reflect only the variants observed in these patients and not a comprehensive assessment of all genes or known Mendelian conditions, and (3) RICTOR-associated disease (if present) might present with different phenotypes or be too rare to appear in this cohort. For these reasons the content is not relevant for establishing that RICTOR is likely associated with at least one Mendelian disease.


[Read Paper](https://www.semanticscholar.org/paper/e5289119c3a34a33fd81c6fcc009413eb467af22)


### Clinical and genetic findings in a cohort of Chinese patients with autosomal recessive spinocerebellar ataxia

**Why Not Relevant**: The paper does not mention RICTOR anywhere in the provided content; it focuses on genes identified as causes of autosomal recessive spinocerebellar ataxia (SCAR) in a specific cohort. The study reports 19 variants in 11 genes (SETX, NPC1, CYP27A1, SYNE1, GBA2, SACS, CLN6, SCARB2, NEU1, ATP7B, NHLRC1) and does not present any data, variants, functional studies, or mechanistic discussion involving RICTOR. Therefore it provides no direct evidence supporting RICTOR as being associated with a Mendelian disease. The only weakly relevant point is that a large targeted panel (>4000 OMIM genes) was used, which implies the study could have detected known Mendelian genes if present in this phenotype and cohort; however, absence of RICTOR among reported findings is not informative because (1) the cohort is small and restricted to young-onset SCAR (so genes causing other Mendelian phenotypes would not be expected), (2) it is not stated whether RICTOR was included or well-covered in the panel, and (3) failure to observe RICTOR variants in this specific sample does not exclude RICTOR being a Mendelian gene in other contexts. For these reasons the paper is not relevant evidence for or against the claim that RICTOR is likely associated with at least one Mendelian disease.


[Read Paper](https://www.semanticscholar.org/paper/b1299c62cd43bcaef8f79fbcc9dbe246358f3323)


### A de novo PAK1 likely pathogenic variant and a de novo terminal 1q microdeletion in a Chinese girl with global developmental delay, severe intellectual disability, and seizures

**Why Not Relevant**: The paper content provided discusses a novel PAK1 variant associated with a terminal 1q microdeletion and states that the patient's main clinical features are believed to be attributable to the PAK1 variant. There is no mention of the gene RICTOR, no data about RICTOR variants, no mechanistic discussion involving RICTOR, nor any analyses that would support or refute RICTOR's association with Mendelian disease. Because the paper's focus is PAK1 and a 1q microdeletion in a single patient, it cannot provide evidence for the claim that RICTOR is likely associated with at least one Mendelian disease. Additional reasons for irrelevance: (1) no experimental, clinical, genetic, or statistical data about RICTOR are presented; (2) the single-case focus and attribution to PAK1 limit any negative inference about other genes in the region; (3) no functional or segregation evidence relevant to RICTOR is given. To evaluate the claim about RICTOR, papers reporting pathogenic RICTOR variants, segregation with disease, functional disruption of RICTOR, or curated gene-disease associations (e.g., OMIM, ClinGen) would be required; this paper does not provide such information.


[Read Paper](https://www.semanticscholar.org/paper/8ac30d308f85987b8cf4644520ee11940136ebaa)


### A de novo variant in ZBTB18 gene caused autosomal dominant non-syndromic intellectual disability 22 syndrome: A case report and literature review

**Why Not Relevant**: The paper describes a case of autosomal dominant non-syndromic intellectual disability 22 caused by pathogenic variation in ZBTB18 and discusses clinical features, diagnosis, and the mechanistic consequence (premature termination) of the ZBTB18 variant. There is no mention of the gene RICTOR, no data (genetic, clinical, or mechanistic) involving RICTOR, and nothing that links RICTOR to any Mendelian phenotype. Therefore this paper provides no direct or mechanistic evidence for the claim that RICTOR is likely associated with at least one Mendelian disease.


[Read Paper](https://www.semanticscholar.org/paper/09bd993a331155206894ef8f2ad7afb75e164664)


### A novel AUTS2 variant in a patient with global developmental delay and intellectual disability

**Why Not Relevant**: This paper is entirely focused on AUTS2 haploinsufficiency and reports a novel de novo loss-of-function variant in the AUTS2 gene. It does not mention RICTOR, mTOR, mTORC2, TORC2, or any other genes or pathways plausibly linking RICTOR to disease. Example sentences from the paper that demonstrate its scope (verbatim) include: "AUTS2 haploinsufficiency causes a neurodevelopmental disorder known as AUTS2, which is characterized by global developmental delay, intellectual disability, autistic features, congenital brain anomalies, and other malformations." and "A novel, heterozygous variant (c.1606C>T) in AUTS2 gene was identified." Because there is no discussion, data, or mechanistic analysis involving RICTOR in this text, the paper provides no direct evidence (no case reports, no genetic association, no segregation, no population or case-control data) and no mechanistic evidence (no pathway analyses, no functional experiments linking RICTOR perturbation to the reported phenotype) supporting the claim that RICTOR is associated with a Mendelian disease. Additional points limiting any indirect inference: (1) the study is a single-case report of an AUTS2 variant, so its conclusions apply only to AUTS2; (2) the authors themselves state further functional experiments are required to validate the AUTS2 variant's impact, underscoring the narrow scope and provisional nature of the findings; (3) without any mention of RICTOR or related biology, one cannot extrapolate an association to RICTOR from this paper. Therefore relevance to the claim that "RICTOR is likely associated with at least one Mendelian disease" is effectively zero.


[Read Paper](https://www.semanticscholar.org/paper/f5e84e7bb3b42239a53cc7e5c457342cb38c6277)


### Molecular Genetics of Autosomal Recessive Mental Disorders in Consanguineous Pakistani Families

**Why Not Relevant**: The paper does not mention the gene RICTOR anywhere in the provided content and provides no direct or mechanistic data about RICTOR. The study describes mapped loci, candidate genes (OR52I2, OR52E6, PGAP2, DRD4, HOMER2, PSMB11, PANK2, etc.), and conclusions about genetic heterogeneity in several Pakistani consanguineous families, but none of these findings implicate or evaluate RICTOR. Therefore the paper offers no evidence for or against the claim that RICTOR is associated with a Mendelian disease. Absence of evidence in this study is not evidence of absence: the negative relevance here reflects that the study simply did not examine or report on RICTOR, was limited to a small set of families and loci, and focused on autosomal recessive mental retardation and Hallervorden–Spatz Syndrome in a specific population. These limitations mean one cannot generalize from this paper to the broader question of RICTOR's involvement in Mendelian disease.


[Read Paper](https://www.semanticscholar.org/paper/58d88a43df50573eeda5e35768169882a023e3dc)


## Search Queries Used

- RICTOR de novo germline variant neurodevelopmental disorder intellectual disability developmental delay exome sequencing case report

- RICTOR gene Mendelian disease monogenic inherited autosomal dominant autosomal recessive single gene disorder human genetics

- RICTOR mTORC2 loss of function gain of function AKT phosphorylation PI3K signaling cytoskeleton neuronal migration animal model mouse zebrafish functional study

- RICTOR ClinVar OMIM ClinGen pathogenic variant germline copy number deletion duplication case series cohort study somatic cancer amplification


## Bibliometric Configuration

- Use Bibliometrics: True

- Author Impact Weight: 0.4

- Citation Impact Weight: 0.4

- Venue Impact Weight: 0.2
