## Supplementary material for "MENDELSEEK: An algorithm that predicts Mendelian Genes and elucidates what makes them special": RO60.txt

# Claim: The gene RO60 is likely associated with at least one Mendelian disease.

**Status**: processed

**Overall Rating**: -1

**Explanation**:

No relevant papers were found that support or refute this claim.


## Relevant Papers


## Other Reviewed Papers


### The Burden of Candidate Pathogenic Variants for Kidney and Genitourinary Disorders Emerging From Exome Sequencing

**Why Not Relevant**: The provided paper content focuses on identification, curation, and variant-prevalence analysis of genes associated specifically with Mendelian kidney and genitourinary disorders. Nowhere in the supplied text is the gene symbol "RO60" mentioned, nor are non–kidney/genitourinary gene–disease associations discussed. The excerpts establish that the authors compiled a specialist list of 625 genes from OMIM and Orphanet and analyzed variant prevalence in those genes in a cohort of 7,974 ostensibly healthy adults. Because the paper's stated scope is limited to kidney and genitourinary Mendelian disorders and the provided content does not include the full curated gene list or any mention of RO60, there is no direct evidence in this document that RO60 is associated with any Mendelian disease. This absence of mention in the provided text is not definitive proof that RO60 has no Mendelian association (absence of evidence is not evidence of absence): RO60 could be absent from the study's gene set because it is not a kidney/genitourinary gene, or it could be present in the full gene list that was not included in the excerpt. To evaluate the claim about RO60, one would need either (a) the paper's complete gene list to check whether RO60 was included and, if so, any variant findings; or (b) other literature/databases (OMIM, ClinVar, HGMD, peer-reviewed association studies) that specifically report RO60 disease associations or mechanistic evidence. Given the lack of any mention or data about RO60 in the provided content, the paper as supplied provides essentially no relevant evidence for or against the claim.


[Read Paper](https://www.semanticscholar.org/paper/a75e297208c04992568e607d19aa9b9aaebd09d5)


### From case studies to community knowledge base: MSeqDR provides a platform for the curation and genomic analysis of mitochondrial diseases

**Why Not Relevant**: The provided paper content is an overview of the MSeqDR resource and examples of how novel mitochondrial disease genes (e.g., PMPCA) were discovered and curated. The text does not mention the gene RO60, nor does it present any patient data, variant annotations, functional experiments, or literature citations linking RO60 to a Mendelian disease. All evidence in the excerpt is general or procedural (about database scope, curation, and data sharing) and therefore cannot be used as direct or mechanistic evidence that RO60 is associated with at least one Mendelian disease. It is possible that RO60 could be included in MSeqDR-LSDB or ClinVar, but this paper does not state that; establishing RO60 as a Mendelian disease gene would require gene-specific case reports, functional studies, or curated pathogenic variants, none of which are present in the supplied text.


[Read Paper](https://www.semanticscholar.org/paper/6f85c595adaca6963c30fea73bc37266fa78be56)


### TTC7A Variants Previously Described to Cause Enteropathy Are Observed on a Single Haplotype and Appear Non-pathogenic in Pediatric Inflammatory Bowel Disease Patients

**Why Not Relevant**: The provided paper content discusses TTC7A (Tetratricopeptide Repeat Domain 7A) and its variants in the context of polygenic inflammatory bowel disease (IBD). It makes no mention of the gene RO60, no mention of any diseases associated with RO60, and no mechanistic pathways connecting TTC7A to RO60. Therefore the paper content supplied is not relevant to evaluating whether RO60 is associated with any Mendelian disease. Additionally, the content provided is minimal (a single sentence summary) and contains no data, methods, results, or discussion about RO60, so it cannot provide direct or mechanistic evidence for the claim.


[Read Paper](https://www.semanticscholar.org/paper/3818a966b5d6e8b0eda65fdef043ac0727fa429c)


### The simplest explanation does not have to be preferred: co-occurrence of pathogenic variants in cancer-predisposing genes

**Why Not Relevant**: The provided paper content does not mention the gene RO60, Mendelian disease, or any specific gene-level genetic evidence. The only sentence supplied is: "The frequency and cancer phenotypes associated with MINAS by reviewing the recent literature and by checking the UK 10,000 Genomes Project data from participants with a phenotype of tumor predisposition syndromes or multiple primary tumors, and if they are additive reflecting each CSG independently or synergistic resulting in more severe phenotypes are reported." This text (1) refers to MINAS and cancer phenotypes and to cancer susceptibility genes (CSGs) in general, (2) describes a literature review and an analysis of UK 10,000 Genomes Project participants with tumor predisposition phenotypes, and (3) addresses whether effects are additive or synergistic. It contains no direct evidence (no variants, no segregation data, no case reports, no statistical associations) concerning RO60 and provides no mechanistic discussion linking RO60 to Mendelian disease. Because the excerpt contains no mention of RO60 or of Mendelian inheritance patterns, it cannot be used to support or refute the claim that "The gene RO60 is likely associated with at least one Mendelian disease." Note that absence of mention in this brief excerpt is not evidence that RO60 is or is not associated with a Mendelian disease in the wider literature; it only means this paper content, as supplied, provides no relevant evidence. To evaluate the claim one would need papers reporting (for example) pathogenic RO60 variants with segregation in families, case series describing rare RO60-related syndromes, burden/association statistics implicating RO60 in a Mendelian phenotype, or mechanistic studies showing that RO60 loss/gain causes a Mendelian disease phenotype.


[Read Paper](https://www.semanticscholar.org/paper/a1372a664bd2c33a0bda638975a8b26d112457fc)


### Case report: Preimplantation genetic testing for infantile GM1 gangliosidosis

**Why Not Relevant**: The provided paper content focuses exclusively on GM1 gangliosidosis and pathogenic variants in the GLB1 gene. There is no mention of the gene RO60, no association analyses involving RO60, and no mechanistic discussion that links RO60 to GM1 gangliosidosis or any other Mendelian phenotype. Although the text states that "next-generation sequencing" was used, it does not specify whether this was a targeted GLB1 panel, whole-exome sequencing, or whole-genome sequencing; the concrete variant reporting and the subsequent PGT-M workflow indicate the study's aims and results center on GLB1. Because the excerpted content contains no data, observations, or discussion about RO60, it neither provides supporting evidence (no positive association, variants, or mechanistic link) nor definitive refutation (the paper does not state that RO60 was examined and found negative). Therefore, based on the provided content, the paper is not relevant to the claim that RO60 is likely associated with at least one Mendelian disease.


[Read Paper](https://www.semanticscholar.org/paper/ce981bc89c565bb4d97baff441baaf8695595448)


## Search Queries Used

- RO60 TROVE2 pathogenic variant case report Mendelian disease monogenic mutation ClinVar OMIM

- RO60 TROVE2 gnomAD loss of function constraint pLI LOEUF haploinsufficiency intolerance rare variants population frequency

- RO60 TROVE2 Ro60 Y RNA RNA binding protein knockout mouse phenotype loss of function zebrafish neurodevelopmental defects RNA quality control

- Ro60 SSA Ro60 TROVE2 Sjogren syndrome systemic lupus erythematosus autoantigen genome wide association study GWAS review systematic review meta analysis monogenic versus complex disease


## Bibliometric Configuration

- Use Bibliometrics: True

- Author Impact Weight: 0.4

- Citation Impact Weight: 0.4

- Venue Impact Weight: 0.2
