## Supplementary material for "MENDELSEEK: An algorithm that predicts Mendelian Genes and elucidates what makes them special": ROCK1.txt

# Claim: The gene ROCK1 is likely associated with at least one Mendelian disease.

**Status**: processed

**Overall Rating**: -1

**Explanation**:

No relevant papers were found that support or refute this claim.


## Relevant Papers


## Other Reviewed Papers


### Diagnostic yield of patients with undiagnosed intellectual disability, global developmental delay and multiples congenital anomalies using karyotype, microarray analysis, whole exome sequencing from Central Brazil

**Why Not Relevant**: The provided paper content does not mention ROCK1 by name, does not list the specific genes included in the 1,252-gene panel, and does not report gene-level results or mechanistic data that would implicate ROCK1 in a Mendelian disease. While the study used an exome-based ID gene panel (which might, in principle, include ROCK1), absence of any explicit reference to ROCK1 or to variants in ROCK1 means there is neither direct evidence (observed pathogenic variants in ROCK1 causing disease in these patients) nor mechanistic evidence (functional studies, pathway analyses, or literature citations) supporting the claim. Additional limitations that prevent inferring support for the claim from this paper: the gene panel composition is unspecified, the sample size for the exome analysis is small (19 trios), and no supplementary tables or gene lists are provided in the excerpt. To evaluate the claim one would need either (a) explicit reporting of ROCK1 being included and observed to harbor pathogenic variants segregating with disease in this cohort, or (b) referenced mechanistic/functional data linking ROCK1 to a Mendelian phenotype. Neither is present in the provided content. Recommended next steps: inspect the full paper (including supplementary tables) for gene lists or variant tables, search other literature or disease databases (OMIM, ClinVar, HGMD) for ROCK1 associations, or contact the authors for gene-level results.


[Read Paper](https://www.semanticscholar.org/paper/23767daf16f48d9a91172aa0fc624f07b869a9c8)


### Case Report: Novel Homozygous Likely Pathogenic SCN1A Variant With Autosomal Recessive Inheritance and Review of the Literature

**Why Not Relevant**: This paper is not relevant to the claim that ROCK1 is likely associated with at least one Mendelian disease. All content provided concerns variants in the SCN1A gene and their association with neurodevelopmental disorders and epilepsies (e.g., Dravet syndrome, GEFS+). The paper describes clinical and genetic findings for homozygous or compound heterozygous SCN1A variants, genotype–phenotype correlations for SCN1A domain locations, and inheritance patterns specific to SCN1A. It contains no mention of the ROCK1 gene, no data on ROCK1 sequence variants, no case reports or segregation analyses involving ROCK1, and no mechanistic discussion linking ROCK1 to Mendelian disease phenotypes. Therefore it provides neither direct evidence (patient/variant/segregation data) nor mechanistic evidence (functional studies, pathway links) supporting or refuting an association between ROCK1 and any Mendelian disorder. To evaluate the claim about ROCK1 would require papers reporting pathogenic ROCK1 variants with clinical phenotypes, family segregation, functional follow-up, or credible population/association evidence — none of which are present in this manuscript.


[Read Paper](https://www.semanticscholar.org/paper/06c4ac652678bddf09a9cc2d1428cbc3a11edf4c)


### A single center experience with publicly funded clinical exome sequencing for neurodevelopmental disorders or multiple congenital anomalies

**Why Not Relevant**: The provided paper content consists of a single general statement about the utility of clinical exome sequencing for individuals with GDD/ID and/or MCA. It contains no mention of the ROCK1 gene, no variant-level data, no case reports, no statistical associations, no segregation or de novo occurrence information, and no mechanistic or functional data concerning ROCK1. Because establishing that a gene is 'likely associated with at least one Mendelian disease' requires gene-specific evidence (case observations with pathogenic variants and appropriate clinical correlation, segregation, recurrence, population/variant frequency analysis, or functional/mechanistic validation), none of which are present here, this paper content is not relevant to the claim. To evaluate the claim one would need excerpts such as identified pathogenic or likely pathogenic ROCK1 variants with associated phenotypes, experimental evidence of variant impact on protein/gene function, or robust genetic/clinical association data; such information is absent in the supplied text.


[Read Paper](https://www.semanticscholar.org/paper/8ec4c0c03c573ad4019a6ee63b9197179a32b173)


### Clinical case report: mosaic ANK3 pathogenic variant in a patient with autism spectrum disorder and neurodevelopmental delay

**Why Not Relevant**: The provided paper content exclusively discusses ankyrins and ANK3. For example: "Ankyrins are a family of proteins that link integral membrane proteins to the underlying spectrin-actin cytoskeleton and play a key role in activities such as cell motility, activation, proliferation, cell–cell contact, and the maintenance of specialized membrane domains." and "Ankyrin 3 (ANK3) is one of the three major subtypes of the ankyrin protein family." The abstract goes on to describe clinical findings for ANK3: "To date, pathogenic variants in ANK3 have been reported in individuals with neuropsychiatric, cognitive, and neurodevelopmental disorders... Here, we report a patient with speech delay, autism spectrum disorder, and a language disorder in which a de novo nonsense ANK3 alteration was discovered by exome sequencing." Nowhere in the provided text is the gene ROCK1 mentioned, nor are there any data, analyses, mechanism descriptions, or references that implicate ROCK1 in Mendelian disease. Because the paper's content is confined to ANK3 and related ankyrin biology, it provides no direct evidence (no variants, association data, clinical cases, or segregation evidence) that ROCK1 is associated with any Mendelian disease. Mechanistically, the paper describes roles of ankyrins in linking membrane proteins to the spectrin-actin cytoskeleton and their importance in the central nervous system; these mechanisms pertain to ANK3 function and do not establish a mechanistic link to ROCK1. Limitations: the assessment is limited to the supplied paper content; if the full paper or supplementary materials (not provided here) mention ROCK1, that content was not available for review. Based on the available text, the paper cannot be used to support or refute the claim about ROCK1 being associated with at least one Mendelian disease.


[Read Paper](https://www.semanticscholar.org/paper/8d26866252dcf55922aa6ce59017540fbbc29c46)


### Case Report: A new case of YARS1-associated autosomal recessive disorder with compound heterozygous and concurrent 47, XXY

**Why Not Relevant**: The provided paper content exclusively discusses YARS1 (Tyrosyl tRNA synthetase) and its variants and associated phenotypes. There is no mention of ROCK1 anywhere in the text, no data (genetic, clinical, or functional) about ROCK1, nor any mechanistic pathway that links YARS1 biology to ROCK1. ROCK1 is a distinct gene encoding Rho-associated coiled-coil containing protein kinase 1 (a serine/threonine kinase), belonging to a different protein family and functional class than aminoacyl-tRNA synthetases. Because the paper provides only evidence about YARS1 and no direct or mechanistic information about ROCK1, it cannot be used to support or refute the claim that ROCK1 is likely associated with at least one Mendelian disease. Therefore the relevance to the ROCK1 claim is effectively zero.


[Read Paper](https://www.semanticscholar.org/paper/e2d6155bb31e3150a3094a55fea567b66910329d)


### Identification of a novel METTL23 gene variant in a patient with an intellectual development disorder: a literature review and case report

**Why Not Relevant**: This paper discusses METTL23 and METTL23-related autosomal recessive intellectual developmental disorder. It does not mention ROCK1, nor does it provide genetic, clinical, or mechanistic data involving ROCK1. The claim concerns whether ROCK1 is likely associated with at least one Mendelian disease; because the paper focuses exclusively on METTL23 (a different gene) and provides no direct or mechanistic evidence linking METTL23 to ROCK1 or describing shared pathways implicating ROCK1 in Mendelian disease, it offers no support for or against the claim. Specific reasons: (1) No occurrence of the string "ROCK1" or any synonyms appears in the provided content. (2) All reported variant, phenotype, and mechanistic details refer to METTL23. (3) There are no experimental data, patient findings, splicing/functional assays, interaction data, or literature citations in the excerpt that would allow a justified inference about ROCK1. Therefore the paper is not relevant to evaluating whether ROCK1 is associated with a Mendelian disease.


[Read Paper](https://www.semanticscholar.org/paper/be57a7832833d4ed117190f4b4c47915c4ab8d54)


### Case report: A study on the de novo KMT2D variant of Kabuki syndrome with Goodpasture’s syndrome by whole exome sequencing

**Why Not Relevant**: This paper does not provide evidence for or against the claim that ROCK1 is associated with a Mendelian disease. The content is a case report of Kabuki syndrome with a de novo KMT2D missense variant (c.15143G > A, p. R5048H) and describes clinical features including pulmonary hemorrhage and positive anti-GBM antibodies; it explicitly names KMT2D and KDM6A as the causative genes. Nowhere in the provided text is the gene ROCK1 mentioned, nor are there genetic, phenotypic, or mechanistic data linking ROCK1 to the reported phenotype(s). Therefore there is no direct evidence (e.g., pathogenic variants in ROCK1 segregating with disease) nor mechanistic evidence (e.g., pathway connections, functional assays implicating ROCK1) in this paper to support the claim. Additional limitations relevant to the paper itself: it is a single-case report focused on KMT2D-associated Kabuki syndrome and a coincident Goodpasture-like presentation, so even findings about KMT2D cannot be extrapolated to ROCK1 without separate data. Because the paper contains no information about ROCK1, it cannot inform the claim.


[Read Paper](https://www.semanticscholar.org/paper/2c51a28e8f2b66dffe79e534a6cd02098b38f6d0)


### Case report: Diagnosis of ADCY5-related dyskinesia explaining the entire phenotype in a patient with atypical citrullinemia type I

**Why Not Relevant**: This paper does not provide any evidence for or against the claim that ROCK1 is associated with a Mendelian disease. The case study discusses pathogenic variants in ASS1 (citrullinemia type 1, MIM #215700) and a de novo pathogenic variant in ADCY5 (ADCY5-related dyskinesia, MIM #606703), and it describes the ADCY5-related mechanism (disruption of the enzymatic conversion of ATP to cAMP) and related clinical/therapeutic consequences. Nowhere in the provided text is the gene ROCK1 mentioned, nor are there experiments, sequence data, clinical observations, or mechanistic discussions that involve ROCK1. Therefore the paper contains no direct evidence (no patient variants, no segregation, no case reports) and no mechanistic evidence (no functional studies, pathways, or interactions) relevant to ROCK1. Absence of mention in this single paper is not evidence that ROCK1 is not associated with a Mendelian disease; it simply means this paper is not informative for that claim. For assessment of ROCK1 gene-disease associations, query disease-gene resources (OMIM, ClinVar, HGMD, ClinGen) or search the broader literature for studies that specifically report ROCK1 variants, segregation, or functional data.


[Read Paper](https://www.semanticscholar.org/paper/1c99698fe2dd4a343b6d726a01d1ad32698ab5b8)


### De novo HDAC3 variants leading to epigenetic machinery dysfunction are associated with a neurodevelopmental disorder

**Why Not Relevant**: The paper exclusively studies HDAC3: its de novo missense variants, associated neurodevelopmental phenotype, and mechanistic effects (reduced deacetylase activity, disrupted protein interactions, impaired nuclear localization). There is no mention of ROCK1, no data on ROCK1 sequence variants, function, patient phenotypes, or mechanistic overlap presented that would support or refute an association between ROCK1 and any Mendelian disease. Therefore the content is not relevant to the claim that ROCK1 is likely associated with at least one Mendelian disease.


[Read Paper](https://www.semanticscholar.org/paper/524b09f9e39f306b6871f816044577dccb53abd6)


### A novel AUTS2 variant in a patient with global developmental delay and intellectual disability

**Why Not Relevant**: The paper exclusively concerns AUTS2 haploinsufficiency and a novel de novo truncating variant in the AUTS2 gene (c.1606C>T, p.Gln536*) causing an AUTS2 neurodevelopmental syndrome. There is no mention of the gene ROCK1, no data on ROCK1 sequence variants, segregation, phenotype association, functional studies, or mechanistic pathways involving ROCK1. Therefore this paper provides no direct evidence that ROCK1 is associated with any Mendelian disease, nor does it provide mechanistic evidence that would bear on ROCK1. Furthermore, because the study is a single-case report about AUTS2, it cannot be used to infer associations for unrelated genes. Excerpt pages: null.


[Read Paper](https://www.semanticscholar.org/paper/f5e84e7bb3b42239a53cc7e5c457342cb38c6277)


## Search Queries Used

- ROCK1 pathogenic variant human Mendelian disorder case report clinical phenotype autosomal dominant autosomal recessive

- Rho associated coiled coil containing protein kinase 1 ROCK1 OMIM ClinVar HGMD monogenic disorder gene disease association

- ROCK1 loss of function haploinsufficiency dominant negative knockout mouse phenotype developmental craniofacial cardiac abnormalities mechanistic study

- ROCK1 de novo variant exome sequencing trio intellectual disability congenital anomalies neurodevelopmental disorder rare variant burden study


## Bibliometric Configuration

- Use Bibliometrics: True

- Author Impact Weight: 0.4

- Citation Impact Weight: 0.4

- Venue Impact Weight: 0.2
