## Supplementary material for "MENDELSEEK: An algorithm that predicts Mendelian Genes and elucidates what makes them special": ROCK2.txt

# Claim: The gene ROCK2 is likely associated with at least one Mendelian disease.

**Status**: processed

**Overall Rating**: -1

**Explanation**:

No relevant papers were found that support or refute this claim.


## Relevant Papers


## Other Reviewed Papers


### The Burden of Candidate Pathogenic Variants for Kidney and Genitourinary Disorders Emerging From Exome Sequencing

**Why Not Relevant**: The supplied paper content describes a study that compiled and curated genes associated with Mendelian kidney and genitourinary disorders and outlines methods for identifying and filtering candidate pathogenic variants. However, the provided excerpts do not mention the gene ROCK2 by name, nor do they provide gene-level results or specific associations for individual genes. Therefore the paper content as provided offers only methodological and contextual information relevant to how Mendelian gene-disease associations are assessed, not direct or gene-specific evidence about whether ROCK2 is likely associated with a Mendelian disease. To evaluate the claim about ROCK2 specifically would require either (a) the curated gene list or supplementary tables showing whether ROCK2 was included and annotated as disease-associated, or (b) reported variant-level findings or clinical correlations implicating ROCK2. None of that gene-specific information is present in the supplied text, so the relevance of this paper to the ROCK2 claim is minimal.


[Read Paper](https://www.semanticscholar.org/paper/a75e297208c04992568e607d19aa9b9aaebd09d5)


### Precision Medicine in Internal Medicine

**Why Not Relevant**: The paper is a general review of genetic testing approaches, diagnostic sequencing workflows, and variant-interpretation issues. It contains no mention of ROCK2, no experimental data, clinical observations, segregation analyses, functional studies, or case reports that would directly support or refute an association between ROCK2 and any Mendelian disease. The extracted sentences provide only general context (e.g., capabilities of exome sequencing, typical variant counts, and the fraction of genes already linked to Mendelian disorders as of 2019). Those statements can inform background probability or explain how an association might be discovered, but they do not constitute gene-specific direct or mechanistic evidence for ROCK2. Therefore the content is not relevant for concluding that ROCK2 is likely associated with at least one Mendelian disease. Additional gene-specific sources (case reports, ClinVar/OMIM entries for ROCK2, segregation and functional studies, or large-scale gene–disease curation resources) would be required to evaluate the claim.


[Read Paper](https://www.semanticscholar.org/paper/c7199e8a5226f8d9d55777da0e15e91797bb459a)


### 54 Novel formin homology 2 domain containing 3 (FHOD3) mutations associated with the pathogenesis of hypertrophic cardiomyopathy (HCM) in an irish population

**Why Not Relevant**: The paper's content exclusively addresses FHOD3 variants and their potential role in hypertrophic cardiomyopathy (HCM). There is no mention of the gene ROCK2, no data (genetic, clinical, or mechanistic) concerning ROCK2, and no discussion of diseases associated with ROCK2. Therefore this paper provides no direct evidence that ROCK2 is associated with any Mendelian disease, nor does it provide mechanistic pathways that could plausibly link ROCK2 to HCM or to other Mendelian disorders. Absence of ROCK2 in the text means the paper cannot support or refute the claim. Note: absence of evidence in this paper is not evidence of absence overall; it only indicates that this source is irrelevant to evaluating the claim about ROCK2. To assess the claim properly, one would need studies that directly report pathogenic variants in ROCK2 segregating with Mendelian phenotypes, functional/mechanistic studies implicating ROCK2 in disease-relevant pathways, or curated clinical-genetic database entries (e.g., OMIM, ClinVar) that associate ROCK2 with Mendelian disease.


[Read Paper](https://www.semanticscholar.org/paper/bc661c80165d8f75b313be1eb974a0d70d965ac1)


### Novel de Novo Nonsense Variants in AGO3 and KHSRP: Insights into Global Developmental Delay and Autism Spectrum Disorders through Whole Genome Analysis

**Why Not Relevant**: This paper does not provide evidence for or against the claim that ROCK2 is likely associated with at least one Mendelian disease. The study is a single-case trio whole-genome analysis that identifies novel de novo nonsense variants in AGO3 and KHSRP and explicitly states those two genes are not currently associated with OMIM phenotypes. Nowhere in the provided text is ROCK2 mentioned, discussed mechanistically, or tested. Therefore there is no direct genetic, clinical, or mechanistic evidence bearing on ROCK2 in this paper. Absence of mention in this paper is not evidence that ROCK2 is or is not associated with Mendelian disease; it only means this paper is not relevant to that claim. Additional reasons for low relevance: the study sample is a single proband (limiting generalizability), the identified variants are in other genes, and the authors themselves call for further study to establish causality for the reported genes—indicating the paper's limited scope and lack of established gene–disease associations that could inform ROCK2 status.


[Read Paper](https://www.semanticscholar.org/paper/5aba59259e6535c76aa2fd80054cb774eb3d7452)


### Novel SYNGAP1 Variant in an Adult Individual Affected by Intellectual Disability and Epilepsy: A Cold Case Solved through Whole-Exome Sequencing

**Why Not Relevant**: This paper focuses entirely on a diagnostic whole-exome sequencing case that identifies a de novo variant in SYNGAP1 as the likely cause of the proband's neurodevelopmental phenotype. There is no mention of the gene ROCK2, no reported ROCK2 variants, no phenotype descriptions attributed to ROCK2, and no mechanistic discussion connecting ROCK2 to the proband's condition or to SYNGAP1. Because the content contains zero data or discussion about ROCK2, it provides no evidence either supporting or refuting the claim that ROCK2 is associated with a Mendelian disease. Absence of evidence in this paper does not disprove the claim, but it means this paper is irrelevant for assessing ROCK2's Mendelian disease associations.


[Read Paper](https://www.semanticscholar.org/paper/e2f8d50ff5458adc25bc78983e545080ac7d5e62)


### De novo HDAC3 variants leading to epigenetic machinery dysfunction are associated with a neurodevelopmental disorder

**Why Not Relevant**: The provided paper content exclusively studies HDAC3 (histone deacetylase 3): its de novo variants, associated neurodevelopmental phenotype, functional assays (HDAC activity, western blot), proteomics (CoREST and NCoR interactions), and localization (immunofluorescence). There is no mention of ROCK2, its variants, expression, pathways involving ROCK2, clinical phenotypes linked to ROCK2, or any resources (e.g., OMIM) evaluated for ROCK2. Because the paper neither investigates nor references ROCK2, it provides no direct genetic or clinical evidence supporting or refuting the claim that ROCK2 is likely associated with at least one Mendelian disease. Mechanistically, the paper's molecular pathways (histone deacetylation, CoREST/NCoR complexes, nuclear localization) do not invoke ROCK2 or Rho-associated kinase signaling, so no mechanistic bridge to implicate ROCK2 is provided. In sum, the article is irrelevant to the claim: it neither provides supporting data nor mechanistic rationale regarding ROCK2 and Mendelian disease.


[Read Paper](https://www.semanticscholar.org/paper/524b09f9e39f306b6871f816044577dccb53abd6)


## Search Queries Used

- ROCK2 Rho associated coiled coil containing protein kinase 2 pathogenic variant likely pathogenic variant Mendelian disease case report family segregation

- ROCK2 ClinVar OMIM GeneReviews Orphanet gene disease association review Mendelian single gene disorder

- ROCK2 de novo variant developmental delay intellectual disability neurodevelopmental disorder exome sequencing rare disease cohort

- ROCK2 loss of function haploinsufficiency gain of function dominant negative knockout mouse phenotype embryonic lethal cardiovascular development neuronal migration actin cytoskeleton functional study


## Bibliometric Configuration

- Use Bibliometrics: True

- Author Impact Weight: 0.4

- Citation Impact Weight: 0.4

- Venue Impact Weight: 0.2
