## Supplementary material for "MENDELSEEK: An algorithm that predicts Mendelian Genes and elucidates what makes them special": RPL3.txt

# Claim: The gene RPL3 is likely associated with at least one Mendelian disease.

**Status**: processed

**Overall Rating**: 0

**Explanation**:

Supporting evidence:
The provided papers describe Diamond-Blackfan anemia (DBA) as a prototypical ribosomopathy caused by heterozygous loss-of-function mutations or deletions in over 20 ribosomal protein genes. That general statement establishes that mutations in ribosomal protein genes can and do cause Mendelian (usually autosomal dominant, haploinsufficiency) disorders. RPL3 encodes a core component of the large (60S) ribosomal subunit (ribosomal protein L3), so, on mechanistic grounds, it is biologically plausible that damaging germline variants in RPL3 could perturb ribosome biogenesis or function and thereby produce a Mendelian phenotype analogous to other ribosomal protein–linked diseases such as DBA.

Caveats or contradictory evidence:
Neither of the excerpts mentions RPL3 specifically as an established DBA gene or as causative for any Mendelian disease. The review excerpt notes >20 ribosomal protein genes as DBA causes but gives RPS19 as an example and does not list RPL3. The second paper focuses on an alternative mechanism (OXPHOS disruption) that leads to ribosomal defects and erythroid failure without changes in ribosomal protein composition, and does not provide evidence of RPL3 mutations in patients. The absence of any explicit mention of RPL3 in these two papers—one a review on genetics of DBA and the other an investigation of alternative DBA mechanisms—means there is no direct experimental or genetic evidence in the provided material linking RPL3 to a Mendelian disorder. Absence of a mention in a genetic review that summarizes known DBA genes suggests RPL3 is not among the established DBA genes (at least as of those papers), which weakens the claim.

Analysis of mechanisms and deeper implications:
Ribosomal protein–related Mendelian diseases (ribosomopathies) typically arise via haploinsufficiency of specific ribosomal protein genes that disturb ribosome biogenesis, activate nucleolar stress pathways (e.g., p53), and lead to tissue-specific phenotypes such as erythroid failure. RPL3 is an essential component of the 60S subunit and participates in peptide bond formation and ribosome function; damaging variants could plausibly interfere with these processes. However, the mere mechanistic plausibility does not substitute for genetic evidence: many essential ribosomal proteins are intolerant of variation and might cause embryonic lethality when disrupted (thus not observed as segregating Mendelian disease), while others can be haploinsufficient and produce an identifiable syndrome. Determining whether RPL3 variants cause Mendelian disease requires population and familial genetic data (segregation, de novo occurrences, enrichment of damaging variants in cases vs controls) or functional studies showing pathogenicity of specific variants—none of which are provided here.

Which rating is most appropriate:
Given the available excerpts, there is mechanistic plausibility because DBA and other inherited ribosomopathies are caused by ribosomal protein genes. However, the provided papers do not present any direct genetic, clinical, or functional evidence implicating RPL3 in a Mendelian disease, and the DBA review’s omission of RPL3 from its discussion of known causative genes suggests it is not an established DBA gene. Therefore the correct assessment based solely on the supplied evidence is that there is no evidence supporting the claim that RPL3 is associated with at least one Mendelian disease. The most appropriate rating is "No Evidence."


**Final Reasoning**:

I reviewed the supplied excerpts: they establish that ribosomal protein genes commonly underlie Mendelian ribosomopathies like DBA, which makes RPL3 a biologically plausible candidate. However, neither paper names RPL3 as a disease gene or provides genetic/clinical/functional data linking RPL3 variants to a Mendelian disorder. A review summarizing known DBA genes that does not list RPL3 implies it is not an established DBA gene in the literature cited. Mechanistic plausibility alone is insufficient to conclude that RPL3 is ‘‘likely’’ disease-associated; concrete genetic evidence (segregation, case reports, case-control enrichment, or demonstrated pathogenic variants) is required and absent in the provided material. Therefore the appropriate rating based on the provided evidence is "No Evidence."


## Relevant Papers


### Hematopoietic cell transplantation and gene therapy for Diamond-Blackfan anemia: state of the art and science

**Authors**: S. Bhoopalan (H-index: 8), M. Wlodarski (H-index: 42)

**Relevance**: 0.35

**Bibliometric Impact**: 0.3224


**Excerpts**:

- Diamond-Blackfan anemia (DBA) is one of the most common inherited causes of bone marrow failure in children.

- As a prototypical ribosomopathy, DBA is caused by heterozygous loss-of-function mutations or deletions in over 20 ribosomal protein genes, with RPS19 being involved in 25% of patients.

- In this review, we summarize the results of HCT studies and review advances and potential future directions in hematopoietic stem cell-based therapies for DBA.


**Explanations**:

- This sentence provides clinical and genetic-context background: DBA is an inherited (Mendelian) disorder. It is relevant because the claim concerns whether a ribosomal protein gene (RPL3) is likely associated with at least one Mendelian disease. However, this sentence is contextual only and does not provide gene-specific or mechanistic evidence about RPL3 itself. Strength: establishes that Mendelian disease caused by ribosomal protein genes exists. Limitation: no mention of which ribosomal protein genes are implicated.

- This sentence is the key mechanistic statement in the paper: it explicitly says DBA is caused by heterozygous loss-of-function mutations or deletions in over 20 ribosomal protein genes and gives an example (RPS19). This provides mechanistic evidence that loss-of-function in ribosomal protein genes can produce a Mendelian disease phenotype (DBA), thereby increasing the plausibility that other ribosomal protein genes (such as RPL3) might also be disease-associated. Strength: directly supports the general mechanism (ribosomopathy via heterozygous LOF in ribosomal protein genes). Limitations: it does not list the 20+ genes, does not mention RPL3 by name, and does not provide primary genetic data implicating any specific additional gene. Therefore it is indirect evidence for RPL3 specifically.

- This sentence indicates the article is a review of clinical and therapeutic studies for DBA. It is relevant because review articles synthesize prior genetic and mechanistic findings, but as presented here it does not supply primary or gene-specific data about RPL3. Strength: signals that the source is a review (so conclusions are secondary synthesis). Limitation: being a review means the excerpted content may summarize many studies but the provided text does not include details or citations that would confirm whether RPL3 is among the implicated genes.


[Read Paper](https://www.semanticscholar.org/paper/aed51f3a418368efd390afc674bca103ff00f1cd)


### Oxidative phosphorylation pathway disruption is an alternative pathological mechanism leading to Diamond-Blackfan anemia

**Authors**: Rudan Xiao (H-index: 2), X. Fang (H-index: 1)

**Relevance**: 0.2

**Bibliometric Impact**: 0.132


**Excerpts**:

- Diamond-Blackfan anemia (DBA) is a rare congenital disorder characterized by the failure of erythroid progenitor differentiation; however, the molecular mechanisms leading to erythroid defects remain unclear.

- By analyzing the transcriptomic profiles of bone marrow from patients with DBA (n = 10), we identified the dysfunction of the oxidative phosphorylation (OXPHOS) pathway as a possible cause of DBA.

- We established a DBA cell model using differentiating hematopoietic stem progenitor cells in which the OXPHOS pathway was suppressed to completely recapitulate the defects in erythroid progenitor differentiation, ribosome biogenesis, and heme biosynthesis, which are representative characteristics of patients with DBA.

- Disruption of the OXPHOS pathway led to ribosomal defects and associated erythroid defects via abolishment of the Ran GTPase activating protein RanGAP1, which is pivotal in the RNA transport pathway.

- The composition of the ribosomal proteins in the established DBA cells was unchanged, but an overall reduction in ribosomal protein levels was observed, leading to an alteration in the translation of a subset of transcripts specific to erythropoiesis.

- We revealed that the OXPHOS pathway participates in erythropoiesis, particularly at an early stage, and reinforced the relationship between the OXPHOS pathway and erythropoiesis.

- Coenzyme Q10, an activator of OXPHOS, largely rescued the erythroid defects in DBA cells.

- Our results reveal that OXPHOS repression is an alternative pathological mechanism leading to DBA, demonstrating its potential as a therapeutic pathway.

- Key Points Oxidative phosphorylation (OXPHOS) pathway disruption is an alternative pathological mechanism underlying Diamond-Blackfan anemia (DBA).

- Key Points Suppression of OXPHOS leads to defects in erythropoiesis and ribosomal biogenesis via the RanGAP1 protein.


**Explanations**:

- Context: This sentence defines the disease studied (DBA) and frames the paper's aim. Relation to claim: indirect context only. Strength/Limitations: It establishes the clinical phenotype (erythroid progenitor differentiation failure) but does not mention molecular causes or any specific ribosomal protein genes (such as RPL3). This excerpt is not direct evidence for RPL3 involvement.

- Context: Describes the transcriptomic analysis and sample size (n = 10). Relation to claim: indirect mechanistic relevance. Strength/Limitations: Identifies a pathway (OXPHOS) implicated in DBA from patient data, which is relevant because DBA is often linked to ribosomal dysfunction; however, this sentence gives no gene-level (e.g., RPL3) association or genetic evidence. The sample size is modest (n = 10), limiting statistical power for gene-specific claims.

- Context: Summary of the cellular model recapitulating DBA features. Relation to claim: mechanistic relevance. Strength/Limitations: Shows that suppression of OXPHOS in a cell model reproduces defects in ribosome biogenesis (a process involving RPL3), providing mechanistic plausibility that perturbing ribosomal function leads to DBA-like phenotypes. Limitation: the experiment manipulates a pathway (OXPHOS) rather than testing mutations in a specific ribosomal protein gene like RPL3, so it does not provide direct evidence that RPL3 mutations cause a Mendelian disease.

- Context: Mechanistic link between OXPHOS disruption and ribosomal/erythroid defects via RanGAP1. Relation to claim: mechanistic. Strength/Limitations: Demonstrates a specific mechanistic cascade (OXPHOS -> RanGAP1 loss -> ribosomal defects) that could explain how ribosomal dysfunction arises in DBA. This strengthens the plausibility that ribosomal protein perturbations underlie disease, but it remains indirect regarding any single ribosomal protein gene (e.g., RPL3). It does not present genetic or clinical data implicating RPL3 mutations.

- Context: Observations about ribosomal protein levels and translational consequences in the model. Relation to claim: mechanistic and partially functional. Strength/Limitations: Directly reports altered ribosomal protein abundance (overall reduction) and downstream effects on translation of erythropoiesis-specific transcripts, which is mechanistically relevant because RPL3 encodes a core ribosomal protein. However, the excerpt states that composition was unchanged and reports only global reduction — it does not single out RPL3, nor does it provide mutation or inheritance data tying RPL3 to a Mendelian disease.

- Context: Summarizes the role of OXPHOS in erythropoiesis. Relation to claim: contextual mechanistic relevance. Strength/Limitations: Reinforces the pathway-level connection between metabolism and ribosome-dependent erythropoiesis, which again makes ribosomal genes plausible contributors to pathology. Limitation: no gene-specific evidence for RPL3 or genetic association data.

- Context: Therapeutic rescue experiment. Relation to claim: mechanistic/functional. Strength/Limitations: Rescue of defects by Coenzyme Q10 supports causality of OXPHOS dysfunction in the model and suggests reversibility of ribosomal/erythroid defects caused by pathway suppression. This supports the idea that perturbation of ribosomal function can be disease-relevant, but it does not provide evidence that RPL3 is mutated in a Mendelian manner in patients.

- Context: Overall conclusion that OXPHOS repression can cause DBA. Relation to claim: indirect support. Strength/Limitations: Establishes an alternative disease mechanism leading to DBA, reinforcing that ribosomal dysfunction can be pathologic. Limitation: still pathway-level and not gene-specific; no mention of RPL3 or of inheritance patterns linking RPL3 to Mendelian disease.

- Context: Key point summarizing mechanism linking OXPHOS disruption to DBA. Relation to claim: mechanistic relevance. Strength/Limitations: Concise restatement of pathway-level finding; relevant to the general hypothesis that ribosomal dysfunction can cause DBA but provides no direct genetic evidence implicating RPL3.

- Context: Key point summarizing suppression of OXPHOS effects via RanGAP1. Relation to claim: mechanistic relevance. Strength/Limitations: Reinforces the mechanistic chain from OXPHOS to ribosomal biogenesis defects via RanGAP1. Helpful for plausibility that ribosomal proteins (including RPL3) could be involved in disease phenotypes, but does not give gene-specific or Mendelian association data for RPL3.


[Read Paper](https://www.semanticscholar.org/paper/4bd9fa4990db4f1027be5093d472efcbc46d25d0)


## Other Reviewed Papers


### Perspectives of current understanding and therapeutics of Diamond-Blackfan anemia

**Why Not Relevant**: The paper excerpt mentions RPS19 (a ribosomal protein) and gene therapy for RPS19-deficient Diamond-Blackfan anemia, but does not mention RPL3 or provide any genetic, clinical, or experimental data about RPL3. While it offers a general mechanistic point that mutations in ribosomal protein genes can cause Mendelian disease (DBA), that is only indirect and speculative with respect to RPL3. There is no direct association, no variant data, no patient phenotypes linked to RPL3, and no experimental manipulation of RPL3 described. Because the excerpt contains no direct or specific evidence about RPL3, its relevance to the claim that "RPL3 is likely associated with at least one Mendelian disease" is minimal. Consequently the evidence in this paper is insufficient to support or refute the claim.


[Read Paper](https://www.semanticscholar.org/paper/623633e8f787ac7d1cd01ef6d9ad5a777fb3a09a)


### Predictors of the utility of clinical exome sequencing as a first-tier genetic test in patients with Mendelian phenotypes: results from a referral center study on 603 consecutive cases

**Why Not Relevant**: The provided paper content contains a single summary sentence about overall diagnostic yield from clinical exome sequencing (CES) and predictors of CES utility: "Based on a large dataset of consecutive patients with various Mendelian phenotypes referred for CES as a first-tier genetic test, a diagnostic yield of 40% and several independent predictors of CES utility that might improve CES diagnostic efficiency are reported." This sentence does not mention the gene RPL3, does not report any gene-level findings, does not report specific variants, and provides no mechanistic information linking RPL3 to any Mendelian phenotype. Therefore, there is no direct evidence (no reported association, no variant pathogenicity, no segregation data) and no mechanistic evidence (no functional studies, pathways, or biological rationale) in the provided content to support or refute the claim that RPL3 is likely associated with at least one Mendelian disease. 

Limitations and caveats: (1) Only a single summary sentence was provided rather than the full paper. The full text might contain gene-level tables, supplementary lists, or case reports that mention RPL3; those could be relevant but are not present here. (2) Aggregate statistics (e.g., a 40% diagnostic yield) do not allow inference about any particular gene unless gene-specific results are reported. (3) Without specific excerpts that name RPL3 or describe relevant variants or mechanisms, neither direct nor mechanistic support can be extracted. If you can provide the full text or the sections listing identified genes/variants or supplementary tables, I can re-evaluate for evidence about RPL3.


[Read Paper](https://www.semanticscholar.org/paper/3e4076a587328718af716949d2e787a464220fa6)


### Novel LSS variants in alopecia and intellectual disability syndrome: New case report and clinical spectrum of LSS‐related rare disease traits

**Why Not Relevant**: The provided paper content exclusively discusses the gene LSS and pathogenic biallelic variants in LSS being associated with multiple Mendelian rare disease traits (including APMR4). There is no mention of the gene RPL3, no genetic findings, no mechanistic data, and no clinical associations involving RPL3 in the text. Therefore the paper provides no direct or mechanistic evidence that RPL3 is associated with any Mendelian disease. Because the topic and data center entirely on LSS, the content is not relevant to evaluating the claim about RPL3.


[Read Paper](https://www.semanticscholar.org/paper/fe90a5bb55278bc80588741eed5def6cf4f1aa82)


### Case report: Discovery of a de novo FAM111B pathogenic variant in a patient with an APECED-like clinical phenotype

**Why Not Relevant**: The paper focuses on AIRE and FAM111B as genetic causes of the syndromes discussed and reports a de novo pathogenic variant in FAM111B in a single patient. The text does not mention RPL3, provide any association study, case reports, functional data, or mechanistic discussion related to RPL3. Performing exome sequencing in this patient does not imply evaluation or reporting of RPL3-specific results relevant to the broader claim. Therefore the paper provides no direct or mechanistic evidence for or against the claim that RPL3 is likely associated with at least one Mendelian disease. Limitations preventing any inference include: (1) absence of any mention of RPL3 or related findings, (2) single-patient focus on different genes, and (3) lack of population-level, segregation, or functional data pertaining to RPL3. Because of these points, the content is not relevant to assessing the claim.


[Read Paper](https://www.semanticscholar.org/paper/6ed49c6ffda4dbf2de15474aa07c9326d5636a34)


### Exome sequencing identified a novel Col6α1 mutation in an Iranian patient with Ullrich congenital muscular dystrophy: a case report

**Why Not Relevant**: The paper content provided reports a clinical/genetic finding linking a homozygous c.2551_2562 mutation in COL6α1 to a UCMD phenotype in a 4-year-old patient. There is no mention of the gene RPL3, no genetic or functional data concerning RPL3, and no mechanistic discussion connecting COL6α1 and RPL3. Because the content is entirely focused on a different gene and disease (COL6α1 and UCMD), it provides no relevant direct or mechanistic evidence for or against the claim that RPL3 is likely associated with at least one Mendelian disease.


[Read Paper](https://www.semanticscholar.org/paper/e65cf4aab08675ea16a9bc4216bff41793dabf49)


### A Phase I/II, Intra-Patient Dose-Escalation Study of Bitopertin, a Selective Inhibitor of GlyT1 and Heme Synthesis, for Steroid-Refractory Diamond-Blackfan Anemia

**Why Not Relevant**: The provided paper content does not mention RPL3 by name, nor does it report any genetic sequencing, variant identification, case reports, segregation analyses, functional assays of RPL3 variants, or statistical association implicating RPL3 in a Mendelian disease. The paper discusses Diamond-Blackfan anemia (DBA) and notes that causative mutations, when identified, are usually in ribosomal protein (RP) genes; this is a class-level statement that increases the plausibility that other RP genes could cause Mendelian disease, but it is circumstantial. Without gene-specific data (variants in RPL3, affected individuals with RPL3 mutations, functional impact of such mutations, inheritance pattern, or clinical correlation), the content cannot be taken as direct evidence that RPL3 is associated with at least one Mendelian disease. Therefore the relevance to the specific claim (RPL3 likely associated with a Mendelian disease) is very low.


[Read Paper](https://www.semanticscholar.org/paper/de953e06a827835256cc2aeca8cd2c99f4d31c6b)


### Novel Mouse Model That Recapitulates the Hematologic Phenotype of Diamond Blackfan Anemia

**Why Not Relevant**: Relevance is very low because the paper does not mention RPL3 at all and presents no human genetic, segregation, mutation, or functional evidence specific to RPL3. The manuscript focuses on RPL11: human RPL11 mutations in DBA and an inducible Rpl11 haploinsufficient mouse model that phenocopies DBA. The paper does, however, provide class-level mechanistic evidence that loss-of-function mutations in ribosomal protein genes can cause Mendelian ribosomopathies (e.g., DBA). That class-level statement increases biological plausibility that other ribosomal protein genes, potentially including RPL3, could be disease-causing, but plausibility is not the same as evidence. Specific limitations preventing this paper from supporting the claim about RPL3 include: (1) absence of any mention of RPL3 sequence variants, patient data, or functional perturbation of RPL3; (2) all gene-specific experimental data pertain to RPL11 only; (3) mouse-model and small-sample (n values given) results pertain to Rpl11 and may not generalize to all RP genes; and (4) even when class-level mechanistic inference is possible, gene-specific differences (expression, redundancy, essentiality, interactors) can make RPL3 behavior different from RPL11. Therefore this paper does not provide direct or sufficient mechanistic evidence to conclude that RPL3 is likely associated with at least one Mendelian disease.


[Read Paper](https://www.semanticscholar.org/paper/9a5b100b5d60beb5063671656aba08b24f646b09)


## Search Queries Used

- RPL3 pathogenic variant Mendelian disease human case report clinical exome sequencing

- ribosomal protein L3 RPL3 ribosomopathy congenital disorder bone marrow failure Diamond Blackfan anemia

- RPL3 loss of function haploinsufficiency missense splice mutation functional study knockout mouse zebrafish developmental phenotype

- RPL3 ClinVar OMIM gnomAD pLI loss of function intolerance de novo variant trio sequencing Mendelian gene discovery


## Usage Statistics

- Prompt Tokens: 0

- Completion Tokens: 0

- Total Tokens: 0

- Estimated Cost: $0.0433


## Bibliometric Configuration

- Use Bibliometrics: True

- Author Impact Weight: 0.4

- Citation Impact Weight: 0.4

- Venue Impact Weight: 0.2
