## Supplementary material for "MENDELSEEK: An algorithm that predicts Mendelian Genes and elucidates what makes them special": RPL18A.txt

# Claim: The gene RPL18A is likely associated with at least one Mendelian disease.

**Status**: processed

**Overall Rating**: 3

**Explanation**:

Supporting evidence.
The provided paper reviews ribosomopathies and emphasizes that mutations in ribosomal proteins or ribosome biogenesis factors underlie a number of Mendelian, tissue-specific disorders (for example, Diamond–Blackfan anemia). It describes mechanisms (ribosome haploinsufficiency, altered translational control, activation of p53 pathways) by which loss or dysfunction of ribosomal proteins can produce disease phenotypes. Because RPL18A encodes a component of the ribosome (60S ribosomal protein L18a), the paper provides a plausible, mechanism-based rationale that mutations in ribosomal protein genes can cause Mendelian diseases. That group-level evidence therefore supports the general plausibility that another ribosomal protein gene such as RPL18A could be associated with at least one Mendelian disorder.

Caveats or contradictory evidence.
The excerpted material does not mention RPL18A specifically and contains no direct evidence (case reports, genetic association, OMIM entry, or functional studies) linking RPL18A to any Mendelian phenotype. Not all ribosomal proteins are known to cause Mendelian disease; only a subset (e.g., RPS19, RPL5, RPL11 and others) have well-documented associations. Some ribosomal protein perturbations may be embryonic lethal or tolerated due to redundancy, and thus never present as an inheritable, viable Mendelian syndrome. Therefore the provided paper supports a class effect (ribosomal protein mutations can cause Mendelian disease) but does not supply gene-level evidence for RPL18A specifically.

Analysis of mechanisms and deeper implications.
Mechanistically, ribosomal protein haploinsufficiency can lead to specific tissue vulnerabilities through altered translational control and p53-mediated responses; such mechanisms would apply to RPL18A in principle because it contributes to the large ribosomal subunit. If heterozygous loss-of-function of RPL18A produced selective translational defects in sensitive lineages (e.g., erythroid progenitors), one could expect a ribosomopathy-like Mendelian phenotype. Conversely, if RPL18A is essential for early development such that loss-of-function causes embryonic lethality or if its function is buffered by paralogs/compensatory mechanisms, pathogenic variants might be absent from liveborn individuals and thus absent from clinical reports. Therefore mechanistic plausibility exists but outcome depends on gene-specific biology (essentiality, dosage sensitivity, expression patterns) that is not established in the excerpt.

Which rating is most appropriate.
Given the evidence at hand, the claim is supported at the level of general plausibility (ribosomal proteins as a class cause Mendelian diseases) but lacks any direct, gene-specific supporting evidence for RPL18A. There is no contradicting paper-level evidence showing RPL18A is not associated with disease, but absence of gene-level evidence is an important limitation. Balancing the plausible mechanism against the lack of direct observations, the most accurate characterization is that the evidence is mixed: the class-level data make the claim plausible, but the provided literature does not document an actual Mendelian association for RPL18A. Therefore "Mixed Evidence" best reflects the current state based on the supplied material.


**Final Reasoning**:

I reaffirm the rating "Mixed Evidence." The reviewed paper gives a strong, general argument that mutations in ribosomal protein genes cause Mendelian disorders and outlines plausible mechanisms (haploinsufficiency, translational specificity, p53 activation) that would apply to ribosomal proteins including RPL18A. However, the excerpts contain no gene-specific evidence linking RPL18A to any Mendelian disease, and not all ribosomal proteins are known disease genes — some may be lethal when mutated or functionally redundant. Because the evidence is indirect and lacks documented cases, a judgment that the claim is definitively true would be premature, but the claim is not contradicted. Thus the balance of the provided evidence is mixed: plausible but unproven for RPL18A specifically.


## Relevant Papers


### Ribosomopathies: There’s strength in numbers

**Authors**: Eric W. Mills (H-index: 9), R. Green (H-index: 71)

**Relevance**: 0.45

**Bibliometric Impact**: 0.5409714285714287


**Excerpts**:

- Ribosomopathies are t issuespecific disorders that result from mutations in ribosomal proteins or ribosome biogenesis factors.

- This group broadly comprises two categories: (i) disorders caused by single-copy mutations in specific ribosomal proteins, and (ii) disorders associated with defects in ribosome biogenesis factors.

- Ribosomopathies are a heterogeneous group of human disorders that are in some cases known, and in other cases suspected, to result from ribosome dysfunction.

- These include effects on bone marrow–derived cell lineages and skeletal tissues.

- An increasing number of studies in models of distinct ribosomopathies have revealed that “ribosomal stress” signals converge on the p53 signaling pathway in affected cells and tissues.

- One hypothesis is that ribosome dysfunction (or deficiency) can affect global and messenger RNA (mRNA)–specific translational control, and that certain specific cells or tissues may be more vulnerable to ribosome dysfunction.

- We discuss ribosome homeostasis as an overarching principle that governs the sensitivity of specific cells and tissue types to ribosomal protein mutations.

- Ribosomopathies as a group are diverse in their origins and clinical manifestations; we use the well-described Diamond-Blackfan anemia (DBA) as a specific example to highlight some common features.

- Mathematical models and experimental insights rationalize how even subtle shifts in the availability of ribosomes, such as those created by ribosome haploinsufficiency, can drive messenger RNA–specific effects on protein expression.


**Explanations**:

- This sentence states, in general terms, that mutations in ribosomal proteins can cause tissue-specific disorders (ribosomopathies). It provides class-level direct relevance to the claim because RPL18A encodes a ribosomal protein; however, it is not specific to RPL18A. Strength: clear assertion that ribosomal protein mutations cause disease. Limitation: typographical error present and no gene-specific data or evidence provided—this is a general statement from a review.

- This excerpt explicitly indicates that disorders can be caused by single-copy (likely heterozygous/haploinsufficient) mutations in specific ribosomal proteins. Relevance: mechanistic/direct class-level evidence that haploinsufficiency of ribosomal protein genes can produce Mendelian disorders. Strength: supports biological plausibility that a ribosomal protein gene (such as RPL18A) could be disease-associated via single-copy mutation. Limitation: still not evidence that RPL18A itself is affected—no gene-specific genetic or clinical association data are provided.

- This sentence frames ribosomopathies as known or suspected outcomes of ribosome dysfunction. Relevance: supports the general concept that defects in ribosome components (including ribosomal protein genes) underlie Mendelian disorders. Strength: places ribosomal protein mutations in the context of human disease. Limitation: nonspecific—does not identify particular genes or studies linking RPL18A to disease.

- This sentence describes common phenotypic targets (bone marrow–derived lineages and skeletal tissues) of ribosomopathies. Relevance: provides phenotypic context for ribosomal protein mutation effects, useful for evaluating whether a gene’s dysfunction could plausibly produce a Mendelian phenotype. Limitation: again, no direct link to RPL18A; review-level overview.

- This sentence gives a mechanistic pathway (p53 activation) by which ribosomal dysfunction leads to cell cycle arrest and apoptosis in affected tissues. Relevance: mechanistic evidence strengthening plausibility that mutations in ribosomal proteins can produce disease phenotypes through conserved cellular stress pathways—if RPL18A dysfunction perturbs ribosome function, similar downstream effects could occur. Limitation: this is a summary of multiple studies, not direct evidence that RPL18A mutations activate p53 or cause disease in humans.

- This sentence describes a mechanistic hypothesis about how ribosome dysfunction affects translation in an mRNA-specific manner and how certain tissues may be more vulnerable. Relevance: mechanistic support for how mutation in any ribosomal protein gene (including RPL18A) could yield tissue-restricted Mendelian phenotypes. Limitation: hypothesis/interpretation level rather than direct experimental proof for particular genes.

- This sentence posits ribosome homeostasis as a principle governing sensitivity to ribosomal protein mutations. Relevance: mechanistic and conceptual support that loss or alteration of ribosomal protein genes can have pathogenic consequences; it increases plausibility that a ribosomal protein gene like RPL18A could be implicated in Mendelian disease. Limitation: general principle; no empirical RPL18A-specific data.

- This sentence cites Diamond-Blackfan anemia as a well-described example of a ribosomopathy. Relevance: provides a concrete Mendelian disease known to be caused by mutations in ribosomal proteins, strengthening the class-level inference that other ribosomal protein genes (potentially RPL18A) could be Mendelian-disease-associated. Limitation: does not establish that RPL18A is among the DBA genes or linked to DBA or any other Mendelian disorder.

- This sentence indicates that even subtle ribosome haploinsufficiency can drive mRNA-specific changes in protein expression. Relevance: mechanistic evidence (and modeling/experimental support) that small quantitative changes in ribosomal protein dosage can have biologically significant effects—supporting the plausibility that heterozygous loss-of-function in a ribosomal protein gene could be pathogenic. Limitation: inferential—no direct genetic association or clinical data for RPL18A are provided.


[Read Paper](https://www.semanticscholar.org/paper/3fa757adedfac0659b7c19ee93dd94371925e1dc)


## Other Reviewed Papers


### The Burden of Candidate Pathogenic Variants for Kidney and Genitourinary Disorders Emerging From Exome Sequencing

**Why Not Relevant**: The provided paper excerpt does not mention the gene RPL18A at all, nor does it provide a complete gene list or any gene-specific findings that would allow assessment of whether RPL18A is associated with a Mendelian disease. The content focuses on methods for identifying and curating genes associated with Mendelian kidney and genitourinary disorders and on cohort and variant-filtering procedures. Because the claim concerns RPL18A specifically and the excerpt contains no direct observations, results, or discussion about RPL18A (presence/absence in the curated gene list, observed pathogenic variants, inheritance patterns, mechanistic data, or prior disease reports), this paper excerpt cannot provide direct or substantive mechanistic evidence for or against the claim. Any inference about RPL18A from this text would be speculative. Therefore relevance is very low.


[Read Paper](https://www.semanticscholar.org/paper/a75e297208c04992568e607d19aa9b9aaebd09d5)


### Whole-Exome Sequencing in Adults With Chronic Kidney Disease

**Why Not Relevant**: The provided paper content describes a WES study in adults with chronic kidney disease and details the sequencing, annotation, and variant interpretation pipeline focused primarily on a curated list of 287 OMIM 'nephropathy' genes and the ACMG 59 actionable genes. The excerpts do not mention RPL18A at all, and there are no reported findings, variant calls, or disease associations involving RPL18A in the supplied text. While the methods (WES, annotation against ClinVar/HGMD, ACMG classification) could in principle detect a protein-coding pathogenic variant in RPL18A, the study's scope—prioritizing kidney- and hypertension-related OMIM genes—and the absence of any explicit reference to RPL18A mean the paper provides no direct evidence supporting or refuting the claim that RPL18A is likely associated with at least one Mendelian disease. Additionally, capture limitations and gene-list choices further reduce the ability to infer anything about RPL18A from this paper. For these reasons, relevance to the specific claim about RPL18A is very low.


[Read Paper](https://www.semanticscholar.org/paper/a29bd57964f68ac5f8e1abcafe1112ba488f53f5)


### Frequency of Variants in Mendelian Alzheimer's Disease Genes within the Alzheimer's Disease Sequencing Project (ADSP)

**Why Not Relevant**: The paper exclusively studies three Mendelian Alzheimer's disease genes (PSEN2, PSEN1, APP) using ADSP WGS/WES data and reports variants and age-of-onset associations for those genes. There is no mention of RPL18A, no variant data for RPL18A, and no mechanistic discussion involving RPL18A. Therefore this paper provides no direct genetic evidence or mechanistic support for the claim that RPL18A is likely associated with at least one Mendelian disease. Note that absence of evidence in this paper is not evidence of absence overall: to evaluate the claim about RPL18A one would need to consult other sources (ClinVar, OMIM, primary gene-disease association studies, functional assays, segregation analyses), because this study's design and gene selection preclude any assessment of RPL18A.


[Read Paper](https://www.semanticscholar.org/paper/3760efb72c8ec6adaf553ac773087fd2b16aeafc)


### Novel variants of NEK9 associated with neonatal arthrogryposis: Two case reports and a literature review

**Why Not Relevant**: This paper focuses exclusively on pathogenic variants in NEK9 and their association with neonatal arthrogryposis. The provided content (Objective, Methods, Results, Conclusion) describes trio-whole-exome sequencing and copy number variation analysis that identified three novel pathogenic NEK9 variants in two families and discusses how these NEK9 variants expand the phenotype and gene spectrum of NEK9-associated arthrogryposis. Nowhere in the supplied text is the gene RPL18A mentioned, nor are there any data, results, mechanistic discussions, or incidental findings reported about RPL18A. Therefore the paper provides no direct evidence that RPL18A is associated with any Mendelian disease, and it provides no mechanistic evidence linking RPL18A to disease. Note also that absence of a mention of RPL18A in this paper is not evidence that RPL18A is not associated with Mendelian disease more generally; it only means this paper does not bear on the claim. Additional sources would be required to assess the claim about RPL18A.


[Read Paper](https://www.semanticscholar.org/paper/a69677b41b5f628b50b661893ae39d1b61fa7955)


### De Novo RPS20 Mutations in Diamond Blackfan Anemia

**Why Not Relevant**: The paper does not mention RPL18A anywhere. All evidence concerns other ribosomal protein genes (RPS7, RPS10, RPS17, RPS19, RPS24, RPS26, RPS29, RPL5, RPL11, RPL15, RPL26, RPL31, RPL35A, and RPS20). While the paper provides strong mechanistic and genetic evidence that mutations in certain ribosomal protein genes cause Mendelian diseases (notably Diamond Blackfan anemia) and shows that additional ribosomal protein genes can be implicated (RPS20), there is no direct genetic, clinical, or functional data about RPL18A. Thus the paper offers only indirect, family-level mechanistic plausibility (that other ribosomal proteins can be Mendelian disease genes) but no specific evidence tying RPL18A to any Mendelian disorder. Because of that absence, the paper is largely irrelevant to the specific claim that RPL18A is likely associated with at least one Mendelian disease.


[Read Paper](https://www.semanticscholar.org/paper/a615979f9c2853ae21d1e572a50954ed02561c7a)


### Ribosomopatias. Doenças e disfunção do ribosoma

**Why Not Relevant**: The provided content consists only of high-level section headings from a review on ribosomal abnormalities and ribosomopathies (for example: "DiamondBlackfan anemia", "5q− syndrome", "Ribosomal haploinsufficiency in human disease", etc.). There are no complete sentences, paragraphs, or any mention of the specific gene RPL18A in the supplied text. Therefore the excerpt contains no direct evidence (no reported cases, mutations, genetic linkage, segregation, functional studies, or diagnostic associations) that RPL18A is associated with any Mendelian disease. 

Mechanistic plausibility: The review topic (ribosomal proteins and ribosomopathies) makes it biologically plausible in general that a ribosomal protein gene could cause a Mendelian disease via haploinsufficiency or other mechanisms. However, such a general mechanistic statement is indirect and insufficient to support the claim that RPL18A specifically is likely associated with at least one Mendelian disease. The provided headings do not supply mechanistic details tied to RPL18A (e.g., loss-of-function effects, tissue specificity, p53 activation, model organism phenotypes) nor do they provide evidence of pathogenic variants, inheritance patterns, or clinical phenotypes for RPL18A.

Limitations and caveats: (1) Absence of evidence in the supplied headings is not evidence of absence in the full paper or the literature — the full article could mention RPL18A in sections not included here. (2) The headings indicate that some ribosomal proteins are implicated in Mendelian disorders (e.g., Diamond–Blackfan anemia) but do not identify which ribosomal protein genes are involved; many ribosomal protein genes differ in their disease associations. (3) To assess the claim definitively requires searching the full text for "RPL18A" and consulting genetic-disease resources (OMIM, ClinVar, HGMD) and primary literature for reported pathogenic variants, segregation, frequency data, and functional studies.

Conclusion: Based solely on the provided content, this paper does not provide direct or specific mechanistic evidence either supporting or refuting the claim that RPL18A is likely associated with at least one Mendelian disease.


[Read Paper](https://www.semanticscholar.org/paper/370ea1d0786158a8ae3e1f6fc162fd58c388bb98)


### Hematopoietic Stem Cells Supporting Fetal Erythropoiesis Are Differentially Regulated By Small and Large Ribosomal Subunits

**Why Not Relevant**: Relevance is low because the paper does not mention RPL18A at all. The content focuses on RPS19, RPL5, and in part RPL40 (Uba52), reporting mouse-model and mechanistic evidence that mutations/haploinsufficiency in certain ribosomal protein genes cause Diamond Blackfan anemia. While these results provide indirect mechanistic support for the general plausibility that other ribosomal protein genes (including RPL18A) might underlie Mendelian disorders, there is no direct genetic, clinical, or experimental evidence concerning RPL18A in this paper. Additionally, the authors explicitly note that defects may not be common to all ribosomal proteins, which limits how much one can infer about RPL18A from these data.


[Read Paper](https://www.semanticscholar.org/paper/10c5d5ab5c3f1eac33951613daf8bcb441d5a0c9)


## Search Queries Used

- RPL18A Mendelian disease pathogenic variant ClinVar OMIM case report exome sequencing trio RPL18A variant

- RPL18A ribosomal protein ribosome biogenesis nucleolar stress p53 haploinsufficiency ribosomopathy Diamond Blackfan anemia congenital anemia skeletal anomalies

- RPL18A loss of function intolerance LOEUF pLI gnomAD constraint haploinsufficiency score ClinGen gene disease validity clinical curation

- RPL18A knockout mouse zebrafish morpholino CRISPR functional assay ribosome subunit 60S translation defect embryonic lethality developmental disorder


## Usage Statistics

- Prompt Tokens: 0

- Completion Tokens: 0

- Total Tokens: 0

- Estimated Cost: $0.0421


## Bibliometric Configuration

- Use Bibliometrics: True

- Author Impact Weight: 0.4

- Citation Impact Weight: 0.4

- Venue Impact Weight: 0.2
