## Supplementary material for "MENDELSEEK: An algorithm that predicts Mendelian Genes and elucidates what makes them special": RPS6KB1.txt

# Claim: The gene RPS6KB1 is likely associated with at least one Mendelian disease.

**Status**: processed

**Overall Rating**: -1

**Explanation**:

No relevant papers were found that support or refute this claim.


## Relevant Papers


## Other Reviewed Papers


### Setd5 is required in cardiopharyngeal mesoderm for heart development and its haploinsufficiency is associated with outflow tract defects in mouse

**Why Not Relevant**: The provided paper text makes no mention of the gene RPS6KB1 and provides no direct or indirect evidence linking RPS6KB1 to any Mendelian disease. The study describes identification and characterization of Setd5 (a chromatin modifier) and its haploinsufficiency phenotypes in mouse models, and discusses Tbx1 and congenital heart defects; none of these sentences name RPS6KB1 or describe pathways that implicate RPS6KB1. Because the claim is specific to RPS6KB1, absence of any reference to that gene means this paper cannot support or refute the claim. Limitations: although the paper addresses general concepts (haploinsufficiency, network interactions, mouse embryonic lethal screens) that are methodologically relevant for discovering Mendelian disease genes, these general mechanisms do not provide mechanistic evidence for RPS6KB1 specifically. To assess the claim about RPS6KB1, one would need publications or database entries that explicitly report RPS6KB1 genetic variants segregating with Mendelian phenotypes, functional studies of RPS6KB1 loss-of-function in relevant models, or curated gene–disease annotations (e.g., OMIM, ClinGen, ClinVar, HGMD).


[Read Paper](https://www.semanticscholar.org/paper/640ca11404c6c6b8ba7083b50642fd3062edf14a)


### Case report: De novo mutation of a-galactosidase A in a female patient with end-stage renal disease: report of a case of late diagnosis of Anderson–Fabry disease

**Why Not Relevant**: This paper does not provide evidence for or against the claim that RPS6KB1 is likely associated with at least one Mendelian disease. The manuscript exclusively concerns Anderson–Fabry disease (AFD) and variants in the alpha-galactosidase A gene (GLA). Relevant verbatim sentences from the provided text demonstrate the paper's focus: "Background: Anderson–Fabry disease (AFD) is an X-linked disease that results from reduced activity of the enzyme galactosidase alpha (GLA)." and "Alpha-galactosidase A gene sequence analysis revealed a pathogenic variant of c.947dupT in exon 6, leading to the p. I317NfsTer16 amino acid substitution." and "The international Fabry disease genotype-phenotype database (dbFGP) reports a pathogenic variant c.947dupT in exon 6 that is probably associated with a classical phenotype of Fabry disease." None of the provided text mentions RPS6KB1, its sequence, variants, phenotype associations, or any mechanistic pathway linking RPS6KB1 to Fabry disease or to any Mendelian condition. Therefore: - There is no direct evidence in this paper that RPS6KB1 is associated with a Mendelian disease (no patient genotypes, phenotypes, segregation data, case reports, or statistical associations concerning RPS6KB1). - There is no mechanistic evidence (e.g., biochemical pathways, protein interactions, functional studies) connecting RPS6KB1 to the disease discussed or to Mendelian inheritance in general. Additional caveats about the paper's evidence that are relevant to why it cannot support the claim: - The paper is a single-case report of a pathogenic GLA variant; case reports provide evidence only about the gene(s) they study. - Even strong evidence for GLA causing Fabry disease cannot be generalized to other genes unless those genes are explicitly studied or mechanistically linked. - Because RPS6KB1 (ribosomal protein S6 kinase B1) is a different gene with different functions, absence of any mention in this manuscript means it neither supports nor refutes the claim. In summary, relevance is effectively zero because the study subject, genetic findings, and mechanistic discussion are entirely focused on GLA and Fabry disease and contain no information on RPS6KB1.


[Read Paper](https://www.semanticscholar.org/paper/a7c27f053151b6265c6c2b709b2f2085f8129f0e)


### De novo LAMP2 insertion mutation causes cardiac-only Danon disease: A case report

**Why Not Relevant**: The paper content exclusively addresses Danon disease and a pathogenic mutation in LAMP2 (c.764_765insGA), describing clinical presentation, family segregation (de novo), histopathology, decreased LAMP2 protein expression, and predicted structural effects on LAMP2. There is no mention of RPS6KB1, no genetic association data concerning RPS6KB1, and no mechanistic discussion that implicates RPS6KB1. Because the paper provides neither direct genetic/clinical evidence nor mechanistic links involving RPS6KB1, it does not support or refute the claim that RPS6KB1 is associated with a Mendelian disease. The study type (single-case report) and its focus on a different gene further limit any extrapolation to RPS6KB1.


[Read Paper](https://www.semanticscholar.org/paper/4c783f8dec24c49e2827ef0a67762ff5fd28c8ad)


### FRI387 Reproductive Capacity Of Znhit3 Knockout Mouse Model

**Why Not Relevant**: Relevance is effectively zero because the paper exclusively studies ZNHIT3 (TRIP-3). The text documents that ZNHIT3 is associated with Mendelian disorders (PEHO and MRKH) and describes generation and phenotyping of Znhit3 knockout mice. There is no mention of RPS6KB1, no experimental data concerning RPS6KB1, and no mechanistic links or interaction data connecting ZNHIT3 to RPS6KB1. Therefore this paper provides no direct or mechanistic evidence supporting or refuting the claim that RPS6KB1 is likely associated with a Mendelian disease.


[Read Paper](https://www.semanticscholar.org/paper/31dabd0957ffec78267540944decdfe71d088dec)


### Identification and in silico structural analysis for the first de novo mutation in the cystic fibrosis transmembrane conductance regulator protein in Iran: case report and developmental insight using microsatellite markers

**Why Not Relevant**: The paper exclusively studies cystic fibrosis and variants in the CFTR gene (including one de novo CFTR mutation). There is no mention of RPS6KB1, no genetic, clinical, or mechanistic data involving RPS6KB1, and no analysis that could be mapped to implicate RPS6KB1 in any Mendelian disease. Therefore this paper provides no direct or mechanistic evidence supporting or refuting the claim that RPS6KB1 is likely associated with at least one Mendelian disease. To evaluate that claim, sources explicitly addressing RPS6KB1 (e.g., OMIM, ClinVar, PubMed literature on RPS6KB1 pathogenic variants, GenCC, or functional studies of RPS6KB1) should be consulted.


[Read Paper](https://www.semanticscholar.org/paper/b16e327aff8b8bb2511958c4f9874b3460fa62b5)


### Tuberous sclerosis complex associated lymphangioleiomyomatosis caused by de novo mutation of TSC2 gene in Vietnam: A case report

**Why Not Relevant**: Relevance is effectively zero because the paper content supplied focuses exclusively on a case of pulmonary LAM associated with a de novo frameshift mutation in TSC2. The text contains no mention of the gene RPS6KB1, no data (genetic, clinical, or experimental) implicating RPS6KB1 in any Mendelian disorder, and no mechanistic discussion connecting RPS6KB1 to the disease described. While outside biological knowledge indicates that RPS6KB1 (S6K1) functions downstream of mTOR and that TSC1/TSC2 regulate mTOR signaling, the provided paper content does not state or provide evidence of that pathway or any involvement of RPS6KB1; therefore no direct or mechanistic evidence in this paper supports the claim that RPS6KB1 is likely associated with at least one Mendelian disease.


[Read Paper](https://www.semanticscholar.org/paper/6b7d5d519fb771d9bd7eb969a3b3847dcfdd8dad)


### Exome sequencing in over 63,000 patients in TIMI trials uncovers pathogenic cardiomyopathy variant carriers with high risk for heart failure and cardiovascular death

**Why Not Relevant**: The paper focuses specifically on identifying and analyzing pathogenic/likely pathogenic variants for clinically established cardiomyopathy phenotypes (DCM, HCM, ARVC) using phenotype- and gene-specific curation and ClinVar classifications. The text makes explicit reference to established cardiomyopathy genes (for example, TTN) and to variant carriers for those CMP categories. Nowhere in the provided content is RPS6KB1 mentioned, nor are there descriptions of mechanistic pathways or case/variant-level evidence linking RPS6KB1 to a Mendelian disease. Because the study's design, curation, and reported results are targeted to cardiomyopathy-associated genes and outcomes, the absence of RPS6KB1 in the paper means it offers no direct evidence (e.g., reported pathogenic variants, case series, segregation, statistical association) or mechanistic evidence (e.g., functional data implicating RPS6KB1 in monogenic disease) relevant to the claim. Limitations to this conclusion: absence of mention in the manuscript is not definitive proof that RPS6KB1 has no Mendelian associations — the underlying exome dataset could include RPS6KB1 variants that were not analyzed or reported because the study's pre-specified focus was cardiomyopathy genes. However, given the explicit gene- and phenotype-specific curation approach described, this paper cannot be used to support the claim that RPS6KB1 is likely associated with at least one Mendelian disease.


[Read Paper](https://www.semanticscholar.org/paper/f9e3adf19cb2bfdb8171744405eb1112137a3bb9)


### Descriptive analysis of MC4R gene variants associated with obesity listed on ClinVar

**Why Not Relevant**: The provided paper content exclusively concerns the MC4R gene: its ClinVar-identified variants, variant classification, mutation spectrum, and mechanistic implications for receptor activation and obesity-related phenotypes. The claim concerns RPS6KB1 and whether it is likely associated with at least one Mendelian disease. The text contains no mention of RPS6KB1, no data on variants in RPS6KB1, and no mechanistic discussion relevant to RPS6KB1. Therefore the paper provides no direct or indirect evidence that bears on the claim. Because all excerpts are specific to MC4R, they cannot support or refute any assertion about RPS6KB1; hence the relevance score is 0.0.


[Read Paper](https://www.semanticscholar.org/paper/6bae624c1c06216b5458a99ef3b5e0a4b2d4cd16)


### Novel Mouse Model That Recapitulates the Hematologic Phenotype of Diamond Blackfan Anemia

**Why Not Relevant**: The provided paper content focuses exclusively on Diamond-Blackfan anemia (DBA) and genetic and experimental work involving RPL11 (a ribosomal protein gene). There is no mention of the gene RPS6KB1 anywhere in the text, nor are there data (genetic, clinical, segregation, functional, or mechanistic) concerning RPS6KB1. Therefore the paper provides no direct evidence that RPS6KB1 is associated with any Mendelian disease. Mechanistic evidence in the paper pertains to RPL11 haploinsufficiency and DBA; those mechanisms (ribosomal protein deficiency leading to defective erythropoiesis) cannot be assumed to apply to RPS6KB1, which encodes a serine/threonine kinase (p70 S6 kinase) with distinct functions. Given the absence of any mention or data about RPS6KB1 in the provided content, the relevance of this paper to the claim that RPS6KB1 is likely associated with at least one Mendelian disease is effectively zero. If you want an assessment of RPS6KB1 specifically, please provide papers mentioning RPS6KB1 or I can search the literature/databases for evidence linking RPS6KB1 to Mendelian disease.


[Read Paper](https://www.semanticscholar.org/paper/9a5b100b5d60beb5063671656aba08b24f646b09)


## Search Queries Used

- RPS6KB1 de novo mutation Mendelian disease case report

- RPS6KB1 ClinVar OMIM pathogenic variant gene disease association

- S6K1 RPS6KB1 p70 S6 kinase mTOR pathway neurodevelopmental disorder overgrowth

- RPS6KB1 knockout mouse phenotype loss of function haploinsufficiency congenital syndrome model


## Bibliometric Configuration

- Use Bibliometrics: True

- Author Impact Weight: 0.4

- Citation Impact Weight: 0.4

- Venue Impact Weight: 0.2
