## Supplementary material for "MENDELSEEK: An algorithm that predicts Mendelian Genes and elucidates what makes them special": RXFP2.txt

# Claim: The gene RXFP2 is likely associated with at least one Mendelian disease.

**Status**: processed

**Overall Rating**: -1

**Explanation**:

No relevant papers were found that support or refute this claim.


## Relevant Papers


## Other Reviewed Papers


### Variants of Uncertain Significance and “Missing Pathogenicity”

**Why Not Relevant**: This paper text does not mention RXFP2 or present any data, results, or discussion about that gene. The manuscript excerpt focuses on analysis of variants in the ACMG 59-gene set (with emphasis on 30 cardiac genes), variant interpretation issues, and results from sequencing participants of the NUgene biobank. Because RXFP2 is not named and no variants, phenotypes, segregation, functional studies, or mechanistic discussion concerning RXFP2 are provided, there is no direct evidence in this paper to support or refute the claim that RXFP2 is likely associated with at least one Mendelian disease. The excerpt does contain general statements about challenges in variant interpretation, population frequencies, and the prevalence of predicted-deleterious variants exceeding disease prevalence; however, these are general contextual points about variant interpretation and do not provide mechanistic or gene-specific evidence for RXFP2. Therefore the paper is not relevant for assessing the claim about RXFP2. To evaluate the claim would require data not present here: explicit mention of RXFP2, evidence of rare, segregating variants in affected families, functional assays demonstrating a deleterious effect, or epidemiologic evidence linking RXFP2 variants to a Mendelian phenotype.


[Read Paper](https://www.semanticscholar.org/paper/a2df451199c905611c0053a2f7d8a6a4dfecc14b)


### PD48-03 HOW FAR SHOULD WE EXPLORE HYPOSPADIAS? NEXT-GENERATION SEQUENCING APPLIED TO A LARGE COHORT OF HYPOSPADIAC PATIENTS

**Why Not Relevant**: The provided paper content does not mention the gene RXFP2 or report pathogenic or likely pathogenic variants in RXFP2. The genetic findings reported are focused on other genes (e.g., AR, NR5A1, HSD17B3, MAMLD1, MID1, RARA) and an 18q deletion in a cohort of patients with hypospadias and related disorders. Although an NGS panel of 336 genes was used, the text does not list all genes tested, so absence of RXFP2 in the results could reflect that: (a) RXFP2 was not included in the panel, (b) RXFP2 was included but no variants of interest were found, or (c) RXFP2 variants were found but not reported because they were not judged pathogenic or relevant to the study phenotypes. Therefore, this paper provides no direct evidence supporting an association between RXFP2 and any Mendelian disease; at best it offers limited indirect negative evidence (it does not report RXFP2 among discovered pathogenic genes). Important caveats: (1) absence of mention in this paper is not evidence that RXFP2 is not associated with Mendelian disease in other contexts or cohorts, and (2) the study populations (hypospadias/DSD and other pediatric surgical cohorts) and the panel's focus on exonic variants limit the generalizability of negative findings to all Mendelian phenotypes or variant types.


[Read Paper](https://www.semanticscholar.org/paper/b4d09d5fc960bf77654042a156ee66c8b6327617)


### Abstract LB-372: Robust targeted sequencing strategy identifies highly penetrant mutations in hereditary colorectal cancer cases from Colon Cancer Family Registry

**Why Not Relevant**: The paper is a targeted sequencing study of hereditary colorectal cancer (CRC) that screened 1,046 CRC cases and 1,006 controls for mutations in 40 known or candidate CRC genes and reports deleterious variants in a list of CRC-relevant genes (ABCC1, APC, AXIN2, BMPR1A, CHEK2, CTNNB1, ENG, MUTYH, POLD1, POLE, PTCH1, SH2B3, TGFBR2, TP53, etc.). The abstract contains no mention of RXFP2, no data about variants in RXFP2, and no mechanistic discussion linking RXFP2 to CRC or any other Mendelian disease within this study. Because the study's scope is clearly CRC-focused and the reported results enumerate CRC-related genes while omitting RXFP2, there is no direct or mechanistic evidence here to support the claim that "RXFP2 is likely associated with at least one Mendelian disease." Absence of mention in this abstract does not prove RXFP2 has no Mendelian associations (absence of evidence is not evidence of absence), but for the purposes of evaluating this paper's relevance to the RXFP2 claim, the content provides essentially no supporting information. Therefore relevance is judged very low (<0.1).


[Read Paper](https://www.semanticscholar.org/paper/50c937f4f1eb7ceef5965bb0c11a3b8dae5f3da4)


### University of Southern Denmark Clinician's guide to genes associated with Rett-like phenotypes Investigation of a Danish cohort and review of the literature

**Why Not Relevant**: The paper is a focused review and cohort study of genes associated with Rett syndrome, atypical RTT, and Rett-like phenotypes and reports genes and variants identified in a Danish Rett-like cohort. RXFP2 is not mentioned anywhere in the provided text, nor are any mechanistic data, variant observations, or disease associations for RXFP2 presented. Therefore the paper provides no direct evidence supporting or refuting the claim that RXFP2 is associated with a Mendelian disease. The absence of RXFP2 from gene lists and cohort findings in this Rett-focused study is only weak indirect evidence (limited negative evidence) because: (1) the study is restricted to Rett-like phenotypes and so would not necessarily include genes associated with other Mendelian conditions; (2) absence of reporting does not equal evidence of absence; and (3) the cohort size and the literature surveyed are not comprehensive for all possible Mendelian gene–disease associations. Given these limitations, the paper is not relevant to establishing whether RXFP2 is likely associated with at least one Mendelian disease.


[Read Paper](https://www.semanticscholar.org/paper/0026a99b45e2786572daf5995d7fb96806295eb4)


### Syddansk Universitet Clinician ' s guide to genes associated with Rett-like phenotypes-Investigation of a Danish cohort and review of the literature

**Why Not Relevant**: The provided paper is a review of Rett syndrome / Rett-like disorders and reports findings from a Danish cohort of 35 Rett-like patients. It lists many genes known or newly implicated in Rett-like phenotypes (MECP2, CDKL5, FOXG1, SMC1A, STXBP1, SCN2A, TCF4, KCNB1, etc.) but does not mention RXFP2 anywhere in the supplied text. Therefore the paper provides no direct evidence that RXFP2 is associated with any Mendelian disease, nor does it discuss mechanisms that would plausibly link RXFP2 to a Mendelian phenotype. The absence of RXFP2 in the gene lists and cohort findings is negative evidence only within the scope of Rett-like disorders; it is not definitive proof against any Mendelian association outside the paper's scope. Because the paper neither reports RXFP2 variants nor discusses functional/mechanistic data for RXFP2, its relevance to the claim that "RXFP2 is likely associated with at least one Mendelian disease" is negligible.


[Read Paper](https://www.semanticscholar.org/paper/d14fab69ce67e0cba07e6800afa7e5096cd37492)


## Search Queries Used

- RXFP2 relaxin family peptide receptor 2 mutation familial cryptorchidism undescended testis monogenic Mendelian segregation pathogenic variant

- RXFP2 variant pathogenicity ClinVar case report cohort study next generation sequencing association study rare variant congenital undescended testis

- INSL3 RXFP2 signaling testicular descent gubernaculum Leydig cells knockout mouse model functional assay human loss of function mutation

- RXFP2 relaxin family peptide receptor 2 OMIM ClinVar Mendelian disease review systematic review meta analysis hereditary monogenic disease


## Bibliometric Configuration

- Use Bibliometrics: True

- Author Impact Weight: 0.4

- Citation Impact Weight: 0.4

- Venue Impact Weight: 0.2
