## Supplementary material for "MENDELSEEK: An algorithm that predicts Mendelian Genes and elucidates what makes them special": SEH1L.txt

# Claim: The gene SEH1L is likely associated with at least one Mendelian disease.

**Status**: processed

**Overall Rating**: 0

**Explanation**:

Supporting evidence.
The provided paper excerpts give general context about gene–disease discovery: as of early 2019 only ~18% of protein-coding genes had been linked to Mendelian disorders (OMIM), and exome sequencing can reveal pathogenic variants in genes not previously associated with disease. These points support the general plausibility that many genes not yet associated with disease, including SEH1L, might ultimately be found to cause Mendelian disorders. In other words, the background literature cited makes it reasonable to consider uncharacterized genes as potential disease genes.

Caveats or contradictory evidence.
Crucially, none of the provided excerpts (or the paper title/metadata) mention SEH1L specifically or provide any empirical association between SEH1L and a Mendelian phenotype. The excerpts emphasize the challenges of variant interpretation and that discovery is ongoing, but they do not provide evidence for SEH1L being linked to any monogenic disease. Absence of mention in the provided material is not definitive proof of absence, but it means no direct supportive data are present in the supplied evidence.

Analysis of potential underlying mechanisms, deeper relationships, or second-order implications.
From a biological perspective (drawing on general knowledge), SEH1L encodes a component of the nuclear pore complex (Seh1-like protein) involved in nucleocytoplasmic transport and possibly other cellular processes. Genes encoding core components of essential cellular machineries can present in different ways clinically: loss-of-function may be embryonically lethal (preventing observation of a viable Mendelian syndrome), or hypomorphic/allelic variants may cause recessive or dominant disease. Thus, biologically it is plausible that pathogenic variants in SEH1L could cause disease, but plausible mechanism does not substitute for empirical association data. The supplied paper excerpts do not provide functional studies, case reports, database entries, or segregation evidence linking SEH1L to disease.

Which rating is most appropriate.
Given the evidence provided, there is no direct support that SEH1L is associated with any Mendelian disease. The background statements only indicate that many genes remain to be linked to disease and that exome sequencing can uncover such links; these are general and do not specifically support SEH1L. Therefore the correct assessment based on the supplied evidence is that there is no evidence to support the claim that SEH1L is likely associated with at least one Mendelian disease. The most appropriate rating is "No Evidence."


**Final Reasoning**:

I re-read the logic: the claim is a specific assertion about SEH1L. The provided paper content is general background on the state of gene–disease discovery and does not mention SEH1L or supply any empirical association, case, database entry, or functional/segregation data for SEH1L. While biological plausibility exists in the abstract (many genes could turn out to be disease genes; SEH1L has cellular roles that could in principle be disease-relevant), that plausibility is not evidence of an association. The criteria for ratings require evidence supporting or refuting the claim. Since none of the supplied excerpts provide specific evidence linking SEH1L to a Mendelian disorder, the proper assignment is "No Evidence." I find no flaw in this reasoning and therefore reaffirm the rating.


## Relevant Papers


### Precision Medicine in Internal Medicine

**Authors**: Krzysztof Kiryluk (H-index: 0), W. Chung (H-index: 97)

**Relevance**: 0.15

**Bibliometric Impact**: 0.5692


**Excerpts**:

- As of 1 February 2019, the Online Mendelian Inheritance in Man database indicated that out of 20000 protein-coding genes, only 3652 (approximately 18%) had been found to cause known single-gene disorders. The process of establishing associations of the other genes with diseases remains laborious and continues to evolve (6).

- Because ES involves sequence analysis of all genes in the genome, it can identify mutations in genes that are not suspected on the basis of clinical presentation or are not yet known to cause disease. When initial analysis is unable to establish a diagnosis, ES data can be reinterrogated as new genes for a given condition are discovered and new exome analysis methods are developed.

- Figure 1 summarizes the approximate numbers of genetic variants found in a typical human exome. On average, ES detects approximately 10000 protein-altering variants, including 150 to 180 protein-truncating variants, 20 to 30 known disease-causing (mainly recessive) variants, and 1 to 4 de novo variants that are not present in parents.

- The key challenge relates to interpretation of genetic variants.

- Interpretation of genetic variation detected by sequencing is challenging but continues to improve with data sharing and accelerated gene discovery for monogenic diseases.


**Explanations**:

- Relation to claim: Indirect, quantitative context. This excerpt states that only ~18% of protein-coding genes were, as of the cited date, known to cause single-gene (Mendelian) disorders and explicitly notes that associating the remaining genes with disease is a slow ongoing process. Strength: provides a data-driven prior about how many genes are currently linked to Mendelian disease, which bears on the plausibility that a given gene (SEH1L) is already known to be associated with a Mendelian disease. Limitations: no mention of SEH1L specifically, the statistic is dated (1 Feb 2019) and aggregate — it does not provide evidence for or against SEH1L itself and cannot substitute for gene-specific data.

- Relation to claim: Mechanistic / methodological relevance. Explains that exome sequencing can discover disease-causing mutations in genes not previously associated with disease, and that reanalysis over time can reveal new gene–disease associations. Strength: supports plausibility that an association for SEH1L could be discovered (or already discovered but not yet recognized) via ES and reanalysis. Limitations: still generic — it does not report any ES findings implicating SEH1L, nor does it provide mechanistic functional data specific to SEH1L.

- Relation to claim: Indirect evidence about signal-to-noise and discovery burden. The numbers show many protein-altering variants are typically found per exome, including protein-truncating and de novo variants, which contextualizes the frequency of potentially pathogenic variants that need interpretation. Strength: demonstrates why detecting a variant in SEH1L in a patient is not, by itself, strong proof of causality without additional evidence (segregation, functional data). Limitations: not specific to SEH1L and does not state how often SEH1L variants occur or whether any are pathogenic.

- Relation to claim: Methodological caveat. Emphasizes that the main obstacle to linking genes (including SEH1L) to Mendelian disorders is interpretation of variants. Strength: underscores need for robust criteria (segregation, functional evidence) before claiming a gene–disease association. Limitations: again, generic — no SEH1L-specific interpretation is provided.

- Relation to claim: General statement about progress. Notes that interpretation is improving with data sharing and accelerated gene discovery, implying that gene–disease links (potentially including SEH1L) may be identified over time. Strength: supports the idea that lack of current evidence does not preclude future identification. Limitations: non-specific and not evidence that SEH1L is or is not associated with a Mendelian disease now.


[Read Paper](https://www.semanticscholar.org/paper/c7199e8a5226f8d9d55777da0e15e91797bb459a)


## Other Reviewed Papers


### Trio-based whole exome sequencing in patients with suspected sporadic inborn errors of immunity: A retrospective cohort study

**Why Not Relevant**: The paper is a study of de novo variants in a cohort of 123 patients with suspected inborn errors of immunity and reports candidate DNVs in immune-related genes (NLRP3, RELA, PSMB10, DDX1, KMT2C, FBXW11). The gene SEH1L is not mentioned anywhere in the provided paper content. Therefore this paper provides no direct genetic, mechanistic, or functional evidence supporting or refuting the claim that SEH1L is likely associated with at least one Mendelian disease. The absence of SEH1L in the reported findings may reflect the study's phenotype focus (IEI), cohort composition, and variant filtering/interpretation pipeline; it does not constitute evidence that SEH1L is not disease-associated in other disorders or datasets. Given the complete lack of mention or data on SEH1L in this paper, relevance to the claim is effectively zero.


[Read Paper](https://www.semanticscholar.org/paper/cdde2232daad22b7d92ec15faa38b9776f1e4c15)


### Novel variant of KIF11 associated with MCLMR syndrome

**Why Not Relevant**: This paper describes a novel heterozygous frameshift variant in the KIF11 gene causing a Mendelian syndrome (microcephaly with or without chorioretinopathy, lymphedema, or mental retardation / MLCRD). Nowhere in the provided content is the gene SEH1L mentioned, discussed, or linked to any phenotype. All genetic, clinical, and mechanistic statements refer to KIF11 (for example, the paper states: “We found that our patient carries a novel heterozygous KIF11 frameshift variant c.1858_1859delGT (p.Val620Thrfs *6) resulting in a premature stop codon…” and discusses loss-of-function as the probable disease mechanism). Because the claim concerns SEH1L specifically, and this paper contains no direct evidence (no mention of SEH1L sequence, variation, functional data, clinical segregation, or disease association) nor any mechanistic data that would plausibly connect SEH1L to the described phenotype, it provides no support for or against the claim that SEH1L is associated with a Mendelian disease. Limitations: the paper is a single-case report focused on KIF11; its findings demonstrate how to evaluate a candidate Mendelian gene (de novo truncating variant, absence from population databases, consistent phenotype), but those methods and conclusions cannot be transferred to SEH1L in the absence of any data about SEH1L in this manuscript. To assess the claim about SEH1L, literature or database searches (OMIM, ClinVar, HGMD, primary gene-disease studies) that specifically address SEH1L would be required.


[Read Paper](https://www.semanticscholar.org/paper/598b1c7508865997c295d53a98d38407fbecde29)


### HOXA11 is another monogenic cause of congenital anomalies of the kidney and urinary tract

**Why Not Relevant**: The provided paper content discusses HOXA11 and its possible role as a monogenic cause of CAKUT (congenital anomalies of the kidney and urinary tract). The exact sentence supplied is: "Adding HOXA11 to the list of monogenic causes of CAKUT may hopefully encourage physicians and researchers to look for HOxA11 variants in selected cases for prompt genetic diagnosis and genetic counselling, as well as to study the molecular pathogenesis of HOX a11-related disease in humans withCAKUT." This text does not mention SEH1L, presents no data, no genetic variant reports, no segregation or population evidence, and no mechanistic information about SEH1L. Therefore it provides neither direct evidence (e.g., patient genotypes, statistical associations, reported pathogenic variants) nor mechanistic evidence (e.g., functional studies, pathways implicating SEH1L in disease) relevant to the claim that SEH1L is likely associated with at least one Mendelian disease. Additional reasons for irrelevance and limitations of the provided text: 1) Different gene: HOXA11 is a distinct gene from SEH1L; statements about one gene do not support claims about the other. 2) Speculative language: the sentence is an encouragement to look for HOXA11 variants rather than a report of findings, so it contains no empirical evidence. 3) Lack of methodological or result detail: no cohorts, no variants, no segregation, no functional assays, and no statistical measures are provided. To evaluate the claim about SEH1L you would need papers that explicitly discuss SEH1L variants in affected individuals, segregation with disease, functional/mechanistic data linking SEH1L perturbation to a phenotype, or entries in curated resources (e.g., OMIM, ClinVar) that annotate SEH1L as disease-associated. Because none of that is present in the supplied content, relevance to the SEH1L–Mendelian disease claim is effectively zero.


[Read Paper](https://www.semanticscholar.org/paper/db3d83a93884ccd5d66fca19ab8728ccabfc9ffc)


### Case report: A novel FBXW7 gene variant causes global developmental delay

**Why Not Relevant**: The paper exclusively investigates a neurodevelopmental disorder caused by a pathogenic de novo variant in FBXW7. There is no mention of the gene SEH1L, no reported variants in SEH1L, and no mechanistic discussion linking FBXW7 to SEH1L. Although trio whole-exome sequencing (noted in Methods) could in principle detect exonic variants in many genes including SEH1L, the authors report only the FBXW7 finding; absence of reported SEH1L data in this paper means it provides no evidence for or against an association between SEH1L and any Mendelian disease. Therefore the paper is not relevant to the claim. Caveat: absence of evidence in this paper is not evidence of absence overall — this single study neither supports nor disproves SEH1L involvement in Mendelian disease, it simply does not address SEH1L at all.


[Read Paper](https://www.semanticscholar.org/paper/bb0b7ce4ae459771c428a6c9d1dbc93aecc6bcdc)


### Autosomal Dominant Intellectual Development Disorder-6 (MRD6) Without Seizures Linked to a De Novo Mutation in the grin2b Gene Revealed by Exome Sequencing: A Case Report of a Moroccan Child

**Why Not Relevant**: The paper's content exclusively describes a pathogenic GRIN2B (grin2b) nonsense variant (c.3912C>G, p.Tyr1304Ter) causing autosomal dominant intellectual development disorder-6 (MRD6). It contains no mention of the gene SEH1L, no reported SEH1L variants, no segregation or case reports involving SEH1L, and no mechanistic or pathway analyses linking SEH1L to GRIN2B or to MRD6. Therefore there is no direct evidence in this paper that SEH1L is associated with any Mendelian disease. 

Why this is insufficient to support the claim: Evidence that a different gene (GRIN2B) causes a Mendelian disorder cannot be extrapolated to assert that an unrelated gene (SEH1L) is disease-associated. Supporting the claim that SEH1L is likely associated with a Mendelian disease would require observations such as: (a) pathogenic or likely pathogenic SEH1L variants segregating with disease in families or arising de novo in affected individuals, (b) population frequency data showing rarity consistent with pathogenicity, (c) functional studies showing deleterious impact of SEH1L variants on gene/protein function relevant to disease mechanism, or (d) replication in independent cohorts and/or database entries (OMIM, ClinVar) implicating SEH1L. None of these are present in this paper. 

Conclusion: Relevance is 0.0 because the paper provides no direct or mechanistic evidence about SEH1L. To evaluate the claim, a dedicated search of literature and variant databases (OMIM, ClinVar, HGMD, PubMed) for SEH1L-specific reports is required.


[Read Paper](https://www.semanticscholar.org/paper/67d756e0cc302e5d46fdda9df34403f1f2de9a5a)


### Confirmation of risk of cancer in blepharocheilodontic syndrome

**Why Not Relevant**: The provided paper content does not mention SEH1L or any data about that gene. The single sentence discusses variants causing BCDS1 (blepharocheilodontic syndrome 1) and their localization: "In the light of both studies and the description of 26 patients from 18 families, variants causing BCDS1 were likely to localize in mutational hotspots, corresponding to E-cadherin calcium-binding sites in the EC1-EC2 linker domain and to exon 9–intron 9 junction." This sentence implicates E-cadherin (the CDH1 gene) mutational hotspots as relevant to BCDS1, not SEH1L. Therefore the content provides no direct evidence (no patient data, variants, statistical associations) nor mechanistic evidence linking SEH1L to any Mendelian disease. Limitations: absence of any mention of SEH1L in this excerpt does not prove SEH1L has no Mendelian association, but based on the provided text alone there is nothing to support or refute the claim about SEH1L. Also, the excerpt is limited to a single sentence focusing on CDH1-related mutational hotspots from 26 patients in 18 families, so it is narrowly focused and not generalizable to genes outside that study.


[Read Paper](https://www.semanticscholar.org/paper/9ae4a26ce01b68b3153fcb48cb63fd4860bb3938)


### Case Report: A novel de novo variant of COL1A1 in fetal genetic osteogenesis imperfecta

**Why Not Relevant**: The paper exclusively studies a de novo COL1A1 variant as the likely cause of osteogenesis imperfecta in a single fetal case. There is no mention of the gene SEH1L, no data (genetic, functional, epidemiologic) concerning SEH1L, and no discussion that would allow inference that SEH1L is associated with any Mendelian disease. All mechanistic content refers to collagen type I and Gly-X-Y triple helix disruption, which is unrelated to SEH1L unless an independent functional link is established; none is provided here. Because the paper's focus and evidence concern a different gene (COL1A1) and disease mechanism, it provides effectively no relevant evidence for the claim that SEH1L is likely associated with at least one Mendelian disease.


[Read Paper](https://www.semanticscholar.org/paper/1caeaf6717a5af526be5f1f0dcef41ef4c377699)


## Search Queries Used

- SEH1L pathogenic variant Mendelian disease case report patient family segregation de novo ClinVar OMIM

- exome sequencing rare disease gene discovery SEH1L de novo variant trio Mendelian cohort novel gene

- SEH1L Seh1 like nucleoporin Y complex nuclear pore complex knockout mouse zebrafish knockdown loss of function phenotype development

- gnomAD SEH1L loss of function variants pLI LOEUF constraint loss of function intolerance ClinGen haploinsufficiency dosage sensitivity population frequency


## Usage Statistics

- Prompt Tokens: 0

- Completion Tokens: 0

- Total Tokens: 0

- Estimated Cost: $0.0370


## Bibliometric Configuration

- Use Bibliometrics: True

- Author Impact Weight: 0.4

- Citation Impact Weight: 0.4

- Venue Impact Weight: 0.2
