## Supplementary material for "MENDELSEEK: An algorithm that predicts Mendelian Genes and elucidates what makes them special": SEMA5A.txt

# Claim: The gene SEMA5A is likely associated with at least one Mendelian disease.

**Status**: processed

**Overall Rating**: -1

**Explanation**:

No relevant papers were found that support or refute this claim.


## Relevant Papers


## Other Reviewed Papers


### Biallelic variants identified in 36 Pakistani families and trios with autism spectrum disorder

**Why Not Relevant**: The single provided sentence, "Information on the genetic variability associated with ASD in Pakistan is offered, and a marked enrichment for biallelic variants over that reported in outbreeding populations is demonstrated.", does not mention SEMA5A, does not report any specific genes or variants, and gives no mechanistic or clinical data linking SEMA5A to a Mendelian disease. Therefore the provided content contains no direct evidence (no variant, no segregation, no case report, no statistical association) and no mechanistic evidence (no functional studies, no pathway discussion) relevant to the claim that SEMA5A is likely associated with at least one Mendelian disease. At best, the sentence makes a general point that biallelic (recessive) variants are enriched in this population, which is a population-genetic observation that increases the prior probability that recessive Mendelian disorders might be discovered in consanguineous populations; this is an indirect and nonspecific point and cannot be used to implicate any particular gene such as SEMA5A. Limitations: only a single summary sentence was provided rather than the full paper text, figures, tables, or supplementary data; the full paper might include gene-level results or tables that were not supplied here. To evaluate the claim properly one would need the paper's gene/variant lists, segregation data, statistical tests, and any functional follow-up, or to consult clinical databases (OMIM, ClinVar, HGMD) and the primary literature for reported Mendelian associations for SEMA5A. Because none of that information is present in the supplied content, relevance to the claim is essentially zero.


[Read Paper](https://www.semanticscholar.org/paper/17be98b475c43e8e296ed0d13d54250c91e9c42d)


### Case report: A novel FBXW7 gene variant causes global developmental delay

**Why Not Relevant**: The paper reports a single case study of a neurodevelopmental disorder caused by a pathogenic de novo variant in FBXW7 and does not mention SEMA5A at all. There is no experimental, genetic, clinical, or mechanistic data about SEMA5A in the provided content. Therefore this paper provides no direct evidence that SEMA5A is associated with any Mendelian disease, nor does it provide mechanistic evidence that would make such an association more or less plausible. 

Notes on limitations and interpretation: (1) Absence of SEMA5A in this FBXW7-focused case report does not disprove any existing or potential gene-disease association for SEMA5A; it is simply irrelevant evidence. (2) To evaluate the claim about SEMA5A one would need papers reporting pathogenic variants in SEMA5A segregating with a Mendelian phenotype, curated entries in OMIM/ClinVar, case series, functional assays, or genetic evidence from trio/segregation studies — none of which are present here. (3) This paper is a single-case report (trio-WES with Sanger confirmation) about FBXW7, so it cannot be used to infer associations for other genes. 

Recommended next steps to assess the claim: search gene-disease resources (OMIM, ClinVar), PubMed for SEMA5A case reports, and population/constraint metrics (gnomAD) and functional studies involving SEMA5A. 


[Read Paper](https://www.semanticscholar.org/paper/bb0b7ce4ae459771c428a6c9d1dbc93aecc6bcdc)


### Germline mosaicism in a collagen VI‐related myopathy family: A cause of autosomal recessive inheritance

**Why Not Relevant**: The provided paper content focuses entirely on germline mosaicism of COL6A1 and collagen VI-related myopathy; SEMA5A is not mentioned anywhere in the quoted text. Therefore the paper supplies no direct genetic, clinical, or mechanistic evidence that SEMA5A is associated with any Mendelian disease. Absence of mention in this single paper is not evidence that SEMA5A is not disease-associated generally, but it means this paper cannot be used to support the claim. To assess SEMA5A's association with Mendelian disease would require literature or database evidence (e.g., case reports, ClinVar, OMIM, gene-disease curation) specifically mentioning SEMA5A; that evidence is not present in the supplied text.


[Read Paper](https://www.semanticscholar.org/paper/4572f635bb354974678a75579201d9eedc2c40ea)


### Whole exome sequencing identifies variants in the \(\textit{TNNI3}\) gene in vietnamese patient with restrictive cardiomyopathy - a case report

**Why Not Relevant**: The paper exclusively reports on restrictive cardiomyopathy and identifies pathogenic variants in TNNI3 in the studied patient. It does not mention SEMA5A anywhere, nor does it list the 22 genes referred to, so there is no direct or mechanistic evidence in this text linking SEMA5A to any Mendelian disease. While the paper notes that multiple genes can cause RCM and demonstrates that WES can identify pathogenic variants, absence of any explicit reference to SEMA5A (or data implicating it) means this document cannot be used to support the claim that SEMA5A is likely associated with at least one Mendelian disease. Additional sources that specifically name SEMA5A and provide genetic or mechanistic evidence would be required to evaluate that claim.


[Read Paper](https://www.semanticscholar.org/paper/83a11d71a34fe3ae5618faf4b4a593ecd29e0884)


### P672 A tertiary multicenter cohort of patients with chronic intestinal pseudo-obstruction and Crohn’s disease: a rare association with a high prevalence of monogenic disorders

**Why Not Relevant**: The paper does not mention SEMA5A anywhere in the provided content. It focuses on a small, specific cohort (n=8) of patients with chronic intestinal pseudo-obstruction co-occurring with Crohn's disease, using a targeted panel of 129 predefined genes for monogenic enteropathies. The paper reports pathogenic findings in ACTG2, STAT1 (gain-of-function), and TYMP, but neither lists SEMA5A among examined or discovered genes nor provides mechanistic data relevant to SEMA5A. Critical limitations preventing relevance: (1) no explicit inclusion of SEMA5A in the gene panel is stated, so absence of a finding cannot be interpreted as evidence against SEMA5A involvement; (2) small, highly selected cohort lacks power and generalizability for assessing whether SEMA5A is associated with any Mendelian disease; (3) the study addresses monogenic enteropathies in a specific clinical presentation rather than performing a broad discovery or functional analysis that could implicate SEMA5A. For these reasons the paper provides effectively no direct or mechanistic evidence for or against the claim that SEMA5A is likely associated with at least one Mendelian disease.


[Read Paper](https://www.semanticscholar.org/paper/e4c28f408ab8fa5cc40463e76b61dad7e8f0f6ec)


### Autosomal Dominant Intellectual Development Disorder-6 (MRD6) Without Seizures Linked to a De Novo Mutation in the grin2b Gene Revealed by Exome Sequencing: A Case Report of a Moroccan Child

**Why Not Relevant**: The paper's content exclusively describes a Mendelian disorder (MRD6) caused by a GRIN2B (grin2b) mutation in a single patient and presents molecular validation and pathogenic classification for that GRIN2B variant. Nowhere in the provided text is the gene SEMA5A mentioned, nor are any data, analyses, functional assays, variant calls, population statistics, or mechanistic pathways involving SEMA5A reported. Therefore the paper provides no direct evidence (variant-disease co-segregation, case reports, statistical enrichment) nor mechanistic evidence (biochemical function, pathway perturbation, model organism phenotypes) linking SEMA5A to any Mendelian disease. Because the study is focused on a different gene (GRIN2B) and contains no discussion of SEMA5A, it cannot be used to support the claim that SEMA5A is likely associated with at least one Mendelian disease. Additional reasons limiting relevance: the paper is a single-case report about GRIN2B (limited generalizability), contains no broader gene-level association analyses, and does not present negative or comparative data that would implicate SEMA5A. In short, absence of any mention or data concerning SEMA5A makes the paper irrelevant to the claim.


[Read Paper](https://www.semanticscholar.org/paper/67d756e0cc302e5d46fdda9df34403f1f2de9a5a)


### Trio-based genome sequencing identifies candidate causal genes in lifelong premature ejaculation

**Why Not Relevant**: The provided paper excerpt does not mention SEMA5A anywhere in the text. All quoted sentences refer to other genes (ZNF717, CTDSP2, FAT4) or to the overall burden of mutated genes found in eight trios of lifelong premature ejaculation patients. Therefore the excerpt contains no direct or mechanistic evidence for or against the claim that SEMA5A is likely associated with at least one Mendelian disease. Additional possibilities remain: SEMA5A could appear in parts of the paper not provided (e.g., Table 1 or supplementary materials), or it could be implicated by other studies not included here. Moreover, the study is small (eight trios), focused on lifelong PE, and not designed as a comprehensive screen for genes associated with Mendelian diseases broadly. For these reasons I assign a very low relevance (0.02) rather than zero, reflecting that the excerpted content is essentially non-informative regarding SEMA5A.


[Read Paper](https://www.semanticscholar.org/paper/213b15e38e05ff12c8444735db0324f71381fd39)


## Search Queries Used

- SEMA5A pathogenic variant mutation Mendelian disease case report exome sequencing ClinVar OMIM

- SEMA5A gene association monogenic inherited rare disease autosomal dominant autosomal recessive de novo trio sequencing cohort study

- SEMA5A semaphorin 5A axon guidance neurodevelopmental disorder autism intellectual disability epilepsy knockout mouse zebrafish functional study loss of function

- SEMA5A gene disease validity ClinGen gene curation systematic review semaphorin family SEMA5A SEMA3A SEMA4D constraint metrics gnomAD pLI


## Bibliometric Configuration

- Use Bibliometrics: True

- Author Impact Weight: 0.4

- Citation Impact Weight: 0.4

- Venue Impact Weight: 0.2
