## Supplementary material for "MENDELSEEK: An algorithm that predicts Mendelian Genes and elucidates what makes them special": SERPINE2.txt

# Claim: The gene SERPINE2 is likely associated with at least one Mendelian disease.

**Status**: processed

**Overall Rating**: -1

**Explanation**:

No relevant papers were found that support or refute this claim.


## Relevant Papers


## Other Reviewed Papers


### The Burden of Candidate Pathogenic Variants for Kidney and Genitourinary Disorders Emerging From Exome Sequencing

**Why Not Relevant**: The provided paper content describes methods for identifying and curating genes associated with Mendelian kidney and genitourinary disorders and details cohort and variant-filtering approaches, but it does not mention the gene SERPINE2 by name nor present gene-level results that would indicate whether SERPINE2 is associated with a Mendelian disease. The excerpts show that a curated list of 625 genes exists and that it is available on an external laboratory website or in appendices, but without access to that list or to gene-specific results in the supplied text, no direct or mechanistic evidence concerning SERPINE2 can be extracted. To evaluate the claim that SERPINE2 is likely associated with at least one Mendelian disease would require either (a) the curated gene list or supplementary tables from this paper showing SERPINE2 is included and evidence supporting that inclusion, or (b) independent gene-specific evidence from OMIM, ClinVar, HGMD, peer-reviewed gene-disease curation resources (e.g., ClinGen), or case reports. Because none of that is present in the provided content, relevance to the specific claim is very low.


[Read Paper](https://www.semanticscholar.org/paper/a75e297208c04992568e607d19aa9b9aaebd09d5)


### Explorer Quantifying single nucleotide variant detection sensitivity in exome sequencing

**Why Not Relevant**: The paper is a methodological study on SNV detection sensitivity in targeted (exome) sequencing and does not present any gene-specific genetic, clinical, segregation, population, or functional data concerning SERPINE2. Although the methods and conclusions are relevant in a general sense (they describe how well exome sequencing can detect variants and therefore bear on the ability to discover Mendelian associations), there are no direct observations, analyses, or claims about SERPINE2 or its involvement in any Mendelian disease. Therefore this paper provides no direct or gene‑specific mechanistic evidence to support or refute the claim that SERPINE2 is likely associated with at least one Mendelian disease. Limitations: absence of gene-level variant data, no phenotype correlation, no case reports, no linkage/segregation analyses, and no functional assays related to SERPINE2.


[Read Paper](https://www.semanticscholar.org/paper/3972fcdacd0349501312b50f641e2babb5f9337a)


### Diagnostic clarity of exome sequencing following negative comprehensive panel testing in the neonatal intensive care unit

**Why Not Relevant**: The provided paper does not mention the gene SERPINE2 anywhere in the text. Therefore it contains no direct evidence that SERPINE2 is associated with any Mendelian disease. The excerpts included are relevant only insofar as they provide general, indirect, mechanistic context: the study demonstrates that exome sequencing can identify Mendelian disease genes not present on prior panels and shows how variant classification (e.g., ACMG criteria) is applied at the gene/variant level. However, these do not constitute evidence for SERPINE2 specifically. Because establishing that a particular gene is 'likely associated with at least one Mendelian disease' requires gene-specific observations (e.g., pathogenic variants in multiple unrelated affected individuals, segregation data, functional validation, or strong statistical/clinical associations), the absence of any mention or data regarding SERPINE2 in this paper means it neither supports nor refutes the claim. To assess the claim one would need papers or database entries that specifically report SERPINE2 variants linked to Mendelian phenotypes, segregation evidence, functional studies, or curated gene-disease assertions (e.g., ClinGen, OMIM) — none of which are provided here.


[Read Paper](https://www.semanticscholar.org/paper/62096c8bc6e246796ed40c8a0de20b582381c61f)


### Whole-Exome Sequencing in Adults With Chronic Kidney Disease

**Why Not Relevant**: The provided paper content does not mention the gene SERPINE2 (or any variants in SERPINE2) and contains no results, sentences, or paragraphs that link SERPINE2 to a Mendelian disease. The excerpt is principally background and methods describing whole-exome sequencing in adults with chronic kidney disease, the construction of a 287-gene nephropathy OMIM list, sequencing and variant-calling methods, and variant-interpretation pipelines. Although the study describes screening OMIM genes and returning actionable findings, no gene-level results or mechanistic discussion involving SERPINE2 are present in the supplied text. Therefore, there is no direct evidence (no reported SERPINE2 variants, segregation, pathogenicity assessments, or clinical cases) and no mechanistic evidence (no functional data, pathways, or biological rationale regarding SERPINE2) in the excerpt to support or refute the claim that SERPINE2 is likely associated with at least one Mendelian disease. 

Limitations and caveats: (1) This assessment is limited to the supplied excerpt only. The full paper (including supplements and result sections not provided here) could potentially contain mentions of SERPINE2; that content was not provided for review. (2) The study design (WES of nephropathy patients and a nephropathy OMIM gene list) means many genes were screened, so absence of evidence in this excerpt does not prove SERPINE2 is not implicated elsewhere. Based solely on the provided text, however, there is no relevant evidence for the claim.


[Read Paper](https://www.semanticscholar.org/paper/a29bd57964f68ac5f8e1abcafe1112ba488f53f5)


### Biallelic variants identified in 36 Pakistani families and trios with autism spectrum disorder

**Why Not Relevant**: The supplied paper content is a single summary sentence about autism spectrum disorder genetics in Pakistan and an enrichment for biallelic variants. It does not mention SERPINE2, any specific genes, any variants, Mendelian disease assignments, functional studies, or mechanistic links. While the reported enrichment for biallelic variants might generally increase the likelihood of identifying recessive Mendelian disease genes in that population, this is an entirely non-specific observation and cannot be used to support or refute the claim that SERPINE2 is likely associated with at least one Mendelian disease. Additionally, the excerpt lacks methodological details, results, or gene lists that might reveal whether SERPINE2 was studied or implicated. To assess the claim about SERPINE2, the full text (or at least the gene/variant tables, supplementary materials, or specific gene mentions) would be required; based on the provided content alone, relevance to the SERPINE2 claim is negligible.


[Read Paper](https://www.semanticscholar.org/paper/17be98b475c43e8e296ed0d13d54250c91e9c42d)


### Novel variants of NEK9 associated with neonatal arthrogryposis: Two case reports and a literature review

**Why Not Relevant**: The paper exclusively concerns NEK9 variants and their association with neonatal arthrogryposis. There is no mention of the gene SERPINE2, no experimental data, genetic findings, mechanistic discussion, or literature citations related to SERPINE2. Because the paper contains no information about SERPINE2, it provides neither supporting nor contradicting evidence for the claim that SERPINE2 is likely associated with at least one Mendelian disease. Absence of mention in this paper does not imply anything about SERPINE2's disease associations; it simply means this study is unrelated to the claim.


[Read Paper](https://www.semanticscholar.org/paper/a69677b41b5f628b50b661893ae39d1b61fa7955)


### Whole‐exome sequencing analyses in a Saudi Ischemic Stroke Cohort reveal association signals, and shows polygenic risk scores are related to Modified Rankin Scale Risk

**Why Not Relevant**: The provided paper content does not mention SERPINE2, Mendelian disease(s), or any gene-level findings for specific genes. It only reports that gene burden testing in a Saudi stroke population found statistically significant signals in stroke genes in aggregate and that lower cumulative rare-variant risk correlated with lower mRS. Because the claim concerns whether SERPINE2 is likely associated with at least one Mendelian disease, evidence would need to include explicit statements such as: (a) SERPINE2 variants are enriched in affected individuals versus controls; (b) segregation of SERPINE2 variants with disease in families; (c) functional studies showing that SERPINE2 variants cause a loss- or gain-of-function consistent with a Mendelian phenotype; or (d) clinical case reports linking pathogenic SERPINE2 variants to a defined Mendelian disorder. None of those elements appear in the provided content. Without specific gene-level results or mechanistic data for SERPINE2, one cannot infer an association from the aggregate statement about stroke gene burden. To assess the claim properly, the full paper (including methods, results, tables listing gene-level burden p-values and variant-level annotations) or other literature specifically addressing SERPINE2 would be required.


[Read Paper](https://www.semanticscholar.org/paper/ef1e3cc537505def74437c2048ef722316d024a6)


### Crosstalk between BH4, pain, and dystonia

**Why Not Relevant**: The provided paper content does not mention the gene SERPINE2 anywhere. The excerpt only refers to IMPDH2, GCH1, and SPR in the context of dystonia and dopa-responsive dystonia. Therefore it provides neither direct evidence (no report of variants, segregation, phenotype, or case reports implicating SERPINE2) nor mechanistic evidence (no functional data, pathways, or interactions involving SERPINE2) relevant to the claim that SERPINE2 is likely associated with a Mendelian disease. Additionally, the fragment is truncated and limited in scope, preventing any indirect inference that would implicate SERPINE2.


[Read Paper](https://www.semanticscholar.org/paper/8165567c7df86cf75a17349edfb5017cf505c7d6)


## Search Queries Used

- SERPINE2 pathogenic variant Mendelian disease case report co segregation family clinical exome sequencing ClinVar OMIM HGMD

- SERPINE2 protease nexin 1 PN1 glia derived nexin loss of function knockout mouse phenotype functional assay protease inhibitor serpin extracellular matrix neural development

- SERPINE2 rare variant burden exome sequencing study rare disease cohort trio de novo autosomal dominant autosomal recessive monogenic gene level association

- SERPINE2 association genome wide association study GWAS complex trait expression quantitative trait locus eQTL pulmonary disease emphysema asthma neurodevelopmental disorder serpin family monogenic disease comparison


## Bibliometric Configuration

- Use Bibliometrics: True

- Author Impact Weight: 0.4

- Citation Impact Weight: 0.4

- Venue Impact Weight: 0.2
