## Supplementary material for "MENDELSEEK: An algorithm that predicts Mendelian Genes and elucidates what makes them special": SGK1.txt

# Claim: The gene SGK1 is likely associated with at least one Mendelian disease.

**Status**: processed

**Overall Rating**: -1

**Explanation**:

No relevant papers were found that support or refute this claim.


## Relevant Papers


## Other Reviewed Papers


### Evaluating the current state of Mendelian randomization studies: a protocol for a systematic review on methodological and clinical aspects using neurodegenerative disorders as outcome

**Why Not Relevant**: The provided paper content is a brief description of a systematic review protocol focused on methods in Mendelian randomization (MR) and does not mention the gene SGK1, any specific genes, Mendelian diseases, gene-disease associations, or empirical results. Verbatim from the paper content: "This systematic review protocol utilizes a unique comprehensive data abstraction tool based on recent methodological advancements in the field of MR to develop an in-depth understanding of how various MR methods have been applied, what has been achieved, and what can be done in future to arrive at true causal risk factors." This single-sentence description (and the surrounding context provided) makes clear that the document is methodological (a protocol) rather than presenting primary data, genetic association results, or mechanistic evidence about SGK1. Therefore it provides no direct evidence (no reported associations, statistics, or case descriptions) and no mechanistic evidence (no molecular, physiological, or pathophysiological information) relevant to the claim that "SGK1 is likely associated with at least one Mendelian disease." Additional limitations: the content is extremely short, contains no methods, results, or discussion sections, and does not reference any studies, databases, or gene lists that could be mined for indirect relevance. Because of these factors, the paper content cannot support or refute the claim about SGK1 and Mendelian disease.


[Read Paper](https://www.semanticscholar.org/paper/95ef9e8ea6a0e580d596685d70816fc4ac576bfc)


### Identification of a novel de novo pathogenic variant in GFAP in an Iranian family with Alexander disease by whole-exome sequencing

**Why Not Relevant**: The provided paper content is a single, high-level sentence: "The role of genetic in AxD diagnosing was highlighted and more information was provided to provide more information in order to expand the genetic spectrum of Iranian patients with AxD." This sentence does not mention the gene SGK1, any other specific gene, any Mendelian disease name linked to SGK1, nor any experimental, genetic, statistical, or mechanistic data. Therefore it provides no direct evidence (no associations, no variants, no case reports) and no mechanistic evidence (no expression, pathway, functional, or pathogenicity data) that would support or refute the claim that SGK1 is likely associated with at least one Mendelian disease. Limitations of the provided content: it is a single general statement about genetics and AxD (presumably Alexander disease or another AxD acronym) without methods, results, gene names, variants, patient data, or mechanistic discussion. Because none of the required elements (SGK1 mention, disease association, variant or functional evidence) are present, the content is not relevant to the claim.


[Read Paper](https://www.semanticscholar.org/paper/9b228b684c3371c832698bc6ddcbd35325a8af79)


### Exploring the Impact of Genetics in a Large Cohort of Moebius Patients by Trio Whole Exome Sequencing

**Why Not Relevant**: The paper focuses on Moebius syndrome (MBS) and a trio-WES study of 37 MBS and MBS-like patients, reporting negative or inconclusive results for candidate genes REV3L and PLXND1 and failing to identify causal mutations for MBS in this cohort. The manuscript text provided does not mention the gene SGK1, does not present any genetic, clinical, or functional data about SGK1, nor does it include mechanistic discussion or variant-level evidence that could bear on whether SGK1 is associated with a Mendelian disease. Because there is no analysis, result, or discussion of SGK1, the paper provides no direct evidence for or against the claim and no mechanistic data relevant to SGK1. Therefore the document is not relevant to evaluating whether SGK1 is likely associated with at least one Mendelian disease.


[Read Paper](https://www.semanticscholar.org/paper/226b561a7e847eca27fae7e2558a187686dd8f75)


### The simplest explanation does not have to be preferred: co-occurrence of pathogenic variants in cancer-predisposing genes

**Why Not Relevant**: The supplied text is a single, partial sentence describing a study of MINAS (multiple inherited neoplasia alleles syndrome) frequency and cancer phenotypes using recent literature and UK 10,000 Genomes Project data, and whether effects of cancer susceptibility genes (CSGs) are additive or synergistic. It does not mention the gene SGK1, Mendelian disease(s), any variants, results, mechanisms involving SGK1, or any data that could be interpreted as linking SGK1 to a Mendelian disorder. Therefore the content provides no direct evidence (no reports of pathogenic SGK1 variants or clinical associations) and no mechanistic evidence (no molecular, functional, or pathway information involving SGK1) relevant to the claim. Additional limitations that reduce relevance: the text is fragmentary and lacks methods, results, or statistical detail; it focuses on cancer predisposition syndromes and interactions among CSGs generally, not on specific genes; and absence of SGK1 in this excerpt is not evidence about SGK1 overall—it only means this paper excerpt does not bear on the claim. To evaluate the claim about SGK1 and Mendelian disease would require searching the full paper for any mention of SGK1 or consulting studies that report SGK1 variants segregating with Mendelian phenotypes, functional evidence of pathogenic variants, or curated disease–gene databases.


[Read Paper](https://www.semanticscholar.org/paper/a1372a664bd2c33a0bda638975a8b26d112457fc)


### Autosomal Dominant Intellectual Development Disorder-6 (MRD6) Without Seizures Linked to a De Novo Mutation in the grin2b Gene Revealed by Exome Sequencing: A Case Report of a Moroccan Child

**Why Not Relevant**: The provided paper content exclusively concerns the grin2b (GRIN2B) gene and its role in autosomal dominant intellectual development disorder-6 (MRD6). There is no mention of SGK1, no data on SGK1 variants, and no mechanistic discussion linking SGK1 to Mendelian disease in the text. Therefore the paper provides no direct or mechanistic evidence for or against the claim that SGK1 is likely associated with at least one Mendelian disease. The evidence in the paper is specific to a different gene (GRIN2B) and a single patient/family case; while useful for GRIN2B-related disease assessment, it cannot inform SGK1 disease association without additional data or explicit cross-gene analyses. To evaluate the SGK1 claim, literature or databases that specifically report SGK1 variants, segregation, functional studies, or OMIM annotations should be consulted.


[Read Paper](https://www.semanticscholar.org/paper/67d756e0cc302e5d46fdda9df34403f1f2de9a5a)


### P-540 Strategic implementation of PGT-M for multiple genes post qhole exome sequencing in couples with familial history: a comprehensive study

**Why Not Relevant**: This paper does not provide evidence for or against the claim that SGK1 is likely associated with at least one Mendelian disease. The text is a retrospective description of using WES for preimplantation genetic testing in couples with familial histories of genetic disorders; it does not mention SGK1, does not list gene-level findings, and does not describe any phenotypes, variant calls, or mechanistic data specific to SGK1. Representative verbatim passages from the paper that demonstrate the scope but lack SGK1-specific information include: "WES, followed by segregation analysis, pinpointed reportable pathogenic or likely pathogenic variants in 46 out of 94 families as the root cause of the referral condition." and "Interestingly, 42 of these families exhibited at least one additional common gene mutation in both partners." These sentences show that the study identified pathogenic variants and additional shared mutations at the cohort level but do not identify which genes were involved. Because no gene names, variant identifiers, phenotypic links, segregation details for SGK1, or mechanistic data related to SGK1 are provided, the paper cannot be used as direct evidence (identifying SGK1 variants causing Mendelian disease in patients) nor as mechanistic evidence (biological role of SGK1 causing a Mendelian phenotype). Limitations that prevent using this paper to support the SGK1 claim: (1) absence of any mention of SGK1; (2) aggregated reporting of diagnostic yield without per-gene results or variant lists; (3) no phenotype–genotype correlation data for any specific genes; (4) no functional or mechanistic experiments reported. Given these facts, the paper is not relevant to assessing whether SGK1 is associated with a Mendelian disease.


[Read Paper](https://www.semanticscholar.org/paper/1b721e406f9d31ac5983b146c5b9aa03d58fc49c)


### The application of whole-exome sequencing in the early diagnosis of rare genetic diseases in children: a study from Southeastern China

**Why Not Relevant**: This paper does not provide evidence for or against the claim that SGK1 is likely associated with at least one Mendelian disease. The provided content summarizes a retrospective whole-exome sequencing (WES) diagnostic cohort (144 patients, 61 diagnoses) and reports aggregate findings (diagnostic yield, higher yield for Trio-WES vs singleton-WES, distribution of clinical phenotypes, proportions of autosomal dominant disorders and de novo variants). Nowhere in the supplied text is the gene name "SGK1" (or any variant of that name) mentioned, nor are individual diagnosed genes, specific variants, segregation data, variant pathogenicity assessments, phenotype–genotype correlations, or functional/mechanistic data reported that would implicate SGK1. Therefore there is no direct evidence (no reported SGK1 variants, no clinical cases attributed to SGK1) nor mechanistic evidence (no functional studies, no pathway analyses) in this paper content that bears on the claim.

Limitations and caveats that maintain this non-relevance: (1) The absence of SGK1 in the supplied excerpt does not logically prove SGK1 is not implicated in the full study — it only shows the excerpt contains no gene-level details. If SGK1 were among the diagnosed genes but simply not named in the excerpt, the provided text would still be insufficient to evaluate the claim. (2) The cohort-level statistics (e.g., proportions of inheritance modes, de novo rate) are general and cannot be used to infer anything specific about SGK1 without gene-level data. (3) Mechanistic association of a gene with a Mendelian disease requires variant-level evidence (pathogenic/likely pathogenic variants), segregation or de novo occurrence, and ideally functional validation; none of these are present for SGK1 in the supplied content.

If you want a definitive assessment, I can (a) ask you to provide the full paper (including tables listing diagnosed genes and variants) so I can search for any SGK1 entries and extract verbatim sentences about them, or (b) run a literature review for published associations between SGK1 and Mendelian diseases and summarize genetic and mechanistic evidence.


[Read Paper](https://www.semanticscholar.org/paper/ba9aa0637319a74b3dfdae65099a70edd6c60ebf)


## Search Queries Used

- SGK1 mutation pathogenic variant case report familial segregation Mendelian disease OMIM ClinVar

- SGK1 whole exome sequencing monogenic disorder de novo variant autosomal dominant autosomal recessive rare disease trio sequencing

- serum glucocorticoid regulated kinase 1 SGK1 loss of function gain of function variant functional study knockout mouse phenotype human phenotype

- SGK1 review Mendelian gene ClinGen gene disease validity systematic review meta analysis hereditary disease database


## Bibliometric Configuration

- Use Bibliometrics: True

- Author Impact Weight: 0.4

- Citation Impact Weight: 0.4

- Venue Impact Weight: 0.2
