## Supplementary material for "MENDELSEEK: An algorithm that predicts Mendelian Genes and elucidates what makes them special": SHROOM3.txt

# Claim: The gene SHROOM3 is likely associated with at least one Mendelian disease.

**Status**: processed

**Overall Rating**: -1

**Explanation**:

No relevant papers were found that support or refute this claim.


## Relevant Papers


## Other Reviewed Papers


### Simultaneous identification and prioritization of variants in familial, de novo, and somatic genetic disorders with VariantMaster

**Why Not Relevant**: The provided paper content describes a computational pipeline (VariantMaster) and its application to identify causative variants, including an application to a rare Mendelian disease, but it does not mention the gene SHROOM3 or any gene-level results by name. Therefore the paper provides no direct evidence that SHROOM3 is associated with any Mendelian disease. The excerpts show the methods and that the approach can find causative variants in principle, which is mechanistically relevant (i.e., the tool could detect such an association), but without explicit results naming SHROOM3 there is no support for the specific claim. Limitations affecting this assessment: (1) the supplied content may be a summary/abstract and could omit gene-level results present elsewhere in the full paper; (2) typographical issues in the excerpt ('row (BAM) data') do not alter the absence of SHROOM3-specific findings. To evaluate the claim further would require the full paper (results/tables) or searching primary literature and databases (e.g., OMIM, ClinVar, PubMed) for SHROOM3 disease associations.


[Read Paper](https://www.semanticscholar.org/paper/a56e2dc7369c76b03ad5e9309093beb98a159719)


### Whole Exome Sequencing Identifies Novel De Novo Variants Interacting with Six Gene Networks in Autism Spectrum Disorder

**Why Not Relevant**: The provided paper content focuses on whole-exome sequencing and network analyses of de novo variants in Korean families with autism spectrum disorder. The text does not mention the gene SHROOM3 anywhere in the supplied excerpts and contains no gene-level statements or results that name SHROOM3. There are no experimental, statistical, or mechanistic data about SHROOM3 in the provided content. Consequently, this paper does not provide direct evidence that SHROOM3 is associated with any Mendelian disease, nor does it provide mechanistic evidence implicating SHROOM3. The paper's stated limitations (small cohort size, lack of functional experiments, reliance on network rather than statistical analysis) further reduce its capacity to supply indirect evidence about any specific gene not explicitly reported. Absence of SHROOM3 in these excerpts is not proof that SHROOM3 has no Mendelian association; it only shows this paper does not address the claim.


[Read Paper](https://www.semanticscholar.org/paper/2137cd76f68a615b7688c1f425cb6fa99233fd98)


### Leveraging linkage evidence to identify low-frequency and rare variants on 16p13 associated with blood pressure using TOPMed whole genome sequencing data

**Why Not Relevant**: The paper content provided contains only a general methodological statement about linkage analysis and rare-variant detection in whole-genome sequencing (WGS) data. It contains no mention of the gene SHROOM3, no presentation of results tying SHROOM3 to any phenotype or disease, and no mechanistic discussion of SHROOM3 function or pathogenic variants. Because the claim concerns a specific gene-disease association (SHROOM3 being likely associated with at least one Mendelian disease), a paper that only describes a general approach without reporting gene-specific findings cannot provide supporting or refuting evidence. While the described method could, in principle, be used to discover rare pathogenic variants in SHROOM3, the absence of any data, results, or discussion about SHROOM3 or Mendelian disorders means this paper does not bear on the claim. Therefore relevance is negligible.


[Read Paper](https://www.semanticscholar.org/paper/de84dfd3fb201fde6509a3aa1be506511ad59fea)


### Mutations and mechanism: how PINK1 may contribute to risk of sporadic Parkinson's disease.

**Why Not Relevant**: The provided paper content is a short scientific commentary focused on Parkinson’s disease genetics and specifically discusses genes known to cause autosomal dominant or recessive Parkinson’s disease (SNCA, LRRK2, VPS35, PINK1, PARK7/DJ-1, PARK2/PARKIN, PLA2G6, ATP13A2, and FBXO7) and the question of whether those genes also influence sporadic disease risk. The text makes no mention of SHROOM3, provides no genetic association data, no clinical descriptions linking SHROOM3 to any Mendelian disorder, and no mechanistic discussion involving SHROOM3. Therefore the document contains neither direct evidence (case reports, mutation-disease co-segregation, linkage, or association statistics) nor mechanistic evidence (functional studies, pathways, or molecular interactions) that would support or refute the claim that SHROOM3 is associated with a Mendelian disease. Additionally, this is a commentary about PINK1 and Parkinson’s genetics rather than a primary study or review of SHROOM3, so no indirect inferences about SHROOM3 can be drawn from the content provided. Because the required verbatim mentions, data, or mechanistic descriptions of SHROOM3 are entirely absent, the relevance to the claim is effectively zero.


[Read Paper](https://www.semanticscholar.org/paper/99a394b75e80548eaff92176d5e13d2748cf9bc6)


### Success and Pitfalls of Genetic Testing in Undiagnosed Diseases: Whole Exome Sequencing and Beyond

**Why Not Relevant**: This paper does not provide evidence for or against the claim that SHROOM3 is likely associated with at least one Mendelian disease. The manuscript excerpt describes methodological approaches and three specific case examples (variants in MEIS2, NIPBL, and genes involved in chromatin-remodeling) but does not mention SHROOM3, any variants in SHROOM3, nor mechanistic data linking SHROOM3 to a Mendelian phenotype. Specific reasons this paper is not relevant to the claim: 1) No mention of the gene SHROOM3 in the title, abstract, or provided text; 2) Reported cases and genes are MEIS2, NIPBL, and chromatin-remodeling genes—none are SHROOM3; 3) No experimental results, segregation data, population or constraint metrics, functional assays, methylation signatures, animal models, or literature references related to SHROOM3 are presented; 4) The paper focuses on diagnostic strategies (re-analysis, reverse phenotyping, methylation signatures) rather than establishing novel gene-disease associations for SHROOM3. Note that the absence of SHROOM3 data in this paper is not evidence that SHROOM3 is not associated with a Mendelian disease; it simply means this paper cannot be used to support or refute the claim. To evaluate the claim, one should consult primary gene-disease resources (OMIM, ClinVar, HGMD), constraint and population databases (gnomAD), case reports and cohort studies reporting pathogenic SHROOM3 variants, and functional/animal model studies demonstrating a plausible disease mechanism.


[Read Paper](https://www.semanticscholar.org/paper/9c76f2886b2c4e59e399c5b986c6cf0018a2cdce)


### Identification of ultra-rare genetic variants in Pediatric Acute Onset Neuropsychiatric Syndrome (PANS) by exome and whole genome sequencing

**Why Not Relevant**: The paper does not mention SHROOM3 anywhere in the provided text. It reports candidate mutations in a set of 11 genes (PPM1D, SGCE, PLCG2, NLRC4, CACNA1B, SHANK3, CHK2, GRIN2A, RAG1, GABRG2, and SYNGAP1) identified in a PANS cohort and discusses mechanistic themes for those genes. Because SHROOM3 is neither listed among identified candidate genes nor discussed mechanistically, the paper provides no direct evidence that SHROOM3 is associated with any Mendelian disease. Additionally, the study is disease-focused (PANS) and applied filters for de novo/ultra-rare variants (MAF < 0.001) in mostly exome data, so even as negative evidence this is limited: absence of SHROOM3 here could reflect the study's phenotype focus, cohort composition, variant-type detection limits (e.g., noncoding or structural variants, or variants with higher population frequency), or insufficient power to detect very rare associations. Therefore the paper is not relevant for supporting the claim that SHROOM3 is likely associated with at least one Mendelian disease.


[Read Paper](https://www.semanticscholar.org/paper/12b10580fd211efe9b3cb05b11a539b2e4a7eabc)


### 1684: PATHOGENIC VARIATION ASSOCIATED WITH MONOGENIC DISORDERS IN A SEVERE PEDIATRIC SEPSIS COHORT

**Why Not Relevant**: The paper does not mention the gene SHROOM3 anywhere in the provided text. The study analyzed a pre-defined panel of 96 genes and reports pathogenic variants in a set of specific genes/conditions (atypical HUS, ALPS, cryopyrin-associated periodic syndromes, CVID, FMF, PIK3R1 deficiency, HLH, XIAP-related disease), but SHROOM3 is not among the reported genes or diagnoses. Therefore the paper provides no direct evidence supporting or refuting the claim that SHROOM3 is associated with at least one Mendelian disease. Furthermore, because the study used a limited gene panel and specific variant filters (MAF <5%, HGMD/ACMG pathogenicity criteria, and certain mutation types), the absence of SHROOM3 findings could reflect that SHROOM3 was not included in the panel or that any SHROOM3 variants did not meet reporting thresholds; neither possibility is addressed in the text. In sum, absence of mention in this small, targeted study is insufficient to evaluate the claim. To assess the claim about SHROOM3 one would need studies that specifically include SHROOM3 in their analysis or report SHROOM3 variants with disease segregation/functional evidence.


[Read Paper](https://www.semanticscholar.org/paper/cdd32cc853ff52097298692a62266f79b44b8c53)


### #4317 CLINICAL DIVERSITY OF STEROID-RESISTANT NEPHROTIC SYNDROME CAUSED BY TRPC6 MUTATIONS

**Why Not Relevant**: The paper content supplied pertains to steroid-resistant nephrotic syndrome (SRNS) and reports pathogenic variants found in that SRNS cohort, notably TRPC6, NUP107, PLCE1, COL4A3, COL4A5, and other single-case genes. The text does not mention SHROOM3 anywhere, nor does it provide experimental, genetic, or mechanistic data related to SHROOM3. Therefore the paper provides no direct evidence for or against the claim that SHROOM3 is associated with a Mendelian disease, and offers no mechanistic rationale involving SHROOM3. Note that absence of SHROOM3 in this disease-focused cohort does not prove SHROOM3 is not associated with any Mendelian disorder (limited sample size, disease specificity, and the study's targeted gene findings mean negative inference is not generalizable). Given the complete lack of mention or data about SHROOM3 in the provided text, relevance to the claim is effectively zero.


[Read Paper](https://www.semanticscholar.org/paper/c2747aaff3d6f0c6e29493cdff73cf48fd818d22)


### #3320 GENETIC TESTING FOR MOLECULAR DIAGNOSIS OF NEPHROLITHIASIS AND NEPHROCALCINOSIS IN ADULT PATIENTS: A SINGLE-CENTER COHORT

**Why Not Relevant**: This paper does not mention SHROOM3 explicitly, nor does it provide a full list of the 45 genes on the nephrolithiasis panel. The text reports which genes had pathogenic or likely pathogenic variants in this cohort, and SHROOM3 is not among the genes listed as harboring pathogenic variants—however, omission from the reported positive findings does not constitute affirmative evidence that SHROOM3 is not associated with a Mendelian disease. Because the paper's focus is genes linked to nephrolithiasis and it neither names SHROOM3 nor describes any mechanistic data involving SHROOM3, it provides no direct or mechanistic evidence for or against the claim that SHROOM3 is likely associated with at least one Mendelian disease. Limitations preventing relevance include: (1) lack of explicit statement whether SHROOM3 was included on the panel, (2) small cohort limited to nephrolithiasis phenotypes (so non-renal Mendelian diseases would not be detected), and (3) absence of functional or segregation data regarding SHROOM3. For these reasons the paper is of minimal relevance to the claim.


[Read Paper](https://www.semanticscholar.org/paper/2097cfd6ee6b8857428d8e8895413812313205a9)


## Search Queries Used

- SHROOM3 SHROOM 3 mutation variant Mendelian disease monogenic familial autosomal dominant autosomal recessive pathogenic

- SHROOM3 neural tube defect spina bifida congenital malformation kidney renal development CAKUT congenital anomalies of the kidney and urinary tract

- SHROOM3 exome sequencing whole exome sequencing genome sequencing rare variant burden study segregation linkage study de novo variant family study

- SHROOM3 ClinVar OMIM HGMD pathogenic variant clinical significance loss of function knockout mouse functional study epithelial morphogenesis actin cytoskeleton


## Bibliometric Configuration

- Use Bibliometrics: True

- Author Impact Weight: 0.4

- Citation Impact Weight: 0.4

- Venue Impact Weight: 0.2
