## Supplementary material for "MENDELSEEK: An algorithm that predicts Mendelian Genes and elucidates what makes them special": SLC4A5.txt

# Claim: The gene SLC4A5 is likely associated with at least one Mendelian disease.

**Status**: processed

**Overall Rating**: -1

**Explanation**:

No relevant papers were found that support or refute this claim.


## Relevant Papers


## Other Reviewed Papers


### Genome-Wide Association Study of Blood Pressure Extremes Identifies Variant near UMOD Associated with Hypertension

**Why Not Relevant**: Relevance is essentially zero because the provided paper text investigates and reports associations for the UMOD gene (uromodulin) locus (chromosome 16, rs13333226) in relation to hypertension and related phenotypes. The content contains no mention of SLC4A5, no data about variants in SLC4A5, and no analyses relevant to Mendelian (monogenic) disease causation. Furthermore, the study focuses on common-variant, population-level association statistics (ORs, HRs, combined P values) and functional correlates of a common allele, which are conceptually distinct from evidence that a gene is causally associated with a Mendelian disease (which would require rare, highly penetrant variants segregating in families or functional experiments showing monogenic pathogenicity). Therefore this paper does not provide direct or mechanistic evidence bearing on the claim that SLC4A5 is likely associated with at least one Mendelian disease.


[Read Paper](https://www.semanticscholar.org/paper/e4f8bacddb8dd09701d8050f9136fe14998e5806)


### The Burden of Candidate Pathogenic Variants for Kidney and Genitourinary Disorders Emerging From Exome Sequencing

**Why Not Relevant**: The provided paper content describes the methods used to identify and curate a set of genes associated with Mendelian kidney and genitourinary disorders and the variant-detection/annotation pipeline, but the excerpt does not mention the gene SLC4A5 by name nor present a gene-level list or results that would confirm or refute SLC4A5's association with any Mendelian disease. Consequently, there is no direct evidence in the supplied text that SLC4A5 is associated with a Mendelian disorder. The methodological descriptions are relevant only insofar as they indicate that SLC4A5 would have been captured if present in OMIM/Orphanet and retained after curation, but without the actual curated gene list or variant-level findings included here, one cannot determine SLC4A5's status from this excerpt. To assess the claim definitively, one would need (a) the paper's supplementary gene list (the final 625 genes) or tables reporting per-gene findings, or (b) direct queries of OMIM, ClinVar, HGMD, or the authors' supplementary materials for SLC4A5 entries and evidence of Mendelian association. Also note the paper emphasizes that databases can contain misclassified variants and that manual curation removed genes with insufficient evidence, so even database presence alone would require critical appraisal.


[Read Paper](https://www.semanticscholar.org/paper/a75e297208c04992568e607d19aa9b9aaebd09d5)


### A common genetic variant of the chromogranin A-derived peptide catestatin is associated with atherogenesis and hypertension in a Japanese population.

**Why Not Relevant**: The paper addresses polymorphisms in the CHGA gene (chromogranin A), specifically catestatin (CST) Ser-364 and pancreastatin (PST) Ser-297 variants, and their associations with cardiovascular/metabolic phenotypes in a Japanese cohort. It contains no mention of the gene SLC4A5, no data or discussion linking CHGA to SLC4A5, and no evidence regarding Mendelian (monogenic) disease associations. The study design is a population association study of common SNPs and complex traits (blood pressure, baPWV), not an investigation of rare pathogenic variants or cosegregation in families that would support a Mendelian disease assignment. Therefore the paper provides no direct or mechanistic evidence bearing on the claim that SLC4A5 is likely associated with at least one Mendelian disease. Additional reasons limiting relevance: (1) different gene entirely (CHGA vs SLC4A5); (2) phenotype focus on complex cardiovascular/metabolic traits rather than Mendelian syndromes; (3) no functional or mechanistic experiments linking the studied variants to monogenic disease processes; (4) population-specific association results (N=343) that do not address causality or inheritance patterns typical of Mendelian disease evidence.


[Read Paper](https://www.semanticscholar.org/paper/0d109d9ea746b298331a114774f00253535fa5e8)


### Whole-Exome Sequencing in Adults With Chronic Kidney Disease

**Why Not Relevant**: This paper does not provide any direct or mechanistic evidence about the gene SLC4A5. The provided text describes the study design, methods, and general analytic pipeline for whole-exome sequencing (WES) in adults with chronic kidney disease and indicates that the authors compiled a curated list of 287 OMIM 'nephropathy genes', but the excerpt does not list those genes or report any findings for SLC4A5 specifically. Key reasons the paper is not relevant to the claim that "SLC4A5 is likely associated with at least one Mendelian disease":

- No mention of SLC4A5: The supplied content contains no occurrences of the string "SLC4A5" or any direct report of variants, pathogenicity assessments, or phenotype associations involving SLC4A5.

- No gene-level results shown: Although the methods describe annotation, gene lists, and variant interpretation (including use of ClinVar, HGMD, and an OMIM-derived 287-gene list), the excerpt does not provide the gene list nor per-gene results that would allow assessment of SLC4A5.

- Absence of mechanistic detail specific to SLC4A5: The text explains general WES methodology and curation strategy, but provides no functional studies, mechanistic pathways, or clinical-phenotype correlations that implicate SLC4A5 in Mendelian disease.

- Coverage caveats are generic: The authors note that some exons in their nephropathy gene set are not captured by the capture kit ("Among the 287 nephropathy genes, 29 were identified with at least 1 exon that is not captured by the Agilent kit, representing potential blind spots in the analysis (Supplement Table 3)."), which highlights that even if SLC4A5 had been of interest, absence of a reported finding could reflect technical blind spots; however, because SLC4A5 is not mentioned, we cannot use this paper to support or refute its association with Mendelian disease.

Conclusion: Based on the supplied content, the paper is not relevant to assessing whether SLC4A5 is associated with a Mendelian disease. To evaluate that claim you should consult primary resources that specifically report SLC4A5 variants and associated phenotypes (for example, OMIM, ClinVar, HGMD, PubMed case reports, or larger WES/WGS studies that list per-gene diagnostic findings).


[Read Paper](https://www.semanticscholar.org/paper/a29bd57964f68ac5f8e1abcafe1112ba488f53f5)


### From case studies to community knowledge base: MSeqDR provides a platform for the curation and genomic analysis of mitochondrial diseases

**Why Not Relevant**: The provided paper content describes a community resource (MSeqDR) for mitochondrial disease gene and variant curation, examples of gene discovery workflow (e.g., PMPCA), and the capabilities of the database and tools. It does not mention the gene SLC4A5 anywhere in the text, nor does it present data (genetic, clinical, functional, or statistical) about SLC4A5. Therefore there is no direct evidence (case reports, functional assays, genetic segregation, or curated pathogenic assertions) or mechanistic evidence in this paper that bears on whether SLC4A5 is associated with any Mendelian disease. Because the content is focused on database infrastructure and mitochondrial gene curation broadly, it cannot be used to support or refute the specific claim about SLC4A5. If additional evidence is desired, targeted literature, ClinVar/MSeqDR queries, or primary genetic studies mentioning SLC4A5 would be required.


[Read Paper](https://www.semanticscholar.org/paper/6f85c595adaca6963c30fea73bc37266fa78be56)


### Use of animal models for exome prioritization of rare disease genes

**Why Not Relevant**: The provided paper content contains only a single high-level methodological sentence about the general benefit of comparing patient phenotypes to known disease and model-organism phenotypes. It contains no mention of the gene SLC4A5, no variant data, no phenotype descriptions linked to SLC4A5, and no explicit statements about Mendelian disease associations. Consequently, there is no direct experimental, observational, or mechanistic evidence in the excerpt that bears on whether SLC4A5 is associated with at least one Mendelian disease. The content only implies that the described approach might be useful in general for discovering gene-disease associations, but without any gene-specific results or details this is not evidence for the claim. Given the absence of any SLC4A5-specific information, the relevance to the claim is negligible.


[Read Paper](https://www.semanticscholar.org/paper/dd7890c1b59c7ecd3f43ad5f49d23ac809fa8a05)


### Genome-wide association study on blood pressure traits in the Iranian population suggests ZBED9 as a new locus for hypertension

**Why Not Relevant**: The provided paper content contains a single sentence: "Although there is no strong evidence to support the function of ZBED9 in blood pressure regulation, it provides new insight into the pleiotropic effects of hypertension and other cardiovascular diseases." This sentence discusses ZBED9, blood pressure regulation, and pleiotropy in hypertension/cardiovascular disease. It does not mention the gene SLC4A5, Mendelian disease(s), pathogenic variants, segregation, clinical phenotypes consistent with a Mendelian disorder, or any experimental or genetic evidence linking SLC4A5 to disease. Therefore there is no direct evidence in this content supporting or refuting the claim that "The gene SLC4A5 is likely associated with at least one Mendelian disease." There is also no mechanistic information (e.g., biochemical function, pathway involvement, loss/gain-of-function effects, animal models) about SLC4A5 to evaluate plausibility. Because the content neither addresses the gene nor Mendelian inheritance, it cannot be used to assess the claim. To evaluate the claim would require other sources such as primary genetic association studies, case reports showing segregation, curated gene-disease databases (ClinVar, OMIM, ClinGen), functional studies of SLC4A5 variants, or population/variant pathogenicity analyses.


[Read Paper](https://www.semanticscholar.org/paper/2ee15df8d8a71bd05769b4ebbd2795d65d273e60)


### Integration of multi-omics technologies for molecular diagnosis in ataxia patients

**Why Not Relevant**: This paper does not provide direct or mechanistic evidence regarding the claim that SLC4A5 is likely associated with at least one Mendelian disease. The provided content describes a study of eight patients with episodic ataxia and reports pathogenic variants in SPG7, ATXN2, ELOVL4, and PMPCB (and a potential modifier in ZFYVE26). Nowhere in the abstract/background/methods/results/conclusion text provided is SLC4A5 mentioned, analyzed, or implicated. Therefore the paper offers no direct observations (variant calls, segregation, functional assays) or mechanistic data (gene expression changes, splicing effects, protein function mechanisms) related to SLC4A5. Additional reasons why this paper is not informative for the claim: 

- Scope mismatch: the study focuses on identifying molecular causes of episodic ataxia in a very small cohort (n=8) using multi-omics and long-read sequencing; its gene findings are limited to the genes explicitly named (SPG7, ATXN2, ELOVL4, PMPCB, ZFYVE26). Absence of SLC4A5 in these results does not constitute evidence against SLC4A5 being associated with some Mendelian disease generally — it only shows SLC4A5 was not implicated in this particular small ataxia cohort. 

- Sample size and phenotype limitation: with only eight patients and a specific phenotype (episodic ataxia), the study cannot comprehensively assess all Mendelian disease genes across different phenotypes. Thus, lack of mention of SLC4A5 is expected and uninformative for its overall disease association. 

- No mechanistic interrogation of SLC4A5: the methods (WGS, RNA-seq from PBMCs, long-read sequencing) could in principle detect variants or aberrant transcripts in many genes, but the paper reports findings only for other genes; there are no mechanistic data (e.g., altered transcript, splicing, protein function) regarding SLC4A5. 

In sum, because the paper does not mention or analyze SLC4A5 at all, it provides neither supporting nor refuting evidence for the claim. Note that absence of evidence in this single, small, phenotype-specific study is not evidence that SLC4A5 is not associated with a Mendelian disease; it simply provides no information relevant to the claim.


[Read Paper](https://www.semanticscholar.org/paper/bd3da10cb65efe894fe906b0bf2fed7fb9eb0919)


### TTC7A Variants Previously Described to Cause Enteropathy Are Observed on a Single Haplotype and Appear Non-pathogenic in Pediatric Inflammatory Bowel Disease Patients

**Why Not Relevant**: The provided paper content discusses variants in TTC7A and assessment of TTC7A's contribution to polygenic IBD. The claim concerns whether the gene SLC4A5 is likely associated with at least one Mendelian disease. The excerpt contains no mention of SLC4A5, Mendelian disease associations, or mechanistic links involving SLC4A5. Therefore the paper content as given provides no direct evidence (no observations, case reports, segregation data, variant pathogenicity assessments, or statistical associations) that SLC4A5 is associated with any Mendelian disease. Mechanistic evidence is also absent: there are no descriptions of SLC4A5 function, pathogenic mechanisms, or biological pathways that would link SLC4A5 variation to Mendelian phenotypes. Because the gene names and study aims differ (TTC7A vs SLC4A5; polygenic IBD vs Mendelian disease), the paper is irrelevant to evaluating the claim. Caveat: only a single-sentence summary of the paper was provided; if the full paper contains additional content mentioning SLC4A5, that text was not supplied and could not be evaluated. Based on the supplied content alone, relevance to the claim is effectively zero.


[Read Paper](https://www.semanticscholar.org/paper/3818a966b5d6e8b0eda65fdef043ac0727fa429c)


### The simplest explanation does not have to be preferred: co-occurrence of pathogenic variants in cancer-predisposing genes

**Why Not Relevant**: The provided paper content does not mention SLC4A5 or any data specific to that gene. The only sentence supplied is: "The frequency and cancer phenotypes associated with MINAS by reviewing the recent literature and by checking the UK 10,000 Genomes Project data from participants with a phenotype of tumor predisposition syndromes or multiple primary tumors, and if they are additive reflecting each CSG independently or synergistic resulting in more severe phenotypes are reported." This text discusses MINAS, cancer phenotypes, the UK 10,000 Genomes Project, and cancer susceptibility genes (CSGs) in the context of tumor predisposition syndromes and multiple primary tumors. It provides no direct evidence (no variant, case, cosegregation, statistical association, or clinical description) linking SLC4A5 to any Mendelian disease, nor does it provide mechanistic information about SLC4A5 (e.g., gene function, pathogenic mechanisms, protein effects). Therefore the paper content as provided is not relevant to assessing the claim that SLC4A5 is likely associated with at least one Mendelian disease. Limitations: only a single sentence of the paper was provided; if the full paper contains sections mentioning SLC4A5, those were not included here. Based on the supplied content alone, no support or refutation of the claim can be extracted.


[Read Paper](https://www.semanticscholar.org/paper/a1372a664bd2c33a0bda638975a8b26d112457fc)


### Genetic Association of CYP1A2 Gene Variant (rs762551) with Caffeine Induced-Hypertension Susceptibility and Cytochrome P450 1A2 Protein Analyses

**Why Not Relevant**: The provided paper content does not mention the gene SLC4A5, Mendelian disease(s), or any gene-disease association relevant to SLC4A5. Instead, the entire excerpt discusses CYP1A2 and a specific CYP1A2 variant (15:74749576C>A), its allele frequencies, association testing, and bioinformatics characterization for that gene/protein in a Pakistani case-control pilot study (n = 50). Because SLC4A5 is not named and no data, results, or mechanistic discussion relevant to SLC4A5 appear in the text, there is neither direct evidence (e.g., genetic association, segregation, variant pathogenicity) nor mechanistic evidence (e.g., functional assay, pathway linking SLC4A5 perturbation to a Mendelian phenotype) in this paper to support or refute the claim that SLC4A5 is likely associated with at least one Mendelian disease.

Relevant points from the excerpt that demonstrate this irrelevance (verbatim content is not included in "excerpts" because it does not pertain to SLC4A5): the paper explicitly focuses on "CYP1A2" and describes a variant "15:74749576C>A (C allele lowering CYP1A2 activity)"; reports sample sizes and allele frequencies (e.g., "overall, 8%, 56% and 36% sampled population (n=50) is homozygous wild-type (C/C), heterozygous (C/A) and homozygous mutant (A/A) respectively"); reports a chi-square association test result "χ2(2, N = 50) = 4.244, p = 0.039" and a Hardy Weinberg p-value "0.241"; and lists bioinformatics tools applied to predict properties of the CYP1A2 protein. None of these statements concern SLC4A5 or Mendelian disease associations.

Because the paper contains no information about SLC4A5, no further extraction or mechanistic interpretation relating to SLC4A5 is possible from this document. If you want evidence about SLC4A5 and Mendelian disease associations, please provide papers or data that mention SLC4A5 specifically, or I can search the literature for studies that analyze SLC4A5 and its clinical associations.


[Read Paper](https://www.semanticscholar.org/paper/8f683d5c73ddc4b4eadea9fab31fadcbd85f0291)


### Re-evaluation of whole exome sequencing, including intronic region, in combination with genetic intolerance score for foetal structural anomalies, is helpful for diagnosis, especially in X-linked disorders

**Why Not Relevant**: The provided paper content does not mention the gene SLC4A5 and contains no data, results, or discussion that directly or mechanistically address SLC4A5 or its association with any Mendelian disease. The paper describes methods (WES, CNV analysis, LOEUF, spliceAI) and reports findings for other genes (for example TMEM67 and CASK) and general observations about X-linked genes, but SLC4A5 is not present in the text provided. Therefore there is no direct evidence (case reports, variant pathogenicity, segregation, population frequency, or functional data) nor mechanistic evidence (molecular function, pathway perturbation, animal models) in this paper that would support or refute the claim that SLC4A5 is likely associated with at least one Mendelian disease.

Limitations and context: absence of SLC4A5 in this paper is not evidence that SLC4A5 is not disease-associated — it only means this particular study (138 fetuses with structural abnormalities, diagnostic reanalysis using LOEUF and spliceAI) did not report any findings related to SLC4A5. The study population, phenotype focus (fetal structural abnormalities), and sample size could easily miss disease associations for genes not implicated in this cohort. To evaluate the claim about SLC4A5, a targeted literature and database search (OMIM, ClinVar, HGMD, PubMed, gnomAD constraint metrics, and functional studies) is required.


[Read Paper](https://www.semanticscholar.org/paper/b9004e518cf738650e488c6c4990525732fce092)


### Uncovering the shared genetic contributors to primary and secondary hypertension using whole genome sequencing in a national disease cohort

**Why Not Relevant**: The provided paper content does not mention SLC4A5, nor does it report any association between SLC4A5 and extreme early-onset hypertension (EEHTN) or any Mendelian disease. The study performed WGS and per-gene rare-variant analyses in an EEHTN cohort and reported PKD1 as a significant gene and no other gene-level associations reaching significance; the absence of SLC4A5 from the results is therefore an indirect negative observation only for this specific phenotype and cohort. Important limitations that reduce relevance to the broader claim: (1) the study targeted EEHTN, so it would not detect Mendelian disease associations of SLC4A5 manifesting with other phenotypes; (2) lack of a reported association in this cohort does not prove lack of Mendelian effects elsewhere (absence of evidence is not evidence of absence); (3) detectability depends on variant types, filters, allele-frequency thresholds, statistical power, and phenotype definition—factors that could prevent discovery of a true SLC4A5 Mendelian association in this study. Because the paper provides only indirect, negative, and phenotype-limited evidence and contains no direct or mechanistic data on SLC4A5, its relevance to the claim that "SLC4A5 is likely associated with at least one Mendelian disease" is very low.


[Read Paper](https://www.semanticscholar.org/paper/77cb04eafb3fb7e573bd2726504cc4376c22b2c1)


## Search Queries Used

- SLC4A5 pathogenic variant mutation case report Mendelian monogenic disease clinical genetics OMIM ClinVar HGMD

- solute carrier family 4 member 5 SLC4A5 NBCe2 sodium bicarbonate cotransporter ion transport knockout mouse model kidney acid base homeostasis functional study

- SLC4A5 genetic association GWAS blood pressure hypertension locus common variant population study allele frequency association

- SLC4A5 rare disease exome sequencing undiagnosed rare disorder trio sequencing Mendelian candidate gene pathogenicity loss of function variant


## Bibliometric Configuration

- Use Bibliometrics: True

- Author Impact Weight: 0.4

- Citation Impact Weight: 0.4

- Venue Impact Weight: 0.2
