## Supplementary material for "MENDELSEEK: An algorithm that predicts Mendelian Genes and elucidates what makes them special": SLC4A7.txt

# Claim: The gene SLC4A7 is likely associated with at least one Mendelian disease.

**Status**: processed

**Overall Rating**: 0

**Explanation**:

Supporting evidence.
The provided paper establishes that SLC4A7 (encoding the electroneutral Na/HCO3 cotransporter NBCn1) is a functional effector at a genome-wide association locus for blood pressure. The authors report that a blood-pressure (BP) risk allele at the SLC4A7 locus is associated with open chromatin, allele-specific DNA–protein binding, higher SLC4A7 mRNA and protein expression, increased membrane availability of NBCn1, and measurable changes in Na/HCO3-dependent base flux and steady-state intracellular pH in vascular smooth muscle cells. The paper also notes that knockout models show an altered blood pressure phenotype, indicating that loss of SLC4A7 function influences blood-pressure regulation in experimental systems. These findings demonstrate a clear physiological role for SLC4A7 in a trait relevant to disease.

Caveats or contradictory evidence.
Despite the clear link to blood-pressure regulation, the study concerns common polymorphisms and complex-trait (polygenic) association signals rather than rare, high-penetrance variants segregating in families. The missense polymorphism examined (E326K) showed no discernible functional effect in the assays reported. There is no presentation in the excerpts of human pedigrees, segregation data, de novo/rare variant findings, or clinical descriptions that would establish a Mendelian (monogenic) disease caused by mutations in SLC4A7. Knockout phenotypes in model organisms indicate that perturbation of the gene affects physiology, but such models do not by themselves demonstrate that humans carry rare variants in SLC4A7 that produce a Mendelian disorder. Therefore, the provided evidence supports involvement in a complex trait (blood pressure) but does not provide the kind of genetic/clinical evidence required to claim association with a Mendelian disease.

Analysis of potential underlying mechanisms, deeper relationships, or second-order implications.
Mechanistically, SLC4A7’s role in regulating intracellular pH in vascular smooth muscle provides a plausible pathway by which altered transporter expression or activity could influence vascular tone and blood pressure. This physiological plausibility makes SLC4A7 a credible candidate gene for disorders affecting blood-pressure homeostasis. However, the difference between being a contributor to a complex trait (via common regulatory variation altering expression) and being the locus of rare, highly penetrant mutations that cause a Mendelian syndrome is substantial. Monogenic disease typically requires rare variants with large effect sizes, often disrupting protein function in a way that results in a consistent clinical phenotype with familial segregation. The evidence here documents regulatory effects of common alleles and model-organism phenotypes; it does not document rare, damaging variants in human families or clinical case series consistent with Mendelian inheritance.

Which rating is most appropriate and why.
Given the available information, the most accurate assessment is that there is no evidence in the provided paper that SLC4A7 is associated with a Mendelian disease. The study provides strong support for SLC4A7’s involvement in blood-pressure regulation as a complex trait locus but does not provide the genetic or clinical data required to establish a link to a Mendelian disorder. Thus, based on the excerpts provided, the claim that SLC4A7 is likely associated with at least one Mendelian disease is not supported by the evidence at hand.


**Final Reasoning**:

I reviewed the provided paper excerpt and considered the types of evidence required to establish that a gene is associated with a Mendelian disease. The paper gives solid functional and GWAS-based evidence that SLC4A7 affects blood-pressure biology via common regulatory variation and that perturbation in model organisms alters blood-pressure phenotype. However, there is no presentation of rare, high-penetrance human variants, family segregation data, de novo mutation evidence, or clinical case descriptions that would indicate a Mendelian disorder caused by SLC4A7. The presence of a physiologically relevant role (and knockout phenotypes) increases biological plausibility but does not constitute evidence of Mendelian disease in humans. Therefore, the proper rating, given only the provided evidence, is "No Evidence." If further human genetic or clinical studies (not provided here) identify segregating rare variants or recurrent pathogenic mutations in SLC4A7 causing a consistent syndrome, the assessment should be revisited.


## Relevant Papers


### LBOS 02-04 BLOOD PRESSURE-ASSOCIATED POLYMORPHISMS IN SLC4A7 (SODIUM/BICARBONATE CO-TRANSPORTER NBCn1) ARE LINKED TO GENE EXPRESSION AND INTRACELLULAR pH REGULATION.

**Authors**: F. Ng (H-index: 16), M. Caulfield (H-index: 112)

**Relevance**: 0.15

**Bibliometric Impact**: 0.54


**Excerpts**:

- Chromosome 3p24.1 containing the SLC4A7 gene has been identified by genome-wide association studies as one of the genomic loci that influence blood pressure.

- SLC4A7 encodes electroneutral Na/HCO3 co-transporter (NBCn1) which regulates intracellular pH (pHi) in various tissues including vascular smooth muscle and endothelium, with knockout models demonstrating an altered blood pressure phenotype.

- We conducted a functional study of blood pressure-associated single nucleotide polymorphisms at the SLC4A7 locus in primary cultures of human vascular smooth muscle and endothelial cells.

- FAIRE identified the BP risk allele as associated with preferentially open chromatin, and EMSA identified allele-dependent DNA-protein interactions.

- Consistent with this, the risk allele was associated with increased NBCn1 mRNA, protein expression, availability at the cell membrane and subsequently higher Na/HCO3-dependent, dimethlyamiloride-insensitive net base flux and steady-state pHi compared to protective allele homozygotes.

- However, in the presence of Na/H exchange activity, the genotypic effect of NBCn1 on net acid extrusion and steady-state pHi was detected only in vascular smooth muscle cells but not endothelial cells.

- There was no discernable effect of the missense polymorphism resulting in the amino acid substitution E326K.

- These findings provide an insight into the molecular mechanism underlying the association of variation at the SLC4A7 locus and blood pressure, revealing an important genotypic influence on SLC4A7 expression and pHi regulation in vascular smooth muscle cells.


**Explanations**:

- Relation to claim: This sentence documents that common-variant association (GWAS) links the SLC4A7 locus to blood pressure, a complex trait. Type of evidence: indirect/population-level association evidence, not Mendelian. Strengths/limitations: Supports that variation at SLC4A7 affects human physiology (blood pressure) but does not demonstrate a rare, high-penetrance, familial (Mendelian) disease caused by SLC4A7 variants.

- Relation to claim: Describes SLC4A7's molecular function (NBCn1) and notes that knockout models show altered blood pressure phenotype. Type of evidence: mechanistic (gene function) and animal-model functional evidence. Strengths/limitations: Mechanistically strengthens plausibility that SLC4A7 perturbation can produce a blood-pressure phenotype; however, knockout model phenotypes (presumably in animals) do not equal demonstration of a human Mendelian disease and do not show inheritance or high penetrance in humans.

- Relation to claim: States the study interrogates BP-associated SNPs at SLC4A7 in primary human vascular cells. Type of evidence: mechanistic/functional study context. Strengths/limitations: Shows the paper intentionally examines functional consequences of variants at SLC4A7, which is relevant to pathogenic mechanisms, but these are common SNPs linked to polygenic blood pressure variation rather than rare Mendelian mutations.

- Relation to claim: Reports allele-specific chromatin state and protein-DNA binding for the BP-associated allele. Type of evidence: mechanistic molecular evidence. Strengths/limitations: Demonstrates a regulatory mechanism by which SLC4A7 expression is altered by common alleles; this strengthens biological plausibility that SLC4A7 dosage affects physiology, but it is not evidence for a monogenic Mendelian disorder caused by SLC4A7 variants.

- Relation to claim: Shows the BP-risk allele increases SLC4A7/NBCn1 expression and functional transporter activity in cells. Type of evidence: mechanistic functional evidence. Strengths/limitations: Directly links genetic variation to altered protein levels and cellular function, bolstering plausibility that SLC4A7 perturbations can change physiology. Limitation: effect described for common risk allele (likely modest population effect) and measured in vitro; not evidence of segregating, highly penetrant pathogenic variants causing Mendelian disease.

- Relation to claim: Indicates cell-type specificity of the genotype effect (present in vascular smooth muscle cells but not endothelial cells under certain conditions). Type of evidence: mechanistic/modifier evidence. Strengths/limitations: Refines understanding of where SLC4A7 variation acts (vascular smooth muscle), which is helpful for disease mechanism modeling; does not provide evidence for Mendelian inheritance or monogenic disease phenotypes.

- Relation to claim: Reports that a tested missense polymorphism (E326K) had no discernable effect. Type of evidence: direct functional test of a coding variant. Strengths/limitations: This is direct negative evidence that this particular amino-acid substitution likely does not alter NBCn1 function in the assays used, which weakens the case that common coding variation at least (E326K) confers strong functional perturbation. Limitation: only one missense variant was evaluated; absence of effect here does not exclude other rare, damaging variants causing Mendelian disease.

- Relation to claim: Authors' overall interpretation that variation at SLC4A7 influences blood pressure via altered expression and pHi regulation in vascular smooth muscle. Type of evidence: mechanistic and interpretive. Strengths/limitations: Summarizes functional link between genetic variation and physiologic effect, supporting biological plausibility that SLC4A7 perturbation can affect a clinically relevant phenotype (blood pressure). However, these data concern common-variant, complex-trait associations and cell/animal models rather than human pedigrees, segregation, or identification of rare, high-penetrance pathogenic SLC4A7 alleles required to support a Mendelian-disease claim.


[Read Paper](https://www.semanticscholar.org/paper/48402ac30b91eaee9a90e33d8abc09cad1711ea1)


## Other Reviewed Papers


### Effective variant filtering and expected candidate variant yield in studies of rare human disease

**Why Not Relevant**: The provided paper content is a single, high-level methodological sentence about variant filtering and expected candidate counts by inheritance mode. It contains no mention of the gene SLC4A7, no reported associations between any specific gene and Mendelian disease, and no mechanistic or experimental data that would link SLC4A7 to a Mendelian phenotype. Although methods that improve variant filtering can indirectly aid discovery of disease genes, that is a general statement about capability and not evidence that any particular gene (including SLC4A7) is likely associated with a Mendelian disease. Because the content lacks gene-specific results, annotations, patient data, statistical associations, functional experiments, or mechanistic discussion involving SLC4A7, the paper content as provided is not relevant to the claim.


[Read Paper](https://www.semanticscholar.org/paper/8da5eda5547e3521df7aecc8c8e4b7380a7641fa)


### Identification of a novel pathogenic missense mutation in PRPF31 using whole exome sequencing: a case report

**Why Not Relevant**: This paper concerns a novel missense mutation in PRPF31 causing autosomal-dominant retinitis pigmentosa (adRP) and presents clinical phenotyping, segregation analysis, and cellular localisation experiments for PRPF31 c.590T>C (p.Leu197Pro). The provided text makes no mention of the gene SLC4A7, nor of SLC4 family genes, bicarbonate transporters, or any phenotypes or mechanisms associated with SLC4A7. Therefore it provides neither direct genetic evidence (segregation, case reports, variant pathogenicity) nor mechanistic evidence (biochemical function, cellular assays linking variant to disease) relevant to the claim that SLC4A7 is likely associated with a Mendelian disease. In short, the paper is entirely focused on PRPF31; nothing in the manuscript supports or refutes an association between SLC4A7 and any Mendelian disorder. Consequently the relevance to the claim is effectively zero. If you want evidence for or against SLC4A7 causing a Mendelian disease, papers reporting genetic variants in SLC4A7 with segregation, ClinVar/OMIM annotations, functional studies on SLC4A7 protein function, or population/constraint metrics should be examined instead.


[Read Paper](https://www.semanticscholar.org/paper/b7bd5ced68a256e3065d97070c1ea52154f31a5c)


### Different Interpretations of the Same Genetic Data

**Why Not Relevant**: The provided paper excerpt discusses discordant clinical interpretations of genetic variants across laboratories, using ClinVar and ClinGen data focused on hereditary cancer genes and long QT syndrome genes. Nowhere in the supplied text is the gene SLC4A7 mentioned, nor are any data, variants, functional studies, or gene-disease associations presented that pertain to SLC4A7. Therefore the paper provides no direct evidence (e.g., reported pathogenic variants, case reports, segregation, or statistical association) nor mechanistic evidence (e.g., functional studies, gene-disease pathways) supporting that SLC4A7 is associated with any Mendelian disease. Absence of mention in this paper is not proof that SLC4A7 is not disease-associated, but it means this paper is irrelevant to evaluating the claim. Limitations: the paper analyzes certain gene sets (hereditary cancer and long QT genes) and general patterns of interpretation discordance; it may use ClinVar as a data source that could contain entries for many genes, but the excerpt shows the authors' focus was on specific disease gene lists and on interpretation discordance rather than on discovering or validating gene-disease relationships. To assess the claim about SLC4A7 one would need literature, database entries, or functional/segregation evidence specifically mentioning SLC4A7.


[Read Paper](https://www.semanticscholar.org/paper/523adc8d0ddf6caf2c9fbac87b68ea818a15577b)


### The Brain Gene Registry: a data snapshot

**Why Not Relevant**: The provided paper content does not mention the gene SLC4A7, nor does it mention Mendelian disease(s), pathogenicity, specific variants, genotype-phenotype relationships, or any experimental/clinical findings tying a specific gene to disease. The two sentences only describe the BGR as a resource recruiting participants with clinically reported variants to support research on brain genes and neurodevelopmental disorders. While such a resource could, in principle, contain data relevant to SLC4A7 in the future, the excerpt contains no direct evidence (no case reports, association statistics, segregation data, or functional studies) and no mechanistic information (no molecular function, pathway, or variant effect descriptions) that would support or refute the claim that SLC4A7 is likely associated with at least one Mendelian disease. Key limitations: (1) absence of any mention of SLC4A7 by name or by any synonyms, (2) absence of disease names or descriptions indicating Mendelian inheritance, (3) absence of methods, results, or conclusions linking genes to phenotypes, and (4) no page or figure references; therefore the content is not relevant to the claim.


[Read Paper](https://www.semanticscholar.org/paper/6bfb15c40fc279b33126d895d962795ae504b41b)


### Homozygous nonsense mutation of WNT10B gene in a Moroccan family with split-hand foot malformation identified by exome sequencing: a case report

**Why Not Relevant**: The paper exclusively investigates split-hand foot malformation (SHFM) and identifies a homozygous nonsense variant in WNT10B (p.Arg115*) as the cause of SHFM6 in the studied family. There is no mention of SLC4A7, no genetic data concerning SLC4A7, and no mechanistic pathways discussed that link WNT10B (a Wnt family gene involved in limb development) to SLC4A7 (a solute carrier gene). Therefore this paper provides no direct genetic evidence, no mechanistic evidence, and no contextual support for the claim that SLC4A7 is likely associated with at least one Mendelian disease. Absence of evidence in this paper does not prove SLC4A7 is not associated with Mendelian disease, but it means this paper is not relevant to assessing that claim.


[Read Paper](https://www.semanticscholar.org/paper/194f0bf41aeb43f2c77e85cee133dbef6aae04f2)


### A novel pathogenic CDH3 variant underlying heredity hypotrichosis simplex detected by whole-exome sequencing (WES)—a case report

**Why Not Relevant**: This paper does not provide any direct or mechanistic evidence regarding SLC4A7. The study focuses on hereditary hypotrichosis simplex in a family and reports a novel pathogenic variant in CDH3 (p.Ser223GlyfsTer4) as the plausible causal variant. There is no mention of SLC4A7, no reported variants in SLC4A7, and no experimental or interpretive discussion about SLC4A7 function or disease association. While whole-exome sequencing was used (and WES can in principle detect SLC4A7 variants), the presented results and conclusions concern CDH3 only. Therefore the paper is not relevant to the claim that SLC4A7 is likely associated with at least one Mendelian disease.


[Read Paper](https://www.semanticscholar.org/paper/19c243ea07846bbfa3e076308b66f1d70d8154ca)


### Gene discovery and cellular modelling of rare autosomal recessive neurodevelopmental conditions

**Why Not Relevant**: The paper text provided does not mention the gene SLC4A7 anywhere. All reported gene-disease findings in this study concern HERC2, TUBA1A, KLHL7 and MAL. Therefore there is no direct evidence (e.g., variant association, segregation, functional assays) nor mechanistic evidence in this paper supporting or refuting an association between SLC4A7 and any Mendelian disease. Limitations and caveats: absence of mention in this paper is not evidence that SLC4A7 is not associated with a Mendelian disease—this study examined only 13 families with NDD using WES and focused on variants they prioritized; SLC4A7 could be implicated in other cohorts, phenotypes, non-coding variants, structural variants, or diseases outside the scope of this study. However, with respect to this paper alone, the relevance to the claim is effectively nil.


[Read Paper](https://www.semanticscholar.org/paper/1916e63deed84eba433676327e1d8b5a8d009cf7)


### Novel compound heterozygous variants of the SEC23A gene in a Chinese family with cranio-lenticulo-sutural dysplasia based on data from a large cohort of congenital cataract patients

**Why Not Relevant**: The provided content discusses CLSD/ARCLSD (clinical spectrum, use of TEM, and congenital cataracts) and contains no mention of the gene SLC4A7, its variants, functional studies, patient genotypes, or mechanistic pathways involving SLC4A7. Therefore the excerpt does not support or refute the claim that SLC4A7 is likely associated with at least one Mendelian disease. Additionally, there are no experimental details, statistical associations, or mechanistic descriptions in the excerpt that could be used to infer a link to SLC4A7. Caveat: this assessment is based solely on the single sentence provided; if the full paper includes other sections (methods, results, supplementary data) that were not supplied and that discuss SLC4A7, those could change the relevance, but they are not available for review here.


[Read Paper](https://www.semanticscholar.org/paper/68f926b3596267c271d77af321ead5eb43315c5a)


### A novel mutation in ANOS1 in a Chinese family with Kallmann syndrome: Case report

**Why Not Relevant**: The provided paper content does not mention SLC4A7 at all. The two sentences discuss a novel variant related to Kallmann syndrome and explicitly refer to the ANOS1 gene and CHH patients. While these sentences are relevant to ANOS1 as a Mendelian-disease-associated gene, they provide no direct or mechanistic evidence about SLC4A7. There are no variant identifiers, experimental results, functional assays, segregation data, population statistics, or mechanistic pathways relevant to SLC4A7 in the excerpt. Therefore the paper content as provided is not relevant to the claim that SLC4A7 is likely associated with at least one Mendelian disease.


[Read Paper](https://www.semanticscholar.org/paper/0d61601eb76c8707ec36d8b02a62cb32092b0a68)


### Clingen Coagulation Factor Deficiency Variant Curation Expert Panel: Meeting the Need for Recommendations to Curate Variants in the Coagulation Factor Genes

**Why Not Relevant**: The provided paper content focuses exclusively on variant curation for coagulation factor genes, specifically F8 and F9, and on the activities of the ClinGen Coagulation Factor Deficiency Variant Curation Expert Panel (CFD-VCEP). There is no mention of SLC4A7, no presentation of variants in SLC4A7, no clinical case reports, no genetic linkage or association data, and no functional/mechanistic studies involving SLC4A7. The material consists of methodological and organizational description and illustrative examples (F8/F9 pilot variants, ClinVar curation plans) relevant to coagulation disorders; it does not provide direct genetic or mechanistic evidence that SLC4A7 is associated with any Mendelian disease. The only remotely relevant point is that the curation framework described could in principle be applied to other genes in future, but that is an administrative/methodological statement and does not constitute biological or clinical evidence of disease association. Therefore the content is not relevant to evaluating the claim that SLC4A7 is likely associated with at least one Mendelian disease.


[Read Paper](https://www.semanticscholar.org/paper/9ea8d4ce19622ae1c6b0cdfd40848c93199f6280)


### Genome Alert!: a standardized procedure for genomic variant reinterpretation and automated genotype-phenotype reassessment in clinical routine

**Why Not Relevant**: The paper describes a tool and presents aggregate results showing that Genome Alert! can identify new gene-disease associations and produce clinically actionable reclassifications. However, the provided content contains no mention of the gene SLC4A7 (no gene-level lists, examples, or explicit statements about SLC4A7). Therefore, although the paper demonstrates a mechanism and precedent for discovering Mendelian associations, it supplies no direct or specific evidence that SLC4A7 is associated with any Mendelian disease. Without gene-level data or a list of the 75 high-confidence genes, one cannot conclude from this paper that SLC4A7 is among them. Consequently the relevance to the specific claim about SLC4A7 is very low.


[Read Paper](https://www.semanticscholar.org/paper/e34deed2cf57a676587676cd0c6b782e03832a25)


### LBOS 02-04 BLOOD PRESSURE-ASSOCIATED POLYMORPHISMS IN SLC4A7 (SODIUM/BICARBONATE CO-TRANSPORTER NBCn1) ARE LINKED TO GENE EXPRESSION AND INTRACELLULAR pH REGULATION

**Why Not Relevant**: The paper investigates common-variant (GWAS) associations at the SLC4A7 locus and provides mechanistic cellular data showing how risk alleles influence SLC4A7 expression and intracellular pH regulation in vascular smooth muscle. However, it does not identify or characterize rare, highly penetrant variants segregating in families, describe clinical Mendelian phenotypes attributable to SLC4A7 mutations, or provide case-level genetic/segregation evidence required to establish association with a Mendelian (monogenic) disease. The study even reports that a tested missense polymorphism (E326K) had no discernable effect. Therefore, while the paper offers mechanistic plausibility that perturbing SLC4A7 can alter blood pressure, it provides no direct evidence that SLC4A7 is associated with any human Mendelian disease.


[Read Paper](https://www.semanticscholar.org/paper/558493f53f1c9b0d876f678c328631fc04d7603d)


### Whole-exome sequencing revealed a novel homozygous missense variant in OSGEP gene: a case report of Galloway–Mowat syndrome in Iran

**Why Not Relevant**: The provided paper content does not mention SLC4A7 and therefore provides no direct or mechanistic evidence for the claim that "The gene SLC4A7 is likely associated with at least one Mendelian disease." The only sentence provided is: "A 12-year-old male with intellectual disability, poor speech, seizures, microcephaly, and nephrotic syndrome is described that were in favor of Galloway–Mowat syndrome, born to a healthy Iranian consanguineous parents, and mutational screening revealed a novel homozygote OSGEP gene missense variant." This sentence (and the rest of the supplied content) implicates a different gene, OSGEP, in a clinical presentation consistent with Galloway–Mowat syndrome; it does not mention SLC4A7, any variants in SLC4A7, nor any pathway linking OSGEP to SLC4A7. Therefore there is neither direct genetic evidence (cases, segregation, pathogenic variants) nor mechanistic evidence (functional studies, biochemical pathway links) in this content that would support or refute an association between SLC4A7 and a Mendelian disease. Additional reasons for non-relevance: (1) the report is a single-case description focused on OSGEP, so even if SLC4A7 were mentioned it would be low-level evidence without replication/segregation; (2) no experimental methods, results, or mechanistic data concerning SLC4A7 are provided; and (3) no statistical evidence or population data relevant to SLC4A7 are included. To evaluate the claim about SLC4A7, manuscripts reporting SLC4A7 variants in affected individuals, segregation data, case-control or cohort analyses, or functional/mechanistic studies of SLC4A7 would be required, none of which are present in the supplied content.


[Read Paper](https://www.semanticscholar.org/paper/ea9cceaa84e1b07a4276db70840e43b0ef851bb5)


### Germline mosaicism in a collagen VI‐related myopathy family: A cause of autosomal recessive inheritance

**Why Not Relevant**: The paper exclusively studies COL6A1 (and generally COL6A2/COL6A3) variants and collagen VI-related myopathies, describing germline mosaicism in COL6A1 as the cause of recessive Ullrich congenital muscular dystrophy in the reported family. There is no mention of the gene SLC4A7, no variants of SLC4A7 are reported or tested, and no mechanistic pathways involving SLC4A7 are discussed. Therefore the paper provides no direct evidence (genetic, clinical, or statistical) nor mechanistic plausibility linking SLC4A7 to any Mendelian disease. Because the paper's entire focus is on COL6A1 and related genes, its findings cannot be used to support or refute the claim about SLC4A7.


[Read Paper](https://www.semanticscholar.org/paper/4572f635bb354974678a75579201d9eedc2c40ea)


### [OP.7C.02] BLOOD PRESSURE-ASSOCIATED POLYMORPHISMS IN SLC4A7 (SODIUM/BICARBONATE CO-TRANSPORTER NBCN1) ARE LINKED WITH GENE EXPRESSION AND INTRACELLULAR PH REGULATION

**Why Not Relevant**: The paper provides strong mechanistic and functional evidence that variation at the SLC4A7 locus affects gene regulation, protein expression, and cellular physiology relevant to blood pressure. However, the claim concerns association of SLC4A7 with at least one Mendelian (monogenic, typically high-penetrance) disease. This manuscript focuses on common-variant (GWAS) associations and in vitro/knockout mouse physiological effects; it does not report rare, segregating, high-penetrance variants in human families, clinical case series, loss-of-function human mutations, or genetic evidence meeting typical standards for Mendelian disease association (segregation, de novo occurrence, strong effect sizes in humans). The knockout mouse phenotype is informative for biological plausibility but does not establish a human Mendelian disorder. Therefore the paper is only indirectly relevant (mechanistic plausibility) and does not provide direct genetic or clinical evidence that SLC4A7 is the cause of any human Mendelian disease.


[Read Paper](https://www.semanticscholar.org/paper/efa23fe51610ee64406af0caf502c1526ffbb8f7)


## Search Queries Used

- SLC4A7 Mendelian disease mutation case report family exome sequencing pathogenic variant

- SLC4A7 ClinVar OMIM ClinGen gene disease validity clinical variant report

- SLC4A7 NBCn1 sodium bicarbonate transporter loss of function knockout mouse phenotype pH regulation vascular blood pressure

- exome sequencing rare disease SLC4A7 candidate de novo segregation autosomal dominant autosomal recessive SLC4 family review


## Usage Statistics

- Prompt Tokens: 0

- Completion Tokens: 0

- Total Tokens: 0

- Estimated Cost: $0.0688


## Bibliometric Configuration

- Use Bibliometrics: True

- Author Impact Weight: 0.4

- Citation Impact Weight: 0.4

- Venue Impact Weight: 0.2
