## Supplementary material for "MENDELSEEK: An algorithm that predicts Mendelian Genes and elucidates what makes them special": SLC30A4.txt

# Claim: The gene SLC30A4 is likely associated with at least one Mendelian disease.

**Status**: processed

**Overall Rating**: 0

**Explanation**:

Supporting evidence.
The provided paper presents a broad method showing that information about pathogenic missense variants in paralogous genes can be used to extend ACMG PS1/PM5 criteria (para-PS1/PM5) and thereby increase the number of amino-acid positions that can be classified as likely pathogenic. The authors report mapping millions of variants across 9,990 genes and state that for 36 genes the extended criteria reach a strong evidence level. In principle, this approach could produce new evidence implicating genes that previously lacked direct disease associations if pathogenic variants in their paralogs exist and align to conserved residues.

Caveats or contradictory evidence.
None of the provided excerpts mention SLC30A4 specifically, nor do they list which genes reached the strong evidence threshold. The paper is methodological and reports global performance improvements and counts (e.g., 3.6-fold increase in classifiable amino acids), but it does not by itself provide direct, gene-specific evidence that SLC30A4 is associated with any Mendelian disorder. Without a statement that SLC30A4 was among the genes with pathogenic evidence, the paper does not constitute direct support for the specific claim. Additionally, demonstrating that paralogous pathogenic variants provide moderate-to-strong evidence for variant interpretation is not equivalent to demonstrating that a given gene is causally associated with a monogenic disease — variant interpretation criteria operate at the variant level and require corroborating genetic and clinical evidence to establish gene–disease relationships.

Analysis of potential underlying mechanisms, deeper relationships, or second-order implications.
SLC30A4 (a member of the SLC30A/ZNT zinc transporter family) has paralogs (e.g., SLC30A1–SLC30A10). If pathogenic missense variants are known in one paralog at a conserved residue, transfer of that evidence to SLC30A4 variants could strengthen interpretation of specific SLC30A4 variants under the para-PS1/PM5 framework. Thus, the methodological advance described could plausibly enable discovery or stronger support of SLC30A4 pathogenic variants in the future. However, establishing that the gene itself is associated with a Mendelian disease typically requires case-level segregation, recurrence, functional validation, or curated gene–disease assertions (e.g., in ClinGen). The excerpted paper does not report such gene-level assertions for SLC30A4.

Which rating is most appropriate.
Given the absence of direct, gene-specific evidence in the provided excerpts and the methodological (rather than gene-focused) nature of the paper, the balanced conclusion is that there is no evidence in the provided material that SLC30A4 is associated with at least one Mendelian disease. While the para-PS1/PM5 framework could in principle help implicate SLC30A4 if paralogous pathogenic variants exist, that possibility is speculative without explicit data. Therefore the appropriate rating based on the supplied evidence is "No Evidence."


**Final Reasoning**:

The single supplied paper is a methods/annotation study showing that paralogous pathogenic variants can extend variant interpretation criteria and increase the number of amino-acid positions where PS1/PM5-like evidence can be applied. However, the excerpts do not mention SLC30A4 specifically or provide gene-level assertions that it is associated with a Mendelian disorder. The methodological finding is necessary background but not sufficient to claim SLC30A4 is likely disease-associated. Without direct evidence (case reports, curated gene–disease links, or explicit mention in the paper's gene list), the correct assessment is that there is no evidence in the provided material to support the claim. I reviewed the logic for gaps and confirmed that while the paper supplies a plausible mechanism by which SLC30A4 could be implicated in future studies, it does not itself supply the required gene-specific evidence. Thus the rating remains "No Evidence."


## Relevant Papers


### Pathogenic paralogous variants can be used to apply the ACMG PS1 and PM5 variant interpretation criteria

**Authors**: Tobias Brünger (H-index: 9), D. Lal (H-index: 11)

**Relevance**: 0.12

**Bibliometric Impact**: 0.18000000000000002


**Excerpts**:

- Purpose The majority of missense variants in clinical genetic tests are classified as variants of uncertain significance.

- We hypothesized that incorporation of pathogenic missense variants in conserved residues across paralogous genes can increase the number of variants where ACMG PS1/PM5 criteria can be applied.

- We mapped over 2.5 million pathogenic and general population variants from ClinVar, HGMD, and gnomAD databases onto 9,990 genes and aligned these by gene families.

- Subsequently, we developed a novel framework to extend PS1/PM5 by incorporating pathogenic paralogous variants annotations (para-PS1/PM5).

- We demonstrate that para-PS1/PM5 criteria increase the number of classifiable amino acids 3.6-fold compared to PS1 and PM5.

- Across all gene families with at least two disease-associated genes, the calculated likelihood ratios suggest moderate evidence for pathogenicity.

- Moreover, for 36 genes, the extended para-PS1/PM5 criteria reach strong evidence level.

- Conclusion We show that single pathogenic paralogous variants incorporation at paralogous protein positions increases the applicability of the PS1 and PM5 criteria, likely leading to a reduction of variants of uncertain significance across many monogenic disorders. Future iterations of the ACMG guidelines may consider para-PS1 and para-PM5.


**Explanations**:

- This sentence provides general context about the problem the paper addresses (high prevalence of variants of uncertain significance). Relevance to SLC30A4: indirect. It identifies the clinical interpretation gap that the paper's methods aim to reduce, which could, in principle, affect interpretation of variants in SLC30A4 if applied. Limitation: no SLC30A4-specific data; this is background rather than evidence that SLC30A4 is disease-associated.

- This is the core mechanistic hypothesis of the paper: using pathogenic missense variants at conserved residues in paralogs to broaden PS1/PM5 application. Relevance to SLC30A4: mechanistic and indirect. If SLC30A4 has paralogs with known pathogenic missense variants at conserved positions, this approach could provide evidence linking SLC30A4 variants to Mendelian disease. Limitation: hypothesis-level statement; actual applicability to any specific gene (including SLC30A4) depends on whether the necessary paralogous pathogenic annotations exist and on the quality of alignments.

- This methods sentence documents the scale and data sources used (ClinVar, HGMD, gnomAD; 2.5M variants; 9,990 genes). Relevance to SLC30A4: indirect. Strength: large-scale analysis increases the chance that many gene families (potentially including the SLC family) were examined. Limitation: no gene-level breakdown in this excerpt; presence of SLC30A4 among analyzed genes is not confirmed here, nor are results for specific genes provided.

- This describes the new analytic framework (para-PS1/PM5). Relevance to SLC30A4: mechanistic/technical. The framework is precisely the tool that could generate evidence for or against SLC30A4 being disease-associated by transferring pathogenicity evidence across paralogs. Limitation: description of method only; no demonstration in this excerpt that the framework produced evidence for SLC30A4 specifically or that it avoids potential misannotations when paralogs have differing functional constraints.

- This result shows the method increases the number of classifiable amino acids 3.6-fold relative to using PS1/PM5 alone. Relevance to SLC30A4: indirect but practically meaningful. It indicates the method meaningfully increases the number of variants that can be given pathogenic evidence, implying that applying the method could make some SLC30A4 variants classifiable. Limitations: the fold-change is aggregated across genes; no per-gene breakdown, confidence intervals, or statistical tests are provided here. It does not indicate that SLC30A4 itself reached classification or has disease association.

- This sentence reports that across gene families with ≥2 disease-associated genes, likelihood ratios indicate moderate evidence for pathogenicity. Relevance to SLC30A4: mechanistic/statistical and indirect. If SLC30A4 belongs to a gene family that meets these criteria and has paralogs with disease association, the para-PS1/PM5 approach could supply moderate evidence toward pathogenicity of SLC30A4 variants and thus support SLC30A4 being associated with Mendelian disease. Limitations: no numeric LRs, no per-family or per-gene LRs given here; the strength 'moderate' is aggregated and may not apply to all families. The approach requires existing disease annotations in paralogs, which may be absent for SLC30A4 or its paralogs.

- This reports that for 36 genes the extended criteria reach 'strong evidence' level. Relevance to SLC30A4: indirect. It demonstrates the method can in some cases produce strong evidence for specific genes. Limitation: the excerpt does not name the 36 genes; SLC30A4 is not mentioned, so we cannot infer whether it is among them. Without a gene list, this does not constitute direct evidence for SLC30A4.

- This conclusion summarizes that incorporation of single pathogenic paralogous variants at paralogous positions increases PS1/PM5 applicability and likely reduces VUS across many monogenic disorders. Relevance to SLC30A4: indirect/mechanistic. It supports the plausibility that applying para-PS1/PM5 to SLC30A4 variants could yield pathogenic classifications and thus strengthen an association between SLC30A4 and Mendelian disease if supporting paralogous data exist. Limitations: 'likely' is qualitative; no evidence specific to SLC30A4 is presented here. Also, applicability depends on the availability and correctness of paralogous pathogenic annotations and on evolutionary and functional conservation that may differ between paralogs.


[Read Paper](https://www.semanticscholar.org/paper/0a921a972e8db185945c5510191b7ac52b42e325)


## Other Reviewed Papers


### Whole exome sequencing of familial hypercholesterolaemia patients negative for LDLR/APOB/PCSK9 mutations

**Why Not Relevant**: The paper focuses specifically on discovering novel genetic causes of familial hypercholesterolaemia (FH) by exome sequencing 125 affected individuals and comparing them to 1926 controls. The gene SLC30A4 is not mentioned anywhere in the provided content, and the study's conclusions list different candidate genes (e.g., CH25H and INSIG2) while explicitly stating no major novel FH locus was detected. Therefore the paper provides no direct evidence that SLC30A4 is associated with any Mendelian disease. Moreover, the study’s scope is limited to FH; even if SLC30A4 were associated with a different Mendelian disorder, this study would not address that. Additional limitations reduce indirect informativeness: modest case sample size, disease-specific cohort, allele-frequency and functional-effect filtering, and the fact that lack of detection here does not prove absence of an association—particularly for very rare or locus-heterogeneous Mendelian conditions. For these reasons the paper is of very low relevance to the claim that "SLC30A4 is likely associated with at least one Mendelian disease."


[Read Paper](https://www.semanticscholar.org/paper/7263443ef9fc236db6e7f85afa27611bf139577c)


### A rare loss-of-function variant of ADAM17 is associated with late-onset familial Alzheimer disease

**Why Not Relevant**: The provided paper content discusses a rare nonsynonymous variant rs142946965 (p.R215I) in ADAM17 co-segregating with autosomal-dominant late-onset Alzheimer’s disease in a single family. There is no mention of the gene SLC30A4, no data on variants in SLC30A4, and no mechanistic discussion involving SLC30A4. Verbatim from the provided content: "Using whole-genome sequencing, a single rare nonsynonymous variant (SNV) rs142946965 [p.R215I] in ADAM17 is identified co-segregating with an autosomal-dominant pattern of late-onset AD in one family and a causative association of the identifiedADAM17 variant in the pathogenesis of AD is supported." Because the paper excerpt exclusively addresses ADAM17 and Alzheimer’s disease, it provides neither direct evidence (e.g., genetic co-segregation, case reports, or statistical association) nor mechanistic evidence (e.g., functional assays, pathway analyses) linking SLC30A4 to any Mendelian disease. The absence of any mention of SLC30A4 in the provided content makes it impossible to extract relevant supporting or refuting evidence for the claim that SLC30A4 is likely associated with at least one Mendelian disease. Note that this conclusion applies only to the supplied content; it does not judge other literature or sections of the full paper that were not provided.


[Read Paper](https://www.semanticscholar.org/paper/14fa706015833b1c741911774309d29908a911c5)


### The Burden of Candidate Pathogenic Variants for Kidney and Genitourinary Disorders Emerging From Exome Sequencing

**Why Not Relevant**: The supplied paper excerpt does not mention the gene SLC30A4 anywhere and provides no gene-specific results or discussion that would allow assessment of an association between SLC30A4 and any Mendelian disease. The text describes study aims, cohort characteristics, gene-identification methods (OMIM, Orphanet), curation to a final list of 625 genes, and variant-filtering pipelines, but does not list the genes or report variant findings at the level of individual genes in the provided content. Therefore there is neither direct evidence (e.g., reported pathogenic variants in SLC30A4, statistical association, case reports) nor mechanistic evidence (e.g., functional data, pathogenic mechanism involving SLC30A4) in the excerpt to support or refute the claim that SLC30A4 is likely associated with at least one Mendelian disease.

Limitations and caveats relevant to this assessment:
- The study methodology (use of OMIM/Orphanet, ClinVar/HGMD, and curation to 625 genes) means SLC30A4 could potentially be included among the curated genes or among reported findings in the full paper or appendices, but the provided content does not include the gene list or any gene-specific results, so inclusion cannot be determined from the excerpt.
- Because no gene-level data or mechanistic discussion of SLC30A4 are present, no evaluation of statistical significance, variant pathogenicity, inheritance mode, or functional mechanism is possible from the supplied text.

Recommended next steps to evaluate the claim properly:
1) Provide the full paper or appendices containing the curated gene list and per-gene results so we can check whether SLC30A4 is included and whether any variants in SLC30A4 were observed and how they were classified.
2) If unavailable, search the full text, supplementary materials, or the laboratory Web site referenced in the paper for an explicit mention of SLC30A4 or for the downloadable gene list used in the study.
3) If SLC30A4 is listed, extract the relevant verbatim passages (variant counts, classifications, evidence) for direct and mechanistic evaluation.

Because the provided excerpt contains no SLC30A4-specific information, the relevance to the claim is effectively zero based on the available content.


[Read Paper](https://www.semanticscholar.org/paper/a75e297208c04992568e607d19aa9b9aaebd09d5)


### Precision Medicine in Internal Medicine

**Why Not Relevant**: This paper is a general review of genetic testing strategies, variant interpretation, and resources (e.g., OMIM, ClinVar, ES/GS) and does not mention the gene SLC30A4 or present any data (case reports, cohort analyses, pathogenic variants, segregation evidence, functional studies) that would directly link SLC30A4 to an established Mendelian disease. Therefore it provides no direct evidence for the claim. The included excerpts are contextual/mechanistic only (showing the overall state of gene-disease discovery, the role of ClinVar, and the capability of exome sequencing to discover novel associations). Limitations affecting relevance: the paper's OMIM statistic is dated (1 February 2019) and the review format means it is not intended to catalog gene-specific associations; any relevant evidence about SLC30A4 (if it exists) would need to be sought in gene-specific databases, ClinVar entries, OMIM gene/disease entries, primary case reports, or more recent gene-disease curation resources. Because the paper contains no mention of SLC30A4, the evidence bearing on the claim within this paper is essentially absent.


[Read Paper](https://www.semanticscholar.org/paper/c7199e8a5226f8d9d55777da0e15e91797bb459a)


### From case studies to community knowledge base: MSeqDR provides a platform for the curation and genomic analysis of mitochondrial diseases

**Why Not Relevant**: This paper does not provide any direct or mechanistic evidence regarding the gene SLC30A4. The content is focused on mitochondrial disease genetics, an example involving PMPCA, and the description and goals of the MSeqDR resource (a database and tool suite). Nowhere in the provided text is SLC30A4 named, nor are any data, variants, functional assays, or mechanistic discussions presented that pertain to SLC30A4. The paper therefore supplies no direct observational evidence (e.g., patient variants, segregation, functional rescue) nor any mechanistic rationale linking SLC30A4 to a Mendelian disease. The text does contain general methodological and contextual statements (for example, that massively parallel sequencing and functional follow-up can establish causality for newly identified disease genes, and that MSeqDR curates >1360 nuclear and mitochondrial genes relevant to mitochondrial biology and disease), but these are generic and do not bear on the claim about SLC30A4 specifically. Limitations: absence of mention in this paper is not evidence that SLC30A4 is not associated with Mendelian disease—only that this paper does not address it. To evaluate the claim about SLC30A4 would require searching other primary literature, gene-disease databases (ClinVar, OMIM, ClinGen), and functional studies focused on SLC30A4.


[Read Paper](https://www.semanticscholar.org/paper/6f85c595adaca6963c30fea73bc37266fa78be56)


### Two novel mutations in ALDH18A1 and SPG11 genes found by whole-exome sequencing in spastic paraplegia disease patients in Iran

**Why Not Relevant**: This paper does not provide any direct or mechanistic evidence relevant to the claim about SLC30A4. The manuscript text refers specifically to other genes implicated in hereditary spastic paraplegia (ALDH18A1 and SPG11) and reports pathogenic variants in those genes. Relevant verbatim sentences from the provided content (included here to show scope and focus) are:

"Hereditary spastic paraplegia is a not common inherited neurological disorder with heterogeneous clinical expressions." 

"ALDH18A1 (located on 10q24.1) gene-related spastic paraplegias (SPG9A and SPG9B) are rare metabolic disorders caused by dominant and recessive mutations that have been found recently." 

"Autosomal recessive hereditary spastic paraplegia is a common and clinical type of familial spastic paraplegia linked to the SPG11 locus (locates on 15q21.1)." 

"Here, for the first time, we report two cases, the first one which contains likely pathogenic NM_002860: c.475C>T: p.R159X mutation of the ALDH18A1 and the second one has likely pathogenic NM_001160227.2: c.5454dupA: p.Glu1819Argfs Ter11 mutation of the SPG11 gene and also was identified by the whole-exome sequencing and confirmed by Sanger sequencing." 

How these excerpts relate to the claim and why relevance is zero:
- None of the sentences mention SLC30A4 (gene name, aliases, locus, or variants). Therefore there is no direct evidence in this paper that SLC30A4 is associated with any Mendelian disease.
- The paper reports identified variants and mechanistic/diagnostic sequencing for ALDH18A1 and SPG11, not SLC30A4, so it provides no mechanistic pathway or functional data relevant to SLC30A4.

Limitations and caveats regarding this assessment:
- The conclusion that this paper is irrelevant to SLC30A4 is based solely on the supplied content excerpt. If other sections of the full paper (not provided here) mention SLC30A4, those were not available for review.
- Absence of evidence in this paper is not evidence of absence overall; this single paper neither supports nor disproves the claim about SLC30A4 beyond showing it does not address SLC30A4.

Because the manuscript excerpt contains no mention or data relating to SLC30A4, the relevance to the claim "The gene SLC30A4 is likely associated with at least one Mendelian disease" is 0.0.


[Read Paper](https://www.semanticscholar.org/paper/75a201bc47209dabc2b08a6a6888f29571c37a97)


### Abstract 4289: Whole-exome sequencing study of nevoid basal cell carcinoma syndrome (NBCCS) families to identify disease-causing exonic mutations

**Why Not Relevant**: The paper is an abstract focused on identifying PTCH1 mutations and potential novel NBCCS-associated genes via whole-exome sequencing. It does not mention SLC30A4 anywhere, does not provide a list of the novel genes discovered, and gives no variant-level data or mechanistic links involving SLC30A4. Because there is no direct or indirect reference to SLC30A4 (no variants, no functional/mechanistic discussion, no naming of candidate genes), this paper provides no evidence supporting or refuting the claim that SLC30A4 is likely associated with a Mendelian disease. Additional limitations: (1) the text is an abstract with limited detail, (2) where novel genes are referenced they are unnamed, preventing inference, and (3) the study focus is NBCCS/PTCH1/SHH pathway genes, which are mechanistically unrelated to SLC30A4 unless explicitly shown — which they did not do. For these reasons the relevance to the claim is effectively zero.


[Read Paper](https://www.semanticscholar.org/paper/f86e6deb7ee6eb11676ccc709b78849815be588e)


### Germline Variant Interpretation in Children with Severe Sepsis

**Why Not Relevant**: The provided paper content does not mention the gene SLC30A4 and contains no gene-level data or lists that would allow assessment of whether SLC30A4 is associated with any Mendelian disease. The only sentence provided states: "The authors report that 200/330 of children admitted to the PICU with severe sepsis who underwent exome sequencing and had a clinically significant genetic variant in a gene(s) associated with a known inborn error of immunity (IEI) that may have contributed to the patient’s clinical presentation." This sentence (a) does not name any specific genes, (b) provides no variant-level or gene-level evidence, and (c) offers no mechanistic discussion linking any specific gene to a Mendelian phenotype. Therefore there is neither direct evidence (reported pathogenic/likely pathogenic variants in SLC30A4) nor mechanistic evidence (functional data, pathway analyses, or clinical phenotype–genotype correlation) in the supplied content to support or refute the claim that "SLC30A4 is likely associated with at least one Mendelian disease." 

Limitations and implications: Because the study performed exome sequencing, it is theoretically possible that variants in SLC30A4 were observed but not reported in the supplied excerpt; however, without the full gene/variant list, pathogenicity assessments, segregation or functional data, or any mention of SLC30A4, no inference about SLC30A4 can be made from this content. To evaluate the claim properly, access to the full paper (including supplementary tables listing identified genes and classified variants), or independent resources such as OMIM/ClinVar/literature reporting SLC30A4 as causative for a Mendelian disorder, would be required.


[Read Paper](https://www.semanticscholar.org/paper/8148b944a985ef1eb6ab7a15ba1b7990c783459a)


### Mapping genetic variations to three-dimensional protein structures to enhance variant interpretation: a proposed framework

**Why Not Relevant**: The supplied paper content is a workshop/report and review about integrating genetic-variant data with 3D protein-structure information and describing tools, databases, and a proposed framework for interoperability. Nowhere in the provided text is the gene name SLC30A4 mentioned, nor are any specific gene-level or variant-level results reported that link any particular gene (including SLC30A4) to a Mendelian disease. The paper describes resources (for example, ClinVar, HGMD, ExAC/gnomAD), methods (mapping variants to 3D structures, homology modeling tools, visualization and variant-effect pipelines such as VIPUR, STRUM, CRAVAT), and general principles of variant interpretation, but it does not present direct evidence (case reports, segregation data, variant pathogenicity classifications) or mechanistic experimental data (functional assays, structural mapping for SLC30A4, protein-interaction or biochemical studies) that would support the claim that SLC30A4 is likely associated with at least one Mendelian disease.

Limitations and caveats relevant to this assessment:
- Absence of mention: The provided excerpt contains no mention of SLC30A4, no variant lists, and no disease associations; therefore there is no direct or indirect evidence in this text linking SLC30A4 to Mendelian disease.
- Paper type: The manuscript is a methods/workshop review proposing community standards and tools; such papers typically do not present new gene-disease associations but instead describe approaches that could be used to discover or interpret such associations.
- Resource pointers only: Although the paper cites databases (ClinVar, HGMD, ExAC/gnomAD) and tools that could be queried to investigate whether SLC30A4 has Mendelian associations, the paper itself does not perform or report such queries or analyses for that gene.
- Truncation caveat: Assessment is limited to the provided content. If other parts of the full paper (not provided here) contained gene-specific analyses or supplementary tables, those could change this conclusion; based on only the supplied text, no relevant evidence is present.

Conclusion: The provided paper content does not provide direct evidence nor gene-specific mechanistic evidence supporting the claim that SLC30A4 is likely associated with at least one Mendelian disease. The paper only describes methods and resources that could be used to investigate such a claim, but it does not itself supply data or findings about SLC30A4.


[Read Paper](https://www.semanticscholar.org/paper/ae68aa934d5a5ad3d9009bb8811fa8d5fc4dbb1c)


### THU0024 Whole genome linkage and exome sequencing analyses in an autosomal recessive takayasu arteritis family

**Why Not Relevant**: The paper does not mention SLC30A4 anywhere in the provided content. Its results focus on candidate variants identified in a single consanguineous family affected by Takayasu arteritis, listing ANXA8L, EHBP1L1, MYH14, KCNJ14, SYNGR4, TULP2 and SHANK1 as candidate genes. Therefore the paper provides no direct evidence that SLC30A4 is associated with any Mendelian disease, nor does it present mechanistic data linking SLC30A4 to disease. The absence of SLC30A4 in this study is also weak negative evidence because (1) the analysis was limited to one family and to variants within linkage regions and detectable by exome sequencing, (2) non-coding or structural variants would not necessarily be captured, and (3) a negative result in one disease/family does not rule out Mendelian associations of SLC30A4 in other families or diseases. Given these points, the paper is not relevant for supporting the claim that SLC30A4 is likely associated with at least one Mendelian disease.


[Read Paper](https://www.semanticscholar.org/paper/7df98d3323114688165f9b3dc23781f18758f1ed)


## Search Queries Used

- SLC30A4 ZnT4 solute carrier family 30 member 4 pathogenic variant Mendelian disease monogenic mutation clinical case report family segregation

- SLC30A4 ClinVar OMIM HGMD ClinGen variant annotation pathogenic likely pathogenic clinical significance database

- SLC30A4 ZnT4 knockout mouse rat zinc transporter lactation milk zinc deficiency phenotype loss of function mechanism zinc homeostasis

- SLC30A4 sequencing rare variant exome sequencing whole exome study rare disease familial segregation autosomal recessive autosomal dominant GWAS expression association


## Usage Statistics

- Prompt Tokens: 0

- Completion Tokens: 0

- Total Tokens: 0

- Estimated Cost: $0.0484


## Bibliometric Configuration

- Use Bibliometrics: True

- Author Impact Weight: 0.4

- Citation Impact Weight: 0.4

- Venue Impact Weight: 0.2
