## Supplementary material for "MENDELSEEK: An algorithm that predicts Mendelian Genes and elucidates what makes them special": SMPD2.txt

# Claim: The gene SMPD3 is likely associated with at least one Mendelian disease.

**Status**: processed

**Overall Rating**: -1

**Explanation**:

No relevant papers were found that support or refute this claim.


## Relevant Papers


## Other Reviewed Papers


### NGS based studies on primary immunodeficiencies (PIDs) : causative gene identification, tool development and application

**Why Not Relevant**: The provided paper content does not mention SMPD3 anywhere and focuses on genes implicated in primary immunodeficiency disorders (PIDs) discovered or evaluated in this cohort and analyses. Because SMPD3 is neither listed among the discovered candidate genes nor reported as having pathogenic variants or copy number defects in this study, the paper provides no direct evidence supporting an association between SMPD3 and any Mendelian disease. The negative/inconclusive information is indirect: absence of SMPD3 from reported results. Important caveats: the study is specialized to PIDs and to the specific cohorts, gene panels, and analytical approaches used; absence of SMPD3 here does not prove SMPD3 is not associated with a Mendelian disease in other contexts, cohorts, or phenotypes. Therefore the paper is largely irrelevant to the claim that "SMPD3 is likely associated with at least one Mendelian disease."


[Read Paper](https://www.semanticscholar.org/paper/7786e43fbaf68904111f286747f0cf8968c2dd1e)


## Search Queries Used

- SMPD3 neutral sphingomyelinase 2 SMPD3 mutation human patient case report Mendelian monogenic inherited disorder exome sequencing rare variant

- SMPD3 ClinVar OMIM DECIPHER HGMD pathogenic variant loss of function germline SMPD3 variant interpretation clinical significance

- SMPD3 knockout mouse fro phenotype bone mineralization skeletal dysplasia growth plate chondrocyte sphingolipid ceramide neutral sphingomyelinase 2 functional study

- neutral sphingomyelinase SMPD3 SMPD1 SMPD2 genetic disease review human monogenic sphingomyelinase deficiency rare variant burden GWAS population frequency penetrance expressivity


## Bibliometric Configuration

- Use Bibliometrics: True

- Author Impact Weight: 0.4

- Citation Impact Weight: 0.4

- Venue Impact Weight: 0.2
