## Supplementary material for "MENDELSEEK: An algorithm that predicts Mendelian Genes and elucidates what makes them special": SNCAIP.txt

# Claim: The gene SNCAIP is likely associated with at least one Mendelian disease.

**Status**: processed

**Overall Rating**: 0

**Explanation**:

Supporting evidence.
The provided papers establish that SNCAIP (synphilin-1) is biologically relevant to Parkinson's-disease-related pathways. Nuber et al. report that transgenic overexpression of synphilin-1 in mice produces behavioral and neuropathological alterations, implicating synphilin-1 in in vivo mechanisms of protein aggregation and neuronal cell death. Liu et al. show biochemical and genetic interaction between synphilin-1 and LRRK2: synphilin-1 co-immunoprecipitates with LRRK2, modifies LRRK2 aggregation and kinase activity, and alters LRRK2-driven neurotoxicity in cell and Drosophila models. Cheon et al. demonstrate that co-expression of alpha-synuclein and synphilin-1 promotes Lewy body–like inclusions and increases apoptotic death in human neural stem cells. Together these studies support a consistent functional role for SNCAIP in cellular and animal models relevant to Parkinsonian pathology.

Caveats or contradictory evidence.
None of the excerpts provide human genetic evidence that variants in SNCAIP cause a Mendelian disorder. The experimental data are largely based on overexpression, knockdown, or co-expression in cell culture, transgenic mice, and Drosophila, which speak to function and modifier effects but do not demonstrate segregation of pathogenic SNCAIP variants with disease in human pedigrees, de novo mutations, or other criteria for Mendelian causation. There are no cited reports of autosomal-dominant or autosomal-recessive human diseases directly caused by SNCAIP mutations in the provided material. Thus, while the gene is implicated in disease-relevant biology, direct evidence of SNCAIP as a Mendelian disease gene is absent in these excerpts.

Analysis of potential underlying mechanisms, deeper relationships, or second-order implications.
Mechanistically, SNCAIP interacts with alpha-synuclein and LRRK2—proteins that are themselves implicated in Mendelian forms of Parkinson's disease (e.g., SNCA duplications/mutations, LRRK2 mutations). This positions SNCAIP as a plausible modifier of protein aggregation, kinase activity, and neuronal survival pathways. Such a modifier role could influence penetrance, age at onset, or severity of Mendelian PD caused by other genes, without SNCAIP being the primary Mendelian cause. Also, findings that SNCAIP knockdown increases toxicity and that its expression can be protective argue for a modulatory rather than a primary causative role. Animal and cellular models can highlight pathogenic mechanisms but cannot substitute for human genetic evidence required to establish Mendelian disease association.

Explanation of rating.
Given the absence of human genetic data linking SNCAIP variants to a segregating Mendelian phenotype in the provided papers, the strongest defensible conclusion is that there is no evidence here to support the claim that SNCAIP is associated with at least one Mendelian disease. The literature excerpts do show plausible biological involvement in pathways relevant to Mendelian Parkinsonism, so the gene remains a candidate for modifier effects or a potential future Mendelian association, but the current evidence set does not demonstrate a causal Mendelian relationship. Therefore the appropriate rating is "No Evidence."


**Final Reasoning**:

The provided evidence demonstrates that SNCAIP (synphilin-1) is functionally connected to proteins and processes implicated in Parkinson's disease and can modify disease-relevant phenotypes in model systems. However, establishing a gene as causative for a Mendelian disease requires human genetic evidence (e.g., segregating pathogenic variants in families, recurrent pathogenic variants with appropriate statistical support, or de novo variants with strong functional correlation). The excerpts contain no such human genetic data linking SNCAIP variants to a Mendelian disorder. They instead show functional and modifier roles in models. Because the claim specifically asserts a likely association with at least one Mendelian disease and the provided papers do not present supporting human genetic evidence, the correct classification is "No Evidence." I reviewed the logic and find it consistent: biological implication ≠ proof of Mendelian causation, so absence of human genetic linkage equates to no evidence in support of the Mendelian claim.


## Relevant Papers


### Transgenic overexpression of the alpha-synuclein interacting protein synphilin-1 leads to behavioral and neuropathological alterations in mice

**Authors**: S. Nuber (H-index: 26), O. Riess (H-index: 83)

**Relevance**: 0.15

**Bibliometric Impact**: 0.48071428571428576


**Excerpts**:

- A pathological role of overexpressed synphilin-1 in vivo will help to further elucidate the mechanisms of protein aggregation and neuronal cell death in Parkinson’s disease.


**Explanations**:

- This sentence links synphilin-1 (the protein product of SNCAIP) to pathological processes relevant to Parkinson's disease, providing mechanistic evidence that altered levels of the protein can influence protein aggregation and neuronal death. This is mechanistic, not genetic, evidence: it supports biological plausibility that SNCAIP perturbation can affect disease-related pathways, but it does not provide direct evidence that variants in SNCAIP cause a Mendelian disease. Limitations: the statement is aspirational ("will help to further elucidate"), provides no empirical data, no human genetic or segregation evidence, no description of mutations, effect sizes, or statistical tests, and may reflect model-system overexpression that does not equate to pathogenic germline variants in humans. Therefore the excerpt modestly increases plausibility but is insufficient to support the claim that SNCAIP is likely associated with at least one Mendelian disease.


[Read Paper](https://www.semanticscholar.org/paper/8f9638d2c004ba06a926d991c5ff77b410111977)


### Synphilin-1 attenuates mutant LRRK2-induced neurodegeneration in Parkinson's disease models.

**Authors**: Jingnan Liu (H-index: 11), Wanli W. Smith (H-index: 31)

**Relevance**: 0.25

**Bibliometric Impact**: 0.30915000000000004


**Excerpts**:

- Mutations in leucine-rich repeat kinase 2 (LRRK2) cause autosomal-dominant Parkinsonism with pleomorphic pathology including deposits of aggregated protein and neuronal degeneration.

- Here, using co-immunoprecipitation, we found that LRRK2 interacted with synphilin-1 (SP1), a cytoplasmic protein that interacts with α-synuclein and has implications in PD pathogenesis.

- LRRK2 interacted with the N-terminus of SP1 whereas SP1 predominantly interacted with the C-terminus of LRRK2, including kinase domain.

- Co-expression of SP1 with LRRK2 increased LRRK2-induced cytoplasmic aggregation in cultured cells.

- Moreover, SP1 also attenuated mutant LRRK2-induced toxicity and reduced LRRK2 kinase activity in cultured cells.

- Knockdown of SP1 by siRNA enhanced LRRK2 neuronal toxicity.

- In vivo Drosophila studies, co-expression of SP1 and mutant G2019S-LRRK2 in double transgenic Drosophila increased survival and improved locomotor activity.

- Expression of SP1 protects against G2019S-LRRK2-induced dopamine neuron loss and reduced LRRK2 phosphorylation in double transgenic fly brains.

- Our findings demonstrate that SP1 attenuates mutant LRRK2-induced PD-like phenotypes and plays a neural protective role.


**Explanations**:

- This sentence establishes that mutations in LRRK2 are a cause of an autosomal-dominant (i.e., Mendelian) form of Parkinsonism. It provides important disease context linking LRRK2 to a Mendelian disorder, but it does not mention SNCAIP (which encodes synphilin-1) or provide any direct evidence that SNCAIP itself is mutated in or causative of a Mendelian disease. This is indirect context only: it makes SNCAIP relevant to Mendelian disease discussion because of its interaction partner, but is not direct genetic evidence for SNCAIP.

- This is direct mechanistic evidence from the paper that synphilin-1 (SP1, product of SNCAIP) physically interacts with LRRK2. Because LRRK2 mutations cause Mendelian Parkinsonism, an interaction between LRRK2 and SNCAIP protein provides mechanistic plausibility that SNCAIP could influence LRRK2-linked disease processes. Limitation: a biochemical interaction alone does not demonstrate that SNCAIP variants cause a Mendelian disease in humans.

- This excerpt gives more detailed mechanistic mapping of the interaction domains (N-terminus of SP1 with C-terminus/kinase domain of LRRK2). It strengthens plausibility that SP1 can modulate LRRK2 function (e.g., via the kinase domain). Limitation: domain-level binding supports mechanism but still does not provide genetic evidence that SNCAIP is a Mendelian disease gene.

- This experimental observation shows that co-expression of SP1 modifies a cellular phenotype (increased LRRK2-induced cytoplasmic aggregation). It is mechanistic evidence that SNCAIP expression affects LRRK2-related pathological processes (aggregation). Limitation: overexpression in cultured cells may not reflect endogenous physiological or pathogenic conditions; increased aggregation is a disease-relevant phenotype but not proof that SNCAIP mutations cause Mendelian disease.

- This sentence reports that SP1 reduces mutant LRRK2 toxicity and reduces LRRK2 kinase activity in cells. Mechanistically, this supports a functional role for synphilin-1 in modulating the pathogenic activity of a Mendelian PD protein (LRRK2). Limitation: this is cell-based functional modulation, not human genetic linkage or mutation data implicating SNCAIP as causative in a Mendelian disorder.

- This result — that siRNA knockdown of SP1 enhances LRRK2 neuronal toxicity — provides loss-of-function evidence that reducing synphilin-1 makes LRRK2-related toxicity worse. Mechanistically this suggests SNCAIP has a protective role; if translated to humans, SNCAIP loss-of-function could hypothetically influence disease risk or severity. Limitation: knockdown in cells is an experimental perturbation and is not equivalent to demonstrating that germline SNCAIP mutations produce Mendelian inheritance of disease.

- This in vivo Drosophila result shows that co-expression of SP1 with mutant LRRK2 ameliorates organismal phenotypes (survival and locomotor activity). It is mechanistic/functional evidence in an animal model that SNCAIP can modify LRRK2-driven, Mendelian-disease-relevant phenotypes. Limitation: Drosophila models and transgenic co-expression may not reflect human genetics or causation; rescue by overexpression does not equate to SNCAIP being a Mendelian disease gene.

- This sentence documents protection against dopamine neuron loss and reduced LRRK2 phosphorylation in fly brains when SP1 is expressed, further supporting a functional, mechanistic role for synphilin-1 in modulating LRRK2 pathogenicity. Limitation: again, model-organism functional evidence supports plausibility but is not direct evidence of SNCAIP causing or being linked to a Mendelian disease in humans.

- This concluding statement summarizes the authors' interpretation that SP1 attenuates mutant LRRK2-induced PD-like phenotypes and is neuroprotective. It synthesizes the mechanistic and functional data linking SNCAIP protein activity to a Mendelian PD protein's phenotypes. Limitation: summary of experimental results; it does not present human genetic evidence that SNCAIP mutations cause a Mendelian disease.


[Read Paper](https://www.semanticscholar.org/paper/002189df834330724f4acb87de5253ed0aca3d90)


### Inclusion Body Formation and Apoptotic Cell Death in the Human Neural Stem Cells HB1.F3 Following Gene Transfection of Alpha-Synuclein and Synphilin-1

**Authors**: S. Cheon (H-index: 17), Seung-Up Kim (H-index: 23)

**Relevance**: 0.15

**Bibliometric Impact**: 0.16002352941176473


**Excerpts**:

- Background: The etiology of Parkinson’s disease (PD) has not been established, but familial forms of the disease have some clues for its pathogenesis.

- Autosomal dominantly inherited familial PD induced by aberrations of the alpha-synucein gene has been known as a genetic model of PD and sheds light to the understanding of PD pathogenesis.

- Synphilin-1 is a protein which interacts with alpha-synuclein and constitutes the Lewy body.

- Methods: Immortalized human neural stem cells were transfected with the alpha-synuclein gene and synphilin-1 gene, to define the role of Lewy body inclusions in neuronal cell death.

- Results: Human neural stem cells with Lewy body-like inclusions showed an increased apoptotic cell death compared to those with diffuse alpha-synuclein-positive and synphilin-1-positive reaction after transfection with the alpha-synuclein gene and synphilin-1 gene.

- Tyrosine hydroxylase over-expressing cells produced a high level of levodopa and showed a higher rate of the apoptotic marker.

- Conclusions: These results suggest that the formation of Lewy body-like inclusions by the over-expression of alpha-synuclein and synphilin-1 could be an underlying cause of apoptotic neuronal cell death and the dopaminergic cell might be more susceptible.


**Explanations**:

- Contextual background: this sentence situates the study within Parkinson's disease and notes that familial forms exist. It is contextual and not direct evidence about SNCAIP (the claim). Strength: frames relevance to genetic disease generally. Limitation: does not mention SNCAIP or provide any genetic (Mendelian) evidence.

- Contextual mechanistic/genetic precedent: this sentence states that alpha-synuclein gene aberrations can cause autosomal dominant familial PD (a Mendelian example). It is relevant only insofar as it establishes that proteins interacting with alpha-synuclein could plausibly be involved in Mendelian PD. Strength: provides a genetic precedent (alpha-synuclein) showing Mendelian etiology is possible for PD. Limitation: it refers to SNCA (alpha-synuclein) not SNCAIP (synphilin-1), so it is indirect and not evidence that SNCAIP itself causes a Mendelian disease.

- Mechanistic connection: this is a direct statement that synphilin-1 (product of SNCAIP) interacts with alpha-synuclein and is a component of Lewy bodies. This is mechanistic evidence increasing biological plausibility that SNCAIP could be involved in PD-related pathology. Strength: identifies a molecular interaction and localization relevant to PD pathology. Limitation: an interaction / presence in aggregates does not constitute genetic evidence that SNCAIP variants cause Mendelian disease; no data on pathogenic variants or inheritance are provided.

- Experimental approach: this sentence describes that the authors over-expressed both alpha-synuclein and synphilin-1 in an immortalized human neural stem cell model to study Lewy-body-like inclusions. This is mechanistic/modeling evidence. Strength: shows the study directly interrogates synphilin-1 function in a cellular system. Limitations: over-expression in cell lines may not reflect effects of endogenous SNCAIP mutations in humans; transfection/over-expression models can produce non-physiologic artifacts and do not address inheritance or segregation in families.

- Experimental result (mechanistic): this reports that cells with Lewy body-like inclusions (after co-expression) had increased apoptosis compared with diffuse staining. This provides mechanistic evidence that synphilin-1 (with alpha-synuclein) can contribute to cellular toxicity. Strength: functional evidence that synphilin-1 participates in a pathogenic process relevant to PD. Limitations: result is from an in vitro over-expression system, not from human genetic studies; no data are provided on specific SNCAIP sequence variants or their segregation in pedigrees, so this is not direct evidence of a Mendelian disease association.

- Mechanistic detail relevant to cell-type susceptibility: this indicates that tyrosine hydroxylase over-expressing (dopaminergic-like) cells produce more levodopa and show higher apoptosis markers, suggesting dopaminergic cells may be more vulnerable. Strength: refines mechanistic plausibility for PD-relevant cell types. Limitation: still an in vitro observation under over-expression conditions; does not demonstrate SNCAIP mutation-driven Mendelian inheritance.

- Authors' interpretation (mechanistic inference): the conclusion explicitly proposes that over-expression-driven Lewy-body-like inclusions involving alpha-synuclein and synphilin-1 could underlie apoptotic neuronal death and that dopaminergic cells may be especially susceptible. Strength: summarizes mechanistic link between synphilin-1 and pathogenic processes in PD. Limitation: interpretive and mechanistic only; it does not provide genetic association, familial segregation, mutation identification, linkage or case reports required to support the specific claim that SNCAIP is likely associated with at least one Mendelian disease.


[Read Paper](https://www.semanticscholar.org/paper/04ab5f8834a4dfe4aefada49e3a6cb19d0247c66)


## Other Reviewed Papers


### Alpha-synuclein and Parkinson's disease


**Why Not Relevant**: The provided paper content discusses α-synuclein (SNCA) function and interactions but does not mention SNCAIP (synuclein alpha interacting protein) or any disease associations, Mendelian or otherwise. The claim concerns the gene SNCAIP being likely associated with at least one Mendelian disease. Because the excerpt neither names SNCAIP nor provides genetic association, segregation, pathogenic-variant, clinical, or inheritance evidence, it cannot be interpreted as support or refutation of the claim. Any inference linking this text to SNCAIP would be speculative and unsupported by the quoted content. Therefore the content is not relevant to assessing whether SNCAIP is associated with a Mendelian disease.


[Read Paper](https://www.semanticscholar.org/paper/e55f107f4be98d353e9e0227d24d2ac9d4369001)


### Synphilin-1 attenuates neuronal degeneration in the A53T alpha-synuclein transgenic mouse model.

**Why Not Relevant**: The paper provides mechanistic and in vivo mouse-model evidence that synphilin-1 (SNCAIP) interacts with alpha-synuclein and can modify alpha-synuclein-driven pathology. However, it contains no human genetic evidence (no reports of SNCAIP pathogenic variants, no segregation analyses, no case reports of Mendelian inheritance linked to SNCAIP). The experimental data are based on transgenic overexpression and modifier effects in mice (including double-transgenic contexts with an established SNCA A53T mutation), which inform biological plausibility but do not demonstrate that SNCAIP itself is causative of any Mendelian disease. Therefore the paper is of low relevance to the specific claim that 'SNCAIP is likely associated with at least one Mendelian disease.' Key limitations: lack of human variant data, absence of linkage/segregation or functional studies of patient variants, overexpression mouse models that do not model germline heterozygous pathogenic variants, and no statistical/genetic evidence presented to establish Mendelian association.


[Read Paper](https://www.semanticscholar.org/paper/e3fbcb48c7b08182da556f38bb9ccc1d9589c579)


### Simultaneous identification and prioritization of variants in familial, de novo, and somatic genetic disorders with VariantMaster

**Why Not Relevant**: The paper describes a variant prioritization/analysis tool (VariantMaster) and reports that it was applied to find causative variants in at least one rare Mendelian genetic disease. However, the provided content contains no mention of the gene SNCAIP, no specific variant(s) or gene-level findings, and no experimental or clinical evidence tying SNCAIP to any Mendelian disorder. Therefore the paper provides no direct evidence for or against the claim that SNCAIP is likely associated with at least one Mendelian disease. The only indirect relevance is that the described method could, in principle, be used to discover such an association if SNCAIP variants were present in analyzed cases; but without gene-specific results, this does not support the claim.


[Read Paper](https://www.semanticscholar.org/paper/a56e2dc7369c76b03ad5e9309093beb98a159719)


### The Burden of Candidate Pathogenic Variants for Kidney and Genitourinary Disorders Emerging From Exome Sequencing

**Why Not Relevant**: The provided paper content concerns the identification, curation, and population-level analysis of genes associated with Mendelian kidney and genitourinary disorders and discusses variant interpretation pipelines and database limitations. Nowhere in the supplied text is the gene symbol "SNCAIP" mentioned, nor are gene-level results or lists that include individual gene names provided in the excerpt. Because the claim concerns whether SNCAIP is likely associated with at least one Mendelian disease, and the excerpt contains no direct observations, analyses, or mechanistic discussion specific to SNCAIP, the paper content as provided offers no direct or mechanistic evidence for or against the claim. The only remotely relevant material is methodological: the authors created a curated list of 625 Mendelian kidney/genitourinary genes (accessible elsewhere) and applied variant-evaluation criteria. However, without the gene list or gene-specific results in the supplied content, it is impossible to determine whether SNCAIP was included in the curated set or whether any variants in SNCAIP were observed or interpreted. Therefore the relevance of this excerpt to the SNCAIP-specific claim is effectively zero.


[Read Paper](https://www.semanticscholar.org/paper/a75e297208c04992568e607d19aa9b9aaebd09d5)


### TTC7A Variants Previously Described to Cause Enteropathy Are Observed on a Single Haplotype and Appear Non-pathogenic in Pediatric Inflammatory Bowel Disease Patients

**Why Not Relevant**: The provided paper content addresses variants in TTC7A and the role of TTC7A variation in polygenic IBD. It contains no mention of SNCAIP (gene name, aliases, variants, phenotypes or mechanistic pathways), nor any discussion of Mendelian diseases linked to SNCAIP. Because the topic, gene, and phenotype focus are different (TTC7A and polygenic IBD versus SNCAIP and Mendelian disease), the paper content is not relevant to assessing whether SNCAIP is likely associated with at least one Mendelian disease. Additionally, the supplied text is only a short descriptive sentence rather than full methods, results, or discussion, so there is no hidden or indirect evidence within this content that could be reinterpreted to support the SNCAIP claim.


[Read Paper](https://www.semanticscholar.org/paper/3818a966b5d6e8b0eda65fdef043ac0727fa429c)


### Precision Medicine in Internal Medicine

**Why Not Relevant**: This paper is a general review of genetic testing strategies, variant interpretation, and the state of gene discovery as of 2019. It contains no gene-level data, case reports, variant listings, segregation analyses, functional studies, ClinVar/OMIM entries, or any explicit mention of the gene SNCAIP. Therefore it provides no direct evidence that SNCAIP is associated with any Mendelian disease. The excerpts supplied are background and methodological context (how exome sequencing can discover gene-disease links, the variant burden per exome, and how variants are classified), which bear only indirectly on the plausibility that any given gene (including SNCAIP) could be disease-associated. In short, the paper does not address SNCAIP specifically and cannot support or refute the claim; additional, gene-specific sources (OMIM entries, ClinVar records, peer-reviewed reports of pathogenic SNCAIP variants with segregation and/or functional validation) would be required to evaluate the claim.


[Read Paper](https://www.semanticscholar.org/paper/c7199e8a5226f8d9d55777da0e15e91797bb459a)


### Germline Variant Interpretation in Children with Severe Sepsis

**Why Not Relevant**: The provided paper content (a single summary sentence) reports aggregate results about variants in genes associated with inborn errors of immunity (IEI) in a pediatric sepsis cohort, but it does not mention SNCAIP by name, provide a gene list, or describe findings for non-IEI genes. SNCAIP (synuclein alpha interacting protein) is not typically categorized as an IEI gene; the cohort and analyses appear targeted to immunity-related Mendelian disorders. Consequently, there is no direct evidence (no cases, variant descriptions, segregation data, functional assays, or statistical association) nor mechanistic data in the provided content that link SNCAIP to a Mendelian disease. To evaluate the claim about SNCAIP specifically, one would need either: (a) the paper's gene-level results showing SNCAIP variants with pathogenic/likely pathogenic classification and supporting clinical/segregation/functional evidence, or (b) mechanistic data from the study showing how SNCAIP alteration causes a Mendelian phenotype. Neither is present in the supplied excerpt. Additionally, because only one summary sentence was provided and no methods, tables, or gene lists were available, generalizability and applicability to SNCAIP cannot be assessed. For these reasons the relevance of this paper content to the specific claim about SNCAIP is minimal. Recommended next steps: check the paper's full gene-level supplementary tables for any mention of SNCAIP; consult OMIM, ClinVar, and recent literature for established or candidate Mendelian associations of SNCAIP.


[Read Paper](https://www.semanticscholar.org/paper/8148b944a985ef1eb6ab7a15ba1b7990c783459a)


### OP0309 WES ANALYSIS OF RARE FAMILIES POINTS TO A PATHOGENIC ROLE FOR THE cGAS/STING-TYPE I IFN AXIS IN SSc

**Why Not Relevant**: The provided paper content does not mention the gene SNCAIP anywhere. The study focuses on whole-exome sequencing in five familial systemic sclerosis (SSc) families and highlights candidate genes and pathways (notably cGAS/STING–type I IFN signaling, and specific genes TMEM173/STING, NT5E/CD73, PRKD2). Although the paper states that "Mutations in two of our candidates cause Mendelian diseases," it does not name those genes as SNCAIP. Therefore the paper provides no direct evidence supporting or refuting the claim that SNCAIP is likely associated with at least one Mendelian disease. Absence of any mention of SNCAIP in the text means relevance to the claim is effectively zero for this source. Note: absence of mention in this single paper is not evidence that SNCAIP lacks Mendelian associations generally; it only shows this paper does not address SNCAIP.


[Read Paper](https://www.semanticscholar.org/paper/a5d1937aaf7ad488b3d5d4df3648ee2ca8966c0d)


## Search Queries Used

- SNCAIP synphilin 1 mutation variant family segregation exome sequencing Mendelian disease monogenic case report

- SNCAIP ClinVar OMIM HGMD pathogenic variant clinical annotation gene disease association SNCAIP synphilin 1

- SNCAIP synphilin 1 alpha synuclein interacting protein aggregation Parkinson disease familial parkinsonism autosomal dominant neurodegeneration mechanism

- SNCAIP loss of function intolerance gnomAD pLI constraint haploinsufficiency rare variant burden exome sequencing gene based association


## Usage Statistics

- Prompt Tokens: 0

- Completion Tokens: 0

- Total Tokens: 0

- Estimated Cost: $0.0529


## Bibliometric Configuration

- Use Bibliometrics: True

- Author Impact Weight: 0.4

- Citation Impact Weight: 0.4

- Venue Impact Weight: 0.2
