## Supplementary material for "MENDELSEEK: An algorithm that predicts Mendelian Genes and elucidates what makes them special": SOCS2.txt

# Claim: The gene SOCS2 is likely associated with at least one Mendelian disease.

**Status**: processed

**Overall Rating**: 3

**Explanation**:

Supporting evidence.
The provided papers show robust functional evidence that SOCS2 has potent, single-gene effects on growth-regulatory pathways in mammals. Metcalf et al. (cited in the first excerpt) created Socs2–/– mice which displayed a clear, highly penetrant gigantism phenotype with deregulated growth hormone (GH) and IGF-1 signalling; Greenhalgh et al. (cited) further linked this phenotype mechanistically to increased GH signalling. The second paper describes SOCS2 deletion altering pancreatic islet size and GH/PRL-STAT5 sensitivity and partially protecting mice from toxin-induced beta-cell destruction. Together these studies demonstrate that loss of SOCS2 function in mice produces a reproducible, Mendelian-like phenotype (recessive knockout effect) affecting endocrine growth and metabolism. The gene is mapped to human chromosome 12q, supporting straightforward translation of locus information to humans for variant screening.

Caveats or contradictory evidence.
None of the provided excerpts report a human Mendelian disorder caused by SOCS2 variation. The evidence is limited to rodent experimental knockouts and physiological/biochemical studies; animal-model phenotypes do not guarantee an analogous Mendelian disease in humans. The excerpts do not describe segregation of SOCS2 variants with disease in human families, case reports of pathogenic SOCS2 variants, or population genetics data (e.g., depletion of loss-of-function variants in humans) that would indicate SOCS2 is a human Mendelian disease gene. Thus direct human genetic evidence is absent in the provided material. Additionally, phenotypes in knockout mice can arise from species-specific developmental compensations or differences in physiology, and the mouse phenotype may reflect complete loss-of-function; heterozygous human variation, dominant negative effects, or compensation by paralogs could alter phenotypic outcomes.

Analysis of potential underlying mechanisms, deeper relationships, or second-order implications.
SOCS2 encodes a negative regulator of cytokine signalling (notably JAK-STAT) and appears to be a critical brake on GH/IGF-1 and prolactin signalling. Mechanistically, loss of SOCS2 increases sensitivity to GH/PRL signalling that then drives increased somatic growth and altered islet biology in mice. This mechanistic chain (loss of negative regulation → increased pathway activity → endocrine phenotypes) is exactly the type of single-gene causal mechanism that underlies many Mendelian disorders of signalling pathways (e.g., mutations in receptors, signalling mediators). Therefore, SOCS2 is a biologically plausible candidate for human monogenic disorders affecting growth regulation, GH/PRL sensitivity, or related metabolic phenotypes. However, plausibility alone does not equal demonstrated human genetic causality; a missing link is documented pathogenic human variants that segregate with a Mendelian phenotype. The mouse data do, however, provide strong functional grounds to prioritize SOCS2 in searches for undiagnosed Mendelian growth syndromes.

Explanation of rating.
The preponderance of evidence in the excerpts is experimental and from mouse models, showing that complete loss of SOCS2 function causes clear, single-gene phenotypes in rodents. That makes SOCS2 a strong candidate for causing a Mendelian phenotype. However, the provided corpus contains no direct evidence of SOCS2 causing a Mendelian disease in humans (no human variant-phenotype co-segregation, case reports, or clinical genetic studies are cited). Because the claim as stated—"SOCS2 is likely associated with at least one Mendelian disease"—implicitly refers to human Mendelian disease in typical use, the available evidence is suggestive but not confirmatory. Consequently the appropriate judgement is that evidence is mixed: convincing functional and animal-model support exists, but direct human genetic evidence is lacking in the provided materials.


**Final Reasoning**:

I reviewed the logic of the essay and stand by the assessment. The mouse knockout data provide compelling, single-gene functional evidence that SOCS2 loss produces a Mendelian-like phenotype in an experimental mammalian system; this supports biological plausibility that SOCS2 could underlie a human Mendelian disorder affecting growth or endocrine signalling. However, the provided excerpts contain no direct human genetic evidence (no pathogenic SOCS2 variants reported in families or patients) to confirm association with a human Mendelian disease. Because the claim, in standard interpretation, concerns human Mendelian disease and direct human evidence is absent, the evidence neither fully supports nor refutes the claim. That balance—strong functional/animal-model support but lacking human genetic confirmation—best maps to the "Mixed Evidence" rating.


## Relevant Papers


### Assignment1 of suppressor of cytokine signalling-2 (SOCS2) to porcine chromosome 5 with radiation hybrids

**Authors**: E. Piper (H-index: 15), C. Moran (H-index: 29)

**Relevance**: 0.45

**Bibliometric Impact**: 0.27610526315789474


**Excerpts**:

- SOCS (suppressor of cytokine signalling) proteins, also known as SSI (stat-induced stat-inhibitor) proteins, are negative regulators of cytokine-induced signal transduction.

- Cytokines regulate cellular behaviour by interacting with receptors on the plasma membrane of target cells and activating intracellular signal transduction cascades such as the JAK-STAT pathway (Krebs and Hilton, 2000.)

- Metcalf et al. (2000) generated mice unable to express Socs2, to examine the function of the gene in vivo.

- Socs2–/– mice grew significantly larger than their wildtype littermates (Metcalf et al., 2000).

- Characteristics of deregulated growth hormone and insulin-like growth factor-1 (IGF-1) signalling were observed in the Socs2–/– mice, indicating that SOCS2 may have an essential negative regulatory role in the growth hormone/IGF-1 pathway (Metcalf et al., 2000).

- Greenhalgh et al. (2003) provided compelling evidence that the gigantism displayed in the Socs2–/– phenotype is a result of negative regulation of growth hormone (GH) signalling by creating Socs2–/– mice that were also deficient in endogenous growth hormone production due to a point mutation in the GH-releasing hormone receptor.

- These Socs2–/–Ghrhrlit/lit mice displayed almost double the response to exogenous growth hormone administration as Socs2+/+Ghrhrlit/lit mice.

- Genome sequences show that SOCS2 maps to human chromosome 12q (GenBank accession number NM003877) and mouse chromosome 10 (GenBank accession number NM007706).

- SOCS2 has not previously been mapped in the pig, and this is the first step in further investigation into the gene’s action in a large production animal.


**Explanations**:

- Background mechanistic statement: Establishes that SOCS family proteins function as negative regulators of cytokine signalling. Relevance to the claim: mechanistic plausibility — if SOCS2 negatively regulates signalling, its dysfunction could plausibly produce a defined phenotype. Strengths: gives functional class. Limitations: generic to family members, does not demonstrate SOCS2-specific disease association or human data.

- Background mechanistic statement: Places SOCS activity in the JAK-STAT pathway context, a major signalling pathway implicated in several Mendelian immune and growth disorders. Relevance: mechanistic support that perturbation of SOCS2 could alter conserved signalling with potential pathogenic consequences. Limitations: pathway-level description only; no direct link to human monogenic disease is provided.

- Experimental-methods context: Reports generation of Socs2 loss-of-function mice. Relevance: indicates an in vivo loss-of-function model exists, enabling assessment of whether SOCS2 perturbation causes a discrete phenotype (a necessary piece of evidence when inferring potential Mendelian effects). Strengths: provides experimental system that produced a clear phenotype. Limitations: model organism evidence and complete knock-out may not reflect human allelic states (e.g., heterozygous variants or hypomorphic alleles).

- Direct experimental result (mechanistic/phenotypic): Shows that homozygous knockout mice have a robust, measurable phenotype (significantly larger growth). Relevance: supports that SOCS2 loss causes a strong, reproducible biological effect consistent with a monogenic phenotype in mice. Strengths: phenotypic magnitude is reported as statistically significant in the original study. Limitations: no p-values or effect-size numbers are given here; species differences may limit direct inference to human Mendelian disease; the phenotype arises from homozygous null animals, which may not correspond to human variant presentations.

- Mechanistic/phenotypic linkage: Connects the observed phenotype to deregulated GH/IGF-1 signalling and explicitly indicates SOCS2 as an essential negative regulator of that pathway. Relevance: strengthens biological plausibility that SOCS2 disruption can produce a defined endocrine phenotype (a mechanistic pathway commonly implicated in Mendelian growth disorders). Strengths: links molecular signalling perturbation to the phenotype. Limitations: observations are in mice; causality in humans is not shown; the excerpt reports interpretation ('indicating that') rather than direct human genetic evidence.

- Mechanistic experimental evidence: An interaction experiment where Socs2–/– animals were combined with a GH-deficient background, supporting that the gigantism in Socs2–/– is mediated via GH signalling. Relevance: strengthens the mechanistic chain (SOCS2 → negative GH signalling → growth phenotype), increasing plausibility that SOCS2 variants could cause a Mendelian growth disorder. Strengths: targeted genetic interaction provides compelling mechanistic causality in mice. Limitations: again, mouse model; the specific allele types and doses differ from human genetic variation; no human clinical cases are presented.

- Quantitative experimental result: Reports that the double-mutant mice have roughly doubled response to exogenous GH compared to controls. Relevance: provides functional magnitude of the signalling consequence of Socs2 loss in an experimental setting, bolstering mechanistic plausibility for a strong effect if SOCS2 is disrupted. Strengths: gives direction and relative magnitude of functional effect. Limitations: no absolute numbers, statistics, or translational data; an induced exogenous GH response in mice may not predict human disease manifestations or inheritance patterns.

- Genomic-context statement: Maps SOCS2 to human chromosome 12q and mouse chromosome 10. Relevance: confirms that a human ortholog exists and is locatable in the human genome, a prerequisite for human Mendelian genetics studies. Strengths: provides the human locus identifier (GenBank accession). Limitations: mapping alone does not indicate disease association; no human mutation or phenotype data are reported.

- Contextual/methodological statement: Notes absence of prior mapping in pig and frames the study as an initial step for investigating gene action in livestock. Relevance: minor — indicates the paper's scope is comparative/genomic mapping in animals rather than reporting human disease associations. Limitation: emphasizes that the work is not reporting human Mendelian disease connection.


[Read Paper](https://www.semanticscholar.org/paper/3e0c97da0e20069cef4ac07dde51bdeb53b848d5)


### Conditions associated with increased growth hormone and prolactin sensitivity

**Authors**: Amira Alkharusi (H-index: 8)

**Relevance**: 0.18

**Bibliometric Impact**: 0.064


**Excerpts**:

- A particular focus is put on two different proteins, Suppressors of Cytokine Signaling2 (SOCS2) and Tuberous Sclerosis Complex2 (TSC2) that uniquely impinge on JAK-STAT activation and on mTOR activation.

- Study I. We explored the influence of SOCS2 on glucose metabolism by using a mouse model of diabetes induced by multiple low dose streptozotocin (MLDSTZ).

- Pancreatic islets from untreated SOCS2 mice appeared larger than in wild-type (WT) controls, which could explain the augmented serum insulin levels observed in SOCS2 mice.

- Pancreatic islets, derived from SOCS2 mice showed increased GHR and PRLR staining, which suggest a higher sensitivity to GH/PRL-STAT5 signals in SOCS2 than in WT-derived β-cells.

- Our results suggest that SOCS2 ablation can partly overcome β-cell destruction induced by MLDSTZ.

- In summary, our studies indicate that the tissue sensitivity to GH/PRL is regulated by SOCS2 and TSC2 proteins. Since both SOCS2 and TSC2 have links to different disorders, an increased GH/PRL sensitivity in such conditions could play a functional role.

- I. Alkharusi A, Mirecki-Garrido M, Ma Z, Zadjali F, Flores-Morales A, Nystrm T, Castrillo A, Bjorklund A, Norstedt G, Fernandez-Prez L. Suppressor of cytokine signaling 2 (SOCS2) deletion protects against multiple low dose streptozotocin-induced type 1 diabetes in adult male mice. Horm Mol Biol Clin Investig. 2015 Nov 12, ahead-of-print/hmbci-2015-0036.


**Explanations**:

- Mechanistic relevance: This sentence places SOCS2 in the context of key intracellular signaling pathways (JAK-STAT and mTOR) that are often implicated in monogenic disorders. It supports biological plausibility that SOCS2 perturbation can have large physiological effects. Limitation: naming involvement in signaling pathways is mechanistic but does not constitute evidence that SOCS2 mutations cause a Mendelian disease in humans.

- Context / experimental evidence: States that the authors experimentally manipulated SOCS2 in a mouse model of chemically induced diabetes. This shows the paper studies disease-relevant phenotypes after SOCS2 perturbation, which is relevant mechanistic/disease-model evidence. Limitation: the model is an induced (toxin) model in mice, not a study of naturally occurring human Mendelian variants or inheritance.

- Direct experimental observation (mouse): Shows SOCS2 knockout mice have larger pancreatic islets and higher serum insulin, demonstrating physiological consequences of SOCS2 loss. This is strong mechanistic/disease-relevant evidence that SOCS2 influences pancreatic β-cell biology. Limitation: findings are in mice and demonstrate phenotype after experimental ablation, not documented human Mendelian disease caused by SOCS2 variants.

- Mechanistic evidence: Directly links SOCS2 ablation to increased receptor staining (GHR, PRLR) and suggests altered STAT5 signaling sensitivity. This gives a cellular mechanism by which SOCS2 loss alters hormone signaling and potentially causes disease phenotypes. Limitation: again, this is mechanistic in mouse cells/tissues and does not show causation of a Mendelian disorder in humans.

- Experimental conclusion (mouse model): States that SOCS2 ablation can ameliorate chemically induced β-cell destruction, showing a functional effect of SOCS2 loss on a disease model. This supports biological plausibility that SOCS2 influences disease susceptibility or progression. Limitation: protection in an induced model does not equate to SOCS2 causing a Mendelian disorder.

- Summary / contextual statement: Authors conclude SOCS2 and TSC2 regulate GH/PRL sensitivity and note links of both proteins to disorders, implying clinical relevance. This raises plausibility that SOCS2 alterations could be disease-associated. Limitation: the phrase 'links to different disorders' is nonspecific and does not identify any Mendelian disease directly attributable to SOCS2 in humans.

- Citation / study title: The paper title explicitly reports that SOCS2 deletion "protects against multiple low dose streptozotocin-induced type 1 diabetes in adult male mice," which is direct experimental evidence that SOCS2 loss changes disease phenotype in a mammalian model. Limitation: type 1 diabetes in this context is an induced model in mice and is not presented as a Mendelian disease caused by SOCS2 mutations in humans.


[Read Paper](https://www.semanticscholar.org/paper/4756b3cb45e9eb0d9b7110bff393dd5d817c70e8)


## Other Reviewed Papers


### Case Report: Expressive Speech Disorder in a Family as a Hallmark of 7q31 Deletion Involving the FOXP2 Gene

**Why Not Relevant**: The provided paper excerpt exclusively addresses FOXP2 gene variants and interstitial deletions at 7q31 as a cause of childhood apraxia of speech and related phenotypes. There is no mention of the gene SOCS2, no data on SOCS2 sequence or copy-number variants, no segregation or case reports implicating SOCS2, and no mechanistic discussion that would connect SOCS2 to a Mendelian phenotype. To support the claim that SOCS2 is likely associated with at least one Mendelian disease, one would expect to see (a) identification of rare, likely pathogenic variants in SOCS2 in affected individuals, (b) segregation of such variants with disease in families or consistent case series, (c) functional data showing a pathogenic mechanism attributable to SOCS2 perturbation, or (d) population genetics evidence (e.g., depletion of loss-of-function variation consistent with haploinsufficiency). None of these elements appear in the excerpt. Absence of evidence in this paper does not disprove SOCS2 involvement in Mendelian disease in general, but this paper provides no supporting or mechanistic evidence for the claim; therefore its relevance to the SOCS2 claim is negligible.


[Read Paper](https://www.semanticscholar.org/paper/0e512d4cda610883bca7a1967da817e8f9f22a21)


### Case Report: Identification of a de novo Missense Mutation in the F8 Gene, p.(Phe690Leu)/c.2070C > A, Causing Hemophilia A: A Case Report

**Why Not Relevant**: The paper focuses exclusively on a novel pathogenic mutation in the factor VIII gene (F8) causing hemophilia A and on preimplantation genetic testing for that condition. There is no mention of the SOCS2 gene, no data (genetic, clinical, experimental) concerning SOCS2, and no discussion of mechanisms involving SOCS2. Because the paper contains no direct observations, analyses, or mechanistic reasoning related to SOCS2, it cannot provide support for or against the claim that SOCS2 is associated with a Mendelian disease. The absence of SOCS2 information means this paper is not relevant to evaluating the claim.


[Read Paper](https://www.semanticscholar.org/paper/0af8dec1d6c21aa6c782c6b3779bba787bf4c3f3)


### [A rare variant in the TTR gene (p.E112K) is associated with systemic amyloidosis and a new symptom - skin hyperemia in response to ethanol intake: family segregation analysis, literature review, and a clinical case. Case report].

**Why Not Relevant**: This paper does not provide evidence for or against the claim that SOCS2 is likely associated with at least one Mendelian disease. The provided content exclusively discusses transthyretin amyloidosis (ATTR-amyloidosis) and variation in the TTR gene. For example, the paper states: "Transthyretin amyloidosis (ATTR-amyloidosis) is a systemic disorder associated with extracellular deposition in the tissues and organs of amyloid fibrils, transthyretin-containing insoluble protein-polysaccharide complexes." It further focuses on TTR variants: "To date, more than 140 variants in the TTR gene have been identified; however, most of them are described in single patients and do not have clear evidence of pathogenicity." The article is described as presenting a clinical case of ATTRv-amyloidosis associated with a rare pathogenic variant in the TTR gene and a literature review. Nowhere in the supplied text is SOCS2 mentioned, nor are there genetic, clinical, or mechanistic data relating SOCS2 to any Mendelian phenotype. Because the paper's scope is limited to TTR-related amyloidosis (case report and literature review), its methods, results, and conclusions cannot be used to support or refute associations between SOCS2 and Mendelian disease. Absence of SOCS2 in this paper is not evidence that SOCS2 is not associated with Mendelian disease; it simply means this paper is not relevant to that claim. To evaluate the claim about SOCS2, one would need papers that specifically report pathogenic variants in SOCS2 segregating with Mendelian phenotypes, functional/mechanistic studies of SOCS2 variants, or authoritative databases/curations (e.g., ClinVar, OMIM, gene-disease validity curation) that address SOCS2, none of which are present in the provided content.


[Read Paper](https://www.semanticscholar.org/paper/2567df8ee078bbbd995e907e0465d6337ffeeb26)


### Autosomal Dominant Intellectual Development Disorder-6 (MRD6) Without Seizures Linked to a De Novo Mutation in the grin2b Gene Revealed by Exome Sequencing: A Case Report of a Moroccan Child

**Why Not Relevant**: The paper exclusively reports a clinical case and genetic analysis implicating GRIN2B (encoding an NMDA receptor subunit) in autosomal dominant intellectual development disorder-6 (MRD6). There is no mention of SOCS2, no data (genetic, functional, or epidemiologic) concerning SOCS2, and no mechanistic discussion that would link SOCS2 to the described disease or to Mendelian disorders more generally. Because the content is entirely about a different gene (GRIN2B) and a specific patient-level GRIN2B pathogenic variant, it provides neither direct evidence (no SOCS2 variants, no segregation, no case reports) nor mechanistic evidence (no pathways, interactions, or functional assays involving SOCS2) relevant to the claim that SOCS2 is likely associated with a Mendelian disease. Additionally, the study type (single-patient case report) and its focus on GRIN2B mean that even indirect inference about other genes is not supported. Therefore the relevance to the SOCS2 claim is effectively zero.


[Read Paper](https://www.semanticscholar.org/paper/67d756e0cc302e5d46fdda9df34403f1f2de9a5a)


### O-287 Preimplantation genetic testing for monogenic disease (PGT-M) where results are only based upon analysis of linked polymorphisms/haplotypes risks serious diagnostic errors

**Why Not Relevant**: The provided paper content discusses preimplantation genetic testing (PGT-M) methods, the use of direct mutation testing (DMT) versus linkage-based approaches, and examples in which DMT prevented misdiagnosis. It does not mention the gene name SOCS2, nor does it present any data, results, or discussion linking SOCS2 to a Mendelian disease. The text is methodological and aggregate (covering 312 mutations in 116 genes) but does not list gene identities or report gene-specific associations. Therefore, there is no direct evidence in this paper supporting or refuting the claim that SOCS2 is likely associated with at least one Mendelian disease. Any inference about SOCS2 based on this paper would be unsupported; to evaluate the claim one would need gene-specific studies (case reports, segregation analyses, population genetics, functional studies) or a gene list from this cohort that explicitly includes SOCS2.


[Read Paper](https://www.semanticscholar.org/paper/6491f889424bf8836ab418a0c7b6956290c35d6f)


## Search Queries Used

- SOCS2 mutation case report patient monogenic single gene disorder Mendelian disease family segregation

- Suppressor of cytokine signaling 2 SOCS2 JAK STAT growth hormone signaling IGF1 human mutation growth disorder stature

- SOCS2 loss of function variants ClinVar gnomAD constraint pLI haploinsufficiency pathogenic variants database

- SOCS2 knockout mouse phenotype functional study model organism human disease relevance review OMIM Mendelian


## Usage Statistics

- Prompt Tokens: 0

- Completion Tokens: 0

- Total Tokens: 0

- Estimated Cost: $0.0371


## Bibliometric Configuration

- Use Bibliometrics: True

- Author Impact Weight: 0.4

- Citation Impact Weight: 0.4

- Venue Impact Weight: 0.2
