## Supplementary material for "MENDELSEEK: An algorithm that predicts Mendelian Genes and elucidates what makes them special": SOCS3.txt

# Claim: The gene SOCS3 is likely associated with at least one Mendelian disease.

**Status**: processed

**Overall Rating**: 0

**Explanation**:

No relevant papers were found for this claim.


## Relevant Papers


## Search Queries Used

- SOCS3 germline mutation human case report familial autosomal dominant autosomal recessive Suppressor of cytokine signaling 3 Mendelian disease

- SOCS3 ClinVar OMIM Orphanet gene disease association Suppressor of cytokine signaling 3 pathogenic variant

- Suppressor of cytokine signaling 3 deficiency knockout mouse phenotype model organism cytokine signaling JAK STAT immune dysregulation perinatal lethality growth phenotype

- exome sequencing trio rare variant burden monogenic gene discovery early onset immunodeficiency auto inflammatory disorder SOCS3 Suppressor of cytokine signaling 3 de novo variant
