## Supplementary material for "MENDELSEEK: An algorithm that predicts Mendelian Genes and elucidates what makes them special": SPIB.txt

# Claim: The gene SPIB is likely associated with at least one Mendelian disease.

**Status**: processed

**Overall Rating**: -1

**Explanation**:

No relevant papers were found that support or refute this claim.


## Relevant Papers


## Other Reviewed Papers


### Whole exome sequencing identifies recessive germline mutations in FAM160A1 in familial NK/T cell lymphoma

**Why Not Relevant**: The provided paper content focuses on familial natural killer/T-cell lymphoma (NKTL) and reports novel recessive germline mutations in FAM160A1. There is no mention of the gene SPIB, no genetic association data, no functional or mechanistic discussion involving SPIB, and no evidence that SPIB is implicated in any Mendelian disease. Because the excerpt neither names nor implicates SPIB, and offers only findings about a different gene (FAM160A1) and phenotype (NKTL), it does not provide relevant direct or mechanistic evidence for the claim that SPIB is likely associated with at least one Mendelian disease. Note: this conclusion is limited to the content you provided. If the full article contains sections not included here that mention SPIB, those were not available for review and could change the relevance assessment.


[Read Paper](https://www.semanticscholar.org/paper/08388b16f9f94a55459297e29336a0634b1c17ec)


### From case studies to community knowledge base: MSeqDR provides a platform for the curation and genomic analysis of mitochondrial diseases

**Why Not Relevant**: This paper does not provide any direct or mechanistic evidence relevant to the claim that the gene SPIB is likely associated with at least one Mendelian disease. The text discusses mitochondrial disease, a newly reported nuclear disease gene (PMPCA), and the MSeqDR community resource for mitochondrial disease genes and variants, but SPIB is not mentioned anywhere in the provided content. There are no case reports, variant descriptions, functional assays, protein modeling, segregation data, or curated database entries in this text that reference SPIB or link SPIB to a Mendelian phenotype. The mechanisms described in the paper pertain to mitochondrial biology and the discovery/validation workflow for mitochondrial disease genes (e.g., massively parallel sequencing, protein modeling, functional analyses in patient fibroblasts, cDNA rescue) and do not implicate SPIB or provide mechanistic pathways connecting SPIB to mitochondrial or other Mendelian diseases. While the MSeqDR resource described could in principle contain entries for many genes, including nuclear genes, the paper provides no evidence that SPIB is among those genes or that it has any disease-associated variants. Limitations: because the analysis is restricted to the provided paper content, it is possible that SPIB is associated with disease in other literature or databases not included here. To evaluate the claim further, one should search primary disease databases and literature (e.g., OMIM, ClinVar, ClinGen, PubMed, and MSeqDR itself) for explicit SPIB disease associations, case reports, pathogenic variants, or expert gene-disease validity assessments.


[Read Paper](https://www.semanticscholar.org/paper/6f85c595adaca6963c30fea73bc37266fa78be56)


### Germline and Somatic Whole-Exome Sequencing Identifies New Candidate Genes Involved in Familial Predisposition to Serrated Polyposis Syndrome

**Why Not Relevant**: The paper focuses on identifying candidate germline genes predisposing to serrated polyposis syndrome (familial SPS) using whole-exome sequencing. The provided content explicitly lists the candidate genes identified (ANXA10, ASXL1, CFTR, DOT1L, HIC1, INO80, KLF3, MCM3AP, MCM8, PDLIM2, POLD1, TP53BP1, WNK2 and WRN) and does not mention SPIB anywhere. Therefore the paper provides no direct evidence (positive association, variant data, segregation, functional assay) or mechanistic evidence linking SPIB to any Mendelian disease. Absence of SPIB in this study is weak negative evidence only for SPIB involvement in familial SPS in this cohort and does not constitute evidence for or against SPIB being associated with other Mendelian diseases. Because the paper does not address SPIB at all, its relevance to the claim that "SPIB is likely associated with at least one Mendelian disease" is effectively zero.


[Read Paper](https://www.semanticscholar.org/paper/f986efea9a4a25f8c852540c4e0e82473d2812b8)


### Clingen Coagulation Factor Deficiency Variant Curation Expert Panel: Meeting the Need for Recommendations to Curate Variants in the Coagulation Factor Genes

**Why Not Relevant**: The provided paper excerpt exclusively discusses the Clinical Genome Resource (ClinGen) Coagulation Factor Deficiency Variant Curation Expert Panel (CFD-VCEP), specification of ACMG/AMP guidelines for variant curation, and pilot curation work on the coagulation genes F8 and F9 (Hemophilia A and B). There is no mention of the gene SPIB, no data on SPIB variants, no clinical reports, and no mechanistic discussion involving SPIB. Therefore the paper contains no direct evidence supporting or refuting the claim that SPIB is likely associated with at least one Mendelian disease. Note that absence of evidence in this paper is not evidence of absence: the lack of SPIB-related content here does not inform the true disease-association status of SPIB. To evaluate the claim you would need to consult resources that specifically report on SPIB (e.g., OMIM, ClinVar, ClinGen gene-disease validity curation, PubMed literature on SPIB variants and phenotype correlations).


[Read Paper](https://www.semanticscholar.org/paper/9ea8d4ce19622ae1c6b0cdfd40848c93199f6280)


### Novel Candidate loci and Pathogenic Germline Variants Involved in Familial Hematological Malignancies Revealed by Whole-Exome Sequencing

**Why Not Relevant**: The provided paper content does not mention the gene SPIB anywhere in the Simple Summary or Abstract. The study reports pathogenic or likely pathogenic germline variants in genes such as CHEK2, RAD54L, NFATC2, TC2N, GATA1, MSH4 and PRF1 and discusses whole-exome sequencing of sixteen individuals with familial or personal onco-hematological histories, but there is no direct evidence (variant findings, case reports, association statistics) nor any mechanistic discussion concerning SPIB. Therefore this paper provides no support for or against the claim that SPIB is likely associated with at least one Mendelian disease. Limitations: this conclusion is based solely on the text you supplied (Simple Summary and Abstract). If SPIB appears elsewhere in the full paper (main text, results, supplement) not provided here, that could alter the assessment. Based on the provided content alone, there are no excerpts to extract that bear on SPIB, and no mechanistic or statistical evidence relating SPIB to a Mendelian disorder. For a proper assessment of the claim, consult databases and literature focused on Mendelian gene–disease relationships (OMIM, ClinVar, HGMD, PubMed) for SPIB-specific variant and phenotype evidence.


[Read Paper](https://www.semanticscholar.org/paper/dd17d9125bd5ce91a41cc677b82cd494ab411d99)


### Somatic Mutation Screening in Adult Patients with Myeloid Malignancies Provides Insights into Likely Germline Variants

**Why Not Relevant**: This paper does not provide evidence for or against the claim that SPIB is likely associated with at least one Mendelian disease. The manuscript reports analyses of germline-associated variants in adult patients with AML/MDS and explicitly lists the genes observed (e.g., TP53, RUNX1, KRAS, PTPN11, DDX41, GATA2, CBL, CEBPA, NF1, BRCA1, ASXL1, FLT3, IDH1/2, NPM1, TET2, and splicing factors) but does not mention SPIB anywhere in the provided text. Limitations affecting potential negative inference: the somatic sequencing panel evolved over the study period and the authors excluded variants of uncertain significance; confirmatory germline testing was performed in only five patients. Therefore, absence of SPIB in this paper may reflect that SPIB was not included on the panels used, was not observed in this specific myeloid-neoplasm cohort, or was not among pathogenic/likely pathogenic calls, but it does not constitute evidence that SPIB is not associated with any Mendelian disease. To evaluate the claim about SPIB, one would need literature or databases that specifically report SPIB germline pathogenic variants, clinical case reports, OMIM entries, ClinVar submissions, or functional/mechanistic studies linking SPIB loss or mutation to a Mendelian phenotype.


[Read Paper](https://www.semanticscholar.org/paper/9e17b5c78f81cb7c08b9e3faf01919f391e308ae)


### Whole Genome Sequencing Uncovers a Rare Germline Variant in ATM Associated with Familial Myeloproliferative Neoplasms

**Why Not Relevant**: The supplied paper does not mention the gene SPIB anywhere in the Introduction, Methods, Results, or Conclusions. The study's aim, cohort, methods, and reported findings focus on identifying germline predisposition variants for familial myeloproliferative neoplasms and specifically report a candidate germline ATM variant. Therefore this paper provides no direct evidence that SPIB is associated with any Mendelian disease, nor mechanistic evidence linking SPIB to disease. Absence of SPIB in the report is not evidence that SPIB is not associated with a Mendelian disease — it only indicates SPIB was not implicated in this particular familial MPN WGS study of 67 individuals. Additional reasons the paper is not relevant: the sample is small and disease-specific (familial MPN), analyses and filtering were tailored to identify variants enriched in this cohort relative to gnomAD, and mechanistic follow-up focuses on ATM. To evaluate the claim about SPIB would require studies that specifically interrogate SPIB variants across Mendelian phenotypes, large-scale case series, family segregation analyses, ClinVar/OMIM entries, or functional studies of SPIB, none of which are present in this paper.


[Read Paper](https://www.semanticscholar.org/paper/8d7b0bab45f8b0f6be019446ee1a711b75df3a68)


## Search Queries Used

- SPIB Spi B germline mutation pathogenic Mendelian disease monogenic family case report segregation autosomal dominant autosomal recessive

- SPIB ClinVar OMIM HPO ClinGen pathogenic variant germline gene disease association review

- SPIB Spi B transcription factor ETS family plasmacytoid dendritic cells B cell development knockout mouse loss of function immunodeficiency mechanism

- SPIB somatic mutation lymphoma cancer hematologic malignancy germline predisposition exome sequencing cohort familial rare disease study


## Bibliometric Configuration

- Use Bibliometrics: True

- Author Impact Weight: 0.4

- Citation Impact Weight: 0.4

- Venue Impact Weight: 0.2
