## Supplementary material for "MENDELSEEK: An algorithm that predicts Mendelian Genes and elucidates what makes them special": ST6GAL1.txt

# Claim: The gene ST6GAL1 is likely associated with at least one Mendelian disease.

**Status**: processed

**Overall Rating**: -1

**Explanation**:

No relevant papers were found that support or refute this claim.


## Relevant Papers


## Other Reviewed Papers


### The role of BRCA status on prognosis in patients with triple-negative breast cancer

**Why Not Relevant**: The paper exclusively addresses BRCA1/2 dysfunction (germline/somatic BRCA1/2 mutations, BRCA1 promoter methylation, BRCA1 protein expression) and their prognostic significance in triple-negative breast cancer. There is no mention of the gene ST6GAL1, no genetic association analyses of ST6GAL1, and no mechanistic discussion involving ST6GAL1. Because the claim concerns whether ST6GAL1 is likely associated with at least one Mendelian disease, and the paper contains no information about ST6GAL1 or Mendelian disease associations for any gene other than BRCA1/2, the paper is irrelevant to the claim and provides neither supporting nor refuting evidence. Additionally, the paper's focus on cancer prognostic effects rather than germline Mendelian disease phenotypes further limits any possible indirect inference about ST6GAL1.


[Read Paper](https://www.semanticscholar.org/paper/4202a315a25e63ae40b09de375d30e02652a937b)


### Familial focal segmental glomerulosclerosis associated with an ACTN4 mutation and paternal germline mosaicism.

**Why Not Relevant**: The provided paper content does not mention ST6GAL1 and contains no data (clinical, genetic, or mechanistic) about ST6GAL1. Instead it focuses on ACTN4 and, secondarily, NPHS1. Representative verbatim sentences from the supplied text that demonstrate the paper's topical focus are:

"Mutations in the ACTN4 gene cause focal segmental glomerulosclerosis (FSGS), which shows autosomal dominant inheritance (Online Mendelian Inheritance in Man No. 603278, FSGS1)."

"Here, we report a familial case of FSGS1 in which 2 affected siblings showed unusual clinical, pathological, and genetic features."

"A novel ACTN4 mutation, p.Ser262Phe, was detected in the patients, and their father was found to have a germline mosaicism for the mutation."

"In addition, these siblings also had a heterozygous p.Thr5Met substitution in NPHS1, which encodes nephrin, although the functional significance of this substitution is unclear."

How this bears on the claim: because ST6GAL1 is not mentioned and no experimental or clinical data concerning ST6GAL1 are presented, the paper provides neither direct evidence (e.g., pathogenic variants, segregation with disease, case reports) nor mechanistic evidence (e.g., biochemical function, pathways connecting ST6GAL1 to a Mendelian phenotype) to support or refute the claim that "ST6GAL1 is likely associated with at least one Mendelian disease." 

Limitations relevant to this assessment: the paper is a focused case report/series about ACTN4-related FSGS and a possible NPHS1 variant; absence of ST6GAL1 data in this paper neither proves nor disproves any association for ST6GAL1 in general—it simply means this source is irrelevant to the claim. To evaluate the claim about ST6GAL1, one would need to review other literature or databases (e.g., OMIM, ClinVar, gene-disease curation resources) that specifically address ST6GAL1.


[Read Paper](https://www.semanticscholar.org/paper/16a08fd6ebb15730e4e5a44956e57231edeef62a)


### The First Case of an Infant with Familial A20 Haploinsufficiency in Korea

**Why Not Relevant**: The paper is a case report/series about haploinsufficiency of A20 (HA20) caused by pathogenic variants in the TNFAIP3 gene and does not mention ST6GAL1 anywhere in the provided content. Therefore it contains no direct or mechanistic evidence that ST6GAL1 is associated with a Mendelian disease. The available excerpts establish the gene implicated in the reported Mendelian condition (TNFAIP3), so the paper neither supports nor explores ST6GAL1; absence of mention means the paper is not relevant to the specific claim.


[Read Paper](https://www.semanticscholar.org/paper/2872dffb8e6deffd21f98650628f04ce654d53ff)


### Fanconi anaemia, BRCA2 and familial considerations – follow up on a previous case report

**Why Not Relevant**: The provided paper content exclusively discusses Fanconi anaemia and germline BRCA2 mutations (including family testing and variant interpretation). There is no mention of the gene ST6GAL1, no data (genetic, functional, segregation, or clinical) related to ST6GAL1, and no mechanistic discussion that would link ST6GAL1 to Mendelian disease. Because the paper's scope, methods, and results are restricted to BRCA2 and FA in this family case report, it provides no direct or mechanistic evidence supporting or refuting the claim that ST6GAL1 is associated with at least one Mendelian disease. Additional studies that explicitly examine ST6GAL1 (e.g., segregation in families, pathogenic variants in affected individuals, functional assays showing loss/gain of function causing a Mendelian phenotype) would be required to evaluate the claim.


[Read Paper](https://www.semanticscholar.org/paper/55a84a17c03762d80eac3a4ee4da2c55be58645a)


### British Society for Matrix Biology Autumn 2020 Meeting: “Basement Membranes in Health and Disease”

**Why Not Relevant**: I found no mention of the gene ST6GAL1, nor any human genetic or Mendelian-disease data, in the provided paper content. The text focuses on extracellular matrix biology (giantin, collagen I and IV, integrins, collagen VII, LAMB2, COL4A3/4/5, Col5a2, Col4a1 disease modifiers, etc.), animal models of ECM and basement-membrane disorders, fibrosis mechanisms, and related experimental studies. There are descriptions of knockout phenotypes in animal models, proteomic and RNAi screens, and disease models (e.g., DEB, Alport, Pierson syndrome), but none of these paragraphs references ST6GAL1, sialyltransferases, human patients with ST6GAL1 variants, or mechanistic pathways directly involving ST6GAL1. Therefore the paper provides neither direct evidence (e.g., human pathogenic variants, segregation, phenotype descriptions) nor mechanistic evidence (e.g., biochemical pathways, knockouts, or functional assays) linking ST6GAL1 to any Mendelian disease. Limitations: Because the supplied content appears to be an excerpt or composite of multiple sections, it is possible that ST6GAL1 could be discussed elsewhere in the full source; however, based solely on the text you provided, there is no relevant evidence to support or refute the claim that ST6GAL1 is likely associated with at least one Mendelian disease. Suggested next steps to find relevant evidence: query human-disease databases (OMIM, ClinVar, HGMD), search PubMed for "ST6GAL1 AND (mutation OR variant OR syndrome OR "Mendelian" OR "heritable"), and inspect population-constraint metrics (gnomAD) and any reported clinical case reports or functional studies of ST6GAL1 variants.


[Read Paper](https://www.semanticscholar.org/paper/e27ef31faa9280438adb116ad5e8faaf9838ff15)


## Search Queries Used

- ST6GAL1 germline mutation monogenic disorder Mendelian disease case report familial inherited variant

- ST6GAL1 sialyltransferase 1 beta galactoside alpha 2 6 sialyltransferase congenital disorder of glycosylation inborn error of glycosylation

- ST6GAL1 loss of function knockout mouse phenotype developmental defect immune deficiency growth abnormality zebrafish knockdown model

- ST6GAL1 somatic mutation cancer expression ClinVar OMIM gene disease association review systematic review meta analysis


## Bibliometric Configuration

- Use Bibliometrics: True

- Author Impact Weight: 0.4

- Citation Impact Weight: 0.4

- Venue Impact Weight: 0.2
