## Supplementary material for "MENDELSEEK: An algorithm that predicts Mendelian Genes and elucidates what makes them special": SULT1A1.txt

# Claim: The gene SULT1A1 is likely associated with at least one Mendelian disease.

**Status**: processed

**Overall Rating**: -1

**Explanation**:

No relevant papers were found that support or refute this claim.


## Relevant Papers


## Other Reviewed Papers


### The Burden of Candidate Pathogenic Variants for Kidney and Genitourinary Disorders Emerging From Exome Sequencing

**Why Not Relevant**: The paper focuses on identifying and curating genes associated with Mendelian kidney and genitourinary disorders and on evaluating rates of candidate pathogenic variants in a cohort of ostensibly healthy adults. The text provided does not mention SULT1A1 or present data about that gene. All quoted material relates to the study's gene-selection methods, cohort, variant-annotation approach, and interpretation criteria within a kidney/genitourinary disease context. Because SULT1A1 is neither discussed nor listed in the provided excerpts, the paper provides no direct or mechanistic evidence for or against the claim that SULT1A1 is likely associated with at least one Mendelian disease. Additionally, absence of SULT1A1 from a kidney/genitourinary-specific curated gene list (if it is absent) would not be definitive evidence against its association with Mendelian disease in other phenotypic domains. Therefore the paper is minimally relevant to the claim.


[Read Paper](https://www.semanticscholar.org/paper/a75e297208c04992568e607d19aa9b9aaebd09d5)


### From case studies to community knowledge base: MSeqDR provides a platform for the curation and genomic analysis of mitochondrial diseases

**Why Not Relevant**: The paper's content is focused on mitochondrial diseases, example discovery of PMPCA as a nuclear-encoded mitochondrial disease gene, and the MSeqDR resource for curating mitochondrial disease genes and variants. The text contains no mention of the gene symbol SULT1A1, no presentation of variants, case reports, functional assays, or mechanistic data related to SULT1A1, and no claims that SULT1A1 is associated with any Mendelian disease. Therefore there is no direct evidence supporting or refuting the claim that SULT1A1 is likely associated with at least one Mendelian disease. The paper does describe a database (MSeqDR) that curates many genes and synchronizes with ClinVar, so it is theoretically possible that SULT1A1 data could be present in those resources, but the paper itself does not present or cite any such data. Absence of evidence in this paper is not evidence of absence overall; to evaluate the claim one must examine primary disease-gene reports, locus-specific databases, ClinVar/OMIM entries, or functional studies that specifically address SULT1A1.


[Read Paper](https://www.semanticscholar.org/paper/6f85c595adaca6963c30fea73bc37266fa78be56)


### [Clinical and genetic analysis of three children with Menkes disease due to variants of ATP7A gene].

**Why Not Relevant**: The paper focuses exclusively on Menkes disease and reports pathogenic variants in the ATP7A gene in three patients. Neither the gene name SULT1A1 nor any variants, analyses, mechanistic discussion, or database results concerning SULT1A1 are present in the provided text. Therefore the paper provides no direct genetic evidence or mechanistic data linking SULT1A1 to any Mendelian disease. Note: absence of mention in this paper does not disprove an association between SULT1A1 and Mendelian disease, but this publication neither supports nor evaluates that claim; it is entirely unrelated in its gene and disease focus.


[Read Paper](https://www.semanticscholar.org/paper/2c7af41359206fc3b0a3fce382c29206021a28c3)


### Autosomal Dominant Intellectual Development Disorder-6 (MRD6) Without Seizures Linked to a De Novo Mutation in the grin2b Gene Revealed by Exome Sequencing: A Case Report of a Moroccan Child

**Why Not Relevant**: The paper content exclusively discusses a case of autosomal dominant intellectual developmental disorder linked to a GRIN2B (grin2b) mutation (MRD6). There is no mention of the gene SULT1A1, no data, analyses, or mechanistic discussion related to SULT1A1, and no cross-references suggesting an association between SULT1A1 and any Mendelian disease. Because the claim concerns SULT1A1 specifically, this paper does not provide direct or mechanistic evidence to support or refute that claim. Additionally, the evidence presented is from a single-patient exome study focused on another gene, further limiting any indirect applicability.


[Read Paper](https://www.semanticscholar.org/paper/67d756e0cc302e5d46fdda9df34403f1f2de9a5a)


### #5782 AN EXCEPTIONAL CASE OF A PATIENT WITH TWO RARE GENETIC VARIANTS OF AUTOSOMAL DOMINANT POLYCYSTIC KIDNEY DISEASE AND ALPORT SYNDROME

**Why Not Relevant**: Relevance is extremely low because the paper contains no mention of the gene SULT1A1, no genetic data or clinical observations involving SULT1A1, and no mechanistic discussion linking SULT1A1 to any Mendelian disease. The paper is a single-patient case report focused on COL4A3 and GANAB variants in the context of Alport syndrome and ADPKD. While the provided excerpts show which genes were studied and the clinical-genetic conclusions drawn, none constitute direct or mechanistic evidence for SULT1A1. Important caveat: absence of SULT1A1 in this paper does not imply SULT1A1 is or is not associated with a Mendelian disease in the literature at large; it only means this specific paper provides no supporting or refuting evidence regarding the claim.


[Read Paper](https://www.semanticscholar.org/paper/0337e73775f38924bbc68438e6e09327983393d6)


### #1049 Genetic characteristics of autosomal dominant polycystic kidney disease in a Greek population

**Why Not Relevant**: This paper does not mention SULT1A1 anywhere in the text. It reports genetic findings primarily in PKD1 and PKD2 in a cohort of patients with autosomal dominant polycystic kidney disease, with a few variants in SEC61B and SLC3A1, and some undiagnosed cases despite WES/WGS. Although the study performed broad testing (tNGS of >600 genes and WES/WGS in a subset), it does not report any SULT1A1 variants or discuss SULT1A1-related mechanisms or diseases. Therefore the paper provides no direct evidence that SULT1A1 is associated with any Mendelian disease. Limitations to this conclusion: absence of mention is not definitive proof that SULT1A1 was not tested or that it has no Mendelian associations generally — the cohort is narrowly ascertained for ADPKD and the authors may not list every gene included in the tNGS panel. However, within the context and reported results of this study, there is no supporting or mechanistic evidence regarding SULT1A1 and Mendelian disease.


[Read Paper](https://www.semanticscholar.org/paper/809bc9272642701e2600d56210354f9dc489b442)


## Search Queries Used

- SULT1A1 mutation Mendelian disease monogenic autosomal recessive autosomal dominant case report family segregation

- SULT1A1 pathogenic variant ClinVar OMIM HGMD ClinGen pathogenicity annotation review

- SULT1A1 loss of function knockout mouse zebrafish inborn error of metabolism sulfotransferase deficiency steroid metabolism liver phenotype

- SULT1A1 polymorphism rs9282861 pharmacogenetics association study meta analysis review SULT1A gene family SULT1A2 SULT1A3 Mendelian evidence


## Bibliometric Configuration

- Use Bibliometrics: True

- Author Impact Weight: 0.4

- Citation Impact Weight: 0.4

- Venue Impact Weight: 0.2
