## Supplementary material for "MENDELSEEK: An algorithm that predicts Mendelian Genes and elucidates what makes them special": SV2B.txt

# Claim: The gene SV2B is likely associated with at least one Mendelian disease.

**Status**: processed

**Overall Rating**: -1

**Explanation**:

No relevant papers were found that support or refute this claim.


## Relevant Papers


## Other Reviewed Papers


### De novo mutation of NAXE (APOAIBP)-related early-onset progressive encephalopathy with brain edema and/or leukoencephalopathy-1: A case report

**Why Not Relevant**: The paper exclusively reports clinical and genetic findings relating to NAXE (APOA1BP) variants causing PEBEL1 and does not mention SV2B anywhere in the provided text. All quoted content documents identification of NAXE variants and interpretation of their pathogenicity; there is no direct evidence, mechanistic pathway, genetic association, or discussion regarding SV2B. Therefore this paper provides no support for the claim that SV2B is likely associated with at least one Mendelian disease. The absence of mention in this paper is not definitive proof that SV2B is not disease-associated, but it means this paper offers no relevant evidence for the claim.


[Read Paper](https://www.semanticscholar.org/paper/5a593d0a440a88cd34e0a92d055a797c68cbc328)


### De novo mutation of the TSC2 gene in patient with Tuberous Sclerosis Complex-Associated Neuropsychiatric Disorders (TAND) Phenotype: a case report

**Why Not Relevant**: The provided paper content exclusively addresses tuberous sclerosis complex and mutations in TSC1/TSC2. All genetic data, case description, and conclusions refer to a de novo TSC2 missense variant and related neuropsychiatric manifestations. There is no mention of the gene SV2B, no reported SV2B variants, no functional or mechanistic data bearing on SV2B, and no discussion that would allow inference about SV2B’s involvement in Mendelian disease. Therefore the paper provides no direct evidence (no reported association, no case reports, no segregation data) and no mechanistic evidence (no functional studies, pathways, or biological rationale) relevant to the claim that SV2B is likely associated with at least one Mendelian disease. Absence of evidence in this single paper is not evidence of absence overall; to evaluate the claim about SV2B one would need to search other genetic case reports, variant databases, functional studies, or gene-disease curation resources. Given the complete lack of SV2B content in the provided material, relevance is 0.0.


[Read Paper](https://www.semanticscholar.org/paper/cf2a073be9de76eba9b581cec01dd4c9adc163fb)


### Case Report: A novel de novo variant of COL1A1 in fetal genetic osteogenesis imperfecta

**Why Not Relevant**: This paper is not relevant to the claim that SV2B is likely associated with at least one Mendelian disease. The study reports a single fetal case of osteogenesis imperfecta and identifies a novel de novo variant in COL1A1 (c.1309G>A, p.Gly437Ser) as the likely genetic cause. The manuscript excerpts provided make no mention of SV2B, no variants in SV2B are reported, and no mechanistic or clinical data tie SV2B to the OI phenotype or any other Mendelian condition. The study design (trio whole-exome sequencing with Sanger validation and bioinformatic predictions) is appropriate to discover pathogenic variants, but the authors present COL1A1 as the causal gene in this case and do not present negative or positive findings about SV2B. Therefore, this paper provides neither direct evidence (e.g., SV2B variants segregating with disease, case reports, or association statistics) nor mechanistic evidence (functional assays, biological pathways linking SV2B to a Mendelian phenotype) supporting the claim. 

Caveat: while this paper cannot be used to support the association of SV2B with Mendelian disease, it also cannot be used to definitively disprove such an association in general; that would require targeted studies of SV2B variants across affected individuals, segregation analyses, functional validation, or literature demonstrating lack of pathogenic variants in sufficiently powered cohorts. None of those are present here.


[Read Paper](https://www.semanticscholar.org/paper/1caeaf6717a5af526be5f1f0dcef41ef4c377699)


## Search Queries Used

- SV2B germline mutation Mendelian disease monogenic case report whole exome sequencing de novo variant clinical phenotype

- SV2B ClinVar OMIM DECIPHER gnomAD ExAC pathogenic likely pathogenic loss of function intolerance pLI LOEUF population frequency rare variants

- synaptic vesicle glycoprotein 2B SV2B function synaptic vesicle trafficking neurotransmitter release knockout mouse zebrafish morpholino phenotype seizure neurodevelopmental delay

- SV2 family SV2A SV2C paralog functional redundancy compensatory expression paralog rescue epilepsy levetiracetam drug target absence of phenotype Mendelian disease


## Bibliometric Configuration

- Use Bibliometrics: True

- Author Impact Weight: 0.4

- Citation Impact Weight: 0.4

- Venue Impact Weight: 0.2
