## Supplementary material for "MENDELSEEK: An algorithm that predicts Mendelian Genes and elucidates what makes them special": TACC3.txt

# Claim: The gene NELL1 is likely associated with at least one Mendelian disease.

**Status**: processed

**Overall Rating**: -1

**Explanation**:

No relevant papers were found that support or refute this claim.


## Relevant Papers


## Other Reviewed Papers


### Whole-Exome Sequencing in Adults With Chronic Kidney Disease

**Why Not Relevant**: The provided paper excerpt does not mention the gene NELL1 anywhere and contains no data, variant calls, or discussion specific to NELL1. The study describes using whole-exome sequencing to evaluate Mendelian kidney disease genes and states that the authors compiled a list of 287 OMIM genes implicated in Mendelian kidney disorders (referred to as nephropathy genes), but the actual gene list (Supplement Table 3) is not included in the excerpt. Because NELL1 is neither named nor discussed in the supplied text, there is no direct evidence in this paper excerpt either supporting or refuting the claim that NELL1 is associated with a Mendelian disease. 

Additional context and limitations relevant to interpretation:
- The paper notes technical limitations of WES (“A known limitation of exome sequencing is that some segments of the genome are not amenable to capture (23). Among the 287 nephropathy genes, 29 were identified with at least 1 exon that is not captured by the Agilent kit, representing potential blind spots in the analysis (Supplement Table 3).”), which means even if NELL1 had been considered, negative findings could reflect capture/coverage gaps rather than true absence of pathogenic variation. This sentence is in the excerpt but does not mention NELL1 specifically. 
- The absence of any mention of NELL1 in the excerpt is not evidence that NELL1 is not a Mendelian gene; it only shows this paper (as provided) does not provide supporting data. 

To evaluate the claim properly, one would need to consult resources or publications that explicitly report pathogenic variants in NELL1 segregating with Mendelian phenotypes (e.g., OMIM, ClinVar, ClinGen gene-disease validity curation, or primary case reports), or examine the paper’s full Supplement Table 3 (if available) and the variant results to see whether NELL1 was included and whether pathogenic variants were observed.


[Read Paper](https://www.semanticscholar.org/paper/a29bd57964f68ac5f8e1abcafe1112ba488f53f5)


### Precision Medicine in Internal Medicine

**Why Not Relevant**: The provided paper content does not mention the gene NELL1, nor does it present any case reports, cohort analyses, variant-level data, segregation studies, functional assays, or statistical associations specific to NELL1. The text is a general overview of genetic testing strategies, capabilities of exome/genome sequencing, and challenges in variant interpretation, which is contextually relevant to how one would discover or validate a gene-disease association but supplies no direct or gene-specific mechanistic evidence for NELL1 being associated with any Mendelian disease. Therefore it cannot be used to support or refute the claim about NELL1; at best it explains the general processes and limitations that would govern such an inference. Additional primary literature, database entries (e.g., OMIM, ClinVar), or experimental/clinical reports specifically addressing NELL1 would be required to evaluate the claim.


[Read Paper](https://www.semanticscholar.org/paper/c7199e8a5226f8d9d55777da0e15e91797bb459a)


### BRCA 1 and BRCA 2 Mutation Status and Tumor Characteristics in Male Breast Cancer : A Population-based Study in Italy 1

**Why Not Relevant**: This paper does not mention NELL1 anywhere and exclusively analyzes BRCA1 and BRCA2 in a population-based series of 25 male breast cancer cases from Florence. The experimental focus (protein truncation test, SSCP, sequencing, primer extension assays for BRCA1/BRCA2 polymorphisms, LOH analyses at BRCA loci) is limited to BRCA1/BRCA2; no data, results, or discussion address NELL1, its variation, expression, or mechanistic role. Although the paper discusses general mechanisms relevant to Mendelian tumor-suppressor genes (e.g., loss-of-heterozygosity, transcript imbalance, double-hit/loss-of-function), these mechanistic points are made in the context of BRCA1/BRCA2 and cannot be extrapolated as direct or specific evidence that NELL1 is associated with any Mendelian disease. Additional limitations preventing relevance to the NELL1 claim: the sample size (n=25) and targeted gene panel approach mean only BRCA1/BRCA2 were interrogated; promoter, deep intronic, or other-gene analyses were not performed for genes outside BRCA loci; and no broader genomic or linkage analyses that could implicate other Mendelian genes (such as NELL1) were reported. Therefore this paper provides no direct evidence and no specific mechanistic evidence supporting or refuting the claim that NELL1 is likely associated with at least one Mendelian disease.


[Read Paper](https://www.semanticscholar.org/paper/d293621e8cb928a98993cfc5c513b4907b54bed9)


### TTC7A Variants Previously Described to Cause Enteropathy Are Observed on a Single Haplotype and Appear Non-pathogenic in Pediatric Inflammatory Bowel Disease Patients

**Why Not Relevant**: The provided paper content exclusively addresses TTC7A (Tetratricopeptide Repeat Domain 7A) variants and their role in polygenic IBD, with no mention of NELL1, its variants, clinical phenotypes, segregation data, functional studies, or mechanistic links to Mendelian disease. The claim concerns whether NELL1 is likely associated with at least one Mendelian disease; to evaluate that claim requires evidence such as case reports of NELL1 variants segregating with a Mendelian phenotype, functional validation showing pathogenic effect, or genetic studies linking NELL1 loss/gain of function to a Mendelian disorder. None of those types of evidence are present in this excerpt. Absence of mention of NELL1 in this TTC7A-focused study does not constitute evidence that NELL1 is not associated with a Mendelian disease; it simply means this paper is irrelevant to the claim. To assess the claim one would need to search the literature specifically for NELL1 and terms like "Mendelian", "monogenic", "biallelic", "dominant", "recessive", "segregation", "pathogenic variant", case series, or functional studies implicating NELL1 in a Mendelian phenotype.


[Read Paper](https://www.semanticscholar.org/paper/3818a966b5d6e8b0eda65fdef043ac0727fa429c)


### Abstract 4371: Integrated molecular characterization of pheochromocytoma and paraganglioma including a novel, recurrent and prognostic fusion gene

**Why Not Relevant**: This paper does not provide evidence for or against the claim that NELL1 is likely associated with at least one Mendelian disease. The provided abstract focuses on genomic characterization of pheochromocytoma and paraganglioma (PCC/PGL) and reports germline and somatic findings for a set of known and novel genes (e.g., RET, NF1, VHL, CSDE1, MAML3, BRAF, SDHB, ATRX). NELL1 is not mentioned anywhere in the text. Therefore the study does not supply direct evidence (observations of pathogenic germline variants in NELL1 in affected individuals) nor mechanistic evidence (functional data implicating NELL1 in a Mendelian phenotype). Absence of NELL1 from this PCC/PGL-focused cohort does not constitute evidence that NELL1 is not a Mendelian disease gene: the study is limited to one tumor type, includes 173 patients, and reports genes relevant to PCC/PGL specifically. To evaluate the claim about NELL1 and Mendelian disease association would require studies that (a) report segregating germline variants in NELL1 with Mendelian inheritance in pedigrees, (b) show enrichment of rare pathogenic NELL1 variants among affected individuals across relevant phenotypes, or (c) provide functional/mechanistic evidence linking NELL1 disruption to a heritable disease phenotype. None of those data appear in this paper.


[Read Paper](https://www.semanticscholar.org/paper/30985322b61bd7640c44402a4f336ac592eecba4)


### The simplest explanation does not have to be preferred: co-occurrence of pathogenic variants in cancer-predisposing genes

**Why Not Relevant**: The provided text does not mention NELL1 or any specific gene, nor does it present data about Mendelian diseases. It is a single-sentence fragment describing a general review/analysis of cancer phenotypes associated with MINAS (multiple inherited neoplasia alleles?) and use of the UK 10,000 Genomes Project to assess tumor predisposition syndromes or multiple primary tumors, and whether combined cancer-susceptibility genes (CSGs) act additively or synergistically. There is no empirical result, variant, segregation, genotype–phenotype correlation, functional assay, or statistical association reported for NELL1 or any other specific gene. Therefore this content provides neither direct evidence (e.g., identification of pathogenic NELL1 variants causing a Mendelian phenotype, segregation in families, or case reports) nor mechanistic evidence (e.g., protein function disruption, pathway perturbation) that would support or refute the claim that NELL1 is likely associated with at least one Mendelian disease.

Specific reasons for low relevance:
- No occurrence of the string "NELL1" or any synonym/alias in the provided text.
- No mention of Mendelian disease, monogenic inheritance, pathogenic/likely pathogenic variants, penetrance, or familial segregation.
- No gene-level data (variants, allele frequencies, functional impact) or statistical results linking any gene to disease.
- The text is methodological/contextual (describing review and dataset-checking) without results; thus it cannot support or contradict the claim.

Caveats:
- This assessment is limited to the single sentence provided. The full paper might contain gene-specific analyses (including NELL1) elsewhere; such content was not supplied and therefore could not be evaluated. Absence of evidence in this excerpt is not proof that the full paper lacks relevant information.


[Read Paper](https://www.semanticscholar.org/paper/a1372a664bd2c33a0bda638975a8b26d112457fc)


### PATH-44. THE LANDSCAPE OF SOMATIC MUTATIONS AND COPY NUMBER ALTERATIONS IN PRIMARY GLIOBLASTOMA IN JAPAN

**Why Not Relevant**: The supplied paper content does not mention the gene NELL1 anywhere. The text is a compilation of excerpts from studies of gliomas/GBMs and related tumors, focusing on genes and alterations such as MGMT, EGFR (including EGFRvIII and EGFR fusions), TERT promoter mutations, TP53, MSH2, MSH6, PTEN, CDK4, ALK, ROS1, MET, FGFR3-TACC3, ZMIZ1-FGFR2, BRAF, GOPC-ROS1, MZT2B-ALK, ETV6-NTRK3, FGFR1-TACC1, and MYD88, among others. Although the paper mentions that 13% of patients harbored pathogenic or likely pathogenic germline mutations in various genes, no excerpt names NELL1 or provides data about associations between NELL1 and any Mendelian phenotype. The study designs described are largely cancer-focused (targeted sequencing panels, somatic tumor sequencing, cfDNA from CSF), which primarily detect somatic driver mutations and selected germline variants relevant to cancer predisposition; these approaches and the reported results do not constitute evidence for or against a Mendelian disease association for NELL1. Therefore there is no direct evidence, no mechanistic discussion, and no contextual data in this paper content relevant to the claim that NELL1 is likely associated with at least one Mendelian disease.


[Read Paper](https://www.semanticscholar.org/paper/8efef8f78273fb4c9a760f73d0470645f96ecfc6)


### GENE-36. ACCURATE DETECTION OF TERT PROMOTER MUTATION IN GLIOMAS USING INFINIUM DNA METHYLATION ARRAYS IDENTIFIES NOVEL EPIGENETIC ASSOCIATION

**Why Not Relevant**: The provided paper content (conference abstracts) does not mention the gene NELL1, nor does it present any data on Mendelian (heritable, germline) diseases associated with NELL1. The abstracts focus on somatic driver mutations and tumor biology in gliomas (e.g., TP53, PTEN, EGFR, PDGFR, TERT promoter mutations), methylation-based prediction of TERT promoter mutation, NF2-fusion events in radiation-induced meningiomas, endothelial gene expression predictors of anti-angiogenic response (HGF, CXCL10), and an incomplete abstract title about unanticipated germline cancer susceptibility mutations — none of which include NELL1 or provide mechanistic or genetic evidence tying NELL1 to a Mendelian disorder. Specific reasons for assigning relevance = 0.0: (1) No occurrence of the string “NELL1” or any clear synonym in the provided text; (2) no descriptions of germline mutations implicating specific genes in Mendelian inheritance patterns related to NELL1; (3) all explicit genetic findings reported are somatic cancer drivers or epigenetic predictors unrelated to NELL1. Limitations: the material appears to be a set of conference abstracts and may be incomplete (one abstract is truncated), so it is possible that other parts of the conference proceedings mention NELL1. However, based solely on the text you provided, there is no direct or mechanistic evidence to support or refute the claim that NELL1 is associated with at least one Mendelian disease. Recommended next steps: search authoritative Mendelian-disease databases (OMIM, ClinVar, HGMD), and query PubMed for “NELL1 germline”, “NELL1 Mendelian”, or specific diseases known to involve NELL1 to locate any primary evidence if it exists.


[Read Paper](https://www.semanticscholar.org/paper/a7b62c8de45b10d34c9da87d119854fc11c9e55d)


### Study in Italy Characteristics in Male Breast Cancer : A Population-based Mutation Status and Tumor BRCA 2 and BRCA 1 Updated Version

**Why Not Relevant**: This paper provides no direct or mechanistic evidence regarding the gene NELL1 or its association with any Mendelian disease. The manuscript is focused entirely on BRCA1 and BRCA2 germ-line alterations, loss of heterozygosity, transcript imbalance, and clinicopathological features in a population-based series of 25 male breast cancer cases from Central Italy. There is no mention of NELL1, no data on NELL1 variants, no family segregation analysis, no functional assays involving NELL1, and no discussion that would link NELL1 to Mendelian inheritance or disease mechanisms. 

Although the paper does discuss general mechanistic concepts that are broadly relevant to gene-disease relationships (for example, the "double-hit/loss-of-function mechanism, with conversion to homozygosity often attributable to somatic loss of the wild-type allele", transcript imbalance, and the use of population-based mutation screening), these are presented specifically in the context of BRCA1/BRCA2 and male breast cancer. Such general mechanisms cannot be used to infer an association between NELL1 and a Mendelian disease without data specifically implicating NELL1 (e.g., pathogenic germline variants segregating with disease in families, clinical case series, functional evidence for pathogenicity, or database/epidemiological evidence). 

Therefore the paper is not relevant for evaluating the claim that NELL1 is likely associated with at least one Mendelian disease. To assess that claim, literature or database searches that specifically report NELL1 germline variants, family segregation, functional studies, or curated disease annotations (e.g., OMIM, ClinVar, HGMD, peer-reviewed case reports or gene-discovery studies) would be required.


[Read Paper](https://www.semanticscholar.org/paper/fbafe77cedd6f256d862a089ce92e835a7273225)


## Search Queries Used

- NELL1 mutation pathogenic variant Mendelian monogenic inherited disease familial segregation OMIM ClinVar case report

- NELL1 gene function Neural epidermal growth factor like 1 bone development osteogenesis craniosynostosis suture development extracellular signaling mouse knockout mechanistic study

- NELL1 ClinVar gnomAD LOEUF pLI loss of function intolerance population frequency variants pathogenic likely pathogenic variant interpretation

- NELL1 somatic mutation fusion cancer tumor expression methylation oncology recurrent alterations somatic versus germline


## Bibliometric Configuration

- Use Bibliometrics: True

- Author Impact Weight: 0.4

- Citation Impact Weight: 0.4

- Venue Impact Weight: 0.2
