## Supplementary material for "MENDELSEEK: An algorithm that predicts Mendelian Genes and elucidates what makes them special": TERF1.txt

# Claim: The gene TERF1 is likely associated with at least one Mendelian disease.

**Status**: processed

**Overall Rating**: -1

**Explanation**:

No relevant papers were found that support or refute this claim.


## Relevant Papers


## Other Reviewed Papers


### Whole Exome Sequencing Identifies a Causal RBM20 Mutation in a Large Pedigree With Familial Dilated Cardiomyopathy

**Why Not Relevant**: This paper does not provide evidence for or against the claim that TERF1 is associated with a Mendelian disease. The study reports whole exome sequencing in a large family with autosomal dominant dilated cardiomyopathy and identifies a segregating variant in RBM20 (c.1907 G>A) as the causal mutation. The paper's text (summary and methods/results provided) makes no mention of TERF1, its variants, nor any mechanistic data involving TERF1. Therefore there is no direct evidence (no identified TERF1 variant segregating with disease) and no mechanistic evidence (no functional, biochemical, or pathway data implicating TERF1) in this paper. Key points showing irrelevance: the conclusions state that the identified causal mutation is in RBM20 and the methodological description focuses on filtering and prioritization that led to RBM20. Because TERF1 is never discussed, the study cannot be used to support or refute an association between TERF1 and any Mendelian disorder. Note that absence of TERF1 in this paper is not evidence that TERF1 is not associated with Mendelian disease generally; it only means this paper contains no information bearing on that claim. The paper's limitations with respect to TERF1: (1) TERF1 is not queried or reported here, so there is no data; (2) the study is focused on a single pedigree and disease (familial dilated cardiomyopathy) and thus would only be informative for TERF1 if TERF1 variants appeared in their filtering/segregation results, which they did not. Because the paper contains no TERF1-relevant sentences or paragraphs, no verbatim excerpts are provided.


[Read Paper](https://www.semanticscholar.org/paper/64591300dba1c06e3f7216d58688ad7c6a3694be)


### Whole Exome Sequencing Identifies a Heterozygous Variant in the Cav1.3 Gene CACNA1D Associated with Familial Sinus Node Dysfunction and Focal Idiopathic Epilepsy

**Why Not Relevant**: The paper exclusively studies CACNA1D (encoding the Cav1.3 α1-subunit) and reports genetic and functional evidence linking a CACNA1D variant (p.Arg930His / R930H) to a syndromal phenotype involving sinus node dysfunction, epilepsy and ADHD. There is no mention of the gene TERF1, no data (genetic, functional, clinical, or mechanistic) concerning TERF1, and no pathway-level or homology-based analysis that would link findings about CACNA1D to TERF1. Therefore this paper provides no direct evidence that TERF1 is associated with any Mendelian disease, nor does it provide mechanistic evidence that would plausibly implicate TERF1. Because the subject gene, experimental results, and conclusions all pertain to CACNA1D rather than TERF1, the content is irrelevant to the claim that 'TERF1 is likely associated with at least one Mendelian disease.'


[Read Paper](https://www.semanticscholar.org/paper/66576bd7a92183605cc1096dcf44e7c5c33ae2f9)


### Whole-genome sequencing revealed armadillo repeat containing 5 (ARMC5) mutation in a Chinese family with ACTH-independent macronodular adrenal hyperplasia.

**Why Not Relevant**: The provided paper text does not mention TERF1 anywhere and instead focuses exclusively on ARMC5 as the causative gene for AIMAH/PMAH. Verbatim sentences from the paper illustrating this include: "ARMC5 was the causative gene of two pedigrees confirmed by whole-genome sequencing (WGA) and Sanger sequencing.", "The variant sites of the two families were c.C943T (p.R315W) and c.C1960T (p.R654X), respectively.", "Autosomal dominant inheritance of AIMAH was confirmed by genotypes of one family member.", "In addition, haploid clone of tumor DNA was confirmed by germline mutation and somatic mutation, which suggested the pathogenic mechanism of "two-hit-model."", and "ARMC5 was the causative gene of AIMAH pedigrees."  These sentences show the paper provides direct evidence that ARMC5 (not TERF1) is implicated in a Mendelian form of adrenal hyperplasia (AIMAH/PMAH), including identified germline variants, somatic second hits, and a described inheritance pattern. There is no direct evidence (no mention of TERF1 variants, association studies, functional data, or mechanistic links) connecting TERF1 to any Mendelian disease in the provided content. Limitation: absence of any mention of TERF1 in this single paper is not proof that TERF1 is not associated with Mendelian disease in other studies, but based on the supplied text alone there is no support—direct or mechanistic—for the claim that TERF1 is likely associated with at least one Mendelian disease. If you want evidence for or against TERF1 specifically, I can search databases (OMIM, ClinVar, PubMed) or analyze other papers that mention TERF1.


[Read Paper](https://www.semanticscholar.org/paper/d2cdb21fb1581c8746d51c15790d5e5fba6a09ca)


### A GP1BA Variant in a Czech Family with Monoallelic Bernard-Soulier Syndrome

**Why Not Relevant**: This paper provides no direct or mechanistic evidence relevant to the claim that TERF1 is likely associated with at least one Mendelian disease. The manuscript excerpt is focused on Bernard–Soulier syndrome (BSS) and a heterozygous GP1BA variant (p.C33Y) that segregates with macrothrombocytopenia in the studied family. For example, the paper states: "We have detected a heterozygous GP1BA likely pathogenic variant, causing monoallelic BSS." and "A germline GP1BA gene variant (NM_000173:c.98G > A:p.C33Y), segregating with the macrothrombocytopenia, was detected by whole-exome sequencing." These verbatim sentences demonstrate the study's topic and primary genetic finding. The excerpt further documents functional and structural follow-up on GPIbα: "In silico analysis of the protein structure of the novel GPIbα variant revealed a potential structural defect, which could impact proper protein folding and subsequent binding to VWF. Flow cytometry, immunoblot, and electron microscopy demonstrated further differences between p.C33Y GP1BA carriers and healthy controls." All quoted material concerns GP1BA/GPIbα biology and Bernard–Soulier syndrome; there is no mention of TERF1 (telomeric repeat binding factor 1), telomere biology, or any phenotypes or mechanisms plausibly linking TERF1 to a Mendelian disorder in the provided text.

Given that whole-exome sequencing was performed and the authors report a specific GP1BA variant that segregates with disease in the family, it is reasonable to infer that the authors prioritized and reported the variant they judged causative; if TERF1 had been implicated it would be expected to appear in the results or discussion. Limitations: the provided excerpt is only a portion of the paper (no page numbers were supplied), so if TERF1 were discussed elsewhere in the full text it would not be reflected here—but based on the content supplied there is no evidence for TERF1 involvement. Therefore the relevance of this paper to the claim about TERF1 is effectively zero.


[Read Paper](https://www.semanticscholar.org/paper/bd438ab5f15163f51f856681c5e4bb6fdb69b00e)


### Expanded Assessments of Germline Genetic Variants Identify Additional Risk Loci for Langerhans Cell Histiocytosis

**Why Not Relevant**: The provided paper text does not mention the gene TERF1, nor does it present data (genetic, functional, or clinical) that implicates TERF1 in Langerhans cell histiocytosis (LCH) or any Mendelian disease. The study focuses on somatic activating mutations in MAPK pathway genes (e.g., BRAF V600E), inherited variation in SMAD6, and a GWAS identifying other loci associated with LCH risk. Because TERF1 is not named and no genomic coordinates, linkage signals, variant lists, mechanistic pathways, or clinical phenotypes related to TERF1 are reported, there is no direct or mechanistic evidence in this paper to support or refute the claim that TERF1 is likely associated with at least one Mendelian disease. 

Limitations and context: Absence of evidence in this single LCH-focused study does not prove that TERF1 is not associated with a Mendelian disease; it only means this paper provides no relevant data. To evaluate the claim about TERF1, one should search genetic-disease databases (e.g., OMIM, ClinVar), pedigree-based studies, or publications specifically reporting pathogenic germline variants in TERF1 and associated Mendelian phenotypes. If you would like, I can perform a literature and database search summary for TERF1 and known Mendelian disease associations.


[Read Paper](https://www.semanticscholar.org/paper/3f0ed93d36e60c68c723f245c11e79fe7ed01112)


### Case report: Whole exome sequencing and genome-wide methylation profiling of Czech dysplasia in a Chinese pedigree

**Why Not Relevant**: The paper focuses exclusively on Czech dysplasia and identifies a COL2A1 missense mutation (R275C, c.823C > T) as the implicated variant in affected individuals. TERF1 is not mentioned anywhere in the provided content, and there is no experimental, genetic, or mechanistic data about TERF1. Therefore the paper provides no direct evidence that TERF1 is associated with any Mendelian disease, nor does it offer mechanistic pathways involving TERF1. Limitations: the absence of TERF1 in this report does not prove TERF1 is not associated with any Mendelian disease generally — the study is small (a case and relatives) and disease-specific, and the authors reported the COL2A1 finding from WES without presenting a comprehensive negative variant list. However, for the specific claim about TERF1 being associated with at least one Mendelian disease, this paper supplies no supporting evidence and no plausible mechanistic link.


[Read Paper](https://www.semanticscholar.org/paper/0b4c5d8d1c1628c4b97fb2a617e56922033a0080)


## Search Queries Used

- TERF1 TRF1 telomeric repeat binding factor 1 mutation Mendelian disease case report family segregation pathogenic variant OMIM ClinVar

- TERF1 germline variant whole exome sequencing whole genome sequencing rare variant family pedigree autosomal dominant autosomal recessive Mendelian disorder

- TERF1 shelterin telomere maintenance loss of function knockout mouse zebrafish telomere shortening bone marrow failure pulmonary fibrosis dyskeratosis congenita telomere biology disorder

- TERF1 population variation common variant genome wide association study GWAS no association variant frequency gnomAD ClinVar DECIPHER clinical significance pathogenicity


## Bibliometric Configuration

- Use Bibliometrics: True

- Author Impact Weight: 0.4

- Citation Impact Weight: 0.4

- Venue Impact Weight: 0.2
