## Supplementary material for "MENDELSEEK: An algorithm that predicts Mendelian Genes and elucidates what makes them special": TGFBR3.txt

# Claim: The gene TGFBR3 is likely associated with at least one Mendelian disease.

**Status**: processed

**Overall Rating**: -1

**Explanation**:

No relevant papers were found that support or refute this claim.


## Relevant Papers


## Other Reviewed Papers


### Germline RAD51C mutations confer susceptibility to ovarian cancer

**Why Not Relevant**: The provided paper content concerns RAD51C, not TGFBR3. The single sentence supplied states: "This work sequenced the full coding region and intronexon boundaries of RAD51C in 1,132 probands from families with a history of ovarian cancer occurring with or without breast cancer, and identified 12 mutations that result in premature protein truncation in cases compared to 1 such mutation in controls, which confirmed that the specific variant is rare but can seldom provide conclusive evidence of disease association." This sentence (and the paper it summarizes) addresses mutation findings and disease association for RAD51C in familial ovarian/breast cancer cohorts. It contains no mention of TGFBR3, no data on TGFBR3 variants, and no mechanistic discussion linking RAD51C findings to TGFBR3 function or disease association. Therefore the paper provides neither direct evidence (genetic association data for TGFBR3) nor mechanistic evidence (pathway, interaction, or functional data involving TGFBR3) relevant to the claim that "TGFBR3 is likely associated with at least one Mendelian disease." Any inference that results from RAD51C sequencing would be speculative with respect to TGFBR3 and is not supported by the text provided. Limitations that lead to the relevance score of 0.0: (1) different gene: RAD51C versus TGFBR3; (2) different disease context (familial ovarian/breast cancer study) with no reported links to TGFBR3; (3) absence of mechanistic or interaction data tying RAD51C findings to TGFBR3; (4) only a single sentence of the paper was provided, and that sentence contains no TGFBR3-relevant content. Because of these points, the content does not support or refute the claim about TGFBR3 and cannot be used as evidence for an association between TGFBR3 and any Mendelian disease.


[Read Paper](https://www.semanticscholar.org/paper/569ee9379a17e9a58c1fc5514086529d723d203f)


### A single center experience with publicly funded clinical exome sequencing for neurodevelopmental disorders or multiple congenital anomalies

**Why Not Relevant**: The single provided sentence ('The findings support the utility of clinical ES in a real-world setting, as part of a publicly funded genetic workup for individuals with GDD/ID and/or MCA.') does not mention the gene TGFBR3, any specific genes, any reported pathogenic or likely pathogenic variants, nor any Mendelian disease associations. There is no presentation of results, variant lists, case reports, segregation data, functional studies, or mechanistic discussion related to TGFBR3. Although clinical exome sequencing (ES) can in principle detect variants in many genes including TGFBR3, this paper content as provided contains no direct evidence (e.g., observed variants, statistical association, case descriptions) and no mechanistic evidence (e.g., biochemical pathway, functional assay) linking TGFBR3 to a Mendelian disorder. The excerpt also lacks methodological or sample-size details, so one cannot infer from it whether the authors examined TGFBR3 in their cohort or found any relevant findings. For these reasons the paper content provided is not relevant to the claim that 'TGFBR3 is likely associated with at least one Mendelian disease.'


[Read Paper](https://www.semanticscholar.org/paper/8ec4c0c03c573ad4019a6ee63b9197179a32b173)


### Precision Medicine in Internal Medicine

**Why Not Relevant**: This paper is a general review about genetic testing strategies, exome/genome sequencing, and the challenges of variant interpretation. It does not mention TGFBR3 or present any data (case reports, linkage, segregation, functional studies, variant pathogenicity evidence, or database annotations) specific to TGFBR3. Therefore it provides no direct evidence that TGFBR3 is associated with a Mendelian disease. The excerpts included only offer contextual or mechanistic reasons why previously unassociated genes (in general) might later be found to cause Mendelian disorders, but they cannot be used to assert that TGFBR3 specifically is likely disease-associated. The lack of any TGFBR3-specific content is the primary reason for the very low relevance score. Note also that statements are based on the state of knowledge as of 2019; absence of mention in this paper does not constitute evidence against an association found later or reported elsewhere.


[Read Paper](https://www.semanticscholar.org/paper/c7199e8a5226f8d9d55777da0e15e91797bb459a)


### Heterozygous rare variants in NR2F2 cause a recognizable multiple congenital anomaly syndrome with developmental delays

**Why Not Relevant**: The provided paper content describes evidence for rare, heterozygous NR2F2 variants causing a variable syndrome (including heart defects, developmental delay/intellectual disability, dysmorphic features, feeding difficulties, hypotonia, and genital anomalies) and provides clinical recommendations. It does not mention the gene TGFBR3 (or TGF-beta receptor 3) anywhere in the supplied text, nor does it present genetic, clinical, or mechanistic data linking TGFBR3 to a Mendelian disease. Therefore the paper provides no direct evidence for the claim that TGFBR3 is likely associated with at least one Mendelian disease. 

Mechanistic inference is also unsupported: the content centers on NR2F2 and its associated phenotype, and does not describe molecular pathways, protein interactions, or functional studies that would connect NR2F2 to TGFBR3 or imply pathogenicity of TGFBR3 variants. Without explicit mention of TGFBR3, shared loci, co-segregation, functional perturbation, or pathway-level links in the paper, no mechanistic evidence for TGFBR3 can be extracted. 

Limitations and caveats: absence of mention in this single paper is not evidence that TGFBR3 is not associated with Mendelian disease; it only means this particular paper does not provide supporting evidence. To evaluate the claim about TGFBR3, a targeted literature and database search (e.g., OMIM, ClinVar, HGMD, PubMed) for TGFBR3 and associated phenotypes/variants is required.


[Read Paper](https://www.semanticscholar.org/paper/b387c485f1ff97a98e73a500784107d513934e73)


### Whole-Exome Sequencing in Adults With Chronic Kidney Disease

**Why Not Relevant**: The provided paper content does not mention TGFBR3 (no occurrence of the gene symbol or full gene name). The manuscript fragment describes a study design and methods for whole-exome sequencing in adults with chronic kidney disease, including that the authors "developed a curated priority list of 287 Online Mendelian Inheritance in Man (OMIM) genes implicated in Mendelian forms of kidney disorders and hypertension" and that they reviewed variants in OMIM genes and the ACMG actionable gene list, but the content supplied does not include the gene list itself, any gene-level results, or any discussion of TGFBR3 specifically. Therefore there is no direct evidence in this text supporting or refuting the claim that TGFBR3 is likely associated with at least one Mendelian disease. Mechanistic evidence is likewise absent: the excerpt contains methodological descriptions (sequencing, variant annotation, interpretation workflow) and general statements about diagnostic yield in other contexts, but no molecular, functional, or clinical data concerning TGFBR3. Limitations that prevent establishing relevance from this paper fragment include (1) absence of the gene-level results or the explicit list of the 287 OMIM nephropathy genes in the provided text, (2) no reporting of detected variants tied to specific genes, and (3) no mechanistic experiments or literature review presented here that mention TGFBR3. If you want a definitive assessment of TGFBR3’s association with Mendelian disease, I can (a) search the full paper (if you can provide the full text or supplementary tables listing the 287 genes and results), or (b) query external resources (OMIM, ClinVar, HGMD, PubMed) and summarize evidence for TGFBR3 from the broader literature.


[Read Paper](https://www.semanticscholar.org/paper/a29bd57964f68ac5f8e1abcafe1112ba488f53f5)


### The simplest explanation does not have to be preferred: co-occurrence of pathogenic variants in cancer-predisposing genes

**Why Not Relevant**: The provided paper content is focused on the frequency and phenotypes of MINAS and on reviewing literature and UK 10,000 Genomes Project data for participants with tumor predisposition syndromes or multiple primary tumors, and on whether phenotypes are additive or synergistic across cancer susceptibility genes. It does not mention TGFBR3, present any data, statistics, variant-level findings, or mechanistic discussion specific to TGFBR3, nor does it report any Mendelian disease associations for any individual gene. Because the excerpt contains no gene-specific information, no results, and no mechanistic detail concerning TGFBR3, it cannot be used as evidence for or against the claim that TGFBR3 is likely associated with at least one Mendelian disease. Any inference that TGFBR3 might be included in the analyzed CSGs would be speculative without additional text listing genes or presenting results. Consequently the excerpt is not relevant to the claim.


[Read Paper](https://www.semanticscholar.org/paper/a1372a664bd2c33a0bda638975a8b26d112457fc)


### De novo HDAC3 variants leading to epigenetic machinery dysfunction are associated with a neurodevelopmental disorder

**Why Not Relevant**: The provided paper describes heterozygous de novo variants in HDAC3 and their association with a novel neurodevelopmental syndrome, including detailed genetic, biochemical, proteomic, and cellular evidence for HDAC3 dysfunction (e.g., identified variants D93N, A110T, P201S, L266S, G267S, and R359C; reduced deacetylation activity for L266S and G267S; disrupted interactions with CoREST and NCoR for A110T; and impaired nuclear localization for A110T, G267S, and R359C). Nowhere in the provided text is the gene TGFBR3 mentioned, nor are there genetic, segregation, statistical, or functional data concerning TGFBR3. Therefore this paper provides no direct evidence (no reported variants, no phenotypic correlations, no functional assays) and no mechanistic evidence connecting TGFBR3 to any Mendelian disease. Absence of mention in this paper is not evidence that TGFBR3 is not associated with Mendelian disease; it simply means this paper is irrelevant to the claim. To evaluate the claim about TGFBR3, one would need studies reporting pathogenic variants in TGFBR3 with segregation/occurrence in affected individuals, population-frequency data, and ideally supporting functional/mechanistic experiments — none of which are present in this HDAC3-focused paper.


[Read Paper](https://www.semanticscholar.org/paper/524b09f9e39f306b6871f816044577dccb53abd6)


### Abstract LB-372: Robust targeted sequencing strategy identifies highly penetrant mutations in hereditary colorectal cancer cases from Colon Cancer Family Registry

**Why Not Relevant**: The paper is a short conference abstract describing a targeted next-generation sequencing (NGS) pooling strategy applied to familial colorectal cancer (CRC) cases. Its aims and results are limited to identifying highly penetrant CRC-predisposing mutations within a predefined panel of 40 known or candidate CRC genes. The abstract names many genes in which novel deleterious variants were found (including TGFBR2) but does not mention TGFBR3 at all. Therefore it provides neither direct evidence that TGFBR3 is associated with any Mendelian disease nor mechanistic data about TGFBR3. Additional limitations reducing relevance: (1) the study is disease-specific (familial CRC) and not a general survey of Mendelian disease genes; (2) the gene panel was limited and may not have included TGFBR3, so absence of mention does not constitute evidence against association; (3) the abstract-level format provides limited methodological and variant-level detail (no variant-level pathogenicity evidence for genes beyond those listed); and (4) no functional or mechanistic assays regarding TGFBR3 are reported. For these reasons the paper is essentially not relevant to the claim that TGFBR3 is likely associated with at least one Mendelian disease.


[Read Paper](https://www.semanticscholar.org/paper/50c937f4f1eb7ceef5965bb0c11a3b8dae5f3da4)


### Identification of novel disease-causing variants in rare diseases using trio exome sequencing

**Why Not Relevant**: The provided paper content does not mention TGFBR3 anywhere and reports specific gene findings (GLRA1, DOLK, DMD, ATRN, NKX6-2, MED14, DRG1, GRIN1, GRIN2B) related to the cohorts studied. Therefore it contains no direct evidence that TGFBR3 is associated with any Mendelian disease, nor does it present mechanistic data (e.g., functional studies, pathways, or variant segregation) pertaining to TGFBR3. The absence of TGFBR3 in this small, phenotype-specific set of exome analyses cannot be taken as evidence against TGFBR3 involvement in Mendelian disease more broadly; it simply means this paper does not address the claim. For an assessment of the claim, literature specifically reporting TGFBR3 variants, segregation, functional assays, or disease cohorts should be consulted.


[Read Paper](https://www.semanticscholar.org/paper/8ae081628e0ab2aec44f8fe9990f230956fb40ba)


### Contribution of rare transmitted and de novo variants among 2 , 871 1 congenital heart disease probands

**Why Not Relevant**: The provided paper excerpt contains no mention of the gene TGFBR3, nor any explicit data, gene lists, tables, or supplementary text naming TGFBR3. The manuscript discusses broad results about congenital heart disease (CHD), numbers of implicated genes, methods, and enrichment analyses, but the specific gene TGFBR3 is not present in the supplied content. Without any verbatim mention or data for TGFBR3 (for example, variant counts, enrichment statistics, de novo or recessive genotypes, or mechanistic descriptions involving TGFBR3), this excerpt provides no direct or mechanistic evidence to support or refute the claim that “TGFBR3 is likely associated with at least one Mendelian disease.” It is possible that TGFBR3 could appear in supplementary materials (e.g., Supplementary Excel files) or elsewhere in the full paper, but those materials were not provided here; therefore we cannot infer inclusion from the summary text. To evaluate the claim based on this paper would require explicit sentences, tables, or supplementary entries naming TGFBR3 and describing observed pathogenic variants, segregation, statistical significance, or mechanistic data linking TGFBR3 to a Mendelian phenotype. In their absence, relevance is effectively zero.


[Read Paper](https://www.semanticscholar.org/paper/e5830d9e95469f22518560007746202e62db26a0)


## Search Queries Used

- TGFBR3 germline mutation pathogenic variant Mendelian disease case report familial segregation OMIM ClinVar

- TGFBR3 de novo variant exome sequencing trio rare disease cohort congenital anomaly developmental disorder intellectual disability congenital heart disease

- TGFBR3 loss of function knockout mouse phenotype haploinsufficiency TGF beta signaling SMAD aortic aneurysm connective tissue disorder developmental biology

- TGFBR3 gene disease association review systematic review ClinGen Orphanet HGMD ClinVar gene curation dosage sensitivity haploinsufficiency


## Bibliometric Configuration

- Use Bibliometrics: True

- Author Impact Weight: 0.4

- Citation Impact Weight: 0.4

- Venue Impact Weight: 0.2
