## Supplementary material for "MENDELSEEK: An algorithm that predicts Mendelian Genes and elucidates what makes them special": THBS1.txt

# Claim: The gene THBS4 is likely associated with at least one Mendelian disease.

**Status**: processed

**Overall Rating**: -1

**Explanation**:

No relevant papers were found that support or refute this claim.


## Relevant Papers


## Other Reviewed Papers


### Effective variant filtering and expected candidate variant yield in studies of rare human disease

**Why Not Relevant**: The provided paper content contains only a high-level methodological statement about variant filtering and expected numbers of candidate variants for different inheritance modes. There is no mention of THBS4, no presentation of variants in THBS4, no clinical case reports, no segregation data, no functional/biochemical assays, and no statistical association results linking THBS4 to any Mendelian phenotype. Therefore the content does not provide direct evidence for or against the claim that THBS4 is likely associated with at least one Mendelian disease. It is possible that the full paper (beyond the single supplied sentence) might include gene-level analyses or examples that mention THBS4, but based solely on the supplied content there is no relevant evidence. To evaluate the claim one would need excerpts showing: variant(s) in THBS4 co-segregating with disease in families, de novo pathogenic variants in THBS4 in affected individuals with appropriate filtering and validation, functional studies demonstrating a disease-causing mechanism for THBS4 variants, or curated clinical annotations (e.g., OMIM/ClinVar) linking THBS4 to Mendelian disorders. None of these are present in the provided content, which is why relevance is effectively zero.


[Read Paper](https://www.semanticscholar.org/paper/8da5eda5547e3521df7aecc8c8e4b7380a7641fa)


### Diagnostic power and clinical impact of exome sequencing in a cohort of 500 patients with rare diseases

**Why Not Relevant**: The provided paper excerpt does not mention the gene THBS4 or present any gene-specific variant data, variant lists, or per-gene diagnostic results. The text is a summary of overall exome sequencing (ES) diagnostic yield in a 500-patient rare disease cohort (rates of autosomal dominant/recessive/X-linked findings, number of variants, percent novel, subgroup yields, and secondary findings). Because there is no explicit reference to THBS4, no reported variants in THBS4, no segregation or case-level descriptions, and no functional/mechanistic data tied to THBS4, the excerpt provides no direct or mechanistic evidence for or against the claim that THBS4 is likely associated with at least one Mendelian disease. 

What would be needed to support the claim (not present in this excerpt):
- explicit reporting of pathogenic or likely pathogenic variants in THBS4 in one or more patients with relevant phenotypes, including zygosity and segregation data;
- evidence of recurrent THBS4 variants in unrelated affected individuals or de novo occurrences in dominant disorders;
- functional studies showing that THBS4 variants alter protein function in a way consistent with disease mechanism;
- population-frequency data showing pathogenic variants are rare or absent in controls; and
- clinical correlation and case-level phenotyping linking THBS4 disruption to a Mendelian phenotype.

Because none of these elements appear in the provided content, the excerpt is not informative regarding the claim.


[Read Paper](https://www.semanticscholar.org/paper/65c0b99fbca59c1e752feb6329a60b0eb2b9a59a)


### Expanding the phenotype of a recurrent de novo variant in PACS1 causing intellectual disability

**Why Not Relevant**: This paper does not mention the gene THBS4 anywhere in the provided content. The entire excerpt concerns a recurrent de novo missense variant in PACS1 (c.[607C>T], p.[Arg203Trp]) causing the intellectual disability syndrome MRD17, discussion of PACS1 function, and related clinical/experimental context. There are no data, statements, or mechanistic discussions about THBS4, THBS4 variants, THBS4-related phenotypes, or pathways implicating THBS4 in Mendelian disease. Therefore the paper provides no direct evidence for or against the claim that THBS4 is likely associated with a Mendelian disease. Note that absence of mention in this paper does not disprove any association in other literature; it merely means this paper is not a relevant source of evidence regarding THBS4.


[Read Paper](https://www.semanticscholar.org/paper/e460d89d850693b4fb1e3df12702be0dfb8d7f5b)


### Germline heterozygous exons 8–11 pathogenic BARD1 gene deletion reported for the first time in a family with suspicion of a hereditary colorectal cancer syndrome: more than an incidental finding?

**Why Not Relevant**: The provided paper content contains a single sentence discussing a BARD1 deletion detected in a family with familial colorectal cancer type X (FCCTX) and notes that the clinical criteria for Lynch syndrome are fulfilled without alteration of the DNA mismatch repair (MMR) system. There is no mention of the gene THBS4, no data linking THBS4 to any phenotype, no segregation or association analysis involving THBS4, and no mechanistic discussion that could plausibly connect BARD1 or FCCTX to THBS4. Because the text neither names THBS4 nor provides genetic, clinical, or functional evidence about it, it cannot support or refute the claim that "THBS4 is likely associated with at least one Mendelian disease." Additional limitations: the excerpt is a single sentence with no methods, results, or evidence relevant to THBS4; BARD1 is a distinct gene and findings about it do not imply anything about THBS4 without explicit data or mechanistic rationale; FCCTX describes a familial cancer syndrome but is not by itself proof of a Mendelian relationship for an unrelated gene. For these reasons the content is not relevant (relevance = 0).


[Read Paper](https://www.semanticscholar.org/paper/884a95b299f7238bf2a4a70eaeec5e815a99798c)


### Assessing Utility of Clinical Exome Sequencing in Diagnosis of Rare Idiopathic Neurodevelopmental Disorders in Indian Population

**Why Not Relevant**: The provided paper content does not mention the gene THBS4, nor does it list the specific genes (the 16 genes) or the 21 variants identified in the cohort. While the methods note that a CES panel of 4620 genes was assayed, the excerpt does not specify whether THBS4 was included on that panel or whether any variants in THBS4 were observed. Therefore there is no direct evidence (no reported THBS4 variants or gene-level association) and no mechanistic data concerning THBS4 in the excerpts provided. The small cohort size (19 patients) would also limit power to detect associations with a rare Mendelian disease even if THBS4 were included but unreported. Because of the absence of any mention of THBS4 and lack of gene-level results in the provided text, this paper excerpt is not relevant to assessing the claim that THBS4 is likely associated with at least one Mendelian disease.


[Read Paper](https://www.semanticscholar.org/paper/d6386b554b14a2943bd27e810e563c2304e699ad)


### TTC7A Variants Previously Described to Cause Enteropathy Are Observed on a Single Haplotype and Appear Non-pathogenic in Pediatric Inflammatory Bowel Disease Patients

**Why Not Relevant**: The provided paper content discusses compound heterozygote variants in the Tetratricopeptide Repeat Domain 7A gene (TTC7A) and the possible contribution of TTC7A variation to polygenic inflammatory bowel disease. The claim to be evaluated concerns the gene THBS4 (Thrombospondin-4) and its likelihood of being associated with at least one Mendelian disease. The two gene symbols (TTC7A and THBS4) refer to distinct, unrelated genes with different names, functions, and prior disease associations. The excerpt contains no mention of THBS4, no data on variants in THBS4, and no mechanistic discussion involving THBS4. Therefore this paper, as represented by the provided content, supplies neither direct evidence (e.g., reported pathogenic THBS4 variants segregating with Mendelian disease) nor mechanistic evidence (e.g., functional studies, pathway links, or model organism data) relevant to the claim about THBS4.

Limitations/Notes:
- The assessment is based solely on the single-sentence content you supplied. If the full paper contains additional sections not provided here, those were not available for review and could in principle mention THBS4; my conclusion applies only to the provided excerpt. 
- Because the provided excerpt explicitly names TTC7A and Lawless et al. and focuses on polygenic IBD, it is unlikely (from the excerpt) to contain evidence bearing on THBS4’s involvement in a Mendelian disorder. If you can provide the full text or any sections mentioning THBS4, I can re-evaluate for direct or mechanistic evidence.


[Read Paper](https://www.semanticscholar.org/paper/3818a966b5d6e8b0eda65fdef043ac0727fa429c)


### Exome sequencing reveals neurodevelopmental genes in simplex consanguineous Iranian families with syndromic autism

**Why Not Relevant**: The provided paper content does not mention THBS4 at all and contains no variant, clinical, genetic segregation, functional assay, or statistical evidence concerning THBS4. The single sentence discusses DMD and CHKB and a conceptual point about genes crossing clinical categories (ASD and muscular dystrophy). That general statement cannot be taken as evidence that THBS4 is associated with a Mendelian disease. To support the claim that THBS4 is likely associated with at least one Mendelian disease, one would need gene-specific evidence such as: (a) identification of rare, likely pathogenic variants in THBS4 in affected individuals with consistent phenotypes; (b) segregation of such variants with disease in families; (c) functional assays showing a deleterious effect consistent with disease mechanism; and (d) appropriate population frequency and statistical analyses. None of these are present in the supplied text. Therefore the paper content is not relevant to the claim.


[Read Paper](https://www.semanticscholar.org/paper/9cb23120953e32cc9faf4f47f59f270690fe7487)


### #2412 Monoallelic NEK8-related polycystic kidney disease in an Irish ADPKD cohort

**Why Not Relevant**: The provided paper content does not mention the gene THBS4 anywhere and instead focuses on autosomal dominant polycystic kidney disease and genes such as PKD1, PKD2, NEK8, IFT140, ALG5, DNAJB11, ALG8, and ALG9. All extracted sentences either set the ADPKD context, describe methods, or report findings specific to NEK8 and other listed genes. Therefore the paper provides no direct evidence supporting or refuting the claim that THBS4 is likely associated with at least one Mendelian disease. Additionally, absence of THBS4 among reported findings in this PKD-focused cohort is not strong evidence against a THBS4–Mendelian disease association generally, because the study population, panel composition, and disease focus (PKD-like phenotypes) limit detectability and relevance for other Mendelian conditions.


[Read Paper](https://www.semanticscholar.org/paper/9442a227dce2554b7c52a413419f138a5797381d)


## Search Queries Used

- THBS4 pathogenic variant case report familial segregation Mendelian disease thrombospondin 4

- THBS4 exome sequencing rare disease cohort de novo variant autosomal dominant autosomal recessive thrombospondin 4

- thrombospondin 4 knockout mouse phenotype loss of function model extracellular matrix integrin TGF beta fibrosis cardiac neuropathy skeletal muscle

- THBS4 ClinVar ClinGen gnomAD pLI loss of function intolerance gene disease validity review thrombospondin family THBS1 THBS2 THBS3 THBS4 THBS5


## Bibliometric Configuration

- Use Bibliometrics: True

- Author Impact Weight: 0.4

- Citation Impact Weight: 0.4

- Venue Impact Weight: 0.2
