## Supplementary material for "MENDELSEEK: An algorithm that predicts Mendelian Genes and elucidates what makes them special": TLX1.txt

# Claim: The gene TLX1 is likely associated with at least one Mendelian disease.

**Status**: processed

**Overall Rating**: -1

**Explanation**:

No relevant papers were found that support or refute this claim.


## Relevant Papers


## Other Reviewed Papers


### The Burden of Candidate Pathogenic Variants for Kidney and Genitourinary Disorders Emerging From Exome Sequencing

**Why Not Relevant**: The provided paper excerpt does not mention the gene TLX1 anywhere. The manuscript focuses on identification and curation of genes associated with Mendelian kidney and genitourinary disorders (a curated set of 625 genes), methods for variant filtering and interpretation, and the prevalence of candidate pathogenic variants in a cohort of ostensibly healthy adults. None of the quoted methods, results, or background text provide direct evidence that TLX1 is associated with any Mendelian disease, nor do they provide mechanistic data concerning TLX1 function or pathogenic variants. Without TLX1 being explicitly listed among the curated genes, reported as carrying candidate pathogenic variants in the cohort, or discussed mechanistically, this paper cannot be used to support or refute the claim that TLX1 is likely associated with at least one Mendelian disease. To evaluate that claim would require either: (a) the paper's full gene list or appendices showing TLX1 is included among the 625 curated genes, (b) reported variants in TLX1 from this cohort or from referenced databases (OMIM, ClinVar, HGMD) presented in the paper, or (c) mechanistic or clinical case evidence linking TLX1 to a Mendelian phenotype. If you want, I can (1) search the cited supplementary gene list (the laboratory website/appendix referenced) for TLX1, or (2) look up TLX1 in OMIM, ClinVar, HGMD, and the literature to summarize existing evidence for or against a Mendelian disease association.


[Read Paper](https://www.semanticscholar.org/paper/a75e297208c04992568e607d19aa9b9aaebd09d5)


### Discovery of Novel Recurrent Mutations in Childhood Early T-Cell Precursor Acute Lymphoblastic Leukemia by Whole Genome Sequencing - a Report From the St Jude Children's Research Hospital - Washington University Pediatric Cancer Genome Project

**Why Not Relevant**: The provided paper abstract does not mention the gene TLX1 at all. The study focuses on somatic genomic alterations in early T‑cell precursor acute lymphoblastic leukemia (ETP ALL) and reports recurrent somatic mutations, chromosomal rearrangements, and disrupted hematopoietic regulators (e.g., ETV6, RUNX1, GATA3, EZH2, SUZ12, etc.). The claim concerns whether TLX1 is likely associated with at least one Mendelian (germline) disease. There are three reasons the abstract is not relevant to that claim: (1) TLX1 is not listed among any genes, mutations, or structural variants described in the abstract, so there is no direct evidence for or against an association; (2) the paper addresses somatic alterations in cancer (leukemia) rather than germline variants causing Mendelian disease, so even if TLX1 were mentioned in the broader manuscript, the context would likely be somatic tumor biology rather than inherited Mendelian disorders; (3) the provided content is only the abstract, so absence of TLX1 in this text cannot definitively prove the gene is not discussed elsewhere, but for the content given there is no information to evaluate the claim. Limitations and caveats: (a) The assessment is based solely on the supplied abstract text (no page numbers or full text provided); it is possible TLX1 appears elsewhere in the full paper, though that is unlikely given its absence from the abstract’s comprehensive gene lists; (b) even if a gene is implicated in cancer by somatic mutation, that is not direct evidence of a Mendelian (germline) disease association — separate genetic, familial, and functional evidence would be required to support a Mendelian link. Given these points, the abstract provides no direct or mechanistic evidence that TLX1 is associated with any Mendelian disease.


[Read Paper](https://www.semanticscholar.org/paper/aad509fb9507d1399698ca948aa5fac8639e1fa4)


### Precision Medicine in Internal Medicine

**Why Not Relevant**: This paper is a general review/guidance on genetic testing methods, variant interpretation, and the state of gene–disease discovery as of 2019. It contains no gene-specific data, case reports, cohort analyses, segregation studies, functional assays, statistical associations, or database entries that mention TLX1. Therefore it provides no direct evidence that TLX1 is associated with any Mendelian disease. The few statements that are relevant are high-level (e.g., the fraction of genes known to cause Mendelian disorders, and that ES can reveal previously unknown gene–disease relationships) and only speak to general plausibility and the incompleteness of knowledge; they do not identify TLX1 as disease-associated. For a claim about TLX1 being likely associated with a Mendelian disease one would need gene-specific evidence: OMIM/ClinVar entries linking TLX1 variants to a consistent phenotype, multiple affected individuals with segregating variants, statistical enrichment of deleterious variants in cases vs controls, and/or functional data showing pathogenic mechanisms — none of which are present in this paper. Additionally, the OMIM statistic is dated to 1 February 2019, so even its indirect implications may be outdated relative to subsequent gene discoveries; however, the paper itself does not provide updated TLX1-specific information.


[Read Paper](https://www.semanticscholar.org/paper/c7199e8a5226f8d9d55777da0e15e91797bb459a)


### Nanoparticle delivery of inhibitory signal transducer and activator of transcription 3 G-quartet oligonucleotides blocks tumor growth in HMGA1 transgenic model of T-cell leukemia

**Why Not Relevant**: The paper documents TLX1 as an oncogene with activating (somatic) mutations and as a driver in T-cell acute lymphoblastic leukemia, and it provides mechanistic insight into TLX1's role in transcriptional networks. However, it does not present any evidence of germline TLX1 variants causing inherited disease, familial segregation data, or clinical descriptions of Mendelian phenotypes attributable to TLX1. Because the claim concerns Mendelian disease association (inherited, germline causation), the paper's focus on somatic oncogenic roles makes it largely irrelevant for supporting the claim that TLX1 is likely associated with at least one Mendelian disease.


[Read Paper](https://www.semanticscholar.org/paper/9d30f20c55001982942147529e8919d811f154bd)


### High frequency of cryptic chromosomal rearrangements involving the LMO2 gene in T-cell acute lymphoblastic leukemia

**Why Not Relevant**: The paper focuses on somatic LMO2 rearrangements and their role in T-cell acute lymphoblastic leukemia (T-ALL). Although TLX1 is mentioned as one of several genes for which mRNA levels were measured, the study: (1) reports only somatic tumor expression data (qRT-PCR) and somatic structural rearrangements; (2) contains no analysis of germline TLX1 variants, segregation, penetrance, or clinical phenotypes consistent with Mendelian inheritance; and (3) provides no epidemiologic, genetic linkage, or functional germline evidence linking TLX1 to any inherited Mendelian disease. Therefore the manuscript does not provide direct or mechanistic evidence that TLX1 is associated with a Mendelian disorder. Any inference about TLX1 and Mendelian disease would require germline genetic studies, phenotype descriptions, family segregation data, or functional assays of inherited variants, none of which are present in this paper.


[Read Paper](https://www.semanticscholar.org/paper/97c7cb665f050298aa5eb92d00cee3620db5763a)


### The Genomic Landscape of Childhood T-Lineage Acute Lymphoblastic Leukemia

**Why Not Relevant**: Relevance to the claim is very low because the paper reports somatic TLX1 rearrangements in T-cell acute lymphoblastic leukemia (i.e., tumor-derived, acquired genomic alterations). The claim concerns TLX1 being associated with a Mendelian disease, which requires evidence of germline variants causing an inherited phenotype (family segregation, de novo germline mutations with consistent phenotypes, population genetics, or case reports of inherited disease). This paper provides no germline analyses, no family history, and no discussion of inherited Mendelian phenotypes linked to TLX1 — only recurrent somatic rearrangements in cancer. Therefore, while the paper supports a role for TLX1 in somatic oncogenesis, it does not provide direct or mechanistic evidence that TLX1 is implicated in any Mendelian (germline-inherited) disease.


[Read Paper](https://www.semanticscholar.org/paper/41a49e0576c2754b5691acabeade1470eb678a2d)


### Clingen Coagulation Factor Deficiency Variant Curation Expert Panel: Meeting the Need for Recommendations to Curate Variants in the Coagulation Factor Genes

**Why Not Relevant**: The provided paper content focuses exclusively on coagulation factor genes (primarily F8 and F9), the ClinGen CFD-VCEP effort to specify ACMG/AMP guidelines for those genes, and the curation of coagulation-factor variants. There is no mention of the TLX1 gene, no experimental data, no genetic association analyses, and no mechanistic discussion involving TLX1. Therefore the paper provides no direct evidence supporting or refuting the claim that TLX1 is likely associated with at least one Mendelian disease. The absence of TLX1 in this paper reflects the narrow, gene-specific scope of the study rather than any assessment of TLX1; absence of evidence in this single, out-of-scope paper should not be interpreted as evidence that TLX1 is not associated with Mendelian disease. To evaluate the claim about TLX1 one would need to consult primary genetic studies, OMIM entries, ClinVar/ClinGen curations, or functional/mechanistic studies that explicitly analyze TLX1.


[Read Paper](https://www.semanticscholar.org/paper/9ea8d4ce19622ae1c6b0cdfd40848c93199f6280)


### A New Subtype of T-Cell Acute Leukemia in Very Young Children Is Defined by a Translocation Targeting the C-MYB Oncogene, and a Specific Gene Expression Signature.

**Why Not Relevant**: The paper studies somatic genomic alterations (translocations, duplications) involving the C-MYB locus in T-cell acute lymphoblastic leukemia and classifies cases into T-ALL subtypes, one of which is termed TLX1/HOX11. All evidence concerns somatic oncogenic events in leukemia (demonstrated somatic origin, expression changes, co-occurring somatic mutations) and subtype classification. There are no data on germline variants, familial segregation, congenital phenotypes, or inherited disease consistent with Mendelian inheritance. Therefore the paper provides no direct or indirect evidence that TLX1 is causally associated with any Mendelian disease; at best it shows TLX1 is implicated in somatic T-ALL subtype biology, which is insufficient to support the claim.


[Read Paper](https://www.semanticscholar.org/paper/7ee96a7fd3f82c7e4139559ef60cc957749f2891)


## Search Queries Used

- TLX1 germline variant pathogenic variant mutation inherited monogenic Mendelian disease T cell leukemia homeobox 1 HOX11 case report family segregation exome sequencing

- TLX1 translocation T cell acute lymphoblastic leukemia somatic oncogene HOX11 translocation insertion rearrangement t 10 14 somatic mutation cancer

- TLX1 knockout mouse Tlx1 null spleen agenesis splenic development homeobox transcription factor Hox11 embryonic development phenotype

- TLX1 OMIM ClinVar ClinGen HGMD gene disease association gene disease validity review monogenic disease database curation


## Bibliometric Configuration

- Use Bibliometrics: True

- Author Impact Weight: 0.4

- Citation Impact Weight: 0.4

- Venue Impact Weight: 0.2
