## Supplementary material for "MENDELSEEK: An algorithm that predicts Mendelian Genes and elucidates what makes them special": TMEM62.txt

# Claim: The gene TMEM62 is likely associated with at least one Mendelian disease.

**Status**: processed

**Overall Rating**: -1

**Explanation**:

No relevant papers were found that support or refute this claim.


## Relevant Papers


## Other Reviewed Papers


### Genome-wide meta-analysis identifies 93 risk loci and enables risk prediction equivalent to monogenic forms of venous thromboembolism

**Why Not Relevant**: The provided paper content does not mention TMEM62, nor does it present any gene-level association, variant, phenotype, or mechanistic data related to TMEM62 or to any specific Mendelian disease gene. The only sentence supplied is: "A polygenic score derived from these results identifies individuals at increased VTE risk equivalent to monogenic forms of the disease, and shows that PRS improved individual risk prediction beyond that of genetic and clinical risk factors." This sentence discusses a polygenic risk score (PRS) for venous thromboembolism (VTE) and compares its risk magnitude to that of unspecified "monogenic forms of the disease," but it does not (a) name TMEM62, (b) report an association between TMEM62 and any Mendelian phenotype, or (c) provide mechanistic evidence linking TMEM62 to disease biology. Therefore the content cannot be used to support or refute the claim that "The gene TMEM62 is likely associated with at least one Mendelian disease."

Additional notes on insufficiency: even though the sentence references monogenic forms of VTE, that phrase is generic and most commonly refers to well-established genes (for VTE, e.g., F5 [Factor V Leiden], F2 [prothrombin], PROC, PROS1, SERPINC1, etc.). Without explicit mention of TMEM62 in association analyses, segregation studies, pathogenic variants, functional experiments, or clinical case reports, no direct or mechanistic evidence about TMEM62 can be inferred from this text. To evaluate the claim about TMEM62 one would need gene-specific evidence such as: statistically significant rare-variant or single-variant associations with a Mendelian phenotype, segregation of putatively pathogenic variants in families, functional studies demonstrating a causative effect, or curated clinical variant annotations (e.g., ClinVar, OMIM). None of these are present in the supplied content, so relevance is effectively zero.


[Read Paper](https://www.semanticscholar.org/paper/b63879346cee607462085c91eac9db0b0588c46e)


### Effective variant filtering and expected candidate variant yield in studies of rare human disease

**Why Not Relevant**: The provided paper content does not mention TMEM62 or any gene-level association data. The only sentence given—"Effective guidelines for variant (SNP and INDEL) filtering are explored and the expected number of candidates for de novo dominant, recessive, and autosomal dominant modes of inheritance is reported."—is methodological and summarizes variant-filtering guidance and expected candidate counts by inheritance mode. This provides no direct evidence (no observations, statistical associations, case reports, or database entries) that TMEM62 is associated with any Mendelian disease, nor does it provide mechanistic evidence linking TMEM62 to disease biology. Limitations and caveats: (1) only a single summary sentence from the paper was supplied; it is possible the full paper contains gene lists, supplementary tables, or example disease-gene associations that mention TMEM62, but those are not present in the provided content and therefore cannot be used to support or refute the claim; (2) methodological papers that discuss filtering strategies are indirectly relevant because they describe approaches that could be used to discover gene–disease associations, but they do not constitute evidence that a specific gene (TMEM62) is associated with a Mendelian disorder. Conclusion: based on the supplied content alone, the paper provides no direct or mechanistic evidence for or against the claim that TMEM62 is likely associated with at least one Mendelian disease. To evaluate the claim further, the full paper and additional resources (OMIM, ClinVar, HGMD, peer-reviewed case reports, and functional studies) should be searched for explicit mentions of TMEM62 or data implicating it in Mendelian phenotypes.


[Read Paper](https://www.semanticscholar.org/paper/8da5eda5547e3521df7aecc8c8e4b7380a7641fa)


### Whole exome sequencing reveals inherited and de novo variants in autism spectrum disorder: a trio study from Saudi families

**Why Not Relevant**: The provided paper content does not mention TMEM62 or any specific gene names; it only gives a summary statement that variants were found in 15 ASD-candidate genes (five of which were previously unreported in human disease) and that implicated genes are enriched for neuronal processes. Because the claim concerns a specific gene (TMEM62) and whether it is likely associated with at least one Mendelian disease, relevant evidence would need to (a) name TMEM62 as one of the implicated genes, (b) report evidence of pathogenicity consistent with a Mendelian disease (segregation in families, de novo occurrence with appropriate phenotypic correlation, loss-of-function intolerance and supporting functional data, case-control enrichment with statistical significance), or (c) provide mechanistic functional data specifically implicating TMEM62 in a Mendelian phenotype. None of these items are present in the excerpt. The generic statement that some novel ASD-candidate genes were found and that implicated genes are enriched for neuronal functions is insufficient to infer that TMEM62 is likely associated with a Mendelian disease. Additional information (full gene lists, variant descriptions, inheritance patterns, segregation/phenotype correlation, functional assays, and statistical analyses) would be required to evaluate the claim for TMEM62.


[Read Paper](https://www.semanticscholar.org/paper/30bd7a60fbdb82dd358000834c3adc8c4a5cc869)


### Saturation genome editing-based functional evaluation and clinical classification of BRCA2 single nucleotide variants

**Why Not Relevant**: The provided paper content exclusively describes comprehensive saturation genome editing and functional characterization of BRCA2 single nucleotide variants in the DNA binding domain (exons 15–26), including validation, case-control association with breast and ovarian cancer, loss of heterozygosity in tumors, and clinical classification using a ClinGen/ACMG/AMP-like model. Nowhere in the supplied text is the gene TMEM62 mentioned, discussed, or analyzed. Consequently, there is no direct evidence (statistical association, clinical cases, or variant-level data) relating TMEM62 to any Mendelian disease, nor is there any mechanistic evidence (protein function, pathways, animal models, segregation data) that could plausibly link TMEM62 to a Mendelian phenotype. 

Although the methods used in this paper (saturation genome editing, functional assays, integration with clinical classification frameworks) are generally relevant as approaches that could be applied to other genes such as TMEM62, methodological relevance does not constitute evidence that TMEM62 itself is associated with any Mendelian disease. The results and conclusions are gene-specific to BRCA2 and cannot be extrapolated to TMEM62 without new, gene-specific data. Therefore the paper provides no support for the claim that TMEM62 is likely associated with at least one Mendelian disease. 

Limitations relevant to this assessment: (1) assessment is limited to the supplied paper content; if the full paper contained sections not provided here that mention TMEM62, that material was not available for review; (2) the paper’s general experimental approaches could inform future studies on TMEM62 but do not provide evidence about TMEM62’s disease associations.


[Read Paper](https://www.semanticscholar.org/paper/fa048a5ee952cd621f63179fdfa8381751e1ba2a)


### Non-Invasive Detection of a De Novo Frameshift Variant of STAG2 in a Female Fetus: Escape Genes Influence the Manifestation of X-Linked Diseases in Females

**Why Not Relevant**: The paper's content as provided focuses exclusively on STAG2: clinical case description and molecular findings identifying a de novo frameshift in the X-linked STAG2 gene, together with discussion of STAG2 loss-of-function vs missense variant effects, X-inactivation escape, and associated syndromes (HPE, MKMS). There is no mention of TMEM62, no variants reported in TMEM62, no mechanistic link or pathway analysis connecting STAG2 to TMEM62, and no population, segregation, functional, or case evidence related to TMEM62. Therefore this paper provides no direct evidence that TMEM62 is associated with any Mendelian disease. Absence of evidence in this single paper does not disprove the claim generally, but it means this paper cannot be used to support it. To evaluate the claim one would need studies reporting TMEM62 variants segregating with disease, de novo TMEM62 variants in affected individuals, functional studies showing deleterious effects, or human genetic database evidence linking TMEM62 to a Mendelian phenotype.


[Read Paper](https://www.semanticscholar.org/paper/de6a1dc65b6e99274f77c1a8b88d4171e94e468e)


### Multisite Validation of a Functional Assay to Adjudicate SCN5A Brugada Syndrome–Associated Variants

**Why Not Relevant**: The provided paper content exclusively concerns SCN5A and Brugada syndrome and makes no mention of the gene TMEM62. Relevant lines include: "Brugada syndrome is an inheritable arrhythmia condition that is associated with rare, loss-of-function variants in SCN5A," descriptions of an "in vitro SCN5A-Brugada syndrome automated patch clamp assay," and results about SCN5A variant classification (e.g., "Application of the assay to 4 clinical SCN5A variants of uncertain significance revealed loss-of-function for 3/4 variants, enabling reclassification to likely pathogenic"). Because TMEM62 is not discussed, the paper provides no direct genetic, clinical, or functional evidence that TMEM62 is associated with any Mendelian disease. Furthermore, there is no mechanistic discussion connecting SCN5A biology, electrophysiology assays, or Brugada syndrome pathology to TMEM62 or to any pathway in which TMEM62 is implicated. While the paper demonstrates a generalizable approach (high-throughput functional assays) that could, in principle, be applied to other genes in future studies, that methodological point does not constitute evidence that TMEM62 is associated with Mendelian disease. Therefore, this paper is not relevant for supporting or refuting the claim that "TMEM62 is likely associated with at least one Mendelian disease."


[Read Paper](https://www.semanticscholar.org/paper/fa3d4295e543f9b5fcaed7f61336420d621f5a32)


### Genetic Analysis in a Familial Case With High Bone Mineral Density Suggests Additive Effects at Two Loci

**Why Not Relevant**: The provided paper content contains no mention of the gene TMEM62 and therefore provides no direct or mechanistic evidence that TMEM62 is associated with any Mendelian disease. The excerpt focuses on a whole-exome sequencing study of a family segregating high bone mineral density and describes identified variants and candidate genes VAV3 and ADGRE5 (including mouse knockout phenotypes) as putative high-BMD genes. Specifically, the text reports identification of 38 rare coding variants and discusses VAV3 and ADGRE5 as fitting the phenotype, but does not report any findings implicating TMEM62. Consequently, there is no direct evidence (e.g., a pathogenic variant in TMEM62 segregating with disease) nor mechanistic evidence (e.g., functional data, animal models, pathway links) in the provided content to support the claim about TMEM62.

Limitations and caveats: (1) The absence of TMEM62 in the provided excerpt does not definitively prove the full paper contains no mention of TMEM62 — only that the supplied content does not. (2) The study is family-based and focused on high-BMD phenotypes; even if TMEM62 were mentioned elsewhere in the full paper, that would not necessarily establish TMEM62 as causative of a Mendelian disease without segregation, functional, or replication evidence. (3) The family sample is small (three affected, three unaffected sequenced), which limits generalizability and power to implicate genes beyond those discussed. 

Recommended next steps to evaluate the claim: search the full text of this paper for TMEM62; search locus- and disease-specific databases (OMIM, ClinVar, HGMD), large sequencing resources (gnomAD, ClinGen), and published literature for reported TMEM62 pathogenic variants, segregation in families, and functional/animal-model evidence supporting a Mendelian disease association.


[Read Paper](https://www.semanticscholar.org/paper/de079d062b38ceaec92ab3987723acef25f28dff)


### Clingen Coagulation Factor Deficiency Variant Curation Expert Panel: Meeting the Need for Recommendations to Curate Variants in the Coagulation Factor Genes

**Why Not Relevant**: The provided paper content focuses on development and specification of variant curation guidelines for coagulation factor genes, specifically the ClinGen Coagulation Factor Deficiency Variant Curation Expert Panel (CFD-VCEP) and its initial focus on F8 and F9. There is no mention of the gene TMEM62, no case reports, no variant data, no functional studies, no mechanistic discussion, and no literature review related to TMEM62. Therefore the paper supplies no direct evidence (e.g., observed pathogenic variants in TMEM62, segregation with disease, case reports, population frequency data) nor mechanistic evidence (e.g., molecular function, pathway involvement, animal models) relevant to the claim that TMEM62 is likely associated with at least one Mendelian disease. While the curation methods and framework described could, in principle, be applied to TMEM62 in a different study, that methodological relevance does not constitute evidence for a TMEM62–disease association. Given the complete absence of TMEM62-specific data or discussion, the content is not relevant to supporting or refuting the claim.


[Read Paper](https://www.semanticscholar.org/paper/9ea8d4ce19622ae1c6b0cdfd40848c93199f6280)


### Curation of OCA2 Variants of Uncertain Significance From Chinese Oculocutaneous Albinism Patients Based on Multiplex Assays

**Why Not Relevant**: Relevance is near zero because the provided paper content exclusively concerns OCA2 and variants associated with oculocutaneous albinism type 2. The text repeatedly specifies that the study's aims, experimental assays (MAVEs), variant selection, functional results, and clinical diagnoses all pertain to OCA2. The paper content contains no mention of the gene TMEM62, no experimental data, no mechanistic discussion, and no clinical associations involving TMEM62. Therefore this paper provides neither direct evidence (genetic association, variant segregation, case reports) nor mechanistic evidence (protein function, pathways, interactions) supporting that TMEM62 is associated with any Mendelian disease. Limitations: absence of evidence in this paper is not proof that TMEM62 is not disease-associated — it only means this particular study does not address TMEM62. To evaluate the claim about TMEM62 one must search other primary literature, clinical databases (OMIM, ClinVar), or large-scale genetic association/segregation studies that include TMEM62.


[Read Paper](https://www.semanticscholar.org/paper/c3ce96590826502429e313724256a8c3ad09976e)


### Re-evaluation of whole exome sequencing, including intronic region, in combination with genetic intolerance score for foetal structural anomalies, is helpful for diagnosis, especially in X-linked disorders

**Why Not Relevant**: This paper does not provide evidence for or against the claim that TMEM62 is likely associated with at least one Mendelian disease. The paper mentions other genes (for example TMEM67 and CASK) but contains no mention of TMEM62, no variants reported in TMEM62, and no mechanistic discussion involving TMEM62. Representative verbatim sentences from the paper that show which genes are discussed (and thereby demonstrate the absence of TMEM62) include:

"Using the second step, we diagnosed Meckel-Gruber syndrome by detecting likely pathogenic intron variant in TMEM67." 

"In the third step, we identified a de novo hemizygous pathogenic variant in one severe hydrops fetalis male, which could cause aberrant splicing in CASK." 

"Moreover, we revealed that the LOEUF score of X-linked disease-causing genes was significantly lower than that of autosomal genes among all OMIM-registered genes." 

"Conclusion: We showed that the evaluation of variants, including introns of WES data, in combination with the LOEUF score, could improve the WES diagnostic yield and detect new candidate genes, especially on chromosome X."

How this bears on the claim:
- Direct evidence: None. The paper does not report TMEM62 variants, TMEM62 phenotypes, segregation data, or any clinical cases implicating TMEM62. Therefore there is no direct diagnostic or association evidence for TMEM62 in this study.
- Mechanistic evidence: None specific to TMEM62. The paper describes an approach (reassessing WES including intronic variants and using LOEUF/spliceAI) that could in principle detect new disease genes. However, this methodological discussion does not provide mechanistic data about TMEM62 protein function, pathogenic mechanisms, or experimental results that would implicate TMEM62.

Limitations relevant to the claim:
- Absence of evidence in this paper is not evidence of absence: this single study of 138 fetuses simply did not report TMEM62 findings; it does not rule out TMEM62 as a disease gene in other cohorts or contexts.
- The paper focuses on genes identified within their dataset (not a genome-wide statement about all genes), so lack of mention likely reflects no TMEM62 findings in these samples rather than a deliberate evaluation of TMEM62.

Recommendation: To evaluate the claim about TMEM62, search other sources (OMIM, ClinVar, gnomAD constraint metrics, case reports, functional studies) specifically for TMEM62 (not TMEM67). This paper cannot be used as supporting evidence for TMEM62 association with Mendelian disease.


[Read Paper](https://www.semanticscholar.org/paper/b9004e518cf738650e488c6c4990525732fce092)


### Complexity of familial Mediterranean fever genetics.

**Why Not Relevant**: Relevance is effectively zero because the provided paper content discusses familial Mediterranean fever (FMF), the MEFV gene, inheritance patterns, and possible mechanisms (epigenetic dysregulation, somatic mosaicism, microchimerism) relevant to FMF genetics, but it contains no mention of the gene TMEM62, nor any data, analyses, or mechanistic discussion that implicates TMEM62 in any Mendelian disease. Therefore the content provides neither direct evidence (no association studies, case reports, segregation data, or variant pathogenicity data for TMEM62) nor mechanistic evidence (no functional data, pathway links, protein interactions, or model systems) to support the claim that TMEM62 is likely associated with at least one Mendelian disease. The excerpts above are provided to show the paper's focus and why it does not bear on TMEM62; none change that conclusion.


[Read Paper](https://www.semanticscholar.org/paper/01f0e9e5e4493cb1f5c3fc049a4b6b9b1fb29a0b)


### OP0309 WES ANALYSIS OF RARE FAMILIES POINTS TO A PATHOGENIC ROLE FOR THE cGAS/STING-TYPE I IFN AXIS IN SSc

**Why Not Relevant**: The paper does not mention TMEM62 anywhere in the provided text. It describes whole-exome sequencing of familial systemic sclerosis cases, the filtering strategy used, and highlights other candidate genes (for example TMEM173/STING, NT5E/CD73, PRKD2) including that two candidates cause known Mendelian diseases, but TMEM62 is not named as a candidate or discussed. While the methods (WES and the applied filters) could have detected coding variants in TMEM62 if such variants were present in these families, the absence of any mention of TMEM62 in the study means there is no direct or mechanistic evidence within this paper to support or refute the claim that TMEM62 is associated with a Mendelian disease. Additionally, even if TMEM62 had been absent from the candidate lists here, that would not be strong evidence against TMEM62 having Mendelian disease associations in other cohorts or contexts (limitations include small sample size, disease-specific focus on systemic sclerosis families, WES capture limits, and variant-filtering choices). For these reasons the paper is non-relevant to the specific claim about TMEM62.


[Read Paper](https://www.semanticscholar.org/paper/a5d1937aaf7ad488b3d5d4df3648ee2ca8966c0d)


### Major cardiovascular consequences of familial hypercholesterolemia in Poland from the economic perspective

**Why Not Relevant**: The paper is a focused review/economic analysis of familial hypercholesterolemia (FH) and discusses known FH-causing genes (LDLR, ApoB, PCSK9, LDLRAP1 and a few others). It contains no mention of TMEM62, no data linking TMEM62 to any Mendelian disease, and no experimental, genetic, or mechanistic evidence involving TMEM62. Therefore the paper provides no direct evidence supporting the claim that TMEM62 is likely associated with at least one Mendelian disease. The only information relevant to the claim is negative/indirect (i.e., lists of known FH genes that do not include TMEM62), which is insufficient to establish an association. Absence of TMEM62 in this FH-focused paper does not constitute proof that TMEM62 has no Mendelian disease associations elsewhere; it merely indicates this paper does not support the claim.


[Read Paper](https://www.semanticscholar.org/paper/30ae7d03a107f66f1a6a2c2b54229265a3b502d0)


### [UTILIZATION OF WHOLE EXOME SEQUENCING IN DIAGNOSTICS OF GENETIC DISEASE: RABIN MEDICAL CENTER'S EXPERIENCE].

**Why Not Relevant**: The provided paper content does not mention TMEM62 at all and contains no per-gene results, lists, or identifiers. Although the study reports that it identified novel mutations in genes not previously associated with genetic disease (9/39 patients), it does not name those genes or present segregation, recurrence, or functional evidence that would allow one to evaluate whether TMEM62 is among them or whether any nominated gene meets criteria for a Mendelian disease association. Therefore the content is only weakly relevant: it demonstrates that WES can discover novel candidate disease genes in principle, but provides no direct or mechanistic evidence specific to TMEM62. To support the claim about TMEM62 one would need an explicit statement that TMEM62 harbors pathogenic variants segregating with disease (or multiple unrelated patients with similar phenotypes and TMEM62 variants), plus supporting functional or statistical evidence; none of that is present in the excerpt.


[Read Paper](https://www.semanticscholar.org/paper/7df5093c3d8a10e67b7fe0cc49f03c39e45d2904)


### MOL CLASS OF 2022 SENIOR THESIS ABSTRACTS Identification and characterization of positive regulators governing hepatitis C virus host tropism

**Why Not Relevant**: Based on the provided paper content, there is no mention of the gene TMEM62 (neither the symbol 'TMEM62' nor the name 'transmembrane protein 62' appears). The excerpts discuss hepatitis C host factors, TRIM26 and cyclophilin A, METTL3/METTL14 epitranscriptomic work, a CRISPR dCas13-FTO construct, Vps33/Vps18, lung morphogenesis and WNT signaling, Kurly protein, Prdm1 in Ciona, bacterial defense gene surveys, RTK biosensors, and cerebellar circuit tracing. None of these sections describe human genetic association, family-based (Mendelian) linkage/segregation analyses, pathogenic variants, clinical phenotypes, or cite databases of Mendelian disease (e.g., OMIM/ClinVar/DECIPHER) that would provide direct or mechanistic evidence that TMEM62 is associated with a Mendelian disease.

Reasons for rating and limitations:
- No direct evidence: The text contains no sentences or paragraphs that assert or present data linking TMEM62 to any human disease, Mendelian or otherwise.
- No mechanistic evidence: Although some mechanistic molecular biology is discussed (e.g., methyltransferases METTL3/METTL14, CRISPR tools, host factors for HCV), none of that mechanistic data references TMEM62 or provides a plausible pathway implicating TMEM62 in Mendelian disease.
- Scope and methods mismatch: The described studies are experimental and mechanistic across virology, developmental biology, structural biology, and bacterial genomics, not clinical human genetics or variant/segregation analyses that would identify Mendelian disease genes.
- Possible coverage limitation: My conclusion is based solely on the supplied content. If the full paper (or supplementary materials) includes sections not provided here that mention TMEM62 or human genetic data, those were not available for review. To conclusively rule out relevance, search the full text and supplementary files for 'TMEM62', 'transmembrane protein 62', 'mutation', 'variant', 'familial', 'segregation', 'OMIM', 'ClinVar', or related terms.

Suggested next steps if you want to establish whether TMEM62 is linked to a Mendelian disease:
- Search the full paper PDF and any supplements for 'TMEM62' or synonyms.
- Query human genetic resources: OMIM, ClinVar, HGMD (if available), DECIPHER, and gnomAD constraint metrics for TMEM62.
- Search PubMed for 'TMEM62' + 'mutation' or 'disease' and check recent gene-disease curation efforts (ClinGen).

Because no relevant excerpts were found in the provided content, no verbatim quotes or per-excerpt explanations are included.


[Read Paper](https://www.semanticscholar.org/paper/9571d247a1af65ff123ad6a20084e2ce86f82e58)


### POGZ controls embryonic stem cell self-renewal and pluripotency by association with esBAF and HP1

**Why Not Relevant**: This paper describes experiments and findings about the gene POGZ (its domains, role in embryonic stem cells, association with esBAF/BRG1 and HP1 proteins, and links between loss-of-function POGZ mutations and neurodevelopmental disorders). The text contains no mention of TMEM62, no genetic association data for TMEM62, and no mechanistic data that implicates TMEM62 in disease. Therefore the paper provides neither direct evidence (e.g., variant/disease co-segregation, case reports, statistical association) nor mechanistic evidence (e.g., functional assays, pathway links specific to TMEM62) relevant to the claim that TMEM62 is likely associated with at least one Mendelian disease. Because the subject is a different gene (POGZ), findings about POGZ cannot be used to support or refute an association of TMEM62 with Mendelian disease without additional, independent data. If you would like, I can (a) search the literature for TMEM62-specific genetic or functional evidence, or (b) evaluate another paper that mentions TMEM62 for relevance to the claim.


[Read Paper](https://www.semanticscholar.org/paper/40197ea88cf80a3a8e23984fb47ee9fa046fc6ef)


### Expression of Huntingtons disease markers in eye tissues

**Why Not Relevant**: The provided paper content does not mention TMEM62 anywhere, nor does it present any genetic, clinical, or experimental data linking TMEM62 to a Mendelian disease. The text centers on Huntington's disease (HTT), the MommeD3 mutant (for which a causative mutation is said to have been identified but not named in the excerpt), and other genes/systems (AAK1/SEL-5, p130Cas/BCAR1, ATM). Because there is no explicit statement that TMEM62 is mutated in any of the described phenotypes, no allelic, segregation, variant, functional, or clinical evidence for TMEM62 is provided, and no mechanistic pathway involving TMEM62 is described, the paper cannot be used to support the claim that TMEM62 is likely associated with at least one Mendelian disease. Limitations and caveats: it is possible (but not stated) that the causative gene identified for MommeD3 could be TMEM62; however, without the gene name, variant details, phenotypic segregation, functional validation, or any mention in the text, no inference can be made. Similarly, the presence of next-generation sequencing methods or general statements about identifying causative mutations does not constitute evidence for any specific gene. Therefore the relevance of this paper to the claim is effectively zero based on the provided content.


[Read Paper](https://www.semanticscholar.org/paper/20ae6edb20d81fe8df2617d06f8ba75e11d0ee30)


### Title : Loss of cytoskeletal transport during egress critically attenuates 1 Ectromelia virus infection in vivo 2 3 Running title : Cytoskeletal transport is critical for ECTV infection 4 5

**Why Not Relevant**: This paper contains no mention of the gene TMEM62, no human genetic data, and no discussion of Mendelian diseases. The manuscript focuses entirely on orthopoxvirus biology (ECTV and VACV), the viral A36/A36R protein, viral egress, microtubule- and actin-based transport, and mouse infection models. Example topics covered include A36R function, interactions with kinesin-1, actin comet formation, and attenuation of an A36R-deleted virus in BALB/c and Rag-1-deficient mice. None of these sections provide direct evidence (genetic linkage, mutation-disease co-segregation, pathogenic variants, case reports, or phenotypic descriptions in families or patients) nor mechanistic evidence (molecular function in human cells relevant to disease, pathogenic variant effects, animal models linking TMEM62 perturbation to disease phenotypes) that would support or refute the claim that TMEM62 is associated with a Mendelian disease. Therefore the paper is not relevant to the claim.

If you would like, I can (a) search the literature and human genetic databases (OMIM, ClinVar, HGMD, PubMed, gnomAD) for evidence linking TMEM62 to Mendelian disease, (b) evaluate any specific TMEM62 variant or study you provide, or (c) explain what types of evidence would be needed to support the claim that TMEM62 is associated with a Mendelian disorder.


[Read Paper](https://www.semanticscholar.org/paper/ea511113489f01e40495e01038abaa31018e016e)


## Search Queries Used

- TMEM62 mutation Mendelian disease case report monogenic familial transmembrane protein 62

- TMEM62 ClinVar OMIM ClinGen pathogenic variant loss of function pathogenicity classification

- whole exome sequencing rare disease TMEM62 de novo variant trio sequencing familial autosomal recessive autosomal dominant X linked rare variant burden

- TMEM62 knockout mouse phenotype functional characterization transmembrane protein 62 expression localization protein interaction pathway mechanistic study


## Bibliometric Configuration

- Use Bibliometrics: True

- Author Impact Weight: 0.4

- Citation Impact Weight: 0.4

- Venue Impact Weight: 0.2
