## Supplementary material for "MENDELSEEK: An algorithm that predicts Mendelian Genes and elucidates what makes them special": TMPO.txt

# Claim: The gene TMPO is likely associated with at least one Mendelian disease.

**Status**: processed

**Overall Rating**: 4

**Explanation**:

Supporting evidence.
The strongest direct evidence is the 2005 study reporting a putative TMPO (LAP2) mutation in a familial dilated cardiomyopathy (DCM) pedigree. The authors screened 113 subjects from 88 families and identified a single nonsynonymous change (c.2068C>T; p.R690C) affecting the LAP2α isoform in one familial DCM pedigree. Functional work reported in the paper showed that the R690C substitution significantly impaired the in vitro interaction between mutated LAP2α and the pre–lamin A C-terminus, although localization of the mutant protein in HeLa cells appeared similar to wild type. The authors conclude that LAP2 (TMPO) mutations may represent a rare cause of DCM, implicating TMPO as a candidate disease gene associated with a Mendelian form of cardiomyopathy.

Caveats or contradictory evidence.
The evidence is limited to a single reported family and a single variant. The excerpts do not present detailed segregation data, allele frequency in controls/population databases, or independent replication in additional families or cohorts. The functional data are limited to an in vitro interaction assay and cell localization in HeLa cells; while the interaction deficit is biologically plausible given LAP2α’s binding to A-type lamins (LMNA), the functional work does not by itself prove pathogenicity in vivo. The first cited paper discusses nuclear envelope proteins (emerin, lamin A/C) causing tissue-specific degenerative diseases, which makes TMPO a biologically plausible candidate, but does not itself provide direct evidence that TMPO mutations cause human Mendelian disease. Thus the evidence is suggestive but not definitive, and alternative interpretations (a rare benign/private variant, incomplete segregation, or a modifier role) cannot be excluded based on the provided excerpts.

Analysis of potential mechanisms and implications.
Biologically, TMPO (LAP2) encodes lamina-associated polypeptide 2 isoforms that interact with A-type lamins and localize to the nucleus; disruption of nuclear lamina interactions is a known mechanism in several laminopathies including familial DCM and Emery–Dreifuss muscular dystrophy. The reported R690C change lies in a C-terminal domain known to interact with lamin A/C, and the paper documents a specific impairment of that interaction, providing a plausible mechanistic link to laminopathy-like cardiac phenotypes. If pathogenic, TMPO mutations would fit mechanistically with established nuclear envelope disease pathways. However, absent robust genetic evidence (segregation, recurrence, absence in healthy controls) and more extensive functional validation (e.g., animal models, cardiomyocyte assays), the role of TMPO could be rare or context-dependent (e.g., a low-penetrance allele or genetic modifier), rather than a common Mendelian disease gene.

Which rating is most appropriate.
Given a direct report of a TMPO coding variant in a familial DCM pedigree with accompanying functional evidence and a biologically plausible mechanism linking TMPO to laminopathy-like disease, there is reasonable support for the claim that TMPO is associated with at least one Mendelian disease. However, the evidence is limited in scope (single family, limited functional assays, no reported replication), so the association is not yet definitive or strongly established. Balancing the presence of concrete, relevant positive evidence against significant limitations, the best-fitting rating is "Likely True": TMPO is likely associated with at least one Mendelian disease (rare familial DCM), though further genetic and functional corroboration is needed to make the association robust and widely accepted.


**Final Reasoning**:

I reviewed the logic and evidence: a targeted genetic screen identified a TMPO (LAP2α) missense change in one familial DCM pedigree and demonstrated impaired interaction with lamin A in vitro. This provides a direct, biologically plausible link between TMPO variation and a Mendelian cardiac phenotype. However, the evidence is limited to a single variant/family, with no reported replication or comprehensive segregation/ population-frequency data in the provided excerpts, and functional data are limited to in vitro assays. There is no contradictory paper supplied, but the limitations meaningfully weaken certainty. Therefore the conclusion that TMPO is associated with at least one Mendelian disease is supported but not proven beyond reasonable doubt; "Likely True" most accurately reflects that balance.


## Relevant Papers


### Molecular signatures of Emery-Dreifuss muscular dystrophy.

**Authors**: Matthew A. Wheeler (H-index: 6), Juliet A. Ellis (H-index: 28)

**Relevance**: 0.2

**Bibliometric Impact**: 0.27665


**Excerpts**:

- Mutations in genes encoding the nuclear envelope proteins emerin and lamin A/C lead to a range of tissue-specific degenerative diseases.

- These include dilated cardiomyopathy, limb-girdle muscular dystrophy and X-linked and autosomal dominant EDMD (Emery-Dreifuss muscular dystrophy).

- The molecular mechanisms underlying these disorders are poorly understood; however, recent work using animal models has identified a number of signalling pathways that are altered in response to the deletion of either emerin or lamin A/C or expression of Lmna mutants found in patients with laminopathies.

- A distinguishing feature of patients with EDMD is the association of a dilated cardiomyopathy with conduction defects.

- In the present article, we describe several of the pathways altered in response to an EDMD phenotype, which are known to be key mediators of hypertrophic growth, and focus on a possible role of an emerin-beta-catenin interaction in the pathogenesis of this disease.


**Explanations**:

- How it relates: This sentence provides direct evidence that mutations in genes encoding nuclear envelope proteins (specifically emerin and lamin A/C) cause Mendelian, tissue-specific degenerative diseases. Relevance to the claim: indirect support for plausibility that other nuclear envelope genes (such as TMPO) could likewise be disease-associated. Strength: establishes precedent that nuclear envelope genes can be Mendelian disease genes. Limitations: the sentence names only emerin and lamin A/C; TMPO is not mentioned, so there is no direct evidence linking TMPO to Mendelian disease in this paper.

- How it relates: This lists specific Mendelian clinical phenotypes caused by mutations in nuclear envelope genes. Relevance to the claim: provides contextual examples of diseases arising from nuclear envelope gene mutations, supporting mechanistic plausibility that other nuclear envelope components might be implicated. Strength: concrete examples of Mendelian disorders tied to nuclear envelope gene mutations. Limitations: none of the listed genes are TMPO; the excerpt does not present genetic, clinical, or variant-level evidence for TMPO.

- How it relates: Describes mechanistic investigations (animal models) showing altered signalling pathways after loss/mutation of nuclear envelope components. Relevance to the claim: mechanistic evidence that perturbation of nuclear envelope proteins can cause downstream molecular changes that drive disease, increasing plausibility that mutations in other nuclear envelope genes (e.g., TMPO) could have pathogenic effects. Strength: points to experimental (animal-model) mechanistic work rather than only clinical correlation. Limitations: the experimental work described concerns emerin and lamin A/C (and Lmna mutants); TMPO is not studied here, so mechanistic extrapolation to TMPO is indirect and speculative.

- How it relates: Provides clinical phenotype detail (cardiac involvement in EDMD) associated with nuclear envelope gene mutations. Relevance to the claim: gives phenotype context for nuclear envelope gene–related Mendelian disease, supporting the general concept that nuclear envelope gene mutations have clinically significant, inheritable consequences. Strength: clarifies the types of Mendelian phenotypes seen with such genes. Limitations: again, TMPO is not mentioned; this is contextual rather than gene-specific evidence.

- How it relates: Indicates the paper examines specific signalling pathways and a molecular interaction (emerin-beta-catenin) as pathogenic mechanisms in EDMD. Relevance to the claim: supports the idea that specific molecular interactions involving nuclear envelope proteins can underlie Mendelian disease mechanisms, thereby providing mechanistic plausibility that other nuclear envelope proteins (such as TMPO) might cause disease via analogous interactions. Strength: points to concrete molecular hypotheses and pathway-level mechanisms. Limitations: the molecular focus is on emerin and beta-catenin; there is no experimental or genetic data here about TMPO, so this does not constitute direct evidence linking TMPO to Mendelian disease.


[Read Paper](https://www.semanticscholar.org/paper/e87c6dbc419d466c744d6f6fefc7fdf942cabbb4)


### Thymopoietin (lamina‐associated polypeptide 2) gene mutation associated with dilated cardiomyopathy

**Authors**: M. Taylor (H-index: 5), L. Mestroni (H-index: 63)

**Relevance**: 0.85

**Bibliometric Impact**: 0.4160631578947368


**Excerpts**:

- Thymopoietin or TMPO (indicated by its alternative gene symbol, LAP2, in this work) has been proposed as a candidate disease gene for dilated cardiomyopathy (DCM), since a LAP2 product associates with nucleoplasmic lamins A/C, which are encoded by the DCM gene LMNA.

- We developed a study to screen for genetic mutations in LAP2 in a large collection of DCM patients and families.

- A total of 113 subjects from 88 families (56 with familial DCM (FDC) and 32 with sporadic DCM) were screened for LAP2 mutations using denaturing high‐performance liquid chromatography and sequence analysis.

- We found a single putative mutation affecting the LAP2α isoform in one FDC pedigree.

- The mutation predicts an Arg690Cys substitution (c.2068C>T; p.R690C) located in the C‐terminal domain of the LAP2α protein, a region that is known to interact with lamin A/C.

- RT‐PCR, Western blot analyses, and immunolocalization revealed low‐level LAP2α expression in adult cardiac muscle, and localization to a subset of nuclei.

- Mutated Arg690Cys LAP2α expressed in HeLa cells localized to the nucleoplasm like wild‐type LAP2α, with no effect on peripheral and nucleoplasmic lamin A distribution.

- However, the in vitro interaction of mutated LAP2α with the pre‐lamin A C‐terminus was significantly compromised compared to the wild‐type protein.

- LAP2 mutations may represent a rare cause of DCM.

- The Arg690Cys mutation altered the observed LAP2α interaction with A‐type lamins.

- Our finding implicates a novel nuclear lamina‐associated protein in the pathogenesis of genetic forms of dilated cardiomyopathy.

- Hum Mutat 26(6), 566–574, 2005.  2005 Wiley‐Liss, Inc.


**Explanations**:

- Relation to claim: This statement establishes TMPO/LAP2 as a candidate disease gene for a Mendelian cardiac disorder (DCM) by linking its product to lamin A/C, proteins already implicated in Mendelian DCM (LMNA). Type of evidence: mechanistic/contextual. Strengths/limitations: It provides biologic rationale (protein–protein interaction) supporting plausibility of TMPO involvement in Mendelian disease, but is not direct genetic evidence of causation.

- Relation to claim: Describes study purpose to detect mutations in LAP2 among DCM patients, indicating direct investigation of a potential Mendelian association. Type of evidence: methodological/direct. Strengths/limitations: Shows the authors set out to seek genetic evidence; not a result but important context for interpretation of subsequent findings.

- Relation to claim: Provides sample size and ascertainment (familial and sporadic DCM cases) used to search for LAP2 mutations—relevant to the strength of genetic screening. Type of evidence: methodological. Strengths/limitations: Reasonable cohort size for a candidate gene screen, but still limited power for rare variants; sample description allows assessment of how broadly the finding can generalize.

- Relation to claim: This is direct genetic evidence: identification of a single putative LAP2α mutation in one familial DCM pedigree. Type of evidence: direct. Strengths/limitations: Positive finding supports that TMPO can carry rare variants in Mendelian DCM, but being a single pedigree limits confidence—segregation details, penetrance, replication, and statistical enrichment versus controls are not provided here.

- Relation to claim: Provides exact mutation (Arg690Cys; c.2068C>T; p.R690C) and locates it to a domain that interacts with lamin A/C, which links the variant to a functional region. Type of evidence: mechanistic/direct. Strengths/limitations: Locating the variant to a known interaction domain strengthens plausibility that the variant could be pathogenic; however, presence in a functional domain alone does not prove causality without segregation, frequency in controls, or in vivo functional impact.

- Relation to claim: Shows that LAP2α is expressed at low levels in adult cardiac muscle and localizes to nuclei—important for biological plausibility that LAP2 variants could affect heart function. Type of evidence: mechanistic/contextual. Strengths/limitations: Confirms tissue-relevant expression, supporting plausibility; 'low‐level' expression and localization to only a subset of nuclei could limit expected impact and complicate interpretation of pathogenicity.

- Relation to claim: Tests cellular localization of the mutant protein and reports no detectable change in lamin A distribution in HeLa cells, which argues against a dominant mislocalization mechanism. Type of evidence: mechanistic/functional. Strengths/limitations: Negative effect on localization weakens some pathogenic models, but HeLa cell overexpression may not recapitulate cardiomyocyte context; lack of effect on lamin A distribution does not rule out other pathogenic mechanisms.

- Relation to claim: Functional assay result showing the mutant has significantly reduced binding to pre-lamin A C-terminus—direct mechanistic evidence that the variant impairs interaction with A-type lamins. Type of evidence: mechanistic/functional. Strengths/limitations: This is important functional evidence linking the variant to disrupted interaction with a protein implicated in Mendelian DCM (LMNA). Limitation: described as 'in vitro'—assay conditions, quantitation, and physiological relevance (especially in heart tissue/in vivo) are not detailed here.

- Relation to claim: Authors explicitly state that LAP2 mutations could be a rare cause of DCM—this is an interpretive direct statement supporting the claim that TMPO is associated with at least one Mendelian disease. Type of evidence: interpretive/conclusion. Strengths/limitations: Supports the claim but is cautious ('may represent a rare cause'); the conclusion is based on a single pedigree and limited functional data, so it is suggestive rather than definitive.

- Relation to claim: Reiterates that the specific Arg690Cys variant altered the interaction with A-type lamins—ties the identified variant to a mechanistic effect relevant to disease. Type of evidence: mechanistic. Strengths/limitations: Reinforces functional impairment; again limitations include in vitro assay reliance and lack of in vivo/segregation evidence in this excerpt.

- Relation to claim: Summative conclusion that implicates LAP2 (TMPO) in the pathogenesis of genetic forms of DCM—this directly supports the assertion that TMPO is associated with a Mendelian disease (here, familial DCM). Type of evidence: interpretive/direct. Strengths/limitations: This is the authors' conclusion based on their genetic screen and functional assays; however, because evidence comes from one putative mutation in one family with limited additional data presented here, the association remains provisional and would benefit from replication, segregation analysis, population frequency data, and in vivo functional validation.


[Read Paper](https://www.semanticscholar.org/paper/d222bc0186a0b172e0e034097af7c3bda64d4e0b)


## Search Queries Used

- TMPO mutation pathogenic Mendelian disease case report family segregation exome sequencing trio de novo autosomal dominant familial thymopoietin LAP2

- TMPO thymopoietin LAP2 interaction LMNA lamin A C laminopathy nuclear envelope dilated cardiomyopathy mechanism protein interaction nuclear morphology

- TMPO ClinVar pathogenic likely pathogenic HGMD gnomAD pLI loss of function constraint allele frequency clinical significance TMPO variants

- TMPO knockout mouse knockdown zebrafish CRISPR functional study cell model nuclear envelope cardiac phenotype muscle phenotype thymopoietin LAP2


## Usage Statistics

- Prompt Tokens: 0

- Completion Tokens: 0

- Total Tokens: 0

- Estimated Cost: $0.0183


## Bibliometric Configuration

- Use Bibliometrics: True

- Author Impact Weight: 0.4

- Citation Impact Weight: 0.4

- Venue Impact Weight: 0.2
