## Supplementary material for "MENDELSEEK: An algorithm that predicts Mendelian Genes and elucidates what makes them special": TNFRSF10A.txt

# Claim: The gene TNFRSF10A is likely associated with at least one Mendelian disease.

**Status**: processed

**Overall Rating**: -1

**Explanation**:

No relevant papers were found that support or refute this claim.


## Relevant Papers


## Other Reviewed Papers


### BRCA2 loss‐of‐function germline mutations are associated with esophageal squamous cell carcinoma risk in Chinese

**Why Not Relevant**: The provided paper text contains no mention of TNFRSF10A (by gene name or synonym) and reports findings only for BRCA2 and a small set of other genes (POLQ, MSH2, TTC39B, KDR) in relation to esophageal squamous cell carcinoma (ESCC) genetic susceptibility. There is no direct evidence (no reported variants, associations, pedigrees, functional assays, or clinical phenotypes) relating TNFRSF10A to any Mendelian disease in this excerpt. Mechanistic evidence relevant to TNFRSF10A is also absent: the paper discusses defective homologous recombination and BRCA2-related synthetic lethality with PARP1 inhibitors, which does not implicate TNFRSF10A or its pathway. While the absence of a mention in this paper does not disprove any association between TNFRSF10A and Mendelian disease in other literature, this paper provides neither supporting nor refuting data for the claim. Limitations: the study focuses on ESCC predisposition in Chinese cohorts and on a targeted set of genes; if TNFRSF10A were not part of the investigated genes or not significantly associated, it would not be reported here. In summary, relevance of this paper to the claim about TNFRSF10A is effectively zero because TNFRSF10A is not discussed and no data concerning it are presented.


[Read Paper](https://www.semanticscholar.org/paper/44298aea73f11273c252f3e90fbaadb6bf04d3b8)


### The Burden of Candidate Pathogenic Variants for Kidney and Genitourinary Disorders Emerging From Exome Sequencing

**Why Not Relevant**: The supplied paper content does not mention the gene TNFRSF10A and provides no gene-level data or statements about TNFRSF10A. The manuscript excerpt describes the authors' approach to assembling a curated list of genes associated with Mendelian kidney and genitourinary disorders, methods for variant annotation and filtering, and the prevalence of candidate pathogenic variants in a cohort of ostensibly healthy adults, but no specific results, citations, or mechanistic discussion involving TNFRSF10A are present in the provided text. Therefore there is no direct evidence (case reports, cohort associations, variant pathogenicity data) nor mechanistic evidence (functional studies, described pathogenic variants, or described disease mechanisms) in this content that supports or refutes the claim that TNFRSF10A is likely associated with at least one Mendelian disease. 

Relevant contextual points from the paper that explain why it cannot inform the claim: (1) the study focuses on genes implicated in kidney and genitourinary Mendelian disorders and assembled a final list of 625 genes, but the provided excerpt does not include the gene list or indicate whether TNFRSF10A is among them; (2) the paper uses databases such as OMIM, Orphanet, ClinVar, and HGMD to identify and annotate genes and variants — databases that would be appropriate sources to check for TNFRSF10A — but the excerpt contains no extracted entries or variant-level findings for TNFRSF10A; (3) the cohort and analytic pipelines described (WES/WGS of 7,974 unrelated adults, stringent filtering, evaluation by inheritance mode) are relevant methodologically but do not yield any TNFRSF10A-specific evidence in the text provided. 

If you would like, I can (a) search ClinVar, HGMD, OMIM, and the primary literature for any reported Mendelian disease associations for TNFRSF10A and extract verbatim evidence and mechanistic data, or (b) review the paper's full gene list/appendices (if you can provide them) to see whether TNFRSF10A was included and whether any variants in that gene were observed and how they were classified.


[Read Paper](https://www.semanticscholar.org/paper/a75e297208c04992568e607d19aa9b9aaebd09d5)


### Fucose is on the TRAIL of colon cancer.

**Why Not Relevant**: The paper discusses the role of fucosylation in modulating TRAIL receptor function (DR4/DR5), provides mechanistic and experimental evidence in cancer cell lines and mouse xenografts, and notes structural/biochemical features of DR4 (TNFRSF10A) such as an N-glycosylation site. However, it contains no human genetic data (no germline variants, no segregation or case reports), no clinical descriptions of inherited phenotypes, and no statistical or epidemiological evidence linking TNFRSF10A variants to a Mendelian disorder. The experiments are performed in cancer cell lines (somatic context) and focus on post-translational modification and tumor sensitivity to apoptosis; these contexts do not provide direct evidence that TNFRSF10A is associated with any Mendelian disease. Therefore the paper offers mechanistic relevance to TNFRSF10A function but does not provide the type of genetic or clinical evidence required to support the claim that TNFRSF10A is likely associated with at least one Mendelian disease.


[Read Paper](https://www.semanticscholar.org/paper/7cb823582c4547dc4e7029cf9c7083b772bb8696)


### An investigation of the association of genetic susceptibility risk with somatic mutation burden in breast cancer

**Why Not Relevant**: The provided paper content consists of a single sentence: "An inverse association between common germline risk variants and TSMC is observed, which, if confirmed, could provide new insights into how germline variation informs the understanding of somatic mutation patterns in breast cancer." This sentence makes no mention of the gene TNFRSF10A, Mendelian disease, pathogenic variants, family segregation, clinical phenotypes, or any gene-specific mechanistic evidence. Therefore it provides neither direct evidence (e.g., identification of TNFRSF10A variants causing a Mendelian disorder, segregation data, or clinical case reports) nor mechanistic evidence (e.g., functional impact of TNFRSF10A variants consistent with Mendelian inheritance) relevant to the claim. Additional limitations: the excerpt does not define TSMC, does not present statistical details or variant-level data, and is focused on population-level associations between germline variation and somatic mutation patterns in breast cancer rather than on monogenic disease associations. To evaluate the claim that "TNFRSF10A is likely associated with at least one Mendelian disease" would require explicit evidence such as (a) reports of rare, pathogenic TNFRSF10A variants segregating with a Mendelian phenotype, (b) clinical descriptions linking TNFRSF10A disruption to a specific syndrome or disease in affected families, (c) functional studies showing a mechanism consistent with Mendelian pathogenicity (loss-of-function, dominant-negative, gain-of-function), or (d) database annotations (e.g., OMIM) naming a Mendelian disorder caused by TNFRSF10A. None of these are present in the supplied content, so the relevance to the claim is effectively zero.


[Read Paper](https://www.semanticscholar.org/paper/574e7719c6ed81e0a8379a0abda9265ab723a991)


### Whole-Exome Sequencing in Adults With Chronic Kidney Disease

**Why Not Relevant**: The provided paper content does not mention TNFRSF10A by name and contains no gene-level results or diagnostic findings that implicate TNFRSF10A in a Mendelian disease. The excerpts show the study's scope (curated OMIM/nephropathy genes and ACMG actionable genes) and methods for variant detection and interpretation, but they do not include the gene lists (Supplement Table 3) or results that would allow determination of whether TNFRSF10A was included or found to be associated with Mendelian disease. Without explicit mention of TNFRSF10A or access to the Supplement tables and result sections, this paper does not provide direct or strong mechanistic evidence for or against the claim. For these reasons relevance to the specific claim is very low.


[Read Paper](https://www.semanticscholar.org/paper/a29bd57964f68ac5f8e1abcafe1112ba488f53f5)


### Precision Medicine in Internal Medicine

**Why Not Relevant**: The provided paper content is a general clinical review about genetic testing technologies, variant interpretation, and the current state of gene–disease discovery. It contains no mention of the specific gene TNFRSF10A, no case reports, cohort data, functional studies, linkage/segregation evidence, population allele frequencies, ClinVar entries, or OMIM entries specific to TNFRSF10A. Therefore it supplies no direct evidence that TNFRSF10A is associated with any Mendelian disease. The included excerpts are only indirectly relevant (they describe methods and resources by which such an association might be discovered or evaluated) but do not support or refute the claim about TNFRSF10A itself.


[Read Paper](https://www.semanticscholar.org/paper/c7199e8a5226f8d9d55777da0e15e91797bb459a)


### Patients with coronary heart disease, dilated cardiomyopathy and idiopathic ventricular tachycardia share overlapping patterns of pathogenic variation in cardiac risk genes

**Why Not Relevant**: This paper does not provide evidence for or against the claim that TNFRSF10A is likely associated with at least one Mendelian disease. The study is a targeted sequencing investigation of 96 cardiac genes in 92 patients with ventricular tachycardia (VT) of various etiologies; the authors report findings in specific cardiac-related genes (for example LAMA2, MYBPC3, MYH6, KCNQ1, GAA, DSG2, TTN) and give aggregate variant counts and ACMG classifications. Nowhere in the provided text is TNFRSF10A mentioned, nor are results reported for that gene. Therefore there is no direct genetic association evidence (no reported pathogenic or likely pathogenic variants in TNFRSF10A) and no mechanistic discussion linking TNFRSF10A to Mendelian disease in this paper. Important caveats: (1) absence of mention in this paper is not evidence that TNFRSF10A is not associated with a Mendelian disease — it only shows this particular study did not provide such evidence; (2) the paper does not list the full 96-gene panel in the supplied excerpt, so it is theoretically possible TNFRSF10A was not included in the panel or was included but yielded no noteworthy variants; (3) the study sample is small (92 patients) and focused on cardiac phenotypes, so it would not be a comprehensive study for Mendelian disease associations beyond its targeted scope. Given these points, the relevance of this paper to the specific claim about TNFRSF10A is negligible.


[Read Paper](https://www.semanticscholar.org/paper/3f3b535d385c9b16f6a599120d890be36b90f4b6)


### Whole exome sequencing identifies recessive germline mutations in FAM160A1 in familial NK/T cell lymphoma

**Why Not Relevant**: The provided paper content exclusively discusses FAM160A1 mutations in a familial NKTL case and does not mention TNFRSF10A, its variants, function, pathogenicity, genotype-phenotype correlations, or any mechanistic pathway involving TNFRSF10A. Therefore the paper provides no direct evidence (genetic association, segregation, variant pathogenicity) nor mechanistic evidence (biochemical pathway, functional assay, animal model) supporting or refuting the claim that TNFRSF10A is likely associated with at least one Mendelian disease. Because TNFRSF10A is not addressed at all, the content cannot be used to evaluate the claim; any inference about TNFRSF10A would require additional literature that specifically investigates that gene.


[Read Paper](https://www.semanticscholar.org/paper/08388b16f9f94a55459297e29336a0634b1c17ec)


### Germline variation networks in the PI3K/AKT pathway corresponding to familial high-incidence lung cancer pedigrees

**Why Not Relevant**: The provided paper content focuses solely on germline SNP enrichment in the PI3K/AKT pathway as a predictor of familial susceptibility to lung cancer. It does not mention TNFRSF10A, nor does it discuss Mendelian diseases or single-gene (monogenic) inheritance. Therefore the supplied content is not relevant to the claim that "The gene TNFRSF10A is likely associated with at least one Mendelian disease." Additional reasons: (1) The subject pathway (PI3K/AKT) differs from TNFRSF10A's canonical biology (TNFRSF10A encodes TRAIL receptor 1, part of the TNF receptor superfamily involved in apoptosis signaling), (2) familial susceptibility to lung cancer is a complex-trait topic, not a statement about Mendelian (monogenic) disease associations, and (3) only a brief summary sentence was provided, preventing review of full methods/results where TNFRSF10A might appear. To evaluate the claim about TNFRSF10A, review of primary resources that catalogue Mendelian gene–disease relationships (e.g., OMIM, ClinVar, HGMD) and a targeted search of the full paper text and related literature would be required.


[Read Paper](https://www.semanticscholar.org/paper/a7baf3f9c9c6c8ffe7c7a61f9d754a2fa206d085)


### Clingen Coagulation Factor Deficiency Variant Curation Expert Panel: Meeting the Need for Recommendations to Curate Variants in the Coagulation Factor Genes

**Why Not Relevant**: The paper's content is focused on the development and specification of ACMG/AMP variant curation guidelines for coagulation factor genes, specifically F8 and F9, and on curating reported variants in those genes. The text supplied contains no mention of TNFRSF10A, no data (genetic, clinical or population) about TNFRSF10A variants, and no mechanistic discussion linking TNFRSF10A to any Mendelian phenotype. The provided excerpts instead explicitly identify F8 and F9 as the genes under study and describe their known Mendelian associations (Hemophilia A and B). While the methods and framework described (CFD-VCEP variant curation) could in principle be applied to other genes in the future, the paper does not present direct evidence (case reports, segregation, functional assays, variant-disease associations) nor mechanistic evidence (pathway involvement, molecular function leading to disease phenotype) pertaining to TNFRSF10A. Therefore the paper neither supports nor refutes the claim about TNFRSF10A and Mendelian disease, and its relevance to that claim is negligible.


[Read Paper](https://www.semanticscholar.org/paper/9ea8d4ce19622ae1c6b0cdfd40848c93199f6280)


## Search Queries Used

- TNFRSF10A germline variant Mendelian disease case report pathogenic variant monogenic disease TRAIL receptor 1 DR4

- TNFRSF10A ClinVar OMIM HGMD gene disease association pathogenic germline variant clinical genetics diagnostic panel

- TNFRSF10A TRAIL receptor 1 DR4 apoptosis death receptor signaling loss of function mutation functional study mouse knockout phenotype

- TNFRSF10A germline versus somatic mutation cancer susceptibility association study exome sequencing familial cancer risk polymorphism


## Bibliometric Configuration

- Use Bibliometrics: True

- Author Impact Weight: 0.4

- Citation Impact Weight: 0.4

- Venue Impact Weight: 0.2
