## Supplementary material for "MENDELSEEK: An algorithm that predicts Mendelian Genes and elucidates what makes them special": TNFRSF17.txt

# Claim: The gene TNFSF13 is likely associated with at least one Mendelian disease.

**Status**: processed

**Overall Rating**: 2

**Explanation**:

Supporting evidence:
The provided papers examine APRIL (TNFSF13) in the context of primary antibody deficiency, particularly common variable immunodeficiency (CVID) and IgA deficiency. They report elevated BAFF and APRIL protein levels in patients versus healthy controls, and they explicitly screened TNF family genes including APRIL (TNFSF13) for mutations in affected cohorts. Those screens did not identify causative coding-region mutations in APRIL in the studied patients. Thus the strongest direct evidence in the provided material is negative: APRIL sequence variation was not found to explain Mendelian CVID in these cohorts.

Caveats or contradictory evidence:
Elevated APRIL expression in patients shows that TNFSF13 is biologically relevant to immune function and to the pathophysiology of antibody deficiency disorders, but altered expression is not equivalent to a Mendelian (monogenic) causal relationship. The absence of detected coding mutations in these particular cohorts does not strictly rule out the existence of rare APRIL (TNFSF13) mutations causing Mendelian disease in other families or via noncoding regulatory variants, structural variants, or oligogenic mechanisms. Sample sizes and cohort composition limit detection power for extremely rare, private, or noncoding pathogenic variants. Also, the papers focus on a subset of primary antibody deficiencies and may not have surveyed all phenotypes where TNFSF13 could be causal.

Analysis of mechanisms and second-order implications:
Biologically, APRIL (encoded by TNFSF13) is a B‑cell–stimulating ligand that binds receptors such as TACI (TNFRSF13B) and BCMA (TNFRSF17) and participates in B‑cell survival and class-switch recombination. TACI (a receptor) has documented pathogenic mutations in some CVID patients, showing that perturbation of the APRIL–TACI axis can produce Mendelian immune phenotypes. However, redundancy in the BAFF/APRIL system (overlapping ligands and receptors) could buffer loss-of-function in APRIL itself, making monogenic disease from TNFSF13 loss less likely than from receptor defects. Conversely, a gain-of-function mutation in TNFSF13 theoretically could cause disease, but no such mutations were reported in the provided screens. The regulatory complexity and extracellular ligand nature also make phenotype expressivity and penetrance variable, which complicates simple Mendelian attribution.

Why the rating "Likely False" is appropriate:
The available direct evidence from the provided papers is negative: targeted mutation screens in relevant patient cohorts did not identify causative TNFSF13 coding variants. While the biology implicates APRIL in immune function and receptor mutations in the same pathway (e.g., TACI) do cause Mendelian immunodeficiencies, the lack of observed APRIL mutations in these studies, combined with plausible functional redundancy in the pathway, argues that TNFSF13 is unlikely to be a common cause of Mendelian disease. This does not absolutely exclude very rare or untested mechanisms (noncoding variants, structural changes, or phenotypes outside the studied cohorts), but given the negative screening evidence and the biological considerations, the claim that "TNFSF13 is likely associated with at least one Mendelian disease" is not well supported by the provided literature and is therefore more likely false than true.


**Final Reasoning**:

The primary direct evidence supplied consists of genetic screens of patients with primary antibody deficiency that explicitly included APRIL (TNFSF13) and failed to find causative coding mutations, while showing only expression-level changes. Given that receptor genes in the APRIL/BAFF pathway (e.g., TACI/TNFRSF13B) have established Mendelian links but APRIL itself was not found mutated in these cohorts, the balance of evidence argues against TNFSF13 being a likely cause of a Mendelian disease. Biological plausibility for an effect exists, but redundancy and the negative screening results reduce the likelihood that TNFSF13 is commonly causal for a monogenic disorder. Because absence of evidence in these targeted studies is not an absolute disproof, the correct strength-of-evidence judgement is "Likely False": the claim is unlikely given current data but cannot be definitively refuted for all possible rare mechanisms or unstudied phenotypes.


## Relevant Papers


### Age-related changes in BAFF and APRIL profiles and upregulation of BAFF and APRIL expression in patients with primary antibody deficiency.

**Authors**: R. Jin (H-index: 23), N. Kondo (H-index: 42)

**Relevance**: 0.62

**Bibliometric Impact**: 0.381375


**Excerpts**:

- In some patients with common variable immunodeficiency (CVID) and immunoglobulin (Ig) A deficiency (IgAD), tumor necrosis factor (TNF) family receptor transmembrane activator and calcium-modulator and cyclophilin ligand interactor (TACI) gene mutations have been reported.

- B cells from individuals with TACI mutations do not produce IgG and IgA in response to the TACI ligand a proliferation-inducing ligand (APRIL) which probably suggests impaired isotype switching.

- To clarify the pathogenesis of CVID and IgAD of Japanese patients, we investigated the mutations of TNF family members TACI, APRIL, B-cell activating factor (BAFF), B-cell maturation antigen (BCMA) and BAFF receptor (BAFF-R) genes and the expression levels of BAFF and APRIL in patients with CVID, IgAD and X-linked agammaglobulinaemia (XLA).

- The causative gene mutations of TNF family members in our patients were not detected.

- In healthy subjects, the BAFF and APRIL plasma levels correlated inversely with age.

- The BAFF and APRIL plasma levels of patients with CVID, IgAD and XLA were significantly higher than those of healthy children.

- Elevated BAFF and APRIL expression levels might partially reflect the common immunological feature of primary antibody deficiency.


**Explanations**:

- This sentence provides background linking mutations in a TNF-family receptor gene (TACI) to Mendelian primary immunodeficiency phenotypes (CVID and IgAD). It is indirectly relevant because it demonstrates that TNF-family genes can be causal in Mendelian immune disorders. Strength: establishes precedent that TNF-family gene defects cause Mendelian disease. Limitation: TACI is a receptor (TNFRSF13B), not TNFSF13 (APRIL) itself, so this is circumstantial rather than direct evidence implicating TNFSF13.

- This sentence is mechanistic: it links TACI mutations to a functional B-cell defect (failure to produce IgG and IgA upon stimulation with APRIL). It implicates APRIL (the product of TNFSF13) in isotype switching biology and therefore provides mechanistic plausibility that perturbing APRIL or its pathway could produce immunodeficiency. Strength: direct functional connection between APRIL signaling and antibody isotype production. Limitation: the statement concerns effects of TACI mutations on response to APRIL, not mutations in APRIL (TNFSF13) itself; it shows pathway relevance but does not demonstrate that TNFSF13 mutations cause disease.

- This sentence documents that the investigators specifically examined APRIL (TNFSF13) among other TNF-family genes in patients with CVID, IgAD and XLA. It is direct evidence of an effort to detect Mendelian (causative) variants in TNFSF13 in affected patients. Strength: shows targeted genetic analysis of TNFSF13 in relevant Mendelian phenotypes. Limitation: the sentence states the experimental aim but does not provide results or methods (e.g., sequencing depth, variant interpretation criteria, sample size), so interpretation of negative or positive findings depends on later statements.

- This sentence gives the primary negative result: "The causative gene mutations of TNF family members in our patients were not detected." This directly refutes the presence of causative TNF-family gene mutations (including APRIL/TNFSF13) in this patient cohort. Strength: direct negative finding. Limitations: the statement is cohort- and method-dependent (Japanese patients only, unspecified sample size and sequencing/variant-detection methods). A failure to detect causative mutations in this study does not exclude TNFSF13 as causative of Mendelian disease in other cohorts or by other mutation types (e.g., regulatory variants, structural variants, mosaicism) or by different mechanisms not detectable by the authors' methods.

- This sentence provides physiological context showing that APRIL (TNFSF13 product) levels vary with age in healthy subjects (inverse correlation). This is mechanistic/physiologic evidence relevant to interpreting APRIL measurements in disease cohorts. Strength: informs on normal APRIL regulation, which is useful when evaluating elevated APRIL in patients. Limitation: it does not bear directly on genetic causation.

- This sentence reports that APRIL (and BAFF) plasma levels are elevated in patients with CVID, IgAD and XLA relative to healthy children. It is evidence of association between altered APRIL expression and primary antibody deficiency phenotypes, which is mechanistically relevant because it shows APRIL dysregulation in disease. Strength: supports involvement of APRIL-related pathways in disease states. Limitation: elevated protein expression can be a secondary/compensatory effect rather than evidence of a causal genetic mutation in TNFSF13; this is not genetic evidence of Mendelian causation.

- This interpretive sentence suggests elevated APRIL/BAFF levels may reflect a shared immunological feature of primary antibody deficiency. It reinforces that APRIL pathway perturbation is associated with these conditions, strengthening biological plausibility that TNFSF13-related perturbation might influence disease. Limitation: the authors frame this as a partial reflection/association and do not claim direct genetic causation by TNFSF13 mutations; it remains correlative rather than definitive genetic evidence of Mendelian disease linkage.


[Read Paper](https://www.semanticscholar.org/paper/ced605b6a498e81585c1d0de057439bb7ba5ec6f)


### Screening of functional and positional candidate genes in families with common variable immunodeficiency

**Authors**: U. Salzer (H-index: 54), B. Grimbacher (H-index: 88)

**Relevance**: 0.18

**Bibliometric Impact**: 0.48102500000000004


**Excerpts**:

- It is unable to identify obvious disease causing mutations in the protein coding regions of the analyzed genes in the studied cohort of individuals with CVID.


**Explanations**:

- Relation to claim: This sentence provides negative evidence relevant to the claim because it reports that the study did not find "obvious disease causing mutations" in the protein-coding regions of the analyzed genes in a cohort of individuals with CVID. If TNFSF13 was among the "analyzed genes," this statement would be direct evidence against TNFSF13 being an obvious cause of Mendelian disease in that cohort. Evidence type: direct (negative finding). Strength: The statement is a direct report of sequencing/variant-detection results in a disease cohort, which can directly refute a simple hypothesis that TNFSF13 harbors obvious coding mutations causing the CVID cases studied. Limitations and caveats: The sentence is brief and non-specific — it does not name which genes were analyzed, does not explicitly mention TNFSF13, and gives no details about sample size, sequencing depth, variant-calling methods, or criteria for calling a mutation "obvious" and "disease causing." It also restricts the search to "protein coding regions," so it does not rule out non-coding (regulatory, promoter, intronic, splice regulatory) variants, structural variants not detectable by the methods used, mosaicism, or oligogenic/complex contributions. Finally, absence of evidence in a single cohort (with a specific phenotype, CVID) does not preclude TNFSF13 causing other Mendelian diseases or causing disease in other populations or via mechanisms not assayed. Therefore, this excerpt provides limited negative evidence against the claim but is insufficient to conclude that TNFSF13 is unlikely to be associated with any Mendelian disease.


[Read Paper](https://www.semanticscholar.org/paper/068021af99b7ee6dc3402112a20459721bd2465e)


## Other Reviewed Papers


### The role of costimulation in antibody deficiencies: ICOS and common variable immunodeficiency

**Why Not Relevant**: The paper content provided is entirely focused on ICOS (inducible costimulator) gene mutations and ICOS deficiency as a monogenic cause of common variable immunodeficiency. It contains mechanistic descriptions and clinical consequences for ICOS but does not mention TNFSF13 (APRIL) or present any genetic, clinical, or functional data connecting TNFSF13 to a Mendelian disorder. Therefore the paper provides no direct evidence for the claim that TNFSF13 is likely associated with at least one Mendelian disease. To evaluate the claim about TNFSF13 one would need papers reporting (a) pathogenic TNFSF13 variants segregating with a Mendelian phenotype, (b) case series or database/OMIM entries documenting TNFSF13-related Mendelian disease, and/or (c) functional studies showing variant impact on protein function consistent with the disease mechanism. None of those elements are present in this ICOS-focused summary, so relevance to the TNFSF13 claim is essentially nil.


[Read Paper](https://www.semanticscholar.org/paper/64c7593660d1df3a19650df957817b178c7d5659)


### Evaluation of B‐cell intracellular signaling by monitoring the PI3K‐Akt axis in patients with common variable immunodeficiency and activated phosphoinositide 3‐kinase delta syndrome

**Why Not Relevant**: The provided paper content does not mention TNFSF13 (APRIL) or any cognate names (for example TNFSF13B, BAFF, APRIL) and therefore contains no direct evidence that TNFSF13 is associated with any Mendelian disease. The excerpt focuses on primary antibody deficiencies (PADs), common variable immunodeficiency (CVID), and activated phosphoinositide 3-kinase delta syndrome (APDS) caused by gain-of-function mutations in PIK3CD, and on analyzing the PI3K-Akt-mTOR pathway in B cells. While it discusses monogenic disease in the context of PIK3CD (APDS) and describes intracellular signaling pathways relevant to B-cell homeostasis, it provides no mechanistic statements, experimental data, or conclusions that implicate TNFSF13 in Mendelian disease. Limitations: this assessment is based solely on the supplied excerpt; if the full paper contains additional sections not provided here that discuss TNFSF13, those were not available for review. For evidence about TNFSF13 specifically, consult genetic disease databases (OMIM, ClinVar) and primary studies that explicitly assay TNFSF13 sequence variants or function in Mendelian contexts.


[Read Paper](https://www.semanticscholar.org/paper/a3b15eca7a03a605c6fa23a5104e80bafb17d605)


### Expanding the Role of SH2B3: T Cell Large Granular Lymphocytosis (LGL) and Common Variable Immune Deficiency in the Setting of Biallelic SH2B3 Mutations

**Why Not Relevant**: The provided paper content exclusively discusses the gene SH2B3 (LNK) and a single-patient case with biallelic SH2B3 variants and associated phenotypes (ITP, CVID, T-LGL, MPN-like features). There is no mention of TNFSF13, any variants in TNFSF13, functional data involving TNFSF13, or any mechanistic linkage between SH2B3 and TNFSF13. Because the paper contains no direct or mechanistic evidence concerning TNFSF13, it cannot support or refute the claim that TNFSF13 is likely associated with at least one Mendelian disease. Any inference about TNFSF13 from this paper would be speculative and unsupported by the text. Therefore the relevance to the claim is effectively zero.


[Read Paper](https://www.semanticscholar.org/paper/e1a40a9f39825e2b46d50b76730dd38ea2148355)


## Search Queries Used

- TNFSF13 APRIL pathogenic variant Mendelian disease human loss of function familial segregation primary immunodeficiency

- APRIL mutation common variable immunodeficiency CVID familial antibody deficiency monogenic B cell deficiency TNFSF13

- TNFSF13 APRIL knockout mouse phenotype B cell survival class switch recombination immunoglobulin deficiency loss of function model

- TNFSF13 review Mendelian disease OMIM ClinVar gene nomenclature TNFSF13B BAFF confusion clinical genetics summary


## Usage Statistics

- Prompt Tokens: 0

- Completion Tokens: 0

- Total Tokens: 0

- Estimated Cost: $0.0260


## Bibliometric Configuration

- Use Bibliometrics: True

- Author Impact Weight: 0.4

- Citation Impact Weight: 0.4

- Venue Impact Weight: 0.2
