## Supplementary material for "MENDELSEEK: An algorithm that predicts Mendelian Genes and elucidates what makes them special": TRAPPC3.txt

# Claim: The gene TRAPPC3 is likely associated with at least one Mendelian disease.

**Status**: processed

**Overall Rating**: 0

**Explanation**:

Supporting evidence.
The provided excerpts include a study that systematically annotated upstream open reading frames (uORFs) and translation initiation sites in OMIM-listed disease genes and reported that disruption of uORFs by variants (cataloged in HGMD/ClinVar) can cause Mendelian disorders. That work demonstrates, in general, that non-coding variants (specifically 5′UTR/uORF-disrupting changes) are a credible mechanism for Mendelian disease and that systematic annotation can reveal new disease-relevant variants. The second paper reports an enrichment of genes involved in intracellular protein transport among de novo variants in autism spectrum disorder, supporting the broader plausibility that genes functioning in intracellular trafficking (a category that includes TRAPP complex subunits) can be implicated in human disease.

Caveats or contradictory evidence.
Neither paper provides any direct evidence linking TRAPPC3 specifically to a Mendelian disorder. The uORF/OMIM study lists several genes for which they experimentally validated uORF effects (ETFDH, PAX9, MAST1, HTT, TTN, GLI2, COL2A1, ZIC2), but TRAPPC3 is not mentioned in the excerpts. The ASD trio study enumerates pathogenic or candidate genes discovered in that cohort; TRAPPC3 is not among them. Thus, while the general mechanisms reported are relevant to disease biology, neither excerpt contains a reported pathogenic variant, ClinVar/HGMD entry, OMIM entry, or experimental evidence implicating TRAPPC3 in a Mendelian phenotype.

Analysis of mechanisms and implications.
Biologically, TRAPPC3 encodes a subunit of the TRAPP complex involved in intracellular vesicle trafficking. This class of proteins has precedent for Mendelian disease associations (other TRAPP subunits are implicated in neurodevelopmental and muscular disorders). Mechanistically, loss-of-function or dominant-negative variants in trafficking machinery can plausibly produce Mendelian phenotypes. However, plausibility alone does not constitute evidence of an actual disease-gene relationship. To move from plausibility to supported association requires direct genetic evidence (recurrent pathogenic variants, segregation with disease, ClinVar/HGMD/OMIM entries, functional validation), none of which is shown in the provided excerpts for TRAPPC3.

Rating justification.
Given the available evidence, the excerpts support that genes like TRAPPC3 (intracellular trafficking genes) are plausible disease candidates, but they do not present any direct or specific evidence that TRAPPC3 is associated with a Mendelian disorder. The correct evaluation based on these papers is therefore that there is no evidence in the provided materials to support the claim. The lack of any specific mention of TRAPPC3, absence of reported pathogenic variants in that gene, and absence of OMIM/ClinVar/HGMD entries in the excerpts means the claim is not supported by the supplied literature.


**Final Reasoning**:

I reviewed the provided excerpts and considered mechanistic plausibility and precedent. The uORF study and the ASD exome study provide general, indirect support that noncoding variation and intracellular trafficking genes can be disease-relevant, which makes TRAPPC3 a plausible candidate in principle. However, neither paper contains any explicit observation, variant report, experimental validation, or database annotation tying TRAPPC3 to a Mendelian phenotype. The standards for asserting that a gene is "likely associated with at least one Mendelian disease" require specific genetic or clinical evidence (e.g., pathogenic variants in ClinVar/HGMD, OMIM disease entry, segregation data, recurrent de novo mutations, or functional validation), none of which appears in the provided evidence. Therefore the appropriate rating based solely on the supplied papers is "No Evidence." If additional papers or database entries specific to TRAPPC3 were provided, the rating could be revisited.


## Relevant Papers


### Annotation of uORFs in the OMIM genes allows to reveal pathogenic variants in 5′UTRs

**Authors**: A. Filatova (H-index: 8), M. Skoblov (H-index: 5)

**Relevance**: 0.15

**Bibliometric Impact**: 0.23600000000000004


**Excerpts**:

- An increasing number of studies emphasize the role of non-coding variants in the development of hereditary diseases.

- It was previously shown that variants in 5′-untranslated regions (5′UTRs) can lead to hereditary diseases due to disruption of upstream open reading frames (uORFs).

- Here, we performed a manual annotation of upstream translation initiation sites (TISs) in human disease-associated genes from the OMIM database and revealed ∼4.7 thousand of TISs related to uORFs.

- We revealed the variants from the HGMD and ClinVar databases that disrupt uORFs and thereby could lead to Mendelian disorders.

- Using a luciferase assay, we experimentally validated the translation of uORFs in the ETFDH, PAX9, MAST1, HTT, TTN,GLI2 and COL2A1 genes, as well as existence of N-terminal CDS extension in the ZIC2 gene.

- Besides, we created a tool to annotate the effects of genetic variants located in uORFs. We revealed the variants from the HGMD and ClinVar databases that disrupt uORFs and thereby could lead to Mendelian disorders. We also showed that the distribution of uORFs-affecting variants differs between pathogenic and population variants.


**Explanations**:

- This sentence gives general context that non-coding variants are increasingly implicated in hereditary disease. Relation to claim: indirect background that supports plausibility that non-coding effects (including in any specific gene such as TRAPPC3) can underlie Mendelian phenotypes. Evidence type: mechanistic/contextual. Limitations: general statement, no gene-specific data, no mention of TRAPPC3.

- This sentence states a specific mechanistic route—disruption of 5′UTR uORFs—as a cause of hereditary disease. Relation to claim: mechanistic evidence that if TRAPPC3 harbors pathogenic uORF-disrupting variants, such variants could plausibly cause a Mendelian disorder. Evidence type: mechanistic. Limitations: does not present data on TRAPPC3, does not show that TRAPPC3 actually has such variants or that they are pathogenic.

- This sentence indicates the authors annotated upstream TISs across OMIM disease-associated genes and found ~4.7k TISs related to uORFs. Relation to claim: methodological scope that could include TRAPPC3 if it is an OMIM-listed gene; suggests the authors generated resources that might identify uORF-mediated pathogenicity in many genes. Evidence type: indirect/operational. Limitations: abstract-level statement only; no gene-level output in the excerpt, so whether TRAPPC3 was annotated or found to be implicated is unknown.

- This sentence claims that variants from ClinVar and HGMD that disrupt uORFs were identified and could lead to Mendelian disorders. Relation to claim: indicates the study connects uORF-disrupting variants to Mendelian disease at the variant/database level. Evidence type: indirect; potentially directly relevant if TRAPPC3 harbors such variants in those databases, but the excerpt does not specify which genes are affected. Limitations: no gene-by-gene listing here—no explicit mention of TRAPPC3 or statistical details (counts, pathogenicity classifications), so cannot conclude TRAPPC3 is implicated based solely on this sentence.

- This sentence lists genes for which the authors experimentally validated uORF translation (ETFDH, PAX9, MAST1, HTT, TTN,GLI2, COL2A1) and a CDS extension in ZIC2. Relation to claim: demonstrates experimental validation of the mechanism in specific genes, strengthening the biological plausibility that uORF disruption can cause Mendelian disease. Evidence type: mechanistic/experimental. Limitations: TRAPPC3 is not among the validated genes; absence of TRAPPC3 here means no direct experimental evidence from this paper supporting TRAPPC3's association with Mendelian disease.

- These sentences describe the creation of an annotation tool and reiterate that the authors found uORF-disrupting variants in ClinVar/HGMD and observed distributional differences between pathogenic and population variants. Relation to claim: methodological and statistical context indicating the study systematically links uORF disruption to pathogenic variants across databases. Evidence type: mechanistic/statistical. Limitations: abstract-level summary; no gene-specific output provided here, so whether TRAPPC3 is among the genes with such variants is not indicated. Furthermore, the excerpt lacks details (numbers, effect sizes, criteria) necessary to assess the strength of the variant-level associations.


[Read Paper](https://www.semanticscholar.org/paper/5d45c0cdf801505df45fc93a466a2a4101a12871)


### Characterization of de novo variants in exomes of individuals with autism spectrum disorder

**Authors**: C. G. Lopes (H-index: 6), M. R. Bueno (H-index: 8)

**Relevance**: 0.15

**Bibliometric Impact**: 0.136


**Excerpts**:

- The main objective of this essay was to contribute to the characterization of the genetic architecture of autism spectrum disorder (ASD) based on an analysis of a Brazilian series, which is still little studied.

- Sixty-three trios were evaluated, composed of parents and probands diagnosed with ASD treated at the Human Genome and Stem Cell Studies Center (CEGH-CEL, USP).

- Whole-exome sequencing was performed through a collaboration with Mount Sinai, New York, United States (collaboration with the Autism Sequencing ConsortiumACS).

- Nine de novo pathogenic or potentially pathogenic variants were identified in candidate genes: for TEA: six in SFARI genes (four pathogenic variants in NF1, TLK2, DNAH17, BRSK2 genes, and two probably pathogenic variants in ARHGAP5 and HUWE1) and three in genes of neurodevelopment of DECIPHER (Deciphering Developmental Disorders Study, 2015) (two pathogenic variants in the ERLIN2, ST3GAL3 genes and one probably pathogenic in COL11A1).

- When performing the gene enrichment analysis of genes with pathogenic or potentially pathogenic variants, we observed the enrichment of genes for intracellular protein transport.

- This study suggests that de novo variants are also an essential mechanism for the etiology of ASD in Brazil, explaining the genetic architecture of 9.5% of cases.


**Explanations**:

- Context/background: Describes the study's purpose (characterizing genetic architecture of ASD in a Brazilian cohort). Relevance to the TRAPPC3 claim: contextual only — it establishes the paper's focus on genetic causes of ASD but contains no mention of TRAPPC3. Strengths/limitations: useful context but provides no direct or mechanistic evidence for TRAPPC3.

- Method/sample size: States the number of trios (63) analyzed. Relevance: indicates modest sample size which limits power to detect rare disease associations (including potential TRAPPC3 Mendelian associations). Strength/limitations: small cohort reduces chance of observing rare causal variants like those in TRAPPC3; does not provide evidence for or against TRAPPC3 itself.

- Method/assay: Notes use of whole-exome sequencing in trios. Relevance: shows appropriate technology to detect coding de novo variants, the class of evidence relevant to Mendelian disease gene discovery. Strength/limitations: WES can detect coding SNVs/indels but may miss noncoding variation, copy-number variants, or some structural variants; nonetheless, the excerpt contains no information about TRAPPC3 variants specifically.

- Results/gene list: Lists the nine de novo pathogenic or likely pathogenic variants and the specific genes observed (NF1, TLK2, DNAH17, BRSK2, ARHGAP5, HUWE1, ERLIN2, ST3GAL3, COL11A1). Relevance: direct evidence about which genes were implicated in these ASD cases. Strength/limitations: this is direct experimental evidence but TRAPPC3 is not among the reported genes, so this excerpt provides negative information (i.e., TRAPPC3 was not identified here). The small sample and limited number of identified variants mean absence here is not strong evidence against TRAPPC3 being a Mendelian disease gene in general.

- Mechanistic enrichment result: Reports enrichment of genes for intracellular protein transport among genes with pathogenic or potentially pathogenic variants. Relevance: mechanistic — TRAPPC3 is known (from external sources) to participate in intracellular protein trafficking as part of TRAPP complexes, so the reported pathway-level enrichment could be circumstantially consistent with involvement of other intracellular transport genes in neurodevelopmental phenotypes. Strength/limitations: the paper provides an association at the pathway level but does not name TRAPPC3, provide a variant in it, or give statistical details (e.g., p-values, effect sizes) in the excerpt. Thus this is indirect, suggestive mechanistic context only; it neither confirms nor robustly supports TRAPPC3 as a Mendelian disease gene.

- Interpretation/significance: States that de novo variants explain 9.5% of cases in this cohort and that de novo variants are an essential mechanism for ASD in Brazil. Relevance: provides overall significance of de novo coding variants in ASD etiology, indicating that trio WES can identify Mendelian/near-Mendelian contributions to neurodevelopmental disorders. Strength/limitations: reinforces that the study design can discover Mendelian disease genes, but it does not provide any specific evidence about TRAPPC3 and the 9.5% figure is cohort-specific and modest; absence of TRAPPC3 in the reported findings limits its direct relevance to the claim.


[Read Paper](https://www.semanticscholar.org/paper/ac00b973ad036a15626b0e5f721516a43f7c0909)


## Other Reviewed Papers


### Genotypic and phenotypic spectra of NBEA-related neurodevelopmental disorder with epilepsy: a case series and literature review

**Why Not Relevant**: The provided paper content does not mention TRAPPC3 at all. The single sentence supplied states: "The clinical data, including demographics, clinical features, auxiliary examination, and neuroimaging, were collected retrospectively from three patients who were admitted in the Department of Pediatrics, Xiangya Hospital of Central South University because of epileptic encephalopathy of unknown etiology and three novel NBEA variants were identified." This sentence concerns the gene NBEA and epileptic encephalopathy in three patients. It provides no direct evidence (no observations, variants, segregation, functional data, or disease associations) regarding TRAPPC3, nor does it provide mechanistic information that would link NBEA findings to TRAPPC3. Therefore the paper content as provided cannot support or refute the claim that TRAPPC3 is likely associated with at least one Mendelian disease. Limitations underlying this conclusion: (1) absence of the gene name TRAPPC3 in the excerpt means no direct observational evidence; (2) no mechanistic pathway or interaction data are presented that could plausibly connect NBEA variants to TRAPPC3-mediated pathology; (3) the excerpt is a retrospective clinical description of three patients (small sample) focused on a different gene, so it cannot be used to infer associations for other genes. To evaluate the claim about TRAPPC3 would require papers reporting TRAPPC3 variants in Mendelian disease cases (segregation, de novo occurrence, case-control enrichment, or functional studies), none of which are present in the provided text.


[Read Paper](https://www.semanticscholar.org/paper/1763b06ea7b8b1561a04aaf97102159eb2caf359)


### Exploring the Impact of Genetics in a Large Cohort of Moebius Patients by Trio Whole Exome Sequencing

**Why Not Relevant**: This paper does not mention TRAPPC3 at all and therefore provides no direct or mechanistic evidence for or against the claim that TRAPPC3 is likely associated with a Mendelian disease. The study focuses on Moebius syndrome (MBS) and reports whole-exome sequencing results in a cohort of 37 MBS or MBS-like patients, specifically noting the absence of de novo variants in REV3L and PLXND1 and the failure of WES to identify causative mutations for MBS in this cohort. Because TRAPPC3 is not assessed, discussed, or functionally investigated in this manuscript, nothing in the paper can be used to support or refute an association between TRAPPC3 and any Mendelian disorder. Limitations relevant to using this paper as indirect evidence: (1) lack of any mention or analysis of TRAPPC3 means absence of evidence in this paper is not evidence of absence for TRAPPC3-disease associations; (2) the study population is specific to MBS cases, so even if TRAPPC3 were implicated in other Mendelian phenotypes, this cohort would not necessarily reveal that; (3) WES has technical and coverage limits, but since TRAPPC3 was not targeted or reported, those limits are irrelevant here. For assessing the claim about TRAPPC3, one should consult studies or databases that explicitly analyze TRAPPC3 variation, functional assays, segregation in families, or clinical reports linking TRAPPC3 variants to Mendelian phenotypes (e.g., gene-specific case reports, OMIM, ClinVar, or larger exome/genome association/cohort studies).


[Read Paper](https://www.semanticscholar.org/paper/226b561a7e847eca27fae7e2558a187686dd8f75)


### MSeqDR Quick‐Mitome (QM): Combining Phenotype‐Guided Variant Interpretation and Machine Learning Classifiers to Aid Primary Mitochondrial Disease Genetic Diagnosis

**Why Not Relevant**: The provided paper content describes the MSeqDR Quick‑Mitome (QM) platform and an XGBoost‑based Mitochondrial Disease Variant (MDV) classifier for automated variant interpretation in primary mitochondrial diseases. The text does not mention the gene TRAPPC3, report any variants in TRAPPC3, present case reports, segregation data, statistical associations, or functional/mechanistic experiments involving TRAPPC3. Therefore there is no direct evidence (e.g., observed pathogenic TRAPPC3 variants causing Mendelian phenotypes) nor mechanistic evidence (e.g., protein function disruption, cellular/animal phenotypes) in this paper content to support or refute the claim that TRAPPC3 is associated with a Mendelian disease. While the platform described could in principle be used to analyze TRAPPC3 variants, that capability is not evidence of association. In summary, the paper is not relevant to the claim because it contains no data or statements about TRAPPC3, and thus cannot be used to evaluate whether TRAPPC3 is likely associated with a Mendelian disease.


[Read Paper](https://www.semanticscholar.org/paper/d12bb508040b75453f2a34e64f096b0cc1f15b01)


### [Clinical and genetic analysis of three children with Menkes disease due to variants of ATP7A gene].

**Why Not Relevant**: The paper reports genetic findings implicating ATP7A variants in Menkes disease in three children and contains detailed variant validation and pathogenicity interpretation for ATP7A. There is no mention of the gene TRAPPC3, no data (genetic, clinical, or mechanistic) linking TRAPPC3 to the phenotypes studied, and no discussion that would allow inference of TRAPPC3's involvement. Therefore this paper provides no direct or mechanistic evidence for the claim that TRAPPC3 is likely associated with at least one Mendelian disease. Note: absence of TRAPPC3 in this paper is not proof that TRAPPC3 has no Mendelian associations generally — it only means this specific paper is irrelevant to that claim. To evaluate the claim about TRAPPC3 would require literature or database searches (e.g., OMIM, ClinVar, HGMD) or studies that specifically report TRAPPC3 variants segregating with Mendelian phenotypes or provide mechanistic links to disease.


[Read Paper](https://www.semanticscholar.org/paper/2c7af41359206fc3b0a3fce382c29206021a28c3)


### Letter regarding “ SCN9A variant in a family of mixed breed dogs with congenital insensitivity to pain”—Navigating the pathogenicity of candidate gene mutations: Spotlight on paralog Nav genes

**Why Not Relevant**: The submitted paper content does not mention TRAPPC3, nor does it present any data (genetic variant, case report, functional assay, population frequency, segregation, or clinical phenotype) related to TRAPPC3. The entire excerpt concerns a different gene (SCN9A) and a canine sodium-channelopathy (Nav1.7) and discusses variant interpretation in that context. Therefore there is no direct evidence (no TRAPPC3 variants, no patients/cases, no phenotype linkage) and no mechanistic evidence (no discussion of TRAPPC3 molecular function, pathways, or pathogenic mechanisms) in this paper that would support or refute the claim that TRAPPC3 is likely associated with at least one Mendelian disease.

Additional detail supporting this conclusion:
- The text repeatedly references SCN9A/Nav1.7, congenital insensitivity to pain, and variant c.C2761T (p.R921C) in SCN9A; it never names TRAPPC3. That establishes the paper's scope is unrelated to the queried gene.
- The paper's methodological and interpretive content (comparisons to controls, in-silico predictions, ACMG-AMP guidance, need for functional follow-up) are general points about variant assessment. While those points are applicable as guidance for evaluating variants in any gene, they do not constitute evidence that TRAPPC3 is associated with a Mendelian disease.
- No mechanistic pathways described (ion channel pore loops, S4 voltage sensor, arginine substitutions) relate to TRAPPC3 biology (TRAPPC3 is a subunit of the TRAPP complex involved in vesicle trafficking), so mechanistic extrapolation is unsupported by the text.

Limitations and caveats:
- If the intent was to find any indirect relevance, note that the paper's discussion of variant interpretation standards (ACMG-AMP) could inform how one would evaluate TRAPPC3 variants in human or veterinary genetics, but that is procedural guidance only and not evidence of gene-disease association.
- Because no TRAPPC3-specific data are present, no assessment of statistical or practical significance, segregation, functional assay quality, or generalizability for TRAPPC3 can be made from this paper.

Conclusion: Relevance to the claim is effectively zero because the paper does not contain any TRAPPC3-related content or evidence.


[Read Paper](https://www.semanticscholar.org/paper/39dfc4195578b71ad782115a3a552b9b9e2dae03)


### Deciphering the Non-Coding Genome in Autism Spectrum Disorders (ASD): A Study of De Novo and Rare Inherited Variants through Targeted Sequencing in Regulatory Regions

**Why Not Relevant**: The provided paper content does not mention the gene TRAPPC3 anywhere and therefore contains no direct or mechanistic evidence bearing on the claim that "TRAPPC3 is likely associated with at least one Mendelian disease." The paper is focused on non-coding regulatory variation in Autism Spectrum Disorder (ASD). Example sentences from the supplied text that illustrate the paper's scope (verbatim) include: "This study aims to analyze the impact of non-coding mutations within regulatory regions in Autism Spectrum Disorder (ASD)." and "We discovered that 28% of de novo variants and 25% of inherited variants with high regulatory potential were found in patients with negative results from whole-exome sequencing (WES) and microarray analyses, as assessed by Sei." Another sentence emphasizes a mechanistic focus unrelated to TRAPPC3: "Notably, the most important observation in this study is the implication of a global dysregulation of CTCF suggesting a potential mechanistic impact on the chromatin architecture." The paper also states: "We have found that the most high-impact regulatory variants-whether de novo or inherited-are linked to genes not previously associated with ASD." However, no gene names are provided in the excerpt, and TRAPPC3 is not named.

Because (a) TRAPPC3 is not mentioned, (b) the study centers on non-coding regulatory variation in ASD rather than on gene-level Mendelian disease associations, and (c) the types of evidence required to support a Mendelian association (e.g., multiple unrelated families with concordant rare coding or loss-of-function variants, consistent segregations, de novo protein-truncating variants with replication, functional assays showing pathogenic loss/gain of function, or established ClinVar/OMIM entries) are not present in the supplied text, the relevance of this paper to the claim is effectively zero. Note also that even if the paper had implicated a gene via non-coding regulatory variants in ASD, that would not by itself establish a Mendelian disease association; establishing a Mendelian gene-disease relationship typically requires coding-variant segregation and functional/replication evidence not provided here.

If you can provide the full paper or sections that list implicated gene names (e.g., tables of prioritized target genes) or any supplementary lists, I can re-check for TRAPPC3 and re-evaluate. As it stands, there is no evidence in the provided content to support or refute the claim about TRAPPC3.


[Read Paper](https://www.semanticscholar.org/paper/649aeab19738420d26b974bc80eaee323f848ea6)


## Search Queries Used

- TRAPPC3 Bet3 mutation loss of function pathogenic patient biallelic compound heterozygous Mendelian disease

- TRAPPC3 whole exome sequencing trio de novo rare variant candidate gene Mendelian disorder case series

- TRAPPC3 ClinVar OMIM HGMD gene disease association pathogenic variant review clinical annotation

- TRAPPC3 Bet3 trafficking protein particle complex vesicle trafficking Golgi autophagy neuronal development mouse knockout phenotype


## Usage Statistics

- Prompt Tokens: 0

- Completion Tokens: 0

- Total Tokens: 0

- Estimated Cost: $0.0365


## Bibliometric Configuration

- Use Bibliometrics: True

- Author Impact Weight: 0.4

- Citation Impact Weight: 0.4

- Venue Impact Weight: 0.2
