## Supplementary material for "MENDELSEEK: An algorithm that predicts Mendelian Genes and elucidates what makes them special": TRPM8.txt

# Claim: The gene TRPM8 is likely associated with at least one Mendelian disease.

**Status**: processed

**Overall Rating**: -1

**Explanation**:

No relevant papers were found that support or refute this claim.


## Relevant Papers


## Other Reviewed Papers


### The Burden of Candidate Pathogenic Variants for Kidney and Genitourinary Disorders Emerging From Exome Sequencing

**Why Not Relevant**: The provided paper content focuses specifically on identifying and evaluating genes and variants associated with Mendelian kidney and genitourinary disorders. The excerpts show the study's scope (a curated list of 625 kidney/genitourinary genes) and the methods used to identify and annotate variants (OMIM, Orphanet, ClinVar, HGMD, ExAC). Nowhere in the provided text is TRPM8 mentioned, nor are mechanisms or disease associations for TRPM8 described. Because the paper's analyses and conclusions are confined to genes associated with kidney and genitourinary phenotypes, and the supplied excerpt does not include the full gene list, there is no direct or mechanistic evidence in this content that TRPM8 is associated with any Mendelian disease. It remains possible (but unproven from this excerpt) that TRPM8 could be among the 625 genes; however, without the explicit gene list or any mention of TRPM8, this paper cannot be used to support the claim. Therefore the relevance to the claim is minimal.


[Read Paper](https://www.semanticscholar.org/paper/a75e297208c04992568e607d19aa9b9aaebd09d5)


### Novel mutation in the TGFBI gene in a Moroccan family with atypical corneal dystrophy: a case report

**Why Not Relevant**: The paper content provided discusses a novel mutation in the TGFBI gene and its phenotype–genotype correlation in a family. It does not mention the TRPM8 gene, any variants in TRPM8, nor any phenotypes or molecular mechanisms involving TRPM8. Because the claim concerns TRPM8 being likely associated with at least one Mendelian disease, and the provided paper contains no information about TRPM8, it cannot support or refute the claim. Absence of evidence in this paper is not evidence against the claim, but the paper is simply not relevant: it studies a different gene (TGFBI) in a small family, so no direct or mechanistic inference about TRPM8 can be drawn from it.


[Read Paper](https://www.semanticscholar.org/paper/80ec4cae25175caaf4a121cf8f04baa72b130d54)


### De novo single-nucleotide and copy number variation in discordant monozygotic twins reveals disease-related genes

**Why Not Relevant**: The provided paper content does not mention TRPM8, Mendelian disease, monogenic inheritance, or any mechanism connecting TRPM8 to disease. The single supplied sentence concerns a de novo variant in RASD2 and a rare deletion overlapping ARHGAP11B in twins with suspected autism spectrum disorder or with schizotypal personality disorder/schizophrenia. Those are complex neuropsychiatric phenotypes and genes distinct from TRPM8. Based on the supplied text, there is no direct evidence (no observations, variants, segregation, or case reports) implicating TRPM8 in any Mendelian disease, nor is there mechanistic evidence (e.g., functional studies, pathway links) connecting TRPM8 to the reported phenotypes. Caveats: only a short summary/sentence from the paper was provided; it is possible the full paper includes other data or mentions other genes (including TRPM8), but nothing in the provided content addresses TRPM8. Consequently, relevance to the claim is effectively zero given the available text.


[Read Paper](https://www.semanticscholar.org/paper/9d76583342c700aa74f7d738435b29dd9cce5936)


### Clinical and Molecular Findings in a Turkish Family Who Had a (c.879-1G>A) Splicing Variant in PSEN1 Gene with A Rare Condition: The Variant Alzheimer's Disease with Spastic Paraparesis.

**Why Not Relevant**: The paper exclusively addresses PSEN1 variants in variant Alzheimer's disease (varAD) and presents clinical, methodological, and genetic findings related to PSEN1 (including identification of a heterozygous splicing variant c.869-1G>A). There is no mention of TRPM8, no data about TRPM8 variation, and no mechanistic discussion involving TRPM8. Therefore this paper provides no direct or mechanistic evidence supporting or refuting the claim that TRPM8 is associated with at least one Mendelian disease.


[Read Paper](https://www.semanticscholar.org/paper/000d2a196e26bf52a3d2278b63f7a6a2c095d903)


### Activated leukocyte cell adhesion molecule/cluster of differentiation 166 rs10933819 (G>A) variant is associated with familial intracranial aneurysms

**Why Not Relevant**: The paper's content exclusively reports a family-based investigation that identifies and validates a pathogenic ALCAM (CD166) variant associated with intracranial aneurysm in a single pedigree. There is no mention of TRPM8, no reported variants in TRPM8, and no mechanistic discussion that links TRPM8 to the phenotype studied. While the study used whole-exome sequencing and a variant-prioritization pipeline (which in principle could detect TRPM8 variants), the text provided documents only the prioritized ALCAM variant and downstream validation; therefore it contains no direct or mechanistic evidence to support or refute the claim that TRPM8 is likely associated with a Mendelian disease. Absence of evidence in this paper does not disprove the claim, but it means this paper is not relevant for assessing TRPM8's Mendelian disease associations.


[Read Paper](https://www.semanticscholar.org/paper/4f88a77fb34a64d233d7a947efdf3f7770827c97)


### Case Report: A Disease Phenotype of Rett Syndrome and Neurofibromatosis Resulting from A Bilocus Variant Combination

**Why Not Relevant**: The provided paper content contains a single clinical-summary sentence that does not mention TRPM8, does not report any genetic variants, and does not present experimental, segregation, or functional data linking a specific gene to a Mendelian disease. The claim concerns whether TRPM8 is likely associated with at least one Mendelian disease; to evaluate that claim one needs direct evidence such as identification of pathogenic TRPM8 variants in individuals with a consistent phenotype, segregation of such variants in affected families, statistically significant enrichment of TRPM8 variants in cases versus controls, or mechanistic functional data demonstrating how TRPM8 perturbation causes disease. None of these types of data are present in the excerpt. Although the sentence references "multilocus variations" and a possible protein/cellular-level relationship, this is generic and does not implicate TRPM8 specifically, nor does it address Mendelian (single-gene) causation. Therefore the content is not relevant to assessing the claim.


[Read Paper](https://www.semanticscholar.org/paper/993edabc0eeab1343cda0a7dbad248110f7e87ad)


### Rare Variant Genetic Association Study for Transplant-Associated Thrombotic Microangiopathy (TA-TMA) Via Whole Exome Sequencing

**Why Not Relevant**: This paper does not provide evidence for or against the claim about TRPM8. The study focuses on germline rare variants associated with transplant-associated thrombotic microangiopathy (TA-TMA) and examines five a priori pathways (complement regulation, VWF and coagulation, VWF clearance, ADAMTS13 mimics/interacting proteins, and angiopoietin family/endothelial activation). The manuscript text supplied does not mention TRPM8, Mendelian diseases associated with TRPM8, nor any analysis or results concerning TRPM8. Key relevant methodological details show the analysis was pathway-focused: "we defined qualifying variants..." and "we then focused on the exome profiles of 5 a priori selected genetic pathways: complement regulation (17 genes), VWF and coagulation (7 genes), VWF clearance (10 genes), ADAMTS13 mimics or interacting proteins (10 genes), and angiopoietin family and endothelial activation (7 genes)." Because TRPM8 is not listed among these pathways/genes and no TRPM8-specific results or discussion are present, there is no direct evidence (variant associations, case reports, or statistical signals) supporting or refuting TRPM8 as a Mendelian-disease gene in this paper.

Limitations relevant to interpreting the absence of TRPM8 findings: the study used whole-exome sequencing (germline variant detection 40x) which could in principle detect TRPM8 coding variants, but the analysis was intentionally restricted to variants within the five pre-specified pathways and to "qualifying variants" (MAF <1%, certain functional classes, and in-silico/ClinVar pathogenicity filters). Therefore, absence of TRPM8 in the paper does not constitute evidence that TRPM8 is not associated with any Mendelian disease—TRPM8 was simply not analyzed or reported here. In summary, relevance to the claim is nil because TRPM8 is not addressed; the paper neither supports nor contradicts the claim.


[Read Paper](https://www.semanticscholar.org/paper/a28f55f5071fa816fb5c466832a38b225dfaef02)


### Novel mutation in the TGFBI gene in a Moroccan family with atypical corneal dystrophy: a case report

**Why Not Relevant**: The paper content provided exclusively concerns a novel mutation in the TGFBI gene and its association with Thiel-Behnke corneal dystrophy. There is no mention of the gene TRPM8, no experimental data, clinical observations, variant reporting, or biological mechanisms pertaining to TRPM8, nor any cross-reference or analysis that would connect TGFBI findings to TRPM8. Because the content addresses a different gene and disease, it neither supports nor refutes the claim that TRPM8 is associated with a Mendelian disease; it is simply unrelated. Additionally, the provided text is a single summary sentence without methodological detail, statistical evidence, or mechanistic discussion that might hypothetically allow indirect inference. For these reasons the relevance to the claim is effectively zero.


[Read Paper](https://www.semanticscholar.org/paper/f901921467669d44be4f58c9350f9f390c9bf428)


### OA30 Genetic analysis of whole exome sequencing in a cohort of children with refractory JIA reveals genetic risk factors for rare juvenile diseases

**Why Not Relevant**: The paper reports whole-exome sequencing of 99 children with refractory juvenile idiopathic arthritis and identifies deleterious variants in several genes linked to monogenic diseases (notably ADAR, ATP7B, MVK, and HLA-DRB1). Nowhere in the provided text is TRPM8 mentioned, nor are any variants, mechanistic data, or clinical associations involving TRPM8 described. The methods describe variant annotation against PanelApp, arthritis-associated genes, and a paediatric monogenic gene list, but the results and discussion only present findings for other genes. Therefore this paper provides no direct genetic, clinical, or mechanistic evidence that TRPM8 is associated with a Mendelian disease. Caveats: absence of TRPM8 in this paper is weak negative evidence — it could reflect that (a) no qualifying TRPM8 variants were detected in this specific, disease-focused cohort; (b) TRPM8 was not included on the annotation panels used or failed upstream quality/pathogenicity filters; or (c) any TRPM8 variants present did not meet the rarity/pathogenicity thresholds and so were not reported. Additionally, the cohort is small and disease-specific (refractory JIA), so even if TRPM8 were associated with a Mendelian disease, this study would not necessarily be expected to detect it. In sum, relevance to the claim is effectively nil: the paper neither supports nor meaningfully refutes the claim about TRPM8.


[Read Paper](https://www.semanticscholar.org/paper/5ceb90bf884c2beab8d50226ab822bc8d21c2783)


### Identifying pathogenic variants in rare pediatric neurological diseases using exome sequencing

**Why Not Relevant**: The provided paper content does not mention TRPM8 or any gene-level results; it only summarizes a methods-focused analysis: "This study retrospectively analyzed the utility of four annotation tools in identifying 271 pathogenic single nucleotide and small insertion/deletion variants and found that a combination of multiple annotation tools and appropriate evaluation can improve the diagnosis of rare diseases." Because the excerpt contains no gene names, no list of the 271 variants, no diseases linked to specific genes, and no mechanistic information about TRPM8, it provides no direct evidence that TRPM8 is associated with any Mendelian disease. Methodologically, the study appears to evaluate annotation tools and aggregate variant classification performance rather than to perform gene-disease discovery or provide segregation, functional, or clinical phenotype data required to establish a Mendelian association. Even if a TRPM8 variant were among the 271 variants, a single annotated "pathogenic" classification in an annotation-tool benchmarking set would not by itself establish that TRPM8 is causally linked to a Mendelian disease without additional evidence (segregation, functional studies, replicated case series, phenotype correlation, etc.). Therefore, on the available text the paper is irrelevant to the specific claim that "TRPM8 is likely associated with at least one Mendelian disease." The absence of mention of TRPM8 in the summary cannot be taken as evidence against the claim, only that this paper provides no supporting evidence. Additional information required to evaluate the claim would include: (1) whether TRPM8 variants were included among the 271 variants and how they were classified, (2) gene-level and disease-level annotations and phenotype correlations, (3) any functional or segregation data for TRPM8 variants, and (4) methodological details on variant ascertainment and inclusion criteria. Without those details, the study neither supports nor refutes the claim.


[Read Paper](https://www.semanticscholar.org/paper/ab130c4f5e6708f688acbdb584daf59359a913d8)


## Search Queries Used

- TRPM8 mutation Mendelian disease case report familial monogenic pathogenic variant segregation study clinical phenotype

- TRPM8 rare variant exome sequencing whole exome sequencing rare disease cohort de novo mutation ClinVar gnomAD pathogenic variant frequency

- TRPM8 transient receptor potential melastatin 8 ion channelopathy thermosensation cold receptor loss of function gain of function electrophysiology patch clamp knockout mouse phenotype peripheral neuropathy cold allodynia congenital insensitivity to pain

- TRPM8 gene disease association review OMIM ClinGen ClinVar gene curation Mendelian gene list systematic review meta analysis monogenic disease


## Bibliometric Configuration

- Use Bibliometrics: True

- Author Impact Weight: 0.4

- Citation Impact Weight: 0.4

- Venue Impact Weight: 0.2
