## Supplementary material for "MENDELSEEK: An algorithm that predicts Mendelian Genes and elucidates what makes them special": UBQLN1.txt

# Claim: The gene UBQLN1 is likely associated with at least one Mendelian disease.

**Status**: processed

**Overall Rating**: -1

**Explanation**:

No relevant papers were found that support or refute this claim.


## Relevant Papers


## Other Reviewed Papers


### Frequency of carriers for rare recessive Mendelian diseases in a Brazilian cohort of 320 patients

**Why Not Relevant**: This paper does not provide direct or mechanistic evidence regarding the claim that UBQLN1 is likely associated with at least one Mendelian disease. The text supplied describes a study of carrier frequencies for many autosomal recessive disorders in a Brazilian whole-exome sequencing (WES) cohort, but it does not mention UBQLN1, report any variants in UBQLN1, or discuss mechanisms linking UBQLN1 to Mendelian phenotypes. Key points from the supplied content show the paper's scope and why it is not informative for the UBQLN1 claim: it summarizes carrier counts and disease groups (metabolic disorders, epilepsy, intellectual disability, etc.), states the number of variants and genes evaluated (e.g., 351 variants in 278 genes), and discusses estimating aggregate recessive-disease frequencies and carrier-panel sensitivity. None of these statements identify UBQLN1 or present gene-level evidence relevant to UBQLN1 association with Mendelian disease. Therefore the paper cannot be used to support or refute the claim about UBQLN1. Limitations and caveats: (1) If the full paper (beyond the provided excerpt) contains a gene list or supplementary tables, UBQLN1 could be present there; that would change relevance, but no such information was provided here. (2) The paper focuses on carrier-frequency estimates and aggregate burdens rather than establishing gene–disease causality; even if UBQLN1 were listed among genes with rare variants in this cohort, that would be insufficient to demonstrate it is causally associated with a Mendelian disease without segregation, functional, or case-level clinical evidence. (3) WES-based carrier-frequency studies detect variants but typically do not perform the analyses required to establish Mendelian causation (e.g., de novo occurrence, cosegregation, functional validation), so even presence of UBQLN1 variants in the cohort would be weak mechanistic evidence by itself.


[Read Paper](https://www.semanticscholar.org/paper/e50781767181334f904da3f5b9220656083732f8)


### Identifying genomic variant associated with long QT syndrome type 2 in an ecuadorian mestizo individual: a case report

**Why Not Relevant**: The provided paper is a clinical case report of Long QT Syndrome type 2 and discusses pathogenic variation in KCNH2 and variants of uncertain significance in AKAP9 and TTN. The text does not mention the gene UBQLN1 anywhere, presents no genetic, statistical, clinical, or functional data about UBQLN1, and does not discuss pathways or mechanisms involving UBQLN1. Therefore the paper contains no direct evidence supporting or refuting the claim that UBQLN1 is likely associated with at least one Mendelian disease. Important caveats: absence of any mention of UBQLN1 in this single, phenotype-specific case report does not prove UBQLN1 lacks Mendelian disease associations (absence of evidence is not evidence of absence). Also, the sequencing methods and gene sets screened are not fully specified in the excerpt, so UBQLN1 may not have been assessed in this study. Nonetheless, based on the paper content provided, there is no relevant evidence related to the claim.


[Read Paper](https://www.semanticscholar.org/paper/aa65df486c4f1720d86feb42ddc72ed25ab57431)


### Novel guanidinoacetate methyltransferase (GAMT) mutation associated with cerebral creatine deficiency syndrome in a Syrian child: a case report

**Why Not Relevant**: The paper is a case report focused exclusively on GAMT (guanidinoacetate methyltransferase) deficiency and reports a novel GAMT variant. Nowhere in the provided text is UBQLN1 mentioned, nor are there data, analyses, or mechanistic discussions that concern UBQLN1. Therefore this paper provides no direct evidence (genetic association, case reports, linkage, segregation, or variant pathogenicity) nor mechanistic evidence (biochemical pathway, protein function, animal models) relevant to the claim that UBQLN1 is associated with a Mendelian disease. Because the manuscript addresses a different gene and disease, its relevance to the claim is effectively zero. To evaluate the claim about UBQLN1, sources that specifically mention UBQLN1 (e.g., OMIM entries, ClinVar, locus-specific studies, case reports, or functional studies on UBQLN1 variants) should be consulted.


[Read Paper](https://www.semanticscholar.org/paper/bb275b43543e4cfefc6e2d8bc3071c5a400658ed)


### Case report: Preimplantation genetic testing for infantile GM1 gangliosidosis

**Why Not Relevant**: The supplied paper text does not mention UBQLN1 and focuses entirely on GLB1 and GM1 gangliosidosis; it therefore provides no direct or mechanistic evidence relevant to the claim that UBQLN1 is associated with a Mendelian disease. Relevant verbatim sentences from the supplied text that demonstrate the paper's focus include:

Ganglioside-monosialic acid (GM1) gangliosidosis (ICD-10: E75.1; OMIM: 230500, 230600, 230650) is a rare autosomal recessive hereditary disease, lysosomal storage disorder caused by mutations in the GLB1 gene that lead to the absence or insufficiency of β-galactosidase.

The analysis revealed two variants in the heterozygous state: a frameshift variant c.699delG (rs1452318343, ClinVar ID 928700) in exon 6 and a missense variant c.809A>C (rs371546950, ClinVar ID 198727) in exon 8 of the GLB1 gene.

Additional content describes preimplantation genetic testing and embryo genotypes specifically for GLB1 variants (e.g., PGT-M, trophectoderm biopsy, embryo transfer), with no mention of UBQLN1 or mechanistic links to UBQLN1. Thus, based on the provided content, this paper neither supports nor refutes the claim about UBQLN1 because it contains no information about that gene. Limitation: this conclusion is restricted to the text you provided; if UBQLN1 appears elsewhere in the full paper (not included here), that material was not available for review.


[Read Paper](https://www.semanticscholar.org/paper/ce981bc89c565bb4d97baff441baaf8695595448)


### Autosomal Dominant Intellectual Development Disorder-6 (MRD6) Without Seizures Linked to a De Novo Mutation in the grin2b Gene Revealed by Exome Sequencing: A Case Report of a Moroccan Child

**Why Not Relevant**: The provided paper text exclusively discusses GRIN2B (grin2b) and autosomal dominant intellectual development disorder-6 (MRD6), including a specific de novo nonsense variant c.3912C>G (p.Tyr1304Ter) in GRIN2B, its pathogenic classification (PVS1, PS2, PM2, PP3, PP5), absence from gnomAD, and the clinical phenotype. Nowhere in the supplied content is the gene UBQLN1 mentioned, nor are any variants, mechanistic data, segregation, or phenotype correlations relating to UBQLN1 provided. Therefore this paper provides no direct evidence (no reported pathogenic variants, no case reports, no linkage or segregation data) nor mechanistic evidence (no functional studies, no pathway links) that UBQLN1 is associated with any Mendelian disease. Limitations/caveats: absence of evidence in this single paper does not constitute evidence of absence—UBQLN1 could be associated with Mendelian disease in other studies—but the supplied document contains no information relevant to the claim and cannot be used to support it. The paper does provide strong, specific evidence for GRIN2B involvement in MRD6 (de novo pathogenic stop variant, ACMG criteria applied), which further demonstrates that the content is gene-specific and not about UBQLN1.


[Read Paper](https://www.semanticscholar.org/paper/67d756e0cc302e5d46fdda9df34403f1f2de9a5a)


## Search Queries Used

- UBQLN1 mutation pathogenic variant Mendelian disease family segregation case report ClinVar OMIM

- UBQLN1 rare variant whole exome sequencing inherited disease neurodevelopmental intellectual disability epilepsy myopathy burden test

- UBQLN1 loss of function haploinsufficiency ubiquitin proteasome autophagy ER associated degradation protein aggregation neurodegeneration familial disease

- UBQLN1 gnomAD loss of function constraint pLI observed expected LoF ClinVar DECIPHER gene disease association no pathogenic variants


## Bibliometric Configuration

- Use Bibliometrics: True

- Author Impact Weight: 0.4

- Citation Impact Weight: 0.4

- Venue Impact Weight: 0.2
