## Supplementary material for "MENDELSEEK: An algorithm that predicts Mendelian Genes and elucidates what makes them special": UBR4.txt

# Claim: The gene UBR2 is likely associated with at least one Mendelian disease.

**Status**: processed

**Overall Rating**: -1

**Explanation**:

No relevant papers were found that support or refute this claim.


## Relevant Papers


## Other Reviewed Papers


### A Novel Nonsense Variant in GRM1 Causes Autosomal Recessive Spinocerebellar Ataxia 13 in a Consanguineous Pakistani Family

**Why Not Relevant**: This paper provides no direct or mechanistic evidence regarding the claim that UBR2 is likely associated with at least one Mendelian disease. The study focuses exclusively on a novel pathogenic variant in GRM1 as the cause of SCAR13 in a consanguineous Pakistani family; UBR2 is not mentioned anywhere in the provided content. Key verbatim sentences showing the paper's focus include:

"Background and objectives: Autosomal recessive spinocerebellar ataxia-13 (SCAR13) is an ultra-rare disorder characterized by slowly progressive cerebellar ataxia, cognitive deficiencies, and skeletal and oculomotor abnormalities." 

"The objective of this case report is to expand the clinical and molecular spectrum of SCAR13." 

"We investigated a consanguineous Pakistani family with four patients partially presenting with clinical features of SCAR13 using whole exome sequencing." 

"Whole exome sequencing identified a novel pathogenic nonsense homozygous variant, Gly240*, in the gene GRM1 as a cause of SCAR13 that segregates with the recessive disease." 

"We report a novel homozygous nonsense variant in the GRM1 gene in four Pakistani patients presenting with clinical features that partially overlap with the already reported phenotype of SCAR13." 

"So far, only four variants in GRM1 have been reported, in families of Roma, Iranian, and Tunisian origins." 

These excerpts demonstrate that the paper's data, analyses, and conclusions are about GRM1 and SCAR13. Because UBR2 is not studied, mentioned, or discussed, there is neither direct evidence (e.g., variants in UBR2 segregating with disease) nor mechanistic evidence (e.g., functional assays, pathway analyses implicating UBR2) in this paper to support or refute the claim. 

Limitations relevant to assessing UBR2 from this paper: the study is a case report of a single consanguineous family and reports WES findings that implicate GRM1; it does not provide genome-wide variant lists, gene burden analyses, functional assays of other genes, or discussion of UBR2. Therefore no inference about UBR2 can be made from this manuscript.


[Read Paper](https://www.semanticscholar.org/paper/9c91287e0257315d66c1495b68280f77248198bb)


### A pathogenic variant in RAB32 causes autosomal dominant Parkinson’s disease and activates LRRK2 kinase

**Why Not Relevant**: This paper does not provide evidence for or against the claim that UBR2 is likely associated with at least one Mendelian disease. The study focuses on RAB GTPases and specifically identifies RAB32 c.213C>G (Ser71Arg) as co-segregating with autosomal dominant parkinsonism and functionally activating LRRK2 kinase. Nowhere in the provided content is the gene UBR2 mentioned, nor are any data, variants, segregation analyses, functional assays, or mechanistic pathways involving UBR2 reported. Therefore there is no direct evidence (genetic association, segregation, case descriptions) or mechanistic evidence (biochemical interaction, pathway perturbation) in this paper relevant to UBR2. Limitations: absence of mention in this paper does not mean UBR2 is not associated with Mendelian disease in other literature; whole-exome sequencing was performed in the study population, so UBR2 variants may have been observed but not reported because they were not implicated — however, that is speculative and not supported by the text provided. Because the paper contains no UBR2-specific content, it cannot be used to support or refute the claim about UBR2.


[Read Paper](https://www.semanticscholar.org/paper/a36d28e828ede7c87c650af5f5117982b9749e7b)


### Autosomal Dominant Intellectual Development Disorder-6 (MRD6) Without Seizures Linked to a De Novo Mutation in the grin2b Gene Revealed by Exome Sequencing: A Case Report of a Moroccan Child

**Why Not Relevant**: The provided paper content exclusively discusses a GRIN2B (grin2b) nonsense variant (c.3912C>G, p.Tyr1304Ter) causing autosomal dominant intellectual developmental disorder-6 (MRD6, OMIM 613970). There is no mention of the gene UBR2, no data on UBR2 variation, and no mechanistic discussion involving UBR2. Because the text contains no direct or mechanistic evidence linking UBR2 to any Mendelian disease, it cannot be used to support or refute the claim that UBR2 is likely associated with at least one Mendelian disease. (Note: the paper appears to provide strong, but gene-specific, evidence implicating GRIN2B in MRD6—including de novo status, ACMG classification, absence from gnomAD, and predicted deleterious effect of the variant—but none of that bears on UBR2.)


[Read Paper](https://www.semanticscholar.org/paper/67d756e0cc302e5d46fdda9df34403f1f2de9a5a)


### Whole-exome sequencing of pathogenic genes in a family with congenital heart disease: A case report

**Why Not Relevant**: The provided paper content does not mention the gene UBR2 at all. All quoted sentences either describe the study's background, methods (whole-exome sequencing), the number of variants found, or explicitly name the implicated genes (AMER1 and KCNE1). Therefore there is no direct evidence in this paper supporting that UBR2 is associated with any Mendelian disease. Indirectly, because whole-exome sequencing was performed on a small CHD family and UBR2 was not reported among the identified variants, the paper provides only extremely weak negative evidence concerning UBR2 in this particular family; it does not address UBR2 in other cohorts or disease contexts. Additional limitations that prevent this study from informing the claim include: (1) single-family design and small sample size, so absence of UBR2 here cannot exclude UBR2–disease associations elsewhere; (2) WES can miss noncoding, structural, or copy-number variants and the excerpt provides no information on analyses for those variant types; and (3) the paper focuses on congenital heart disease specifically, whereas UBR2 could be associated with other Mendelian phenotypes not investigated here. For these reasons relevance to the claim is negligible.


[Read Paper](https://www.semanticscholar.org/paper/2afe5ddc8ce353c6339a8bb041306ea8d115518f)


### Whole exome sequencing identified a pathogenic mutation of COL2A1 causing Stickler syndrome in a Vietnamese family

**Why Not Relevant**: The paper exclusively reports a family with Stickler syndrome attributable to a pathogenic COL2A1 variant (c.C2818T/p.R940X) and contains no mention, data, analyses, or discussion of the UBR2 gene. Therefore it provides no direct genetic evidence linking UBR2 to any Mendelian disease, nor any mechanistic data implicating UBR2. The absence of UBR2 in this report does not disprove any association between UBR2 and Mendelian disease in general, but this paper cannot be used to support or refute the claim that UBR2 is likely associated with at least one Mendelian disease.


[Read Paper](https://www.semanticscholar.org/paper/d6709617c6a6cdcef9daa16c2b4fb1ef4156dad2)


## Search Queries Used

- UBR2 pathogenic variant human Mendelian disease case report family segregation whole exome sequencing autosomal recessive autosomal dominant

- UBR2 ClinVar OMIM gnomAD loss of function constraint pLI haploinsufficiency pathogenic variants UBR2 disease association

- UBR2 E3 ubiquitin ligase N recognin 2 N end rule pathway functional study knockout mouse spermatogenesis infertility developmental phenotype model organism

- UBR family UBR1 UBR2 UBR3 UBR4 Mendelian gene discovery gene panel undiagnosed disease program candidate gene sequencing congenital disorder


## Bibliometric Configuration

- Use Bibliometrics: True

- Author Impact Weight: 0.4

- Citation Impact Weight: 0.4

- Venue Impact Weight: 0.2
