## Supplementary material for "MENDELSEEK: An algorithm that predicts Mendelian Genes and elucidates what makes them special": UBR5.txt

# Claim: The gene UBR5 is likely associated with at least one Mendelian disease.

**Status**: processed

**Overall Rating**: -1

**Explanation**:

No relevant papers were found that support or refute this claim.


## Relevant Papers


## Other Reviewed Papers


### Simultaneous identification and prioritization of variants in familial, de novo, and somatic genetic disorders with VariantMaster

**Why Not Relevant**: The supplied paper text is primarily methodological: it describes a variant-prioritization tool (VariantMaster), its algorithmic features, and general proof-of-concept applications. Although the authors state they identified causative variants in a rare Mendelian genetic disease, the excerpt does not name any genes or present any gene-specific results. There is no mention of UBR5 anywhere in the provided content, nor are there variant- or gene-level data, statistics, or case descriptions that would support or refute an association between UBR5 and a Mendelian disease. Therefore the paper content does not provide direct evidence for the claim. Mechanistically, the methods described could detect a pathogenic UBR5 variant if present in analyzed samples, but absence of reported findings about UBR5 means no substantive evidence is available here to evaluate the claim.


[Read Paper](https://www.semanticscholar.org/paper/a56e2dc7369c76b03ad5e9309093beb98a159719)


### The Burden of Candidate Pathogenic Variants for Kidney and Genitourinary Disorders Emerging From Exome Sequencing

**Why Not Relevant**: The provided paper content does not contain any direct evidence (e.g., reported pathogenic variants, statistical associations, case reports) or mechanistic discussion (e.g., functional data, molecular pathway linking) concerning the gene UBR5. The study focuses on assembling and annotating a curated list of genes associated with Mendelian kidney and genitourinary disorders (final n = 625) and on methods for filtering and interpreting variants in a control cohort, but the specific gene list and any gene-level results are not included in the text you provided. The excerpted material instead describes methods, cohort characteristics, and where the gene list can be accessed externally. Because no mention of UBR5 occurs in the provided content and no gene-specific findings or mechanistic data for UBR5 are present, this paper excerpt cannot be used to support or refute the claim that "UBR5 is likely associated with at least one Mendelian disease." To evaluate that claim using this study, one would need (a) the full curated gene list (available via the cited laboratory website) to see whether UBR5 was included as a gene associated with a Mendelian phenotype in their curation, and (b) any gene-level variant results or annotations (e.g., ClinVar/HGMD entries, observed pathogenic/truncating variants) for UBR5. Absent those elements in the provided text, relevance to the UBR5 claim is negligible.


[Read Paper](https://www.semanticscholar.org/paper/a75e297208c04992568e607d19aa9b9aaebd09d5)


### A Novel Missense Variant in the Gene PPP2R5D Causes a Rare Neurodevelopmental Disorder with Increased Phenotype

**Why Not Relevant**: The provided paper content exclusively concerns pathogenic variants in the PPP2R5D gene and a PPP2R5D-related neurodevelopmental disorder. There is no mention of the gene UBR5, no reported variants in UBR5, and no mechanistic discussion involving UBR5. Therefore the paper provides no direct genetic evidence (e.g., observed pathogenic variants in UBR5 in affected individuals) nor any mechanistic evidence (e.g., functional studies implicating UBR5 in disease pathways) relevant to the claim that UBR5 is likely associated with at least one Mendelian disease. Because none of the extracted sentences reference UBR5, the relevance to the claim is effectively zero.


[Read Paper](https://www.semanticscholar.org/paper/598838d6722d4bd259802a761ff462c7dcfbc8e1)


### A Novel De Novo Mutation of the DHX30 Gene in a Patient With Neurodevelopmental Disorder, Severe Motor Impairment, and Absent Language (NEDMIAL)

**Why Not Relevant**: The provided paper content exclusively discusses DHX30 and a novel de novo DHX30 mutation (p.Pro796Leu) causing a Mendelian neurodevelopmental disorder (NEDMIAL). There is no mention of UBR5, no data on UBR5 variants, no mechanistic pathways involving UBR5, and no epidemiological or genetic association information about UBR5. Therefore the paper provides no direct evidence supporting or refuting the claim that UBR5 is likely associated with at least one Mendelian disease. If additional parts of the full text (not provided here) mention UBR5, those would need to be supplied to reassess relevance; based solely on the supplied content, relevance is zero.


[Read Paper](https://www.semanticscholar.org/paper/0cc1d790c4d04bc8dabc0ded99cba5398b32ea1b)


### Prenatal Diagnosis of PPP2R1A-Related Neurodevelopmental Disorders Using Whole Exome Sequencing: Clinical Report and Review of Literature

**Why Not Relevant**: The paper exclusively studies PPP2R1A-related neurodevelopmental disorder and reports de novo PPP2R1A variants in two prenatal cases. There is no mention of the gene UBR5, no genetic, epidemiologic, or mechanistic data concerning UBR5, and no discussion of interactions between PPP2R1A and UBR5 that could allow indirect inference. Because the claim concerns whether UBR5 is likely associated with at least one Mendelian disease, and this paper provides only evidence for PPP2R1A, it is not relevant to the claim. The absence of any reference to UBR5 (or to pathways implicating UBR5) means neither direct nor mechanistic support can be extracted from this content.


[Read Paper](https://www.semanticscholar.org/paper/7595f38ac95f91f2d03163020f3687301e6f88d0)


### Seizure remissions and cycles in pediatric focal epilepsy

**Why Not Relevant**: The provided paper excerpt does not mention the gene UBR5, any variants in UBR5, nor any phenotypes or mechanistic data related to UBR5. The text discusses hereditary spastic paraplegias (HSPs) in general and names several established HSP genes (for example, SPG4, SPG11, SPG3A), reports diagnostic yields from exome/genome sequencing, and comments on inheritance patterns and de novo variants, but contains no direct evidence (no case reports, no segregation, no variant pathogenicity data) nor mechanistic evidence (no functional studies, pathways, or protein function descriptions) linking UBR5 to a Mendelian disease. Therefore, based on the provided content, this paper provides no support for the claim that UBR5 is likely associated with at least one Mendelian disease. Important caveat: this assessment is restricted to the exact excerpt you provided. If UBR5 appears elsewhere in the full paper (main text, tables, figures, or supplementary materials) that material was not available for review here; I recommend searching the full text and supplements, and consulting genetic-disease databases (OMIM, ClinVar, HGMD) and the primary literature for UBR5-specific evidence.


[Read Paper](https://www.semanticscholar.org/paper/22c9ae6cdc4251b4d768112d9656a176366618d0)


### Germline mosaicism in a collagen VI‐related myopathy family: A cause of autosomal recessive inheritance

**Why Not Relevant**: This paper contains no direct or mechanistic information about the gene UBR5. The content exclusively discusses germline mosaicism involving COL6A1 (and mentions COL6A2 and COL6A3) as causes of collagen VI-related myopathy and recurrence of Ullrich congenital muscular dystrophy in a family. There is no occurrence of the string "UBR5" or any discussion of UBR5-related biology, variants, functional assays, patient phenotypes linked to UBR5, or any pathway implicating UBR5 in a Mendelian disorder. Therefore this paper provides neither supporting nor contradicting evidence for the claim that UBR5 is likely associated with at least one Mendelian disease. Limitations: absence of UBR5 in this paper does not constitute evidence that UBR5 is not associated with Mendelian disease—only that this particular study is irrelevant to the claim. To evaluate the claim one should examine literature and databases (e.g., OMIM, ClinVar, HGMD, peer-reviewed studies) specifically about UBR5.


[Read Paper](https://www.semanticscholar.org/paper/4572f635bb354974678a75579201d9eedc2c40ea)


### Clingen Coagulation Factor Deficiency Variant Curation Expert Panel: Meeting the Need for Recommendations to Curate Variants in the Coagulation Factor Genes

**Why Not Relevant**: The provided paper content focuses exclusively on coagulation factor genes (notably F8 and F9), the ClinGen Coagulation Factor Deficiency Variant Curation Expert Panel, and development/specification of ACMG/AMP-based variant-interpretation guidelines for coagulation factor deficiencies. The excerpted text makes no mention of the gene UBR5, presents no genetic or mechanistic data about UBR5, and contains no analyses (population, segregation, functional studies) that could be used to support or refute an association between UBR5 and any Mendelian disease. Therefore this paper does not provide direct or mechanistic evidence relevant to the claim that UBR5 is likely associated with at least one Mendelian disease. Caveat: this assessment is based solely on the supplied content; if the full manuscript or supplementary materials (not provided here) contain additional sections that discuss UBR5, those were not available for review and could change the relevance assessment.


[Read Paper](https://www.semanticscholar.org/paper/9ea8d4ce19622ae1c6b0cdfd40848c93199f6280)


### Autosomal Dominant Intellectual Development Disorder-6 (MRD6) Without Seizures Linked to a De Novo Mutation in the grin2b Gene Revealed by Exome Sequencing: A Case Report of a Moroccan Child

**Why Not Relevant**: The provided paper content exclusively describes a Mendelian disorder (MRD6) caused by mutations in the GRIN2B (grin2b) gene and presents a case report identifying a de novo nonsense mutation in GRIN2B. There is no mention of the gene UBR5, no data on UBR5 variants, and no mechanistic discussion linking UBR5 to any disease. Therefore this paper provides no direct or mechanistic evidence that UBR5 is associated with a Mendelian disease. Absence of evidence in this single GRIN2B-focused report does not constitute evidence that UBR5 is not associated with disease; it simply means this paper is irrelevant to the claim about UBR5.


[Read Paper](https://www.semanticscholar.org/paper/67d756e0cc302e5d46fdda9df34403f1f2de9a5a)


### Molecular and phenotypical findings of a novel de novo SYNGAP1 gene variant in an 11-year-old Iranian boy with intellectual disability.

**Why Not Relevant**: The paper exclusively reports on SYNGAP1: it identifies and interprets a de novo nonsense variant in SYNGAP1, describes SYNGAP1’s function (Ras GTPase-activating protein), and discusses phenotypes associated with SYNGAP1 variants. There is no mention of the gene UBR5, no analysis of UBR5 sequence or function, and no data that could be used to support or refute an association between UBR5 and any Mendelian disease. Because UBR5 is a different gene with distinct sequence, function, and disease relationships, evidence about SYNGAP1 cannot be used to infer an association for UBR5. Therefore the paper is not relevant to the claim that UBR5 is likely associated with at least one Mendelian disease.


[Read Paper](https://www.semanticscholar.org/paper/3f4b2eee9f7ad16af6b2e8b2a6417e1d72f20aed)


### #4317 CLINICAL DIVERSITY OF STEROID-RESISTANT NEPHROTIC SYNDROME CAUSED BY TRPC6 MUTATIONS

**Why Not Relevant**: This paper does not provide evidence that UBR5 is associated with any Mendelian disease. UBR5 is not mentioned anywhere in the provided text; the paper focuses on steroid-resistant nephrotic syndrome (SRNS) and reports specific genes (TRPC6, NUP107, PLCE1, COL4A3, COL4A5, etc.) found in this SRNS cohort. Although the authors used whole-exome sequencing or targeted panels (so in principle variants in many genes could be detected), the manuscript neither reports UBR5 variants nor discusses UBR5 mechanistically. Limitations that temper this negative finding: (1) the study is disease-specific (SRNS) and relatively small (n = 39), so failure to observe UBR5 variants in this cohort does not rule out UBR5 involvement in other Mendelian disorders or in other cohorts; (2) targeted panels may not have included UBR5 and exome capture can have incomplete coverage of some genes; (3) stringent filtering and reliance on existing pathogenic annotations (ACMG, ClinVar, Varsome) could miss novel gene–disease relationships. Overall, the paper provides no direct or mechanistic evidence for UBR5 being associated with a Mendelian disease, only indirect (limited) negative evidence from a small, phenotype-focused sequencing study.


[Read Paper](https://www.semanticscholar.org/paper/c2747aaff3d6f0c6e29493cdff73cf48fd818d22)


## Search Queries Used

- UBR5 pathogenic variant Mendelian disease monogenic disorder case report exome sequencing family trio de novo autosomal dominant autosomal recessive ClinVar OMIM

- UBR5 de novo mutation developmental disorder intellectual disability neurodevelopmental disorder exome sequencing rare variant burden gene association Deciphering Developmental Disorders

- UBR5 ubiquitin ligase HECT domain E3 ubiquitin ligase loss of function haploinsufficiency knockout mouse zebrafish developmental phenotype embryonic lethal neurodevelopment

- UBR5 review gene disease association OMIM ClinVar ClinGen gene curation gene disease validity systematic review meta analysis


## Bibliometric Configuration

- Use Bibliometrics: True

- Author Impact Weight: 0.4

- Citation Impact Weight: 0.4

- Venue Impact Weight: 0.2
