## Supplementary material for "MENDELSEEK: An algorithm that predicts Mendelian Genes and elucidates what makes them special": URB2.txt

# Claim: The gene URB2 is likely associated with at least one Mendelian disease.

**Status**: processed

**Overall Rating**: 0

**Explanation**:

Supporting evidence.
The provided paper describes a clear Mendelian disease (North American Indian Childhood Cirrhosis, NAIC) caused by a homozygous missense mutation in CIRH1A (the human homolog of yeast Utp4). CIRH1A is a nucleolar protein that is part of the t-Utp subcomplex of the small subunit (SSU) processome and is required for ribosomal RNA processing and small subunit assembly. The paper shows an in vivo zebrafish model in which loss of cirh1a leads to biliary defects mediated by p53 activation, supporting that defects in SSU processome components can produce a congenital (Mendelian) disease phenotype.

Caveats or contradictory evidence.
The excerpted paper does not mention URB2 at all. All specific genetic and mechanistic data concern CIRH1A/Utp4 and the modeled disease NAIC. No variants, phenotypes, linkage, cohort studies, case reports, functional assays, or animal models linking URB2 to any Mendelian disorder are presented in the provided material. Thus there is no direct evidence in these excerpts that URB2 is associated with any human genetic disease.

Analysis of potential underlying mechanisms, deeper relationships, or second-order implications.
The mechanistic theme—mutations in factors of the SSU processome and other ribosome biogenesis components causing Mendelian 'ribosomopathies'—is well supported by the CIRH1A example in the excerpt. URB2 (a conserved nucleolar/ribosome biogenesis factor in many organisms) occupies a biological role in the same broad pathway (ribosome assembly/SSU processome) as CIRH1A. By analogy, genes encoding proteins with similar roles have in other cases been implicated in Mendelian disorders (e.g., other ribosomopathies). This functional similarity makes it biologically plausible that pathogenic variants in URB2 could cause a Mendelian disease. However, plausibility by analogy is not evidence of actual association: without observed pathogenic variants and phenotype correlation, mechanistic plausibility alone cannot demonstrate that URB2 is a disease gene.

Which rating is most appropriate and why.
Given that the only provided paper documents a different SSU processome gene (CIRH1A) causing a Mendelian disorder and contains no data on URB2, the correct assessment is that there is no evidence in the supplied materials to support the claim that URB2 is associated with at least one Mendelian disease. Although mechanistic analogy makes the claim plausible and worth investigating, the prompt asks for evaluation based on the provided evidence; that evidence does not link URB2 to disease. Therefore the most appropriate rating is "No Evidence."


**Final Reasoning**:

I reviewed the provided excerpt carefully. The paper supplies a robust example of a Mendelian disease caused by mutation in CIRH1A (Utp4), a nucleolar SSU processome component, and discusses ribosomopathy mechanisms. However, URB2 is not mentioned anywhere in the excerpt, and no experimental, clinical, genetic, or epidemiological data connecting URB2 to any Mendelian phenotype are provided. While functional analogy makes URB2 a plausible candidate gene for future study, plausibility is not evidence of association. Because the evaluation must be based on the provided evidence, the correct conclusion is that there is no evidence in the supplied material that URB2 is associated with a Mendelian disease. I therefore reaffirm the rating "No Evidence."


## Relevant Papers


### p53-Mediated Biliary Defects Caused by Knockdown of cirh1a, the Zebrafish Homolog of the Gene Responsible for North American Indian Childhood Cirrhosis

**Authors**: B. Wilkins (H-index: 30), M. Pack (H-index: 35)

**Relevance**: 0.3

**Bibliometric Impact**: 0.40130909090909095


**Excerpts**:

- North American Indian Childhood Cirrhosis (NAIC) is a rare, autosomal recessive, progressive cholestatic disease of infancy affecting the Cree-Ojibway first Nations of Quebec.

- All NAIC patients are homozygous for a missense mutation (R565W) in CIRH1A, the human homolog of the yeast nucleolar protein Utp4.

- Utp4 is part of the t-Utp subcomplex of the small subunit (SSU) processome, a ribonucleoprotein complex required for ribosomal RNA processing and small subunit assembly.

- NAIC has thus been proposed to be a primary ribosomal disorder (ribosomopathy); however, investigation of the pathophysiologic mechanism of this disease has been hindered by lack of an animal model.

- Here, using a morpholino oligonucleotide (MO)-based loss-of-function strategy, we have generated a model of NAIC in the zebrafish, Danio rerio.

- Zebrafish Cirhin shows substantial homology to the human homolog, and cirh1a mRNA is expressed in developing hepatocytes and biliary epithelial cells.

- Injection of two independent MOs directed against cirh1a at the one-cell stage causes defects in canalicular and biliary morphology in 5 dpf larvae.

- In addition, 5 dpf Cirhin-deficient larvae have dose-dependent defects in hepatobiliary function, as assayed by the metabolism of an ingested fluorescent lipid reporter.

- Previous yeast and in vitro studies have shown that defects in ribosome biogenesis cause stabilization and nuclear accumulation of p53, which in turn causes p53-mediated cell cycle arrest and/or apoptosis.

- In accordance with this hypothesis, transcriptional targets of p53 are upregulated in Cirhin-deficient zebrafish embryos, and defects in biliary function seen in Cirhin-deficient larvae are completely abrogated by mutation of tp53.

- Our data provide the first in vivo evidence of a role for Cirhin in biliary development, and support the hypothesis that congenital defects affecting ribosome biogenesis can activate a cellular stress response mediated by p53.


**Explanations**:

- This sentence establishes that NAIC is an autosomal recessive (Mendelian) disease. It provides context that the paper studies a defined Mendelian disorder, which is relevant because the claim concerns whether a particular gene is associated with at least one Mendelian disease. This is contextual (neither direct nor mechanistic evidence about URB2 specifically). Limitation: does not name any gene.

- This is direct evidence in the paper that a single-gene recessive mutation (R565W) in CIRH1A underlies NAIC in affected patients. It therefore demonstrates that CIRH1A (also called the human homolog of yeast Utp4) is causally associated with a Mendelian disease. Strength: explicit genotype–phenotype assertion. Limitation: the excerpt names CIRH1A/UTP4, not URB2; whether URB2 is the same gene or an alias is not stated in the paper content provided, so this is direct evidence only for CIRH1A/UTP4, not explicitly for URB2.

- This sentence explains the molecular function/location of Utp4 (part of the SSU processome involved in ribosomal RNA processing). This is mechanistic evidence that defects in this protein family can plausibly cause disease via disruption of ribosome biogenesis. Strength: links the gene product to a conserved essential cellular process. Limitation: mechanistic, not a direct association to URB2; again depends on whether URB2 encodes a related/identical protein.

- This places NAIC in the category of ribosomopathies and notes an unmet need for animal models. It supports the broader mechanistic hypothesis that impaired ribosome biogenesis can cause Mendelian disease phenotypes. Limitation: conceptual; does not mention URB2.

- Describes generation of an in vivo model to study the gene function tied to the human disease. This is experimental context showing the authors tested gene loss-of-function to recapitulate disease features, strengthening causal inference for the studied gene (CIRH1A). Limitation: again pertains to cirh1a/Cirhin, not URB2 specifically.

- Reports sequence homology and expression of cirh1a (Cirhin) in relevant tissues (developing hepatocytes and biliary epithelial cells). This supports functional relevance of the gene to the disease-affected tissue (mechanistic/contextual). Limitation: applies to cirh1a/Cirhin; the paper text does not say URB2.

- Demonstrates that loss of cirh1a function in zebrafish causes structural biliary defects, providing in vivo functional evidence that perturbation of this gene disrupts a phenotype analogous to the human disease. This is direct functional evidence for cirh1a as disease-relevant (strong experimental support). Limitation: morpholino knockdown is a transient knockdown method with known off-target and developmental toxicity possibilities; evidence pertains to cirh1a, not URB2.

- Shows dose-dependent functional defects in hepatobiliary physiology in Cirhin-deficient larvae, strengthening the phenotype–gene relationship (functional/dose-response evidence). Limitation: same as above—cirh1a-specific and based on zebrafish MO approach; translational relevance depends on conservation and on-target specificity.

- Summarizes prior mechanistic work linking ribosome biogenesis defects to p53 stabilization and cell-cycle/apoptotic responses. This provides a plausible molecular pathway by which defects in nucleolar/SSU processome components (like Utp4/CIRH1A) could produce tissue-specific pathology. Limitation: mechanistic generalization from yeast/in vitro to vertebrate disease requires demonstration; not direct evidence for URB2.

- Provides evidence that in the zebrafish model the p53 pathway is activated by Cirhin deficiency and that tp53 mutation suppresses the biliary phenotype. This is mechanistic in vivo evidence linking ribosome biogenesis disruption to a p53-mediated cellular stress response causing the observed phenotype; it strengthens biological plausibility that mutations in SSU processome components can cause Mendelian disease via p53 activation. Limitation: specific to cirh1a/Cirhin; inference to URB2 depends on gene identity or shared pathway membership.

- The authors conclude they present in vivo evidence that Cirhin is required for biliary development and that congenital defects in ribosome biogenesis can activate p53-mediated stress responses. This is a summary tying the experimental findings to the hypothesis that ribosome biogenesis genes can cause Mendelian disease phenotypes (mechanistic plus disease relevance). Limitation: conclusion concerns Cirhin/CIRH1A; the paper text does not mention URB2 explicitly.


[Read Paper](https://www.semanticscholar.org/paper/33e5e535fbfeb60592fe68c846ebc7a04219bd97)


## Other Reviewed Papers


### Genetic and phenotypic analysis of 101 patients with developmental delay or intellectual disability using whole‐exome sequencing

**Why Not Relevant**: The provided paper content does not mention the gene URB2 anywhere and contains no gene-specific results or variant data that could be mapped to URB2. The text is an abstract-like summary of a whole-exome sequencing study in 101 Japanese patients with developmental delay/intellectual disability and reports overall diagnostic yield, numbers of pathogenic variants, CNVs, and associations with clinical features (autism spectrum disorder and external ear anomalies). There are no verbatim sentences or paragraphs that name URB2, describe variants in URB2, report segregation or functional evidence for URB2, or provide a mechanistic discussion specific to URB2. Therefore this paper provides no direct evidence for or against the claim that URB2 is likely associated with at least one Mendelian disease.

Limitations and caveats relevant to this assessment:
- Absence of mention of URB2 in this paper does not prove that URB2 is not associated with Mendelian disease; it only means this paper does not provide evidence either way. A gene-specific association could be established in other studies.
- The paper is focused on aggregate diagnostic yield and general findings (e.g., number of pathogenic variants, enrichment for certain phenotypes) and does not list per-case gene-level results in the provided excerpt; if the full paper contains a Supplement or tables with gene lists, those were not included here and could potentially mention URB2. Based on the provided content alone, however, no support or mechanistic plausibility for URB2 can be extracted.
- The paper does include a general statement that genetic factors associated with gene expression regulation may be involved ("...which potentially suggests the involvement of genetic factors associated with gene expression regulation."), but this is a broad, non-gene-specific remark and cannot be used as mechanistic evidence for URB2 without additional, gene-specific data linking URB2 to those processes and to disease phenotypes.


[Read Paper](https://www.semanticscholar.org/paper/c4a6b545539bc97495aa04ca6401c2b071178038)


### Diagnostic yield of patients with undiagnosed intellectual disability, global developmental delay and multiples congenital anomalies using karyotype, microarray analysis, whole exome sequencing from Central Brazil

**Why Not Relevant**: The provided paper content does not mention URB2 anywhere and reports no gene-level results or variant calls for URB2. All gene-related statements are general (about exome sequencing and diagnostic yield) and summary-level diagnostic statistics; no specific genes, variants, segregation evidence, functional experiments, or clinical descriptions linked to URB2 are given. Therefore the paper provides no direct evidence that URB2 is associated with any Mendelian disease. It only provides indirect, methodological context (that exome sequencing can discover new genes for ID), which slightly increases the plausibility that a gene could be discovered but does not support URB2 specifically. To evaluate the claim about URB2 one would need excerpts reporting (verbatim) identification of URB2 variants in affected individuals, segregation with disease in families, classification of variants as pathogenic/likely pathogenic, and ideally functional or mechanistic data linking URB2 disruption to disease phenotypes. The absence of any such content here is why relevance to the specific claim is very low.


[Read Paper](https://www.semanticscholar.org/paper/23767daf16f48d9a91172aa0fc624f07b869a9c8)


### Frequency of Variants in Mendelian Alzheimer's Disease Genes within the Alzheimer's Disease Sequencing Project (ADSP)

**Why Not Relevant**: The paper exclusively studies three Mendelian Alzheimer's disease genes (PSEN2, PSEN1, and APP) in the ADSP WGS/WES datasets and reports variant discovery, carrier rates, and age-of-onset associations for those genes. The text provided contains no mention of the gene URB2, no data (genetic, clinical, or mechanistic) about URB2, and no discussion that would link findings in PSEN1/PSEN2/APP to URB2. While the paper demonstrates that loss-of-function and damaging missense variants in specific AD genes can be associated with disease, that general observation does not constitute evidence that URB2 is associated with any Mendelian disease. Establishing URB2 as likely associated with a Mendelian disease would require direct genetic, clinical, or functional data on URB2 (absent here). Therefore the paper provides no relevant supporting or refuting evidence for the claim about URB2.


[Read Paper](https://www.semanticscholar.org/paper/3760efb72c8ec6adaf553ac773087fd2b16aeafc)


### Case report: A novel de novo deletion mutation of DYRK1A is associated with intellectual developmental disorder, autosomal dominant 7

**Why Not Relevant**: This paper provides no direct or mechanistic evidence relevant to the claim that URB2 is likely associated with a Mendelian disease. The entire reported study concerns a pathogenic de novo deletion in the DYRK1A gene causing DYRK1A syndrome, and URB2 is not mentioned anywhere in the provided content. Therefore there are no verbatim sentences or paragraphs in this paper that support or refute an association between URB2 and any Mendelian disease. Specific reasons: (1) gene mismatch — the paper identifies and characterizes a DYRK1A variant (a 9,424 bp de novo heterozygous deletion affecting DYRK1A exons) as causative in the reported patient; this provides evidence only about DYRK1A, not URB2; (2) no mechanistic connection — the paper does not describe biological pathways, interactions, or functional data linking DYRK1A to URB2 or implicating URB2 in the described phenotype; (3) no data on URB2 variation — there are no sequencing results, variant calls, genotype–phenotype correlations, functional assays, or literature citations pertaining to URB2; and (4) study scope — the paper is a single-case exome finding focused on DYRK1A syndrome, so even indirect inference about other genes would be unsupported. Given these points, the paper cannot be used to support or refute the claim about URB2's involvement in Mendelian disease.


[Read Paper](https://www.semanticscholar.org/paper/98eb2bb2850f75cb23ca3954609b1a07001ab461)


### Identification of variants in genes associated with hypertrophic cardiomyopathy in Mexican patients

**Why Not Relevant**: The provided paper content does not mention the gene URB2 or any specific gene names. The single sentence given—"Five variants that were not previously reported in public databases are found in Mexican patients diagnosed with hypertrophic cardiomyopathy, especially those of uncertain significance in patients with symptoms of the condition."—reports the existence of five novel variants in patients with hypertrophic cardiomyopathy (HCM) but gives no information about which genes harbor those variants, whether URB2 is among them, or any segregation, functional, or statistical data linking any particular gene to a Mendelian disease. Therefore the excerpt provides no direct evidence (no mention of URB2, no genotype–phenotype co-segregation, no case-control enrichment, no diagnostic-level classification) and no mechanistic evidence (no functional assays, molecular pathways, or biological role of URB2).

Why this fails to support the claim that "URB2 is likely associated with at least one Mendelian disease":
- Absence of gene name: The text does not list gene(s) affected; URB2 is not named. Without identifying URB2 as one of the genes with novel variants, there is no link. 
- No case-level or segregation data: The excerpt does not describe whether variants segregate with disease in families, nor their inheritance patterns—critical for Mendelian association. 
- No functional/mechanistic data: There are no experimental results (e.g., protein function perturbation, model organism phenotypes, biochemical assays) that would support a mechanistic role for URB2 in disease. 
- Variant classification is uncertain: The excerpt explicitly states the variants are "of uncertain significance," indicating the authors themselves do not claim pathogenicity sufficient to establish a Mendelian disease association.

What would be required in this or another paper to support the claim: explicit identification of URB2 as the gene harboring variants in affected individuals; evidence of segregation with disease in multiple unrelated families or de novo occurrences; statistical excess of rare damaging variants in cases vs controls; experimental evidence that URB2 perturbation causes relevant phenotypes (cellular or organismal), and clinical/diagnostic classification (e.g., pathogenic or likely pathogenic according to ACMG criteria). Without any of those elements in the provided content, the relevance to the claim is effectively zero.

Recommended next steps: search the full paper for any mention of URB2 or specific gene lists; query databases (OMIM, ClinVar, PubMed) for URB2 associations; and if variants in URB2 are reported elsewhere, evaluate segregation, functional data, and variant classifications before inferring a Mendelian disease association.


[Read Paper](https://www.semanticscholar.org/paper/56c7d110dd7a0d150666b83aac1c846edc16f4da)


### Expanding phenotype of MED13-associated syndrome presenting novel de novo missense variant in a patient with multiple congenital anomalies

**Why Not Relevant**: The provided paper content concerns a de novo MED13 p.Pro835Ser missense variant and proposes expanding the MED13-associated phenotype. There is no mention of the gene URB2, no genetic or clinical data concerning URB2, and no mechanistic discussion involving URB2. Therefore the paper content as provided offers no direct or mechanistic evidence supporting the claim that URB2 is likely associated with a Mendelian disease. Note as a caveat that this assessment is based solely on the short excerpt supplied; the full manuscript might discuss other genes, but no such information was provided here.


[Read Paper](https://www.semanticscholar.org/paper/9b2cc0996fdf5fb478e07dbf06be71de50a057ee)


### A novel AUTS2 variant in a patient with global developmental delay and intellectual disability

**Why Not Relevant**: The provided paper content concerns the AUTS2 gene and a novel de novo loss-of-function (c.1606C>T; p.Gln536*) variant causing AUTS2 haploinsufficiency and a neurodevelopmental syndrome. It makes no mention of the gene URB2, presents no data (genetic, clinical, or mechanistic) about URB2, and provides no comparative or functional evidence that would link URB2 to Mendelian disease. Therefore the paper provides no direct evidence that URB2 is associated with any Mendelian disorder. Additionally, the mechanisms discussed (haploinsufficiency of AUTS2 leading to neurodevelopmental phenotypes) are specific to AUTS2 and cannot be extrapolated to URB2 without further data. Because the gene names are different (AUTS2 vs URB2) and no homology, interaction, or co-occurrence data are provided, this paper is not relevant to the claim. If you intended URB2 but meant AUTS2, this paper is directly relevant to AUTS2; otherwise, please provide literature or data mentioning URB2 for a relevant assessment.


[Read Paper](https://www.semanticscholar.org/paper/f5e84e7bb3b42239a53cc7e5c457342cb38c6277)


### The simplest explanation does not have to be preferred: co-occurrence of pathogenic variants in cancer-predisposing genes

**Why Not Relevant**: The provided paper content contains a single sentence describing the study scope (investigating MINAS and cancer phenotypes using literature review and UK 10,000 Genomes Project data, and assessing additive vs synergistic effects of cancer susceptibility genes). It does not mention the gene URB2, nor does it present any gene-specific results, variant annotations, phenotype associations, mechanistic data, or statistical findings that could be mapped to URB2. Without any occurrence of URB2 or related evidence in the excerpt, the paper content cannot be used to support or refute the claim that URB2 is likely associated with at least one Mendelian disease. To evaluate that claim, sources that explicitly mention URB2 (e.g., OMIM, ClinVar, published case reports, or gene-disease association studies) would be required.


[Read Paper](https://www.semanticscholar.org/paper/a1372a664bd2c33a0bda638975a8b26d112457fc)


### Ribosomopatias. Doenças e disfunção do ribosoma

**Why Not Relevant**: The supplied content consists only of section headings (e.g., “Identification of ribosomal abnormalities in human disease”, “Overview of ribosome biogenesis”, “Clinical syndromes”, etc.) and contains no complete sentences, paragraphs, results, or citations mentioning the gene URB2. Because there are no verbatim sentences or paragraphs to extract, I cannot identify any direct evidence (reports of pathogenic URB2 variants, case descriptions, segregation, functional assays) nor any explicit mechanistic statements about URB2 in this document fragment.

Mechanistic relevance: The topics listed (ribosome biogenesis, ribosomopathies such as Diamond–Blackfan anemia, Schwachman–Diamond syndrome, dyskeratosis congenita, cellular consequences of ribosomal haploinsufficiency, p53 role, etc.) are broadly relevant to genes involved in ribosome assembly. If URB2 is itself a ribosome biogenesis factor, those topics would be mechanistically relevant to assessing plausibility that URB2 variants could cause a Mendelian ribosomopathy. However, that inference is speculative because the provided text does not state that URB2 is discussed or implicated.

Limitations preventing assessment:
- No verbatim sentences or paragraphs were supplied to extract or evaluate.
- No mention of URB2, no data, no methods, no results, and no conclusions are present in the fragment.
- Without explicit text linking URB2 to disease (clinical data, genetic evidence, functional studies), the paper fragment cannot provide direct or mechanistic evidence for the claim.

Recommended next steps to evaluate the claim:
1. Search the full text of this review/research article for “URB2” (and synonyms) to see if the gene is discussed and to extract pertinent sentences/paragraphs.
2. If the paper does discuss URB2, extract verbatim sentences/paragraphs that report clinical cases, genetic evidence, functional assays, or mechanistic links to ribosome biogenesis.
3. Independently query databases for disease associations: OMIM, ClinVar, HGMD, PubMed, gnomAD (constraint metrics), and GeneReviews for URB2.
4. Look for primary research articles that report segregation, de novo occurrence, functional impact on ribosome biogenesis, or animal models for URB2.

Because the provided fragment lacks substantive text, I assign low relevance (0.05) rather than zero to reflect that the paper’s topic area is potentially relevant in principle but that no usable evidence is present in the supplied content.


[Read Paper](https://www.semanticscholar.org/paper/370ea1d0786158a8ae3e1f6fc162fd58c388bb98)


## Search Queries Used

- URB2 gene ClinVar OMIM URB2 Mendelian disease case report

- URB2 de novo mutation exome sequencing developmental delay intellectual disability congenital disorder trio

- URB2 ribosome biogenesis nucleolar UTP complex preribosomal processing ribosomopathy

- URB2 loss of function constraint pLI gnomAD intolerance haploinsufficiency rare variant burden Mendelian gene discovery


## Usage Statistics

- Prompt Tokens: 0

- Completion Tokens: 0

- Total Tokens: 0

- Estimated Cost: $0.0360


## Bibliometric Configuration

- Use Bibliometrics: True

- Author Impact Weight: 0.4

- Citation Impact Weight: 0.4

- Venue Impact Weight: 0.2
