## Supplementary material for "MENDELSEEK: An algorithm that predicts Mendelian Genes and elucidates what makes them special": USP4.txt

# Claim: The gene USP4 is likely associated with at least one Mendelian disease.

**Status**: processed

**Overall Rating**: -1

**Explanation**:

No relevant papers were found that support or refute this claim.


## Relevant Papers


## Other Reviewed Papers


### The Burden of Candidate Pathogenic Variants for Kidney and Genitourinary Disorders Emerging From Exome Sequencing

**Why Not Relevant**: The provided paper content does not mention USP4 by name, nor does it present gene-specific results or lists that allow determining whether USP4 was included among genes associated with Mendelian disease. The excerpts describe methods, databases, cohort characteristics, and general variant-interpretation principles relevant to identifying and validating gene-disease associations, but they contain no direct evidence (e.g., reported pathogenic variants, case reports, statistical associations, functional studies) linking USP4 to any Mendelian disorder. Because the required gene-specific information is absent, the paper content cannot be used to support or refute the claim that USP4 is likely associated with at least one Mendelian disease.


[Read Paper](https://www.semanticscholar.org/paper/a75e297208c04992568e607d19aa9b9aaebd09d5)


### Improved Pathogenic Variant Localization using a Hierarchical Model of Sub-regional Intolerance

**Why Not Relevant**: The provided paper content describes a statistical/methodological advance for estimating sub-regional intolerance within genes and demonstrates that these regional intolerance metrics correlate with pathogenic variation in sets of known Mendelian (OMIM) genes. However, the excerpt contains no mention of the specific gene USP4, no gene-level results or lists, and no case/control or variant-level evidence implicating USP4 in any Mendelian disease. Therefore there is no direct evidence in the provided text to support or refute the claim that USP4 is likely associated with at least one Mendelian disease. The paper's results provide only a general mechanistic and methodological rationale that could be applied to USP4 if gene-specific data were presented, but without such data the relevance of this paper to the USP4-specific claim is minimal.


[Read Paper](https://www.semanticscholar.org/paper/b1ed717153e0081117b9755ef170e01d9e0f5a93)


### The RET functional variant c 587T>C is not associated with susceptibility to sporadic medullary thyroid cancer.

**Why Not Relevant**: The provided paper text focuses entirely on medullary thyroid carcinoma (MTC), RET proto-oncogene variants (e.g., G691S, S904S, S836S, IVS1-126G>T, and c*587T>C), and their possible roles in sporadic MTC and Hirschsprung disease. There is no mention of the gene USP4, no data concerning USP4 sequence variation, expression, function, or disease associations, and no mechanistic discussion related to USP4 (for example, deubiquitination activity, substrates, or pathways in which USP4 participates). Key representative sentences from the text that illustrate the paper's focus include: "While germline gain-of-function mutations in the RET (rearranged during transfection) proto-oncogene (OMIM 164761) were shown to be responsible for approximately 92% of MEN 2 cases (2,3), very little is known about the mechanism causing the majority of sporadic tumors." and "a recent report strongly suggests a functional effect of c*587T>C on mRNA stability (13). Those authors demonstrated that c*587C lowers RET mRNA degradation in human neuroblastoma cells, ultimately leading to an increase of transcription product and probably an increase in the amount of total RET protein at the cell membrane (13)." These illustrate the paper's RET-centric genetic and mechanistic analyses and confirm absence of USP4-related content. Therefore this paper provides no direct evidence (no reported USP4 variants, genotype–phenotype correlations, segregation data, or case reports) and no mechanistic evidence (no biochemical, cellular, or animal model data) that would support or refute the claim that "USP4 is likely associated with at least one Mendelian disease." 

Limitations and caveats: absence of any mention of USP4 in this paper does not constitute evidence that USP4 is not associated with a Mendelian disease. It only indicates this particular study is irrelevant to the claim. A proper assessment of the claim requires literature specifically addressing USP4 (genetic studies, ClinVar/OMIM entries, case reports, functional studies, or large-scale sequencing datasets).


[Read Paper](https://www.semanticscholar.org/paper/a98882c7e21a23cbbf7068ba7f4038654a1b1d71)


### The simplest explanation does not have to be preferred: co-occurrence of pathogenic variants in cancer-predisposing genes

**Why Not Relevant**: The provided paper content contains no mention of USP4, no gene-level results, no clinical cases linking USP4 to a Mendelian phenotype, and no mechanistic data about USP4 function. The excerpt only describes the general aims and datasets (MINAS, literature review, UK 10,000 Genomes Project, tumor predisposition phenotypes, and whether multiple cancer susceptibility genes act additively or synergistically). Without any explicit reference to USP4 or presentation of evidence (genetic association, segregation, functional studies, pathogenic variant descriptions), the content cannot be used to support or refute the claim that USP4 is likely associated with at least one Mendelian disease. It is possible the full paper contains relevant information, but based solely on the provided text there is no basis to evaluate USP4; therefore relevance is effectively zero.


[Read Paper](https://www.semanticscholar.org/paper/a1372a664bd2c33a0bda638975a8b26d112457fc)


### Case report: A study on the de novo KMT2D variant of Kabuki syndrome with Goodpasture’s syndrome by whole exome sequencing

**Why Not Relevant**: This paper provides no direct or mechanistic evidence relating to the gene USP4. The study focuses on a single patient with Kabuki syndrome in whom whole-exome sequencing identified a de novo KMT2D missense variant (c.15143G > A, p. R5048H). The text discusses clinical features (facial dysmorphism, skeletal anomalies, growth retardation), pulmonary hemorrhage, a developing anti-GBM antibody positivity and treatment response, and notes KMT2D and KDM6A as the main causative genes for Kabuki syndrome. Nowhere in the provided content is USP4 mentioned, nor are there any data (genetic, expression, functional, or mechanistic) concerning USP4. There are no variants reported in USP4, no experimental assays involving USP4, and no discussion of pathways implicating USP4 in Mendelian disease. Because the paper does not address USP4 at all, it cannot provide evidence for or against the claim that USP4 is likely associated with at least one Mendelian disease. Additional reasons why the paper cannot inform the claim: (1) it is a single-case report centered on KMT2D as the causal gene, so its findings are specific to that gene and phenotype; (2) the methodology described (trio whole-exome sequencing) could in principle detect USP4 variants, but none are reported—absence of reporting implies no relevant findings for USP4 in this study; (3) no mechanistic links (e.g., shared pathways, protein interactions, or functional overlap) between KMT2D/KDM6A and USP4 are presented. In summary, relevance is effectively zero because USP4 is neither mentioned nor investigated in this paper, so the paper provides neither supporting nor refuting evidence regarding USP4 and Mendelian disease association.


[Read Paper](https://www.semanticscholar.org/paper/2c51a28e8f2b66dffe79e534a6cd02098b38f6d0)


### Trio-Based Whole-Exome Sequencing Identifies a De novo EFNB1 Mutation as a Genetic Cause in Female Infant With Brain Anomaly and Developmental Delay

**Why Not Relevant**: The paper exclusively reports a clinical case and genetic analysis implicating EFNB1 (an X-linked gene) in a patient with brain anomaly and global developmental delay. There is no mention of USP4 anywhere in the provided content, no genetic data, no mechanistic discussion, and no analyses that could be interpreted as evidence for or against an association between USP4 and any Mendelian disease. Therefore this paper provides no direct or mechanistic evidence relevant to the claim that USP4 is likely associated with at least one Mendelian disease. The absence of USP4 discussion in multiple explicit statements about the implicated gene and variant means the paper is not relevant to evaluating the claim.


[Read Paper](https://www.semanticscholar.org/paper/939ec66d3caac7b206eab2e4073583eff41bfa48)


### Combined exome and whole transcriptome sequencing identifies a de novo intronic SRCAP variant causing DEHMBA syndrome with severe sleep disorder.

**Why Not Relevant**: The supplied paper content does not mention the gene USP4, does not report any genetic findings linking USP4 to disease, and provides no mechanistic data about USP4 function or pathogenicity. The excerpt only discusses the utility of a combined ES-WTS approach generally and a speculative clinical complication of DEHMBA syndrome. Because there is no information about USP4 in the provided content, the paper (as given) is not relevant to the claim that USP4 is likely associated with at least one Mendelian disease. Note: if additional portions of the paper (methods, results, supplementary data) were available and contained USP4-related data, they were not included here; this assessment is limited to the provided excerpt.


[Read Paper](https://www.semanticscholar.org/paper/22ade7a766f116133e03729baaf0b3b717e6cd33)


### c.1103T>C (p.Ile368Th) de novo Variant in Synaptotagmin 1 (SYT1) Gene is Pathogenic, Leading to an Ultra-Rare Neurodevelopmental Disorder: The Baker-Gordon Syndrome

**Why Not Relevant**: The provided paper content (abstract) focuses exclusively on Baker-Gordon Syndrome caused by mutations in SYT1 and describes a specific SYT1 pathogenic variant and SYT1's mechanistic role in synaptic transmission. There is no mention of USP4, no data on USP4 variants, and no mechanistic discussion linking USP4 to any Mendelian phenotype. Therefore this paper provides no direct evidence nor mechanistic evidence relevant to the claim that USP4 is likely associated with at least one Mendelian disease. Caveat: only the abstract text was provided; while highly unlikely, the full paper or supplementary material could theoretically mention USP4 in another context. Based on the provided content, however, there is no support for the claim. If you want, I can search the broader literature (databases, OMIM, ClinVar, gnomAD, PubMed) for evidence linking USP4 to Mendelian disease.


[Read Paper](https://www.semanticscholar.org/paper/ee225e1b916fbf3fda1d1b5451fe2038e281b8c8)


### Transgenic models to study TGF-[beta] function in hematopoiesis

**Why Not Relevant**: The paper's content focuses on TGF-β signaling (Smad4, type I/II receptors), mouse transgenic/knockout models, megakaryocyte-specific Cre drivers, and the development of a lentiviral siRNA system validated on p53. Nowhere in the provided text is the gene USP4 mentioned, nor are there any human genetic studies, mutation reports, linkage/segregation data, case reports, or functional assays involving USP4 that would support association with a Mendelian disease. Therefore the paper provides no direct evidence and no mechanistic evidence specific to USP4; it is not relevant to the claim that USP4 is likely associated with at least one Mendelian disease.


[Read Paper](https://www.semanticscholar.org/paper/761d3f0c12010d9f6d541782278502bdda78786f)


## Search Queries Used

- USP4 pathogenic variant Mendelian disease ClinVar OMIM germline mutation

- USP4 de novo mutation whole exome sequencing trio rare variant monogenic disorder

- USP4 loss of function knockout mouse phenotype deubiquitinase TGF beta Wnt signaling developmental disorder functional study

- USP family deubiquitinase Mendelian disease review systematic review ClinGen genotype phenotype correlation somatic versus germline phenocopy


## Bibliometric Configuration

- Use Bibliometrics: True

- Author Impact Weight: 0.4

- Citation Impact Weight: 0.4

- Venue Impact Weight: 0.2
