## Supplementary material for "MENDELSEEK: An algorithm that predicts Mendelian Genes and elucidates what makes them special": USP39.txt

# Claim: The gene USP39 is likely associated with at least one Mendelian disease.

**Status**: processed

**Overall Rating**: -1

**Explanation**:

No relevant papers were found that support or refute this claim.


## Relevant Papers


## Other Reviewed Papers


### The Burden of Candidate Pathogenic Variants for Kidney and Genitourinary Disorders Emerging From Exome Sequencing

**Why Not Relevant**: The provided excerpt of the paper does not mention the gene USP39 by name, nor does it present a gene-level list or results that include USP39. All quoted material pertains to study design, curation methods, cohort description, and general variant-interpretation criteria focused on Mendelian kidney and genitourinary disorder genes. Because no statements, data, or analyses about USP39 appear in the supplied text, the paper content as provided offers no direct or mechanistic evidence that USP39 is associated with any Mendelian disease. It remains possible that USP39 could be present in the full curated gene list or in appendices/supplements not included in the supplied content; however, based solely on the text you provided, no relevant evidence for the claim is available.


[Read Paper](https://www.semanticscholar.org/paper/a75e297208c04992568e607d19aa9b9aaebd09d5)


### Genetic Diagnostic Evaluation of Trio-Based Whole Exome Sequencing Among Children With Diagnosed or Suspected Autism Spectrum Disorder

**Why Not Relevant**: This paper does not mention USP39 and provides no data (genetic, clinical, or mechanistic) about USP39. The study reports whole exome sequencing results in 80 simplex families with ASD and lists seven presumed causative genes (CHD8, AFF2, ADNP, POGZ, SHANK3, IL1RAPL1, and PTEN) identified in the cohort; USP39 is not among them. There are no verbatim sentences, paragraphs, figures, or tables in the provided text that reference USP39, its variants, expression, function, or disease associations. Therefore the paper gives neither direct evidence (e.g., pathogenic USP39 variants segregating with disease or de novo LOF variants in affected individuals) nor mechanistic evidence (e.g., functional studies, pathway links, animal models) to support or refute the claim that USP39 is likely associated with at least one Mendelian disease. 

Limitations and caveats relevant to this assessment:
- Absence of evidence in this paper is not evidence of absence: the fact that USP39 is not reported here does not disprove any potential association described elsewhere. 
- The cohort and study focus (ASD patients, CNV-negative simplex families, sample size = 80) constrain what genes are likely to be observed; a gene associated with other phenotypes or very rare Mendelian disorders might simply not appear in this dataset. 

Recommended next steps to evaluate the claim independently (not provided in this paper): search curated disease databases (OMIM, ClinVar, HGMD), population constraint and variant databases (gnomAD), the primary literature for USP39 functional and genetic studies, and model organism phenotypes for USP39 loss-of-function to look for evidence of Mendelian disease association.


[Read Paper](https://www.semanticscholar.org/paper/e83590642a0ed51634cc31d0b28fac2e3ec75069)


### Expanding the phenotype of a recurrent de novo variant in PACS1 causing intellectual disability

**Why Not Relevant**: This paper does not provide any direct or mechanistic evidence related to the claim that USP39 is associated with a Mendelian disease. The manuscript exclusively discusses a recurrent de novo missense variant in PACS1 (c.[607C>T], p.[Arg203Trp]) and the delineation of an intellectual disability syndrome (MRD17) caused by that PACS1 variant. The gene symbol 'USP39' does not appear in the provided text; no sequence variants, functional data, genotype–phenotype correlations, or population/segregation evidence concerning USP39 are presented. Mechanistic evidence given (e.g., PACS1 expression in embryonic brain, proximity of the variant to a CK2-binding motif, and zebrafish antisense experiments suggesting a dominant-negative effect on SOX10-positive neural crest cells) all pertain to PACS1, not USP39. Therefore the paper is irrelevant to the claim about USP39. Limitations: because the paper focuses on PACS1 and MRD17 in a very small number of patients, it cannot be used to infer associations for other genes (including USP39), nor does it provide any negative evidence (e.g., USP39 was screened and found wild type) that would bear on USP39. No verbatim excerpts about USP39 can be provided because the gene is not mentioned.


[Read Paper](https://www.semanticscholar.org/paper/e460d89d850693b4fb1e3df12702be0dfb8d7f5b)


### Diagnostic yield of patients with undiagnosed intellectual disability, global developmental delay and multiples congenital anomalies using karyotype, microarray analysis, whole exome sequencing from Central Brazil

**Why Not Relevant**: The provided paper content does not mention the gene USP39 anywhere, nor does it present gene-level results or mechanistic data for specific genes beyond aggregated diagnostic yields. Because the claim concerns whether USP39 is likely associated with at least one Mendelian disease, evidence would need to include (a) identification of pathogenic or likely pathogenic variants in USP39 in affected individuals with segregation data, (b) functional/mechanistic data linking USP39 disruption to disease biology, or (c) population/replication data supporting a gene-disease relationship. None of these are present in the supplied text. The only indirectly relevant sentence is that the ID gene panel included 1,252 genes; however, the paper excerpt does not state whether USP39 was included in the panel nor report any findings for it. Additionally, the exome/panel cohort for gene discovery was small (19 trios), limiting power to detect or rule out rare Mendelian associations. For these reasons the manuscript excerpt provides no direct or mechanistic evidence to support or refute the claim that USP39 is likely associated with at least one Mendelian disease.


[Read Paper](https://www.semanticscholar.org/paper/23767daf16f48d9a91172aa0fc624f07b869a9c8)


### Precision Medicine in Internal Medicine

**Why Not Relevant**: This paper is a general review of genetic testing strategies, variant interpretation, and the state of gene-disease knowledge; it does not mention USP39, report variants in USP39, present cases with USP39 mutations, provide segregation data, functional studies, or any direct mechanistic evidence linking USP39 to a Mendelian phenotype. Therefore it provides no direct evidence to support or refute the claim that "USP39 is likely associated with at least one Mendelian disease." The included sentences provide background about how many genes are known to cause Mendelian disorders and how exome sequencing can discover new gene-disease relationships, but these are generic and insufficient to evaluate USP39 specifically. To assess the claim about USP39, one would need gene-specific evidence such as: (a) published clinical case reports or case series describing pathogenic or likely pathogenic variants in USP39 that segregate with disease; (b) entries in ClinVar or OMIM implicating USP39; (c) population-constraint metrics (e.g., gnomAD LOEUF/pLI) suggesting intolerance to loss-of-function; (d) functional/biochemical data showing how USP39 variants alter protein function consistent with disease mechanisms; and (e) replication in independent cohorts. None of these are present in the supplied text, so relevance to the USP39 claim is effectively nil.


[Read Paper](https://www.semanticscholar.org/paper/c7199e8a5226f8d9d55777da0e15e91797bb459a)


### Prenatal Diagnosis of PPP2R1A-Related Neurodevelopmental Disorders Using Whole Exome Sequencing: Clinical Report and Review of Literature

**Why Not Relevant**: This paper describes PPP2R1A-related neurodevelopmental disorder and reports prenatal diagnosis of PPP2R1A variants in two fetuses. For example, the paper states: "PPP2R1A-related neurodevelopmental disorder (NDD) is expressed with autosomal dominant inheritance and is typically caused by a pathogenic de novo PPP2R1A mutation." Nowhere in the provided content is the gene USP39 mentioned, discussed, or linked mechanistically to PPP2R1A or to any Mendelian phenotype. Therefore the paper provides no direct evidence (no case reports, variants, segregation, clinical phenotypes, or database annotations) that USP39 is associated with a Mendelian disease. It also provides no mechanistic evidence (protein function, pathway involvement, model organism phenotypes, biochemical effects, or interaction data) that would plausibly connect USP39 to a Mendelian disorder. Because there is no mention or data about USP39, one cannot infer support or contradiction for the claim from this paper. Limitations: the absence of evidence in this paper does not disprove an association between USP39 and Mendelian disease overall; it simply means this paper is irrelevant to that claim. To evaluate the claim about USP39 one should examine sources that specifically report USP39 variants in affected individuals, functional studies of USP39 perturbation, entries in clinical variant databases (e.g., ClinVar, OMIM, DECIPHER), and population constraint metrics (e.g., gnomAD).


[Read Paper](https://www.semanticscholar.org/paper/7595f38ac95f91f2d03163020f3687301e6f88d0)


### Delineation of New Disorders and Phenotypic Expansion of Known Disorders Through Whole Exome Sequencing

**Why Not Relevant**: The provided paper content contains only a single, general sentence about the clinical utility of whole-exome sequencing and does not mention the gene USP39, any variants, phenotypes, associations, experiments, statistical results, or mechanistic pathways. There is therefore no direct evidence (e.g., observed pathogenic variants, segregation, case reports, or cohort analyses) and no mechanistic evidence (e.g., functional studies, protein function, pathway perturbation) related to USP39 or any Mendelian disease. Because the excerpt lacks any gene- or disease-specific information, it is not possible to evaluate the claim from this content.


[Read Paper](https://www.semanticscholar.org/paper/85771f116f3fd8c7d13f8db8d66ee4ada827a124)


### Clingen Coagulation Factor Deficiency Variant Curation Expert Panel: Meeting the Need for Recommendations to Curate Variants in the Coagulation Factor Genes

**Why Not Relevant**: The provided paper content concerns the ClinGen Coagulation Factor Deficiency Variant Curation Expert Panel (CFD-VCEP) and development/specification of ACMG/AMP guidelines for interpreting variants in coagulation factor genes—specifically F8 and F9. The excerpt contains no mention of the gene USP39, no genetic association data, no case reports, no segregation analyses, no functional studies, and no mechanistic discussion pertaining to USP39. Because there is no mention or data regarding USP39, the paper provides no direct evidence (e.g., variants in affected individuals, segregation, population frequency, ClinVar entries) nor mechanistic evidence (e.g., biochemical function perturbation, model organism phenotypes, pathway involvement) supporting the claim that USP39 is associated with at least one Mendelian disease. The paper's focus on F8/F9 and on procedural guideline development means it is not generalizable to support or refute associations for genes outside that scope; statements that the panel will later address "other coagulation factor genes" do not constitute evidence that USP39 is among those genes or linked to disease. Therefore relevance to the USP39–Mendelian disease claim is effectively zero.


[Read Paper](https://www.semanticscholar.org/paper/9ea8d4ce19622ae1c6b0cdfd40848c93199f6280)


### Mapping genetic variations to three-dimensional protein structures to enhance variant interpretation: a proposed framework

**Why Not Relevant**: The paper is a workshop report/review focused on methods and community needs for integrating genetic variant data with three-dimensional protein structure to improve variant interpretation. It does not mention the gene USP39, does not report analyses of USP39 variants, and provides no empirical data (population frequencies, segregation, case reports, functional assays, or statistical associations) linking USP39 to any Mendelian disease. All relevant sentences in the paper are general background or methodological—useful for explaining how one might evaluate whether USP39 is disease-associated, but they do not constitute direct or specific evidence for the claim. To evaluate the claim about USP39, one must search primary gene-specific sources (OMIM, ClinVar, HGMD, PubMed) or perform gene-focused analyses (case series, segregation studies, functional assays, or structural mapping of USP39 variants).


[Read Paper](https://www.semanticscholar.org/paper/ae68aa934d5a5ad3d9009bb8811fa8d5fc4dbb1c)


## Search Queries Used

- USP39 ubiquitin specific peptidase 39 Usp39 mutation variant loss of function Mendelian monogenic rare disease case report familial segregating

- USP39 ClinVar OMIM DECIPHER gnomAD pathogenic variant clinical significance gene disease association review

- USP39 knockout mouse embryonic lethal developmental phenotype spliceosome pre mRNA splicing zebrafish loss of function functional study

- USP39 whole exome sequencing trio de novo rare variant gene discovery intellectual disability congenital anomaly neurodevelopmental disorder cohort study


## Bibliometric Configuration

- Use Bibliometrics: True

- Author Impact Weight: 0.4

- Citation Impact Weight: 0.4

- Venue Impact Weight: 0.2
