## Supplementary material for "MENDELSEEK: An algorithm that predicts Mendelian Genes and elucidates what makes them special": VAMP8.txt

# Claim: The gene VAMP8 is likely associated with at least one Mendelian disease.

**Status**: processed

**Overall Rating**: -1

**Explanation**:

No relevant papers were found that support or refute this claim.


## Relevant Papers


## Other Reviewed Papers


### Evaluation of in silico algorithms for use with ACMG/AMP clinical variant interpretation guidelines

**Why Not Relevant**: The provided paper content contains a single, general sentence about the performance and usage of recently developed predictive algorithms in the clinical genetics literature: "Recently developed algorithms with high predictive power and robust to variables such as disease mechanism, gene constraint, and mode of inheritance are identified, although poorer performing algorithms are more frequently used based on review of the clinical genetics literature (2011–2017)." This sentence does not mention the gene VAMP8, any specific gene, any specific Mendelian disease, variant-level evidence, association statistics, case reports, linkage/segregation data, functional studies, or any other information that would directly or mechanistically link VAMP8 to a Mendelian disorder. While the sentence refers to algorithmic tools that might in general be used to predict gene-disease associations, it provides no output from those algorithms, no examples of genes they identify, and no direct or mechanistic evidence concerning VAMP8. Therefore the content cannot be used to support or refute the claim that "The gene VAMP8 is likely associated with at least one Mendelian disease." Any inference that VAMP8 is implicated would require additional, gene-specific data (e.g., variant pathogenicity predictions for VAMP8, genetic co-segregation, population frequency and constraint metrics for VAMP8, functional data), none of which are present. Limitations: the sentence is high-level and methodological (about algorithm performance and usage), temporally limited to a literature review period (2011–2017), and contains no gene- or disease-specific findings; thus it is not relevant evidence for the claim.


[Read Paper](https://www.semanticscholar.org/paper/c2c29313dd550e729c7694b83f63622ebf4334e7)


### Genomic Landscape and Mutational Spectrum of ADAMTS Family Genes in Mendelian Disorders Based on Gene Evidence Review for Variant Interpretation

**Why Not Relevant**: The provided paper focuses exclusively on the ADAMTS family of genes and their variant landscape and Mendelian disease associations. It does not mention VAMP8, provide any data on VAMP8 variants, nor describe mechanisms linking VAMP8 to Mendelian disease. Therefore it provides no direct or mechanistic evidence supporting or refuting the claim that VAMP8 is likely associated with at least one Mendelian disease. Note that absence of mention in this ADAMTS-focused paper is not evidence that VAMP8 is or is not disease-associated; it simply falls outside the paper's scope. To evaluate the claim about VAMP8, one should examine primary genetic-disease resources (ClinVar, HGMD), gene-specific literature on VAMP8 (also known as vesicle-associated membrane protein 8), and functional/mechanistic studies relating VAMP8 perturbation to Mendelian phenotypes.


[Read Paper](https://www.semanticscholar.org/paper/523306c9d896a6a598eabd7692caaaee427426fc)


### 1453-P: Adaption of the ACMG/AMP Variant Interpretation Guidelines for GCK, HNF1A, HNF4A-MODY: Recommendations from the ClinGen Monogenic Diabetes Expert Panel

**Why Not Relevant**: Relevance is very low because the paper focuses specifically on variant interpretation for monogenic diabetes and on three genes (GCK, HNF1A, HNF4A). There is no mention of VAMP8, no reported variants, phenotypes, functional data, or mechanistic discussion relating to VAMP8, nor any claim about VAMP8 being associated with a Mendelian disease. Therefore the paper provides neither direct evidence (no data or associations reported for VAMP8) nor mechanistic evidence (no functional or pathway discussion involving VAMP8) to support or refute the claim. Note that absence of VAMP8 in this paper does not disprove the claim generally; it only means this paper is not a source of evidence about VAMP8.


[Read Paper](https://www.semanticscholar.org/paper/ad1eb507390d767b7a0ee240e7f8201d33bdabd7)


### Autosomal Dominant Intellectual Development Disorder-6 (MRD6) Without Seizures Linked to a De Novo Mutation in the grin2b Gene Revealed by Exome Sequencing: A Case Report of a Moroccan Child

**Why Not Relevant**: The paper exclusively reports a GRIN2B (grin2b) pathogenic variant causing autosomal dominant intellectual development disorder-6 (MRD6) in a single patient and discusses GRIN2B's role in neurodevelopment. The provided content contains no mention of VAMP8, no experiments or analyses involving VAMP8, and no mechanistic pathways that implicate VAMP8. Therefore this paper provides no direct evidence that VAMP8 is associated with any Mendelian disease, nor does it provide mechanistic evidence that would make such an association more or less plausible. Absence of VAMP8 in this paper does not disprove any association elsewhere, but it means this paper is not relevant to the claim.


[Read Paper](https://www.semanticscholar.org/paper/67d756e0cc302e5d46fdda9df34403f1f2de9a5a)


### #5782 AN EXCEPTIONAL CASE OF A PATIENT WITH TWO RARE GENETIC VARIANTS OF AUTOSOMAL DOMINANT POLYCYSTIC KIDNEY DISEASE AND ALPORT SYNDROME

**Why Not Relevant**: The paper does not mention VAMP8 anywhere in the provided content. All genetic and mechanistic discussion concerns COL4A3, PKD1, PKD2, and GANAB in the context of Alport syndrome and ADPKD. Therefore this paper provides no direct evidence that VAMP8 is associated with any Mendelian disease, nor does it provide mechanistic data involving VAMP8. Absence of mention in this single case report is not proof that VAMP8 is uninvolved in any Mendelian disease overall, but it means this paper neither supports nor refutes the claim about VAMP8. Additional literature specifically examining VAMP8 would be required to assess the claim.


[Read Paper](https://www.semanticscholar.org/paper/0337e73775f38924bbc68438e6e09327983393d6)


### Prevalence of the cancer-associated germline variants in Russian adults and long-living individuals: using the ACMG recommendations and computational interpreters for pathogenicity assessment

**Why Not Relevant**: The provided paper content does not mention the gene VAMP8 anywhere, nor does it provide a list of the 28 ACMG-recommended cancer-associated genes that were analyzed. The study is focused on cancer-associated germline variants in a predefined set of genes and reports aggregate and methodological results; without gene-level statements including VAMP8, there is no direct evidence that VAMP8 is associated with any Mendelian disease. Mechanistic evidence is also absent — the excerpts contain no molecular, functional, or disease-mechanism discussion related to VAMP8. A small nonzero relevance value is retained only because the paper involves germline variant analysis and pathogenicity assessment methods, which are generally relevant to assessing gene–disease associations; however, based on the provided content alone, there is effectively no usable evidence for or against the specific claim about VAMP8. To evaluate the claim properly would require either (a) the full list of genes analyzed (to confirm whether VAMP8 was included) and any gene-level variant findings, or (b) other studies that explicitly discuss VAMP8 and its association with Mendelian phenotypes.


[Read Paper](https://www.semanticscholar.org/paper/3d0d4b23c56fdedc897c23bf34ad7662df386963)


### #1049 Genetic characteristics of autosomal dominant polycystic kidney disease in a Greek population

**Why Not Relevant**: This paper provides no direct evidence for or against the claim that VAMP8 is likely associated with at least one Mendelian disease. VAMP8 is not mentioned anywhere in the provided text. The study is focused on genetic causes of autosomal dominant polycystic kidney disease (ADPKD) and reports findings predominantly in PKD1 and PKD2, with only three non-PKD gene findings (SEC61B and SLC3A1). Although the authors performed targeted sequencing of over 600 clinically relevant genes and in some unresolved cases used WES/WGS, the paper does not (a) list whether VAMP8 was included in the panel, (b) report any VAMP8 variants, or (c) discuss VAMP8 in mechanistic terms. Therefore the manuscript offers only weak, indirect negative information (i.e., no VAMP8 association found/reported for ADPKD in this cohort), which is insufficient to support the claim that VAMP8 is likely associated with a Mendelian disease. Limitations that prevent drawing a stronger conclusion include the study’s disease-specific focus (ADPKD), relatively small cohort size for detecting rare gene-disease associations beyond ADPKD, incomplete reporting of the full gene panel content, and the fact that absence of a reported VAMP8 finding does not exclude VAMP8 associations with other phenotypes or in other populations.


[Read Paper](https://www.semanticscholar.org/paper/809bc9272642701e2600d56210354f9dc489b442)


## Search Queries Used

- VAMP8 mutation Mendelian disease case report family segregation monogenic autosomal recessive autosomal dominant

- VAMP8 pathogenic variants ClinVar OMIM ACMG variant interpretation pathogenicity review

- VAMP8 SNARE vesicle fusion exocytosis membrane trafficking platelet secretion immune secretion neuronal secretion pancreatic beta cell functional study human mutation

- VAMP8 whole exome sequencing trio undiagnosed rare disease candidate gene rare variant burden study knockout mouse phenotype loss of function zebrafish Caenorhabditis elegans model organism


## Bibliometric Configuration

- Use Bibliometrics: True

- Author Impact Weight: 0.4

- Citation Impact Weight: 0.4

- Venue Impact Weight: 0.2
