## Supplementary material for "MENDELSEEK: An algorithm that predicts Mendelian Genes and elucidates what makes them special": VPS54.txt

# Claim: The gene VPS54 is likely associated with at least one Mendelian disease.

**Status**: processed

**Overall Rating**: 3

**Explanation**:

Supporting evidence.
The provided paper shows that Vps54 is an essential subunit of the GARP complex with conserved cellular roles in endosome-to–TGN trafficking, and that genetic perturbation of Vps54 produces clear, reproducible monogenic phenotypes in multiple model organisms. Specifically, the excerpt notes that reduction of Vps54 in the wobbler mouse causes neurodegeneration (the wobbler mouse is a well-characterized single-gene model of motor neuron disease), and that both null mutants and motor-neuron-specific knockdown of the Vps54 ortholog in Drosophila cause neuromuscular junction (NMJ) overgrowth and postsynaptic composition defects. The Drosophila work further ties the phenotypes to a defined mechanism — altered Rab7-mediated endosomal trafficking and mislocalization of TGN SNARE components — strengthening the biological plausibility that loss or dysfunction of VPS54 can cause a cell-autonomous, deleterious phenotype in neurons. Together, these data provide strong cross-species functional evidence that VPS54 disruption can produce a Mendelian-like neurodegenerative phenotype in animals.

Caveats or contradictory evidence.
All evidence provided is from model organisms (mouse and Drosophila); no human genetic data are provided in the excerpts. A pathogenic effect in mouse or fly does not by itself prove that mutations in the human VPS54 gene cause a Mendelian disease in people: interspecies differences in redundancy, expression, or compensatory mechanisms can alter whether a variant produces disease in humans. The excerpt does not report any human segregation, de novo occurrence, allele frequency, or clinical genetic evidence tying VPS54 variants to a heritable human disorder. Absent such human genetic data, the association of VPS54 with a human Mendelian condition remains a plausible hypothesis but unconfirmed.

Analysis of mechanisms and implications.
The mechanistic findings are coherent: VPS54 is part of the GARP tethering complex, and its loss perturbs trafficking between endosomes and the trans-Golgi network, alters SNARE localization (Syntaxin-16), and disrupts Rab7-dependent endosomal pathways. Neurons — particularly motor neurons with long axons — are known to be highly sensitive to defects in intracellular trafficking, so a single-gene lesion that impairs these processes plausibly produces neurodegeneration. The cross-species consistency (mouse and fly) increases confidence that VPS54 has a conserved, dosage-sensitive role in neuronal health. Second-order implications are that VPS54 is a strong candidate gene for human motor neuron or other neurodevelopmental/neurodegenerative Mendelian disorders, meriting focused human genetic screening (segregation studies, burden testing, evaluation of rare loss-of-function variants) and functional follow-up of any human variants identified.

Which rating is most appropriate.
Given the strong functional and genetic evidence from animal models that loss of Vps54 causes monogenic neurodegenerative phenotypes, VPS54 is plausibly a Mendelian disease gene. However, the provided evidence lacks any direct human genetic confirmation. Because the claim as stated — "The gene VPS54 is likely associated with at least one Mendelian disease" — is most naturally interpreted as asserting an association with a human Mendelian disorder, the available evidence is supportive but indirect. Therefore the most accurate characterization is "Mixed Evidence": animal-model data strongly support biological plausibility and candidacy, but without human genetic data the claim that VPS54 is associated with a human Mendelian disease is unproven.


**Final Reasoning**:

The excerpts present robust functional genetics from mouse and Drosophila showing that disruption of Vps54 produces monogenic neurodegenerative phenotypes and define a plausible cellular mechanism (defective endosome-to–TGN trafficking, Rab7-dependent pathways). This is strong evidence that VPS54 loss can cause Mendelian-like disease phenotypes in animals. However, the claim as typically interpreted refers to human Mendelian disease, and no human genetic evidence (segregation, pathogenic variants, case reports) was provided. Thus the evidence supports VPS54 as a strong candidate Mendelian gene but does not confirm a human disease association. Given the weight of supportive cross-species functional data but absence of direct human genetic confirmation, the balanced judgement is "Mixed Evidence."


## Relevant Papers


### Vps54 regulates Drosophila neuromuscular junction development and controls postsynaptic density composition via a Rab7-dependent mechanism

**Authors**: Prajal H. Patel (H-index: 6), S. Barbee (H-index: 11)

**Relevance**: 0.7

**Bibliometric Impact**: 0.2081


**Excerpts**:

- Vps54 is a subunit of the Golgi-associated retrograde protein (GARP) complex, which is involved in tethering endosome-derived vesicles to the trans-Golgi network (TGN).

- In the wobbler mouse, a model for human motor neuron (MN) disease, reduction in the levels of Vps54 causes neurodegeneration.

- To better understand the role of Vps54 in MNs, we have disrupted expression of the Vps54 ortholog in Drosophila and examined the impact on the larval neuromuscular junction (NMJ).

- Here, we show that both null mutants and MN-specific knockdown of Vps54 leads to NMJ overgrowth.

- Reduction of Vps54 partially disrupts localization of the t-SNARE, Syntaxin-16, to the TGN but has no impact on endosomal pools.

- Presynaptic knockdown of Vps54 in MNs combined with overexpression of the small GTPases Rab5, Rab7, or Rab11 suppresses the Vps54 NMJ phenotype.

- Conversely, knockdown of Vps54 combined with overexpression of dominant negative Rab7 causes axonal and behavioral abnormalities including a decrease in postysynaptic Dlg and GluRIIB levels without any effect on GluRIIA.

- Taken together, these data suggest that Vps54 controls larval MN axon development and postsynaptic density composition by modulating Rab7-mediated endosomal trafficking.


**Explanations**:

- This sentence defines Vps54's molecular role in the GARP complex and places it in a conserved intracellular trafficking pathway (TGN tethering of endosome-derived vesicles). Relation to claim: mechanistic. Strength: establishes a biologically plausible role for Vps54 such that perturbation could cause cellular dysfunction relevant to disease. Limitation: descriptive molecular role only — no disease linkage or phenotype described here.

- This sentence reports that reduction of Vps54 in the wobbler mouse causes neurodegeneration and frames that mouse as a model for human motor neuron disease. Relation to claim: indirect/direct animal-model evidence supporting that loss or reduction of Vps54 function can produce a Mendelian-like disease phenotype in an organism. Strength: provides in vivo evidence that Vps54 perturbation leads to a clear neurodegenerative phenotype. Limitation: this is a mouse model; it does not by itself demonstrate that VPS54 mutations cause a Mendelian human disease (no human genetic evidence presented). Also, the phrasing indicates a reduction in levels rather than a specific Mendelian variant with segregation data.

- This sentence explains the experimental approach in Drosophila to probe Vps54 function in motor neurons. Relation to claim: contextual/mechanistic. Strength: shows the authors tested conserved function across species, which supports plausibility that VPS54 dysfunction can be pathogenic. Limitation: again, model organism experiment — not direct human genetic evidence; ortholog function may not fully recapitulate human disease mechanisms.

- This reports that loss-of-function (null mutants) and MN-specific knockdown produce a quantifiable synaptic phenotype (NMJ overgrowth) in Drosophila. Relation to claim: mechanistic/phenotype evidence. Strength: demonstrates that Vps54 loss causes clear developmental/neurobiological defects in vivo, strengthening the argument that VPS54 is dosage-sensitive and disruption can lead to disease-relevant phenotypes. Limitation: Drosophila phenotype (NMJ overgrowth) is not a Mendelian disease diagnosis in humans; translation to human pathology is inferential.

- This sentence identifies a molecular consequence of Vps54 reduction: partial mislocalization of Syntaxin-16 to the TGN, while endosomal pools are unaffected. Relation to claim: mechanistic. Strength: provides a specific trafficking defect linking Vps54 loss to disrupted Golgi/TGN SNARE localization, offering a cellular mechanism that could underlie neurodegeneration. Limitation: effect is 'partial' and limited to Syntaxin-16 localization; this is mechanistic insight but does not demonstrate causality for a human Mendelian phenotype.

- This reports genetic interaction data showing that overexpression of Rab5, Rab7, or Rab11 can suppress the NMJ phenotype caused by presynaptic Vps54 knockdown. Relation to claim: mechanistic. Strength: supports a role for Vps54 in endosomal/GTPase-mediated trafficking pathways and shows that modulation of these pathways can modify the phenotype, strengthening pathway-level plausibility for disease causation. Limitation: suppression in fly presynaptic context is experimental and does not equate to human pathogenicity or to Mendelian inheritance evidence.

- This sentence describes that combining Vps54 knockdown with dominant negative Rab7 overexpression causes axonal and behavioral abnormalities and specific changes in postsynaptic protein levels (decrease in Dlg and GluRIIB; no effect on GluRIIA). Relation to claim: mechanistic and phenotype evidence. Strength: links Vps54 deficiency with specific neuronal and behavioral phenotypes and altered postsynaptic composition, reinforcing biological plausibility that VPS54 dysfunction can produce neurologic disease phenotypes. Limitation: again, evidence is from a model organism and an engineered genetic interaction; it does not provide human Mendelian variant evidence, and the precise behavioral/biochemical changes in larvae may not map directly to human disease phenotypes.

- This concluding sentence summarizes that Vps54 controls axon development and postsynaptic composition via Rab7-mediated endosomal trafficking. Relation to claim: mechanistic synthesis. Strength: ties together molecular and phenotypic data into a coherent pathway (Vps54 → Rab7-mediated trafficking → axonal/postsynaptic defects) that plausibly explains how VPS54 perturbation could cause neurodegenerative or neuromuscular disease. Limitation: the conclusion is based on invertebrate and mouse model data; it remains inferential with regard to human Mendelian disease causation because no human genetic/segregation data are presented in the provided content.


[Read Paper](https://www.semanticscholar.org/paper/d97c6d773d85383708eb492f538dee077b222f61)


## Other Reviewed Papers


### Exome sequencing analysis on products of conception: a cohort study to evaluate clinical utility and genetic etiology for pregnancy loss

**Why Not Relevant**: The provided paper content contains only a single general sentence about the clinical utility of exome sequencing for identifying monogenic causes of pregnancy loss. It does not mention VPS54, any specific gene, any variant, or any Mendelian condition. There is no direct evidence (no reported VPS54 variants, case reports, or statistical associations) and no mechanistic information (no discussion of VPS54 protein function, pathways, or pathogenic mechanisms) relevant to the claim. Because the content contains no gene-specific data, its relevance to the claim that 'VPS54 is likely associated with at least one Mendelian disease' is effectively zero.


[Read Paper](https://www.semanticscholar.org/paper/d3ea672fb689541e4ecbd007c0e7c5182f1b2cd9)


### Whole-exome sequencing analysis on products of conception: A cohort study to evaluate clinical utility and genetic etiology for pregnancy loss

**Why Not Relevant**: The provided paper content does not mention the gene VPS54, nor does it provide a list of the specific genes or variant-level findings from the WES cohort. All quoted material is high-level (purpose, methods, aggregated results, and broad categories of affected genes) and therefore only demonstrates that WES can detect Mendelian disease variants in pregnancy loss samples. Because the abstract does not name VPS54, present variants in VPS54, describe segregation, functional assays, or published disease associations involving VPS54, there is no direct or mechanistic evidence in this text to support or refute the claim that "VPS54 is likely associated with at least one Mendelian disease." It is possible that VPS54 variants were present among the detected findings in the full paper or supplement, but that information is not included in the provided content. For these reasons the relevance score is very low (<0.1): the content is methodologically and conceptually relevant to how one would discover gene–disease associations, but contains no specific evidence about VPS54 itself.


[Read Paper](https://www.semanticscholar.org/paper/3da1de97a0b0630a9c688c74e4535a1767739a38)


### Clinical application of next generation sequencing for Mendelian disease diagnosis in the Iranian population

**Why Not Relevant**: The provided content is a single, high-level summary sentence stating that the paper presents a large mutational spectrum of Mendelian disease variants in the Iranian population. It does not mention VPS54, any variant identifiers, phenotypes, statistical associations, functional/biochemical data, or mechanistic explanations linking VPS54 to disease. Therefore, based solely on the text supplied, the paper provides no direct or mechanistic evidence for or against the claim that VPS54 is associated with at least one Mendelian disease. To evaluate the claim from this study would require the full paper and supplementary materials (gene/variant-level tables, methods, variant pathogenicity assertions, and any functional follow-up). As an alternative, one could query locus-specific and clinical databases (OMIM, ClinVar, HGMD) or broader literature for VPS54 disease associations if the full paper cannot be provided.


[Read Paper](https://www.semanticscholar.org/paper/7dc010a5154823dced32a42f93c9bc87c614b498)


### Complex PKD1 Genetics in Early-Onset Cystic Kidney Disease

**Why Not Relevant**: The provided paper content exclusively addresses PKD1 variants, their phase (in trans), pathogenicity interpretation (including missense VUS vs truncating variants), potential additional kidney-related genes (GANAB, DNAJB11, HNF1B, IFT140, PKHD1, ALG8, ALG9, DZIP1L, COL4A3/4/5, etc.), and the clinical/genetic context of cystic kidney disease. There is no mention of VPS54 anywhere in the text. Consequently, the paper provides no direct genetic, clinical, or mechanistic evidence linking VPS54 to any Mendelian disease. The mechanistic discussions relate to polycystin function and PKD1 biology and do not generalize to provide evidence about VPS54. Because the study did not evaluate VPS54 (and did not list it among considered genes), we cannot infer from this paper that VPS54 is associated with a Mendelian disorder; there is simply no relevant data or discussion to support or refute the claim.


[Read Paper](https://www.semanticscholar.org/paper/93aa5634e2918da0adf9d4d1273325fd11dae0af)


### Assessment of disease-associated missense variants in RYR2 on transcript splicing

**Why Not Relevant**: The provided paper content exclusively concerns RYR2 variants and their potential to alter splicing in catecholaminergic polymorphic ventricular tachycardia (CPVT). The text discusses RYR2, CALM1, CASQ2, TRDN, TECRL and methods for predicting and assaying spliceogenic effects in RYR2 missense variants. Nowhere in the provided content is the gene VPS54 mentioned, nor are there any data, analyses, mechanistic discussions, or conclusions that involve VPS54. Therefore the paper provides neither direct evidence (e.g., genetic association, case reports, segregation data, pathogenic variants) nor mechanistic evidence (e.g., biochemical function, pathway linking VPS54 to disease, loss/gain-of-function models) relevant to the claim that VPS54 is likely associated with at least one Mendelian disease. Absence of evidence in this paper does not argue for or against VPS54’s disease association generally; it only means this specific paper gives no information on VPS54. To evaluate the claim about VPS54, literature and database searches (OMIM, ClinVar, HGMD, PubMed, gnomAD constraint metrics, functional studies) specific to VPS54 would be required.


[Read Paper](https://www.semanticscholar.org/paper/ebe071e4b0c26bb341f5b5ec48d725ef296118da)


## Search Queries Used

- VPS54 pathogenic variant human monogenic disease case report autosomal recessive autosomal dominant missense nonsense frameshift loss of function clinical genetics

- VPS54 exome sequencing rare variant burden diagnostic exome trios cohort study Mendelian disorder gene discovery clinical sequencing

- VPS54 wobbler mouse motor neuron degeneration GARP complex vesicle trafficking Golgi retrograde transport model organism functional study neurodegeneration

- VPS54 OMIM ClinVar review GARP complex VPS52 VPS53 human disease review gene curation gene constraint gnomAD loss of function intolerance


## Usage Statistics

- Prompt Tokens: 0

- Completion Tokens: 0

- Total Tokens: 0

- Estimated Cost: $0.0270


## Bibliometric Configuration

- Use Bibliometrics: True

- Author Impact Weight: 0.4

- Citation Impact Weight: 0.4

- Venue Impact Weight: 0.2
