## Supplementary material for "MENDELSEEK: An algorithm that predicts Mendelian Genes and elucidates what makes them special": WWP2.txt

# Claim: The gene WWP2 is likely associated with at least one Mendelian disease.

**Status**: processed

**Overall Rating**: -1

**Explanation**:

No relevant papers were found that support or refute this claim.


## Relevant Papers


## Other Reviewed Papers


### Missense variant in CCDC22 causes X-linked recessive intellectual disability with features of Ritscher-Schinzel/3C syndrome

**Why Not Relevant**: The provided paper content describes identification of a variant in CCDC22 as causal for an X-linked phenotype in a family. It contains no mention of the gene WWP2, no experimental or clinical evidence involving WWP2, and no mechanistic discussion that links CCDC22 findings to WWP2. Therefore the paper does not provide support for (or substantive evidence against) the claim that WWP2 is likely associated with at least one Mendelian disease. Absence of evidence in this single paper is not evidence of absence overall; to evaluate the WWP2 claim one would need papers that specifically study WWP2 variants, segregation with disease, functional assays, or established clinical associations.


[Read Paper](https://www.semanticscholar.org/paper/2644517cb053beb7fb2e4cde91791ba284ce557f)


### Heterozygous mutations affecting the protein kinase domain of CDK13 cause a syndromic form of developmental delay and intellectual disability

**Why Not Relevant**: The provided paper text exclusively addresses pathogenic variation in CDK13 and the resulting syndromic intellectual disability/congenital heart disease phenotype. There is no mention of WWP2, no data (genetic, clinical, or functional) on WWP2, and no discussion that would allow inference or transfer of evidence to WWP2. While the paper contains both direct genetic evidence and mechanistic interpretation (e.g., missense mutations in the kinase domain likely acting via a dominant negative mechanism) for CDK13 being a Mendelian-disease-associated gene, none of that evidence pertains to WWP2. Therefore the paper does not provide support for the claim that WWP2 is likely associated with at least one Mendelian disease. Any attempt to link WWP2 to disease would require independent genetic or functional evidence not present in this manuscript.


[Read Paper](https://www.semanticscholar.org/paper/504e280add9da26029c53b70b733fcb4eff04e23)


### Diagnostic yield of patients with undiagnosed intellectual disability, global developmental delay and multiples congenital anomalies using karyotype, microarray analysis, whole exome sequencing from Central Brazil

**Why Not Relevant**: This paper does not provide direct or mechanistic evidence regarding the claim that WWP2 is likely associated with at least one Mendelian disease. Nowhere in the provided text is WWP2 mentioned, nor are gene-level results or lists of implicated genes reported. The study describes using an ID gene panel of 1,252 genes and reports overall diagnostic yields, but without a gene list or per-gene findings one cannot determine whether WWP2 was included in the panel, whether any WWP2 variants were observed, or whether any mechanistic data (e.g., functional studies, pathway analyses) relating WWP2 to disease were produced. Therefore the content is only loosely relevant as methodological context (it shows that large gene panels can find pathogenic variants), but it does not support or refute an association between WWP2 and Mendelian disease. Important caveats: absence of mention of WWP2 in this paper is not evidence that WWP2 is not disease-associated—this cohort and the subset analyzed by exome (19 trios) are small, and the authors may have omitted gene-level details in the provided text. To evaluate the claim about WWP2, consult gene-disease resources (OMIM, ClinGen, ClinVar, HGMD), primary reports that specifically mention WWP2 variants in patients, and functional/mechanistic studies of WWP2. If needed, request full supplementary materials or gene lists from this study to check whether WWP2 was included and whether any variants were observed.


[Read Paper](https://www.semanticscholar.org/paper/23767daf16f48d9a91172aa0fc624f07b869a9c8)


### Prominent and Regressive Brain Developmental Disorders Associated with Nance-Horan Syndrome

**Why Not Relevant**: The paper content you provided exclusively discusses Nance-Horan syndrome and pathogenic variants in the NHS gene. There is no mention of the gene WWP2, no genetic or functional data concerning WWP2, and no discussion of WWP2 in any mechanistic context. Therefore this paper does not provide evidence for or against the claim that WWP2 is likely associated with at least one Mendelian disease. Absence of WWP2 information in this paper does not disprove the claim; it only means the paper is not relevant to evaluating that claim.


[Read Paper](https://www.semanticscholar.org/paper/d2d8619c882943bdd94ddc65f420190f03bf28b6)


### The Pathogenic RET Val804Met Variant in Acromegaly: A New Clinical Phenotype?

**Why Not Relevant**: The provided paper content exclusively discusses genetic findings related to RET pathogenic variants in somatotropinomas (acromegaly) and does not mention the gene WWP2, any variants in WWP2, or any phenotypes or mechanistic data related to WWP2. Key sentences focus on RET: for example, reporting "the first acromegaly patient carrying a RET pathogenic variant: c.2410G>A (rs79658334), p.Val804Met," investigation of clinical significance of this RET variant, and review of RET oncogenic mechanisms. Because there is no mention of WWP2, no germline or somatic variants in WWP2, no mechanistic pathway involving WWP2, and no phenotype linked to WWP2 in the provided text, the paper provides neither direct evidence nor mechanistic evidence for or against the claim that WWP2 is associated with a Mendelian disease. Note that absence of mention in this paper does not disprove the claim; it only means this paper is not a source of evidence regarding WWP2.


[Read Paper](https://www.semanticscholar.org/paper/e1f9ea673f7e10424a0c3260e54b7649179f9004)


### Case report: A rare case of pyruvate kinase deficiency and Crigler-Najjar syndrome type II with a novel pathogenic variant of PKLR and UGT1A1 mutation

**Why Not Relevant**: The paper is a clinical case report focused entirely on a single patient with Pyruvate Kinase Deficiency (PKLR) and Crigler-Najjar syndrome (UGT1A1). The text provided names and mechanistic detail only for PKLR and UDP-glucuronosyltransferase/UGT1A1 and reports specific variants in those genes. Nowhere in the provided content is the gene WWP2 mentioned, studied, or discussed at the genetic, clinical, or mechanistic level. Therefore the paper provides no direct evidence (e.g., variant cosegregation, population enrichment, functional assays) nor mechanistic evidence (pathway overlap, molecular function linking WWP2 to disease phenotype) supporting or refuting the claim that WWP2 is likely associated with at least one Mendelian disease. The only possible, very weak negative inference is that WWP2 was not implicated in this particular case; however, absence of mention in a single case report is not informative about WWP2's broader disease associations. In short, the paper is not relevant to assessing WWP2's association with Mendelian disease because it does not address WWP2 at all.


[Read Paper](https://www.semanticscholar.org/paper/5109dccd5fcfea16c12d3477e55fc868643504f1)


### The first pineoblastoma case report of a patient with Sotos syndrome harboring NSD1 germline mutation

**Why Not Relevant**: The provided paper content describes a single clinical case of Sotos syndrome attributed to a novel de novo germline NSD1 likely pathogenic variant. There is no mention of the gene WWP2, no data linking WWP2 to the phenotype, and no mechanistic discussion that would tie WWP2 to Mendelian disease in this text. Because the excerpt exclusively concerns NSD1 and Sotos syndrome, it provides neither direct evidence (genetic association, segregation, case reports implicating WWP2) nor mechanistic evidence (functional studies, pathway connections) supporting the claim that WWP2 is likely associated with at least one Mendelian disease. Conclusion is limited to the provided content; if the full paper contains additional sections not supplied here, those were not available for review.


[Read Paper](https://www.semanticscholar.org/paper/96dc04c6134726797978ce76de5ce0e5e5a930f9)


### [A clinical case of reverse left ventricular remodeling in patient with pathogenic TTN mutation. Case report].

**Why Not Relevant**: The provided paper content discusses dilated cardiomyopathy and genetic mutations primarily in the titin gene (TTN). It contains no mention of the gene WWP2, no data (genetic, functional, clinical, or epidemiological) about WWP2, and no mechanistic discussion that links WWP2 to DCM or any other Mendelian disorder. Therefore this paper provides no direct evidence for or against the claim that WWP2 is likely associated with at least one Mendelian disease. Note that absence of evidence in this single paper is not evidence of absence for WWP2 overall; a lack of mention here simply means this source is not informative for the claim. To evaluate the claim about WWP2 one would need to consult studies that specifically analyze WWP2 variants, functional data, segregation in families, case reports, or curated gene–disease resources (e.g., OMIM, ClinGen, HGMD).


[Read Paper](https://www.semanticscholar.org/paper/60d1854470eefd43d337c3a90dc9927f877e022b)


### Case report: Salivary duct carcinoma in a patient with a germline CDH1 pathogenic variant - expanding the spectrum of hereditary cancer predisposition syndromes

**Why Not Relevant**: The provided paper content focuses entirely on CDH1 germline and somatic variants in a single case of salivary duct carcinoma and contains no mention of the gene WWP2, nor any genetic, epidemiologic, or mechanistic data pertaining to WWP2. Therefore it provides neither direct evidence (e.g., cases, segregation, pathogenic variants in WWP2 causing a Mendelian phenotype) nor mechanistic evidence (e.g., biochemical pathways, functional assays implicating WWP2) that would support or refute the claim that WWP2 is likely associated with at least one Mendelian disease. Additional reasons for non-relevance: (1) study type is a single-case clinical report about CDH1, not a genetic association study; (2) the molecular findings and mechanistic discussion are specific to CDH1 biallelic inactivation and tumor predisposition; (3) there are no data (variant lists, exome results mentioning WWP2, family segregation, functional experiments) that could be reinterpreted as evidence for WWP2. Given the complete absence of WWP2 in the text and the narrow CDH1 focus, the paper does not inform the claim about WWP2 and is therefore rated as having zero relevance.


[Read Paper](https://www.semanticscholar.org/paper/ce73436b92f6b462eeb27adc0bd46ffc929aa25e)


## Search Queries Used

- WWP2 pathogenic variant Mendelian disease case report germline mutation patient clinical phenotype

- WWP2 rare variants exome sequencing cohort developmental disorders intellectual disability congenital anomalies gene burden study

- WWP2 WW domain containing E3 ubiquitin protein ligase 2 knockout mouse zebrafish loss of function developmental phenotype mechanism ubiquitination substrate

- WWP2 ClinVar OMIM ClinGen Decipher gnomAD pLI loss of function constraint gene disease association review


## Bibliometric Configuration

- Use Bibliometrics: True

- Author Impact Weight: 0.4

- Citation Impact Weight: 0.4

- Venue Impact Weight: 0.2
