## Supplementary material for "MENDELSEEK: An algorithm that predicts Mendelian Genes and elucidates what makes them special": XRCC5.txt

# Claim: The gene XRCC5 is likely associated with at least one Mendelian disease.

**Status**: processed

**Overall Rating**: 4

**Explanation**:

Supporting evidence.
The provided papers describe XRCC5 (Ku80) as a core non-homologous end joining (NHEJ) factor required for repair of DNA double-strand breaks and for V(D)J recombination. Multiple excerpts state that loss of Ku (Ku80) compromises rejoining of radiation-induced DSBs, increases end degradation, alters repair pathway usage, and produces pronounced hypersensitivity to ionizing radiation. Critically, one excerpt explicitly says that “deficiency of either of these proteins leads to severe clinical immunodeficiency in mice and humans (21–25).” That phrasing places Ku80 (XRCC5) among the NHEJ core proteins whose deficiency is linked to severe immunodeficiency phenotypes, which are classical Mendelian disorders arising from defective V(D)J recombination.

Caveats or contradictory evidence.
The papers are largely mechanistic and experimental (cellular and mouse KO) studies of NHEJ components; they do not present primary human genetic case reports or variant-to-disease segregation data for XRCC5 in the excerpts provided. The specific human evidence is asserted by citation but not reproduced in the excerpts; therefore the assertion that humans are affected may be referencing work on other NHEJ genes as well, or aggregated evidence across the pathway. Also, Ku80 knockout in mice is noted to be viable (unlike XRCC4 KO which is embryonic lethal), which suggests species- and context-dependent differences in phenotypic severity. Thus, while strong mechanistic links exist between XRCC5 function and processes that, when defective, produce Mendelian immunodeficiencies, the excerpts do not provide direct genetic proof (e.g., pedigrees, pathogenic variants, inheritance pattern) tying specific XRCC5 mutations to a named Mendelian syndrome.

Analysis of potential underlying mechanisms and implications.
XRCC5 encodes Ku80, which heterodimerizes with Ku70 to bind DNA ends and recruit the DNA-PKcs/Artemis and the XRCC4/LigaseIV/XLF complex for classical NHEJ. Disruption of Ku80 impairs V(D)J recombination by preventing proper end recognition and joining of antigen receptor gene segments, a canonical mechanism that produces severe combined immunodeficiency (SCID) or related immunodeficiencies when defective (examples: DCLRE1C/Artemis, LIG4, NHEJ1/XLF). Given this conserved mechanism and the fact that defects in multiple other NHEJ genes are established Mendelian causes of immunodeficiency and radiosensitivity syndromes, XRCC5 loss-of-function is mechanistically well positioned to produce a Mendelian recessive immunodeficiency phenotype in humans. The viability difference in mice between XRCC4 and Ku80 knockouts suggests phenotypic variability depending on the exact molecular lesion, redundancy, or compensation, but does not undermine the causal plausibility for human disease.

Which rating is most appropriate.
On balance, the mechanistic evidence directly links XRCC5 function to processes whose disruption causes Mendelian immunodeficiencies, and the excerpts even state that deficiency of these core NHEJ proteins leads to severe clinical immunodeficiency in humans. However, the provided material stops short of showing primary human genetic evidence (specific pathogenic XRCC5 variants, pedigrees, or clinical case descriptions). Given the strong mechanistic rationale, pathway-level precedent (other NHEJ genes cause Mendelian disease), and the explicit statement in the excerpts that humans can be affected, the claim that XRCC5 is likely associated with at least one Mendelian disease is well supported but not established beyond doubt by the supplied excerpts. Therefore the most appropriate rating is "Likely True."


**Final Reasoning**:

The supplied excerpts establish that XRCC5/Ku80 is an essential NHEJ protein required for V(D)J recombination and radiation-induced DSB repair; they note that deficiency in NHEJ core proteins produces severe clinical immunodeficiency in mice and humans. This provides strong mechanistic and pathway-level evidence that loss of XRCC5 function can produce Mendelian immunodeficiency. The primary limitation is that the excerpts do not include direct human genetic case data tying specific XRCC5 variants to a named Mendelian syndrome. Taking the explicit statement about human clinical immunodeficiency together with the robust mechanistic context and precedent from other NHEJ genes, the preponderance of evidence supports the claim. Hence the rating is "Likely True."


## Relevant Papers


### Distinct roles of XRCC4 and Ku80 in non-homologous end-joining of endonuclease- and ionizing radiation-induced DNA double-strand breaks

**Authors**: Leonie Schulte-Uentrop (H-index: 1), J. Dahm-Daphi (H-index: 31)

**Relevance**: 0.45

**Bibliometric Impact**: 0.28990000000000005


**Excerpts**:

- Non-homologous end-joining (NHEJ) of DNA double-strand breaks (DSBs) is mediated by two protein complexes comprising Ku80/Ku70/DNA-PKcs/Artemis and XRCC4/LigaseIV/XLF.

- Loss of Ku or XRCC4/LigaseIV function compromises the rejoining of radiation-induced DSBs and leads to defective V(D)J recombination.

- We found that the frequency of NHEJ was reduced by more than 20-fold in XRCC4−/− compared to XRCC4+/+ cells, while a Ku80 knock-out reduced the rejoining efficiency by only 1.4-fold.

- In contrast, lack of either XRCC4 or Ku80 increased end degradation and shifted repair towards a mode that used longer terminal microhomologies for rejoining.

- However, both proteins proved to be essential for the repair of radiation-induced DSBs.

- The remarkably different phenotype of XRCC4- and Ku80-deficient cells with regard to the repair of enzyme-induced DSBs mirrors the embryonic lethality of XRCC4 knock-out mice as opposed to the viability of the Ku80 knock-out.

- Thus, I-SceI-induced breaks may resemble DSBs arising during normal DNA metabolism and mouse development. The removal of these breaks likely has different genetic requirements than the repair of radiation-induced DSBs.


**Explanations**:

- Mechanistic relevance: This sentence places Ku80 (the protein product of XRCC5) explicitly in the canonical NHEJ machinery, establishing a direct mechanistic role for XRCC5 in repairing DNA double-strand breaks. Strength: identifies biological function directly relevant to disorders of DNA repair. Limitation: descriptive mechanistic statement from cellular/biochemical context; does not by itself demonstrate that XRCC5 variation causes Mendelian disease in humans.

- Mechanistic relevance with disease implication: States that loss of Ku (XRCC5) function compromises rejoining of radiation-induced DSBs and causes defective V(D)J recombination. Strength: links Ku80 deficiency to impaired V(D)J recombination, a process whose defects are known to underlie Mendelian immunodeficiency syndromes (mechanistic plausibility). Limitation: phraseology refers to 'loss of Ku' broadly (likely in experimental systems); no human clinical cases or direct demonstration of Mendelian inheritance are provided in this paper.

- Experimental/mechanistic data: Provides quantitative experimental evidence that complete loss of XRCC4 drastically reduces NHEJ (>20-fold) while Ku80 loss produces a much smaller reduction (1.4-fold) in rejoining efficiency for I-SceI-induced breaks. Strength: gives comparative functional impact of knocking out XRCC5 (Ku80) versus XRCC4, informing expectation about phenotypic severity. Limitation: data are from murine fibroblast reporter assays with enzyme-induced breaks; the excerpt lacks sample sizes, statistical tests, and human data, so extrapolation to Mendelian disease causation in humans is indirect.

- Mechanistic relevance: Shows that both XRCC4 and Ku80 deficiencies alter repair pathway choice (increased end degradation and microhomology usage), indicating that XRCC5 disruption changes DSB processing in ways that could produce genomic instability. Strength: mechanistic pathway supporting plausibility that XRCC5 defects could lead to disease phenotypes tied to faulty DNA repair. Limitation: observed in the experimental system used; effect magnitude and phenotypic consequences in whole organisms or humans are not provided.

- Mechanistic relevance: States both proteins are essential for repair of radiation-induced DSBs. Strength: underscores that XRCC5 is required for at least some types of DSB repair, which increases plausibility that XRCC5 loss-of-function could have pathological consequences. Limitation: 'essential' here is presented in the context of their assays/experiments; again no direct human clinical evidence or inheritance pattern is reported.

- Organismal phenotype context: Notes that differing cellular phenotypes mirror organismal outcomes—XRCC4 knock-out is embryonic lethal while Ku80 knock-out mice are viable. Strength: provides in vivo evidence that complete loss of Ku80 is compatible with life in mice, which informs expectations about the severity and potential expressivity of XRCC5-related phenotypes. Limitation: species differences may affect transferability to humans; viability in knockout mice does not rule out Mendelian disease with non-lethal but clinically significant phenotypes, nor does it address heterozygous vs homozygous states.

- Contextual limitation and generalizability: Authors caution that I-SceI-induced breaks may mimic certain endogenous DSBs and that genetic requirements differ between enzyme-induced and radiation-induced breaks. Strength: helps define which types of DNA lesions the experimental findings most closely apply to, supporting nuanced mechanistic interpretation. Limitation: implies that results may not generalize to all DSB types or to in vivo human biology, limiting the strength of inference about human Mendelian disease causation.


[Read Paper](https://www.semanticscholar.org/paper/49c21c411726c8fd183d10767c3c4ba7ec01cbde)


### Deficiency of XLF and PAXX prevents DNA double-strand break repair by non-homologous end joining in lymphocytes

**Authors**: Putzer J. Hung (H-index: 7), Andrea L Bredemeyer (H-index: 31)

**Relevance**: 0.3

**Bibliometric Impact**: 0.2544571428571429


**Excerpts**:

- ABSTRACT Non-homologous end joining (NHEJ) is a major DNA double-strand break (DSB) repair pathway that functions in all phases of the cell cycle.

- NHEJ repairs genotoxic and physiological DSBs, such as those generated by ionizing radiation and during V(D)J recombination at antigen receptor loci, respectively.

- DNA end joining by NHEJ relies on the core factors Ku70, Ku80, XRCC4, and DNA Ligase IV.

- The XRCC4-like factor (XLF) participates in NHEJ through its interaction with XRCC4, and XLF deficiency in humans leads to immunodeficiency and increased sensitivity to ionizing radiation.

- However, XLF is dispensable for NHEJ-mediated DSB repair during V(D)J recombination in murine lymphocytes, where it may have redundant functions with other DSB repair factors.

- Paralog of XRCC4 and XLF (PAXX) is a recently identified NHEJ factor that has structural similarity to XRCC4 and XLF.

- Here we show that PAXX is also dispensable for NHEJ during V(D)J recombination and during the repair of genotoxic DSBs in lymphocytes.

- However, a combined deficiency of PAXX and XLF blocks NHEJ with a severity comparable to that observed in DNA Ligase IV-deficient cells.

- Similar to XLF, PAXX interacts with Ku through its C-terminal region, and mutations that disrupt Ku binding prevent PAXX from promoting NHEJ in XLF-deficient lymphocytes.

- Our findings suggest that the PAXX and XLF proteins may have redundant functions during NHEJ.


**Explanations**:

- General mechanistic context: This sentence establishes the biological pathway (NHEJ) in which the XRCC5 protein product (Ku80) operates. It is mechanistic background relevant to evaluating whether disruption of components (including XRCC5) could cause Mendelian disease. Limitation: this is general pathway description and contains no genetic or phenotypic data about XRCC5 variants or human disease.

- Mechanistic context linking NHEJ activity to physiologic processes (V(D)J recombination) whose disruption produces immunodeficiency phenotypes. This increases plausibility that mutations in NHEJ components (including XRCC5) could produce Mendelian immunodeficiency. Limitation: again descriptive; no direct data on XRCC5 variants or human cases.

- Direct identification of Ku80 as a core NHEJ factor (Ku80 is encoded by XRCC5). This is mechanistic evidence that XRCC5 is essential to an important DNA-repair pathway; loss-of-function in such a core factor is biologically plausible to cause a Mendelian disorder. Limitation: the sentence does not report genetic variation, human phenotypes, or disease association for XRCC5 itself.

- This sentence provides a direct human-disease example where deficiency of an NHEJ factor (XLF) causes an inherited immunodeficiency and radiation sensitivity. This is strong, direct evidence that pathogenic variants in NHEJ genes can produce Mendelian disease in humans, thus supporting the plausibility that other NHEJ genes (like XRCC5) could likewise underlie Mendelian disorders. Limitation: the sentence refers to XLF, not XRCC5; it is an analogy rather than direct evidence for XRCC5.

- This sentence reports that XLF is dispensable in murine V(D)J recombination because of redundancy, highlighting that loss of a single NHEJ factor may be compensated by others. Mechanistic implication: redundancy among NHEJ factors could mask phenotypes from heterozygous or even homozygous XRCC5 variants depending on context and species, lowering the likelihood that XRCC5 variation always produces a Mendelian phenotype. Limitation: this is murine data about XLF, not XRCC5 in humans.

- Mechanistic context: introduces PAXX as a newly identified NHEJ factor related to XRCC4 and XLF, illustrating the network complexity and paralogy within NHEJ. This supports consideration of redundancy and functional overlap in the pathway relevant to assessing XRCC5's disease potential. Limitation: PAXX is distinct from XRCC5; no genetic disease data for XRCC5 is provided.

- Experimental result: PAXX is dispensable for NHEJ during V(D)J recombination and genotoxic DSB repair in lymphocytes, indicating single-gene loss may be tolerated because of redundancy. Mechanistic implication for XRCC5: if XRCC5 function were similarly redundant in certain contexts, XRCC5 mutations might not always produce an overt Mendelian phenotype. Limitation: result pertains to PAXX, not XRCC5; experimental system is lymphocytes and (implied) murine models.

- Mechanistic result showing that combined loss of PAXX and XLF produces a severe NHEJ defect comparable to Ligase IV deficiency. Relevance to XRCC5: this emphasizes that combined deficiencies can unmask severe phenotypes, suggesting that XRCC5 pathogenicity could depend on genetic background or combined defects. Limitation: the observation is about PAXX and XLF; it does not test XRCC5 variants or human disease.

- Mechanistic detail: PAXX and XLF interact with Ku (the complex that includes Ku80/XRCC5); mutations disrupting Ku binding impair PAXX function in XLF-deficient cells. This directly implicates interactions with Ku/Ku80 as functionally important, strengthening the mechanistic link between Ku80 (XRCC5 product) and NHEJ function. Limitation: functional interaction data does not substitute for human genetic association; no XRCC5 mutation phenotypes are reported.

- Synthesis: the authors conclude redundancy between PAXX and XLF in NHEJ. For the XRCC5 claim, this highlights that NHEJ factor redundancy is an important modifier of phenotypic outcome—i.e., XRCC5 loss might or might not cause a Mendelian disorder depending on redundancy and context. Limitation: conclusion refers to PAXX/XLF redundancy rather than XRCC5-specific data.


[Read Paper](https://www.semanticscholar.org/paper/57dd5f9cbe9b99eb868510061539ae0093eae2df)


### „Distinct roles of XRCC4 and Ku80 in non-homologous end-joining of endonuclease- and ionizing radiation-induced DNA double-strand breaks“

**Authors**: Leonie Schulte-Uentrop (H-index: 1), Lena Schliecker (H-index: 1)

**Relevance**: 0.7

**Bibliometric Impact**: 0.008


**Excerpts**:

- Non-homologous end-joining (NHEJ) of DNA doublestrand breaks (DSBs) is mediated by two protein complexes comprising Ku80/Ku70/DNA-PKcs/ Artemis and XRCC4/LigaseIV/XLF.

- Loss of Ku or XRCC4/LigaseIV function compromises the rejoining of radiation-induced DSBs and leads to defective V(D)J recombination.

- We found that the frequency of NHEJ was reduced by more than 20-fold in XRCC4!/! compared to XRCC4+/+ cells, while a Ku80 knock-out reduced the rejoining efficiency by only 1.4-fold.

- In contrast, lack of either XRCC4 or Ku80 increased end degradation and shifted repair towards a mode that used longer terminal microhomologies for rejoining.

- Cells deficient in any of the NHEJ core proteins display pronounced hypersensitivity to ionizing radiation (IR) and a reduced ability to rejoin IR-induced DSBs (17).

- The NHEJ core proteins are also required for V(D)J recombination and class-switch recombination (18–20).

- Deficiency of either of these proteins leads to severe clinical immunodeficiency in mice and humans (21–25).

- Genetic knock-out of XRCC4 but not of Ku80 in mice leads to embryonic death, suggesting that XRCC4 function is critical for the removal of DSBs that arise during development (22,23).

- Thus, I-SceI-induced breaks may resemble DSBs arising during normal DNA metabolism and mouse development.


**Explanations**:

- This sentence identifies Ku80 as a core component of the NHEJ machinery (Ku80/Ku70/DNA-PKcs/Artemis), directly linking the Ku80 protein (encoded by XRCC5) to an essential DNA repair pathway. This is mechanistic evidence: it shows that Ku80 participates in a fundamental cellular process whose disruption can plausibly lead to disease. Limitation: it is a statement of function/mechanism rather than evidence of disease in humans.

- This sentence states that loss of Ku (Ku80) function compromises DSB rejoining and causes defective V(D)J recombination. Mechanistic evidence: defective V(D)J recombination is a known cause of immunodeficiency syndromes (Mendelian). Limitation: the sentence describes functional consequences (in cells/animals) but does not itself report human genetic data tying XRCC5 variants to a Mendelian disease.

- This result quantifies the functional impact of complete XRCC4 loss versus Ku80 loss on NHEJ frequency in the experimental reporter system (mouse cells). Mechanistic/experimental evidence: it shows that Ku80 loss produces a measurable (albeit modest) defect in chromosomal DSB rejoining, supporting biological plausibility that XRCC5 disruption can perturb genome maintenance. Limitations: these are mouse cell assays and reflect knockout phenotypes in vitro; they do not provide human clinical genetic evidence and the magnitude of effect for Ku80 was small (1.4-fold), which may influence penetrance/severity considerations.

- This sentence describes that both XRCC4 and Ku80 deficiencies change repair outcomes (increased end degradation and longer microhomology use). Mechanistic evidence: it indicates how Ku80 loss alters DNA end processing, which could produce the kinds of genomic rearrangements underlying disease (e.g., immunodeficiency, cancer). Limitation: contextual/experimental (reporter assay) rather than direct human genotype–phenotype data.

- This sentence documents that cells lacking NHEJ core proteins are hypersensitive to ionizing radiation and have reduced ability to rejoin IR-induced DSBs. Mechanistic evidence: supports that disruption of NHEJ components (including Ku80/XRCC5) has strong cellular consequences that can lead to pathology. Limitation: this is cellular phenotype data; it does not by itself prove a Mendelian disease association in humans.

- This sentence states that NHEJ core proteins are required for V(D)J and class-switch recombination. Mechanistic evidence: since V(D)J recombination is essential for lymphocyte development and adaptive immunity, disruption of NHEJ components (including Ku80/XRCC5) plausibly causes Mendelian immunodeficiencies. Limitation: the sentence is mechanistic/pathway-level and does not provide direct human genetic association data.

- This sentence asserts that deficiency of these proteins causes severe clinical immunodeficiency in mice and humans. This is the closest statement to direct disease association: it links NHEJ protein deficiency to clinical immunodeficiency in humans, implying that mutations in NHEJ genes (potentially including XRCC5/Ku80) have been implicated in Mendelian immunodeficiency. Strength: cites prior human/mouse clinical literature (refs 21–25). Limitations: the paper does not itself present primary human genetic data for XRCC5; the phrase "either of these proteins" is context-dependent (refers to NHEJ proteins previously discussed) and does not explicitly name XRCC5/XRCC5 as the causal gene in humans in this manuscript—association is inferred via cited literature rather than demonstrated here.

- This sentence highlights that complete loss of XRCC4 in mice is embryonic lethal whereas Ku80 knock-out is not. Interpretative mechanistic evidence: it shows differing in vivo importance of different NHEJ components during development—suggesting that loss of Ku80 (XRCC5) may be tolerated to an extent in mice, which affects interpretation of how strongly XRCC5 mutations might present as an obvious Mendelian syndrome. Limitation: mouse viability differences do not directly predict human disease manifestations or penetrance for XRCC5 variants.

- This concluding interpretive sentence indicates that the experimentally induced breaks may model physiologic DSBs arising during development and normal DNA metabolism. Mechanistic relevance: supports extrapolation that the observed Ku80-dependent repair phenotypes could relate to developmental processes whose disruption leads to disease. Limitation: speculative—bridges an experimental model to in vivo developmental contexts but is not direct human genetic evidence.


[Read Paper](https://www.semanticscholar.org/paper/a5468e845c30da72f8ad16588ee676db59729b54)


## Other Reviewed Papers


### SHLD1 is dispensable for 53BP1-dependent V(D)J recombination but critical for productive class switch recombination

**Why Not Relevant**: The provided paper content discusses SHLD1 and XLF roles in lymphocyte development and class-switch recombination but contains no mention of XRCC5, any XRCC5 variants, patient phenotypes, or experimental manipulations of XRCC5. Therefore it provides neither direct genetic/clinical evidence nor explicit mechanistic evidence implicating XRCC5 in a Mendelian disease. Although the biological processes referenced (RAG-mediated V(D)J recombination and class-switch recombination) are mediated in part by DNA double-strand break repair pathways (e.g., classical NHEJ), which conceptually could involve XRCC5, the excerpt contains no data or statements about XRCC5 itself. For these reasons the paper content as provided is not relevant to assessing whether XRCC5 is likely associated with at least one Mendelian disease.


[Read Paper](https://www.semanticscholar.org/paper/de715755e75971bc765e70acd8327c9bc803e852)


## Search Queries Used

- XRCC5 mutation human case report Mendelian disease Ku80 X ray repair cross complementing 5 biallelic homozygous pathogenic variant autosomal recessive

- XRCC5 Ku80 non homologous end joining NHEJ V D J recombination DNA double strand break repair immunodeficiency lymphocyte development radiosensitivity

- XRCC5 germline variant ClinVar OMIM HGMD loss of function frameshift nonsense splice exome sequencing family segregation monogenic Mendelian disease

- XRCC5 knockout mouse phenotype immunodeficiency radiosensitivity tumor predisposition growth retardation cellular V D J recombination assay complementation


## Usage Statistics

- Prompt Tokens: 0

- Completion Tokens: 0

- Total Tokens: 0

- Estimated Cost: $0.0290


## Bibliometric Configuration

- Use Bibliometrics: True

- Author Impact Weight: 0.4

- Citation Impact Weight: 0.4

- Venue Impact Weight: 0.2
