## Supplementary material for "MENDELSEEK: An algorithm that predicts Mendelian Genes and elucidates what makes them special": XRCC6.txt

# Claim: The gene XRCC6 is likely associated with at least one Mendelian disease.

**Status**: processed

**Overall Rating**: 4

**Explanation**:

Supporting evidence.
The excerpts identify XRCC6/Ku70 as one of the core non-homologous end-joining (NHEJ) factors (Ku70, Ku80, DNA-PKcs, LIG4, XRCC4, XLF) and describe clear consequences of loss of these factors in model systems. In mice, absence of Ku70 (Xrcc6) and other NHEJ core proteins produces immunodeficiency due to blocked V(D)J recombination in B and T lymphocytes, ionizing radiation sensitivity, chromosomal instability, premature cellular senescence, and increased cancer (pro-B lymphoma) when combined with p53 deficiency. The second paper highlights that defects in factors that participate in end-processing or end-ligation of V(D)J recombination (including KU70/80) are drivers of disease, notably severe combined immunodeficiency (SCID). Together these points provide strong mechanistic and phenotypic support that perturbing XRCC6 function produces syndromic phenotypes consistent with Mendelian disorders affecting lymphocyte development, genomic stability, and cancer predisposition.

Caveats or contradictory evidence.
The provided excerpts document mouse phenotypes and general statements about NHEJ factors, but do not present direct human genetic evidence (i.e., reported pathogenic germline XRCC6 variants segregating with a Mendelian disorder in pedigrees). Mouse knockout phenotypes do not always translate into an identifiable human Mendelian disease—some knockouts are embryonic lethal, others produce phenotypes that are clinically heterogeneous, and some genes show functional redundancy or partial compensation in humans. Also, the second paper lists multiple NHEJ factors causally implicated in human immunodeficiency (e.g., LIG4, XRCC4, XLF) but does not explicitly document confirmed human Mendelian disease caused by XRCC6 variants. Thus the evidence here is indirect and inferential rather than direct clinical genetic proof.

Analysis of potential underlying mechanisms, deeper relationships, or second-order implications.
XRCC6 encodes Ku70, which dimerizes with Ku80 to recognize DNA double-strand break ends and recruit downstream NHEJ machinery. Loss-of-function of XRCC6 would be expected to compromise V(D)J recombination, producing failure of B- and T-cell receptor gene assembly and a block in lymphocyte development—a classic mechanistic route to a recessive SCID-like immunodeficiency. In addition, defective NHEJ causes genomic instability and radiosensitivity, predisposing to malignancy and premature cellular senescence/aging phenotypes; these outcomes are consistent with Mendelian syndromes caused by other DSB-repair genes. Given the central, nonredundant biochemical role of the Ku heterodimer in NHEJ, biallelic inactivating variants in XRCC6 are mechanistically plausible causes of a recessive Mendelian disease characterized by immune deficiency and genome instability. Heterozygous carriers could be clinically unaffected or show milder phenotypes depending on residual protein function and dosage sensitivity.

Which rating is most appropriate.
Balancing the evidence: the mechanistic role of XRCC6, the clear and reproducible knockout phenotypes in mice that mirror human Mendelian syndromes caused by other NHEJ factors, and the explicit linking of NHEJ factor defects to SCID provide strong circumstantial support that XRCC6 is likely to be associated with at least one Mendelian disease. The primary limitation is the absence of direct human genetic case reports in the provided material. That makes the support strong but not definitive. Therefore the most appropriate qualitative judgment from the provided evidence is that the claim is "Likely True": XRCC6 is plausibly and probably associated with a Mendelian disorder (most plausibly a recessive immunodeficiency and/or genome-instability syndrome), but direct human-gene-to-disease evidence is not present in the supplied excerpts.


**Final Reasoning**:

I reviewed the logic and evidence: (1) XRCC6 is a core NHEJ gene (Ku70) required for V(D)J recombination; (2) Loss of XRCC6 in mice causes immunodeficiency and other Mendelian-like phenotypes (radiation sensitivity, genomic instability, cancer predisposition); (3) defects in analogous NHEJ genes are established causes of human Mendelian disorders such as SCID. The chain of inference from conserved mechanistic role + animal knockout phenotypes + precedent from related genes yields a strong likelihood that XRCC6 can underlie a Mendelian disease in humans. The main weakness is lack of direct human genetic/clinical case evidence in the provided papers. That weakness reduces the confidence from "Highly Supported" to "Likely True," but does not overturn the substantial mechanistic and phenotypic support. I therefore affirm the "Likely True" rating.


## Relevant Papers


### Análisis de la deficiencia de la DNA polimerasa [mu] en la unión de extremos no homólogos. Implicaciones en envejecimiento y desarrollo tumoral

**Authors**: B. González (H-index: 5)

**Relevance**: 0.6

**Bibliometric Impact**: 0.04000000000000001


**Excerpts**:

- The two principal mechanisms for DSBs repair are Homologous Recombination (HR) and Non Homologous End Joining (NHEJ).

- Mice lacking any of the NHEJ core factors (Ku70, Ku80, DNA-PKcs, LigIV, Xrcc4 and XLF) show immunodeficiency due to blocked recombination of gene segments in B and T lymphocytes receptors (a NHEJ mediated process).

- These mice also exhibit ionizing radiation sensitivity and some of them (Ku80-/- and DNA-PKcs-/-) have premature aging phenotypes.

- NHEJ deficient mouse embryonic fibroblasts (MEFs) enter senescence prematurely due to persistent DNA damage and chromosomal instability.

- The combined deficiency of a NHEJ core factor and p53 results in pro-B lymphoma development, due to the survival of chromosomically aberrant lymphocytes, many of which carry specific translocations between the IgH enhancer locus and C-myc .

- All together, these data demonstrate that pol ? participates in general NHEJ pathway: 1) Its abscence affects the hematopoietic system and non hematopoietic tissues, as shown by the higher chromosome instability and/or ?H2AX levels in medula osea, spleen, liver and MEFs ?-/- under genotoxic stress.


**Explanations**:

- Context-setting (mechanistic). This sentence defines the two major DSB repair pathways and places NHEJ as the relevant pathway in which XRCC6 (Ku70) functions. It is mechanistic background that supports plausibility: if XRCC6 encodes a core NHEJ factor, defects could reasonably produce disease related to defective DSB repair. Limitation: this is general background and does not provide direct evidence linking XRCC6 to a Mendelian disease.

- Directly relevant mechanistic evidence. This sentence explicitly names Ku70 among the 'NHEJ core factors' whose loss in mice causes immunodeficiency due to blocked V(D)J recombination. Because XRCC6 encodes Ku70, this provides strong mechanistic evidence that loss of XRCC6 function can produce a severe, Mendelian-like phenotype (immunodeficiency) in a mammalian model. Limitations: the evidence is from mouse knockouts, not human patients; the sentence lists multiple core factors together without quantifying the phenotype for each gene specifically; no human inheritance or clinical data are provided here.

- Mechanistic/phenotypic evidence in model organisms. This sentence shows that loss of NHEJ components yields radiation sensitivity and premature aging in some cases. It supports the claim by demonstrating that disruption of NHEJ core proteins (which include Ku70/XRCC6) causes clear, deleterious organismal phenotypes. Limitation: Ku80 and DNA-PKcs are singled out for premature aging; Ku70 is not specifically singled out for premature aging here, and again data are from mice rather than humans.

- Mechanistic cellular evidence. This sentence indicates that NHEJ deficiency causes cellular phenotypes (premature senescence, persistent DNA damage, chromosomal instability) consistent with a gene whose loss would yield a severe, heritable disorder. This supports plausibility that XRCC6 loss-of-function could cause Mendelian disease affecting tissues dependent on genomic stability. Limitation: this is an in vitro/primary-cell observation in mice and is not direct human clinical evidence.

- Mechanistic evidence about genetic interactions and oncogenic outcomes. This sentence shows that combining loss of a NHEJ core factor with p53 deficiency drives lymphoma due to chromosomal instability and oncogenic translocations. It strengthens the mechanistic argument that defective NHEJ (which would include XRCC6 dysfunction) can produce profound, heritable phenotypes (here in mice) and specific disease manifestations (lymphoma). Limitation: this is an interaction phenotype in mice and depends on p53 loss; it does not demonstrate that XRCC6 mutation alone causes a Mendelian disease in humans.

- Direct statement about the role of the polymerase under study in general NHEJ and tissue effects. Although focused on pol ?, this sentence documents that disruption of general NHEJ components elevates chromosome instability and DNA-damage markers across multiple tissues (bone marrow, spleen, liver, MEFs) under genotoxic stress. This supports the broader mechanistic argument that NHEJ core factors (including Ku70/XRCC6) are essential for genome maintenance and that their loss can produce systemic, disease-relevant phenotypes. Limitation: this is indirect for XRCC6 specifically (the paper's experimental focus is pol ?) and the observations are under genotoxic stress in mice.


[Read Paper](https://www.semanticscholar.org/paper/8811cbccaef1f193f2b205f2687afe2b30cee904)


### The Lysine Histone Methyltransferase SETD2 Is Required for Appropriate Immunoglobulin VDJ Recombination

**Authors**: S. H. Chu (H-index: 9), S. Armstrong (H-index: 79)

**Relevance**: 0.62

**Bibliometric Impact**: 0.552


**Excerpts**:

- Rearrangement of the immunoglobulin heavy chain locus (IgH) through non-homologous end-joining (NHEJ)-mediated VDJ recombination is a requisite step in normal lymphocyte development.

- Loss or defects of these factors can impair end-processing (DNA-PKcs, ARTEMIS) or end-ligation (KU70/80, XRCC4, XLF, LIGIV) of rearrangement and are drivers of disease, notably, severe combined immunodeficiency (SCID).

- While heterozygous mice had no overt phenotype, mice with homozygous loss of Setd2(Setd2KO) displayed aberrant hematopoiesis, with a significant depletion of lymphoid primed progenitor population and B/T-cells in hematopoietic tissues.

- Setd2 loss early in hematopoiesis lead to a profound block of B-cell development at the proB cell stage and an early block in T-cell development, in a compartment where TCRβ rearrangements initiate.

- These results phenocopy defects or loss of NHEJ end-ligation factors and suggest that Setd2 and H3K36me3 could play a pivotal role in the end-ligation steps of NHEJ.

- Defects in end-ligation and resolution of VDJ intermediates have previously been shown to result in the formation of aberrant coding-signal hybrid joints.

- Remarkably, upon induction in Setd2KO v-Abl lines, we detected the formation of hybrid joints by both PCR and by the loss of a GFP-based coding-joint formation signal from a reporter expressed in these cells.


**Explanations**:

- Contextual/Mechanistic. This sentence establishes the biological process (VDJ recombination) that depends on classical NHEJ. It is mechanistically relevant because XRCC6 (KU70) is a core NHEJ component; the statement frames why defects in NHEJ genes can produce immunological disease phenotypes. Limitation: it does not mention XRCC6/KU70 explicitly and is background rather than data on XRCC6.

- Directly relevant background claim. This sentence explicitly lists KU70/80 among end-ligation factors and states that defects in these factors are drivers of disease, specifically SCID. This is the most direct statement in the paper connecting KU70/80 (the protein complex that includes the XRCC6 gene product, Ku70) to Mendelian disease. Strength: clear assertion linking KU70/80 dysfunction to disease. Limitation: it is background text; the paper does not present new genetic or clinical data on XRCC6 mutations in human Mendelian disease, nor does it cite specific human cases or genetic evidence within this excerpt.

- Phenotypic evidence (experimental) / Indirect. This sentence shows that loss of SETD2 causes lymphoid depletion and aberrant hematopoiesis in mice, i.e., an immunodeficiency-like phenotype resulting from perturbation of a pathway that includes NHEJ factors. Mechanistic relevance: supports the view that disruption of factors that affect NHEJ or its regulation can produce Mendelian-like immunodeficiency phenotypes. Limitation: this is about Setd2, not XRCC6; it is circumstantial evidence that perturbing the NHEJ pathway causes disease.

- Mechanistic/Phenotypic. This sentence indicates that Setd2 loss blocks B- and T-cell development at stages where VDJ (TCRβ) rearrangement occurs, implicating defective VDJ recombination. Relevance: reinforces that defects in proteins affecting VDJ recombination can produce profound immunological defects. Limitation: again, pertains to Setd2 and H3K36me3; does not provide XRCC6-specific genetic evidence.

- Mechanistic inference. The authors state that loss of Setd2 phenocopies defects of NHEJ end-ligation factors and therefore suggest SETD2/H3K36me3 act in end-ligation. This links the observed phenotype to the same mechanistic node where XRCC6 (Ku70) operates. Strength: provides mechanistic plausibility that disruption of end-ligation causes disease; limitation: this is inference/phenocopy rather than demonstration that XRCC6 itself causes disease.

- Mechanistic consequence. This sentence summarizes known consequences of end-ligation defects (formation of aberrant coding-signal hybrid joints). It supports the pathogenic mechanism by which defects in end-ligation factors (including KU70/80) can lead to genome instability and dysfunctional lymphocytes. Limitation: general mechanistic statement, not XRCC6-specific evidence.

- Experimental result (mechanistic). This sentence describes that Setd2KO cells form hybrid joints upon induction, an outcome characteristic of defective end-ligation. Relevance: provides experimental evidence that perturbing components or regulators of the end-ligation machinery produces molecular lesions consistent with disease-causing defects that would be expected if KU70/80 (XRCC6) were defective. Limitation: result pertains to Setd2KO; no direct data on XRCC6 sequence variants, patient phenotypes, or Mendelian segregation.


[Read Paper](https://www.semanticscholar.org/paper/f5fdd43dda812ad4a1898f2d2a9b5f682c364730)


## Other Reviewed Papers


### Somatic genetic rescue in Mendelian haematopoietic diseases

**Why Not Relevant**: The provided paper content does not mention XRCC6, nor does it present any gene-specific data, results, or discussion that could be used to support or refute the claim that "The gene XRCC6 is likely associated with at least one Mendelian disease." The entire supplied text is a single, general sentence: "A growing number of studies of Mendelian haematopoietic disorders indicate that somatic genetic events can offset the pathogenic effect of germline mutations at the cellular level, leading to genetic mosaicism and, in some cases, resulting in a milder disease phenotype." This sentence: (1) is background commentary about the role of somatic genetic events and mosaicism in modifying Mendelian haematopoietic disorders, (2) contains no mention of XRCC6 or any other specific gene, (3) provides no experimental methods, data, statistics, or case reports that could link XRCC6 to a Mendelian phenotype, and (4) therefore supplies neither direct evidence (e.g., observations of pathogenic XRCC6 germline variants segregating with disease) nor mechanistic evidence specific to XRCC6 (e.g., functional assays, pathways implicating XRCC6 in a Mendelian disease mechanism). The sentence is general mechanistic context about somatic variation potentially modifying germline pathogenicity, which could be conceptually relevant if independent evidence established XRCC6 germline pathogenicity and showed somatic events modulating its effect, but by itself it does not provide any such evidence. Important limitations of the provided content: it is a single general statement with no experimental detail, no data, no mention of XRCC6, and therefore cannot be used to assess the claim.


[Read Paper](https://www.semanticscholar.org/paper/2bdb86e118f234bb10c7b432ccad62f0c6be1839)


### Familial focal segmental glomerulosclerosis associated with an ACTN4 mutation and paternal germline mosaicism.

**Why Not Relevant**: The provided paper excerpt does not mention XRCC6 anywhere. It exclusively discusses ACTN4 as the causal gene for focal segmental glomerulosclerosis (FSGS1), describes a novel ACTN4 mutation (p.Ser262Phe), notes autosomal dominant inheritance, reports germline mosaicism in the father, and mentions a heterozygous NPHS1 p.Thr5Met variant of unclear significance. Because XRCC6 is not referenced, this paper provides neither direct genetic evidence (variant–disease associations) nor mechanistic data (biochemical pathways, functional assays, animal models) relevant to the claim that XRCC6 is likely associated with at least one Mendelian disease. Absence of mention in this paper does not disprove an association elsewhere, but it means this paper cannot be used to support or refute the claim. Therefore relevance is 0.0.


[Read Paper](https://www.semanticscholar.org/paper/16a08fd6ebb15730e4e5a44956e57231edeef62a)


### Trio-based whole exome sequencing in patients with suspected sporadic inborn errors of immunity: A retrospective cohort study

**Why Not Relevant**: Relevance is essentially zero because the paper focuses on de novo variants in a cohort of patients with suspected inborn errors of immunity and reports candidate DNVs in specific immune-related genes (NLRP3, RELA, PSMB10, DDX1, KMT2C, FBXW11). XRCC6 is not mentioned anywhere in the provided content, so there is no direct evidence (case reports, genetic findings) or mechanistic evidence (functional studies) from this paper linking XRCC6 to any Mendelian disease. Important caveats: absence of XRCC6 in this paper is not evidence that XRCC6 is not disease-associated — the study's design (IEI-focused, DNV-centric, 123-person cohort, trio WES) could easily miss XRCC6 associations arising from inherited variants, non-coding changes, structural variants, different clinical phenotypes outside IEI, or larger cohorts. Therefore this paper provides no support for the claim, but it also does not constitute a definitive refutation.


[Read Paper](https://www.semanticscholar.org/paper/cdde2232daad22b7d92ec15faa38b9776f1e4c15)


### Fanconi anaemia, BRCA2 and familial considerations – follow up on a previous case report

**Why Not Relevant**: The paper discusses BRCA2 mutations and their relationship to Fanconi anaemia and familial breast/ovarian cancer. There is no mention of XRCC6 (no occurrences of the gene name, protein, or synonyms) nor any data, analyses, or mechanistic discussion involving XRCC6. Therefore the paper provides no direct or mechanistic evidence for or against the claim that XRCC6 is likely associated with a Mendelian disease. The excerpts above are provided to show the paper's focus on BRCA2 and FA and to explain why it is not relevant to XRCC6.


[Read Paper](https://www.semanticscholar.org/paper/55a84a17c03762d80eac3a4ee4da2c55be58645a)


### Germline APC Mutation and Familial Barrett Esophagus: Causal or Coincidence?

**Why Not Relevant**: This paper does not mention XRCC6 (or its protein product Ku70) anywhere in the provided content. The article focuses on familial Barrett esophagus, esophageal adenocarcinoma, and germline mutations in the APC gene causing FAP/AFAP, as well as Wnt/β-catenin pathway alterations and APC promoter hypermethylation. While the paper discusses Mendelian inheritance in the context of APC-related disorders and possible autosomal dominant susceptibility alleles for familial Barrett esophagus, there is no direct experimental, clinical, genetic, or mechanistic evidence presented that links XRCC6 to any Mendelian disease. Therefore it provides neither direct evidence (e.g., reported pathogenic variants, segregation with disease, case reports) nor mechanistic evidence (e.g., pathway interactions implicating XRCC6 in the diseases discussed) relevant to the claim. Because XRCC6 is not mentioned, no verbatim excerpts can be extracted that would support or refute the claim. Limitations: assessment is restricted to the supplied text; if other sections of the paper (not provided) mention XRCC6, they were not available for review. Based on the provided content alone, the paper is irrelevant to the claim that XRCC6 is likely associated with at least one Mendelian disease.


[Read Paper](https://www.semanticscholar.org/paper/3d4cf4dffb9bfae5c0410596be5f39acbb54d989)


### What a ‘Ku’incidence!: parallel discoveries of a new DNA repair factor

**Why Not Relevant**: The paper content provided mentions identification of XLS/PAXX as a new component of the NHEJ pathway and notes NHEJ is the main double-strand break repair mechanism in G1 and early S phase. However, it does not mention XRCC6 (Ku70) by name, does not present any human genetic or clinical data, and contains no information about Mendelian disease associations, pathogenic variants, segregation, phenotypes, or disease mechanism for XRCC6. Because the claim concerns a likely association between XRCC6 and at least one Mendelian disease, and the paper contains neither direct genetic/disease evidence nor explicit discussion of XRCC6, the content is not relevant (relevance < 0.1) for supporting or refuting the claim beyond providing very general pathway context.


[Read Paper](https://www.semanticscholar.org/paper/e5dd3134dd31e154919e78b9a377d1cb04afe504)


### Case Report: A novel de novo variant of COL1A1 in fetal genetic osteogenesis imperfecta

**Why Not Relevant**: This paper does not provide any direct or mechanistic evidence relevant to the claim that XRCC6 is likely associated with at least one Mendelian disease. The study exclusively reports identification and analysis of a de novo COL1A1 variant (c.1309G>A, p.Gly437Ser) in a fetus with osteogenesis imperfecta (OI) and uses trio-WES and Sanger sequencing to validate that variant. Representative verbatim text from the paper shows the focus on COL1A1 and OI: "Both WES and Sanger sequencing identified a novel de novo variant of COL1A1 (c. 1309G>A, p. Gly437Ser) in a fetus with OI." and "Conclusion This study revealed that the c.1309G>A (p. Gly437Ser) variant in the COL1A1 gene may be the genetic cause of fetal OI in this case." Nowhere in the provided content is XRCC6 mentioned, nor are DNA-repair pathways, Ku70/Ku80, or any mechanistic links that would connect COL1A1 findings to XRCC6. Because of this complete absence of mention or analysis of XRCC6 (no genetic variants reported in XRCC6, no functional assays, no clinical correlations, and no mechanistic discussion involving XRCC6), the paper cannot be used to support or refute the claim. Note: absence of evidence in this paper is not evidence that XRCC6 is or is not associated with a Mendelian disease; it simply means this paper is irrelevant to that specific gene. If you want evidence regarding XRCC6, consult gene-disease resources (OMIM, ClinVar, HGMD), population databases (gnomAD), and PubMed articles specifically mentioning XRCC6/Ku70 and clinical phenotypes.


[Read Paper](https://www.semanticscholar.org/paper/1caeaf6717a5af526be5f1f0dcef41ef4c377699)


### The simplest explanation does not have to be preferred: co-occurrence of pathogenic variants in cancer-predisposing genes

**Why Not Relevant**: The provided paper content consists of a single methodological/introductory sentence about reviewing MINAS frequency and cancer phenotypes using recent literature and UK 10,000 Genomes Project data, and about whether phenotypes are additive or synergistic across cancer susceptibility genes (CSGs). It does not mention the gene XRCC6, any specific genes, any genetic variants, inheritance patterns, segregation data, functional assays, statistical associations, or Mendelian disease phenotypes. Therefore there is no direct evidence (e.g., pathogenic XRCC6 variants segregating with disease, case reports, case-control enrichment) and no mechanistic evidence (e.g., loss-of-function in XRCC6 producing a Mendelian phenotype) in the provided content to support or refute the claim that XRCC6 is likely associated with at least one Mendelian disease.

Limitations and caveats: (1) The assessment is limited to the exact text supplied. The full paper (not provided here) might contain XRCC6-specific data elsewhere; that cannot be evaluated. (2) The excerpt is general and methodological; it only indicates that the authors reviewed cancer-susceptibility genes and MINAS in UK10K data, but without gene-level results or quotations, no inference about XRCC6 is possible. (3) To support the claim, one would need verbatim sentences reporting XRCC6-specific findings (e.g., identified pathogenic variants, segregation evidence, functional impact consistent with a Mendelian mechanism, or statistically significant enrichment in affected individuals). None of those are present in the supplied content.


[Read Paper](https://www.semanticscholar.org/paper/a1372a664bd2c33a0bda638975a8b26d112457fc)


### The application of whole-exome sequencing in the early diagnosis of rare genetic diseases in children: a study from Southeastern China

**Why Not Relevant**: The provided paper content does not mention the gene XRCC6, any variants in XRCC6, nor any phenotypes or mechanistic data tied to XRCC6. The text is a high-level summary of a retrospective whole-exome sequencing (WES) study reporting overall diagnostic rates, differences between Trio- and singleton-WES, phenotype groups (e.g., global developmental delay, muscular hypotonia), inheritance mode distributions (autosomal dominant predominance, de novo mutation fraction), and sex chromosome findings. None of these sentences or statistics provide direct or mechanistic evidence that XRCC6 is associated with any Mendelian disease. To support the claim that XRCC6 is likely associated with at least one Mendelian disease, the paper would need to include verbatim evidence such as (a) identification of pathogenic or likely pathogenic variants in XRCC6 in one or more affected individuals, (b) segregation data showing co-segregation with disease in families, (c) functional or mechanistic studies demonstrating how XRCC6 variants disrupt protein function relevant to the phenotype, or (d) population/variant filtering and statistical enrichment analyses implicating XRCC6. None of those elements are present in the provided content, so the paper cannot be used as evidence for or against the claim.


[Read Paper](https://www.semanticscholar.org/paper/ba9aa0637319a74b3dfdae65099a70edd6c60ebf)


## Search Queries Used

- XRCC6 Ku70 germline mutation Mendelian disorder case report clinical phenotype

- XRCC6 Ku70 whole exome sequencing rare disease trio Mendelian study de novo variant

- XRCC6 Ku70 non homologous end joining DNA repair knockout mouse phenotype immunodeficiency cancer predisposition telomere maintenance

- XRCC6 ClinVar OMIM ClinGen pathogenic variant review gene disease association gnomad constraint pLI


## Usage Statistics

- Prompt Tokens: 0

- Completion Tokens: 0

- Total Tokens: 0

- Estimated Cost: $0.0413


## Bibliometric Configuration

- Use Bibliometrics: True

- Author Impact Weight: 0.4

- Citation Impact Weight: 0.4

- Venue Impact Weight: 0.2
